# Cortical Thickness Trajectories across the Lifespan: Data from 17,075 healthy individuals aged 3-90 years

**DOI:** 10.1101/2020.05.05.077834

**Authors:** Sophia Frangou, Amirhossein Modabbernia, Gaelle E Doucet, Efstathios Papachristou, Steven CR Williams, Ingrid Agartz, Moji Aghajani, Theophilus N Akudjedu, Anton Albajes-Eizagirre, Dag Alnæs, Kathryn I Alpert, Micael Andersson, Nancy Andreasen, Ole A Andreassen, Philip Asherson, Tobias Banaschewski, Nuria Bargallo, Sarah Baumeister, Ramona Baur-Streubel, Alessandro Bertolino, Aurora Bonvino, Dorret I Boomsma, Stefan Borgwardt, Josiane Bourque, Daniel Brandeis, Alan Breier, Henry Brodaty, Rachel M Brouwer, Jan K Buitelaar, Geraldo F Busatto, Randy L Buckner, Vincent Calhoun, Erick J Canales-Rodríguez, Dara M Cannon, Xavier Caseras, Francisco X Castellanos, Simon Cervenka, Tiffany M Chaim-Avancini, Christopher RK Ching, Vincent P Clark, Patricia Conrod, Annette Conzelmann, Benedicto Crespo-Facorro, Fabrice Crivello, Eveline AM Crone, Anders M Dale, Cristopher Davey, Eco JC de Geus, Lieuwe de Haan, Greig I de Zubicaray, Anouk den Braber, Erin W Dickie, Annabella Di Giorgio, Nhat Trung Doan, Erlend S Dørum, Stefan Ehrlich, Susanne Erk, Thomas Espeseth, Helena Fatouros-Bergman, Simon E Fisher, Jean-Paul Fouche, Barbara Franke, Thomas Frodl, Paola Fuentes-Claramonte, David C Glahn, Ian H Gotlib, Hans-Jörgen Grabe, Oliver Grimm, Nynke A Groenewold, Dominik Grotegerd, Oliver Gruber, Patricia Gruner, Rachel E Gur, Ruben C Gur, Ben J Harrison, Catharine A Hartman, Sean N Hatton, Andreas Heinz, Dirk J Heslenfeld, Derrek P Hibar, Ian B Hickie, Beng-Choon Ho, Pieter J Hoekstra, Sarah Hohmann, Avram J Holmes, Martine Hoogman, Norbert Hosten, Fleur M Howells, Hilleke E Hulshoff Pol, Chaim Huyser, Neda Jahanshad, Anthony James, Jiyang Jiang, Erik G Jönsson, John A Joska, Rene Kahn, Andrew Kalnin, Ryota Kanai, Sim Kang, Marieke Klein, Tatyana P Klushnik, Laura Koenders, Sanne Koops, Bernd Krämer, Jonna Kuntsi, Jim Lagopoulos, Luisa Lázaro, Irina Lebedeva, Won Hee Lee, Klaus-Peter Lesch, Christine Lochner, Marise WJ Machielsen, Sophie Maingault, Nicholas G Martin, Ignacio Martínez-Zalacaín, David Mataix-Cols, Bernard Mazoyer, Colm McDonald, Brenna C McDonald, Andrew M McIntosh, Katie L McMahon, Genevieve McPhilemy, José M Menchón, Sarah E Medland, Andreas Meyer-Lindenberg, Jilly Naaijen, Pablo Najt, Tomohiro Nakao, Jan E Nordvik, Lars Nyberg, Jaap Oosterlaan, Víctor Ortiz-García de la Foz, Yannis Paloyelis, Paul Pauli, Giulio Pergola, Edith Pomarol-Clotet, Maria J Portella, Steven G Potkin, Joaquim Radua, Andreas Reif, Joshua L Roffman, Pedro GP Rosa, Matthew D Sacchet, Perminder S Sachdev, Raymond Salvador, Pascual Sánchez-Juan, Salvador Sarró, Theodore D Satterthwaite, Andrew J Saykin, Mauricio H Serpa, Lianne Schmaal, Knut Schnell, Gunter Schumann, Jordan W Smoller, Iris Sommer, Carles Soriano-Mas, Dan J Stein, Lachlan T Strike, Suzanne C Swagerman, Christian K Tamnes, Henk S Temmingh, Sophia I Thomopoulos, Alexander S Tomyshev, Diana Tordesillas-Gutiérrez, Julian N Trollor, Jessica A Turner, Anne Uhlmann, Odille A van den Heuvel, Dennis van den Meer, Nic JA van der Wee, Neeltje EM van Haren, Dennis van ’t Ent, Theo GM van Erp, Ilya M Veer, Dick J Veltman, Henry Völzke, Henrik Walter, Esther Walton, Lei Wang, Yang Wang, Thomas H Wassink, Bernd Weber, Wei Wen, John D West, Lars T Westlye, Heather Whalley, Lara M Wierenga, Katharina Wittfeld, Daniel H Wolf, Margaret J Wright, Kun Yang, Yulyia Yoncheva, Marcus V Zanetti, Georg C Ziegler, Paul M Thompson, Danai Dima

**Affiliations:** Department of Psychiatry, Icahn School of Medicine at Mount Sinai, USA; Department of Psychiatry, Djavad Mowafaghian Centre for Brain Health, University of British Columbia, Canada; Boys Town National Research Hospital, University of Nebraska Medical Center, USA; Psychology and Human Development, Institute of Education, University College London, UK; Department of Neuroimaging, Institute of Psychiatry, Psychology and Neuroscience, King’s College London, UK; Norwegian Centre for Mental Disorders Research (NORMENT), Institute of Clinical Medicine, University of Oslo, Norway; Department of Psychiatric Research, Diakonhjemmet Hospital, Norway; Centre for Psychiatric Research, Department of Clinical Neuroscience, Karolinska Institutet, Sweden; Department of Psychiatry, Amsterdam University Medical Centre, Vrije Universiteit, Netherlands; Clinical Neuroimaging Laboratory, Centre for Neuroimaging and Cognitive Genomics and NCBES Galway Neuroscience Centre, National University of Ireland, Ireland; FIDMAG Germanes Hospitalàries, Spain; Mental Health Research Networking Center (CIBERSAM), Spain; Division of Mental Health and Addiction, Institute of Clinical Medicine, University of Oslo, Norway; Radiologics, INC, USA; Umeå Center for Functional Brain Imaging, Umeå University, Sweden; Department of Integrative Medical Biology, Umeå University, Sweden; Department of Psychiatry, Carver College of Medicine, University of Iowa, USA; Social, Genetic and Developmental Psychiatry Centre, Institute of Psychiatry, Psychology and Neuroscience, King’s College London, London, UK; Department of Child and Adolescent Psychiatry and Psychotherapy, Central Institute of Mental Health, Heidelberg University, Germany; Imaging Diagnostic Centre, Barcelona University Clinic, Spain; Department of Psychology, Biological Psychology, Clinical Psychology and Psychotherapy, University of Würzburg, Germany; Department of Basic Medical Science, Neuroscience and Sense Organs, University of Bari Aldo Moro, Italy; Department of Biological Psychology, Vrije Universiteit, Netherlands; Department of Psychiatry & Psychotherapy, University of Lübeck, Germany; Department of Psychiatry, University of Pennsylvania, Philadelphia, USA; Department of Radiology and Imaging Sciences, Indiana University School of Medicine, USA; Centre for Healthy Brain Ageing, University of New South Wales, Australia; Rudolf Magnus Institute of Neuroscience, University Medical Center Utrecht, Netherlands; Donders Center of Medical Neurosciences, Radboud University, Netherlands; Donders Centre for Cognitive Neuroimaging, Radboud University, Netherlands; Donders Institute for Brain, Cognition and Behaviour, Radboud University, Netherlands; Laboratory of Psychiatric Neuroimaging, Departamento e Instituto de Psiquiatria, Hospital das Clinicas HCFMUSP, Faculdade de Medicina, Universidade de São Paulo, Brazil; Department of Psychology, Center for Brain Science, Harvard University, USA; Department of Psychiatry, Massachusetts General Hospital, USA; College of Arts and Sciences, Georgia State University, USA; Computer Science, Math, Neuroscience and Physics at Georgia State University, USA; Electrical and Computer Engineering and Biomedical Engineering, Georgia Institute of Technology, USA; Neurology, Radiology, Psychiatry and Biomedical Engineering Emory University, USA; MRC Centre for Neuropsychiatric Genetics and Genomics, Cardiff University, UK; Department of Child and Adolescent Psychiatry, New York University, USA; Stockholm Health Care Services, Stockholm County Council, Karolinska University Hospital, Sweden; Imaging Genetics Center, Mark and Mary Stevens Neuroimaging and Informatics Institute, Keck School of Medicine, University of Southern California, Marina del Rey, USA; Department of Psychology, University of New Mexico, USA; Mind Research Network, USA; Department of Psychiatry, Université de Montréal, Canada; Department of Child and Adolescent Psychiatry, Psychosomatics and Psychotherapy, University of Tübingen, Germany; HU Virgen del Rocio, IBiS, University of Sevilla, Spain; Groupe d’Imagerie Neurofonctionnelle, Institut des Maladies Neurodégénératives, UMR5293, Université de Bordeaux, France; Faculteit der Sociale Wetenschappen, Instituut Psychologie, Universiteit Leiden, Netherlands; Center for Multimodal Imaging and Genetics, University of California San Diego, USA; Department of Neurosciences, University of California San Diego USA; Department of Radiology, University of California San Diego, USA; Centre for Youth Mental Health, University of Melbourne, Australia; Orygen, Australia; Academisch Medisch Centrum, Universiteit van Amsterdam, Netherlands; Faculty of Health, Institute of Health and Biomedical Innovation, Queensland University of Technology, Australia; Kimel Family Translational Imaging Genetics Research Laboratory, University of Toronto, Canada; Campbell Family Mental Health Research Institute, The Centre for Addiction and Mental Health, University of Toronto, Canada; Department of Experimental and Clinical Medicine, Università Politecnica delle Marche, Italy; Department of Psychology, University of Oslo, Norway; Sunnaas Rehabilitation Hospital HT, Nesodden, Norway; Klinik und Poliklinik für Kinder und Jugendpsychiatrie und Psychotherapie, Universitätsklinikum Carl Gustav Carus an der TU Dresden, Germany; Division of Mind and Brain Research, Department of Psychiatry and Psychotherapy, Charité-Universitätsmedizin Berlin, Germany; Language and Genetics Department, Max Planck Institute for Psycholinguistics, Netherlands; Department of Psychiatry and Mental Health, University of Cape Town, South Africa; Department of Psychiatry and Psychotherapy, Otto von Guericke University Magdeburg, Germany; Department of Psychiatry, Tommy Fuss Center for Neuropsychiatric Disease Research Boston Children’s Hospital, Harvard Medical School, USA; Department of Psychology, Stanford University, USA; Department of Psychiatry and Psychotherapy, University Medicine Greifswald, University of Greifswald, Germany; Department for Psychiatry, Psychosomatics and Psychotherapy, Universitätsklinikum Frankfurt, Goethe Universitat, Germany; Neuroscience Institute, University of Cape Town, South Africa; Department of Psychiatry and Psychotherapy, University of Muenster, Germany; Section for Experimental Psychopathology and Neuroimaging, Department of General Psychiatry, Heidelberg University, Heidelberg, Germany; Department of Psychiatry, Yale University, USA; Learning Based Recovery Center, VA Connecticut Health System, USA; Lifespan Brain Institute, Perelman School of Medicine, University of Pennsylvania, USA; Children’s Hospital of Philadelphia, University of Pennsylvania, USA; Melbourne Neuropsychiatry Center, University of Melbourne, Australia; Interdisciplinary Center Psychopathology and Emotion regulation, University Medical Center Groningen, University of Groningen, Netherlands; Brain and Mind Centre, University of Sydney, Australia; Departments of Experimental and Clinical Psychology, Vrije Universiteit Amsterdam, Netherlands; Personalized Healthcare, Genentech, Inc., USA; Department of Psychiatry, University Medical Center Groningen, University of Groningen, Netherlands; Department of Psychology, Yale University, USA; Radboud University Medical Center, The Netherlands; Norbert Institute of Diagnostic Radiology and Neuroradiology, University Medicine Greifswald, University of Greifswald, Germany; Bascule, Academic Centre for Children and Adolescent Psychiatry, Netherlands; Department of Psychiatry, Oxford University, UK; Depart of Radiology, The Ohio State University College of Medicine, USA; Department of Neuroinformatics, Araya, Inc., Japan; Institute of Mental Health, Singapore; Laboratory of Neuroimaging and Multimodal Analysis, Mental Health Research Center, Russian Academy of Medical Sciences, Russia; Sunshine Coast Mind and Neuroscience, Thompson Institute, University of the Sunshine Coast, Australia; Department of Child and Adolescent Psychiatry and Psychology, Barcelona University Clinic, Spain; August Pi i Sunyer Biomedical Research Institut (IDIBAPS), Spain; Mental Health Research Center, Moscow, Russia; Department of Psychiatry, Psychosomatics and Psychotherapy, Julius-Maximilians Universität Würzburg, Germany; SA MRC Unit on Risk and Resilience in Mental Disorders, Department of Psychiatry, Stellenbosch University, South Africa; Queensland Institute of Medical Research, Berghofer Medical Research Institute, Australia; Department of Psychiatry, Bellvitge University Hospital-IDIBELL, University of Barcelona, Spain; Division of Psychiatry, University of Edinburgh, UK; Herston Imaging Research Facility, Queensland University of Technology, Australia; Department of Psychiatry and Psychotherapy, Central Institute of Mental Health, Heidelberg University, Germany; Department of Clinical Medicine, Kyushu University, Japan; CatoSenteret Rehabilitation Hospital, Norway; Department of Clinical Neuropsychology, Amsterdam University Medical Centre, Vrije Universiteit Amsterdam, Netherlands; Department of Psychiatry, University Hospital “Marques de Valdecilla”, Instituto de Investigación Valdecilla (IDIVAL), Spain; Instituto de Salud Carlos III, Spain; Centre of Mental Health, University of Würzburg, Germany; Department of Psychiatry, Hospital de la Santa Creu i Sant Pau, Institut d’Investigació Biomèdica Sant Pau, Universitat Autònoma de Barcelona, Spain; Department of Psychiatry, University of California at Irvine, USA; Department of Psychosis Studies, Institute of Psychiatry, Psychology & Neuroscience, King’s College London, UK; Center for Depression, Anxiety, and Stress Research, McLean Hospital, Harvard Medical School, USA; Centro de Investigacion Biomedica en Red en Enfermedades Neurodegenerativas (CIBERNED), Spain; Department of Psychiatry and Psychotherapy, University Medical Center Göttingen, Germany; Centre for Population Neuroscience and Precision Medicine, Institute of Psychiatry, Psychology & Neuroscience, King’s College London, UK; Center for Genomic Medicine, Massachusetts General Hospital, USA; Department of Biomedical Sciences of Cells and Systems, Rijksuniversiteit Groningen, University Medical Center Groningen, Netherlands; Queensland Brain Institute, University of Queensland, Australia; PROMENTA Research Center, Department of Psychology, University of Oslo, Norway; Neuroimaging Unit, Technological Facilities, Valdecilla Biomedical Research Institute IDIVAL, Spain; School of Mental Health and Neuroscience, Faculty of Health, Medicine and Life Sciences, Maastricht University, Netherlands; Department of Psychiatry, Leiden University Medical Center, Netherlands; Leiden Institute for Brain and Cognition, Netherlands; Department of Child and Adolescent Psychiatry/Psychology, Erasmus University Medical Center, Sophia Children’s Hospital, Rotterdam, The Netherlands; Center for the Neurobiology of Learning and Memory, University of California Irvine, USA; Institute of Community Medicine, University Medicine, Greifswald, University of Greifswald, Germany; MRC Integrative Epidemiology Unit, Population Health Sciences Bristol Medical School, Bristol University, UK; Department of Psychiatry and Behavioral Sciences, Feinberg School of Medicine, Northwestern University, USA; Department of Radiology, Medical College of Wisconsin, USA; Institute for Experimental Epileptology and Cognition Research, University of Bonn, Germany; Developmental and Educational Psychology Unit, Institute of Psychology, Leiden University, Netherlands; German Center for Neurodegenerative Diseases (DZNE), Rostock/Greifswald Site, University of Greifswald, Germany; National High Magnetic Field Laboratory, Florida State University, USA; Division of Molecular Psychiatry, Center of Mental Health, University of Würzburg, Germany; Department of Psychology, School of Arts and Social Sciences, City, University of London, UK

**Keywords:** Cortical Thickness, Development, Aging, Trajectories

## Abstract

Delineating age-related cortical trajectories in healthy individuals is critical given the association of cortical thickness with cognition and behaviour. Previous research has shown that deriving robust estimates of age-related brain morphometric changes requires large-scale studies. In response, we conducted a large-scale analysis of cortical thickness in 17,075 individuals aged 3-90 years by pooling data through the Lifespan Working group of the Enhancing Neuroimaging Genetics through Meta-Analysis (ENIGMA) Consortium. We used fractional polynomial (FP) regression to characterize age-related trajectories in cortical thickness, and we computed normalized growth centiles using the parametric Lambda, Mu, and Sigma (LMS) method. Inter-individual variability was estimated using meta-analysis and one-way analysis of variance. Overall, cortical thickness peaked in childhood and had a steep decrease during the first 2-3 decades of life; thereafter, it showed a gradual monotonic decrease which was steeper in men than in women particularly in middle-life. Notable exceptions to this general pattern were entorhinal, temporopolar and anterior cingulate cortices. Inter-individual variability was largest in temporal and frontal regions across the lifespan. Age and its FP combinations explained up to 59% variance in cortical thickness. These results reconcile uncertainties about age-related trajectories of cortical thickness; the centile values provide estimates of normative variance in cortical thickness, and may assist in detecting abnormal deviations in cortical thickness, and associated behavioural, cognitive and clinical outcomes.

## Introduction

In the last two decades, there has been a steady increase in the number of studies of age-related changes in brain morphometry (Ducharme, et al., 2015; Giedd and Rapoport, 2010; Good, et al., 2001; Hasan, et al., 2016; Kaup, et al., 2011; Mutlu, et al., 2013; Pomponio et al., 2019; Raznahan, et al., 2011; Salat, et al., 2004; Shaw, et al., 2008; Sowell, et al., 2007; Sowell, et al., 2003; Sowell, et al., 2004; Storsve, et al., 2014; Tamnes, et al., 2010; Thambisetty, et al., 2010; Vaidya, et al., 2007; Walhovd, et al., 2017; Wierenga, et al., 2014) as a means to understand the genetic and environmental influences on the human brain (Fjell and Walhovd, 2010; Raz, et al., 2005; Walhovd, et al., 2017). Here we focus specifically on cortical thickness, as assessed using magnetic resonance imaging (MRI), as this measure has established associations with behaviour and cognition in healthy populations (Burgaleta, et al., 2014; Fjell and Walhovd, 2010; Hedden and Gabrieli, 2004; Kharitonova, et al., 2013; Mills, et al., 2014; Shaw, et al., 2006a) and with disease mechanisms implicated in neuropsychiatric disorders (Boedhoe, et al., 2018; Hibar, et al., 2018; Rapoport, et al., 2001; Schmaal, et al., 2017; Shaw, et al., 2006b; Thompson, et al., 2007; Thormodsen, et al., 2013; van Erp, et al., 2016; van Rooij, et al.; Walton, et al., 2017; Whelan, et al., 2018).

Structural MRI is the most widely used neuroimaging method in research and clinical settings because of its excellent safety profile, even in repeat administration, ease of data acquisition and high patient acceptability. Thus, establishing the typical patterns of age-related trajectories in cortical thickness could be a significant first step in the translational application of neuroimaging. The value of reference data is firmly established in medicine where deviations from the expected range are used to trigger further investigations or interventions. Classic examples are the growth charts developed by the World Health Organization (http://www.who.int/childgrowth/en/) and US National Center for Health Statistics (https://www.cdc.gov/growthcharts/cdc_charts.htm) to monitor child development and the body mass index (BMI) which has been instrumental in informing scientific models and public health policies relating to cardio-metabolic health (Johnson, et al., 2015).

There is significant uncertainty about the shape and inter-individual variability of age-related trajectories. Prior studies have reported linear and non-linear associations between age and cortical thickness (e.g., (Amlien, et al., 2016; Brouwer, et al., 2017; Brown and Jernigan, 2012; Brown, et al., 2012; Mills, et al., 2014; Mutlu, et al., 2013; Raznahan, et al., 2011; Shaw, et al., 2006a; Shaw, et al., 2008; Sowell, et al., 2003; van Soelen, et al., 2012; Wierenga, et al., 2014) that may be influenced by sex (Coffey, et al., 1998; Paus, 2010; Raz, et al., 2010). The present study harnesses the power of the Enhancing Neuroimaging Genetics through Meta-Analysis (ENIGMA) Consortium, a multinational collaborative network of researchers organized into working groups that conduct large-scale analyses integrating data from over 250 institutions (Grasby, et al., 2018; Thompson, et al., 2017; Thompson, et al., 2014). Within ENIGMA, the focus of the Lifespan Working group is to delineate age-related trajectories of brain morphometry extracted from MRI images using standardized protocols and unified quality control procedures harmonized and validated across all participating sites. Moreover, the ENIGMA Lifespan dataset is the largest sample of healthy individuals available worldwide that offers the most comprehensive coverage of the human lifespan. This distinguishes the ENIGMA Lifespan dataset from other imaging samples, such as the UK Biobank (http://www.ukbiobank.ac.uk) which only includes individuals over 40 years of age. In the present study, we used MRI data from 17,075 healthy participants aged 3-90 years to define age-related trajectories and centile values for regional cortical thickness in the entire sample and for each sex. We estimated regional inter-individual variability because it represents a major source of inter-study variation on age-related effects (Dickie, et al., 2013; Raz, et al., 2010). Based on prior literature, our initial hypotheses were that cortical thickness in most regions would have an inverse U-shaped trajectory with variable rates of decline between late childhood and old age that would be influenced by sex.

## Materials and Methods

### Study Samples

De-identified demographic and cortical thickness data from 83 worldwide samples (Figure 1) were pooled to create the dataset analysed in this study. The pooled sample comprised 17,075 participants (52% female) aged 3-90 years (Table 1). All participants had been screened to exclude psychiatric disorders, medical and neurological morbidity and cognitive impairment. Information on the screening protocols and eligibility criteria is provided in Supplemental Table S1.

**Table 1.**
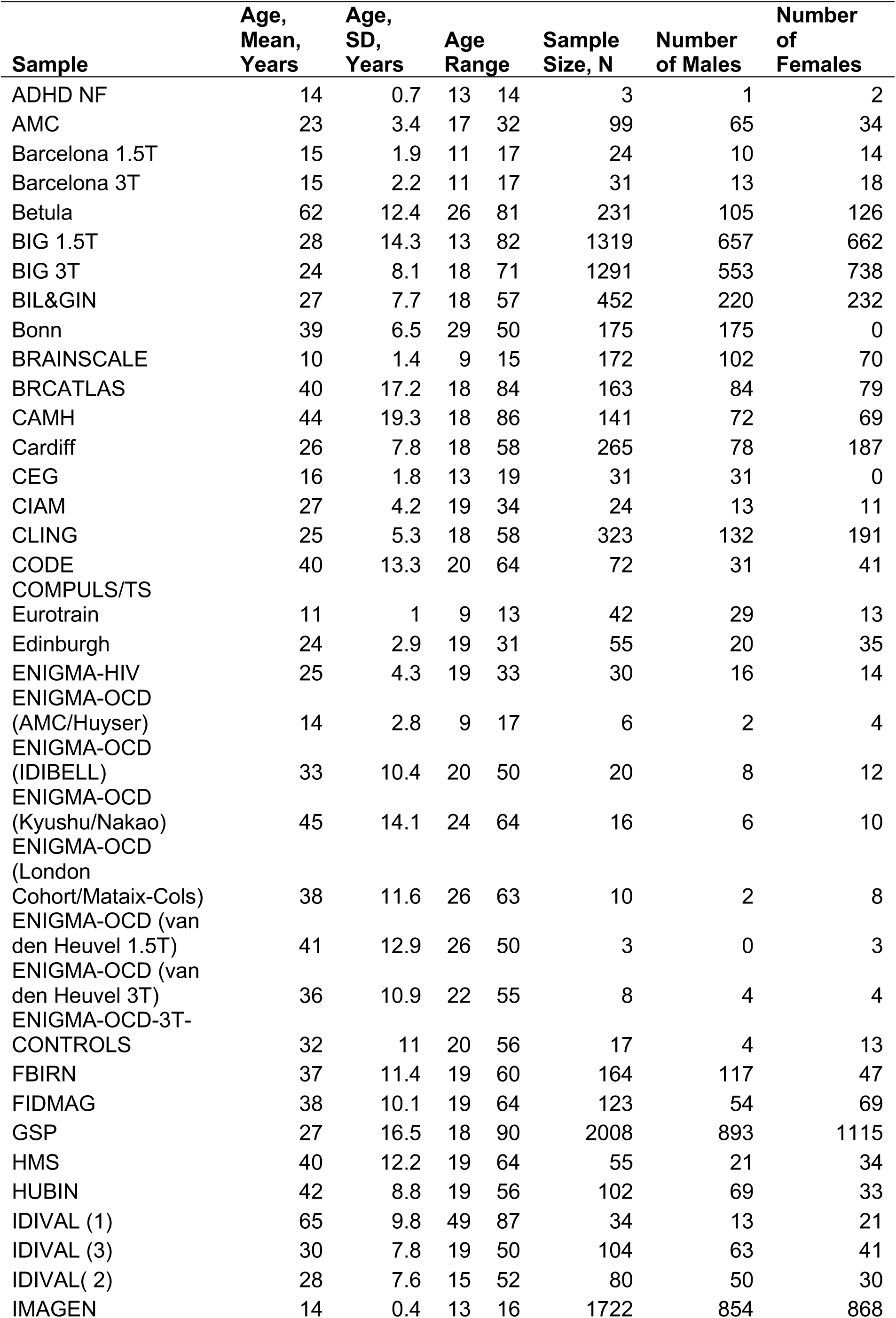
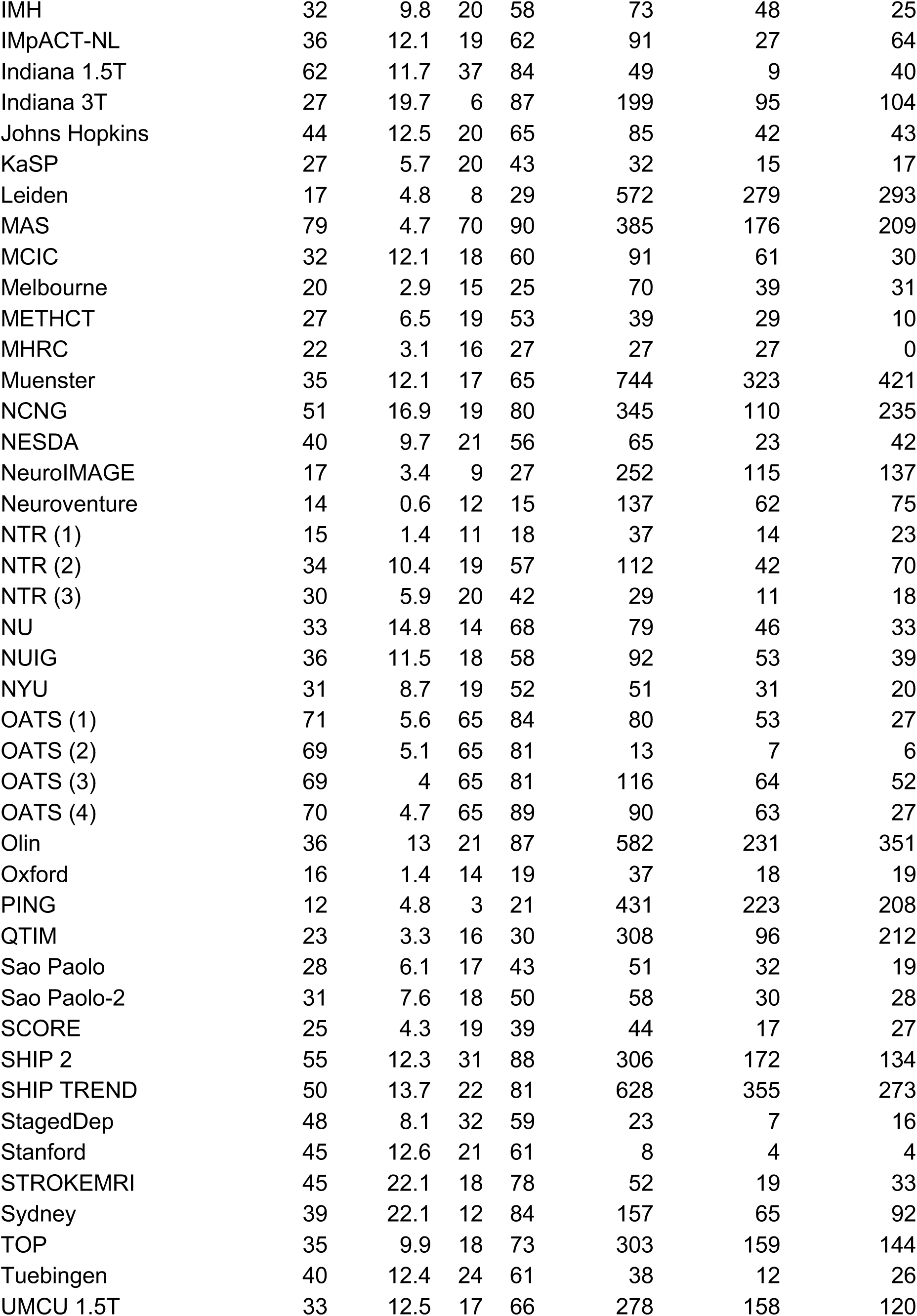
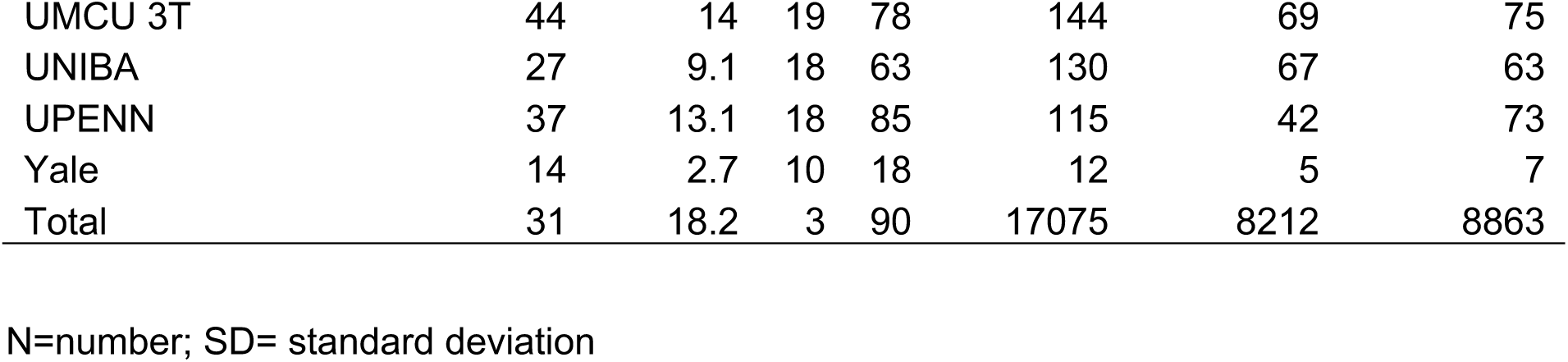
Characteristics of the included samples.

**Figure 1.**
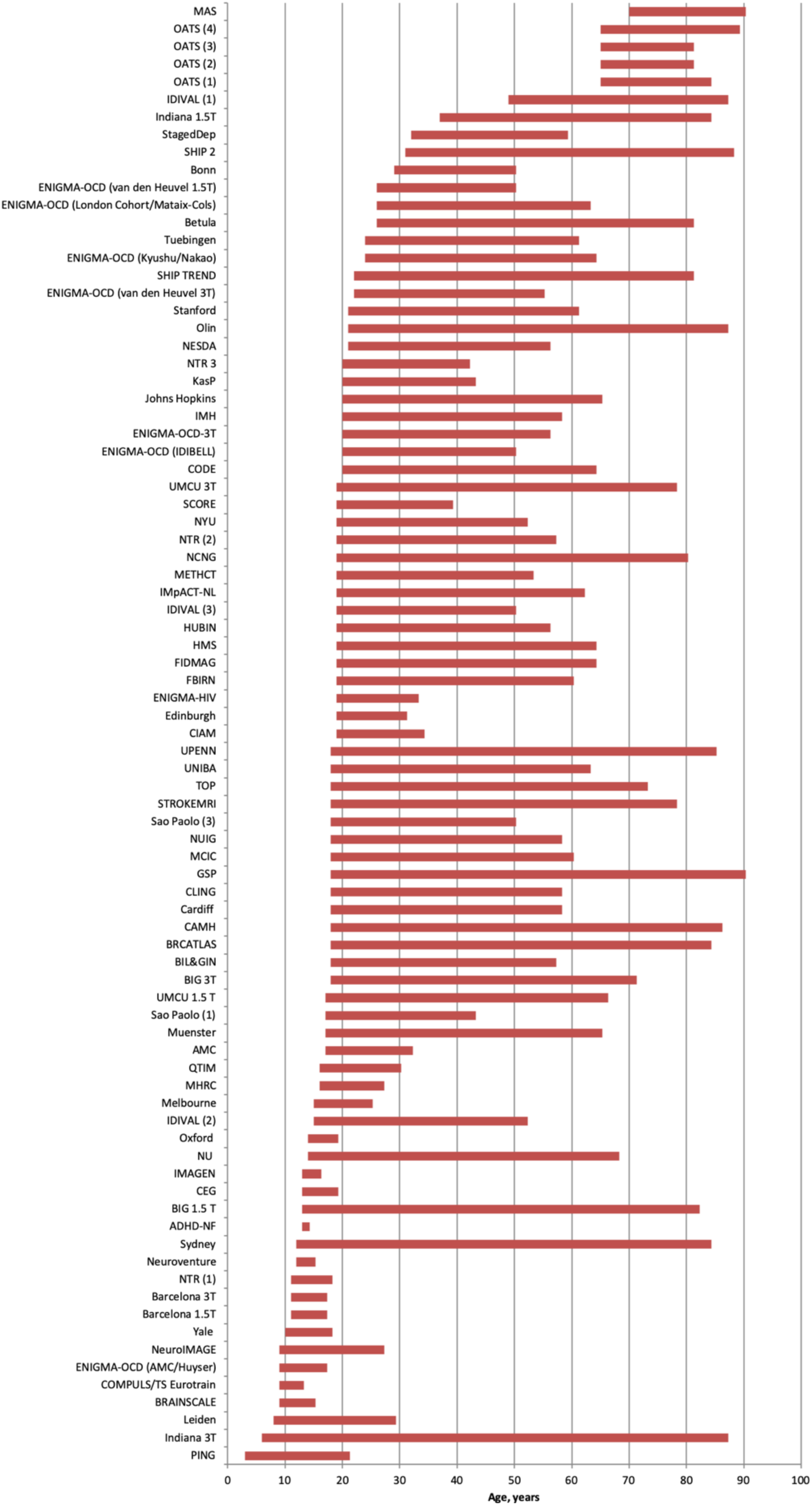
Age range for each sample. Abbreviations are explained in Table 1; further details of each sample are provided in the supplemental material.

### Image acquisition and processing

Prior to pooling the data used in this study, researchers at each participating institution (a) used the ENIGMA MRI analysis protocols, which are based on FreeSurfer (http://surfer.nmr.mgh.harvard.edu) (Fischl, 2012; Fischl, et al., 2002), to extract cortical thickness of 68 regions from high-resolution T1-weighted MRI brain scans collected at their site; (b) inspected all images by overlaying the cortical parcellations on the participants’ anatomical scans; (c) excluded improperly segmented scans and outliers identified using five median absolute deviations (MAD) of that of the median value. Information on scanner vendor, magnetic field strengths, FreeSurfer version and acquisition parameters for each sample provided by the participating institutions is detailed in Supplemental Table S1.

### Analysis of age-related trajectories in cortical thickness

We modeled the effect of age on regional cortical thickness using higher order fractional polynomial (FP) regression analyses (Royston and Altman, 1994; Sauerbrei, et al., 2006) implemented in STATA software version 14.0 (Stata Corp., College Station, TX). FP regression is one of the most flexible methods to study the effect of continuous variables on a response variable (Royston and Altman, 1994; Sauerbrei, et al., 2006). FP allows for testing a broad family of shapes and multiple turning points while simultaneously providing a good fit at the extremes of the covariates (Royston and Altman, 1994). Prior to FP regression analysis, cortical thickness values were harmonized between sites using the ComBat method in R (Fortin, et al., 2018; Fortin, et al., 201 Radua et al., 2020; this issue). Originally developed to adjust for batch effect in genetic studies, ComBat uses an empirical Bayes method to adjust for inter-site (inter-scanner) variability in the data, while preserving variability related to the variables of interest. As the effect of site and thus scanner had been adjusted for using ComBat, we only included sex as a covariate in the regression models. Additionally, standard errors were adjusted for the effect of site in the FP regression. We centred the data from each brain region so that the intercept of an FP was zero for all covariates. We used a predefined set of power terms (−2, -1, -0.5, 0.5, 1, 2, 3) and the natural logarithm function, and up to four power combinations to identify the best fitting model. FP for age is written as age^(p1,p2,…p6)^′β where p in *age*^(*p*1,*p*2,…*p*6)^ refers to regular powers except *age*^(0)^ which refers to ln(age). Powers can be repeated in FP; each time a power s repeated, it is multiplied by another ln(age). As an example:

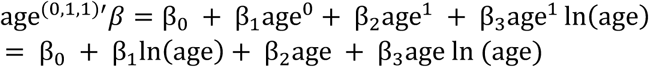

494 models were trained for each region. Model comparison was performed using a partial *F*-test and the lowest degree model with the smallest *P-value* was selected as the optimal model. Following permutation, critical alpha value was set at 0.01 to decrease the probability of over-fitting. The age at maximum cortical thickness for each cortical region was the maximum fitted value of the corresponding optimal FP model.

Further, we divided the dataset into three age-groups corresponding to early (3-29 years), middle (30-59 years) and late life (60-90 years). Within each age-group, we calculated Pearson’s correlation coefficient between age and regional cortical thickness. Finally, we used the *cocor* package in R to obtain P-values for the differences in correlation coefficients between males and females in each age-group.

### Inter-individual Variation in Cortical Thickness

The residuals of the FP regression models for each cortical region were normally distributed. Using one-way analysis of variance we extracted the residual variance around the optimal fitted FP regression model so as to identify age-group differences in inter-individual variation for each cortical region. Separately for each age-group (t), we calculated the mean age-related variance of each cortical region using 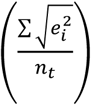 where *e*^2^ denotes the squared residual variance of that region around the best fitting FP regression line for each individual (i) of that age-group, and *n* the number of observations in that age-group. Because the square root of the squared residuals was positively skewed, we applied a natural logarithm transformation to the calculated variance. To account for multiple comparisons (68 regions assessed in three age-groups), statistical inference was based on a Bonferroni adjusted *p*-value of 0.0007 as a cut-off for a significant *F-*test. To confirm that the sample effect did not drive the inter-individual variability analyses, we also conducted a meta-analysis of the standard deviation of the regional cortical thickness in each age-group, following previously validated methodology (Senior, et al., 2016). To test whether inter-individual variability is a function of surface area (and possibly measurement error by FreeSurfer) we plotted SD values of each region against their corresponding average surface area.

### Centile Values of Cortical Thickness

We calculated the centiles (0.4, 1, 2.5, 5, 10, 25, 50, 75, 90, 95, 97.5, 99, 99.6) for each regional cortical thickness measure by sex and hemisphere as normalized growth centiles using parametric Lambda (λ), Mu (μ), Sigma (σ) (LMS) method (Cole and Green, 1992) in the Generalised Additive Models for Location, Scale and Shape (GAMLSS) package in R (http://cran.r-project.org/web/packages/gamlss/index.html) (Rigby and Stasinopoulos, 2005; Stasinopoulos and Rigby, 2007). LMS is considered a powerful method for estimating centile curves based on the distribution of a response variable at each covariate value (in this case age). GAMLSS uses a penalized maximum likelihood function to estimate parameters of smoothness (effective degrees of freedom) which are then used to estimate the λ, μ and σ parameters (Indrayan, 2014). The goodness of fit for these parameters in the GAMLSS algorithm is established by minimizing the Generalized Akaike Information Criterion (GAIC) index.

## Results

### Age-related trajectories in cortical thickness

Figure 2 shows characteristic trajectories for cortical regions in each lobe, while the trajectories of all cortical regions are provided in Supplemental File S1. For most regions, cortical thickness showed a steep decrease until the 3^rd^ decade of life, followed by a monotonic gradual decline thereafter (Supplemental Table S2). However, both entorhinal and temporopolar cortices showed an inverse U-shaped relation with age bilaterally while in the anterior cingulate cortex (ACC), cortical thickness showed an attenuated U-shaped trajectory. In general, age and its FP combinations explained up to 59% of the variance in mean cortical thickness (Supplemental Table S2). Age explained the smallest proportion of the variance for entorhinal (1-2%) and temporopolar (2-3%) cortices, whereas it explained the largest proportion of variance for superior frontal and precuneus gyri (50-52%). We observed some significant sex differences in the slopes of age-related regional cortical thickness reduction. In general, in the early-life group (3-29 years), the slopes for mean cortical thickness were not meaningfully different for males (r=-0.59) than females (r=-0.56). Similarly, in the middle-life group (30-59 years) the slopes for mean cortical thickness were steeper for men (r =-0.39 to -0.38) than for women (r=-0.27). In the late-life group (61-90 years) there was no meaningful difference between men (r-range= -0.30 to -0.29) and women (r-range= =-0.33 to -0.31) because the slopes of regional cortical thickness reduction became less pronounced in men while slightly increasing in women. At the regional level, the slope of cortical thinning in the early-life group was greater (P<0.0007) in males than in females in the bilateral cuneus, lateral occipital, lingual, superior parietal, postcentral, and paracentral, precuneus, and pericalcarine gyri. In middle-life age-group, the slope of cortical thinning was greater (P<0.0002) in men than in women in the bilateral *pars orbitalis* and *pars triangularis* as well as left isthmus of the cingulate, *pars opercularis*, precuneus, rostral middle frontal, and supramarginal, and right fusiform, inferior temporal, inferior parietal, lateral occipital, lateral orbitofrontal, rostral anterior cingulate, superior frontal, supramarginal regions and the insula (Figure 3, Supplemental Table S3, Supplemental Figure S1).

**Figure 2.**
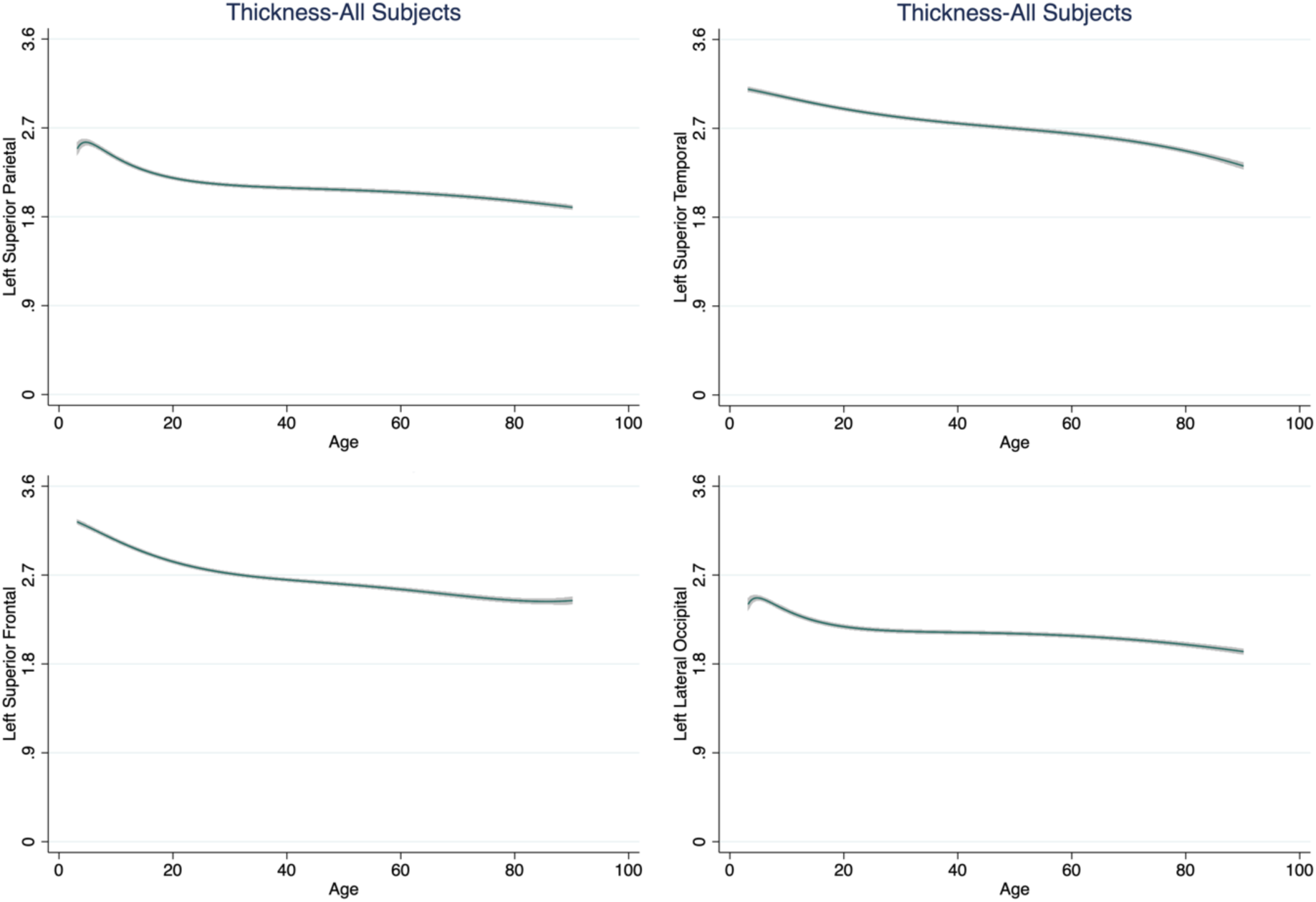
Illustrative age-related trajectories of cortical thickness. We present exemplars from each lobe as derived from fractional polynomial analyses of the entire dataset. Age-related trajectories of thickness for all cortical regions (for the entire dataset and separately for males and females) are given in the supplementary material.

**Figure 3.**
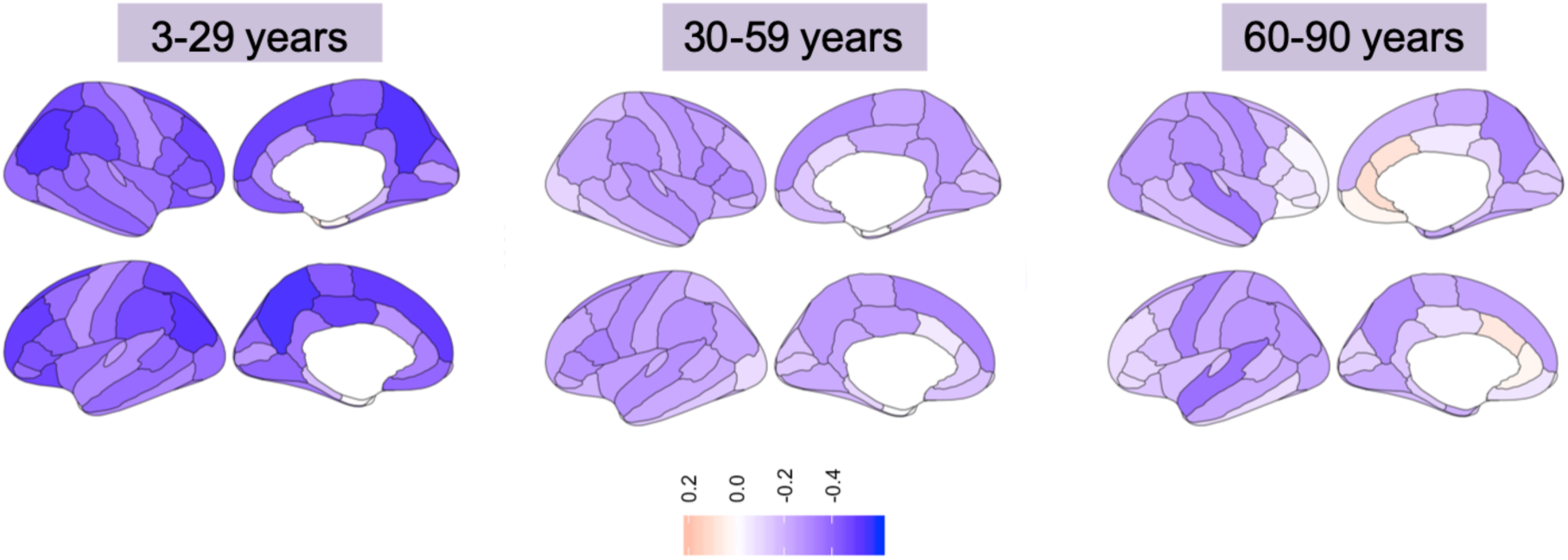
Correlation between age and cortical thickness across age-groups. Left panel: early life age-group (3-29 years); Middle panel: middle life age-group (30-59 years); Right panel: late life age-group (60-90 years).

### Inter-individual Variation in Cortical Thickness

Details of the inter-individual variation for all cortical regions in each group are provided in Supplemental Table S4, Supplemental Figure S2, and Figure 4. Across age-groups, the inter-individual variability in most cortical regions as measured by pooled SD was between0.1 and 0.2 mm. Higher levels of inter-individual variation were also observed but were mainly apparent bilaterally for the entorhinal, parahippocampal, transverse temporal, temporopolar, frontopolar, anterior and isthmus of the cingulate cortex, and the *pars orbitalis*. The meta-analysis conducted as per (Senior, et al., 2016) confirmed the replicability of these findings in each age-group (early, middle and late life). We observed a nonlinear association between regional cortical surface area and inter-individual variability in that variability was typically higher in regions with smaller surface areas (Supplementary Figure S3).

**Figure 4.**
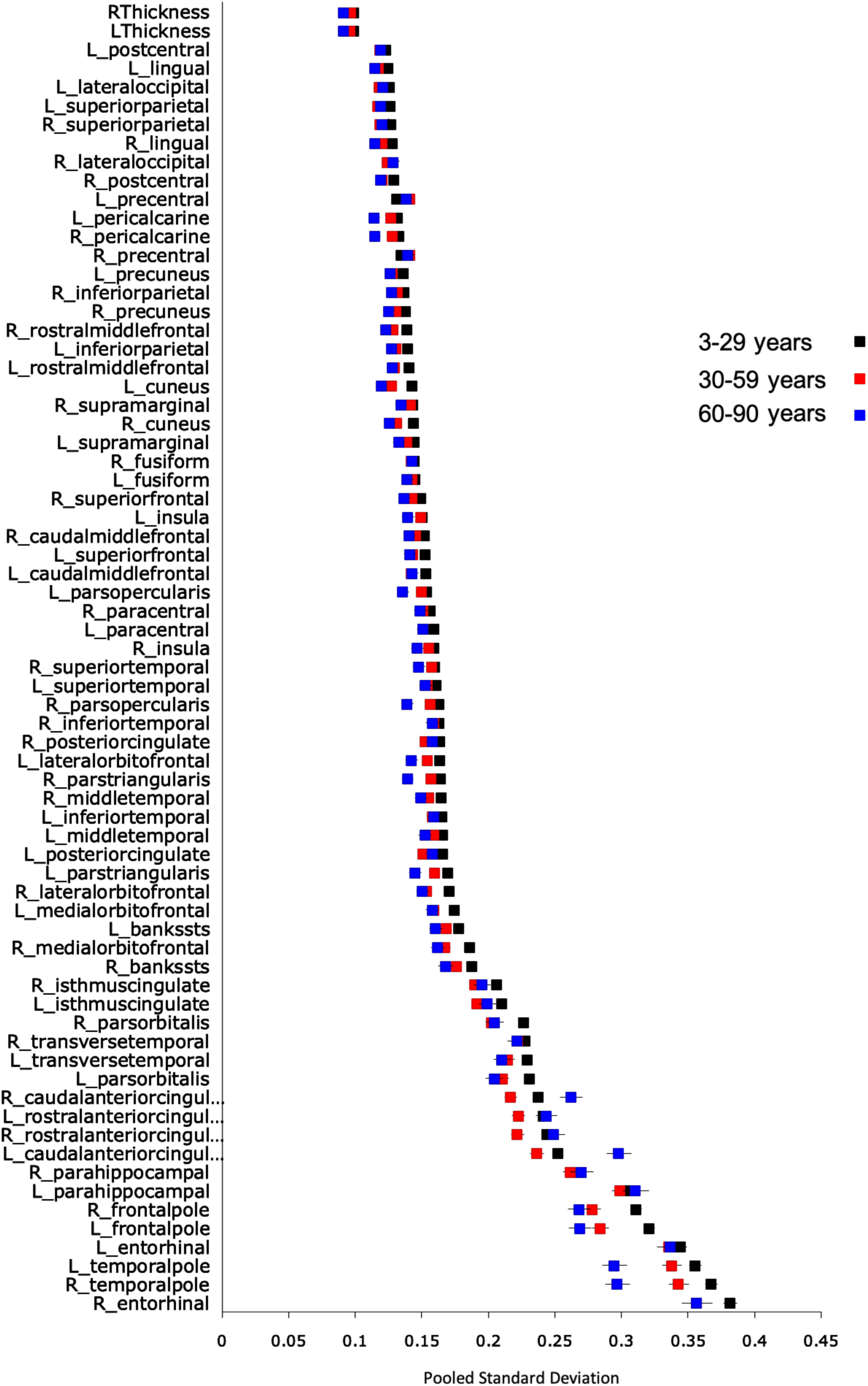
Meta-analysis of the pooled standard deviation in the entire dataset.

### Centile Curves of Cortical Thickness

Representative centiles curves for each lobe are presented in Figure 5. Centile values for the thickness of each cortical region, stratified by sex and hemisphere, are provided in Supplemental Tables S5-Table S7 and Supplemental File S2.

**Figure 5.**
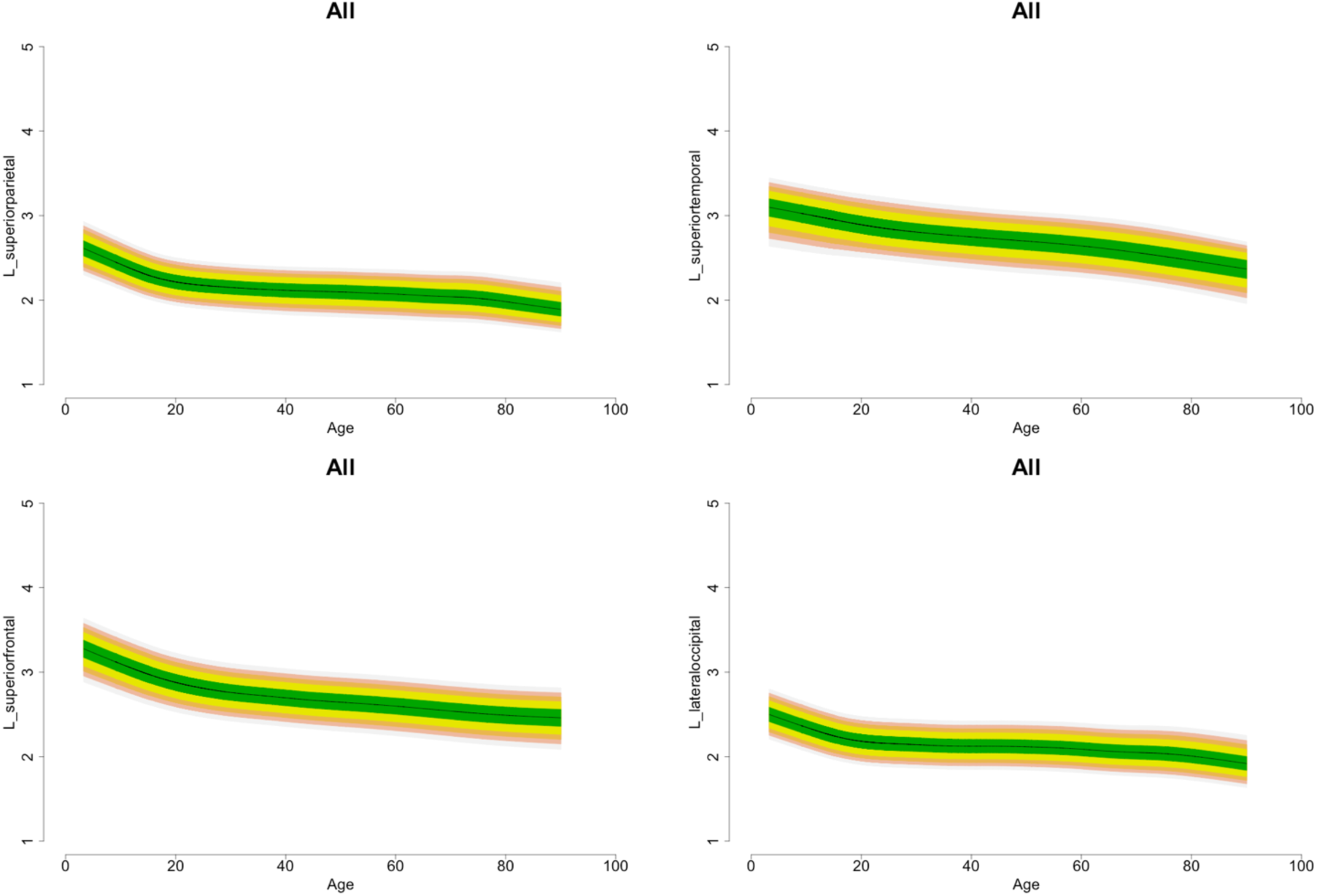
Illustrative Normative Centile Curves of Cortical Thickness. We present exemplar sets of centile curves for each lobe as derived from LMS of the entire dataset. Normative Centile Curves of thickness for all cortical regions (for the entire dataset and separately for males and females) are given in the supplementary material.

## Discussion

In the present study, we provide the most comprehensive characterisation of lifetime trajectories of regional cortical thickness based on multiple analytic methods (i.e., FP analysis, meta-analysis and centile calculations) and the largest dataset of cortical thickness measures available from healthy individuals aged 3 to 90 years. In addition to sample size, the study benefited from the standardised and validated protocols for data extraction and quality control that are common to all ENIGMA sites and have supported all published ENIGMA structural MRI studies (Hibar, et al., 2018; Schmaal, et al., 2017; Walton, et al., 2017; Whelan, et al., 2018).

As predicted, most regional cortical thickness measures reached their maximum value between 3-10 years of age, showed a steep decrease during the second and third decades of life and an attenuated or plateaued slope until later life. This pattern was independent of the hemisphere and sex. A recent review (Walhovd, et al., 2017) has highlighted contradictions between studies that report an increase in cortical thickness during early childhood and studies that report a decrease in cortical thickness during the same period. The results from our large-scale analysis help reconcile previous findings as we show that the median age at maximum thickness for most cortical regions is in the lower bound of the age range we examined here.

In the entorhinal and temporopolar regions, cortical thickness remained largely stable until 7^th^-8^th^ decades of life when it started to decline. Although the FreeSurfer estimation of cortical thickness in these regions is often considered suboptimal (compared to the rest of the brain), we note that our findings are consistent with a prior multicentre study of 1,660 healthy individuals (Hasan, et al., 2016). Further, the current study supports results from the National Institutes of Health MRI study of 384 individuals that found no significant change in the bilateral entorhinal and medial temporopolar cortex between the ages of 4-22 years (Ducharme, et al., 2016). A further study of 207 healthy adults aged 23-87 years also showed no significant cortical thinning in the entorhinal cortex until the 6^th^ decade of life (Storsve, et al., 2014). These observations suggest that the cortex of the entorhinal and temporopolar regions is largely preserved across the lifespan in healthy individuals. Both these regions contribute to episodic memory and the temporopolar region is also involved in semantic memory (Horel, et al., 1984; Nakamura, et al., 1994; Rolls, 2017). Degenerative changes of the temporopolar cortex have been reliably associated with semantic dementia, which is characterised by loss of conceptual knowledge about real-world items (Hodges and Patterson, 2007). The integrity and resting metabolic rate of the temporopolar cortex decrease with age (Allen, et al., 2005; Eberling, et al., 1995; Fjell, et al., 2009; Insausti, et al., 1998), and lower perfusion rates in this region correlate with cognitive impairment in patients with Alzheimer’s disease (AD) (Alegret, et al., 2010). Entorhinal cortical thickness is a reliable marker of episodic memory performance (Dickerson, et al., 2009; Fjell and Walhovd, 2010) and entorhinal cortex volume and metabolism are reduced in patients with Alzheimer’s Disease and mild cognitive impairment (Dickerson, et al., 2009; Hedden and Gabrieli, 2004). We therefore infer that “accelerated” entorhinal and temporopolar cortical thinning may be a marker of age-related cognitive decline; as they grow older, individuals at risk of cognitive decline may show a gradual shift in the distribution of the cortical thickness of these regions to the left which aligns with the exponential age-related increase in the incidence of AD in later decades of life (Mayeux and Stern, 2012).

The thickness of the ACC showed an attenuated U-shaped association with age. This observation replicates an earlier finding in 178 healthy individuals aged 7-87 years, which also found a U-shaped relationship between ACC thickness and age (Sowell, et al., 2007). The U-shaped age trajectory of ACC thickness might explain divergent findings in previous studies that have reported age-related increases (Abe, et al., 2008; Salat, et al., 2004), age-related reductions or no change (Brickman, et al., 2007; Ducharme, et al., 2016; Fjell and Walhovd, 2010; Good, et al., 2001; Vaidya, et al., 2007).

A consistently higher degree of inter-individual variation was observed in the most rostral frontal regions (frontopolar cortex and *pars orbitalis*), in the ACC and in several temporal regions (entorhinal, parahippocampal, temporopolar and transverse temporal cortex). To some degree, greater variability in several of these regions may reflect variability in measurement associated with their smaller size (Supplementary Figure S3). Nevertheless, the pattern observed suggests that greater inter-individual variability may be a feature of proisocortical and periallocortical regions (in the cingulate and temporal cortices) that are anatomically connected to prefrontal isocortical regions, and particularly the frontopolar cortex. This isocortical region of the prefrontal cortex is considered evolutionarily important based on its connectivity and function compared both to other human cortical regions and corresponding cortical regions in non-human primates (Ongur, et al., 2003; Semendeferi, et al., 2011). The frontopolar region has several microstructural characteristics, such as a higher number and greater width of minicolumns and greater inter-neuron space, which are conducive to facilitating neuronal connectivity (Semendeferi, et al., 2011). According to the popular ‘gateway’ hypothesis, the lateral frontopolar cortex implements processing of external information (‘stimulus-oriented’ processing) while the medial frontopolar cortex attends to self-generated or maintained representations (“stimulus-independent” processing) (Burgess, et al., 2007). Stimulus-oriented processing in the frontopolar cortex is focused on multitasking and goal-directed planning while stimulus-independent processing involves mainly mentalising and social cognition (Gilbert, et al., 2010). The other regions (entorhinal, parahippocampal, cingulate, and temporopolar) with high inter-individual variation in cortical thickness are periallocortical and proisocortical regions that are functionally connected to the medial frontopolar cortex (Gilbert, et al., 2010; Moayedi, et al., 2015). Notably, the periallocortex and proisocortex are considered transitional zones between the phylogenetically older allocortex and the more evolved isocortex (Galaburda and Sanides, 1980). Specifically, the entorhinal cortex is perialiocortical (Insausti, et al., 2017; Insausti, et al., 1995), the cingulate and parahippocampal cortices are proisocortical and the cortex of the temporopolar region is mixed (Blaizot, et al., 2010; Petrides and Pandya, 2012). Considered together, these regions are core nodes of the default mode network (DMN; Raichle et al., 2001). At present, it is unclear whether this higher inter-individual variation in the cortical thickness of the DMN nodes is associated with functional variation, but this is an important question for future studies.

The results presented here are based on the largest available brain MRI dataset worldwide covering the human lifespan. However, none of the pooled samples in the current study was longitudinal. We fully appreciate that longitudinal studies are considered preferable to cross-sectional designs when aiming to define age-related brain morphometric trajectories. However, a longitudinal study of this size over nine decades of life is not feasible. In addition to problems with participant recruitment and retention, such a lengthy study would have involved changes in scanner types, magnetic field strengths and acquisition protocols in line with necessary upgrades and technological advances. We took several steps to mitigate against site effects. First, we ensured that we used age-overlapping datasets throughout. Second, standardised analyses and quality control protocols were used to extract cortical thickness measures at all participating institutions. Third, we estimated and controlled for the contribution of site and scanner using ComBat prior to conducting our analysis. The validity of the findings reported here is reinforced by their alignment with the results from short-term longitudinal studies of cortical thickness (Shaw, et al., 2006b; Sowell, et al., 2004; Storsve, et al., 2014; Tamnes, et al., 2010; Thambisetty, et al., 2010; Wierenga, et al., 2014). The generalizability of our findings for the older age-group is qualified by our selection of individuals who appear to be ageing successfully in terms of cognitive function and absence of significant medical morbidity. Nevertheless, despite the efforts to include only healthy older individuals, the observed pattern of brain aging may still be influenced by subclinical mental or medical conditions. For example, vascular risk factors (e.g., hypertension) are prevalent in older individuals and have been associated with decline in the age-sensitive regions identified here (Raz et al., 2005). Thus we cannot conclusively exclude the possibility that such factors may have contributed to our results. Cellular studies show that the number of neurons, the extent of dendritic arborisation, and amount of glial support explain most of the variability in cortical thickness (Burgaleta, et al., 2014; la Fougere, et al., 2011; Rakic, 1988; Thompson, et al., 2007). MRI lacks the resolution to assess microstructural tissue properties but provides an estimate of cortical thickness based on the MR signal (Walhovd, et al., 2017). Nevertheless, there is remarkable similarity between MRI-derived thickness maps and post-mortem data (Sowell, et al., 2004; von Economo, 1929).

The findings of the current study suggest several avenues of further research. MRI-derived measures of cortical thickness do not provide information on the mechanisms that underlie the observed age-related trajectories. However, the centile values across the lifespan, provided here, could be used to study factors that may lead to deviations in cortical thickness way from the expected age-appropriate range. Such factors may be genetic, epigenetic, hormonal, socioeconomic or related to physical traits and health and lifestyle choices. Additionally, the results of the current study provide a new avenue for investigating the functional correlates, either cognitive or behavioral, of age-related changes and inter-individual variation in regional cortical thickness.

In summary, we performed a large-scale analysis using data from 17,075 individuals to investigate the lifespan trajectories of cortical thickness in healthy individuals. Our results may shed light on the uncertainties regarding age-related developmental trajectories for cortical thickness. Estimated centile values and inter-individual variability measures have the potential to provide scientists and clinicians with new tools to detect morphometric deviations and investigating associated behavioural and cognitive phenotypes.

## Acknowledgments

This study presents independent research funded by multiple agencies. The funding sources had no role in the study design, data collection, analysis, and interpretation of the data. The views expressed in the manuscript are those of the authors and do not necessarily represent those of any of the funding agencies. Dr. Dima received funding from the National Institute for Health Research (NIHR) Biomedical Research Centre at South London and Maudsley NHS Foundation Trust and King’s College London, the Psychiatry Research Trust and 2014 NARSAD Young Investigator Award. Dr. Frangou received support from the National Institutes of Health (R01 MH104284) the European Community’s Seventh Framework Programme (FP7/2007-2013) (grant agreement n°602450). FBIRN data collection and analysis was supported by the National Center for Research Resources at the National Institutes of Health (grant numbers: NIH 1 U24 RR021992 (Function Biomedical Informatics Research Network) and NIH 1 U24 RR025736-01 (Biomedical Informatics Research Network Coordinating Center; http://www.birncommunity.org). FBIRN data was processed by the UCI High Performance Computing cluster supported by the National Center for Research Resources and the National Center for Advancing Translational Sciences, National Institutes of Health, through Grant UL1 TR000153. Betula sample: Data collection was supported by a grant from Knut and Alice Wallenberg Foundation (KAW). Indiana sample: Brenna McDonald acknowledges the support in part by grants to BCM from Siemens Medical Solutions, from the members of the Partnership for Pediatric Epilepsy Research, which includes the American Epilepsy Society, the Epilepsy Foundation, the Epilepsy Therapy Project, Fight Against Childhood Epilepsy and Seizures (F.A.C.E.S.), and Parents Against Childhood Epilepsy (P.A.C.E.), from the Indiana State Department of Health Spinal Cord and Brain Injury Fund Research Grant Program, and by a Project Development Team within the ICTSI NIH/NCRR Grant Number RR025761. Andrew Saykin received support from U.S. National Institutes of Health grants R01 AG19771, P30 AG10133 and R01 CA101318. For the QTIM sample: We are grateful to the twins for their generosity of time and willingness to participate in our study. We also thank the many research assistants, radiographers, and other staff at QIMR Berghofer Medical Research Institute and the Centre for Advanced Imaging, University of Queensland. QTIM was funded by the Australian National Health and Medical Research Council (Project Grants No. 496682 and 1009064) and US National Institute of Child Health and Human Development (RO1HD050735). Lachlan Strike was supported by a University of Queensland PhD scholarship. The TOP study was supported by the European Community’s Seventh Framework Programme (FP7/2007-2013), grant agreement n°602450. The Southern and Eastern Norway Regional Health Authority supported Lars T. Westlye (grant no. 2014-097) and STROKEMRI (grant no. 2013-054). For the HUBIN sample: HUBIN was supported by the Swedish Research Council (K2007-62X-15077-04-1, K2008-62P-20597-01-3. K2010-62X-15078-07-2, K2012-61X-15078-09-3), the regional agreement on medical training and clinical research between Stockholm County Council, and the Karolinska Institutet, and the Knut and Alice Wallenberg Foundation. P.M.T., N.J., M.J.W., S.E.M., O.A.A., D.A.R., L.S., D.J.V., T.G.M. v.E., D.G., and D.P.H. were supported in part by a Consortium grant (U54 EB020403 to P.M.T.) from the NIH Institutes contributing to the Big Data to Knowledge (BD2K) Initiative.

## Data Availability Statement

The ENIGMA Lifespan Working Group welcomes expression of interest from researchers in the field who wish to use the ENIGMA samples. Data sharing is possible subsequent to consent for the principal investigators of the contributing datasets. Requests should be directed to the corresponding authors.

## Abbreviations of studies

ADHD-NF: Attention Deficit Hyperactivity Disorder-Neurofeedback Study;
AMC: Amsterdam Medisch Centrum;
Basel: University of Basel;
Barcelona: University of Barcelona;
Betula: Swedish longitudinal study on aging, memory, and dementia;
BIG: Brain Imaging Genetics;
BIL&GIN: a multimodal multidimensional database for investigating hemispheric specialization;
Bonn: University of Bonn;
BrainSCALE: Brain Structure and Cognition: an Adolescence Longitudinal twin study;
CAMH: Centre for Addiction and Mental Health;
Cardiff: Cardiff University;
CEG: Cognitive-experimental and Genetic study of ADHD and Control Sibling Pairs;
CIAM: Cortical Inhibition and Attentional Modulation study;
CLiNG: Clinical Neuroscience Göttingen;
CODE: formerly Cognitive Behavioral Analysis System of Psychotherapy (CBASP) study;
Edinburgh: The University of Edinburgh;
ENIGMA-HIV: Enhancing NeuroImaging Genetics through Meta-Analysis-Human Immunodeficiency Virus Working Group;
ENIGMA-OCD: Enhancing NeuroImaging Genetics through Meta-Analysis-Obsessive Compulsive Disorder Working Group;
FBIRN: Function Biomedical Informatics Research Network;
FIDMAG: Fundación para la Investigación y Docencia Maria Angustias Giménez;
GSP: Brain Genomics Superstruct Project;
HMS: Homburg Multidiagnosis Study;
HUBIN: Human Brain Informatics;
IDIVAL: Valdecilla Biomedical Research Institute;
IMAGEN: the IMAGEN Consortium;
IMH: Institute of Mental Health, Singapore;
IMpACT: The International Multicentre persistent ADHD Genetics Collaboration;
Indiana: Indiana University School of Medicine;
Johns Hopkins: Johns Hopkins University;
KaSP: The Karolinska Schizophrenia Project;
Leiden: Leiden University;
MAS: Memory and Ageing Study;
MCIC: MIND Clinical Imaging Consortium formed by the Mental Illness and Neuroscience Discovery (MIND) Institute now the Mind Research Network;
Melbourne: University of Melbourne;
Meth-CT: study of methamphetamine users, University of Cape Town;
MHRC: Mental Health Research Center;
Muenster: Muenster University;
NESDA: The Netherlands Study of Depression and Anxiety;
NeuroIMAGE: Dutch part of the International Multicenter ADHD Genetics (IMAGE) study; Neuroventure: the imaging part of the Co-Venture Trial funded by the Canadian Institutes of Health Research (CIHR);
NCNG: Norwegian Cognitive NeuroGenetics sample;
NTR: Netherlands Twin Register;
NU: Northwestern University;
NUIG: National University of Ireland Galway;
NYU: New York University;
OATS: Older Australian Twins Study;
Olin: Olin Neuropsychiatric Research Center;
Oxford: Oxford University;
QTIM: Queensland Twin Imaging;
Sao Paulo: University of Sao Paulo;
SCORE: University of Basel Study;
SHIP-2 and SHIP TREND: Study of Health in Pomerania;
Staged-Dep: Stages of Depression Study;
Stanford: Stanford University;
StrokeMRI: Stroke Magnetic Resonance Imaging;
Sydney: University of Sydney;
TOP: Tematisk Område Psykoser (Thematically Organized Psychosis Research);
TS-EUROTRAIN: European-Wide Investigation and Training Network on the Etiology and Pathophysiology of Gilles de la Tourette Syndrome;
Tuebingen: University of Tuebingen;
UMCU: Universitair Medisch Centrum Utrecht;
UNIBA: University of Bari Aldo Moro;
UPENN: University of Pennsylvania;
Yale: Yale University

