## Supplementary Figures and Tables for "Cortical Thickness Trajectories across the Lifespan: Data from 17,075 healthy individuals aged 3-90 years"

Online Supplement

**Supplementary Figures**

**Figure S1.** Correlation between age and cortical thickness across age-groups and stratified by sex

**Figure S2.** Meta-analysis of pooled standard deviation stratified by sex

**Figure S3.** Pooled Standard deviation of cortical regions as a function of surface area

**Supplementary Tables**

**Table S1.** Screening Process and Eligibility Criteria, Scanner, Image Acquisition Parameters and Image Segmentation Software

**Table S2.** Variance Explained by Age in fractional polynomial model

**Table S3:** Pearson's Correlation Coefficient between Age and Cortical Thickness

**Table S4.** Inter-individual variations in cortical thickness

**Table S5.** Centile Values for Cortical Thickness

**Table S6.** Centile Values for Cortical Thickness in Males

**Table S7.** Centile Values for Cortical Thickness in Females

**Figure S1.** Correlation between age and cortical thickness across age-groups and stratified by sex

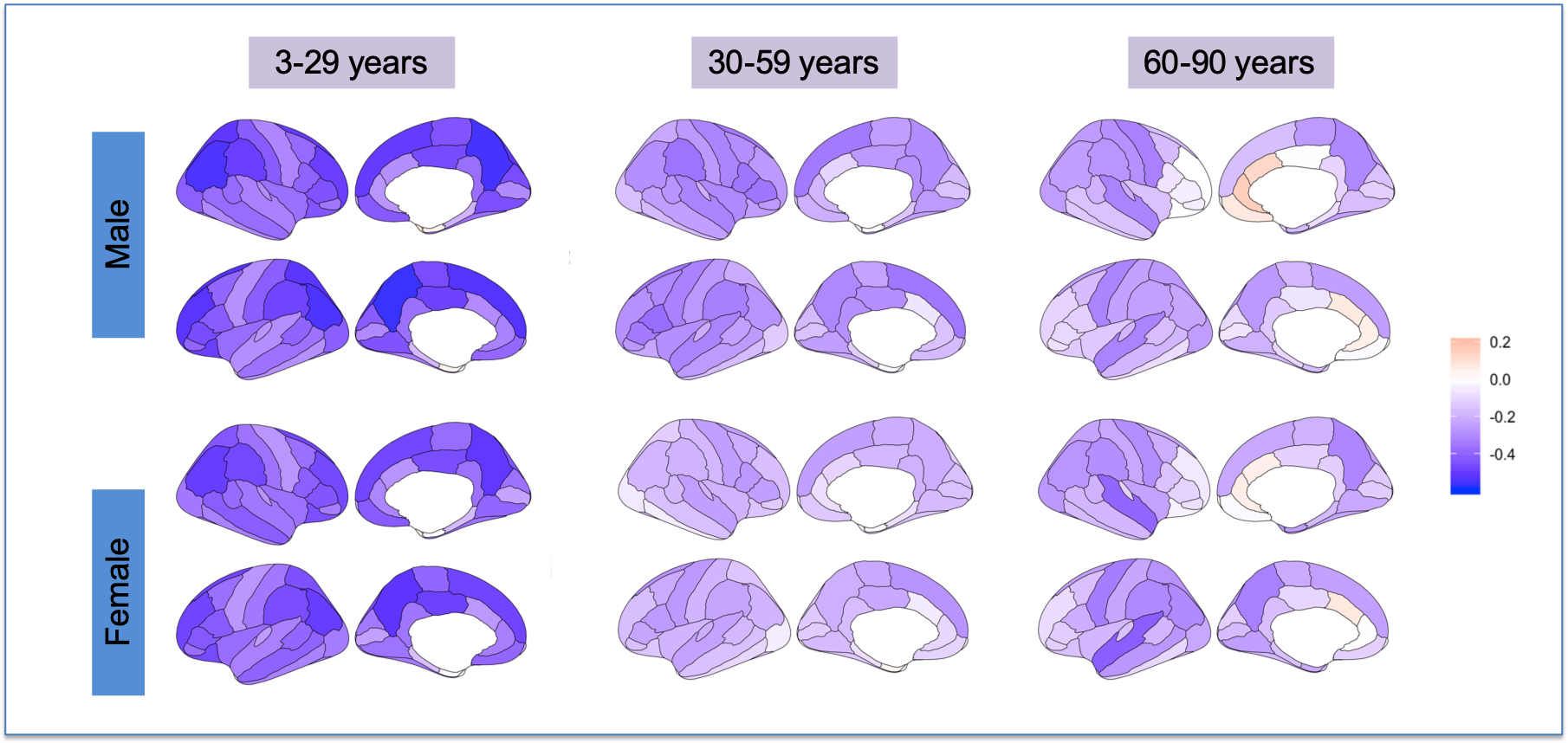

**Figure S2.** Meta-analysis of pooled standard deviation stratified by sex

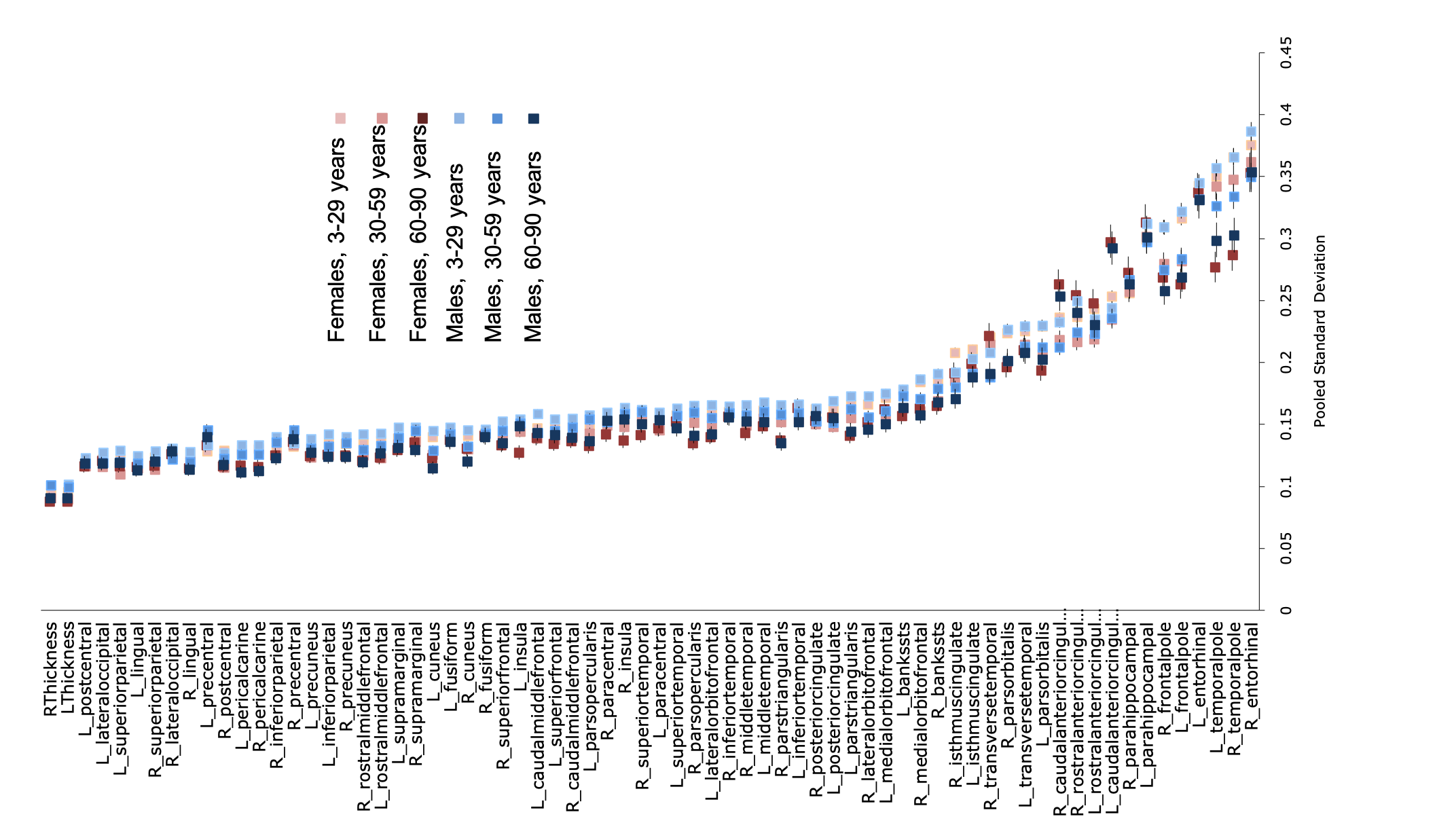

**Figure S3.** Pooled Standard deviation of cortical regions as a function of surface area

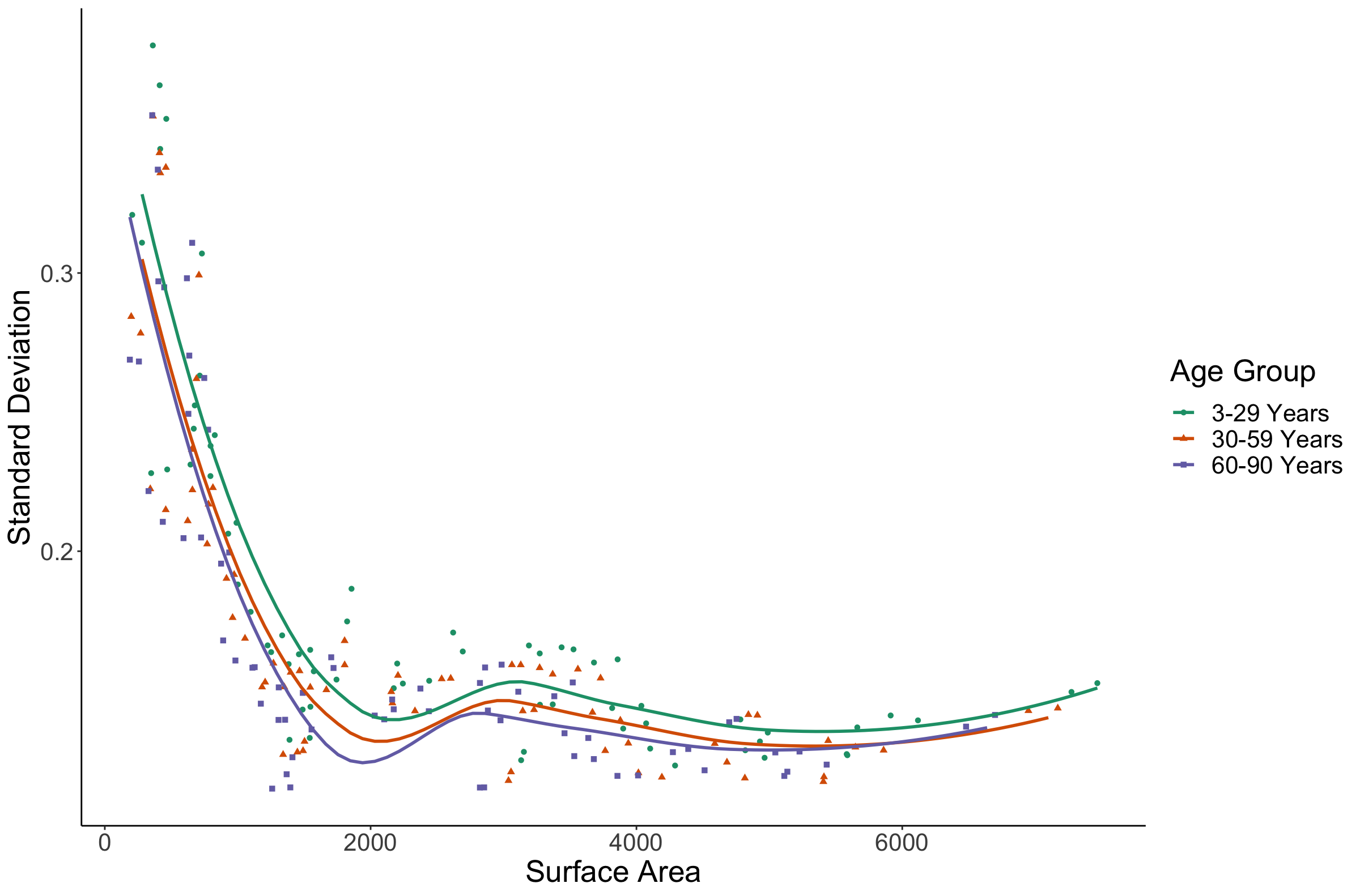

**Table S1.** Screening Process and Eligibility Criteria, Scanner, Image Acquisition Parameters and Image Segmentation Software

| **Sample** | **Screening Process** | **Eligibility Criteria** | **Magnet strength/ Scanner Vendor** | **Acquisition parameters** | **Freesurfer version** |
| --- | --- | --- | --- | --- | --- |
| **ADHD-NF** | KSADS | No head trauma, no neurological and psychiatric history, no lifetime alcohol or substance abuse, no previous or current use of psychotropic medication, IQ>75. | 3T Siemens Tim Trio | T1-weighted 3D MPRAGE; TR/TE/TI/FA=2300ms/3030ms/900ms/9^o^;  Image matrix= 256 × 256; 192 sagittal slices; voxel size=1mm^3^ | 5.3 |
| **AMC** | Personal Interview | No head trauma, no medical, neurological or psychiatric history, no lifetime alcohol or substance abuse, no previous or current use of psychotropic medication, IQ>75. No history of any psychiatric disorders in 1st degree family members. | 3T Philips Intera | T1-weighted 3D MPRAGE; TE range= 3.5-4.6ms, TR range= 9-9.663ms, FA= | 5.1 |
| **Barcelona 1.5T** | KSADS | No head trauma, no medical, neurological or psychiatric history, no lifetime alcohol or substance abuse, no previous or current use of psychotropic medication, IQ>75. No history of any psychiatric disorders in 1st degree family members. | 1.5 T General Electric Signa | T1-weighted; image matrix = 256 x 256; 128 slices; voxel size 1 x 1 x 1 mm^3^. | 5.3 |
| **Barcelona 3T** | KSADS | No head trauma, no medical, neurological or psychiatric history, no lifetime alcohol or substance abuse, no previous or current use of psychotropic medication, IQ>75. No history of any psychiatric disorders in 1st degree family members. | 3 T Siemens MAGNETOM TIM Trio | T1-weighted; image matrix = 256 x 256; 240 slices; voxel size 1 x 1 x 1 mm^3^. | 5.3 |
| **Betula** | Personal Interview | No head trauma, no medical, neurological or psychiatric history, no lifetime alcohol or substance abuse, no previous or current use of psychotropic medication. | 3T General Electric Discovery  MR750 | T1-weighted MPRAGE; TR/TE/TI/FA=8.1240 ms/3.2000 ms/450ms/ 12°; image matrix = 256x256 | 5.3 |
| **BIG** | Questionnaire for psychiatric history | No head trauma, no medical, neurological or psychiatric history, no lifetime alcohol or substance abuse, no previous or current use of psychotropic medication, IQ>70. No history of any psychiatric disorders in 1st or 2^nd^ degree family members. | 1,5 T Siemens Sonata and Avanto and 3 T Siemens Trio, TimTrio and Skyra | T1-weighted 3D MPRAGE; TR/TE/TI/sagittal slices = 1940-2730 ms/850-110 ms/2.92-4.58 ms; 176-192 sagittal slices; voxel size= 1.0x1.0x1.0 mm^3^ | 5.3 |
| **BIL&GIN** | Personal Interview | No head trauma, no current neurological or psychiatric disorders, no current use of psychotropic medication, IQ>70. | 3T Phillips ACHIEVA | T1 - weighted 3D; TR/TE/TI/FA=20 ms/4.6 ms/800ms/ 10°; turbo field echo factor = 65; sense factor = 2; matrix size = 256x256x180mm 3 ; voxel size= 1.0x1.0x1.0 mm^3^ | 5.3 |
| **Bonn** | Personal interview | No head trauma, no medical, neurological or psychiatric history, no previous or current use of psychotropic medication. | 3T Siemens Trio | TR/TE/FA= 1570-1660ms/2.75-3.42ms/8-9° | NA |
| **BrainSCALE** | Personal interview | No head trauma, no medical, neurological or psychiatric history, no lifetime alcohol or substance abuse, no previous or current use of psychotropic medication. | 1.5T Philips Achieva | T1-weighted 3D SPGR; TR/TE/ FA= 30 ms/4.6 ms/30°; image matrix=256x256; 160**–**180 contiguous coronal slices; voxel size=1 x 1 x 1.2 mm^3^ | 5.1 |
| **BRCATLAS** | Telephone interview | No head trauma, no medical, neurological or psychiatric history, no lifetime alcohol or substance abuse, no mild cognitive impairment, no previous or current use of psychotropic medication, IQ>75. | 3T GE Signa | T1-weighted 3D; TR/TE/TI/FA= 6.9 ms/2.8 ms/650 ms/8°; Image matrix = 256 x 256 x 180mm^3^; voxel size=1mm^3^ |  |
| **CAMH** | SCID | No head trauma, no neurological or psychiatric history, no alcohol or substance abuse preceding 6 months, no previous or current use of psychotropic medication, IQ>75. No history of any psychotic disorders in 1st degree family members. | 1.5 T GE (echospeed) | 124 Axial inversion recovery–prepared spoiled gradient recall images, 1.5-mm-thick slice acquisition TE/TR/TI/FA=5.3ms/12.3ms/300.0ms/20°. | 5.3 |
| **Cardiff** | MINI | No head trauma, no medical history, including neurological and psychiatric history, no alcohol or substance abuse in the preceding 6 months, no previous or current use of psychotropic medication. | 3T General Electric Signa | T1-weighted 3D FSPGR; TR/TE/TI/FA=7.9ms/3.0ms/450ms/20^o^;  Image matrix= 256 × 192 x 172; voxel size=1mm^3^ | 5.3 |
| **CODE (1-5)** | SCID | No head trauma, no medical, neurological or psychiatric history, no lifetime alcohol or substance abuse, no previous or current use of psychotropic medication, IQ>75. No history of any psychiatric disorders in 1st degree family members. | 3T Siemens Trio (4 CODE sites); 3T Philips Achieva (1 CODE site) | Siemens: T1 mprage, 1mm isotropic voxels, 12 channel head coil, TR=1900ms, TE=2.52ms, 170/192 slices.  Philips: T1 3D-TFE, 1mm isotropic voxels, 8 channel head coil, TR=8.3ms, TE=3.8ms, 170 slices | 5.3 |
| **CEG** | Teacher and parent  Conners' | No head trauma, no medical history, including neurological and psychiatric history, no alcohol or substance abuse in the preceding 6 months, no previous or current use of psychotropic medication. IQ>75 | 3T General Electric Signa | T1-weighted 3D SPGR; TR/TE/ FA=2000ms/30ms/90^o^;  Image matrix= 128 × 128; 43 slices | 5.3 |
| **CIAM** | SCID | No head trauma or psychiatric history, no previous or current use of psychotropic medication, IQ>75. | 3T Siemens Allegra | T1-weighted 3D MPRAGE; TR/TE/TI/FA= 2530 ms/1.53, 3.21, 4.89, 6.57/2.91 ms/ ms/7^o^; image matrix= 256x256; 128 sagittal slices; voxel size= 1.3x1.0x1.3 mm^3^ | 5.3 |
| **CLiNG** | Personal Interview | No head trauma, no medical, neurological or psychiatric history, no lifetime alcohol or substance abuse, no previous or current use of psychotropic medication, IQ>75. No history of any psychiatric disorders in 1st degree family members. | 3T Siemens Tim Trio | T1-weighted 3D MPRAGE; TR/TE/TI/FA=2250 ms/3.26 ms/900 ms/9°; image matrix = 256 x 256; 192 sagittal slices; voxel size= 1 mm^3^ | 5.3 |
| **COMPULS/TS EUROTRAIN** | KSADS | No head trauma, no medical, neurological or psychiatric history, no alcohol or substance abuse preceding 6 months, no previous or current use of psychotropic medication, IQ>75. No history of any psychiatric disorders in 1^st^ or 2^nd^ degree family members. | 3T Siemens Tim Trio and Prisma | T1-weighted 3D MPRAGE; TR/TE/FA=2300 ms/2.98 ms/9°; image matrix = 256 x 256; 176 sagittal slices; voxel size= 1x1x1.2 mm^3^ | 5.3 |
| **Edinburgh** | SCID-NP | No head trauma, no medical, neurological or psychiatric history, no previous or  current use of psychotropic medication, IQ>75. No history of any psychiatric  disorders in 1st and 2nd degree family members. | 1.5T Siemens Magnetom Essenza 1.5T General Electric Signa | T1-weighted MPRAGE; TR/TE/TI/FA =10 ms/4 ms/200 ms/8o; 128  contiguous coronal slices; voxel size=1.25 x 1.25 x 1.20 mm  T1-weighted MPRAGE; TR/TE/TI/FA =10 ms/4 ms/500 ms/8o; image matrix  = 192 x 192; 180 coronal slices; voxel size=1.25 x 1.25 x 1.2 mm3 | 5.3 |
| **ENIGMA-HIV** | MINI | No head trauma, no medical, neurological or psychiatric history, no alcohol or substance abuse preceding 6 months, no previous or current use of psychotropic medication, IQ>75. No mild cognitive impairment | 3T Siemens Allegra | T1-weighted MPRAGE; TR/TE/TI/FA=2400 ms/2.38 ms/1000 ms/ 8°; 162 slices; voxel size= 1 mm^3^ | 5.1 |
| **ENIGMA-OCD (3T OCD control)** | MINI-Plus | No head trauma, no medical, neurological or psychiatric history, no lifetime alcohol or substance abuse, no previous or current use of psychotropic medication, IQ>75. No cognitive impairment | 3T Siemens Allegra | T1-weighted 3D MPRAGE; TR/TE/TI/FA=2300 ms/3.93 ms/ 1100 ms/12^o^; image matrix =256×240; 160 contiguous sagittal slices; voxel size=1.3 x 1 x 1 mm^3^ | 5.3 |
| **ENIGMA-OCD (van den Heuvel 1.5T)** | SCID-I | No medical or psychiatric history | 1.5T Siemens Sonata | T1-weighted 3D MPRAGE; TR/TE/TI/FA=2700 ms/4 ms/ 950 ms/8^o^; image matrix =256×192; 160 slices; voxel size= 1 mm^3^ | 5.3 |
| **ENIGMA-OCD (van den Heuvel 3T)** | SCID-I | No medical or psychiatric history | 3T General Electric Signa | T1-weighted 3D MPRAGE; image matrix =256×256; 172 slices; voxel size= 1x0.977x0.977 mm^3^ | 5.3 |
| **ENIGMA-OCD (Huyser)** | Personal Interview | No head trauma, no medical, neurological or psychiatric history, no lifetime alcohol or substance abuse, no previous or current use of psychotropic medication, IQ>75. | 3T Phillips Intera | T1-weighted 3D MPRAGE; TR/TE/ FA=9.69 ms/4.60 ms/8^o^; image matrix =256×256; 182 slices; voxel size=1 x 1 x 1.2 mm^3^ | 5.3 |
| **ENIGMA-OCD (Mataix-Cols)** | SCID | No head trauma, no neurological or psychiatric history, no lifetime alcohol or substance abuse, no previous or current use of psychotropic medication, IQ>75. | 1.5T General Electric Signa | T1-weighted 3D SPGR; TR/TE/FA= 14.8 ms/ 1.7 ms/ 20º; image matrix=  256 x 256 x 124; voxel size: 0.94 x 0.94 x 1.50 mm | 5.3 |
| **ENIGMA-OCD (Nakao)** | Personal Interview | No head trauma, no neurological or psychiatric history, no lifetime alcohol or substance abuse. | 3T Phillips Achieva | T1-weighted 3D TFE; TR/TE/TI/FA=8.2 ms/3.8ms/1026 ms/8^o^; image matrix =240×240; 190 slices; voxel size=1 mm^3^ | 5.3 |
| **ENIGMA-OCD (IDIBELL)** | SCID-I/NP | No head trauma, no medical, neurological or psychiatric history, no alcohol or substance abuse in the preceding 6 months, no previous or current use of psychotropic medication, IQ>70. No history of any psychiatric disorders in 1st or 2^nd^ degree family members | 1.5 T General Electric Signa | T1-weighted 3D FSPGR; TR/TE/ FA=11.8ms/4.2ms/90^o^;  Image matrix= 256 × 256 x 130; voxel size=1.2mm^3^ | 5.3 |
| **FBIRN** | SCID-I/NP | No head trauma, no medical, neurological or psychiatric history, no alcohol or substance abuse in the preceding 5 years, no previous or current use of psychotropic medication, IQ>75. No history of any Axis-I psychotic disorders in 1st degree family members. | 3T Siemens Tim ^Trio^ or General Electric Discovery MR750 | T1-weighted SPGR; TR/TE/TI/FA=2300 ms/2.94 ms/1100 ms/9^o^; image matrix=256×256x160; voxel size=0.86x0.86x1.2mm^3^; sagittal plane acquisition | 5.1 |
| **FIDMAG** | Personal interview; structured interview in part of the sample | No head trauma, no medical, neurological or psychiatric history, no lifetime alcohol or substance abuse, no previous or current use of psychotropic medication, IQ>70. | 1.5 T General Electric Signa | T1-weighted MPRAGE; TR/TE/FA=2000 ms/4 ms/ 9^o^; image matrix=512 x 512; 180 contiguous sagittal slices; voxel size=0.56 x 0.56 x 1 mm3 | 5.3 |
| **GSP** | Structured phone screen and study specific self-report battery and clinical screen | No head trauma, no medical, neurological or psychiatric history, no lifetime alcohol or substance abuse, no current use of psychotropic medication, normal brain anatomy following brain scan. | 3T Siemens Tim Trio | T1-weighted 3D multi-echo MPRAGE; TR/TE/TI/FA =2200 ms/1.54-7 ms/  1100/7 ^o^; voxel size=1.2x1.2x1.2 mm | 4.5 |
| **HUBIN** | SCID-I | No head trauma, no medical, neurological or psychiatric history, no lifetime alcohol or substance abuse, no previous or current use of psychotropic medication, IQ>75. No history of any psychiatric disorders in 1st degree family members. | 1.5 T General Electric Signa | T1-weighted SPGR; TR/TE/FA= 24 ms/6 ms/35 ^o^; 124 coronal slices; voxel size 0.86 x 0.86 x 1.50 mm3. | 5.3 |
| **HMS** | Personal Interview | No head trauma, no medical, neurological or psychiatric history, no lifetime alcohol or substance abuse, no previous or current use of psychotropic medication, IQ>75. No history of any psychiatric disorders in 1st degree family members. | 1.5T Siemens Magnetom Sonata | T1-weighted 3D MPRAGE; TR/TE/TI/FA=1900 ms/4.0 ms/700 ms/15°; image matrix = 256 x 256; 176 consecutive sagittal slices; voxel size=1 mm^3^ | 5.3 |
| **IDIVAL (1) + (2)** | CASH | No head trauma, no medical, neurological or psychiatric history, no lifetime alcohol or substance abuse, no previous or current use of psychotropic medication, IQ>75. No history of any psychiatric disorders in 1st degree family members. | 3T Siemens Alegra, Phillips Achieva  1.5T General Electric Signa | T1-weighted SPGR; TR/TE/FA=24 ms/5 ms/ 5^o^; image matrix=256x192; T1-weighted SPGR; TR/TE/FA=3000 ms/3.9 ms/8^o^; image matrix=256x256;  voxel size=1mm^3^; sagittal plane acquisition | 5.3 |
| **IDIVAL (3)** | Personal Interview | No lifetime history of Axis I psychiatric disorders, no mild cognitive | 3T Phillips Achieva | T1-weighted SPGR; TR/TE/FA=3000 ms/4.6 ms/8^o^; image matrix=321x312; voxel size=1mm^3^; sagittal plane acquisition | 5.3 |
| **IMAGEN** | DAWBA questionnaire clinician interview | No head trauma, no medical, neurological or psychiatric history, no previous or current use of psychotropic medication. IQ>75 | 3T Siemens Verio and TimTrio, Philips Achieva, General Electric Signa Excite, and Signa HDx | T1-weighted 3D MPRAGE; TR/TE/TI/FA=2300ms/3030ms/900ms/9^o^;  Image matrix= 256 × 256;192 sagittal slices; voxel size=1mm^3^ | 5.3 |
| **IMH** | SCID-I/NP | No head trauma, no medical history, neurological or psychiatric history, no lifetime alcohol or substance abuse, as well as no previous or current use of psychotropic medication, IQ>75. No cognitive impairment | 3T Phillips Achieva | T1-weighted 3D MPRAGE; TR/TE/FA= 7.2ms/ 3.8ms/8^o^; image matrix=256 x 256; 180 axial slices; voxel size=0.9mm^3^ | 5.3 |
| **IMpACT** | SCID-I (and SCID-II) | No head trauma, no medical, neurological or psychiatric history, no alcohol or substance abuse in the preceding 6 months, no previous or current use of psychotropic medication, IQ>70. No history of any psychiatric disorders in 1st or 2^nd^ degree family members. | 1.5 T Siemens | T1-weighted 3D-MPRAGE; TR/TE/TI/FA =2730 ms/2.95 ms/1000 ms; 176 consecutive sagittal slices; voxel size= 1 mm^3^ | 3.5 |
| **Indiana 1.5T** | Personal Interview | No head trauma, no medical, neurological or psychiatric history, no lifetime alcohol or substance abuse, no previous or current use of psychotropic medication, IQ>75. | 1.5T General Electric Signa Horizon LX | T1-weighted 3D SPGR; TR/TE/FA=25 ms/3 ms/ 45^o^; image matrix= 256 x  256; 124 contiguous coronal slices | 5.1 |
| **Indiana 3T** | Personal interview  Structured phone screen | No head trauma, no medical, neurological or psychiatric history, no alcohol or substance abuse in the preceding 6 months, no previous or current use of psychotropic medication, IQ>75. | 3T Siemens Skyra | T1-weighted MPRAGE; TR/TE/FA=2300 ms/2.95 ms/ 9^o^; image matrix=256  x 240; 176 contiguous sagittal slices | 5.1 |
| **Johns Hopkins** | Personal Interview | No head trauma, no medical, neurological or psychiatric history, no alcohol or substance abuse preceding 6 months, never prescribed with psychotropic medication, IQ>75. No mild cognitive impairment | 1.5T General Electric Signa | T1-weighted SPGR; TR/TE/FA=35 ms/5 ms/ 45^o^; image matrix=256x256; 124 slices | 5.3 |
| **KaSP** | MINI | No head trauma, no medical, neurological or psychiatric history, no lifetime alcohol or substance abuse, no previous or current use of psychotropic medication, IQ>75. No history of any psychiatric disorders in 1^st^ or 2^nd^ degree family members. | 3T General Electric | T1-weighted SPGR; TR/TI/FA=7.904 ms/450 ms/12 ^o^; image matrix= 256 x 256 mm^3^; 145 sagittal slices ; voxel size=0.934 x 0.934 x 1.2 mm3 | 5.3 |
| **Leiden** | Self-report | No psychiatric or neurological disorders, no use of psychotropic medications | 3T Philips Achieva | T1-weighted 3D SPGR; TR/TE = 9.76 ms/4.59 ms; image matrix=256x256; 160**–**180 contiguous coronal slices; voxel size=0.875x 0.875 x 1.2 mm^3^ | 5.3 |
| **MAS** | Personal interview | No head trauma, no diagnosis of dementia, schizophrenia, bipolar disorder no psychotic symptoms, no neurological disorder, no mild cognitive impairment, IQ>75. | 3T Philips Achieva Quasar Dual | TR/TE = 6.39 ms/2.9 ms; 190 coronal slices; voxel size = 1mm^3^ | 5.3 |
| **MCIC** | SCID, SCID-I/NP, CASH | No head trauma, no medical, neurological or psychiatric history, no lifetime alcohol or substance abuse, no previous or current use of psychotropic medication, IQ>75. | 1.5T Siemens Sonata-3T Siemens Trio | T1-weighted MPRAGE sequence; TR/TE/TI/FA=2530 ms/4.76 ms/1100  ms/20^o^; image matrix=256×256×128 cm; voxel size=0.625 mm^3^ | 5.3 |
| **Melbourne** | SCID-I | No head trauma, no neurological or psychiatric history, no lifetime alcohol or substance abuse, no previous or current use of psychotropic medication. No history of any psychiatric disorders in 1^st^ or 2^nd^ degree family members. | 3T GE Signa Excite | 3D BRAVO sequence 140; TR/TE/FA=7900 ms/3000 ms/13º; FOV=256 mm; matrix=256 x 256 | 5.3 |
| **Meth-CT** | SCID DSM-IV | No head trauma, no medical, neurological or psychiatric history, no lifetime alcohol or substance abuse, no previous or current use of psychotropic medication, IQ>70. | 3T Siemens Allegra | T1-weighted 3D MPRAGE; TR/graded TE/FA=2530 ms/ 1.53, 3.21, 4.89, 6.57 ms/ 7^o^; 160 contiguous sagittal slices; voxel size=1 x 1 *x 1 mm3 | 5.3 |
| **MHRC** | Personal interview | No head trauma, no medical history, including neurological and psychiatric history. No Family History of neurological or psychiatric disorders | 3Т Philips Achieva | T1-weighted TFE; TR/TE/ FA=8.2ms/3.7ms/8^o;^; voxel size=0.83 x 0.83 x 1 mm^3^ | 5.3 |
| **Muenster** | SCID | No head trauma, no medical, neurological or psychiatric history, no lifetime alcohol or substance abuse, no previous or current use of psychotropic medication, IQ>75. | 3T Phillips Intera | T1 weighted TFE: TR/TE/FA= 7.4 ms/3.4 ms/9°; image matrix = 256x204x160mm^3^; voxel size=0.5mm^3^; sagittal plane acquisition | 5.3 |
| **NCNG** | Personal interview | No head trauma, no medical, neurological or psychiatric history, no lifetime alcohol or substance abuse, no mild cognitive impairment, no previous or current use of  psychotropic medication, IQ>84. | 1.5T Siemens Avanto  1.5T Siemens Sonata | T1-weighted 3D MPRAGE; TR/TE/TI/FA= 2400 ms/3.61 ms/1000 ms/8°; image matrix=192x192; 160 sagittal slices; voxel size=1.25 mm^3^  T1-weighted 3D MPRAGE; TR/TE/TI/FA= 2730 ms/3.43 ms/1000 ms/7°; image matrix=256x256; 128 sagittal slices; voxel size=1 mm^3^ | 4.5 |
| **NESDA** | CIDI | No lifetime history of Axis-I diagnoses, no lifetime medical or neurological morbidity including hypertension, no lifetime substance dependence, no substance abuse in the preceding year, no medication use. | 3T Philips Achieva  SENSE-6 to 8 channel head coil | T1-weighted 3D MPRAGE; TR/TE/FA= 9 ms/3.5 ms/8^o^; image matrix=256x256; 170 sagittal slices; voxel size=1mm^3^ | 5.0 |
| **NeuroIMAGE** | KSADS-PL | No head trauma, no mild cognitive impairment, neurological or psychiatric history, no previous or current use of psychotropic medication, IQ>75. No history of any psychiatric disorders in 1st and 2nd degree family members. | 1.5 T Siemens AVANTO (Donders Centre for Cognitive Neuroimaging)  1.5 T Siemens SONATA (VU University Amsterdam) | MPRAGE 176 sagittal slices, repetition time=2,730ms, echo time=2.95ms, voxel size=1.0x1.0x1.0mm, field of view=256 mm | 5.3 |
| **Neuroventure** | DAWBA and BSI | No head trauma, no medical, neurological or psychiatric history, no lifetime alcohol or substance abuse, no previous or current use of psychotropic medication, IQ>75. | 3T SIEMENS TrioTim | T1-weighted 3D MPRAGE; TR/TE/ FA= 2300 ms/2.96 ms/9^o^; image matrix= 256x256; voxel size= 1.0x1.0x1.0 mm^3^ | 5.3 |
| **NTR (1)** | DISC-IV | No head trauma, no medical, neurological or psychiatric history, no lifetime alcohol or substance abuse, no mild cognitive impairment, no previous or current use of psychotropic medication, IQ>75. | 1.5T Siemens Sonata | T1-weighted 3D MPRAGE; TR/TE/TI/FA=1900 ms/3.93 ms/1100 ms/ 15°; image matrix=256 x 224; 160 sagittal slices; voxel size=1 mm^3^ | 5.1 |
| **NTR (2)** | MINI, BDI, STAI, STAS, YBOCS | No head trauma, no previous or current use of psychotropic medication, normal IQ. | 3T Philips Intera | T1-weighted 3D MPRAGE; TR/TE/FA=9.64 ms/4.60 ms/8 ^o^; image matrix=256 x 256; 182 coronal slices; voxel size=1 x1x1.2 mm^3^ | 5.1 |
| **NTR (3)** | CIDI, MADRS, BDI, STAI | No current psychiatric disorder, no current use of psychotropic medication, normal IQ. | 1.5 T Siemens Sonata | T1-weighted 3D MPRAGE; TR/TE/TI/FA= 15 ms/7 ms/300 ms/8°; image matrix=256x176; 160 coronal slices; voxel size=1x1x1.5 mm^3^ | 5.1 |
| **NU** | SCID | No head trauma, no medical, neurological or psychiatric history, no lifetime alcohol or substance abuse, no previous or current use of psychotropic medication, IQ>75. No history of any psychiatric disorders in 1st degree family members. | 1.5T SIEMENS Vision | T1-weighted 3D MPRAGE; TR/TE/TI/FA=2200 ms/4.13 ms/766 ms/13°; voxel size =0.8mm^3^; axial plane acquisition. | 5.3 |
| **NUIG** | SCID | No head trauma, no neurological or psychiatric history, no alcohol or substance abuse preceding 6 months, no previous or current use of psychotropic medication, IQ>75. No history of any psychiatric disorders in 1st degree family members. | Siemens Magnetom Symphony 1.5T | 3D, T1-weighted MPRAGE 4 channel head coil, FOV 230mm, TR/TE/: 1140ms/4.38ms, matrix size 256 x 256, interpolated to 512 x 512, yielding an in-plane voxel size of 0.45mm x 0.45mm^2^, slice thickness 0.9mm. | 5.1 |
| **NYU** | SCID-NP for DSM-IV | No head trauma, no medical history, including neurological and psychiatric history, no lifetime alcohol or substance abuse, no previous or current use of psychotropic medication. IQ>75. | 3T Siemens Allegra | T1-weighted 3D MPRAGE; TR/TE/TI/FA=2530ms/3.25ms/1100ms/7^o^ | 5.3 |
| **OATS (1-4)** | Personal interview | No head trauma, no current diagnosis of a psychotic disorder, no neurological disorder, no malignancy (other than skin cancer) or other severe medical comorbidity, no mild cognitive impairment, IQ>75. | 1.5T Philips Gyroscan, Siemens Magnetom Avanto, Siemens Sonata; 3T Philips Achieva Quasar Dual, a | T1-weighted 3D acquisition; TR/TE/TI/FA=15370 ms/3.24 ms/780 ms/8^o^; 144 slices; voxel size=1 x 1 x 1.5 mm^3^ | 5.3 |
| **OLIN** | SCID I | No head trauma, no medical, neurological or psychiatric history, no alcohol or substance abuse preceding 6 months, never prescribed with psychotropic medication, IQ>75. | 3T Siemens Alegra | T1-weighted 3D MPRAGE; TR/TE/TI/FA= 2300 ms/2.91 ms/900 ms/9^o^; image matrix= 256x240x192; 160 sagittal slices; voxel size= 1.0x1.0x1.2 mm^3^ | 5.1 |
| **PING** | Personal interview | No lifetime history of major developmental, psychiatric, or neurological disorders, brain injury, or other medical conditions that affect development. Individuals born  earlier than 36 weeks of gestational age were excluded. | 3T Philips Achieva  3T GE SIGNA  3T Siemens TrioTim  3T Siemens TrioTim  3T General Electric Discovery  MR750 | T1-weighted 3D IR-GRE; TR/TE/TI/FA= 8.1 ms/3.5 ms/640 ms/9^o^ | 5.3 |
| **QTIM** | CIDI | No head trauma, no medical history, neurological and psychiatric history, no alcohol or substance abuse in the preceding 6 months, no antidepressant medication or medication affecting cognition. | 4T Bruckner | T1-weighted 3D MPRAGE: TR/TE/TI/FA = 1500 ms/3.35 ms/ 700 ms/ 8°; image matrix= 256 × 256 × 256 or 256 × 256 × 240; 256 coronal slices; voxel size= 0.9 mm^3^ | 5.1 |
| **Oxford** | KSADS | No head trauma, no medical, neurological or psychiatric history, no lifetime alcohol or substance abuse, no previous or current use of psychotropic medication, IQ>75. | 1.5T Siemens Sonata | T1-weighted 3D MPRAGE; TR/TE =12 ms/5.6 ms; image matrix =256×240x 208 mm^3^; voxel size=1 mm^3^ | 5.3 |
| **Sao Paulo (1)** | SCID | No head trauma, neurological or psychiatric history, no lifetime alcohol or substance abuse. | 1.5T Siemens Espree | T1-weighted 3D MPRAGE; TR/TE/TI/FA=2400 ms/3.65 ms/ 0 ms/8^o^; 160 contiguous sagittal slices; voxel size=1.3 x 1.3x 1.2 mm3 | 5.3 |
| **Sao Paulo (3)** | SCID | No head trauma, neurological or psychiatric history, no lifetime alcohol or substance abuse. IQ>75 | 1.5T General Electric Signa | T1-weighted FSPGR  ; TR/TE/TI/FA=21.7 ms/52 ms /20^o^; 124 axial slices; voxel size= 0.86 x 0.86 x 1.5 mm3 | 5.3 |
| **SCORE** | BPRS | No head trauma, no medical, neurological or psychiatric history, no lifetime history of alcohol or substance abuse, no previous or current use of psychotropic medication, IQ>75. No history of any psychiatric disorders in 1st degree family members. | 3T Siemens Magnetom Verio | T1-weighted 3D-MPRAGE; TR/TE/TI/FA =2000 ms/3.37 ms/1000 ms/8^o^; image matrix=256x256x176; 176 consecutive sagittal slices; voxel size= 1 mm^3^ | 6.0 |
| **SHIP-2** | Personal Interview | No head trauma, no neurological and psychiatric history, no alcohol or substance abuse in the preceding 6 months, no previous or current use of psychotropic medication. IQ>75. | 1.5T Siemens Avanto | T1-weighted 3D MPRAGE; TR/TE/ FA=1900ms/3.4ms/15^o^; voxel size=1mm^3^ | 5.3 |
| **SHIP-TREND** | Personal Interview | No head trauma, no neurological and psychiatric history, no alcohol or substance abuse in the preceding 6 months, no previous or current use of psychotropic medication. IQ>75. | 1.5T Siemens Avanto | T1-weighted 3D MPRAGE; TR/TE/ FA=1900ms/3.4ms/15^o^; voxel size=1 mm^3^ | 5.3 |
| **Stages-Dep** | SCID-I | No head trauma, no medical, neurological or psychiatric history, no lifetime alcohol or substance abuse, no previous or current use of psychotropic medication, IQ>75. No history of any psychiatric disorders in 1st degree family members. | 3T Phillips Achieva | T1-weighted 3D-MPRAGE; TR/TE/TI/FA =6.7 ms/3.2 ms/200 ms/88^o^; °; image matrix = 288 x 288; 170 consecutive sagittal slices; voxel size= 0.896×0.896×1.2 mm^3^ | 5.1 |
| **Stanford** | SCID | No head trauma, no medical, neurological or psychiatric history, no lifetime alcohol or substance abuse, no previous or current use of psychotropic medication. no mild cognitive impairment. | 1.5T General Electric Signa Excite | T1-weighted SPGR; TR/TE/TI/FA=8.3-10.3 ms/1.7-3.0 ms/300 ms/15^o^; image matrix= 256 x 192; 176 contiguous sagittal slices; voxel size=0.86x0.86x1.5 mm^3^; sagittal plan acquisition | 5.3 |
| **StrokeMRI** | Personal interview | No head trauma, no medical, neurological or psychiatric history, no lifetime alcohol or substance abuse, no previous or current use of psychotropic medication, IQ>75. | 3T General Electric Signa HDxt | T1-weighted FSPGR; TR/TE/TI/FA=7.8 s/2.956 ms/450 ms/12°; 170 slices; voxel size= 1.0x1.0x1.2 mm | 5.3 |
| **Sydney** | SCID | No head trauma, no medical history, neurological or psychiatric history, no alcohol or substance abuse preceding 6 months, as well as no previous or current use of psychotropic medication, IQ>75. | 3T General Electric Discovery MR750 | T1-weighted 3D MPRAGE; TR/TE/FA= 7264ms/ 2784ms/15^o^; image matrix  =256 x 256 x 196; voxel size=0.9mm^3^ | 5.1 |
| **TOP** | PRIME-MD | No head trauma, no organic or other psychotic disorder (ICD codes 290-299), no substance abuse in the preceding 6 months, no previous or current use of psychotropic medication, IQ>75. No history of any psychiatric disorders in 1st degree family members. | 1.5T Siemens Magnetom Sonata | T1-weighted SPGR; TR/TE/TI/FA=2730 ms/3.93 ms/1000 ms/71^o^; voxel size = 1.33x0.94x1mm^3^; sagittal plane acquisition | 5.3 |
| **Tuebingen** | SCID I and II | No head trauma, no medical history, neurological or psychiatric history, no lifetime alcohol or substance abuse as well as no previous or current use of psychotropic medication, IQ>75. No history of any psychiatric disorders in 1st degree family members. | 1.5T Siemens Avanto | T1-weighted 3D MPRAGE; TR/TE/FA= 2250ms/ 3.93ms/8^o^; image matrix  =256 x 256; voxel size=1mm^3^ | 5.3 |
| **UMCU** | CASH | No head trauma, no medical, neurological or psychiatric history, no lifetime alcohol or substance abuse, no previous or current use of psychotropic medication, IQ>75. No history of any psychiatric disorders in 1st degree family members. | 1.5T Philips Intera and Achieva | T1-weighted 3D FFE; TE/TR/FA= 4.6 ms/0 ms**/ 0˚;** 160-180 contiguous coronal slices; voxel size=1x1x1.2 mm^3^ | 5.1 |
| **UNIBA** | SCID-NP | No head trauma, no medical, neurological or psychiatric history, no lifetime alcohol or substance abuse, no previous or current use of psychotropic medication, IQ>75. No history of any psychiatric disorders in 1st degree family members. | 3T General Electric | T1-weighted 3D SPGR; TE/FA = min full/ 6°; image matrix= 256×256 x124 | 5.3 |
| **UPENN** | SCID | No head trauma, no medical history, including neurological and psychiatric history, no alcohol or substance abuse preceding 6 months, no previous or current use of psychotropic medication, IQ>75. No history of any psychiatric disorders in 1st degree family members. | 3T Siemens Tim Trio | T1-weighted 3D MPRAGE; TR/TE/TI/FA=1810 ms/3.51 ms/1100 ms/9^o^; image matrix= 256 × 192;160 axial slices | 5.3 |
| **Yale** | KSADS-PL | No head trauma, neurological or psychiatric history, no alcohol or substance abuse in the preceding 6 months, no previous or current use of psychotropic medication, IQ>75. | 3T General Electric Signa | T1-weighted 3D MPRAGE; image matrix =256×256; voxel size=0.976 x 0.976 x 1 mm3 | 5.3 |

Abbreviations of Terms: BDI = Behavioural Descriptive Interview; BSI = Brief Symptom Inventory; CASH = Comprehensive assessment of symptoms and history; CDR = Clinical Dementia Rating; CIDI = Composite International Diagnostic Interview; DAWBA = Development and Well-Being Assessment; DISC-IV = Diagnostic Interview Schedule for Children; DSM = Diagnostic and Statistical Manual of Mental Disorders (DSM); FA=flip angle; FSPGR=fast spoiled gradient echo sequence; GRE=spoiled gradient echo sequence; ICD= International Classification of Diseases; IR= inversion recovery; KSADS-PL= Kiddie Schedule for Affective Disorders and Schizophrenia-Present and Lifetime; MADRS = Montgomery-Asberg Depression Rating Scale; MINI = Mini International Neuropsychiatric Interview; MMSE = Mini Mental State Exam; PRIME-MD = Primary Care Evaluation of Mental Disorder; SCID = Structured Clinical Interview for DSM Disorders; SCID-I/NP = SCID Non-Patient version; SPGR=spoiled gradient recalled sequence; STAI = State-Trait Anxiety Inventory; STAS = State-trait anger scale; TE=echo time; TI=inversion time; TR=repetition time; TFE=turbo field echo sequence; YBOCS = Yale-Brown Obsessive Compulsive Scale

Abbreviations of studies: **ADHD-NF = Attention Deficit Hyperactivity Disorder- Neurofeedback Study; AMC = Amsterdam Medisch Centrum;** **Basel = University of Basel**; **Barcelona = University of Barcelona;** **Betula = Swedish longitudinal study on aging, memory, and dementia**; **BIG = Brain Imaging Genetics; BIL&GIN = a multimodal multidimensional database for investigating hemispheric specialization; Bonn = University of Bonn; BrainSCALE=Brain Structure and Cognition: an Adolescence Longitudinal twin study; CAMH = Centre for Addiction and Mental Health; Cardiff = Cardiff University; CEG = Cognitive-experimental and Genetic study of ADHD and Control Sibling Pairs; CIAM = Cortical Inhibition and Attentional Modulation study; CLiNG = Clinical Neuroscience Göttingen**; **CODE =** **formerly** **Cognitive Behavioral Analysis System of Psychotherapy (CBASP) study; Edinburgh = The University of Edinburgh**; **ENIGMA-HIV = Enhancing NeuroImaging Genetics through Meta-Analysis-Human Immunodeficiency Virus Working Group; ENIGMA-OCD = Enhancing NeuroImaging Genetics through Meta-Analysis- Obsessive Compulsive Disorder Working Group;** **FBIRN = Function Biomedical Informatics Research Network;** **FIDMAG = Fundación para la Investigación y Docencia Maria Angustias Giménez**; **GSP = Brain Genomics Superstruct Project; HMS = Homburg Multidiagnosis Study**; **HUBIN = Human Brain Informatics; IDIVAL = Valdecilla Biomedical Research Institute; IMAGEN = the IMAGEN Consortium; IMH=Institute of Mental Health, Singapore; IMpACT = The International Multicentre persistent ADHD Genetics Collaboration; Indiana = Indiana University School of Medicine; Johns Hopkins = Johns Hopkins University; KaSP= The Karolinska Schizophrenia Project; Leiden = Leiden University; MAS = Memory and Ageing Study;** **MCIC = MIND Clinical Imaging Consortium formed by the Mental Illness and Neuroscience Discovery (MIND) Institute now the Mind Research Network;** **Melbourne = University of Melbourne**; **Meth-CT = study of methamphetamine users, University of Cape Town; MHRC = Mental Health Research Center; Muenster = Muenster University; NESDA = The Netherlands Study of Depression and Anxiety**; **NeuroIMAGE = Dutch part of the International Multicenter ADHD Genetics (IMAGE) study;** **Neuroventure: the imaging part of the Co-Venture Trial funded by the Canadian Institutes of Health Research (CIHR); NCNG = Norwegian Cognitive NeuroGenetics sample; NTR = Netherlands Twin Register**; **NU = Northwestern University; NUIG = National University of Ireland Galway**; **NYU = New York University**; **OATS = Older Australian Twins Study**; **Olin = Olin Neuropsychiatric Research Center;** **Oxford =Oxford University;** **QTIM = Queensland Twin Imaging; Sao Paulo = University of Sao Paulo; SCORE = University of Basel Study; SHIP-2 and SHIP TREND = Study of Health in Pomerania; Staged-Dep= Stages of Depression Study;** **Stanford = Stanford University**; **StrokeMRI = Stroke Magnetic Resonance Imaging; Sydney = University of Sydney; TOP = Tematisk Område Psykoser (Thematically Organized Psychosis Research); TS-EUROTRAIN = European-Wide Investigation and Training Network on the Etiology and Pathophysiology of Gilles de la Tourette Syndrome; Tuebingen = University of Tuebingen; UMCU = Universitair Medisch Centrum Utrecht**; **UNIBA = University of Bari Aldo Moro**; **UPENN=University of Pennsylvania; Yale = Yale University**

**Table S2.** Variance Explained by Age in fractional polynomial model

| **Cortical Region** | **R-Squared** |
| --- | --- |
| **LThickness** | 0.59 |
| **RThickness** | 0.59 |
| **L_bankssts** | 0.35 |
| **L_caudalanteriorcingulate** | 0.13 |
| **L_caudalmiddlefrontal** | 0.38 |
| **L_cuneus** | 0.29 |
| **L_entorhinal** | 0.01 |
| **L_fusiform** | 0.28 |
| **L_inferiorparietal** | 0.46 |
| **L_inferiortemporal** | 0.22 |
| **L_isthmuscingulate** | 0.33 |
| **L_lateraloccipital** | 0.25 |
| **L_lateralorbitofrontal** | 0.41 |
| **L_lingual** | 0.37 |
| **L_medialorbitofrontal** | 0.29 |
| **L_middletemporal** | 0.38 |
| **L_parahippocampal** | 0.11 |
| **L_paracentral** | 0.37 |
| **L_parsopercularis** | 0.46 |
| **L_parsorbitalis** | 0.32 |
| **L_parstriangularis** | 0.44 |
| **L_pericalcarine** | 0.19 |
| **L_postcentral** | 0.36 |
| **L_posteriorcingulate** | 0.43 |
| **L_precentral** | 0.41 |
| **L_precuneus** | 0.51 |
| **L_rostralanteriorcingulate** | 0.23 |
| **L_rostralmiddlefrontal** | 0.44 |
| **L_superiorfrontal** | 0.52 |
| **L_superiorparietal** | 0.39 |
| **L_superiortemporal** | 0.4 |
| **L_supramarginal** | 0.47 |
| **L_frontalpole** | 0.15 |
| **L_temporalpole** | 0.02 |
| **L_transversetemporal** | 0.27 |
| **L_insula** | 0.38 |
| **R_bankssts** | 0.34 |
| **R_caudalanteriorcingulate** | 0.14 |
| **R_caudalmiddlefrontal** | 0.37 |
| **R_cuneus** | 0.31 |
| **R_entorhinal** | 0.02 |
| **R_fusiform** | 0.32 |
| **R_inferiorparietal** | 0.48 |
| **R_inferiortemporal** | 0.23 |
| **R_isthmuscingulate** | 0.33 |
| **R_lateraloccipital** | 0.28 |
| **R_lateralorbitofrontal** | 0.36 |
| **R_lingual** | 0.38 |
| **R_medialorbitofrontal** | 0.3 |
| **R_middletemporal** | 0.37 |
| **R_parahippocampal** | 0.13 |
| **R_paracentral** | 0.37 |
| **R_parsopercularis** | 0.44 |
| **R_parsorbitalis** | 0.29 |
| **R_parstriangularis** | 0.44 |
| **R_pericalcarine** | 0.17 |
| **R_postcentral** | 0.36 |
| **R_posteriorcingulate** | 0.4 |
| **R_precentral** | 0.4 |
| **R_precuneus** | 0.5 |
| **R_rostralanteriorcingulate** | 0.18 |
| **R_rostralmiddlefrontal** | 0.4 |
| **R_superiorfrontal** | 0.5 |
| **R_superiorparietal** | 0.39 |
| **R_superiortemporal** | 0.41 |
| **R_supramarginal** | 0.48 |
| **R_frontalpole** | 0.17 |
| **R_temporalpole** | 0.03 |
| **R_transversetemporal** | 0.28 |
| **R_insula** | 0.38 |

**Table S3-Pearson's Correlation Coefficient between Age and Cortical Thickness**

|  | **All** | | | **Males** | | | **Females** | | |
| --- | --- | --- | --- | --- | --- | --- | --- | --- | --- |
| **Region** | **3-29 years** | **30-59 years** | **60-90 years** | **3-29 years** | **30-59 years** | **60-90 years** | **3-29 years** | **30-59 years** | **60-90 years** |
| **LThickness** | -0.572 | -0.327 | -0.313 | -0.586 | -0.376 | -0.3 | -0.558 | -0.274 | -0.342 |
| **RThickness** | -0.575 | -0.328 | -0.291 | -0.587 | -0.386 | -0.285 | -0.564 | -0.266 | -0.314 |
| **L_bankssts** | -0.401 | -0.216 | -0.176 | -0.403 | -0.244 | -0.17 | -0.398 | -0.182 | -0.187 |
| **L_caudalanteriorcingulate** | -0.278 | -0.056 | 0.073 | -0.3 | -0.067 | 0.066 | -0.268 | -0.054 | 0.073 |
| **L_caudalmiddlefrontal** | -0.383 | -0.258 | -0.163 | -0.395 | -0.308 | -0.174 | -0.373 | -0.217 | -0.163 |
| **L_cuneus** | -0.385 | -0.159 | -0.11 | -0.423 | -0.18 | -0.096 | -0.348 | -0.139 | -0.126 |
| **L_entorhinal** | 0.008 | -0.004 | -0.198 | 0.037 | -0.023 | -0.174 | -0.018 | 0.029 | -0.22 |
| **L_fusiform** | -0.343 | -0.182 | -0.173 | -0.34 | -0.218 | -0.174 | -0.345 | -0.14 | -0.18 |
| **L_inferiorparietal** | -0.52 | -0.23 | -0.263 | -0.543 | -0.277 | -0.233 | -0.499 | -0.192 | -0.304 |
| **L_inferiortemporal** | -0.31 | -0.144 | -0.092 | -0.29 | -0.175 | -0.088 | -0.327 | -0.103 | -0.103 |
| **L_isthmuscingulate** | -0.362 | -0.224 | -0.147 | -0.37 | -0.286 | -0.137 | -0.357 | -0.168 | -0.166 |
| **L_lateraloccipital** | -0.381 | -0.088 | -0.233 | -0.419 | -0.132 | -0.252 | -0.342 | -0.049 | -0.226 |
| **L_lateralorbitofrontal** | -0.48 | -0.191 | -0.146 | -0.493 | -0.229 | -0.136 | -0.468 | -0.145 | -0.162 |
| **L_lingual** | -0.404 | -0.216 | -0.224 | -0.445 | -0.249 | -0.198 | -0.364 | -0.176 | -0.249 |
| **L_medialorbitofrontal** | -0.396 | -0.153 | -0.067 | -0.393 | -0.178 | -0.012 | -0.398 | -0.113 | -0.12 |
| **L_middletemporal** | -0.385 | -0.218 | -0.227 | -0.365 | -0.261 | -0.19 | -0.404 | -0.162 | -0.27 |
| **L_parahippocampal** | -0.171 | -0.105 | -0.129 | -0.181 | -0.13 | -0.102 | -0.167 | -0.082 | -0.159 |
| **L_paracentral** | -0.427 | -0.225 | -0.198 | -0.462 | -0.236 | -0.183 | -0.395 | -0.223 | -0.217 |
| **L_parsopercularis** | -0.424 | -0.325 | -0.164 | -0.422 | -0.38 | -0.12 | -0.425 | -0.258 | -0.212 |
| **L_parsorbitalis** | -0.386 | -0.214 | -0.093 | -0.399 | -0.269 | -0.076 | -0.377 | -0.161 | -0.122 |
| **L_parstriangularis** | -0.447 | -0.29 | -0.109 | -0.463 | -0.344 | -0.1 | -0.432 | -0.232 | -0.128 |
| **L_pericalcarine** | -0.274 | -0.142 | -0.122 | -0.306 | -0.159 | -0.068 | -0.244 | -0.13 | -0.172 |
| **L_postcentral** | -0.382 | -0.226 | -0.226 | -0.419 | -0.267 | -0.243 | -0.349 | -0.193 | -0.229 |
| **L_posteriorcingulate** | -0.487 | -0.254 | -0.083 | -0.485 | -0.29 | -0.069 | -0.49 | -0.221 | -0.104 |
| **L_precentral** | -0.278 | -0.29 | -0.321 | -0.287 | -0.326 | -0.319 | -0.271 | -0.266 | -0.336 |
| **L_precuneus** | -0.543 | -0.262 | -0.294 | -0.567 | -0.311 | -0.265 | -0.518 | -0.207 | -0.33 |
| **L_rostralanteriorcingulate** | -0.326 | -0.129 | 0.032 | -0.327 | -0.158 | 0.052 | -0.334 | -0.103 | 0.005 |
| **L_rostralmiddlefrontal** | -0.5 | -0.227 | -0.092 | -0.521 | -0.285 | -0.11 | -0.48 | -0.167 | -0.089 |
| **L_superiorfrontal** | -0.503 | -0.31 | -0.239 | -0.531 | -0.354 | -0.25 | -0.48 | -0.279 | -0.244 |
| **L_superiorparietal** | -0.484 | -0.176 | -0.266 | -0.519 | -0.216 | -0.265 | -0.451 | -0.145 | -0.282 |
| **L_superiortemporal** | -0.312 | -0.285 | -0.37 | -0.294 | -0.32 | -0.325 | -0.328 | -0.24 | -0.417 |
| **L_supramarginal** | -0.47 | -0.279 | -0.269 | -0.481 | -0.333 | -0.244 | -0.46 | -0.232 | -0.309 |
| **L_frontalpole** | -0.244 | -0.096 | -0.02 | -0.258 | -0.14 | -0.025 | -0.235 | -0.057 | -0.019 |
| **L_temporalpole** | 0.003 | -0.035 | -0.135 | 0.018 | -0.066 | -0.125 | -0.012 | 0.004 | -0.155 |
| **L_transversetemporal** | -0.262 | -0.206 | -0.169 | -0.273 | -0.213 | -0.189 | -0.259 | -0.209 | -0.161 |
| **L_insula** | -0.381 | -0.251 | -0.165 | -0.382 | -0.291 | -0.142 | -0.378 | -0.192 | -0.188 |
| **R_bankssts** | -0.385 | -0.234 | -0.195 | -0.384 | -0.279 | -0.199 | -0.385 | -0.18 | -0.196 |
| **R_caudalanteriorcingulate** | -0.279 | -0.102 | 0.09 | -0.291 | -0.111 | 0.127 | -0.28 | -0.11 | 0.049 |
| **R_caudalmiddlefrontal** | -0.379 | -0.248 | -0.186 | -0.386 | -0.293 | -0.174 | -0.373 | -0.212 | -0.209 |
| **R_cuneus** | -0.415 | -0.169 | -0.145 | -0.462 | -0.18 | -0.159 | -0.368 | -0.162 | -0.138 |
| **R_entorhinal** | 0.024 | -0.002 | -0.212 | 0.05 | -0.009 | -0.178 | -0.001 | 0.013 | -0.252 |
| **R_fusiform** | -0.367 | -0.2 | -0.235 | -0.368 | -0.248 | -0.245 | -0.366 | -0.146 | -0.232 |
| **R_inferiorparietal** | -0.52 | -0.235 | -0.255 | -0.542 | -0.29 | -0.249 | -0.499 | -0.187 | -0.276 |
| **R_inferiortemporal** | -0.32 | -0.121 | -0.153 | -0.308 | -0.179 | -0.14 | -0.33 | -0.049 | -0.176 |
| **R_isthmuscingulate** | -0.372 | -0.24 | -0.11 | -0.376 | -0.286 | -0.11 | -0.369 | -0.2 | -0.122 |
| **R_lateraloccipital** | -0.382 | -0.109 | -0.254 | -0.414 | -0.165 | -0.252 | -0.349 | -0.058 | -0.265 |
| **R_lateralorbitofrontal** | -0.447 | -0.207 | -0.017 | -0.44 | -0.254 | -0.005 | -0.453 | -0.15 | -0.038 |
| **R_lingual** | -0.426 | -0.223 | -0.192 | -0.455 | -0.261 | -0.166 | -0.398 | -0.178 | -0.221 |
| **R_medialorbitofrontal** | -0.426 | -0.177 | 0.032 | -0.427 | -0.206 | 0.072 | -0.425 | -0.14 | -0.012 |
| **R_middletemporal** | -0.403 | -0.213 | -0.168 | -0.381 | -0.25 | -0.156 | -0.421 | -0.164 | -0.188 |
| **R_parahippocampal** | -0.183 | -0.11 | -0.127 | -0.197 | -0.148 | -0.085 | -0.177 | -0.076 | -0.176 |
| **R_paracentral** | -0.411 | -0.229 | -0.227 | -0.443 | -0.249 | -0.26 | -0.381 | -0.212 | -0.199 |
| **R_parsopercularis** | -0.411 | -0.315 | -0.113 | -0.405 | -0.355 | -0.07 | -0.414 | -0.266 | -0.157 |
| **R_parsorbitalis** | -0.342 | -0.193 | -0.054 | -0.356 | -0.261 | -0.045 | -0.335 | -0.13 | -0.078 |
| **R_parstriangularis** | -0.442 | -0.304 | -0.078 | -0.445 | -0.358 | -0.046 | -0.44 | -0.247 | -0.115 |
| **R_pericalcarine** | -0.269 | -0.155 | -0.125 | -0.303 | -0.157 | -0.119 | -0.237 | -0.159 | -0.131 |
| **R_postcentral** | -0.39 | -0.221 | -0.245 | -0.424 | -0.264 | -0.284 | -0.362 | -0.186 | -0.226 |
| **R_posteriorcingulate** | -0.462 | -0.224 | -0.058 | -0.466 | -0.264 | -0.004 | -0.459 | -0.186 | -0.118 |
| **R_precentral** | -0.292 | -0.278 | -0.304 | -0.313 | -0.318 | -0.329 | -0.274 | -0.249 | -0.294 |
| **R_precuneus** | -0.532 | -0.258 | -0.307 | -0.552 | -0.298 | -0.309 | -0.511 | -0.21 | -0.313 |
| **R_rostralanteriorcingulate** | -0.294 | -0.142 | 0.104 | -0.273 | -0.19 | 0.151 | -0.317 | -0.083 | 0.055 |
| **R_rostralmiddlefrontal** | -0.482 | -0.213 | -0.023 | -0.494 | -0.263 | 0.001 | -0.47 | -0.166 | -0.057 |
| **R_superiorfrontal** | -0.487 | -0.294 | -0.189 | -0.503 | -0.348 | -0.167 | -0.476 | -0.247 | -0.223 |
| **R_superiorparietal** | -0.475 | -0.167 | -0.239 | -0.513 | -0.221 | -0.258 | -0.438 | -0.123 | -0.236 |
| **R_superiortemporal** | -0.341 | -0.29 | -0.357 | -0.328 | -0.322 | -0.322 | -0.352 | -0.247 | -0.396 |
| **R_supramarginal** | -0.483 | -0.281 | -0.277 | -0.494 | -0.364 | -0.275 | -0.473 | -0.196 | -0.291 |
| **R_frontalpole** | -0.263 | -0.111 | -0.025 | -0.277 | -0.165 | -0.025 | -0.256 | -0.064 | -0.034 |
| **R_temporalpole** | 0.077 | -0.02 | -0.208 | 0.107 | -0.059 | -0.158 | 0.052 | 0.026 | -0.263 |
| **R_transversetemporal** | -0.285 | -0.196 | -0.157 | -0.297 | -0.202 | -0.211 | -0.278 | -0.19 | -0.115 |
| **R_insula** | -0.371 | -0.26 | -0.199 | -0.372 | -0.307 | -0.159 | -0.368 | -0.197 | -0.241 |

**Table S4. Inter-individual variations in cortical thickness**

| **Cortical Region- Both sexes** | **Mean Inter-individual variation (SD)** | | |  |
| --- | --- | --- | --- | --- |
|  | **3-29 years** | **30-59 years** | **60-90 years** | **Unadjusted P value for F test** |
| **LThickness** | 0.07 (0.06) | 0.07 ( 0.06) | 0.07 (0.06) | 0.1460 |
| **RThickness** | 0.07 ( 0.06) | 0.07 (0.06) | 0.08 ( 0.06) | 0.1097 |
| **L_bankssts** | 0.13 ( 0.11) | 0.13 ( 0.10) | 0.14 (0.10) | 0.1348 |
| **L_caudalanteriorcingulate** | 0.20 ( 0.15) | 0.20 (0.15) | 0.22 (0.17) | 0.0002 |
| **L_caudalmiddlefrontal** | 0.11 (0.09) | 0.11 (0.09) | 0.12 (0.09) | 0.5413 |
| **L_cuneus** | 0.11 (0.08) | 0.10 (0.08) | 0.10 (0.08) | 0.0025 |
| **L_entorhinal** | 0.27 (0.21) | 0.27 ( 0.21) | 0.28 (0.22) | 0.7225 |
| **L_fusiform** | 0.11 ( 0.09) | 0.11 (0.09) | 0.12 ( 0.09) | 0.1722 |
| **L_inferiorparietal** | 0.10 ( 0.08) | 0.10 (0.08) | 0.11 (0.08) | 0.2081 |
| **L_inferiortemporal** | 0.13 ( 0.10) | 0.13 ( 0.10) | 0.14 (0.10) | 0.0449 |
| **L_isthmuscingulate** | 0.16 (0.12) | 0.16 (0.12) | 0.16 (0.13) | 0.5258 |
| **L_lateraloccipital** | 0.10 (0.07) | 0.10 (0.07) | 0.10 ( 0.08) | 0.2683 |
| **L_lateralorbitofrontal** | 0.12 (0.10) | 0.12 (0.09) | 0.13 (0.10) | 0.6571 |
| **L_lingual** | 0.10 (0.07) | 0.09 (0.07) | 0.09 (0.07) | 0.5093 |
| **L_medialorbitofrontal** | 0.13 (0.10) | 0.13 (0.10) | 0.13 (0.11) | 0.9359 |
| **L_middletemporal** | 0.13 (0.10) | 0.12 (0.10) | 0.13 (0.10) | 0.0033 |
| **L_parahippocampal** | 0.24 (0.18) | 0.25 (0.18) | 0.25 (0.18) | 0.2011 |
| **L_paracentral** | 0.12 (0.09) | 0.12 (0.09) | 0.13 ( 0.10) | 0.1578 |
| **L_parsopercularis** | 0.12 (0.09) | 0.12 (0.09) | 0.12 (0.09) | 0.4775 |
| **L_parsorbitalis** | 0.18 (0.13) | 0.17 (0.13) | 0.17 (0.14) | 0.0847 |
| **L_parstriangularis** | 0.13 (0.10) | 0.13 (0.10) | 0.13 (0.10) | 0.3120 |
| **L_pericalcarine** | 0.10 (0.08) | 0.10 (0.08) | 0.10 (0.08) | 0.1714 |
| **L_postcentral** | 0.09 (0.07) | 0.09 (0.07) | 0.10 (0.07) | 0.0441 |
| **L_posteriorcingulate** | 0.13 (0.09) | 0.12 (0.09) | 0.13 (0.10) | 0.0227 |
| **L_precentral** | 0.10 (0.08) | 0.11 (0.08) | 0.11 (0.09) | 0 |
| **L_precuneus** | 0.10 (0.08) | 0.10 (0.08) | 0.10 (0.08) | 0.7903 |
| **L_rostralanteriorcingulate** | 0.19 (0.14) | 0.18 (0.14) | 0.19 (0.14) | 0.0664 |
| **L_rostralmiddlefrontal** | 0.11 (0.08) | 0.10 (0.08) | 0.11 (0.09) | 0.5133 |
| **L_superiorfrontal** | 0.11 (0.09) | 0.11 (0.09) | 0.12 (0.09) | 0.6952 |
| **L_superiorparietal** | 0.10 (0.07) | 0.10 (0.07) | 0.10 (0.08) | 0.9745 |
| **L_superiortemporal** | 0.12 (0.10) | 0.12 (0.09) | 0.13 (0.10) | 0.0815 |
| **L_supramarginal** | 0.11 (0.08) | 0.11 (0.08) | 0.11 (0.09) | 0.3212 |
| **L_frontalpole** | 0.25 (0.19) | 0.23 (0.18) | 0.24 (0.19) | 0.0005 |
| **L_temporalpole** | 0.28 (0.21) | 0.28 (0.21) | 0.27 (0.22) | 0.0379 |
| **L_transversetemporal** | 0.18 (0.13) | 0.18 (0.13) | 0.18 (0.13) | 0.6329 |
| **L_insula** | 0.12 (0.09) | 0.12 (0.09) | 0.13 (0.10) | 0.3535 |
| **R_bankssts** | 0.14 (0.11) | 0.14 (0.11) | 0.14 (0.11) | 0.2366 |
| **R_caudalanteriorcingulate** | 0.18 (0.14) | 0.18 (0.14) | 0.20 (0.16) | 0.0001 |
| **R_caudalmiddlefrontal** | 0.12 (0.09) | 0.12 (0.09) | 0.12 (0.09) | 0.8999 |
| **R_cuneus** | 0.11 (0.08) | 0.11 (0.08) | 0.11 (0.08) | 0.0112 |
| **R_entorhinal** | 0.30 (0.22) | 0.30 (0.22) | 0.30 (0.23) | 0.1436 |
| **R_fusiform** | 0.11 (0.09) | 0.11 (0.09) | 0.12 (0.09) | 0.0303 |
| **R_inferiorparietal** | 0.10 (0.08) | 0.10 (0.08) | 0.11 (0.08) | 0.1918 |
| **R_inferiortemporal** | 0.13 (0.10) | 0.13 (0.10) | 0.14 (0.10) | 0.0857 |
| **R_isthmuscingulate** | 0.16 (0.12) | 0.15 (0.12) | 0.16 (0.12) | 0.0196 |
| **R_lateraloccipital** | 0.10 (0.07) | 0.10 (0.08) | 0.11 (0.08) | 0.8078 |
| **R_lateralorbitofrontal** | 0.13 (0.10) | 0.12 (0.10) | 0.13 (0.10) | 0.0027 |
| **R_lingual** | 0.10 (0.07) | 0.10 (0.07) | 0.10 (0.07) | 0.1938 |
| **R_medialorbitofrontal** | 0.14 (0.11) | 0.14 (0.11) | 0.14 (0.11) | 0.0545 |
| **R_middletemporal** | 0.13 (0.10) | 0.12 (0.10) | 0.13 (0.10) | 0.0009 |
| **R_parahippocampal** | 0.21 (0.16) | 0.21 (0.16) | 0.21 (0.16) | 0.3844 |
| **R_paracentral** | 0.12 (0.09) | 0.12 (0.09) | 0.13 (0.09) | 0.0248 |
| **R_parsopercularis** | 0.12 (0.09) | 0.12 (0.09) | 0.12 (0.09) | 0.6346 |
| **R_parsorbitalis** | 0.17 (0.13) | 0.16 (0.13) | 0.17 (0.13) | 0.0043 |
| **R_parstriangularis** | 0.12 (0.09) | 0.12 (0.09) | 0.12 (0.09) | 0.4239 |
| **R_pericalcarine** | 0.10 (0.08) | 0.10 (0.08) | 0.10 (0.08) | 0.0415 |
| **R_postcentral** | 0.10 (0.08) | 0.10 (0.08) | 0.10 (0.08) | 0.3740 |
| **R_posteriorcingulate** | 0.12 (0.10) | 0.12 (0.09) | 0.13 (0.10) | 0.0076 |
| **R_precentral** | 0.10 (0.08) | 0.11 (0.08) | 0.12 (0.09) | 0.0000 |
| **R_precuneus** | 0.10(0.08) | 0.10 (0.08) | 0.10 (0.08) | 0.6332 |
| **R_rostralanteriorcingulate** | 0.19 (0.15) | 0.18 (0.14) | 0.20 (0.15) | 0.0376 |
| **R_rostralmiddlefrontal** | 0.11 (0.08) | 0.10 (0.08) | 0.11 (0.08) | 0.2797 |
| **R_superiorfrontal** | 0.11 (0.09) | 0.11 (0.09) | 0.12 (0.09) | 0.6076 |
| **R_superiorparietal** | 0.10 (0.07) | 0.10 (0.07) | 0.10 (0.08) | 0.7408 |
| **R_superiortemporal** | 0.12 (0.10) | 0.12 (0.09) | 0.13 (0.09) | 0.0476 |
| **R_supramarginal** | 0.11 (0.08) | 0.11 (0.09) | 0.11 (0.09) | 0.0463 |
| **R_frontalpole** | 0.24 (0.18) | 0.23 (0.17) | 0.24 (0.18) | 0.0515 |
| **R_temporalpole** | 0.29 (0.22) | 0.28 (0.22) | 0.29 (0.22) | 0.0013 |
| **R_transversetemporal** | 0.18 (0.13) | 0.18 (0.13) | 0.19 (0.14) | 0.0427 |
| **R_insula** | 0.12 (0.10) | 0.12 (0.10) | 0.13 (0.10) | 0.1187 |

**SD: Standard Deviation; no significant difference was observed between males and females in inter-individual variance in either age groups**

**Table S5. Centile Values for Cortical Thickness**

|  | **Age** | **C0.4** | **C1** | **C2.5** | **C5** | **C10** | **C25** | **C50** | **C75** | **C90** | **C95** | **C97.5** | **C99** | **C99.6** |
| --- | --- | --- | --- | --- | --- | --- | --- | --- | --- | --- | --- | --- | --- | --- |
| **LThickness** | 3 | 2.574216 | 2.627425 | 2.680705 | 2.722176 | 2.766101 | 2.832661 | 2.900059 | 2.963892 | 3.020608 | 3.055015 | 3.08551 | 3.122091 | 3.155958 |
|  | 5 | 2.538326 | 2.588923 | 2.64007 | 2.680195 | 2.722973 | 2.788275 | 2.854874 | 2.918241 | 2.974653 | 3.00888 | 3.039199 | 3.07553 | 3.109115 |
|  | 10 | 2.447806 | 2.492656 | 2.538978 | 2.575984 | 2.616043 | 2.678285 | 2.742885 | 2.805084 | 2.860783 | 2.894622 | 2.92458 | 2.960419 | 2.99346 |
|  | 15 | 2.361206 | 2.401106 | 2.44313 | 2.477275 | 2.514781 | 2.574082 | 2.636753 | 2.697911 | 2.753119 | 2.786773 | 2.816599 | 2.852281 | 2.885151 |
|  | 20 | 2.298128 | 2.334287 | 2.373015 | 2.404954 | 2.440498 | 2.497607 | 2.559009 | 2.619745 | 2.675061 | 2.708933 | 2.739011 | 2.775037 | 2.808237 |
|  | 30 | 2.222697 | 2.256642 | 2.293526 | 2.324325 | 2.358969 | 2.415319 | 2.47659 | 2.537546 | 2.593053 | 2.62692 | 2.656862 | 2.692518 | 2.725139 |
|  | 40 | 2.169817 | 2.204566 | 2.242394 | 2.274013 | 2.309583 | 2.36735 | 2.429851 | 2.491467 | 2.546881 | 2.580292 | 2.609549 | 2.644013 | 2.675166 |
|  | 50 | 2.12603 | 2.162181 | 2.201505 | 2.234334 | 2.271199 | 2.330852 | 2.39496 | 2.457551 | 2.513187 | 2.54638 | 2.575205 | 2.608855 | 2.638969 |
|  | 60 | 2.081535 | 2.119081 | 2.159873 | 2.193878 | 2.231991 | 2.29346 | 2.359143 | 2.422777 | 2.478826 | 2.511998 | 2.540626 | 2.573818 | 2.603305 |
|  | 70 | 2.034683 | 2.07162 | 2.112019 | 2.145892 | 2.184053 | 2.245974 | 2.31255 | 2.377325 | 2.434498 | 2.468346 | 2.497547 | 2.531373 | 2.56138 |
|  | 80 | 1.995061 | 2.029207 | 2.067151 | 2.099433 | 2.1363 | 2.197199 | 2.264085 | 2.330456 | 2.390004 | 2.42565 | 2.456617 | 2.492724 | 2.52495 |
|  | 90 | 1.954851 | 1.98544 | 2.020066 | 2.050053 | 2.08489 | 2.143819 | 2.210509 | 2.278703 | 2.341571 | 2.379955 | 2.413751 | 2.453677 | 2.48978 |
| **RThickness** | 3 | 2.593615 | 2.642276 | 2.692291 | 2.732058 | 2.774887 | 2.840863 | 2.90835 | 2.972032 | 3.027685 | 3.060764 | 3.089548 | 3.123336 | 3.153848 |
|  | 5 | 2.551742 | 2.598631 | 2.647091 | 2.685801 | 2.727661 | 2.792459 | 2.859093 | 2.92225 | 2.977633 | 3.010622 | 3.039364 | 3.073143 | 3.103678 |
|  | 10 | 2.450658 | 2.493423 | 2.538209 | 2.574395 | 2.613916 | 2.675846 | 2.7404 | 2.802304 | 2.857082 | 2.889901 | 2.918598 | 2.952435 | 2.983116 |
|  | 15 | 2.360692 | 2.399656 | 2.441005 | 2.474808 | 2.512111 | 2.571336 | 2.634002 | 2.694901 | 2.749375 | 2.782248 | 2.811126 | 2.845322 | 2.876457 |
|  | 20 | 2.295709 | 2.331728 | 2.370402 | 2.402355 | 2.437958 | 2.495189 | 2.556628 | 2.617132 | 2.671855 | 2.705129 | 2.734503 | 2.769452 | 2.801417 |
|  | 30 | 2.216045 | 2.251019 | 2.288803 | 2.320189 | 2.355328 | 2.412138 | 2.473488 | 2.534161 | 2.589162 | 2.622626 | 2.652165 | 2.68729 | 2.719388 |
|  | 40 | 2.158664 | 2.195518 | 2.235127 | 2.267865 | 2.304336 | 2.362877 | 2.42547 | 2.486679 | 2.541518 | 2.574567 | 2.603532 | 2.637716 | 2.668703 |
|  | 50 | 2.115501 | 2.153948 | 2.195153 | 2.229115 | 2.266841 | 2.327142 | 2.391225 | 2.453439 | 2.508751 | 2.541867 | 2.57075 | 2.604659 | 2.635224 |
|  | 60 | 2.07552 | 2.11457 | 2.156484 | 2.191074 | 2.229533 | 2.291052 | 2.356434 | 2.419838 | 2.476081 | 2.509675 | 2.538914 | 2.573161 | 2.603947 |
|  | 70 | 2.031973 | 2.069603 | 2.11038 | 2.144318 | 2.182342 | 2.243758 | 2.30975 | 2.374359 | 2.432092 | 2.466735 | 2.496968 | 2.532462 | 2.564435 |
|  | 80 | 1.997237 | 2.032437 | 2.071117 | 2.10373 | 2.140711 | 2.201394 | 2.267846 | 2.33407 | 2.394148 | 2.430576 | 2.462586 | 2.500412 | 2.534701 |
|  | 90 | 1.957392 | 1.989816 | 2.025979 | 2.056898 | 2.09243 | 2.151808 | 2.218326 | 2.286124 | 2.348888 | 2.387508 | 2.421785 | 2.462694 | 2.500147 |
| **L_bankssts** | 3 | 2.429632 | 2.508717 | 2.59415 | 2.664559 | 2.742087 | 2.862459 | 2.982549 | 3.101039 | 3.216674 | 3.289537 | 3.354638 | 3.432282 | 3.502859 |
|  | 5 | 2.386275 | 2.463954 | 2.547945 | 2.617234 | 2.69361 | 2.812399 | 2.931231 | 3.048484 | 3.162615 | 3.234416 | 3.298507 | 3.374883 | 3.444255 |
|  | 10 | 2.275775 | 2.34987 | 2.430169 | 2.496574 | 2.569968 | 2.684616 | 2.800077 | 2.914016 | 3.024211 | 3.093258 | 3.154746 | 3.227867 | 3.29416 |
|  | 15 | 2.166407 | 2.236954 | 2.313582 | 2.377107 | 2.447504 | 2.557951 | 2.669925 | 2.780435 | 2.886636 | 2.952914 | 3.011797 | 3.081676 | 3.144913 |
|  | 20 | 2.089784 | 2.157849 | 2.231949 | 2.293526 | 2.361948 | 2.469764 | 2.579798 | 2.688406 | 2.792122 | 2.856591 | 2.913733 | 2.981406 | 3.042534 |
|  | 30 | 2.015452 | 2.081121 | 2.152935 | 2.212904 | 2.279893 | 2.38637 | 2.496479 | 2.605188 | 2.707713 | 2.770938 | 2.826715 | 2.892502 | 2.95171 |
|  | 40 | 1.96689 | 2.031005 | 2.101433 | 2.160528 | 2.226892 | 2.333294 | 2.444765 | 2.554854 | 2.657411 | 2.720159 | 2.775259 | 2.839985 | 2.898028 |
|  | 50 | 1.926667 | 1.989501 | 2.058828 | 2.117279 | 2.183268 | 2.289987 | 2.403238 | 2.515122 | 2.618101 | 2.680615 | 2.735258 | 2.799189 | 2.856314 |
|  | 60 | 1.883352 | 1.944803 | 2.012906 | 2.070599 | 2.136077 | 2.242885 | 2.357684 | 2.471141 | 2.574335 | 2.636493 | 2.690577 | 2.753602 | 2.809718 |
|  | 70 | 1.840206 | 1.900282 | 1.967153 | 2.024072 | 2.089011 | 2.19586 | 2.312157 | 2.427143 | 2.530514 | 2.592299 | 2.645814 | 2.707931 | 2.763043 |
|  | 80 | 1.796837 | 1.855531 | 1.92115 | 1.977267 | 2.041629 | 2.148443 | 2.26616 | 2.382603 | 2.486093 | 2.547473 | 2.600399 | 2.66159 | 2.715693 |
|  | 90 | 1.737627 | 1.794424 | 1.858197 | 1.912993 | 1.976168 | 2.081921 | 2.199913 | 2.316687 | 2.41931 | 2.479712 | 2.531561 | 2.591274 | 2.643888 |
| **L_caudalanteriorcingulate** | 3 | 2.413618 | 2.513054 | 2.620928 | 2.71092 | 2.812122 | 2.976886 | 3.156686 | 3.336797 | 3.502394 | 3.604373 | 3.695225 | 3.804343 | 3.905136 |
|  | 5 | 2.369966 | 2.467668 | 2.573717 | 2.662235 | 2.761834 | 2.924118 | 3.10142 | 3.279278 | 3.443056 | 3.544045 | 3.634105 | 3.742387 | 3.842523 |
|  | 10 | 2.267532 | 2.361163 | 2.462932 | 2.547991 | 2.643827 | 2.8003 | 2.971764 | 3.144384 | 3.30396 | 3.402685 | 3.49095 | 3.597365 | 3.696067 |
|  | 15 | 2.182457 | 2.272665 | 2.370859 | 2.453049 | 2.545788 | 2.697537 | 2.864349 | 3.032917 | 3.189382 | 3.286513 | 3.37358 | 3.478846 | 3.57678 |
|  | 20 | 2.119157 | 2.206601 | 2.301964 | 2.381928 | 2.472316 | 2.620597 | 2.784171 | 2.950125 | 3.104802 | 3.201149 | 3.287733 | 3.392699 | 3.490639 |
|  | 30 | 2.039291 | 2.122242 | 2.213189 | 2.28983 | 2.376872 | 2.520582 | 2.680401 | 2.843873 | 2.997399 | 3.093579 | 3.180364 | 3.286013 | 3.385013 |
|  | 40 | 2.001267 | 2.081142 | 2.169372 | 2.244235 | 2.329792 | 2.472201 | 2.632047 | 2.796871 | 2.952622 | 3.050566 | 3.139139 | 3.247168 | 3.348563 |
|  | 50 | 1.982083 | 2.061408 | 2.14967 | 2.22505 | 2.3117 | 2.456953 | 2.621175 | 2.791351 | 2.952508 | 3.053857 | 3.145433 | 3.256957 | 3.361404 |
|  | 60 | 1.965346 | 2.047307 | 2.138914 | 2.217445 | 2.307989 | 2.460214 | 2.632564 | 2.810865 | 2.978922 | 3.08402 | 3.178507 | 3.292883 | 3.399251 |
|  | 70 | 1.933245 | 2.020924 | 2.118945 | 2.202944 | 2.299693 | 2.4619 | 2.644466 | 2.831586 | 3.005866 | 3.11366 | 3.209712 | 3.324837 | 3.430726 |
|  | 80 | 1.906909 | 2.004709 | 2.113565 | 2.206415 | 2.312821 | 2.489804 | 2.686616 | 2.885372 | 3.067501 | 3.178603 | 3.276564 | 3.392653 | 3.498129 |
|  | 90 | 1.88595 | 1.998553 | 2.122804 | 2.227908 | 2.34738 | 2.543795 | 2.758813 | 2.972234 | 3.164419 | 3.280029 | 3.380926 | 3.499217 | 3.605484 |
| **L_caudalmiddlefrontal** | 3 | 2.332929 | 2.422606 | 2.512801 | 2.582994 | 2.656848 | 2.766179 | 2.870975 | 2.966611 | 3.049915 | 3.098502 | 3.139827 | 3.186899 | 3.227859 |
|  | 5 | 2.326167 | 2.410798 | 2.497032 | 2.564841 | 2.636787 | 2.744325 | 2.848447 | 2.944225 | 3.028146 | 3.07729 | 3.119199 | 3.167053 | 3.208792 |
|  | 10 | 2.30208 | 2.3758 | 2.453074 | 2.515277 | 2.582583 | 2.685557 | 2.787806 | 2.883803 | 2.969239 | 3.019816 | 3.063256 | 3.113196 | 3.157041 |
|  | 15 | 2.265792 | 2.330124 | 2.399212 | 2.45601 | 2.518618 | 2.616637 | 2.716534 | 2.812424 | 2.899291 | 2.95137 | 2.996479 | 3.048763 | 3.095034 |
|  | 20 | 2.22425 | 2.281134 | 2.343422 | 2.395542 | 2.453931 | 2.547286 | 2.644824 | 2.740543 | 2.828867 | 2.88254 | 2.92946 | 2.984342 | 3.033351 |
|  | 30 | 2.152156 | 2.202819 | 2.25924 | 2.307212 | 2.361781 | 2.450862 | 2.546373 | 2.642284 | 2.732463 | 2.788056 | 2.83715 | 2.895161 | 2.947502 |
|  | 40 | 2.100779 | 2.150685 | 2.206443 | 2.254002 | 2.308278 | 2.397298 | 2.49335 | 2.590186 | 2.681355 | 2.737629 | 2.787375 | 2.846226 | 2.899392 |
|  | 50 | 2.046383 | 2.099085 | 2.157568 | 2.207135 | 2.263366 | 2.354873 | 2.452692 | 2.550159 | 2.640705 | 2.696043 | 2.744634 | 2.801735 | 2.85298 |
|  | 60 | 1.985082 | 2.042442 | 2.105321 | 2.158017 | 2.217178 | 2.312155 | 2.412072 | 2.509871 | 2.599057 | 2.652833 | 2.699624 | 2.754123 | 2.802605 |
|  | 70 | 1.94062 | 1.998752 | 2.062446 | 2.11581 | 2.175714 | 2.271904 | 2.373169 | 2.472132 | 2.562042 | 2.616118 | 2.663096 | 2.717736 | 2.766277 |
|  | 80 | 1.912415 | 1.967104 | 2.027741 | 2.079112 | 2.137392 | 2.232325 | 2.33406 | 2.434969 | 2.52766 | 2.58387 | 2.632984 | 2.690437 | 2.741775 |
|  | 90 | 1.90218 | 1.95197 | 2.008108 | 2.056442 | 2.112143 | 2.204879 | 2.307057 | 2.411022 | 2.508581 | 2.568712 | 2.621856 | 2.684747 | 2.741605 |
| **L_cuneus** | 3 | 1.919166 | 1.959809 | 2.006501 | 2.047504 | 2.095772 | 2.178878 | 2.274957 | 2.375166 | 2.469062 | 2.527004 | 2.578355 | 2.639374 | 2.694838 |
|  | 5 | 1.859584 | 1.89952 | 1.94542 | 1.985745 | 2.033238 | 2.115066 | 2.20976 | 2.308636 | 2.401388 | 2.458676 | 2.509481 | 2.569895 | 2.624847 |
|  | 10 | 1.727075 | 1.765463 | 1.809629 | 1.848472 | 1.89427 | 1.973313 | 2.065011 | 2.161032 | 2.251375 | 2.30731 | 2.357004 | 2.416208 | 2.470167 |
|  | 15 | 1.624333 | 1.661601 | 1.704513 | 1.742286 | 1.786866 | 1.863927 | 1.953536 | 2.047639 | 2.136451 | 2.191581 | 2.240657 | 2.299245 | 2.352764 |
|  | 20 | 1.561371 | 1.59815 | 1.640515 | 1.677827 | 1.721891 | 1.798151 | 1.887007 | 1.980567 | 2.069143 | 2.124276 | 2.173458 | 2.23231 | 2.286204 |
|  | 30 | 1.501126 | 1.537475 | 1.579286 | 1.616077 | 1.659507 | 1.734688 | 1.822448 | 1.915221 | 2.003559 | 2.058858 | 2.108422 | 2.168053 | 2.222998 |
|  | 40 | 1.473445 | 1.509628 | 1.551012 | 1.587255 | 1.629879 | 1.7034 | 1.789083 | 1.879882 | 1.9669 | 2.021789 | 2.071328 | 2.131424 | 2.187341 |
|  | 50 | 1.45078 | 1.487563 | 1.529192 | 1.565326 | 1.607511 | 1.67973 | 1.763512 | 1.852496 | 1.938567 | 1.993497 | 2.043618 | 2.105234 | 2.163478 |
|  | 60 | 1.430242 | 1.467939 | 1.510182 | 1.54654 | 1.588699 | 1.660371 | 1.743188 | 1.831388 | 1.917526 | 1.973159 | 2.024486 | 2.088442 | 2.149872 |
|  | 70 | 1.417014 | 1.454665 | 1.496869 | 1.533211 | 1.575378 | 1.647156 | 1.730295 | 1.819149 | 1.906309 | 1.962833 | 2.015156 | 2.080598 | 2.143725 |
|  | 80 | 1.394571 | 1.430998 | 1.472118 | 1.507753 | 1.54933 | 1.620587 | 1.703665 | 1.792799 | 1.880287 | 1.936939 | 1.989266 | 2.054512 | 2.117193 |
|  | 90 | 1.349261 | 1.383782 | 1.423021 | 1.457237 | 1.497378 | 1.566628 | 1.647884 | 1.735386 | 1.821312 | 1.876862 | 1.928055 | 1.991682 | 2.052551 |
| **L_entorhinal** | 3 | 2.209098 | 2.369608 | 2.538679 | 2.675785 | 2.825564 | 3.058625 | 3.296166 | 3.519988 | 3.715541 | 3.830089 | 3.927952 | 4.040022 | 4.138134 |
|  | 5 | 2.233764 | 2.387302 | 2.550572 | 2.684008 | 2.830734 | 3.060815 | 3.297282 | 3.521727 | 3.71902 | 3.835066 | 3.934473 | 4.048595 | 4.148744 |
|  | 10 | 2.289715 | 2.428522 | 2.579132 | 2.704346 | 2.844078 | 3.067174 | 3.301087 | 3.527144 | 3.728929 | 3.848878 | 3.952338 | 4.071894 | 4.177474 |
|  | 15 | 2.336906 | 2.464631 | 2.605353 | 2.723942 | 2.857895 | 3.075077 | 3.306853 | 3.534597 | 3.740879 | 3.864772 | 3.972363 | 4.09751 | 4.208736 |
|  | 20 | 2.368315 | 2.488706 | 2.622679 | 2.736612 | 2.866389 | 3.079108 | 3.309078 | 3.537917 | 3.747588 | 3.874558 | 3.985435 | 4.115097 | 4.230947 |
|  | 30 | 2.37524 | 2.490842 | 2.620181 | 2.730724 | 2.857223 | 3.065835 | 3.293008 | 3.520824 | 3.731171 | 3.859264 | 3.971542 | 4.103325 | 4.221494 |
|  | 40 | 2.361328 | 2.477727 | 2.607653 | 2.718448 | 2.844962 | 3.052975 | 3.278635 | 3.504262 | 3.712184 | 3.838615 | 3.94932 | 4.079119 | 4.195385 |
|  | 50 | 2.358652 | 2.475959 | 2.60672 | 2.718083 | 2.845083 | 3.053518 | 3.27911 | 3.504324 | 3.711733 | 3.837788 | 3.94812 | 4.077429 | 4.193205 |
|  | 60 | 2.333627 | 2.45686 | 2.593438 | 2.709131 | 2.840405 | 3.0544 | 3.284096 | 3.511753 | 3.720219 | 3.846385 | 3.956497 | 4.085179 | 4.200066 |
|  | 70 | 2.218737 | 2.359112 | 2.511636 | 2.638569 | 2.780303 | 3.006626 | 3.243756 | 3.473738 | 3.680566 | 3.804137 | 3.911055 | 4.03496 | 4.144674 |
|  | 80 | 1.94149 | 2.133774 | 2.329561 | 2.484113 | 2.649285 | 2.899768 | 3.148065 | 3.377898 | 3.577053 | 3.693063 | 3.791807 | 3.904477 | 4.002764 |
|  | 90 | 1.501325 | 1.798911 | 2.080887 | 2.286137 | 2.490415 | 2.776243 | 3.037841 | 3.265665 | 3.454416 | 3.561245 | 3.650568 | 3.750834 | 3.836968 |
| **L_fusiform** | 3 | 2.438677 | 2.546236 | 2.649149 | 2.726232 | 2.804938 | 2.917641 | 3.022081 | 3.114958 | 3.194343 | 3.240057 | 3.278623 | 3.322217 | 3.359873 |
|  | 5 | 2.426503 | 2.52574 | 2.622522 | 2.696064 | 2.771991 | 2.88203 | 2.985231 | 3.077816 | 3.157435 | 3.203469 | 3.242406 | 3.286524 | 3.32472 |
|  | 10 | 2.386098 | 2.468318 | 2.551869 | 2.617418 | 2.686833 | 2.790354 | 2.890347 | 2.982084 | 3.062239 | 3.109086 | 3.148987 | 3.194494 | 3.23414 |
|  | 15 | 2.348224 | 2.414468 | 2.484643 | 2.541643 | 2.603803 | 2.699818 | 2.796183 | 2.887405 | 2.969059 | 3.017593 | 3.05939 | 3.107567 | 3.149972 |
|  | 20 | 2.327912 | 2.38211 | 2.441522 | 2.491285 | 2.547086 | 2.636419 | 2.729904 | 2.821735 | 2.906496 | 2.958017 | 3.003065 | 3.055766 | 3.102838 |
|  | 30 | 2.261045 | 2.316916 | 2.377854 | 2.42867 | 2.485433 | 2.575886 | 2.670093 | 2.761899 | 2.845737 | 2.896314 | 2.940319 | 2.991561 | 3.037122 |
|  | 40 | 2.208745 | 2.265885 | 2.328023 | 2.379711 | 2.43733 | 2.528949 | 2.624194 | 2.71654 | 2.8002 | 2.850393 | 2.893912 | 2.944421 | 2.989191 |
|  | 50 | 2.193674 | 2.24679 | 2.305393 | 2.354802 | 2.410589 | 2.50084 | 2.596688 | 2.691213 | 2.777859 | 2.830308 | 2.876067 | 2.92951 | 2.977182 |
|  | 60 | 2.162369 | 2.216205 | 2.2756 | 2.325684 | 2.382253 | 2.473854 | 2.571308 | 2.667256 | 2.754772 | 2.807567 | 2.853533 | 2.907116 | 2.954828 |
|  | 70 | 2.116368 | 2.173198 | 2.23544 | 2.287582 | 2.346129 | 2.440244 | 2.539558 | 2.636251 | 2.723275 | 2.77528 | 2.820277 | 2.872421 | 2.918586 |
|  | 80 | 2.101499 | 2.151426 | 2.207323 | 2.255133 | 2.309894 | 2.400348 | 2.499094 | 2.598144 | 2.689415 | 2.744925 | 2.793545 | 2.850577 | 2.901692 |
|  | 90 | 2.087844 | 2.129222 | 2.176794 | 2.218568 | 2.267697 | 2.352007 | 2.448839 | 2.550965 | 2.649435 | 2.711489 | 2.767255 | 2.834419 | 2.89628 |
| **L_inferiorparietal** | 3 | 2.492339 | 2.569601 | 2.649285 | 2.712541 | 2.780161 | 2.882063 | 2.981507 | 3.073734 | 3.155231 | 3.203221 | 3.244287 | 3.291327 | 3.332475 |
|  | 5 | 2.4466 | 2.519665 | 2.595678 | 2.656461 | 2.721845 | 2.821129 | 2.918844 | 3.01002 | 3.090888 | 3.138636 | 3.179568 | 3.226538 | 3.267697 |
|  | 10 | 2.329546 | 2.393275 | 2.460882 | 2.515863 | 2.575886 | 2.668723 | 2.762041 | 2.850471 | 2.92967 | 2.976764 | 3.017332 | 3.064106 | 3.105286 |
|  | 15 | 2.217993 | 2.2739 | 2.334231 | 2.384052 | 2.4392 | 2.526037 | 2.615186 | 2.701034 | 2.778739 | 2.825308 | 2.865642 | 2.912395 | 2.953778 |
|  | 20 | 2.144175 | 2.194636 | 2.249849 | 2.296028 | 2.347759 | 2.430521 | 2.51715 | 2.601836 | 2.679263 | 2.726019 | 2.766731 | 2.814176 | 2.856397 |
|  | 30 | 2.072465 | 2.119238 | 2.171013 | 2.214805 | 2.2644 | 2.344977 | 2.431016 | 2.51621 | 2.594494 | 2.64197 | 2.683448 | 2.731959 | 2.775295 |
|  | 40 | 2.031783 | 2.078732 | 2.130793 | 2.174913 | 2.224993 | 2.306675 | 2.394413 | 2.481233 | 2.560442 | 2.608251 | 2.6499 | 2.698489 | 2.741794 |
|  | 50 | 1.991785 | 2.038903 | 2.091242 | 2.135686 | 2.18625 | 2.269045 | 2.358506 | 2.446978 | 2.527127 | 2.575276 | 2.617105 | 2.66578 | 2.709065 |
|  | 60 | 1.948682 | 1.994913 | 2.04651 | 2.090532 | 2.140865 | 2.223906 | 2.31457 | 2.404578 | 2.48588 | 2.534638 | 2.576961 | 2.626189 | 2.669952 |
|  | 70 | 1.912742 | 1.957208 | 2.007223 | 2.050227 | 2.099787 | 2.182509 | 2.274245 | 2.366163 | 2.449361 | 2.499349 | 2.542814 | 2.593465 | 2.638592 |
|  | 80 | 1.870688 | 1.911055 | 1.957116 | 1.997294 | 2.044273 | 2.124377 | 2.21578 | 2.3095 | 2.395626 | 2.447985 | 2.493901 | 2.547884 | 2.596418 |
|  | 90 | 1.809278 | 1.844264 | 1.884877 | 1.920932 | 1.963868 | 2.039134 | 2.128393 | 2.223225 | 2.312903 | 2.368669 | 2.418394 | 2.477877 | 2.532338 |
| **L_inferiortemporal** | 3 | 2.416212 | 2.541526 | 2.658998 | 2.745926 | 2.834362 | 2.96214 | 3.084515 | 3.194017 | 3.285956 | 3.339225 | 3.384853 | 3.437644 | 3.484674 |
|  | 5 | 2.412815 | 2.526478 | 2.63553 | 2.717675 | 2.802393 | 2.926609 | 3.047255 | 3.156297 | 3.248427 | 3.301973 | 3.347904 | 3.401096 | 3.448506 |
|  | 10 | 2.393415 | 2.484165 | 2.575725 | 2.647452 | 2.72376 | 2.839578 | 2.956005 | 3.063987 | 3.156802 | 3.211252 | 3.258181 | 3.312723 | 3.361461 |
|  | 15 | 2.369798 | 2.442404 | 2.51891 | 2.581055 | 2.649217 | 2.756495 | 2.86861 | 2.975969 | 3.070469 | 3.126719 | 3.175619 | 3.232876 | 3.284378 |
|  | 20 | 2.351647 | 2.410965 | 2.475656 | 2.529838 | 2.590925 | 2.690498 | 2.798847 | 2.906434 | 3.003978 | 3.0632 | 3.115334 | 3.1771 | 3.23328 |
|  | 30 | 2.313403 | 2.36641 | 2.42534 | 2.475565 | 2.5331 | 2.6288 | 2.735339 | 2.843186 | 2.94235 | 3.003043 | 3.056703 | 3.120488 | 3.178639 |
|  | 40 | 2.265494 | 2.319952 | 2.380381 | 2.431763 | 2.490454 | 2.587571 | 2.694745 | 2.801972 | 2.89923 | 2.958048 | 3.009568 | 3.070192 | 3.124854 |
|  | 50 | 2.23727 | 2.29301 | 2.354886 | 2.407495 | 2.467557 | 2.566794 | 2.675933 | 2.784536 | 2.882361 | 2.941144 | 2.992372 | 3.05231 | 3.106017 |
|  | 60 | 2.191307 | 2.248208 | 2.311454 | 2.365278 | 2.426762 | 2.528367 | 2.640017 | 2.750866 | 2.850365 | 2.909945 | 2.961717 | 3.022092 | 3.075988 |
|  | 70 | 2.163978 | 2.216739 | 2.276356 | 2.327888 | 2.387637 | 2.488392 | 2.601916 | 2.717409 | 2.823323 | 2.887716 | 2.944235 | 3.010789 | 3.070766 |
|  | 80 | 2.153029 | 2.20132 | 2.256873 | 2.305744 | 2.363407 | 2.463104 | 2.579188 | 2.701427 | 2.817249 | 2.889424 | 2.953879 | 3.03111 | 3.10195 |
|  | 90 | 2.103198 | 2.15039 | 2.205081 | 2.253552 | 2.311167 | 2.411862 | 2.530815 | 2.658037 | 2.780414 | 2.857566 | 2.927041 | 3.010997 | 3.088682 |
| **L_isthmuscingulate** | 3 | 2.517676 | 2.571316 | 2.633631 | 2.688987 | 2.754939 | 2.870597 | 3.007785 | 3.153909 | 3.292748 | 3.379385 | 3.456814 | 3.549643 | 3.634816 |
|  | 5 | 2.475823 | 2.529112 | 2.591037 | 2.646063 | 2.711649 | 2.826745 | 2.963413 | 3.108947 | 3.246999 | 3.33304 | 3.409873 | 3.501917 | 3.586304 |
|  | 10 | 2.372766 | 2.425182 | 2.486133 | 2.540337 | 2.605004 | 2.718684 | 2.854021 | 2.998039 | 3.134094 | 3.218629 | 3.293966 | 3.384039 | 3.466462 |
|  | 15 | 2.272548 | 2.324126 | 2.384138 | 2.437544 | 2.501313 | 2.613589 | 2.747568 | 2.890029 | 3.024081 | 3.107122 | 3.180981 | 3.269119 | 3.349619 |
|  | 20 | 2.185671 | 2.236608 | 2.295894 | 2.348677 | 2.411737 | 2.522879 | 2.655715 | 2.796816 | 2.929104 | 3.010827 | 3.08338 | 3.169803 | 3.248597 |
|  | 30 | 2.08162 | 2.132221 | 2.191069 | 2.243421 | 2.305921 | 2.415964 | 2.547304 | 2.686323 | 2.815957 | 2.895707 | 2.966302 | 3.050149 | 3.126366 |
|  | 40 | 2.021657 | 2.07272 | 2.131918 | 2.1844 | 2.246817 | 2.356035 | 2.485219 | 2.621273 | 2.74812 | 2.826123 | 2.89513 | 2.977018 | 3.051373 |
|  | 50 | 1.962316 | 2.015517 | 2.076898 | 2.131024 | 2.195002 | 2.3058 | 2.434853 | 2.570156 | 2.697231 | 2.775763 | 2.845427 | 2.928278 | 3.003644 |
|  | 60 | 1.893863 | 1.950992 | 2.016504 | 2.073877 | 2.141156 | 2.256093 | 2.387243 | 2.5242 | 2.654603 | 2.735944 | 2.808493 | 2.895183 | 2.974364 |
|  | 70 | 1.820424 | 1.881931 | 1.951983 | 2.012862 | 2.083625 | 2.202722 | 2.335566 | 2.473688 | 2.607198 | 2.691312 | 2.766772 | 2.85739 | 2.940523 |
|  | 80 | 1.754312 | 1.820282 | 1.894904 | 1.959263 | 2.033434 | 2.156501 | 2.290838 | 2.429838 | 2.56593 | 2.652372 | 2.730278 | 2.824197 | 2.910644 |
|  | 90 | 1.704355 | 1.775831 | 1.856107 | 1.924807 | 2.003298 | 2.131703 | 2.268902 | 2.410134 | 2.550056 | 2.639575 | 2.720577 | 2.818551 | 2.908976 |
| **L_lateraloccipital** | 3 | 2.171733 | 2.212244 | 2.257966 | 2.297405 | 2.342985 | 2.41934 | 2.504368 | 2.590275 | 2.668946 | 2.716616 | 2.758299 | 2.807139 | 2.850884 |
|  | 5 | 2.128572 | 2.168781 | 2.21416 | 2.253299 | 2.298525 | 2.374259 | 2.45854 | 2.543746 | 2.621923 | 2.669358 | 2.71087 | 2.759548 | 2.80318 |
|  | 10 | 2.02365 | 2.063083 | 2.107586 | 2.145961 | 2.190288 | 2.264448 | 2.346844 | 2.430282 | 2.507206 | 2.554039 | 2.595112 | 2.643373 | 2.686713 |
|  | 15 | 1.928805 | 1.967519 | 2.011208 | 2.048875 | 2.09237 | 2.165076 | 2.245729 | 2.327542 | 2.403337 | 2.449642 | 2.490341 | 2.538261 | 2.581379 |
|  | 20 | 1.867702 | 1.906149 | 1.949537 | 1.986939 | 2.030114 | 2.102222 | 2.182088 | 2.263242 | 2.338797 | 2.385116 | 2.425918 | 2.474059 | 2.517462 |
|  | 30 | 1.826015 | 1.864724 | 1.908403 | 1.946042 | 1.989456 | 2.061832 | 2.141726 | 2.223175 | 2.299747 | 2.347013 | 2.388834 | 2.438382 | 2.483232 |
|  | 40 | 1.809579 | 1.848232 | 1.891836 | 1.929389 | 1.972664 | 2.044657 | 2.123834 | 2.20479 | 2.281629 | 2.32938 | 2.371815 | 2.422294 | 2.468168 |
|  | 50 | 1.801254 | 1.8402 | 1.884126 | 1.921937 | 1.965473 | 2.037756 | 2.116962 | 2.198197 | 2.276054 | 2.324774 | 2.368261 | 2.420214 | 2.467619 |
|  | 60 | 1.769811 | 1.809124 | 1.853468 | 1.89163 | 1.935543 | 2.008336 | 2.08784 | 2.169667 | 2.248898 | 2.29884 | 2.343632 | 2.397387 | 2.446654 |
|  | 70 | 1.728426 | 1.768176 | 1.813029 | 1.85163 | 1.896033 | 1.96954 | 2.049589 | 2.1323 | 2.213247 | 2.264663 | 2.31101 | 2.366905 | 2.418379 |
|  | 80 | 1.68075 | 1.721162 | 1.766795 | 1.806081 | 1.851271 | 1.926016 | 2.007219 | 2.091496 | 2.174917 | 2.228339 | 2.276761 | 2.335468 | 2.389819 |
|  | 90 | 1.593271 | 1.633483 | 1.678935 | 1.718092 | 1.763144 | 1.837627 | 1.918385 | 2.002617 | 2.086995 | 2.141502 | 2.191199 | 2.251804 | 2.308236 |
| **L_lateralorbitofrontal** | 3 | 2.630192 | 2.746347 | 2.857114 | 2.940158 | 3.025459 | 3.149833 | 3.269632 | 3.376735 | 3.466062 | 3.517379 | 3.560998 | 3.61101 | 3.655101 |
|  | 5 | 2.591642 | 2.694766 | 2.795714 | 2.872924 | 2.953476 | 3.072969 | 3.190114 | 3.296358 | 3.385955 | 3.437805 | 3.482083 | 3.533073 | 3.578216 |
|  | 10 | 2.490154 | 2.569491 | 2.651256 | 2.71643 | 2.786764 | 2.895309 | 3.006355 | 3.110818 | 3.201555 | 3.255126 | 3.301474 | 3.355519 | 3.403953 |
|  | 15 | 2.395726 | 2.45975 | 2.52813 | 2.58435 | 2.646707 | 2.746322 | 2.852405 | 2.955984 | 3.048881 | 3.10499 | 3.154283 | 3.212633 | 3.265718 |
|  | 20 | 2.329494 | 2.38471 | 2.445013 | 2.495623 | 2.552849 | 2.646656 | 2.749802 | 2.85377 | 2.949774 | 3.009032 | 3.061886 | 3.125417 | 3.184129 |
|  | 30 | 2.234377 | 2.287253 | 2.345298 | 2.394254 | 2.449875 | 2.541668 | 2.643496 | 2.747103 | 2.843656 | 2.903682 | 2.957502 | 3.022545 | 3.082998 |
|  | 40 | 2.181001 | 2.234855 | 2.293758 | 2.343269 | 2.39934 | 2.491474 | 2.593135 | 2.696034 | 2.79148 | 2.850619 | 2.903519 | 2.967305 | 3.02645 |
|  | 50 | 2.158484 | 2.209528 | 2.265783 | 2.313413 | 2.367739 | 2.457905 | 2.558715 | 2.662196 | 2.75951 | 2.82046 | 2.875409 | 2.942209 | 3.004686 |
|  | 60 | 2.123784 | 2.175963 | 2.233346 | 2.281836 | 2.337038 | 2.428427 | 2.530293 | 2.634553 | 2.732354 | 2.7935 | 2.848562 | 2.91542 | 2.977881 |
|  | 70 | 2.064203 | 2.118566 | 2.178132 | 2.228292 | 2.285208 | 2.379015 | 2.482997 | 2.588841 | 2.687635 | 2.749177 | 2.804455 | 2.871407 | 2.933791 |
|  | 80 | 2.033177 | 2.0865 | 2.145401 | 2.195392 | 2.25255 | 2.347781 | 2.454858 | 2.565528 | 2.670391 | 2.736491 | 2.796379 | 2.869574 | 2.938433 |
|  | 90 | 2.007866 | 2.057482 | 2.113085 | 2.160952 | 2.216465 | 2.310878 | 2.420045 | 2.536415 | 2.65019 | 2.723748 | 2.791668 | 2.876379 | 2.957832 |
| **L_lingual** | 3 | 2.069481 | 2.111305 | 2.158487 | 2.199237 | 2.246483 | 2.326268 | 2.416559 | 2.509139 | 2.594942 | 2.647633 | 2.69426 | 2.749668 | 2.800112 |
|  | 5 | 2.019745 | 2.060746 | 2.106985 | 2.146912 | 2.193196 | 2.271346 | 2.359787 | 2.450492 | 2.534598 | 2.586272 | 2.632019 | 2.686409 | 2.735955 |
|  | 10 | 1.905172 | 1.944291 | 1.988374 | 2.026415 | 2.070491 | 2.144884 | 2.229072 | 2.315469 | 2.395676 | 2.44502 | 2.488755 | 2.540824 | 2.588328 |
|  | 15 | 1.811411 | 1.84905 | 1.891427 | 1.927969 | 1.970287 | 2.041677 | 2.12246 | 2.205414 | 2.282526 | 2.330033 | 2.372192 | 2.422458 | 2.468396 |
|  | 20 | 1.75238 | 1.789247 | 1.830713 | 1.866439 | 1.907786 | 1.977501 | 2.056378 | 2.137427 | 2.212871 | 2.259423 | 2.30079 | 2.35019 | 2.39542 |
|  | 30 | 1.702101 | 1.738859 | 1.780103 | 1.815571 | 1.856556 | 1.925569 | 2.003623 | 2.083941 | 2.158939 | 2.205378 | 2.246774 | 2.296391 | 2.342013 |
|  | 40 | 1.666882 | 1.703903 | 1.745322 | 1.780855 | 1.821841 | 1.890741 | 1.968628 | 2.048911 | 2.124161 | 2.170956 | 2.212828 | 2.26324 | 2.309834 |
|  | 50 | 1.634011 | 1.671425 | 1.713136 | 1.748816 | 1.789879 | 1.858764 | 1.936585 | 2.016961 | 2.092643 | 2.139952 | 2.182479 | 2.233959 | 2.281844 |
|  | 60 | 1.599796 | 1.637676 | 1.679725 | 1.715567 | 1.756702 | 1.825533 | 1.903224 | 1.983657 | 2.059813 | 2.10772 | 2.151027 | 2.203803 | 2.253273 |
|  | 70 | 1.558938 | 1.597264 | 1.639585 | 1.675501 | 1.71658 | 1.785097 | 1.862341 | 1.942531 | 2.018963 | 2.067412 | 2.111509 | 2.165684 | 2.216949 |
|  | 80 | 1.51354 | 1.552394 | 1.595019 | 1.630999 | 1.671971 | 1.740026 | 1.816611 | 1.896359 | 1.972963 | 2.021964 | 2.066931 | 2.122718 | 2.176116 |
|  | 90 | 1.476821 | 1.516704 | 1.560108 | 1.596495 | 1.637705 | 1.705784 | 1.782191 | 1.862008 | 1.939376 | 1.989403 | 2.035763 | 2.093961 | 2.150441 |
| **L_medialorbitofrontal** | 3 | 2.556011 | 2.618736 | 2.687132 | 2.744526 | 2.809515 | 2.916593 | 3.03581 | 3.158524 | 3.275045 | 3.348895 | 3.416199 | 3.49909 | 3.577808 |
|  | 5 | 2.47608 | 2.53723 | 2.603969 | 2.660017 | 2.723528 | 2.82827 | 2.945009 | 3.065279 | 3.179555 | 3.252009 | 3.318056 | 3.399412 | 3.476684 |
|  | 10 | 2.304397 | 2.36221 | 2.425446 | 2.478661 | 2.539073 | 2.638939 | 2.750544 | 2.865788 | 2.975476 | 3.045092 | 3.108588 | 3.186842 | 3.261198 |
|  | 15 | 2.184716 | 2.24037 | 2.301379 | 2.352823 | 2.411332 | 2.508284 | 2.616928 | 2.729381 | 2.836607 | 2.904737 | 2.96692 | 3.0436 | 3.116499 |
|  | 20 | 2.10973 | 2.16419 | 2.224015 | 2.274558 | 2.332146 | 2.427792 | 2.535261 | 2.646764 | 2.753287 | 2.821053 | 2.88295 | 2.95933 | 3.031991 |
|  | 30 | 2.027735 | 2.080744 | 2.139182 | 2.188716 | 2.245329 | 2.339741 | 2.446338 | 2.557431 | 2.663957 | 2.731896 | 2.794054 | 2.870878 | 2.944075 |
|  | 40 | 1.990795 | 2.042887 | 2.100454 | 2.149365 | 2.205392 | 2.299116 | 2.405356 | 2.516528 | 2.623536 | 2.691981 | 2.754734 | 2.832461 | 2.906688 |
|  | 50 | 1.968225 | 2.019675 | 2.076606 | 2.125041 | 2.1806 | 2.273743 | 2.379663 | 2.490928 | 2.598486 | 2.667534 | 2.731021 | 2.809904 | 2.885496 |
|  | 60 | 1.937735 | 1.988687 | 2.045107 | 2.093145 | 2.148303 | 2.240937 | 2.346593 | 2.458046 | 2.566326 | 2.636156 | 2.700596 | 2.780996 | 2.858398 |
|  | 70 | 1.919802 | 1.971193 | 2.028143 | 2.076678 | 2.132468 | 2.226355 | 2.333819 | 2.447737 | 2.559077 | 2.631272 | 2.698191 | 2.782098 | 2.863331 |
|  | 80 | 1.888765 | 1.94075 | 1.998418 | 2.047623 | 2.104264 | 2.199819 | 2.309654 | 2.426759 | 2.542017 | 2.61723 | 2.687305 | 2.775683 | 2.861808 |
|  | 90 | 1.860959 | 1.913638 | 1.972139 | 2.022118 | 2.079736 | 2.177195 | 2.28972 | 2.410436 | 2.530145 | 2.608801 | 2.682494 | 2.776027 | 2.867833 |
| **L_middletemporal** | 3 | 2.540565 | 2.688508 | 2.822847 | 2.919641 | 3.015803 | 3.150073 | 3.272208 | 3.377633 | 3.464163 | 3.512756 | 3.553154 | 3.598239 | 3.636741 |
|  | 5 | 2.547843 | 2.678793 | 2.801315 | 2.891689 | 2.983068 | 3.112999 | 3.233192 | 3.338395 | 3.425765 | 3.475192 | 3.516467 | 3.562715 | 3.602354 |
|  | 10 | 2.546653 | 2.645232 | 2.744073 | 2.820878 | 2.901674 | 3.021493 | 3.136903 | 3.241537 | 3.331149 | 3.38286 | 3.426571 | 3.476092 | 3.518976 |
|  | 15 | 2.528759 | 2.604821 | 2.685274 | 2.750559 | 2.821726 | 2.931687 | 3.042219 | 3.146528 | 3.239203 | 3.294023 | 3.341098 | 3.39522 | 3.442746 |
|  | 20 | 2.504543 | 2.565659 | 2.632752 | 2.689022 | 2.752196 | 2.85348 | 2.959642 | 3.064186 | 3.160992 | 3.219971 | 3.27162 | 3.332134 | 3.386263 |
|  | 30 | 2.430661 | 2.485546 | 2.546719 | 2.59875 | 2.65793 | 2.754427 | 2.857584 | 2.961722 | 3.060943 | 3.122682 | 3.177532 | 3.242714 | 3.301848 |
|  | 40 | 2.362436 | 2.422799 | 2.489101 | 2.544699 | 2.607058 | 2.706737 | 2.810568 | 2.913511 | 3.010691 | 3.07068 | 3.123649 | 3.186185 | 3.242526 |
|  | 50 | 2.319542 | 2.381603 | 2.449565 | 2.506371 | 2.569859 | 2.670762 | 2.774964 | 2.878229 | 2.976463 | 3.037401 | 3.091362 | 3.155229 | 3.212899 |
|  | 60 | 2.250707 | 2.315086 | 2.385277 | 2.443731 | 2.508865 | 2.612068 | 2.718409 | 2.822924 | 2.92097 | 2.981197 | 3.034189 | 3.096529 | 3.152489 |
|  | 70 | 2.195188 | 2.255438 | 2.322038 | 2.378285 | 2.441892 | 2.544993 | 2.654722 | 2.763321 | 2.863335 | 2.924079 | 2.977195 | 3.039364 | 3.094938 |
|  | 80 | 2.128966 | 2.189206 | 2.256054 | 2.312812 | 2.377447 | 2.483633 | 2.599125 | 2.712322 | 2.812485 | 2.871695 | 2.922623 | 2.981354 | 3.033143 |
|  | 90 | 2.070378 | 2.130002 | 2.19659 | 2.253562 | 2.319049 | 2.42844 | 2.550392 | 2.669016 | 2.770023 | 2.828184 | 2.877432 | 2.933445 | 2.982225 |
| **L_parahippocampal** | 3 | 2.230403 | 2.32555 | 2.433144 | 2.526471 | 2.635407 | 2.822045 | 3.037481 | 3.247534 | 3.421147 | 3.519046 | 3.600958 | 3.693184 | 3.772799 |
|  | 5 | 2.208479 | 2.302791 | 2.409592 | 2.502356 | 2.610771 | 2.796834 | 3.012065 | 3.222392 | 3.396589 | 3.494954 | 3.577327 | 3.670147 | 3.750337 |
|  | 10 | 2.154338 | 2.246528 | 2.351306 | 2.442622 | 2.549689 | 2.734245 | 2.948915 | 3.15992 | 3.335614 | 3.435181 | 3.51875 | 3.61312 | 3.694814 |
|  | 15 | 2.100325 | 2.190329 | 2.293001 | 2.382792 | 2.48842 | 2.671324 | 2.885302 | 3.09692 | 3.274117 | 3.37492 | 3.459732 | 3.555721 | 3.638999 |
|  | 20 | 2.049096 | 2.13704 | 2.237721 | 2.326072 | 2.430346 | 2.611725 | 2.825151 | 3.037535 | 3.216398 | 3.318546 | 3.404705 | 3.502448 | 3.587438 |
|  | 30 | 1.973321 | 2.058119 | 2.155797 | 2.242016 | 2.344356 | 2.523786 | 2.737097 | 2.951767 | 3.134482 | 3.239593 | 3.328667 | 3.430164 | 3.518794 |
|  | 40 | 1.918217 | 2.001046 | 2.096778 | 2.181555 | 2.282498 | 2.460257 | 2.672791 | 2.888127 | 3.072659 | 3.179319 | 3.269982 | 3.373587 | 3.464309 |
|  | 50 | 1.880444 | 1.9643 | 2.061115 | 2.146753 | 2.248604 | 2.427639 | 2.641178 | 2.857193 | 3.042221 | 3.149136 | 3.239994 | 3.343797 | 3.434667 |
|  | 60 | 1.84351 | 1.930917 | 2.03138 | 2.119856 | 2.22462 | 2.407611 | 2.624049 | 2.841347 | 3.026421 | 3.132951 | 3.22325 | 3.326158 | 3.416027 |
|  | 70 | 1.786847 | 1.878484 | 1.983174 | 2.07484 | 2.182764 | 2.369756 | 2.588632 | 2.806307 | 2.990375 | 3.095813 | 3.184904 | 3.286129 | 3.374265 |
|  | 80 | 1.709817 | 1.80627 | 1.915607 | 2.010638 | 2.121738 | 2.312358 | 2.532748 | 2.749517 | 2.931293 | 3.034841 | 3.122021 | 3.220737 | 3.306405 |
|  | 90 | 1.625217 | 1.727949 | 1.843197 | 1.942415 | 2.057378 | 2.25227 | 2.474308 | 2.689883 | 2.868911 | 2.970246 | 3.055218 | 3.151067 | 3.233942 |
| **L_paracentral** | 3 | 2.345259 | 2.400971 | 2.463306 | 2.516577 | 2.577522 | 2.677951 | 2.787157 | 2.896927 | 2.998882 | 3.061229 | 3.116027 | 3.180508 | 3.238469 |
|  | 5 | 2.307584 | 2.362433 | 2.423862 | 2.476405 | 2.536571 | 2.635841 | 2.743956 | 2.852791 | 2.954006 | 3.015962 | 3.070453 | 3.134616 | 3.192333 |
|  | 10 | 2.208085 | 2.260646 | 2.319648 | 2.370232 | 2.428282 | 2.524359 | 2.629416 | 2.73557 | 2.834613 | 2.89539 | 2.948938 | 3.012103 | 3.069025 |
|  | 15 | 2.105224 | 2.155423 | 2.211904 | 2.260433 | 2.316247 | 2.408905 | 2.510625 | 2.613793 | 2.710364 | 2.769771 | 2.822205 | 2.884169 | 2.94011 |
|  | 20 | 2.030361 | 2.079219 | 2.134261 | 2.181614 | 2.236141 | 2.326825 | 2.426606 | 2.528016 | 2.623099 | 2.681668 | 2.73341 | 2.794613 | 2.849922 |
|  | 30 | 1.940465 | 1.990411 | 2.046457 | 2.094489 | 2.149597 | 2.24078 | 2.340469 | 2.441063 | 2.534689 | 2.592041 | 2.642511 | 2.701979 | 2.755509 |
|  | 40 | 1.881324 | 1.934448 | 1.993613 | 2.043955 | 2.101317 | 2.195345 | 2.296953 | 2.398235 | 2.491383 | 2.547935 | 2.597398 | 2.655325 | 2.707151 |
|  | 50 | 1.822403 | 1.880521 | 1.944466 | 1.998264 | 2.058917 | 2.156952 | 2.261099 | 2.363154 | 2.455526 | 2.510958 | 2.559059 | 2.614958 | 2.664587 |
|  | 60 | 1.77558 | 1.835703 | 1.901641 | 1.956955 | 2.019152 | 2.119345 | 2.225373 | 2.328835 | 2.422086 | 2.477878 | 2.526195 | 2.582238 | 2.631904 |
|  | 70 | 1.747078 | 1.802093 | 1.86342 | 1.915653 | 1.975232 | 2.073055 | 2.179013 | 2.284698 | 2.381786 | 2.44069 | 2.49219 | 2.552486 | 2.60642 |
|  | 80 | 1.730431 | 1.777561 | 1.831371 | 1.878282 | 1.933023 | 2.025822 | 2.130549 | 2.239442 | 2.343439 | 2.408432 | 2.466455 | 2.535834 | 2.599233 |
|  | 90 | 1.70803 | 1.749091 | 1.796848 | 1.839272 | 1.889745 | 1.977807 | 2.081201 | 2.193482 | 2.305532 | 2.378102 | 2.444632 | 2.526437 | 2.603435 |
| **L_parsopercularis** | 3 | 2.426956 | 2.525592 | 2.623193 | 2.697932 | 2.77526 | 2.886751 | 2.989463 | 3.083878 | 3.170369 | 3.222386 | 3.267407 | 3.319455 | 3.365333 |
|  | 5 | 2.421748 | 2.510342 | 2.600131 | 2.670233 | 2.743944 | 2.852304 | 2.954352 | 3.049239 | 3.136338 | 3.188839 | 3.234367 | 3.287115 | 3.333717 |
|  | 10 | 2.396359 | 2.465935 | 2.539854 | 2.59997 | 2.665503 | 2.766369 | 2.866705 | 2.962813 | 3.051562 | 3.105404 | 3.152352 | 3.207075 | 3.255738 |
|  | 15 | 2.358629 | 2.414498 | 2.476013 | 2.52775 | 2.585989 | 2.679701 | 2.77837 | 2.875895 | 2.966447 | 3.02173 | 3.070214 | 3.127104 | 3.178069 |
|  | 20 | 2.316498 | 2.363142 | 2.415825 | 2.46128 | 2.513798 | 2.601643 | 2.699142 | 2.7983 | 2.890571 | 2.947123 | 2.996935 | 3.055704 | 3.108692 |
|  | 30 | 2.219475 | 2.262794 | 2.312153 | 2.355145 | 2.405341 | 2.490743 | 2.58791 | 2.687333 | 2.778611 | 2.834059 | 2.882657 | 2.939758 | 2.991064 |
|  | 40 | 2.12697 | 2.175896 | 2.230669 | 2.277516 | 2.331177 | 2.419832 | 2.516657 | 2.613713 | 2.702989 | 2.757217 | 2.804677 | 2.860309 | 2.910133 |
|  | 50 | 2.061573 | 2.113531 | 2.17121 | 2.220083 | 2.275467 | 2.36532 | 2.460777 | 2.556531 | 2.647208 | 2.703366 | 2.753091 | 2.811985 | 2.865231 |
|  | 60 | 2.013068 | 2.066711 | 2.12607 | 2.176169 | 2.232669 | 2.32353 | 2.41867 | 2.514426 | 2.607043 | 2.665221 | 2.717191 | 2.77924 | 2.835761 |
|  | 70 | 1.967207 | 2.021765 | 2.082088 | 2.132963 | 2.190299 | 2.282422 | 2.378782 | 2.475638 | 2.569169 | 2.627851 | 2.680227 | 2.742709 | 2.799577 |
|  | 80 | 1.945987 | 1.99586 | 2.051896 | 2.099964 | 2.155138 | 2.246416 | 2.346071 | 2.447206 | 2.542514 | 2.601413 | 2.653544 | 2.715304 | 2.771192 |
|  | 90 | 1.933268 | 1.977303 | 2.027803 | 2.072097 | 2.12421 | 2.213988 | 2.317979 | 2.42486 | 2.522033 | 2.580666 | 2.631857 | 2.691807 | 2.745521 |
| **L_parsorbitalis** | 3 | 2.531465 | 2.652035 | 2.780364 | 2.88543 | 3.001316 | 3.184351 | 3.375004 | 3.555021 | 3.709762 | 3.799633 | 3.876151 | 3.963641 | 4.040237 |
|  | 5 | 2.522573 | 2.630552 | 2.746571 | 2.842329 | 2.948691 | 3.118149 | 3.296403 | 3.46621 | 3.613262 | 3.699109 | 3.77245 | 3.856582 | 3.930477 |
|  | 10 | 2.462514 | 2.549287 | 2.644043 | 2.723386 | 2.812664 | 2.957292 | 3.112458 | 3.263047 | 3.395592 | 3.473874 | 3.541279 | 3.6192 | 3.688167 |
|  | 15 | 2.350735 | 2.429439 | 2.516226 | 2.589557 | 2.672779 | 2.809164 | 2.957621 | 3.103816 | 3.234237 | 3.312045 | 3.379513 | 3.458057 | 3.528077 |
|  | 20 | 2.235948 | 2.31348 | 2.399634 | 2.472965 | 2.556783 | 2.695522 | 2.848518 | 3.001253 | 3.139306 | 3.222501 | 3.295159 | 3.380366 | 3.456902 |
|  | 30 | 2.190817 | 2.257607 | 2.332325 | 2.396359 | 2.470072 | 2.593405 | 2.731523 | 2.871895 | 3.001175 | 3.080292 | 3.150186 | 3.233151 | 3.30864 |
|  | 40 | 2.116404 | 2.18637 | 2.264376 | 2.331054 | 2.407669 | 2.535697 | 2.679189 | 2.825612 | 2.961408 | 3.045122 | 3.119538 | 3.208511 | 3.290135 |
|  | 50 | 2.062334 | 2.135474 | 2.215714 | 2.283315 | 2.359987 | 2.486067 | 2.624979 | 2.764914 | 2.893723 | 2.972911 | 3.043279 | 3.127485 | 3.204897 |
|  | 60 | 1.949795 | 2.033653 | 2.123835 | 2.198491 | 2.281882 | 2.416546 | 2.562271 | 2.707375 | 2.840367 | 2.922196 | 2.99511 | 3.082754 | 3.163824 |
|  | 70 | 1.908166 | 1.990117 | 2.078044 | 2.150749 | 2.231988 | 2.363631 | 2.507575 | 2.653659 | 2.791123 | 2.877876 | 2.956806 | 3.053943 | 3.146197 |
|  | 80 | 1.911007 | 1.985697 | 2.066384 | 2.133632 | 2.209492 | 2.334606 | 2.475837 | 2.625964 | 2.775691 | 2.875412 | 2.970218 | 3.092853 | 3.216064 |
|  | 90 | 2.033707 | 2.093099 | 2.156414 | 2.208637 | 2.267145 | 2.363391 | 2.473241 | 2.593913 | 2.721026 | 2.810752 | 2.900607 | 3.024446 | 3.15877 |
| **L_parstriangularis** | 3 | 2.514795 | 2.57609 | 2.643253 | 2.699745 | 2.763678 | 2.868377 | 2.982877 | 3.097014 | 3.200748 | 3.263796 | 3.319318 | 3.385103 | 3.444923 |
|  | 5 | 2.466646 | 2.527027 | 2.593277 | 2.649066 | 2.71227 | 2.815911 | 2.92941 | 3.04267 | 3.145672 | 3.208291 | 3.26344 | 3.32878 | 3.388188 |
|  | 10 | 2.348794 | 2.40692 | 2.470904 | 2.524942 | 2.586325 | 2.687307 | 2.798284 | 2.909329 | 3.010492 | 3.07204 | 3.12626 | 3.190501 | 3.2489 |
|  | 15 | 2.235064 | 2.291003 | 2.352777 | 2.405101 | 2.464692 | 2.563047 | 2.671527 | 2.780385 | 2.879745 | 2.940254 | 2.993578 | 3.05677 | 3.114215 |
|  | 20 | 2.148751 | 2.20314 | 2.263394 | 2.314577 | 2.373021 | 2.469803 | 2.57694 | 2.684776 | 2.783409 | 2.843542 | 2.896564 | 2.959418 | 3.016566 |
|  | 30 | 2.048298 | 2.100821 | 2.159368 | 2.209382 | 2.266786 | 2.362478 | 2.469206 | 2.57732 | 2.676672 | 2.737407 | 2.791037 | 2.854682 | 2.912593 |
|  | 40 | 1.989907 | 2.04075 | 2.097752 | 2.146707 | 2.203176 | 2.297921 | 2.40439 | 2.51296 | 2.613246 | 2.674746 | 2.72915 | 2.793814 | 2.852727 |
|  | 50 | 1.942445 | 1.991646 | 2.04711 | 2.094988 | 2.150478 | 2.244172 | 2.350255 | 2.459177 | 2.560349 | 2.622616 | 2.67782 | 2.743564 | 2.803564 |
|  | 60 | 1.890089 | 1.937518 | 1.991262 | 2.03788 | 2.092162 | 2.184391 | 2.289607 | 2.398414 | 2.500087 | 2.562917 | 2.618765 | 2.685434 | 2.746415 |
|  | 70 | 1.85137 | 1.897245 | 1.949479 | 1.994998 | 2.048236 | 2.139241 | 2.243846 | 2.352811 | 2.45528 | 2.518883 | 2.575582 | 2.643456 | 2.705705 |
|  | 80 | 1.835052 | 1.87994 | 1.931287 | 1.976232 | 2.029024 | 2.119807 | 2.224946 | 2.335294 | 2.439765 | 2.504925 | 2.563203 | 2.633189 | 2.697576 |
|  | 90 | 1.817218 | 1.861152 | 1.91163 | 1.956003 | 2.008344 | 2.098885 | 2.204548 | 2.316314 | 2.422893 | 2.489722 | 2.549713 | 2.622018 | 2.688783 |
| **L_pericalcarine** | 3 | 1.570709 | 1.601103 | 1.637002 | 1.66947 | 1.708926 | 1.780361 | 1.869364 | 1.971026 | 2.07627 | 2.146978 | 2.213917 | 2.29942 | 2.383574 |
|  | 5 | 1.533097 | 1.563367 | 1.59905 | 1.631261 | 1.670326 | 1.740846 | 1.828343 | 1.927802 | 2.030231 | 2.098739 | 2.163365 | 2.245581 | 2.326131 |
|  | 10 | 1.447637 | 1.477806 | 1.513175 | 1.544928 | 1.583227 | 1.65181 | 1.735981 | 1.830501 | 1.926625 | 1.990252 | 2.049791 | 2.12487 | 2.197704 |
|  | 15 | 1.379939 | 1.410461 | 1.446005 | 1.477708 | 1.515707 | 1.583145 | 1.664959 | 1.755738 | 1.847006 | 1.906887 | 1.962562 | 2.032293 | 2.099453 |
|  | 20 | 1.335836 | 1.367255 | 1.403566 | 1.435724 | 1.474004 | 1.541318 | 1.622076 | 1.710747 | 1.799111 | 1.856739 | 1.910111 | 1.976709 | 2.040621 |
|  | 30 | 1.29037 | 1.323763 | 1.361845 | 1.395163 | 1.434393 | 1.50245 | 1.582964 | 1.670526 | 1.757443 | 1.814152 | 1.866789 | 1.932726 | 1.996355 |
|  | 40 | 1.259715 | 1.293431 | 1.331693 | 1.365033 | 1.404165 | 1.471857 | 1.551886 | 1.639229 | 1.726609 | 1.784133 | 1.837962 | 1.906057 | 1.97254 |
|  | 50 | 1.243647 | 1.27653 | 1.31399 | 1.346751 | 1.385339 | 1.452416 | 1.532204 | 1.619815 | 1.70795 | 1.76621 | 1.820887 | 1.890265 | 1.958217 |
|  | 60 | 1.231129 | 1.263028 | 1.299597 | 1.331763 | 1.369844 | 1.436451 | 1.516167 | 1.604021 | 1.692466 | 1.750854 | 1.805544 | 1.874738 | 1.942248 |
|  | 70 | 1.211173 | 1.241806 | 1.277193 | 1.308529 | 1.345846 | 1.411564 | 1.490694 | 1.57811 | 1.66595 | 1.723697 | 1.777539 | 1.845245 | 1.9108 |
|  | 80 | 1.196088 | 1.225276 | 1.259276 | 1.289614 | 1.325996 | 1.390638 | 1.469219 | 1.556663 | 1.644891 | 1.702964 | 1.7571 | 1.825102 | 1.89081 |
|  | 90 | 1.177836 | 1.205452 | 1.237868 | 1.267003 | 1.302188 | 1.365308 | 1.442961 | 1.530395 | 1.61954 | 1.678663 | 1.734063 | 1.804017 | 1.87197 |
| **L_postcentral** | 3 | 2.068496 | 2.11042 | 2.156931 | 2.19653 | 2.24189 | 2.317477 | 2.40211 | 2.488682 | 2.569426 | 2.619524 | 2.664312 | 2.718223 | 2.768068 |
|  | 5 | 2.031497 | 2.07299 | 2.119008 | 2.158174 | 2.203026 | 2.277743 | 2.361381 | 2.446929 | 2.526725 | 2.576243 | 2.620523 | 2.673834 | 2.723139 |
|  | 10 | 1.939309 | 1.979698 | 2.02445 | 2.062508 | 2.10606 | 2.178558 | 2.259656 | 2.342588 | 2.419964 | 2.468003 | 2.51098 | 2.562756 | 2.610678 |
|  | 15 | 1.856912 | 1.896354 | 1.940014 | 1.977112 | 2.019537 | 2.090099 | 2.168979 | 2.249623 | 2.324884 | 2.371632 | 2.413476 | 2.463919 | 2.510644 |
|  | 20 | 1.803901 | 1.842987 | 1.886209 | 1.922904 | 1.964836 | 2.034521 | 2.112366 | 2.191933 | 2.266206 | 2.312365 | 2.353702 | 2.403568 | 2.449795 |
|  | 30 | 1.744846 | 1.784214 | 1.827657 | 1.864471 | 1.906475 | 1.976157 | 2.053881 | 2.133285 | 2.207445 | 2.253582 | 2.294946 | 2.344911 | 2.39131 |
|  | 40 | 1.705654 | 1.745767 | 1.789932 | 1.827286 | 1.869834 | 1.940286 | 2.018745 | 2.098853 | 2.173715 | 2.22034 | 2.26219 | 2.312818 | 2.359915 |
|  | 50 | 1.668046 | 1.708978 | 1.753938 | 1.791884 | 1.83503 | 1.906329 | 1.985594 | 2.066479 | 2.14211 | 2.189271 | 2.231653 | 2.283006 | 2.33087 |
|  | 60 | 1.627145 | 1.668856 | 1.714553 | 1.753035 | 1.796706 | 1.868719 | 1.948631 | 2.030124 | 2.106372 | 2.153977 | 2.196815 | 2.248807 | 2.297364 |
|  | 70 | 1.586723 | 1.629263 | 1.675739 | 1.714784 | 1.759004 | 1.831753 | 1.912325 | 1.994432 | 2.071307 | 2.119368 | 2.162678 | 2.215335 | 2.264619 |
|  | 80 | 1.538196 | 1.581378 | 1.628417 | 1.667834 | 1.712377 | 1.785477 | 1.866267 | 1.948536 | 2.025618 | 2.073878 | 2.11743 | 2.170483 | 2.220252 |
|  | 90 | 1.481563 | 1.525171 | 1.572526 | 1.612098 | 1.656712 | 1.729736 | 1.810259 | 1.892192 | 1.969017 | 2.01719 | 2.060733 | 2.11388 | 2.16386 |
| **L_posteriorcingulate** | 3 | 2.641938 | 2.693659 | 2.751887 | 2.802117 | 2.860335 | 2.958757 | 3.070613 | 3.186235 | 3.294609 | 3.361887 | 3.421957 | 3.494064 | 3.560458 |
|  | 5 | 2.597263 | 2.648514 | 2.706212 | 2.755982 | 2.813664 | 2.911167 | 3.021949 | 3.136421 | 3.243667 | 3.310218 | 3.369619 | 3.440897 | 3.506503 |
|  | 10 | 2.484791 | 2.534814 | 2.591125 | 2.639694 | 2.695973 | 2.791071 | 2.89905 | 3.010518 | 3.114831 | 3.179494 | 3.237163 | 3.306301 | 3.369872 |
|  | 15 | 2.373934 | 2.422705 | 2.477601 | 2.524943 | 2.579789 | 2.672429 | 2.777543 | 2.885947 | 2.987273 | 3.05002 | 3.105932 | 3.172902 | 3.234419 |
|  | 20 | 2.281319 | 2.32913 | 2.382938 | 2.429334 | 2.483071 | 2.573797 | 2.676661 | 2.782635 | 2.881568 | 2.942766 | 2.997252 | 3.062454 | 3.122286 |
|  | 30 | 2.169091 | 2.216222 | 2.269243 | 2.314935 | 2.367824 | 2.457024 | 2.557978 | 2.661744 | 2.75836 | 2.817988 | 2.870981 | 2.934274 | 2.992231 |
|  | 40 | 2.108026 | 2.155465 | 2.208803 | 2.254738 | 2.307869 | 2.397362 | 2.498449 | 2.602092 | 2.698326 | 2.757575 | 2.810136 | 2.872788 | 2.930036 |
|  | 50 | 2.058412 | 2.106664 | 2.16088 | 2.207537 | 2.26146 | 2.352167 | 2.454417 | 2.558988 | 2.655811 | 2.715283 | 2.767945 | 2.830596 | 2.887721 |
|  | 60 | 2.019362 | 2.069073 | 2.124889 | 2.172885 | 2.228308 | 2.32141 | 2.42614 | 2.532971 | 2.63161 | 2.692054 | 2.745481 | 2.80892 | 2.866645 |
|  | 70 | 1.978013 | 2.029437 | 2.087127 | 2.136692 | 2.193869 | 2.289773 | 2.397408 | 2.506907 | 2.607714 | 2.669337 | 2.723708 | 2.788142 | 2.846651 |
|  | 80 | 1.947285 | 2.00095 | 2.061092 | 2.112708 | 2.172184 | 2.27177 | 2.383259 | 2.496344 | 2.600134 | 2.66342 | 2.719154 | 2.785071 | 2.844802 |
|  | 90 | 1.928049 | 1.984548 | 2.047786 | 2.101989 | 2.164364 | 2.268597 | 2.384962 | 2.502618 | 2.610249 | 2.675704 | 2.733234 | 2.801139 | 2.862536 |
| **L_precentral** | 3 | 2.215852 | 2.307635 | 2.393975 | 2.458002 | 2.523229 | 2.617657 | 2.708441 | 2.790232 | 2.859574 | 2.900134 | 2.935147 | 2.976012 | 3.012775 |
|  | 5 | 2.223767 | 2.309963 | 2.392371 | 2.454242 | 2.51788 | 2.610982 | 2.701446 | 2.783639 | 2.853763 | 2.894945 | 2.930584 | 2.972277 | 3.009865 |
|  | 10 | 2.235904 | 2.310414 | 2.38422 | 2.441184 | 2.501076 | 2.590903 | 2.680469 | 2.763585 | 2.835658 | 2.878436 | 2.915703 | 2.959574 | 2.999361 |
|  | 15 | 2.232481 | 2.297874 | 2.364498 | 2.417114 | 2.473502 | 2.560006 | 2.648399 | 2.732168 | 2.806036 | 2.850375 | 2.889283 | 2.935403 | 2.977511 |
|  | 20 | 2.211455 | 2.270563 | 2.331971 | 2.381282 | 2.434894 | 2.518603 | 2.605853 | 2.690006 | 2.765297 | 2.81094 | 2.851253 | 2.899338 | 2.943513 |
|  | 30 | 2.137284 | 2.194599 | 2.25459 | 2.303071 | 2.356074 | 2.439383 | 2.526829 | 2.611631 | 2.687774 | 2.734015 | 2.774891 | 2.823671 | 2.868494 |
|  | 40 | 2.055954 | 2.11856 | 2.183311 | 2.235098 | 2.291191 | 2.378305 | 2.468413 | 2.554521 | 2.630771 | 2.676586 | 2.716777 | 2.764363 | 2.807731 |
|  | 50 | 1.991578 | 2.05868 | 2.127587 | 2.182364 | 2.241382 | 2.332419 | 2.425802 | 2.514284 | 2.591993 | 2.638382 | 2.678884 | 2.726603 | 2.769867 |
|  | 60 | 1.937351 | 2.003981 | 2.072883 | 2.127982 | 2.187643 | 2.280215 | 2.375751 | 2.466687 | 2.546777 | 2.594652 | 2.636474 | 2.685759 | 2.730442 |
|  | 70 | 1.88236 | 1.944783 | 2.010347 | 2.06348 | 2.121682 | 2.213277 | 2.309299 | 2.401938 | 2.484396 | 2.534024 | 2.577562 | 2.629069 | 2.675937 |
|  | 80 | 1.816631 | 1.876348 | 1.939694 | 1.991474 | 2.048625 | 2.139415 | 2.23559 | 2.329205 | 2.413101 | 2.463811 | 2.508411 | 2.561297 | 2.609516 |
|  | 90 | 1.743043 | 1.802765 | 1.866237 | 1.918205 | 1.975642 | 2.067017 | 2.163924 | 2.258272 | 2.342767 | 2.393784 | 2.438608 | 2.491689 | 2.540014 |
| **L_precuneus** | 3 | 2.523557 | 2.576882 | 2.634597 | 2.682623 | 2.73648 | 2.823749 | 2.918263 | 3.012006 | 3.097213 | 3.149162 | 3.195083 | 3.249768 | 3.299812 |
|  | 5 | 2.465 | 2.517442 | 2.574232 | 2.621511 | 2.67455 | 2.760534 | 2.853695 | 2.946113 | 3.030113 | 3.081317 | 3.126571 | 3.180449 | 3.22974 |
|  | 10 | 2.318309 | 2.368476 | 2.422876 | 2.468217 | 2.519132 | 2.601767 | 2.69139 | 2.780343 | 2.861188 | 2.910449 | 2.953967 | 3.005746 | 3.053083 |
|  | 15 | 2.183034 | 2.231074 | 2.283236 | 2.326763 | 2.375691 | 2.45519 | 2.541503 | 2.627217 | 2.705118 | 2.75257 | 2.794472 | 2.844303 | 2.889826 |
|  | 20 | 2.091723 | 2.138523 | 2.189404 | 2.231909 | 2.279733 | 2.357524 | 2.442067 | 2.526065 | 2.602403 | 2.648887 | 2.689918 | 2.738686 | 2.783208 |
|  | 30 | 2.001854 | 2.048348 | 2.098969 | 2.141306 | 2.188985 | 2.266607 | 2.350998 | 2.4348 | 2.510851 | 2.557083 | 2.597831 | 2.646178 | 2.690227 |
|  | 40 | 1.950344 | 1.997213 | 2.048306 | 2.091083 | 2.139296 | 2.217846 | 2.303271 | 2.388048 | 2.464874 | 2.511502 | 2.55254 | 2.60115 | 2.645353 |
|  | 50 | 1.912482 | 1.959052 | 2.010021 | 2.052844 | 2.101263 | 2.180461 | 2.266964 | 2.353112 | 2.431364 | 2.478915 | 2.520787 | 2.570401 | 2.615522 |
|  | 60 | 1.872206 | 1.917388 | 1.967173 | 2.009264 | 2.057131 | 2.136024 | 2.222971 | 2.310279 | 2.390127 | 2.438869 | 2.481916 | 2.533061 | 2.579692 |
|  | 70 | 1.832542 | 1.875881 | 1.924018 | 1.965021 | 2.01198 | 2.090114 | 2.177225 | 2.265674 | 2.347349 | 2.397546 | 2.442079 | 2.49522 | 2.543879 |
|  | 80 | 1.776016 | 1.817047 | 1.86299 | 1.902425 | 1.947922 | 2.024387 | 2.110716 | 2.199469 | 2.282346 | 2.333697 | 2.379506 | 2.434468 | 2.485071 |
|  | 90 | 1.718529 | 1.757311 | 1.801074 | 1.838921 | 1.882905 | 1.957582 | 2.042998 | 2.131982 | 2.216102 | 2.268701 | 2.315923 | 2.372945 | 2.425787 |
| **L_rostralanteriorcingulate** | 3 | 2.581834 | 2.730356 | 2.880349 | 2.997526 | 3.121224 | 3.305014 | 3.481739 | 3.644682 | 3.788953 | 3.874037 | 3.946914 | 4.030469 | 4.103624 |
|  | 5 | 2.554954 | 2.689965 | 2.828829 | 2.938905 | 3.056495 | 3.233625 | 3.40645 | 3.567552 | 3.711277 | 3.79649 | 3.869733 | 3.95399 | 4.028001 |
|  | 10 | 2.486616 | 2.595496 | 2.711715 | 2.806753 | 2.911016 | 3.073221 | 3.237248 | 3.394499 | 3.537656 | 3.623769 | 3.698509 | 3.7853 | 3.862244 |
|  | 15 | 2.433751 | 2.525145 | 2.625359 | 2.709315 | 2.803463 | 2.954142 | 3.111683 | 3.266949 | 3.411299 | 3.49949 | 3.576858 | 3.667657 | 3.749006 |
|  | 20 | 2.3925 | 2.47331 | 2.563466 | 2.640238 | 2.727681 | 2.870623 | 3.02406 | 3.1787 | 3.324962 | 3.415509 | 3.495691 | 3.590687 | 3.676618 |
|  | 30 | 2.295194 | 2.369126 | 2.452453 | 2.524129 | 2.606603 | 2.743434 | 2.89325 | 3.046265 | 3.191848 | 3.282442 | 3.362994 | 3.458852 | 3.545974 |
|  | 40 | 2.226622 | 2.300549 | 2.383858 | 2.455531 | 2.538046 | 2.675153 | 2.825733 | 2.97898 | 3.123324 | 3.212509 | 3.291446 | 3.384976 | 3.469628 |
|  | 50 | 2.200714 | 2.27202 | 2.352928 | 2.423024 | 2.504306 | 2.640847 | 2.793071 | 2.949353 | 3.096803 | 3.188075 | 3.268999 | 3.365076 | 3.452234 |
|  | 60 | 2.210207 | 2.277088 | 2.353881 | 2.421214 | 2.500266 | 2.635538 | 2.790213 | 2.952216 | 3.10703 | 3.203867 | 3.290406 | 3.394012 | 3.488834 |
|  | 70 | 2.205999 | 2.270598 | 2.345305 | 2.411302 | 2.489405 | 2.624725 | 2.782208 | 2.949199 | 3.109677 | 3.210556 | 3.301071 | 3.409925 | 3.510039 |
|  | 80 | 2.213792 | 2.277312 | 2.351181 | 2.416827 | 2.495017 | 2.631887 | 2.793568 | 2.966665 | 3.133515 | 3.238706 | 3.333334 | 3.447477 | 3.552813 |
|  | 90 | 2.234035 | 2.297658 | 2.371948 | 2.438263 | 2.517638 | 2.657709 | 2.825149 | 3.005561 | 3.179435 | 3.289089 | 3.387801 | 3.506992 | 3.617134 |
| **L_rostralmiddlefrontal** | 3 | 2.497281 | 2.56067 | 2.629329 | 2.685872 | 2.74785 | 2.842737 | 2.934354 | 3.027081 | 3.125518 | 3.19118 | 3.252054 | 3.327249 | 3.397956 |
|  | 5 | 2.441046 | 2.500569 | 2.565487 | 2.619365 | 2.678939 | 2.7715 | 2.863059 | 2.95574 | 3.051773 | 3.114869 | 3.172826 | 3.243817 | 3.310052 |
|  | 10 | 2.308166 | 2.359669 | 2.416706 | 2.46486 | 2.51915 | 2.606315 | 2.69721 | 2.789239 | 2.879678 | 2.937126 | 2.988819 | 3.05097 | 3.107971 |
|  | 15 | 2.18685 | 2.233029 | 2.284692 | 2.328808 | 2.379197 | 2.461904 | 2.551171 | 2.641555 | 2.727343 | 2.780605 | 2.827871 | 2.883998 | 2.934889 |
|  | 20 | 2.108318 | 2.151939 | 2.200983 | 2.243099 | 2.291515 | 2.371861 | 2.460048 | 2.549327 | 2.632612 | 2.683724 | 2.728766 | 2.781915 | 2.829829 |
|  | 30 | 2.030693 | 2.072242 | 2.119195 | 2.159747 | 2.206677 | 2.28545 | 2.373411 | 2.462412 | 2.543927 | 2.593328 | 2.636532 | 2.687166 | 2.73253 |
|  | 40 | 1.988094 | 2.031625 | 2.080509 | 2.122432 | 2.170562 | 2.250269 | 2.3375 | 2.425687 | 2.507976 | 2.558479 | 2.602981 | 2.655484 | 2.702806 |
|  | 50 | 1.935921 | 1.984153 | 2.037534 | 2.082582 | 2.133359 | 2.214889 | 2.299972 | 2.385898 | 2.469852 | 2.52296 | 2.570616 | 2.627761 | 2.680031 |
|  | 60 | 1.880735 | 1.932821 | 1.989924 | 2.037608 | 2.090722 | 2.174327 | 2.258883 | 2.344207 | 2.43011 | 2.485499 | 2.535777 | 2.59669 | 2.652934 |
|  | 70 | 1.832784 | 1.882445 | 1.937471 | 1.98397 | 2.03647 | 2.121025 | 2.20971 | 2.299168 | 2.385924 | 2.44053 | 2.489376 | 2.547777 | 2.60105 |
|  | 80 | 1.836312 | 1.878506 | 1.926749 | 1.968977 | 2.018617 | 2.104216 | 2.203653 | 2.303951 | 2.391693 | 2.443184 | 2.487346 | 2.538208 | 2.583056 |
|  | 90 | 1.867191 | 1.900471 | 1.940193 | 1.976701 | 2.022122 | 2.10839 | 2.221986 | 2.336549 | 2.424864 | 2.471828 | 2.509813 | 2.551384 | 2.586411 |
| **L_superiorfrontal** | 3 | 2.822843 | 2.8901 | 2.96067 | 3.017836 | 3.080486 | 3.179343 | 3.283752 | 3.385837 | 3.478383 | 3.535067 | 3.58551 | 3.646149 | 3.702317 |
|  | 5 | 2.776398 | 2.842119 | 2.911229 | 2.967322 | 3.028903 | 3.126292 | 3.229423 | 3.330508 | 3.422344 | 3.478678 | 3.528862 | 3.589251 | 3.645246 |
|  | 10 | 2.663736 | 2.725761 | 2.791338 | 2.844822 | 2.903799 | 2.997598 | 3.097597 | 3.196236 | 3.286346 | 3.341843 | 3.391414 | 3.451229 | 3.506842 |
|  | 15 | 2.559603 | 2.618234 | 2.680549 | 2.731617 | 2.788175 | 2.87864 | 2.975745 | 3.072155 | 3.160738 | 3.215522 | 3.264598 | 3.323986 | 3.379366 |
|  | 20 | 2.475586 | 2.531579 | 2.591365 | 2.640564 | 2.695263 | 2.783197 | 2.878162 | 2.973003 | 3.060603 | 3.114986 | 3.163833 | 3.2231 | 3.278519 |
|  | 30 | 2.368253 | 2.421974 | 2.479606 | 2.527238 | 2.580405 | 2.66632 | 2.759681 | 2.853466 | 2.940528 | 2.994771 | 3.043609 | 3.10301 | 3.158687 |
|  | 40 | 2.302935 | 2.356685 | 2.414406 | 2.462155 | 2.515492 | 2.60176 | 2.69558 | 2.789865 | 2.877388 | 2.931904 | 2.980973 | 3.040627 | 3.096513 |
|  | 50 | 2.24613 | 2.30077 | 2.359411 | 2.407888 | 2.462004 | 2.549436 | 2.644369 | 2.739576 | 2.827751 | 2.882563 | 2.93182 | 2.991601 | 3.047501 |
|  | 60 | 2.19088 | 2.246977 | 2.307071 | 2.356664 | 2.411929 | 2.501002 | 2.597395 | 2.693706 | 2.78256 | 2.837618 | 2.88698 | 2.946737 | 3.002461 |
|  | 70 | 2.126314 | 2.183635 | 2.244938 | 2.295446 | 2.351646 | 2.442024 | 2.539537 | 2.636639 | 2.725909 | 2.781068 | 2.830413 | 2.890012 | 2.945453 |
|  | 80 | 2.067338 | 2.12639 | 2.189382 | 2.241161 | 2.298642 | 2.390794 | 2.489813 | 2.587981 | 2.677833 | 2.733157 | 2.78252 | 2.841979 | 2.897128 |
|  | 90 | 2.022818 | 2.084501 | 2.150055 | 2.203755 | 2.26318 | 2.358041 | 2.459415 | 2.559346 | 2.65031 | 2.706077 | 2.755681 | 2.815235 | 2.870284 |
| **L_superiorparietal** | 3 | 2.259242 | 2.304818 | 2.355407 | 2.398442 | 2.447632 | 2.529133 | 2.619284 | 2.709776 | 2.792176 | 2.842182 | 2.886103 | 2.93794 | 2.984832 |
|  | 5 | 2.209726 | 2.254473 | 2.304157 | 2.346435 | 2.394773 | 2.474898 | 2.56358 | 2.652655 | 2.733817 | 2.783098 | 2.826398 | 2.877522 | 2.923788 |
|  | 10 | 2.085647 | 2.128281 | 2.175655 | 2.215998 | 2.262159 | 2.338759 | 2.423667 | 2.509093 | 2.58706 | 2.634462 | 2.676153 | 2.725425 | 2.770063 |
|  | 15 | 1.971975 | 2.012672 | 2.057927 | 2.096495 | 2.140656 | 2.214018 | 2.295457 | 2.377528 | 2.452557 | 2.498233 | 2.538444 | 2.586015 | 2.629157 |
|  | 20 | 1.898774 | 1.938548 | 1.982777 | 2.020472 | 2.063636 | 2.135352 | 2.214985 | 2.295268 | 2.368697 | 2.41342 | 2.452806 | 2.499418 | 2.541709 |
|  | 30 | 1.828255 | 1.869389 | 1.914868 | 1.953418 | 1.997333 | 2.069781 | 2.149518 | 2.229203 | 2.301515 | 2.345305 | 2.383719 | 2.429007 | 2.469939 |
|  | 40 | 1.782763 | 1.826414 | 1.8743 | 1.914595 | 1.960182 | 2.034696 | 2.115777 | 2.195909 | 2.267913 | 2.311209 | 2.349009 | 2.393365 | 2.433269 |
|  | 50 | 1.755423 | 1.800427 | 1.849685 | 1.891047 | 1.937755 | 2.013913 | 2.096551 | 2.17802 | 2.251081 | 2.294959 | 2.333238 | 2.378127 | 2.418486 |
|  | 60 | 1.729419 | 1.774019 | 1.823012 | 1.864293 | 1.911061 | 1.987666 | 2.071274 | 2.154197 | 2.228989 | 2.274101 | 2.313578 | 2.360016 | 2.401903 |
|  | 70 | 1.703236 | 1.746076 | 1.793507 | 1.833772 | 1.879719 | 1.95573 | 2.039753 | 2.124181 | 2.201266 | 2.248192 | 2.289522 | 2.33846 | 2.382897 |
|  | 80 | 1.658541 | 1.698266 | 1.742718 | 1.780843 | 1.824789 | 1.898529 | 1.981568 | 2.066649 | 2.145791 | 2.194662 | 2.238145 | 2.290168 | 2.337915 |
|  | 90 | 1.586663 | 1.622589 | 1.663226 | 1.698454 | 1.739497 | 1.809448 | 1.889894 | 1.974229 | 2.054481 | 2.104934 | 2.150413 | 2.205569 | 2.256919 |
| **L_superiortemporal** | 3 | 2.549748 | 2.639232 | 2.730713 | 2.80272 | 2.879035 | 2.992537 | 3.101207 | 3.202247 | 3.293573 | 3.348113 | 3.395163 | 3.449434 | 3.4972 |
|  | 5 | 2.533104 | 2.620408 | 2.710047 | 2.780866 | 2.85617 | 2.968645 | 3.076892 | 3.177742 | 3.268813 | 3.32318 | 3.370075 | 3.424167 | 3.471779 |
|  | 10 | 2.492583 | 2.573993 | 2.658608 | 2.726167 | 2.798681 | 2.908302 | 3.015354 | 3.115749 | 3.206317 | 3.260378 | 3.307025 | 3.36086 | 3.40828 |
|  | 15 | 2.463351 | 2.535189 | 2.611553 | 2.673732 | 2.741644 | 2.846605 | 2.951798 | 3.052124 | 3.143342 | 3.198127 | 3.245611 | 3.300662 | 3.349379 |
|  | 20 | 2.446536 | 2.506953 | 2.573068 | 2.628344 | 2.690206 | 2.788924 | 2.891744 | 2.992741 | 3.086389 | 3.14348 | 3.193486 | 3.252077 | 3.304482 |
|  | 30 | 2.384603 | 2.438841 | 2.499117 | 2.55028 | 2.608409 | 2.703219 | 2.804884 | 2.906317 | 3.000496 | 3.05804 | 3.108563 | 3.167937 | 3.221223 |
|  | 40 | 2.316272 | 2.37282 | 2.435245 | 2.487934 | 2.547525 | 2.644282 | 2.747689 | 2.849229 | 2.94097 | 2.995959 | 3.043646 | 3.099037 | 3.148196 |
|  | 50 | 2.260421 | 2.317473 | 2.38044 | 2.43361 | 2.49382 | 2.5919 | 2.69736 | 2.800253 | 2.891537 | 2.94558 | 2.992092 | 3.045746 | 3.093062 |
|  | 60 | 2.215261 | 2.2685 | 2.328016 | 2.378914 | 2.437293 | 2.534199 | 2.64104 | 2.74656 | 2.840035 | 2.895346 | 2.942966 | 2.997946 | 3.04649 |
|  | 70 | 2.14741 | 2.197464 | 2.253994 | 2.30284 | 2.35947 | 2.455019 | 2.562723 | 2.670154 | 2.765006 | 2.821022 | 2.869218 | 2.924856 | 2.973996 |
|  | 80 | 2.037762 | 2.091109 | 2.150762 | 2.201854 | 2.260627 | 2.358877 | 2.468473 | 2.575371 | 2.666812 | 2.719628 | 2.764436 | 2.815487 | 2.860016 |
|  | 90 | 1.892661 | 1.957514 | 2.02751 | 2.085618 | 2.150637 | 2.255642 | 2.368216 | 2.47288 | 2.557964 | 2.605456 | 2.644898 | 2.688966 | 2.726702 |
| **L_supramarginal** | 3 | 2.481802 | 2.58111 | 2.678427 | 2.752237 | 2.827853 | 2.935208 | 3.031796 | 3.120864 | 3.204775 | 3.256085 | 3.300912 | 3.353151 | 3.399517 |
|  | 5 | 2.453605 | 2.544144 | 2.634582 | 2.704273 | 2.776654 | 2.881226 | 2.977382 | 3.066583 | 3.149923 | 3.200671 | 3.244923 | 3.296422 | 3.342092 |
|  | 10 | 2.377995 | 2.450825 | 2.526605 | 2.587112 | 2.651993 | 2.749768 | 2.844618 | 2.934033 | 3.016028 | 3.065482 | 3.108418 | 3.158239 | 3.202339 |
|  | 15 | 2.304674 | 2.364196 | 2.428238 | 2.480978 | 2.539213 | 2.630607 | 2.724057 | 2.813795 | 2.894896 | 2.943445 | 2.985457 | 3.034116 | 3.077148 |
|  | 20 | 2.251711 | 2.301859 | 2.357264 | 2.404077 | 2.457094 | 2.543401 | 2.636025 | 2.726776 | 2.808087 | 2.85655 | 2.898431 | 2.946926 | 2.98984 |
|  | 30 | 2.186734 | 2.229324 | 2.277545 | 2.319311 | 2.367835 | 2.449926 | 2.542751 | 2.63592 | 2.718889 | 2.768188 | 2.810776 | 2.860126 | 2.903872 |
|  | 40 | 2.127618 | 2.171268 | 2.220581 | 2.263186 | 2.312545 | 2.395648 | 2.488964 | 2.582626 | 2.666626 | 2.716783 | 2.760237 | 2.810721 | 2.855576 |
|  | 50 | 2.066849 | 2.114661 | 2.168038 | 2.21357 | 2.265592 | 2.35123 | 2.444337 | 2.537265 | 2.622427 | 2.674012 | 2.719072 | 2.771787 | 2.818907 |
|  | 60 | 2.007931 | 2.059636 | 2.116765 | 2.164953 | 2.219322 | 2.306997 | 2.399425 | 2.491551 | 2.578382 | 2.631957 | 2.679268 | 2.735146 | 2.785522 |
|  | 70 | 1.955815 | 2.006994 | 2.063719 | 2.111731 | 2.16611 | 2.254359 | 2.348291 | 2.441934 | 2.529388 | 2.583024 | 2.630221 | 2.685791 | 2.735748 |
|  | 80 | 1.902502 | 1.950629 | 2.004537 | 2.050708 | 2.103723 | 2.191782 | 2.288862 | 2.385384 | 2.471979 | 2.523677 | 2.568437 | 2.620385 | 2.666481 |
|  | 90 | 1.860343 | 1.904606 | 1.954899 | 1.998665 | 2.04983 | 2.137424 | 2.23825 | 2.338451 | 2.424497 | 2.47433 | 2.516707 | 2.565121 | 2.60748 |
| **L_frontalpole** | 3 | 2.415261 | 2.54188 | 2.68102 | 2.798266 | 2.93106 | 3.148313 | 3.384705 | 3.617931 | 3.826799 | 3.951934 | 4.060837 | 4.188157 | 4.30221 |
|  | 5 | 2.359738 | 2.479778 | 2.612322 | 2.724503 | 2.852081 | 3.061944 | 3.291829 | 3.520126 | 3.725789 | 3.849538 | 3.957552 | 4.084205 | 4.197996 |
|  | 10 | 2.244375 | 2.35121 | 2.4704 | 2.572259 | 2.689168 | 2.883897 | 3.100572 | 3.319164 | 3.518949 | 3.640458 | 3.74731 | 3.873537 | 3.987807 |
|  | 15 | 2.176709 | 2.275495 | 2.386554 | 2.482166 | 2.592693 | 2.778648 | 2.988266 | 3.202609 | 3.401033 | 3.522892 | 3.63079 | 3.759143 | 3.876171 |
|  | 20 | 2.123003 | 2.216819 | 2.322809 | 2.414498 | 2.520998 | 2.701414 | 2.906669 | 3.118638 | 3.316778 | 3.439385 | 3.548538 | 3.679114 | 3.798864 |
|  | 30 | 2.037698 | 2.126269 | 2.226682 | 2.313856 | 2.41549 | 2.588647 | 2.787254 | 2.994316 | 3.189807 | 3.31177 | 3.421016 | 3.552553 | 3.674022 |
|  | 40 | 1.995298 | 2.082822 | 2.181922 | 2.267867 | 2.36799 | 2.538456 | 2.73396 | 2.937984 | 3.130987 | 3.251662 | 3.359961 | 3.490655 | 3.611659 |
|  | 50 | 1.986646 | 2.074924 | 2.174608 | 2.260854 | 2.361115 | 2.53139 | 2.726208 | 2.929245 | 3.121293 | 3.241451 | 3.349381 | 3.47979 | 3.600722 |
|  | 60 | 1.919024 | 2.006766 | 2.105687 | 2.191158 | 2.290413 | 2.458814 | 2.651421 | 2.852336 | 3.042784 | 3.162234 | 3.269763 | 3.400024 | 3.521186 |
|  | 70 | 1.856692 | 1.944435 | 2.043561 | 2.129401 | 2.229335 | 2.399577 | 2.595537 | 2.801616 | 2.998765 | 3.123407 | 3.236306 | 3.373999 | 3.50302 |
|  | 80 | 1.832715 | 1.921291 | 2.021965 | 2.109689 | 2.21248 | 2.38934 | 2.595868 | 2.816788 | 3.03202 | 3.170179 | 3.29678 | 3.453117 | 3.601592 |
|  | 90 | 1.800393 | 1.888677 | 1.989756 | 2.078505 | 2.183338 | 2.365983 | 2.583225 | 2.820796 | 3.057875 | 3.213179 | 3.357735 | 3.539309 | 3.714986 |
| **L_temporalpole** | 3 | 2.319029 | 2.54195 | 2.757523 | 2.92218 | 3.095229 | 3.357352 | 3.622122 | 3.850172 | 4.019933 | 4.109565 | 4.18166 | 4.260002 | 4.325445 |
|  | 5 | 2.361728 | 2.569386 | 2.7739 | 2.932377 | 3.100757 | 3.358696 | 3.622018 | 3.851289 | 4.023775 | 4.115441 | 4.189446 | 4.270122 | 4.337707 |
|  | 10 | 2.454466 | 2.630432 | 2.811399 | 2.956293 | 3.114099 | 3.3622 | 3.621906 | 3.854171 | 4.033696 | 4.130723 | 4.209821 | 4.296782 | 4.370187 |
|  | 15 | 2.52443 | 2.678639 | 2.842415 | 2.976836 | 3.126103 | 3.365782 | 3.622053 | 3.85705 | 4.043616 | 4.146184 | 4.230641 | 4.324315 | 4.404023 |
|  | 20 | 2.567714 | 2.709558 | 2.863008 | 2.99082 | 3.134429 | 3.368021 | 3.621084 | 3.857639 | 4.049862 | 4.157158 | 4.246309 | 4.345987 | 4.431427 |
|  | 30 | 2.567091 | 2.709445 | 2.86316 | 2.990828 | 3.133709 | 3.364341 | 3.611384 | 3.843798 | 4.036988 | 4.146485 | 4.238277 | 4.341701 | 4.430958 |
|  | 40 | 2.527533 | 2.683115 | 2.847486 | 2.981354 | 3.128459 | 3.360022 | 3.60046 | 3.824322 | 4.012246 | 4.119469 | 4.209666 | 4.311562 | 4.399682 |
|  | 50 | 2.506482 | 2.666387 | 2.833961 | 2.969347 | 3.116888 | 3.346151 | 3.579973 | 3.798006 | 3.984674 | 4.092554 | 4.183969 | 4.287888 | 4.378248 |
|  | 60 | 2.474916 | 2.633448 | 2.800298 | 2.93539 | 3.082642 | 3.310804 | 3.541787 | 3.759549 | 3.951257 | 4.064082 | 4.160727 | 4.271643 | 4.368924 |
|  | 70 | 2.411249 | 2.569124 | 2.736443 | 2.872524 | 3.021188 | 3.251451 | 3.483371 | 3.704987 | 3.906193 | 4.027032 | 4.131821 | 4.253419 | 4.361158 |
|  | 80 | 2.320336 | 2.482004 | 2.653676 | 2.793347 | 2.945741 | 3.180695 | 3.414942 | 3.640914 | 3.851956 | 3.981068 | 4.094297 | 4.22703 | 4.345738 |
|  | 90 | 2.23392 | 2.397871 | 2.572613 | 2.715039 | 2.870434 | 3.109233 | 3.345123 | 3.575126 | 3.796472 | 3.934589 | 4.057205 | 4.202557 | 4.333908 |
| **L_transversetemporal** | 3 | 2.11739 | 2.190679 | 2.27366 | 2.345477 | 2.428783 | 2.56916 | 2.7268 | 2.886137 | 3.030946 | 3.118226 | 3.194294 | 3.283161 | 3.36254 |
|  | 5 | 2.087803 | 2.160828 | 2.243474 | 2.314973 | 2.397875 | 2.537493 | 2.694168 | 2.852411 | 2.996129 | 3.082708 | 3.15814 | 3.246233 | 3.324893 |
|  | 10 | 2.015781 | 2.08818 | 2.170026 | 2.240758 | 2.322686 | 2.460463 | 2.614784 | 2.770357 | 2.911406 | 2.996269 | 3.070141 | 3.156339 | 3.233241 |
|  | 15 | 1.948562 | 2.020466 | 2.101659 | 2.171747 | 2.252842 | 2.389011 | 2.54124 | 2.694406 | 2.833027 | 2.916321 | 2.988763 | 3.073219 | 3.148502 |
|  | 20 | 1.889166 | 1.960834 | 2.041663 | 2.111355 | 2.1919 | 2.326934 | 2.477594 | 2.628879 | 2.765551 | 2.847564 | 2.91883 | 3.00184 | 3.075769 |
|  | 30 | 1.787799 | 1.859624 | 1.940415 | 2.009899 | 2.090009 | 2.223866 | 2.372589 | 2.521309 | 2.65516 | 2.735264 | 2.804741 | 2.885522 | 2.957338 |
|  | 40 | 1.709995 | 1.78296 | 1.86479 | 1.934971 | 2.01567 | 2.150026 | 2.298633 | 2.446584 | 2.579221 | 2.658374 | 2.726896 | 2.806421 | 2.876989 |
|  | 50 | 1.648131 | 1.723117 | 1.806933 | 1.878596 | 1.960758 | 2.097012 | 2.246994 | 2.395617 | 2.528309 | 2.607263 | 2.675479 | 2.754498 | 2.824486 |
|  | 60 | 1.591315 | 1.668896 | 1.755289 | 1.828897 | 1.913017 | 2.05192 | 2.204024 | 2.354003 | 2.487327 | 2.566415 | 2.634609 | 2.713447 | 2.783142 |
|  | 70 | 1.52973 | 1.610092 | 1.699197 | 1.77482 | 1.860931 | 2.002451 | 2.156552 | 2.307695 | 2.441444 | 2.520532 | 2.588583 | 2.667098 | 2.736371 |
|  | 80 | 1.460492 | 1.543654 | 1.635407 | 1.712932 | 1.800852 | 1.944595 | 2.100162 | 2.251883 | 2.385502 | 2.464252 | 2.531864 | 2.609711 | 2.678258 |
|  | 90 | 1.386409 | 1.472475 | 1.566886 | 1.646253 | 1.735855 | 1.881509 | 2.038101 | 2.189903 | 2.322922 | 2.401049 | 2.467976 | 2.54487 | 2.612438 |
| **L_insula** | 3 | 2.900816 | 3.014069 | 3.121494 | 3.201572 | 3.283216 | 3.400434 | 3.510044 | 3.605977 | 3.684911 | 3.729302 | 3.766245 | 3.807516 | 3.842799 |
|  | 5 | 2.881321 | 2.982558 | 3.081249 | 3.156333 | 3.234073 | 3.347517 | 3.455269 | 3.550832 | 3.630344 | 3.675376 | 3.713016 | 3.755229 | 3.791447 |
|  | 10 | 2.828031 | 2.906924 | 2.988219 | 3.052793 | 3.121971 | 3.226824 | 3.330293 | 3.425235 | 3.506608 | 3.553598 | 3.593347 | 3.638421 | 3.677493 |
|  | 15 | 2.785636 | 2.849808 | 2.918607 | 2.975118 | 3.037417 | 3.135143 | 3.235241 | 3.330432 | 3.414744 | 3.464533 | 3.50726 | 3.556368 | 3.599488 |
|  | 20 | 2.75197 | 2.807521 | 2.868564 | 2.91983 | 2.977497 | 3.070302 | 3.168207 | 3.264242 | 3.35194 | 3.404869 | 3.450951 | 3.504655 | 3.552454 |
|  | 30 | 2.660769 | 2.714214 | 2.773332 | 2.823275 | 2.879742 | 2.971164 | 3.068202 | 3.16461 | 3.254301 | 3.309153 | 3.357325 | 3.413933 | 3.464725 |
|  | 40 | 2.586254 | 2.641778 | 2.702951 | 2.754418 | 2.812356 | 2.905537 | 3.003506 | 3.100602 | 3.191358 | 3.247019 | 3.295976 | 3.353574 | 3.405304 |
|  | 50 | 2.534499 | 2.58914 | 2.649654 | 2.700801 | 2.758606 | 2.851986 | 2.950565 | 3.049345 | 3.143285 | 3.201619 | 3.25335 | 3.314699 | 3.370227 |
|  | 60 | 2.501303 | 2.560613 | 2.625769 | 2.680401 | 2.741652 | 2.839454 | 2.941095 | 3.042084 | 3.138042 | 3.197539 | 3.250219 | 3.312574 | 3.368889 |
|  | 70 | 2.439082 | 2.504503 | 2.575462 | 2.634241 | 2.699375 | 2.801691 | 2.90582 | 3.007924 | 3.104447 | 3.164015 | 3.216559 | 3.2785 | 3.334198 |
|  | 80 | 2.395701 | 2.461611 | 2.533281 | 2.592765 | 2.658765 | 2.762522 | 2.868036 | 2.972137 | 3.071819 | 3.133881 | 3.188937 | 3.254189 | 3.313165 |
|  | 90 | 2.338484 | 2.402909 | 2.473399 | 2.532217 | 2.597773 | 2.701341 | 2.807101 | 2.912597 | 3.015397 | 3.080207 | 3.138183 | 3.207451 | 3.270555 |
| **R_bankssts** | 3 | 2.458923 | 2.550296 | 2.64509 | 2.721116 | 2.803669 | 2.932275 | 3.065752 | 3.193653 | 3.30712 | 3.375363 | 3.435257 | 3.506195 | 3.570857 |
|  | 5 | 2.424884 | 2.512184 | 2.603426 | 2.677064 | 2.757448 | 2.883452 | 3.015044 | 3.141693 | 3.254316 | 3.322104 | 3.3816 | 3.452041 | 3.516204 |
|  | 10 | 2.337799 | 2.416018 | 2.499152 | 2.56722 | 2.64245 | 2.762121 | 2.889007 | 3.012495 | 3.12303 | 3.189737 | 3.248324 | 3.317675 | 3.380781 |
|  | 15 | 2.253443 | 2.323981 | 2.400068 | 2.463177 | 2.533722 | 2.647507 | 2.769944 | 2.890483 | 2.999191 | 3.065037 | 3.122958 | 3.191575 | 3.254019 |
|  | 20 | 2.188866 | 2.253726 | 2.324546 | 2.383928 | 2.450954 | 2.560371 | 2.679666 | 2.798363 | 2.906187 | 2.971745 | 3.029518 | 3.098039 | 3.16043 |
|  | 30 | 2.114009 | 2.174762 | 2.241877 | 2.298737 | 2.363499 | 2.470369 | 2.588153 | 2.706182 | 2.813702 | 2.879057 | 2.936563 | 3.004594 | 3.066318 |
|  | 40 | 2.053418 | 2.115189 | 2.183446 | 2.241259 | 2.307051 | 2.415384 | 2.534212 | 2.652404 | 2.759058 | 2.823323 | 2.879477 | 2.945398 | 3.004699 |
|  | 50 | 1.996798 | 2.059765 | 2.129312 | 2.18817 | 2.255074 | 2.364977 | 2.485003 | 2.603644 | 2.709897 | 2.773487 | 2.828757 | 2.89326 | 2.950913 |
|  | 60 | 1.948422 | 2.011744 | 2.081796 | 2.141154 | 2.208685 | 2.319686 | 2.440876 | 2.560466 | 2.667254 | 2.730969 | 2.786203 | 2.85047 | 2.907718 |
|  | 70 | 1.905386 | 1.968597 | 2.038717 | 2.098274 | 2.166167 | 2.278019 | 2.400389 | 2.52126 | 2.629172 | 2.693503 | 2.749217 | 2.813962 | 2.871545 |
|  | 80 | 1.847223 | 1.910299 | 1.980365 | 2.039944 | 2.107925 | 2.220027 | 2.342744 | 2.463938 | 2.572039 | 2.636408 | 2.692095 | 2.756724 | 2.814115 |
|  | 90 | 1.781283 | 1.844361 | 1.914443 | 1.97404 | 2.042039 | 2.154139 | 2.276765 | 2.397715 | 2.505419 | 2.569451 | 2.624775 | 2.688893 | 2.745741 |
| **R_caudalanteriorcingulate** | 3 | 2.399764 | 2.476966 | 2.56247 | 2.635317 | 2.719054 | 2.860067 | 3.02184 | 3.194086 | 3.363498 | 3.474034 | 3.577026 | 3.706943 | 3.83358 |
|  | 5 | 2.353896 | 2.429965 | 2.51425 | 2.586081 | 2.668668 | 2.807759 | 2.967286 | 3.137002 | 3.303703 | 3.412319 | 3.513399 | 3.640723 | 3.764629 |
|  | 10 | 2.248123 | 2.321622 | 2.40314 | 2.472665 | 2.552639 | 2.687353 | 2.841752 | 3.005672 | 3.166148 | 3.270349 | 3.367034 | 3.488398 | 3.606027 |
|  | 15 | 2.162941 | 2.234477 | 2.313888 | 2.381659 | 2.459647 | 2.591024 | 2.741476 | 2.900869 | 3.056397 | 3.157043 | 3.250159 | 3.366648 | 3.479111 |
|  | 20 | 2.101098 | 2.171316 | 2.249323 | 2.315932 | 2.392605 | 2.521761 | 2.669531 | 2.825743 | 2.977663 | 3.075647 | 3.166042 | 3.278752 | 3.387155 |
|  | 30 | 2.020355 | 2.08907 | 2.165518 | 2.230865 | 2.306129 | 2.432908 | 2.577718 | 2.730193 | 2.877586 | 2.972071 | 3.058786 | 3.166259 | 3.268914 |
|  | 40 | 1.98433 | 2.052984 | 2.129552 | 2.195133 | 2.270786 | 2.398404 | 2.544236 | 2.697557 | 2.845263 | 2.939582 | 3.025844 | 3.132315 | 3.233521 |
|  | 50 | 1.950772 | 2.019571 | 2.096584 | 2.162762 | 2.239319 | 2.368888 | 2.517396 | 2.673753 | 2.824326 | 2.920344 | 3.00802 | 3.116008 | 3.218384 |
|  | 60 | 1.92813 | 1.997965 | 2.076478 | 2.144209 | 2.222837 | 2.356489 | 2.51036 | 2.672868 | 2.829589 | 2.929536 | 3.020759 | 3.133013 | 3.239292 |
|  | 70 | 1.910096 | 1.981643 | 2.062424 | 2.132381 | 2.213879 | 2.353011 | 2.51393 | 2.684457 | 2.849207 | 2.954323 | 3.050246 | 3.168216 | 3.279796 |
|  | 80 | 1.921084 | 1.995971 | 2.080855 | 2.154627 | 2.240845 | 2.388622 | 2.560257 | 2.742691 | 2.919217 | 3.031881 | 3.134667 | 3.260996 | 3.380357 |
|  | 90 | 1.970441 | 2.050608 | 2.14182 | 2.221362 | 2.314612 | 2.475059 | 2.662176 | 2.86167 | 3.05502 | 3.178476 | 3.291093 | 3.429434 | 3.560032 |
| **R_caudalmiddlefrontal** | 3 | 2.317082 | 2.410027 | 2.501266 | 2.571205 | 2.644308 | 2.753031 | 2.860003 | 2.95746 | 3.040075 | 3.088103 | 3.129263 | 3.176846 | 3.21915 |
|  | 5 | 2.310932 | 2.397062 | 2.483088 | 2.549931 | 2.620561 | 2.726901 | 2.832859 | 2.930372 | 3.013652 | 3.062292 | 3.104091 | 3.152533 | 3.195699 |
|  | 10 | 2.287129 | 2.359412 | 2.434274 | 2.49419 | 2.559077 | 2.659635 | 2.762976 | 2.860559 | 2.94556 | 2.995834 | 3.039375 | 3.090194 | 3.135775 |
|  | 15 | 2.252397 | 2.314141 | 2.379919 | 2.433868 | 2.493556 | 2.588542 | 2.689117 | 2.786625 | 2.87339 | 2.925443 | 2.970936 | 3.024486 | 3.072911 |
|  | 20 | 2.211719 | 2.266358 | 2.325718 | 2.375269 | 2.430979 | 2.52149 | 2.619691 | 2.717063 | 2.805361 | 2.859028 | 2.906333 | 2.962474 | 3.013645 |
|  | 30 | 2.130273 | 2.181753 | 2.238218 | 2.28576 | 2.339625 | 2.42799 | 2.524883 | 2.621781 | 2.710162 | 2.764041 | 2.811599 | 2.868087 | 2.919591 |
|  | 40 | 2.069268 | 2.122665 | 2.18104 | 2.230023 | 2.285323 | 2.375539 | 2.473652 | 2.570799 | 2.658454 | 2.711405 | 2.757821 | 2.812545 | 2.862043 |
|  | 50 | 2.021591 | 2.075943 | 2.135368 | 2.185224 | 2.241482 | 2.333153 | 2.43259 | 2.530654 | 2.618683 | 2.671611 | 2.717831 | 2.772097 | 2.820956 |
|  | 60 | 1.980099 | 2.035485 | 2.096051 | 2.146861 | 2.204181 | 2.297497 | 2.398513 | 2.497811 | 2.586573 | 2.639734 | 2.686013 | 2.740161 | 2.788727 |
|  | 70 | 1.942695 | 1.997558 | 2.057825 | 2.108587 | 2.166056 | 2.260016 | 2.362186 | 2.462942 | 2.553167 | 2.607233 | 2.654298 | 2.709343 | 2.758678 |
|  | 80 | 1.893796 | 1.949015 | 2.009742 | 2.060938 | 2.118935 | 2.213807 | 2.316957 | 2.418572 | 2.509394 | 2.563707 | 2.610905 | 2.665996 | 2.71526 |
|  | 90 | 1.82703 | 1.886143 | 1.950521 | 2.004304 | 2.064709 | 2.162376 | 2.267014 | 2.36857 | 2.458087 | 2.511061 | 2.55676 | 2.609707 | 2.656695 |
| **R_cuneus** | 3 | 1.94035 | 1.9857 | 2.037482 | 2.082676 | 2.13554 | 2.225699 | 2.328574 | 2.434322 | 2.53198 | 2.591558 | 2.643924 | 2.705624 | 2.761212 |
|  | 5 | 1.880894 | 1.925066 | 1.975539 | 2.01962 | 2.071217 | 2.159302 | 2.259941 | 2.363533 | 2.459323 | 2.517819 | 2.56927 | 2.629934 | 2.684628 |
|  | 10 | 1.74783 | 1.789366 | 1.836909 | 1.878503 | 1.927275 | 2.010743 | 2.106424 | 2.205259 | 2.296959 | 2.3531 | 2.402568 | 2.461 | 2.513778 |
|  | 15 | 1.643646 | 1.683156 | 1.728461 | 1.768168 | 1.814808 | 1.894835 | 1.986887 | 2.082321 | 2.171177 | 2.225722 | 2.273876 | 2.330862 | 2.382435 |
|  | 20 | 1.576949 | 1.615236 | 1.659218 | 1.697834 | 1.743275 | 1.821448 | 1.911684 | 2.005586 | 2.093334 | 2.147349 | 2.195128 | 2.251785 | 2.303164 |
|  | 30 | 1.517329 | 1.554401 | 1.597131 | 1.634774 | 1.679222 | 1.756072 | 1.845383 | 1.939009 | 2.027127 | 2.081673 | 2.130112 | 2.187783 | 2.240299 |
|  | 40 | 1.488086 | 1.52396 | 1.565428 | 1.602066 | 1.645458 | 1.720814 | 1.808924 | 1.901913 | 1.990017 | 2.04484 | 2.093708 | 2.152115 | 2.205516 |
|  | 50 | 1.469198 | 1.503992 | 1.54432 | 1.580048 | 1.622482 | 1.696486 | 1.783524 | 1.875986 | 1.964172 | 2.019334 | 2.068692 | 2.127917 | 2.182288 |
|  | 60 | 1.455468 | 1.48956 | 1.529185 | 1.564392 | 1.606331 | 1.679801 | 1.766757 | 1.859788 | 1.949158 | 2.005382 | 2.055904 | 2.116793 | 2.17295 |
|  | 70 | 1.436084 | 1.46957 | 1.508608 | 1.543399 | 1.584975 | 1.658163 | 1.745377 | 1.839414 | 1.930474 | 1.988132 | 2.040188 | 2.10324 | 2.161699 |
|  | 80 | 1.406451 | 1.439161 | 1.477411 | 1.511606 | 1.552606 | 1.625143 | 1.712202 | 1.806849 | 1.899291 | 1.958234 | 2.01173 | 2.076885 | 2.137648 |
|  | 90 | 1.376191 | 1.408091 | 1.445506 | 1.47906 | 1.519422 | 1.591195 | 1.677969 | 1.773115 | 1.866888 | 1.927126 | 1.982106 | 2.049469 | 2.112696 |
| **R_entorhinal** | 3 | 2.260363 | 2.416479 | 2.582717 | 2.719838 | 2.873207 | 3.122961 | 3.395293 | 3.642734 | 3.832354 | 3.934113 | 4.016765 | 4.107386 | 4.183726 |
|  | 5 | 2.284518 | 2.437203 | 2.600788 | 2.73634 | 2.888469 | 3.137005 | 3.408839 | 3.657517 | 3.84994 | 3.953866 | 4.038591 | 4.131782 | 4.210513 |
|  | 10 | 2.342913 | 2.488172 | 2.645894 | 2.777924 | 2.927224 | 3.172919 | 3.443476 | 3.695174 | 3.89477 | 4.004337 | 4.094513 | 4.194525 | 4.27965 |
|  | 15 | 2.394701 | 2.534205 | 2.687278 | 2.816459 | 2.963428 | 3.206672 | 3.47591 | 3.73036 | 3.937086 | 4.052423 | 4.148262 | 4.255457 | 4.347396 |
|  | 20 | 2.424817 | 2.560289 | 2.709981 | 2.836987 | 2.982046 | 3.22292 | 3.490193 | 3.746048 | 3.958422 | 4.078638 | 4.1794 | 4.29297 | 4.391063 |
|  | 30 | 2.424834 | 2.555635 | 2.700953 | 2.824712 | 2.966365 | 3.201653 | 3.462228 | 3.715302 | 3.931515 | 4.056338 | 4.162207 | 4.282805 | 4.387979 |
|  | 40 | 2.437513 | 2.562663 | 2.702409 | 2.821905 | 2.959096 | 3.187588 | 3.441119 | 3.689995 | 3.906537 | 4.033122 | 4.141315 | 4.265424 | 4.374367 |
|  | 50 | 2.460586 | 2.581662 | 2.71723 | 2.833432 | 2.967119 | 3.190336 | 3.438695 | 3.683589 | 3.897896 | 4.023673 | 4.131445 | 4.255359 | 4.364372 |
|  | 60 | 2.438826 | 2.564526 | 2.704514 | 2.82392 | 2.960688 | 3.187752 | 3.438709 | 3.684266 | 3.897451 | 4.021883 | 4.128127 | 4.249879 | 4.356648 |
|  | 70 | 2.301576 | 2.446291 | 2.60415 | 2.736389 | 2.885458 | 3.128083 | 3.390269 | 3.640648 | 3.852862 | 3.974734 | 4.077731 | 4.19464 | 4.296235 |
|  | 80 | 1.9954 | 2.184405 | 2.380486 | 2.538275 | 2.710409 | 2.980211 | 3.260477 | 3.517919 | 3.728386 | 3.846458 | 3.944825 | 4.055031 | 4.149643 |
|  | 90 | 1.550417 | 1.823425 | 2.087692 | 2.287517 | 2.494839 | 2.802649 | 3.106188 | 3.372308 | 3.581156 | 3.695397 | 3.789165 | 3.892839 | 3.980777 |
| **R_fusiform** | 3 | 2.59874 | 2.672242 | 2.747102 | 2.806192 | 2.869467 | 2.966351 | 3.064937 | 3.157736 | 3.238846 | 3.28713 | 3.32922 | 3.378743 | 3.423592 |
|  | 5 | 2.552424 | 2.629118 | 2.706425 | 2.766931 | 2.831267 | 2.928957 | 3.027488 | 3.11956 | 3.199595 | 3.247075 | 3.288379 | 3.336887 | 3.380743 |
|  | 10 | 2.421509 | 2.51057 | 2.596959 | 2.66255 | 2.730603 | 2.831091 | 2.929555 | 3.019456 | 3.096282 | 3.14138 | 3.180366 | 3.225895 | 3.266849 |
|  | 15 | 2.347756 | 2.428917 | 2.50902 | 2.570668 | 2.635338 | 2.732085 | 2.828308 | 2.917437 | 2.994633 | 3.040423 | 3.080303 | 3.12724 | 3.169804 |
|  | 20 | 2.351466 | 2.409534 | 2.47073 | 2.520505 | 2.575267 | 2.662154 | 2.754609 | 2.845775 | 2.929158 | 2.980606 | 3.026643 | 3.082314 | 3.13421 |
|  | 30 | 2.281914 | 2.340318 | 2.402226 | 2.4528 | 2.508596 | 2.597213 | 2.691134 | 2.782704 | 2.864987 | 2.914858 | 2.958823 | 3.011094 | 3.058896 |
|  | 40 | 2.211558 | 2.27365 | 2.339308 | 2.392775 | 2.451506 | 2.543974 | 2.640355 | 2.732031 | 2.81191 | 2.858979 | 2.899557 | 2.946626 | 2.988517 |
|  | 50 | 2.222243 | 2.27068 | 2.324547 | 2.37043 | 2.422909 | 2.509843 | 2.605795 | 2.701651 | 2.788332 | 2.84058 | 2.886215 | 2.939733 | 2.987805 |
|  | 60 | 2.179837 | 2.232081 | 2.289886 | 2.338859 | 2.394545 | 2.485953 | 2.585471 | 2.683259 | 2.770123 | 2.821704 | 2.866253 | 2.917877 | 2.963655 |
|  | 70 | 2.139762 | 2.189765 | 2.245703 | 2.293573 | 2.348517 | 2.439808 | 2.540615 | 2.640927 | 2.730925 | 2.784707 | 2.83133 | 2.885534 | 2.933737 |
|  | 80 | 2.10115 | 2.146627 | 2.198352 | 2.243328 | 2.295754 | 2.384748 | 2.485725 | 2.588985 | 2.683939 | 2.741708 | 2.792398 | 2.852034 | 2.905692 |
|  | 90 | 2.050449 | 2.091568 | 2.139033 | 2.180917 | 2.230469 | 2.316416 | 2.416797 | 2.522683 | 2.623023 | 2.685493 | 2.741214 | 2.807867 | 2.86887 |
| **R_inferiorparietal** | 3 | 2.454437 | 2.550891 | 2.64364 | 2.713463 | 2.78515 | 2.888741 | 2.986082 | 3.072083 | 3.143798 | 3.184471 | 3.218488 | 3.256657 | 3.289415 |
|  | 5 | 2.424199 | 2.5112 | 2.596915 | 2.662633 | 2.731057 | 2.831439 | 2.927176 | 3.012794 | 3.084891 | 3.126038 | 3.160586 | 3.199489 | 3.232988 |
|  | 10 | 2.339662 | 2.408512 | 2.479773 | 2.536575 | 2.597592 | 2.69033 | 2.782051 | 2.866675 | 2.939801 | 2.982258 | 3.018292 | 3.059275 | 3.094901 |
|  | 15 | 2.250801 | 2.307118 | 2.367513 | 2.417134 | 2.471846 | 2.557673 | 2.645566 | 2.729281 | 2.80366 | 2.847677 | 2.885501 | 2.929029 | 2.967294 |
|  | 20 | 2.182032 | 2.231351 | 2.285374 | 2.330614 | 2.381365 | 2.462758 | 2.54827 | 2.631807 | 2.707802 | 2.753541 | 2.793289 | 2.839528 | 2.880611 |
|  | 30 | 2.09386 | 2.142684 | 2.196309 | 2.241317 | 2.291902 | 2.373181 | 2.458702 | 2.542725 | 2.619893 | 2.666646 | 2.707449 | 2.755107 | 2.797612 |
|  | 40 | 2.041816 | 2.092347 | 2.147694 | 2.194029 | 2.245981 | 2.329195 | 2.416419 | 2.50182 | 2.580027 | 2.627309 | 2.668513 | 2.71657 | 2.759369 |
|  | 50 | 2.012702 | 2.061118 | 2.114644 | 2.159866 | 2.211035 | 2.294089 | 2.382704 | 2.470204 | 2.550235 | 2.598614 | 2.640792 | 2.690025 | 2.733923 |
|  | 60 | 1.974504 | 2.019708 | 2.070255 | 2.113462 | 2.162955 | 2.244829 | 2.334535 | 2.423806 | 2.504529 | 2.55298 | 2.595066 | 2.64405 | 2.687628 |
|  | 70 | 1.931382 | 1.972901 | 2.019938 | 2.060716 | 2.108157 | 2.188643 | 2.280061 | 2.371485 | 2.451984 | 2.499437 | 2.540227 | 2.58728 | 2.628816 |
|  | 80 | 1.897934 | 1.934476 | 1.976669 | 2.014004 | 2.058436 | 2.136653 | 2.230171 | 2.324814 | 2.405738 | 2.452448 | 2.49212 | 2.537429 | 2.577084 |
|  | 90 | 1.865461 | 1.896578 | 1.933304 | 1.966592 | 2.007286 | 2.082173 | 2.177322 | 2.275691 | 2.357926 | 2.404531 | 2.443715 | 2.488103 | 2.526697 |
| **R_inferiortemporal** | 3 | 2.573664 | 2.656434 | 2.742114 | 2.810642 | 2.884779 | 2.999407 | 3.116635 | 3.226482 | 3.321218 | 3.376733 | 3.424457 | 3.479701 | 3.528802 |
|  | 5 | 2.531456 | 2.617255 | 2.705279 | 2.77516 | 2.850285 | 2.965559 | 3.082449 | 3.191149 | 3.284301 | 3.338645 | 3.385225 | 3.438989 | 3.48664 |
|  | 10 | 2.428313 | 2.521393 | 2.614802 | 2.687651 | 2.764829 | 2.881228 | 2.997063 | 3.103036 | 3.192637 | 3.244425 | 3.288542 | 3.339162 | 3.383763 |
|  | 15 | 2.382848 | 2.465009 | 2.549467 | 2.616625 | 2.688916 | 2.799988 | 2.912727 | 3.017577 | 3.107363 | 3.159686 | 3.204487 | 3.256135 | 3.301843 |
|  | 20 | 2.393156 | 2.45328 | 2.518682 | 2.573315 | 2.634746 | 2.73446 | 2.842298 | 2.948587 | 3.044181 | 3.101827 | 3.152314 | 3.2118 | 3.265586 |
|  | 30 | 2.330314 | 2.388341 | 2.451824 | 2.50513 | 2.565351 | 2.663692 | 2.770775 | 2.87695 | 2.97288 | 3.030892 | 3.081783 | 3.141832 | 3.196193 |
|  | 40 | 2.279066 | 2.340506 | 2.407243 | 2.462903 | 2.525378 | 2.626491 | 2.735334 | 2.841968 | 2.937211 | 2.9943 | 3.044067 | 3.102411 | 3.154875 |
|  | 50 | 2.262421 | 2.322355 | 2.387959 | 2.443059 | 2.505301 | 2.606876 | 2.717274 | 2.826378 | 2.924521 | 2.983625 | 3.0353 | 3.096044 | 3.150804 |
|  | 60 | 2.227361 | 2.288282 | 2.355009 | 2.41108 | 2.474443 | 2.577866 | 2.690252 | 2.801226 | 2.900912 | 2.960859 | 3.013207 | 3.074657 | 3.129965 |
|  | 70 | 2.193909 | 2.25032 | 2.312982 | 2.366333 | 2.427373 | 2.528675 | 2.641022 | 2.754174 | 2.857602 | 2.920577 | 2.976031 | 3.041659 | 3.101206 |
|  | 80 | 2.184579 | 2.232892 | 2.287855 | 2.33575 | 2.391814 | 2.487922 | 2.599128 | 2.716227 | 2.827903 | 2.898138 | 2.961423 | 3.038093 | 3.109357 |
|  | 90 | 2.182158 | 2.224122 | 2.272798 | 2.31606 | 2.367744 | 2.459101 | 2.569473 | 2.691589 | 2.814234 | 2.894728 | 2.96964 | 3.063617 | 3.154338 |
| **R_isthmuscingulate** | 3 | 2.488597 | 2.560976 | 2.638168 | 2.701777 | 2.772823 | 2.888626 | 3.017853 | 3.154273 | 3.290099 | 3.38072 | 3.467122 | 3.579366 | 3.692781 |
|  | 5 | 2.43879 | 2.508184 | 2.582864 | 2.644892 | 2.714643 | 2.829179 | 2.957721 | 3.093467 | 3.227915 | 3.316941 | 3.401201 | 3.509666 | 3.618066 |
|  | 10 | 2.313392 | 2.37699 | 2.446719 | 2.505589 | 2.572714 | 2.684632 | 2.811719 | 2.946067 | 3.077767 | 3.163666 | 3.243777 | 3.345021 | 3.44399 |
|  | 15 | 2.198248 | 2.257975 | 2.324397 | 2.381174 | 2.446596 | 2.556949 | 2.683397 | 2.817226 | 2.947473 | 3.031506 | 3.109039 | 3.205718 | 3.298706 |
|  | 20 | 2.113175 | 2.170474 | 2.234878 | 2.290439 | 2.354963 | 2.464739 | 2.591371 | 2.72551 | 2.855359 | 2.938457 | 3.014517 | 3.108414 | 3.197638 |
|  | 30 | 2.024413 | 2.07868 | 2.140475 | 2.194385 | 2.257577 | 2.366166 | 2.492318 | 2.625869 | 2.754038 | 2.835096 | 2.908452 | 2.997752 | 3.081195 |
|  | 40 | 1.96982 | 2.02243 | 2.082741 | 2.135653 | 2.197958 | 2.30552 | 2.430815 | 2.563235 | 2.689544 | 2.768819 | 2.840058 | 2.926042 | 3.00558 |
|  | 50 | 1.912825 | 1.965327 | 2.025748 | 2.07893 | 2.141721 | 2.250413 | 2.377229 | 2.51114 | 2.63844 | 2.718 | 2.789212 | 2.874756 | 2.95344 |
|  | 60 | 1.854386 | 1.907489 | 1.968739 | 2.02275 | 2.086615 | 2.19733 | 2.326611 | 2.463037 | 2.592456 | 2.673132 | 2.745175 | 2.831472 | 2.910584 |
|  | 70 | 1.809215 | 1.863314 | 1.925776 | 1.980901 | 2.04612 | 2.159235 | 2.291303 | 2.430529 | 2.562358 | 2.644374 | 2.717487 | 2.80489 | 2.884829 |
|  | 80 | 1.758094 | 1.814025 | 1.878657 | 1.935736 | 2.003302 | 2.120543 | 2.257453 | 2.401723 | 2.538199 | 2.623012 | 2.698544 | 2.788733 | 2.87111 |
|  | 90 | 1.721878 | 1.780932 | 1.849223 | 1.90957 | 1.98104 | 2.105119 | 2.250061 | 2.40278 | 2.547175 | 2.636851 | 2.716666 | 2.811901 | 2.898813 |
| **R_lateraloccipital** | 3 | 2.221443 | 2.263544 | 2.311119 | 2.352241 | 2.39991 | 2.480251 | 2.570665 | 2.662475 | 2.746468 | 2.79742 | 2.842063 | 2.894528 | 2.941704 |
|  | 5 | 2.17828 | 2.219881 | 2.266867 | 2.307461 | 2.354499 | 2.433742 | 2.522887 | 2.613397 | 2.696211 | 2.74646 | 2.790499 | 2.842272 | 2.888846 |
|  | 10 | 2.075177 | 2.115623 | 2.161232 | 2.200582 | 2.246127 | 2.322755 | 2.408868 | 2.496272 | 2.576282 | 2.624871 | 2.667492 | 2.717654 | 2.762838 |
|  | 15 | 1.986733 | 2.026324 | 2.070879 | 2.109253 | 2.153604 | 2.228106 | 2.311724 | 2.396572 | 2.474302 | 2.521566 | 2.563074 | 2.612002 | 2.656156 |
|  | 20 | 1.928447 | 1.967835 | 2.012049 | 2.050046 | 2.093881 | 2.16737 | 2.249721 | 2.333258 | 2.409865 | 2.456521 | 2.49756 | 2.546029 | 2.589874 |
|  | 30 | 1.875235 | 1.915875 | 1.961184 | 1.999893 | 2.044326 | 2.118393 | 2.200979 | 2.284588 | 2.361354 | 2.408241 | 2.449612 | 2.49867 | 2.543267 |
|  | 40 | 1.846876 | 1.889444 | 1.936576 | 1.976601 | 2.022302 | 2.09802 | 2.18197 | 2.266706 | 2.344507 | 2.392113 | 2.434213 | 2.484287 | 2.529983 |
|  | 50 | 1.826895 | 1.870403 | 1.918491 | 1.959267 | 2.005773 | 2.082741 | 2.168033 | 2.254183 | 2.333428 | 2.382021 | 2.425076 | 2.476402 | 2.523364 |
|  | 60 | 1.799346 | 1.841894 | 1.88916 | 1.929428 | 1.975558 | 2.052343 | 2.138017 | 2.225094 | 2.305602 | 2.355135 | 2.399116 | 2.451646 | 2.499796 |
|  | 70 | 1.758885 | 1.79908 | 1.844153 | 1.882888 | 1.927618 | 2.00286 | 2.087837 | 2.175133 | 2.256507 | 2.306825 | 2.351631 | 2.405277 | 2.454546 |
|  | 80 | 1.723043 | 1.76066 | 1.803293 | 1.840298 | 1.883435 | 1.956914 | 2.04116 | 2.128924 | 2.211683 | 2.263249 | 2.309386 | 2.364867 | 2.416024 |
|  | 90 | 1.645173 | 1.679481 | 1.71875 | 1.753159 | 1.793641 | 1.863477 | 1.944833 | 2.030937 | 2.113284 | 2.165117 | 2.211812 | 2.26834 | 2.32081 |
| **R_lateralorbitofrontal** | 3 | 2.595008 | 2.695363 | 2.795215 | 2.872573 | 2.954138 | 3.076725 | 3.198833 | 3.311454 | 3.408063 | 3.464756 | 3.513675 | 3.570634 | 3.62166 |
|  | 5 | 2.548074 | 2.639867 | 2.732698 | 2.805563 | 2.883221 | 3.001413 | 3.120765 | 3.232155 | 3.328639 | 3.385637 | 3.435036 | 3.492799 | 3.544762 |
|  | 10 | 2.438647 | 2.513197 | 2.591247 | 2.654323 | 2.723252 | 2.831432 | 2.944543 | 3.053498 | 3.150439 | 3.208803 | 3.260035 | 3.320694 | 3.375948 |
|  | 15 | 2.349814 | 2.411783 | 2.478485 | 2.533748 | 2.595526 | 2.695393 | 2.803617 | 2.911547 | 3.010627 | 3.071669 | 3.126114 | 3.191626 | 3.252291 |
|  | 20 | 2.286435 | 2.340536 | 2.399876 | 2.449921 | 2.506822 | 2.600984 | 2.706133 | 2.814308 | 2.91661 | 2.981099 | 3.039579 | 3.111163 | 3.178654 |
|  | 30 | 2.198609 | 2.249901 | 2.306455 | 2.354396 | 2.409181 | 2.500501 | 2.603476 | 2.710556 | 2.812919 | 2.878011 | 2.937419 | 3.010643 | 3.080192 |
|  | 40 | 2.130826 | 2.185976 | 2.246057 | 2.2964 | 2.353285 | 2.446612 | 2.549676 | 2.654491 | 2.752528 | 2.813806 | 2.869033 | 2.936205 | 2.999113 |
|  | 50 | 2.102853 | 2.155784 | 2.213828 | 2.262766 | 2.318387 | 2.410358 | 2.512915 | 2.618209 | 2.717537 | 2.780012 | 2.836561 | 2.905641 | 2.970625 |
|  | 60 | 2.098833 | 2.147924 | 2.20248 | 2.249064 | 2.302665 | 2.392822 | 2.495578 | 2.603483 | 2.70747 | 2.773952 | 2.834841 | 2.910136 | 2.981882 |
|  | 70 | 2.062649 | 2.114391 | 2.171745 | 2.220587 | 2.276617 | 2.37041 | 2.476527 | 2.586937 | 2.692236 | 2.758953 | 2.819626 | 2.894073 | 2.964403 |
|  | 80 | 2.029625 | 2.088597 | 2.153299 | 2.207844 | 2.269788 | 2.371968 | 2.485273 | 2.600547 | 2.707997 | 2.774825 | 2.834765 | 2.90724 | 2.974638 |
|  | 90 | 2.024677 | 2.087915 | 2.157308 | 2.215805 | 2.28222 | 2.391684 | 2.512836 | 2.635717 | 2.749802 | 2.820493 | 2.883707 | 2.959882 | 3.030455 |
| **R_lingual** | 3 | 2.191826 | 2.229774 | 2.273138 | 2.311016 | 2.355365 | 2.431116 | 2.51775 | 2.607047 | 2.68975 | 2.740324 | 2.784856 | 2.83742 | 2.88487 |
|  | 5 | 2.125239 | 2.162553 | 2.205208 | 2.242481 | 2.286138 | 2.360746 | 2.446133 | 2.534208 | 2.615839 | 2.665785 | 2.709778 | 2.761728 | 2.808642 |
|  | 10 | 1.9779 | 2.013853 | 2.054993 | 2.090975 | 2.133161 | 2.205354 | 2.288125 | 2.373667 | 2.453095 | 2.501761 | 2.544669 | 2.595386 | 2.641233 |
|  | 15 | 1.862439 | 1.89744 | 1.937529 | 1.972628 | 2.013816 | 2.084403 | 2.165483 | 2.249443 | 2.32755 | 2.375474 | 2.417772 | 2.467819 | 2.513107 |
|  | 20 | 1.793611 | 1.828256 | 1.867975 | 1.90278 | 1.943663 | 2.013818 | 2.094544 | 2.178295 | 2.256346 | 2.304302 | 2.346668 | 2.396845 | 2.442296 |
|  | 30 | 1.739659 | 1.774146 | 1.813737 | 1.848479 | 1.889341 | 1.959601 | 2.040665 | 2.125006 | 2.203824 | 2.252355 | 2.295293 | 2.346225 | 2.392432 |
|  | 40 | 1.706181 | 1.73996 | 1.77877 | 1.812855 | 1.852981 | 1.922061 | 2.0019 | 2.085123 | 2.163041 | 2.211085 | 2.253637 | 2.304164 | 2.350054 |
|  | 50 | 1.678692 | 1.711867 | 1.750017 | 1.783549 | 1.823058 | 1.891161 | 1.970007 | 2.052349 | 2.129585 | 2.177277 | 2.219561 | 2.269824 | 2.315522 |
|  | 60 | 1.647627 | 1.680657 | 1.718685 | 1.752151 | 1.791629 | 1.859802 | 1.938919 | 2.021762 | 2.099669 | 2.147872 | 2.19067 | 2.241618 | 2.28801 |
|  | 70 | 1.60914 | 1.641968 | 1.679813 | 1.713161 | 1.752553 | 1.820709 | 1.900018 | 1.983302 | 2.061847 | 2.110552 | 2.153864 | 2.205508 | 2.252612 |
|  | 80 | 1.576237 | 1.608917 | 1.646641 | 1.679927 | 1.719298 | 1.787553 | 1.867191 | 1.951067 | 2.030401 | 2.079707 | 2.123625 | 2.176079 | 2.224005 |
|  | 90 | 1.529637 | 1.561885 | 1.59916 | 1.632095 | 1.671106 | 1.738876 | 1.818172 | 1.901949 | 1.981431 | 2.030949 | 2.075133 | 2.127998 | 2.176388 |
| **R_medialorbitofrontal** | 3 | 2.599925 | 2.660059 | 2.727616 | 2.785766 | 2.853009 | 2.966283 | 3.09436 | 3.22596 | 3.348535 | 3.424236 | 3.491565 | 3.572054 | 3.645842 |
|  | 5 | 2.502594 | 2.561508 | 2.62763 | 2.684499 | 2.75022 | 2.860878 | 2.985994 | 3.114654 | 3.234681 | 3.308936 | 3.375078 | 3.454287 | 3.527048 |
|  | 10 | 2.295393 | 2.351999 | 2.415338 | 2.46968 | 2.532363 | 2.637733 | 2.756833 | 2.879564 | 2.994558 | 3.066045 | 3.129992 | 3.206956 | 3.278066 |
|  | 15 | 2.153059 | 2.208919 | 2.271177 | 2.32442 | 2.385681 | 2.488424 | 2.604467 | 2.724311 | 2.837168 | 2.907731 | 2.971174 | 3.047993 | 3.119468 |
|  | 20 | 2.064933 | 2.121305 | 2.183826 | 2.237075 | 2.298144 | 2.400246 | 2.515407 | 2.634601 | 2.747483 | 2.818528 | 2.882786 | 2.961143 | 3.034653 |
|  | 30 | 1.967573 | 2.025285 | 2.08849 | 2.141749 | 2.202302 | 2.302652 | 2.415262 | 2.532196 | 2.644236 | 2.715769 | 2.781326 | 2.862546 | 2.940173 |
|  | 40 | 1.927427 | 1.986968 | 2.051183 | 2.104584 | 2.164637 | 2.26303 | 2.372618 | 2.486669 | 2.597286 | 2.669027 | 2.735744 | 2.819887 | 2.902018 |
|  | 50 | 1.901079 | 1.961542 | 2.026051 | 2.079199 | 2.138508 | 2.234901 | 2.341715 | 2.453109 | 2.562168 | 2.633745 | 2.70105 | 2.787082 | 2.872398 |
|  | 60 | 1.876794 | 1.936171 | 1.999835 | 2.05252 | 2.111546 | 2.207942 | 2.315284 | 2.427602 | 2.537727 | 2.610013 | 2.677955 | 2.764731 | 2.850688 |
|  | 70 | 1.863764 | 1.92131 | 1.984164 | 2.037038 | 2.09712 | 2.19684 | 2.309457 | 2.427907 | 2.543526 | 2.618711 | 2.688687 | 2.776941 | 2.863023 |
|  | 80 | 1.871661 | 1.927807 | 1.99036 | 2.043909 | 2.105686 | 2.210013 | 2.329632 | 2.456163 | 2.579091 | 2.658231 | 2.731105 | 2.821756 | 2.908692 |
|  | 90 | 1.852912 | 1.907558 | 1.969377 | 2.02302 | 2.085637 | 2.19285 | 2.317326 | 2.449757 | 2.578175 | 2.660337 | 2.735463 | 2.828043 | 2.915796 |
| **R_middletemporal** | 3 | 2.748278 | 2.844401 | 2.941897 | 3.018272 | 3.09905 | 3.219339 | 3.33534 | 3.44166 | 3.534524 | 3.588785 | 3.634991 | 3.687681 | 3.733576 |
|  | 5 | 2.701777 | 2.798662 | 2.896547 | 2.972999 | 3.053676 | 3.173536 | 3.288894 | 3.394313 | 3.486038 | 3.5395 | 3.584958 | 3.636723 | 3.681756 |
|  | 10 | 2.591561 | 2.688793 | 2.786419 | 2.862321 | 2.942155 | 3.0604 | 3.173963 | 3.277223 | 3.366388 | 3.418105 | 3.461951 | 3.511751 | 3.554973 |
|  | 15 | 2.541358 | 2.621334 | 2.704859 | 2.771916 | 2.844356 | 2.955092 | 3.065147 | 3.167795 | 3.257933 | 3.310828 | 3.356017 | 3.407721 | 3.452915 |
|  | 20 | 2.538954 | 2.5964 | 2.659984 | 2.713748 | 2.774611 | 2.873398 | 2.978701 | 3.083104 | 3.179436 | 3.238024 | 3.289299 | 3.349367 | 3.403107 |
|  | 30 | 2.460249 | 2.513345 | 2.572709 | 2.623407 | 2.681371 | 2.77681 | 2.88049 | 2.984569 | 3.081088 | 3.140045 | 3.191822 | 3.252704 | 3.307389 |
|  | 40 | 2.376208 | 2.438291 | 2.505957 | 2.5624 | 2.625543 | 2.726594 | 2.832729 | 2.93526 | 3.026589 | 3.080775 | 3.127442 | 3.181286 | 3.228756 |
|  | 50 | 2.367288 | 2.422102 | 2.483171 | 2.535184 | 2.594544 | 2.692208 | 2.798431 | 2.903721 | 2.998902 | 3.056007 | 3.105584 | 3.163252 | 3.214518 |
|  | 60 | 2.300794 | 2.358857 | 2.423115 | 2.477537 | 2.539363 | 2.640614 | 2.750317 | 2.857351 | 2.951606 | 3.007132 | 3.054783 | 3.109615 | 3.157863 |
|  | 70 | 2.261181 | 2.31375 | 2.373059 | 2.424254 | 2.483544 | 2.583416 | 2.69574 | 2.807664 | 2.906512 | 2.964897 | 3.015134 | 3.073128 | 3.124346 |
|  | 80 | 2.226267 | 2.27571 | 2.332269 | 2.381793 | 2.440022 | 2.540446 | 2.657111 | 2.775044 | 2.87864 | 2.939613 | 2.992007 | 3.052456 | 3.105846 |
|  | 90 | 2.160647 | 2.215375 | 2.277419 | 2.331325 | 2.394294 | 2.502121 | 2.626369 | 2.748488 | 2.851114 | 2.909653 | 2.958969 | 3.014832 | 3.063323 |
| **R_parahippocampal** | 3 | 2.259482 | 2.355013 | 2.460732 | 2.550285 | 2.652068 | 2.818907 | 2.999963 | 3.176936 | 3.333052 | 3.425154 | 3.504279 | 3.59543 | 3.675729 |
|  | 5 | 2.238003 | 2.332956 | 2.438092 | 2.527196 | 2.628514 | 2.794694 | 2.975169 | 3.151699 | 3.307521 | 3.39949 | 3.478523 | 3.569594 | 3.649845 |
|  | 10 | 2.185137 | 2.278641 | 2.382313 | 2.470288 | 2.570441 | 2.734967 | 2.913987 | 3.089413 | 3.244508 | 3.33615 | 3.414962 | 3.505843 | 3.585984 |
|  | 15 | 2.134459 | 2.226507 | 2.328718 | 2.41557 | 2.514571 | 2.677481 | 2.855108 | 3.029514 | 3.183973 | 3.275352 | 3.354001 | 3.444765 | 3.524865 |
|  | 20 | 2.089574 | 2.180302 | 2.281208 | 2.367081 | 2.465102 | 2.626699 | 2.803295 | 2.977068 | 3.131259 | 3.222603 | 3.301292 | 3.392183 | 3.472462 |
|  | 30 | 2.015546 | 2.104449 | 2.203612 | 2.288228 | 2.385059 | 2.545237 | 2.721009 | 2.894663 | 3.049291 | 3.141122 | 3.220363 | 3.312039 | 3.393141 |
|  | 40 | 1.954144 | 2.042756 | 2.141716 | 2.226256 | 2.323105 | 2.483548 | 2.659927 | 2.83449 | 2.990169 | 3.082728 | 3.162656 | 3.255195 | 3.337118 |
|  | 50 | 1.910303 | 2.001165 | 2.102483 | 2.188914 | 2.287801 | 2.451334 | 2.630736 | 2.807937 | 2.965696 | 3.059378 | 3.140211 | 3.233725 | 3.316448 |
|  | 60 | 1.866725 | 1.961939 | 2.067618 | 2.15739 | 2.259697 | 2.428018 | 2.611535 | 2.791741 | 2.951367 | 3.045822 | 3.127132 | 3.220984 | 3.303824 |
|  | 70 | 1.804321 | 1.904516 | 2.014961 | 2.108201 | 2.213862 | 2.386432 | 2.572962 | 2.754659 | 2.914508 | 3.008646 | 3.08943 | 3.182397 | 3.264217 |
|  | 80 | 1.731063 | 1.836974 | 1.952689 | 2.04962 | 2.158699 | 2.33528 | 2.524204 | 2.706526 | 2.865683 | 2.958914 | 3.038643 | 3.130094 | 3.210324 |
|  | 90 | 1.653178 | 1.765922 | 1.887756 | 1.988854 | 2.101689 | 2.282503 | 2.473749 | 2.656441 | 2.814596 | 2.906719 | 2.985216 | 3.074945 | 3.153406 |
| **R_paracentral** | 3 | 2.353051 | 2.407347 | 2.467859 | 2.519505 | 2.578689 | 2.676984 | 2.785821 | 2.894882 | 2.99377 | 3.053489 | 3.105715 | 3.167037 | 3.222183 |
|  | 5 | 2.315116 | 2.369029 | 2.429083 | 2.480316 | 2.539002 | 2.636428 | 2.744254 | 2.852275 | 2.950215 | 3.009367 | 3.061102 | 3.121861 | 3.176513 |
|  | 10 | 2.217686 | 2.27056 | 2.329376 | 2.379493 | 2.436839 | 2.531922 | 2.637029 | 2.742251 | 2.837638 | 2.895259 | 2.945673 | 3.004906 | 3.058218 |
|  | 15 | 2.121752 | 2.173596 | 2.231177 | 2.280174 | 2.336174 | 2.428895 | 2.531249 | 2.63363 | 2.72642 | 2.782483 | 2.831549 | 2.889226 | 2.941171 |
|  | 20 | 2.047671 | 2.099016 | 2.155944 | 2.20431 | 2.259513 | 2.350766 | 2.45134 | 2.551837 | 2.642888 | 2.697907 | 2.746074 | 2.80272 | 2.85377 |
|  | 30 | 1.962673 | 2.015089 | 2.0729 | 2.121789 | 2.177358 | 2.268744 | 2.368917 | 2.468604 | 2.558703 | 2.613098 | 2.660711 | 2.716721 | 2.767228 |
|  | 40 | 1.90225 | 1.957384 | 2.017701 | 2.068336 | 2.125513 | 2.218772 | 2.320078 | 2.420152 | 2.510145 | 2.564332 | 2.611704 | 2.667387 | 2.717583 |
|  | 50 | 1.850575 | 1.909297 | 1.972912 | 2.025854 | 2.085175 | 2.181016 | 2.284063 | 2.385038 | 2.475358 | 2.529601 | 2.576969 | 2.632619 | 2.682784 |
|  | 60 | 1.803752 | 1.865271 | 1.931429 | 1.986141 | 2.047117 | 2.14502 | 2.249664 | 2.351847 | 2.443172 | 2.498065 | 2.546068 | 2.602576 | 2.65365 |
|  | 70 | 1.751961 | 1.814149 | 1.880859 | 1.935921 | 1.997207 | 2.095532 | 2.200723 | 2.303793 | 2.396433 | 2.45244 | 2.501657 | 2.559917 | 2.612908 |
|  | 80 | 1.70153 | 1.763231 | 1.829426 | 1.88409 | 1.944991 | 2.04293 | 2.148234 | 2.252228 | 2.346647 | 2.404267 | 2.455282 | 2.516179 | 2.572082 |
|  | 90 | 1.647558 | 1.708673 | 1.77421 | 1.828334 | 1.888675 | 1.985923 | 2.09103 | 2.195738 | 2.29191 | 2.351242 | 2.404241 | 2.468132 | 2.527433 |
| **R_parsopercularis** | 3 | 2.515322 | 2.581776 | 2.652634 | 2.710797 | 2.775197 | 2.8778 | 2.986619 | 3.092324 | 3.186581 | 3.243227 | 3.292801 | 3.351239 | 3.404165 |
|  | 5 | 2.487495 | 2.552319 | 2.621716 | 2.678883 | 2.742381 | 2.843949 | 2.952146 | 3.057643 | 3.151983 | 3.20878 | 3.258536 | 3.317242 | 3.370451 |
|  | 10 | 2.419076 | 2.479985 | 2.545832 | 2.600552 | 2.661812 | 2.760781 | 2.867409 | 2.972407 | 3.067024 | 3.124267 | 3.174561 | 3.234058 | 3.28811 |
|  | 15 | 2.352594 | 2.409712 | 2.472062 | 2.524331 | 2.583321 | 2.679616 | 2.784626 | 2.889164 | 2.984206 | 3.042043 | 3.093048 | 3.15359 | 3.208768 |
|  | 20 | 2.289652 | 2.343421 | 2.402628 | 2.452664 | 2.509556 | 2.603339 | 2.706805 | 2.810921 | 2.906431 | 2.964907 | 3.016677 | 3.078353 | 3.134761 |
|  | 30 | 2.188069 | 2.238639 | 2.294829 | 2.342711 | 2.397569 | 2.488898 | 2.590801 | 2.694359 | 2.790072 | 2.848938 | 2.90119 | 2.963576 | 3.020733 |
|  | 40 | 2.11093 | 2.161765 | 2.218268 | 2.266414 | 2.321555 | 2.413245 | 2.515274 | 2.618506 | 2.713378 | 2.771421 | 2.822722 | 2.883685 | 2.939246 |
|  | 50 | 2.051797 | 2.103295 | 2.160502 | 2.209209 | 2.26493 | 2.35739 | 2.459894 | 2.563074 | 2.657319 | 2.714665 | 2.765134 | 2.824827 | 2.878954 |
|  | 60 | 2.010345 | 2.061928 | 2.119309 | 2.168217 | 2.224213 | 2.317187 | 2.420251 | 2.523864 | 2.618285 | 2.675597 | 2.72593 | 2.785316 | 2.839014 |
|  | 70 | 1.975506 | 2.02701 | 2.084417 | 2.13343 | 2.189626 | 2.283072 | 2.386782 | 2.491064 | 2.586017 | 2.643579 | 2.69407 | 2.753553 | 2.807239 |
|  | 80 | 1.950518 | 2.001807 | 2.059139 | 2.108215 | 2.164612 | 2.258659 | 2.363343 | 2.468827 | 2.56498 | 2.623284 | 2.674418 | 2.734635 | 2.788951 |
|  | 90 | 1.929384 | 1.980129 | 2.037075 | 2.085994 | 2.142398 | 2.23686 | 2.342523 | 2.449449 | 2.547229 | 2.606632 | 2.658785 | 2.720253 | 2.77573 |
| **R_parsorbitalis** | 3 | 2.537747 | 2.638274 | 2.745845 | 2.834268 | 2.932064 | 3.08684 | 3.248022 | 3.399662 | 3.529172 | 3.60388 | 3.667126 | 3.738969 | 3.801402 |
|  | 5 | 2.499471 | 2.594562 | 2.697229 | 2.78228 | 2.876991 | 3.028172 | 3.187146 | 3.338016 | 3.46779 | 3.543015 | 3.606897 | 3.679677 | 3.743103 |
|  | 10 | 2.40783 | 2.491158 | 2.582933 | 2.660324 | 2.747907 | 2.890636 | 3.044413 | 3.193658 | 3.324486 | 3.401319 | 3.467126 | 3.54271 | 3.609101 |
|  | 15 | 2.32669 | 2.400773 | 2.483716 | 2.554737 | 2.636272 | 2.771722 | 2.921119 | 3.069434 | 3.202052 | 3.281043 | 3.349337 | 3.428495 | 3.49865 |
|  | 20 | 2.262196 | 2.32993 | 2.406666 | 2.473118 | 2.550246 | 2.680327 | 2.826582 | 2.974602 | 3.109295 | 3.190555 | 3.261425 | 3.344275 | 3.418329 |
|  | 30 | 2.178101 | 2.240882 | 2.312579 | 2.375156 | 2.448354 | 2.573173 | 2.715542 | 2.861791 | 2.996741 | 3.079009 | 3.151277 | 3.236371 | 3.312985 |
|  | 40 | 2.119369 | 2.182854 | 2.255094 | 2.317921 | 2.391151 | 2.515394 | 2.656164 | 2.799766 | 2.931401 | 3.011251 | 3.081153 | 3.163177 | 3.236769 |
|  | 50 | 2.069143 | 2.134619 | 2.208781 | 2.272991 | 2.3475 | 2.473124 | 2.614305 | 2.757122 | 2.88702 | 2.965359 | 3.033664 | 3.113496 | 3.184835 |
|  | 60 | 1.996926 | 2.065296 | 2.14241 | 2.208904 | 2.285758 | 2.414622 | 2.558422 | 2.702847 | 2.833346 | 2.911668 | 2.979732 | 3.059022 | 3.129645 |
|  | 70 | 1.937357 | 2.007441 | 2.086539 | 2.154784 | 2.233708 | 2.366145 | 2.514079 | 2.6628 | 2.797298 | 2.878071 | 2.948296 | 3.030137 | 3.103062 |
|  | 80 | 1.902272 | 1.974632 | 2.056466 | 2.127211 | 2.209182 | 2.347091 | 2.501644 | 2.657532 | 2.798931 | 2.884031 | 2.958126 | 3.044602 | 3.121768 |
|  | 90 | 1.884662 | 1.960368 | 2.046146 | 2.120434 | 2.206656 | 2.352062 | 2.515501 | 2.680841 | 2.831213 | 2.92189 | 3.000945 | 3.093327 | 3.175869 |
| **R_parstriangularis** | 3 | 2.520745 | 2.575372 | 2.636011 | 2.687643 | 2.746766 | 2.845158 | 2.954968 | 3.066691 | 3.170158 | 3.233927 | 3.290631 | 3.358473 | 3.420777 |
|  | 5 | 2.473318 | 2.527497 | 2.587622 | 2.638805 | 2.697406 | 2.794914 | 2.903737 | 3.014479 | 3.11708 | 3.180344 | 3.236621 | 3.303982 | 3.365878 |
|  | 10 | 2.359557 | 2.412661 | 2.471552 | 2.521657 | 2.578999 | 2.674381 | 2.780827 | 2.889208 | 2.989727 | 3.05178 | 3.107035 | 3.173252 | 3.234178 |
|  | 15 | 2.255864 | 2.308035 | 2.365849 | 2.41501 | 2.471246 | 2.564754 | 2.669104 | 2.775409 | 2.874112 | 2.935117 | 2.989496 | 3.054742 | 3.114861 |
|  | 20 | 2.173002 | 2.224606 | 2.281749 | 2.330308 | 2.38583 | 2.478112 | 2.581088 | 2.686051 | 2.783618 | 2.843994 | 2.897873 | 2.9626 | 3.022328 |
|  | 30 | 2.068048 | 2.119435 | 2.176238 | 2.224439 | 2.279491 | 2.370899 | 2.472866 | 2.576903 | 2.673825 | 2.733955 | 2.787733 | 2.85251 | 2.912466 |
|  | 40 | 1.997741 | 2.049213 | 2.105995 | 2.154097 | 2.208961 | 2.29994 | 2.40137 | 2.504946 | 2.601652 | 2.661805 | 2.715728 | 2.780861 | 2.84134 |
|  | 50 | 1.947077 | 1.998794 | 2.055711 | 2.103834 | 2.158637 | 2.249372 | 2.350448 | 2.453738 | 2.550396 | 2.610682 | 2.664857 | 2.730487 | 2.791637 |
|  | 60 | 1.895979 | 1.94783 | 2.004748 | 2.052766 | 2.107353 | 2.197572 | 2.297974 | 2.400648 | 2.496959 | 2.557202 | 2.611483 | 2.677449 | 2.739138 |
|  | 70 | 1.856676 | 1.909181 | 1.966656 | 2.015029 | 2.069915 | 2.160457 | 2.261111 | 2.364124 | 2.461007 | 2.521801 | 2.576737 | 2.64373 | 2.706635 |
|  | 80 | 1.838373 | 1.892417 | 1.951399 | 2.000915 | 2.056983 | 2.149285 | 2.251781 | 2.356774 | 2.455806 | 2.518166 | 2.574697 | 2.643898 | 2.709166 |
|  | 90 | 1.819186 | 1.87493 | 1.935567 | 1.986332 | 2.043685 | 2.137891 | 2.242371 | 2.3495 | 2.450864 | 2.514936 | 2.573221 | 2.644866 | 2.712766 |
| **R_pericalcarine** | 3 | 1.554511 | 1.589002 | 1.629215 | 1.665056 | 1.707894 | 1.783328 | 1.873269 | 1.970325 | 2.064403 | 2.124026 | 2.177896 | 2.243197 | 2.303796 |
|  | 5 | 1.517011 | 1.550879 | 1.590384 | 1.625612 | 1.667737 | 1.741972 | 1.830577 | 1.926304 | 2.019207 | 2.078143 | 2.13143 | 2.196074 | 2.25611 |
|  | 10 | 1.433118 | 1.465605 | 1.503545 | 1.537418 | 1.577974 | 1.649581 | 1.735278 | 1.828144 | 1.918551 | 1.976046 | 2.028127 | 2.09143 | 2.150341 |
|  | 15 | 1.369984 | 1.401505 | 1.43836 | 1.471305 | 1.510803 | 1.580674 | 1.664523 | 1.75567 | 1.844686 | 1.901443 | 1.952954 | 2.01569 | 2.074198 |
|  | 20 | 1.330328 | 1.361358 | 1.397682 | 1.430193 | 1.469218 | 1.538387 | 1.621619 | 1.712377 | 1.801299 | 1.858143 | 1.909834 | 1.972917 | 2.031877 |
|  | 30 | 1.293395 | 1.32405 | 1.360008 | 1.392255 | 1.431048 | 1.500031 | 1.583425 | 1.674851 | 1.764932 | 1.822782 | 1.875569 | 1.940225 | 2.000892 |
|  | 40 | 1.275058 | 1.305133 | 1.34045 | 1.372161 | 1.410357 | 1.478411 | 1.560919 | 1.651679 | 1.741428 | 1.79924 | 1.852112 | 1.917034 | 1.978113 |
|  | 50 | 1.256656 | 1.285945 | 1.320366 | 1.3513 | 1.388594 | 1.455137 | 1.535988 | 1.625158 | 1.713587 | 1.770685 | 1.823002 | 1.887372 | 1.948065 |
|  | 60 | 1.243367 | 1.272083 | 1.305861 | 1.336247 | 1.372918 | 1.438457 | 1.51828 | 1.606575 | 1.694412 | 1.751282 | 1.803498 | 1.867888 | 1.928749 |
|  | 70 | 1.231123 | 1.259354 | 1.292595 | 1.322528 | 1.358693 | 1.423443 | 1.502509 | 1.590242 | 1.677822 | 1.734691 | 1.787023 | 1.851716 | 1.913027 |
|  | 80 | 1.210706 | 1.23819 | 1.270579 | 1.29977 | 1.335073 | 1.398374 | 1.475848 | 1.562053 | 1.648373 | 1.704572 | 1.756394 | 1.820601 | 1.881602 |
|  | 90 | 1.174934 | 1.201245 | 1.232272 | 1.260255 | 1.294122 | 1.354923 | 1.429476 | 1.512622 | 1.596096 | 1.650563 | 1.700878 | 1.763339 | 1.822811 |
| **R_postcentral** | 3 | 2.067263 | 2.104844 | 2.147285 | 2.184084 | 2.227045 | 2.300737 | 2.386766 | 2.479196 | 2.570057 | 2.628971 | 2.683448 | 2.751462 | 2.816907 |
|  | 5 | 2.024849 | 2.062363 | 2.104667 | 2.14129 | 2.18398 | 2.257039 | 2.342053 | 2.433055 | 2.522165 | 2.579756 | 2.632877 | 2.699017 | 2.762467 |
|  | 10 | 1.923356 | 1.960755 | 2.002764 | 2.038988 | 2.081045 | 2.152595 | 2.235174 | 2.322754 | 2.407692 | 2.462152 | 2.512084 | 2.573848 | 2.632676 |
|  | 15 | 1.836511 | 1.873987 | 1.915913 | 1.951923 | 1.993562 | 2.063989 | 2.144621 | 2.22938 | 2.31084 | 2.362689 | 2.409965 | 2.468099 | 2.523118 |
|  | 20 | 1.776183 | 1.814176 | 1.856512 | 1.892731 | 1.934451 | 2.004617 | 2.084346 | 2.167465 | 2.246693 | 2.29679 | 2.342246 | 2.397855 | 2.450194 |
|  | 30 | 1.707171 | 1.747084 | 1.791226 | 1.828716 | 1.871589 | 1.942964 | 2.022981 | 2.105216 | 2.182513 | 2.230858 | 2.274377 | 2.327177 | 2.376437 |
|  | 40 | 1.660515 | 1.702309 | 1.748266 | 1.787084 | 1.831241 | 1.904211 | 1.985242 | 2.067703 | 2.144496 | 2.192187 | 2.2349 | 2.286453 | 2.334292 |
|  | 50 | 1.622474 | 1.665545 | 1.712764 | 1.752533 | 1.797648 | 1.871925 | 1.954024 | 2.037183 | 2.114297 | 2.162035 | 2.204698 | 2.256075 | 2.303643 |
|  | 60 | 1.59001 | 1.633336 | 1.680869 | 1.720931 | 1.766412 | 1.841367 | 1.924336 | 2.008515 | 2.086706 | 2.13518 | 2.178544 | 2.230824 | 2.279286 |
|  | 70 | 1.553501 | 1.595586 | 1.641991 | 1.681293 | 1.72612 | 1.800488 | 1.883518 | 1.968529 | 2.048192 | 2.097916 | 2.14262 | 2.196794 | 2.247281 |
|  | 80 | 1.511616 | 1.551672 | 1.596149 | 1.634074 | 1.67762 | 1.750548 | 1.832994 | 1.918543 | 1.999772 | 2.051 | 2.097405 | 2.154085 | 2.207349 |
|  | 90 | 1.461613 | 1.499455 | 1.541761 | 1.578077 | 1.620056 | 1.691043 | 1.772351 | 1.857939 | 1.940383 | 1.992982 | 2.041039 | 2.10027 | 2.156476 |
| **R_posteriorcingulate** | 3 | 2.55784 | 2.626711 | 2.696912 | 2.752484 | 2.812469 | 2.906662 | 3.008921 | 3.116988 | 3.227994 | 3.305303 | 3.382124 | 3.487181 | 3.600097 |
|  | 5 | 2.519773 | 2.585463 | 2.653097 | 2.70712 | 2.765884 | 2.858931 | 2.960539 | 3.067801 | 3.177061 | 3.252318 | 3.326316 | 3.426205 | 3.531901 |
|  | 10 | 2.421569 | 2.480921 | 2.543363 | 2.594203 | 2.650407 | 2.740962 | 2.841039 | 2.9464 | 3.0518 | 3.122703 | 3.190877 | 3.280398 | 3.372044 |
|  | 15 | 2.325157 | 2.379912 | 2.438527 | 2.486981 | 2.541231 | 2.629823 | 2.728614 | 2.832353 | 2.934632 | 3.00215 | 3.065928 | 3.147868 | 3.229603 |
|  | 20 | 2.244669 | 2.296195 | 2.35213 | 2.39893 | 2.451859 | 2.539209 | 2.637296 | 2.740072 | 2.84024 | 2.905388 | 2.966073 | 3.042717 | 3.117631 |
|  | 30 | 2.141138 | 2.18902 | 2.242048 | 2.287181 | 2.338945 | 2.42563 | 2.523873 | 2.626439 | 2.724738 | 2.787319 | 2.844461 | 2.9149 | 2.981796 |
|  | 40 | 2.087178 | 2.133931 | 2.18633 | 2.231376 | 2.283459 | 2.371372 | 2.471418 | 2.575443 | 2.673929 | 2.735715 | 2.791381 | 2.858904 | 2.921829 |
|  | 50 | 2.043824 | 2.09018 | 2.142559 | 2.187895 | 2.240606 | 2.330086 | 2.432269 | 2.538345 | 2.638102 | 2.70016 | 2.755635 | 2.822294 | 2.883725 |
|  | 60 | 2.007019 | 2.05283 | 2.104952 | 2.150343 | 2.203398 | 2.294019 | 2.398092 | 2.506429 | 2.608261 | 2.671454 | 2.727784 | 2.795214 | 2.857057 |
|  | 70 | 1.979506 | 2.023794 | 2.074623 | 2.11925 | 2.171818 | 2.262545 | 2.368046 | 2.479142 | 2.584551 | 2.650364 | 2.709247 | 2.779966 | 2.845013 |
|  | 80 | 1.972261 | 2.014413 | 2.063301 | 2.106671 | 2.158288 | 2.248709 | 2.355943 | 2.47124 | 2.58284 | 2.653589 | 2.717576 | 2.795273 | 2.867549 |
|  | 90 | 1.980455 | 2.02076 | 2.067975 | 2.110286 | 2.161171 | 2.251713 | 2.361468 | 2.48244 | 2.602569 | 2.680323 | 2.751745 | 2.83991 | 2.923381 |
| **R_precentral** | 3 | 2.109848 | 2.23386 | 2.344744 | 2.423372 | 2.500201 | 2.604728 | 2.69622 | 2.776046 | 2.845212 | 2.885321 | 2.919261 | 2.957698 | 2.990935 |
|  | 5 | 2.125428 | 2.238911 | 2.342703 | 2.417671 | 2.491971 | 2.594599 | 2.685771 | 2.76606 | 2.835973 | 2.876654 | 2.911157 | 2.950315 | 2.984246 |
|  | 10 | 2.154418 | 2.24545 | 2.333689 | 2.400314 | 2.468629 | 2.566529 | 2.65677 | 2.738157 | 2.809971 | 2.852148 | 2.88814 | 2.929231 | 2.965039 |
|  | 15 | 2.163461 | 2.238165 | 2.314102 | 2.373613 | 2.436483 | 2.529702 | 2.618772 | 2.701095 | 2.77479 | 2.818523 | 2.856103 | 2.899296 | 2.937187 |
|  | 20 | 2.151248 | 2.215297 | 2.282526 | 2.33665 | 2.395158 | 2.48437 | 2.57232 | 2.655413 | 2.730776 | 2.775934 | 2.814998 | 2.860194 | 2.900101 |
|  | 30 | 2.085331 | 2.144105 | 2.206856 | 2.258162 | 2.314422 | 2.401869 | 2.490175 | 2.57457 | 2.651062 | 2.696918 | 2.736626 | 2.782628 | 2.823315 |
|  | 40 | 2.003877 | 2.067403 | 2.134448 | 2.188757 | 2.247879 | 2.339098 | 2.430684 | 2.51692 | 2.593297 | 2.638382 | 2.677049 | 2.721452 | 2.760404 |
|  | 50 | 1.944908 | 2.011588 | 2.081753 | 2.13849 | 2.200215 | 2.29554 | 2.391579 | 2.48143 | 2.55978 | 2.605568 | 2.644602 | 2.689192 | 2.728121 |
|  | 60 | 1.900646 | 1.963285 | 2.030435 | 2.08565 | 2.14665 | 2.242808 | 2.342158 | 2.436308 | 2.51845 | 2.566496 | 2.607503 | 2.654415 | 2.695441 |
|  | 70 | 1.844304 | 1.901484 | 1.964021 | 2.0164 | 2.07528 | 2.170312 | 2.271417 | 2.368896 | 2.454325 | 2.504474 | 2.547399 | 2.59666 | 2.639885 |
|  | 80 | 1.780902 | 1.835264 | 1.895392 | 1.946307 | 2.004161 | 2.099003 | 2.201976 | 2.302198 | 2.389865 | 2.441279 | 2.485283 | 2.535797 | 2.58015 |
|  | 90 | 1.703481 | 1.759277 | 1.820895 | 1.873026 | 1.932258 | 2.029478 | 2.135353 | 2.237708 | 2.325848 | 2.376992 | 2.420482 | 2.470118 | 2.513468 |
| **R_precuneus** | 3 | 2.491503 | 2.554392 | 2.620034 | 2.672953 | 2.730683 | 2.821184 | 2.915904 | 3.007544 | 3.089685 | 3.13952 | 3.183548 | 3.236067 | 3.284308 |
|  | 5 | 2.43634 | 2.498445 | 2.563228 | 2.615429 | 2.672356 | 2.761585 | 2.855002 | 2.945474 | 3.026703 | 3.076067 | 3.119742 | 3.171922 | 3.219939 |
|  | 10 | 2.298771 | 2.358844 | 2.421403 | 2.471747 | 2.526603 | 2.612546 | 2.702594 | 2.790036 | 2.868885 | 2.917014 | 2.959753 | 3.01103 | 3.058439 |
|  | 15 | 2.170349 | 2.228386 | 2.288746 | 2.337275 | 2.390122 | 2.472904 | 2.55974 | 2.644308 | 2.720908 | 2.767876 | 2.80974 | 2.860181 | 2.907041 |
|  | 20 | 2.0814 | 2.138021 | 2.196919 | 2.244284 | 2.295882 | 2.376776 | 2.461773 | 2.544769 | 2.620201 | 2.666599 | 2.708061 | 2.75816 | 2.804847 |
|  | 30 | 1.992235 | 2.047678 | 2.105627 | 2.152418 | 2.203558 | 2.284006 | 2.368734 | 2.451453 | 2.526434 | 2.572391 | 2.613323 | 2.662587 | 2.708287 |
|  | 40 | 1.944041 | 1.998247 | 2.055497 | 2.102124 | 2.153434 | 2.234695 | 2.3206 | 2.404255 | 2.479439 | 2.525049 | 2.5653 | 2.613221 | 2.65712 |
|  | 50 | 1.910093 | 1.961523 | 2.016777 | 2.062433 | 2.113276 | 2.194844 | 2.281971 | 2.367045 | 2.443124 | 2.488892 | 2.528947 | 2.576146 | 2.618856 |
|  | 60 | 1.878067 | 1.925892 | 1.978264 | 2.022261 | 2.071961 | 2.153032 | 2.241034 | 2.327803 | 2.405614 | 2.452336 | 2.493087 | 2.540864 | 2.583807 |
|  | 70 | 1.838729 | 1.882801 | 1.93192 | 1.973843 | 2.021879 | 2.101648 | 2.18994 | 2.278366 | 2.358489 | 2.406845 | 2.449113 | 2.498715 | 2.543295 |
|  | 80 | 1.785518 | 1.825759 | 1.871314 | 1.910768 | 1.956603 | 2.034126 | 2.121838 | 2.21149 | 2.294096 | 2.344508 | 2.388877 | 2.441274 | 2.488634 |
|  | 90 | 1.729705 | 1.766561 | 1.808841 | 1.845932 | 1.889567 | 1.964677 | 2.051591 | 2.142474 | 2.227973 | 2.280947 | 2.328057 | 2.384257 | 2.435569 |
| **R_rostralanteriorcingulate** | 3 | 2.623742 | 2.725493 | 2.836429 | 2.928762 | 3.031521 | 3.193838 | 3.360064 | 3.524352 | 3.681107 | 3.778512 | 3.864839 | 3.967081 | 4.059447 |
|  | 5 | 2.569132 | 2.668423 | 2.776836 | 2.867197 | 2.967905 | 3.127305 | 3.290992 | 3.452961 | 3.607445 | 3.703428 | 3.788498 | 3.889257 | 3.980296 |
|  | 10 | 2.448994 | 2.542822 | 2.645647 | 2.731655 | 2.82785 | 2.980888 | 3.139125 | 3.296177 | 3.44584 | 3.538806 | 3.621207 | 3.71883 | 3.807068 |
|  | 15 | 2.366145 | 2.456004 | 2.554837 | 2.637799 | 2.730916 | 2.879819 | 3.034851 | 3.189204 | 3.33618 | 3.427462 | 3.50838 | 3.604273 | 3.690983 |
|  | 20 | 2.307956 | 2.394828 | 2.490717 | 2.571493 | 2.662477 | 2.808721 | 2.962054 | 3.115206 | 3.260942 | 3.351443 | 3.431683 | 3.526804 | 3.612855 |
|  | 30 | 2.213989 | 2.295825 | 2.386794 | 2.463964 | 2.551503 | 2.693679 | 2.844868 | 2.996899 | 3.141423 | 3.231172 | 3.310784 | 3.405235 | 3.490772 |
|  | 40 | 2.160401 | 2.238788 | 2.326528 | 2.401474 | 2.487093 | 2.627616 | 2.779207 | 2.932734 | 3.078601 | 3.169213 | 3.249645 | 3.345167 | 3.431783 |
|  | 50 | 2.131681 | 2.207577 | 2.293106 | 2.366665 | 2.451291 | 2.591662 | 2.745313 | 2.902105 | 3.051067 | 3.14366 | 3.225927 | 3.323748 | 3.412584 |
|  | 60 | 2.127036 | 2.201327 | 2.285601 | 2.358571 | 2.443111 | 2.584841 | 2.742292 | 2.904253 | 3.058196 | 3.153979 | 3.23918 | 3.340642 | 3.432948 |
|  | 70 | 2.141592 | 2.214946 | 2.298696 | 2.371697 | 2.456865 | 2.601188 | 2.763948 | 2.932791 | 3.093437 | 3.193528 | 3.282688 | 3.389053 | 3.486021 |
|  | 80 | 2.169386 | 2.242234 | 2.325931 | 2.399366 | 2.485638 | 2.633419 | 2.802641 | 2.979762 | 3.148552 | 3.253904 | 3.347913 | 3.460297 | 3.562998 |
|  | 90 | 2.213322 | 2.286164 | 2.37037 | 2.444728 | 2.532692 | 2.685015 | 2.862162 | 3.049335 | 3.228095 | 3.339919 | 3.439907 | 3.55973 | 3.669532 |
| **R_rostralmiddlefrontal** | 3 | 2.496267 | 2.545748 | 2.600996 | 2.648274 | 2.702639 | 2.793522 | 2.895306 | 2.998908 | 3.094597 | 3.153336 | 3.205364 | 3.267313 | 3.323878 |
|  | 5 | 2.428153 | 2.476487 | 2.530449 | 2.576625 | 2.629722 | 2.718494 | 2.817942 | 2.919218 | 3.012824 | 3.070323 | 3.12128 | 3.18199 | 3.237461 |
|  | 10 | 2.274456 | 2.320216 | 2.371291 | 2.414989 | 2.465236 | 2.549263 | 2.64346 | 2.739515 | 2.828452 | 2.883176 | 2.931742 | 2.989694 | 3.042737 |
|  | 15 | 2.150055 | 2.193795 | 2.242599 | 2.284346 | 2.332347 | 2.412631 | 2.502692 | 2.594648 | 2.679943 | 2.732518 | 2.779244 | 2.835089 | 2.886298 |
|  | 20 | 2.071189 | 2.113823 | 2.161369 | 2.202027 | 2.248767 | 2.326941 | 2.414679 | 2.504362 | 2.587692 | 2.63914 | 2.684929 | 2.739742 | 2.790094 |
|  | 30 | 1.996765 | 2.038926 | 2.085862 | 2.12594 | 2.17196 | 2.248854 | 2.335129 | 2.42341 | 2.505626 | 2.556521 | 2.60192 | 2.656416 | 2.706634 |
|  | 40 | 1.957164 | 1.99989 | 2.04731 | 2.087694 | 2.133962 | 2.211074 | 2.297404 | 2.385674 | 2.467939 | 2.518939 | 2.564503 | 2.619304 | 2.669926 |
|  | 50 | 1.92039 | 1.964298 | 2.01282 | 2.053983 | 2.100985 | 2.178999 | 2.26599 | 2.354709 | 2.437327 | 2.488567 | 2.534385 | 2.589562 | 2.640622 |
|  | 60 | 1.875756 | 1.921431 | 1.971629 | 2.014006 | 2.062183 | 2.141723 | 2.22993 | 2.319545 | 2.402846 | 2.454497 | 2.500703 | 2.556398 | 2.608008 |
|  | 70 | 1.831959 | 1.880202 | 1.932894 | 1.977129 | 2.027172 | 2.109303 | 2.199829 | 2.291416 | 2.376387 | 2.429066 | 2.476215 | 2.533112 | 2.58592 |
|  | 80 | 1.816646 | 1.869102 | 1.925995 | 1.973463 | 2.026871 | 2.113952 | 2.209301 | 2.305343 | 2.394283 | 2.449424 | 2.498818 | 2.558505 | 2.614014 |
|  | 90 | 1.800967 | 1.858514 | 1.920429 | 1.971719 | 2.02907 | 2.121884 | 2.222756 | 2.323862 | 2.417306 | 2.475246 | 2.527195 | 2.590072 | 2.648678 |
| **R_superiorfrontal** | 3 | 2.775616 | 2.843232 | 2.914896 | 2.973356 | 3.037656 | 3.139003 | 3.244677 | 3.345096 | 3.432411 | 3.483746 | 3.527914 | 3.579031 | 3.624408 |
|  | 5 | 2.730233 | 2.795721 | 2.865374 | 2.922371 | 2.985241 | 3.084703 | 3.188881 | 3.28832 | 3.375138 | 3.426336 | 3.470481 | 3.521681 | 3.567232 |
|  | 10 | 2.622025 | 2.682565 | 2.747495 | 2.801031 | 2.860497 | 2.955447 | 3.056047 | 3.153181 | 3.238898 | 3.289858 | 3.334047 | 3.385595 | 3.431729 |
|  | 15 | 2.525909 | 2.58205 | 2.642739 | 2.693147 | 2.749524 | 2.840387 | 2.937806 | 3.033022 | 3.118026 | 3.169012 | 3.213505 | 3.265744 | 3.312809 |
|  | 20 | 2.450414 | 2.503198 | 2.560629 | 2.608622 | 2.662616 | 2.750358 | 2.84545 | 2.939468 | 3.024351 | 3.075715 | 3.120825 | 3.174138 | 3.222504 |
|  | 30 | 2.350618 | 2.40109 | 2.456198 | 2.50242 | 2.554622 | 2.639969 | 2.733318 | 2.826658 | 2.911983 | 2.964164 | 3.010362 | 3.065438 | 3.115874 |
|  | 40 | 2.283504 | 2.33496 | 2.390908 | 2.437667 | 2.490325 | 2.576166 | 2.669877 | 2.763631 | 2.849584 | 2.902346 | 2.94922 | 3.005333 | 3.056968 |
|  | 50 | 2.22801 | 2.281595 | 2.339456 | 2.387527 | 2.441388 | 2.528683 | 2.623487 | 2.718107 | 2.804906 | 2.858306 | 2.905864 | 2.962984 | 3.015758 |
|  | 60 | 2.171182 | 2.226792 | 2.286433 | 2.335695 | 2.390624 | 2.479177 | 2.574941 | 2.670424 | 2.75822 | 2.812439 | 2.860909 | 2.919395 | 2.973732 |
|  | 70 | 2.118807 | 2.174603 | 2.234338 | 2.28362 | 2.338549 | 2.427176 | 2.523378 | 2.620023 | 2.709848 | 2.765905 | 2.816455 | 2.878051 | 2.935915 |
|  | 80 | 2.080935 | 2.136173 | 2.195355 | 2.244244 | 2.298841 | 2.387307 | 2.484157 | 2.582737 | 2.675915 | 2.734982 | 2.788924 | 2.855597 | 2.919231 |
|  | 90 | 2.045679 | 2.100487 | 2.159213 | 2.207762 | 2.262063 | 2.350404 | 2.447981 | 2.548751 | 2.645848 | 2.708533 | 2.766646 | 2.839708 | 2.910795 |
| **R_superiorparietal** | 3 | 2.248143 | 2.292097 | 2.341567 | 2.384136 | 2.433236 | 2.515318 | 2.606558 | 2.697948 | 2.780423 | 2.829875 | 2.87282 | 2.922812 | 2.967306 |
|  | 5 | 2.194658 | 2.238118 | 2.286988 | 2.329002 | 2.377414 | 2.458216 | 2.547832 | 2.637542 | 2.718593 | 2.767227 | 2.80948 | 2.858683 | 2.902486 |
|  | 10 | 2.067295 | 2.109575 | 2.157013 | 2.197699 | 2.244461 | 2.322193 | 2.40791 | 2.493589 | 2.571223 | 2.617895 | 2.658484 | 2.705789 | 2.747931 |
|  | 15 | 1.957192 | 1.998578 | 2.0449 | 2.084528 | 2.12995 | 2.205131 | 2.28753 | 2.369765 | 2.444514 | 2.489544 | 2.528749 | 2.574482 | 2.615254 |
|  | 20 | 1.88414 | 1.925446 | 1.971549 | 2.010872 | 2.0558 | 2.12979 | 2.210302 | 2.290521 | 2.363732 | 2.407949 | 2.446503 | 2.491529 | 2.531711 |
|  | 30 | 1.808654 | 1.851899 | 1.899788 | 1.940297 | 1.986168 | 2.060638 | 2.140023 | 2.218843 | 2.291772 | 2.336212 | 2.375155 | 2.420829 | 2.461738 |
|  | 40 | 1.764756 | 1.810377 | 1.86056 | 1.902711 | 1.950079 | 2.02605 | 2.105605 | 2.184424 | 2.25834 | 2.303771 | 2.34378 | 2.390904 | 2.433269 |
|  | 50 | 1.73902 | 1.785387 | 1.836412 | 1.879291 | 1.927503 | 2.004903 | 2.086074 | 2.166485 | 2.241768 | 2.287987 | 2.328665 | 2.376548 | 2.419573 |
|  | 60 | 1.710154 | 1.75556 | 1.805813 | 1.848301 | 1.896395 | 1.974444 | 2.057619 | 2.140109 | 2.216291 | 2.262652 | 2.303244 | 2.350815 | 2.393391 |
|  | 70 | 1.677564 | 1.721108 | 1.769704 | 1.811162 | 1.858559 | 1.936734 | 2.022042 | 2.106783 | 2.183453 | 2.229485 | 2.269471 | 2.31601 | 2.35741 |
|  | 80 | 1.635101 | 1.675852 | 1.721826 | 1.761512 | 1.807479 | 1.884945 | 1.972137 | 2.058921 | 2.135345 | 2.180395 | 2.219109 | 2.263749 | 2.303132 |
|  | 90 | 1.575687 | 1.613162 | 1.655929 | 1.693313 | 1.737227 | 1.812976 | 1.901065 | 1.988952 | 2.064207 | 2.107699 | 2.144644 | 2.18682 | 2.223701 |
| **R_superiortemporal** | 3 | 2.512247 | 2.645429 | 2.764798 | 2.850029 | 2.934396 | 3.053014 | 3.164406 | 3.263875 | 3.348568 | 3.398623 | 3.4423 | 3.493962 | 3.541178 |
|  | 5 | 2.508073 | 2.630968 | 2.743367 | 2.824915 | 2.906632 | 3.023006 | 3.133534 | 3.232868 | 3.317617 | 3.367673 | 3.411287 | 3.462761 | 3.509672 |
|  | 10 | 2.492711 | 2.592968 | 2.689469 | 2.762302 | 2.83754 | 2.948206 | 3.056475 | 3.155594 | 3.240843 | 3.29125 | 3.335098 | 3.386679 | 3.433459 |
|  | 15 | 2.480284 | 2.559616 | 2.640203 | 2.70369 | 2.771574 | 2.875378 | 2.98094 | 3.080365 | 3.167416 | 3.219337 | 3.264675 | 3.318126 | 3.366643 |
|  | 20 | 2.467682 | 2.531153 | 2.598564 | 2.653727 | 2.714673 | 2.811626 | 2.914531 | 3.014932 | 3.105144 | 3.159793 | 3.207943 | 3.265144 | 3.317405 |
|  | 30 | 2.403113 | 2.459065 | 2.520066 | 2.571144 | 2.62872 | 2.722546 | 2.824625 | 2.925982 | 3.01789 | 3.073701 | 3.122843 | 3.181079 | 3.234062 |
|  | 40 | 2.319256 | 2.38173 | 2.448965 | 2.504561 | 2.566455 | 2.665507 | 2.770589 | 2.871972 | 2.961192 | 3.014056 | 3.059759 | 3.112881 | 3.160226 |
|  | 50 | 2.254675 | 2.319003 | 2.388204 | 2.445379 | 2.508952 | 2.610423 | 2.71752 | 2.820063 | 2.909452 | 2.961961 | 3.007046 | 3.059056 | 3.105026 |
|  | 60 | 2.220378 | 2.277458 | 2.340376 | 2.393508 | 2.453763 | 2.552383 | 2.6595 | 2.764689 | 2.858228 | 2.91388 | 2.962034 | 3.017969 | 3.067711 |
|  | 70 | 2.160972 | 2.213429 | 2.27214 | 2.322426 | 2.380212 | 2.476452 | 2.583178 | 2.690016 | 2.786549 | 2.844599 | 2.895168 | 2.954274 | 3.007142 |
|  | 80 | 2.056238 | 2.111792 | 2.173459 | 2.225857 | 2.285596 | 2.383983 | 2.491494 | 2.597452 | 2.691764 | 2.747826 | 2.796266 | 2.852413 | 2.902204 |
|  | 90 | 1.932744 | 1.995724 | 2.06415 | 2.121154 | 2.184962 | 2.287515 | 2.396293 | 2.500451 | 2.590823 | 2.643562 | 2.688568 | 2.740101 | 2.785249 |
| **R_supramarginal** | 3 | 2.531651 | 2.618007 | 2.703238 | 2.768828 | 2.8376 | 2.940285 | 3.041869 | 3.135053 | 3.214686 | 3.261318 | 3.301509 | 3.348263 | 3.390117 |
|  | 5 | 2.492358 | 2.574409 | 2.656149 | 2.71952 | 2.786364 | 2.886864 | 2.987024 | 3.079481 | 3.15889 | 3.205547 | 3.24585 | 3.292832 | 3.334977 |
|  | 10 | 2.396109 | 2.468721 | 2.542592 | 2.600848 | 2.663173 | 2.75846 | 2.855177 | 2.945884 | 3.024806 | 3.07159 | 3.112237 | 3.159887 | 3.202866 |
|  | 15 | 2.314028 | 2.377241 | 2.443055 | 2.495984 | 2.553573 | 2.643464 | 2.73687 | 2.826346 | 2.905599 | 2.953171 | 2.994848 | 3.044105 | 3.088894 |
|  | 20 | 2.257579 | 2.31215 | 2.370343 | 2.418146 | 2.471162 | 2.555967 | 2.646691 | 2.73603 | 2.817108 | 2.866638 | 2.910555 | 2.963085 | 3.011433 |
|  | 30 | 2.173671 | 2.224493 | 2.279328 | 2.324862 | 2.375869 | 2.458561 | 2.548492 | 2.638504 | 2.721415 | 2.772627 | 2.818388 | 2.873554 | 2.924738 |
|  | 40 | 2.09716 | 2.153281 | 2.213001 | 2.261965 | 2.316173 | 2.402685 | 2.494968 | 2.585583 | 2.667598 | 2.717597 | 2.761867 | 2.81474 | 2.863329 |
|  | 50 | 2.046566 | 2.103471 | 2.164084 | 2.213825 | 2.268937 | 2.356986 | 2.451027 | 2.543476 | 2.627235 | 2.678334 | 2.723598 | 2.777685 | 2.827411 |
|  | 60 | 2.014585 | 2.065917 | 2.121683 | 2.168284 | 2.2208 | 2.306621 | 2.400887 | 2.496173 | 2.584735 | 2.639805 | 2.689244 | 2.749128 | 2.804961 |
|  | 70 | 1.953571 | 2.004425 | 2.059855 | 2.106318 | 2.158833 | 2.244995 | 2.340113 | 2.436749 | 2.526991 | 2.583306 | 2.633991 | 2.695542 | 2.753082 |
|  | 80 | 1.872455 | 1.928702 | 1.989149 | 2.039158 | 2.09498 | 2.185041 | 2.282377 | 2.379165 | 2.467753 | 2.5222 | 2.570677 | 2.628898 | 2.682704 |
|  | 90 | 1.803694 | 1.866591 | 1.933104 | 1.987337 | 2.047082 | 2.141819 | 2.242096 | 2.339816 | 2.427651 | 2.480923 | 2.527919 | 2.583842 | 2.635039 |
| **R_frontalpole** | 3 | 2.550007 | 2.675522 | 2.811713 | 2.925417 | 3.053488 | 3.262815 | 3.493196 | 3.727137 | 3.946075 | 4.083157 | 4.206936 | 4.357855 | 4.499615 |
|  | 5 | 2.453288 | 2.571784 | 2.700966 | 2.809271 | 2.931708 | 3.132714 | 3.354922 | 3.581233 | 3.793291 | 3.92606 | 4.04588 | 4.191834 | 4.328748 |
|  | 10 | 2.265792 | 2.370671 | 2.486246 | 2.584081 | 2.695632 | 2.880678 | 3.087439 | 3.299589 | 3.499083 | 3.62405 | 3.736741 | 3.873778 | 4.001998 |
|  | 15 | 2.172331 | 2.269341 | 2.377255 | 2.469383 | 2.57523 | 2.752473 | 2.952486 | 3.159208 | 3.354365 | 3.476754 | 3.587099 | 3.721155 | 3.846369 |
|  | 20 | 2.113738 | 2.20531 | 2.307995 | 2.396298 | 2.498418 | 2.670833 | 2.867113 | 3.071325 | 3.264847 | 3.386373 | 3.49595 | 3.628995 | 3.753111 |
|  | 30 | 2.035699 | 2.119259 | 2.214077 | 2.2965 | 2.39276 | 2.55729 | 2.747077 | 2.946499 | 3.13654 | 3.256091 | 3.363879 | 3.494597 | 3.616267 |
|  | 40 | 2.009939 | 2.089405 | 2.180256 | 2.259767 | 2.353196 | 2.514098 | 2.701158 | 2.898786 | 3.087566 | 3.206322 | 3.313284 | 3.442762 | 3.562952 |
|  | 50 | 1.994515 | 2.073158 | 2.163466 | 2.242812 | 2.336363 | 2.498116 | 2.686838 | 2.886539 | 3.077178 | 3.196877 | 3.304459 | 3.434321 | 3.554444 |
|  | 60 | 1.919907 | 1.998025 | 2.087937 | 2.167086 | 2.260544 | 2.422354 | 2.611231 | 2.810843 | 3.000837 | 3.11973 | 3.226268 | 3.354412 | 3.472455 |
|  | 70 | 1.852883 | 1.932217 | 2.023641 | 2.104192 | 2.199357 | 2.364149 | 2.556338 | 2.758981 | 2.951175 | 3.071014 | 3.178077 | 3.306407 | 3.42416 |
|  | 80 | 1.826984 | 1.91053 | 2.006845 | 2.091713 | 2.191959 | 2.365412 | 2.567316 | 2.779538 | 2.980004 | 3.104534 | 3.215453 | 3.34796 | 3.469094 |
|  | 90 | 1.80491 | 1.893809 | 1.996255 | 2.08647 | 2.19294 | 2.376863 | 2.590342 | 2.813864 | 3.024054 | 3.154113 | 3.269604 | 3.40712 | 3.532385 |
| **R_temporalpole** | 3 | 2.351655 | 2.529676 | 2.712691 | 2.859075 | 3.018229 | 3.267521 | 3.527089 | 3.760576 | 3.943848 | 4.043928 | 4.125997 | 4.216681 | 4.293571 |
|  | 5 | 2.379023 | 2.557247 | 2.740653 | 2.887439 | 3.04708 | 3.297141 | 3.557438 | 3.791856 | 3.976313 | 4.077201 | 4.16001 | 4.251583 | 4.329279 |
|  | 10 | 2.44771 | 2.62639 | 2.810708 | 2.958442 | 3.11923 | 3.371097 | 3.633075 | 3.869699 | 4.057042 | 4.159923 | 4.244562 | 4.338343 | 4.418051 |
|  | 15 | 2.511457 | 2.690276 | 2.875152 | 3.023532 | 3.185126 | 3.438238 | 3.701282 | 3.939549 | 4.129361 | 4.234018 | 4.320316 | 4.416124 | 4.497697 |
|  | 20 | 2.556872 | 2.734457 | 2.918453 | 3.066315 | 3.227438 | 3.479785 | 3.741784 | 3.97978 | 4.170553 | 4.276166 | 4.363454 | 4.460554 | 4.543372 |
|  | 30 | 2.586936 | 2.758197 | 2.93633 | 3.079806 | 3.2363 | 3.481281 | 3.735079 | 3.966904 | 4.155049 | 4.260054 | 4.347244 | 4.444623 | 4.527972 |
|  | 40 | 2.585695 | 2.754447 | 2.930153 | 3.071691 | 3.22596 | 3.466896 | 3.715472 | 3.943463 | 4.130648 | 4.235909 | 4.323693 | 4.422103 | 4.506614 |
|  | 50 | 2.58228 | 2.751804 | 2.928225 | 3.070187 | 3.224663 | 3.465112 | 3.711875 | 3.93895 | 4.12746 | 4.234239 | 4.323661 | 4.424269 | 4.510944 |
|  | 60 | 2.540668 | 2.712435 | 2.890805 | 3.033999 | 3.189403 | 3.430211 | 3.675735 | 3.902215 | 4.092226 | 4.200599 | 4.291719 | 4.394588 | 4.48348 |
|  | 70 | 2.409624 | 2.588235 | 2.77249 | 2.919565 | 3.078342 | 3.322569 | 3.569232 | 3.796804 | 3.989468 | 4.100008 | 4.193266 | 4.298855 | 4.39033 |
|  | 80 | 2.233842 | 2.430012 | 2.629884 | 2.78783 | 2.956895 | 3.214142 | 3.470632 | 3.706728 | 3.908082 | 4.024167 | 4.12237 | 4.233817 | 4.330562 |
|  | 90 | 2.03603 | 2.255087 | 2.475005 | 2.646733 | 2.828741 | 3.102341 | 3.371331 | 3.618187 | 3.83019 | 3.95297 | 4.057102 | 4.175536 | 4.278539 |
| **R_transversetemporal** | 3 | 2.171763 | 2.254228 | 2.345775 | 2.423546 | 2.512169 | 2.657931 | 2.816753 | 2.972576 | 3.110474 | 3.192007 | 3.262154 | 3.343074 | 3.414456 |
|  | 5 | 2.136463 | 2.217963 | 2.308516 | 2.385503 | 2.473298 | 2.617841 | 2.775527 | 2.930416 | 3.067627 | 3.148815 | 3.218698 | 3.299352 | 3.370532 |
|  | 10 | 2.051774 | 2.130929 | 2.219068 | 2.294153 | 2.379941 | 2.521539 | 2.676493 | 2.829158 | 2.964758 | 3.045144 | 3.114424 | 3.19448 | 3.265217 |
|  | 15 | 1.975285 | 2.0523 | 2.138248 | 2.21162 | 2.295617 | 2.434631 | 2.587257 | 2.738108 | 2.872475 | 2.95229 | 3.021171 | 3.100868 | 3.17138 |
|  | 20 | 1.911378 | 1.986641 | 2.070829 | 2.142856 | 2.225485 | 2.362618 | 2.513699 | 2.663529 | 2.797384 | 2.877065 | 2.945929 | 3.025719 | 3.096408 |
|  | 30 | 1.810337 | 1.883685 | 1.966014 | 2.036678 | 2.117994 | 2.253512 | 2.403591 | 2.553188 | 2.687442 | 2.767622 | 2.837068 | 2.917705 | 2.989296 |
|  | 40 | 1.7269 | 1.800545 | 1.883242 | 1.954249 | 2.03599 | 2.172286 | 2.323323 | 2.47397 | 2.609245 | 2.690067 | 2.76009 | 2.841418 | 2.913642 |
|  | 50 | 1.661071 | 1.737395 | 1.822872 | 1.896085 | 1.980168 | 2.119926 | 2.274199 | 2.427493 | 2.564686 | 2.646458 | 2.717191 | 2.799217 | 2.871949 |
|  | 60 | 1.598623 | 1.679278 | 1.769138 | 1.845737 | 1.933315 | 2.07802 | 2.236602 | 2.393088 | 2.532295 | 2.614914 | 2.686177 | 2.768589 | 2.841469 |
|  | 70 | 1.525913 | 1.611516 | 1.706226 | 1.786449 | 1.877639 | 2.027173 | 2.189577 | 2.348485 | 2.488825 | 2.571697 | 2.642939 | 2.725066 | 2.797467 |
|  | 80 | 1.44315 | 1.534308 | 1.634267 | 1.718271 | 1.813084 | 1.96716 | 2.132751 | 2.293232 | 2.433826 | 2.51639 | 2.587112 | 2.66836 | 2.739749 |
|  | 90 | 1.355399 | 1.453382 | 1.559579 | 1.647939 | 1.7468 | 1.905725 | 2.074454 | 2.236208 | 2.376662 | 2.458648 | 2.528603 | 2.608675 | 2.678784 |
| **R_insula** | 3 | 2.911675 | 3.018808 | 3.122691 | 3.201548 | 3.283317 | 3.403773 | 3.521017 | 3.626755 | 3.715574 | 3.766857 | 3.810594 | 3.86091 | 3.905419 |
|  | 5 | 2.88319 | 2.980241 | 3.076269 | 3.150313 | 3.228051 | 3.344205 | 3.458982 | 3.563856 | 3.65291 | 3.704723 | 3.74914 | 3.800495 | 3.846155 |
|  | 10 | 2.814394 | 2.892648 | 2.973187 | 3.037311 | 3.106462 | 3.213161 | 3.322472 | 3.425697 | 3.51589 | 3.569459 | 3.616036 | 3.670655 | 3.719914 |
|  | 15 | 2.764091 | 2.830358 | 2.900386 | 2.957446 | 3.020265 | 3.119812 | 3.225139 | 3.327815 | 3.420202 | 3.476307 | 3.52586 | 3.584906 | 3.639047 |
|  | 20 | 2.726083 | 2.785543 | 2.849323 | 2.902018 | 2.9608 | 3.055654 | 3.158426 | 3.261192 | 3.356028 | 3.414793 | 3.46747 | 3.531222 | 3.590651 |
|  | 30 | 2.634917 | 2.692994 | 2.755173 | 2.806491 | 2.86373 | 2.956253 | 3.05708 | 3.159007 | 3.254516 | 3.314578 | 3.369075 | 3.435947 | 3.499256 |
|  | 40 | 2.544275 | 2.608813 | 2.676405 | 2.731113 | 2.791095 | 2.886059 | 2.987333 | 3.088108 | 3.181706 | 3.240383 | 3.293601 | 3.358962 | 3.420968 |
|  | 50 | 2.472384 | 2.544995 | 2.619188 | 2.677974 | 2.741264 | 2.839355 | 2.94181 | 3.042421 | 3.135406 | 3.193736 | 3.246781 | 3.312209 | 3.374641 |
|  | 60 | 2.429615 | 2.511034 | 2.59229 | 2.655401 | 2.722232 | 2.823925 | 2.928453 | 3.030395 | 3.124834 | 3.184495 | 3.239163 | 3.307246 | 3.372963 |
|  | 70 | 2.366076 | 2.454761 | 2.54145 | 2.607618 | 2.6767 | 2.780265 | 2.885568 | 2.988246 | 3.084342 | 3.145901 | 3.203043 | 3.275315 | 3.346329 |
|  | 80 | 2.304482 | 2.399345 | 2.490462 | 2.558976 | 2.629646 | 2.734323 | 2.840074 | 2.94381 | 3.042662 | 3.107336 | 3.168506 | 3.247588 | 3.327253 |
|  | 90 | 2.222086 | 2.327209 | 2.425585 | 2.497925 | 2.571185 | 2.677638 | 2.783761 | 2.888081 | 2.989215 | 3.056853 | 3.122119 | 3.208517 | 3.297923 |

**Table S6. Centile Values for Cortical Thickness in Males**

|  | **Age** | **C0.4** | **C1** | **C2.5** | **C5** | **C10** | **C25** | **C50** | **C75** | **C90** | **C95** | **C97.5** | **C99** | **C99.6** |
| --- | --- | --- | --- | --- | --- | --- | --- | --- | --- | --- | --- | --- | --- | --- |
| **LThickness** | 3 | 2.574152 | 2.624302 | 2.676905 | 2.719161 | 2.764666 | 2.833491 | 2.900358 | 2.963871 | 3.022896 | 3.058803 | 3.090138 | 3.126665 | 3.159132 |
|  | 5 | 2.535605 | 2.584207 | 2.635401 | 2.676679 | 2.721286 | 2.789067 | 2.8553 | 2.918391 | 2.977029 | 3.012714 | 3.043868 | 3.080203 | 3.112517 |
|  | 10 | 2.440685 | 2.4856 | 2.533392 | 2.572288 | 2.614688 | 2.679871 | 2.744503 | 2.806533 | 2.864219 | 2.899366 | 2.930086 | 2.965966 | 2.997928 |
|  | 15 | 2.351068 | 2.3925 | 2.43704 | 2.473634 | 2.513884 | 2.57652 | 2.639594 | 2.700644 | 2.757507 | 2.792216 | 2.822606 | 2.858168 | 2.889914 |
|  | 20 | 2.282306 | 2.320883 | 2.362749 | 2.397459 | 2.435969 | 2.49662 | 2.55864 | 2.6192 | 2.67572 | 2.710297 | 2.740629 | 2.776201 | 2.808031 |
|  | 30 | 2.207082 | 2.243482 | 2.283345 | 2.316691 | 2.354021 | 2.413599 | 2.475629 | 2.536619 | 2.593298 | 2.627901 | 2.658235 | 2.693803 | 2.725635 |
|  | 40 | 2.154756 | 2.191654 | 2.232032 | 2.265801 | 2.303615 | 2.364054 | 2.4272 | 2.488978 | 2.545651 | 2.579948 | 2.609849 | 2.64473 | 2.675801 |
|  | 50 | 2.104245 | 2.142618 | 2.184411 | 2.21922 | 2.258069 | 2.319936 | 2.384369 | 2.446841 | 2.503342 | 2.537198 | 2.566527 | 2.60054 | 2.630665 |
|  | 60 | 2.049799 | 2.089755 | 2.133034 | 2.168907 | 2.208775 | 2.271953 | 2.337414 | 2.400362 | 2.456683 | 2.49018 | 2.519059 | 2.552399 | 2.581799 |
|  | 70 | 2.012924 | 2.052998 | 2.096486 | 2.132599 | 2.172808 | 2.236691 | 2.303108 | 2.367075 | 2.424289 | 2.458313 | 2.487648 | 2.521518 | 2.551392 |
|  | 80 | 1.965123 | 2.003093 | 2.04471 | 2.07959 | 2.118759 | 2.181696 | 2.248024 | 2.312699 | 2.371147 | 2.406167 | 2.436516 | 2.471732 | 2.502947 |
|  | 90 | 1.903604 | 1.938798 | 1.977823 | 2.010887 | 2.048401 | 2.109517 | 2.175027 | 2.239956 | 2.299501 | 2.335567 | 2.367055 | 2.403861 | 2.436724 |
| **RThickness** | 3 | 2.60364 | 2.645351 | 2.69058 | 2.728032 | 2.769513 | 2.834628 | 2.900832 | 2.96564 | 3.026767 | 3.064423 | 3.097596 | 3.136649 | 3.171718 |
|  | 5 | 2.559735 | 2.600985 | 2.645719 | 2.682767 | 2.723812 | 2.788277 | 2.853882 | 2.918072 | 2.978504 | 3.015686 | 3.048416 | 3.086922 | 3.121478 |
|  | 10 | 2.453154 | 2.493277 | 2.5368 | 2.572861 | 2.612839 | 2.675709 | 2.73984 | 2.802511 | 2.861239 | 2.89726 | 2.92891 | 2.96608 | 2.999383 |
|  | 15 | 2.355186 | 2.39429 | 2.436718 | 2.471886 | 2.510898 | 2.572327 | 2.635131 | 2.696428 | 2.7536 | 2.788559 | 2.819217 | 2.85516 | 2.887312 |
|  | 20 | 2.279728 | 2.318159 | 2.359867 | 2.394452 | 2.432839 | 2.49336 | 2.555372 | 2.615819 | 2.671933 | 2.706139 | 2.736079 | 2.771121 | 2.802417 |
|  | 30 | 2.198221 | 2.236449 | 2.27795 | 2.312388 | 2.350653 | 2.411125 | 2.473353 | 2.533846 | 2.589477 | 2.623178 | 2.652563 | 2.686836 | 2.717349 |
|  | 40 | 2.144582 | 2.183089 | 2.224903 | 2.25962 | 2.298233 | 2.359392 | 2.422582 | 2.483833 | 2.539632 | 2.573223 | 2.6024 | 2.636313 | 2.666409 |
|  | 50 | 2.096918 | 2.135829 | 2.178084 | 2.213184 | 2.252258 | 2.314273 | 2.378593 | 2.440751 | 2.49684 | 2.530392 | 2.559424 | 2.593052 | 2.6228 |
|  | 60 | 2.048432 | 2.087754 | 2.130452 | 2.165931 | 2.205457 | 2.268309 | 2.333731 | 2.396752 | 2.453077 | 2.486557 | 2.515415 | 2.548725 | 2.578099 |
|  | 70 | 2.009775 | 2.049728 | 2.093101 | 2.129146 | 2.169329 | 2.233336 | 2.300181 | 2.364356 | 2.421163 | 2.454713 | 2.48352 | 2.516656 | 2.545784 |
|  | 80 | 1.958049 | 1.998406 | 2.042199 | 2.078595 | 2.119188 | 2.183949 | 2.251788 | 2.316684 | 2.373576 | 2.406959 | 2.435512 | 2.468242 | 2.496922 |
|  | 90 | 1.889333 | 1.929758 | 1.9736 | 2.010031 | 2.050675 | 2.115606 | 2.183813 | 2.248815 | 2.305246 | 2.338145 | 2.366173 | 2.398191 | 2.426158 |
| **L_bankssts** | 3 | 2.412015 | 2.500207 | 2.593437 | 2.668649 | 2.749697 | 2.871556 | 2.987661 | 3.101111 | 3.214719 | 3.287343 | 3.352731 | 3.431192 | 3.502861 |
|  | 5 | 2.365758 | 2.452031 | 2.543385 | 2.617209 | 2.696906 | 2.817072 | 2.932058 | 3.04444 | 3.156552 | 3.228059 | 3.292356 | 3.369422 | 3.439747 |
|  | 10 | 2.252469 | 2.334069 | 2.420833 | 2.491249 | 2.567609 | 2.683559 | 2.79571 | 2.905382 | 3.013769 | 3.082512 | 3.144124 | 3.217765 | 3.284798 |
|  | 15 | 2.148981 | 2.226312 | 2.30888 | 2.376176 | 2.449481 | 2.561579 | 2.67117 | 2.778403 | 2.883403 | 2.949627 | 3.00879 | 3.079307 | 3.143341 |
|  | 20 | 2.076614 | 2.150841 | 2.230419 | 2.295554 | 2.366823 | 2.47658 | 2.585029 | 2.691214 | 2.794242 | 2.858863 | 2.916409 | 2.984809 | 3.04677 |
|  | 30 | 2.002616 | 2.07323 | 2.149553 | 2.212551 | 2.282103 | 2.390735 | 2.500363 | 2.607854 | 2.710301 | 2.773856 | 2.830094 | 2.896575 | 2.956509 |
|  | 40 | 1.954633 | 2.02261 | 2.096673 | 2.158319 | 2.226986 | 2.335761 | 2.447849 | 2.55793 | 2.661038 | 2.724313 | 2.779955 | 2.845377 | 2.904079 |
|  | 50 | 1.910136 | 1.975638 | 2.047572 | 2.107942 | 2.175789 | 2.284791 | 2.39946 | 2.512278 | 2.616186 | 2.679273 | 2.734407 | 2.798891 | 2.856482 |
|  | 60 | 1.858772 | 1.921608 | 1.991153 | 2.05 | 2.116722 | 2.225443 | 2.342179 | 2.457262 | 2.561542 | 2.62419 | 2.678608 | 2.741922 | 2.79821 |
|  | 70 | 1.823809 | 1.884573 | 1.952344 | 2.010158 | 2.07629 | 2.185586 | 2.305337 | 2.423656 | 2.529192 | 2.591938 | 2.646114 | 2.708822 | 2.764319 |
|  | 80 | 1.774916 | 1.833186 | 1.89867 | 1.954986 | 2.019973 | 2.12891 | 2.250683 | 2.371295 | 2.477254 | 2.539606 | 2.593123 | 2.654755 | 2.709057 |
|  | 90 | 1.694883 | 1.749702 | 1.811771 | 1.865579 | 1.92822 | 2.034725 | 2.156164 | 2.276768 | 2.381181 | 2.442002 | 2.4939 | 2.553368 | 2.605535 |
| **L_caudalanteriorcingulate** | 3 | 2.413271 | 2.518408 | 2.629441 | 2.719975 | 2.819884 | 2.979315 | 3.15078 | 3.32252 | 3.482703 | 3.583329 | 3.674679 | 3.786931 | 3.893446 |
|  | 5 | 2.367937 | 2.470842 | 2.579757 | 2.668733 | 2.767082 | 2.924324 | 3.093741 | 3.263611 | 3.42209 | 3.521618 | 3.611933 | 3.722844 | 3.828003 |
|  | 10 | 2.257856 | 2.355412 | 2.459217 | 2.544413 | 2.638962 | 2.790841 | 2.955225 | 3.120502 | 3.274811 | 3.371669 | 3.459477 | 3.567159 | 3.66907 |
|  | 15 | 2.158955 | 2.251729 | 2.350959 | 2.432772 | 2.523927 | 2.671039 | 2.830996 | 2.992289 | 3.143031 | 3.237622 | 3.32331 | 3.428269 | 3.527445 |
|  | 20 | 2.08729 | 2.176474 | 2.272355 | 2.351766 | 2.440596 | 2.584636 | 2.742 | 2.90118 | 3.050148 | 3.143627 | 3.228266 | 3.331846 | 3.429594 |
|  | 30 | 2.017469 | 2.102626 | 2.195109 | 2.272398 | 2.359546 | 2.502229 | 2.659676 | 2.820084 | 2.97076 | 3.065404 | 3.151074 | 3.255813 | 3.354486 |
|  | 40 | 1.977169 | 2.059999 | 2.150792 | 2.227301 | 2.31421 | 2.457808 | 2.617797 | 2.781966 | 2.936809 | 3.034211 | 3.122394 | 3.23015 | 3.331554 |
|  | 50 | 1.955166 | 2.037501 | 2.128442 | 2.2056 | 2.293783 | 2.440579 | 2.605414 | 2.775512 | 2.936418 | 3.03771 | 3.12939 | 3.241319 | 3.346492 |
|  | 60 | 1.935535 | 2.018713 | 2.111121 | 2.189933 | 2.280417 | 2.431874 | 2.602871 | 2.779935 | 2.947614 | 3.053112 | 3.148492 | 3.264738 | 3.373714 |
|  | 70 | 1.926012 | 2.011675 | 2.107243 | 2.189049 | 2.283267 | 2.441536 | 2.620783 | 2.806615 | 2.982452 | 3.092874 | 3.192503 | 3.313612 | 3.426782 |
|  | 80 | 1.923511 | 2.012887 | 2.11291 | 2.198757 | 2.297845 | 2.464683 | 2.65394 | 2.850122 | 3.03539 | 3.151427 | 3.255862 | 3.382426 | 3.500267 |
|  | 90 | 1.912461 | 2.00564 | 2.110199 | 2.200139 | 2.304142 | 2.479583 | 2.67883 | 2.885271 | 3.079824 | 3.201357 | 3.310474 | 3.442326 | 3.564668 |
| **L_caudalmiddlefrontal** | 3 | 2.35314 | 2.435224 | 2.520217 | 2.587863 | 2.660269 | 2.769344 | 2.875421 | 2.974726 | 3.064355 | 3.117867 | 3.164042 | 3.217332 | 3.264269 |
|  | 5 | 2.338514 | 2.41751 | 2.499889 | 2.565849 | 2.636818 | 2.744409 | 2.849797 | 2.948955 | 3.038714 | 3.092422 | 3.138837 | 3.192484 | 3.239808 |
|  | 10 | 2.299226 | 2.371127 | 2.447347 | 2.509248 | 2.576686 | 2.680538 | 2.784112 | 2.882826 | 2.972884 | 3.027086 | 3.074121 | 3.128706 | 3.177053 |
|  | 15 | 2.253516 | 2.31901 | 2.389466 | 2.447443 | 2.511361 | 2.611311 | 2.712814 | 2.810858 | 2.901064 | 2.955707 | 3.003338 | 3.058868 | 3.108276 |
|  | 20 | 2.204115 | 2.264332 | 2.329894 | 2.384442 | 2.445201 | 2.541512 | 2.640952 | 2.7382 | 2.828375 | 2.883329 | 2.931441 | 2.987777 | 3.038126 |
|  | 30 | 2.123859 | 2.17903 | 2.239796 | 2.290912 | 2.348458 | 2.441042 | 2.53848 | 2.634924 | 2.724757 | 2.779723 | 2.827997 | 2.884717 | 2.935595 |
|  | 40 | 2.070149 | 2.124837 | 2.185144 | 2.235951 | 2.293251 | 2.38574 | 2.483583 | 2.580307 | 2.669721 | 2.724155 | 2.771816 | 2.827662 | 2.877628 |
|  | 50 | 2.009767 | 2.066336 | 2.128358 | 2.180349 | 2.238736 | 2.33253 | 2.431306 | 2.527927 | 2.61584 | 2.668768 | 2.714778 | 2.768322 | 2.815916 |
|  | 60 | 1.95426 | 2.013771 | 2.078473 | 2.132317 | 2.192409 | 2.288246 | 2.388438 | 2.485184 | 2.571673 | 2.623111 | 2.667474 | 2.718724 | 2.763958 |
|  | 70 | 1.921141 | 1.981953 | 2.047925 | 2.102739 | 2.163855 | 2.26129 | 2.363237 | 2.46117 | 2.547793 | 2.598943 | 2.642865 | 2.693402 | 2.737844 |
|  | 80 | 1.878883 | 1.938052 | 2.002557 | 2.056409 | 2.11674 | 2.213603 | 2.315906 | 2.414459 | 2.501304 | 2.552466 | 2.596347 | 2.646792 | 2.691121 |
|  | 90 | 1.838679 | 1.895023 | 1.956957 | 2.009071 | 2.0679 | 2.163378 | 2.265629 | 2.364833 | 2.452251 | 2.503765 | 2.54797 | 2.598825 | 2.643556 |
| **L_cuneus** | 3 | 1.890486 | 1.930861 | 1.977325 | 2.018196 | 2.066385 | 2.149533 | 2.245914 | 2.346692 | 2.441324 | 2.499805 | 2.55168 | 2.613375 | 2.669496 |
|  | 5 | 1.841877 | 1.881737 | 1.927625 | 1.968003 | 2.01563 | 2.097849 | 2.193222 | 2.29302 | 2.386798 | 2.444782 | 2.496237 | 2.557455 | 2.613164 |
|  | 10 | 1.72985 | 1.768521 | 1.813082 | 1.852329 | 1.898663 | 1.97876 | 2.071836 | 2.169414 | 2.261273 | 2.318149 | 2.368669 | 2.428835 | 2.483641 |
|  | 15 | 1.634524 | 1.672201 | 1.715658 | 1.753967 | 1.799236 | 1.877595 | 1.968812 | 2.064622 | 2.154979 | 2.211002 | 2.260814 | 2.320193 | 2.374338 |
|  | 20 | 1.563448 | 1.600436 | 1.643133 | 1.680805 | 1.725357 | 1.802569 | 1.892595 | 1.987318 | 2.076797 | 2.132346 | 2.18178 | 2.240763 | 2.294595 |
|  | 30 | 1.497364 | 1.533653 | 1.575587 | 1.612622 | 1.656466 | 1.732562 | 1.821465 | 1.915205 | 2.003938 | 2.059112 | 2.108267 | 2.166985 | 2.220639 |
|  | 40 | 1.466997 | 1.502425 | 1.543374 | 1.579551 | 1.622392 | 1.696783 | 1.783747 | 1.875507 | 1.96243 | 2.016507 | 2.064706 | 2.122306 | 2.17496 |
|  | 50 | 1.449927 | 1.484477 | 1.524413 | 1.559695 | 1.601478 | 1.674038 | 1.758872 | 1.848401 | 1.933226 | 1.986008 | 2.033059 | 2.089294 | 2.14071 |
|  | 60 | 1.430519 | 1.464437 | 1.503652 | 1.538306 | 1.579357 | 1.650671 | 1.734096 | 1.822192 | 1.905711 | 1.957707 | 2.004073 | 2.059511 | 2.110218 |
|  | 70 | 1.422145 | 1.456154 | 1.4955 | 1.53029 | 1.571528 | 1.643235 | 1.727224 | 1.816037 | 1.90035 | 1.952894 | 1.999786 | 2.055895 | 2.107257 |
|  | 80 | 1.39321 | 1.42701 | 1.466143 | 1.500774 | 1.541856 | 1.613374 | 1.697277 | 1.786154 | 1.870671 | 1.923413 | 1.970524 | 2.026952 | 2.078657 |
|  | 90 | 1.341086 | 1.373946 | 1.412017 | 1.445731 | 1.485753 | 1.555497 | 1.637432 | 1.724356 | 1.80714 | 1.858861 | 1.905099 | 1.960529 | 2.011365 |
| **L_entorhinal** | 3 | 2.244337 | 2.3929 | 2.552161 | 2.683108 | 2.827711 | 3.055242 | 3.289378 | 3.514085 | 3.715613 | 3.835742 | 3.939492 | 4.059487 | 4.165514 |
|  | 5 | 2.262382 | 2.406464 | 2.561858 | 2.690292 | 2.832763 | 3.058194 | 3.291633 | 3.516878 | 3.719758 | 3.841059 | 3.94603 | 4.067672 | 4.175356 |
|  | 10 | 2.304661 | 2.438836 | 2.585498 | 2.708147 | 2.845632 | 3.066063 | 3.297804 | 3.524383 | 3.730666 | 3.854946 | 3.963045 | 4.088932 | 4.200914 |
|  | 15 | 2.342714 | 2.468968 | 2.608462 | 2.726254 | 2.859468 | 3.075517 | 3.305755 | 3.533605 | 3.743135 | 3.870291 | 3.981435 | 4.111489 | 4.227728 |
|  | 20 | 2.373051 | 2.49391 | 2.628407 | 2.742736 | 2.87284 | 3.085593 | 3.314592 | 3.54326 | 3.755111 | 3.884378 | 3.997792 | 4.130994 | 4.250485 |
|  | 30 | 2.396628 | 2.514327 | 2.645695 | 2.757678 | 2.88546 | 3.095207 | 3.322063 | 3.549392 | 3.760426 | 3.889397 | 4.00268 | 4.135884 | 4.255521 |
|  | 40 | 2.393161 | 2.511665 | 2.643465 | 2.755457 | 2.882875 | 3.091244 | 3.315637 | 3.539305 | 3.745703 | 3.871298 | 3.981299 | 4.110283 | 4.225814 |
|  | 50 | 2.383397 | 2.502378 | 2.634463 | 2.746511 | 2.873806 | 3.081603 | 3.304952 | 3.526916 | 3.730954 | 3.854774 | 3.963025 | 4.089742 | 4.203053 |
|  | 60 | 2.32695 | 2.453995 | 2.593759 | 2.711356 | 2.843963 | 3.058377 | 3.286298 | 3.510193 | 3.713674 | 3.836153 | 3.942652 | 4.066666 | 4.176992 |
|  | 70 | 2.169973 | 2.325118 | 2.490209 | 2.625195 | 2.773653 | 3.00641 | 3.245435 | 3.472823 | 3.67372 | 3.792286 | 3.894061 | 4.011121 | 4.114025 |
|  | 80 | 1.822767 | 2.065508 | 2.299957 | 2.477266 | 2.660501 | 2.928483 | 3.18496 | 3.415186 | 3.609392 | 3.720565 | 3.814177 | 3.919944 | 4.011361 |
|  | 90 | 1.399331 | 1.729184 | 2.05897 | 2.298929 | 2.532101 | 2.845833 | 3.120941 | 3.351641 | 3.536604 | 3.639184 | 3.723925 | 3.81803 | 3.898083 |
| **L_fusiform** | 3 | 2.495004 | 2.581206 | 2.667177 | 2.733896 | 2.804353 | 2.910496 | 3.016672 | 3.115222 | 3.200453 | 3.250852 | 3.294607 | 3.345901 | 3.392196 |
|  | 5 | 2.470432 | 2.55349 | 2.636855 | 2.701883 | 2.770842 | 2.87524 | 2.98022 | 3.078091 | 3.163029 | 3.213369 | 3.257137 | 3.308516 | 3.354949 |
|  | 10 | 2.405356 | 2.481108 | 2.558272 | 2.619199 | 2.684465 | 2.784467 | 2.886345 | 2.982393 | 3.066494 | 3.116636 | 3.160399 | 3.211957 | 3.258715 |
|  | 15 | 2.340066 | 2.409312 | 2.480805 | 2.537896 | 2.59965 | 2.695386 | 2.794202 | 2.888437 | 2.971726 | 3.021699 | 3.065495 | 3.117296 | 3.164456 |
|  | 20 | 2.295909 | 2.360212 | 2.427373 | 2.48154 | 2.540639 | 2.633251 | 2.730014 | 2.823307 | 2.906515 | 2.956754 | 3.000964 | 3.053464 | 3.101448 |
|  | 30 | 2.254223 | 2.314028 | 2.377284 | 2.42887 | 2.48571 | 2.575891 | 2.671464 | 2.764796 | 2.84893 | 2.900098 | 2.945339 | 2.99931 | 3.048858 |
|  | 40 | 2.213688 | 2.272305 | 2.334539 | 2.385461 | 2.441733 | 2.531334 | 2.626661 | 2.720044 | 2.804404 | 2.855769 | 2.901211 | 2.955445 | 3.005248 |
|  | 50 | 2.178079 | 2.237202 | 2.299942 | 2.35125 | 2.407917 | 2.49806 | 2.593813 | 2.687418 | 2.771769 | 2.823015 | 2.868275 | 2.922188 | 2.971595 |
|  | 60 | 2.133484 | 2.193756 | 2.257525 | 2.309533 | 2.366828 | 2.457652 | 2.55369 | 2.647112 | 2.730883 | 2.781575 | 2.826215 | 2.879226 | 2.927649 |
|  | 70 | 2.105651 | 2.167164 | 2.232117 | 2.284996 | 2.343152 | 2.435124 | 2.532067 | 2.626036 | 2.709987 | 2.760636 | 2.805136 | 2.857855 | 2.905888 |
|  | 80 | 2.059929 | 2.121539 | 2.186573 | 2.239501 | 2.297688 | 2.389647 | 2.486467 | 2.580165 | 2.663713 | 2.714029 | 2.758175 | 2.810396 | 2.857894 |
|  | 90 | 1.997132 | 2.058274 | 2.122795 | 2.175288 | 2.232977 | 2.324094 | 2.419925 | 2.512527 | 2.594944 | 2.644497 | 2.687916 | 2.739202 | 2.785775 |
| **L_inferiorparietal** | 3 | 2.519245 | 2.585924 | 2.656934 | 2.714795 | 2.777966 | 2.875356 | 2.972365 | 3.065523 | 3.151865 | 3.204375 | 3.250234 | 3.303764 | 3.35143 |
|  | 5 | 2.467082 | 2.532193 | 2.601664 | 2.658375 | 2.720401 | 2.816271 | 2.912104 | 3.004219 | 3.089464 | 3.141263 | 3.186481 | 3.239246 | 3.286221 |
|  | 10 | 2.337262 | 2.39849 | 2.464127 | 2.517952 | 2.577087 | 2.669085 | 2.761861 | 2.851258 | 2.933682 | 2.983662 | 3.027246 | 3.078067 | 3.123286 |
|  | 15 | 2.212892 | 2.270402 | 2.332344 | 2.38337 | 2.439683 | 2.527869 | 2.617595 | 2.704279 | 2.783917 | 2.832111 | 2.874097 | 2.923018 | 2.966524 |
|  | 20 | 2.121234 | 2.175878 | 2.235008 | 2.28394 | 2.338189 | 2.423706 | 2.511503 | 2.596559 | 2.674434 | 2.721471 | 2.76241 | 2.81008 | 2.852455 |
|  | 30 | 2.04469 | 2.096254 | 2.152574 | 2.199608 | 2.252231 | 2.336311 | 2.424222 | 2.509899 | 2.587852 | 2.634767 | 2.675532 | 2.722943 | 2.765055 |
|  | 40 | 2.009599 | 2.058986 | 2.113423 | 2.159299 | 2.211103 | 2.295013 | 2.384391 | 2.472072 | 2.551401 | 2.598993 | 2.640284 | 2.688262 | 2.730853 |
|  | 50 | 1.970403 | 2.017373 | 2.069612 | 2.114033 | 2.164657 | 2.247797 | 2.338039 | 2.4272 | 2.507472 | 2.555492 | 2.597102 | 2.645414 | 2.688288 |
|  | 60 | 1.923412 | 1.967684 | 2.017352 | 2.05996 | 2.108963 | 2.190567 | 2.280841 | 2.370719 | 2.451288 | 2.499363 | 2.540978 | 2.58927 | 2.632117 |
|  | 70 | 1.913535 | 1.955923 | 2.003876 | 2.045368 | 2.09352 | 2.174825 | 2.266503 | 2.358522 | 2.440702 | 2.48963 | 2.531946 | 2.581032 | 2.624585 |
|  | 80 | 1.858614 | 1.89818 | 1.9433 | 1.982668 | 2.02876 | 2.107669 | 2.198366 | 2.290176 | 2.37191 | 2.420476 | 2.462451 | 2.51113 | 2.55433 |
|  | 90 | 1.761872 | 1.797911 | 1.839325 | 1.875752 | 1.918773 | 1.99344 | 2.080921 | 2.170267 | 2.249598 | 2.296658 | 2.33731 | 2.384454 | 2.426305 |
| **L_inferiortemporal** | 3 | 2.400443 | 2.530164 | 2.650054 | 2.737727 | 2.826144 | 2.953086 | 3.075036 | 3.186234 | 3.282734 | 3.340607 | 3.391636 | 3.452653 | 3.509055 |
|  | 5 | 2.400369 | 2.516715 | 2.627108 | 2.709507 | 2.793931 | 2.917188 | 3.037372 | 3.147872 | 3.243969 | 3.301513 | 3.352123 | 3.412419 | 3.467897 |
|  | 10 | 2.3853 | 2.47679 | 2.568515 | 2.640023 | 2.715853 | 2.830798 | 2.946845 | 3.055842 | 3.151414 | 3.208617 | 3.25875 | 3.318129 | 3.372326 |
|  | 15 | 2.363782 | 2.437833 | 2.515359 | 2.578009 | 2.64647 | 2.753901 | 2.8662 | 2.974322 | 3.070463 | 3.128312 | 3.179066 | 3.239129 | 3.293808 |
|  | 20 | 2.347052 | 2.409138 | 2.476297 | 2.532144 | 2.594721 | 2.69598 | 2.805382 | 2.913534 | 3.011451 | 3.070937 | 3.123375 | 3.185631 | 3.242414 |
|  | 30 | 2.311723 | 2.366313 | 2.426875 | 2.478382 | 2.537255 | 2.634854 | 2.742981 | 2.851824 | 2.951311 | 3.011905 | 3.065283 | 3.128487 | 3.185873 |
|  | 40 | 2.267048 | 2.321704 | 2.382584 | 2.434518 | 2.493997 | 2.592702 | 2.701865 | 2.811109 | 2.910026 | 2.969697 | 3.021838 | 3.083009 | 3.137969 |
|  | 50 | 2.224799 | 2.282575 | 2.346626 | 2.400993 | 2.462932 | 2.564884 | 2.676298 | 2.786222 | 2.884262 | 2.942666 | 2.993223 | 3.051949 | 3.104154 |
|  | 60 | 2.164991 | 2.224904 | 2.29105 | 2.346965 | 2.410405 | 2.514193 | 2.626663 | 2.736585 | 2.833682 | 2.891076 | 2.940477 | 2.997518 | 3.047908 |
|  | 70 | 2.151296 | 2.20637 | 2.268141 | 2.321133 | 2.382093 | 2.483687 | 2.596276 | 2.708687 | 2.809818 | 2.870361 | 2.922907 | 2.984058 | 3.038489 |
|  | 80 | 2.135947 | 2.187269 | 2.245555 | 2.296162 | 2.355063 | 2.454827 | 2.56768 | 2.682701 | 2.788128 | 2.852105 | 2.908142 | 2.973945 | 3.03304 |
|  | 90 | 2.081564 | 2.131442 | 2.188327 | 2.237918 | 2.295866 | 2.394564 | 2.507011 | 2.62245 | 2.72896 | 2.793906 | 2.850978 | 2.918213 | 2.978786 |
| **L_isthmuscingulate** | 3 | 2.509356 | 2.561547 | 2.622359 | 2.676532 | 2.741252 | 2.855168 | 2.990948 | 3.137204 | 3.278519 | 3.367867 | 3.448466 | 3.546021 | 3.636413 |
|  | 5 | 2.469957 | 2.521617 | 2.58184 | 2.63552 | 2.699698 | 2.812814 | 2.947922 | 3.093396 | 3.23353 | 3.321932 | 3.401558 | 3.497794 | 3.586838 |
|  | 10 | 2.372781 | 2.423144 | 2.481923 | 2.534393 | 2.597237 | 2.708364 | 2.841774 | 2.985269 | 3.122469 | 3.208535 | 3.28577 | 3.378782 | 3.464538 |
|  | 15 | 2.278206 | 2.327385 | 2.384846 | 2.436204 | 2.497814 | 2.607078 | 2.738839 | 2.880405 | 3.014825 | 3.098707 | 3.173723 | 3.26376 | 3.3465 |
|  | 20 | 2.195838 | 2.244131 | 2.300596 | 2.351109 | 2.411773 | 2.519586 | 2.650016 | 2.789965 | 2.922055 | 3.004109 | 3.077268 | 3.164818 | 3.245039 |
|  | 30 | 2.094524 | 2.142253 | 2.198025 | 2.247892 | 2.307757 | 2.414108 | 2.54271 | 2.680118 | 2.808814 | 2.888283 | 2.958845 | 3.042939 | 3.119669 |
|  | 40 | 2.02396 | 2.072455 | 2.12886 | 2.179033 | 2.238911 | 2.344237 | 2.469741 | 2.603047 | 2.728418 | 2.806041 | 2.875048 | 2.957352 | 3.032477 |
|  | 50 | 1.955099 | 2.006319 | 2.065433 | 2.117556 | 2.179127 | 2.285545 | 2.409027 | 2.539439 | 2.664172 | 2.742306 | 2.812246 | 2.896159 | 2.973155 |
|  | 60 | 1.880434 | 1.936186 | 1.999957 | 2.055611 | 2.120572 | 2.230551 | 2.354162 | 2.48406 | 2.611269 | 2.692238 | 2.765412 | 2.853952 | 2.935816 |
|  | 70 | 1.810943 | 1.872169 | 1.941642 | 2.001724 | 2.071119 | 2.186505 | 2.312589 | 2.444474 | 2.576273 | 2.661278 | 2.738702 | 2.833027 | 2.920771 |
|  | 80 | 1.736884 | 1.803958 | 1.87954 | 1.944396 | 2.018639 | 2.140234 | 2.270006 | 2.40512 | 2.542229 | 2.631503 | 2.713259 | 2.813324 | 2.90678 |
|  | 90 | 1.659812 | 1.733407 | 1.815756 | 1.885868 | 1.965423 | 2.093832 | 2.227802 | 2.366584 | 2.509307 | 2.602973 | 2.689129 | 2.794959 | 2.894095 |
| **L_lateraloccipital** | 3 | 2.180289 | 2.220133 | 2.264951 | 2.303497 | 2.347938 | 2.422197 | 2.504705 | 2.587212 | 2.661471 | 2.705912 | 2.744458 | 2.789277 | 2.829121 |
|  | 5 | 2.137853 | 2.177719 | 2.222561 | 2.261127 | 2.305592 | 2.37989 | 2.462442 | 2.544993 | 2.619291 | 2.663756 | 2.702323 | 2.747164 | 2.78703 |
|  | 10 | 2.032335 | 2.072253 | 2.117154 | 2.155772 | 2.200296 | 2.274693 | 2.357353 | 2.440014 | 2.514411 | 2.558935 | 2.597552 | 2.642454 | 2.682372 |
|  | 15 | 1.93102 | 1.970991 | 2.015952 | 2.054621 | 2.099203 | 2.173699 | 2.256469 | 2.339239 | 2.413734 | 2.458317 | 2.496986 | 2.541946 | 2.581917 |
|  | 20 | 1.856178 | 1.896201 | 1.941221 | 1.979941 | 2.024582 | 2.099175 | 2.182054 | 2.264932 | 2.339526 | 2.384167 | 2.422886 | 2.467906 | 2.507929 |
|  | 30 | 1.813264 | 1.853391 | 1.898526 | 1.937346 | 1.982102 | 2.056888 | 2.13998 | 2.223072 | 2.297858 | 2.342614 | 2.381433 | 2.426569 | 2.466696 |
|  | 40 | 1.790054 | 1.830282 | 1.875532 | 1.914449 | 1.959318 | 2.034293 | 2.117595 | 2.200897 | 2.275872 | 2.320741 | 2.359659 | 2.404909 | 2.445137 |
|  | 50 | 1.779693 | 1.820023 | 1.865386 | 1.904402 | 1.949384 | 2.024547 | 2.108059 | 2.191571 | 2.266735 | 2.311717 | 2.350732 | 2.396096 | 2.436425 |
|  | 60 | 1.741305 | 1.781736 | 1.827215 | 1.866329 | 1.911426 | 1.986779 | 2.070503 | 2.154227 | 2.229581 | 2.274677 | 2.313791 | 2.35927 | 2.399701 |
|  | 70 | 1.715134 | 1.755669 | 1.801264 | 1.840478 | 1.88569 | 1.961236 | 2.045174 | 2.129112 | 2.204659 | 2.24987 | 2.289085 | 2.33468 | 2.375215 |
|  | 80 | 1.661643 | 1.702282 | 1.747995 | 1.78731 | 1.832639 | 1.90838 | 1.992534 | 2.076688 | 2.15243 | 2.197758 | 2.237073 | 2.282786 | 2.323425 |
|  | 90 | 1.540737 | 1.581481 | 1.627312 | 1.666729 | 1.712174 | 1.788111 | 1.872482 | 1.956854 | 2.032791 | 2.078236 | 2.117653 | 2.163484 | 2.204228 |
| **L_lateralorbitofrontal** | 3 | 2.662779 | 2.756058 | 2.851105 | 2.926131 | 3.006378 | 3.128665 | 3.251521 | 3.364551 | 3.460298 | 3.51561 | 3.562667 | 3.616548 | 3.66389 |
|  | 5 | 2.604455 | 2.695169 | 2.78782 | 2.861095 | 2.939594 | 3.059452 | 3.18013 | 3.291381 | 3.385786 | 3.440392 | 3.486889 | 3.540175 | 3.587036 |
|  | 10 | 2.472634 | 2.555353 | 2.640755 | 2.7089 | 2.782452 | 2.895774 | 3.011028 | 3.118243 | 3.209924 | 3.263243 | 3.308807 | 3.361212 | 3.40746 |
|  | 15 | 2.381053 | 2.450086 | 2.523413 | 2.583363 | 2.649452 | 2.753973 | 2.86349 | 2.96816 | 3.059726 | 3.113832 | 3.160561 | 3.214858 | 3.263264 |
|  | 20 | 2.333926 | 2.389219 | 2.449963 | 2.501173 | 2.559247 | 2.654594 | 2.759147 | 2.863585 | 2.958621 | 3.016413 | 3.067312 | 3.127617 | 3.182441 |
|  | 30 | 2.242378 | 2.293835 | 2.35083 | 2.399262 | 2.454618 | 2.546522 | 2.648814 | 2.752648 | 2.848646 | 2.907754 | 2.960286 | 3.02311 | 3.08079 |
|  | 40 | 2.178825 | 2.233876 | 2.294268 | 2.345128 | 2.402773 | 2.497403 | 2.601295 | 2.705381 | 2.800526 | 2.858645 | 2.910024 | 2.971157 | 3.027002 |
|  | 50 | 2.164031 | 2.210928 | 2.263604 | 2.309001 | 2.361638 | 2.450911 | 2.553274 | 2.660749 | 2.763638 | 2.828813 | 2.887979 | 2.960348 | 3.028408 |
|  | 60 | 2.106942 | 2.156994 | 2.212753 | 2.260427 | 2.315284 | 2.407339 | 2.511483 | 2.619353 | 2.72134 | 2.785347 | 2.843076 | 2.913223 | 2.978749 |
|  | 70 | 2.030956 | 2.090625 | 2.155429 | 2.209524 | 2.270362 | 2.369317 | 2.476958 | 2.584147 | 2.681892 | 2.741612 | 2.794474 | 2.857504 | 2.915254 |
|  | 80 | 1.999531 | 2.059092 | 2.123901 | 2.178108 | 2.239197 | 2.338883 | 2.447856 | 2.557037 | 2.657279 | 2.718883 | 2.773658 | 2.839287 | 2.899735 |
|  | 90 | 1.987032 | 2.039672 | 2.098068 | 2.147828 | 2.204939 | 2.300571 | 2.408756 | 2.521221 | 2.628315 | 2.696056 | 2.757572 | 2.832927 | 2.903975 |
| **L_lingual** | 3 | 2.06873 | 2.108064 | 2.152694 | 2.191429 | 2.236525 | 2.313024 | 2.399905 | 2.489044 | 2.571462 | 2.621883 | 2.666336 | 2.718918 | 2.766522 |
|  | 5 | 2.025556 | 2.064384 | 2.108427 | 2.146646 | 2.191133 | 2.266591 | 2.352291 | 2.440245 | 2.521606 | 2.571407 | 2.615333 | 2.667318 | 2.71441 |
|  | 10 | 1.922488 | 1.960121 | 2.002773 | 2.039759 | 2.082791 | 2.155753 | 2.238619 | 2.323723 | 2.402553 | 2.450875 | 2.493549 | 2.544127 | 2.590021 |
|  | 15 | 1.831024 | 1.867652 | 1.909119 | 1.945046 | 1.986819 | 2.057602 | 2.137984 | 2.220596 | 2.297231 | 2.344285 | 2.3859 | 2.435307 | 2.480228 |
|  | 20 | 1.764284 | 1.800357 | 1.841138 | 1.87643 | 1.917427 | 1.986836 | 2.065633 | 2.14667 | 2.221971 | 2.268293 | 2.309332 | 2.358152 | 2.402643 |
|  | 30 | 1.703859 | 1.740242 | 1.781202 | 1.816526 | 1.857445 | 1.926533 | 2.004851 | 2.0855 | 2.160734 | 2.207238 | 2.248615 | 2.298093 | 2.343458 |
|  | 40 | 1.667327 | 1.704541 | 1.746156 | 1.781843 | 1.822993 | 1.892144 | 1.9703 | 2.050865 | 2.126408 | 2.173408 | 2.215482 | 2.266164 | 2.313037 |
|  | 50 | 1.631489 | 1.669854 | 1.712317 | 1.748412 | 1.789733 | 1.858646 | 1.93613 | 2.016072 | 2.09156 | 2.138965 | 2.181774 | 2.233894 | 2.282709 |
|  | 60 | 1.58479 | 1.624838 | 1.668476 | 1.705078 | 1.746517 | 1.814818 | 1.89098 | 1.969623 | 2.044638 | 2.092392 | 2.136077 | 2.190108 | 2.241669 |
|  | 70 | 1.545548 | 1.588911 | 1.635072 | 1.673017 | 1.715258 | 1.783636 | 1.858905 | 1.936685 | 2.011989 | 2.06091 | 2.10653 | 2.164299 | 2.220982 |
|  | 80 | 1.49352 | 1.542534 | 1.59294 | 1.633141 | 1.676768 | 1.745473 | 1.819597 | 1.896258 | 1.972155 | 2.022987 | 2.071779 | 2.135787 | 2.201271 |
|  | 90 | 1.41906 | 1.478237 | 1.536124 | 1.580268 | 1.626373 | 1.696001 | 1.76882 | 1.844206 | 1.921431 | 1.975603 | 2.029929 | 2.105108 | 2.187009 |
| **L_medialorbitofrontal** | 3 | 2.451144 | 2.533081 | 2.618154 | 2.686504 | 2.76098 | 2.878085 | 3.002255 | 3.125578 | 3.24035 | 3.312577 | 3.378342 | 3.459508 | 3.53696 |
|  | 5 | 2.395353 | 2.473046 | 2.554224 | 2.619806 | 2.691617 | 2.805209 | 2.926443 | 3.047494 | 3.160582 | 3.231909 | 3.296936 | 3.377278 | 3.454016 |
|  | 10 | 2.268076 | 2.336543 | 2.409137 | 2.468556 | 2.534379 | 2.640025 | 2.754612 | 2.870589 | 2.980036 | 3.049492 | 3.113047 | 3.191826 | 3.267298 |
|  | 15 | 2.172365 | 2.233711 | 2.299618 | 2.354212 | 2.415349 | 2.514843 | 2.624475 | 2.736975 | 2.844293 | 2.912872 | 2.975898 | 3.054343 | 3.12979 |
|  | 20 | 2.114669 | 2.171146 | 2.232497 | 2.283835 | 2.341868 | 2.437457 | 2.544272 | 2.655253 | 2.762178 | 2.830952 | 2.894422 | 2.973731 | 3.050305 |
|  | 30 | 2.049857 | 2.101739 | 2.158868 | 2.20727 | 2.262606 | 2.355068 | 2.460037 | 2.570504 | 2.677862 | 2.747227 | 2.81138 | 2.891659 | 2.969236 |
|  | 40 | 2.009539 | 2.060901 | 2.117743 | 2.166106 | 2.221586 | 2.314597 | 2.420355 | 2.531417 | 2.638726 | 2.707581 | 2.770863 | 2.849454 | 2.924724 |
|  | 50 | 1.965895 | 2.018879 | 2.077476 | 2.127267 | 2.184273 | 2.279446 | 2.386789 | 2.498164 | 2.604149 | 2.671205 | 2.732133 | 2.806834 | 2.877351 |
|  | 60 | 1.914588 | 1.969367 | 2.02983 | 2.081087 | 2.139598 | 2.236771 | 2.345385 | 2.456692 | 2.56106 | 2.626225 | 2.684818 | 2.755827 | 2.822004 |
|  | 70 | 1.887822 | 1.943786 | 2.005617 | 2.058059 | 2.117922 | 2.217238 | 2.327901 | 2.440666 | 2.545583 | 2.610604 | 2.668712 | 2.738649 | 2.803324 |
|  | 80 | 1.87461 | 1.930663 | 1.992863 | 2.045818 | 2.106462 | 2.207438 | 2.320302 | 2.435439 | 2.542456 | 2.608642 | 2.667664 | 2.738505 | 2.803796 |
|  | 90 | 1.869936 | 1.925902 | 1.988328 | 2.041722 | 2.103118 | 2.205858 | 2.321277 | 2.439424 | 2.549398 | 2.617412 | 2.678025 | 2.750693 | 2.81756 |
| **L_middletemporal** | 3 | 2.609694 | 2.726865 | 2.840901 | 2.927422 | 3.016642 | 3.145751 | 3.266607 | 3.375135 | 3.468785 | 3.523053 | 3.569015 | 3.621156 | 3.666347 |
|  | 5 | 2.597483 | 2.705839 | 2.813165 | 2.895687 | 2.981662 | 3.10748 | 3.226568 | 3.334481 | 3.428283 | 3.482904 | 3.529307 | 3.582098 | 3.627977 |
|  | 10 | 2.560605 | 2.651088 | 2.744104 | 2.817747 | 2.896294 | 3.014325 | 3.129119 | 3.235554 | 3.32985 | 3.385472 | 3.433117 | 3.487743 | 3.535567 |
|  | 15 | 2.522184 | 2.599116 | 2.680573 | 2.74668 | 2.818685 | 2.929624 | 3.040469 | 3.145738 | 3.240951 | 3.297932 | 3.347206 | 3.404211 | 3.454555 |
|  | 20 | 2.48922 | 2.556377 | 2.629099 | 2.6893 | 2.756038 | 2.861156 | 2.968843 | 3.073535 | 3.170245 | 3.229008 | 3.280344 | 3.340325 | 3.393813 |
|  | 30 | 2.42728 | 2.485336 | 2.549585 | 2.60385 | 2.665137 | 2.764051 | 2.868353 | 2.972745 | 3.071913 | 3.13344 | 3.187969 | 3.252597 | 3.311056 |
|  | 40 | 2.383631 | 2.440307 | 2.503315 | 2.55676 | 2.617365 | 2.715698 | 2.820042 | 2.925271 | 3.026084 | 3.089033 | 3.145071 | 3.211781 | 3.272394 |
|  | 50 | 2.323118 | 2.38233 | 2.447817 | 2.503088 | 2.56546 | 2.665979 | 2.771729 | 2.8776 | 2.97846 | 3.041156 | 3.096785 | 3.162785 | 3.222543 |
|  | 60 | 2.229923 | 2.294351 | 2.364726 | 2.423435 | 2.48896 | 2.593011 | 2.700517 | 2.806408 | 2.905903 | 2.967097 | 3.020988 | 3.084442 | 3.141456 |
|  | 70 | 2.15984 | 2.227722 | 2.301442 | 2.362614 | 2.430549 | 2.537715 | 2.647552 | 2.755053 | 2.855612 | 2.917246 | 2.971392 | 3.034988 | 3.091986 |
|  | 80 | 2.090288 | 2.160334 | 2.236218 | 2.299041 | 2.36866 | 2.478161 | 2.589984 | 2.699194 | 2.801285 | 2.863823 | 2.918736 | 2.983201 | 3.040945 |
|  | 90 | 2.019658 | 2.091933 | 2.170026 | 2.234523 | 2.305837 | 2.417671 | 2.53145 | 2.642327 | 2.745909 | 2.809322 | 2.864977 | 2.930278 | 2.988741 |
| **L_parahippocampal** | 3 | 2.285893 | 2.365317 | 2.457658 | 2.540009 | 2.6389 | 2.815671 | 3.031411 | 3.249377 | 3.431363 | 3.534547 | 3.621247 | 3.719321 | 3.804414 |
|  | 5 | 2.255085 | 2.334857 | 2.427559 | 2.510185 | 2.60934 | 2.786382 | 3.00212 | 3.220098 | 3.4024 | 3.505893 | 3.592918 | 3.691423 | 3.776941 |
|  | 10 | 2.179204 | 2.259814 | 2.353377 | 2.436655 | 2.536428 | 2.714067 | 2.929705 | 3.147616 | 3.330623 | 3.434843 | 3.522642 | 3.622187 | 3.708734 |
|  | 15 | 2.106015 | 2.187355 | 2.281659 | 2.365485 | 2.46576 | 2.643816 | 2.85917 | 3.076829 | 3.260351 | 3.365174 | 3.453638 | 3.554091 | 3.641548 |
|  | 20 | 2.040552 | 2.122627 | 2.217689 | 2.302094 | 2.402926 | 2.581552 | 2.796909 | 3.0146 | 3.198793 | 3.304271 | 3.393426 | 3.494801 | 3.583168 |
|  | 30 | 1.959128 | 2.04251 | 2.138951 | 2.224441 | 2.326373 | 2.506349 | 2.72234 | 2.940702 | 3.126366 | 3.233071 | 3.323456 | 3.426424 | 3.516331 |
|  | 40 | 1.914969 | 1.996991 | 2.091858 | 2.175952 | 2.27622 | 2.45326 | 2.665736 | 2.880512 | 3.06307 | 3.167968 | 3.25681 | 3.358008 | 3.446359 |
|  | 50 | 1.889601 | 1.968987 | 2.061009 | 2.142787 | 2.240582 | 2.414135 | 2.623871 | 2.835843 | 3.014707 | 3.116926 | 3.20322 | 3.301236 | 3.386591 |
|  | 60 | 1.867485 | 1.945227 | 2.035652 | 2.116324 | 2.213234 | 2.386546 | 2.59818 | 2.812068 | 2.990669 | 3.091935 | 3.177023 | 3.273272 | 3.356781 |
|  | 70 | 1.818665 | 1.896389 | 1.987128 | 2.06842 | 2.166548 | 2.343484 | 2.561901 | 2.782709 | 2.965175 | 3.067808 | 3.153636 | 3.25032 | 3.333895 |
|  | 80 | 1.743972 | 1.823361 | 1.916417 | 2.000158 | 2.101763 | 2.28656 | 2.517271 | 2.750645 | 2.941521 | 3.048024 | 3.136662 | 3.236096 | 3.321731 |
|  | 90 | 1.658943 | 1.740496 | 1.836488 | 1.923275 | 2.029141 | 2.223429 | 2.468806 | 2.717196 | 2.918271 | 3.029544 | 3.1217 | 3.22464 | 3.312961 |
| **L_paracentral** | 3 | 2.329469 | 2.388236 | 2.453423 | 2.508748 | 2.57173 | 2.675159 | 2.787604 | 2.897656 | 2.994812 | 3.05215 | 3.101416 | 3.158173 | 3.208173 |
|  | 5 | 2.29184 | 2.349435 | 2.413433 | 2.467841 | 2.529875 | 2.631963 | 2.743243 | 2.852432 | 2.949043 | 3.006151 | 3.055272 | 3.111921 | 3.161878 |
|  | 10 | 2.196278 | 2.250913 | 2.311892 | 2.36395 | 2.423541 | 2.52214 | 2.630338 | 2.737204 | 2.832315 | 2.888773 | 2.937472 | 2.99379 | 3.043589 |
|  | 15 | 2.09984 | 2.151513 | 2.209442 | 2.259104 | 2.316183 | 2.411148 | 2.516085 | 2.620446 | 2.713906 | 2.769635 | 2.817851 | 2.873777 | 2.923374 |
|  | 20 | 2.020043 | 2.069501 | 2.125143 | 2.173005 | 2.228196 | 2.320439 | 2.422955 | 2.5255 | 2.617818 | 2.673077 | 2.721012 | 2.776754 | 2.826316 |
|  | 30 | 1.931142 | 1.980374 | 2.03577 | 2.083429 | 2.138394 | 2.230278 | 2.332425 | 2.434629 | 2.526663 | 2.581762 | 2.629565 | 2.68516 | 2.734597 |
|  | 40 | 1.864762 | 1.918034 | 1.977381 | 2.027957 | 2.085758 | 2.181186 | 2.285629 | 2.388523 | 2.479898 | 2.534054 | 2.580719 | 2.634631 | 2.682257 |
|  | 50 | 1.795296 | 1.856104 | 1.922556 | 1.978197 | 2.040754 | 2.141828 | 2.249611 | 2.353208 | 2.443265 | 2.495846 | 2.540707 | 2.592043 | 2.636973 |
|  | 60 | 1.746697 | 1.809862 | 1.87864 | 1.936046 | 2.000402 | 2.104007 | 2.214027 | 2.319375 | 2.410668 | 2.463857 | 2.509175 | 2.560966 | 2.606237 |
|  | 70 | 1.73677 | 1.791229 | 1.852192 | 1.904385 | 1.964297 | 2.063807 | 2.173538 | 2.282449 | 2.379813 | 2.437797 | 2.487923 | 2.546017 | 2.597498 |
|  | 80 | 1.733385 | 1.776728 | 1.826902 | 1.871294 | 1.923944 | 2.015568 | 2.122998 | 2.236716 | 2.344773 | 2.41216 | 2.472325 | 2.54435 | 2.610308 |
|  | 90 | 1.697761 | 1.733157 | 1.775076 | 1.813057 | 1.859256 | 1.942876 | 2.046693 | 2.164307 | 2.28451 | 2.364186 | 2.438715 | 2.532544 | 2.623302 |
| **L_parsopercularis** | 3 | 2.390015 | 2.500442 | 2.604359 | 2.681476 | 2.760082 | 2.87383 | 2.982803 | 3.080166 | 3.161667 | 3.208737 | 3.248944 | 3.295318 | 3.336487 |
|  | 5 | 2.388315 | 2.48796 | 2.584104 | 2.656823 | 2.732045 | 2.842673 | 2.950413 | 3.047964 | 3.130469 | 3.178443 | 3.219601 | 3.26727 | 3.309756 |
|  | 10 | 2.371668 | 2.450647 | 2.530873 | 2.594069 | 2.66162 | 2.7648 | 2.869426 | 2.967471 | 3.05272 | 3.103231 | 3.147101 | 3.198513 | 3.24487 |
|  | 15 | 2.34202 | 2.406404 | 2.474326 | 2.529586 | 2.590338 | 2.686428 | 2.787847 | 2.886445 | 2.974918 | 3.028525 | 3.075791 | 3.132012 | 3.183466 |
|  | 20 | 2.303778 | 2.358756 | 2.418214 | 2.467696 | 2.523245 | 2.613573 | 2.712208 | 2.81137 | 2.903116 | 2.959989 | 3.010942 | 3.072538 | 3.129858 |
|  | 30 | 2.202544 | 2.254616 | 2.311295 | 2.358753 | 2.412345 | 2.500205 | 2.59717 | 2.695753 | 2.787966 | 2.84562 | 2.897596 | 2.960839 | 3.020097 |
|  | 40 | 2.099608 | 2.158034 | 2.220431 | 2.271764 | 2.328783 | 2.420214 | 2.518382 | 2.615491 | 2.704067 | 2.758413 | 2.806762 | 2.864803 | 2.918434 |
|  | 50 | 2.024438 | 2.085393 | 2.150001 | 2.202795 | 2.261078 | 2.3538 | 2.452431 | 2.549153 | 2.636717 | 2.690159 | 2.737535 | 2.794209 | 2.846395 |
|  | 60 | 1.981362 | 2.040206 | 2.102983 | 2.154584 | 2.211863 | 2.303654 | 2.40218 | 2.499684 | 2.588713 | 2.643403 | 2.692112 | 2.750658 | 2.804836 |
|  | 70 | 1.954341 | 2.011429 | 2.072714 | 2.12338 | 2.179922 | 2.27119 | 2.370056 | 2.468818 | 2.559804 | 2.616081 | 2.666451 | 2.727304 | 2.783917 |
|  | 80 | 1.937437 | 1.990848 | 2.048856 | 2.097336 | 2.152002 | 2.241495 | 2.3402 | 2.440654 | 2.534861 | 2.593938 | 2.64734 | 2.712526 | 2.773829 |
|  | 90 | 1.920821 | 1.970874 | 2.025814 | 2.072201 | 2.125025 | 2.212717 | 2.311228 | 2.413473 | 2.511241 | 2.573504 | 2.630429 | 2.700757 | 2.767749 |
| **L_parsorbitalis** | 3 | 2.515258 | 2.636424 | 2.766401 | 2.873544 | 2.992426 | 3.181596 | 3.380307 | 3.569348 | 3.732847 | 3.828197 | 3.909596 | 4.002897 | 4.084775 |
|  | 5 | 2.505685 | 2.615038 | 2.733163 | 2.83112 | 2.940383 | 3.115392 | 3.300618 | 3.478046 | 3.632402 | 3.72279 | 3.800164 | 3.889088 | 3.96733 |
|  | 10 | 2.463335 | 2.549264 | 2.643317 | 2.722234 | 2.811201 | 2.955674 | 3.111105 | 3.262329 | 3.395702 | 3.474578 | 3.542551 | 3.621187 | 3.690837 |
|  | 15 | 2.359476 | 2.435547 | 2.519512 | 2.590518 | 2.671161 | 2.803446 | 2.947585 | 3.089645 | 3.216446 | 3.292116 | 3.357738 | 3.43414 | 3.502252 |
|  | 20 | 2.22354 | 2.300994 | 2.387094 | 2.460403 | 2.544216 | 2.68299 | 2.83606 | 2.98888 | 3.126995 | 3.210211 | 3.282874 | 3.368067 | 3.444567 |
|  | 30 | 2.19854 | 2.264205 | 2.337659 | 2.400604 | 2.473054 | 2.594242 | 2.729893 | 2.867663 | 2.994442 | 3.071971 | 3.14042 | 3.221619 | 3.295449 |
|  | 40 | 2.115336 | 2.183793 | 2.260219 | 2.325628 | 2.400874 | 2.52681 | 2.668217 | 2.812758 | 2.946993 | 3.029819 | 3.103487 | 3.19161 | 3.272492 |
|  | 50 | 2.022341 | 2.09711 | 2.179097 | 2.248133 | 2.32639 | 2.45497 | 2.596462 | 2.738782 | 2.869574 | 2.94987 | 3.021149 | 3.106349 | 3.184579 |
|  | 60 | 1.871767 | 1.963496 | 2.061275 | 2.141582 | 2.230639 | 2.373098 | 2.525499 | 2.675552 | 2.811676 | 2.894795 | 2.968464 | 3.056538 | 3.137554 |
|  | 70 | 1.858445 | 1.945549 | 2.038685 | 2.115454 | 2.200977 | 2.338998 | 2.489131 | 2.640668 | 2.782511 | 2.871655 | 2.952516 | 3.051709 | 3.145596 |
|  | 80 | 1.905934 | 1.977611 | 2.054984 | 2.119427 | 2.192078 | 2.311803 | 2.446811 | 2.590173 | 2.733006 | 2.828057 | 2.918368 | 3.035114 | 3.152328 |
|  | 90 | 2.045266 | 2.097577 | 2.153052 | 2.198589 | 2.249374 | 2.3324 | 2.426414 | 2.528821 | 2.635767 | 2.710709 | 2.78532 | 2.88745 | 2.997334 |
| **L_parstriangularis** | 3 | 2.535206 | 2.589138 | 2.649803 | 2.701978 | 2.762133 | 2.862649 | 2.974329 | 3.086009 | 3.186525 | 3.24668 | 3.298855 | 3.35952 | 3.413452 |
|  | 5 | 2.48649 | 2.540353 | 2.600941 | 2.65305 | 2.713128 | 2.813515 | 2.925053 | 3.036591 | 3.136979 | 3.197057 | 3.249166 | 3.309754 | 3.363617 |
|  | 10 | 2.361682 | 2.415374 | 2.475769 | 2.527711 | 2.587598 | 2.687666 | 2.798849 | 2.910032 | 3.010101 | 3.069987 | 3.12193 | 3.182325 | 3.236017 |
|  | 15 | 2.237199 | 2.29072 | 2.350923 | 2.4027 | 2.462397 | 2.562147 | 2.672976 | 2.783805 | 2.883555 | 2.943252 | 2.995029 | 3.055232 | 3.108753 |
|  | 20 | 2.143737 | 2.197088 | 2.2571 | 2.308713 | 2.36822 | 2.467654 | 2.578131 | 2.688609 | 2.788042 | 2.847549 | 2.899162 | 2.959174 | 3.012525 |
|  | 30 | 2.037145 | 2.09016 | 2.149792 | 2.20108 | 2.260211 | 2.359016 | 2.468797 | 2.578577 | 2.677382 | 2.736513 | 2.787801 | 2.847433 | 2.900448 |
|  | 40 | 1.975497 | 2.028179 | 2.087437 | 2.138402 | 2.197161 | 2.295346 | 2.404436 | 2.513526 | 2.61171 | 2.67047 | 2.721435 | 2.780693 | 2.833374 |
|  | 50 | 1.915293 | 1.967645 | 2.026532 | 2.077179 | 2.135571 | 2.233141 | 2.341549 | 2.449957 | 2.547528 | 2.60592 | 2.656566 | 2.715454 | 2.767806 |
|  | 60 | 1.838157 | 1.890182 | 1.948702 | 1.999033 | 2.057061 | 2.154022 | 2.261754 | 2.369486 | 2.466448 | 2.524476 | 2.574806 | 2.633326 | 2.685352 |
|  | 70 | 1.815875 | 1.867576 | 1.925731 | 1.975747 | 2.033413 | 2.12977 | 2.23683 | 2.343889 | 2.440246 | 2.497912 | 2.547928 | 2.606083 | 2.657784 |
|  | 80 | 1.794852 | 1.84623 | 1.904022 | 1.953726 | 2.011032 | 2.106787 | 2.213178 | 2.31957 | 2.415325 | 2.472631 | 2.522335 | 2.580127 | 2.631505 |
|  | 90 | 1.724557 | 1.775614 | 1.833045 | 1.882439 | 1.939387 | 2.034545 | 2.140271 | 2.245998 | 2.341156 | 2.398103 | 2.447497 | 2.504928 | 2.555985 |
| **L_pericalcarine** | 3 | 1.564779 | 1.592008 | 1.624482 | 1.654172 | 1.690705 | 1.758266 | 1.845425 | 1.949853 | 2.064645 | 2.146353 | 2.227693 | 2.338167 | 2.45542 |
|  | 5 | 1.532409 | 1.55991 | 1.592618 | 1.622436 | 1.65901 | 1.72631 | 1.81246 | 1.914631 | 2.025523 | 2.103482 | 2.180234 | 2.283032 | 2.390205 |
|  | 10 | 1.456342 | 1.484721 | 1.518208 | 1.54849 | 1.585314 | 1.65217 | 1.736042 | 1.833007 | 1.935114 | 2.004902 | 2.071969 | 2.159202 | 2.246948 |
|  | 15 | 1.389975 | 1.419592 | 1.454217 | 1.485235 | 1.522591 | 1.589422 | 1.671521 | 1.7641 | 1.858944 | 1.922225 | 1.981867 | 2.05773 | 2.132091 |
|  | 20 | 1.338577 | 1.369775 | 1.405881 | 1.437902 | 1.476078 | 1.543378 | 1.624423 | 1.713824 | 1.803381 | 1.862053 | 1.916585 | 1.984903 | 2.050756 |
|  | 30 | 1.282665 | 1.316749 | 1.355525 | 1.389362 | 1.429087 | 1.497672 | 1.578213 | 1.664985 | 1.750201 | 1.805282 | 1.85603 | 1.919078 | 1.979361 |
|  | 40 | 1.252349 | 1.287527 | 1.327198 | 1.361548 | 1.401606 | 1.470244 | 1.550328 | 1.636433 | 1.721243 | 1.776359 | 1.827427 | 1.891336 | 1.952993 |
|  | 50 | 1.232277 | 1.26717 | 1.306454 | 1.340433 | 1.38004 | 1.447938 | 1.527383 | 1.613317 | 1.698718 | 1.754719 | 1.807007 | 1.873029 | 1.937389 |
|  | 60 | 1.216224 | 1.250091 | 1.288317 | 1.321469 | 1.360224 | 1.426973 | 1.505633 | 1.591488 | 1.67769 | 1.734731 | 1.788376 | 1.856669 | 1.923858 |
|  | 70 | 1.207462 | 1.240211 | 1.277339 | 1.309681 | 1.347656 | 1.413473 | 1.491688 | 1.577838 | 1.665133 | 1.723327 | 1.778369 | 1.848877 | 1.918727 |
|  | 80 | 1.2009 | 1.231671 | 1.266821 | 1.297673 | 1.334179 | 1.398186 | 1.475489 | 1.562235 | 1.651908 | 1.712716 | 1.771015 | 1.84685 | 1.923308 |
|  | 90 | 1.191837 | 1.220068 | 1.252613 | 1.281448 | 1.315908 | 1.377256 | 1.453031 | 1.540435 | 1.633678 | 1.698731 | 1.762595 | 1.848032 | 1.937105 |
| **L_postcentral** | 3 | 2.070135 | 2.113243 | 2.161507 | 2.202733 | 2.249812 | 2.326983 | 2.409996 | 2.494541 | 2.575987 | 2.627089 | 2.672752 | 2.727331 | 2.777137 |
|  | 5 | 2.030671 | 2.073237 | 2.120895 | 2.161606 | 2.2081 | 2.284322 | 2.366333 | 2.449844 | 2.530249 | 2.580677 | 2.625727 | 2.67956 | 2.728672 |
|  | 10 | 1.935252 | 1.976496 | 2.022678 | 2.062131 | 2.107195 | 2.181097 | 2.260659 | 2.341642 | 2.4195 | 2.468283 | 2.511834 | 2.563844 | 2.611265 |
|  | 15 | 1.852404 | 1.892548 | 1.937499 | 1.975905 | 2.019779 | 2.091751 | 2.169279 | 2.248158 | 2.323884 | 2.371283 | 2.41357 | 2.464039 | 2.510028 |
|  | 20 | 1.793998 | 1.833536 | 1.87781 | 1.915641 | 1.958864 | 2.029789 | 2.10623 | 2.183967 | 2.25849 | 2.305087 | 2.346631 | 2.396183 | 2.441307 |
|  | 30 | 1.735489 | 1.775062 | 1.819378 | 1.857249 | 1.900528 | 1.971584 | 2.048246 | 2.126133 | 2.200578 | 2.247029 | 2.288388 | 2.337654 | 2.382463 |
|  | 40 | 1.69114 | 1.73106 | 1.775764 | 1.81397 | 1.857641 | 1.929374 | 2.006841 | 2.085463 | 2.160385 | 2.207034 | 2.248511 | 2.297854 | 2.342674 |
|  | 50 | 1.653191 | 1.693613 | 1.738878 | 1.777564 | 1.82179 | 1.894467 | 1.973022 | 2.052657 | 2.128312 | 2.175314 | 2.217047 | 2.266626 | 2.311603 |
|  | 60 | 1.606336 | 1.647045 | 1.692627 | 1.731583 | 1.776122 | 1.849338 | 1.928539 | 2.008733 | 2.084678 | 2.131756 | 2.173495 | 2.223013 | 2.267875 |
|  | 70 | 1.567753 | 1.608959 | 1.655089 | 1.694512 | 1.739586 | 1.813704 | 1.893936 | 1.975068 | 2.051653 | 2.099021 | 2.140954 | 2.190632 | 2.235577 |
|  | 80 | 1.51215 | 1.553397 | 1.599563 | 1.639011 | 1.684112 | 1.758292 | 1.838638 | 1.919773 | 1.996109 | 2.043212 | 2.084847 | 2.134102 | 2.178603 |
|  | 90 | 1.443814 | 1.484715 | 1.530479 | 1.569576 | 1.614274 | 1.687798 | 1.767473 | 1.847812 | 1.923144 | 1.969515 | 2.010441 | 2.058786 | 2.102401 |
| **L_posteriorcingulate** | 3 | 2.541025 | 2.62215 | 2.704361 | 2.769085 | 2.838565 | 2.946727 | 3.062526 | 3.182567 | 3.302917 | 3.384829 | 3.464645 | 3.571281 | 3.682749 |
|  | 5 | 2.519775 | 2.592573 | 2.668019 | 2.728605 | 2.794747 | 2.899614 | 3.013398 | 3.131277 | 3.247686 | 3.325305 | 3.399482 | 3.496252 | 3.594608 |
|  | 10 | 2.445044 | 2.504924 | 2.569484 | 2.623148 | 2.683452 | 2.782089 | 2.891548 | 3.004792 | 3.113762 | 3.183905 | 3.248734 | 3.329922 | 3.40854 |
|  | 15 | 2.355669 | 2.408529 | 2.466874 | 2.516359 | 2.572909 | 2.667067 | 2.772882 | 2.882225 | 2.985844 | 3.051179 | 3.110392 | 3.182795 | 3.250945 |
|  | 20 | 2.272924 | 2.321823 | 2.376541 | 2.423494 | 2.477669 | 2.568792 | 2.671908 | 2.778346 | 2.878279 | 2.940519 | 2.996276 | 3.063495 | 3.125712 |
|  | 30 | 2.169296 | 2.215413 | 2.267614 | 2.312842 | 2.365438 | 2.454613 | 2.556022 | 2.660475 | 2.75765 | 2.817476 | 2.870502 | 2.933606 | 2.991135 |
|  | 40 | 2.10508 | 2.151099 | 2.203329 | 2.248679 | 2.301499 | 2.391172 | 2.493147 | 2.597953 | 2.695046 | 2.754548 | 2.807077 | 2.8693 | 2.925722 |
|  | 50 | 2.048724 | 2.095468 | 2.148493 | 2.194505 | 2.248059 | 2.338873 | 2.441955 | 2.547653 | 2.64532 | 2.705042 | 2.757677 | 2.819915 | 2.87624 |
|  | 60 | 2.001018 | 2.049208 | 2.103715 | 2.150889 | 2.205664 | 2.29827 | 2.403047 | 2.510217 | 2.609101 | 2.669537 | 2.722798 | 2.78579 | 2.842826 |
|  | 70 | 1.965818 | 2.016882 | 2.074205 | 2.12349 | 2.180394 | 2.275971 | 2.383478 | 2.493153 | 2.594441 | 2.656524 | 2.711415 | 2.77661 | 2.835953 |
|  | 80 | 1.925399 | 1.982966 | 2.046207 | 2.09958 | 2.160249 | 2.260421 | 2.371576 | 2.484661 | 2.589991 | 2.655434 | 2.714075 | 2.784895 | 2.850663 |
|  | 90 | 1.839063 | 1.918699 | 2.000306 | 2.065162 | 2.135271 | 2.244915 | 2.361709 | 2.480201 | 2.594625 | 2.669474 | 2.739927 | 2.830288 | 2.920376 |
| **L_precentral** | 3 | 2.198532 | 2.303087 | 2.398892 | 2.468526 | 2.538389 | 2.637879 | 2.732009 | 2.81579 | 2.886209 | 2.927187 | 2.962452 | 3.003502 | 3.04034 |
|  | 5 | 2.206545 | 2.302948 | 2.393086 | 2.459623 | 2.527174 | 2.624613 | 2.71799 | 2.801951 | 2.873072 | 2.914668 | 2.950583 | 2.992516 | 3.030258 |
|  | 10 | 2.218714 | 2.298508 | 2.376562 | 2.436214 | 2.498441 | 2.590963 | 2.682445 | 2.766851 | 2.839812 | 2.883066 | 2.92074 | 2.965098 | 3.005351 |
|  | 15 | 2.218485 | 2.286267 | 2.354907 | 2.408848 | 2.466426 | 2.554381 | 2.643946 | 2.728729 | 2.803575 | 2.848597 | 2.88819 | 2.935247 | 2.978349 |
|  | 20 | 2.201656 | 2.262084 | 2.324609 | 2.374649 | 2.428914 | 2.513436 | 2.601437 | 2.686446 | 2.762804 | 2.809305 | 2.850543 | 2.899961 | 2.945606 |
|  | 30 | 2.120634 | 2.181167 | 2.243819 | 2.293975 | 2.348377 | 2.433137 | 2.521408 | 2.606686 | 2.683285 | 2.729926 | 2.771283 | 2.820837 | 2.866598 |
|  | 40 | 2.022023 | 2.091449 | 2.161693 | 2.216859 | 2.275718 | 2.365582 | 2.45703 | 2.543525 | 2.61981 | 2.665657 | 2.705949 | 2.753801 | 2.797595 |
|  | 50 | 1.957984 | 2.0286 | 2.10028 | 2.156729 | 2.217101 | 2.309545 | 2.403926 | 2.493453 | 2.572598 | 2.62024 | 2.662153 | 2.711979 | 2.757625 |
|  | 60 | 1.909716 | 1.977268 | 2.046826 | 2.102268 | 2.162182 | 2.255095 | 2.351329 | 2.443805 | 2.526457 | 2.576595 | 2.620932 | 2.673908 | 2.722689 |
|  | 70 | 1.856455 | 1.921392 | 1.988882 | 2.043109 | 2.102121 | 2.19444 | 2.291023 | 2.384693 | 2.469074 | 2.520546 | 2.566235 | 2.62103 | 2.671674 |
|  | 80 | 1.773937 | 1.842201 | 1.912394 | 1.968282 | 2.02862 | 2.122082 | 2.218745 | 2.311491 | 2.394257 | 2.4444 | 2.4887 | 2.541578 | 2.590215 |
|  | 90 | 1.66381 | 1.74356 | 1.822739 | 1.884002 | 1.948584 | 2.045809 | 2.143241 | 2.234142 | 2.313376 | 2.360594 | 2.401849 | 2.450559 | 2.494876 |
| **L_precuneus** | 3 | 2.505666 | 2.564177 | 2.627739 | 2.680501 | 2.739128 | 2.831718 | 2.92682 | 3.019667 | 3.105921 | 3.158537 | 3.20462 | 3.258592 | 3.306833 |
|  | 5 | 2.448954 | 2.506281 | 2.568607 | 2.620385 | 2.677966 | 2.769012 | 2.86268 | 2.95418 | 3.039139 | 3.090951 | 3.136324 | 3.189462 | 3.236954 |
|  | 10 | 2.309305 | 2.363687 | 2.422934 | 2.472253 | 2.527212 | 2.614375 | 2.704419 | 2.792507 | 2.874195 | 2.923981 | 2.967567 | 3.018604 | 3.064216 |
|  | 15 | 2.177297 | 2.228868 | 2.285169 | 2.332132 | 2.384574 | 2.468001 | 2.554545 | 2.639336 | 2.717876 | 2.765713 | 2.807583 | 2.856604 | 2.900412 |
|  | 20 | 2.080716 | 2.130276 | 2.184494 | 2.229814 | 2.280528 | 2.361454 | 2.445761 | 2.528489 | 2.605034 | 2.651629 | 2.692403 | 2.740135 | 2.782791 |
|  | 30 | 1.996213 | 2.044265 | 2.097058 | 2.141373 | 2.191174 | 2.271147 | 2.35518 | 2.437916 | 2.514312 | 2.56077 | 2.601409 | 2.648976 | 2.691487 |
|  | 40 | 1.94559 | 1.992883 | 2.045066 | 2.089059 | 2.138713 | 2.218968 | 2.304045 | 2.388112 | 2.465601 | 2.512685 | 2.55386 | 2.602055 | 2.645133 |
|  | 50 | 1.903027 | 1.949701 | 2.001427 | 2.045225 | 2.094882 | 2.175674 | 2.2621 | 2.347832 | 2.426741 | 2.474656 | 2.516554 | 2.565598 | 2.609449 |
|  | 60 | 1.845925 | 1.891566 | 1.942373 | 1.985585 | 2.0348 | 2.115421 | 2.202467 | 2.289178 | 2.368896 | 2.41728 | 2.45959 | 2.509127 | 2.553437 |
|  | 70 | 1.812776 | 1.857924 | 1.908408 | 1.951539 | 2.000891 | 2.082302 | 2.17104 | 2.259837 | 2.341406 | 2.390904 | 2.434193 | 2.484897 | 2.530275 |
|  | 80 | 1.749457 | 1.793308 | 1.842564 | 1.88484 | 1.933443 | 2.014192 | 2.103071 | 2.192443 | 2.274501 | 2.324295 | 2.367858 | 2.41891 | 2.464632 |
|  | 90 | 1.668069 | 1.710113 | 1.757557 | 1.798467 | 1.845727 | 1.924818 | 2.012751 | 2.101632 | 2.183231 | 2.23276 | 2.276111 | 2.326951 | 2.372523 |
| **L_rostralanteriorcingulate** | 3 | 2.469105 | 2.651175 | 2.824461 | 2.954167 | 3.087055 | 3.279321 | 3.461051 | 3.621565 | 3.754517 | 3.829673 | 3.892448 | 3.962838 | 4.023236 |
|  | 5 | 2.464805 | 2.624937 | 2.781824 | 2.901881 | 3.026969 | 3.211213 | 3.388373 | 3.547109 | 3.680157 | 3.755938 | 3.819529 | 3.891133 | 3.952813 |
|  | 10 | 2.432433 | 2.552633 | 2.677672 | 2.777909 | 2.886248 | 3.052463 | 3.219004 | 3.373715 | 3.507472 | 3.585221 | 3.651296 | 3.726572 | 3.792132 |
|  | 15 | 2.392559 | 2.488462 | 2.59225 | 2.678277 | 2.773951 | 2.925865 | 3.083891 | 3.236004 | 3.371816 | 3.452521 | 3.522091 | 3.602417 | 3.673282 |
|  | 20 | 2.361756 | 2.443816 | 2.534781 | 2.61183 | 2.69922 | 2.841514 | 2.993907 | 3.145127 | 3.284231 | 3.368672 | 3.442499 | 3.528915 | 3.606177 |
|  | 30 | 2.293307 | 2.367617 | 2.451106 | 2.522693 | 2.604796 | 2.740347 | 2.887771 | 3.037757 | 3.180348 | 3.269 | 3.347763 | 3.4414 | 3.526413 |
|  | 40 | 2.216694 | 2.293446 | 2.379236 | 2.452398 | 2.535817 | 2.672269 | 2.818683 | 2.967076 | 3.109035 | 3.197629 | 3.276498 | 3.370408 | 3.455773 |
|  | 50 | 2.175239 | 2.251957 | 2.337828 | 2.411159 | 2.494884 | 2.632109 | 2.779741 | 2.929614 | 3.07307 | 3.162645 | 3.242421 | 3.337458 | 3.423891 |
|  | 60 | 2.174039 | 2.244598 | 2.324739 | 2.394214 | 2.474796 | 2.610094 | 2.760668 | 2.916082 | 3.064511 | 3.157198 | 3.239851 | 3.338525 | 3.428522 |
|  | 70 | 2.195131 | 2.258141 | 2.331048 | 2.395521 | 2.471948 | 2.604853 | 2.760511 | 2.924683 | 3.079769 | 3.176036 | 3.261699 | 3.3639 | 3.457164 |
|  | 80 | 2.229044 | 2.282973 | 2.346768 | 2.404567 | 2.474966 | 2.603023 | 2.76331 | 2.938675 | 3.104403 | 3.207504 | 3.299626 | 3.410203 | 3.511907 |
|  | 90 | 2.257555 | 2.304269 | 2.360569 | 2.412666 | 2.477695 | 2.60112 | 2.765887 | 2.953247 | 3.131109 | 3.242254 | 3.342149 | 3.463041 | 3.5754 |
| **L_rostralmiddlefrontal** | 3 | 2.470763 | 2.538584 | 2.611342 | 2.670889 | 2.735988 | 2.835978 | 2.934201 | 3.030742 | 3.125676 | 3.185742 | 3.23952 | 3.303756 | 3.362209 |
|  | 5 | 2.417797 | 2.482039 | 2.551373 | 2.608457 | 2.671244 | 2.768559 | 2.865397 | 2.960876 | 3.054158 | 3.112968 | 3.165525 | 3.228214 | 3.285198 |
|  | 10 | 2.291564 | 2.347867 | 2.409491 | 2.46095 | 2.518378 | 2.609365 | 2.702788 | 2.795659 | 2.885059 | 2.940958 | 2.990704 | 3.049851 | 3.103487 |
|  | 15 | 2.178781 | 2.228632 | 2.283872 | 2.330586 | 2.383409 | 2.4688 | 2.559025 | 2.649503 | 2.735601 | 2.789091 | 2.836544 | 2.892838 | 2.94381 |
|  | 20 | 2.103207 | 2.148618 | 2.199443 | 2.242871 | 2.292517 | 2.374153 | 2.462537 | 2.551872 | 2.636111 | 2.688174 | 2.734245 | 2.788804 | 2.838148 |
|  | 30 | 2.026595 | 2.068985 | 2.116806 | 2.158016 | 2.205572 | 2.284968 | 2.372867 | 2.462105 | 2.54508 | 2.5959 | 2.640639 | 2.693391 | 2.740923 |
|  | 40 | 1.97107 | 2.017973 | 2.07014 | 2.114425 | 2.164705 | 2.246494 | 2.333686 | 2.421372 | 2.50453 | 2.556094 | 2.601794 | 2.655974 | 2.705009 |
|  | 50 | 1.901696 | 1.955043 | 2.013225 | 2.061603 | 2.115317 | 2.199648 | 2.284933 | 2.369971 | 2.453575 | 2.506577 | 2.554148 | 2.611149 | 2.663215 |
|  | 60 | 1.849004 | 1.902787 | 1.961444 | 2.010211 | 2.064341 | 2.149258 | 2.234997 | 2.320588 | 2.405065 | 2.458763 | 2.50704 | 2.56498 | 2.617982 |
|  | 70 | 1.825475 | 1.872495 | 1.924992 | 1.969721 | 2.020682 | 2.103988 | 2.193347 | 2.283721 | 2.369844 | 2.423448 | 2.471085 | 2.527716 | 2.579114 |
|  | 80 | 1.825077 | 1.86459 | 1.909985 | 1.949866 | 1.996856 | 2.07795 | 2.172074 | 2.269439 | 2.35887 | 2.413241 | 2.460952 | 2.517102 | 2.567656 |
|  | 90 | 1.844939 | 1.878276 | 1.917623 | 1.953237 | 1.996645 | 2.075932 | 2.175813 | 2.282019 | 2.376411 | 2.432445 | 2.480977 | 2.537503 | 2.587974 |
| **L_superiorfrontal** | 3 | 2.795698 | 2.86471 | 2.938768 | 2.99959 | 3.06655 | 3.171146 | 3.277365 | 3.379337 | 3.472059 | 3.527775 | 3.576094 | 3.632159 | 3.681818 |
|  | 5 | 2.753038 | 2.820121 | 2.89234 | 2.951818 | 3.017461 | 3.120307 | 3.225087 | 3.326073 | 3.418306 | 3.473901 | 3.522214 | 3.578381 | 3.628225 |
|  | 10 | 2.647646 | 2.710126 | 2.777913 | 2.834125 | 2.896542 | 2.995067 | 3.096269 | 3.194789 | 3.285798 | 3.3411 | 3.389418 | 3.445878 | 3.496234 |
|  | 15 | 2.543933 | 2.602099 | 2.665671 | 2.718741 | 2.77802 | 2.872294 | 2.96994 | 3.065985 | 3.155767 | 3.210794 | 3.259146 | 3.315963 | 3.366914 |
|  | 20 | 2.45617 | 2.510835 | 2.570965 | 2.621454 | 2.678151 | 2.768927 | 2.863664 | 2.957764 | 3.046751 | 3.101754 | 3.150363 | 3.207803 | 3.259599 |
|  | 30 | 2.354229 | 2.405979 | 2.463259 | 2.511627 | 2.566218 | 2.654163 | 2.746546 | 2.839363 | 2.928537 | 2.984296 | 3.033962 | 3.093101 | 3.146837 |
|  | 40 | 2.289009 | 2.341082 | 2.398685 | 2.447287 | 2.502085 | 2.590177 | 2.682381 | 2.775148 | 2.864848 | 2.921181 | 2.971495 | 3.031559 | 3.086264 |
|  | 50 | 2.219424 | 2.27319 | 2.332386 | 2.382098 | 2.437879 | 2.526919 | 2.619211 | 2.711636 | 2.801061 | 2.857211 | 2.907335 | 2.967127 | 3.021532 |
|  | 60 | 2.156339 | 2.21313 | 2.275166 | 2.326865 | 2.384442 | 2.47538 | 2.568337 | 2.660635 | 2.749666 | 2.8054 | 2.855028 | 2.914063 | 2.967618 |
|  | 70 | 2.100126 | 2.160658 | 2.226174 | 2.280299 | 2.340075 | 2.433399 | 2.527392 | 2.61986 | 2.708746 | 2.764204 | 2.813451 | 2.871861 | 2.924679 |
|  | 80 | 2.020444 | 2.08464 | 2.153379 | 2.209604 | 2.271122 | 2.365965 | 2.459996 | 2.551602 | 2.639319 | 2.693851 | 2.742139 | 2.799234 | 2.850696 |
|  | 90 | 1.955067 | 2.024118 | 2.097127 | 2.156167 | 2.220094 | 2.31731 | 2.412101 | 2.5035 | 2.590654 | 2.64463 | 2.692285 | 2.748454 | 2.798916 |
| **L_superiorparietal** | 3 | 2.256538 | 2.302051 | 2.353067 | 2.396747 | 2.44682 | 2.529596 | 2.620009 | 2.71104 | 2.795482 | 2.847084 | 2.892423 | 2.945765 | 2.993712 |
|  | 5 | 2.206435 | 2.25134 | 2.301658 | 2.344727 | 2.394082 | 2.475626 | 2.564627 | 2.654209 | 2.737316 | 2.788105 | 2.832731 | 2.885234 | 2.932424 |
|  | 10 | 2.083355 | 2.126724 | 2.175281 | 2.216808 | 2.264355 | 2.342805 | 2.428269 | 2.514219 | 2.593983 | 2.642736 | 2.685574 | 2.735972 | 2.781268 |
|  | 15 | 1.967046 | 2.008947 | 2.055821 | 2.095872 | 2.141687 | 2.217173 | 2.299245 | 2.381711 | 2.45826 | 2.505052 | 2.546166 | 2.594534 | 2.638 |
|  | 20 | 1.881428 | 1.922632 | 1.968655 | 2.00792 | 2.052767 | 2.126494 | 2.206413 | 2.28656 | 2.360903 | 2.406317 | 2.4462 | 2.493091 | 2.535205 |
|  | 30 | 1.809269 | 1.852421 | 1.900269 | 1.940803 | 1.98678 | 2.061635 | 2.141774 | 2.221323 | 2.294575 | 2.33907 | 2.377987 | 2.423555 | 2.464306 |
|  | 40 | 1.760795 | 1.806404 | 1.856616 | 1.898863 | 1.946471 | 2.023281 | 2.104579 | 2.184555 | 2.257757 | 2.302017 | 2.340601 | 2.385627 | 2.425757 |
|  | 50 | 1.7302 | 1.776539 | 1.827559 | 1.870488 | 1.918857 | 1.996862 | 2.079349 | 2.160596 | 2.235208 | 2.280422 | 2.319894 | 2.366015 | 2.407172 |
|  | 60 | 1.701338 | 1.746268 | 1.796089 | 1.838284 | 1.886114 | 1.963856 | 2.046818 | 2.129441 | 2.206251 | 2.253214 | 2.29446 | 2.342939 | 2.38645 |
|  | 70 | 1.686807 | 1.729657 | 1.777661 | 1.818709 | 1.865666 | 1.942925 | 2.026602 | 2.111365 | 2.191605 | 2.241334 | 2.285418 | 2.337715 | 2.385092 |
|  | 80 | 1.634761 | 1.674283 | 1.719046 | 1.75773 | 1.80244 | 1.877054 | 1.959322 | 2.0444 | 2.126753 | 2.178676 | 2.225269 | 2.281228 | 2.332564 |
|  | 90 | 1.54307 | 1.578624 | 1.6193 | 1.654803 | 1.696243 | 1.766376 | 1.845125 | 1.92837 | 2.010935 | 2.064011 | 2.112319 | 2.171195 | 2.226043 |
| **L_superiortemporal** | 3 | 2.563263 | 2.648741 | 2.737069 | 2.807047 | 2.881414 | 2.991776 | 3.096191 | 3.195266 | 3.289486 | 3.347621 | 3.398774 | 3.458834 | 3.51256 |
|  | 5 | 2.545344 | 2.628722 | 2.71526 | 2.784091 | 2.857505 | 2.966984 | 3.071215 | 3.170329 | 3.264426 | 3.322441 | 3.373474 | 3.433386 | 3.486981 |
|  | 10 | 2.501089 | 2.579474 | 2.661701 | 2.727738 | 2.798808 | 2.906089 | 3.009847 | 3.109067 | 3.202885 | 3.260627 | 3.311389 | 3.37097 | 3.424274 |
|  | 15 | 2.459092 | 2.532777 | 2.610882 | 2.674213 | 2.742988 | 2.848099 | 2.951409 | 3.050804 | 3.144461 | 3.202022 | 3.252604 | 3.311975 | 3.365106 |
|  | 20 | 2.421869 | 2.491229 | 2.565495 | 2.626284 | 2.692897 | 2.795988 | 2.898987 | 2.998738 | 3.092455 | 3.149988 | 3.200537 | 3.259881 | 3.313014 |
|  | 30 | 2.360995 | 2.423924 | 2.492403 | 2.549336 | 2.612691 | 2.712931 | 2.816101 | 2.91712 | 3.011306 | 3.068913 | 3.119456 | 3.178757 | 3.231851 |
|  | 40 | 2.309463 | 2.368511 | 2.433461 | 2.48805 | 2.54948 | 2.64834 | 2.752546 | 2.855196 | 2.949783 | 3.007228 | 3.057441 | 3.116181 | 3.168651 |
|  | 50 | 2.256788 | 2.31326 | 2.375844 | 2.428859 | 2.489027 | 2.587177 | 2.692677 | 2.796845 | 2.891417 | 2.948306 | 2.997761 | 3.055344 | 3.10657 |
|  | 60 | 2.195427 | 2.250096 | 2.310984 | 2.362851 | 2.422088 | 2.51975 | 2.626392 | 2.731577 | 2.825411 | 2.881196 | 2.929353 | 2.98508 | 3.034383 |
|  | 70 | 2.12502 | 2.178512 | 2.238252 | 2.289317 | 2.347889 | 2.445219 | 2.55278 | 2.658408 | 2.750757 | 2.804907 | 2.851266 | 2.904515 | 2.951307 |
|  | 80 | 2.043495 | 2.096122 | 2.154962 | 2.20535 | 2.263308 | 2.360173 | 2.468194 | 2.573536 | 2.663637 | 2.715676 | 2.759822 | 2.810116 | 2.853982 |
|  | 90 | 1.959613 | 2.011614 | 2.06976 | 2.119602 | 2.17704 | 2.273475 | 2.38179 | 2.486532 | 2.574114 | 2.623912 | 2.665761 | 2.713041 | 2.753965 |
| **L_supramarginal** | 3 | 2.487489 | 2.584098 | 2.680049 | 2.753595 | 2.829579 | 2.938497 | 3.03748 | 3.129406 | 3.216423 | 3.269826 | 3.316601 | 3.371245 | 3.419866 |
|  | 5 | 2.452703 | 2.541985 | 2.631992 | 2.701862 | 2.77486 | 2.881033 | 2.979331 | 3.071059 | 3.157211 | 3.20987 | 3.255904 | 3.309609 | 3.357347 |
|  | 10 | 2.366053 | 2.440025 | 2.517092 | 2.578682 | 2.644757 | 2.744341 | 2.840868 | 2.932103 | 3.016256 | 3.067207 | 3.111546 | 3.163106 | 3.208835 |
|  | 15 | 2.288052 | 2.350248 | 2.41687 | 2.471499 | 2.531559 | 2.625218 | 2.720154 | 2.811167 | 2.893848 | 2.9435 | 2.986542 | 3.036465 | 3.080668 |
|  | 20 | 2.230731 | 2.284519 | 2.343427 | 2.392774 | 2.448178 | 2.537229 | 2.631183 | 2.722608 | 2.804818 | 2.853914 | 2.896376 | 2.94556 | 2.989086 |
|  | 30 | 2.165817 | 2.211859 | 2.263488 | 2.307758 | 2.358646 | 2.443324 | 2.536901 | 2.630018 | 2.713492 | 2.7633 | 2.806414 | 2.85644 | 2.900823 |
|  | 40 | 2.107718 | 2.153234 | 2.204383 | 2.248313 | 2.298864 | 2.383013 | 2.475934 | 2.569192 | 2.654229 | 2.705584 | 2.750379 | 2.802732 | 2.8495 |
|  | 50 | 2.045444 | 2.093629 | 2.147351 | 2.193075 | 2.245143 | 2.330265 | 2.421704 | 2.513672 | 2.600248 | 2.653671 | 2.700881 | 2.756706 | 2.807111 |
|  | 60 | 1.982198 | 2.033437 | 2.090068 | 2.137814 | 2.191613 | 2.278043 | 2.368466 | 2.45925 | 2.54671 | 2.601491 | 2.650331 | 2.708533 | 2.761453 |
|  | 70 | 1.940339 | 1.992444 | 2.049994 | 2.098501 | 2.153165 | 2.241078 | 2.333298 | 2.42552 | 2.513439 | 2.568109 | 2.616623 | 2.674182 | 2.726296 |
|  | 80 | 1.877377 | 1.92877 | 1.985659 | 2.033755 | 2.08818 | 2.176417 | 2.270259 | 2.363551 | 2.450275 | 2.503292 | 2.549843 | 2.604542 | 2.653619 |
|  | 90 | 1.802345 | 1.852989 | 1.909117 | 1.956667 | 2.010635 | 2.098675 | 2.193311 | 2.286799 | 2.371754 | 2.422893 | 2.467369 | 2.519175 | 2.565283 |
| **L_frontalpole** | 3 | 2.416308 | 2.529325 | 2.656072 | 2.764455 | 2.888286 | 3.091199 | 3.309054 | 3.531394 | 3.746902 | 3.882647 | 4.004222 | 4.14982 | 4.282907 |
|  | 5 | 2.368392 | 2.478465 | 2.602143 | 2.708116 | 2.829466 | 3.029029 | 3.24444 | 3.464515 | 3.6771 | 3.810713 | 3.930229 | 4.073207 | 4.203777 |
|  | 10 | 2.257529 | 2.360758 | 2.477296 | 2.577659 | 2.693227 | 2.885014 | 3.094836 | 3.309776 | 3.515644 | 3.644332 | 3.75908 | 3.895987 | 4.020717 |
|  | 15 | 2.168054 | 2.265501 | 2.376035 | 2.47172 | 2.582531 | 2.768142 | 2.974026 | 3.18552 | 3.386339 | 3.511164 | 3.622106 | 3.754098 | 3.874056 |
|  | 20 | 2.102794 | 2.195558 | 2.301277 | 2.393265 | 2.500403 | 2.681548 | 2.885275 | 3.095153 | 3.292731 | 3.414845 | 3.523019 | 3.651354 | 3.767698 |
|  | 30 | 2.021824 | 2.109652 | 2.210221 | 2.298159 | 2.40112 | 2.576642 | 2.776356 | 2.983197 | 3.177592 | 3.297638 | 3.403959 | 3.530107 | 3.644506 |
|  | 40 | 1.971563 | 2.061442 | 2.163862 | 2.252875 | 2.356296 | 2.530099 | 2.723349 | 2.924342 | 3.119406 | 3.242598 | 3.353258 | 3.48628 | 3.608406 |
|  | 50 | 1.923217 | 2.017372 | 2.123882 | 2.215611 | 2.320988 | 2.494435 | 2.680753 | 2.875275 | 3.072435 | 3.20071 | 3.318126 | 3.46177 | 3.595861 |
|  | 60 | 1.872846 | 1.964911 | 2.069468 | 2.159869 | 2.264128 | 2.436714 | 2.623509 | 2.819673 | 3.019271 | 3.149571 | 3.269154 | 3.415857 | 3.553213 |
|  | 70 | 1.837721 | 1.924341 | 2.023978 | 2.111358 | 2.213777 | 2.387981 | 2.584599 | 2.792856 | 2.999131 | 3.131462 | 3.251657 | 3.397772 | 3.533463 |
|  | 80 | 1.811164 | 1.893096 | 1.988709 | 2.073934 | 2.17571 | 2.354417 | 2.566124 | 2.792762 | 3.010549 | 3.147404 | 3.270177 | 3.417799 | 3.55354 |
|  | 90 | 1.786918 | 1.864365 | 1.956082 | 2.039214 | 2.140445 | 2.324266 | 2.553236 | 2.801397 | 3.032926 | 3.175316 | 3.301408 | 3.451291 | 3.587692 |
| **L_temporalpole** | 3 | 2.247413 | 2.487923 | 2.71777 | 2.891565 | 3.072836 | 3.345378 | 3.618902 | 3.852735 | 4.025367 | 4.116052 | 4.188788 | 4.267637 | 4.333363 |
|  | 5 | 2.290637 | 2.515635 | 2.734054 | 2.901402 | 3.077725 | 3.345588 | 3.617007 | 3.851414 | 4.026264 | 4.118699 | 4.193105 | 4.274014 | 4.341643 |
|  | 10 | 2.38853 | 2.579379 | 2.772508 | 2.925277 | 3.090138 | 3.346819 | 3.612985 | 3.848783 | 4.029392 | 4.126459 | 4.205333 | 4.291803 | 4.364609 |
|  | 15 | 2.473655 | 2.637772 | 2.810077 | 2.950225 | 3.104723 | 3.350775 | 3.611632 | 3.848801 | 4.035593 | 4.137769 | 4.221652 | 4.314444 | 4.39321 |
|  | 20 | 2.544717 | 2.6899 | 2.846385 | 2.976332 | 3.121974 | 3.358182 | 3.613286 | 3.850909 | 4.0433 | 4.150438 | 4.239332 | 4.338596 | 4.423582 |
|  | 30 | 2.611111 | 2.744532 | 2.89018 | 3.012281 | 3.150043 | 3.374644 | 3.617941 | 3.849277 | 4.043349 | 4.154014 | 4.247137 | 4.352426 | 4.443595 |
|  | 40 | 2.587584 | 2.731733 | 2.886062 | 3.013118 | 3.153987 | 3.378018 | 3.613137 | 3.834561 | 4.022584 | 4.130675 | 4.222027 | 4.325667 | 4.415652 |
|  | 50 | 2.485651 | 2.653336 | 2.827156 | 2.966318 | 3.116788 | 3.348304 | 3.58173 | 3.798133 | 3.983345 | 4.09035 | 4.180989 | 4.283974 | 4.373471 |
|  | 60 | 2.340015 | 2.534978 | 2.731296 | 2.88472 | 3.047163 | 3.290394 | 3.527652 | 3.745734 | 3.935054 | 4.045401 | 4.139314 | 4.246421 | 4.339788 |
|  | 70 | 2.229085 | 2.432516 | 2.637242 | 2.796932 | 2.965454 | 3.215986 | 3.457231 | 3.680861 | 3.880546 | 3.999038 | 4.100947 | 4.218251 | 4.321355 |
|  | 80 | 2.16699 | 2.362192 | 2.562499 | 2.720899 | 2.889585 | 3.141997 | 3.385141 | 3.615028 | 3.828298 | 3.958031 | 4.071311 | 4.203501 | 4.321167 |
|  | 90 | 2.128324 | 2.309564 | 2.500258 | 2.653894 | 2.819722 | 3.070802 | 3.314009 | 3.54977 | 3.778351 | 3.921554 | 4.04894 | 4.200171 | 4.336986 |
| **L_transversetemporal** | 3 | 2.123358 | 2.194756 | 2.275982 | 2.346606 | 2.4289 | 2.568455 | 2.726463 | 2.887515 | 3.03502 | 3.124436 | 3.202672 | 3.294429 | 3.37671 |
|  | 5 | 2.089309 | 2.160402 | 2.24123 | 2.311466 | 2.393258 | 2.531846 | 2.688583 | 2.848158 | 2.994153 | 3.082582 | 3.159913 | 3.250558 | 3.331796 |
|  | 10 | 2.007034 | 2.07742 | 2.157315 | 2.226632 | 2.307228 | 2.44349 | 2.597161 | 2.753155 | 2.895487 | 2.981523 | 3.056656 | 3.144603 | 3.223312 |
|  | 15 | 1.932632 | 2.002535 | 2.081747 | 2.150357 | 2.230002 | 2.364348 | 2.515413 | 2.668301 | 2.807412 | 2.891327 | 2.964506 | 3.050045 | 3.126492 |
|  | 20 | 1.871323 | 1.941186 | 2.020208 | 2.088534 | 2.167713 | 2.300957 | 2.450326 | 2.60103 | 2.737765 | 2.820076 | 2.891752 | 2.975417 | 3.050086 |
|  | 30 | 1.773955 | 1.844685 | 1.92437 | 1.993007 | 2.072255 | 2.204938 | 2.352724 | 2.500873 | 2.634506 | 2.714608 | 2.784158 | 2.865111 | 2.937155 |
|  | 40 | 1.69822 | 1.770796 | 1.852195 | 1.922009 | 2.00229 | 2.135958 | 2.283815 | 2.431031 | 2.563016 | 2.641785 | 2.709976 | 2.789119 | 2.859351 |
|  | 50 | 1.636113 | 1.711375 | 1.795352 | 1.867033 | 1.949089 | 2.084888 | 2.233992 | 2.381386 | 2.5127 | 2.590716 | 2.658053 | 2.735978 | 2.804931 |
|  | 60 | 1.573858 | 1.652159 | 1.739013 | 1.812749 | 1.896735 | 2.034808 | 2.185206 | 2.332756 | 2.463351 | 2.540583 | 2.60704 | 2.683721 | 2.751379 |
|  | 70 | 1.509755 | 1.591444 | 1.681439 | 1.757373 | 1.843378 | 1.983747 | 2.135338 | 2.282877 | 2.412578 | 2.48892 | 2.55441 | 2.629749 | 2.696034 |
|  | 80 | 1.432869 | 1.517738 | 1.610491 | 1.688204 | 1.775671 | 1.917289 | 2.068824 | 2.215073 | 2.34274 | 2.417522 | 2.481474 | 2.554825 | 2.619174 |
|  | 90 | 1.34312 | 1.430875 | 1.525884 | 1.604844 | 1.693086 | 1.834706 | 1.984743 | 2.128268 | 2.25265 | 2.325151 | 2.386955 | 2.457631 | 2.519456 |
| **L_insula** | 3 | 2.872207 | 2.993616 | 3.107227 | 3.190806 | 3.274875 | 3.393058 | 3.500212 | 3.594879 | 3.676362 | 3.723455 | 3.763249 | 3.808277 | 3.847196 |
|  | 5 | 2.856058 | 2.966239 | 3.071846 | 3.150957 | 3.231644 | 3.346796 | 3.4528 | 3.547374 | 3.629211 | 3.676684 | 3.716896 | 3.762502 | 3.802008 |
|  | 10 | 2.812298 | 2.900261 | 2.989097 | 3.058398 | 3.13139 | 3.239418 | 3.342719 | 3.437274 | 3.520305 | 3.568968 | 3.610472 | 3.657853 | 3.699161 |
|  | 15 | 2.7757 | 2.847475 | 2.923046 | 2.984085 | 3.050302 | 3.151866 | 3.252907 | 3.348089 | 3.433202 | 3.483739 | 3.527221 | 3.577288 | 3.621305 |
|  | 20 | 2.746198 | 2.806755 | 2.872506 | 2.927097 | 2.987816 | 3.083994 | 3.183381 | 3.27979 | 3.367726 | 3.420713 | 3.466767 | 3.520333 | 3.567901 |
|  | 30 | 2.676538 | 2.72908 | 2.787487 | 2.837083 | 2.893466 | 2.98554 | 3.084475 | 3.183056 | 3.274166 | 3.329659 | 3.378286 | 3.435327 | 3.486432 |
|  | 40 | 2.609363 | 2.6612 | 2.719046 | 2.768374 | 2.824716 | 2.917444 | 3.018241 | 3.11872 | 3.210593 | 3.266156 | 3.31464 | 3.371304 | 3.421905 |
|  | 50 | 2.546055 | 2.59858 | 2.657208 | 2.707239 | 2.764464 | 2.858945 | 2.962231 | 3.064641 | 3.156808 | 3.211941 | 3.259722 | 3.315215 | 3.364481 |
|  | 60 | 2.500547 | 2.555654 | 2.616929 | 2.66906 | 2.728551 | 2.826601 | 2.933717 | 3.038718 | 3.131255 | 3.185804 | 3.232645 | 3.286582 | 3.334079 |
|  | 70 | 2.459787 | 2.516498 | 2.579504 | 2.633099 | 2.694288 | 2.795341 | 2.906169 | 3.014084 | 3.107639 | 3.162169 | 3.208672 | 3.261885 | 3.308475 |
|  | 80 | 2.419088 | 2.474511 | 2.536502 | 2.589594 | 2.650647 | 2.752594 | 2.86609 | 2.977072 | 3.072638 | 3.128089 | 3.17526 | 3.229135 | 3.27623 |
|  | 90 | 2.358289 | 2.412558 | 2.473565 | 2.526096 | 2.586854 | 2.689244 | 2.80468 | 2.917796 | 3.014378 | 3.070091 | 3.117329 | 3.171129 | 3.218048 |
| **R_bankssts** | 3 | 2.486483 | 2.56228 | 2.645342 | 2.714815 | 2.792509 | 2.916096 | 3.043857 | 3.17061 | 3.291331 | 3.366286 | 3.432701 | 3.51136 | 3.582431 |
|  | 5 | 2.446965 | 2.521564 | 2.603384 | 2.671881 | 2.748561 | 2.870725 | 2.99731 | 3.122931 | 3.242357 | 3.316425 | 3.382011 | 3.459644 | 3.529756 |
|  | 10 | 2.349923 | 2.421578 | 2.500339 | 2.566427 | 2.640592 | 2.759218 | 2.882857 | 3.005637 | 3.121842 | 3.193709 | 3.257244 | 3.332345 | 3.400088 |
|  | 15 | 2.258417 | 2.327295 | 2.403168 | 2.466979 | 2.538765 | 2.654039 | 2.774886 | 2.894977 | 3.008134 | 3.077921 | 3.139518 | 3.212226 | 3.277731 |
|  | 20 | 2.183186 | 2.249812 | 2.323359 | 2.38535 | 2.455258 | 2.56795 | 2.686767 | 2.804914 | 2.915742 | 2.983898 | 3.043956 | 3.114747 | 3.178444 |
|  | 30 | 2.101469 | 2.166 | 2.23748 | 2.297955 | 2.366439 | 2.477601 | 2.596025 | 2.713814 | 2.823233 | 2.890095 | 2.948791 | 3.01775 | 3.079616 |
|  | 40 | 2.046454 | 2.110153 | 2.18088 | 2.240885 | 2.309057 | 2.420338 | 2.539928 | 2.658733 | 2.767889 | 2.834102 | 2.891972 | 2.95969 | 3.020223 |
|  | 50 | 1.986051 | 2.049035 | 2.119085 | 2.178641 | 2.246479 | 2.357743 | 2.478224 | 2.597646 | 2.706081 | 2.771333 | 2.828087 | 2.894208 | 2.953077 |
|  | 60 | 1.932495 | 1.995248 | 2.065112 | 2.124595 | 2.192484 | 2.304273 | 2.426101 | 2.546467 | 2.654393 | 2.718785 | 2.774498 | 2.839102 | 2.896368 |
|  | 70 | 1.891533 | 1.954724 | 2.0251 | 2.085069 | 2.153611 | 2.266832 | 2.390882 | 2.512934 | 2.620937 | 2.684793 | 2.739738 | 2.803133 | 2.85907 |
|  | 80 | 1.82995 | 1.893093 | 1.963401 | 2.023333 | 2.091899 | 2.205444 | 2.330406 | 2.452754 | 2.559548 | 2.622101 | 2.675617 | 2.737047 | 2.790992 |
|  | 90 | 1.757837 | 1.820644 | 1.89054 | 1.950124 | 2.018336 | 2.131538 | 2.256608 | 2.37841 | 2.483267 | 2.544107 | 2.595858 | 2.654955 | 2.706607 |
| **R_caudalanteriorcingulate** | 3 | 2.368118 | 2.461048 | 2.559026 | 2.638914 | 2.727296 | 2.869676 | 3.026668 | 3.19107 | 3.354196 | 3.463011 | 3.566865 | 3.702021 | 3.838908 |
|  | 5 | 2.322316 | 2.412467 | 2.507906 | 2.586004 | 2.67267 | 2.812746 | 2.967558 | 3.129635 | 3.289977 | 3.396508 | 3.497799 | 3.629008 | 3.761167 |
|  | 10 | 2.212462 | 2.296424 | 2.386152 | 2.460185 | 2.542913 | 2.677626 | 2.827302 | 2.983896 | 3.137755 | 3.239048 | 3.334527 | 3.456899 | 3.578611 |
|  | 15 | 2.117332 | 2.196326 | 2.281451 | 2.352199 | 2.431739 | 2.56211 | 2.707622 | 2.859755 | 3.008322 | 3.10534 | 3.196089 | 3.311311 | 3.424641 |
|  | 20 | 2.054494 | 2.130219 | 2.212425 | 2.281185 | 2.358905 | 2.487017 | 2.630569 | 2.780547 | 2.926212 | 3.02065 | 3.108386 | 3.218859 | 3.326451 |
|  | 30 | 1.990063 | 2.062571 | 2.142252 | 2.209606 | 2.286406 | 2.414174 | 2.558231 | 2.708515 | 2.853121 | 2.945737 | 3.030802 | 3.136429 | 3.237615 |
|  | 40 | 1.950382 | 2.021693 | 2.100804 | 2.168219 | 2.245602 | 2.375234 | 2.522044 | 2.674967 | 2.820994 | 2.913614 | 2.997913 | 3.101438 | 3.199328 |
|  | 50 | 1.913961 | 1.985091 | 2.064578 | 2.132732 | 2.211356 | 2.343745 | 2.494142 | 2.650549 | 2.798965 | 2.892359 | 2.976747 | 3.079475 | 3.175609 |
|  | 60 | 1.884697 | 1.95665 | 2.037502 | 2.107147 | 2.187793 | 2.324089 | 2.479232 | 2.640303 | 2.792327 | 2.887376 | 2.972752 | 3.075951 | 3.171728 |
|  | 70 | 1.875335 | 1.94951 | 2.033202 | 2.105539 | 2.189526 | 2.331831 | 2.493989 | 2.662035 | 2.819905 | 2.918075 | 3.005827 | 3.111286 | 3.208505 |
|  | 80 | 1.894318 | 1.972462 | 2.060893 | 2.137509 | 2.226627 | 2.377864 | 2.550253 | 2.728542 | 2.895334 | 2.998571 | 3.090477 | 3.2004 | 3.301173 |
|  | 90 | 1.938176 | 2.022029 | 2.11711 | 2.199617 | 2.295693 | 2.458869 | 2.644795 | 2.836659 | 3.015457 | 3.125675 | 3.223451 | 3.339923 | 3.446207 |
| **R_caudalmiddlefrontal** | 3 | 2.392723 | 2.455349 | 2.523488 | 2.580305 | 2.643928 | 2.746166 | 2.854449 | 2.957826 | 3.047153 | 3.099081 | 3.143256 | 3.193662 | 3.237654 |
|  | 5 | 2.370569 | 2.431933 | 2.498876 | 2.554826 | 2.617616 | 2.718798 | 2.826317 | 2.92928 | 3.018481 | 3.070429 | 3.114673 | 3.165215 | 3.209373 |
|  | 10 | 2.314877 | 2.373205 | 2.437238 | 2.491066 | 2.551794 | 2.650335 | 2.755916 | 2.85781 | 2.946672 | 2.998661 | 3.043076 | 3.09396 | 3.138542 |
|  | 15 | 2.257345 | 2.312817 | 2.374071 | 2.425841 | 2.484541 | 2.580423 | 2.683983 | 2.784693 | 2.873104 | 2.925072 | 2.969604 | 3.020776 | 3.06574 |
|  | 20 | 2.201679 | 2.254815 | 2.313762 | 2.363798 | 2.420763 | 2.51432 | 2.616044 | 2.715608 | 2.803508 | 2.855384 | 2.899957 | 2.951306 | 2.996542 |
|  | 30 | 2.114911 | 2.166399 | 2.223664 | 2.272389 | 2.327989 | 2.419586 | 2.519561 | 2.61778 | 2.704781 | 2.756249 | 2.800542 | 2.851648 | 2.896739 |
|  | 40 | 2.051886 | 2.10457 | 2.162935 | 2.212413 | 2.268678 | 2.36094 | 2.461067 | 2.558894 | 2.645129 | 2.695968 | 2.739619 | 2.789872 | 2.834113 |
|  | 50 | 1.992423 | 2.04714 | 2.107387 | 2.158172 | 2.215618 | 2.309161 | 2.409821 | 2.507377 | 2.592773 | 2.642868 | 2.68574 | 2.734942 | 2.778124 |
|  | 60 | 1.948612 | 2.005663 | 2.068117 | 2.120486 | 2.179436 | 2.274821 | 2.376694 | 2.47473 | 2.560029 | 2.609859 | 2.652386 | 2.701064 | 2.743678 |
|  | 70 | 1.91316 | 1.972672 | 2.03747 | 2.091541 | 2.15214 | 2.249641 | 2.353085 | 2.452027 | 2.537674 | 2.587529 | 2.629981 | 2.678466 | 2.72082 |
|  | 80 | 1.851477 | 1.913385 | 1.98029 | 2.035754 | 2.097548 | 2.196229 | 2.300023 | 2.398526 | 2.483238 | 2.532331 | 2.574014 | 2.621491 | 2.662856 |
|  | 90 | 1.770679 | 1.83553 | 1.904853 | 1.961777 | 2.024672 | 2.124073 | 2.227405 | 2.324448 | 2.407196 | 2.454875 | 2.495208 | 2.540989 | 2.580743 |
| **R_cuneus** | 3 | 1.931459 | 1.976865 | 2.028592 | 2.073635 | 2.126204 | 2.215567 | 2.317096 | 2.420985 | 2.51651 | 2.574595 | 2.625532 | 2.685408 | 2.739227 |
|  | 5 | 1.879102 | 1.92357 | 1.97427 | 2.018451 | 2.070053 | 2.157867 | 2.257778 | 2.360164 | 2.454439 | 2.511825 | 2.562186 | 2.621428 | 2.674718 |
|  | 10 | 1.75771 | 1.799987 | 1.848282 | 1.890451 | 1.939797 | 2.024006 | 2.120165 | 2.219087 | 2.310505 | 2.366306 | 2.41537 | 2.473199 | 2.525319 |
|  | 15 | 1.651694 | 1.692035 | 1.738211 | 1.77861 | 1.82598 | 1.90705 | 1.999977 | 2.095958 | 2.185001 | 2.23951 | 2.287536 | 2.344259 | 2.395488 |
|  | 20 | 1.572646 | 1.611565 | 1.656207 | 1.695341 | 1.741322 | 1.820246 | 1.911068 | 2.005266 | 2.093004 | 2.146878 | 2.194447 | 2.250751 | 2.301714 |
|  | 30 | 1.507019 | 1.544716 | 1.588122 | 1.626321 | 1.671379 | 1.749161 | 1.839359 | 1.933687 | 2.022254 | 2.076974 | 2.125501 | 2.183196 | 2.235657 |
|  | 40 | 1.478022 | 1.514449 | 1.556532 | 1.59369 | 1.637669 | 1.713971 | 1.803063 | 1.896939 | 1.985741 | 2.040926 | 2.090071 | 2.148751 | 2.202346 |
|  | 50 | 1.462266 | 1.497376 | 1.538055 | 1.574078 | 1.616843 | 1.691374 | 1.778946 | 1.871871 | 1.960395 | 2.015716 | 2.065183 | 2.124495 | 2.178904 |
|  | 60 | 1.444981 | 1.478758 | 1.517999 | 1.552844 | 1.59433 | 1.666948 | 1.752795 | 1.844521 | 1.932522 | 1.987828 | 2.037486 | 2.097286 | 2.152393 |
|  | 70 | 1.440189 | 1.472832 | 1.510845 | 1.544685 | 1.585078 | 1.656065 | 1.740462 | 1.831229 | 1.9189 | 1.974302 | 2.024249 | 2.084655 | 2.140574 |
|  | 80 | 1.413554 | 1.444525 | 1.480668 | 1.512914 | 1.551496 | 1.619545 | 1.700871 | 1.788871 | 1.874417 | 1.928761 | 1.977949 | 2.037688 | 2.093238 |
|  | 90 | 1.357576 | 1.38632 | 1.41993 | 1.449977 | 1.486005 | 1.549765 | 1.626341 | 1.709686 | 1.791212 | 1.843271 | 1.890577 | 1.948271 | 2.002163 |
| **R_entorhinal** | 3 | 2.223935 | 2.396854 | 2.576551 | 2.721878 | 2.881759 | 3.136913 | 3.409043 | 3.65288 | 3.838754 | 3.938243 | 4.018919 | 4.107233 | 4.181514 |
|  | 5 | 2.248906 | 2.415512 | 2.590492 | 2.733145 | 2.891046 | 3.144674 | 3.416912 | 3.66291 | 3.852253 | 3.954227 | 4.037216 | 4.128346 | 4.20521 |
|  | 10 | 2.308054 | 2.460921 | 2.625256 | 2.761673 | 2.914825 | 3.164612 | 3.436919 | 3.688279 | 3.886622 | 3.995178 | 4.084358 | 4.1831 | 4.26701 |
|  | 15 | 2.363209 | 2.504966 | 2.660228 | 2.791069 | 2.939766 | 3.185615 | 3.45748 | 3.713788 | 3.921266 | 4.036742 | 4.132558 | 4.239593 | 4.331289 |
|  | 20 | 2.40774 | 2.541712 | 2.690301 | 2.816828 | 2.961859 | 3.203946 | 3.474361 | 3.733881 | 3.948927 | 4.070502 | 4.172335 | 4.287056 | 4.386106 |
|  | 30 | 2.448077 | 2.57572 | 2.718305 | 2.840365 | 2.980774 | 3.215644 | 3.478085 | 3.734299 | 3.953427 | 4.080016 | 4.187446 | 4.309901 | 4.416772 |
|  | 40 | 2.476381 | 2.597777 | 2.733898 | 2.850736 | 2.985341 | 3.210536 | 3.461741 | 3.709811 | 3.926959 | 4.054428 | 4.163671 | 4.289304 | 4.399854 |
|  | 50 | 2.486614 | 2.604998 | 2.737902 | 2.85207 | 2.983652 | 3.203744 | 3.449013 | 3.692243 | 3.907072 | 4.033967 | 4.143135 | 4.269122 | 4.380346 |
|  | 60 | 2.411265 | 2.539638 | 2.682105 | 2.803216 | 2.941463 | 3.169829 | 3.420556 | 3.665305 | 3.878212 | 4.00265 | 4.108974 | 4.230883 | 4.337832 |
|  | 70 | 2.210252 | 2.370984 | 2.542915 | 2.684565 | 2.841944 | 3.093502 | 3.359801 | 3.610089 | 3.820031 | 3.93978 | 4.040542 | 4.154442 | 4.253027 |
|  | 80 | 1.819614 | 2.065937 | 2.306591 | 2.491 | 2.684467 | 2.974715 | 3.263062 | 3.519247 | 3.724099 | 3.837463 | 3.931123 | 4.035259 | 4.124023 |
|  | 90 | 1.363601 | 1.693788 | 2.023251 | 2.266029 | 2.507248 | 2.844581 | 3.156463 | 3.417433 | 3.616481 | 3.723553 | 3.810565 | 3.905913 | 3.986121 |
| **R_fusiform** | 3 | 2.72771 | 2.763265 | 2.804303 | 2.840608 | 2.88379 | 2.959666 | 3.05068 | 3.150719 | 3.250664 | 3.316051 | 3.376794 | 3.452893 | 3.526293 |
|  | 5 | 2.66715 | 2.708427 | 2.755138 | 2.795652 | 2.842878 | 2.923423 | 3.01614 | 3.113431 | 3.206089 | 3.264377 | 3.316943 | 3.380722 | 3.44013 |
|  | 10 | 2.475338 | 2.543934 | 2.614962 | 2.671778 | 2.733245 | 2.828279 | 2.925476 | 3.016582 | 3.095204 | 3.141308 | 3.180967 | 3.226908 | 3.267775 |
|  | 15 | 2.32384 | 2.412784 | 2.499067 | 2.564589 | 2.632567 | 2.732828 | 2.830679 | 2.919322 | 2.994222 | 3.03771 | 3.074967 | 3.118042 | 3.156356 |
|  | 20 | 2.337394 | 2.399285 | 2.463897 | 2.516006 | 2.572887 | 2.662186 | 2.755935 | 2.847072 | 2.929272 | 2.979431 | 3.023952 | 3.077332 | 3.126644 |
|  | 30 | 2.281596 | 2.34113 | 2.403856 | 2.454861 | 2.510949 | 2.599846 | 2.694241 | 2.787012 | 2.871504 | 2.923431 | 2.969749 | 3.025563 | 3.077387 |
|  | 40 | 2.210911 | 2.273483 | 2.339384 | 2.39289 | 2.451559 | 2.543873 | 2.640356 | 2.732809 | 2.814284 | 2.862843 | 2.905103 | 2.954647 | 2.999275 |
|  | 50 | 2.216781 | 2.265044 | 2.318638 | 2.364244 | 2.416382 | 2.502778 | 2.598325 | 2.694174 | 2.781367 | 2.834228 | 2.880619 | 2.935317 | 2.984744 |
|  | 60 | 2.147666 | 2.204576 | 2.266524 | 2.318228 | 2.376219 | 2.469717 | 2.569376 | 2.665404 | 2.749322 | 2.798604 | 2.840864 | 2.88951 | 2.932376 |
|  | 70 | 2.117295 | 2.172006 | 2.232259 | 2.283072 | 2.340591 | 2.434395 | 2.535628 | 2.63415 | 2.720831 | 2.771915 | 2.815792 | 2.866346 | 2.91091 |
|  | 80 | 2.087325 | 2.132916 | 2.184647 | 2.229524 | 2.281715 | 2.370021 | 2.469791 | 2.571365 | 2.664384 | 2.7208 | 2.770197 | 2.828186 | 2.880251 |
|  | 90 | 2.015666 | 2.055137 | 2.10076 | 2.141072 | 2.188829 | 2.271839 | 2.369072 | 2.471977 | 2.569821 | 2.630901 | 2.68549 | 2.750924 | 2.81094 |
| **R_inferiorparietal** | 3 | 2.520629 | 2.590973 | 2.664279 | 2.722811 | 2.785484 | 2.879495 | 2.969781 | 3.055555 | 3.13617 | 3.185561 | 3.228846 | 3.279492 | 3.324662 |
|  | 5 | 2.472053 | 2.540574 | 2.612241 | 2.669661 | 2.731346 | 2.824313 | 2.914172 | 2.999655 | 3.079716 | 3.12867 | 3.171528 | 3.221636 | 3.266297 |
|  | 10 | 2.354323 | 2.418431 | 2.486091 | 2.540767 | 2.599993 | 2.690321 | 2.779057 | 2.863775 | 2.942451 | 2.990329 | 3.032141 | 3.080931 | 3.124353 |
|  | 15 | 2.242605 | 2.302476 | 2.366231 | 2.418192 | 2.474951 | 2.562566 | 2.65006 | 2.733924 | 2.811184 | 2.857987 | 2.898763 | 2.94626 | 2.98847 |
|  | 20 | 2.153748 | 2.209975 | 2.270369 | 2.320004 | 2.374672 | 2.460082 | 2.546783 | 2.630234 | 2.706522 | 2.75253 | 2.792522 | 2.839022 | 2.880291 |
|  | 30 | 2.072681 | 2.124383 | 2.180765 | 2.227801 | 2.280397 | 2.364448 | 2.452459 | 2.537796 | 2.614573 | 2.660431 | 2.700088 | 2.746011 | 2.786634 |
|  | 40 | 2.030882 | 2.07974 | 2.133681 | 2.179248 | 2.230875 | 2.315057 | 2.405713 | 2.494081 | 2.572228 | 2.618391 | 2.658069 | 2.703784 | 2.744053 |
|  | 50 | 1.99341 | 2.039401 | 2.090726 | 2.134572 | 2.184845 | 2.268375 | 2.360708 | 2.451087 | 2.529606 | 2.575444 | 2.614579 | 2.659417 | 2.698724 |
|  | 60 | 1.951452 | 1.993997 | 2.041951 | 2.083349 | 2.131359 | 2.212589 | 2.30465 | 2.3951 | 2.472311 | 2.516841 | 2.554598 | 2.597604 | 2.635115 |
|  | 70 | 1.930488 | 1.969799 | 2.014526 | 2.053529 | 2.099261 | 2.178021 | 2.269462 | 2.359622 | 2.435277 | 2.478378 | 2.514666 | 2.555753 | 2.591405 |
|  | 80 | 1.887928 | 1.923619 | 1.964595 | 2.000674 | 2.043435 | 2.118376 | 2.207443 | 2.295577 | 2.368312 | 2.409243 | 2.443461 | 2.481972 | 2.515215 |
|  | 90 | 1.822321 | 1.854309 | 1.891351 | 1.924274 | 1.963705 | 2.034008 | 2.119477 | 2.204349 | 2.273275 | 2.311588 | 2.343388 | 2.378962 | 2.409509 |
| **R_inferiortemporal** | 3 | 2.525628 | 2.621596 | 2.718815 | 2.795169 | 2.876419 | 2.999178 | 3.120658 | 3.230005 | 3.320142 | 3.370922 | 3.413269 | 3.460699 | 3.501366 |
|  | 5 | 2.491359 | 2.587267 | 2.684279 | 2.760383 | 2.841293 | 2.963415 | 3.084141 | 3.192721 | 3.282172 | 3.332547 | 3.374546 | 3.421576 | 3.461893 |
|  | 10 | 2.415578 | 2.508609 | 2.603008 | 2.677249 | 2.756342 | 2.876001 | 2.994589 | 3.101469 | 3.189661 | 3.23938 | 3.28086 | 3.327337 | 3.367202 |
|  | 15 | 2.392543 | 2.468317 | 2.548492 | 2.613736 | 2.685243 | 2.797098 | 2.911994 | 3.01872 | 3.108891 | 3.160518 | 3.204009 | 3.25318 | 3.295716 |
|  | 20 | 2.400952 | 2.457493 | 2.520441 | 2.57405 | 2.635275 | 2.736259 | 2.846637 | 2.955227 | 3.051532 | 3.108553 | 3.157653 | 3.214338 | 3.264378 |
|  | 30 | 2.343767 | 2.397571 | 2.457943 | 2.509743 | 2.569328 | 2.668574 | 2.778397 | 2.887771 | 2.985844 | 3.044377 | 3.095051 | 3.153863 | 3.206052 |
|  | 40 | 2.293118 | 2.350279 | 2.413949 | 2.4682 | 2.53019 | 2.632503 | 2.744432 | 2.854646 | 2.952471 | 3.010427 | 3.060352 | 3.118014 | 3.168937 |
|  | 50 | 2.256504 | 2.313806 | 2.377742 | 2.432308 | 2.494753 | 2.598037 | 2.711324 | 2.823167 | 2.922669 | 2.981718 | 3.032643 | 3.091524 | 3.14358 |
|  | 60 | 2.191503 | 2.253951 | 2.32277 | 2.380827 | 2.44655 | 2.553682 | 2.669116 | 2.781133 | 2.879299 | 2.936934 | 2.986285 | 3.042954 | 3.092716 |
|  | 70 | 2.166547 | 2.223898 | 2.288069 | 2.342982 | 2.405987 | 2.510562 | 2.625776 | 2.74002 | 2.84206 | 2.902788 | 2.955263 | 3.01605 | 3.069893 |
|  | 80 | 2.163827 | 2.211734 | 2.266799 | 2.315175 | 2.372141 | 2.470245 | 2.583664 | 2.70191 | 2.812618 | 2.880876 | 2.941323 | 3.013088 | 3.078253 |
|  | 90 | 2.15528 | 2.196719 | 2.245255 | 2.288728 | 2.340963 | 2.433711 | 2.545669 | 2.668296 | 2.789097 | 2.866706 | 2.937563 | 3.024432 | 3.10604 |
| **R_isthmuscingulate** | 3 | 2.464882 | 2.534255 | 2.61006 | 2.673857 | 2.746379 | 2.866807 | 3.002874 | 3.146018 | 3.28573 | 3.376562 | 3.461081 | 3.567643 | 3.671548 |
|  | 5 | 2.416464 | 2.483971 | 2.557994 | 2.620479 | 2.691692 | 2.810285 | 2.944581 | 3.085914 | 3.223631 | 3.312941 | 3.395835 | 3.500023 | 3.601232 |
|  | 10 | 2.297768 | 2.361028 | 2.430938 | 2.490357 | 2.558471 | 2.672637 | 2.802587 | 2.939475 | 3.072388 | 3.158104 | 3.237225 | 3.335982 | 3.431116 |
|  | 15 | 2.191801 | 2.251559 | 2.318051 | 2.3749 | 2.440398 | 2.550801 | 2.677042 | 2.810152 | 2.939027 | 3.021756 | 3.097765 | 3.192088 | 3.28231 |
|  | 20 | 2.114827 | 2.172168 | 2.236347 | 2.291503 | 2.355332 | 2.463452 | 2.587585 | 2.718605 | 2.845168 | 2.926103 | 3.000177 | 3.091649 | 3.178623 |
|  | 30 | 2.024324 | 2.079155 | 2.141121 | 2.194827 | 2.257429 | 2.364343 | 2.48794 | 2.618671 | 2.744565 | 2.824615 | 2.897449 | 2.986717 | 3.070818 |
|  | 40 | 1.957314 | 2.010828 | 2.071757 | 2.124908 | 2.187214 | 2.294311 | 2.418828 | 2.550826 | 2.677736 | 2.758137 | 2.831003 | 2.919851 | 3.003024 |
|  | 50 | 1.887628 | 1.940117 | 2.000225 | 2.052929 | 2.114988 | 2.22222 | 2.347497 | 2.480615 | 2.608541 | 2.689411 | 2.762517 | 2.851355 | 2.934162 |
|  | 60 | 1.830628 | 1.882687 | 1.942577 | 1.995306 | 2.05762 | 2.165763 | 2.29264 | 2.427793 | 2.557722 | 2.639774 | 2.71384 | 2.803655 | 2.887142 |
|  | 70 | 1.795735 | 1.84817 | 1.90872 | 1.962211 | 2.025616 | 2.136066 | 2.266148 | 2.405073 | 2.538766 | 2.623181 | 2.699328 | 2.791562 | 2.877159 |
|  | 80 | 1.751687 | 1.804362 | 1.865381 | 1.91944 | 1.983686 | 2.095972 | 2.22869 | 2.370816 | 2.507799 | 2.594332 | 2.672384 | 2.766887 | 2.854525 |
|  | 90 | 1.708798 | 1.761831 | 1.82343 | 1.87814 | 1.943311 | 2.057557 | 2.193052 | 2.338566 | 2.479087 | 2.56794 | 2.648117 | 2.745208 | 2.835237 |
| **R_lateraloccipital** | 3 | 2.231998 | 2.272896 | 2.318899 | 2.358465 | 2.404081 | 2.480304 | 2.564994 | 2.649683 | 2.725906 | 2.771523 | 2.811088 | 2.857092 | 2.89799 |
|  | 5 | 2.190386 | 2.231309 | 2.277342 | 2.316932 | 2.362577 | 2.438848 | 2.52359 | 2.608333 | 2.684603 | 2.730249 | 2.769839 | 2.815871 | 2.856795 |
|  | 10 | 2.086956 | 2.127943 | 2.174048 | 2.2137 | 2.259416 | 2.335806 | 2.420682 | 2.505557 | 2.581947 | 2.627663 | 2.667316 | 2.71342 | 2.754407 |
|  | 15 | 1.988753 | 2.029804 | 2.07598 | 2.115695 | 2.161482 | 2.237992 | 2.323 | 2.408007 | 2.484517 | 2.530305 | 2.570019 | 2.616195 | 2.657247 |
|  | 20 | 1.915189 | 1.956304 | 2.002552 | 2.042328 | 2.088187 | 2.164816 | 2.249956 | 2.335096 | 2.411724 | 2.457583 | 2.497359 | 2.543607 | 2.584723 |
|  | 30 | 1.865245 | 1.906487 | 1.952878 | 1.992777 | 2.038777 | 2.115641 | 2.201044 | 2.286446 | 2.36331 | 2.409311 | 2.449209 | 2.4956 | 2.536842 |
|  | 40 | 1.838243 | 1.879612 | 1.926145 | 1.966166 | 2.012307 | 2.089408 | 2.175072 | 2.260737 | 2.337837 | 2.383979 | 2.424 | 2.470533 | 2.511901 |
|  | 50 | 1.815803 | 1.857299 | 1.903977 | 1.944121 | 1.990406 | 2.067745 | 2.153675 | 2.239604 | 2.316944 | 2.363228 | 2.403373 | 2.45005 | 2.491547 |
|  | 60 | 1.780398 | 1.822025 | 1.868849 | 1.90912 | 1.95555 | 2.033132 | 2.119331 | 2.205531 | 2.283113 | 2.329543 | 2.369814 | 2.416637 | 2.458264 |
|  | 70 | 1.744483 | 1.786243 | 1.833215 | 1.873614 | 1.920191 | 1.998019 | 2.084492 | 2.170965 | 2.248793 | 2.295371 | 2.335769 | 2.382742 | 2.424501 |
|  | 80 | 1.685765 | 1.727658 | 1.77478 | 1.815307 | 1.862033 | 1.940109 | 2.026858 | 2.113606 | 2.191683 | 2.238408 | 2.278936 | 2.326058 | 2.36795 |
|  | 90 | 1.577307 | 1.619333 | 1.666605 | 1.707262 | 1.754136 | 1.832462 | 1.919487 | 2.006512 | 2.084838 | 2.131712 | 2.172369 | 2.219641 | 2.261667 |
| **R_lateralorbitofrontal** | 3 | 2.604286 | 2.696168 | 2.790077 | 2.864443 | 2.944312 | 3.06707 | 3.192556 | 3.311182 | 3.415252 | 3.477365 | 3.531606 | 3.595537 | 3.653536 |
|  | 5 | 2.551617 | 2.638457 | 2.727899 | 2.799186 | 2.87617 | 2.995284 | 3.117949 | 3.234663 | 3.3376 | 3.399259 | 3.453229 | 3.516984 | 3.574951 |
|  | 10 | 2.431011 | 2.505857 | 2.584501 | 2.648269 | 2.718174 | 2.828372 | 2.944295 | 3.056739 | 3.157524 | 3.218577 | 3.272417 | 3.336482 | 3.39515 |
|  | 15 | 2.341742 | 2.404118 | 2.471323 | 2.527058 | 2.589426 | 2.690409 | 2.80011 | 2.90985 | 3.010939 | 3.073405 | 3.129249 | 3.196613 | 3.259164 |
|  | 20 | 2.289817 | 2.342258 | 2.400072 | 2.449071 | 2.505052 | 2.598322 | 2.703406 | 2.812544 | 2.916716 | 2.982865 | 3.043171 | 3.117407 | 3.187819 |
|  | 30 | 2.208031 | 2.258843 | 2.315048 | 2.362831 | 2.417577 | 2.509129 | 2.61272 | 2.720713 | 2.824093 | 2.889861 | 2.949884 | 3.023845 | 3.094057 |
|  | 40 | 2.124906 | 2.181913 | 2.243937 | 2.295833 | 2.35437 | 2.45011 | 2.555289 | 2.661498 | 2.760001 | 2.821107 | 2.875852 | 2.942001 | 3.003503 |
|  | 50 | 2.090405 | 2.143609 | 2.202158 | 2.251669 | 2.308084 | 2.401625 | 2.506152 | 2.613499 | 2.714598 | 2.778031 | 2.835311 | 2.905078 | 2.970478 |
|  | 60 | 2.072681 | 2.121877 | 2.176714 | 2.223663 | 2.277804 | 2.369092 | 2.473342 | 2.582853 | 2.688245 | 2.755481 | 2.816931 | 2.892719 | 2.964702 |
|  | 70 | 2.044216 | 2.095953 | 2.153381 | 2.20234 | 2.258551 | 2.352702 | 2.459203 | 2.569842 | 2.675066 | 2.741535 | 2.801826 | 2.87558 | 2.945012 |
|  | 80 | 2.030538 | 2.085974 | 2.147149 | 2.199001 | 2.258185 | 2.356471 | 2.466342 | 2.578976 | 2.684647 | 2.750663 | 2.81005 | 2.88206 | 2.949213 |
|  | 90 | 2.044808 | 2.100973 | 2.163153 | 2.21601 | 2.276509 | 2.377334 | 2.490511 | 2.606959 | 2.716522 | 2.785092 | 2.846844 | 2.921794 | 2.991747 |
| **R_lingual** | 3 | 2.203351 | 2.239055 | 2.27948 | 2.314508 | 2.355238 | 2.424262 | 2.502655 | 2.583224 | 2.657963 | 2.703849 | 2.744428 | 2.792599 | 2.836391 |
|  | 5 | 2.139404 | 2.175085 | 2.21549 | 2.250509 | 2.29124 | 2.3603 | 2.4388 | 2.519572 | 2.594597 | 2.640712 | 2.681531 | 2.730035 | 2.774176 |
|  | 10 | 1.994796 | 2.030521 | 2.070993 | 2.106088 | 2.146936 | 2.216283 | 2.295285 | 2.376816 | 2.452815 | 2.499674 | 2.541253 | 2.59079 | 2.636003 |
|  | 15 | 1.876022 | 1.911953 | 1.95267 | 1.987994 | 2.029134 | 2.099066 | 2.178912 | 2.261574 | 2.338916 | 2.386764 | 2.429332 | 2.480196 | 2.526769 |
|  | 20 | 1.794963 | 1.83114 | 1.872132 | 1.907699 | 1.949136 | 2.019631 | 2.100265 | 2.183977 | 2.262581 | 2.31137 | 2.35489 | 2.407045 | 2.454955 |
|  | 30 | 1.739193 | 1.775566 | 1.816697 | 1.85233 | 1.893803 | 1.964325 | 2.04508 | 2.129202 | 2.208622 | 2.258193 | 2.302621 | 2.356156 | 2.405643 |
|  | 40 | 1.70823 | 1.743889 | 1.784047 | 1.818719 | 1.858964 | 1.927223 | 2.005292 | 2.086755 | 2.164012 | 2.21249 | 2.256147 | 2.309057 | 2.358296 |
|  | 50 | 1.679169 | 1.714266 | 1.753567 | 1.787338 | 1.826385 | 1.892355 | 1.967639 | 2.046313 | 2.121315 | 2.168682 | 2.211593 | 2.263971 | 2.313128 |
|  | 60 | 1.634102 | 1.670368 | 1.710723 | 1.745219 | 1.784943 | 1.851805 | 1.928021 | 2.007971 | 2.084855 | 2.133892 | 2.178709 | 2.233992 | 2.286516 |
|  | 70 | 1.595269 | 1.633213 | 1.675071 | 1.710592 | 1.751261 | 1.81933 | 1.896728 | 1.978244 | 2.057456 | 2.108602 | 2.155871 | 2.214964 | 2.271998 |
|  | 80 | 1.561465 | 1.601024 | 1.644129 | 1.680326 | 1.72141 | 1.789564 | 1.866647 | 1.948082 | 2.028145 | 2.080601 | 2.129742 | 2.192198 | 2.25367 |
|  | 90 | 1.490557 | 1.531394 | 1.57517 | 1.61141 | 1.652058 | 1.718656 | 1.793372 | 1.872541 | 1.951508 | 2.004212 | 2.054451 | 2.119679 | 2.185533 |
| **R_medialorbitofrontal** | 3 | 2.542465 | 2.608054 | 2.681421 | 2.744332 | 2.816836 | 2.9385 | 3.075559 | 3.216117 | 3.347042 | 3.428001 | 3.500114 | 3.586503 | 3.665909 |
|  | 5 | 2.459296 | 2.523204 | 2.594613 | 2.655788 | 2.726242 | 2.84438 | 2.977419 | 3.113903 | 3.241169 | 3.319965 | 3.390233 | 3.474528 | 3.552133 |
|  | 10 | 2.272551 | 2.332797 | 2.399907 | 2.45725 | 2.523152 | 2.63343 | 2.757478 | 2.884853 | 3.00397 | 3.077981 | 3.144193 | 3.223925 | 3.297659 |
|  | 15 | 2.138236 | 2.196226 | 2.260582 | 2.315402 | 2.378247 | 2.483144 | 2.600967 | 2.722063 | 2.835679 | 2.906556 | 2.970198 | 3.047175 | 3.118731 |
|  | 20 | 2.061655 | 2.11902 | 2.182408 | 2.236208 | 2.297701 | 2.400033 | 2.514769 | 2.632803 | 2.743954 | 2.813609 | 2.876418 | 2.952771 | 3.024169 |
|  | 30 | 1.973499 | 2.03169 | 2.095351 | 2.148924 | 2.209734 | 2.31021 | 2.422371 | 2.537977 | 2.647748 | 2.71726 | 2.780545 | 2.858376 | 2.932154 |
|  | 40 | 1.926025 | 1.986338 | 2.05164 | 2.106109 | 2.167489 | 2.26815 | 2.379992 | 2.495501 | 2.606137 | 2.676968 | 2.742111 | 2.823212 | 2.901205 |
|  | 50 | 1.882039 | 1.944268 | 2.011039 | 2.066306 | 2.12819 | 2.229017 | 2.340599 | 2.456073 | 2.567574 | 2.639682 | 2.706617 | 2.790888 | 2.873002 |
|  | 60 | 1.837189 | 1.900529 | 1.968067 | 2.023669 | 2.085655 | 2.186201 | 2.2972 | 2.412306 | 2.524185 | 2.597115 | 2.665307 | 2.751913 | 2.837168 |
|  | 70 | 1.829328 | 1.894243 | 1.96322 | 2.019838 | 2.082804 | 2.184706 | 2.297104 | 2.413903 | 2.527996 | 2.602795 | 2.673097 | 2.76293 | 2.851999 |
|  | 80 | 1.848049 | 1.91511 | 1.986217 | 2.044477 | 2.109181 | 2.213768 | 2.329123 | 2.449247 | 2.567082 | 2.644695 | 2.717941 | 2.811994 | 2.905776 |
|  | 90 | 1.835781 | 1.903767 | 1.975732 | 2.034612 | 2.099932 | 2.205425 | 2.321801 | 2.443251 | 2.562868 | 2.642001 | 2.716969 | 2.813673 | 2.910611 |
| **R_middletemporal** | 3 | 2.770596 | 2.856591 | 2.945957 | 3.017277 | 3.09377 | 3.209224 | 3.321657 | 3.427205 | 3.522863 | 3.580125 | 3.629616 | 3.686813 | 3.737256 |
|  | 5 | 2.71015 | 2.801617 | 2.895235 | 2.969029 | 3.047379 | 3.164286 | 3.276827 | 3.381207 | 3.474629 | 3.530078 | 3.577738 | 3.632538 | 3.680633 |
|  | 10 | 2.567614 | 2.67166 | 2.774418 | 2.853216 | 2.935099 | 3.054474 | 3.166878 | 3.268708 | 3.357599 | 3.40949 | 3.453632 | 3.503906 | 3.547634 |
|  | 15 | 2.515727 | 2.60486 | 2.695836 | 2.767455 | 2.843497 | 2.957193 | 3.06724 | 3.168664 | 3.257894 | 3.310281 | 3.355024 | 3.406182 | 3.450857 |
|  | 20 | 2.531037 | 2.593207 | 2.661206 | 2.718044 | 2.781656 | 2.883223 | 2.989167 | 3.092974 | 3.188625 | 3.24669 | 3.297413 | 3.356698 | 3.409601 |
|  | 30 | 2.46462 | 2.518503 | 2.57878 | 2.630274 | 2.689156 | 2.78608 | 2.891273 | 2.99716 | 3.095968 | 3.156588 | 3.209977 | 3.27292 | 3.329602 |
|  | 40 | 2.359566 | 2.427005 | 2.499855 | 2.560147 | 2.627129 | 2.73342 | 2.844018 | 2.94967 | 3.042632 | 3.097319 | 3.144162 | 3.19793 | 3.245102 |
|  | 50 | 2.362569 | 2.415976 | 2.476077 | 2.527773 | 2.587364 | 2.686837 | 2.797132 | 2.907865 | 3.008491 | 3.06912 | 3.121932 | 3.183581 | 3.238595 |
|  | 60 | 2.280958 | 2.337381 | 2.400427 | 2.454323 | 2.516121 | 2.618676 | 2.731722 | 2.84323 | 2.941789 | 3.000017 | 3.0501 | 3.107871 | 3.158836 |
|  | 70 | 2.2451 | 2.297707 | 2.357245 | 2.408797 | 2.468688 | 2.570036 | 2.684715 | 2.799498 | 2.901109 | 2.961231 | 3.01303 | 3.072904 | 3.125856 |
|  | 80 | 2.216746 | 2.26445 | 2.319299 | 2.367561 | 2.424568 | 2.523487 | 2.639308 | 2.758031 | 2.864242 | 2.927592 | 2.982506 | 3.046399 | 3.103296 |
|  | 90 | 2.171072 | 2.216417 | 2.268992 | 2.315663 | 2.371323 | 2.469394 | 2.58673 | 2.708377 | 2.81713 | 2.881978 | 2.938215 | 3.00371 | 3.062116 |
| **R_parahippocampal** | 3 | 2.291323 | 2.378816 | 2.47713 | 2.561602 | 2.658902 | 2.821279 | 3.001402 | 3.181239 | 3.342868 | 3.439497 | 3.52325 | 3.620565 | 3.707021 |
|  | 5 | 2.261806 | 2.349266 | 2.447499 | 2.531868 | 2.62901 | 2.791035 | 2.970648 | 3.149857 | 3.310825 | 3.407018 | 3.49037 | 3.587191 | 3.673184 |
|  | 10 | 2.18983 | 2.277246 | 2.375317 | 2.459456 | 2.556235 | 2.717429 | 2.895811 | 3.07349 | 3.232843 | 3.327967 | 3.410332 | 3.505939 | 3.590794 |
|  | 15 | 2.122637 | 2.210146 | 2.308203 | 2.392234 | 2.488784 | 2.649363 | 2.826746 | 3.003118 | 3.161054 | 3.255227 | 3.336706 | 3.431217 | 3.51504 |
|  | 20 | 2.065392 | 2.153338 | 2.251759 | 2.336 | 2.432683 | 2.593238 | 2.770261 | 2.945957 | 3.103035 | 3.19659 | 3.277473 | 3.371221 | 3.454308 |
|  | 30 | 1.985821 | 2.075824 | 2.176268 | 2.26202 | 2.360198 | 2.522711 | 2.70119 | 2.877661 | 3.034909 | 3.128345 | 3.208999 | 3.302341 | 3.384944 |
|  | 40 | 1.920529 | 2.012836 | 2.115537 | 2.202972 | 2.302815 | 2.467516 | 2.647647 | 2.825047 | 2.982583 | 3.075963 | 3.156441 | 3.249434 | 3.331605 |
|  | 50 | 1.866044 | 1.960998 | 2.066291 | 2.155659 | 2.257427 | 2.424686 | 2.606817 | 2.785451 | 2.943521 | 3.036988 | 3.117408 | 3.21019 | 3.292049 |
|  | 60 | 1.817854 | 1.915711 | 2.023829 | 2.115294 | 2.219137 | 2.389145 | 2.573423 | 2.753393 | 2.91207 | 3.005659 | 3.086051 | 3.178655 | 3.260231 |
|  | 70 | 1.768923 | 1.869868 | 1.980949 | 2.074583 | 2.180545 | 2.353305 | 2.539663 | 2.720856 | 2.880016 | 2.973648 | 3.053943 | 3.146286 | 3.227506 |
|  | 80 | 1.718151 | 1.822541 | 1.936894 | 2.032905 | 2.141172 | 2.316905 | 2.505496 | 2.688009 | 2.847707 | 2.941407 | 3.021621 | 3.113721 | 3.1946 |
|  | 90 | 1.660451 | 1.768295 | 1.885833 | 1.984085 | 2.094454 | 2.272736 | 2.463022 | 2.646276 | 2.80598 | 2.899428 | 2.979287 | 3.070826 | 3.151084 |
| **R_paracentral** | 3 | 2.351828 | 2.407611 | 2.469448 | 2.521986 | 2.581965 | 2.681181 | 2.790702 | 2.900412 | 3.000126 | 3.060559 | 3.11359 | 3.176121 | 3.232637 |
|  | 5 | 2.313098 | 2.368638 | 2.430145 | 2.482359 | 2.541921 | 2.640358 | 2.748917 | 2.857592 | 2.956339 | 3.016183 | 3.068703 | 3.130642 | 3.186637 |
|  | 10 | 2.215259 | 2.27014 | 2.330762 | 2.382108 | 2.440563 | 2.536938 | 2.642963 | 2.748929 | 2.845145 | 2.903456 | 2.954644 | 3.015044 | 3.069689 |
|  | 15 | 2.116518 | 2.170676 | 2.230336 | 2.280743 | 2.338006 | 2.432173 | 2.535505 | 2.638603 | 2.732152 | 2.788851 | 2.838643 | 2.897431 | 2.950665 |
|  | 20 | 2.031456 | 2.08521 | 2.144246 | 2.193995 | 2.250381 | 2.342852 | 2.444047 | 2.544836 | 2.636234 | 2.691638 | 2.740314 | 2.797828 | 2.84996 |
|  | 30 | 1.947844 | 2.003175 | 2.063534 | 2.114097 | 2.171112 | 2.264048 | 2.365153 | 2.465487 | 2.556371 | 2.611499 | 2.659991 | 2.717394 | 2.769553 |
|  | 40 | 1.882984 | 1.940754 | 2.003274 | 2.055286 | 2.113586 | 2.207959 | 2.309948 | 2.410767 | 2.502015 | 2.557426 | 2.606251 | 2.664188 | 2.717001 |
|  | 50 | 1.834609 | 1.895847 | 1.961497 | 2.015669 | 2.075971 | 2.172805 | 2.276676 | 2.378934 | 2.471439 | 2.52771 | 2.577406 | 2.636566 | 2.690712 |
|  | 60 | 1.783163 | 1.848488 | 1.917738 | 1.974335 | 2.036824 | 2.136249 | 2.242 | 2.345655 | 2.439421 | 2.496598 | 2.547247 | 2.607787 | 2.663479 |
|  | 70 | 1.720063 | 1.789962 | 1.863071 | 1.922143 | 1.986746 | 2.088436 | 2.195561 | 2.300079 | 2.394673 | 2.452549 | 2.504018 | 2.565852 | 2.623093 |
|  | 80 | 1.636914 | 1.71168 | 1.788621 | 1.849948 | 1.916264 | 2.019351 | 2.126763 | 2.231045 | 2.325542 | 2.383617 | 2.435518 | 2.49827 | 2.556813 |
|  | 90 | 1.535317 | 1.615442 | 1.696287 | 1.759679 | 1.827315 | 1.930924 | 2.037528 | 2.140483 | 2.233975 | 2.291767 | 2.343735 | 2.407066 | 2.466713 |
| **R_parsopercularis** | 3 | 2.448905 | 2.535293 | 2.622459 | 2.690767 | 2.763528 | 2.874457 | 2.987289 | 3.094156 | 3.188693 | 3.245701 | 3.295947 | 3.35582 | 3.410826 |
|  | 5 | 2.428174 | 2.510808 | 2.594925 | 2.661324 | 2.73248 | 2.841711 | 2.953576 | 3.060028 | 3.154436 | 3.211411 | 3.261629 | 3.321447 | 3.376363 |
|  | 10 | 2.376177 | 2.450153 | 2.52705 | 2.58883 | 2.656026 | 2.760989 | 2.870407 | 2.975894 | 3.070175 | 3.127262 | 3.177633 | 3.237643 | 3.292699 |
|  | 15 | 2.327761 | 2.392966 | 2.462269 | 2.519032 | 2.581823 | 2.681946 | 2.788679 | 2.893473 | 2.988368 | 3.046262 | 3.097556 | 3.158871 | 3.215272 |
|  | 20 | 2.283904 | 2.341064 | 2.403135 | 2.454966 | 2.513313 | 2.608447 | 2.712489 | 2.81698 | 2.913316 | 2.972779 | 3.025848 | 3.089706 | 3.148812 |
|  | 30 | 2.197107 | 2.247683 | 2.303692 | 2.351305 | 2.405781 | 2.496457 | 2.597939 | 2.701838 | 2.798953 | 2.859357 | 2.91348 | 2.978799 | 3.039376 |
|  | 40 | 2.103979 | 2.157462 | 2.216328 | 2.266046 | 2.322528 | 2.415489 | 2.517756 | 2.620253 | 2.713828 | 2.770872 | 2.821203 | 2.880943 | 2.935356 |
|  | 50 | 2.037214 | 2.090798 | 2.149908 | 2.199909 | 2.256768 | 2.350367 | 2.453153 | 2.55571 | 2.648702 | 2.705008 | 2.754407 | 2.812665 | 2.865346 |
|  | 60 | 2.003273 | 2.052906 | 2.108466 | 2.156108 | 2.210973 | 2.302803 | 2.405643 | 2.510099 | 2.606192 | 2.664925 | 2.716755 | 2.778198 | 2.834023 |
|  | 70 | 1.968695 | 2.019692 | 2.076789 | 2.12574 | 2.18208 | 2.276233 | 2.381328 | 2.487536 | 2.584616 | 2.643605 | 2.695416 | 2.756521 | 2.81172 |
|  | 80 | 1.918614 | 1.976862 | 2.041034 | 2.095204 | 2.156617 | 2.257106 | 2.366239 | 2.473419 | 2.568762 | 2.625508 | 2.674626 | 2.731702 | 2.782483 |
|  | 90 | 1.844221 | 1.915497 | 1.991555 | 2.053946 | 2.122862 | 2.231873 | 2.345573 | 2.45303 | 2.545457 | 2.599156 | 2.644887 | 2.697188 | 2.742987 |
| **R_parsorbitalis** | 3 | 2.520086 | 2.619652 | 2.727277 | 2.816529 | 2.916019 | 3.075038 | 3.242523 | 3.401715 | 3.538832 | 3.618389 | 3.685993 | 3.76306 | 3.830262 |
|  | 5 | 2.48345 | 2.577647 | 2.680219 | 2.765835 | 2.861827 | 3.016389 | 3.180572 | 3.337838 | 3.474172 | 3.553625 | 3.621334 | 3.698729 | 3.766396 |
|  | 10 | 2.392963 | 2.475284 | 2.566429 | 2.643663 | 2.731463 | 2.875404 | 3.031608 | 3.184254 | 3.318864 | 3.398251 | 3.466434 | 3.544959 | 3.614113 |
|  | 15 | 2.313934 | 2.386084 | 2.467203 | 2.536938 | 2.617301 | 2.751502 | 2.900495 | 3.049371 | 3.183272 | 3.263364 | 3.332809 | 3.413526 | 3.485261 |
|  | 20 | 2.258474 | 2.322878 | 2.396223 | 2.460065 | 2.53454 | 2.661049 | 2.804618 | 2.951329 | 3.086036 | 3.167852 | 3.239539 | 3.323731 | 3.399337 |
|  | 30 | 2.189428 | 2.247933 | 2.315233 | 2.374398 | 2.44411 | 2.564251 | 2.703244 | 2.848214 | 2.983961 | 3.06765 | 3.141752 | 3.229709 | 3.309552 |
|  | 40 | 2.119077 | 2.179724 | 2.249102 | 2.309756 | 2.380823 | 2.502301 | 2.641298 | 2.784564 | 2.917181 | 2.998221 | 3.069531 | 3.153639 | 3.229495 |
|  | 50 | 2.037376 | 2.103579 | 2.178399 | 2.24304 | 2.317894 | 2.443731 | 2.584624 | 2.726608 | 2.855298 | 2.932709 | 3.000086 | 3.078695 | 3.148818 |
|  | 60 | 1.93177 | 2.005327 | 2.087151 | 2.156792 | 2.23629 | 2.367353 | 2.510571 | 2.651475 | 2.776473 | 2.850508 | 2.914277 | 2.987921 | 3.052957 |
|  | 70 | 1.891364 | 1.967433 | 2.052009 | 2.123958 | 2.206058 | 2.34134 | 2.489078 | 2.634352 | 2.76317 | 2.839445 | 2.905131 | 2.980976 | 3.047943 |
|  | 80 | 1.863843 | 1.938006 | 2.021198 | 2.09256 | 2.174631 | 2.311302 | 2.462511 | 2.613088 | 2.748106 | 2.828692 | 2.898461 | 2.97944 | 3.051305 |
|  | 90 | 1.831553 | 1.90462 | 1.987131 | 2.058359 | 2.140776 | 2.279169 | 2.433886 | 2.589555 | 2.730436 | 2.815086 | 2.888706 | 2.974532 | 3.051034 |
| **R_parstriangularis** | 3 | 2.512439 | 2.567629 | 2.629008 | 2.68137 | 2.741445 | 2.841707 | 2.954048 | 3.06886 | 3.175677 | 3.241757 | 3.300678 | 3.371379 | 3.436509 |
|  | 5 | 2.465161 | 2.519797 | 2.580526 | 2.632309 | 2.691695 | 2.790762 | 2.901725 | 3.015117 | 3.120635 | 3.185933 | 3.244177 | 3.314094 | 3.378534 |
|  | 10 | 2.351801 | 2.405114 | 2.464286 | 2.514678 | 2.572409 | 2.668601 | 2.776242 | 2.886214 | 2.988604 | 3.052021 | 3.108637 | 3.176673 | 3.239464 |
|  | 15 | 2.250343 | 2.302542 | 2.360391 | 2.409589 | 2.465889 | 2.559583 | 2.664322 | 2.771305 | 2.870966 | 2.932751 | 2.987961 | 3.054384 | 3.115771 |
|  | 20 | 2.171956 | 2.223528 | 2.280588 | 2.329048 | 2.384437 | 2.476494 | 2.579296 | 2.684272 | 2.782123 | 2.842845 | 2.897158 | 2.962583 | 3.023136 |
|  | 30 | 2.070014 | 2.121528 | 2.178321 | 2.226407 | 2.281227 | 2.372076 | 2.473294 | 2.576596 | 2.67301 | 2.732967 | 2.786711 | 2.851623 | 2.911894 |
|  | 40 | 1.997183 | 2.049245 | 2.106414 | 2.15465 | 2.20948 | 2.300054 | 2.400699 | 2.50335 | 2.599293 | 2.6591 | 2.712836 | 2.77793 | 2.838588 |
|  | 50 | 1.930182 | 1.982814 | 2.040348 | 2.088703 | 2.143487 | 2.233654 | 2.333549 | 2.435358 | 2.530665 | 2.590232 | 2.643894 | 2.709115 | 2.770133 |
|  | 60 | 1.870078 | 1.923396 | 1.981385 | 2.029909 | 2.084681 | 2.174458 | 2.27359 | 2.374536 | 2.469201 | 2.528544 | 2.582163 | 2.647573 | 2.70904 |
|  | 70 | 1.838159 | 1.893016 | 1.952342 | 2.001741 | 2.057271 | 2.147874 | 2.247545 | 2.348949 | 2.444238 | 2.504172 | 2.558506 | 2.625065 | 2.687925 |
|  | 80 | 1.822495 | 1.879578 | 1.940921 | 1.991721 | 2.048562 | 2.140833 | 2.241924 | 2.344674 | 2.441455 | 2.50256 | 2.558165 | 2.6266 | 2.691595 |
|  | 90 | 1.804548 | 1.864066 | 1.927575 | 1.979849 | 2.038039 | 2.131965 | 2.234403 | 2.338421 | 2.436662 | 2.498957 | 2.555886 | 2.626324 | 2.693642 |
| **R_pericalcarine** | 3 | 1.537751 | 1.569822 | 1.607734 | 1.642081 | 1.683942 | 1.760269 | 1.856188 | 1.960087 | 2.055986 | 2.114582 | 2.166313 | 2.227689 | 2.283497 |
|  | 5 | 1.507149 | 1.538751 | 1.576102 | 1.609931 | 1.651146 | 1.726238 | 1.820498 | 1.922701 | 2.017329 | 2.075288 | 2.126539 | 2.187438 | 2.242896 |
|  | 10 | 1.436492 | 1.467016 | 1.503078 | 1.535718 | 1.575447 | 1.647694 | 1.738127 | 1.836425 | 1.928158 | 1.984684 | 2.03487 | 2.094738 | 2.149468 |
|  | 15 | 1.37909 | 1.408781 | 1.443845 | 1.475561 | 1.514131 | 1.584137 | 1.671514 | 1.76673 | 1.856295 | 1.911826 | 1.96133 | 2.020622 | 2.075039 |
|  | 20 | 1.338045 | 1.367222 | 1.401663 | 1.432798 | 1.470626 | 1.539155 | 1.624442 | 1.717614 | 1.805961 | 1.861076 | 1.910415 | 1.969749 | 2.024427 |
|  | 30 | 1.295164 | 1.324045 | 1.358106 | 1.388856 | 1.426147 | 1.493439 | 1.576676 | 1.668046 | 1.756079 | 1.811683 | 1.861879 | 1.922744 | 1.979299 |
|  | 40 | 1.275179 | 1.304089 | 1.338149 | 1.368852 | 1.406007 | 1.472772 | 1.554802 | 1.645227 | 1.733729 | 1.790313 | 1.841819 | 1.904792 | 1.963795 |
|  | 50 | 1.256049 | 1.284892 | 1.318833 | 1.349378 | 1.38626 | 1.452239 | 1.532715 | 1.621767 | 1.710285 | 1.767562 | 1.820135 | 1.884946 | 1.946191 |
|  | 60 | 1.240718 | 1.269516 | 1.303361 | 1.333767 | 1.370397 | 1.435622 | 1.514573 | 1.602244 | 1.690745 | 1.748701 | 1.802342 | 1.869031 | 1.932598 |
|  | 70 | 1.228002 | 1.256753 | 1.290498 | 1.32076 | 1.357128 | 1.421575 | 1.498961 | 1.585167 | 1.673532 | 1.732092 | 1.786746 | 1.855276 | 1.921181 |
|  | 80 | 1.20962 | 1.23815 | 1.271588 | 1.301518 | 1.337397 | 1.400666 | 1.476 | 1.560166 | 1.647767 | 1.706512 | 1.7618 | 1.831731 | 1.899596 |
|  | 90 | 1.188541 | 1.216773 | 1.249816 | 1.279335 | 1.314634 | 1.376568 | 1.449677 | 1.531586 | 1.618154 | 1.676902 | 1.732668 | 1.803831 | 1.873549 |
| **R_postcentral** | 3 | 2.069264 | 2.104238 | 2.144048 | 2.178828 | 2.219736 | 2.290644 | 2.37454 | 2.465944 | 2.556998 | 2.616649 | 2.672227 | 2.742178 | 2.810075 |
|  | 5 | 2.027296 | 2.062488 | 2.102439 | 2.137249 | 2.178085 | 2.248599 | 2.331609 | 2.421552 | 2.510659 | 2.568776 | 2.622743 | 2.690417 | 2.75584 |
|  | 10 | 1.927031 | 1.962873 | 2.003268 | 2.03822 | 2.078939 | 2.148561 | 2.229469 | 2.31594 | 2.400466 | 2.455014 | 2.505269 | 2.567762 | 2.627628 |
|  | 15 | 1.84018 | 1.876912 | 1.917994 | 1.953276 | 1.994084 | 2.063159 | 2.142402 | 2.22598 | 2.306669 | 2.358249 | 2.405447 | 2.463722 | 2.519129 |
|  | 20 | 1.773898 | 1.811921 | 1.854102 | 1.890052 | 1.931324 | 2.000485 | 2.078834 | 2.16045 | 2.238366 | 2.287767 | 2.332713 | 2.387888 | 2.440036 |
|  | 30 | 1.695734 | 1.737699 | 1.783452 | 1.821818 | 1.865202 | 1.936453 | 2.01525 | 2.095509 | 2.170692 | 2.217744 | 2.260188 | 2.311862 | 2.360306 |
|  | 40 | 1.6414 | 1.687516 | 1.736982 | 1.777858 | 1.823469 | 1.897133 | 1.977092 | 2.057278 | 2.131561 | 2.177754 | 2.21928 | 2.2697 | 2.31687 |
|  | 50 | 1.592734 | 1.641643 | 1.693511 | 1.735946 | 1.782894 | 1.857963 | 1.938676 | 2.01918 | 2.09368 | 2.14009 | 2.181918 | 2.232886 | 2.280789 |
|  | 60 | 1.554525 | 1.604288 | 1.656822 | 1.699647 | 1.746901 | 1.822315 | 1.90347 | 1.984862 | 2.060926 | 2.1088 | 2.15233 | 2.205918 | 2.256871 |
|  | 70 | 1.526161 | 1.574333 | 1.625361 | 1.66711 | 1.71337 | 1.787748 | 1.86884 | 1.951704 | 2.030957 | 2.08191 | 2.12904 | 2.188185 | 2.245641 |
|  | 80 | 1.475047 | 1.519715 | 1.567331 | 1.606544 | 1.650306 | 1.721519 | 1.800738 | 1.883967 | 1.966316 | 2.020931 | 2.072745 | 2.139665 | 2.206828 |
|  | 90 | 1.40403 | 1.444979 | 1.488825 | 1.525116 | 1.565866 | 1.632936 | 1.709113 | 1.791664 | 1.876689 | 1.935283 | 1.992704 | 2.069754 | 2.150635 |
| **R_posteriorcingulate** | 3 | 2.569016 | 2.637746 | 2.708486 | 2.764959 | 2.826314 | 2.923146 | 3.02803 | 3.137149 | 3.246019 | 3.319475 | 3.390428 | 3.484195 | 3.580955 |
|  | 5 | 2.525294 | 2.591929 | 2.660868 | 2.716154 | 2.776455 | 2.872025 | 2.97587 | 3.083909 | 3.191349 | 3.263514 | 3.332917 | 3.42415 | 3.517702 |
|  | 10 | 2.414883 | 2.476763 | 2.54155 | 2.594054 | 2.65183 | 2.74429 | 2.845481 | 2.950762 | 3.054684 | 3.123767 | 3.189554 | 3.274986 | 3.361335 |
|  | 15 | 2.308588 | 2.366414 | 2.427602 | 2.477651 | 2.53316 | 2.622751 | 2.721426 | 2.824092 | 2.924773 | 2.991097 | 3.05371 | 3.134154 | 3.214434 |
|  | 20 | 2.226961 | 2.281807 | 2.34039 | 2.388704 | 2.44266 | 2.530402 | 2.62758 | 2.728691 | 2.827282 | 2.891713 | 2.952074 | 3.028898 | 3.10471 |
|  | 30 | 2.133774 | 2.185334 | 2.241283 | 2.288062 | 2.340909 | 2.427922 | 2.525187 | 2.626393 | 2.724149 | 2.787195 | 2.845513 | 2.918586 | 2.989366 |
|  | 40 | 2.071828 | 2.121749 | 2.176595 | 2.222945 | 2.275781 | 2.363623 | 2.462516 | 2.565421 | 2.664092 | 2.727076 | 2.784765 | 2.856175 | 2.924347 |
|  | 50 | 2.021818 | 2.071018 | 2.125601 | 2.172118 | 2.225515 | 2.314963 | 2.416223 | 2.521591 | 2.622045 | 2.685655 | 2.743469 | 2.814358 | 2.881268 |
|  | 60 | 1.971799 | 2.020751 | 2.075476 | 2.122419 | 2.176601 | 2.267897 | 2.37169 | 2.479689 | 2.582189 | 2.646685 | 2.704952 | 2.775864 | 2.842205 |
|  | 70 | 1.942792 | 1.992398 | 2.048185 | 2.096286 | 2.152039 | 2.246411 | 2.354052 | 2.466042 | 2.571949 | 2.638257 | 2.697875 | 2.770008 | 2.837018 |
|  | 80 | 1.927495 | 1.978444 | 2.03601 | 2.085841 | 2.143792 | 2.242226 | 2.354779 | 2.47186 | 2.582257 | 2.651102 | 2.712766 | 2.787028 | 2.855634 |
|  | 90 | 1.90406 | 1.956441 | 2.015841 | 2.06742 | 2.127555 | 2.229972 | 2.347295 | 2.469305 | 2.584073 | 2.655412 | 2.719117 | 2.79555 | 2.86585 |
| **R_precentral** | 3 | 2.253454 | 2.327446 | 2.403087 | 2.4626 | 2.525624 | 2.619193 | 2.708494 | 2.791615 | 2.867128 | 2.912364 | 2.951457 | 2.996616 | 3.036414 |
|  | 5 | 2.240368 | 2.313755 | 2.388895 | 2.44809 | 2.510844 | 2.604134 | 2.693296 | 2.776374 | 2.851899 | 2.897166 | 2.936298 | 2.981519 | 3.021385 |
|  | 10 | 2.206895 | 2.278768 | 2.352642 | 2.411025 | 2.473088 | 2.56565 | 2.654436 | 2.737383 | 2.81292 | 2.858254 | 2.897479 | 2.942847 | 2.982876 |
|  | 15 | 2.170107 | 2.240445 | 2.313013 | 2.370546 | 2.431868 | 2.523625 | 2.611957 | 2.694699 | 2.770185 | 2.815548 | 2.854834 | 2.900313 | 2.940477 |
|  | 20 | 2.129654 | 2.198525 | 2.269834 | 2.32654 | 2.387141 | 2.478106 | 2.565988 | 2.648526 | 2.72396 | 2.769351 | 2.808698 | 2.854289 | 2.894587 |
|  | 30 | 2.05154 | 2.118519 | 2.18832 | 2.244136 | 2.304073 | 2.394578 | 2.482606 | 2.565697 | 2.641892 | 2.687858 | 2.727772 | 2.774102 | 2.815125 |
|  | 40 | 1.982717 | 2.049359 | 2.119208 | 2.175342 | 2.235887 | 2.327817 | 2.417801 | 2.503145 | 2.581656 | 2.629134 | 2.670434 | 2.718454 | 2.761046 |
|  | 50 | 1.923478 | 1.990523 | 2.061206 | 2.118302 | 2.180166 | 2.27464 | 2.367733 | 2.456478 | 2.538406 | 2.588085 | 2.631381 | 2.681817 | 2.726638 |
|  | 60 | 1.866587 | 1.933342 | 2.004193 | 2.061764 | 2.124471 | 2.220874 | 2.316605 | 2.408421 | 2.49356 | 2.545359 | 2.590608 | 2.643442 | 2.690504 |
|  | 70 | 1.804179 | 1.869317 | 1.938974 | 1.995956 | 2.058396 | 2.155125 | 2.252035 | 2.345654 | 2.432936 | 2.486257 | 2.532969 | 2.587666 | 2.636525 |
|  | 80 | 1.73508 | 1.797862 | 1.865532 | 1.921282 | 1.982766 | 2.078803 | 2.175954 | 2.270561 | 2.359314 | 2.413789 | 2.461669 | 2.517918 | 2.568329 |
|  | 90 | 1.66354 | 1.723807 | 1.789287 | 1.843627 | 1.903958 | 1.999015 | 2.096171 | 2.191611 | 2.281769 | 2.3374 | 2.386477 | 2.444347 | 2.496405 |
| **R_precuneus** | 3 | 2.451295 | 2.531038 | 2.608563 | 2.667476 | 2.728734 | 2.819998 | 2.911814 | 2.999965 | 3.080856 | 3.131737 | 3.178275 | 3.236171 | 3.29203 |
|  | 5 | 2.404941 | 2.481426 | 2.556449 | 2.613883 | 2.67396 | 2.764031 | 2.855091 | 2.942602 | 3.022677 | 3.072824 | 3.118499 | 3.175038 | 3.22927 |
|  | 10 | 2.285403 | 2.354741 | 2.424123 | 2.478124 | 2.535374 | 2.62244 | 2.711461 | 2.797231 | 2.875248 | 2.923649 | 2.967334 | 3.020817 | 3.071462 |
|  | 15 | 2.166632 | 2.229978 | 2.294473 | 2.3454 | 2.400033 | 2.484183 | 2.571116 | 2.655126 | 2.73122 | 2.77808 | 2.820066 | 2.871008 | 2.918739 |
|  | 20 | 2.076188 | 2.135286 | 2.19634 | 2.245145 | 2.298031 | 2.380385 | 2.466231 | 2.549428 | 2.624542 | 2.67052 | 2.711468 | 2.760778 | 2.80657 |
|  | 30 | 1.985152 | 2.041726 | 2.101139 | 2.149285 | 2.202029 | 2.285071 | 2.372246 | 2.456562 | 2.531876 | 2.57736 | 2.617373 | 2.66486 | 2.708221 |
|  | 40 | 1.92593 | 1.982833 | 2.043047 | 2.092136 | 2.146151 | 2.231502 | 2.321101 | 2.407237 | 2.483294 | 2.528661 | 2.568151 | 2.614448 | 2.656139 |
|  | 50 | 1.882088 | 1.937332 | 1.996489 | 2.045202 | 2.099248 | 2.185423 | 2.276576 | 2.36445 | 2.441882 | 2.487864 | 2.527706 | 2.574143 | 2.615667 |
|  | 60 | 1.849257 | 1.899353 | 1.95402 | 1.99979 | 2.051319 | 2.134966 | 2.225163 | 2.31344 | 2.392006 | 2.438894 | 2.479608 | 2.527116 | 2.569606 |
|  | 70 | 1.836451 | 1.87957 | 1.927691 | 1.96882 | 2.016016 | 2.094567 | 2.181795 | 2.269492 | 2.349275 | 2.397587 | 2.439921 | 2.489731 | 2.534622 |
|  | 80 | 1.799951 | 1.83576 | 1.876571 | 1.91216 | 1.953806 | 2.025026 | 2.106887 | 2.192088 | 2.272079 | 2.321643 | 2.365759 | 2.418474 | 2.46672 |
|  | 90 | 1.749228 | 1.779117 | 1.813751 | 1.844459 | 1.881 | 1.945044 | 2.021148 | 2.103294 | 2.183251 | 2.23422 | 2.280533 | 2.337072 | 2.389992 |
| **R_rostralanteriorcingulate** | 3 | 2.588018 | 2.694566 | 2.81041 | 2.906601 | 3.01345 | 3.181889 | 3.354114 | 3.52358 | 3.684132 | 3.783402 | 3.871098 | 3.974641 | 4.067905 |
|  | 5 | 2.536871 | 2.640518 | 2.75342 | 2.847346 | 2.951878 | 3.117141 | 3.286803 | 3.453919 | 3.611928 | 3.709512 | 3.795666 | 3.897338 | 3.988881 |
|  | 10 | 2.4195 | 2.516485 | 2.622624 | 2.711336 | 2.810539 | 2.968509 | 3.132335 | 3.294121 | 3.446344 | 3.540083 | 3.622716 | 3.720115 | 3.807727 |
|  | 15 | 2.331141 | 2.422826 | 2.523626 | 2.608265 | 2.703364 | 2.855892 | 3.01568 | 3.173904 | 3.322065 | 3.413044 | 3.493123 | 3.5874 | 3.672124 |
|  | 20 | 2.27713 | 2.365012 | 2.462067 | 2.543933 | 2.636352 | 2.78565 | 2.943644 | 3.100524 | 3.246743 | 3.336279 | 3.414971 | 3.507507 | 3.59059 |
|  | 30 | 2.214431 | 2.296735 | 2.388431 | 2.466472 | 2.555399 | 2.701134 | 2.858502 | 3.015665 | 3.160846 | 3.249261 | 3.326743 | 3.417652 | 3.499128 |
|  | 40 | 2.163187 | 2.240638 | 2.327658 | 2.40237 | 2.488292 | 2.631134 | 2.788518 | 2.946659 | 3.091518 | 3.179273 | 3.255964 | 3.345748 | 3.426081 |
|  | 50 | 2.127241 | 2.200631 | 2.283765 | 2.355751 | 2.439295 | 2.580183 | 2.738567 | 2.898744 | 3.044313 | 3.132052 | 3.208523 | 3.297865 | 3.377674 |
|  | 60 | 2.109981 | 2.180143 | 2.260251 | 2.330196 | 2.412103 | 2.552212 | 2.712914 | 2.876545 | 3.024163 | 3.112704 | 3.189677 | 3.279429 | 3.359483 |
|  | 70 | 2.141905 | 2.210572 | 2.289573 | 2.359114 | 2.441273 | 2.583822 | 2.75063 | 2.9217 | 3.074983 | 3.166494 | 3.245855 | 3.33822 | 3.420488 |
|  | 80 | 2.184889 | 2.252437 | 2.330727 | 2.400191 | 2.482978 | 2.628665 | 2.802586 | 2.982299 | 3.142327 | 3.237441 | 3.319737 | 3.415349 | 3.500399 |
|  | 90 | 2.204579 | 2.270323 | 2.347069 | 2.415692 | 2.498184 | 2.645415 | 2.824719 | 3.011465 | 3.176821 | 3.27469 | 3.359184 | 3.45719 | 3.544268 |
| **R_rostralmiddlefrontal** | 3 | 2.502741 | 2.547457 | 2.598653 | 2.64347 | 2.696076 | 2.786325 | 2.890212 | 2.997284 | 3.095773 | 3.15571 | 3.208326 | 3.270267 | 3.32604 |
|  | 5 | 2.435744 | 2.47985 | 2.530291 | 2.57439 | 2.626081 | 2.71455 | 2.816037 | 2.920549 | 3.016874 | 3.075572 | 3.127141 | 3.187887 | 3.242614 |
|  | 10 | 2.281209 | 2.323943 | 2.372671 | 2.415136 | 2.464731 | 2.549105 | 2.645048 | 2.743647 | 2.834982 | 2.890827 | 2.939984 | 2.997981 | 3.050303 |
|  | 15 | 2.152182 | 2.193925 | 2.241377 | 2.282592 | 2.330546 | 2.411626 | 2.502986 | 2.596679 | 2.683929 | 2.737465 | 2.784682 | 2.840483 | 2.890895 |
|  | 20 | 2.069081 | 2.110672 | 2.157799 | 2.198587 | 2.245861 | 2.325277 | 2.413917 | 2.504628 | 2.589575 | 2.641889 | 2.688124 | 2.74286 | 2.792382 |
|  | 30 | 1.985749 | 2.028548 | 2.076742 | 2.118173 | 2.165834 | 2.244933 | 2.331655 | 2.420022 | 2.503596 | 2.555391 | 2.60133 | 2.655871 | 2.705335 |
|  | 40 | 1.940458 | 1.984765 | 2.034427 | 2.076914 | 2.125538 | 2.205582 | 2.292324 | 2.380329 | 2.463843 | 2.515698 | 2.561724 | 2.616393 | 2.66598 |
|  | 50 | 1.896427 | 1.94148 | 1.991873 | 2.034903 | 2.084062 | 2.164806 | 2.252084 | 2.340244 | 2.423406 | 2.474812 | 2.520302 | 2.574173 | 2.622891 |
|  | 60 | 1.845334 | 1.890359 | 1.940712 | 1.983717 | 2.032881 | 2.113796 | 2.201608 | 2.289902 | 2.372125 | 2.422495 | 2.466814 | 2.519015 | 2.56598 |
|  | 70 | 1.822575 | 1.867946 | 1.918726 | 1.962155 | 2.011905 | 2.094131 | 2.184017 | 2.273961 | 2.356342 | 2.406233 | 2.449815 | 2.500809 | 2.546404 |
|  | 80 | 1.819222 | 1.865363 | 1.91706 | 1.961345 | 2.012195 | 2.096649 | 2.189715 | 2.282366 | 2.365729 | 2.415596 | 2.458827 | 2.509058 | 2.55368 |
|  | 90 | 1.827735 | 1.87499 | 1.927987 | 1.97346 | 2.025799 | 2.113149 | 2.210173 | 2.306241 | 2.391122 | 2.441264 | 2.484397 | 2.534166 | 2.57809 |
| **R_superiorfrontal** | 3 | 2.779173 | 2.837784 | 2.902397 | 2.956825 | 3.018215 | 3.11736 | 3.222381 | 3.325932 | 3.42106 | 3.478721 | 3.529068 | 3.587907 | 3.640422 |
|  | 5 | 2.733793 | 2.791752 | 2.855647 | 2.909469 | 2.970172 | 3.068199 | 3.172016 | 3.274397 | 3.3685 | 3.425561 | 3.475394 | 3.533645 | 3.585645 |
|  | 10 | 2.622469 | 2.678808 | 2.740916 | 2.793229 | 2.852225 | 2.94747 | 3.048294 | 3.147769 | 3.239322 | 3.294887 | 3.343441 | 3.400226 | 3.450943 |
|  | 15 | 2.515095 | 2.569843 | 2.630198 | 2.681032 | 2.738355 | 2.830877 | 2.928773 | 3.025404 | 3.114458 | 3.168555 | 3.215854 | 3.271202 | 3.320658 |
|  | 20 | 2.426943 | 2.480471 | 2.53948 | 2.58918 | 2.645219 | 2.735648 | 2.831286 | 2.925733 | 3.012892 | 3.065887 | 3.112249 | 3.166529 | 3.215056 |
|  | 30 | 2.326817 | 2.379497 | 2.437575 | 2.486488 | 2.541631 | 2.630576 | 2.724566 | 2.817475 | 2.903449 | 2.955821 | 3.001693 | 3.055457 | 3.103571 |
|  | 40 | 2.261995 | 2.314559 | 2.372514 | 2.421322 | 2.476342 | 2.565054 | 2.658721 | 2.751407 | 2.837413 | 2.889905 | 2.935936 | 2.989947 | 3.038334 |
|  | 50 | 2.203939 | 2.256499 | 2.314457 | 2.363271 | 2.418291 | 2.506975 | 2.600542 | 2.693228 | 2.779478 | 2.832221 | 2.878529 | 2.932927 | 2.981714 |
|  | 60 | 2.145999 | 2.198516 | 2.256436 | 2.305222 | 2.360208 | 2.44881 | 2.542227 | 2.634869 | 2.721326 | 2.7743 | 2.82087 | 2.875639 | 2.924812 |
|  | 70 | 2.110867 | 2.163867 | 2.222333 | 2.271582 | 2.327092 | 2.416517 | 2.51074 | 2.604293 | 2.691854 | 2.745615 | 2.792936 | 2.848656 | 2.89874 |
|  | 80 | 2.064121 | 2.117284 | 2.175945 | 2.225367 | 2.281072 | 2.370796 | 2.465279 | 2.559206 | 2.64738 | 2.701628 | 2.749441 | 2.80581 | 2.856538 |
|  | 90 | 2.007637 | 2.060669 | 2.119203 | 2.168527 | 2.224127 | 2.313668 | 2.40791 | 2.501717 | 2.590048 | 2.644507 | 2.692572 | 2.749308 | 2.800429 |
| **R_superiorparietal** | 3 | 2.229755 | 2.275453 | 2.326571 | 2.370229 | 2.420123 | 2.502145 | 2.590948 | 2.680549 | 2.764752 | 2.816668 | 2.862534 | 2.916769 | 2.965748 |
|  | 5 | 2.181733 | 2.226981 | 2.27756 | 2.320726 | 2.370026 | 2.450994 | 2.538551 | 2.626789 | 2.709617 | 2.76064 | 2.805691 | 2.858929 | 2.906976 |
|  | 10 | 2.065352 | 2.109504 | 2.158763 | 2.200727 | 2.248568 | 2.32695 | 2.411447 | 2.496337 | 2.575793 | 2.624631 | 2.667685 | 2.718482 | 2.764254 |
|  | 15 | 1.95455 | 1.99772 | 2.045775 | 2.086626 | 2.133102 | 2.209029 | 2.290588 | 2.372236 | 2.448405 | 2.495105 | 2.536202 | 2.584604 | 2.628139 |
|  | 20 | 1.869476 | 1.912535 | 1.960295 | 2.000757 | 2.046638 | 2.12126 | 2.200973 | 2.280335 | 2.353998 | 2.398989 | 2.438476 | 2.484858 | 2.526464 |
|  | 30 | 1.787425 | 1.834201 | 1.885358 | 1.928132 | 1.976041 | 2.052692 | 2.132957 | 2.211347 | 2.282854 | 2.32597 | 2.363478 | 2.407152 | 2.44599 |
|  | 40 | 1.735354 | 1.785807 | 1.840361 | 1.885506 | 1.935598 | 2.014771 | 2.096503 | 2.175273 | 2.246298 | 2.288766 | 2.325502 | 2.368044 | 2.405671 |
|  | 50 | 1.707027 | 1.756874 | 1.811032 | 1.856046 | 1.90619 | 1.985853 | 2.068582 | 2.148763 | 2.221423 | 2.265026 | 2.302837 | 2.346726 | 2.385635 |
|  | 60 | 1.682535 | 1.729088 | 1.780362 | 1.823516 | 1.872146 | 1.950582 | 2.033518 | 2.115292 | 2.190544 | 2.236213 | 2.276117 | 2.322786 | 2.364465 |
|  | 70 | 1.666059 | 1.710299 | 1.75956 | 1.801447 | 1.849113 | 1.927007 | 2.010702 | 2.094539 | 2.17282 | 2.220846 | 2.263126 | 2.312943 | 2.357768 |
|  | 80 | 1.612174 | 1.653598 | 1.70015 | 1.740085 | 1.785919 | 1.861705 | 1.944349 | 2.02838 | 2.107962 | 2.157315 | 2.201095 | 2.253075 | 2.300209 |
|  | 90 | 1.522901 | 1.560881 | 1.60392 | 1.641144 | 1.684211 | 1.75623 | 1.835913 | 1.918169 | 1.997222 | 2.046811 | 2.091161 | 2.144258 | 2.192819 |
| **R_superiortemporal** | 3 | 2.510083 | 2.644494 | 2.7641 | 2.848969 | 2.932577 | 3.049679 | 3.159725 | 3.258849 | 3.344608 | 3.396154 | 3.441773 | 3.496608 | 3.54763 |
|  | 5 | 2.505821 | 2.62983 | 2.742475 | 2.82373 | 2.904799 | 3.019848 | 3.129186 | 3.228215 | 3.313913 | 3.365294 | 3.410631 | 3.464913 | 3.515182 |
|  | 10 | 2.489702 | 2.591622 | 2.688986 | 2.762028 | 2.837142 | 2.9472 | 3.05479 | 3.153761 | 3.239722 | 3.291093 | 3.336189 | 3.389795 | 3.438987 |
|  | 15 | 2.475215 | 2.558144 | 2.641402 | 2.706372 | 2.77532 | 2.879919 | 2.985642 | 3.085084 | 3.17243 | 3.224794 | 3.270744 | 3.325244 | 3.375064 |
|  | 20 | 2.460211 | 2.528843 | 2.600631 | 2.658606 | 2.721934 | 2.821319 | 2.925345 | 3.025809 | 3.11555 | 3.169794 | 3.217565 | 3.274338 | 3.326268 |
|  | 30 | 2.396916 | 2.457055 | 2.521901 | 2.575661 | 2.635715 | 2.732462 | 2.836348 | 2.938328 | 3.029984 | 3.085335 | 3.133909 | 3.191304 | 3.24339 |
|  | 40 | 2.30593 | 2.372773 | 2.444006 | 2.5024 | 2.566911 | 2.669161 | 2.776458 | 2.878992 | 2.96854 | 3.021336 | 3.066838 | 3.11958 | 3.166463 |
|  | 50 | 2.232836 | 2.302048 | 2.375761 | 2.436133 | 2.502741 | 2.608022 | 2.7179 | 2.822048 | 2.91208 | 2.964666 | 3.009652 | 3.061366 | 3.106921 |
|  | 60 | 2.18904 | 2.25031 | 2.317313 | 2.373483 | 2.43675 | 2.539363 | 2.649586 | 2.756666 | 2.850984 | 2.906716 | 2.954717 | 3.010222 | 3.059361 |
|  | 70 | 2.129233 | 2.185595 | 2.24828 | 2.301647 | 2.362615 | 2.463337 | 2.573882 | 2.683383 | 2.781364 | 2.83986 | 2.890565 | 2.949534 | 3.002015 |
|  | 80 | 2.025463 | 2.085732 | 2.152175 | 2.208269 | 2.271837 | 2.375678 | 2.488005 | 2.597613 | 2.694305 | 2.75141 | 2.800534 | 2.857226 | 2.907282 |
|  | 90 | 1.898035 | 1.96899 | 2.045074 | 2.107732 | 2.177154 | 2.2873 | 2.402404 | 2.511134 | 2.604405 | 2.65841 | 2.704259 | 2.756499 | 2.802047 |
| **R_supramarginal** | 3 | 2.536959 | 2.61554 | 2.694838 | 2.756959 | 2.823054 | 2.92346 | 3.024697 | 3.119153 | 3.201046 | 3.249496 | 3.291547 | 3.340803 | 3.385205 |
|  | 5 | 2.496353 | 2.572623 | 2.6499 | 2.710641 | 2.775452 | 2.874256 | 2.974298 | 3.068026 | 3.149603 | 3.198011 | 3.240116 | 3.289546 | 3.334208 |
|  | 10 | 2.397409 | 2.468237 | 2.540694 | 2.598108 | 2.659798 | 2.754679 | 2.851774 | 2.943715 | 3.024552 | 3.072902 | 3.115195 | 3.165139 | 3.210544 |
|  | 15 | 2.304958 | 2.370808 | 2.438787 | 2.493074 | 2.551809 | 2.642952 | 2.737253 | 2.82756 | 2.907832 | 2.956256 | 2.998879 | 3.049541 | 3.095915 |
|  | 20 | 2.231229 | 2.293019 | 2.357326 | 2.409047 | 2.465368 | 2.553514 | 2.645708 | 2.735012 | 2.815295 | 2.864165 | 2.907468 | 2.959297 | 3.00709 |
|  | 30 | 2.149556 | 2.207496 | 2.268338 | 2.317674 | 2.371819 | 2.457493 | 2.548466 | 2.638132 | 2.720248 | 2.771016 | 2.816532 | 2.871704 | 2.923279 |
|  | 40 | 2.091518 | 2.148647 | 2.208719 | 2.257507 | 2.311152 | 2.396344 | 2.487412 | 2.578064 | 2.662118 | 2.714678 | 2.762234 | 2.820468 | 2.875521 |
|  | 50 | 2.029904 | 2.086573 | 2.146111 | 2.194447 | 2.247605 | 2.332147 | 2.422919 | 2.514011 | 2.599431 | 2.653429 | 2.702727 | 2.763709 | 2.822021 |
|  | 60 | 1.977658 | 2.034221 | 2.093536 | 2.141625 | 2.194475 | 2.27857 | 2.369179 | 2.460809 | 2.547719 | 2.603287 | 2.654502 | 2.71855 | 2.780553 |
|  | 70 | 1.932198 | 1.990218 | 2.050699 | 2.099487 | 2.152888 | 2.237533 | 2.328584 | 2.420936 | 2.509189 | 2.566104 | 2.618964 | 2.685672 | 2.750931 |
|  | 80 | 1.861271 | 1.921381 | 1.983431 | 2.033061 | 2.086998 | 2.17182 | 2.262494 | 2.354354 | 2.442489 | 2.49968 | 2.553118 | 2.621056 | 2.688101 |
|  | 90 | 1.794935 | 1.858029 | 1.922396 | 1.973353 | 2.028251 | 2.113752 | 2.204411 | 2.296028 | 2.384226 | 2.441802 | 2.495923 | 2.565246 | 2.634264 |
| **R_frontalpole** | 3 | 2.509697 | 2.620112 | 2.743264 | 2.848542 | 2.969488 | 3.171453 | 3.397416 | 3.627343 | 3.839806 | 3.97034 | 4.086065 | 4.224012 | 4.350146 |
|  | 5 | 2.433525 | 2.541099 | 2.661055 | 2.763591 | 2.881384 | 3.078119 | 3.298358 | 3.5227 | 3.730309 | 3.858043 | 3.971416 | 4.106736 | 4.23065 |
|  | 10 | 2.264459 | 2.365823 | 2.478779 | 2.575289 | 2.686144 | 2.871352 | 3.078973 | 3.291037 | 3.48804 | 3.609702 | 3.718017 | 3.847752 | 3.967015 |
|  | 15 | 2.146838 | 2.244216 | 2.35263 | 2.445202 | 2.551503 | 2.729128 | 2.928504 | 3.132713 | 3.323189 | 3.44129 | 3.546786 | 3.67362 | 3.790714 |
|  | 20 | 2.077271 | 2.172511 | 2.278457 | 2.368872 | 2.472669 | 2.646147 | 2.841125 | 3.041388 | 3.228941 | 3.345692 | 3.450328 | 3.576601 | 3.693672 |
|  | 30 | 2.002674 | 2.094168 | 2.19606 | 2.283125 | 2.383222 | 2.550937 | 2.740211 | 2.93567 | 3.119882 | 3.235193 | 3.338988 | 3.464845 | 3.582142 |
|  | 40 | 1.971767 | 2.059115 | 2.156965 | 2.241025 | 2.338142 | 2.501869 | 2.687896 | 2.881032 | 3.063676 | 3.178182 | 3.281306 | 3.406362 | 3.522873 |
|  | 50 | 1.947478 | 2.031336 | 2.126043 | 2.208002 | 2.303315 | 2.465309 | 2.650912 | 2.844731 | 3.028507 | 3.14374 | 3.247427 | 3.372947 | 3.489592 |
|  | 60 | 1.880124 | 1.960008 | 2.051 | 2.130359 | 2.223299 | 2.382643 | 2.566896 | 2.760593 | 2.944875 | 3.060502 | 3.164484 | 3.290181 | 3.406722 |
|  | 70 | 1.846258 | 1.924704 | 2.014772 | 2.093901 | 2.187196 | 2.348525 | 2.536847 | 2.736316 | 2.926997 | 3.046887 | 3.154765 | 3.285156 | 3.40595 |
|  | 80 | 1.824844 | 1.902991 | 1.993341 | 2.073234 | 2.168006 | 2.333217 | 2.52789 | 2.735796 | 2.935783 | 3.061982 | 3.17576 | 3.313485 | 3.441204 |
|  | 90 | 1.784983 | 1.862163 | 1.95195 | 2.031812 | 2.127085 | 2.294449 | 2.493511 | 2.707985 | 2.915805 | 3.047584 | 3.16675 | 3.311388 | 3.44584 |
| **R_temporalpole** | 3 | 2.358135 | 2.547963 | 2.740953 | 2.893334 | 3.056514 | 3.305488 | 3.555197 | 3.783118 | 3.973395 | 4.081584 | 4.172365 | 4.274657 | 4.362885 |
|  | 5 | 2.378382 | 2.568717 | 2.762249 | 2.91507 | 3.07873 | 3.328447 | 3.578904 | 3.807504 | 3.998343 | 4.106849 | 4.197895 | 4.300482 | 4.388964 |
|  | 10 | 2.430169 | 2.621617 | 2.816365 | 2.970191 | 3.134957 | 3.386399 | 3.638608 | 3.86881 | 4.06098 | 4.170238 | 4.261911 | 4.365202 | 4.454287 |
|  | 15 | 2.48391 | 2.676118 | 2.871761 | 3.026361 | 3.192006 | 3.444857 | 3.698533 | 3.930096 | 4.123409 | 4.233317 | 4.325535 | 4.429438 | 4.519049 |
|  | 20 | 2.532982 | 2.724801 | 2.920233 | 3.074771 | 3.240427 | 3.493416 | 3.747331 | 3.979172 | 4.172744 | 4.282807 | 4.375157 | 4.479212 | 4.568954 |
|  | 30 | 2.581554 | 2.766879 | 2.956218 | 3.106238 | 3.267288 | 3.513611 | 3.761173 | 3.987422 | 4.176438 | 4.283945 | 4.374168 | 4.475839 | 4.563536 |
|  | 40 | 2.577073 | 2.757783 | 2.942467 | 3.088823 | 3.24595 | 3.486272 | 3.727775 | 3.948446 | 4.132765 | 4.237583 | 4.325538 | 4.424644 | 4.510118 |
|  | 50 | 2.542432 | 2.726487 | 2.913782 | 3.061719 | 3.220133 | 3.461728 | 3.703803 | 3.924486 | 4.108493 | 4.213018 | 4.300667 | 4.399367 | 4.484441 |
|  | 60 | 2.441802 | 2.637818 | 2.835156 | 2.989766 | 3.154301 | 3.403544 | 3.651636 | 3.876651 | 4.063576 | 4.169513 | 4.258223 | 4.35799 | 4.443884 |
|  | 70 | 2.267686 | 2.486138 | 2.702065 | 2.868895 | 3.044562 | 3.307686 | 3.566778 | 3.799867 | 3.992381 | 4.101099 | 4.191946 | 4.293921 | 4.381564 |
|  | 80 | 2.036931 | 2.292854 | 2.539856 | 2.726932 | 2.920894 | 3.20671 | 3.483844 | 3.730364 | 3.93238 | 4.045926 | 4.140541 | 4.246483 | 4.337328 |
|  | 90 | 1.775265 | 2.074892 | 2.360133 | 2.572099 | 2.788175 | 3.10068 | 3.398361 | 3.65979 | 3.872157 | 3.990903 | 4.089551 | 4.199713 | 4.293947 |
| **R_transversetemporal** | 3 | 2.232135 | 2.305492 | 2.388006 | 2.458973 | 2.540793 | 2.677511 | 2.829415 | 2.981318 | 3.118036 | 3.199856 | 3.270823 | 3.353337 | 3.426694 |
|  | 5 | 2.1866 | 2.260042 | 2.342653 | 2.413702 | 2.495618 | 2.632496 | 2.784577 | 2.936658 | 3.073535 | 3.155451 | 3.226501 | 3.309112 | 3.382554 |
|  | 10 | 2.074914 | 2.148571 | 2.231423 | 2.30268 | 2.384836 | 2.522113 | 2.674639 | 2.827164 | 2.964442 | 3.046597 | 3.117855 | 3.200707 | 3.274364 |
|  | 15 | 1.972705 | 2.046577 | 2.129671 | 2.201137 | 2.283532 | 2.421211 | 2.574182 | 2.727153 | 2.864832 | 2.947227 | 3.018693 | 3.101787 | 3.175659 |
|  | 20 | 1.89192 | 1.966008 | 2.049345 | 2.121019 | 2.203655 | 2.341736 | 2.495153 | 2.648571 | 2.786652 | 2.869288 | 2.940962 | 3.024299 | 3.098387 |
|  | 30 | 1.789145 | 1.863667 | 1.947491 | 2.019585 | 2.102704 | 2.241593 | 2.395908 | 2.550224 | 2.689113 | 2.772232 | 2.844326 | 2.92815 | 3.002671 |
|  | 40 | 1.707223 | 1.782183 | 1.8665 | 1.939018 | 2.022626 | 2.162331 | 2.317554 | 2.472777 | 2.612483 | 2.696091 | 2.768608 | 2.852926 | 2.927885 |
|  | 50 | 1.652854 | 1.728258 | 1.813076 | 1.886024 | 1.970128 | 2.110662 | 2.266806 | 2.42295 | 2.563485 | 2.647589 | 2.720537 | 2.805355 | 2.880759 |
|  | 60 | 1.608432 | 1.684287 | 1.769612 | 1.842995 | 1.927602 | 2.068976 | 2.226053 | 2.38313 | 2.524504 | 2.609111 | 2.682495 | 2.767819 | 2.843674 |
|  | 70 | 1.551865 | 1.628175 | 1.714012 | 1.787836 | 1.872951 | 2.015174 | 2.173193 | 2.331213 | 2.473436 | 2.55855 | 2.632375 | 2.718211 | 2.794522 |
|  | 80 | 1.457982 | 1.534753 | 1.621108 | 1.695378 | 1.781006 | 1.924088 | 2.083061 | 2.242035 | 2.385116 | 2.470745 | 2.545015 | 2.63137 | 2.708141 |
|  | 90 | 1.33713 | 1.414366 | 1.501244 | 1.575963 | 1.66211 | 1.806058 | 1.965994 | 2.125931 | 2.269878 | 2.356025 | 2.430745 | 2.517623 | 2.594858 |
| **R_insula** | 3 | 2.995009 | 3.062811 | 3.134866 | 3.193874 | 3.259136 | 3.363174 | 3.474045 | 3.582871 | 3.681394 | 3.741494 | 3.794738 | 3.85838 | 3.916918 |
|  | 5 | 2.954926 | 3.022212 | 3.093689 | 3.152207 | 3.216914 | 3.320058 | 3.430002 | 3.537993 | 3.635868 | 3.695637 | 3.748639 | 3.812057 | 3.87046 |
|  | 10 | 2.861343 | 2.927466 | 2.997633 | 3.055034 | 3.118474 | 3.219573 | 3.327398 | 3.433496 | 3.529926 | 3.588983 | 3.641479 | 3.704464 | 3.762648 |
|  | 15 | 2.783224 | 2.848526 | 2.917745 | 2.974319 | 3.036809 | 3.136363 | 3.242596 | 3.347314 | 3.442768 | 3.501402 | 3.553653 | 3.616524 | 3.674791 |
|  | 20 | 2.720244 | 2.785187 | 2.85392 | 2.91003 | 2.971952 | 3.070533 | 3.175741 | 3.279601 | 3.374529 | 3.433007 | 3.485246 | 3.548282 | 3.60689 |
|  | 30 | 2.619267 | 2.685226 | 2.754611 | 2.810962 | 2.872886 | 2.971024 | 3.075394 | 3.178376 | 3.272738 | 3.331084 | 3.383391 | 3.446785 | 3.506029 |
|  | 40 | 2.533741 | 2.602779 | 2.674652 | 2.732503 | 2.795592 | 2.894699 | 2.99922 | 3.101848 | 3.195773 | 3.253916 | 3.306139 | 3.369601 | 3.429115 |
|  | 50 | 2.462998 | 2.53696 | 2.612901 | 2.673306 | 2.738519 | 2.83977 | 2.945333 | 3.048216 | 3.142088 | 3.2002 | 3.252457 | 3.316089 | 3.375932 |
|  | 60 | 2.40843 | 2.489219 | 2.570789 | 2.634761 | 2.70301 | 2.807537 | 2.915079 | 3.018997 | 3.113478 | 3.171962 | 3.224617 | 3.288872 | 3.349484 |
|  | 70 | 2.345933 | 2.434953 | 2.523069 | 2.591051 | 2.662605 | 2.770529 | 2.879957 | 2.984735 | 3.079665 | 3.138441 | 3.191443 | 3.25629 | 3.317675 |
|  | 80 | 2.277268 | 2.376807 | 2.472975 | 2.54573 | 2.621103 | 2.732807 | 2.84421 | 2.9498 | 3.045106 | 3.104138 | 3.157467 | 3.222906 | 3.285094 |
|  | 90 | 2.198451 | 2.311725 | 2.417939 | 2.496413 | 2.576203 | 2.692063 | 2.805449 | 2.911702 | 3.007214 | 3.066404 | 3.119985 | 3.185947 | 3.248898 |

**Table S7. Centile Values for Cortical Thickness in Females**

|  | **age** | **C0.4** | **C1** | **C2.5** | **C5** | **C10** | **C25** | **C50** | **C75** | **C90** | **C95** | **C97.5** | **C99** | **C99.6** |
| --- | --- | --- | --- | --- | --- | --- | --- | --- | --- | --- | --- | --- | --- | --- |
| **LThickness** | 5 | 2.555366 | 2.599829 | 2.645627 | 2.682174 | 2.721779 | 2.783714 | 2.849191 | 2.91438 | 2.975519 | 3.014349 | 3.049998 | 3.094427 | 3.137301 |
|  | 10 | 2.466464 | 2.504508 | 2.544944 | 2.578088 | 2.614812 | 2.673618 | 2.736897 | 2.799976 | 2.85824 | 2.894452 | 2.927021 | 2.966609 | 3.00371 |
|  | 15 | 2.379005 | 2.413151 | 2.45019 | 2.481082 | 2.515803 | 2.572267 | 2.633736 | 2.695087 | 2.751231 | 2.785658 | 2.816223 | 2.852794 | 2.886431 |
|  | 20 | 2.314381 | 2.346261 | 2.381305 | 2.410864 | 2.444401 | 2.49949 | 2.559932 | 2.620326 | 2.675289 | 2.708711 | 2.738144 | 2.77301 | 2.804701 |
|  | 30 | 2.239063 | 2.269048 | 2.302479 | 2.331021 | 2.363731 | 2.418054 | 2.478168 | 2.538334 | 2.592794 | 2.625629 | 2.654305 | 2.687925 | 2.718107 |
|  | 40 | 2.192684 | 2.222321 | 2.255558 | 2.284074 | 2.316891 | 2.371635 | 2.432428 | 2.493314 | 2.548303 | 2.581339 | 2.610091 | 2.643656 | 2.673633 |
|  | 50 | 2.159525 | 2.1895 | 2.223189 | 2.252146 | 2.285517 | 2.34127 | 2.403244 | 2.4653 | 2.521269 | 2.554833 | 2.583996 | 2.617971 | 2.648244 |
|  | 60 | 2.130022 | 2.160662 | 2.195111 | 2.224728 | 2.258865 | 2.315893 | 2.379254 | 2.442636 | 2.49972 | 2.533906 | 2.563576 | 2.598099 | 2.628815 |
|  | 70 | 2.069397 | 2.100419 | 2.135278 | 2.165228 | 2.199727 | 2.2573 | 2.321173 | 2.384958 | 2.442301 | 2.476592 | 2.506322 | 2.540875 | 2.571581 |
|  | 80 | 2.02281 | 2.054506 | 2.090083 | 2.120619 | 2.155754 | 2.214302 | 2.279127 | 2.343725 | 2.40168 | 2.436285 | 2.466252 | 2.501043 | 2.531926 |
|  | 90 | 1.979117 | 2.011576 | 2.047963 | 2.079155 | 2.115002 | 2.174633 | 2.240511 | 2.306008 | 2.364646 | 2.399603 | 2.429842 | 2.46491 | 2.496005 |
| **RThickness** | 5 | 2.567658 | 2.607334 | 2.650473 | 2.68641 | 2.726608 | 2.791079 | 2.859166 | 2.923954 | 2.979748 | 3.012099 | 3.039569 | 3.070855 | 3.098108 |
|  | 10 | 2.464401 | 2.500486 | 2.540123 | 2.573452 | 2.611056 | 2.672059 | 2.73737 | 2.800321 | 2.855138 | 2.887168 | 2.914505 | 2.945792 | 2.973175 |
|  | 15 | 2.375155 | 2.408254 | 2.444942 | 2.476055 | 2.511441 | 2.569468 | 2.632421 | 2.693882 | 2.748009 | 2.779891 | 2.807246 | 2.838716 | 2.8664 |
|  | 20 | 2.312366 | 2.343314 | 2.377873 | 2.40739 | 2.441188 | 2.497131 | 2.558534 | 2.619182 | 2.67315 | 2.705179 | 2.732801 | 2.764736 | 2.792968 |
|  | 30 | 2.237487 | 2.266484 | 2.299115 | 2.32719 | 2.359572 | 2.413711 | 2.473906 | 2.534147 | 2.588403 | 2.620889 | 2.649076 | 2.681861 | 2.711018 |
|  | 40 | 2.191798 | 2.220548 | 2.252963 | 2.280906 | 2.313194 | 2.367318 | 2.427701 | 2.488341 | 2.543135 | 2.576023 | 2.604607 | 2.637909 | 2.667576 |
|  | 50 | 2.161125 | 2.190242 | 2.223093 | 2.251428 | 2.284192 | 2.33916 | 2.400557 | 2.462289 | 2.518131 | 2.551677 | 2.580849 | 2.614856 | 2.645167 |
|  | 60 | 2.131637 | 2.161549 | 2.195286 | 2.224379 | 2.258008 | 2.314406 | 2.377366 | 2.440634 | 2.497837 | 2.532187 | 2.56205 | 2.596852 | 2.627865 |
|  | 70 | 2.06734 | 2.0983 | 2.133129 | 2.163089 | 2.197635 | 2.255372 | 2.319537 | 2.383719 | 2.441498 | 2.476083 | 2.506084 | 2.54097 | 2.571989 |
|  | 80 | 2.0203 | 2.053075 | 2.089805 | 2.121281 | 2.157443 | 2.217571 | 2.283959 | 2.349924 | 2.408945 | 2.444114 | 2.474528 | 2.509787 | 2.541041 |
|  | 90 | 1.974998 | 2.009942 | 2.048925 | 2.082187 | 2.120241 | 2.183151 | 2.252104 | 2.320118 | 2.380572 | 2.416423 | 2.447324 | 2.483034 | 2.514588 |
| **L_bankssts** | 5 | 2.401344 | 2.471978 | 2.549761 | 2.615123 | 2.688548 | 2.806071 | 2.928521 | 3.0507 | 3.167454 | 3.240148 | 3.304695 | 3.381307 | 3.450688 |
|  | 10 | 2.291201 | 2.3588 | 2.433316 | 2.496004 | 2.56652 | 2.679645 | 2.797945 | 2.915938 | 3.0282 | 3.097895 | 3.159672 | 3.232882 | 3.299086 |
|  | 15 | 2.183187 | 2.247799 | 2.319093 | 2.37914 | 2.446774 | 2.555526 | 2.669666 | 2.783466 | 2.891266 | 2.957999 | 3.017047 | 3.086914 | 3.150004 |
|  | 20 | 2.10439 | 2.166865 | 2.235869 | 2.294053 | 2.359676 | 2.465437 | 2.576836 | 2.687861 | 2.792574 | 2.857211 | 2.914305 | 2.981753 | 3.042572 |
|  | 30 | 2.029251 | 2.089873 | 2.156963 | 2.213661 | 2.277777 | 2.381579 | 2.491698 | 2.601357 | 2.703889 | 2.766817 | 2.822207 | 2.88744 | 2.946095 |
|  | 40 | 1.982626 | 2.042274 | 2.108415 | 2.164435 | 2.227951 | 2.331246 | 2.4416 | 2.551399 | 2.653179 | 2.715289 | 2.769769 | 2.833732 | 2.891081 |
|  | 50 | 1.9479 | 2.006999 | 2.072656 | 2.128391 | 2.191745 | 2.295242 | 2.406576 | 2.517253 | 2.61897 | 2.680685 | 2.734632 | 2.797773 | 2.854225 |
|  | 60 | 1.914275 | 1.972937 | 2.038232 | 2.09378 | 2.157084 | 2.260958 | 2.373461 | 2.485195 | 2.587009 | 2.648429 | 2.701933 | 2.764361 | 2.82002 |
|  | 70 | 1.863147 | 1.920903 | 1.98531 | 2.04022 | 2.102956 | 2.206351 | 2.319087 | 2.430942 | 2.532002 | 2.592616 | 2.645237 | 2.706445 | 2.760862 |
|  | 80 | 1.822114 | 1.879309 | 1.943208 | 1.9978 | 2.060329 | 2.163833 | 2.277431 | 2.390025 | 2.490893 | 2.551045 | 2.603084 | 2.663429 | 2.71693 |
|  | 90 | 1.784656 | 1.841413 | 1.904936 | 1.959321 | 2.021767 | 2.12558 | 2.240254 | 2.353793 | 2.45465 | 2.514451 | 2.566006 | 2.625609 | 2.678304 |
| **L_caudalanteriorcingulate** | 5 | 2.364944 | 2.464126 | 2.571899 | 2.661909 | 2.763195 | 2.928071 | 3.107633 | 3.286686 | 3.450201 | 3.550226 | 3.638844 | 3.74461 | 3.841616 |
|  | 10 | 2.280404 | 2.374022 | 2.476288 | 2.562108 | 2.659097 | 2.817854 | 2.99184 | 3.166288 | 3.326277 | 3.424404 | 3.51148 | 3.61555 | 3.711117 |
|  | 15 | 2.210971 | 2.299919 | 2.397575 | 2.479903 | 2.573335 | 2.727096 | 2.896656 | 3.067601 | 3.225057 | 3.321898 | 3.407977 | 3.511008 | 3.605749 |
|  | 20 | 2.153444 | 2.238529 | 2.332379 | 2.411836 | 2.502363 | 2.652099 | 2.818196 | 2.986532 | 3.142233 | 3.238254 | 3.323743 | 3.426221 | 3.520579 |
|  | 30 | 2.066977 | 2.14669 | 2.235288 | 2.310827 | 2.39745 | 2.541942 | 2.703799 | 2.869275 | 3.023387 | 3.118843 | 3.204054 | 3.306436 | 3.400899 |
|  | 40 | 2.02022 | 2.09768 | 2.184275 | 2.258501 | 2.344039 | 2.487637 | 2.64968 | 2.81641 | 2.972449 | 3.069384 | 3.15606 | 3.260348 | 3.356678 |
|  | 50 | 1.996717 | 2.074851 | 2.162499 | 2.237859 | 2.324943 | 2.471634 | 2.637759 | 2.809131 | 2.969732 | 3.069533 | 3.158759 | 3.266066 | 3.36511 |
|  | 60 | 1.987518 | 2.070008 | 2.162489 | 2.241941 | 2.333663 | 2.487877 | 2.66196 | 2.840753 | 3.00744 | 3.110547 | 3.202398 | 3.312425 | 3.41354 |
|  | 70 | 1.95196 | 2.041251 | 2.140853 | 2.225999 | 2.323799 | 2.487023 | 2.669413 | 2.85463 | 3.025335 | 3.129962 | 3.22254 | 3.332656 | 3.433101 |
|  | 80 | 1.929636 | 2.029898 | 2.140735 | 2.234675 | 2.341684 | 2.518224 | 2.712556 | 2.906826 | 3.083202 | 3.190062 | 3.283838 | 3.394444 | 3.494459 |
|  | 90 | 1.93121 | 2.048016 | 2.175487 | 2.282258 | 2.402561 | 2.598154 | 2.809612 | 3.017252 | 3.202701 | 3.313696 | 3.410278 | 3.523225 | 3.624474 |
| **L_caudalmiddlefrontal** | 5 | 2.342883 | 2.424314 | 2.50759 | 2.573313 | 2.643315 | 2.748589 | 2.851447 | 2.945852 | 3.027539 | 3.075 | 3.11529 | 3.161119 | 3.200955 |
|  | 10 | 2.319931 | 2.390032 | 2.463923 | 2.523723 | 2.588782 | 2.689135 | 2.789927 | 2.884548 | 2.967846 | 3.016821 | 3.058719 | 3.106727 | 3.148753 |
|  | 15 | 2.28307 | 2.344053 | 2.409907 | 2.464345 | 2.524696 | 2.620012 | 2.718361 | 2.812854 | 2.897592 | 2.948069 | 2.991631 | 3.041965 | 3.08639 |
|  | 20 | 2.242529 | 2.296326 | 2.355556 | 2.405397 | 2.461566 | 2.552211 | 2.648185 | 2.742561 | 2.828849 | 2.880979 | 2.926402 | 2.979383 | 3.026582 |
|  | 30 | 2.183634 | 2.229547 | 2.28127 | 2.325761 | 2.376982 | 2.462109 | 2.555658 | 2.650912 | 2.740664 | 2.796144 | 2.84527 | 2.903509 | 2.956253 |
|  | 40 | 2.140052 | 2.183774 | 2.233417 | 2.276452 | 2.326387 | 2.410331 | 2.503998 | 2.600673 | 2.692744 | 2.750149 | 2.801302 | 2.862344 | 2.918006 |
|  | 50 | 2.098609 | 2.143726 | 2.194839 | 2.239056 | 2.29026 | 2.376116 | 2.471622 | 2.569624 | 2.662208 | 2.719581 | 2.770489 | 2.830982 | 2.88591 |
|  | 60 | 2.04561 | 2.093741 | 2.14788 | 2.194397 | 2.247908 | 2.33682 | 2.434599 | 2.533506 | 2.625449 | 2.681735 | 2.731257 | 2.789606 | 2.842136 |
|  | 70 | 1.981101 | 2.031689 | 2.088258 | 2.13659 | 2.191896 | 2.283145 | 2.382637 | 2.482169 | 2.573527 | 2.628933 | 2.677373 | 2.734092 | 2.784838 |
|  | 80 | 1.937197 | 1.99015 | 2.049129 | 2.099338 | 2.156601 | 2.250675 | 2.352742 | 2.454113 | 2.546329 | 2.601896 | 2.650268 | 2.706674 | 2.756936 |
|  | 90 | 1.912281 | 1.96761 | 2.029093 | 2.081327 | 2.140792 | 2.238278 | 2.343809 | 2.448153 | 2.542471 | 2.599049 | 2.648158 | 2.705265 | 2.756016 |
| **L_cuneus** | 5 | 1.857743 | 1.898959 | 1.946338 | 1.987966 | 2.036989 | 2.121427 | 2.219062 | 2.320872 | 2.41621 | 2.474999 | 2.527065 | 2.588887 | 2.645029 |
|  | 10 | 1.71678 | 1.755325 | 1.799697 | 1.838739 | 1.884784 | 1.964261 | 2.056426 | 2.152831 | 2.243383 | 2.299352 | 2.349005 | 2.408061 | 2.461785 |
|  | 15 | 1.613987 | 1.650628 | 1.692869 | 1.73009 | 1.774052 | 1.850099 | 1.938547 | 2.031361 | 2.118813 | 2.172997 | 2.221151 | 2.278526 | 2.330816 |
|  | 20 | 1.564689 | 1.600543 | 1.641937 | 1.678464 | 1.72167 | 1.796572 | 1.883946 | 1.97593 | 2.062876 | 2.116881 | 2.164961 | 2.222352 | 2.274756 |
|  | 30 | 1.510156 | 1.545065 | 1.585477 | 1.621233 | 1.663646 | 1.737474 | 1.824082 | 1.915829 | 2.003092 | 2.057558 | 2.106219 | 2.164515 | 2.217945 |
|  | 40 | 1.484347 | 1.518563 | 1.558265 | 1.593479 | 1.635352 | 1.708513 | 1.794784 | 1.886707 | 1.974648 | 2.029794 | 2.079229 | 2.138662 | 2.193334 |
|  | 50 | 1.461224 | 1.494694 | 1.533619 | 1.568222 | 1.609467 | 1.681791 | 1.767507 | 1.859361 | 1.947754 | 2.003442 | 2.053535 | 2.113976 | 2.169787 |
|  | 60 | 1.44966 | 1.482622 | 1.52104 | 1.55527 | 1.596165 | 1.668131 | 1.753855 | 1.846251 | 1.935695 | 1.992318 | 2.043434 | 2.105341 | 2.162733 |
|  | 70 | 1.431684 | 1.463995 | 1.501735 | 1.535435 | 1.575792 | 1.647065 | 1.732397 | 1.824912 | 1.915023 | 1.972354 | 2.024304 | 2.087469 | 2.146274 |
|  | 80 | 1.412539 | 1.444165 | 1.481182 | 1.514309 | 1.554071 | 1.624543 | 1.709347 | 1.801839 | 1.892496 | 1.950473 | 2.003213 | 2.067606 | 2.12782 |
|  | 90 | 1.369731 | 1.400153 | 1.435834 | 1.467834 | 1.506332 | 1.574806 | 1.65763 | 1.748513 | 1.83817 | 1.895817 | 1.94847 | 2.01304 | 2.073706 |
| **L_entorhinal** | 5 | 2.277819 | 2.415695 | 2.564898 | 2.688831 | 2.827251 | 3.049207 | 3.284073 | 3.508461 | 3.702695 | 3.815815 | 3.912188 | 4.022332 | 4.118627 |
|  | 10 | 2.318771 | 2.449179 | 2.591976 | 2.711814 | 2.846886 | 3.065964 | 3.300839 | 3.527906 | 3.726395 | 3.842769 | 3.942344 | 4.056619 | 4.156921 |
|  | 15 | 2.352732 | 2.476527 | 2.61347 | 2.729447 | 2.861249 | 3.077306 | 3.311848 | 3.541222 | 3.743693 | 3.863208 | 3.965925 | 4.084304 | 4.188636 |
|  | 20 | 2.370339 | 2.488287 | 2.619839 | 2.73209 | 2.860547 | 3.073056 | 3.30629 | 3.536766 | 3.742042 | 3.863978 | 3.969213 | 4.090981 | 4.198721 |
|  | 30 | 2.360185 | 2.470129 | 2.593946 | 2.700556 | 2.823604 | 3.029513 | 3.25871 | 3.488319 | 3.695304 | 3.819319 | 3.926966 | 4.052223 | 4.163662 |
|  | 40 | 2.34955 | 2.457345 | 2.57912 | 2.68428 | 2.805995 | 3.010459 | 3.239143 | 3.469326 | 3.677713 | 3.802951 | 3.911886 | 4.038899 | 4.152126 |
|  | 50 | 2.348903 | 2.459241 | 2.583697 | 2.691017 | 2.81506 | 3.023035 | 3.255088 | 3.488109 | 3.698614 | 3.824929 | 3.934686 | 4.062527 | 4.176376 |
|  | 60 | 2.336728 | 2.453463 | 2.584334 | 2.696539 | 2.825522 | 3.040189 | 3.277531 | 3.513739 | 3.725436 | 3.851745 | 3.961077 | 4.087949 | 4.200522 |
|  | 70 | 2.264416 | 2.390027 | 2.529185 | 2.647201 | 2.781496 | 3.002027 | 3.241944 | 3.477066 | 3.684996 | 3.807894 | 3.913612 | 4.035553 | 4.143115 |
|  | 80 | 2.112771 | 2.250641 | 2.400184 | 2.524668 | 2.663985 | 2.887975 | 3.125765 | 3.353653 | 3.55145 | 3.666866 | 3.765319 | 3.877978 | 3.97659 |
|  | 90 | 1.935754 | 2.091965 | 2.256085 | 2.389146 | 2.534778 | 2.76274 | 2.997756 | 3.217354 | 3.404125 | 3.511641 | 3.602567 | 3.705777 | 3.795421 |
| **L_fusiform** | 5 | 2.421028 | 2.525882 | 2.626087 | 2.701033 | 2.77743 | 2.886513 | 2.987145 | 3.076757 | 3.153859 | 3.198436 | 3.236127 | 3.278809 | 3.315737 |
|  | 10 | 2.402839 | 2.481035 | 2.561009 | 2.624058 | 2.691068 | 2.791355 | 2.888488 | 2.978086 | 3.056993 | 3.103348 | 3.142952 | 3.188246 | 3.227808 |
|  | 15 | 2.369371 | 2.431215 | 2.497354 | 2.551519 | 2.611013 | 2.70372 | 2.797668 | 2.887441 | 2.968505 | 3.016986 | 3.058907 | 3.107412 | 3.150265 |
|  | 20 | 2.338439 | 2.391264 | 2.449251 | 2.49789 | 2.55251 | 2.640143 | 2.732128 | 2.822562 | 2.905924 | 2.956554 | 3.000805 | 3.052557 | 3.098771 |
|  | 30 | 2.274175 | 2.32624 | 2.383565 | 2.431804 | 2.486174 | 2.573934 | 2.666891 | 2.758256 | 2.841682 | 2.892039 | 2.935891 | 2.987013 | 3.032533 |
|  | 40 | 2.216722 | 2.270956 | 2.330364 | 2.380146 | 2.436068 | 2.526043 | 2.621136 | 2.713578 | 2.796407 | 2.845754 | 2.888372 | 2.937673 | 2.981251 |
|  | 50 | 2.205115 | 2.25563 | 2.311793 | 2.359534 | 2.413921 | 2.503186 | 2.599989 | 2.695615 | 2.78175 | 2.833294 | 2.877966 | 2.929839 | 2.975878 |
|  | 60 | 2.199267 | 2.246923 | 2.300572 | 2.346745 | 2.400005 | 2.48903 | 2.587947 | 2.687165 | 2.776982 | 2.830954 | 2.877889 | 2.932596 | 2.981347 |
|  | 70 | 2.150393 | 2.19907 | 2.253799 | 2.300863 | 2.355136 | 2.445914 | 2.546978 | 2.647641 | 2.73738 | 2.790726 | 2.836802 | 2.890167 | 2.937436 |
|  | 80 | 2.127077 | 2.171939 | 2.223083 | 2.267677 | 2.319835 | 2.408925 | 2.510947 | 2.614616 | 2.707922 | 2.763801 | 2.812334 | 2.868876 | 2.919271 |
|  | 90 | 2.125553 | 2.163055 | 2.206882 | 2.246082 | 2.293164 | 2.3769 | 2.478373 | 2.587108 | 2.68929 | 2.75258 | 2.808913 | 2.87623 | 2.937838 |
| **L_inferiorparietal** | 5 | 2.486679 | 2.552233 | 2.620259 | 2.674827 | 2.7341 | 2.826542 | 2.922768 | 3.015471 | 3.098332 | 3.14853 | 3.192849 | 3.245689 | 3.294213 |
|  | 10 | 2.352434 | 2.411107 | 2.472923 | 2.523155 | 2.578313 | 2.665407 | 2.757121 | 2.846056 | 2.925651 | 2.973775 | 3.016141 | 3.066449 | 3.11241 |
|  | 15 | 2.233847 | 2.286997 | 2.343731 | 2.390351 | 2.442037 | 2.52456 | 2.612397 | 2.698138 | 2.775038 | 2.821494 | 2.862315 | 2.910651 | 2.954641 |
|  | 20 | 2.159905 | 2.209313 | 2.262641 | 2.30689 | 2.356359 | 2.436128 | 2.521873 | 2.606117 | 2.68188 | 2.727648 | 2.767821 | 2.815297 | 2.858386 |
|  | 30 | 2.092466 | 2.137918 | 2.187799 | 2.229797 | 2.277354 | 2.355226 | 2.440242 | 2.524665 | 2.600966 | 2.647086 | 2.687518 | 2.735177 | 2.778266 |
|  | 40 | 2.060078 | 2.103929 | 2.152566 | 2.193906 | 2.241107 | 2.319175 | 2.405275 | 2.49137 | 2.569425 | 2.616614 | 2.65794 | 2.706559 | 2.75039 |
|  | 50 | 2.029722 | 2.072735 | 2.120804 | 2.161937 | 2.209185 | 2.287905 | 2.375389 | 2.463349 | 2.543317 | 2.591693 | 2.634043 | 2.683815 | 2.728611 |
|  | 60 | 1.999275 | 2.041815 | 2.089633 | 2.130764 | 2.178234 | 2.257795 | 2.346782 | 2.4367 | 2.5187 | 2.568369 | 2.611865 | 2.662977 | 2.708952 |
|  | 70 | 1.937168 | 1.978758 | 2.025729 | 2.066308 | 2.113327 | 2.192537 | 2.281647 | 2.372131 | 2.454936 | 2.505188 | 2.549237 | 2.601031 | 2.647633 |
|  | 80 | 1.889257 | 1.930039 | 1.976321 | 2.016489 | 2.063234 | 2.142441 | 2.232178 | 2.323899 | 2.408295 | 2.459699 | 2.504859 | 2.558069 | 2.606034 |
|  | 90 | 1.831781 | 1.871401 | 1.916592 | 1.956001 | 2.00208 | 2.080665 | 2.170427 | 2.262918 | 2.348639 | 2.40112 | 2.447385 | 2.502077 | 2.551538 |
| **L_inferiortemporal** | 5 | 2.441356 | 2.541915 | 2.642847 | 2.721559 | 2.804857 | 2.929941 | 3.052965 | 3.163167 | 3.253685 | 3.304567 | 3.346941 | 3.394343 | 3.43494 |
|  | 10 | 2.425003 | 2.503619 | 2.586581 | 2.653942 | 2.727629 | 2.842633 | 2.960465 | 3.069684 | 3.161804 | 3.214486 | 3.258835 | 3.308944 | 3.352264 |
|  | 15 | 2.392721 | 2.456848 | 2.526872 | 2.585453 | 2.651251 | 2.757403 | 2.870364 | 2.978689 | 3.072648 | 3.127415 | 3.174086 | 3.227431 | 3.274064 |
|  | 20 | 2.360118 | 2.414855 | 2.476007 | 2.52826 | 2.588128 | 2.687301 | 2.796286 | 2.90408 | 3.000136 | 3.057206 | 3.106461 | 3.163455 | 3.21388 |
|  | 30 | 2.318247 | 2.366813 | 2.421902 | 2.46967 | 2.525187 | 2.619006 | 2.724779 | 2.832152 | 2.930147 | 2.989403 | 3.041167 | 3.101781 | 3.156053 |
|  | 40 | 2.275817 | 2.325889 | 2.382426 | 2.431232 | 2.487705 | 2.582546 | 2.688607 | 2.795371 | 2.892047 | 2.950165 | 3.000728 | 3.059698 | 3.112283 |
|  | 50 | 2.253934 | 2.304685 | 2.362014 | 2.411522 | 2.46883 | 2.565126 | 2.672889 | 2.781443 | 2.879805 | 2.938963 | 2.99045 | 3.050515 | 3.104095 |
|  | 60 | 2.231536 | 2.281467 | 2.338241 | 2.387586 | 2.445069 | 2.54253 | 2.652881 | 2.765396 | 2.86851 | 2.931054 | 2.985807 | 3.050058 | 3.107709 |
|  | 70 | 2.185095 | 2.234206 | 2.290461 | 2.339714 | 2.397509 | 2.496538 | 2.610239 | 2.727899 | 2.837262 | 2.904311 | 2.963451 | 3.03338 | 3.096611 |
|  | 80 | 2.1562 | 2.204209 | 2.25977 | 2.308918 | 2.367201 | 2.468623 | 2.587558 | 2.71351 | 2.833277 | 2.908021 | 2.974793 | 3.054782 | 3.128092 |
|  | 90 | 2.124211 | 2.171613 | 2.226962 | 2.276372 | 2.335523 | 2.439939 | 2.564873 | 2.700232 | 2.831983 | 2.915763 | 2.99165 | 3.083884 | 3.169717 |
| **L_isthmuscingulate** | 5 | 2.46208 | 2.519695 | 2.586346 | 2.645281 | 2.715138 | 2.836611 | 2.978902 | 3.129386 | 3.272273 | 3.361336 | 3.440828 | 3.535963 | 3.623063 |
|  | 10 | 2.358683 | 2.415194 | 2.480524 | 2.538249 | 2.606621 | 2.725377 | 2.864262 | 3.010875 | 3.149824 | 3.236302 | 3.313402 | 3.405566 | 3.489844 |
|  | 15 | 2.260337 | 2.315803 | 2.379878 | 2.436452 | 2.503409 | 2.619568 | 2.75519 | 2.898086 | 3.033251 | 3.117244 | 3.192043 | 3.281352 | 3.362919 |
|  | 20 | 2.175122 | 2.22981 | 2.292938 | 2.348632 | 2.414493 | 2.52861 | 2.661615 | 2.80148 | 2.933514 | 3.015432 | 3.088298 | 3.175196 | 3.254462 |
|  | 30 | 2.068177 | 2.12283 | 2.185813 | 2.241285 | 2.306768 | 2.41993 | 2.551336 | 2.688949 | 2.81832 | 2.898323 | 2.969317 | 3.053773 | 3.130616 |
|  | 40 | 2.010799 | 2.066743 | 2.131095 | 2.187665 | 2.254315 | 2.369162 | 2.501995 | 2.640491 | 2.770122 | 2.850012 | 2.92073 | 3.004647 | 3.080799 |
|  | 50 | 1.958962 | 2.016452 | 2.082446 | 2.14034 | 2.208404 | 2.325319 | 2.459963 | 2.59969 | 2.729872 | 2.809814 | 2.880399 | 2.963939 | 3.039546 |
|  | 60 | 1.909397 | 1.968631 | 2.036472 | 2.095847 | 2.16549 | 2.284706 | 2.421365 | 2.562477 | 2.693313 | 2.773358 | 2.843848 | 2.927054 | 3.002153 |
|  | 70 | 1.841005 | 1.901519 | 1.970646 | 2.030992 | 2.10159 | 2.221991 | 2.359321 | 2.500373 | 2.63049 | 2.709789 | 2.779432 | 2.861413 | 2.935202 |
|  | 80 | 1.783806 | 1.846091 | 1.917035 | 1.978791 | 2.050832 | 2.173192 | 2.312008 | 2.453784 | 2.583873 | 2.662838 | 2.731995 | 2.813176 | 2.886038 |
|  | 90 | 1.754969 | 1.82025 | 1.894365 | 1.958674 | 2.033458 | 2.159912 | 2.302541 | 2.447342 | 2.579469 | 2.659338 | 2.729086 | 2.81073 | 2.883798 |
| **L_lateraloccipital** | 5 | 2.118075 | 2.15819 | 2.203675 | 2.243093 | 2.288872 | 2.366122 | 2.453011 | 2.541365 | 2.622459 | 2.671692 | 2.714811 | 2.765424 | 2.810845 |
|  | 10 | 2.013502 | 2.052012 | 2.095673 | 2.133498 | 2.177403 | 2.251402 | 2.334462 | 2.419091 | 2.49723 | 2.544872 | 2.586709 | 2.635942 | 2.680232 |
|  | 15 | 1.926503 | 1.963711 | 2.005889 | 2.04242 | 2.0848 | 2.156144 | 2.236057 | 2.317645 | 2.393431 | 2.439836 | 2.480699 | 2.52891 | 2.572387 |
|  | 20 | 1.87876 | 1.9154 | 1.95693 | 1.992888 | 2.034585 | 2.104695 | 2.183062 | 2.263237 | 2.338163 | 2.38424 | 2.424929 | 2.473059 | 2.516572 |
|  | 30 | 1.841825 | 1.878443 | 1.919943 | 1.955857 | 1.997461 | 2.067257 | 2.144951 | 2.224767 | 2.300283 | 2.347133 | 2.38874 | 2.438222 | 2.483191 |
|  | 40 | 1.827413 | 1.864454 | 1.906428 | 1.942737 | 1.984761 | 2.055107 | 2.13309 | 2.213546 | 2.290628 | 2.338882 | 2.381988 | 2.433541 | 2.480648 |
|  | 50 | 1.817108 | 1.854684 | 1.897266 | 1.934088 | 1.976672 | 2.047805 | 2.12634 | 2.207721 | 2.286697 | 2.336596 | 2.381442 | 2.435391 | 2.484971 |
|  | 60 | 1.795481 | 1.833397 | 1.876368 | 1.913517 | 1.956449 | 2.028023 | 2.106729 | 2.18866 | 2.26922 | 2.320605 | 2.36708 | 2.423332 | 2.475343 |
|  | 70 | 1.747273 | 1.784967 | 1.827698 | 1.864633 | 1.907292 | 1.978281 | 2.056037 | 2.137361 | 2.218406 | 2.270609 | 2.318134 | 2.376031 | 2.429906 |
|  | 80 | 1.71481 | 1.752599 | 1.795451 | 1.832489 | 1.875245 | 1.946273 | 2.023772 | 2.10522 | 2.187515 | 2.241058 | 2.29014 | 2.350339 | 2.406739 |
|  | 90 | 1.65483 | 1.692066 | 1.734307 | 1.770818 | 1.812948 | 1.882823 | 1.958774 | 2.038995 | 2.121198 | 2.175239 | 2.225132 | 2.286765 | 2.344929 |
| **L_lateralorbitofrontal** | 5 | 2.647113 | 2.733181 | 2.820004 | 2.888024 | 2.960455 | 3.070827 | 3.18302 | 3.28921 | 3.38309 | 3.439675 | 3.48953 | 3.548919 | 3.603459 |
|  | 10 | 2.514902 | 2.586546 | 2.66101 | 2.72084 | 2.785946 | 2.887798 | 2.994379 | 3.09781 | 3.19108 | 3.248035 | 3.298636 | 3.359384 | 3.415594 |
|  | 15 | 2.402662 | 2.464732 | 2.530696 | 2.58475 | 2.644612 | 2.740361 | 2.843145 | 2.945213 | 3.039021 | 3.097054 | 3.149053 | 3.211995 | 3.270708 |
|  | 20 | 2.323904 | 2.380651 | 2.44187 | 2.492724 | 2.549747 | 2.642441 | 2.743853 | 2.846335 | 2.941921 | 3.001658 | 3.055545 | 3.121201 | 3.182843 |
|  | 30 | 2.232096 | 2.285179 | 2.343155 | 2.391855 | 2.447017 | 2.537835 | 2.638605 | 2.741628 | 2.838502 | 2.899314 | 2.954298 | 3.021402 | 3.084475 |
|  | 40 | 2.181864 | 2.233876 | 2.290865 | 2.338862 | 2.393338 | 2.48318 | 2.582892 | 2.684586 | 2.779741 | 2.83915 | 2.89261 | 2.957492 | 3.018085 |
|  | 50 | 2.165807 | 2.216993 | 2.273246 | 2.320737 | 2.374739 | 2.463946 | 2.56299 | 2.663804 | 2.757741 | 2.816117 | 2.868432 | 2.93162 | 2.990305 |
|  | 60 | 2.156942 | 2.206241 | 2.260802 | 2.30716 | 2.360177 | 2.448389 | 2.54709 | 2.648162 | 2.742692 | 2.80153 | 2.854285 | 2.918002 | 2.977148 |
|  | 70 | 2.10439 | 2.151532 | 2.204291 | 2.249588 | 2.301906 | 2.390115 | 2.490412 | 2.594689 | 2.69349 | 2.755543 | 2.811508 | 2.879488 | 2.94294 |
|  | 80 | 2.068077 | 2.113927 | 2.165922 | 2.211135 | 2.264007 | 2.354703 | 2.460133 | 2.572253 | 2.680743 | 2.749965 | 2.813096 | 2.890653 | 2.963895 |
|  | 90 | 2.031747 | 2.076118 | 2.127106 | 2.172025 | 2.225241 | 2.318265 | 2.429165 | 2.550373 | 2.670888 | 2.749458 | 2.822267 | 2.913237 | 3.000716 |
| **L_lingual** | 5 | 2.019893 | 2.064336 | 2.11309 | 2.154199 | 2.200911 | 2.278081 | 2.36393 | 2.451715 | 2.534075 | 2.585607 | 2.632054 | 2.68852 | 2.741351 |
|  | 10 | 1.891561 | 1.933008 | 1.978736 | 2.017486 | 2.061704 | 2.135102 | 2.217084 | 2.301029 | 2.379668 | 2.42873 | 2.472819 | 2.526215 | 2.575944 |
|  | 15 | 1.794764 | 1.834045 | 1.877604 | 1.914679 | 1.957146 | 2.027938 | 2.107303 | 2.188682 | 2.264836 | 2.312238 | 2.354729 | 2.406028 | 2.453618 |
|  | 20 | 1.7434 | 1.781558 | 1.824062 | 1.86038 | 1.90212 | 1.971969 | 2.050543 | 2.131225 | 2.206673 | 2.253547 | 2.29548 | 2.345971 | 2.39266 |
|  | 30 | 1.699285 | 1.736221 | 1.777666 | 1.813309 | 1.854502 | 1.923876 | 2.002368 | 2.083178 | 2.158683 | 2.205463 | 2.247181 | 2.297208 | 2.343232 |
|  | 40 | 1.668628 | 1.704334 | 1.744626 | 1.779448 | 1.819865 | 1.888274 | 1.966032 | 2.046269 | 2.121217 | 2.167568 | 2.208814 | 2.258132 | 2.303339 |
|  | 50 | 1.643556 | 1.678338 | 1.717764 | 1.751975 | 1.79182 | 1.859544 | 1.936832 | 2.016772 | 2.09147 | 2.137625 | 2.178644 | 2.227601 | 2.272368 |
|  | 60 | 1.621617 | 1.65612 | 1.695382 | 1.729569 | 1.769514 | 1.837671 | 1.915767 | 1.996774 | 2.072571 | 2.11941 | 2.161021 | 2.210644 | 2.255965 |
|  | 70 | 1.571751 | 1.605833 | 1.644753 | 1.678753 | 1.718596 | 1.786838 | 1.865362 | 1.94709 | 2.023731 | 2.071142 | 2.113275 | 2.163524 | 2.209407 |
|  | 80 | 1.52133 | 1.555184 | 1.593969 | 1.627953 | 1.667891 | 1.73655 | 1.815896 | 1.898797 | 1.976764 | 2.025079 | 2.068057 | 2.119351 | 2.166212 |
|  | 90 | 1.49134 | 1.525495 | 1.564745 | 1.599234 | 1.639875 | 1.71 | 1.791402 | 1.876804 | 1.957402 | 2.007461 | 2.052054 | 2.105343 | 2.154083 |
| **L_medialorbitofrontal** | 5 | 2.461426 | 2.521226 | 2.587652 | 2.644273 | 2.709217 | 2.817653 | 2.939423 | 3.06443 | 3.181479 | 3.254342 | 3.319648 | 3.39847 | 3.471564 |
|  | 10 | 2.298149 | 2.354034 | 2.416177 | 2.469209 | 2.530108 | 2.631986 | 2.746716 | 2.864901 | 2.975978 | 3.045345 | 3.107669 | 3.183089 | 3.253228 |
|  | 15 | 2.183816 | 2.236966 | 2.296132 | 2.34668 | 2.404797 | 2.502206 | 2.612219 | 2.725944 | 2.833244 | 2.900472 | 2.961027 | 3.034508 | 3.103046 |
|  | 20 | 2.112034 | 2.163478 | 2.220805 | 2.269837 | 2.326279 | 2.421063 | 2.528424 | 2.639806 | 2.745314 | 2.811644 | 2.871547 | 2.944445 | 3.01265 |
|  | 30 | 2.026726 | 2.076124 | 2.131291 | 2.178583 | 2.23316 | 2.325178 | 2.430046 | 2.539673 | 2.644403 | 2.710726 | 2.770962 | 2.844719 | 2.914196 |
|  | 40 | 1.987858 | 2.036229 | 2.090372 | 2.136904 | 2.19075 | 2.281938 | 2.386565 | 2.496873 | 2.603265 | 2.671199 | 2.733298 | 2.809874 | 2.882572 |
|  | 50 | 1.972934 | 2.02072 | 2.074346 | 2.120561 | 2.174202 | 2.265492 | 2.371036 | 2.483385 | 2.592933 | 2.663551 | 2.728587 | 2.809453 | 2.886929 |
|  | 60 | 1.957275 | 2.004334 | 2.057289 | 2.103061 | 2.156364 | 2.247569 | 2.353909 | 2.468334 | 2.581295 | 2.654912 | 2.723301 | 2.809164 | 2.892325 |
|  | 70 | 1.944418 | 1.990761 | 2.043053 | 2.088389 | 2.141363 | 2.232519 | 2.339758 | 2.456504 | 2.573342 | 2.65042 | 2.722733 | 2.814537 | 2.904582 |
|  | 80 | 1.912088 | 1.957246 | 2.008333 | 2.052755 | 2.104836 | 2.194968 | 2.301989 | 2.419947 | 2.53975 | 2.619853 | 2.695836 | 2.793527 | 2.890753 |
|  | 90 | 1.878236 | 1.922189 | 1.972035 | 2.015499 | 2.066626 | 2.155611 | 2.262277 | 2.381374 | 2.504266 | 2.587653 | 2.667729 | 2.772174 | 2.877889 |
| **L_middletemporal** | 5 | 2.519374 | 2.667693 | 2.800446 | 2.895034 | 2.988297 | 3.117768 | 3.235271 | 3.335444 | 3.415921 | 3.460523 | 3.497328 | 3.538139 | 3.572794 |
|  | 10 | 2.555687 | 2.655708 | 2.754769 | 2.83109 | 2.910924 | 3.02883 | 3.142273 | 3.243847 | 3.328886 | 3.377255 | 3.417796 | 3.463386 | 3.502602 |
|  | 15 | 2.546132 | 2.618837 | 2.69609 | 2.75907 | 2.828061 | 2.935491 | 3.044696 | 3.147631 | 3.237918 | 3.29089 | 3.336162 | 3.387999 | 3.433355 |
|  | 20 | 2.514413 | 2.571508 | 2.634747 | 2.688257 | 2.748875 | 2.847369 | 2.952507 | 3.056819 | 3.153059 | 3.211589 | 3.262818 | 3.322834 | 3.376534 |
|  | 30 | 2.43014 | 2.482679 | 2.541549 | 2.591894 | 2.649484 | 2.744203 | 2.846695 | 2.950742 | 3.049745 | 3.11133 | 3.166058 | 3.231135 | 3.290225 |
|  | 40 | 2.339422 | 2.40367 | 2.473274 | 2.530939 | 2.594943 | 2.696001 | 2.799976 | 2.900828 | 2.993192 | 3.048997 | 3.097582 | 3.154173 | 3.204495 |
|  | 50 | 2.329195 | 2.390001 | 2.456466 | 2.511995 | 2.574124 | 2.673319 | 2.77683 | 2.878086 | 2.971055 | 3.027355 | 3.076463 | 3.133781 | 3.184863 |
|  | 60 | 2.320619 | 2.372544 | 2.430815 | 2.480768 | 2.538108 | 2.633103 | 2.737191 | 2.842036 | 2.939063 | 2.998263 | 3.05023 | 3.111322 | 3.166196 |
|  | 70 | 2.245635 | 2.29512 | 2.351238 | 2.399904 | 2.456499 | 2.552317 | 2.66071 | 2.77013 | 2.868595 | 2.927541 | 2.978705 | 3.038261 | 3.091283 |
|  | 80 | 2.149445 | 2.205644 | 2.268675 | 2.32284 | 2.385382 | 2.490585 | 2.608959 | 2.724364 | 2.822156 | 2.878273 | 2.9257 | 2.979557 | 3.026396 |
|  | 90 | 2.069442 | 2.130446 | 2.198698 | 2.257304 | 2.325047 | 2.439558 | 2.56947 | 2.693811 | 2.794839 | 2.851152 | 2.897922 | 2.950218 | 2.995064 |
| **L_parahippocampal** | 5 | 2.184429 | 2.292312 | 2.411344 | 2.512057 | 2.626608 | 2.815213 | 3.021778 | 3.220871 | 3.39038 | 3.488006 | 3.570668 | 3.664667 | 3.746498 |
|  | 10 | 2.144173 | 2.246621 | 2.360729 | 2.458166 | 2.570013 | 2.756663 | 2.964697 | 3.166527 | 3.337559 | 3.435737 | 3.518726 | 3.61298 | 3.694959 |
|  | 15 | 2.102394 | 2.199766 | 2.30919 | 2.403451 | 2.512615 | 2.697193 | 2.906467 | 3.110928 | 3.283558 | 3.382389 | 3.46582 | 3.560487 | 3.642775 |
|  | 20 | 2.059336 | 2.152329 | 2.257657 | 2.3491 | 2.455843 | 2.638465 | 2.848728 | 3.055645 | 3.230053 | 3.329763 | 3.413887 | 3.509312 | 3.592254 |
|  | 30 | 1.981171 | 2.068351 | 2.16821 | 2.255871 | 2.359348 | 2.539316 | 2.751013 | 2.962231 | 3.14104 | 3.243534 | 3.330181 | 3.428679 | 3.514487 |
|  | 40 | 1.918534 | 2.004261 | 2.102857 | 2.189727 | 2.292612 | 2.472297 | 2.684749 | 2.898747 | 3.082197 | 3.188281 | 3.278461 | 3.381502 | 3.471705 |
|  | 50 | 1.873812 | 1.963668 | 2.066555 | 2.156765 | 2.263019 | 2.446897 | 2.661567 | 2.877491 | 3.06447 | 3.173389 | 3.266379 | 3.373027 | 3.466698 |
|  | 60 | 1.825965 | 1.92361 | 2.034245 | 2.130229 | 2.242059 | 2.432438 | 2.649897 | 2.866207 | 3.053704 | 3.163021 | 3.256359 | 3.363379 | 3.457323 |
|  | 70 | 1.761901 | 1.866543 | 1.983759 | 2.084341 | 2.20027 | 2.394603 | 2.612217 | 2.826034 | 3.010714 | 3.118135 | 3.209688 | 3.314456 | 3.406228 |
|  | 80 | 1.681822 | 1.791261 | 1.912525 | 2.015544 | 2.133171 | 2.327854 | 2.542471 | 2.750943 | 2.929892 | 3.033545 | 3.121636 | 3.222161 | 3.309973 |
|  | 90 | 1.600389 | 1.714991 | 1.840417 | 1.945818 | 2.064993 | 2.259769 | 2.471341 | 2.674345 | 2.847058 | 2.946506 | 3.0307 | 3.126428 | 3.209755 |
| **L_paracentral** | 5 | 2.329178 | 2.379705 | 2.436991 | 2.486568 | 2.543977 | 2.640136 | 2.746818 | 2.856955 | 2.962524 | 3.028635 | 3.087705 | 3.158367 | 3.222949 |
|  | 10 | 2.218107 | 2.267187 | 2.322804 | 2.370912 | 2.426594 | 2.519806 | 2.623155 | 2.729677 | 2.831522 | 2.895179 | 2.951982 | 3.019842 | 3.08178 |
|  | 15 | 2.113386 | 2.1611 | 2.215137 | 2.261853 | 2.315897 | 2.406313 | 2.506494 | 2.609577 | 2.707881 | 2.769204 | 2.823852 | 2.889051 | 2.948481 |
|  | 20 | 2.043171 | 2.090252 | 2.143539 | 2.189581 | 2.242816 | 2.331821 | 2.430366 | 2.531591 | 2.627871 | 2.687813 | 2.741158 | 2.804718 | 2.862575 |
|  | 30 | 1.963223 | 2.010395 | 2.063711 | 2.10972 | 2.162856 | 2.251567 | 2.349628 | 2.449993 | 2.544941 | 2.603819 | 2.656072 | 2.71816 | 2.774525 |
|  | 40 | 1.91609 | 1.964187 | 2.018467 | 2.065243 | 2.119194 | 2.209119 | 2.308339 | 2.409503 | 2.504678 | 2.563453 | 2.61547 | 2.677106 | 2.732906 |
|  | 50 | 1.870826 | 1.920005 | 1.975413 | 2.023086 | 2.077992 | 2.169342 | 2.269924 | 2.372063 | 2.467608 | 2.526364 | 2.578213 | 2.63948 | 2.69479 |
|  | 60 | 1.828012 | 1.878466 | 1.935201 | 1.983929 | 2.039961 | 2.132991 | 2.235183 | 2.338514 | 2.434605 | 2.49344 | 2.54521 | 2.606207 | 2.661119 |
|  | 70 | 1.77183 | 1.823311 | 1.881077 | 1.930592 | 1.987423 | 2.081563 | 2.184701 | 2.288515 | 2.38447 | 2.442961 | 2.494274 | 2.554559 | 2.608675 |
|  | 80 | 1.72056 | 1.773321 | 1.832378 | 1.882886 | 1.940739 | 2.03632 | 2.140724 | 2.245306 | 2.341364 | 2.399652 | 2.450631 | 2.510349 | 2.563801 |
|  | 90 | 1.691305 | 1.746183 | 1.807438 | 1.859693 | 1.919407 | 2.017775 | 2.124864 | 2.231586 | 2.328974 | 2.387793 | 2.439078 | 2.498975 | 2.552431 |
| **L_parsopercularis** | 5 | 2.546599 | 2.596623 | 2.652816 | 2.701085 | 2.756666 | 2.84938 | 2.952163 | 3.054716 | 3.146827 | 3.201869 | 3.249559 | 3.304953 | 3.354149 |
|  | 10 | 2.466078 | 2.513873 | 2.567778 | 2.614261 | 2.667988 | 2.758083 | 2.858644 | 2.959677 | 3.051002 | 3.10583 | 3.15349 | 3.209026 | 3.258505 |
|  | 15 | 2.388546 | 2.434267 | 2.486029 | 2.530826 | 2.582794 | 2.670386 | 2.768804 | 2.868362 | 2.958931 | 3.013566 | 3.061213 | 3.116915 | 3.166706 |
|  | 20 | 2.320407 | 2.364398 | 2.414366 | 2.457753 | 2.508248 | 2.593752 | 2.690408 | 2.788806 | 2.878856 | 2.933421 | 2.981157 | 3.037138 | 3.087337 |
|  | 30 | 2.220054 | 2.262048 | 2.309957 | 2.351735 | 2.400567 | 2.483767 | 2.578594 | 2.67597 | 2.765824 | 2.820614 | 2.868759 | 2.925469 | 2.976551 |
|  | 40 | 2.154955 | 2.196421 | 2.243818 | 2.285226 | 2.333717 | 2.41656 | 2.511321 | 2.609004 | 2.699474 | 2.754795 | 2.803501 | 2.860989 | 2.912878 |
|  | 50 | 2.109314 | 2.151077 | 2.198836 | 2.240581 | 2.28949 | 2.373104 | 2.468831 | 2.5676 | 2.659156 | 2.715176 | 2.764521 | 2.822789 | 2.875404 |
|  | 60 | 2.070352 | 2.113093 | 2.161917 | 2.204546 | 2.254434 | 2.339579 | 2.43684 | 2.536945 | 2.629516 | 2.686052 | 2.735784 | 2.794429 | 2.847313 |
|  | 70 | 2.014729 | 2.058655 | 2.108698 | 2.152276 | 2.203138 | 2.289609 | 2.387873 | 2.48845 | 2.580959 | 2.637227 | 2.68658 | 2.744607 | 2.796777 |
|  | 80 | 1.968403 | 2.014024 | 2.065815 | 2.110759 | 2.163034 | 2.251468 | 2.351304 | 2.452787 | 2.545518 | 2.601641 | 2.650696 | 2.708173 | 2.759665 |
|  | 90 | 1.929064 | 1.976753 | 2.030675 | 2.077286 | 2.13129 | 2.22215 | 2.323995 | 2.426756 | 2.520008 | 2.576154 | 2.625055 | 2.682147 | 2.733111 |
| **L_parsorbitalis** | 5 | 2.600049 | 2.686846 | 2.782932 | 2.864375 | 2.95702 | 3.109163 | 3.274856 | 3.437661 | 3.582212 | 3.667997 | 3.742042 | 3.827785 | 3.903756 |
|  | 10 | 2.462518 | 2.544886 | 2.636268 | 2.713885 | 2.802363 | 2.948092 | 3.107435 | 3.264679 | 3.404893 | 3.488386 | 3.560629 | 3.644497 | 3.719001 |
|  | 15 | 2.345993 | 2.4247 | 2.512191 | 2.58665 | 2.671695 | 2.812177 | 2.966396 | 3.119264 | 3.256196 | 3.338033 | 3.409032 | 3.491685 | 3.565327 |
|  | 20 | 2.258057 | 2.33388 | 2.418316 | 2.490302 | 2.572676 | 2.709127 | 2.85952 | 3.009287 | 3.14409 | 3.224973 | 3.295352 | 3.377538 | 3.451005 |
|  | 30 | 2.162968 | 2.234466 | 2.314291 | 2.382537 | 2.460873 | 2.591286 | 2.736134 | 2.881762 | 3.014228 | 3.094423 | 3.16468 | 3.247323 | 3.321782 |
|  | 40 | 2.136529 | 2.205364 | 2.282258 | 2.34807 | 2.423738 | 2.550153 | 2.69149 | 2.834951 | 2.966971 | 3.047739 | 3.119084 | 3.203775 | 3.280846 |
|  | 50 | 2.110877 | 2.177689 | 2.252154 | 2.315802 | 2.388957 | 2.511333 | 2.64883 | 2.789709 | 2.92106 | 3.002437 | 3.075067 | 3.162297 | 3.242723 |
|  | 60 | 2.06476 | 2.131262 | 2.20496 | 2.267684 | 2.339575 | 2.459678 | 2.59505 | 2.735121 | 2.867797 | 2.951334 | 3.026923 | 3.11915 | 3.20573 |
|  | 70 | 1.990838 | 2.059029 | 2.133834 | 2.19698 | 2.268913 | 2.38852 | 2.523472 | 2.664643 | 2.801101 | 2.888917 | 2.969908 | 3.070959 | 3.168303 |
|  | 80 | 1.945978 | 2.019852 | 2.099553 | 2.165893 | 2.24063 | 2.363642 | 2.502072 | 2.648646 | 2.794163 | 2.890684 | 2.982145 | 3.100012 | 3.217931 |
|  | 90 | 1.908248 | 1.992477 | 2.081035 | 2.153098 | 2.232775 | 2.36148 | 2.505031 | 2.659037 | 2.817518 | 2.927217 | 3.035321 | 3.18145 | 3.336186 |
| **L_parstriangularis** | 5 | 2.468507 | 2.529935 | 2.597617 | 2.654573 | 2.718717 | 2.821974 | 2.930724 | 3.038392 | 3.138657 | 3.199997 | 3.253865 | 3.317155 | 3.373924 |
|  | 10 | 2.347053 | 2.405522 | 2.470289 | 2.525078 | 2.587103 | 2.687701 | 2.794706 | 2.901265 | 3.000635 | 3.061516 | 3.115048 | 3.178034 | 3.234618 |
|  | 15 | 2.237088 | 2.292851 | 2.354953 | 2.407765 | 2.467871 | 2.566104 | 2.671668 | 2.777445 | 2.876264 | 2.936912 | 2.990319 | 3.053261 | 3.109909 |
|  | 20 | 2.152268 | 2.205864 | 2.265872 | 2.317173 | 2.375871 | 2.472554 | 2.577543 | 2.683435 | 2.782577 | 2.843546 | 2.897326 | 2.960825 | 3.01809 |
|  | 30 | 2.058656 | 2.108845 | 2.165602 | 2.214613 | 2.271269 | 2.366023 | 2.471073 | 2.578444 | 2.679447 | 2.741829 | 2.797052 | 2.862514 | 2.921807 |
|  | 40 | 2.007699 | 2.054521 | 2.107916 | 2.154424 | 2.208671 | 2.300653 | 2.404598 | 2.51213 | 2.613671 | 2.676614 | 2.732509 | 2.799004 | 2.85947 |
|  | 50 | 1.976072 | 2.02005 | 2.070553 | 2.114866 | 2.166957 | 2.256367 | 2.359173 | 2.466657 | 2.568417 | 2.631665 | 2.687969 | 2.755148 | 2.81644 |
|  | 60 | 1.944931 | 1.986776 | 2.035128 | 2.077831 | 2.12839 | 2.216155 | 2.31873 | 2.427024 | 2.52975 | 2.593737 | 2.650819 | 2.7191 | 2.781584 |
|  | 70 | 1.89091 | 1.930703 | 1.976944 | 2.018033 | 2.067005 | 2.152935 | 2.254952 | 2.363667 | 2.466964 | 2.531427 | 2.589043 | 2.658129 | 2.721526 |
|  | 80 | 1.873676 | 1.912414 | 1.957674 | 1.998129 | 2.04666 | 2.132724 | 2.236506 | 2.348162 | 2.454463 | 2.52094 | 2.580482 | 2.652058 | 2.717941 |
|  | 90 | 1.883983 | 1.92223 | 1.967155 | 2.007544 | 2.056307 | 2.143709 | 2.250779 | 2.36713 | 2.478196 | 2.547833 | 2.610359 | 2.685747 | 2.755382 |
| **L_pericalcarine** | 5 | 1.532669 | 1.565618 | 1.60411 | 1.63849 | 1.679675 | 1.752448 | 1.839655 | 1.93432 | 2.026657 | 2.08548 | 2.138836 | 2.203785 | 2.264327 |
|  | 10 | 1.44085 | 1.47227 | 1.509015 | 1.541871 | 1.581273 | 1.651016 | 1.734795 | 1.825989 | 1.915195 | 1.972155 | 2.02391 | 2.087024 | 2.145969 |
|  | 15 | 1.373091 | 1.403459 | 1.439012 | 1.470837 | 1.509047 | 1.576798 | 1.658386 | 1.747447 | 1.834824 | 1.890749 | 1.941655 | 2.003851 | 2.062056 |
|  | 20 | 1.338346 | 1.368339 | 1.40349 | 1.43499 | 1.472853 | 1.540107 | 1.621295 | 1.710173 | 1.797627 | 1.853738 | 1.904905 | 1.967541 | 2.026277 |
|  | 30 | 1.305277 | 1.335093 | 1.370101 | 1.401534 | 1.439394 | 1.506851 | 1.588648 | 1.678656 | 1.767703 | 1.825092 | 1.8776 | 1.942107 | 2.002832 |
|  | 40 | 1.273788 | 1.303144 | 1.337665 | 1.368708 | 1.406158 | 1.473054 | 1.554465 | 1.644431 | 1.733843 | 1.791684 | 1.844758 | 1.910161 | 1.971934 |
|  | 50 | 1.258095 | 1.287133 | 1.321321 | 1.352101 | 1.389284 | 1.455837 | 1.537071 | 1.62716 | 1.717035 | 1.775362 | 1.829013 | 1.895304 | 1.958096 |
|  | 60 | 1.249836 | 1.278644 | 1.312596 | 1.343199 | 1.380209 | 1.446575 | 1.5278 | 1.618171 | 1.708648 | 1.767541 | 1.821838 | 1.889096 | 1.952981 |
|  | 70 | 1.218005 | 1.245977 | 1.278973 | 1.308743 | 1.344785 | 1.409522 | 1.488949 | 1.577585 | 1.666616 | 1.724732 | 1.77843 | 1.845104 | 1.908601 |
|  | 80 | 1.196801 | 1.224069 | 1.256259 | 1.285323 | 1.320539 | 1.383877 | 1.461742 | 1.548849 | 1.636585 | 1.69399 | 1.747129 | 1.813246 | 1.876354 |
|  | 90 | 1.178247 | 1.204838 | 1.236247 | 1.264626 | 1.299038 | 1.361 | 1.437311 | 1.522869 | 1.60926 | 1.665909 | 1.718439 | 1.783923 | 1.84656 |
| **L_postcentral** | 5 | 2.022735 | 2.064007 | 2.109826 | 2.148864 | 2.193621 | 2.268322 | 2.352182 | 2.438265 | 2.518883 | 2.569083 | 2.61409 | 2.668432 | 2.718849 |
|  | 10 | 1.935938 | 1.975949 | 2.020356 | 2.058184 | 2.101547 | 2.17391 | 2.255139 | 2.338531 | 2.416651 | 2.46531 | 2.508949 | 2.561658 | 2.610579 |
|  | 15 | 1.859236 | 1.898159 | 1.941347 | 1.978129 | 2.020287 | 2.090627 | 2.169581 | 2.250645 | 2.326606 | 2.373936 | 2.416396 | 2.467698 | 2.515331 |
|  | 20 | 1.811404 | 1.849818 | 1.89243 | 1.928714 | 1.970294 | 2.039659 | 2.117513 | 2.197457 | 2.272389 | 2.319093 | 2.361004 | 2.41166 | 2.458714 |
|  | 30 | 1.756723 | 1.794954 | 1.837343 | 1.873422 | 1.914752 | 1.983677 | 2.061027 | 2.14047 | 2.214974 | 2.261443 | 2.303167 | 2.353634 | 2.40055 |
|  | 40 | 1.725138 | 1.763675 | 1.80638 | 1.842711 | 1.884316 | 1.953677 | 2.031502 | 2.11145 | 2.186472 | 2.233295 | 2.275363 | 2.326282 | 2.37366 |
|  | 50 | 1.689694 | 1.728444 | 1.771362 | 1.807859 | 1.849638 | 1.919264 | 1.997374 | 2.077632 | 2.152987 | 2.200051 | 2.242362 | 2.293613 | 2.34134 |
|  | 60 | 1.659113 | 1.698184 | 1.741433 | 1.778193 | 1.820259 | 1.890337 | 1.96894 | 2.049721 | 2.125612 | 2.173044 | 2.215713 | 2.267436 | 2.315645 |
|  | 70 | 1.619486 | 1.658655 | 1.701989 | 1.738803 | 1.780914 | 1.851042 | 1.929686 | 2.010525 | 2.086516 | 2.134045 | 2.176827 | 2.228728 | 2.277146 |
|  | 80 | 1.580278 | 1.61954 | 1.662951 | 1.699814 | 1.741963 | 1.812127 | 1.890794 | 1.971674 | 2.04775 | 2.095366 | 2.138254 | 2.190325 | 2.238946 |
|  | 90 | 1.533652 | 1.572802 | 1.616064 | 1.65278 | 1.694745 | 1.764574 | 1.842849 | 1.923341 | 1.999098 | 2.046549 | 2.089318 | 2.141285 | 2.189853 |
| **L_posteriorcingulate** | 5 | 2.618153 | 2.665606 | 2.719987 | 2.767623 | 2.823555 | 2.919489 | 3.029814 | 3.144206 | 3.250761 | 3.316208 | 3.374014 | 3.442464 | 3.504455 |
|  | 10 | 2.502221 | 2.548631 | 2.601778 | 2.6483 | 2.702885 | 2.796405 | 2.903794 | 3.01496 | 3.118342 | 3.18176 | 3.237722 | 3.303927 | 3.363827 |
|  | 15 | 2.389834 | 2.435207 | 2.487129 | 2.532544 | 2.585789 | 2.676911 | 2.781384 | 2.889349 | 2.989586 | 3.050994 | 3.105134 | 3.16912 | 3.226955 |
|  | 20 | 2.296715 | 2.341366 | 2.392421 | 2.437043 | 2.489316 | 2.578669 | 2.68095 | 2.786463 | 2.884257 | 2.944088 | 2.996786 | 3.059009 | 3.115194 |
|  | 30 | 2.179815 | 2.224295 | 2.275069 | 2.31937 | 2.371178 | 2.459513 | 2.560286 | 2.663861 | 2.759514 | 2.817873 | 2.869174 | 2.929625 | 2.984098 |
|  | 40 | 2.117444 | 2.162861 | 2.214607 | 2.259673 | 2.312276 | 2.401722 | 2.503388 | 2.607471 | 2.703229 | 2.761483 | 2.812586 | 2.872679 | 2.926714 |
|  | 50 | 2.071077 | 2.117849 | 2.171028 | 2.217247 | 2.271084 | 2.362359 | 2.465693 | 2.571042 | 2.667578 | 2.726126 | 2.777377 | 2.837516 | 2.891473 |
|  | 60 | 2.042693 | 2.091352 | 2.146551 | 2.194416 | 2.250046 | 2.344054 | 2.450032 | 2.557596 | 2.655745 | 2.715084 | 2.766912 | 2.827592 | 2.881913 |
|  | 70 | 1.995404 | 2.045643 | 2.102488 | 2.151657 | 2.208661 | 2.304654 | 2.412375 | 2.521191 | 2.620043 | 2.679608 | 2.731516 | 2.792149 | 2.846304 |
|  | 80 | 1.963336 | 2.015696 | 2.074772 | 2.125728 | 2.184645 | 2.28348 | 2.393846 | 2.504772 | 2.605074 | 2.665305 | 2.717667 | 2.778689 | 2.833063 |
|  | 90 | 1.951018 | 2.006266 | 2.0684 | 2.12183 | 2.183422 | 2.286317 | 2.400609 | 2.514869 | 2.617681 | 2.679198 | 2.732548 | 2.794572 | 2.849705 |
| **L_precentral** | 5 | 2.266306 | 2.340905 | 2.413721 | 2.469256 | 2.527132 | 2.613318 | 2.699282 | 2.78011 | 2.851957 | 2.895737 | 2.934739 | 2.981842 | 3.025807 |
|  | 10 | 2.263098 | 2.331013 | 2.399061 | 2.452072 | 2.508271 | 2.593531 | 2.679973 | 2.7619 | 2.834699 | 2.878849 | 2.917963 | 2.964859 | 3.008248 |
|  | 15 | 2.246898 | 2.309409 | 2.373408 | 2.424163 | 2.478763 | 2.562953 | 2.649577 | 2.732321 | 2.805908 | 2.850395 | 2.889647 | 2.936444 | 2.979442 |
|  | 20 | 2.219022 | 2.277256 | 2.337918 | 2.386731 | 2.43988 | 2.522959 | 2.609523 | 2.69279 | 2.766931 | 2.811651 | 2.85098 | 2.897662 | 2.940311 |
|  | 30 | 2.155866 | 2.209483 | 2.266557 | 2.313332 | 2.365046 | 2.447271 | 2.534251 | 2.618514 | 2.693454 | 2.738402 | 2.777681 | 2.823915 | 2.865723 |
|  | 40 | 2.100046 | 2.15321 | 2.210274 | 2.257359 | 2.309694 | 2.393329 | 2.482027 | 2.567741 | 2.643427 | 2.688438 | 2.727477 | 2.773024 | 2.813791 |
|  | 50 | 2.0498 | 2.104747 | 2.163782 | 2.212514 | 2.266667 | 2.353078 | 2.444341 | 2.531899 | 2.608468 | 2.653592 | 2.692441 | 2.737395 | 2.777267 |
|  | 60 | 1.993852 | 2.049713 | 2.109882 | 2.159643 | 2.215013 | 2.303442 | 2.396782 | 2.486083 | 2.563809 | 2.609389 | 2.64847 | 2.693478 | 2.733184 |
|  | 70 | 1.924803 | 1.979282 | 2.038408 | 2.08763 | 2.142712 | 2.231278 | 2.325405 | 2.415893 | 2.494847 | 2.541177 | 2.580892 | 2.626597 | 2.666868 |
|  | 80 | 1.861315 | 1.913203 | 1.970142 | 2.018015 | 2.072065 | 2.15995 | 2.254537 | 2.346456 | 2.427317 | 2.475003 | 2.515997 | 2.563289 | 2.605041 |
|  | 90 | 1.80501 | 1.854362 | 1.909118 | 1.955619 | 2.008609 | 2.095817 | 2.191014 | 2.284726 | 2.368039 | 2.417516 | 2.460236 | 2.509716 | 2.553561 |
| **L_precuneus** | 5 | 2.505547 | 2.546653 | 2.592892 | 2.632659 | 2.678508 | 2.755121 | 2.840242 | 2.925364 | 3.001976 | 3.047826 | 3.087593 | 3.133831 | 3.174938 |
|  | 10 | 2.34059 | 2.381745 | 2.428038 | 2.467853 | 2.513757 | 2.59046 | 2.675683 | 2.760906 | 2.837609 | 2.883513 | 2.923328 | 2.969621 | 3.010776 |
|  | 15 | 2.194676 | 2.23588 | 2.282228 | 2.32209 | 2.368049 | 2.444844 | 2.530168 | 2.615492 | 2.692287 | 2.738245 | 2.778107 | 2.824456 | 2.86566 |
|  | 20 | 2.104241 | 2.145495 | 2.191898 | 2.231808 | 2.277821 | 2.354707 | 2.440133 | 2.525559 | 2.602445 | 2.648459 | 2.688368 | 2.734772 | 2.776025 |
|  | 30 | 2.009297 | 2.050649 | 2.097162 | 2.137167 | 2.183289 | 2.260358 | 2.345987 | 2.431615 | 2.508684 | 2.554807 | 2.594811 | 2.641325 | 2.682676 |
|  | 40 | 1.964671 | 2.00612 | 2.052744 | 2.092842 | 2.139074 | 2.216324 | 2.302155 | 2.387986 | 2.465236 | 2.511467 | 2.551566 | 2.59819 | 2.639639 |
|  | 50 | 1.931887 | 1.973434 | 2.020168 | 2.060362 | 2.106703 | 2.184136 | 2.270171 | 2.356205 | 2.433638 | 2.479979 | 2.520173 | 2.566907 | 2.608454 |
|  | 60 | 1.907664 | 1.949311 | 1.996157 | 2.036447 | 2.082899 | 2.160519 | 2.246759 | 2.333 | 2.410619 | 2.457071 | 2.497362 | 2.544208 | 2.585855 |
|  | 70 | 1.846537 | 1.888285 | 1.935245 | 1.975634 | 2.022199 | 2.100008 | 2.186459 | 2.27291 | 2.350719 | 2.397284 | 2.437672 | 2.484633 | 2.526381 |
|  | 80 | 1.781684 | 1.823536 | 1.870612 | 1.9111 | 1.95778 | 2.035781 | 2.122445 | 2.20911 | 2.28711 | 2.33379 | 2.374278 | 2.421355 | 2.463206 |
|  | 90 | 1.726747 | 1.768703 | 1.815896 | 1.856484 | 1.90328 | 1.981474 | 2.068354 | 2.155233 | 2.233427 | 2.280223 | 2.320811 | 2.368004 | 2.40996 |
| **L_rostralanteriorcingulate** | 5 | 2.514769 | 2.681307 | 2.835619 | 2.94788 | 3.060505 | 3.222804 | 3.384911 | 3.547885 | 3.712814 | 3.828321 | 3.944335 | 4.10528 | 4.280963 |
|  | 10 | 2.560389 | 2.663627 | 2.770456 | 2.85607 | 2.949241 | 3.095885 | 3.252457 | 3.410373 | 3.560758 | 3.657587 | 3.747459 | 3.860835 | 3.971734 |
|  | 15 | 2.501343 | 2.586872 | 2.67952 | 2.756754 | 2.843621 | 2.985316 | 3.140827 | 3.298227 | 3.445015 | 3.53665 | 3.619203 | 3.71961 | 3.813673 |
|  | 20 | 2.42325 | 2.502974 | 2.590961 | 2.665503 | 2.750495 | 2.891254 | 3.047707 | 3.206674 | 3.354132 | 3.445292 | 3.526608 | 3.624284 | 3.714422 |
|  | 30 | 2.301928 | 2.374661 | 2.456515 | 2.527058 | 2.60869 | 2.746225 | 2.901527 | 3.060568 | 3.207957 | 3.298528 | 3.378745 | 3.474184 | 3.561207 |
|  | 40 | 2.260277 | 2.327423 | 2.403861 | 2.470416 | 2.548134 | 2.680525 | 2.831674 | 2.987572 | 3.132402 | 3.221316 | 3.299893 | 3.393058 | 3.477602 |
|  | 50 | 2.247568 | 2.31207 | 2.386001 | 2.45078 | 2.526872 | 2.657482 | 2.807901 | 2.964197 | 3.110167 | 3.20004 | 3.279573 | 3.373954 | 3.459634 |
|  | 60 | 2.249919 | 2.314313 | 2.388509 | 2.45385 | 2.53098 | 2.664281 | 2.819125 | 2.981394 | 3.13409 | 3.228607 | 3.312548 | 3.412496 | 3.503526 |
|  | 70 | 2.22336 | 2.288368 | 2.363656 | 2.430296 | 2.509358 | 2.646999 | 2.808437 | 2.979347 | 3.14174 | 3.242997 | 3.333385 | 3.441566 | 3.540609 |
|  | 80 | 2.206238 | 2.272685 | 2.350049 | 2.41889 | 2.501006 | 2.645103 | 2.815945 | 2.99895 | 3.174852 | 3.285518 | 3.384941 | 3.504718 | 3.615121 |
|  | 90 | 2.202927 | 2.271481 | 2.351744 | 2.42357 | 2.509749 | 2.662299 | 2.845359 | 3.044103 | 3.237726 | 3.360848 | 3.472321 | 3.607699 | 3.733532 |
| **L_rostralmiddlefrontal** | 5 | 2.459017 | 2.517096 | 2.580075 | 2.631985 | 2.688907 | 2.77604 | 2.86002 | 2.945325 | 3.03673 | 3.098109 | 3.155273 | 3.226209 | 3.293218 |
|  | 10 | 2.323537 | 2.370723 | 2.423314 | 2.468014 | 2.518777 | 2.601228 | 2.688689 | 2.77763 | 2.864276 | 2.919036 | 2.968179 | 3.02714 | 3.081128 |
|  | 15 | 2.199476 | 2.241534 | 2.289065 | 2.330101 | 2.377548 | 2.457004 | 2.54535 | 2.635235 | 2.718794 | 2.769974 | 2.815039 | 2.868193 | 2.916107 |
|  | 20 | 2.116078 | 2.157407 | 2.204198 | 2.244677 | 2.291588 | 2.370459 | 2.458678 | 2.548427 | 2.631345 | 2.681917 | 2.726332 | 2.778597 | 2.825609 |
|  | 30 | 2.035012 | 2.076113 | 2.122636 | 2.162875 | 2.209503 | 2.287886 | 2.375551 | 2.464661 | 2.546859 | 2.596935 | 2.640881 | 2.692556 | 2.739005 |
|  | 40 | 2.005554 | 2.045517 | 2.090852 | 2.130167 | 2.175864 | 2.253106 | 2.340219 | 2.428659 | 2.509362 | 2.558159 | 2.600784 | 2.650696 | 2.695386 |
|  | 50 | 1.975155 | 2.017174 | 2.06443 | 2.105015 | 2.151675 | 2.229097 | 2.314039 | 2.40014 | 2.480695 | 2.530232 | 2.573943 | 2.625587 | 2.672202 |
|  | 60 | 1.924422 | 1.972641 | 2.025851 | 2.070595 | 2.120814 | 2.200829 | 2.283269 | 2.366705 | 2.449641 | 2.502708 | 2.550668 | 2.608558 | 2.661839 |
|  | 70 | 1.850273 | 1.902485 | 1.959637 | 2.007265 | 2.06018 | 2.143073 | 2.226201 | 2.310254 | 2.395954 | 2.451676 | 2.502526 | 2.564437 | 2.62187 |
|  | 80 | 1.863322 | 1.905181 | 1.953052 | 1.994956 | 2.044203 | 2.129066 | 2.227528 | 2.327128 | 2.414839 | 2.466557 | 2.511047 | 2.562428 | 2.607852 |
|  | 90 | 1.901978 | 1.928987 | 1.962251 | 1.993953 | 2.035118 | 2.11915 | 2.238733 | 2.359754 | 2.446554 | 2.48962 | 2.523034 | 2.558331 | 2.58717 |
| **L_superiorfrontal** | 5 | 2.804955 | 2.864523 | 2.927878 | 2.979857 | 3.037523 | 3.130111 | 3.230249 | 3.330836 | 3.424642 | 3.483463 | 3.536746 | 3.602032 | 3.66376 |
|  | 10 | 2.686056 | 2.743301 | 2.804383 | 2.854635 | 2.910513 | 3.000448 | 3.097892 | 3.195781 | 3.286922 | 3.343939 | 3.395473 | 3.458447 | 3.5178 |
|  | 15 | 2.580576 | 2.635812 | 2.69493 | 2.743694 | 2.798034 | 2.885694 | 2.980838 | 3.076424 | 3.165282 | 3.22075 | 3.270779 | 3.331758 | 3.389059 |
|  | 20 | 2.497212 | 2.550935 | 2.608601 | 2.656283 | 2.709528 | 2.795609 | 2.88919 | 2.983215 | 3.070494 | 3.124863 | 3.173805 | 3.233317 | 3.28908 |
|  | 30 | 2.390645 | 2.442706 | 2.498886 | 2.545552 | 2.597858 | 2.682764 | 2.775344 | 2.86838 | 2.954512 | 3.007964 | 3.055907 | 3.113946 | 3.168048 |
|  | 40 | 2.32661 | 2.378027 | 2.433778 | 2.480277 | 2.532573 | 2.617769 | 2.710918 | 2.804542 | 2.891016 | 2.9445 | 2.992318 | 3.049979 | 3.103478 |
|  | 50 | 2.28021 | 2.331466 | 2.387282 | 2.434007 | 2.486715 | 2.572865 | 2.667286 | 2.762207 | 2.849698 | 2.90365 | 2.951748 | 3.009544 | 3.062947 |
|  | 60 | 2.235259 | 2.286475 | 2.342464 | 2.389487 | 2.442678 | 2.529871 | 2.625647 | 2.72195 | 2.810554 | 2.865047 | 2.913504 | 2.97155 | 3.024986 |
|  | 70 | 2.164421 | 2.215069 | 2.270629 | 2.317432 | 2.370503 | 2.45773 | 2.553735 | 2.65029 | 2.738983 | 2.793403 | 2.841686 | 2.899363 | 2.952284 |
|  | 80 | 2.122673 | 2.173487 | 2.229405 | 2.276635 | 2.330309 | 2.418739 | 2.516244 | 2.61433 | 2.704304 | 2.759395 | 2.808175 | 2.866303 | 2.919481 |
|  | 90 | 2.091088 | 2.142376 | 2.198976 | 2.246896 | 2.301465 | 2.391562 | 2.491071 | 2.591196 | 2.682928 | 2.738992 | 2.788547 | 2.847467 | 2.901229 |
| **L_superiorparietal** | 5 | 2.237876 | 2.27749 | 2.322049 | 2.360373 | 2.404558 | 2.478388 | 2.56042 | 2.642451 | 2.716281 | 2.760466 | 2.79879 | 2.843349 | 2.882963 |
|  | 10 | 2.097155 | 2.136491 | 2.180738 | 2.218792 | 2.262666 | 2.335978 | 2.417432 | 2.498887 | 2.572199 | 2.616073 | 2.654127 | 2.698373 | 2.737709 |
|  | 15 | 1.975267 | 2.014325 | 2.058259 | 2.096045 | 2.139609 | 2.212404 | 2.293284 | 2.374164 | 2.446958 | 2.490523 | 2.528309 | 2.572243 | 2.611301 |
|  | 20 | 1.906801 | 1.945571 | 1.989181 | 2.026687 | 2.069931 | 2.142188 | 2.222471 | 2.302754 | 2.375011 | 2.418254 | 2.455761 | 2.499371 | 2.538141 |
|  | 30 | 1.84834 | 1.886517 | 1.929459 | 1.966392 | 2.008973 | 2.080124 | 2.159179 | 2.238233 | 2.309384 | 2.351965 | 2.388898 | 2.43184 | 2.470017 |
|  | 40 | 1.820659 | 1.858493 | 1.901051 | 1.937653 | 1.979853 | 2.050367 | 2.128713 | 2.207059 | 2.277573 | 2.319773 | 2.356375 | 2.398933 | 2.436768 |
|  | 50 | 1.802226 | 1.840136 | 1.882779 | 1.919455 | 1.961739 | 2.032394 | 2.110897 | 2.1894 | 2.260055 | 2.302339 | 2.339015 | 2.381658 | 2.419568 |
|  | 60 | 1.782245 | 1.820726 | 1.86401 | 1.901237 | 1.944157 | 2.015875 | 2.095558 | 2.175242 | 2.246959 | 2.28988 | 2.327106 | 2.370391 | 2.408871 |
|  | 70 | 1.730574 | 1.770101 | 1.814563 | 1.852803 | 1.896891 | 1.97056 | 2.052412 | 2.134264 | 2.207933 | 2.252021 | 2.290261 | 2.334723 | 2.374251 |
|  | 80 | 1.671816 | 1.712609 | 1.758495 | 1.797959 | 1.843459 | 1.919487 | 2.003959 | 2.088432 | 2.16446 | 2.209959 | 2.249423 | 2.295309 | 2.336102 |
|  | 90 | 1.582929 | 1.625062 | 1.672456 | 1.713217 | 1.760212 | 1.838738 | 1.925986 | 2.013235 | 2.091761 | 2.138756 | 2.179517 | 2.226911 | 2.269044 |
| **L_superiortemporal** | 5 | 2.478062 | 2.583253 | 2.686159 | 2.764586 | 2.845794 | 2.964 | 3.075536 | 3.175588 | 3.261133 | 3.310426 | 3.352042 | 3.399125 | 3.439838 |
|  | 10 | 2.49369 | 2.574969 | 2.659246 | 2.726506 | 2.798806 | 2.90869 | 3.017234 | 3.117881 | 3.205781 | 3.257171 | 3.30097 | 3.350974 | 3.394591 |
|  | 15 | 2.487442 | 2.552382 | 2.622562 | 2.680623 | 2.745042 | 2.84691 | 2.952207 | 3.053375 | 3.143948 | 3.197843 | 3.244324 | 3.298003 | 3.345356 |
|  | 20 | 2.465679 | 2.519951 | 2.580249 | 2.631437 | 2.689629 | 2.784708 | 2.887018 | 2.988663 | 3.081943 | 3.138479 | 3.187867 | 3.245632 | 3.297245 |
|  | 30 | 2.394379 | 2.443859 | 2.499473 | 2.547233 | 2.602171 | 2.693512 | 2.794142 | 2.895472 | 2.988674 | 3.045304 | 3.094891 | 3.153055 | 3.205192 |
|  | 40 | 2.325697 | 2.378424 | 2.437122 | 2.487097 | 2.544135 | 2.638071 | 2.740475 | 2.841477 | 2.931721 | 2.985427 | 3.031818 | 3.085527 | 3.133065 |
|  | 50 | 2.270116 | 2.325623 | 2.387017 | 2.438999 | 2.498057 | 2.594841 | 2.699862 | 2.801981 | 2.891203 | 2.94348 | 2.988194 | 3.039491 | 3.084502 |
|  | 60 | 2.240268 | 2.291288 | 2.348636 | 2.397959 | 2.454876 | 2.550259 | 2.656819 | 2.762407 | 2.8553 | 2.910014 | 2.957 | 3.011134 | 3.058849 |
|  | 70 | 2.169433 | 2.216331 | 2.26976 | 2.316338 | 2.370832 | 2.464049 | 2.571095 | 2.679115 | 2.774798 | 2.831447 | 2.880294 | 2.936817 | 2.986871 |
|  | 80 | 2.049483 | 2.100227 | 2.157409 | 2.206756 | 2.263951 | 2.360613 | 2.469978 | 2.577637 | 2.670024 | 2.723504 | 2.768953 | 2.820827 | 2.866159 |
|  | 90 | 1.90021 | 1.960724 | 2.026925 | 2.082547 | 2.145469 | 2.248539 | 2.360907 | 2.466728 | 2.553433 | 2.602074 | 2.642602 | 2.688021 | 2.727031 |
| **L_supramarginal** | 5 | 2.465945 | 2.558176 | 2.64568 | 2.710895 | 2.777646 | 2.875302 | 2.971688 | 3.062788 | 3.145385 | 3.196932 | 3.243842 | 3.301916 | 3.357672 |
|  | 10 | 2.409651 | 2.477613 | 2.547007 | 2.601925 | 2.660915 | 2.751778 | 2.845318 | 2.934947 | 3.015091 | 3.063815 | 3.10701 | 3.158791 | 3.20666 |
|  | 15 | 2.331006 | 2.387029 | 2.446707 | 2.495654 | 2.549816 | 2.636055 | 2.72749 | 2.816341 | 2.895646 | 2.943362 | 2.985164 | 3.0345 | 3.079245 |
|  | 20 | 2.266479 | 2.315896 | 2.369938 | 2.415284 | 2.466458 | 2.549838 | 2.640251 | 2.729356 | 2.809272 | 2.857281 | 2.899177 | 2.948329 | 2.992545 |
|  | 30 | 2.193275 | 2.237351 | 2.286773 | 2.329178 | 2.377994 | 2.459498 | 2.550177 | 2.641261 | 2.723826 | 2.773598 | 2.817031 | 2.867884 | 2.91345 |
|  | 40 | 2.144594 | 2.188242 | 2.237452 | 2.279881 | 2.328929 | 2.411227 | 2.503223 | 2.595881 | 2.67991 | 2.730511 | 2.774602 | 2.826118 | 2.872153 |
|  | 50 | 2.0996 | 2.144203 | 2.194411 | 2.237624 | 2.287487 | 2.370912 | 2.463775 | 2.556841 | 2.640794 | 2.691132 | 2.734853 | 2.785763 | 2.831093 |
|  | 60 | 2.062317 | 2.107469 | 2.158261 | 2.201949 | 2.252325 | 2.336518 | 2.430094 | 2.52371 | 2.608008 | 2.658478 | 2.702267 | 2.753199 | 2.798495 |
|  | 70 | 1.99365 | 2.039221 | 2.090416 | 2.134393 | 2.185036 | 2.269522 | 2.363201 | 2.45669 | 2.540679 | 2.590877 | 2.634378 | 2.684913 | 2.7298 |
|  | 80 | 1.929588 | 1.976544 | 2.02911 | 2.074113 | 2.125769 | 2.211559 | 2.306142 | 2.399996 | 2.483876 | 2.533818 | 2.576985 | 2.627005 | 2.671324 |
|  | 90 | 1.882287 | 1.931159 | 1.985658 | 2.032147 | 2.085321 | 2.173212 | 2.269536 | 2.364556 | 2.449032 | 2.499139 | 2.542338 | 2.592269 | 2.636399 |
| **L_frontalpole** | 5 | 2.119581 | 2.311976 | 2.501494 | 2.646754 | 2.799017 | 3.029688 | 3.271171 | 3.520429 | 3.774032 | 3.950524 | 4.12622 | 4.367127 | 4.626595 |
|  | 10 | 2.27929 | 2.374786 | 2.483892 | 2.579183 | 2.690749 | 2.881345 | 3.099478 | 3.324659 | 3.5337 | 3.661837 | 3.774893 | 3.908662 | 4.029748 |
|  | 15 | 2.211718 | 2.302682 | 2.407026 | 2.498477 | 2.605875 | 2.790006 | 3.001435 | 3.220057 | 3.422965 | 3.547178 | 3.656601 | 3.785797 | 3.902433 |
|  | 20 | 2.153752 | 2.241943 | 2.343208 | 2.432049 | 2.536482 | 2.715771 | 2.921989 | 3.135583 | 3.334124 | 3.455795 | 3.563056 | 3.689787 | 3.804275 |
|  | 30 | 2.067096 | 2.151103 | 2.247734 | 2.332657 | 2.432655 | 2.604756 | 2.803352 | 3.009754 | 3.202228 | 3.320467 | 3.424876 | 3.548446 | 3.660268 |
|  | 40 | 2.027737 | 2.109535 | 2.203785 | 2.286755 | 2.38462 | 2.553462 | 2.748933 | 2.952788 | 3.143512 | 3.260967 | 3.364865 | 3.488047 | 3.599715 |
|  | 50 | 2.037638 | 2.119303 | 2.213538 | 2.296621 | 2.394771 | 2.564495 | 2.761607 | 2.96789 | 3.161554 | 3.281143 | 3.387134 | 3.513048 | 3.627429 |
|  | 60 | 1.973629 | 2.054267 | 2.146959 | 2.228428 | 2.324454 | 2.490206 | 2.682749 | 2.884978 | 3.076125 | 3.195027 | 3.301081 | 3.42801 | 3.544305 |
|  | 70 | 1.871499 | 1.962543 | 2.063573 | 2.149729 | 2.248765 | 2.415256 | 2.605269 | 2.805688 | 3.000273 | 3.125646 | 3.241237 | 3.385282 | 3.523739 |
|  | 80 | 1.823213 | 1.932435 | 2.048909 | 2.144821 | 2.25186 | 2.426197 | 2.620874 | 2.82714 | 3.033895 | 3.172828 | 3.306106 | 3.480461 | 3.657999 |
|  | 90 | 1.804108 | 1.913851 | 2.030596 | 2.12653 | 2.23342 | 2.407298 | 2.601573 | 2.808207 | 3.016807 | 3.15806 | 3.294488 | 3.474413 | 3.659366 |
| **L_temporalpole** | 5 | 2.384281 | 2.591767 | 2.795209 | 2.952165 | 3.118186 | 3.370776 | 3.626431 | 3.849333 | 4.018761 | 4.109428 | 4.182908 | 4.263264 | 4.330763 |
|  | 10 | 2.508758 | 2.673915 | 2.845762 | 2.984526 | 3.136582 | 3.377026 | 3.629966 | 3.858292 | 4.037041 | 4.134444 | 4.214225 | 4.302295 | 4.376907 |
|  | 15 | 2.579636 | 2.722703 | 2.87672 | 3.004522 | 3.14772 | 3.380047 | 3.631199 | 3.864216 | 4.051343 | 4.154976 | 4.240684 | 4.336119 | 4.417618 |
|  | 20 | 2.596269 | 2.732821 | 2.881476 | 3.005968 | 3.146517 | 3.376526 | 3.627457 | 3.863248 | 4.055446 | 4.162935 | 4.252355 | 4.35245 | 4.438342 |
|  | 30 | 2.517176 | 2.670038 | 2.832849 | 2.96661 | 3.115036 | 3.352465 | 3.604602 | 3.838237 | 4.028555 | 4.135 | 4.223525 | 4.322561 | 4.407479 |
|  | 40 | 2.467556 | 2.63397 | 2.807389 | 2.94726 | 3.099964 | 3.33917 | 3.587052 | 3.814155 | 3.999446 | 4.103219 | 4.189564 | 4.286174 | 4.369002 |
|  | 50 | 2.573835 | 2.711047 | 2.85927 | 2.982302 | 3.119775 | 3.340795 | 3.575958 | 3.798422 | 3.98647 | 4.094251 | 4.185198 | 4.288254 | 4.377649 |
|  | 60 | 2.668625 | 2.778014 | 2.900656 | 3.005838 | 3.126817 | 3.328443 | 3.551897 | 3.77373 | 3.971133 | 4.088334 | 4.189471 | 4.306506 | 4.41008 |
|  | 70 | 2.645077 | 2.749072 | 2.866729 | 2.968449 | 3.08627 | 3.284248 | 3.505566 | 3.729624 | 3.934819 | 4.059153 | 4.167869 | 4.295251 | 4.409342 |
|  | 80 | 2.500474 | 2.622476 | 2.757594 | 2.872024 | 3.001905 | 3.213935 | 3.442243 | 3.667516 | 3.871399 | 3.993807 | 4.100107 | 4.223769 | 4.333704 |
|  | 90 | 2.292347 | 2.450422 | 2.617994 | 2.754514 | 2.904129 | 3.137506 | 3.375547 | 3.601545 | 3.801749 | 3.920093 | 4.021757 | 4.138757 | 4.241651 |
| **L_transversetemporal** | 5 | 2.090048 | 2.164861 | 2.249014 | 2.32139 | 2.404835 | 2.544268 | 2.699188 | 2.854109 | 2.993542 | 3.076987 | 3.149363 | 3.233515 | 3.308329 |
|  | 10 | 2.03031 | 2.104225 | 2.187368 | 2.258876 | 2.341319 | 2.479078 | 2.632139 | 2.785199 | 2.922959 | 3.005402 | 3.076909 | 3.160052 | 3.233968 |
|  | 15 | 1.969332 | 2.042478 | 2.124756 | 2.195519 | 2.277105 | 2.413431 | 2.564899 | 2.716367 | 2.852692 | 2.934278 | 3.005041 | 3.087319 | 3.160465 |
|  | 20 | 1.909181 | 1.981765 | 2.06341 | 2.133629 | 2.214587 | 2.349864 | 2.500167 | 2.650469 | 2.785747 | 2.866705 | 2.936924 | 3.018568 | 3.091152 |
|  | 30 | 1.806624 | 1.878549 | 1.959452 | 2.029033 | 2.109256 | 2.243304 | 2.392242 | 2.54118 | 2.675228 | 2.755451 | 2.825032 | 2.905935 | 2.97786 |
|  | 40 | 1.731964 | 1.803603 | 1.884185 | 1.95349 | 2.033394 | 2.16691 | 2.315256 | 2.463602 | 2.597118 | 2.677022 | 2.746327 | 2.826909 | 2.898548 |
|  | 50 | 1.674724 | 1.746603 | 1.827455 | 1.896993 | 1.977164 | 2.111128 | 2.259972 | 2.408815 | 2.542779 | 2.622951 | 2.692488 | 2.77334 | 2.845219 |
|  | 60 | 1.629634 | 1.702335 | 1.784112 | 1.854445 | 1.935534 | 2.07103 | 2.221576 | 2.372123 | 2.507619 | 2.588708 | 2.659041 | 2.740818 | 2.813519 |
|  | 70 | 1.572392 | 1.646482 | 1.729822 | 1.801499 | 1.884137 | 2.022223 | 2.175646 | 2.329069 | 2.467155 | 2.549794 | 2.62147 | 2.70481 | 2.778901 |
|  | 80 | 1.503541 | 1.579506 | 1.664955 | 1.738445 | 1.823174 | 1.964754 | 2.122059 | 2.279364 | 2.420944 | 2.505673 | 2.579164 | 2.664612 | 2.740577 |
|  | 90 | 1.430546 | 1.508668 | 1.596543 | 1.67212 | 1.759256 | 1.904856 | 2.066629 | 2.228401 | 2.374001 | 2.461137 | 2.536714 | 2.624589 | 2.702712 |
| **L_insula** | 5 | 2.928054 | 2.998159 | 3.073162 | 3.134799 | 3.202963 | 3.310888 | 3.423387 | 3.528313 | 3.616429 | 3.666674 | 3.708906 | 3.756563 | 3.797724 |
|  | 10 | 2.860649 | 2.922395 | 2.989254 | 3.044748 | 3.10662 | 3.205443 | 3.309315 | 3.407892 | 3.492743 | 3.541929 | 3.583689 | 3.631245 | 3.672665 |
|  | 15 | 2.801911 | 2.858249 | 2.919772 | 2.9712 | 3.028867 | 3.121528 | 3.219435 | 3.313767 | 3.396896 | 3.445858 | 3.487839 | 3.536083 | 3.578459 |
|  | 20 | 2.752954 | 2.807051 | 2.866434 | 2.916282 | 2.972357 | 3.062713 | 3.158332 | 3.251575 | 3.335519 | 3.385692 | 3.42911 | 3.479428 | 3.523981 |
|  | 30 | 2.642746 | 2.696992 | 2.756752 | 2.807028 | 2.863626 | 2.954656 | 3.050383 | 3.145215 | 3.233763 | 3.288031 | 3.33574 | 3.391846 | 3.442214 |
|  | 40 | 2.563881 | 2.620025 | 2.681822 | 2.733709 | 2.791932 | 2.884901 | 2.981356 | 3.077803 | 3.170749 | 3.22895 | 3.280813 | 3.342574 | 3.39868 |
|  | 50 | 2.528379 | 2.585245 | 2.647607 | 2.699726 | 2.75787 | 2.849717 | 2.943273 | 3.037287 | 3.130486 | 3.189954 | 3.243573 | 3.308123 | 3.367363 |
|  | 60 | 2.524735 | 2.584514 | 2.649556 | 2.703451 | 2.763002 | 2.855601 | 2.947603 | 3.03997 | 3.133684 | 3.194353 | 3.249532 | 3.316475 | 3.378342 |
|  | 70 | 2.453916 | 2.520732 | 2.592626 | 2.651511 | 2.715781 | 2.813804 | 2.908379 | 3.002893 | 3.100725 | 3.164798 | 3.223453 | 3.295002 | 3.361433 |
|  | 80 | 2.341972 | 2.42223 | 2.507515 | 2.576486 | 2.650775 | 2.761804 | 2.865704 | 2.969012 | 3.078116 | 3.150378 | 3.216941 | 3.298554 | 3.374656 |
|  | 90 | 2.202265 | 2.301181 | 2.404753 | 2.487306 | 2.574932 | 2.703082 | 2.819203 | 2.933995 | 3.057661 | 3.140445 | 3.21714 | 3.311615 | 3.400045 |
| **R_bankssts** | 5 | 2.539483 | 2.598175 | 2.664194 | 2.720974 | 2.786438 | 2.895826 | 3.017363 | 3.138901 | 3.248288 | 3.313752 | 3.370532 | 3.436551 | 3.495244 |
|  | 10 | 2.410353 | 2.468797 | 2.534536 | 2.591076 | 2.656262 | 2.765186 | 2.886208 | 3.00723 | 3.116154 | 3.18134 | 3.23788 | 3.303619 | 3.362063 |
|  | 15 | 2.289311 | 2.347511 | 2.412975 | 2.469278 | 2.534192 | 2.64266 | 2.763176 | 2.883692 | 2.99216 | 3.057074 | 3.113377 | 3.178842 | 3.237041 |
|  | 20 | 2.202496 | 2.26046 | 2.325661 | 2.381737 | 2.446389 | 2.554419 | 2.674449 | 2.794479 | 2.90251 | 2.967162 | 3.023238 | 3.088438 | 3.146402 |
|  | 30 | 2.110596 | 2.168149 | 2.232887 | 2.288566 | 2.352759 | 2.460024 | 2.579202 | 2.698381 | 2.805646 | 2.869839 | 2.925517 | 2.990256 | 3.047809 |
|  | 40 | 2.060502 | 2.117795 | 2.182239 | 2.237665 | 2.301567 | 2.408345 | 2.526983 | 2.645621 | 2.752399 | 2.816301 | 2.871727 | 2.936171 | 2.993464 |
|  | 50 | 2.022982 | 2.080175 | 2.144508 | 2.199837 | 2.263629 | 2.370222 | 2.488654 | 2.607086 | 2.713679 | 2.777471 | 2.832801 | 2.897133 | 2.954326 |
|  | 60 | 1.990931 | 2.048144 | 2.1125 | 2.167849 | 2.231663 | 2.338293 | 2.456767 | 2.575241 | 2.681871 | 2.745685 | 2.801034 | 2.86539 | 2.922603 |
|  | 70 | 1.940286 | 1.997619 | 2.06211 | 2.117575 | 2.181524 | 2.288378 | 2.407102 | 2.525825 | 2.632679 | 2.696628 | 2.752093 | 2.816584 | 2.873917 |
|  | 80 | 1.885364 | 1.942845 | 2.007501 | 2.06311 | 2.127222 | 2.234352 | 2.353381 | 2.472409 | 2.579539 | 2.643652 | 2.69926 | 2.763916 | 2.821397 |
|  | 90 | 1.826386 | 1.88401 | 1.948828 | 2.004575 | 2.068848 | 2.176244 | 2.29557 | 2.414895 | 2.522292 | 2.586564 | 2.642311 | 2.707129 | 2.764753 |
| **R_caudalanteriorcingulate** | 5 | 2.370855 | 2.437056 | 2.513173 | 2.58017 | 2.659349 | 2.796856 | 2.958501 | 3.131292 | 3.2982 | 3.40409 | 3.500035 | 3.616882 | 3.726023 |
|  | 10 | 2.288592 | 2.352954 | 2.426896 | 2.491936 | 2.56876 | 2.702101 | 2.858801 | 3.026341 | 3.188301 | 3.29115 | 3.384422 | 3.498135 | 3.604486 |
|  | 15 | 2.218519 | 2.281379 | 2.353533 | 2.416953 | 2.491818 | 2.621681 | 2.774237 | 2.937384 | 3.095225 | 3.19556 | 3.286639 | 3.397807 | 3.50192 |
|  | 20 | 2.156492 | 2.218085 | 2.288718 | 2.350752 | 2.423933 | 2.550788 | 2.699754 | 2.859097 | 3.013394 | 3.111584 | 3.200807 | 3.309846 | 3.412116 |
|  | 30 | 2.069983 | 2.130303 | 2.199334 | 2.259856 | 2.331153 | 2.454571 | 2.59939 | 2.754401 | 2.904822 | 3.000794 | 3.08821 | 3.195351 | 3.296187 |
|  | 40 | 2.034059 | 2.094979 | 2.164541 | 2.225415 | 2.297019 | 2.420789 | 2.565933 | 2.721459 | 2.872796 | 2.969668 | 3.058166 | 3.167021 | 3.269902 |
|  | 50 | 2.000228 | 2.06239 | 2.133197 | 2.195037 | 2.267664 | 2.393026 | 2.53998 | 2.697703 | 2.851732 | 2.950734 | 3.041512 | 3.153665 | 3.260213 |
|  | 60 | 1.981238 | 2.045883 | 2.119331 | 2.183343 | 2.258402 | 2.387788 | 2.539459 | 2.702628 | 2.862725 | 2.966161 | 3.061442 | 3.179804 | 3.292969 |
|  | 70 | 1.956565 | 2.024235 | 2.100902 | 2.167565 | 2.245596 | 2.379916 | 2.537396 | 2.707319 | 2.874995 | 2.984006 | 3.08498 | 3.21124 | 3.332886 |
|  | 80 | 1.953694 | 2.025968 | 2.107587 | 2.178369 | 2.261059 | 2.403172 | 2.569833 | 2.750301 | 2.929593 | 3.047023 | 3.156514 | 3.294497 | 3.428659 |
|  | 90 | 1.981295 | 2.06023 | 2.149036 | 2.225812 | 2.31529 | 2.468766 | 2.648777 | 2.844474 | 3.040415 | 3.169857 | 3.291476 | 3.446154 | 3.59817 |
| **R_caudalmiddlefrontal** | 5 | 2.338333 | 2.420057 | 2.501585 | 2.564884 | 2.631808 | 2.733049 | 2.835425 | 2.932305 | 3.01835 | 3.07054 | 3.116796 | 3.172285 | 3.223663 |
|  | 10 | 2.303934 | 2.374882 | 2.447765 | 2.505742 | 2.568293 | 2.665152 | 2.765414 | 2.861883 | 2.948367 | 3.00101 | 3.047703 | 3.103691 | 3.15545 |
|  | 15 | 2.261376 | 2.323821 | 2.389526 | 2.442882 | 2.501476 | 2.594142 | 2.692198 | 2.788132 | 2.875041 | 2.928204 | 2.97545 | 3.032153 | 3.084571 |
|  | 20 | 2.217351 | 2.273436 | 2.333576 | 2.383232 | 2.438569 | 2.52767 | 2.62379 | 2.719269 | 2.806646 | 2.86037 | 2.90823 | 2.965755 | 3.018968 |
|  | 30 | 2.150385 | 2.199835 | 2.254098 | 2.299835 | 2.351748 | 2.43725 | 2.531742 | 2.627353 | 2.715846 | 2.770519 | 2.819294 | 2.877916 | 2.932068 |
|  | 40 | 2.109848 | 2.157084 | 2.209531 | 2.254201 | 2.30537 | 2.390571 | 2.485742 | 2.58268 | 2.67257 | 2.728033 | 2.777394 | 2.836505 | 2.890847 |
|  | 50 | 2.075267 | 2.121873 | 2.173979 | 2.218625 | 2.270027 | 2.356109 | 2.452739 | 2.551339 | 2.642626 | 2.698768 | 2.748562 | 2.807931 | 2.862218 |
|  | 60 | 2.035835 | 2.082977 | 2.135824 | 2.181198 | 2.233516 | 2.321216 | 2.419597 | 2.519664 | 2.611805 | 2.668149 | 2.717879 | 2.776839 | 2.830407 |
|  | 70 | 1.981633 | 2.029658 | 2.083513 | 2.129747 | 2.183027 | 2.272199 | 2.371884 | 2.472728 | 2.564941 | 2.620971 | 2.670172 | 2.728176 | 2.780546 |
|  | 80 | 1.932826 | 1.982114 | 2.037343 | 2.08471 | 2.139223 | 2.230239 | 2.331565 | 2.433493 | 2.52609 | 2.582029 | 2.630932 | 2.688305 | 2.739831 |
|  | 90 | 1.888893 | 1.939816 | 1.996783 | 2.045554 | 2.101573 | 2.19481 | 2.298114 | 2.401428 | 2.494691 | 2.550733 | 2.599529 | 2.656531 | 2.70749 |
| **R_cuneus** | 5 | 1.873333 | 1.917491 | 1.968084 | 2.012386 | 2.06438 | 2.153483 | 2.255809 | 2.361712 | 2.460153 | 2.520509 | 2.573745 | 2.636692 | 2.693609 |
|  | 10 | 1.734401 | 1.775336 | 1.822299 | 1.863478 | 1.911872 | 1.994969 | 2.090649 | 2.189954 | 2.282516 | 2.339385 | 2.389621 | 2.449109 | 2.502982 |
|  | 15 | 1.635838 | 1.674492 | 1.718897 | 1.757886 | 1.803768 | 1.88271 | 1.973848 | 2.068713 | 2.157383 | 2.211978 | 2.26028 | 2.317566 | 2.369527 |
|  | 20 | 1.583544 | 1.621002 | 1.664091 | 1.701975 | 1.746619 | 1.823584 | 1.912681 | 2.005694 | 2.092883 | 2.146686 | 2.19436 | 2.250994 | 2.302448 |
|  | 30 | 1.530561 | 1.566828 | 1.608661 | 1.645541 | 1.689123 | 1.764562 | 1.85238 | 1.944612 | 2.031583 | 2.0855 | 2.133435 | 2.190569 | 2.242659 |
|  | 40 | 1.497365 | 1.532892 | 1.573982 | 1.610307 | 1.653352 | 1.728175 | 1.81577 | 1.908349 | 1.996193 | 2.050917 | 2.09974 | 2.158144 | 2.211592 |
|  | 50 | 1.47154 | 1.506486 | 1.547014 | 1.58294 | 1.625635 | 1.700162 | 1.787925 | 1.881286 | 1.970453 | 2.02629 | 2.076292 | 2.136339 | 2.191513 |
|  | 60 | 1.462817 | 1.497574 | 1.537994 | 1.573924 | 1.616745 | 1.691816 | 1.780752 | 1.876005 | 1.967605 | 2.025279 | 2.077134 | 2.139667 | 2.197375 |
|  | 70 | 1.435565 | 1.469682 | 1.509465 | 1.544927 | 1.587313 | 1.661948 | 1.750916 | 1.846874 | 1.939821 | 1.998683 | 2.051834 | 2.116217 | 2.175916 |
|  | 80 | 1.40974 | 1.443237 | 1.482403 | 1.517413 | 1.559382 | 1.633612 | 1.72266 | 1.819409 | 1.913832 | 1.974 | 2.028579 | 2.095013 | 2.156931 |
|  | 90 | 1.401619 | 1.434905 | 1.473929 | 1.50891 | 1.550967 | 1.625691 | 1.715917 | 1.81469 | 1.91186 | 1.974184 | 2.030998 | 2.100515 | 2.16567 |
| **R_entorhinal** | 5 | 2.337624 | 2.469743 | 2.615681 | 2.739835 | 2.882469 | 3.122632 | 3.39461 | 3.649308 | 3.84841 | 3.956531 | 4.04498 | 4.142589 | 4.22532 |
|  | 10 | 2.391864 | 2.525413 | 2.672806 | 2.797953 | 2.941277 | 3.181022 | 3.450091 | 3.70399 | 3.906988 | 4.018973 | 4.111422 | 4.214253 | 4.302018 |
|  | 15 | 2.433285 | 2.568601 | 2.717759 | 2.844118 | 2.988343 | 3.227948 | 3.494251 | 3.747296 | 3.954148 | 4.070023 | 4.166544 | 4.274739 | 4.367724 |
|  | 20 | 2.441509 | 2.578087 | 2.728477 | 2.855635 | 3.000353 | 3.239372 | 3.502723 | 3.754467 | 3.964302 | 4.083429 | 4.183441 | 4.29632 | 4.393927 |
|  | 30 | 2.39864 | 2.533087 | 2.681489 | 2.807124 | 2.950122 | 3.185846 | 3.444481 | 3.694043 | 3.906591 | 4.029041 | 4.13275 | 4.250718 | 4.353451 |
|  | 40 | 2.399915 | 2.529009 | 2.672495 | 2.794739 | 2.934699 | 3.167246 | 3.424857 | 3.67526 | 3.889467 | 4.013239 | 4.118275 | 4.237981 | 4.342427 |
|  | 50 | 2.435332 | 2.557879 | 2.694927 | 2.812377 | 2.947628 | 3.17423 | 3.42795 | 3.675826 | 3.88768 | 4.010008 | 4.113794 | 4.23207 | 4.335275 |
|  | 60 | 2.473786 | 2.594657 | 2.73006 | 2.846282 | 2.980318 | 3.20534 | 3.457923 | 3.705159 | 3.916707 | 4.038952 | 4.14272 | 4.261032 | 4.36432 |
|  | 70 | 2.41028 | 2.533981 | 2.672368 | 2.79102 | 2.927738 | 3.157045 | 3.414207 | 3.665334 | 3.879451 | 4.002881 | 4.107499 | 4.226621 | 4.330486 |
|  | 80 | 2.200722 | 2.331199 | 2.476783 | 2.60139 | 2.744844 | 2.985505 | 3.255817 | 3.518016 | 3.738431 | 3.86427 | 3.970311 | 4.090431 | 4.194673 |
|  | 90 | 1.93105 | 2.073246 | 2.231069 | 2.365663 | 2.520293 | 2.779561 | 3.071143 | 3.351112 | 3.581728 | 3.711583 | 3.820115 | 3.942167 | 4.047394 |
| **R_fusiform** | 5 | 2.417725 | 2.546046 | 2.662883 | 2.746856 | 2.829623 | 2.942896 | 3.042046 | 3.129774 | 3.207885 | 3.253937 | 3.29328 | 3.338206 | 3.37734 |
|  | 10 | 2.407869 | 2.499007 | 2.589193 | 2.658295 | 2.729872 | 2.833351 | 2.929191 | 3.016937 | 3.09631 | 3.14367 | 3.18447 | 3.231443 | 3.272695 |
|  | 15 | 2.381042 | 2.448698 | 2.519887 | 2.577275 | 2.639322 | 2.733772 | 2.826426 | 2.914565 | 2.995956 | 3.045298 | 3.088284 | 3.138334 | 3.182782 |
|  | 20 | 2.356294 | 2.411317 | 2.471324 | 2.521301 | 2.576987 | 2.665187 | 2.755989 | 2.845286 | 2.929229 | 2.980862 | 3.026328 | 3.07985 | 3.127925 |
|  | 30 | 2.28646 | 2.340389 | 2.399406 | 2.448774 | 2.504088 | 2.592606 | 2.685295 | 2.775913 | 2.858741 | 2.90875 | 2.952292 | 3.003031 | 3.048181 |
|  | 40 | 2.22144 | 2.279102 | 2.341564 | 2.393391 | 2.4511 | 2.542952 | 2.638873 | 2.730697 | 2.811537 | 2.859113 | 2.899877 | 2.946684 | 2.98776 |
|  | 50 | 2.220517 | 2.270544 | 2.326214 | 2.3736 | 2.427684 | 2.516784 | 2.613997 | 2.709676 | 2.794747 | 2.845204 | 2.888697 | 2.938953 | 2.983355 |
|  | 60 | 2.225702 | 2.270026 | 2.320415 | 2.364219 | 2.41528 | 2.502008 | 2.60053 | 2.700526 | 2.791065 | 2.845511 | 2.892915 | 2.94826 | 2.99768 |
|  | 70 | 2.175351 | 2.218446 | 2.267682 | 2.310713 | 2.361163 | 2.447658 | 2.547221 | 2.648726 | 2.740179 | 2.794996 | 2.84264 | 2.898194 | 2.947752 |
|  | 80 | 2.123051 | 2.166836 | 2.21683 | 2.260516 | 2.311759 | 2.399774 | 2.501427 | 2.604221 | 2.695106 | 2.748848 | 2.795158 | 2.848721 | 2.896139 |
|  | 90 | 2.07078 | 2.116665 | 2.168814 | 2.214203 | 2.26727 | 2.358121 | 2.462689 | 2.566713 | 2.656212 | 2.708116 | 2.752291 | 2.802798 | 2.84702 |
| **R_inferiorparietal** | 5 | 2.404034 | 2.499725 | 2.591725 | 2.661196 | 2.732953 | 2.838031 | 2.938996 | 3.026811 | 3.096526 | 3.134873 | 3.166401 | 3.20127 | 3.230823 |
|  | 10 | 2.343748 | 2.410033 | 2.479078 | 2.534529 | 2.594623 | 2.687375 | 2.781273 | 2.867117 | 2.938486 | 2.978909 | 3.012733 | 3.050738 | 3.083421 |
|  | 15 | 2.264297 | 2.315329 | 2.370881 | 2.417205 | 2.469056 | 2.552257 | 2.640131 | 2.724238 | 2.797466 | 2.840247 | 2.876746 | 2.918501 | 2.955024 |
|  | 20 | 2.204296 | 2.248815 | 2.29824 | 2.340189 | 2.387898 | 2.465992 | 2.550353 | 2.633629 | 2.708828 | 2.753899 | 2.792995 | 2.838427 | 2.878773 |
|  | 30 | 2.113208 | 2.160571 | 2.212623 | 2.256325 | 2.305435 | 2.384273 | 2.467038 | 2.548644 | 2.624276 | 2.670384 | 2.710781 | 2.758133 | 2.800508 |
|  | 40 | 2.064192 | 2.114148 | 2.168722 | 2.214233 | 2.264989 | 2.345424 | 2.428197 | 2.509961 | 2.587529 | 2.635543 | 2.677995 | 2.728161 | 2.773388 |
|  | 50 | 2.051005 | 2.096412 | 2.146948 | 2.189837 | 2.238466 | 2.317243 | 2.400488 | 2.485041 | 2.567545 | 2.619732 | 2.666585 | 2.722813 | 2.774306 |
|  | 60 | 2.024088 | 2.066252 | 2.113822 | 2.15477 | 2.201903 | 2.280072 | 2.365516 | 2.453486 | 2.538649 | 2.592327 | 2.640472 | 2.698254 | 2.75122 |
|  | 70 | 1.948306 | 1.989322 | 2.035815 | 2.076108 | 2.122927 | 2.202066 | 2.291365 | 2.38164 | 2.463312 | 2.51239 | 2.55509 | 2.6049 | 2.649336 |
|  | 80 | 1.911438 | 1.949507 | 1.993279 | 2.031876 | 2.077676 | 2.158063 | 2.253794 | 2.349019 | 2.428207 | 2.47301 | 2.510591 | 2.553017 | 2.589746 |
|  | 90 | 1.8889 | 1.922225 | 1.961451 | 1.996975 | 2.040463 | 2.120939 | 2.223347 | 2.324447 | 2.402101 | 2.44342 | 2.476842 | 2.513406 | 2.544189 |
| **R_inferiortemporal** | 5 | 2.511328 | 2.600271 | 2.692045 | 2.76507 | 2.843461 | 2.962673 | 3.080874 | 3.18957 | 3.282685 | 3.336457 | 3.381956 | 3.433571 | 3.478339 |
|  | 10 | 2.467753 | 2.545939 | 2.628453 | 2.695344 | 2.768272 | 2.881208 | 2.995354 | 3.10229 | 3.195503 | 3.249963 | 3.296385 | 3.34941 | 3.3957 |
|  | 15 | 2.421319 | 2.490597 | 2.565117 | 2.626527 | 2.694441 | 2.801452 | 2.911692 | 3.016956 | 3.110391 | 3.165666 | 3.213164 | 3.267832 | 3.315906 |
|  | 20 | 2.380772 | 2.4432 | 2.511385 | 2.568348 | 2.632123 | 2.734186 | 2.841221 | 2.945291 | 3.03928 | 3.095567 | 3.144327 | 3.200883 | 3.250994 |
|  | 30 | 2.324262 | 2.381381 | 2.444533 | 2.497883 | 2.558234 | 2.656122 | 2.760417 | 2.863566 | 2.958333 | 3.015795 | 3.065989 | 3.124682 | 3.177101 |
|  | 40 | 2.285958 | 2.343246 | 2.406585 | 2.460094 | 2.520624 | 2.618807 | 2.723424 | 2.826888 | 2.921931 | 2.979555 | 3.029888 | 3.088741 | 3.141301 |
|  | 50 | 2.283993 | 2.339889 | 2.402127 | 2.455084 | 2.515438 | 2.614449 | 2.721614 | 2.828182 | 2.925588 | 2.984471 | 3.035833 | 3.095834 | 3.149388 |
|  | 60 | 2.284879 | 2.336128 | 2.394125 | 2.444304 | 2.502505 | 2.6006 | 2.710841 | 2.822115 | 2.922902 | 2.983508 | 3.036255 | 3.097799 | 3.152709 |
|  | 70 | 2.251971 | 2.297121 | 2.34919 | 2.395154 | 2.449637 | 2.544719 | 2.656955 | 2.772423 | 2.875578 | 2.937049 | 2.990318 | 3.052289 | 3.107481 |
|  | 80 | 2.230529 | 2.269872 | 2.316142 | 2.357869 | 2.408529 | 2.500502 | 2.615346 | 2.736294 | 2.842834 | 2.905663 | 2.95983 | 3.022629 | 3.078441 |
|  | 90 | 2.219993 | 2.254524 | 2.295875 | 2.333928 | 2.381212 | 2.470525 | 2.58851 | 2.715954 | 2.826892 | 2.89163 | 2.947153 | 3.011295 | 3.068179 |
| **R_isthmuscingulate** | 5 | 2.445998 | 2.51256 | 2.58522 | 2.646345 | 2.715857 | 2.831524 | 2.96296 | 3.10266 | 3.24098 | 3.332178 | 3.418046 | 3.527807 | 3.636541 |
|  | 10 | 2.317394 | 2.379646 | 2.448536 | 2.507186 | 2.574556 | 2.687885 | 2.817696 | 2.955633 | 3.090992 | 3.179131 | 3.261127 | 3.364383 | 3.464849 |
|  | 15 | 2.202138 | 2.261354 | 2.327583 | 2.384484 | 2.450348 | 2.562064 | 2.690791 | 2.827525 | 2.960764 | 3.046686 | 3.125876 | 3.22445 | 3.319035 |
|  | 20 | 2.114678 | 2.17182 | 2.236238 | 2.29196 | 2.356825 | 2.467502 | 2.595547 | 2.731438 | 2.863065 | 2.947278 | 3.024311 | 3.119319 | 3.20948 |
|  | 30 | 2.027371 | 2.081934 | 2.144039 | 2.198194 | 2.261641 | 2.370583 | 2.497001 | 2.63065 | 2.758726 | 2.839626 | 2.91277 | 3.001722 | 3.084751 |
|  | 40 | 1.982105 | 2.035241 | 2.096009 | 2.149197 | 2.211685 | 2.319215 | 2.443956 | 2.575235 | 2.699971 | 2.778036 | 2.848048 | 2.932388 | 3.01025 |
|  | 50 | 1.940315 | 1.993554 | 2.054596 | 2.108129 | 2.171107 | 2.27957 | 2.405284 | 2.53713 | 2.661677 | 2.739151 | 2.808272 | 2.891036 | 2.966915 |
|  | 60 | 1.88643 | 1.940793 | 2.003213 | 2.058014 | 2.122528 | 2.233665 | 2.36237 | 2.497019 | 2.623715 | 2.702212 | 2.772011 | 2.855263 | 2.931254 |
|  | 70 | 1.824875 | 1.88084 | 1.945144 | 2.001624 | 2.068126 | 2.182668 | 2.315178 | 2.453518 | 2.5833 | 2.663477 | 2.7346 | 2.819203 | 2.896195 |
|  | 80 | 1.770357 | 1.828937 | 1.896263 | 1.9554 | 2.025023 | 2.144883 | 2.283383 | 2.4277 | 2.562755 | 2.646 | 2.719708 | 2.807209 | 2.886658 |
|  | 90 | 1.737008 | 1.799547 | 1.871408 | 1.934508 | 2.008762 | 2.13648 | 2.283826 | 2.43703 | 2.580046 | 2.668007 | 2.745761 | 2.837897 | 2.921392 |
| **R_lateraloccipital** | 5 | 2.180928 | 2.221622 | 2.267767 | 2.307768 | 2.354245 | 2.432758 | 2.521238 | 2.611021 | 2.692936 | 2.742459 | 2.785714 | 2.836359 | 2.881699 |
|  | 10 | 2.070088 | 2.109102 | 2.153362 | 2.191746 | 2.236364 | 2.311787 | 2.396854 | 2.483248 | 2.562136 | 2.609858 | 2.651558 | 2.700402 | 2.744149 |
|  | 15 | 1.984685 | 2.022462 | 2.06534 | 2.102543 | 2.145808 | 2.21899 | 2.3016 | 2.385574 | 2.462316 | 2.508768 | 2.549376 | 2.596962 | 2.639601 |
|  | 20 | 1.937895 | 1.975147 | 2.017451 | 2.054172 | 2.096897 | 2.169213 | 2.250918 | 2.334046 | 2.410079 | 2.456132 | 2.496408 | 2.543627 | 2.585956 |
|  | 30 | 1.889618 | 1.926662 | 1.968769 | 2.005356 | 2.047965 | 2.120184 | 2.201924 | 2.285243 | 2.361586 | 2.407886 | 2.448417 | 2.495978 | 2.538652 |
|  | 40 | 1.872573 | 1.910001 | 1.952588 | 1.989628 | 2.032809 | 2.1061 | 2.189208 | 2.274087 | 2.352001 | 2.39932 | 2.440782 | 2.489482 | 2.533221 |
|  | 50 | 1.859187 | 1.897065 | 1.94021 | 1.977774 | 2.02161 | 2.096124 | 2.180782 | 2.267419 | 2.347098 | 2.39556 | 2.438066 | 2.488042 | 2.532974 |
|  | 60 | 1.833586 | 1.871655 | 1.915064 | 1.952899 | 1.997098 | 2.072344 | 2.158005 | 2.245853 | 2.326809 | 2.37612 | 2.419417 | 2.470378 | 2.516243 |
|  | 70 | 1.773284 | 1.810794 | 1.853615 | 1.890978 | 1.934675 | 2.009181 | 2.094178 | 2.181538 | 2.26221 | 2.311427 | 2.354688 | 2.405664 | 2.451594 |
|  | 80 | 1.736799 | 1.774223 | 1.816993 | 1.854355 | 1.898099 | 1.97281 | 2.058225 | 2.146214 | 2.227645 | 2.277407 | 2.321198 | 2.372858 | 2.41946 |
|  | 90 | 1.662316 | 1.698794 | 1.740532 | 1.777036 | 1.819826 | 1.89303 | 1.976911 | 2.063526 | 2.143867 | 2.193048 | 2.23638 | 2.28756 | 2.333788 |
| **R_lateralorbitofrontal** | 5 | 2.563582 | 2.643328 | 2.726566 | 2.793656 | 2.866791 | 2.98119 | 3.100282 | 3.214453 | 3.315542 | 3.376163 | 3.42922 | 3.491846 | 3.548704 |
|  | 10 | 2.45093 | 2.519971 | 2.593391 | 2.653549 | 2.720099 | 2.826179 | 2.939101 | 3.04967 | 3.149409 | 3.210032 | 3.263582 | 3.327373 | 3.38583 |
|  | 15 | 2.356418 | 2.417521 | 2.483467 | 2.538232 | 2.599578 | 2.698996 | 2.807007 | 2.91492 | 3.014073 | 3.075173 | 3.129665 | 3.195213 | 3.255883 |
|  | 20 | 2.284631 | 2.340453 | 2.401333 | 2.452392 | 2.510123 | 2.604888 | 2.709537 | 2.815855 | 2.915104 | 2.977009 | 3.0327 | 3.100292 | 3.163438 |
|  | 30 | 2.193579 | 2.244533 | 2.300638 | 2.348127 | 2.402308 | 2.492381 | 2.593527 | 2.698143 | 2.79754 | 2.860408 | 2.917545 | 2.987641 | 3.053878 |
|  | 40 | 2.145107 | 2.19551 | 2.251024 | 2.298028 | 2.351676 | 2.440921 | 2.541247 | 2.645168 | 2.744077 | 2.806733 | 2.863748 | 2.933791 | 3.000079 |
|  | 50 | 2.120699 | 2.171271 | 2.226984 | 2.274168 | 2.328039 | 2.417705 | 2.518601 | 2.623254 | 2.723021 | 2.786314 | 2.843976 | 2.914906 | 2.98213 |
|  | 60 | 2.113616 | 2.164688 | 2.221036 | 2.268829 | 2.323479 | 2.414652 | 2.51758 | 2.624748 | 2.727332 | 2.792635 | 2.852284 | 2.925867 | 2.995823 |
|  | 70 | 2.077986 | 2.131319 | 2.189983 | 2.239596 | 2.296167 | 2.390178 | 2.495787 | 2.605194 | 2.709428 | 2.775543 | 2.835779 | 2.909889 | 2.980149 |
|  | 80 | 2.052462 | 2.111166 | 2.175209 | 2.228949 | 2.289765 | 2.389777 | 2.50063 | 2.613887 | 2.720374 | 2.787236 | 2.847708 | 2.921544 | 2.990986 |
|  | 90 | 2.037198 | 2.102899 | 2.17388 | 2.232904 | 2.299128 | 2.406778 | 2.524376 | 2.64279 | 2.752639 | 2.82092 | 2.882237 | 2.956562 | 3.025937 |
| **R_lingual** | 5 | 2.108013 | 2.146944 | 2.191608 | 2.230779 | 2.276835 | 2.356006 | 2.447351 | 2.542133 | 2.630246 | 2.684292 | 2.73199 | 2.788431 | 2.839515 |
|  | 10 | 1.960195 | 1.996527 | 2.038255 | 2.074899 | 2.118047 | 2.192412 | 2.278547 | 2.367923 | 2.45068 | 2.501296 | 2.545889 | 2.598571 | 2.646182 |
|  | 15 | 1.849202 | 1.883652 | 1.923254 | 1.958067 | 1.999111 | 2.069999 | 2.152368 | 2.237836 | 2.316711 | 2.36484 | 2.407179 | 2.457133 | 2.502222 |
|  | 20 | 1.793262 | 1.827153 | 1.8661 | 1.900324 | 1.940654 | 2.010253 | 2.091027 | 2.174828 | 2.252243 | 2.299514 | 2.341115 | 2.390217 | 2.434551 |
|  | 30 | 1.736008 | 1.771258 | 1.811504 | 1.846604 | 1.887604 | 1.957287 | 2.03631 | 2.11823 | 2.195618 | 2.243615 | 2.286263 | 2.337034 | 2.383241 |
|  | 40 | 1.693391 | 1.730275 | 1.772114 | 1.808335 | 1.85028 | 1.920511 | 1.998336 | 2.078945 | 2.156807 | 2.20584 | 2.249815 | 2.302611 | 2.351037 |
|  | 50 | 1.668527 | 1.704755 | 1.74599 | 1.781827 | 1.823519 | 1.89388 | 1.972818 | 2.054579 | 2.13255 | 2.181222 | 2.224636 | 2.2765 | 2.323849 |
|  | 60 | 1.653054 | 1.686403 | 1.72485 | 1.758757 | 1.798887 | 1.868667 | 1.950564 | 2.035451 | 2.112826 | 2.159617 | 2.200549 | 2.248592 | 2.291744 |
|  | 70 | 1.618591 | 1.648198 | 1.682876 | 1.714024 | 1.751691 | 1.819689 | 1.903887 | 1.991231 | 2.066686 | 2.110543 | 2.14799 | 2.191002 | 2.228877 |
|  | 80 | 1.59156 | 1.618067 | 1.64959 | 1.678409 | 1.713993 | 1.780604 | 1.867166 | 1.957021 | 2.030963 | 2.072324 | 2.106827 | 2.145649 | 2.179204 |
|  | 90 | 1.575006 | 1.598879 | 1.627708 | 1.654537 | 1.688372 | 1.754077 | 1.843388 | 1.936143 | 2.009085 | 2.048336 | 2.080323 | 2.115581 | 2.145501 |
| **R_medialorbitofrontal** | 5 | 2.500273 | 2.554515 | 2.616454 | 2.6705 | 2.733685 | 2.8413 | 2.963793 | 3.090197 | 3.208105 | 3.280564 | 3.344557 | 3.420299 | 3.488843 |
|  | 10 | 2.306564 | 2.359615 | 2.420119 | 2.472821 | 2.534295 | 2.638527 | 2.756299 | 2.878215 | 2.993501 | 3.06504 | 3.128612 | 3.204291 | 3.273156 |
|  | 15 | 2.165757 | 2.218474 | 2.278535 | 2.330777 | 2.39159 | 2.494278 | 2.609506 | 2.729183 | 2.843905 | 2.915787 | 2.980062 | 3.057028 | 3.127459 |
|  | 20 | 2.072196 | 2.125134 | 2.185396 | 2.237744 | 2.298568 | 2.400885 | 2.514943 | 2.633791 | 2.749233 | 2.822247 | 2.887933 | 2.967044 | 3.039841 |
|  | 30 | 1.9736 | 2.026349 | 2.086264 | 2.138156 | 2.19821 | 2.298438 | 2.408664 | 2.524077 | 2.63887 | 2.712702 | 2.779852 | 2.861574 | 2.937537 |
|  | 40 | 1.938404 | 1.990868 | 2.050282 | 2.101545 | 2.160586 | 2.258234 | 2.363978 | 2.475065 | 2.588104 | 2.661979 | 2.729874 | 2.813332 | 2.891672 |
|  | 50 | 1.925721 | 1.977371 | 2.035791 | 2.086109 | 2.143926 | 2.239108 | 2.341322 | 2.449031 | 2.560288 | 2.63378 | 2.701809 | 2.78602 | 2.865621 |
|  | 60 | 1.921953 | 1.972259 | 2.029456 | 2.079 | 2.136288 | 2.231569 | 2.3355 | 2.445612 | 2.558734 | 2.63325 | 2.702145 | 2.787374 | 2.867923 |
|  | 70 | 1.905042 | 1.953122 | 2.008488 | 2.057137 | 2.114318 | 2.212108 | 2.323521 | 2.442676 | 2.561788 | 2.638881 | 2.70942 | 2.795892 | 2.876953 |
|  | 80 | 1.900832 | 1.94602 | 1.998887 | 2.046189 | 2.102974 | 2.203715 | 2.325172 | 2.456569 | 2.583053 | 2.662804 | 2.734631 | 2.821441 | 2.901772 |
|  | 90 | 1.893651 | 1.935638 | 1.985519 | 2.030948 | 2.086638 | 2.189139 | 2.319733 | 2.462736 | 2.595539 | 2.677065 | 2.749298 | 2.835331 | 2.913885 |
| **R_middletemporal** | 5 | 2.635071 | 2.758893 | 2.875124 | 2.961227 | 3.048861 | 3.175383 | 3.296079 | 3.403202 | 3.49209 | 3.543001 | 3.586201 | 3.635659 | 3.679205 |
|  | 10 | 2.617941 | 2.708925 | 2.800452 | 2.87197 | 2.947868 | 3.062658 | 3.177463 | 3.283276 | 3.373593 | 3.426258 | 3.47144 | 3.523689 | 3.570127 |
|  | 15 | 2.578093 | 2.648671 | 2.723097 | 2.783585 | 2.84996 | 2.954473 | 3.063743 | 3.168408 | 3.260549 | 3.315399 | 3.363082 | 3.418916 | 3.469136 |
|  | 20 | 2.536148 | 2.594513 | 2.657916 | 2.710839 | 2.770335 | 2.866991 | 2.971849 | 3.075811 | 3.170081 | 3.227376 | 3.277878 | 3.337812 | 3.39244 |
|  | 30 | 2.460116 | 2.512699 | 2.570595 | 2.619523 | 2.675159 | 2.766901 | 2.868192 | 2.970242 | 3.064005 | 3.121495 | 3.172453 | 3.233242 | 3.28892 |
|  | 40 | 2.400741 | 2.455933 | 2.516233 | 2.566807 | 2.623886 | 2.716989 | 2.818268 | 2.918644 | 3.009347 | 3.064224 | 3.112391 | 3.169263 | 3.220791 |
|  | 50 | 2.370083 | 2.427328 | 2.489633 | 2.541694 | 2.600229 | 2.695177 | 2.797657 | 2.898314 | 2.988422 | 3.04252 | 3.089731 | 3.145137 | 3.195013 |
|  | 60 | 2.336772 | 2.394021 | 2.456577 | 2.509028 | 2.568178 | 2.664465 | 2.768761 | 2.87144 | 2.963445 | 3.018677 | 3.066856 | 3.123353 | 3.174154 |
|  | 70 | 2.275451 | 2.330981 | 2.392243 | 2.444064 | 2.502978 | 2.599901 | 2.70619 | 2.81201 | 2.907696 | 2.965484 | 3.016079 | 3.075613 | 3.129312 |
|  | 80 | 2.224206 | 2.281544 | 2.344982 | 2.398777 | 2.460071 | 2.561171 | 2.672329 | 2.783183 | 2.88348 | 2.944044 | 2.997046 | 3.059367 | 3.115526 |
|  | 90 | 2.166472 | 2.231382 | 2.30246 | 2.362149 | 2.429526 | 2.539234 | 2.657876 | 2.774193 | 2.877753 | 2.939527 | 2.993125 | 3.055599 | 3.111393 |
| **R_parahippocampal** | 5 | 2.247055 | 2.343496 | 2.449576 | 2.538939 | 2.639987 | 2.804516 | 2.981642 | 3.153471 | 3.304067 | 3.392509 | 3.468262 | 3.555277 | 3.631717 |
|  | 10 | 2.205417 | 2.300029 | 2.404376 | 2.492495 | 2.592361 | 2.755447 | 2.931639 | 3.103129 | 3.253859 | 3.342556 | 3.418627 | 3.506118 | 3.583072 |
|  | 15 | 2.162703 | 2.255387 | 2.357889 | 2.444667 | 2.543243 | 2.704713 | 2.879801 | 3.050808 | 3.201564 | 3.290463 | 3.366814 | 3.454744 | 3.532185 |
|  | 20 | 2.121892 | 2.212672 | 2.313351 | 2.398807 | 2.496114 | 2.656012 | 2.830059 | 3.000671 | 3.151554 | 3.240727 | 3.317426 | 3.405883 | 3.483895 |
|  | 30 | 2.046414 | 2.13391 | 2.231446 | 2.314627 | 2.409765 | 2.567028 | 2.739444 | 2.90963 | 3.061049 | 3.150922 | 3.228444 | 3.318096 | 3.397377 |
|  | 40 | 1.986628 | 2.072683 | 2.168915 | 2.251225 | 2.345627 | 2.502253 | 2.674756 | 2.845783 | 2.998544 | 3.089467 | 3.168039 | 3.259071 | 3.339716 |
|  | 50 | 1.953848 | 2.041431 | 2.139394 | 2.223203 | 2.319346 | 2.478908 | 2.654712 | 2.829081 | 2.984881 | 3.077637 | 3.157808 | 3.250708 | 3.333022 |
|  | 60 | 1.924434 | 2.015815 | 2.117725 | 2.204673 | 2.304163 | 2.468722 | 2.649285 | 2.827666 | 2.986503 | 3.080836 | 3.162237 | 3.256415 | 3.339732 |
|  | 70 | 1.861274 | 1.956456 | 2.062047 | 2.151702 | 2.253833 | 2.42177 | 2.604743 | 2.784302 | 2.943268 | 3.037295 | 3.118217 | 3.211598 | 3.294002 |
|  | 80 | 1.781984 | 1.880552 | 1.989248 | 2.081043 | 2.1851 | 2.355121 | 2.538983 | 2.718161 | 2.875853 | 2.968745 | 3.048473 | 3.14024 | 3.221018 |
|  | 90 | 1.708371 | 1.810679 | 1.922799 | 2.016968 | 2.123191 | 2.295666 | 2.480838 | 2.660103 | 2.817002 | 2.909077 | 2.987908 | 3.078431 | 3.157933 |
| **R_paracentral** | 5 | 2.320171 | 2.370681 | 2.427723 | 2.476971 | 2.533964 | 2.629696 | 2.736781 | 2.844604 | 2.942264 | 3.000984 | 3.052077 | 3.111673 | 3.164823 |
|  | 10 | 2.220972 | 2.27043 | 2.326228 | 2.374356 | 2.429999 | 2.523342 | 2.627576 | 2.732343 | 2.827081 | 2.883974 | 2.933438 | 2.991085 | 3.042455 |
|  | 15 | 2.129762 | 2.178291 | 2.232986 | 2.280114 | 2.334548 | 2.425738 | 2.527385 | 2.629369 | 2.721436 | 2.776658 | 2.824627 | 2.880485 | 2.930219 |
|  | 20 | 2.064394 | 2.112546 | 2.166755 | 2.213416 | 2.267256 | 2.357321 | 2.457532 | 2.557887 | 2.648329 | 2.702507 | 2.74953 | 2.80424 | 2.852909 |
|  | 30 | 1.980651 | 2.029098 | 2.083515 | 2.130252 | 2.184065 | 2.273819 | 2.373308 | 2.472563 | 2.561702 | 2.614964 | 2.661112 | 2.714711 | 2.762312 |
|  | 40 | 1.931553 | 1.981106 | 2.036638 | 2.084227 | 2.138905 | 2.229835 | 2.330254 | 2.430067 | 2.519408 | 2.572661 | 2.618724 | 2.672139 | 2.7195 |
|  | 50 | 1.884894 | 1.935601 | 1.992295 | 2.040776 | 2.096363 | 2.188542 | 2.289982 | 2.390454 | 2.480104 | 2.533419 | 2.579465 | 2.632781 | 2.679983 |
|  | 60 | 1.842425 | 1.894362 | 1.952304 | 2.001751 | 2.058335 | 2.151919 | 2.254566 | 2.355906 | 2.446071 | 2.499582 | 2.545734 | 2.5991 | 2.646283 |
|  | 70 | 1.789932 | 1.84277 | 1.901597 | 1.9517 | 2.00893 | 2.103346 | 2.20659 | 2.308217 | 2.398401 | 2.451824 | 2.497842 | 2.550988 | 2.597923 |
|  | 80 | 1.745691 | 1.799666 | 1.859631 | 1.910605 | 1.968722 | 2.064367 | 2.168642 | 2.270989 | 2.361582 | 2.415152 | 2.461241 | 2.514409 | 2.561308 |
|  | 90 | 1.704192 | 1.75944 | 1.820681 | 1.872632 | 1.931748 | 2.028787 | 2.134253 | 2.237461 | 2.328581 | 2.382363 | 2.42858 | 2.481833 | 2.528755 |
| **R_parsopercularis** | 5 | 2.535721 | 2.586067 | 2.642699 | 2.691406 | 2.747562 | 2.841395 | 2.945651 | 3.049907 | 3.14374 | 3.199896 | 3.248603 | 3.305235 | 3.355581 |
|  | 10 | 2.453394 | 2.503686 | 2.560257 | 2.608911 | 2.665006 | 2.758738 | 2.86288 | 2.967023 | 3.060755 | 3.11685 | 3.165504 | 3.222075 | 3.272367 |
|  | 15 | 2.372583 | 2.422821 | 2.479331 | 2.527932 | 2.583966 | 2.677596 | 2.781627 | 2.885657 | 2.979287 | 3.035322 | 3.083923 | 3.140432 | 3.19067 |
|  | 20 | 2.29536 | 2.345543 | 2.401991 | 2.45054 | 2.506514 | 2.600043 | 2.703961 | 2.807878 | 2.901408 | 2.957381 | 3.00593 | 3.062378 | 3.112562 |
|  | 30 | 2.176298 | 2.226373 | 2.282698 | 2.331142 | 2.386994 | 2.48032 | 2.584012 | 2.687704 | 2.78103 | 2.836882 | 2.885325 | 2.941651 | 2.991725 |
|  | 40 | 2.106649 | 2.156614 | 2.212818 | 2.261155 | 2.316886 | 2.410009 | 2.513475 | 2.616942 | 2.710065 | 2.765795 | 2.814133 | 2.870336 | 2.920302 |
|  | 50 | 2.062044 | 2.111901 | 2.167982 | 2.216215 | 2.271824 | 2.364745 | 2.467986 | 2.571228 | 2.664149 | 2.719758 | 2.767991 | 2.824072 | 2.873929 |
|  | 60 | 2.03102 | 2.080769 | 2.136729 | 2.184858 | 2.240347 | 2.333067 | 2.436086 | 2.539104 | 2.631824 | 2.687314 | 2.735442 | 2.791402 | 2.841151 |
|  | 70 | 1.988481 | 2.038125 | 2.093965 | 2.141991 | 2.197361 | 2.289883 | 2.392682 | 2.495481 | 2.588003 | 2.643373 | 2.691399 | 2.747239 | 2.796883 |
|  | 80 | 1.958882 | 2.00842 | 2.064142 | 2.112066 | 2.167319 | 2.259645 | 2.362226 | 2.464807 | 2.557133 | 2.612386 | 2.66031 | 2.716032 | 2.76557 |
|  | 90 | 1.945346 | 1.994779 | 2.050383 | 2.098206 | 2.153343 | 2.245474 | 2.347838 | 2.450202 | 2.542333 | 2.597469 | 2.645292 | 2.700896 | 2.75033 |
| **R_parsorbitalis** | 5 | 2.489577 | 2.594 | 2.703801 | 2.792726 | 2.889824 | 3.041111 | 3.195937 | 3.339392 | 3.460408 | 3.529641 | 3.587944 | 3.653844 | 3.710843 |
|  | 10 | 2.421664 | 2.507347 | 2.600821 | 2.678967 | 2.766705 | 2.908219 | 3.058826 | 3.203325 | 3.328756 | 3.401917 | 3.464297 | 3.535638 | 3.59804 |
|  | 15 | 2.348199 | 2.422571 | 2.505629 | 2.57658 | 2.65785 | 2.792442 | 2.940324 | 3.08658 | 3.216916 | 3.294359 | 3.361204 | 3.43856 | 3.507008 |
|  | 20 | 2.271403 | 2.340899 | 2.419361 | 2.487084 | 2.565434 | 2.696981 | 2.844028 | 2.991975 | 3.125874 | 3.206333 | 3.276312 | 3.3579 | 3.430629 |
|  | 30 | 2.163444 | 2.231916 | 2.309308 | 2.376177 | 2.453618 | 2.583823 | 2.729633 | 2.8766 | 3.009831 | 3.089985 | 3.159756 | 3.241166 | 3.313796 |
|  | 40 | 2.115015 | 2.183413 | 2.260455 | 2.326802 | 2.403394 | 2.531599 | 2.674359 | 2.817435 | 2.946465 | 3.023797 | 3.090938 | 3.169079 | 3.238615 |
|  | 50 | 2.106012 | 2.169865 | 2.242322 | 2.305169 | 2.378225 | 2.501707 | 2.64093 | 2.782236 | 2.911158 | 2.989087 | 3.057143 | 3.136809 | 3.208112 |
|  | 60 | 2.092737 | 2.151085 | 2.218166 | 2.277102 | 2.346501 | 2.465986 | 2.60403 | 2.747787 | 2.882186 | 2.96494 | 3.038148 | 3.124963 | 3.203692 |
|  | 70 | 2.023399 | 2.081742 | 2.148964 | 2.208152 | 2.277999 | 2.398631 | 2.538578 | 2.684961 | 2.822392 | 2.907284 | 2.982553 | 3.072014 | 3.15333 |
|  | 80 | 1.958004 | 2.026906 | 2.105157 | 2.173082 | 2.252098 | 2.385779 | 2.536672 | 2.689982 | 2.82998 | 2.914655 | 2.988631 | 3.075258 | 3.152818 |
|  | 90 | 1.892952 | 1.981408 | 2.079305 | 2.162244 | 2.256525 | 2.411114 | 2.578956 | 2.743104 | 2.887987 | 2.9735 | 3.046987 | 3.13167 | 3.206295 |
| **R_parstriangularis** | 5 | 2.483912 | 2.536698 | 2.595187 | 2.6449 | 2.701729 | 2.79607 | 2.901014 | 3.007413 | 3.105612 | 3.165973 | 3.219543 | 3.283507 | 3.342126 |
|  | 10 | 2.369417 | 2.421702 | 2.479652 | 2.528924 | 2.585268 | 2.678862 | 2.783083 | 2.88889 | 2.986692 | 3.04689 | 3.100372 | 3.164302 | 3.222965 |
|  | 15 | 2.263015 | 2.314902 | 2.372429 | 2.421359 | 2.477335 | 2.570381 | 2.674106 | 2.779559 | 2.877194 | 2.937374 | 2.990898 | 3.054956 | 3.113813 |
|  | 20 | 2.173797 | 2.225417 | 2.282666 | 2.331377 | 2.387126 | 2.47986 | 2.583353 | 2.688724 | 2.786446 | 2.846767 | 2.900477 | 2.964837 | 3.02405 |
|  | 30 | 2.06617 | 2.117196 | 2.173808 | 2.222 | 2.277188 | 2.369089 | 2.471846 | 2.576737 | 2.674312 | 2.734706 | 2.788597 | 2.853325 | 2.913031 |
|  | 40 | 2.00412 | 2.054183 | 2.10973 | 2.157024 | 2.2112 | 2.301482 | 2.402573 | 2.505989 | 2.602454 | 2.662311 | 2.71583 | 2.780255 | 2.839827 |
|  | 50 | 1.971553 | 2.020964 | 2.075783 | 2.122458 | 2.175937 | 2.265106 | 2.365078 | 2.467557 | 2.563402 | 2.623021 | 2.676434 | 2.740875 | 2.800612 |
|  | 60 | 1.934474 | 1.983276 | 2.037412 | 2.083508 | 2.136333 | 2.224466 | 2.323407 | 2.425046 | 2.52037 | 2.579819 | 2.633192 | 2.697737 | 2.757728 |
|  | 70 | 1.882576 | 1.931812 | 1.986449 | 2.032995 | 2.086369 | 2.175522 | 2.275822 | 2.379164 | 2.476435 | 2.537296 | 2.592077 | 2.658515 | 2.720461 |
|  | 80 | 1.8474 | 1.898595 | 1.955455 | 2.003939 | 2.059595 | 2.152727 | 2.257809 | 2.366497 | 2.469258 | 2.533808 | 2.592089 | 2.663012 | 2.729386 |
|  | 90 | 1.815189 | 1.868902 | 1.928625 | 1.979614 | 2.038225 | 2.136521 | 2.247817 | 2.36345 | 2.473339 | 2.542676 | 2.605499 | 2.682241 | 2.754365 |
| **R_pericalcarine** | 5 | 1.51656 | 1.549854 | 1.588703 | 1.623359 | 1.664816 | 1.73792 | 1.825256 | 1.919721 | 2.011512 | 2.069802 | 2.122547 | 2.186589 | 2.246121 |
|  | 10 | 1.432253 | 1.464326 | 1.501789 | 1.535244 | 1.57531 | 1.646076 | 1.730816 | 1.82271 | 1.912239 | 1.969214 | 2.02085 | 2.083646 | 2.142121 |
|  | 15 | 1.367299 | 1.398523 | 1.435032 | 1.467671 | 1.506803 | 1.576037 | 1.659139 | 1.749497 | 1.83777 | 1.894068 | 1.945173 | 2.007429 | 2.065504 |
|  | 20 | 1.329549 | 1.360467 | 1.396657 | 1.429045 | 1.467918 | 1.53681 | 1.619691 | 1.710048 | 1.798557 | 1.855129 | 1.906564 | 1.969329 | 2.027984 |
|  | 30 | 1.299361 | 1.330314 | 1.366606 | 1.39914 | 1.438258 | 1.507767 | 1.591706 | 1.68361 | 1.774034 | 1.832037 | 1.884916 | 1.949623 | 2.010274 |
|  | 40 | 1.278235 | 1.308772 | 1.344604 | 1.376753 | 1.415442 | 1.484281 | 1.567573 | 1.658974 | 1.749118 | 1.807056 | 1.859952 | 1.924785 | 1.985657 |
|  | 50 | 1.256881 | 1.286677 | 1.321653 | 1.353046 | 1.39084 | 1.458131 | 1.539628 | 1.629164 | 1.717583 | 1.774472 | 1.826456 | 1.890226 | 1.95016 |
|  | 60 | 1.243652 | 1.27294 | 1.307333 | 1.338214 | 1.37541 | 1.441683 | 1.522032 | 1.61042 | 1.697825 | 1.754128 | 1.805622 | 1.868854 | 1.928344 |
|  | 70 | 1.229454 | 1.258403 | 1.292422 | 1.322988 | 1.359832 | 1.425554 | 1.505368 | 1.593342 | 1.680521 | 1.736779 | 1.7883 | 1.851657 | 1.911358 |
|  | 80 | 1.205233 | 1.233621 | 1.267003 | 1.297019 | 1.333228 | 1.397893 | 1.47656 | 1.563448 | 1.649742 | 1.70553 | 1.756694 | 1.819706 | 1.879179 |
|  | 90 | 1.158077 | 1.185256 | 1.217232 | 1.246 | 1.280722 | 1.34279 | 1.4184 | 1.502045 | 1.585265 | 1.639146 | 1.688617 | 1.74962 | 1.807276 |
| **R_postcentral** | 5 | 2.00989 | 2.050353 | 2.096437 | 2.136484 | 2.183025 | 2.261302 | 2.348493 | 2.43986 | 2.529678 | 2.587009 | 2.63892 | 2.701849 | 2.760145 |
|  | 10 | 1.914687 | 1.953324 | 1.997377 | 2.035712 | 2.08034 | 2.155642 | 2.239965 | 2.328212 | 2.414259 | 2.468867 | 2.518125 | 2.577625 | 2.632552 |
|  | 15 | 1.833408 | 1.870497 | 1.912831 | 1.94972 | 1.992739 | 2.065558 | 2.147527 | 2.233194 | 2.316051 | 2.36833 | 2.415312 | 2.47186 | 2.523881 |
|  | 20 | 1.781182 | 1.81731 | 1.858591 | 1.89461 | 1.936687 | 2.008137 | 2.088979 | 2.173349 | 2.254293 | 2.305072 | 2.350534 | 2.405059 | 2.455048 |
|  | 30 | 1.727804 | 1.763047 | 1.803404 | 1.838712 | 1.880094 | 1.950806 | 2.031619 | 2.115709 | 2.195092 | 2.244319 | 2.288066 | 2.340167 | 2.387612 |
|  | 40 | 1.691596 | 1.726319 | 1.766163 | 1.801111 | 1.842208 | 1.912864 | 1.994396 | 2.078969 | 2.157534 | 2.205695 | 2.248179 | 2.298427 | 2.343882 |
|  | 50 | 1.665136 | 1.699555 | 1.739129 | 1.773931 | 1.814986 | 1.885998 | 1.968704 | 2.054209 | 2.132381 | 2.17975 | 2.221231 | 2.269958 | 2.313752 |
|  | 60 | 1.64041 | 1.67458 | 1.713946 | 1.74865 | 1.789721 | 1.861178 | 1.945149 | 2.031654 | 2.109492 | 2.156117 | 2.196651 | 2.243947 | 2.286184 |
|  | 70 | 1.596021 | 1.62955 | 1.66825 | 1.70245 | 1.74305 | 1.814096 | 1.898299 | 1.984719 | 2.061258 | 2.10658 | 2.145698 | 2.191039 | 2.231279 |
|  | 80 | 1.560761 | 1.593853 | 1.632121 | 1.666018 | 1.706379 | 1.777409 | 1.86228 | 1.94904 | 2.024679 | 2.068956 | 2.106898 | 2.15059 | 2.189129 |
|  | 90 | 1.523019 | 1.555637 | 1.593423 | 1.626971 | 1.667033 | 1.737925 | 1.823289 | 1.910189 | 1.98477 | 2.02793 | 2.064652 | 2.106666 | 2.143504 |
| **R_posteriorcingulate** | 5 | 2.5064 | 2.569976 | 2.636119 | 2.689467 | 2.748021 | 2.84186 | 2.945876 | 3.057365 | 3.172638 | 3.253024 | 3.332856 | 3.441884 | 3.558884 |
|  | 10 | 2.421705 | 2.478435 | 2.538873 | 2.58865 | 2.644256 | 2.735031 | 2.836805 | 2.945192 | 3.054479 | 3.128321 | 3.199502 | 3.293188 | 3.389331 |
|  | 15 | 2.337387 | 2.389555 | 2.446129 | 2.493444 | 2.546972 | 2.635484 | 2.735425 | 2.841197 | 2.945742 | 3.01468 | 3.079646 | 3.162816 | 3.24539 |
|  | 20 | 2.259355 | 2.308389 | 2.362277 | 2.40786 | 2.459903 | 2.546748 | 2.64525 | 2.74894 | 2.849854 | 2.915176 | 2.975698 | 3.0516 | 3.125143 |
|  | 30 | 2.149838 | 2.195284 | 2.246149 | 2.28984 | 2.340344 | 2.425657 | 2.523036 | 2.624877 | 2.72209 | 2.783566 | 2.839321 | 2.907469 | 2.971524 |
|  | 40 | 2.106535 | 2.150903 | 2.201103 | 2.244615 | 2.295285 | 2.381516 | 2.480358 | 2.583435 | 2.680853 | 2.741709 | 2.796281 | 2.86208 | 2.922942 |
|  | 50 | 2.071821 | 2.115404 | 2.165091 | 2.208445 | 2.259214 | 2.346158 | 2.446342 | 2.55098 | 2.649632 | 2.710986 | 2.765758 | 2.831418 | 2.891724 |
|  | 60 | 2.05165 | 2.093827 | 2.142314 | 2.184953 | 2.235255 | 2.322245 | 2.423648 | 2.530664 | 2.632383 | 2.695965 | 2.752891 | 2.821299 | 2.884252 |
|  | 70 | 2.020836 | 2.060291 | 2.10612 | 2.146836 | 2.195365 | 2.280544 | 2.381818 | 2.490984 | 2.596894 | 2.664149 | 2.725046 | 2.799076 | 2.868021 |
|  | 80 | 2.006394 | 2.042997 | 2.085983 | 2.124603 | 2.171175 | 2.254387 | 2.355859 | 2.4685 | 2.581222 | 2.654663 | 2.722469 | 2.806645 | 2.886837 |
|  | 90 | 2.025878 | 2.060421 | 2.101389 | 2.138582 | 2.183936 | 2.266424 | 2.369732 | 2.488272 | 2.611438 | 2.694387 | 2.773041 | 2.873674 | 2.972893 |
| **R_precentral** | 5 | 2.13186 | 2.244734 | 2.348036 | 2.422658 | 2.496585 | 2.598559 | 2.688886 | 2.768598 | 2.838472 | 2.879294 | 2.913992 | 2.953448 | 2.987694 |
|  | 10 | 2.159336 | 2.2498 | 2.337531 | 2.403794 | 2.471748 | 2.569138 | 2.658897 | 2.739879 | 2.811396 | 2.853421 | 2.889293 | 2.930257 | 2.965964 |
|  | 15 | 2.168521 | 2.242845 | 2.318391 | 2.37761 | 2.4402 | 2.533119 | 2.622109 | 2.704222 | 2.777337 | 2.82058 | 2.857666 | 2.900218 | 2.937488 |
|  | 20 | 2.158326 | 2.22191 | 2.288664 | 2.342445 | 2.400657 | 2.489668 | 2.577879 | 2.660934 | 2.735401 | 2.779693 | 2.817841 | 2.861806 | 2.900489 |
|  | 30 | 2.106806 | 2.162057 | 2.221607 | 2.270758 | 2.325182 | 2.411035 | 2.499552 | 2.584297 | 2.659843 | 2.704663 | 2.743243 | 2.787718 | 2.826884 |
|  | 40 | 2.047753 | 2.10306 | 2.162784 | 2.212222 | 2.267186 | 2.354594 | 2.445912 | 2.532724 | 2.608119 | 2.652089 | 2.689555 | 2.73236 | 2.769752 |
|  | 50 | 1.998693 | 2.055628 | 2.117077 | 2.167982 | 2.224702 | 2.315411 | 2.411131 | 2.501245 | 2.57736 | 2.620939 | 2.657669 | 2.699232 | 2.735227 |
|  | 60 | 1.943585 | 2.000209 | 2.061704 | 2.112995 | 2.170577 | 2.263815 | 2.363924 | 2.457918 | 2.535627 | 2.579487 | 2.616153 | 2.657354 | 2.692817 |
|  | 70 | 1.881139 | 1.933326 | 1.991039 | 2.040024 | 2.095977 | 2.188882 | 2.291877 | 2.389673 | 2.469745 | 2.514628 | 2.552016 | 2.593917 | 2.62991 |
|  | 80 | 1.836111 | 1.881271 | 1.932564 | 1.977245 | 2.029618 | 2.119902 | 2.224903 | 2.327409 | 2.411716 | 2.459075 | 2.498609 | 2.543029 | 2.581303 |
|  | 90 | 1.799122 | 1.837959 | 1.883177 | 1.923568 | 1.972164 | 2.059323 | 2.166195 | 2.27431 | 2.364366 | 2.415334 | 2.458119 | 2.506478 | 2.548412 |
| **R_precuneus** | 5 | 2.451074 | 2.513024 | 2.576559 | 2.627053 | 2.681539 | 2.76615 | 2.854546 | 2.941104 | 3.020646 | 3.070236 | 3.115091 | 3.170075 | 3.222172 |
|  | 10 | 2.303181 | 2.361349 | 2.421571 | 2.469816 | 2.522221 | 2.604181 | 2.69029 | 2.774708 | 2.85203 | 2.89999 | 2.943158 | 2.995754 | 3.045235 |
|  | 15 | 2.174394 | 2.229673 | 2.287394 | 2.33397 | 2.384866 | 2.464982 | 2.549588 | 2.632635 | 2.708493 | 2.755338 | 2.797318 | 2.848197 | 2.895761 |
|  | 20 | 2.089063 | 2.142598 | 2.198932 | 2.244691 | 2.294969 | 2.374586 | 2.459077 | 2.542123 | 2.617815 | 2.664381 | 2.705956 | 2.756109 | 2.802738 |
|  | 30 | 2.002585 | 2.05321 | 2.107233 | 2.15164 | 2.200922 | 2.279832 | 2.364367 | 2.447747 | 2.523548 | 2.569921 | 2.611085 | 2.660379 | 2.705806 |
|  | 40 | 1.967348 | 2.015066 | 2.066597 | 2.109393 | 2.157301 | 2.234773 | 2.318502 | 2.401429 | 2.476754 | 2.522672 | 2.563267 | 2.611626 | 2.655901 |
|  | 50 | 1.947655 | 1.992526 | 2.041468 | 2.082469 | 2.128711 | 2.204134 | 2.286304 | 2.368037 | 2.442298 | 2.487471 | 2.527298 | 2.574562 | 2.617631 |
|  | 60 | 1.922066 | 1.965522 | 2.013319 | 2.053653 | 2.099431 | 2.174654 | 2.257195 | 2.339648 | 2.414637 | 2.460201 | 2.500301 | 2.547765 | 2.590869 |
|  | 70 | 1.852732 | 1.896598 | 1.945195 | 1.98647 | 2.033582 | 2.111526 | 2.197642 | 2.284062 | 2.362812 | 2.410657 | 2.452729 | 2.502452 | 2.547509 |
|  | 80 | 1.772074 | 1.817466 | 1.868099 | 1.911367 | 1.961028 | 2.043752 | 2.135818 | 2.228722 | 2.313655 | 2.365321 | 2.41076 | 2.464445 | 2.513049 |
|  | 90 | 1.689673 | 1.737043 | 1.790242 | 1.835984 | 1.888783 | 1.977374 | 2.076773 | 2.177759 | 2.270527 | 2.327106 | 2.37693 | 2.435845 | 2.489208 |
| **R_rostralanteriorcingulate** | 5 | 2.337093 | 2.525946 | 2.698431 | 2.823411 | 2.948583 | 3.1268 | 3.295904 | 3.447386 | 3.575853 | 3.651243 | 3.716564 | 3.793171 | 3.862489 |
|  | 10 | 2.394707 | 2.516756 | 2.639648 | 2.735838 | 2.838253 | 2.994436 | 3.153587 | 3.304849 | 3.439296 | 3.520735 | 3.592773 | 3.678962 | 3.758499 |
|  | 15 | 2.403844 | 2.495644 | 2.593024 | 2.672728 | 2.760961 | 2.902236 | 3.054556 | 3.207145 | 3.349151 | 3.438078 | 3.518567 | 3.617114 | 3.710225 |
|  | 20 | 2.363839 | 2.445741 | 2.534108 | 2.60757 | 2.690083 | 2.824783 | 2.973545 | 3.126189 | 3.271451 | 3.363969 | 3.448727 | 3.553812 | 3.654416 |
|  | 30 | 2.196218 | 2.286067 | 2.381295 | 2.459169 | 2.545298 | 2.682988 | 2.831063 | 2.978881 | 3.115872 | 3.201336 | 3.27846 | 3.372574 | 3.461173 |
|  | 40 | 2.140909 | 2.228434 | 2.321798 | 2.398587 | 2.483957 | 2.621338 | 2.770215 | 2.919864 | 3.059346 | 3.146705 | 3.225739 | 3.322425 | 3.413666 |
|  | 50 | 2.141606 | 2.22216 | 2.309617 | 2.382728 | 2.465251 | 2.600772 | 2.751353 | 2.90652 | 3.054481 | 3.148749 | 3.235077 | 3.342012 | 3.444249 |
|  | 60 | 2.146806 | 2.228818 | 2.318187 | 2.393151 | 2.47803 | 2.617985 | 2.774229 | 2.935916 | 3.090627 | 3.189422 | 3.280029 | 3.392422 | 3.500022 |
|  | 70 | 2.145991 | 2.226817 | 2.315553 | 2.390508 | 2.475942 | 2.618089 | 2.778579 | 2.94654 | 3.108934 | 3.213446 | 3.309831 | 3.430079 | 3.545894 |
|  | 80 | 2.15 | 2.232664 | 2.323687 | 2.400784 | 2.488882 | 2.635943 | 2.802613 | 2.977633 | 3.147306 | 3.256693 | 3.357681 | 3.483798 | 3.60538 |
|  | 90 | 2.205908 | 2.29088 | 2.384934 | 2.464994 | 2.556908 | 2.71133 | 2.887752 | 3.074502 | 3.256879 | 3.375103 | 3.484676 | 3.622064 | 3.755071 |
| **R_rostralmiddlefrontal** | 5 | 2.420123 | 2.471549 | 2.528141 | 2.575906 | 2.630108 | 2.719084 | 2.816468 | 2.913329 | 3.000923 | 3.053864 | 3.100255 | 3.154905 | 3.204269 |
|  | 10 | 2.26916 | 2.316783 | 2.369392 | 2.413965 | 2.464738 | 2.548555 | 2.640998 | 2.733737 | 2.818336 | 2.869825 | 2.915176 | 2.968887 | 3.017679 |
|  | 15 | 2.149816 | 2.194525 | 2.244084 | 2.286215 | 2.334375 | 2.414294 | 2.503089 | 2.592923 | 2.675594 | 2.72627 | 2.771143 | 2.824587 | 2.873428 |
|  | 20 | 2.073806 | 2.116718 | 2.16441 | 2.205064 | 2.251669 | 2.32935 | 2.41622 | 2.504785 | 2.586964 | 2.637686 | 2.682834 | 2.736906 | 2.786616 |
|  | 30 | 2.008304 | 2.04935 | 2.095087 | 2.13419 | 2.179172 | 2.254581 | 2.339698 | 2.427531 | 2.510163 | 2.561782 | 2.608162 | 2.664279 | 2.716449 |
|  | 40 | 1.974844 | 2.015328 | 2.060393 | 2.098904 | 2.14321 | 2.217586 | 2.301853 | 2.389392 | 2.472501 | 2.524873 | 2.572268 | 2.630081 | 2.684322 |
|  | 50 | 1.947301 | 1.988662 | 2.034448 | 2.073389 | 2.118007 | 2.192566 | 2.276729 | 2.364078 | 2.447176 | 2.49972 | 2.547433 | 2.605888 | 2.66102 |
|  | 60 | 1.919631 | 1.963396 | 2.011367 | 2.051805 | 2.097769 | 2.173824 | 2.258792 | 2.34632 | 2.429262 | 2.481655 | 2.529249 | 2.587626 | 2.642791 |
|  | 70 | 1.858733 | 1.906552 | 1.958324 | 2.001482 | 2.050053 | 2.129444 | 2.217009 | 2.306372 | 2.390627 | 2.443771 | 2.492054 | 2.551341 | 2.607478 |
|  | 80 | 1.822342 | 1.878532 | 1.938457 | 1.987747 | 2.042563 | 2.13088 | 2.226852 | 2.323776 | 2.414682 | 2.471955 | 2.524023 | 2.588067 | 2.648877 |
|  | 90 | 1.766606 | 1.835098 | 1.906692 | 1.964549 | 2.027908 | 2.128117 | 2.234971 | 2.341458 | 2.440657 | 2.503046 | 2.559792 | 2.629715 | 2.6963 |
| **R_superiorfrontal** | 5 | 2.78963 | 2.844428 | 2.904096 | 2.953966 | 3.01004 | 3.100976 | 3.199046 | 3.295227 | 3.381151 | 3.432637 | 3.477499 | 3.530057 | 3.577276 |
|  | 10 | 2.660054 | 2.713405 | 2.771542 | 2.820172 | 2.8749 | 2.963781 | 3.05984 | 3.154301 | 3.238938 | 3.289782 | 3.334168 | 3.386274 | 3.433193 |
|  | 15 | 2.551753 | 2.603975 | 2.660928 | 2.708608 | 2.762319 | 2.849675 | 2.944302 | 3.037617 | 3.12149 | 3.172009 | 3.216201 | 3.268194 | 3.315121 |
|  | 20 | 2.469048 | 2.520427 | 2.576509 | 2.623503 | 2.676493 | 2.762812 | 2.856544 | 2.949253 | 3.03286 | 3.083362 | 3.127635 | 3.179846 | 3.227088 |
|  | 30 | 2.365511 | 2.415427 | 2.47001 | 2.515836 | 2.567617 | 2.652253 | 2.744635 | 2.836607 | 2.920153 | 2.970934 | 3.015664 | 3.068685 | 3.116928 |
|  | 40 | 2.309895 | 2.358794 | 2.412353 | 2.457401 | 2.508405 | 2.592054 | 2.683847 | 2.775864 | 2.860108 | 2.91166 | 2.957309 | 3.011728 | 3.061549 |
|  | 50 | 2.272985 | 2.321239 | 2.374164 | 2.418749 | 2.469325 | 2.552539 | 2.644349 | 2.737041 | 2.822611 | 2.87536 | 2.922336 | 2.978686 | 3.03063 |
|  | 60 | 2.23278 | 2.281084 | 2.334128 | 2.378881 | 2.42974 | 2.513695 | 2.606842 | 2.701607 | 2.789888 | 2.844748 | 2.893916 | 2.953308 | 3.008479 |
|  | 70 | 2.156673 | 2.205379 | 2.258941 | 2.304207 | 2.355756 | 2.441169 | 2.536538 | 2.634413 | 2.726544 | 2.784331 | 2.836504 | 2.900041 | 2.959592 |
|  | 80 | 2.105671 | 2.155825 | 2.211069 | 2.257849 | 2.311249 | 2.400107 | 2.500051 | 2.603659 | 2.702368 | 2.764954 | 2.821948 | 2.892023 | 2.958402 |
|  | 90 | 2.066658 | 2.118505 | 2.175712 | 2.224258 | 2.279816 | 2.372699 | 2.478025 | 2.588457 | 2.695124 | 2.763604 | 2.826591 | 2.904907 | 2.980027 |
| **R_superiorparietal** | 5 | 2.227762 | 2.267661 | 2.312541 | 2.35114 | 2.395643 | 2.470004 | 2.552626 | 2.635247 | 2.709608 | 2.754111 | 2.79271 | 2.83759 | 2.877489 |
|  | 10 | 2.080463 | 2.120146 | 2.164782 | 2.203171 | 2.247432 | 2.32139 | 2.403562 | 2.485734 | 2.559692 | 2.603953 | 2.642342 | 2.686978 | 2.726661 |
|  | 15 | 1.962709 | 2.002185 | 2.046589 | 2.084778 | 2.128809 | 2.202382 | 2.284127 | 2.365872 | 2.439444 | 2.483475 | 2.521665 | 2.566069 | 2.605545 |
|  | 20 | 1.897214 | 1.9365 | 1.980691 | 2.018697 | 2.062516 | 2.135736 | 2.217088 | 2.29844 | 2.37166 | 2.415479 | 2.453485 | 2.497676 | 2.536962 |
|  | 30 | 1.832107 | 1.871079 | 1.914916 | 1.952618 | 1.996085 | 2.068718 | 2.149419 | 2.230119 | 2.302752 | 2.34622 | 2.383922 | 2.427759 | 2.46673 |
|  | 40 | 1.800382 | 1.839198 | 1.882859 | 1.920411 | 1.963705 | 2.036048 | 2.116426 | 2.196804 | 2.269146 | 2.312441 | 2.349992 | 2.393653 | 2.432469 |
|  | 50 | 1.787012 | 1.825902 | 1.869647 | 1.90727 | 1.950647 | 2.023127 | 2.103659 | 2.18419 | 2.256671 | 2.300048 | 2.337671 | 2.381416 | 2.420305 |
|  | 60 | 1.763988 | 1.803269 | 1.847453 | 1.885454 | 1.929267 | 2.002476 | 2.083816 | 2.165157 | 2.238366 | 2.282178 | 2.320179 | 2.364364 | 2.403644 |
|  | 70 | 1.706279 | 1.746264 | 1.79124 | 1.829922 | 1.87452 | 1.949041 | 2.031839 | 2.114638 | 2.189159 | 2.233757 | 2.272439 | 2.317415 | 2.357399 |
|  | 80 | 1.662813 | 1.703593 | 1.749464 | 1.788915 | 1.8344 | 1.910404 | 1.994849 | 2.079295 | 2.155298 | 2.200783 | 2.240235 | 2.286106 | 2.326886 |
|  | 90 | 1.606641 | 1.648228 | 1.695007 | 1.735239 | 1.781624 | 1.859132 | 1.945249 | 2.031365 | 2.108873 | 2.155258 | 2.19549 | 2.242269 | 2.283856 |
| **R_superiortemporal** | 5 | 2.52687 | 2.636836 | 2.742936 | 2.822846 | 2.904716 | 3.022185 | 3.130968 | 3.22845 | 3.313051 | 3.362235 | 3.403958 | 3.451349 | 3.492463 |
|  | 10 | 2.520975 | 2.607319 | 2.695447 | 2.764806 | 2.838408 | 2.948336 | 3.054524 | 3.152467 | 3.23892 | 3.28978 | 3.333269 | 3.383042 | 3.426543 |
|  | 15 | 2.49854 | 2.568643 | 2.643171 | 2.703886 | 2.770265 | 2.873093 | 2.976593 | 3.074972 | 3.163481 | 3.216274 | 3.26184 | 3.314472 | 3.360889 |
|  | 20 | 2.46972 | 2.529084 | 2.593984 | 2.648217 | 2.708911 | 2.805856 | 2.90709 | 3.006067 | 3.096792 | 3.151671 | 3.199504 | 3.255296 | 3.304985 |
|  | 30 | 2.402822 | 2.455235 | 2.513614 | 2.563289 | 2.619881 | 2.712602 | 2.8127 | 2.912594 | 3.004769 | 3.060856 | 3.109982 | 3.16759 | 3.219193 |
|  | 40 | 2.342016 | 2.395879 | 2.455666 | 2.506403 | 2.564099 | 2.658535 | 2.760554 | 2.861229 | 2.952081 | 3.006512 | 3.053715 | 3.108556 | 3.157249 |
|  | 50 | 2.293523 | 2.347377 | 2.407226 | 2.458102 | 2.516096 | 2.611465 | 2.715279 | 2.817294 | 2.907937 | 2.96167 | 3.007968 | 3.061446 | 3.108674 |
|  | 60 | 2.26241 | 2.312013 | 2.367902 | 2.416067 | 2.471735 | 2.565171 | 2.669693 | 2.774006 | 2.866901 | 2.922088 | 2.969741 | 3.024925 | 3.073803 |
|  | 70 | 2.199074 | 2.245061 | 2.297408 | 2.342994 | 2.396263 | 2.487183 | 2.591255 | 2.696349 | 2.789869 | 2.84542 | 2.893415 | 2.949053 | 2.998404 |
|  | 80 | 2.110066 | 2.15449 | 2.205212 | 2.249541 | 2.301554 | 2.390968 | 2.494382 | 2.598709 | 2.690411 | 2.744409 | 2.790816 | 2.844362 | 2.891654 |
|  | 90 | 2.012378 | 2.055517 | 2.104853 | 2.148063 | 2.198902 | 2.28675 | 2.389121 | 2.492007 | 2.581128 | 2.633059 | 2.677404 | 2.728273 | 2.77296 |
| **R_supramarginal** | 5 | 2.283146 | 2.453985 | 2.594327 | 2.68689 | 2.772789 | 2.886282 | 2.990038 | 3.087269 | 3.180482 | 3.243355 | 3.304801 | 3.38748 | 3.474581 |
|  | 10 | 2.394147 | 2.468812 | 2.543593 | 2.60184 | 2.663606 | 2.757467 | 2.853091 | 2.944537 | 3.026818 | 3.077322 | 3.122508 | 3.177283 | 3.228583 |
|  | 15 | 2.338577 | 2.394744 | 2.454508 | 2.503485 | 2.557658 | 2.643926 | 2.735526 | 2.824826 | 2.904911 | 2.953322 | 2.995894 | 3.046358 | 3.092347 |
|  | 20 | 2.280982 | 2.329466 | 2.382459 | 2.426927 | 2.47714 | 2.559126 | 2.648456 | 2.737182 | 2.817562 | 2.866301 | 2.909155 | 2.95985 | 3.005876 |
|  | 30 | 2.188264 | 2.235163 | 2.286606 | 2.329942 | 2.379093 | 2.459927 | 2.54902 | 2.638823 | 2.721551 | 2.772446 | 2.817698 | 2.871884 | 2.921734 |
|  | 40 | 2.104465 | 2.160441 | 2.219845 | 2.268468 | 2.322274 | 2.408291 | 2.500664 | 2.592566 | 2.677302 | 2.729898 | 2.777158 | 2.834551 | 2.888286 |
|  | 50 | 2.087223 | 2.137663 | 2.193234 | 2.240144 | 2.293332 | 2.380376 | 2.47495 | 2.567901 | 2.650671 | 2.699982 | 2.742698 | 2.79238 | 2.836626 |
|  | 60 | 2.041654 | 2.091673 | 2.147245 | 2.194485 | 2.24835 | 2.337014 | 2.433721 | 2.528711 | 2.612876 | 2.662691 | 2.705584 | 2.755109 | 2.798838 |
|  | 70 | 1.964467 | 2.013828 | 2.068876 | 2.115833 | 2.169548 | 2.258337 | 2.355657 | 2.451671 | 2.537044 | 2.587689 | 2.631355 | 2.681832 | 2.72645 |
|  | 80 | 1.901493 | 1.951976 | 2.008233 | 2.056189 | 2.11101 | 2.201547 | 2.300672 | 2.398359 | 2.485132 | 2.536569 | 2.580896 | 2.632111 | 2.677359 |
|  | 90 | 1.853443 | 1.905242 | 1.962955 | 2.012146 | 2.068371 | 2.161212 | 2.262841 | 2.362981 | 2.451922 | 2.504641 | 2.550071 | 2.602559 | 2.648929 |
| **R_frontalpole** | 5 | 2.382755 | 2.506626 | 2.643371 | 2.75886 | 2.889634 | 3.10255 | 3.331201 | 3.555693 | 3.757469 | 3.878033 | 3.982413 | 4.103493 | 4.21084 |
|  | 10 | 2.27726 | 2.379499 | 2.494832 | 2.59422 | 2.708919 | 2.900477 | 3.112694 | 3.327297 | 3.525208 | 3.645689 | 3.751323 | 3.875385 | 3.986734 |
|  | 15 | 2.212392 | 2.301755 | 2.404026 | 2.4934 | 2.597972 | 2.776026 | 2.978257 | 3.187904 | 3.385665 | 3.508121 | 3.616769 | 3.745897 | 3.863191 |
|  | 20 | 2.161695 | 2.243633 | 2.338211 | 2.421571 | 2.519949 | 2.689574 | 2.885488 | 3.092116 | 3.290193 | 3.414384 | 3.52556 | 3.658903 | 3.781164 |
|  | 30 | 2.079062 | 2.154099 | 2.241379 | 2.318912 | 2.411159 | 2.572154 | 2.761235 | 2.964085 | 3.16163 | 3.287053 | 3.400371 | 3.537587 | 3.664662 |
|  | 40 | 2.036836 | 2.111362 | 2.198126 | 2.275275 | 2.367159 | 2.527783 | 2.71687 | 2.919971 | 3.117738 | 3.243298 | 3.356744 | 3.494129 | 3.621376 |
|  | 50 | 2.014234 | 2.091002 | 2.180206 | 2.259372 | 2.353473 | 2.5175 | 2.709841 | 2.915249 | 3.113833 | 3.239202 | 3.352014 | 3.488059 | 3.613519 |
|  | 60 | 1.957465 | 2.036543 | 2.128085 | 2.209013 | 2.304837 | 2.470918 | 2.66417 | 2.86853 | 3.063919 | 3.186203 | 3.295559 | 3.4266 | 3.546658 |
|  | 70 | 1.871605 | 1.952352 | 2.045386 | 2.127248 | 2.223718 | 2.389784 | 2.581274 | 2.781537 | 2.970707 | 3.088004 | 3.192214 | 3.316265 | 3.429154 |
|  | 80 | 1.856787 | 1.942999 | 2.041758 | 2.128169 | 2.22943 | 2.402366 | 2.599724 | 2.803631 | 2.993788 | 3.110561 | 3.21361 | 3.335458 | 3.445592 |
|  | 90 | 1.865263 | 1.961174 | 2.070007 | 2.164366 | 2.273962 | 2.458837 | 2.666539 | 2.877466 | 3.070825 | 3.188068 | 3.290639 | 3.410887 | 3.518651 |
| **R_temporalpole** | 5 | 2.352571 | 2.536306 | 2.723322 | 2.87196 | 3.033011 | 3.285027 | 3.547827 | 3.781121 | 3.959586 | 4.055401 | 4.133218 | 4.218492 | 4.290269 |
|  | 10 | 2.438778 | 2.619986 | 2.805904 | 2.95451 | 3.116162 | 3.370007 | 3.635456 | 3.872609 | 4.055718 | 4.154618 | 4.235215 | 4.323796 | 4.398551 |
|  | 15 | 2.513563 | 2.691868 | 2.876144 | 3.024202 | 3.18584 | 3.440471 | 3.707403 | 3.94737 | 4.134377 | 4.235992 | 4.319086 | 4.410684 | 4.488188 |
|  | 20 | 2.557885 | 2.731747 | 2.91262 | 3.058629 | 3.218553 | 3.471203 | 3.736634 | 3.9767 | 4.165522 | 4.26874 | 4.353439 | 4.447085 | 4.526533 |
|  | 30 | 2.578606 | 2.741688 | 2.913241 | 3.052834 | 3.206579 | 3.450609 | 3.707839 | 3.943174 | 4.131689 | 4.235975 | 4.322144 | 4.417989 | 4.499741 |
|  | 40 | 2.604519 | 2.760766 | 2.926574 | 3.06235 | 3.212548 | 3.451797 | 3.704548 | 3.938247 | 4.128786 | 4.235419 | 4.324127 | 4.423383 | 4.508496 |
|  | 50 | 2.636427 | 2.787381 | 2.948844 | 3.081843 | 3.229592 | 3.465803 | 3.715978 | 3.949651 | 4.143351 | 4.252946 | 4.344707 | 4.447968 | 4.536974 |
|  | 60 | 2.639749 | 2.78494 | 2.941372 | 3.070948 | 3.215494 | 3.447484 | 3.693923 | 3.926359 | 4.122029 | 4.233874 | 4.328095 | 4.434703 | 4.527054 |
|  | 70 | 2.55725 | 2.695429 | 2.845199 | 2.969837 | 3.109364 | 3.334027 | 3.573264 | 3.800957 | 3.995469 | 4.107749 | 4.202899 | 4.311136 | 4.405358 |
|  | 80 | 2.465188 | 2.59886 | 2.744492 | 2.866177 | 3.002812 | 3.223431 | 3.458811 | 3.684786 | 3.880656 | 3.994831 | 4.092169 | 4.203494 | 4.300892 |
|  | 90 | 2.377509 | 2.507615 | 2.65009 | 2.769624 | 2.904266 | 3.122305 | 3.355425 | 3.581234 | 3.77989 | 3.896862 | 3.997205 | 4.112622 | 4.214135 |
| **R_transversetemporal** | 5 | 2.164763 | 2.237131 | 2.318677 | 2.388927 | 2.470051 | 2.605906 | 2.757272 | 2.909062 | 3.046022 | 3.128138 | 3.19945 | 3.282467 | 3.356361 |
|  | 10 | 2.085483 | 2.157565 | 2.238718 | 2.308573 | 2.389179 | 2.52402 | 2.674052 | 2.824298 | 2.959698 | 3.040806 | 3.111198 | 3.193096 | 3.265949 |
|  | 15 | 2.010694 | 2.082586 | 2.163453 | 2.233003 | 2.31319 | 2.447178 | 2.596048 | 2.744917 | 2.878904 | 2.95909 | 3.028639 | 3.109505 | 3.181396 |
|  | 20 | 1.942469 | 2.014359 | 2.095146 | 2.164563 | 2.244528 | 2.377984 | 2.526041 | 2.673877 | 2.806757 | 2.886203 | 2.955066 | 3.035082 | 3.106173 |
|  | 30 | 1.826163 | 1.898713 | 1.980068 | 2.049835 | 2.130048 | 2.263572 | 2.411228 | 2.558198 | 2.689926 | 2.768524 | 2.836558 | 2.915507 | 2.985557 |
|  | 40 | 1.736782 | 1.811055 | 1.89414 | 1.965228 | 2.046786 | 2.18216 | 2.331331 | 2.479299 | 2.611521 | 2.690242 | 2.758285 | 2.837133 | 2.906998 |
|  | 50 | 1.668872 | 1.745941 | 1.831914 | 1.905286 | 1.989261 | 2.1282 | 2.280707 | 2.43142 | 2.565658 | 2.645397 | 2.714213 | 2.79384 | 2.864292 |
|  | 60 | 1.610199 | 1.690792 | 1.78041 | 1.856669 | 1.943712 | 2.087216 | 2.244063 | 2.398441 | 2.535465 | 2.616661 | 2.686621 | 2.767447 | 2.838851 |
|  | 70 | 1.549078 | 1.633464 | 1.726954 | 1.806242 | 1.896467 | 2.044629 | 2.20581 | 2.363763 | 2.503441 | 2.585995 | 2.657006 | 2.738912 | 2.811156 |
|  | 80 | 1.48519 | 1.573672 | 1.671279 | 1.753742 | 1.847257 | 2.000145 | 2.165616 | 2.32701 | 2.469166 | 2.552956 | 2.624902 | 2.707746 | 2.780699 |
|  | 90 | 1.420284 | 1.513345 | 1.615486 | 1.7014 | 1.798449 | 1.956335 | 2.126254 | 2.291146 | 2.435772 | 2.520774 | 2.593624 | 2.677362 | 2.750976 |
| **R_insula** | 5 | 2.888717 | 2.981402 | 3.075369 | 3.149042 | 3.227131 | 3.344034 | 3.457894 | 3.561347 | 3.649421 | 3.700054 | 3.742765 | 3.791067 | 3.832832 |
|  | 10 | 2.814763 | 2.888624 | 2.966697 | 3.030019 | 3.099023 | 3.205634 | 3.312845 | 3.413716 | 3.502791 | 3.555266 | 3.60021 | 3.651761 | 3.696928 |
|  | 15 | 2.758593 | 2.821015 | 2.888803 | 2.945089 | 3.007686 | 3.106795 | 3.209106 | 3.308496 | 3.399491 | 3.454459 | 3.502313 | 3.55805 | 3.607614 |
|  | 20 | 2.723656 | 2.779841 | 2.84183 | 2.894037 | 2.952842 | 3.047418 | 3.146734 | 3.24565 | 3.339106 | 3.396855 | 3.447893 | 3.508213 | 3.562623 |
|  | 30 | 2.634993 | 2.687376 | 2.745679 | 2.795153 | 2.85123 | 2.941987 | 3.037697 | 3.135137 | 3.230849 | 3.291676 | 3.346456 | 3.412397 | 3.472962 |
|  | 40 | 2.555122 | 2.611514 | 2.673618 | 2.725741 | 2.784136 | 2.876948 | 2.972279 | 3.068509 | 3.16396 | 3.224928 | 3.279959 | 3.346295 | 3.407265 |
|  | 50 | 2.493887 | 2.557271 | 2.625953 | 2.682674 | 2.745193 | 2.842213 | 2.938621 | 3.034548 | 3.130117 | 3.191193 | 3.246263 | 3.312512 | 3.373231 |
|  | 60 | 2.465787 | 2.535591 | 2.610399 | 2.671497 | 2.738078 | 2.839641 | 2.938081 | 3.03546 | 3.133748 | 3.197011 | 3.254263 | 3.323332 | 3.386775 |
|  | 70 | 2.407278 | 2.480366 | 2.558386 | 2.621816 | 2.690563 | 2.794434 | 2.893479 | 2.991708 | 3.092997 | 3.15908 | 3.219387 | 3.292699 | 3.360517 |
|  | 80 | 2.356595 | 2.426972 | 2.502913 | 2.5652 | 2.63317 | 2.736477 | 2.835075 | 2.934744 | 3.041521 | 3.113127 | 3.179718 | 3.262195 | 3.339929 |
|  | 90 | 2.31723 | 2.382451 | 2.453995 | 2.513535 | 2.579323 | 2.680696 | 2.778426 | 2.880109 | 2.99465 | 3.074505 | 3.15091 | 3.248395 | 3.343176 |
