## Supplementary File 1 for "Cortical Thickness Trajectories across the Lifespan: Data from 17,075 healthy individuals aged 3-90 years"

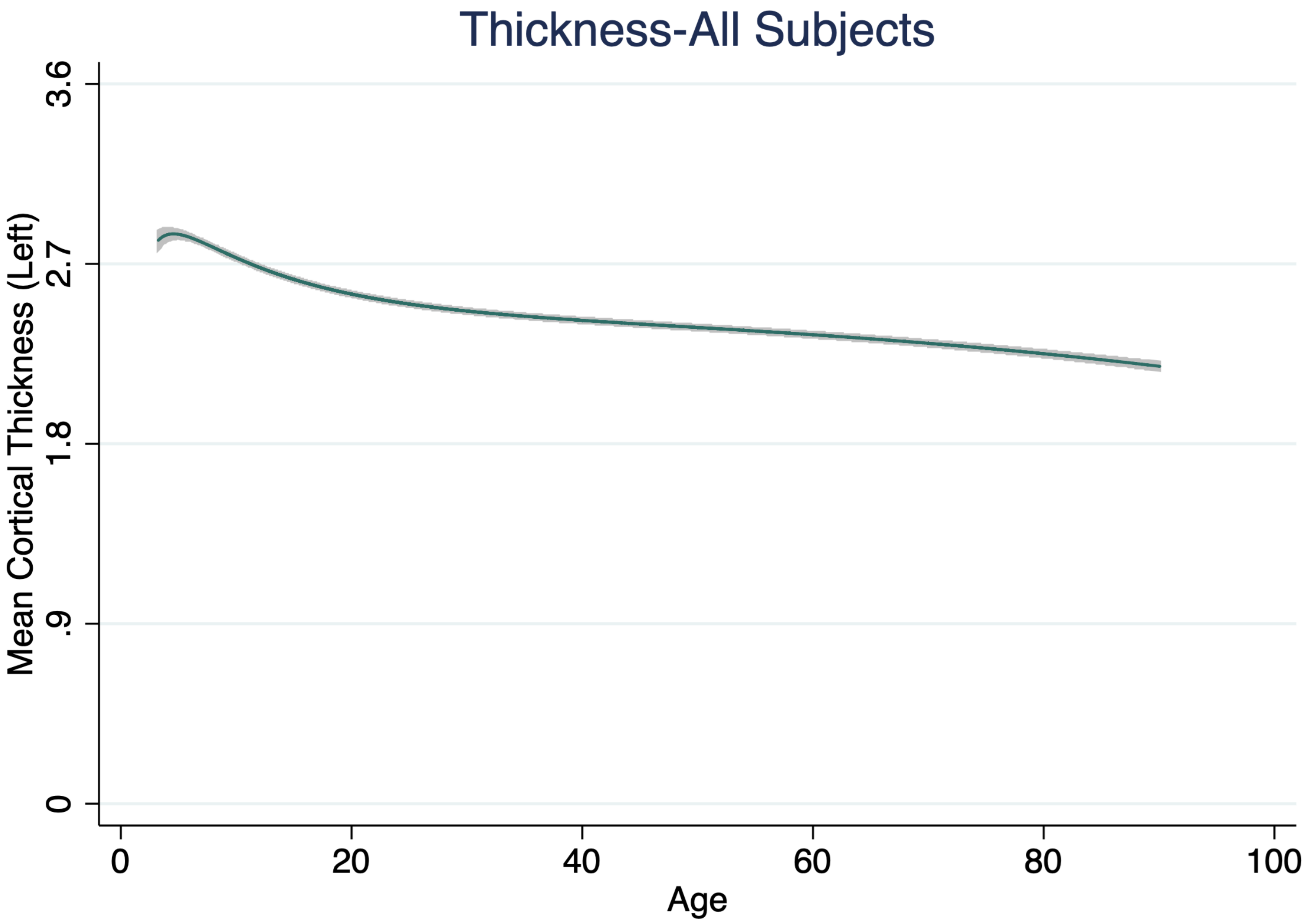
Thickness-All Subjects
Mean Cortical Thickness (Left)
3.6
2.7
1.8
.9
0
0
20
40
60
80
100
Age

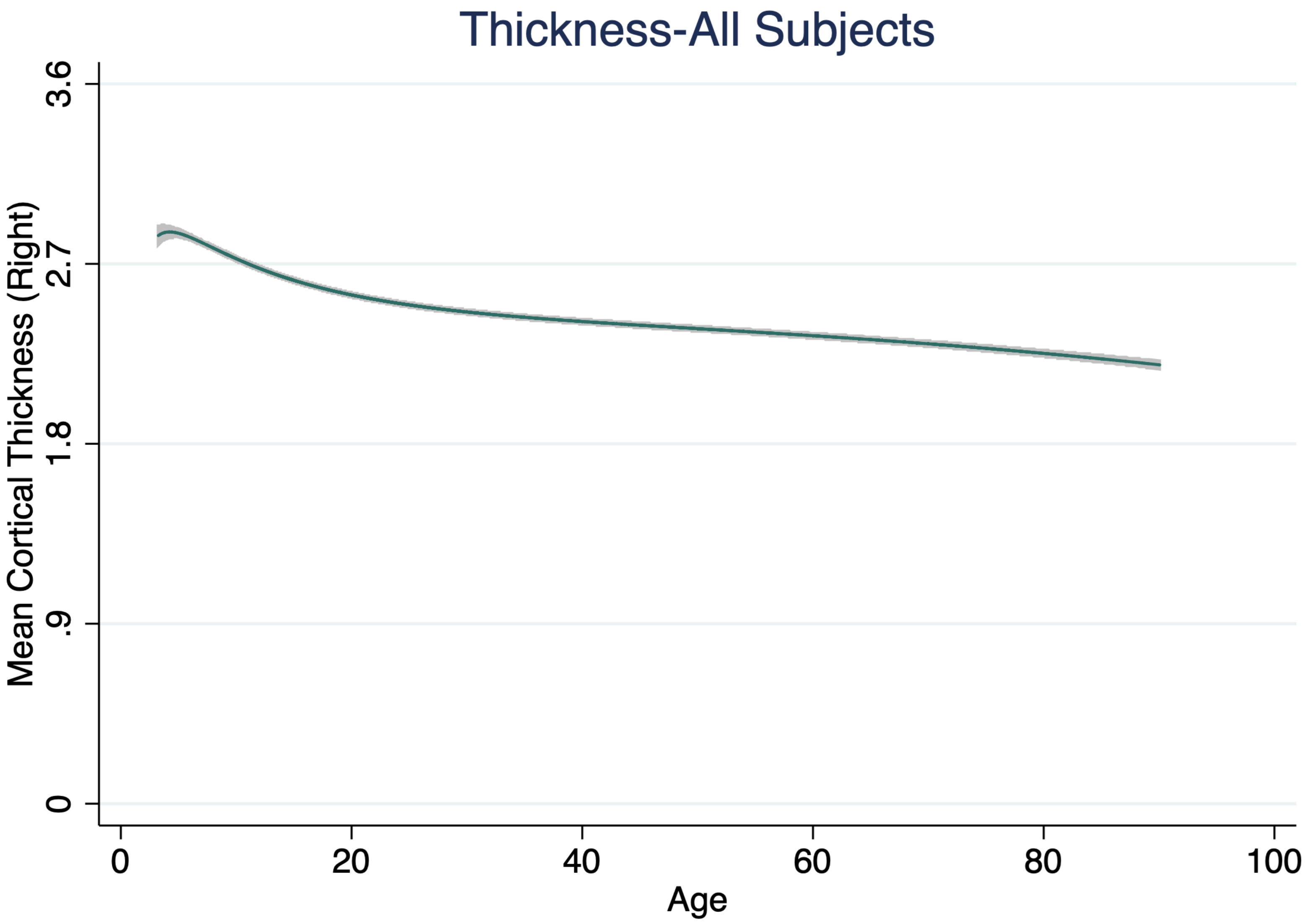

Thickness-All Subjects
Mean Cortical Thickness (Right)
3.6
2.7
1.8
.9
0
0
20
40
60
80
100
Age

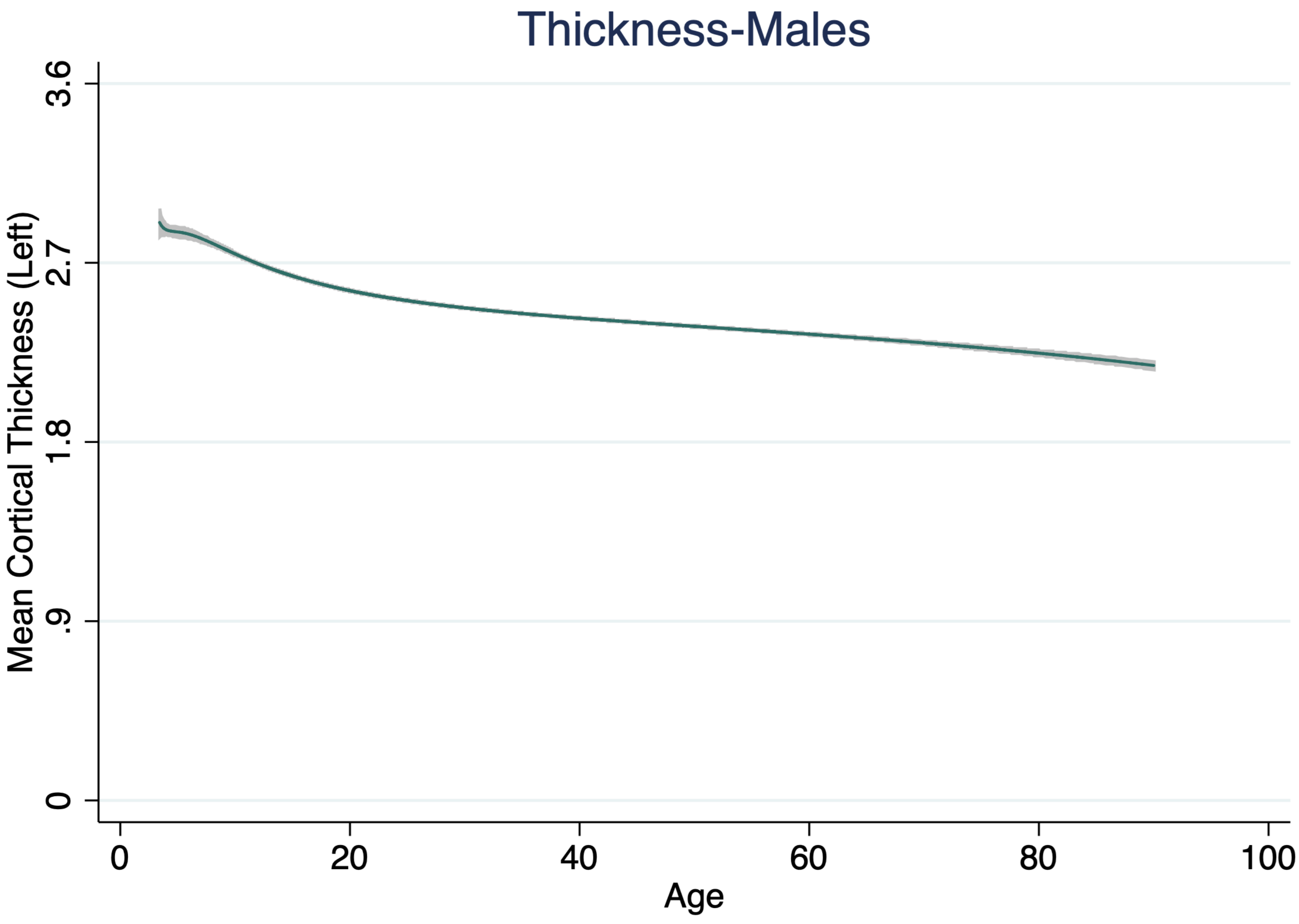

Thickness-Males
Mean Cortical Thickness (Left)
3.6
2.7
1.8
.9
0
0
20
40
60
80
100
Age

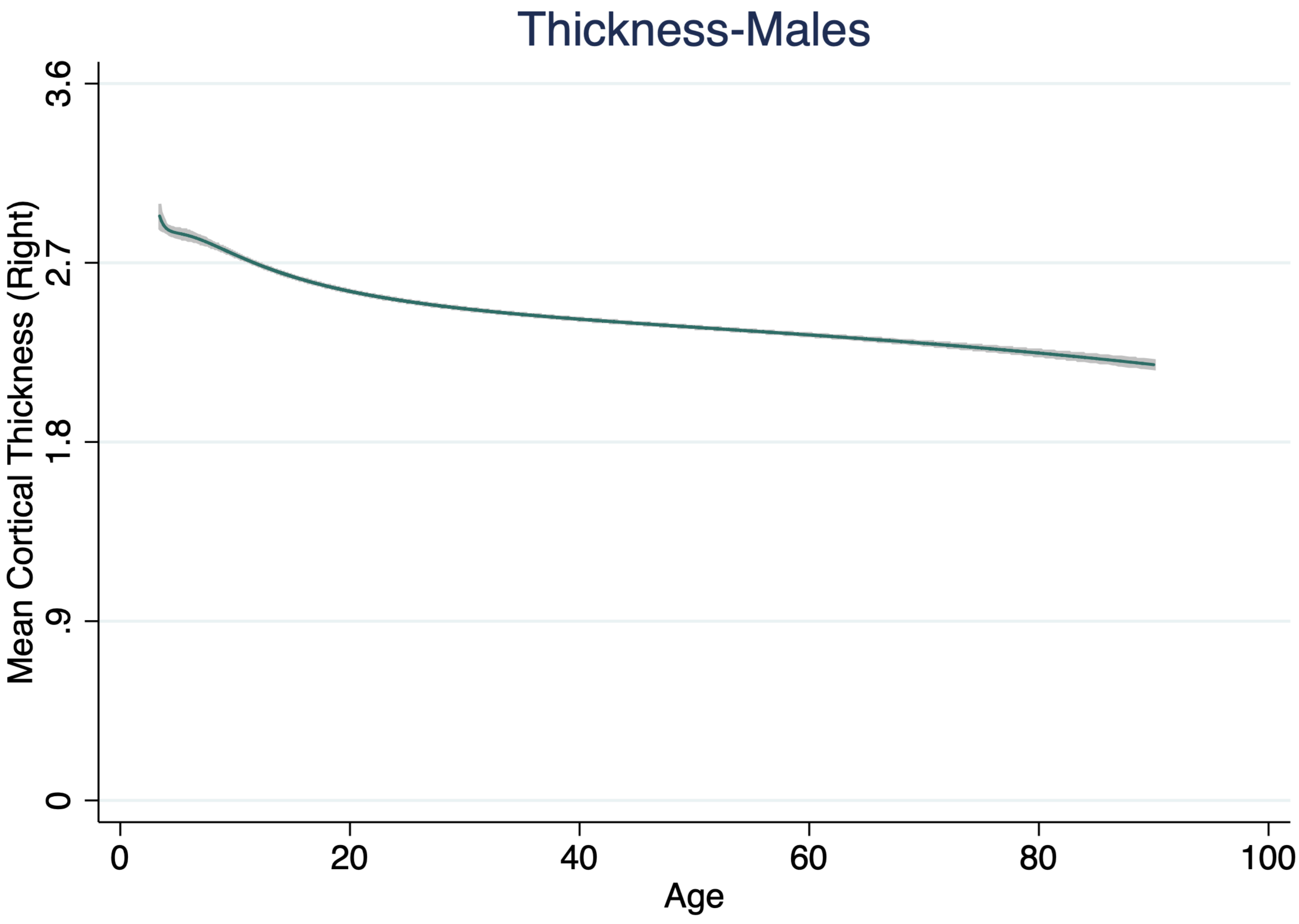
Thickness-Males
Mean Cortical Thickness (Right)
3.6
2.7
1.8
.9
0
0
20
40
60
80
100
Age

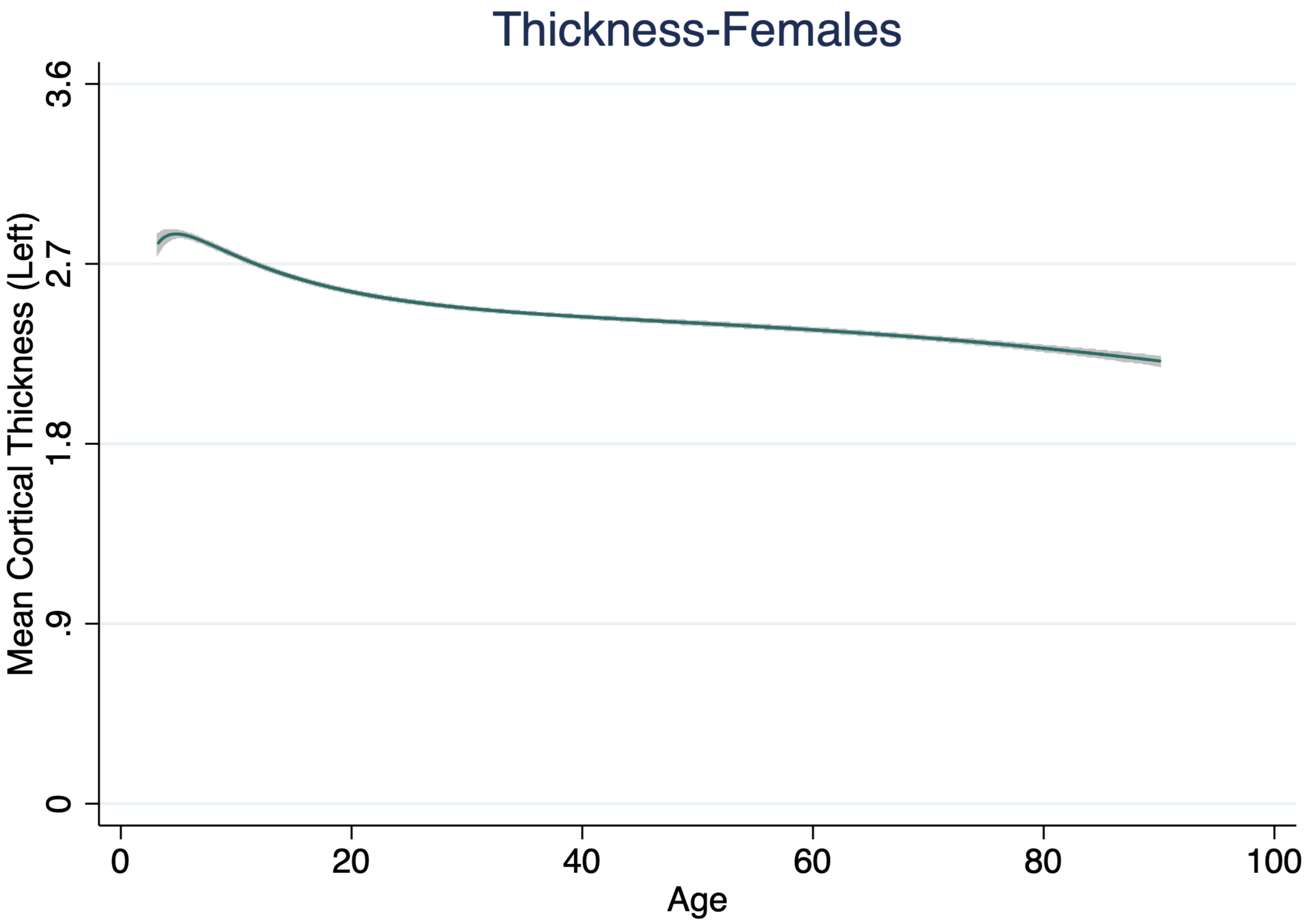

Thickness-Females
Mean Cortical Thickness (Left)
3.6
2.7
1.8
.9
0
0
20
40
60
80
100
Age

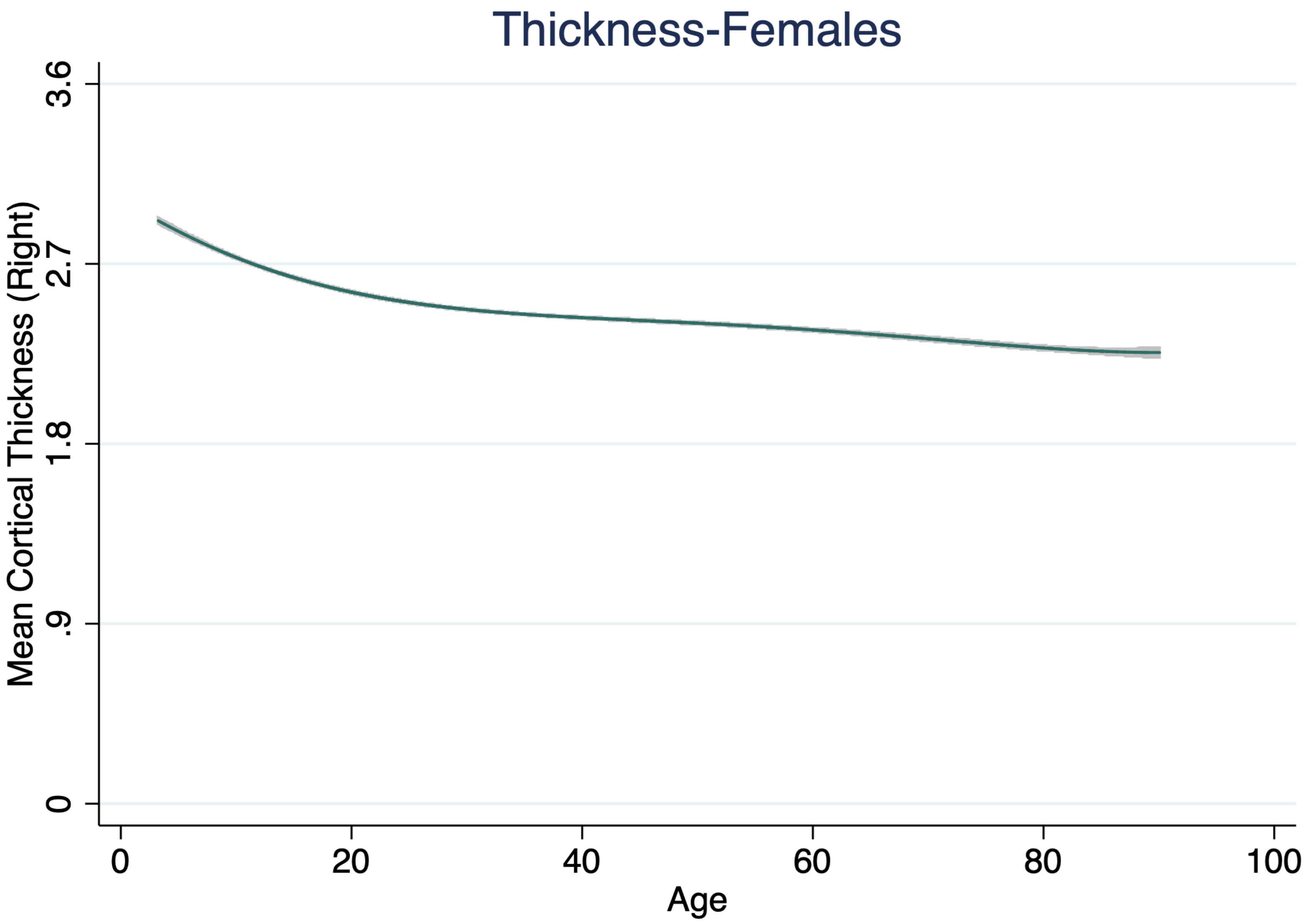
Thickness-Females
Mean Cortical Thickness (Right)
3.6
2.7
1.8
.9
0
0
20
40
60
80
100
Age

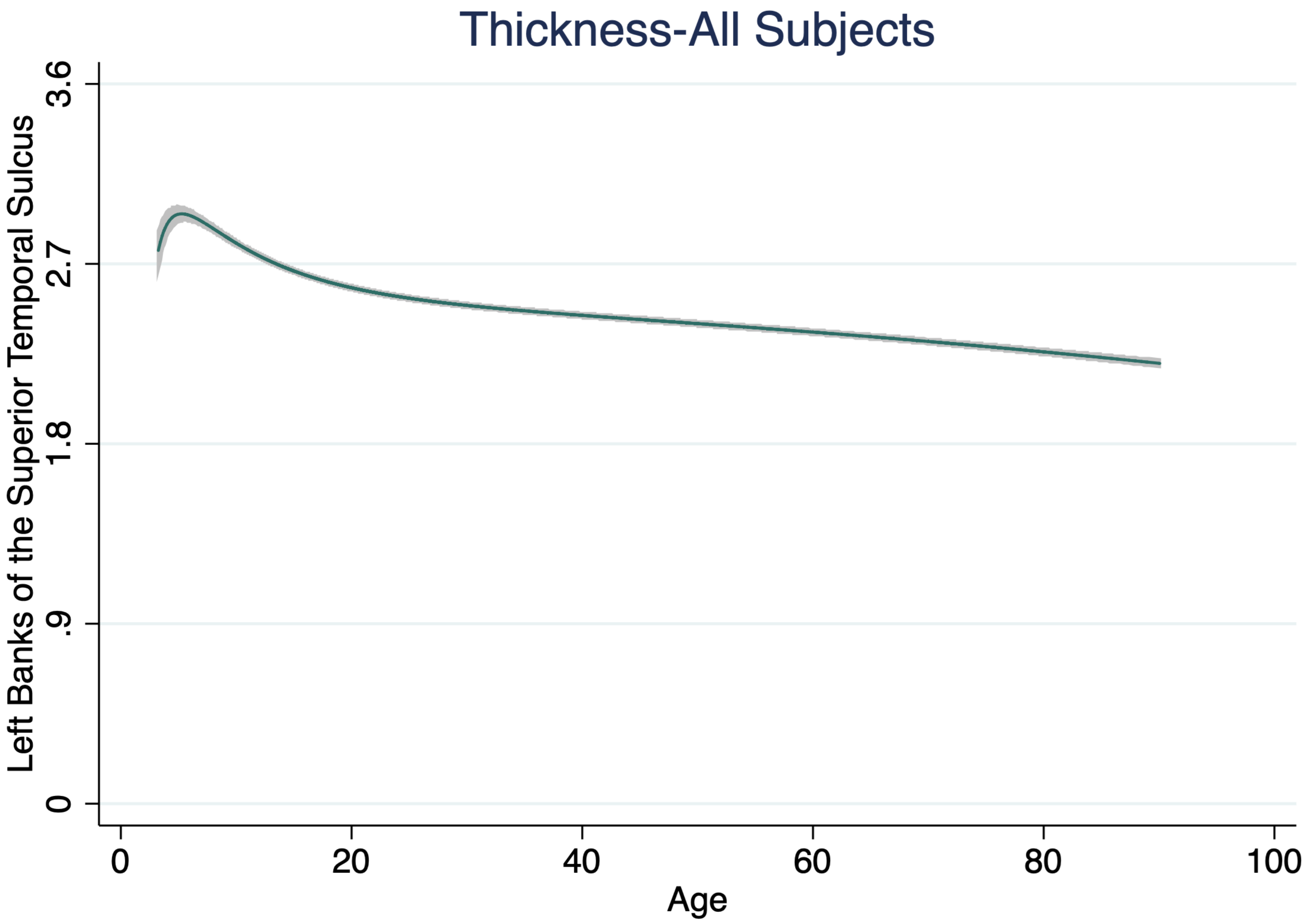
Thickness-All Subjects
Left Banks of the Superior Temporal Sulcus
0
.9
1.8
2.7
3.6
0
20
40
60
80
100
Age

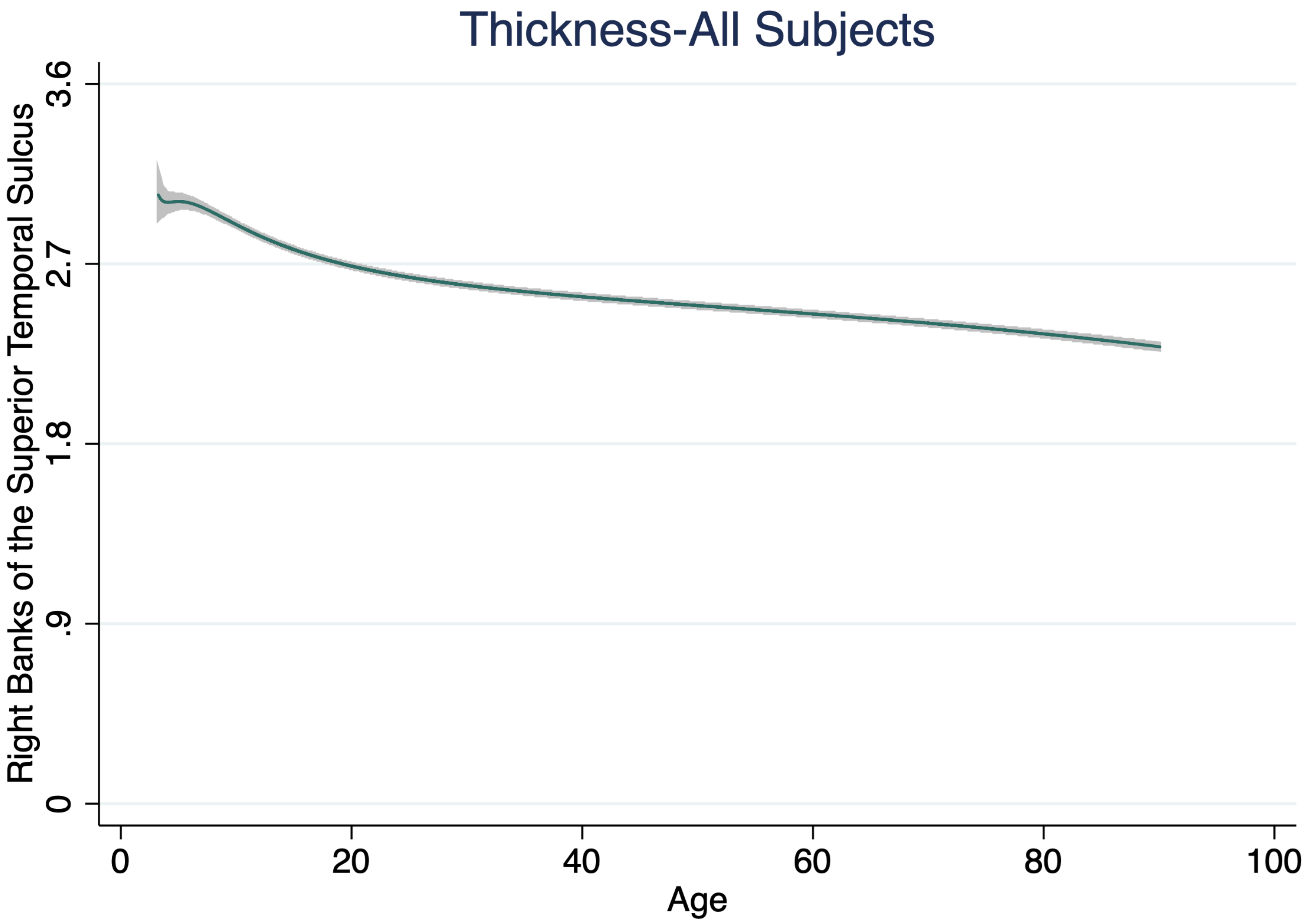

Thickness-All Subjects
Right Banks of the Superior Temporal Sulcus
0
.9
1.8
2.7
3.6
0
20
40
60
80
100
Age

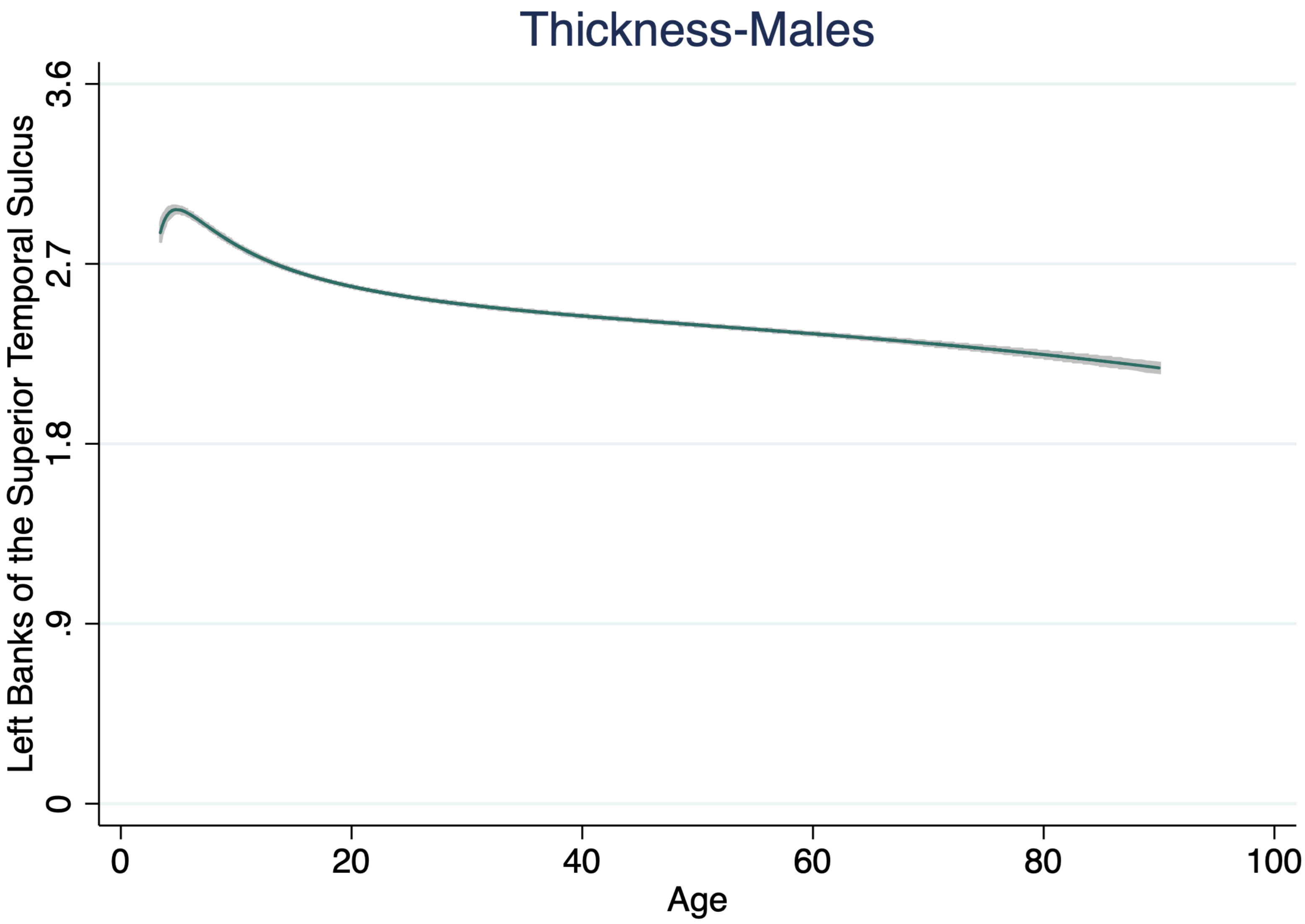

Thickness-Males
Left Banks of the Superior Temporal Sulcus
3.6
2.7
1.8
.9
0
0
20
40
60
80
100
Age

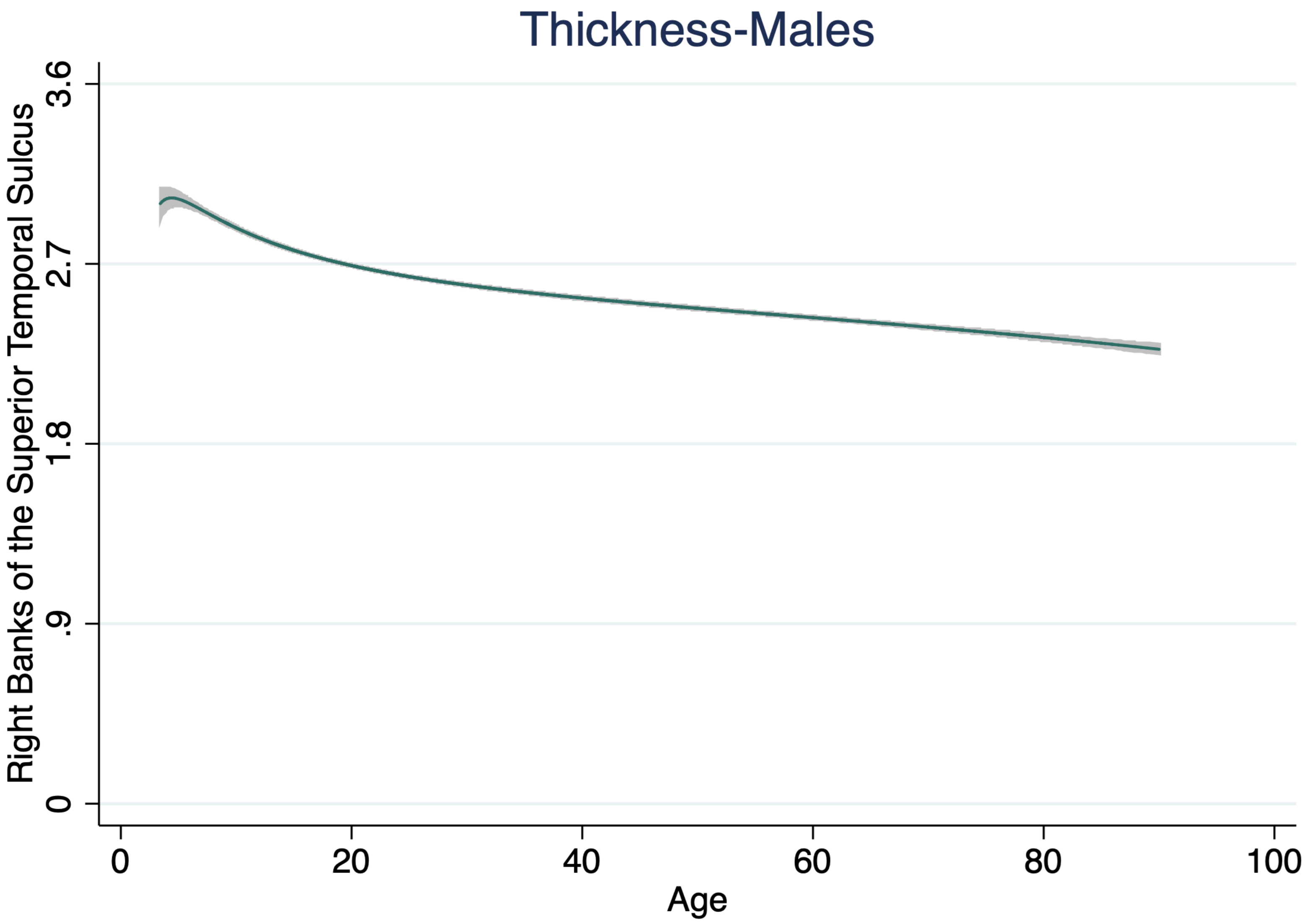
Thickness-Males
Right Banks of the Superior Temporal Sulcus
3.6
2.7
1.8
.9
0
0
20
40
60
80
100
Age

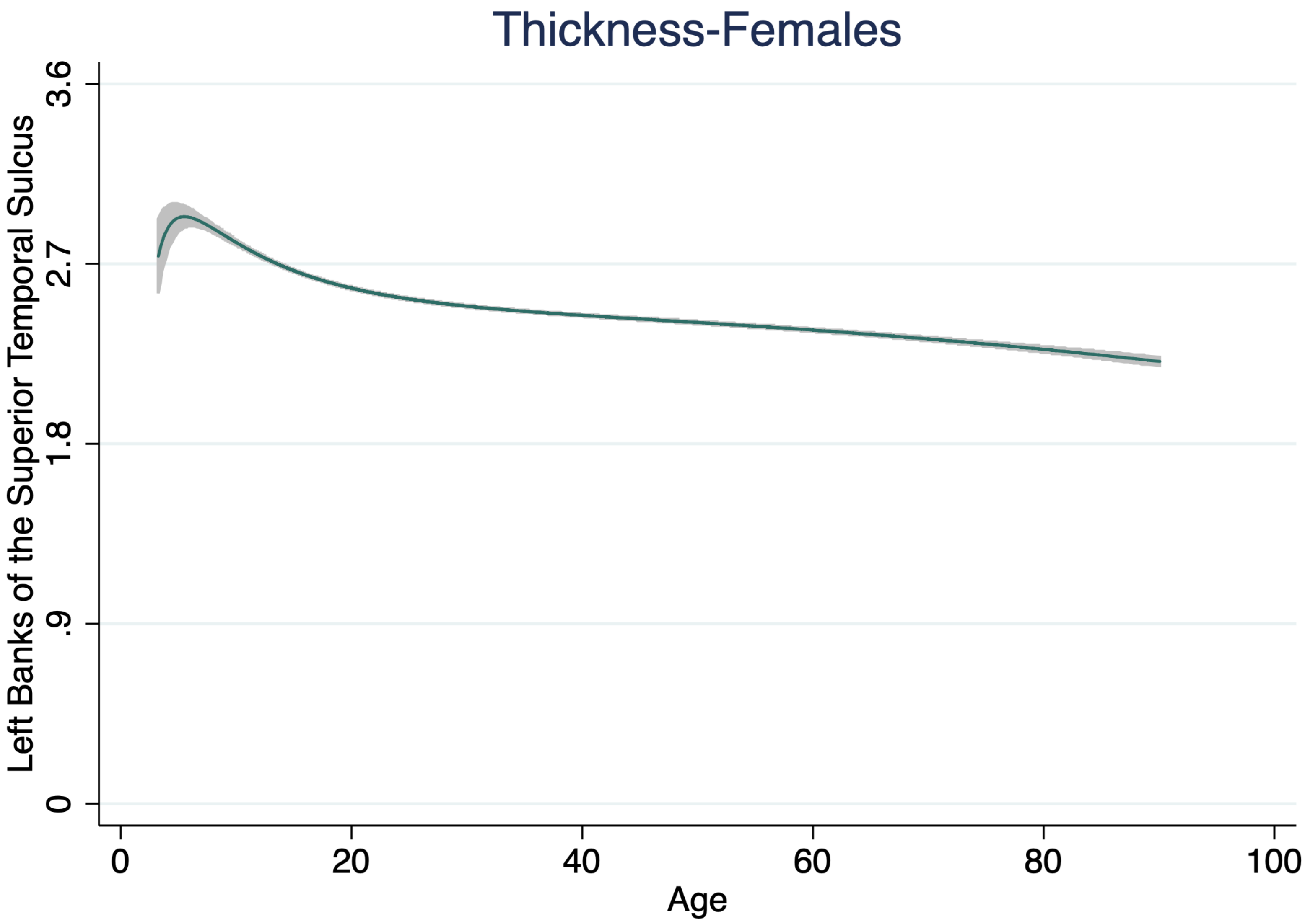
Thickness-Females
Left Banks of the Superior Temporal Sulcus
0
.9
1.8
2.7
3.6
0
20
40
60
80
100
Age

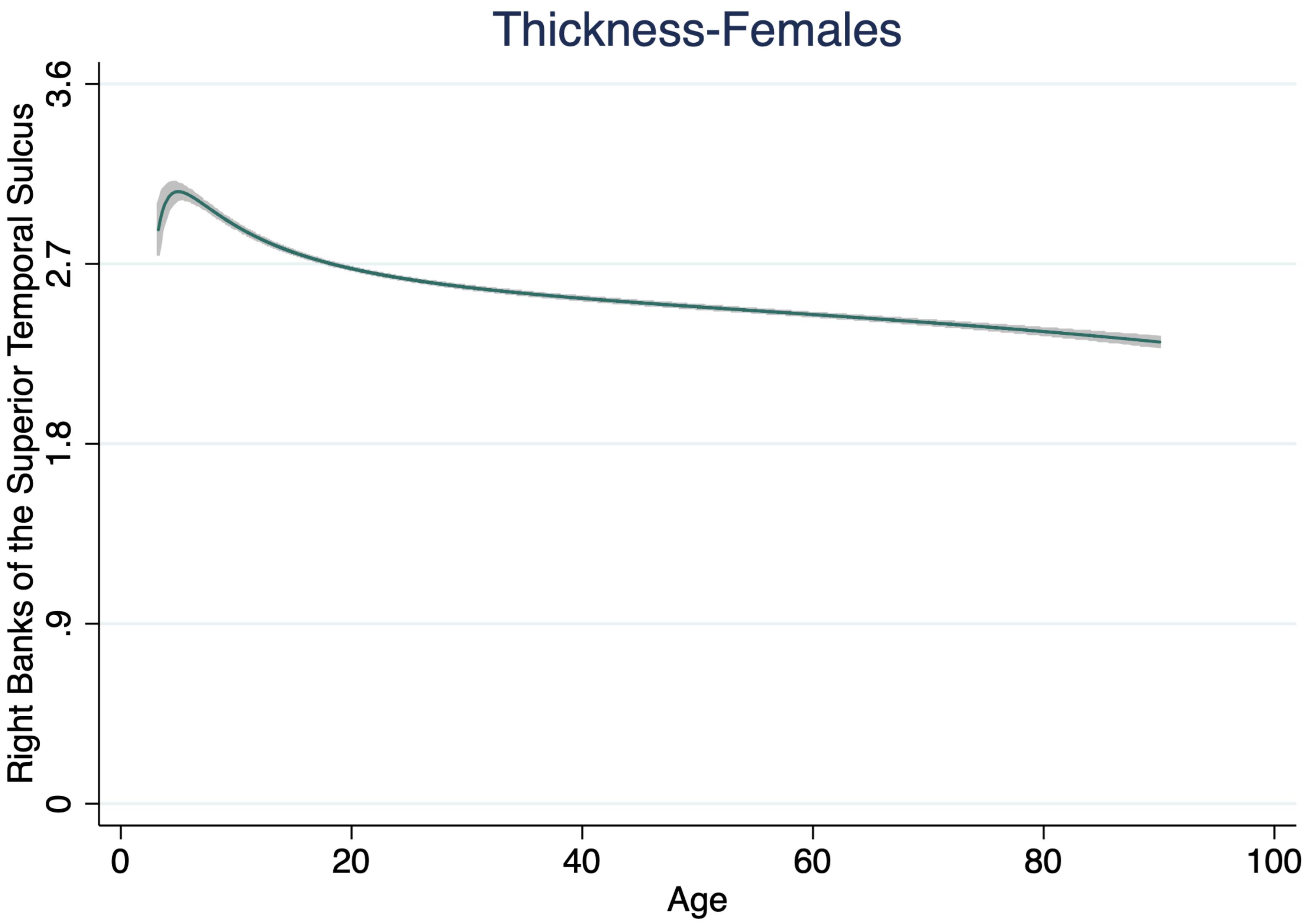

Thickness-Females
Right Banks of the Superior Temporal Sulcus
3.6
2.7
1.8
.9
0
0
20
40
60
80
100
Age

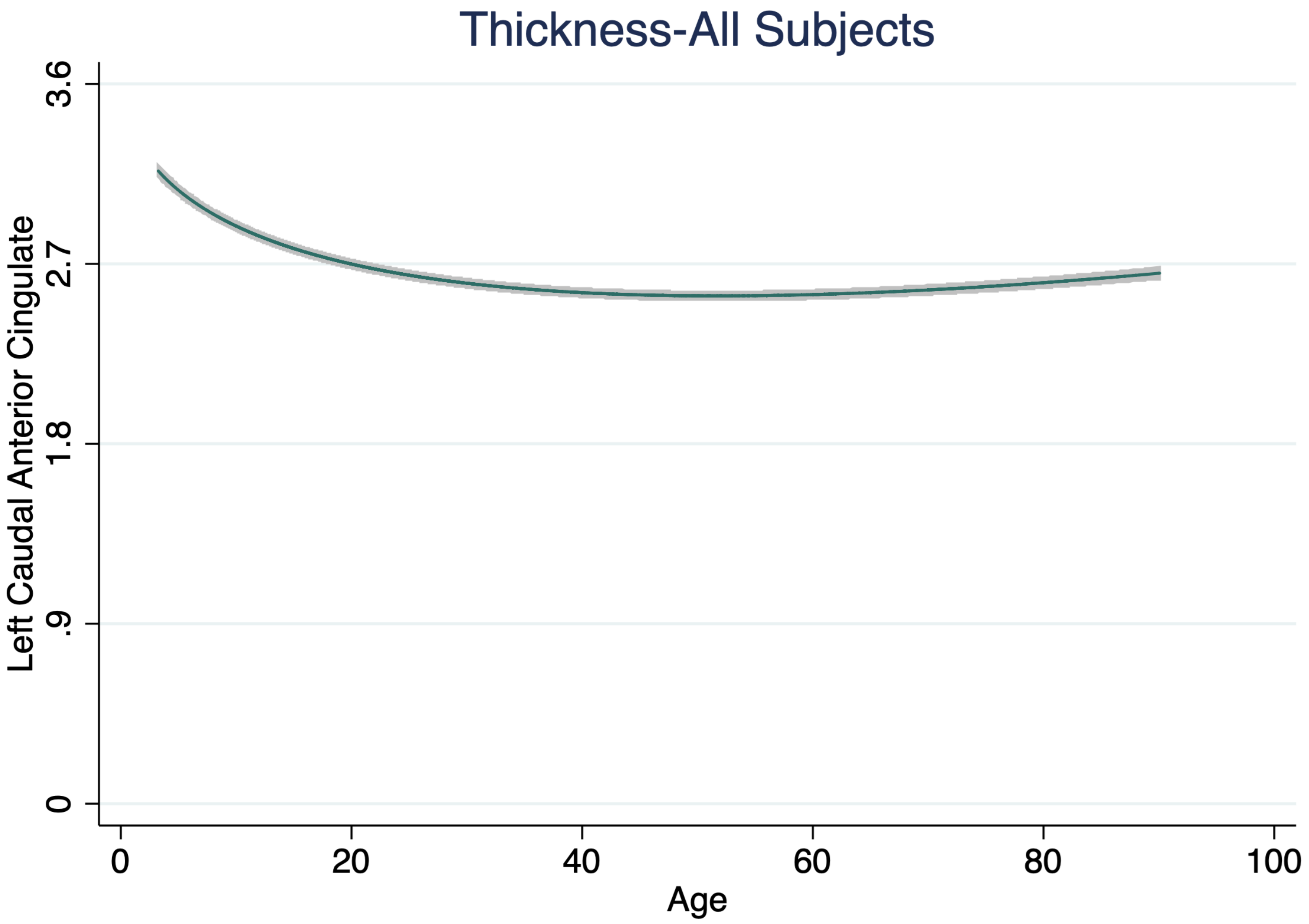

Thickness-All Subjects
Left Caudal Anterior Cingulate
0
.9
1.8
2.7
3.6
0
20
40
60
80
100
Age

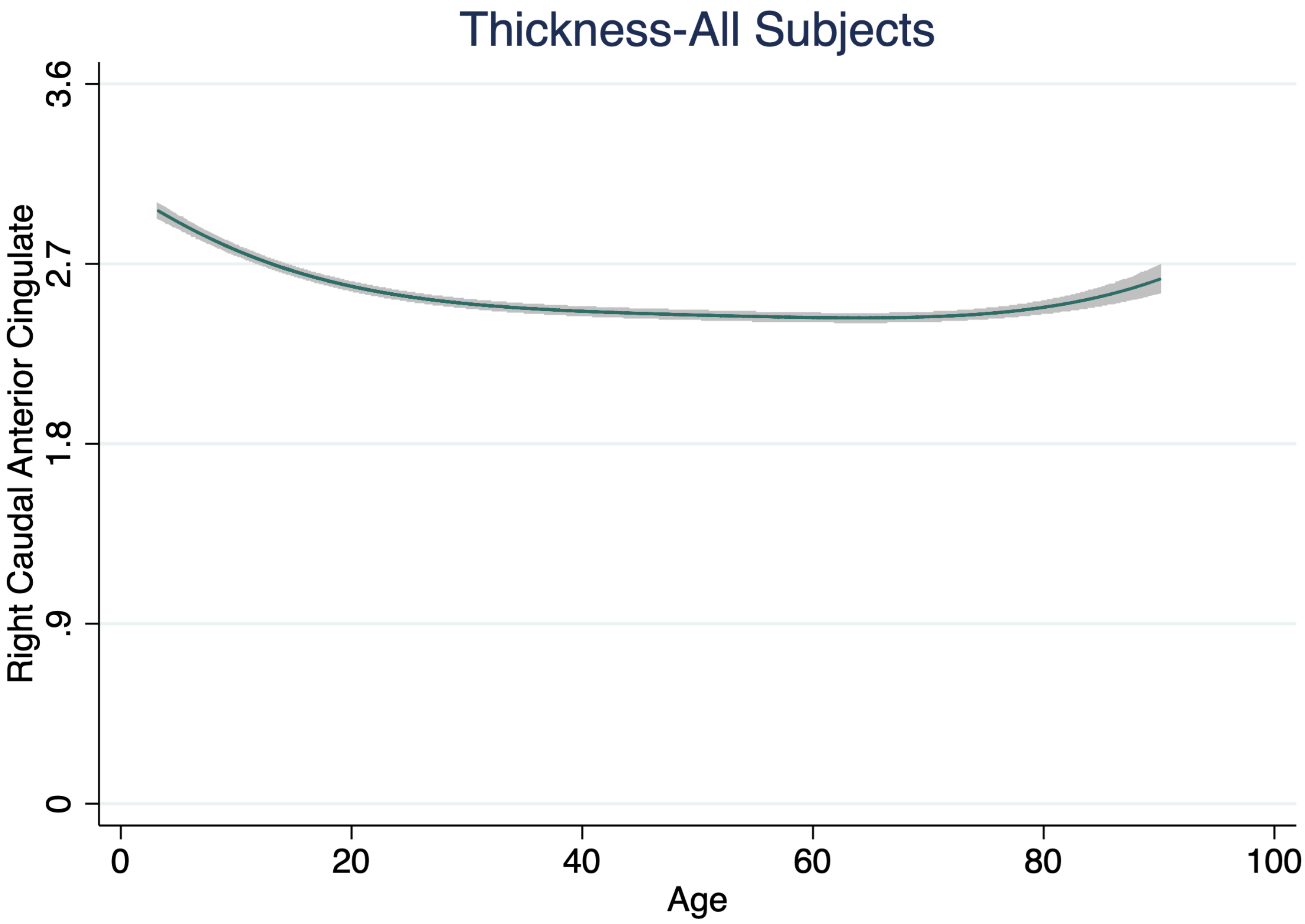

Thickness-All Subjects
Right Caudal Anterior Cingulate
0
.9
1.8
2.7
3.6
0
20
40
60
80
100
Age

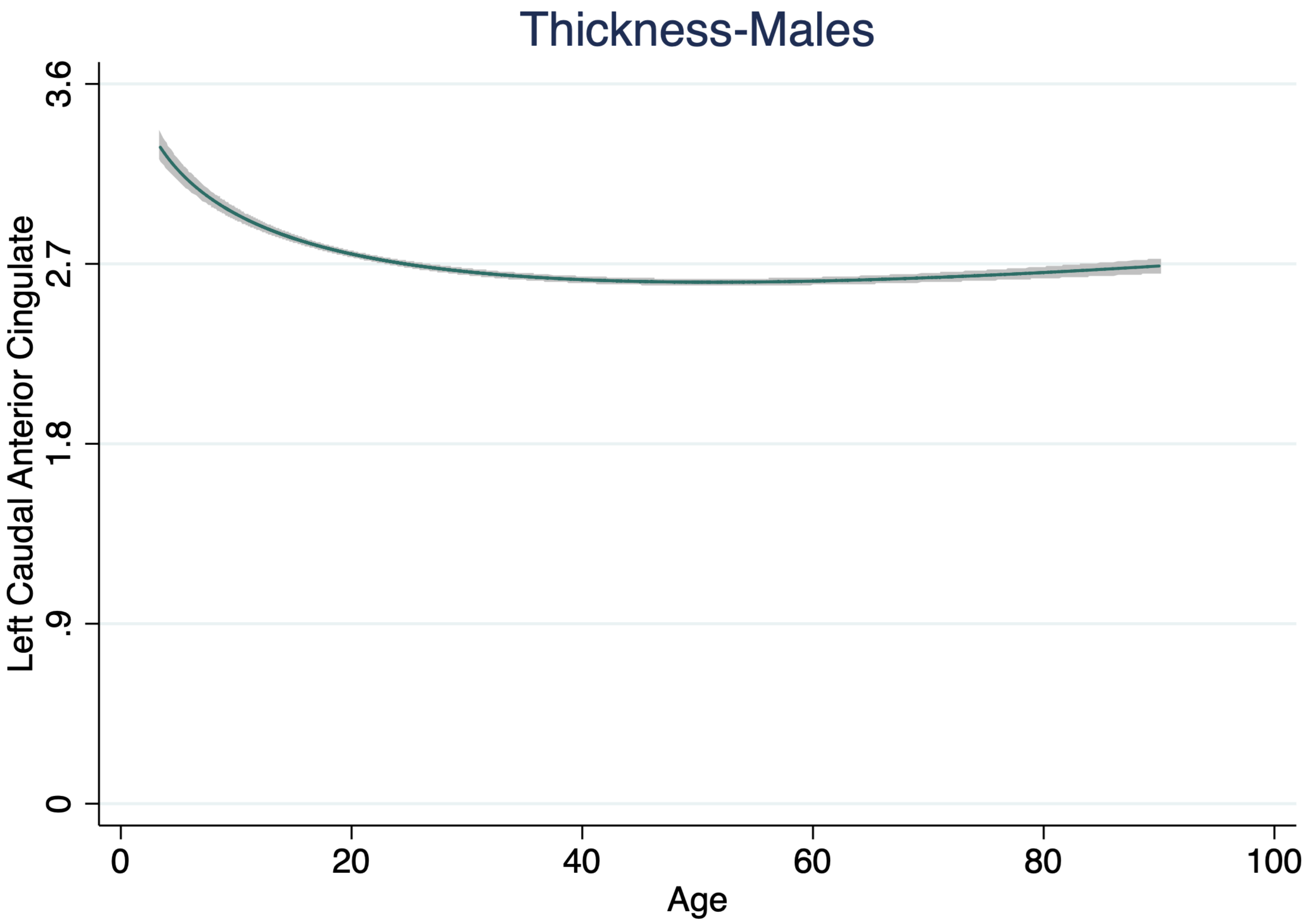
Thickness-Males
Left Caudal Anterior Cingulate
3.6
2.7
1.8
.9
0
0
20
40
60
80
100
Age

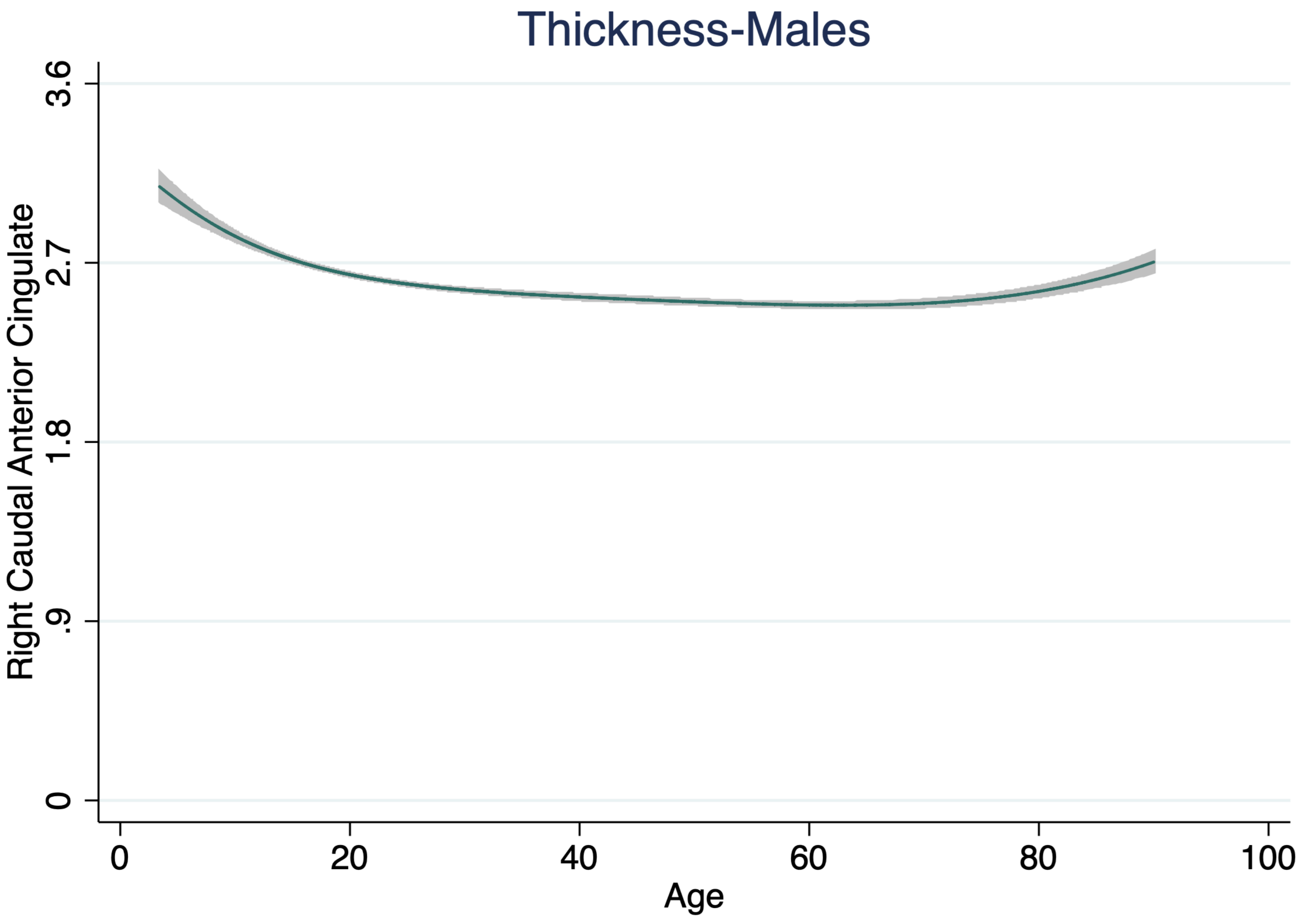

Thickness-Males
Right Caudal Anterior Cingulate
0
.9
1.8
2.7
3.6
0
20
40
60
80
100
Age

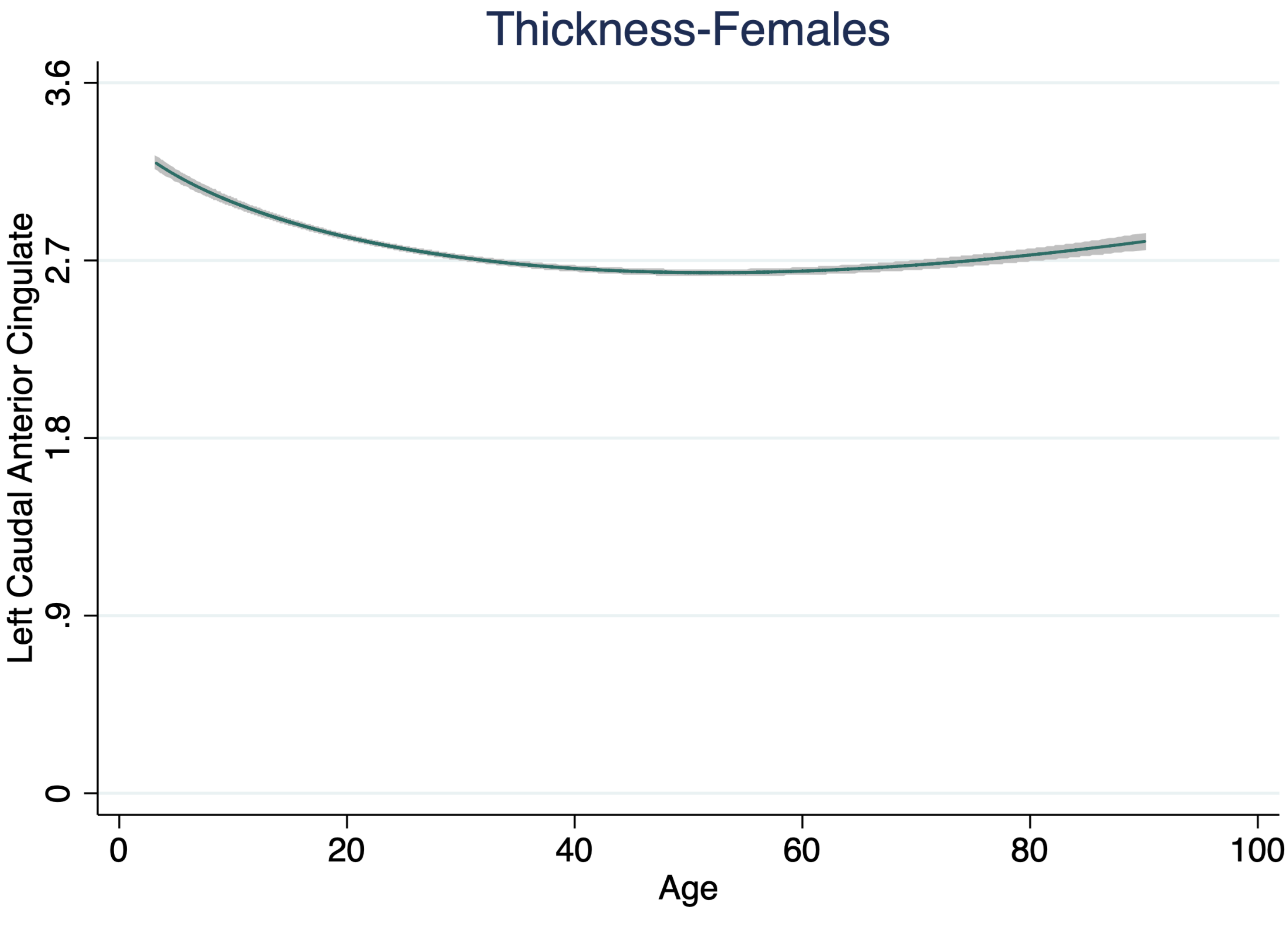

Thickness-Females
Left Caudal Anterior Cingulate
3.6
2.7
1.8
.9
0
0
20
40
60
80
100
Age

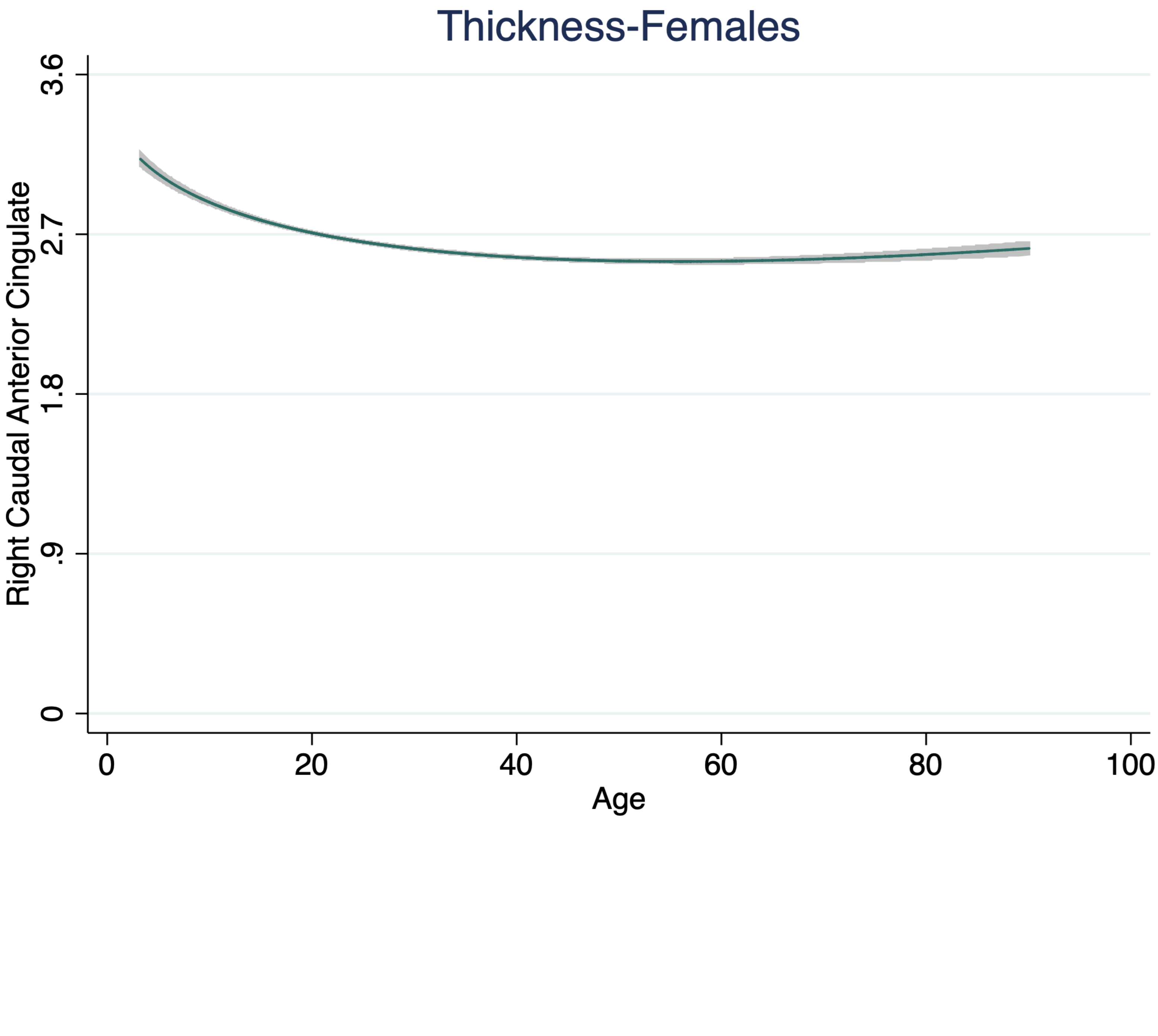
Thickness-Females
Right Caudal Anterior Cingulate
3.6
2.7
1.8
.9
0
0
20
40
60
80
100
Age

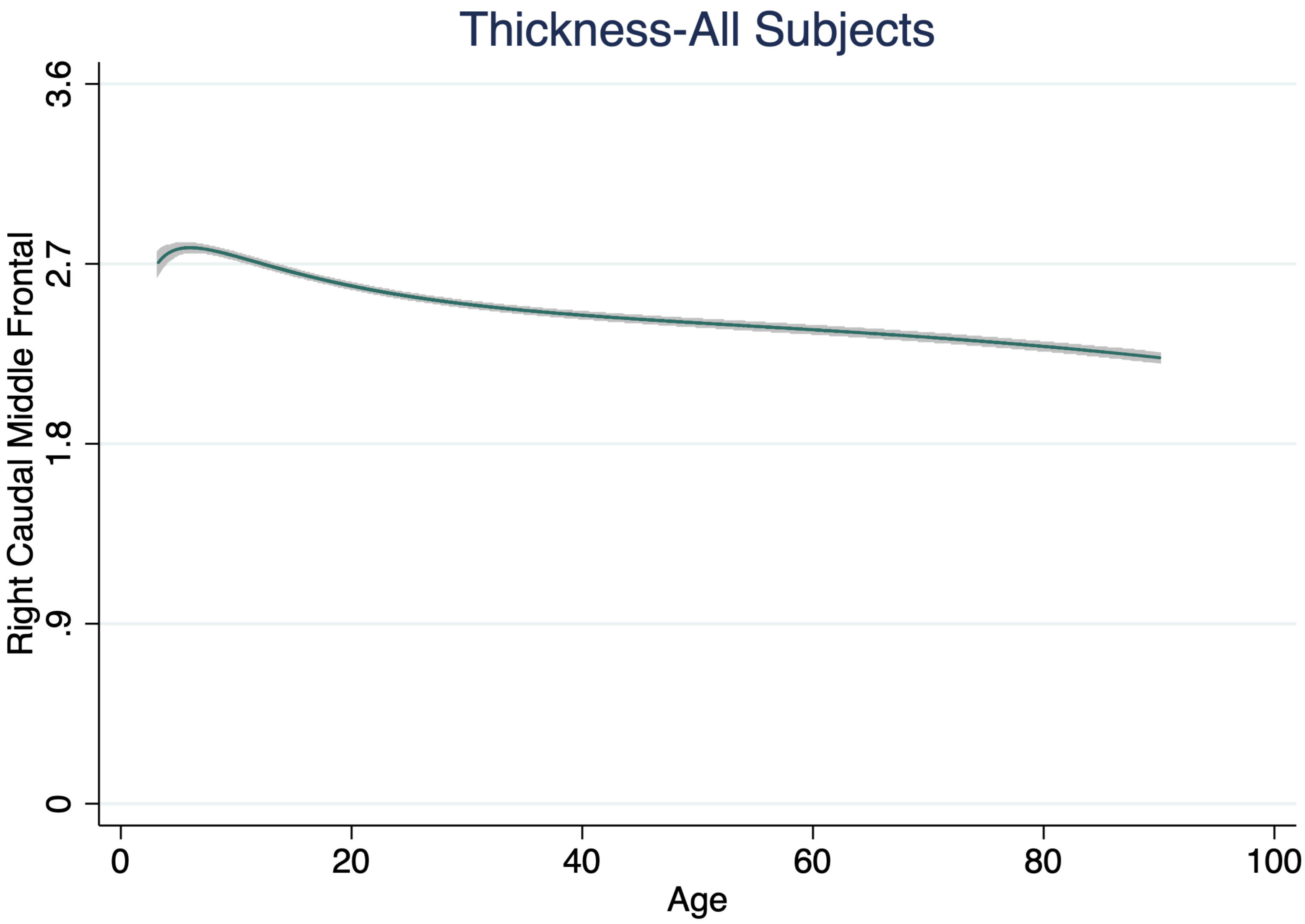
Thickness-All Subjects
Right Caudal Middle Frontal
3.6
2.7
1.8
.9
0
0
20
40
60
80
100
Age

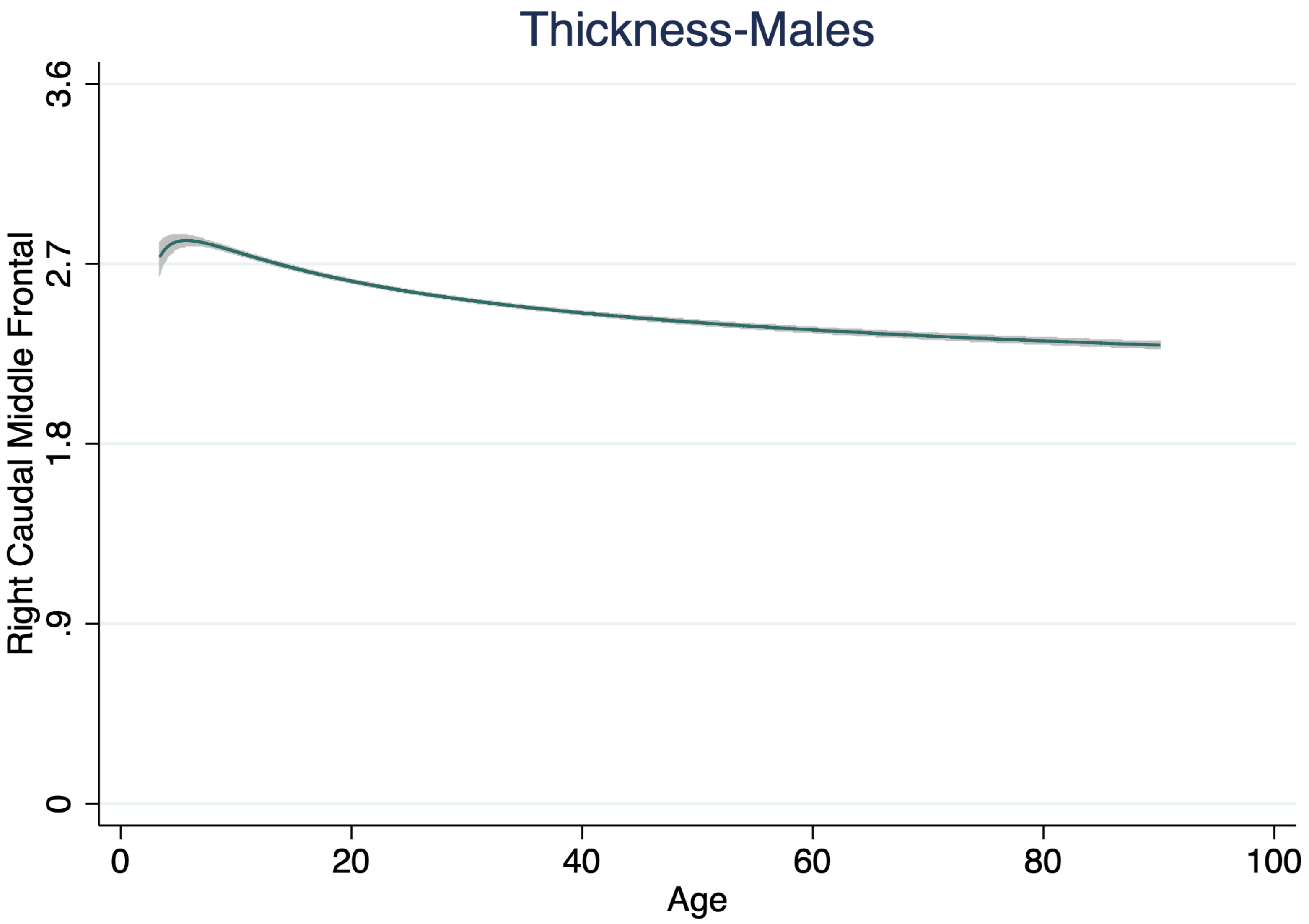

Thickness-Males
Right Caudal Middle Frontal
3.6
2.7
1.8
.9
0
0
20
40
60
80
100
Age

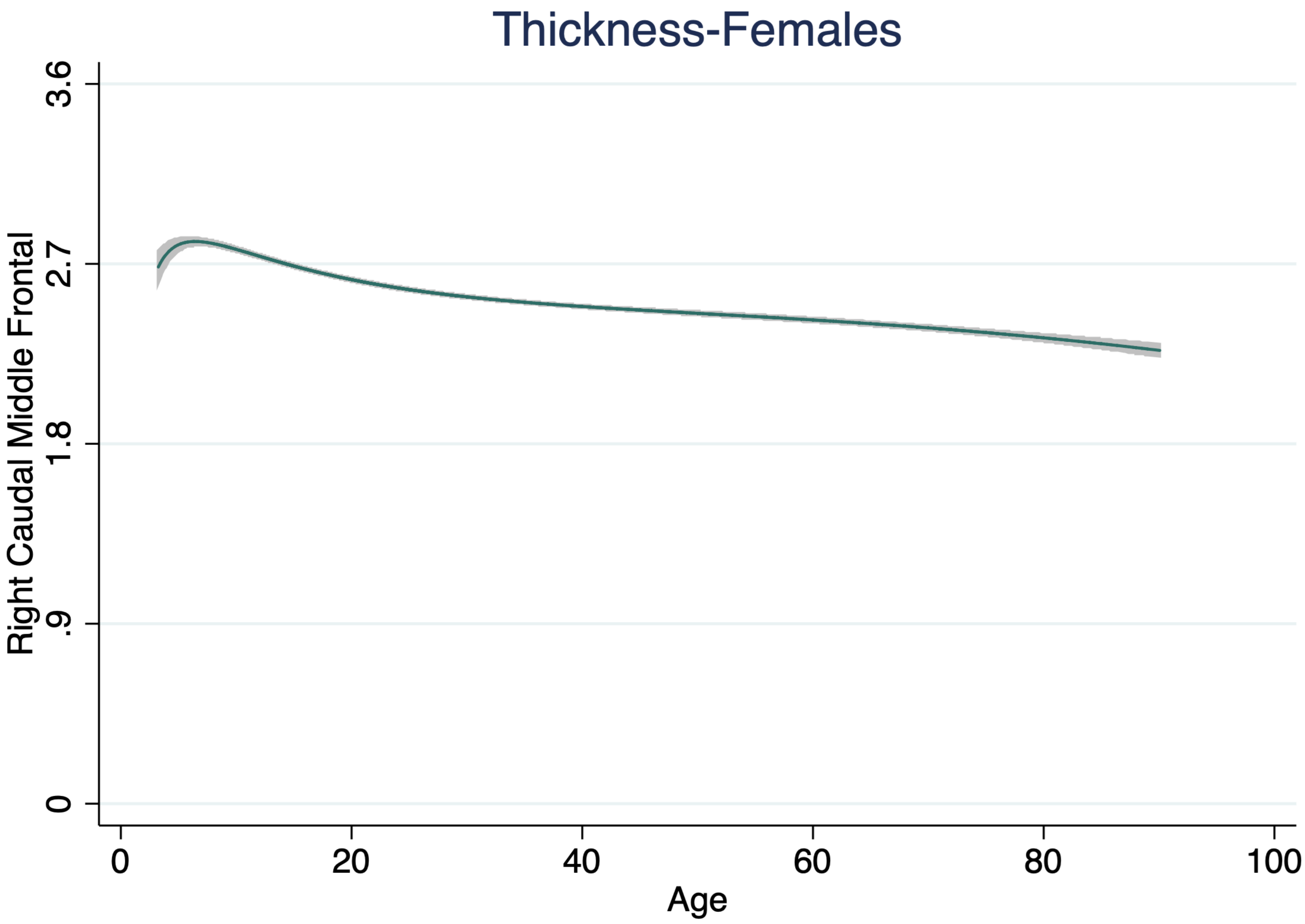

Thickness-Females
Right Caudal Middle Frontal
3.6
2.7
1.8
.9
0
0
20
40
60
80
100
Age

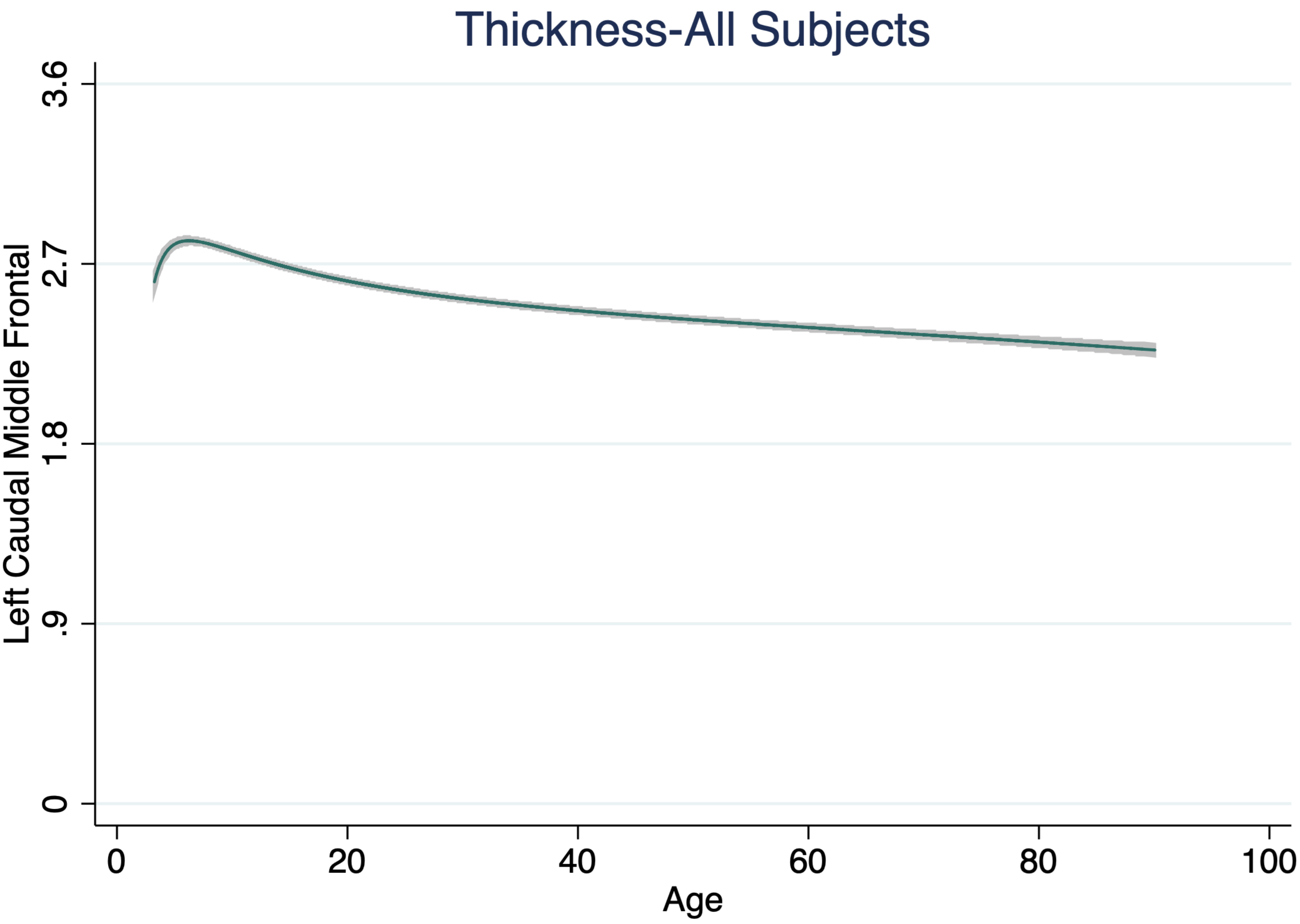

Thickness-All Subjects
Left Caudal Middle Frontal
3.6
2.7
1.8
.9
0
0
20
40
60
80
100
Age

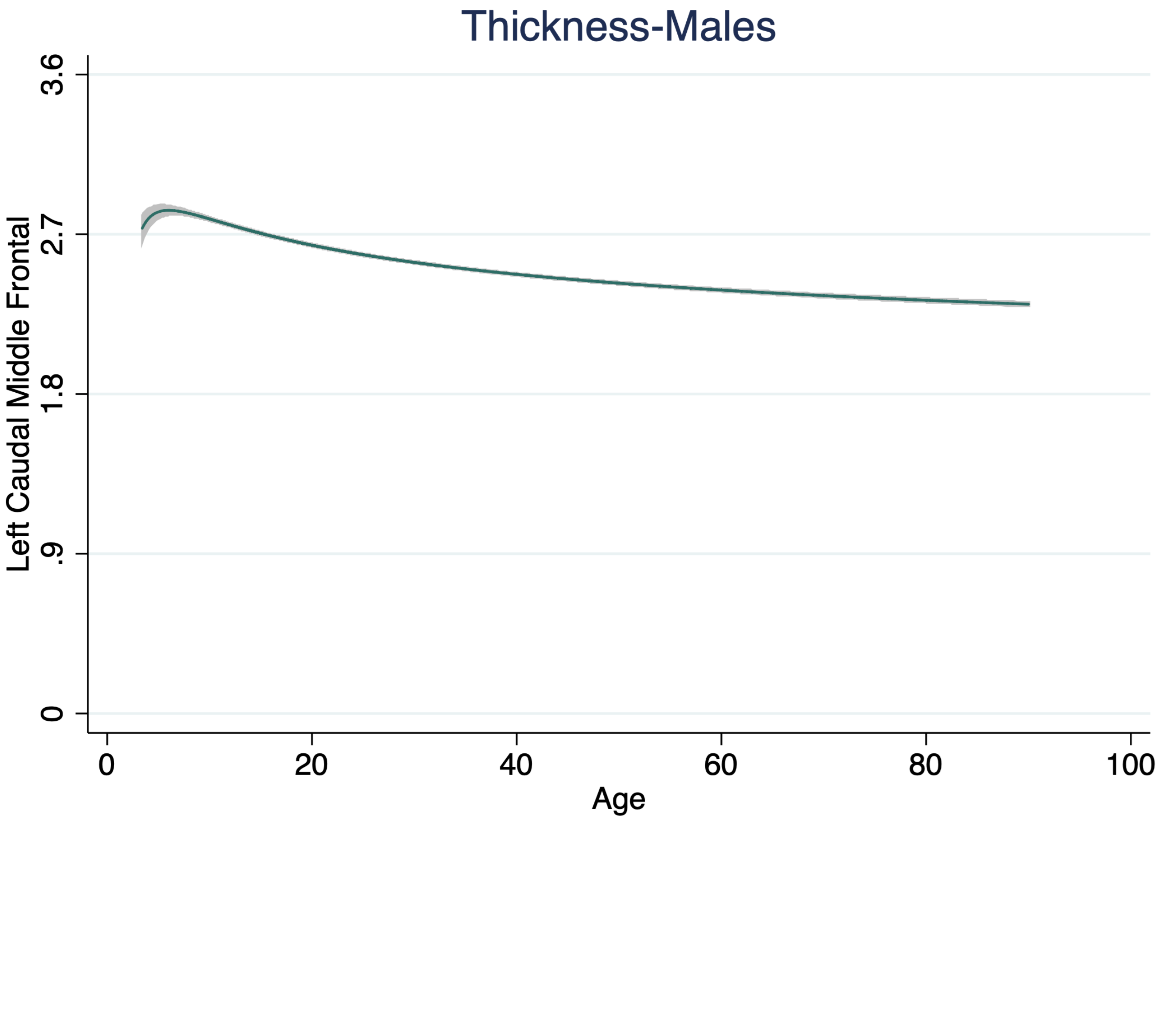

Thickness-Males
Left Caudal Middle Frontal
3.6
2.7
1.8
.9
0
0
20
40
60
80
100
Age

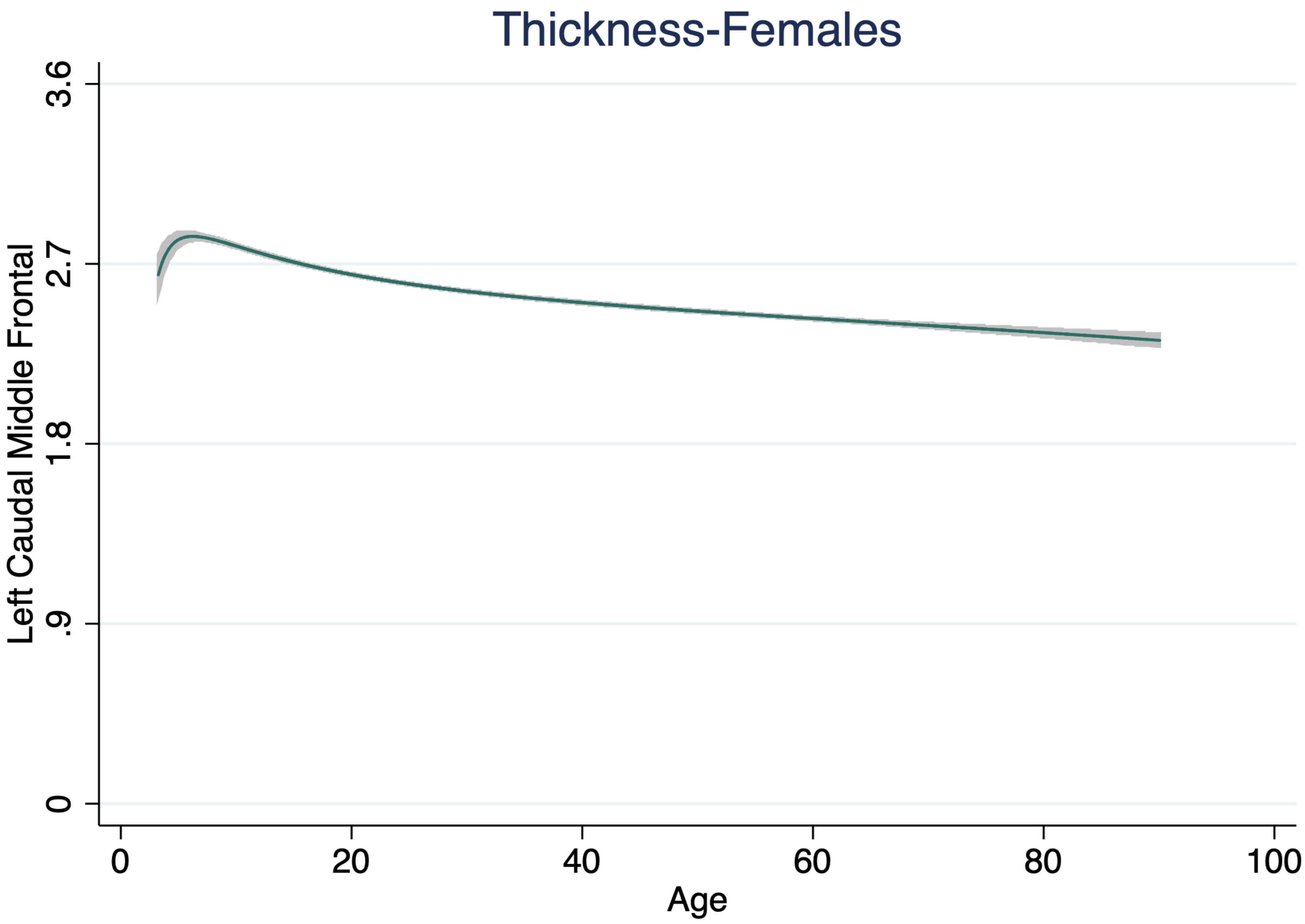

Thickness-Females
Left Caudal Middle Frontal
3.6
2.7
1.8
.9
0
0
20
40
60
80
100
Age

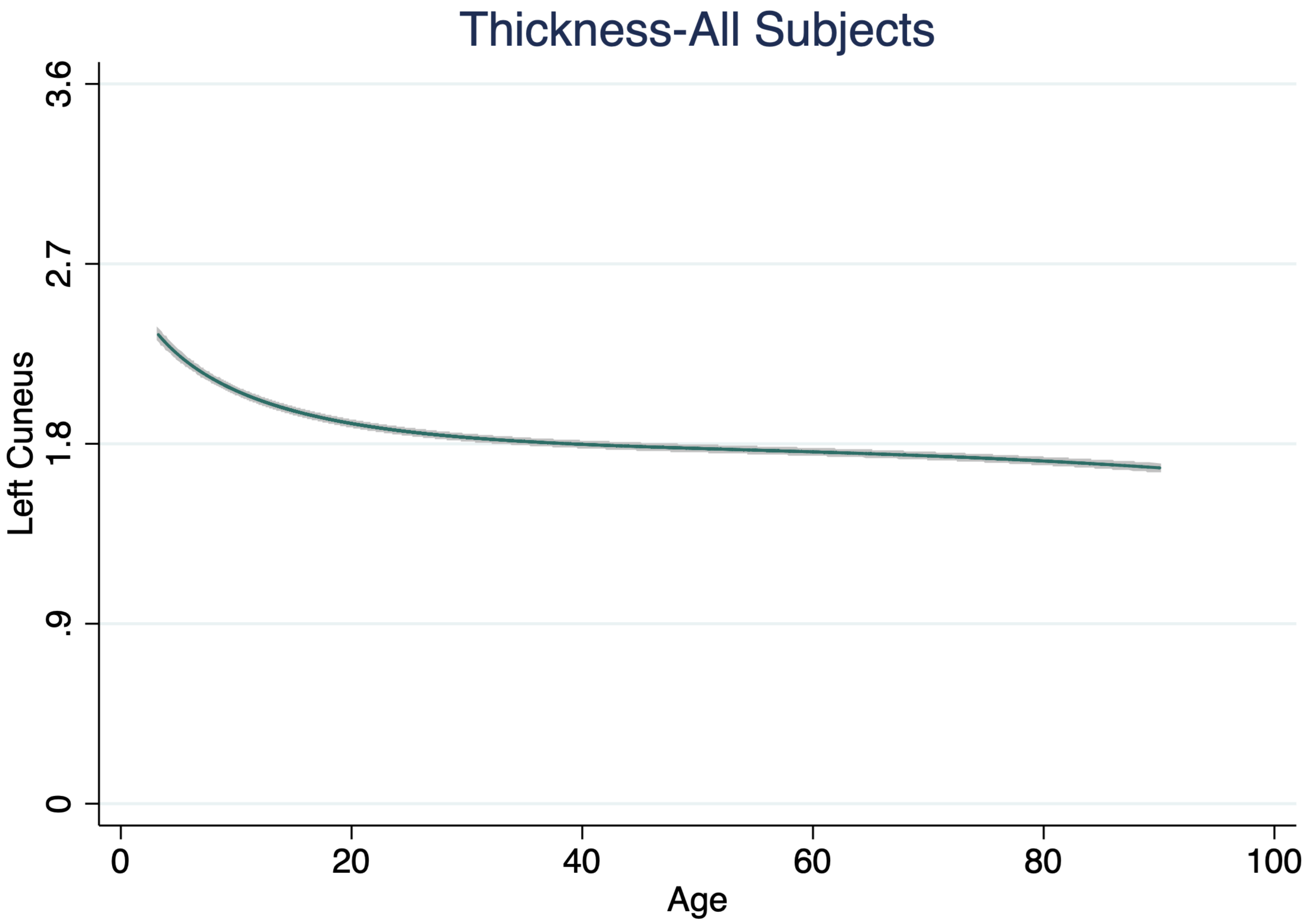
Thickness-All Subjects
Left Cuneus
3.6
2.7
1.8
.9
0
0
20
40
60
80
100
Age

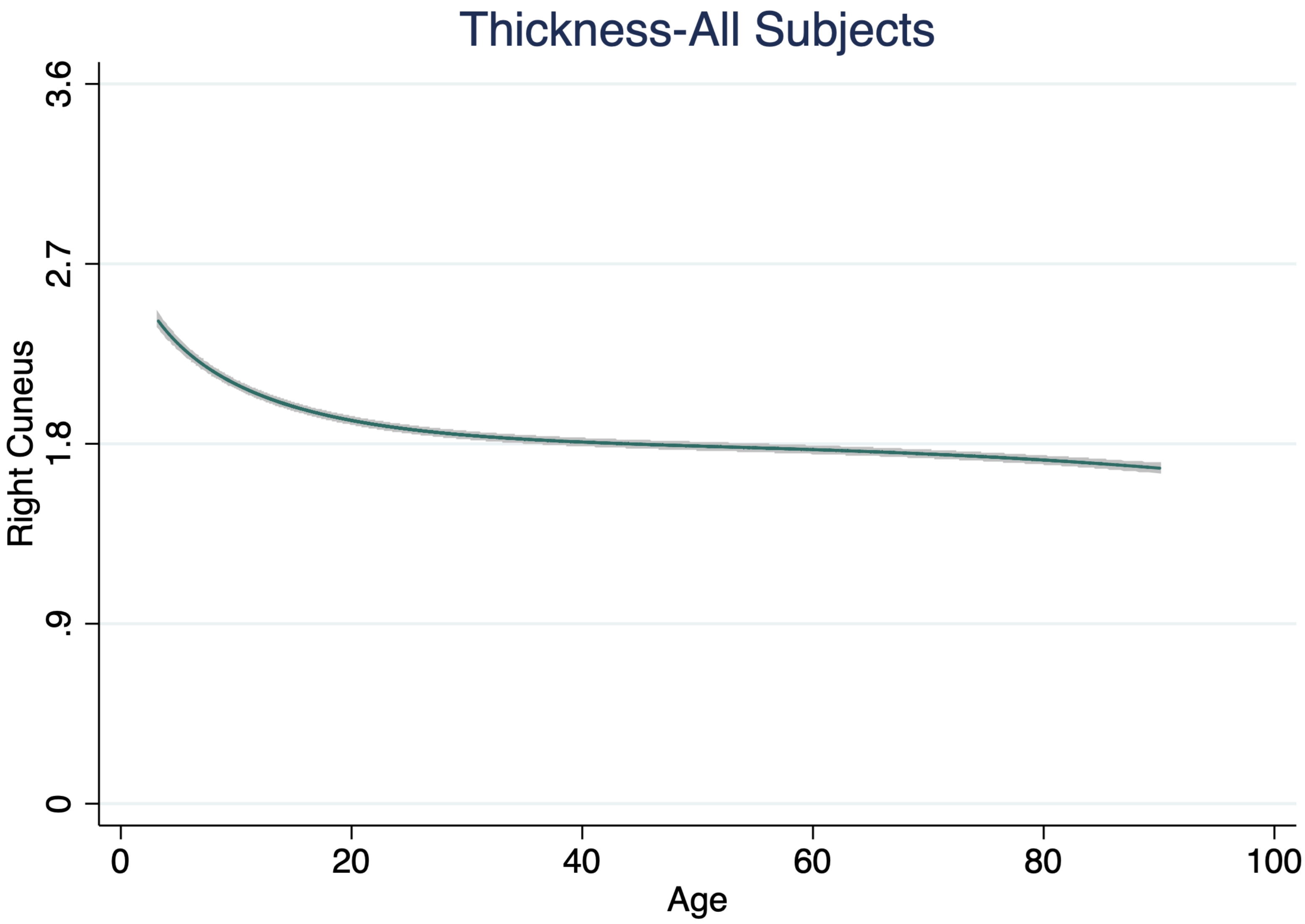
Thickness-All Subjects
Right Cuneus
3.6
2.7
1.8
.9
0
0
20
40
60
80
100
Age

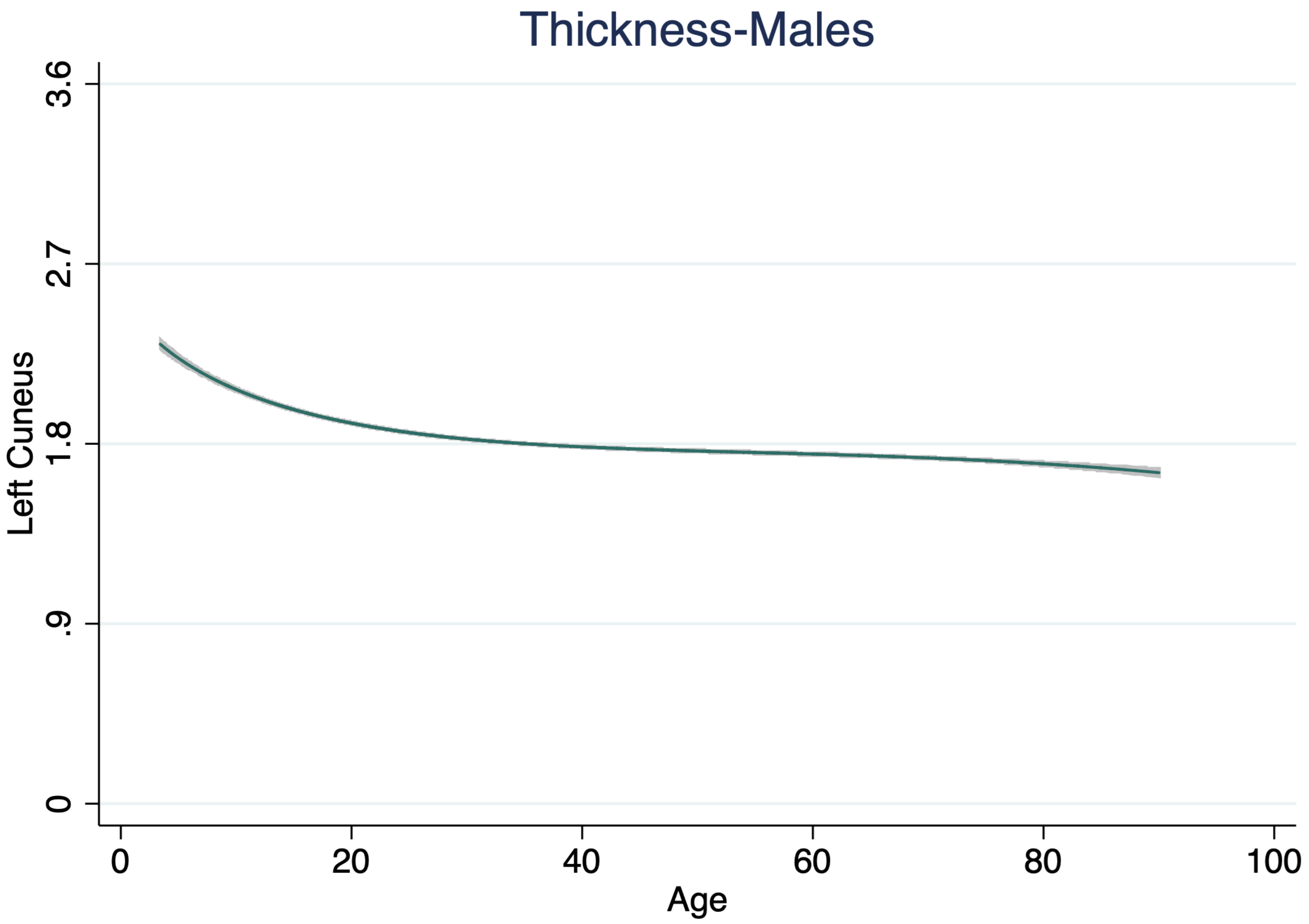

Thickness-Males
Left Cuneus
3.6
2.7
1.8
.9
0
0
20
40
60
80
100
Age

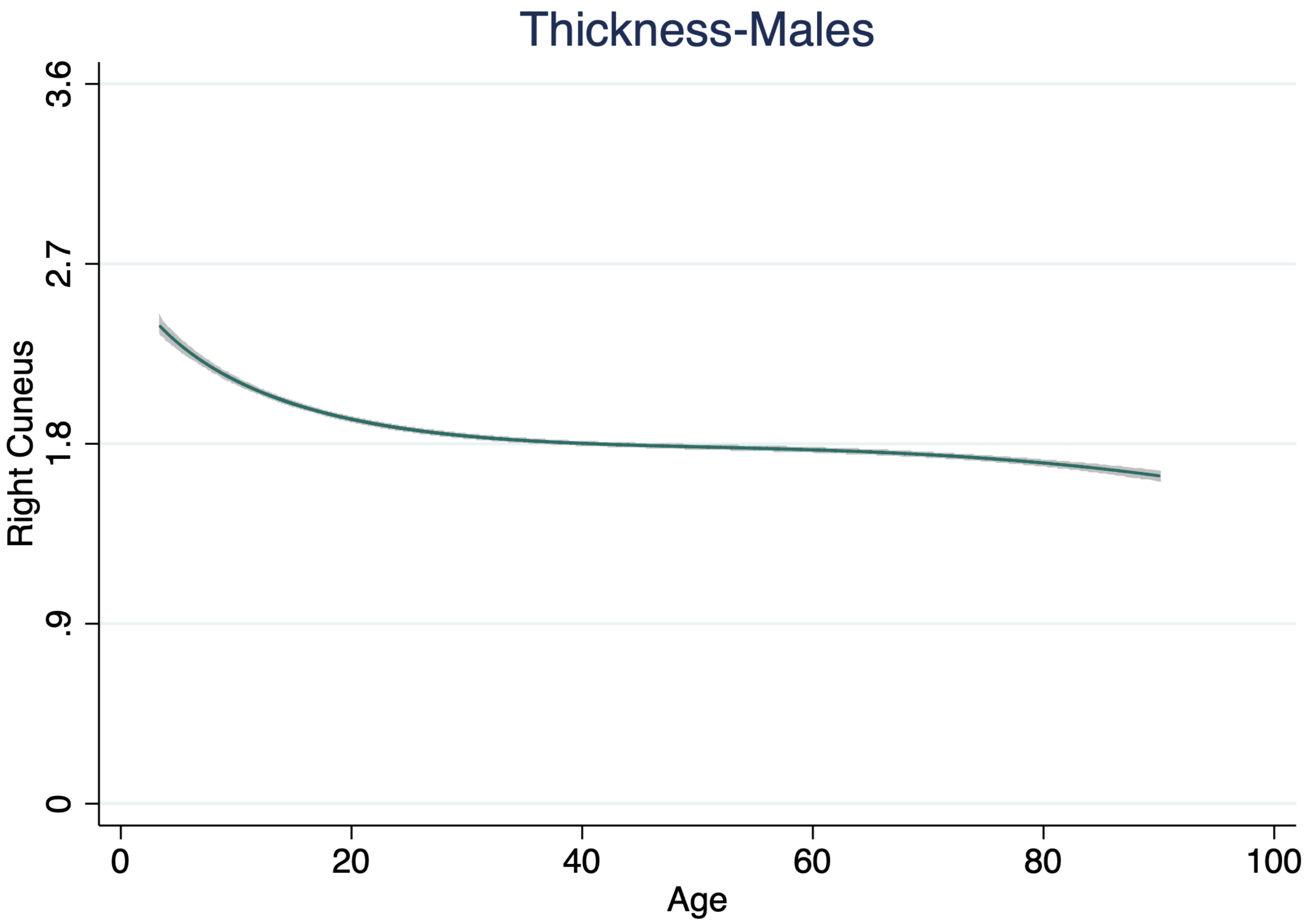

Thickness-Males
Right Cuneus
3.6
2.7
1.8
.9
0
0
20
40
60
80
100
Age

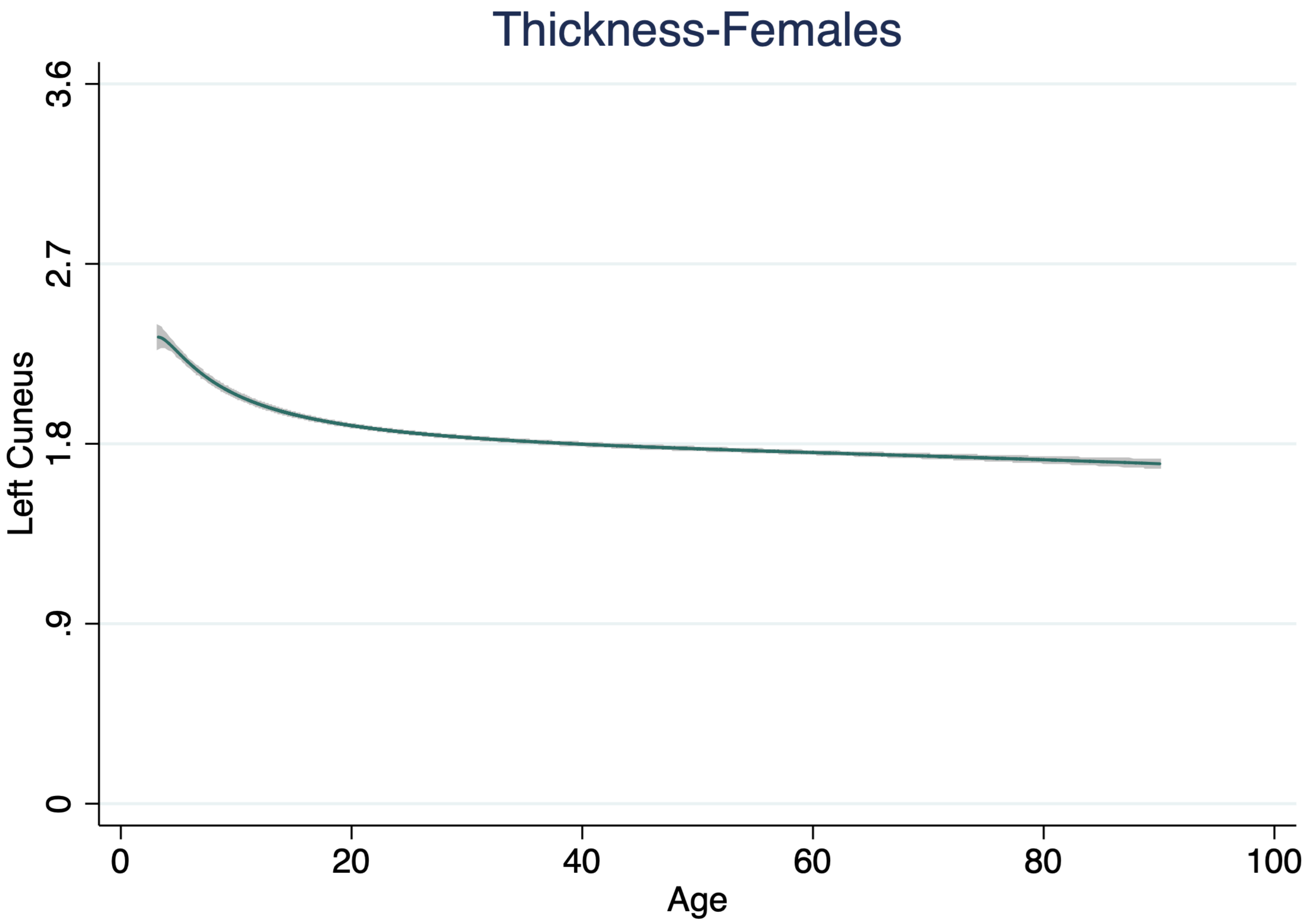

Thickness-Females
Left Cuneus
3.6
2.7
1.8
.9
0
0
20
40
60
80
100
Age

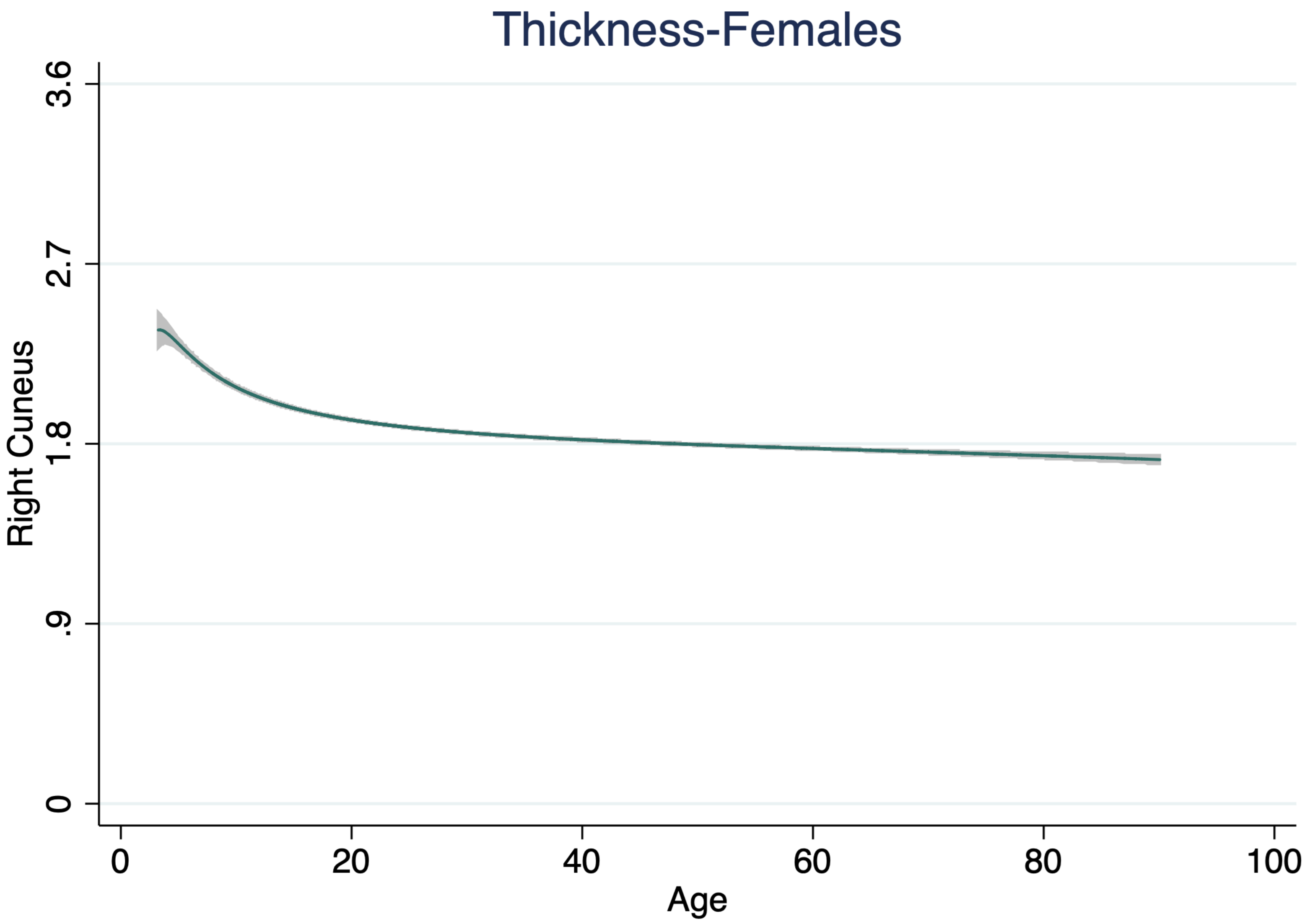

Thickness-Females
Right Cuneus
3.6
2.7
1.8
.9
0
0
20
40
60
80
100
Age

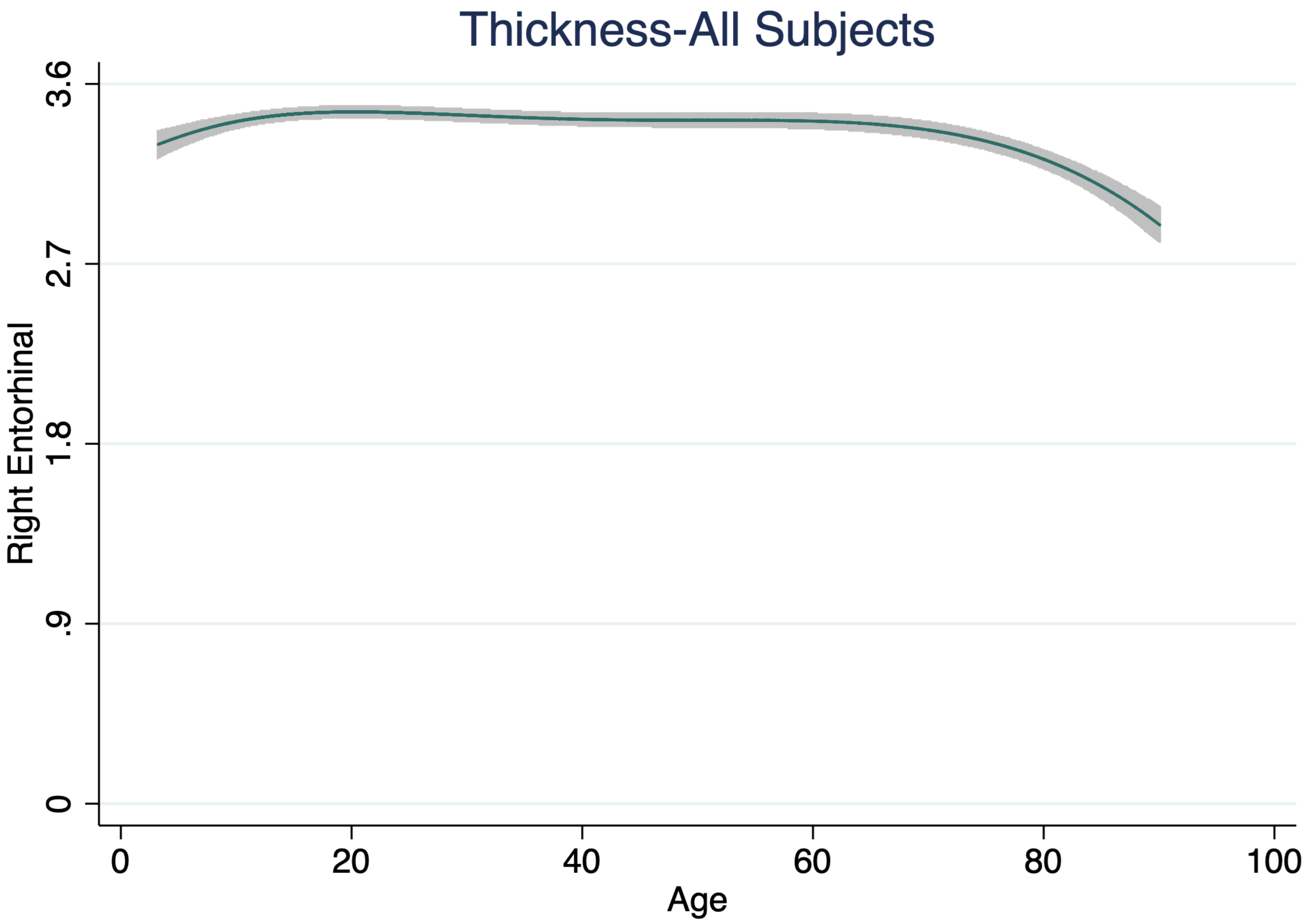

Thickness-All Subjects
Right Entorhinal
3.6
2.7
1.8
.9
0
0
20
40
60
80
100
Age

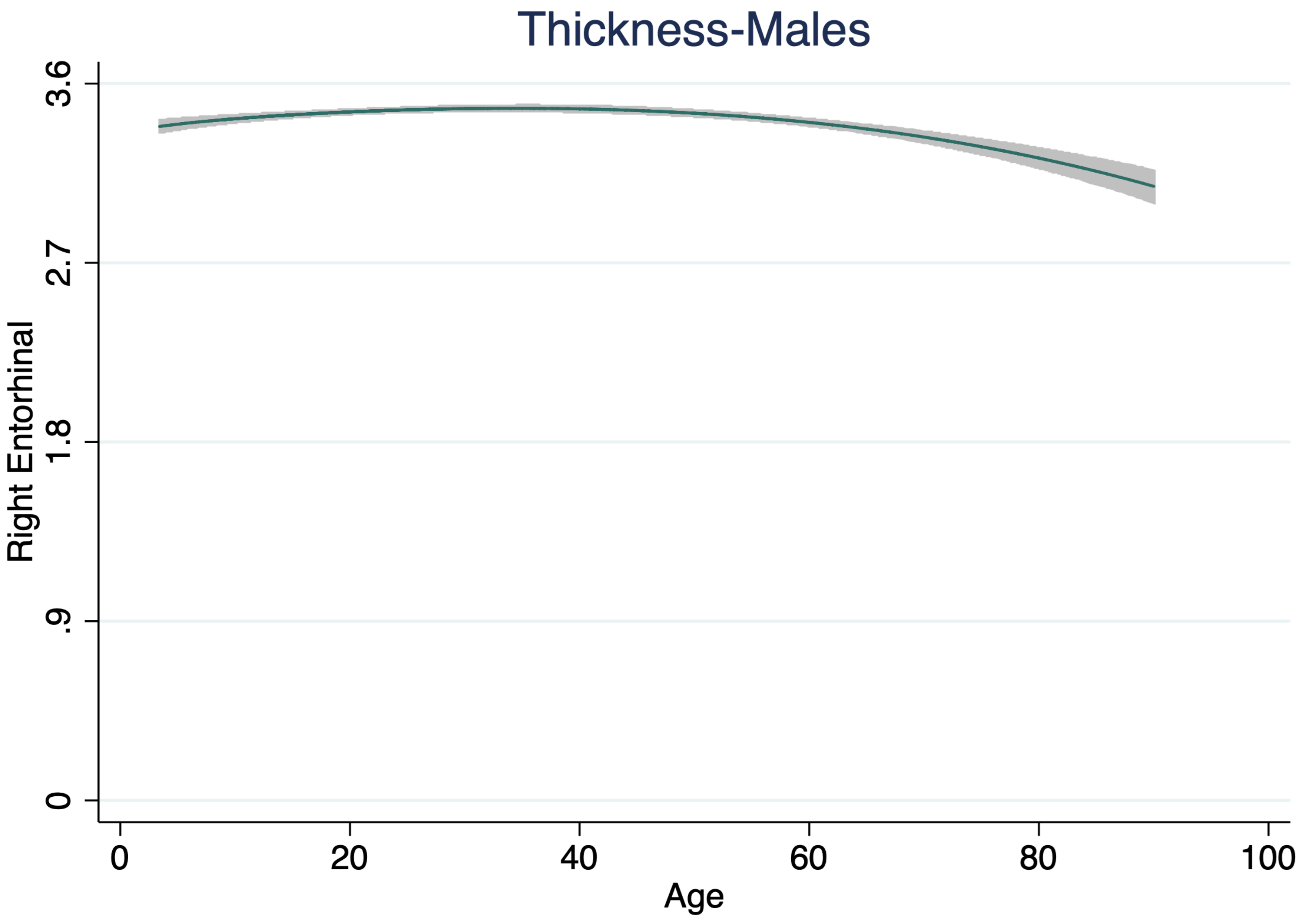

Thickness-Males
Right Entorhinal
3.6
2.7
1.8
.9
0
0
20
40
60
80
100
Age

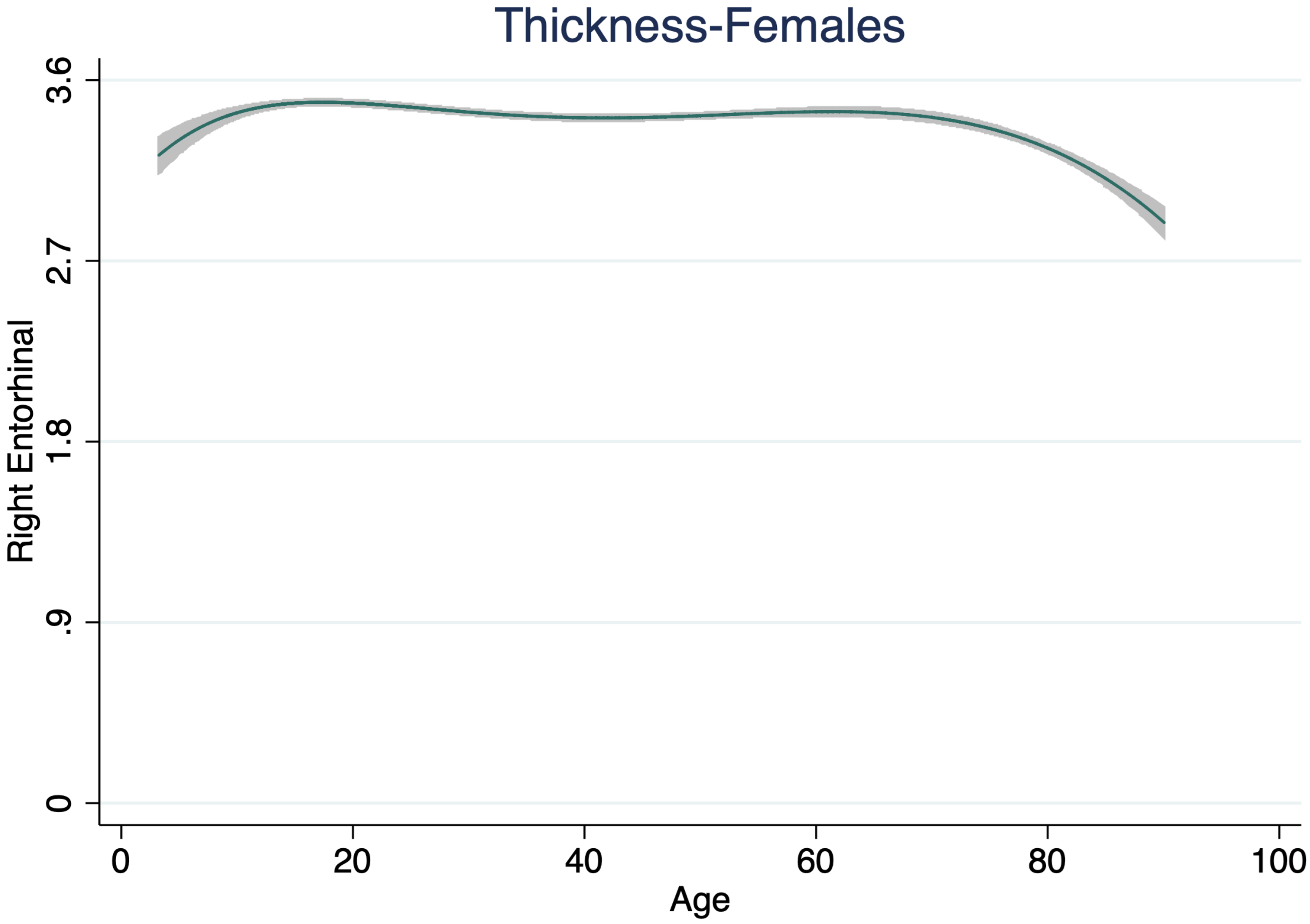

Thickness-Females
Right Entorhinal
3.6
2.7
1.8
.9
0
0
20
40
60
80
100
Age

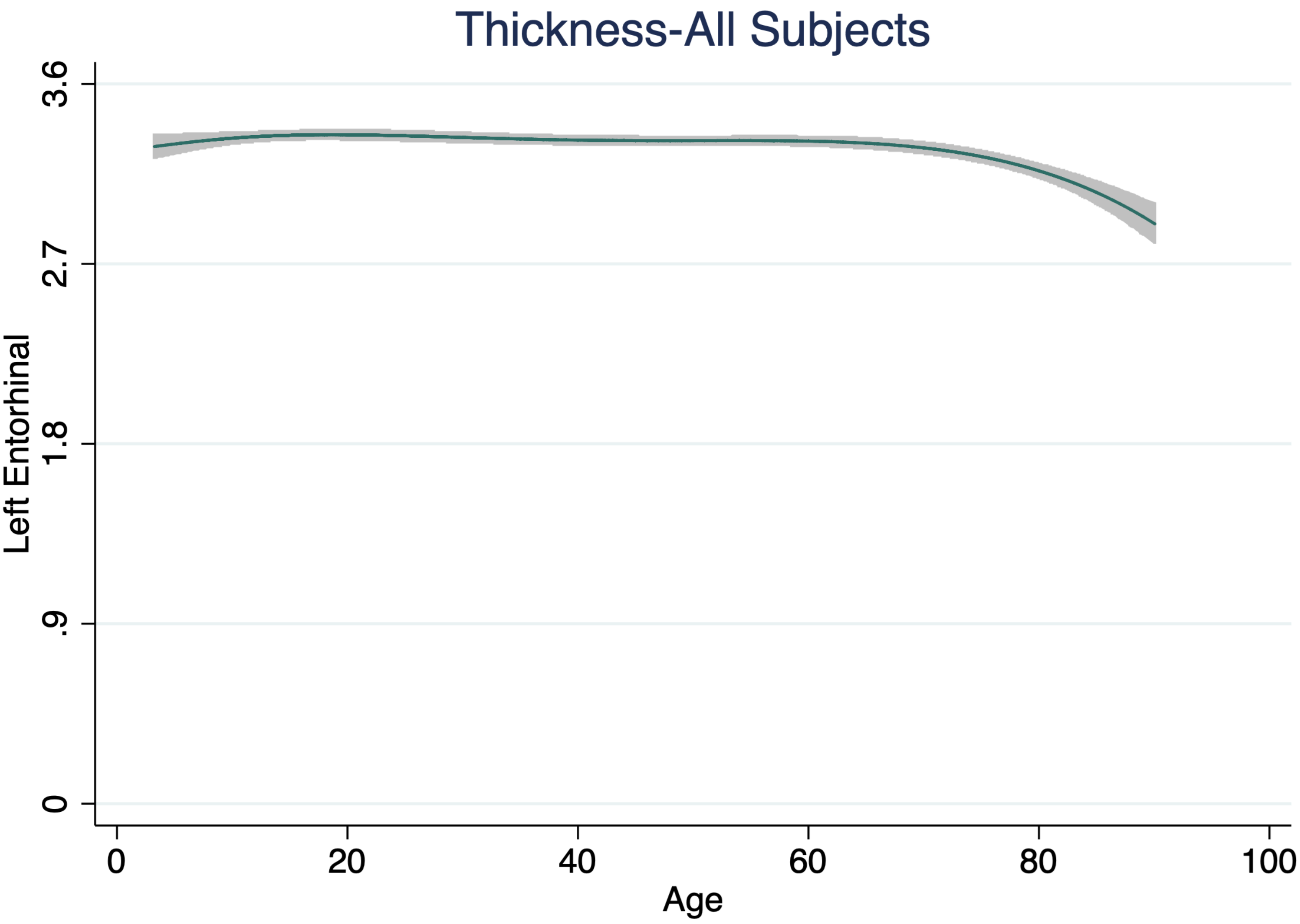

Thickness-All Subjects
Left Entorhinal
3.6
2.7
1.8
.9
0
0
20
40
60
80
100
Age

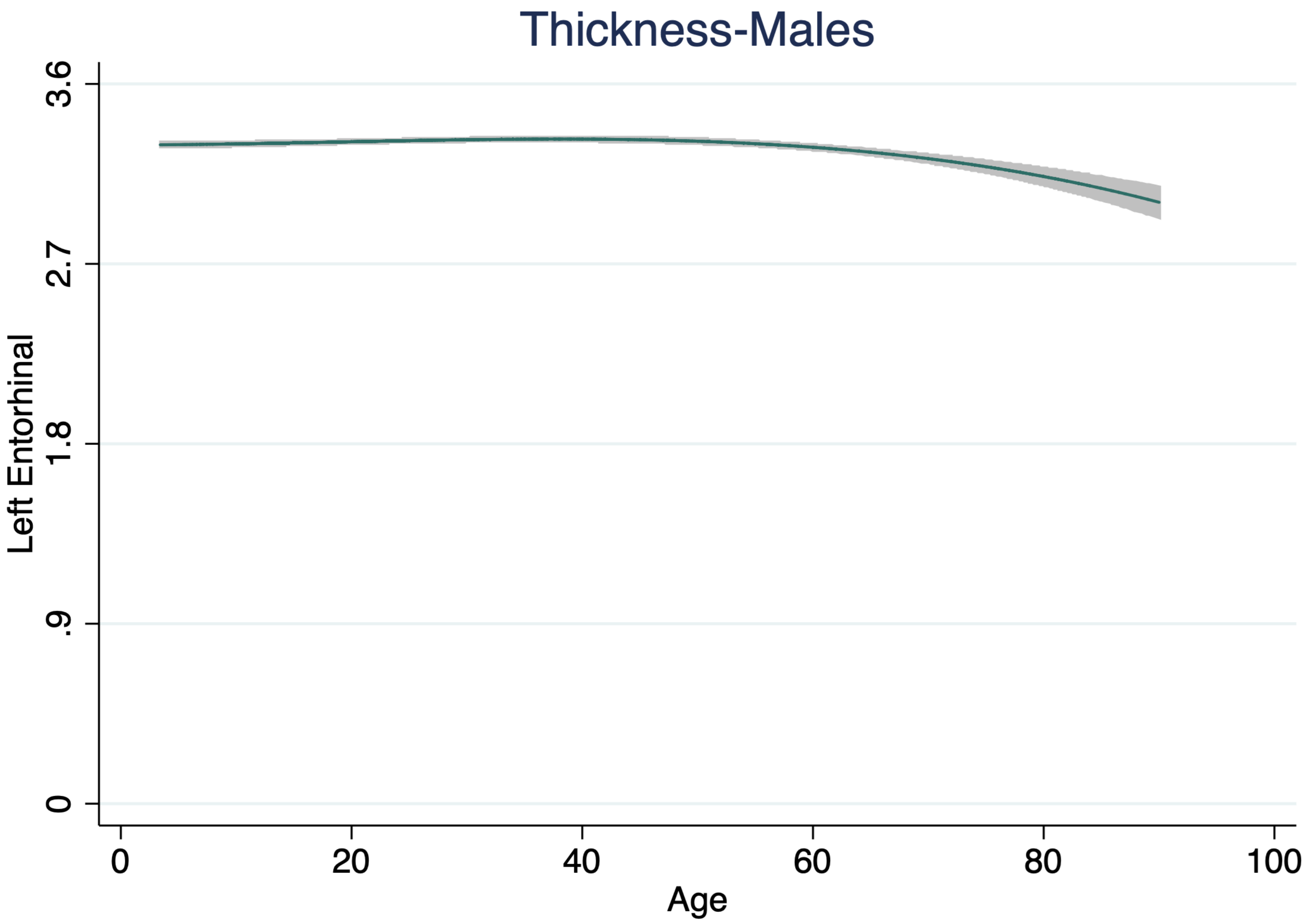
Thickness-Males
3.6
2.7
1.8
.9
0
Left Entorhinal
0
20
40
60
80
100
Age

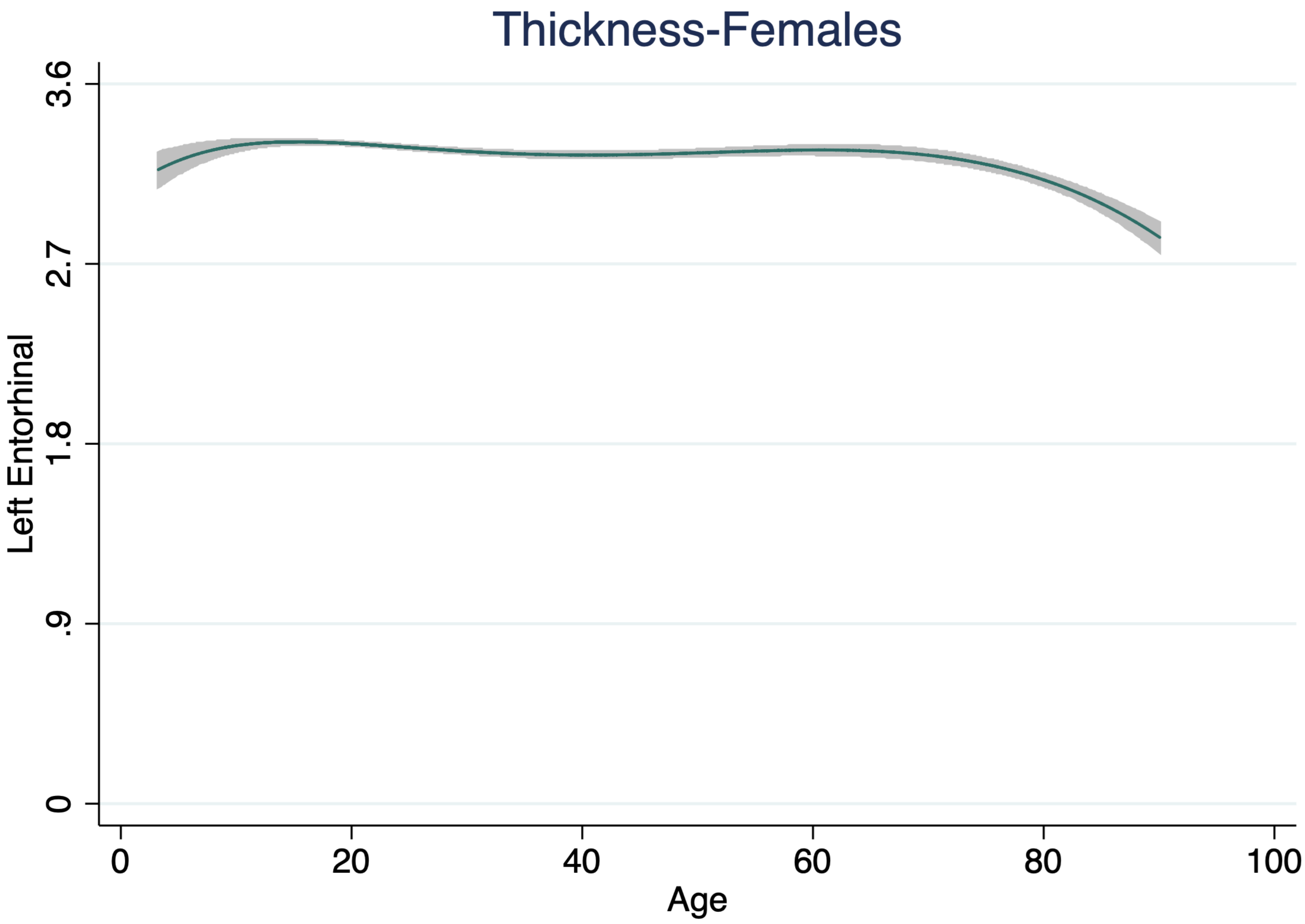

Thickness-Females
Left Entorhinal
3.6
2.7
1.8
.9
0
0
20
40
60
80
100
Age

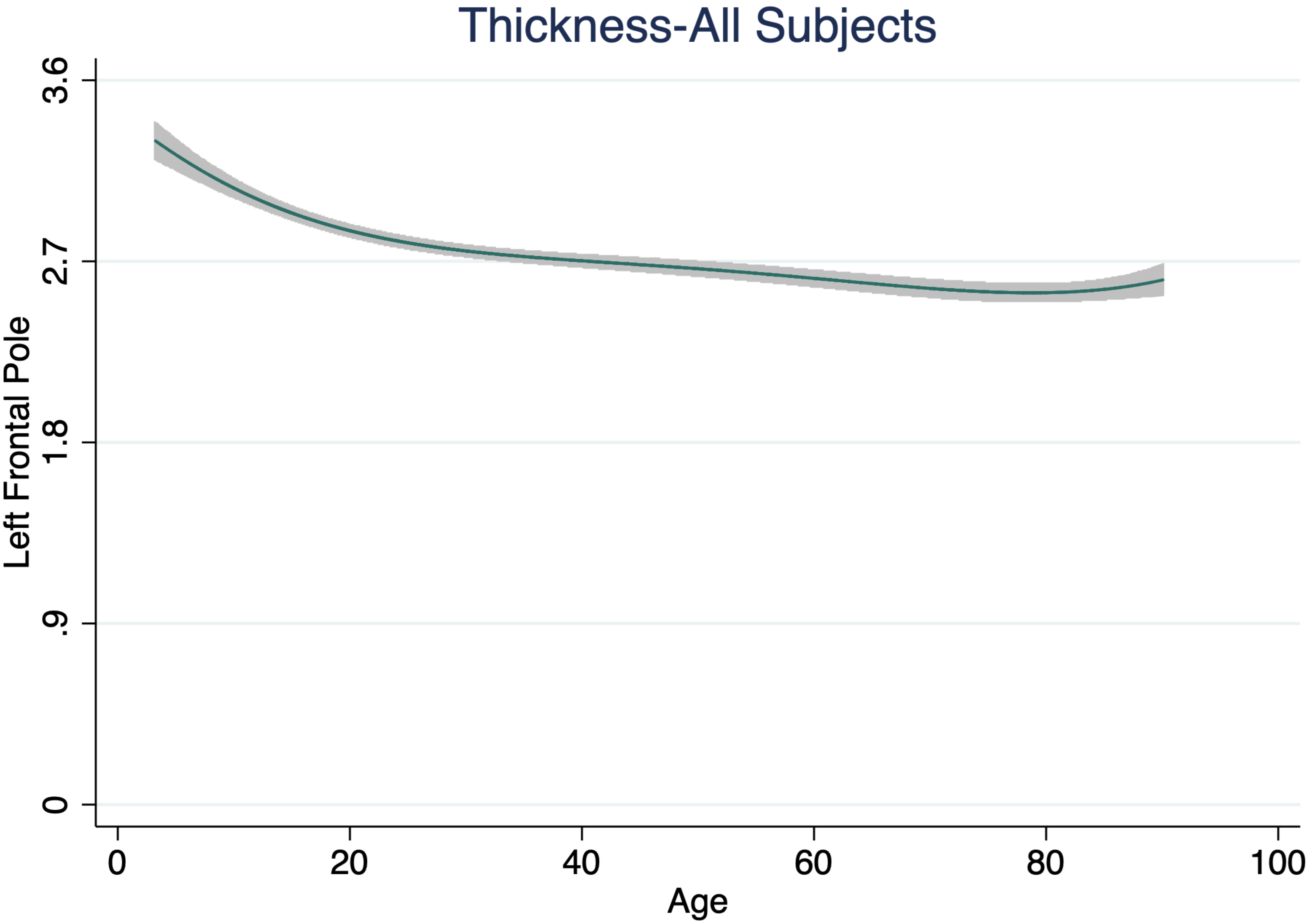

Thickness-All Subjects
Left Frontal Pole
3.6
2.7
1.8
.9
0
0
20
40
60
80
100
Age

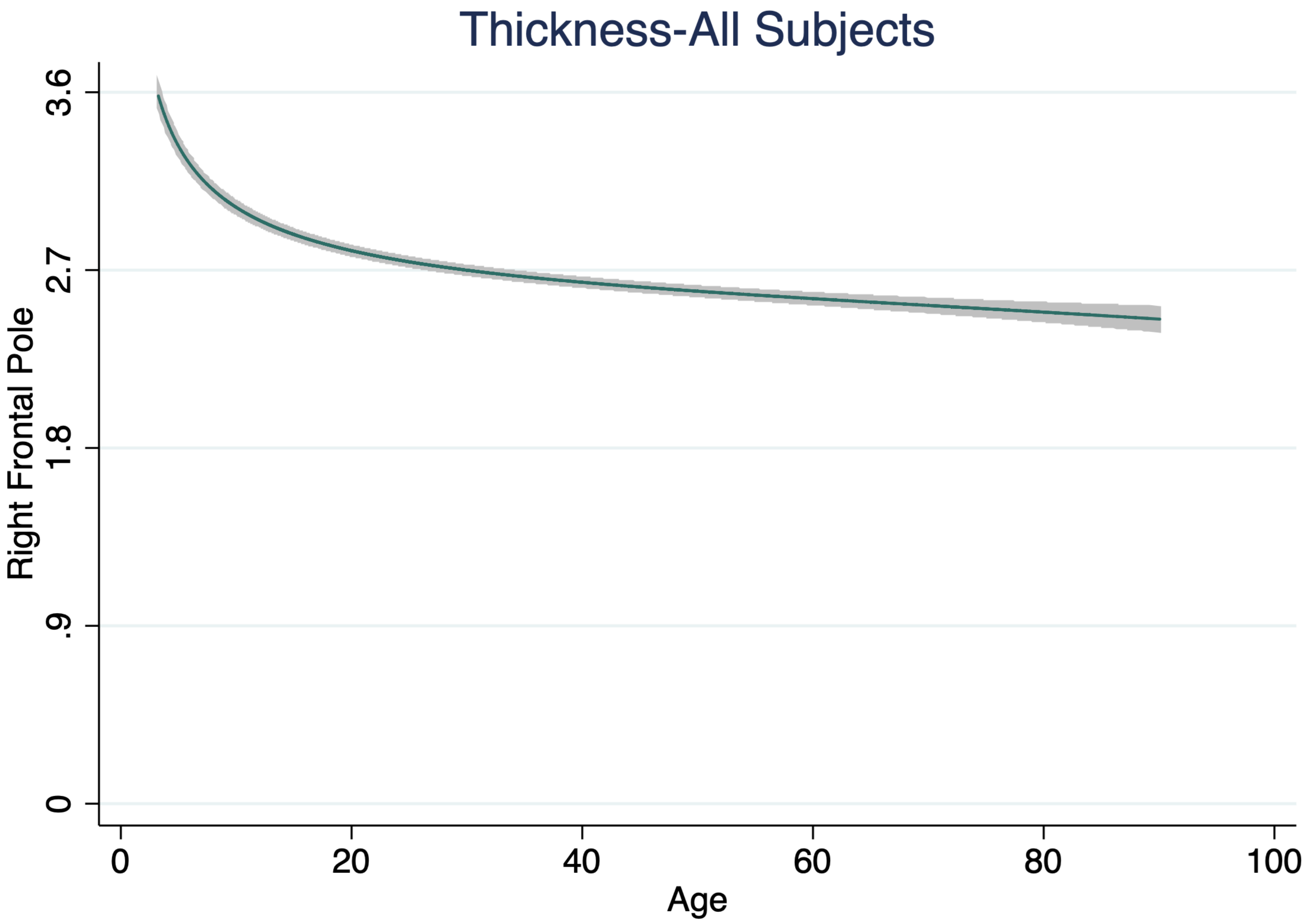
Thickness-All Subjects
Right Frontal Pole
3.6
2.7
1.8
.9
0
0
20
40
60
80
100
Age

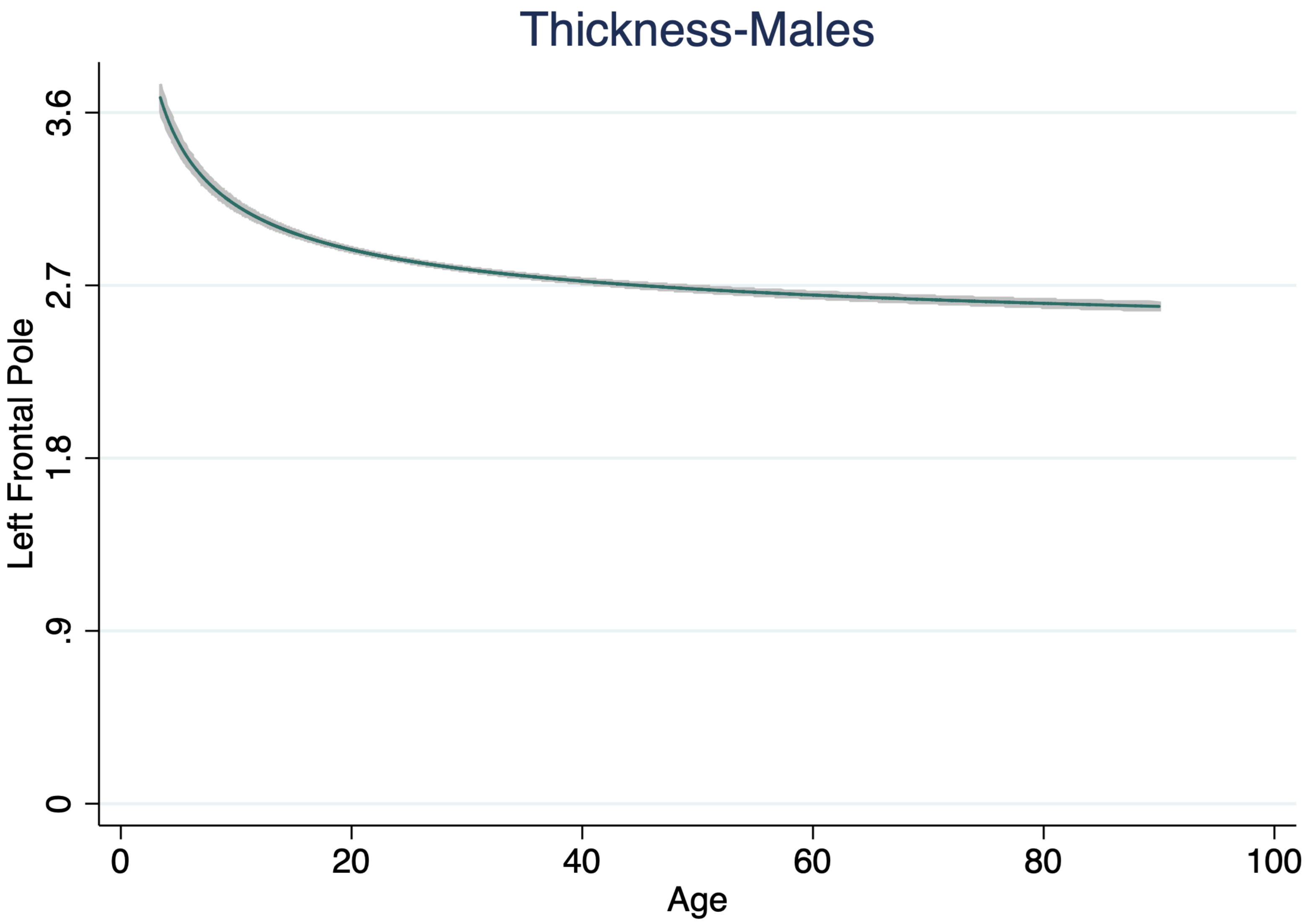

Thickness-Males
Left Frontal Pole
3.6
2.7
1.8
.9
0
0
20
40
60
80
100
Age

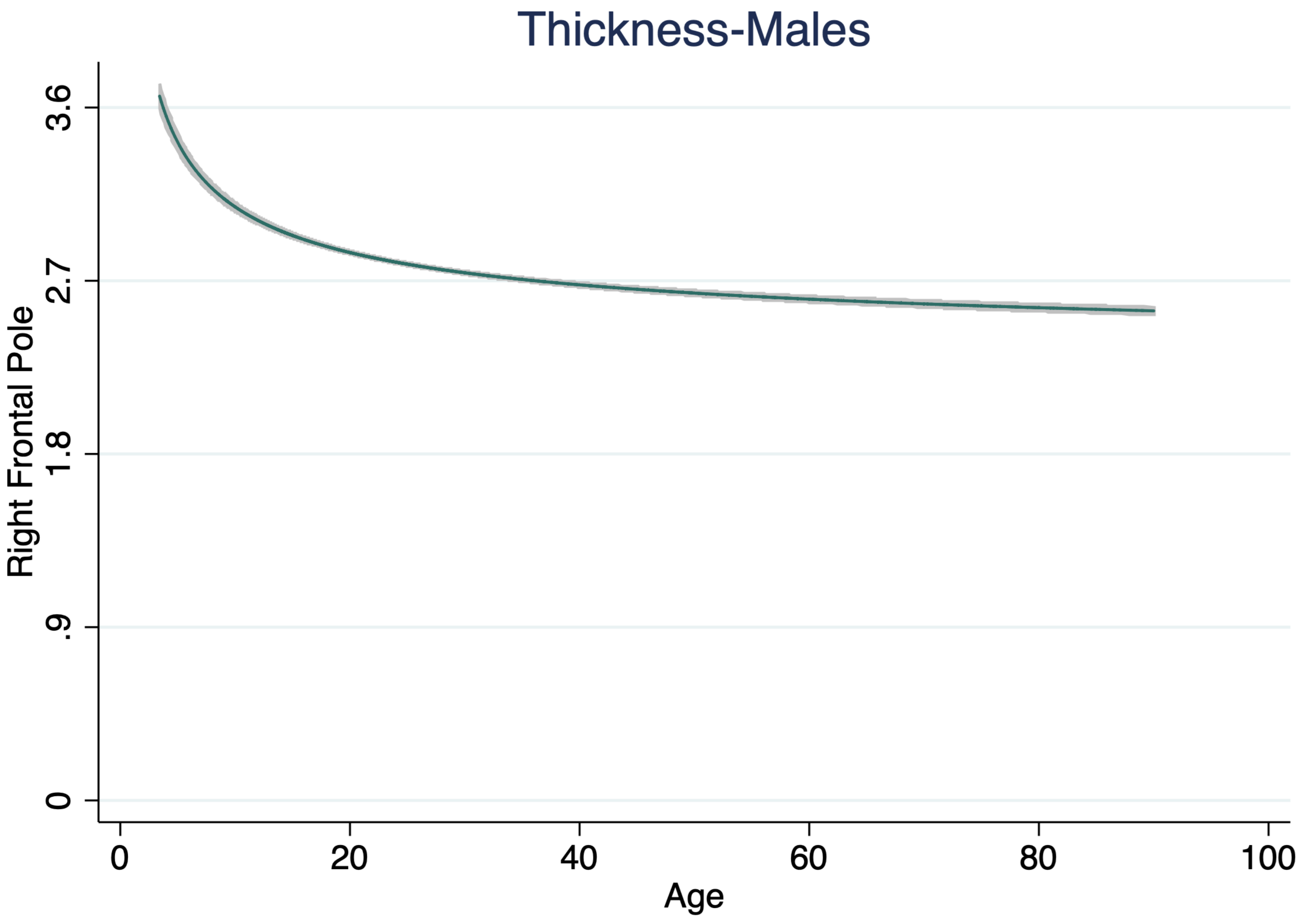
Thickness-Males
Right Frontal Pole
3.6
2.7
1.8
.9
0
0
20
40
60
80
100
Age

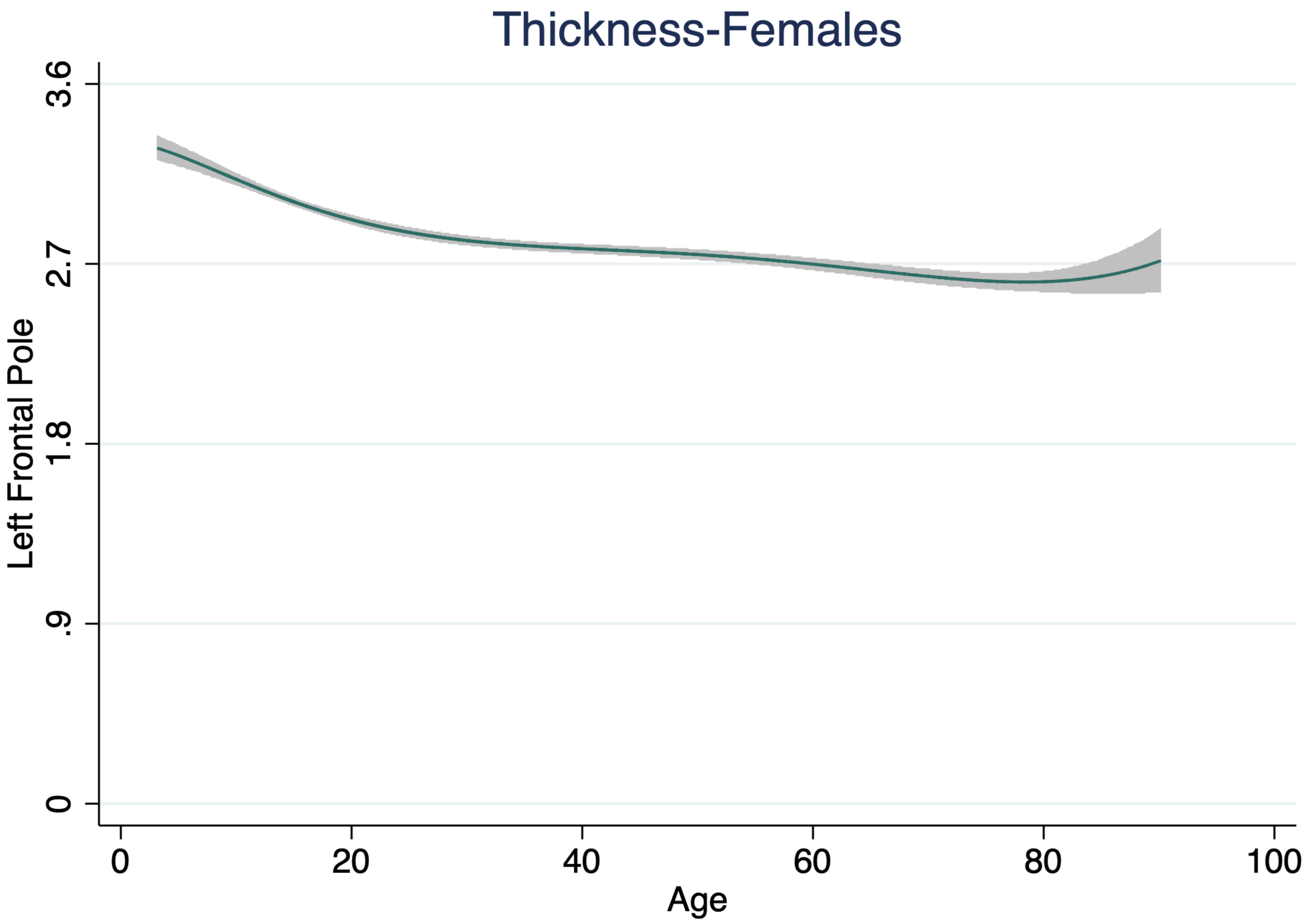

Thickness-Females
Left Frontal Pole
3.6
2.7
1.8
.9
0
0
20
40
60
80
100
Age

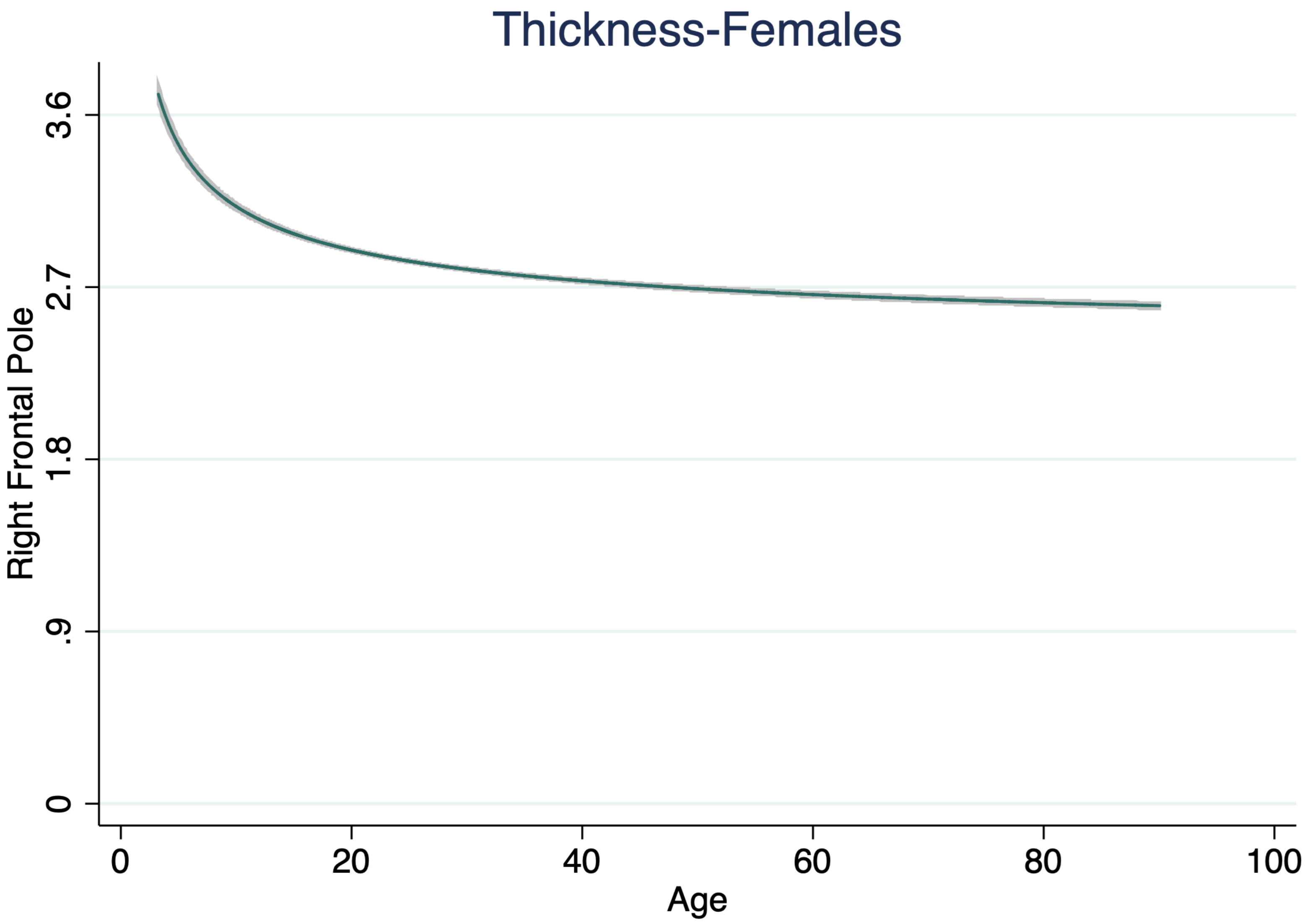

Thickness-Females
Right Frontal Pole
3.6
2.7
1.8
.9
0
0
20
40
60
80
100
Age

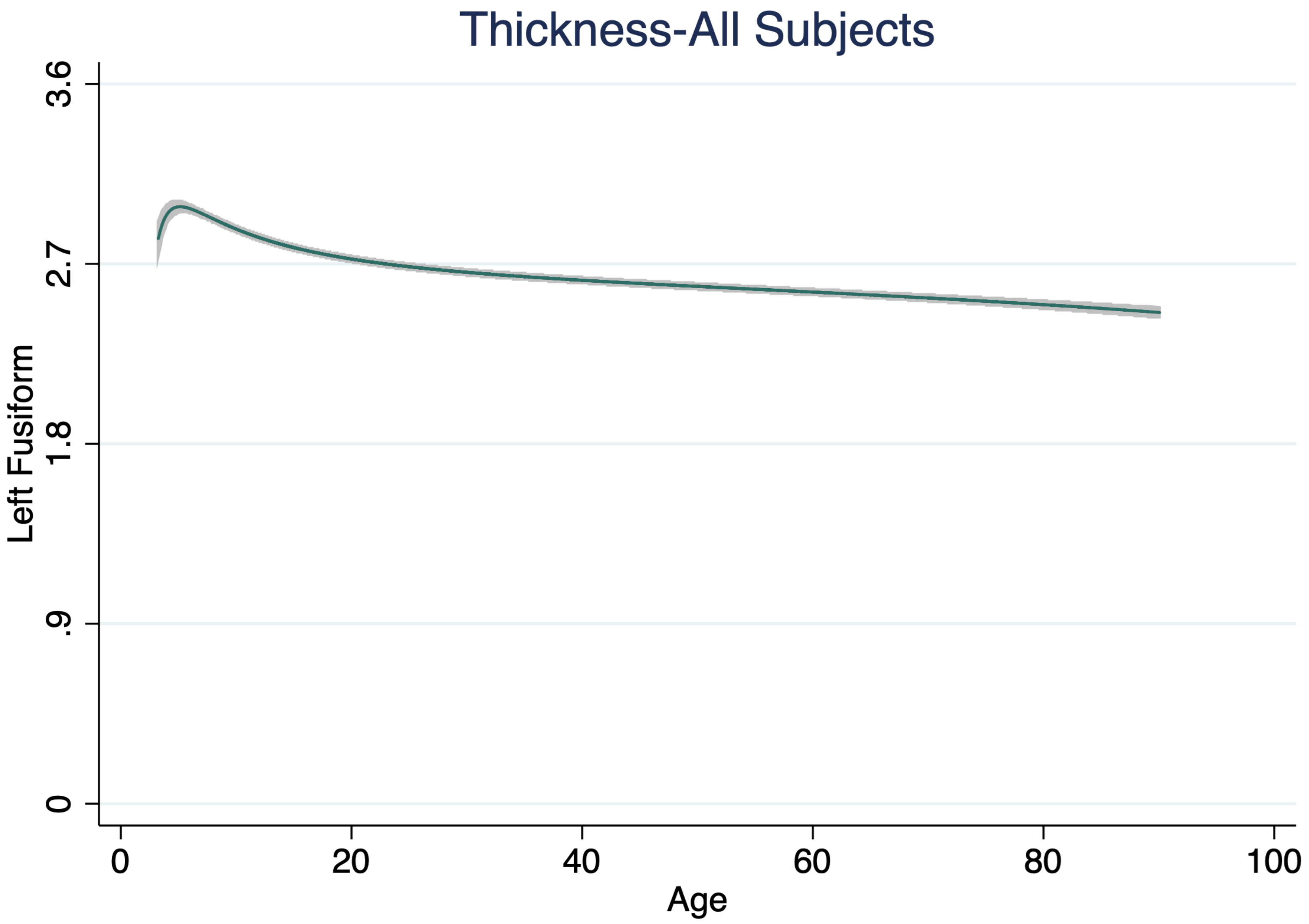

Thickness-All Subjects
Left Fusiform
3.6
2.7
1.8
.9
0
0
20
40
60
80
100
Age

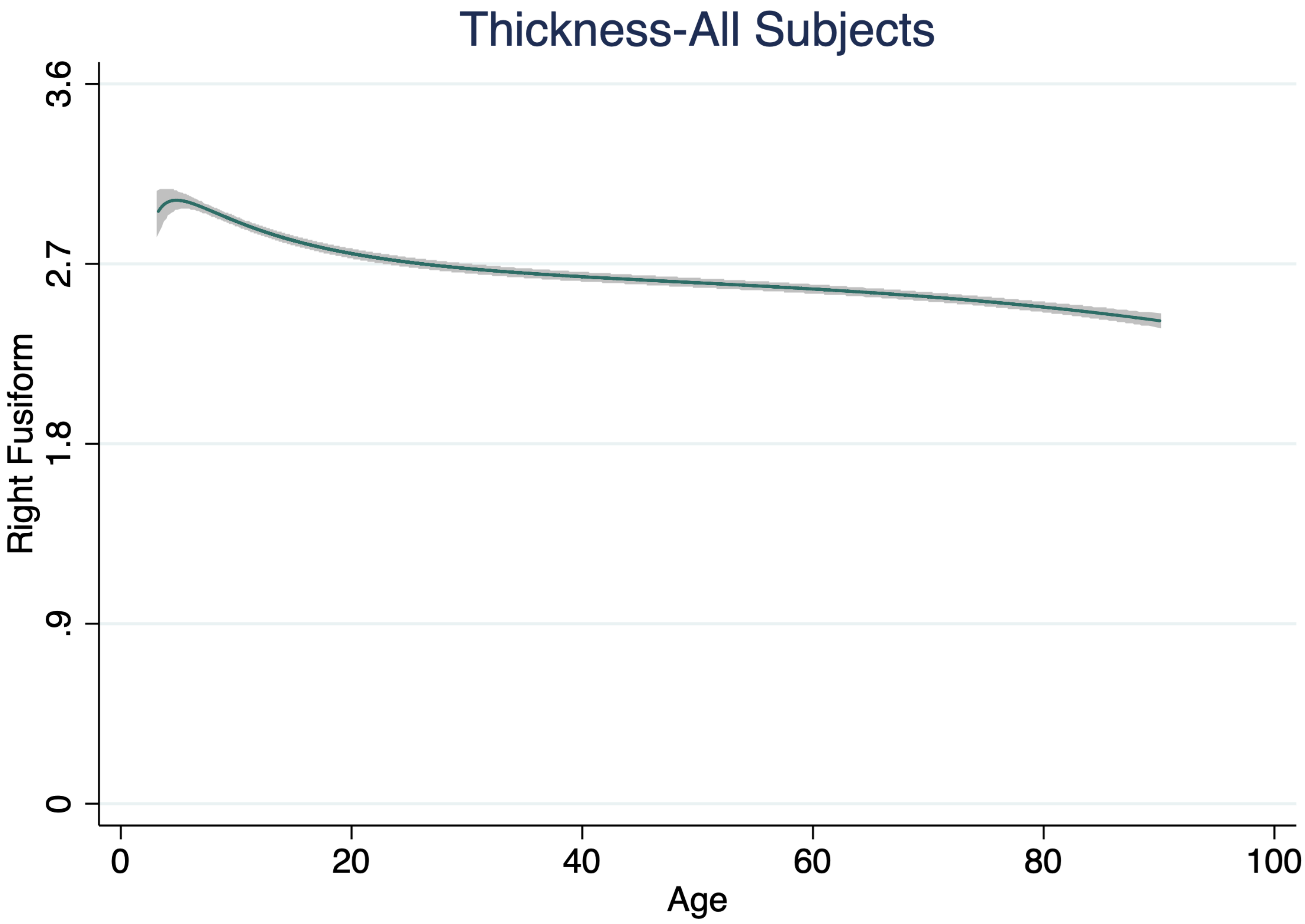
Thickness-All Subjects
Right Fusiform
3.6
2.7
1.8
.9
0
0
20
40
60
80
100
Age

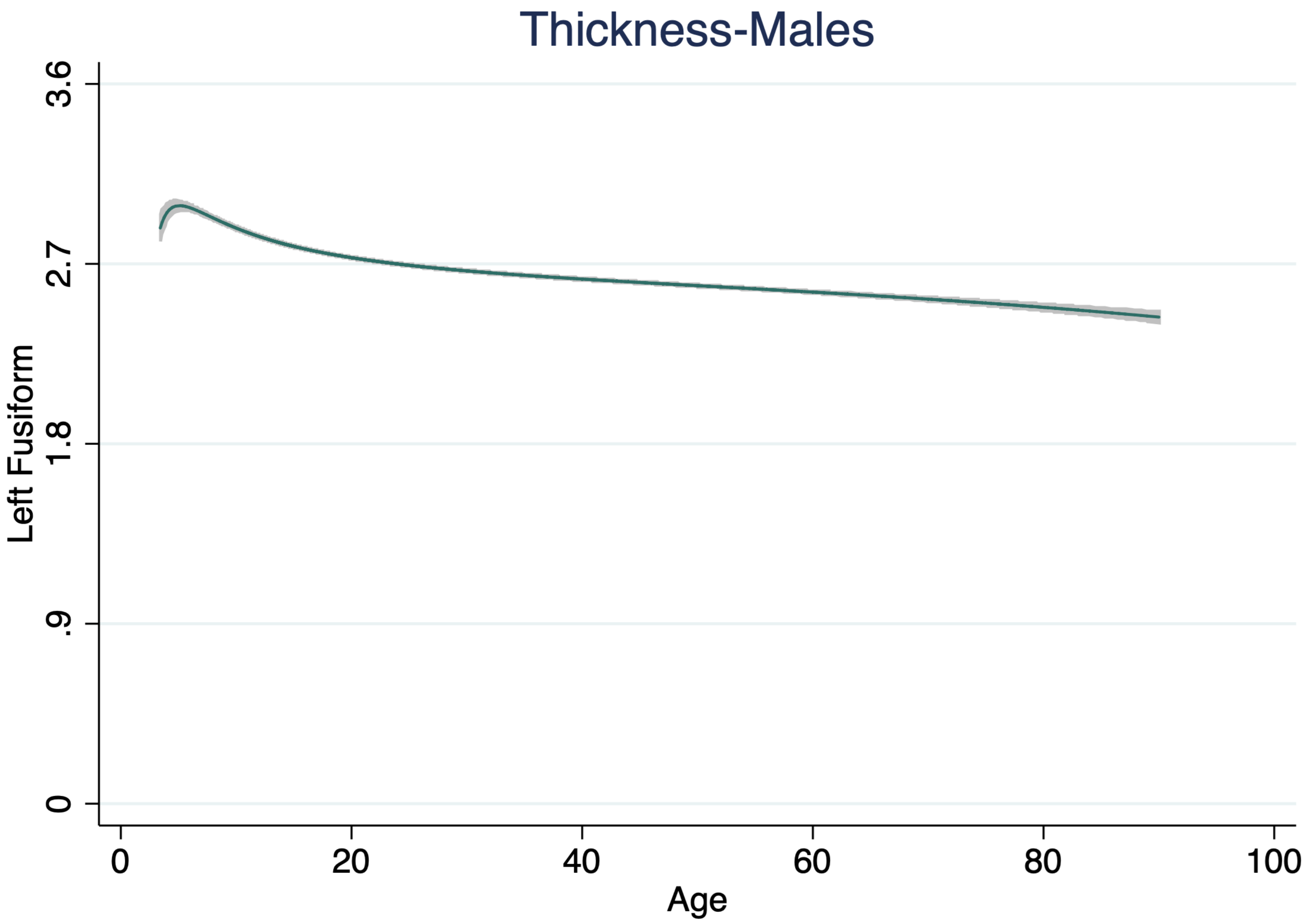

Thickness-Males
Left Fusiform
3.6
2.7
1.8
.9
0
0
20
40
60
80
100
Age

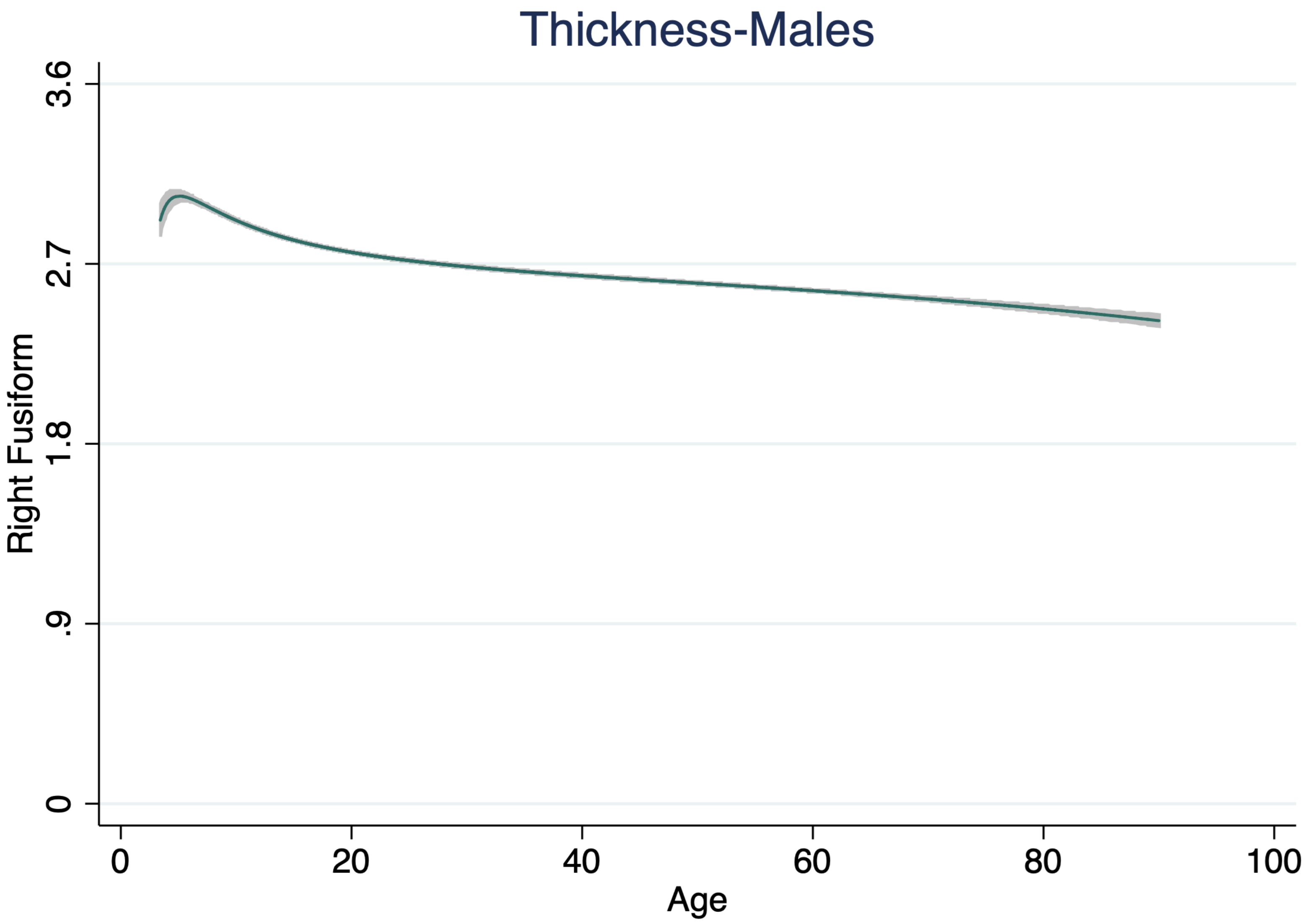

Thickness-Males
Right Fusiform
3.6
2.7
1.8
.9
0
0
20
40
60
80
100
Age

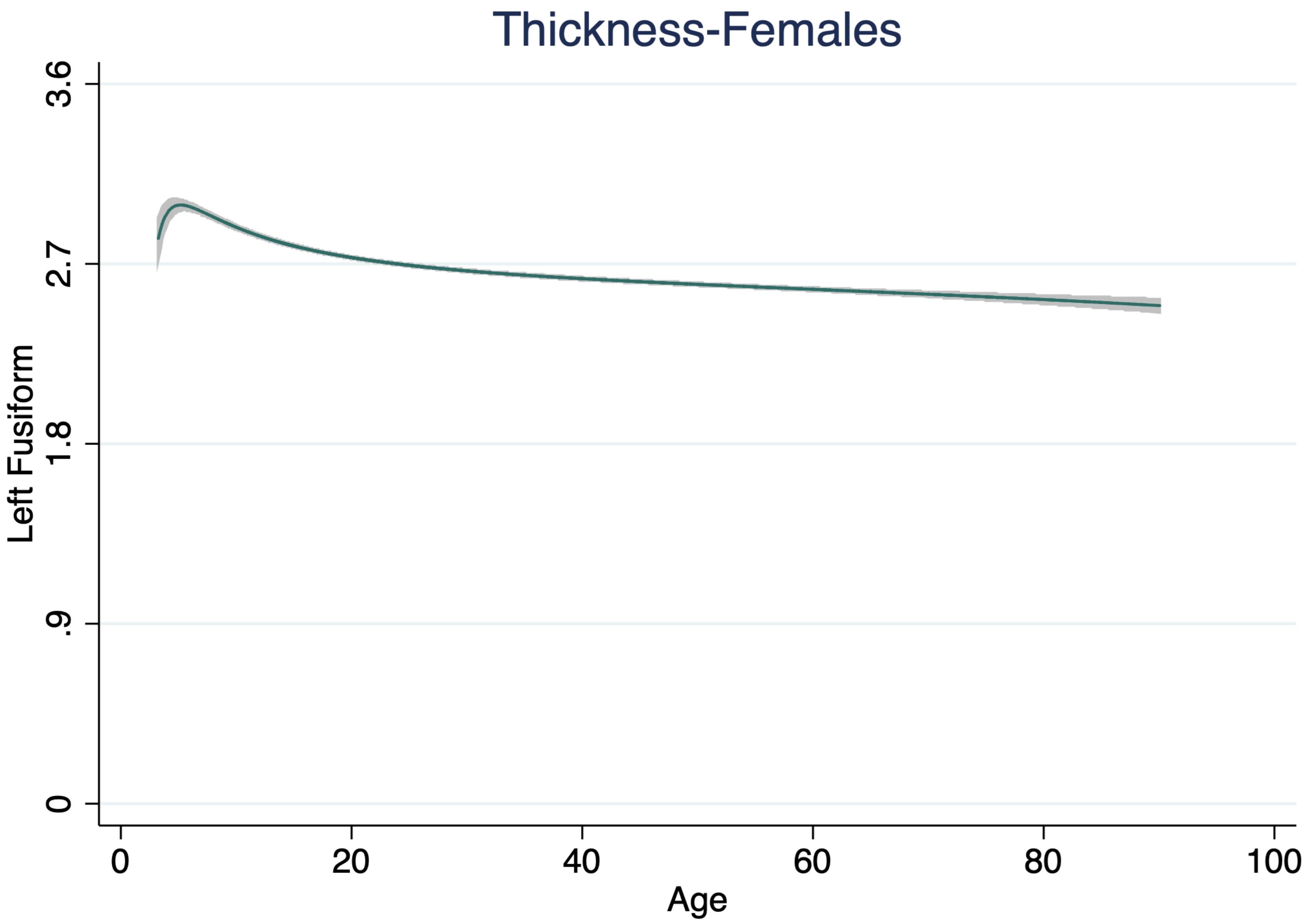
Thickness-Females
Left Fusiform
3.6
2.7
1.8
.9
0
0
20
40
60
80
100
Age

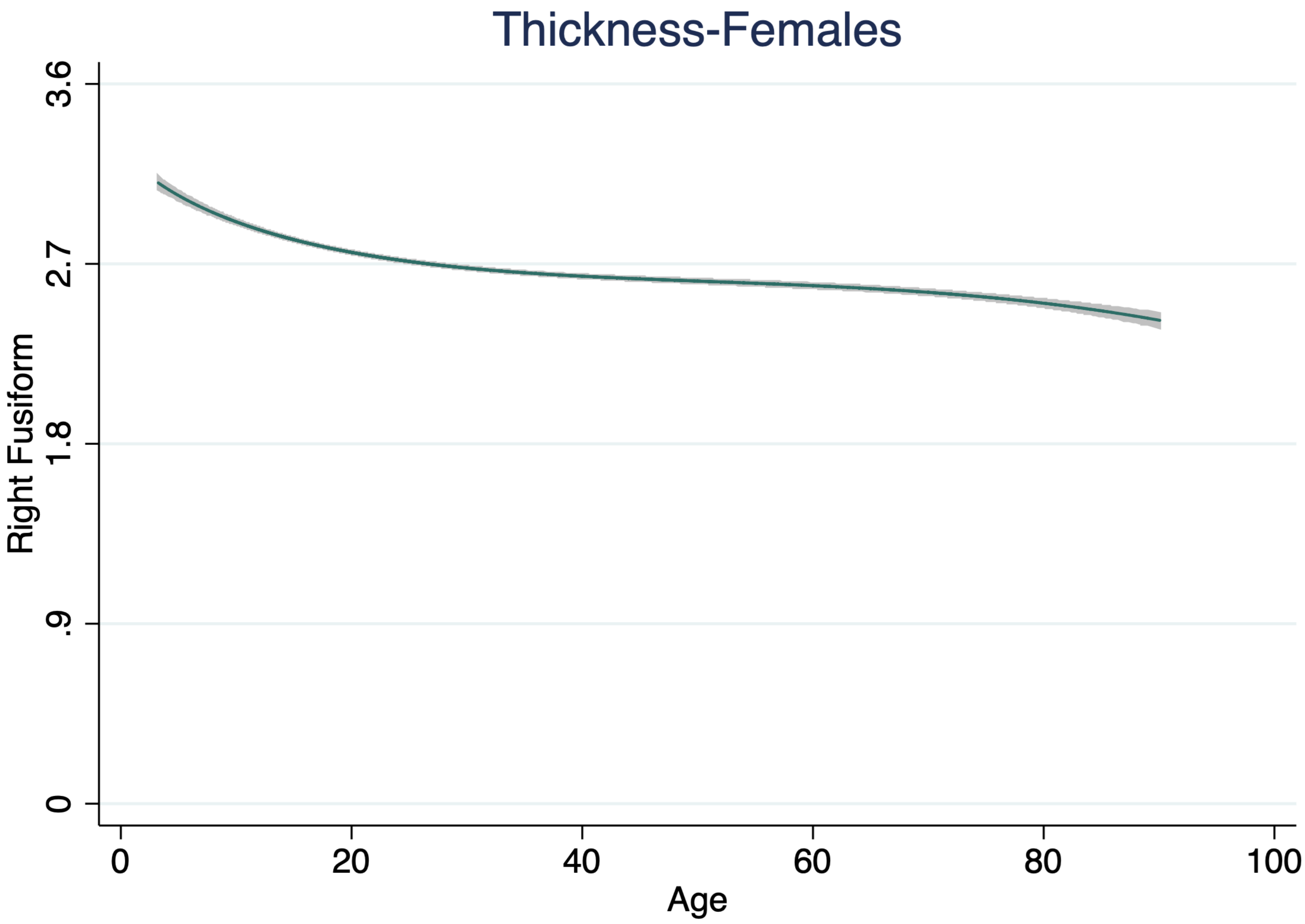
Thickness-Females
Right Fusiform
3.6
2.7
1.8
.9
0
0
20
40
60
80
100
Age

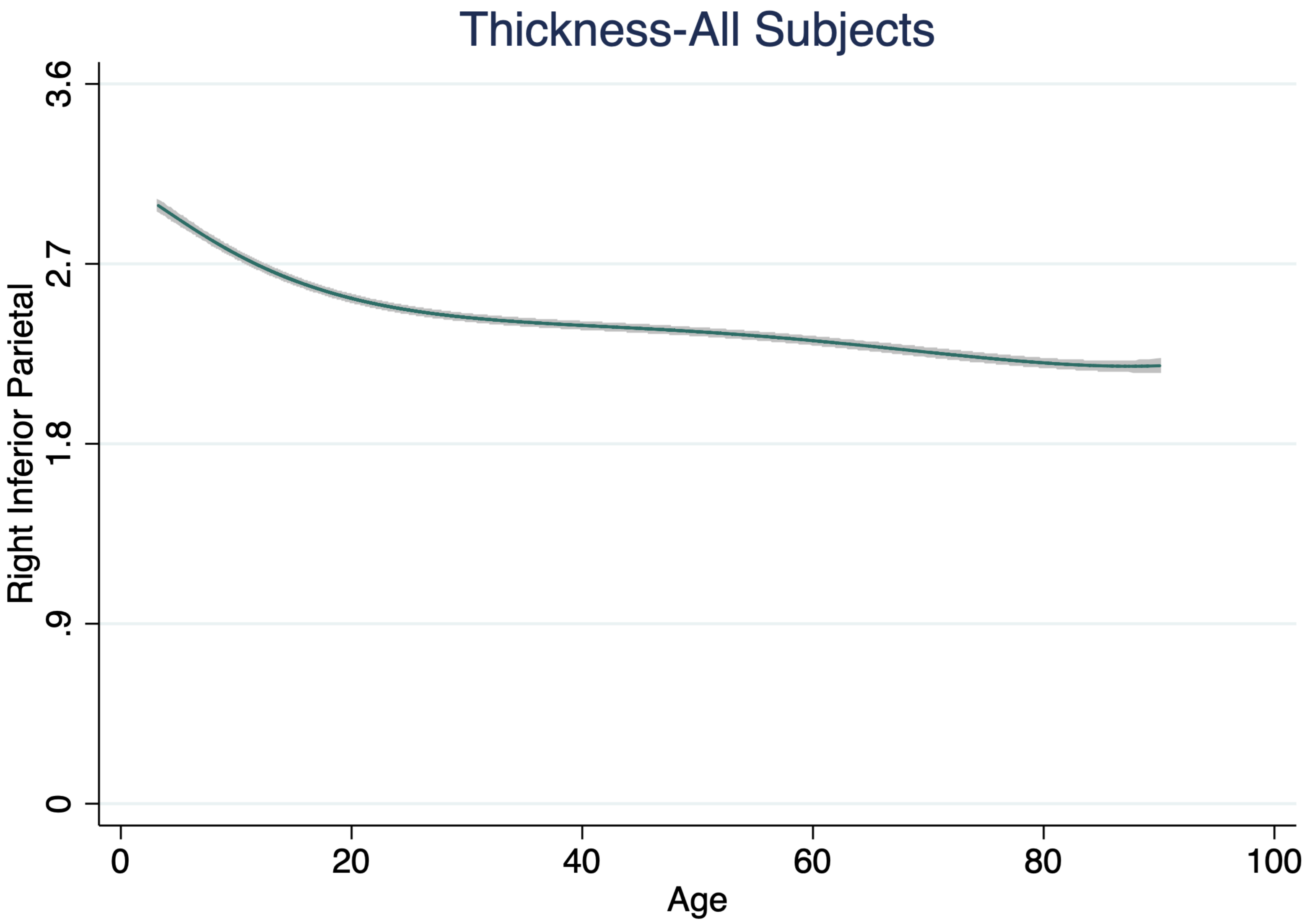

Thickness-All Subjects
Right Inferior Parietal
3.6
2.7
1.8
.9
0
0
20
40
60
80
100
Age

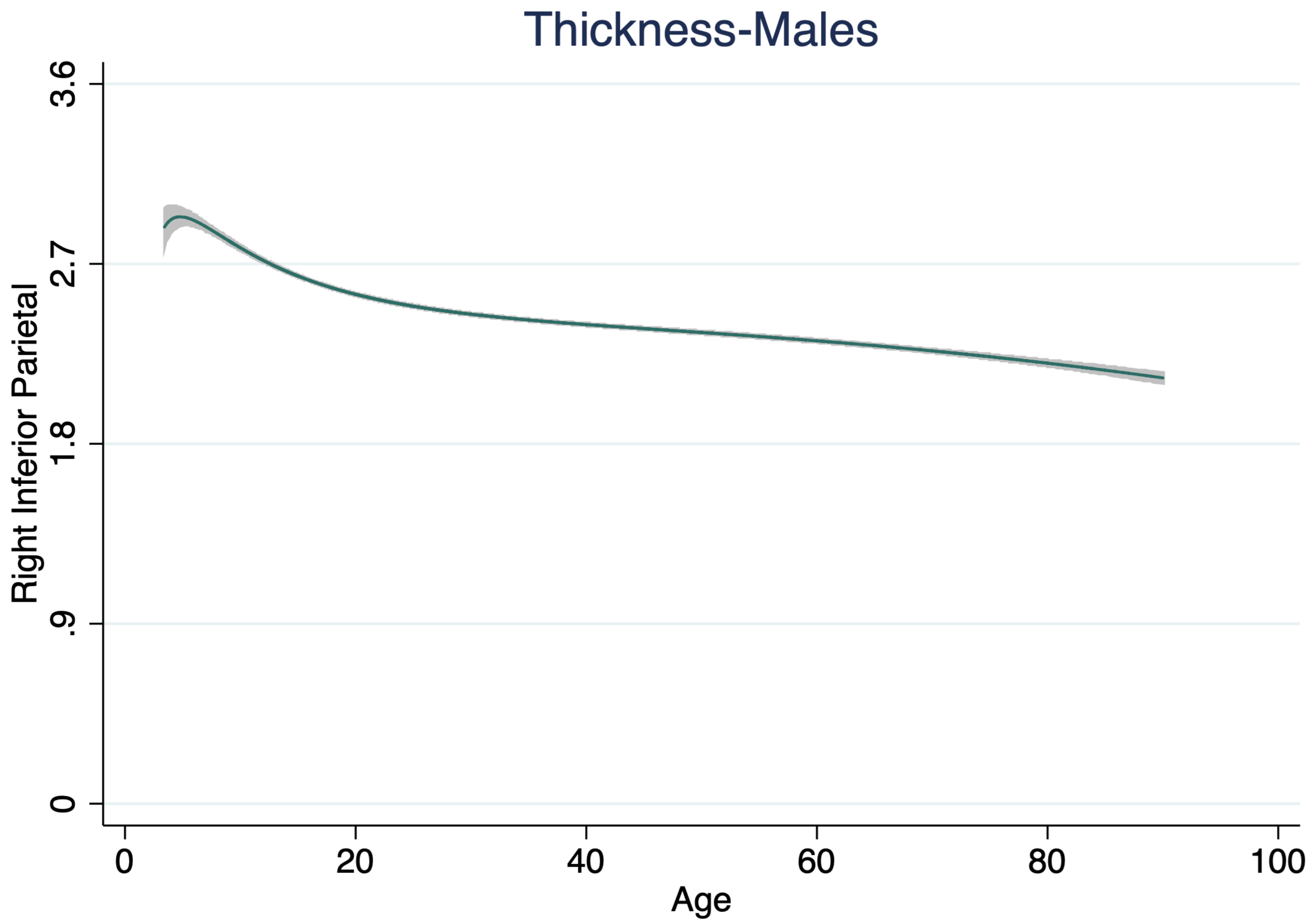
Thickness-Males
Right Inferior Parietal
3.6
2.7
1.8
.9
0
0
20
40
60
80
100
Age

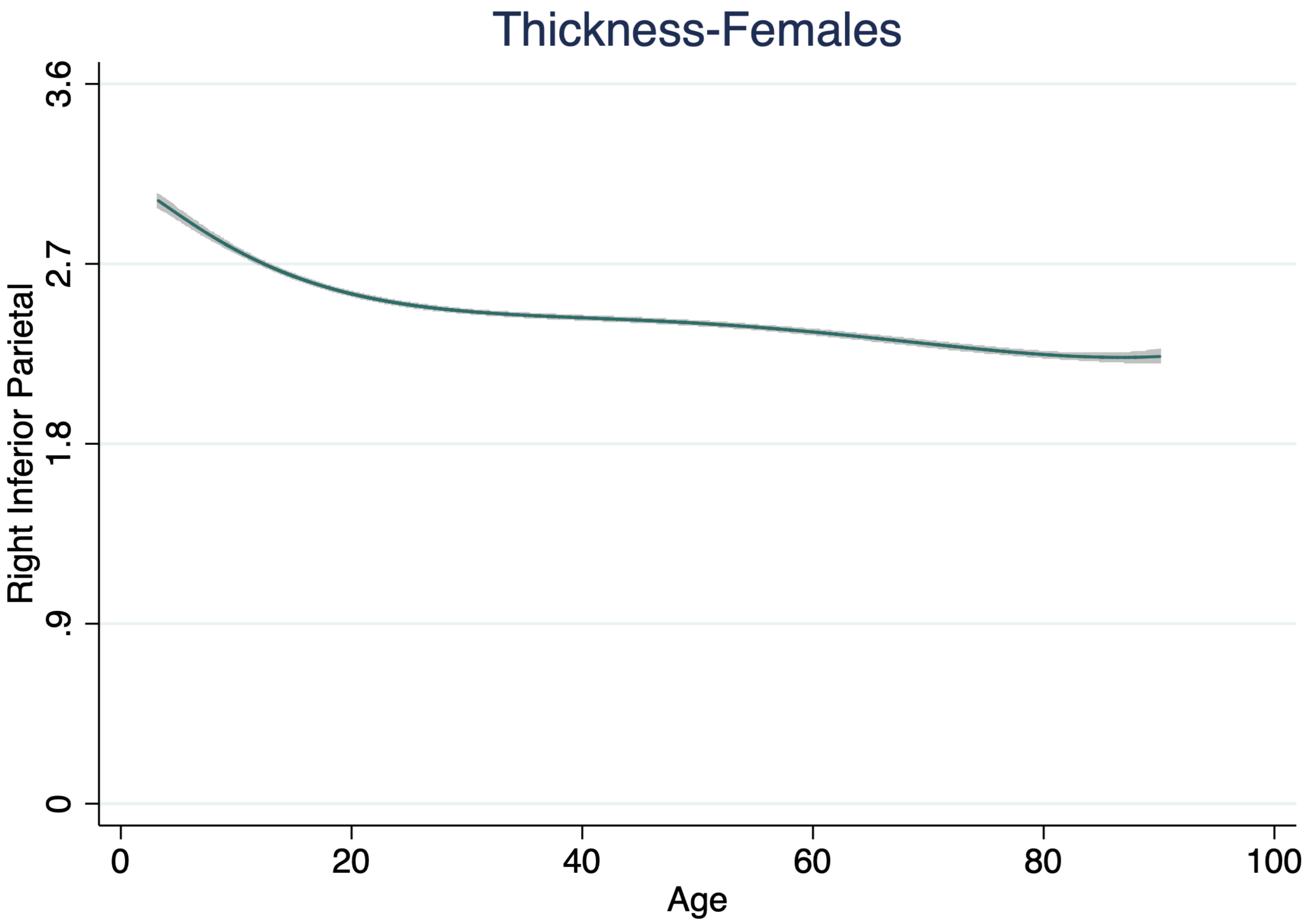

Thickness-Females
Right Inferior Parietal
3.6
2.7
1.8
.9
0
0
20
40
60
80
100
Age

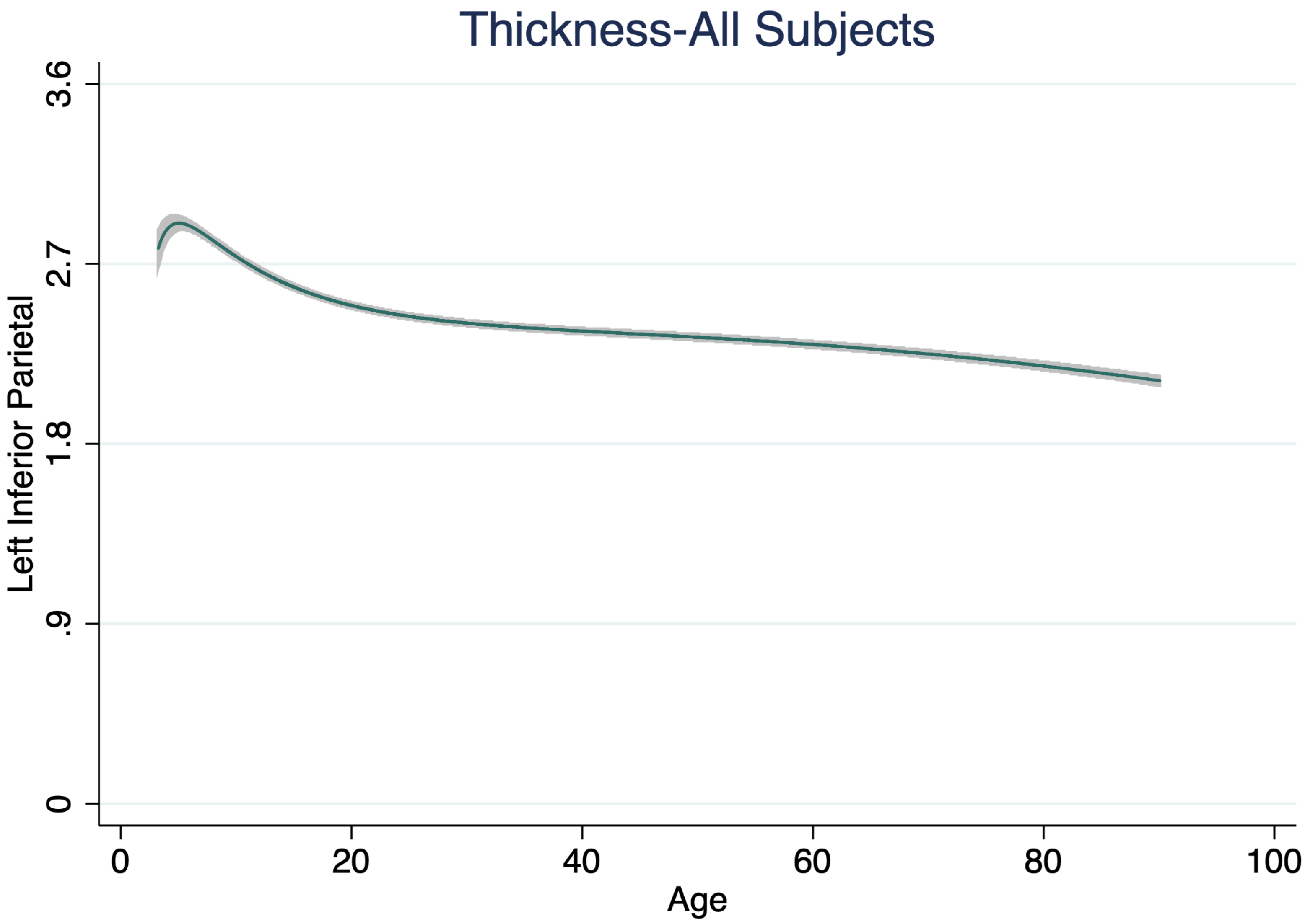
Thickness-All Subjects
Left Inferior Parietal
3.6
2.7
1.8
.9
0
0
20
40
60
80
100
Age

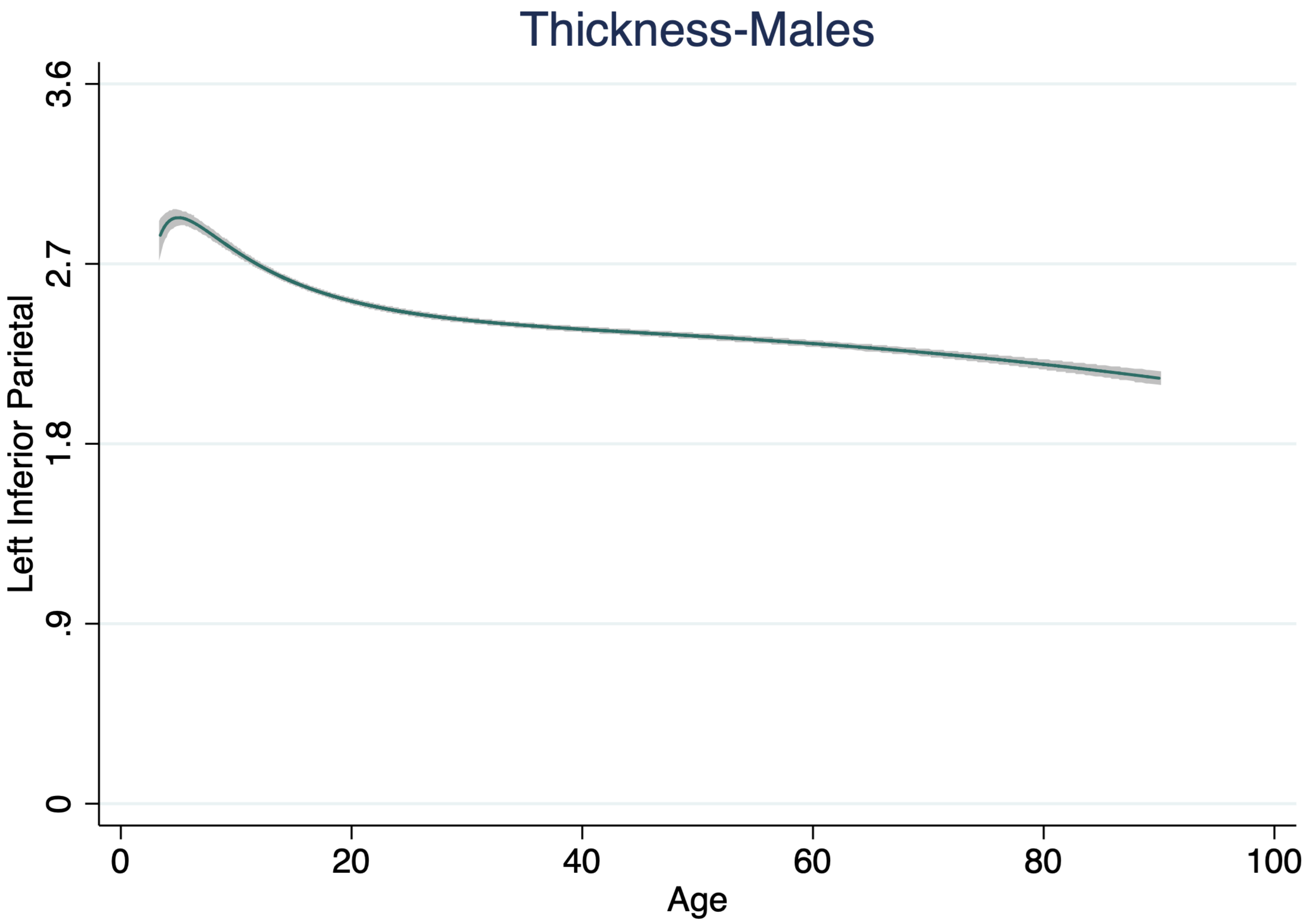

Thickness-Males
Left Inferior Parietal
3.6
2.7
1.8
.9
0
0
20
40
60
80
100
Age

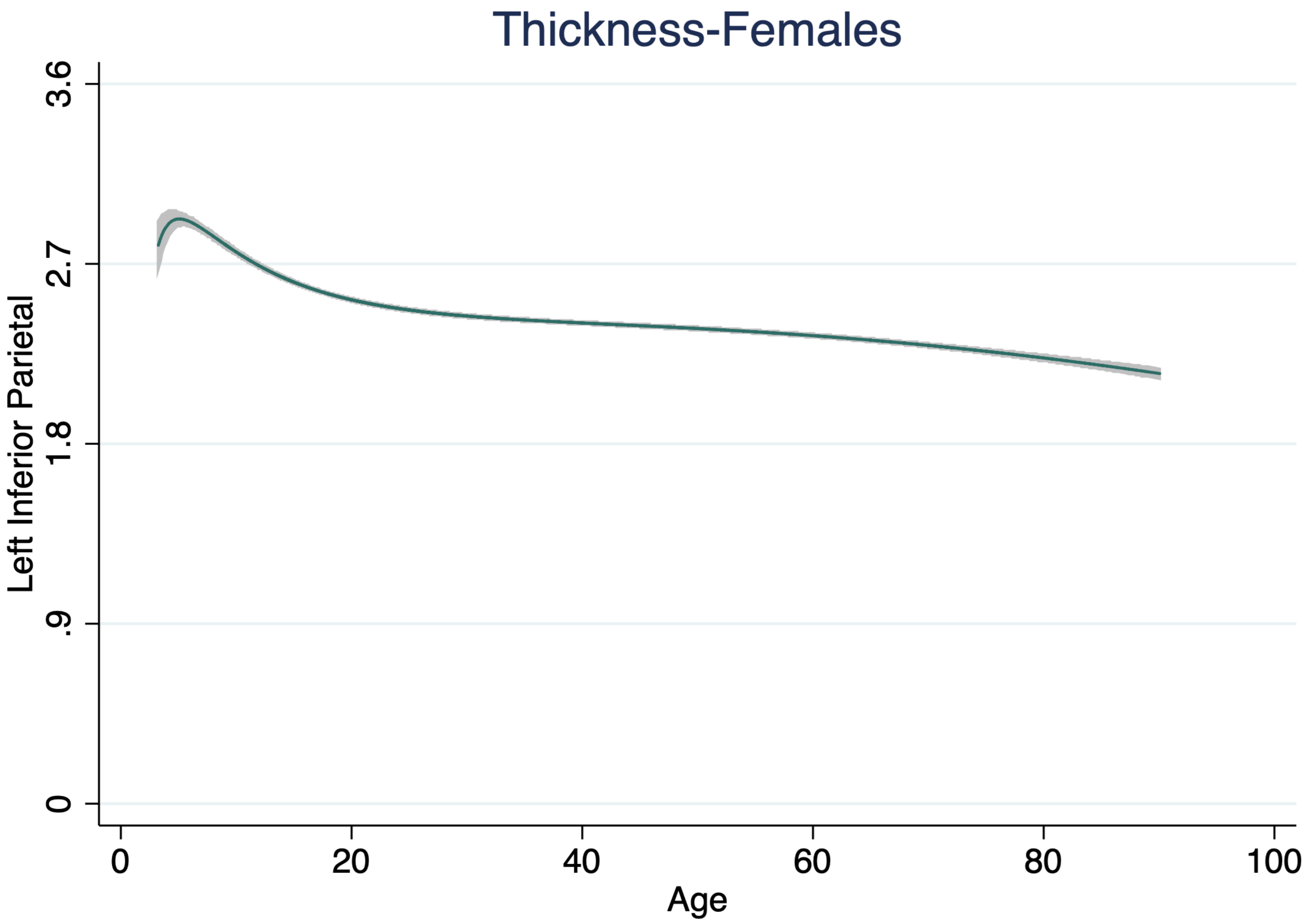

Thickness-Females
Left Inferior Parietal
3.6
2.7
1.8
.9
0
0
20
40
60
80
100
Age

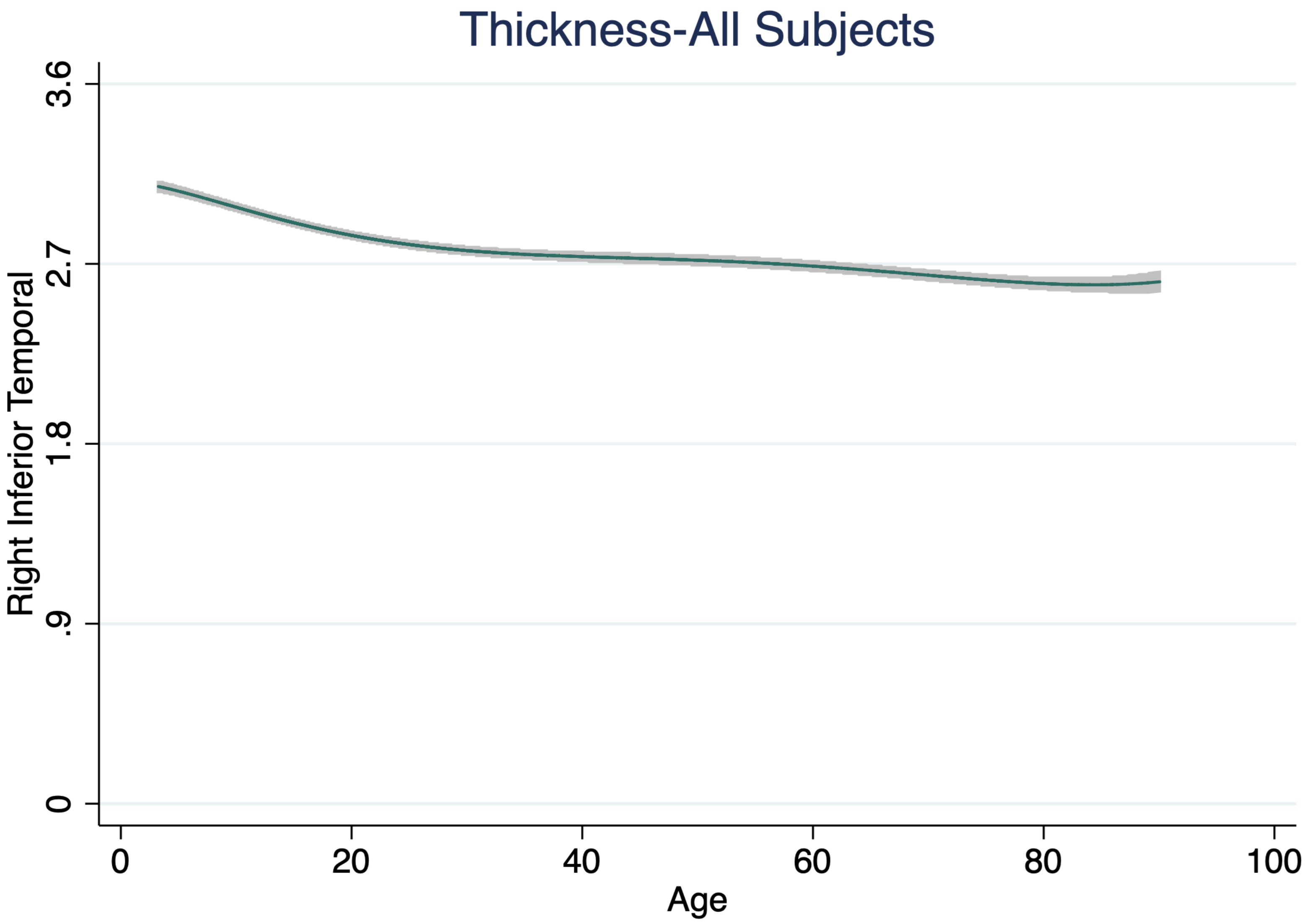

Thickness-All Subjects
Right Inferior Temporal
0
.9
1.8
2.7
3.6
0
20
40
60
80
100
Age

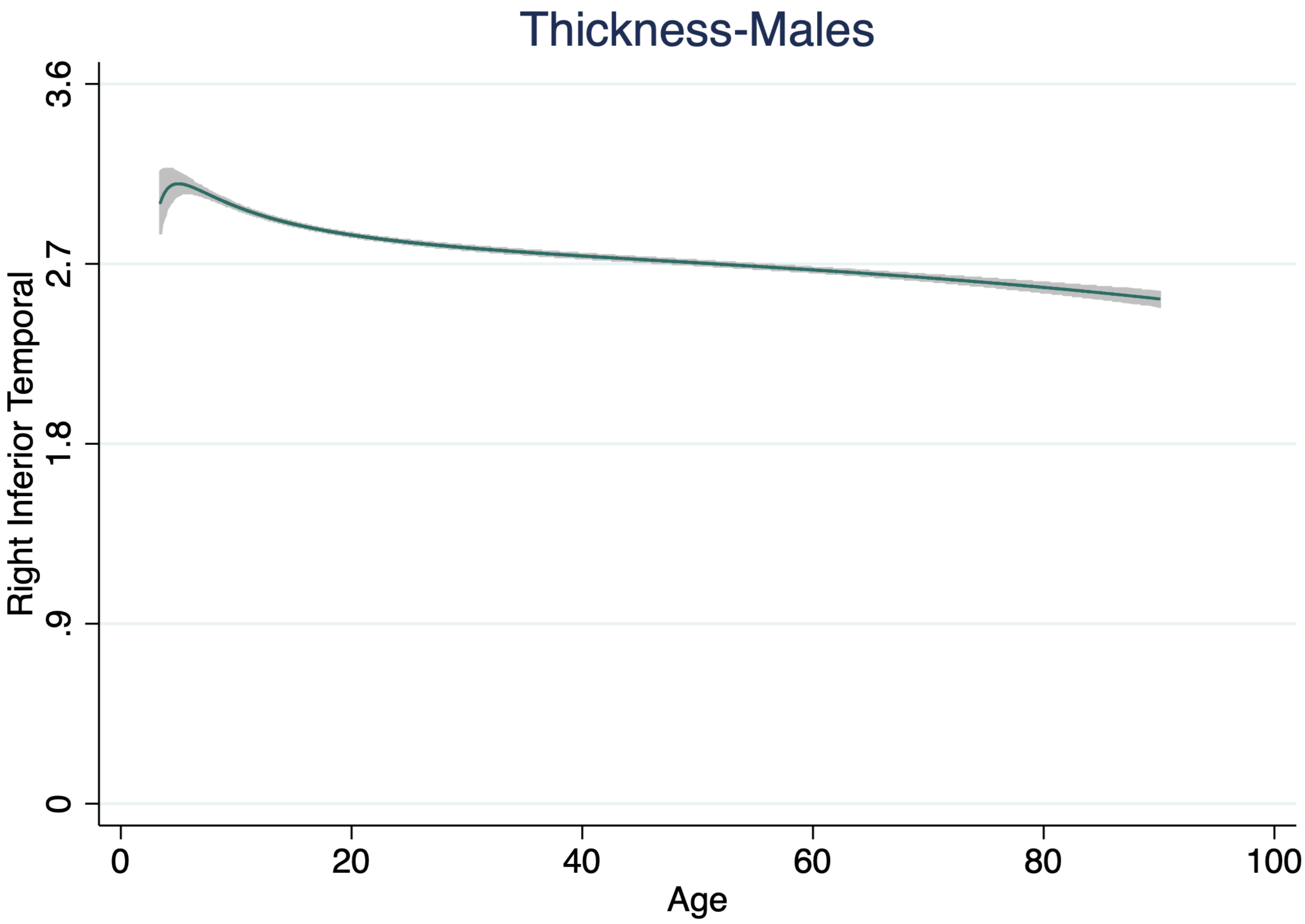

Thickness-Males
3.6
2.7
1.8
.9
0
Right Inferior Temporal
0
20
40
60
80
100
Age

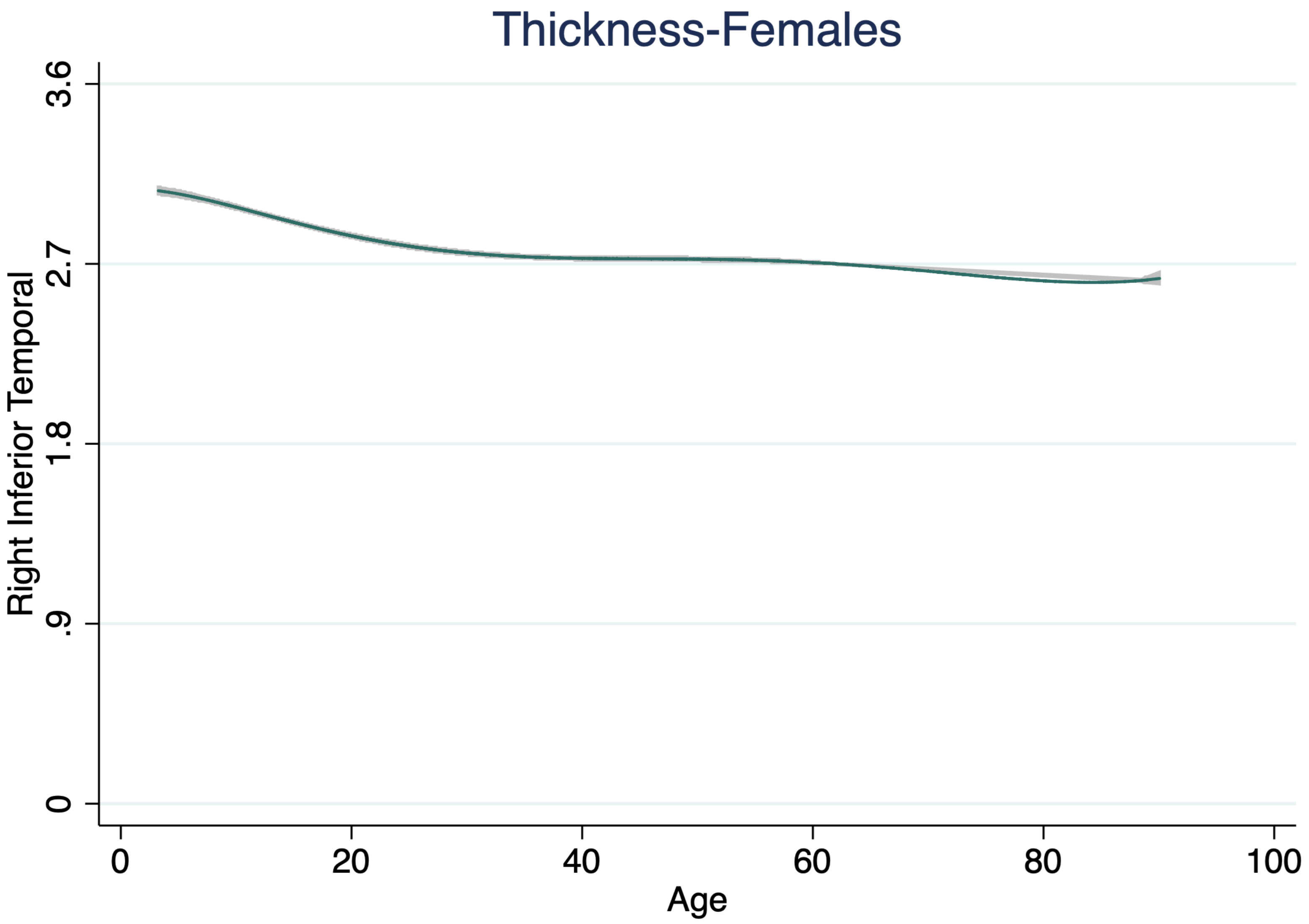

Thickness-Females
3.6
2.7
1.8
.9
0
Right Inferior Temporal
0
20
40
60
80
100
Age

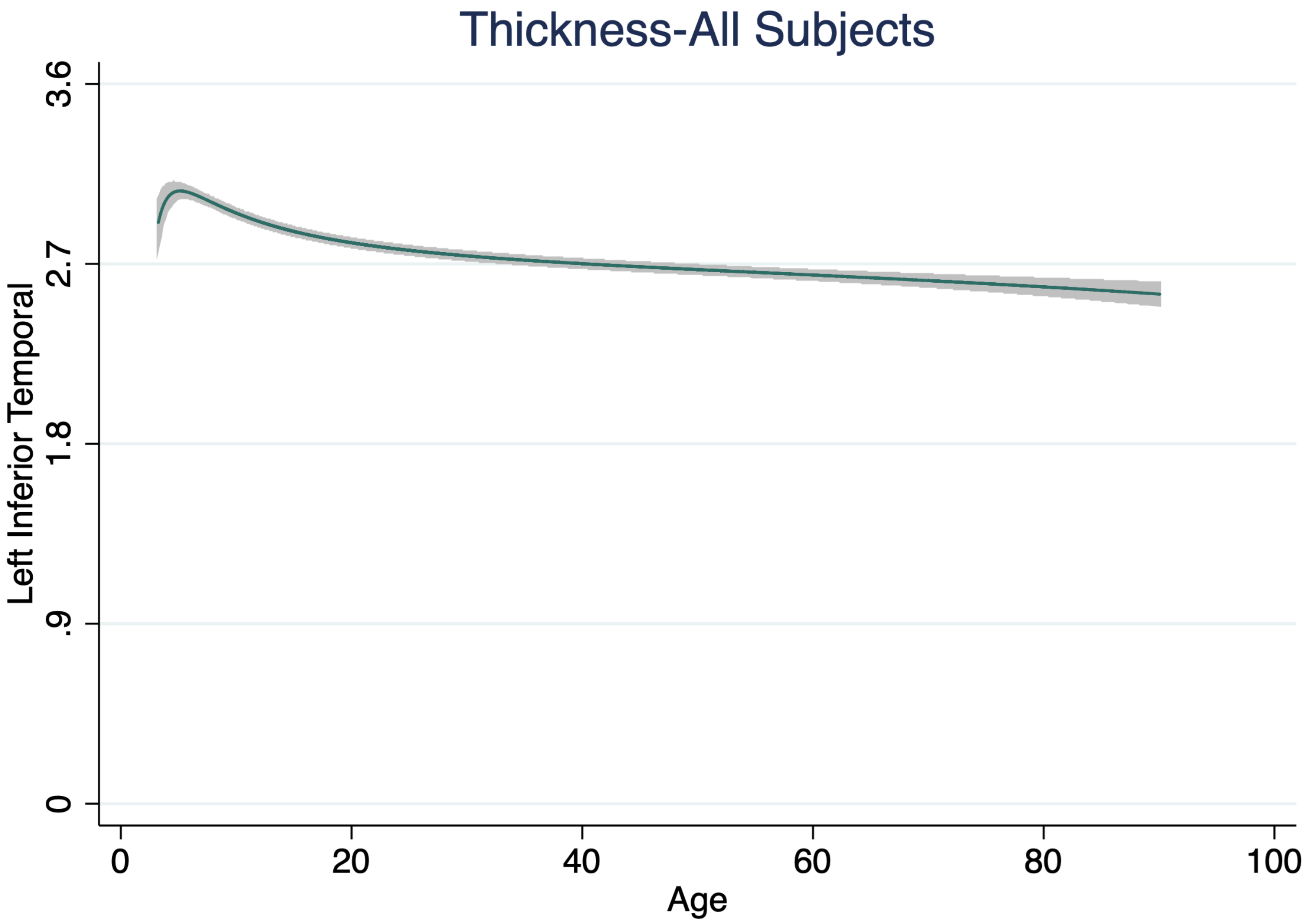
Thickness-All Subjects
Left Inferior Temporal
3.6
2.7
1.8
.9
0
0
20
40
60
80
100
Age

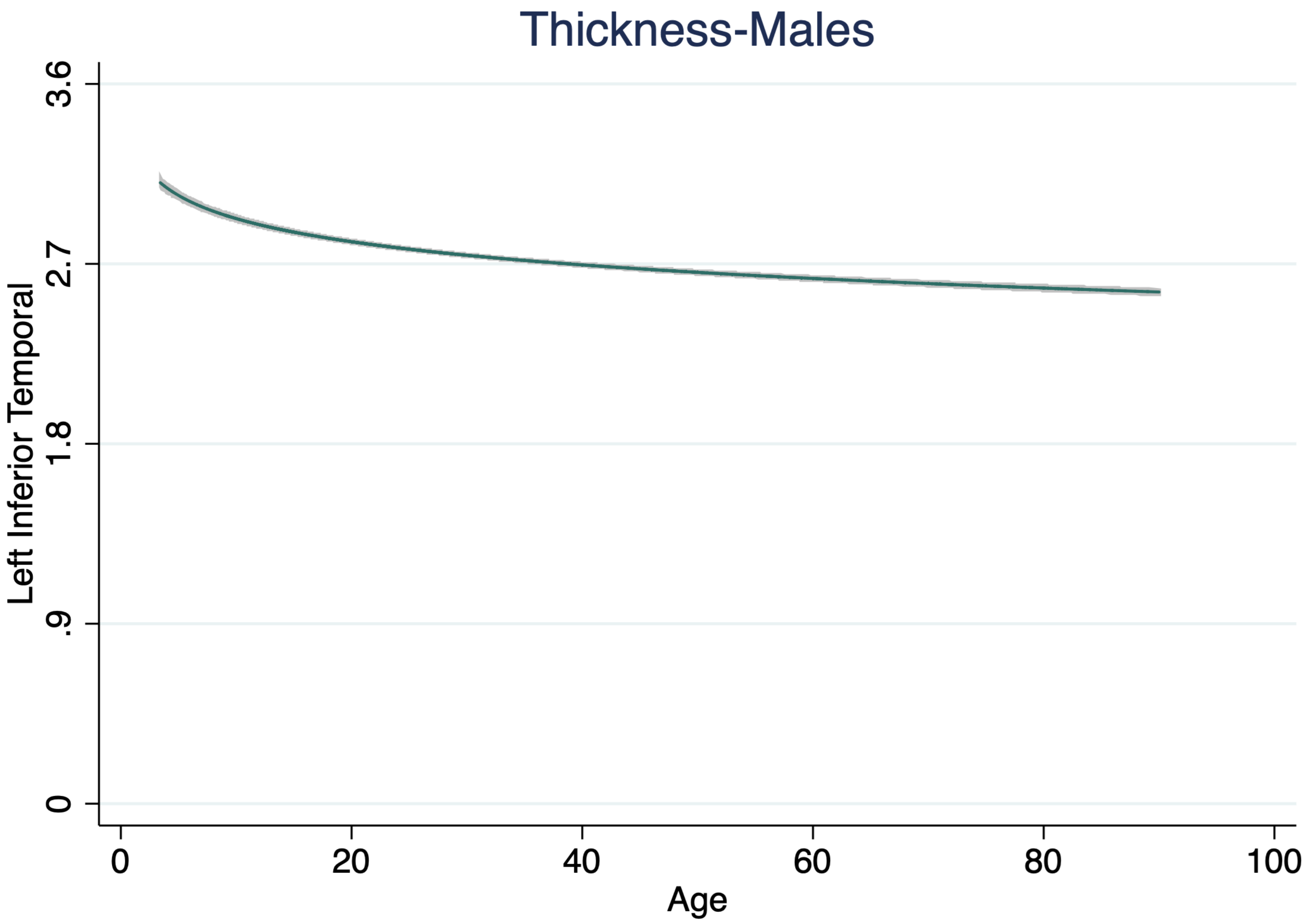
Thickness-Males
Left Inferior Temporal
3.6
2.7
1.8
.9
0
0
20
40
60
80
100
Age

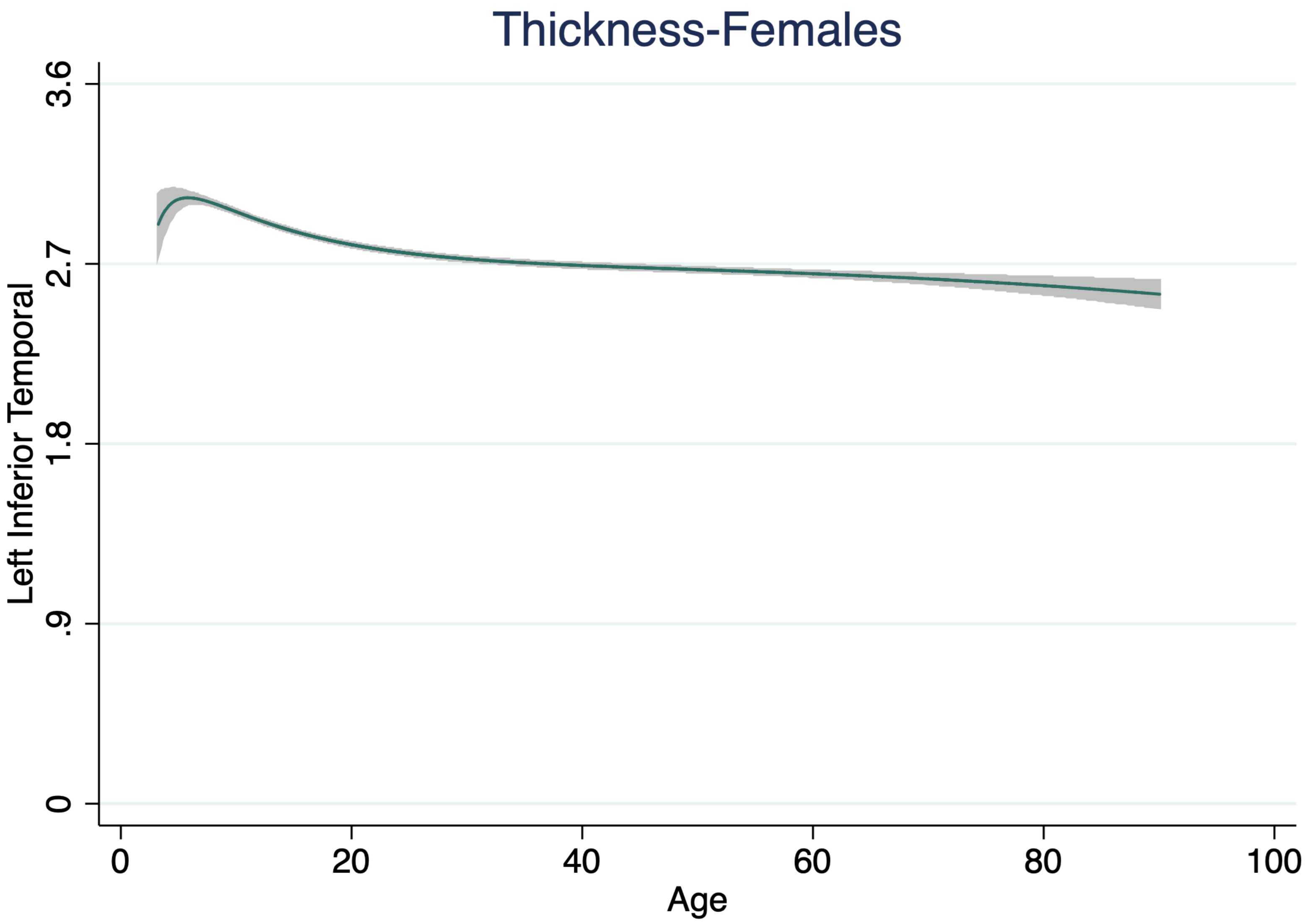
Thickness-Females
Left Inferior Temporal
3.6
2.7
1.8
.9
0
0
20
40
60
80
100
Age

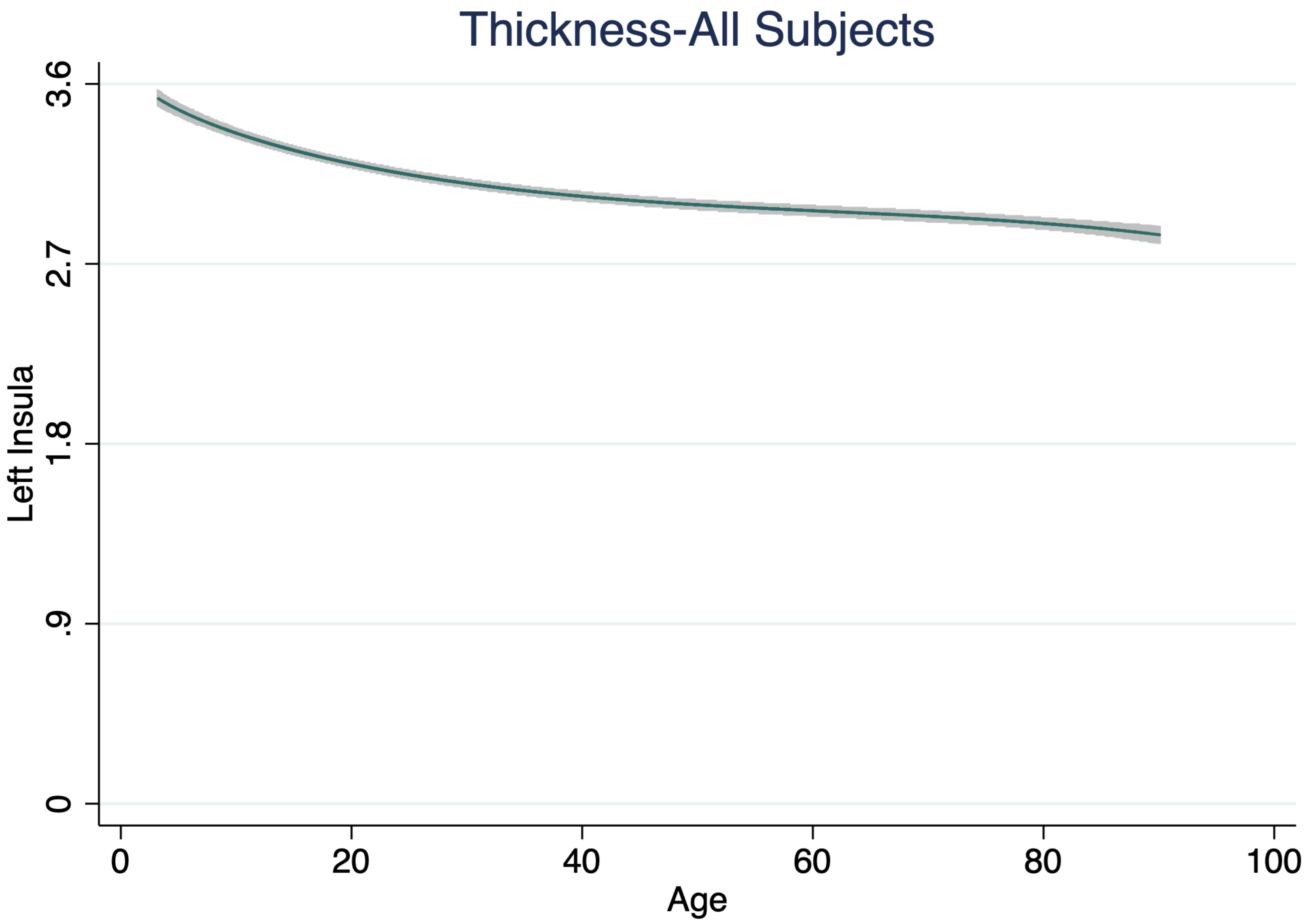
Thickness-All Subjects
3.6
2.7
1.8
.9
0
Left Insula
0
20
40
60
80
100
Age

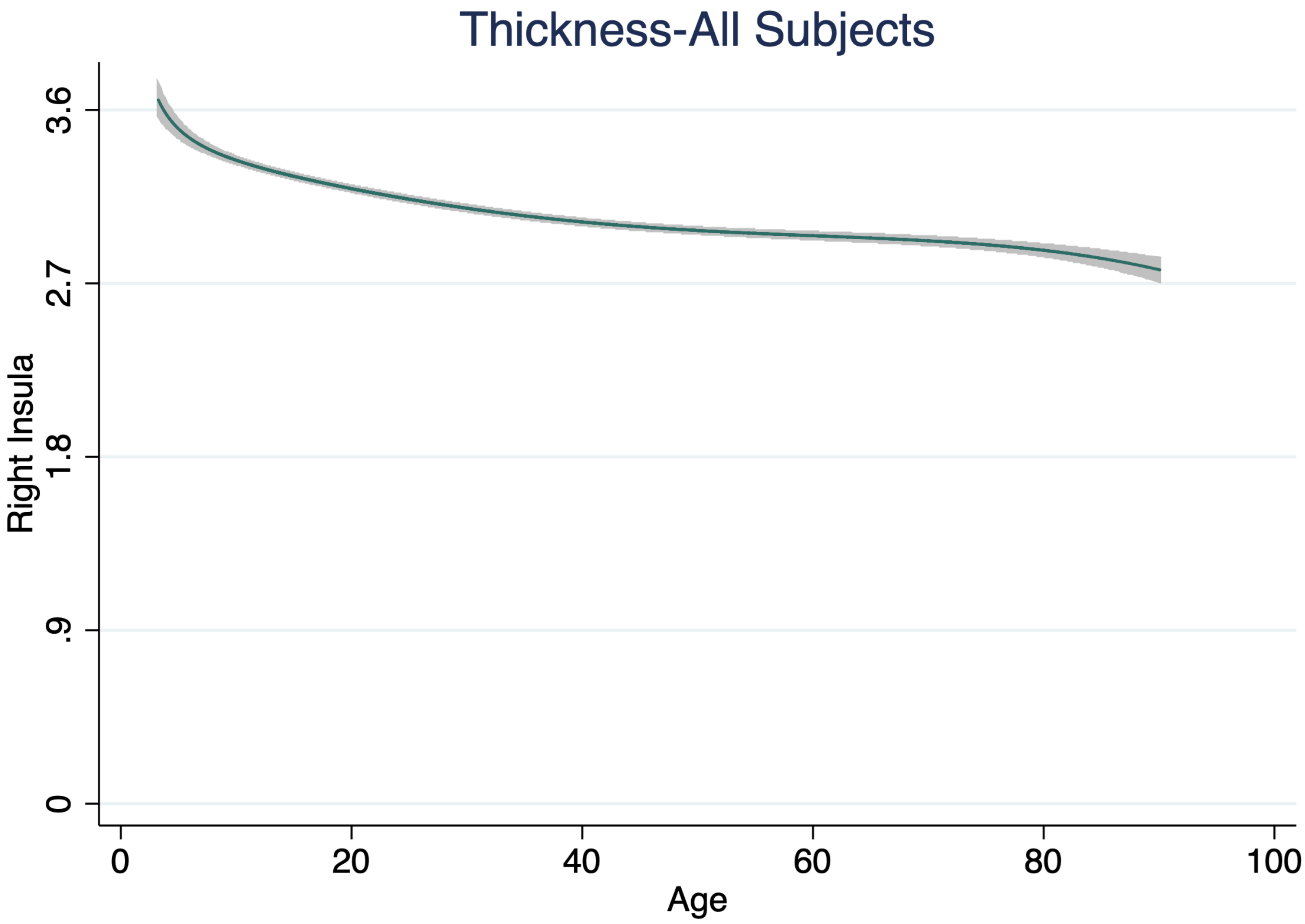
Thickness-All Subjects
Right Insula
3.6
2.7
1.8
.9
0
0
20
40
60
80
100
Age

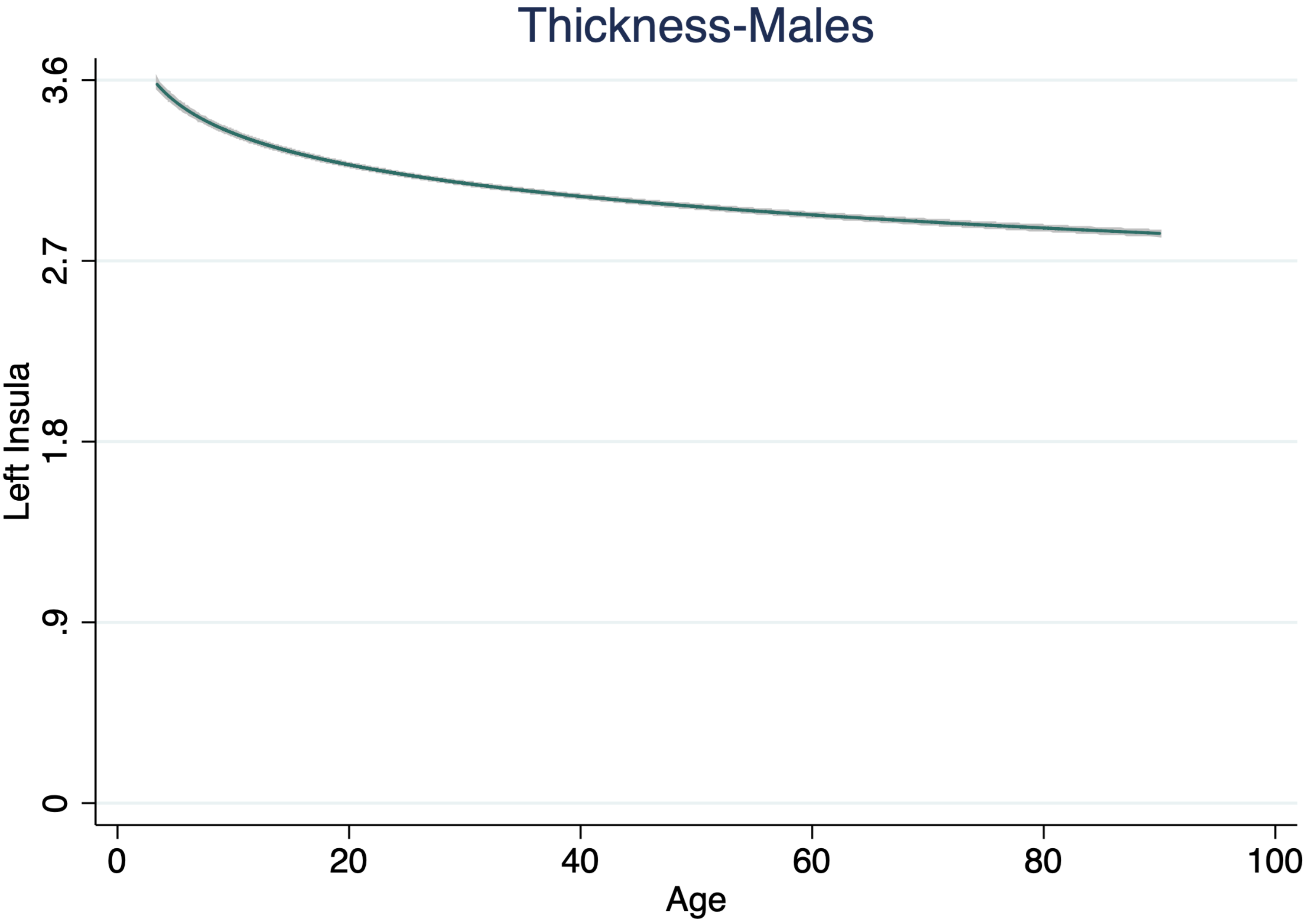

Thickness-Males
3.6
2.7
1.8
.9
0
Left Insula
0
20
40
60
80
100
Age

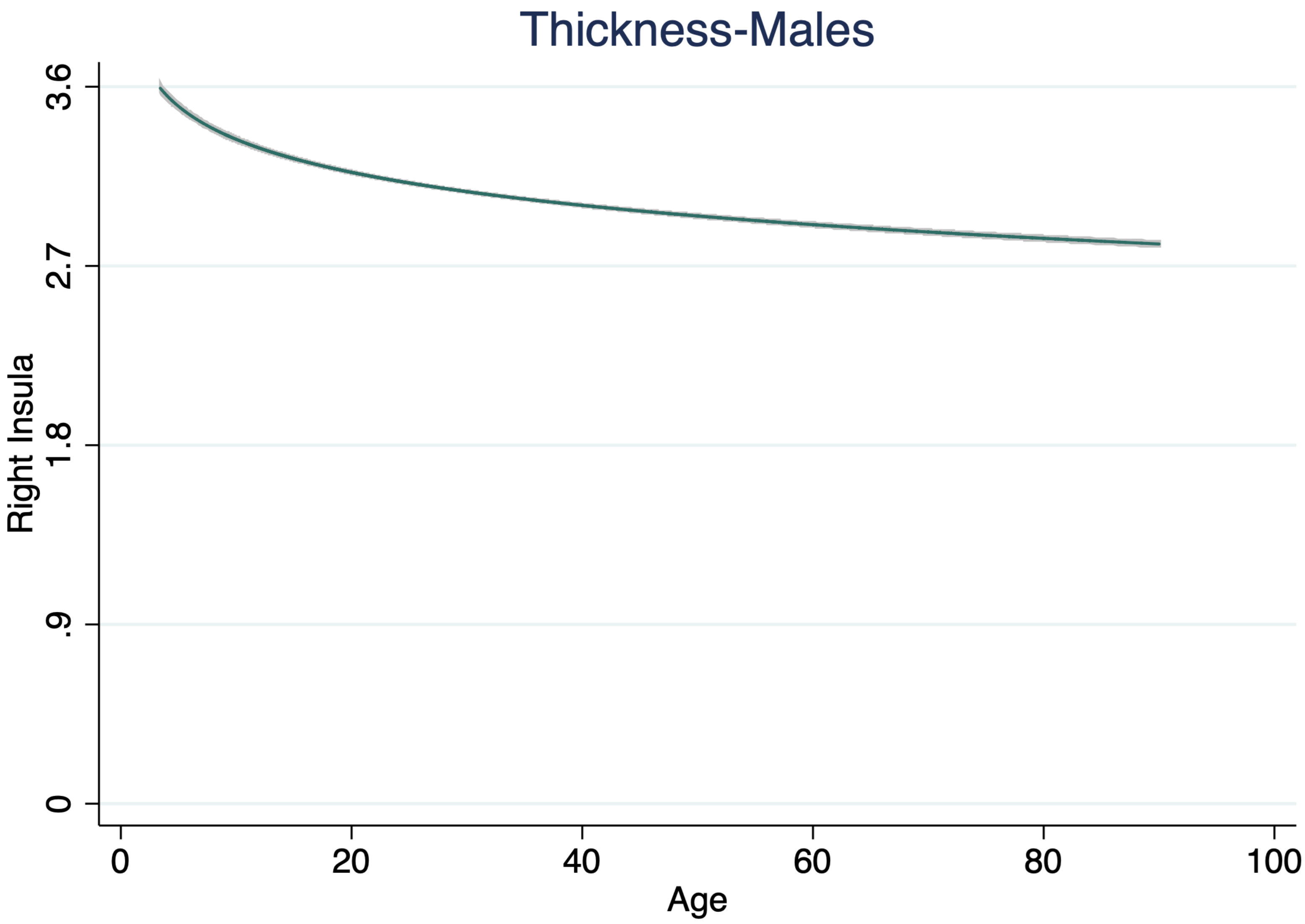

Thickness-Males
Right Insula
3.6
2.7
1.8
.9
0
0
20
40
60
80
100
Age

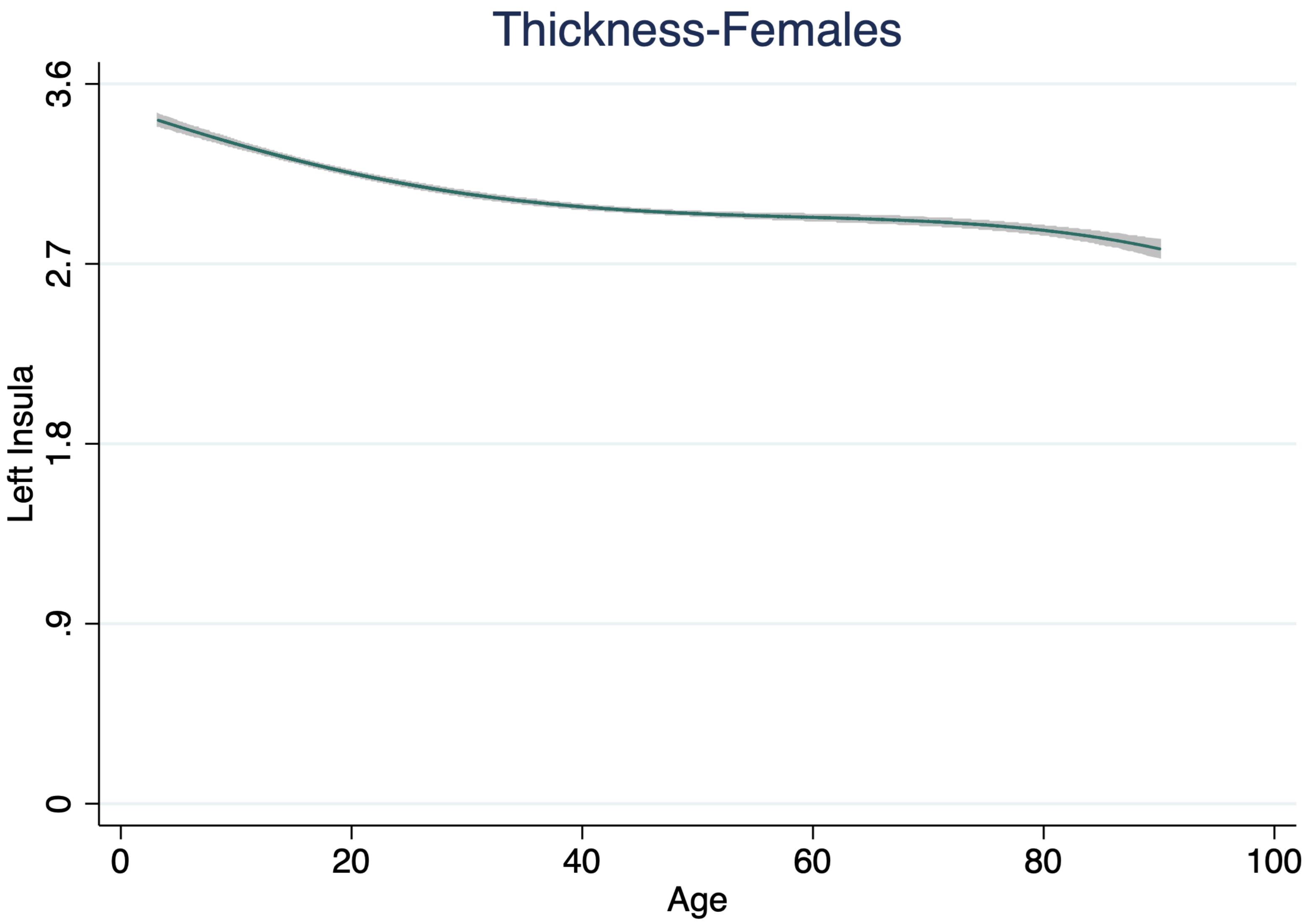
Thickness-Females
3.6
2.7
1.8
.9
0
Left Insula
0
20
40
60
80
100
Age

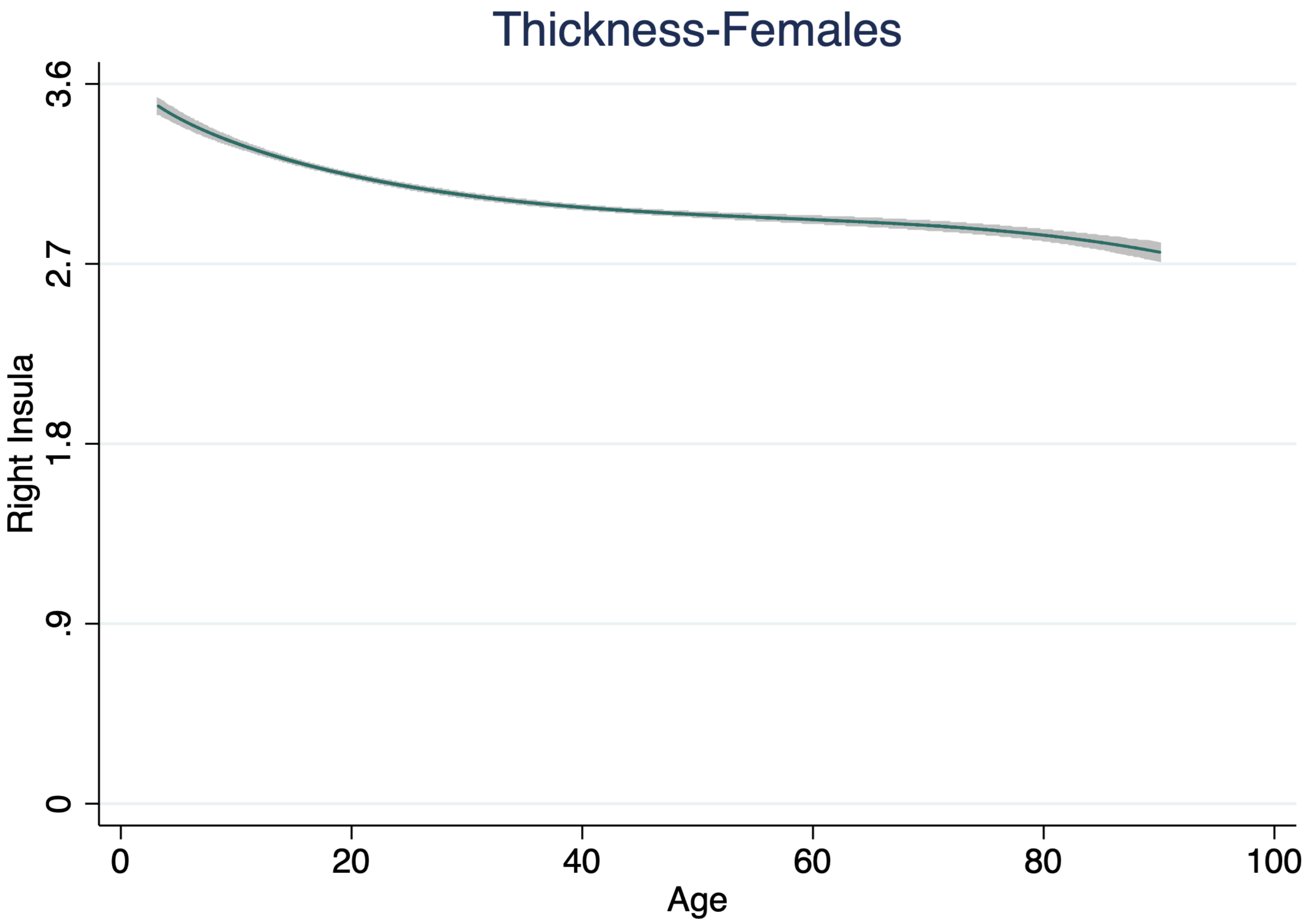

Thickness-Females
3.6
2.7
1.8
.9
0
Right Insula
0
20
40
60
80
100
Age

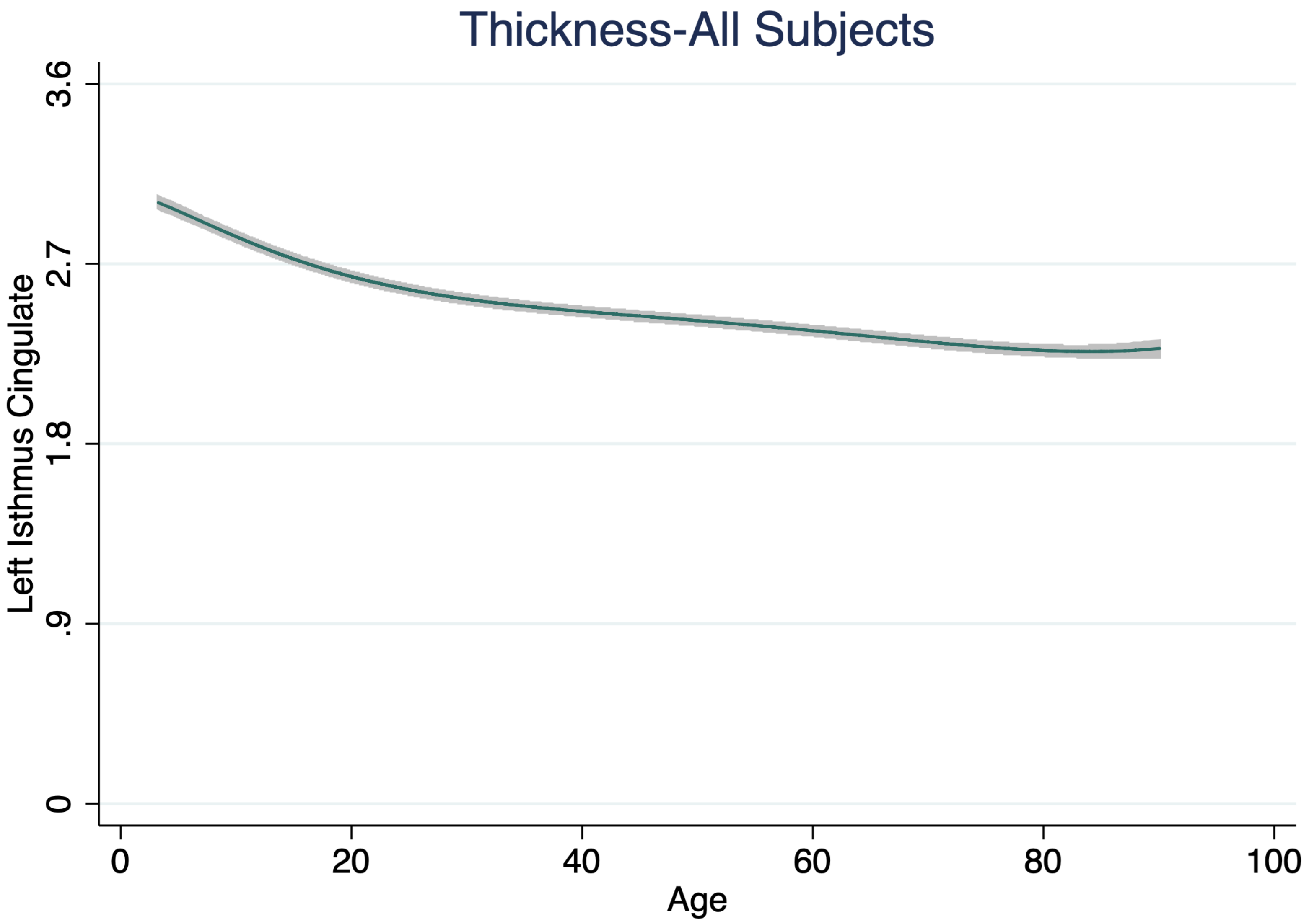

Thickness-All Subjects
Left Isthmus Cingulate
3.6
2.7
1.8
.9
0
0
20
40
60
80
100
Age

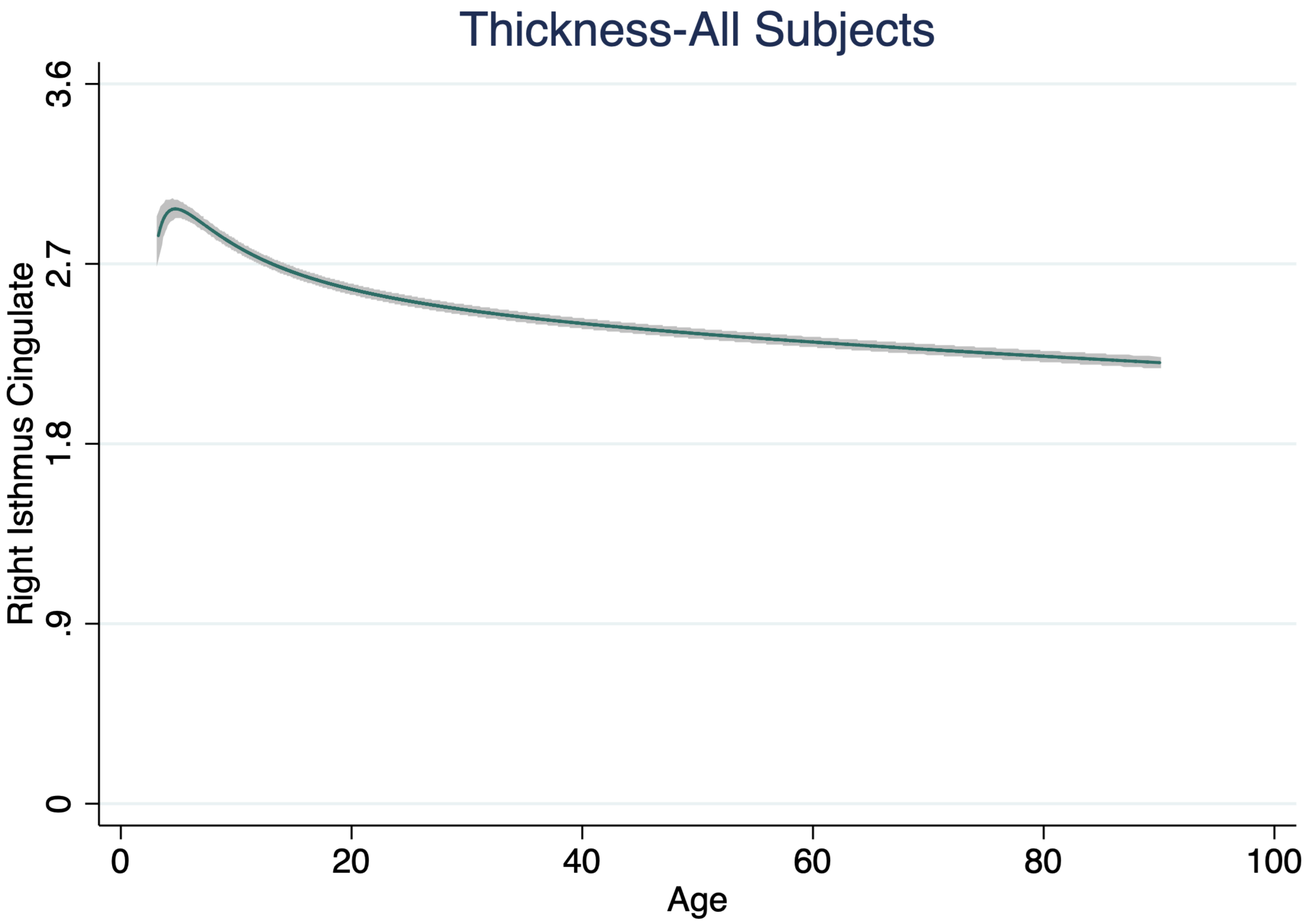
Thickness-All Subjects
Right Isthmus Cingulate
3.6
2.7
1.8
.9
0
0
20
40
60
80
100
Age

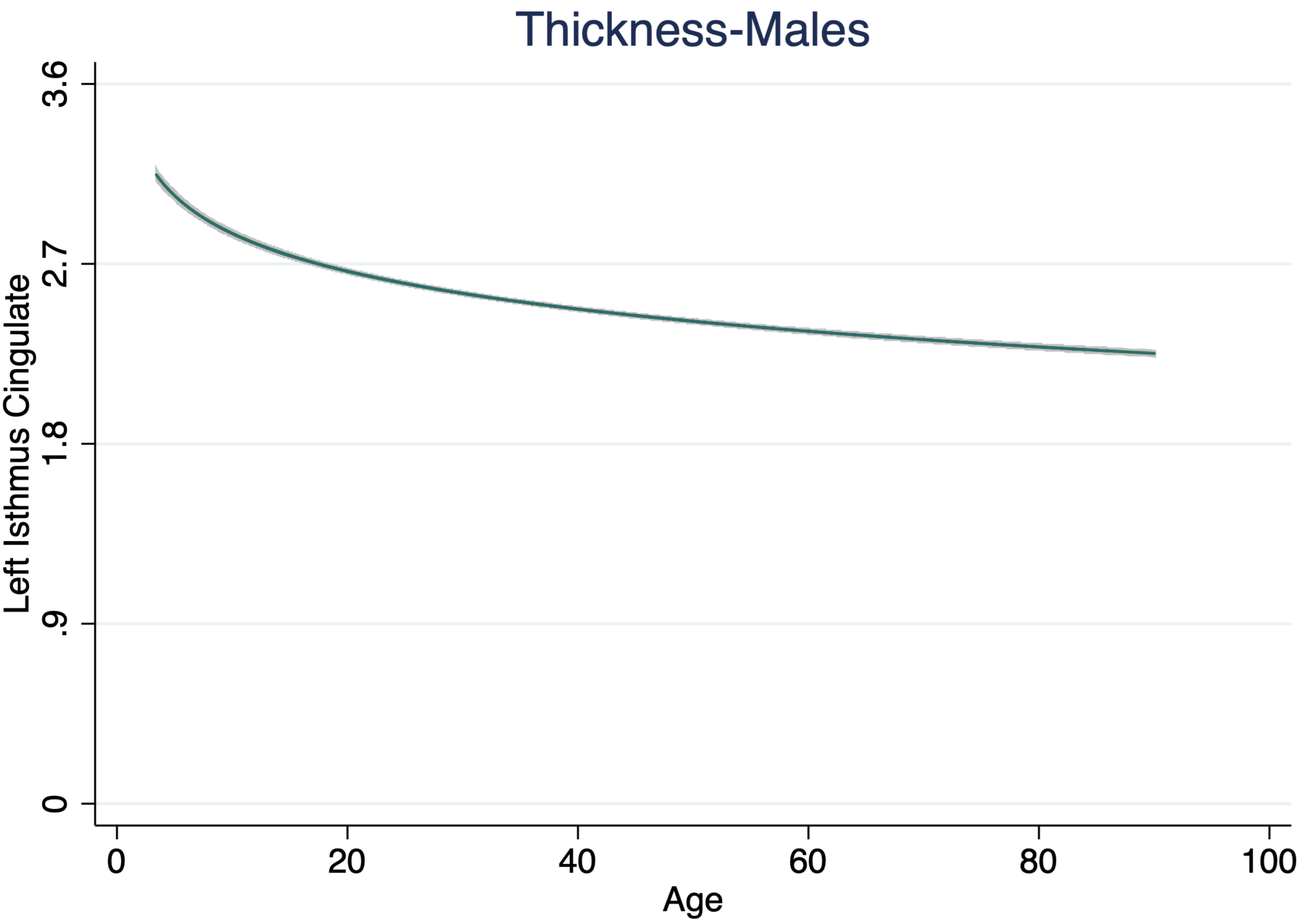

Thickness-Males
Left Isthmus Cingulate
3.6
2.7
1.8
.9
0
0
20
40
60
80
100
Age

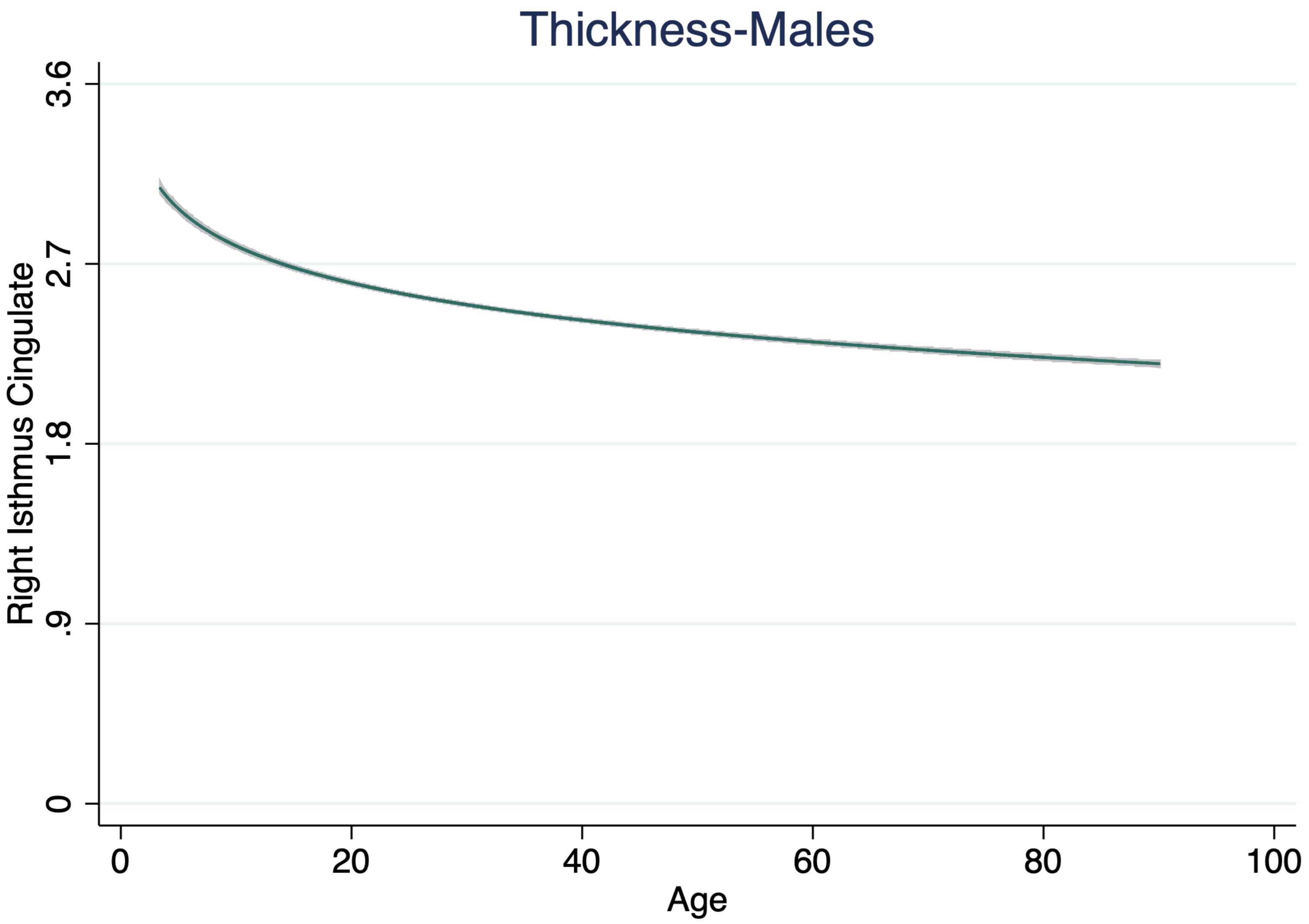

Thickness-Males
Right Isthmus Cingulate
3.6
2.7
1.8
.9
0
0
20
40
60
80
100
Age

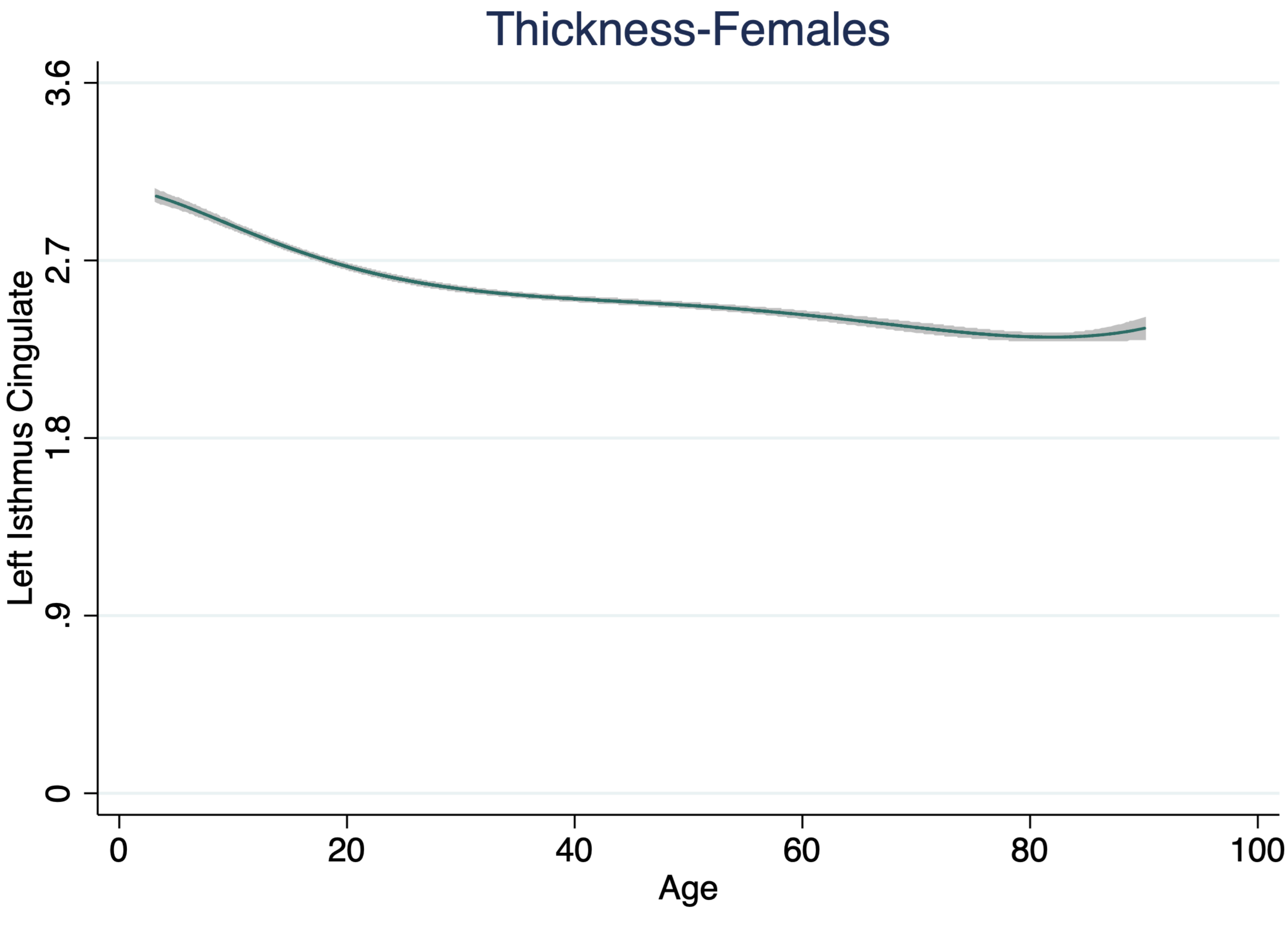

Thickness-Females
Left Isthmus Cingulate
3.6
2.7
1.8
.9
0
0
20
40
60
80
100
Age

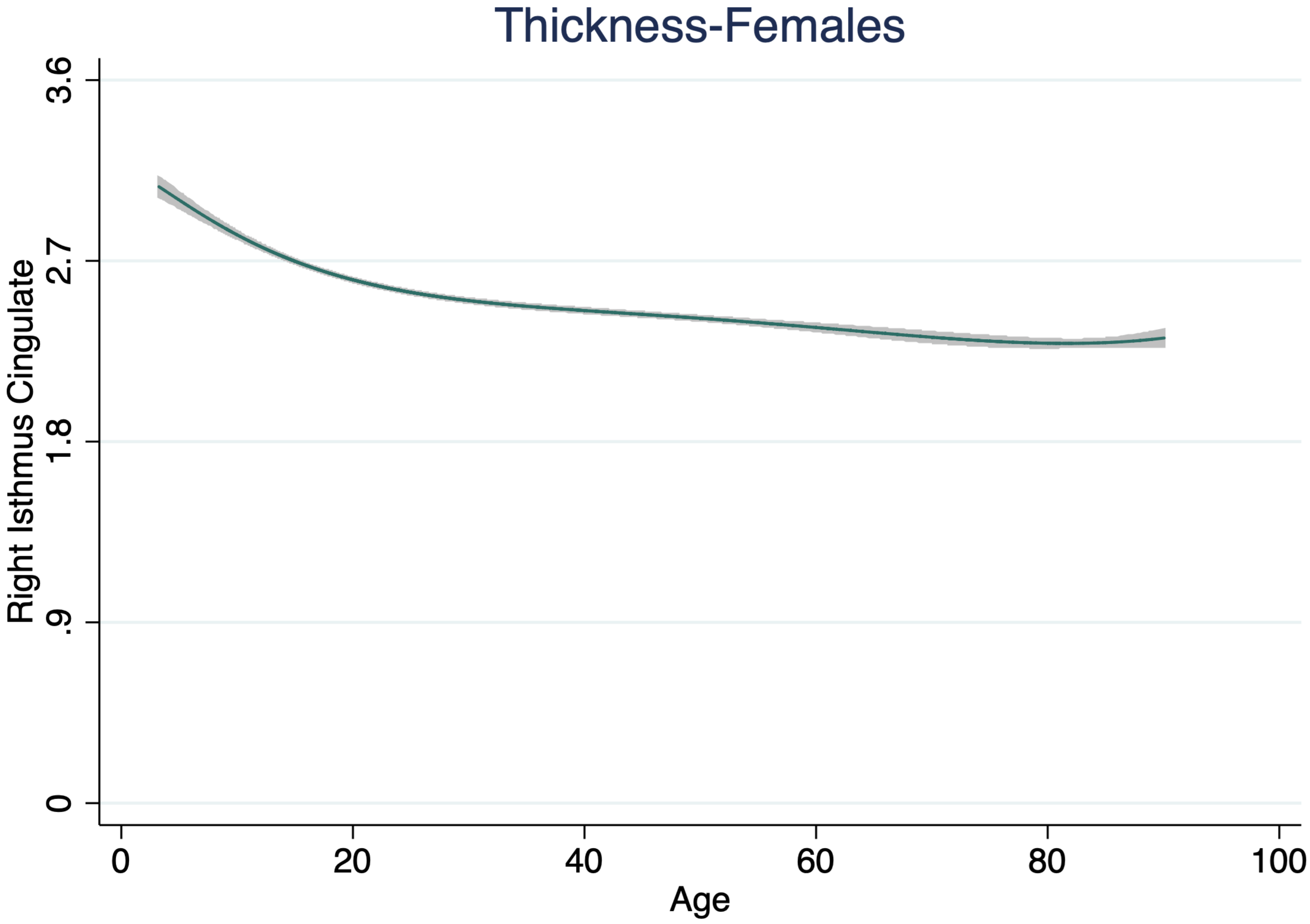

Thickness-Females
Right Isthmus Cingulate
3.6
2.7
1.8
.9
0
0
20
40
60
80
100
Age

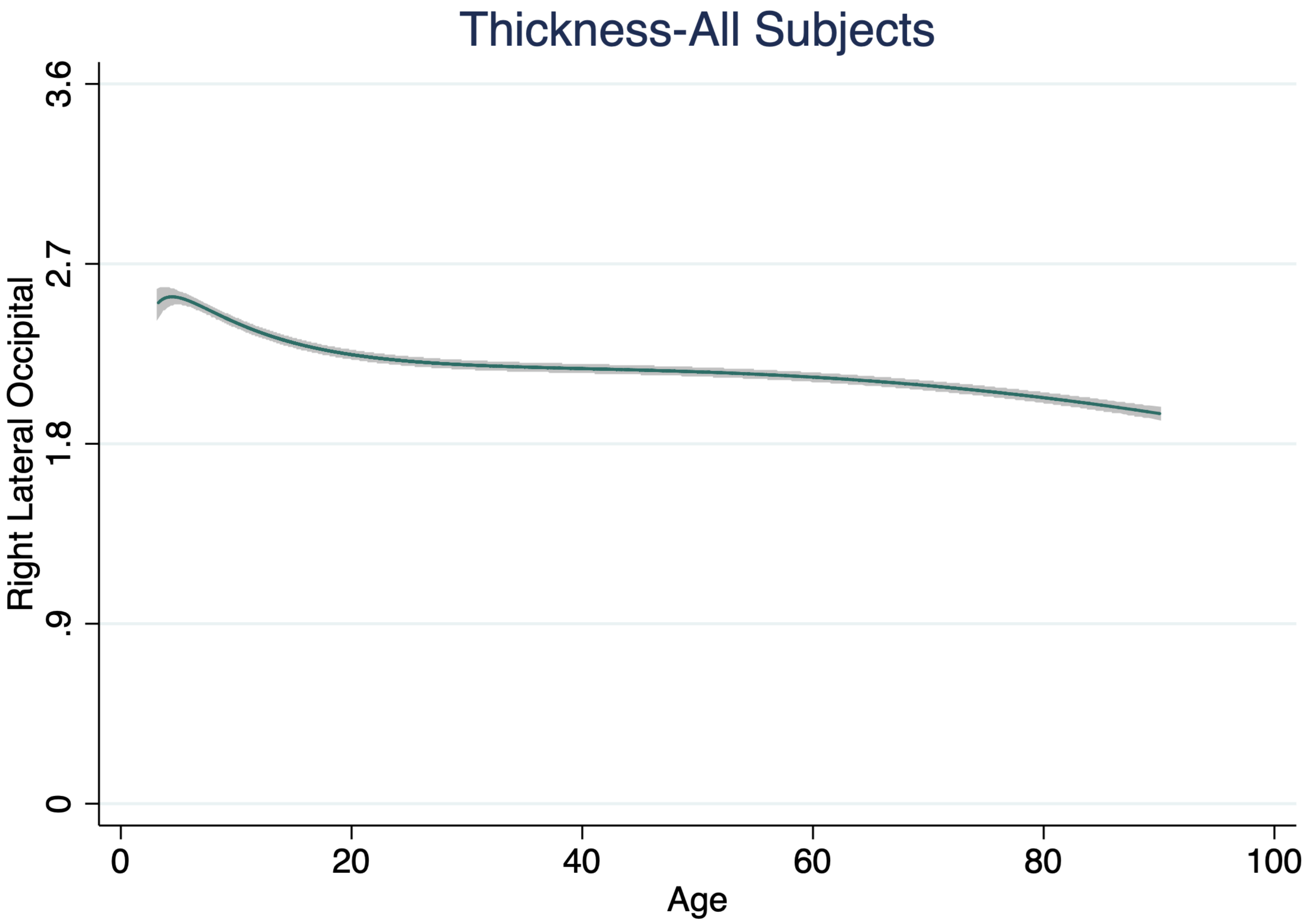
Thickness-All Subjects
Right Lateral Occipital
3.6
2.7
1.8
.9
0
0
20
40
60
80
100
Age

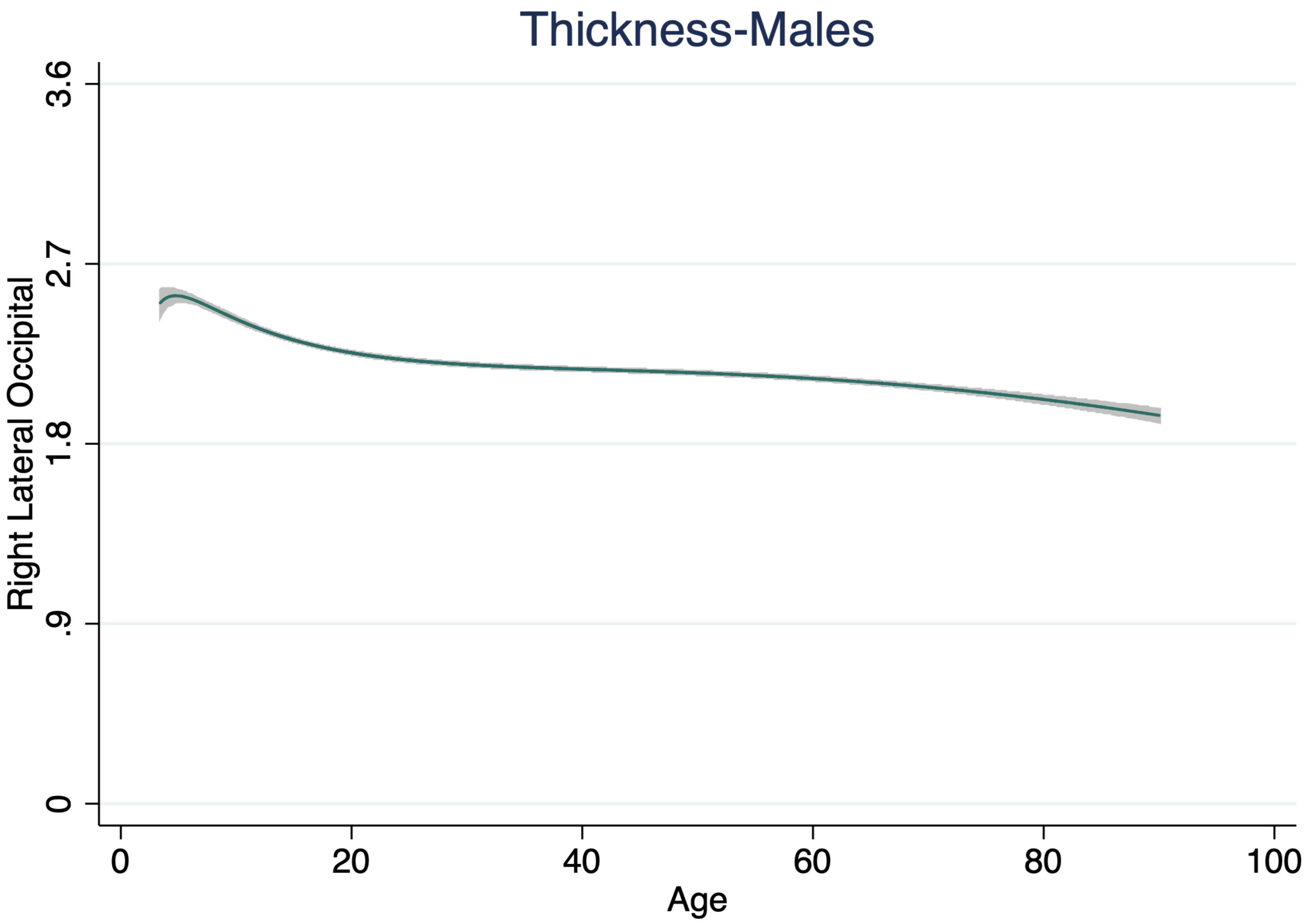

Thickness-Males
Right Lateral Occipital
3.6
2.7
1.8
.9
0
0
20
40
60
80
100
Age

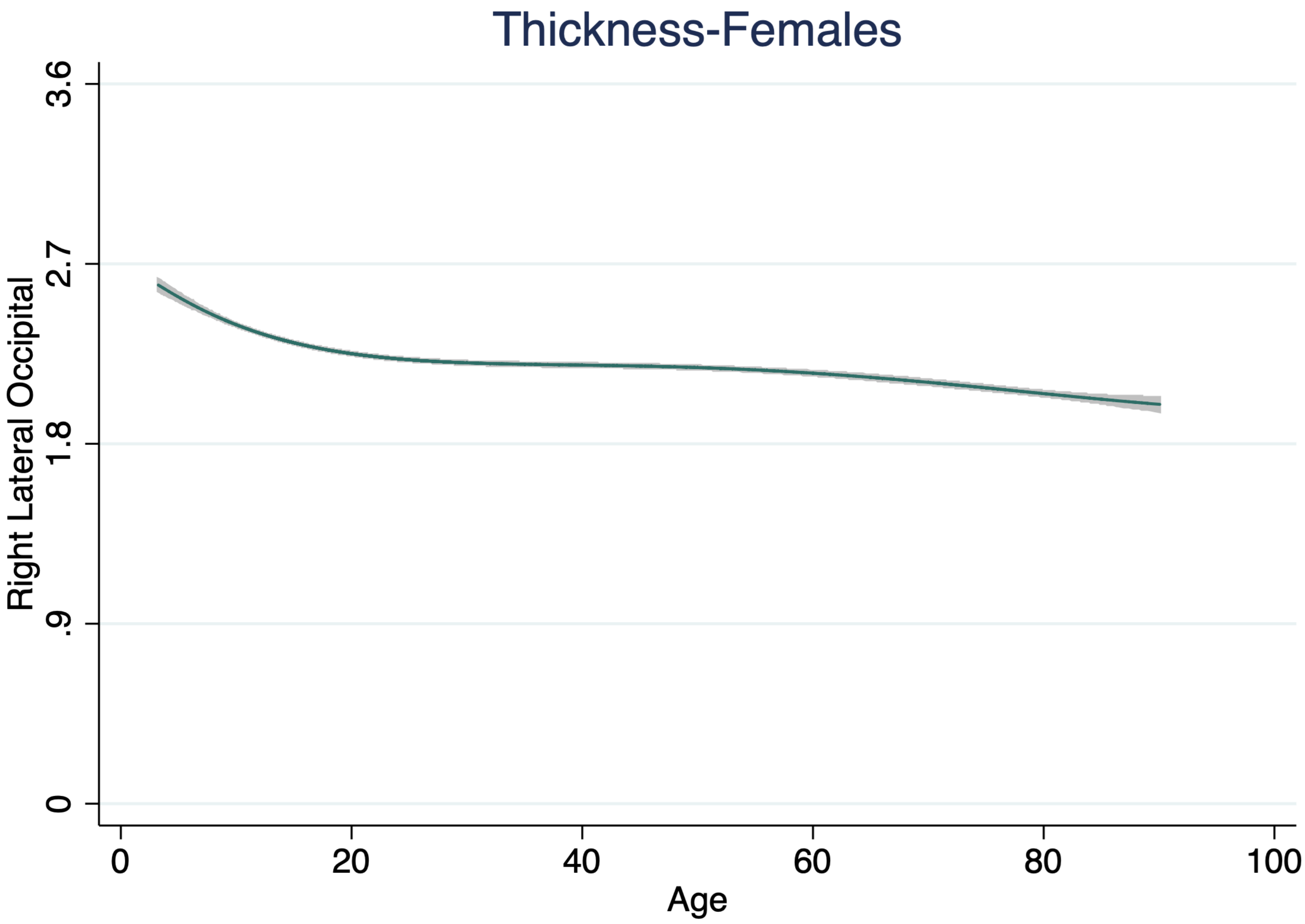
Thickness-Females
Right Lateral Occipital
3.6
2.7
1.8
.9
0
0
20
40
60
80
100
Age

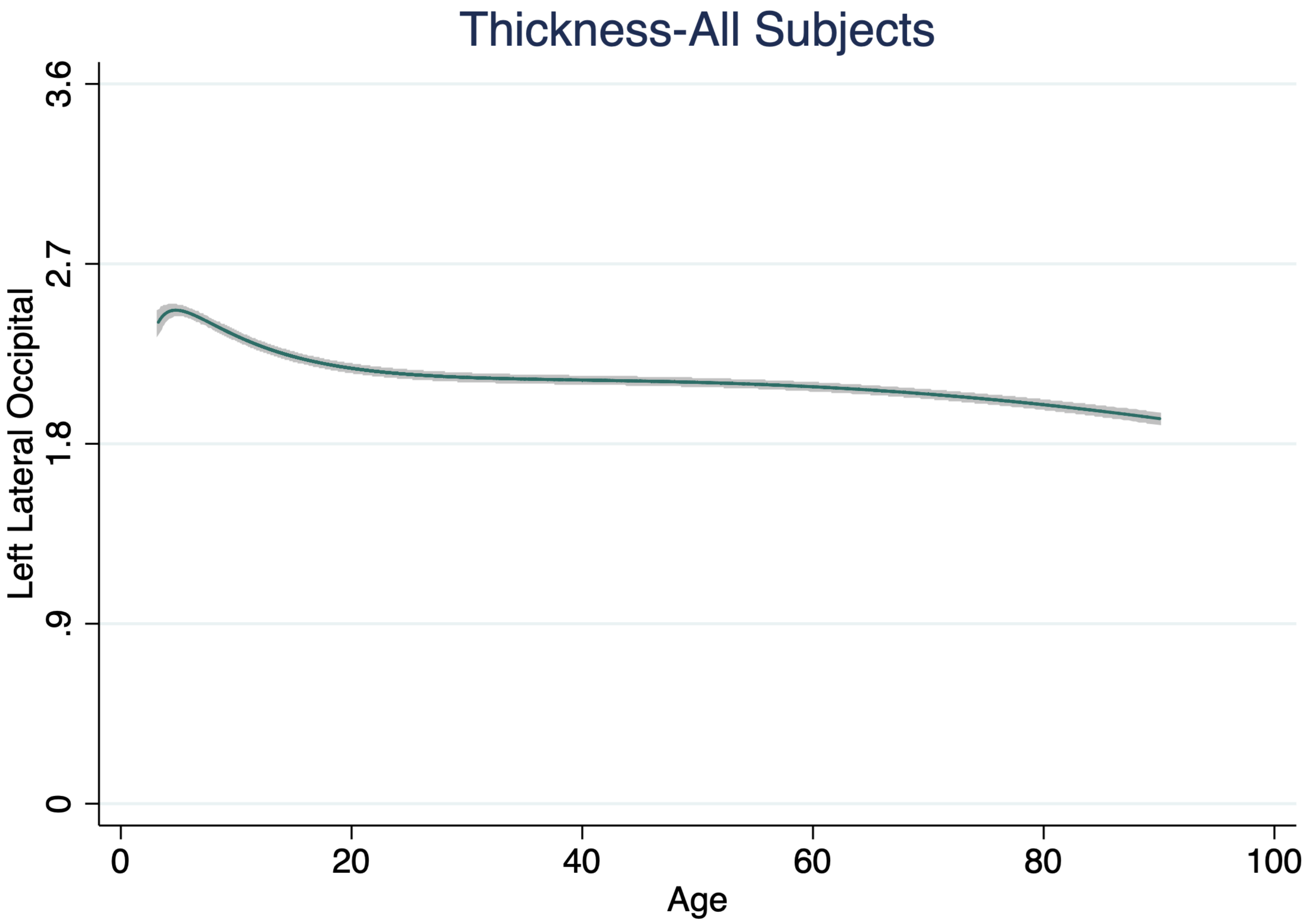

Thickness-All Subjects
Left Lateral Occipital
3.6
2.7
1.8
.9
0
0
20
40
60
80
100
Age

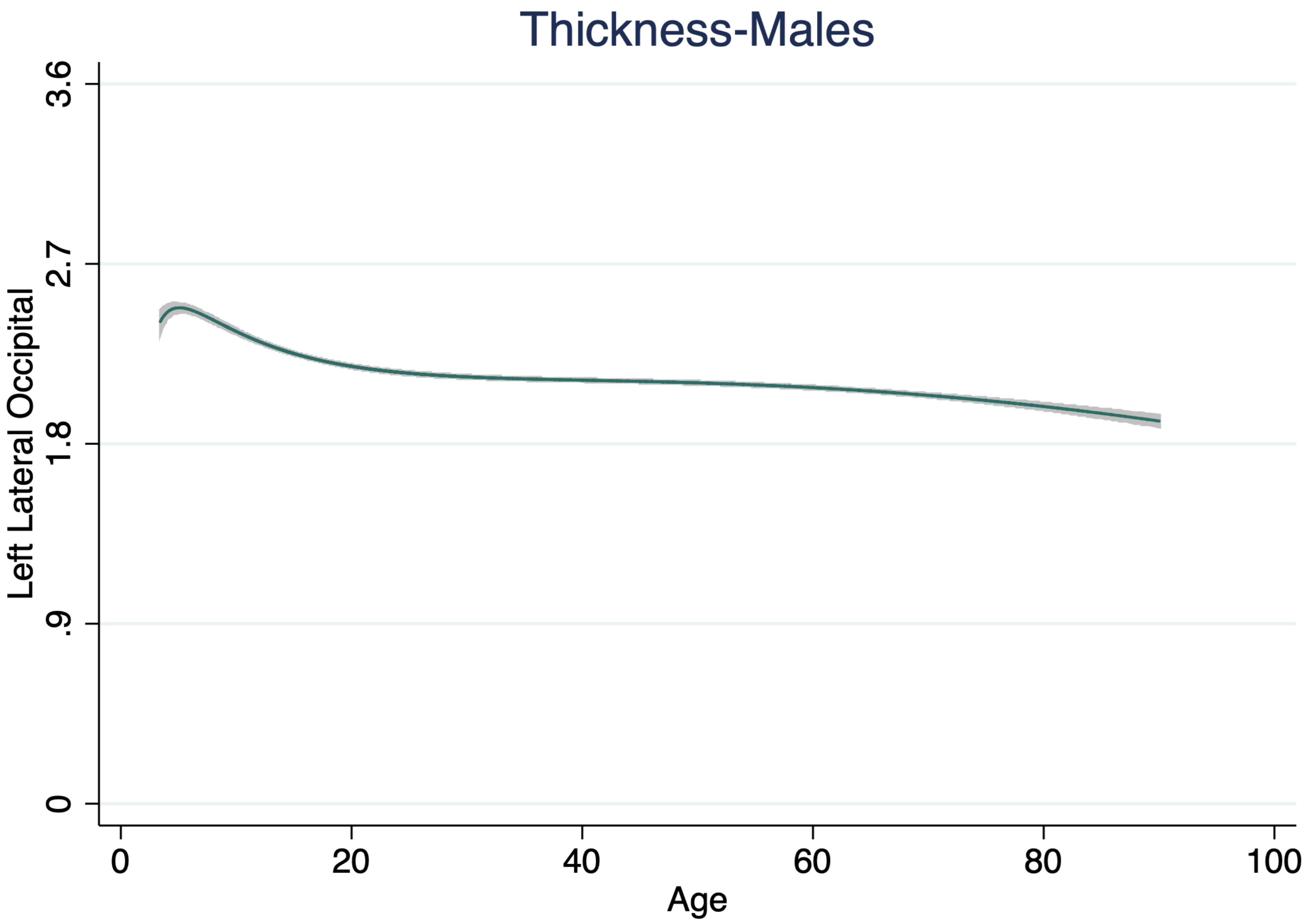

Thickness-Males
Left Lateral Occipital
3.6
2.7
1.8
.9
0
0
20
40
60
80
100
Age

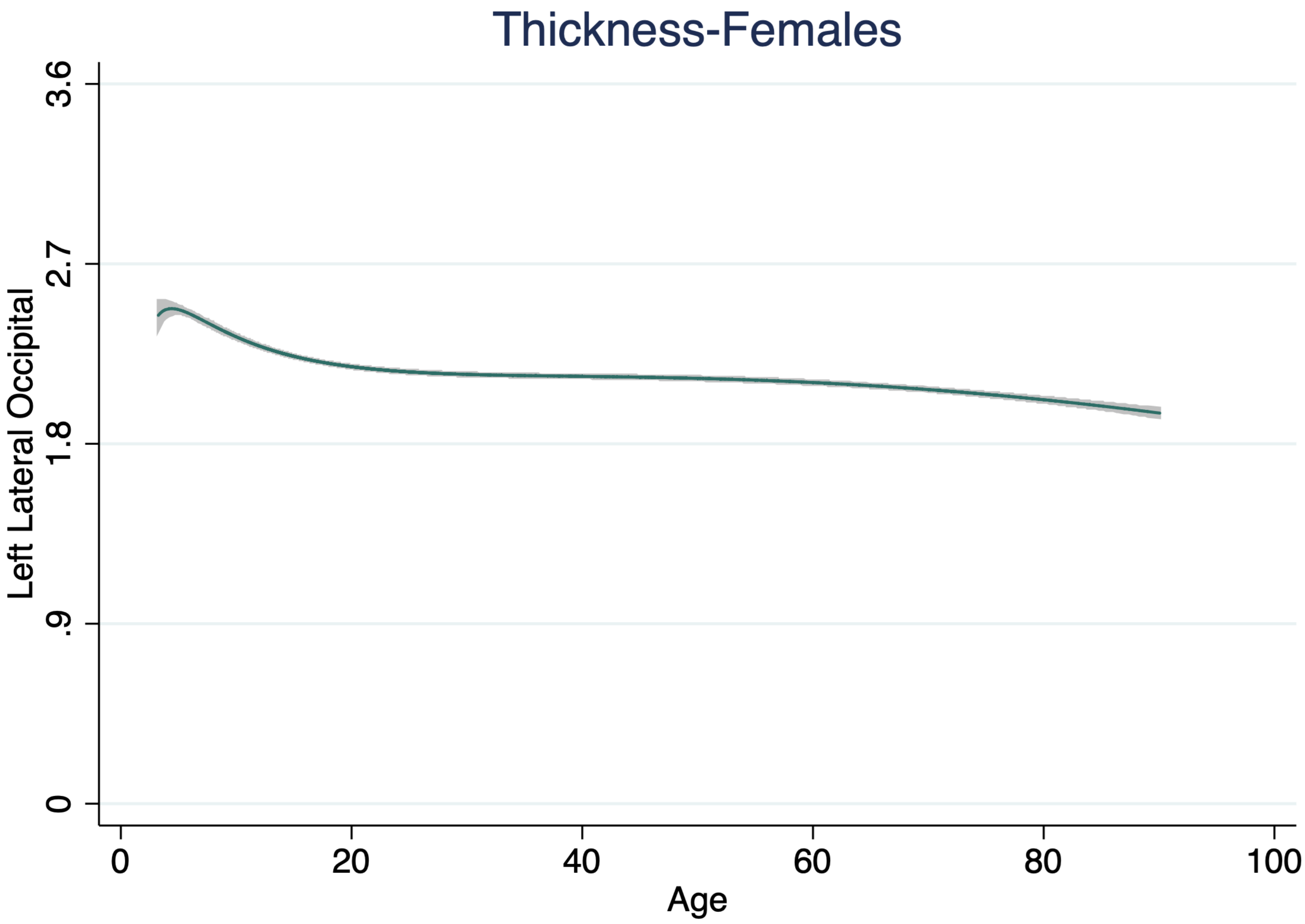
Thickness-Females
Left Lateral Occipital
3.6
2.7
1.8
.9
0
0
20
40
60
80
100
Age

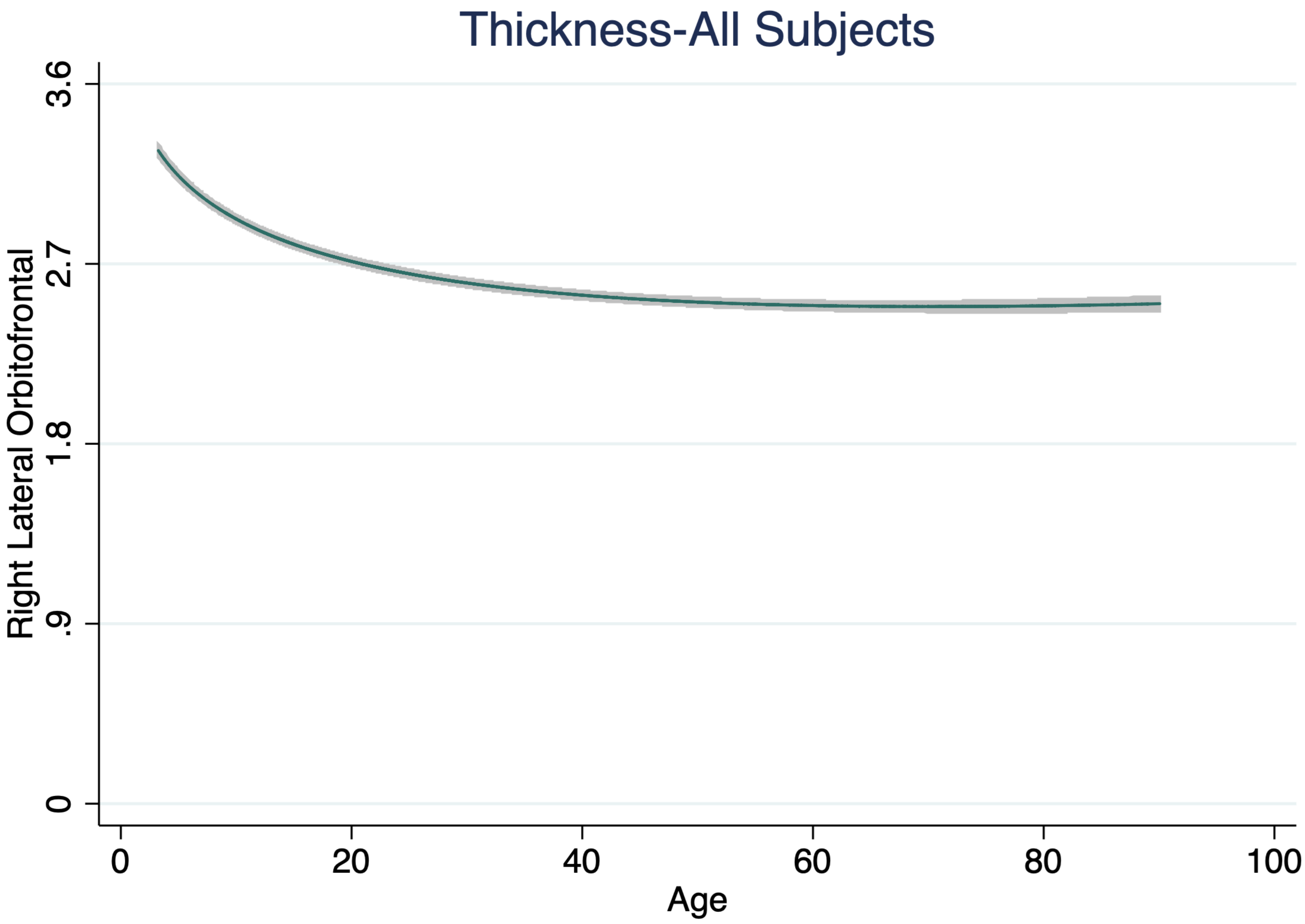
Thickness-All Subjects
Right Lateral Orbitofrontal
3.6
2.7
1.8
.9
0
0
20
40
60
80
100
Age

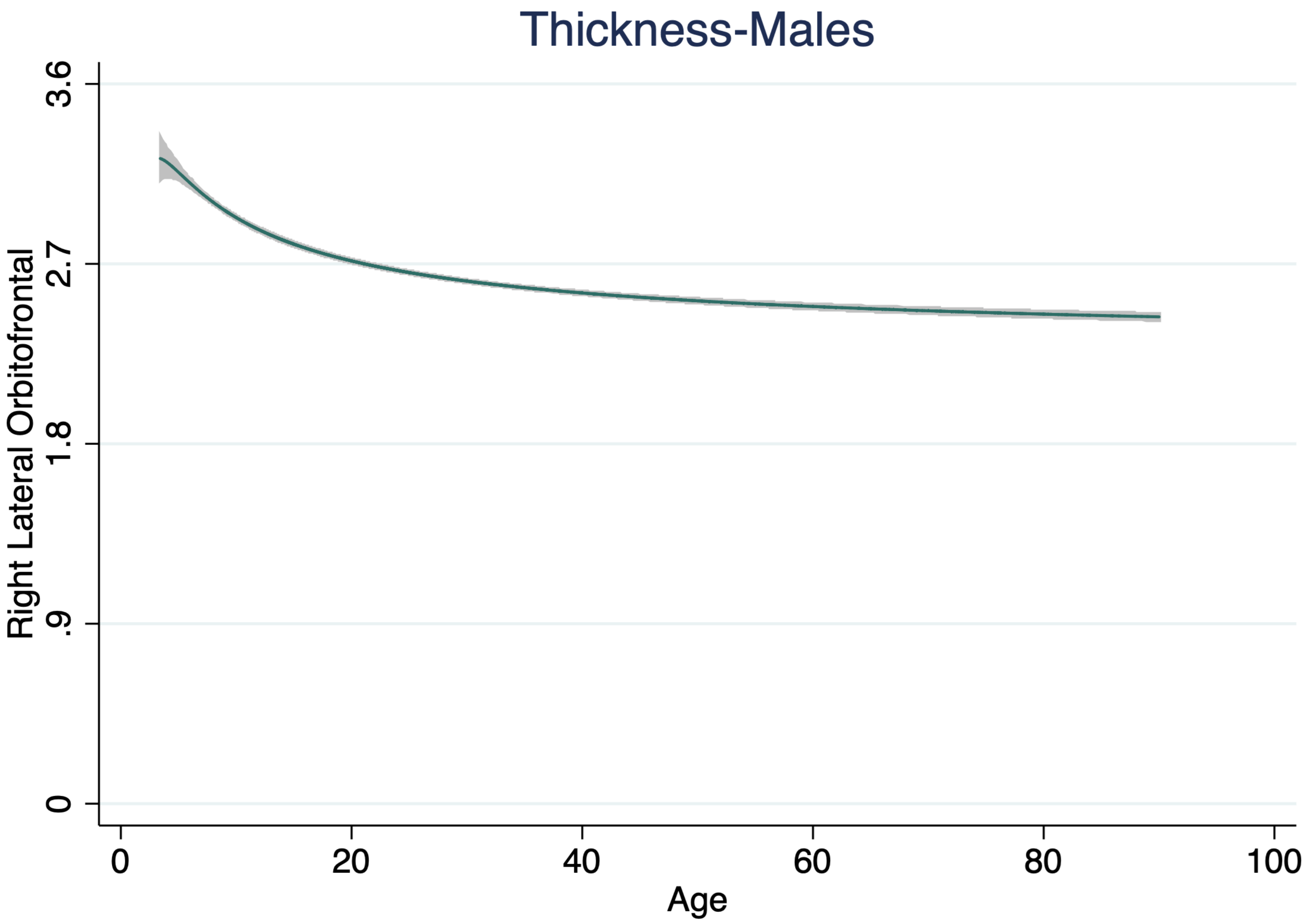

Thickness-Males
Right Lateral Orbitofrontal
3.6
2.7
1.8
.9
0
0
20
40
60
80
100
Age

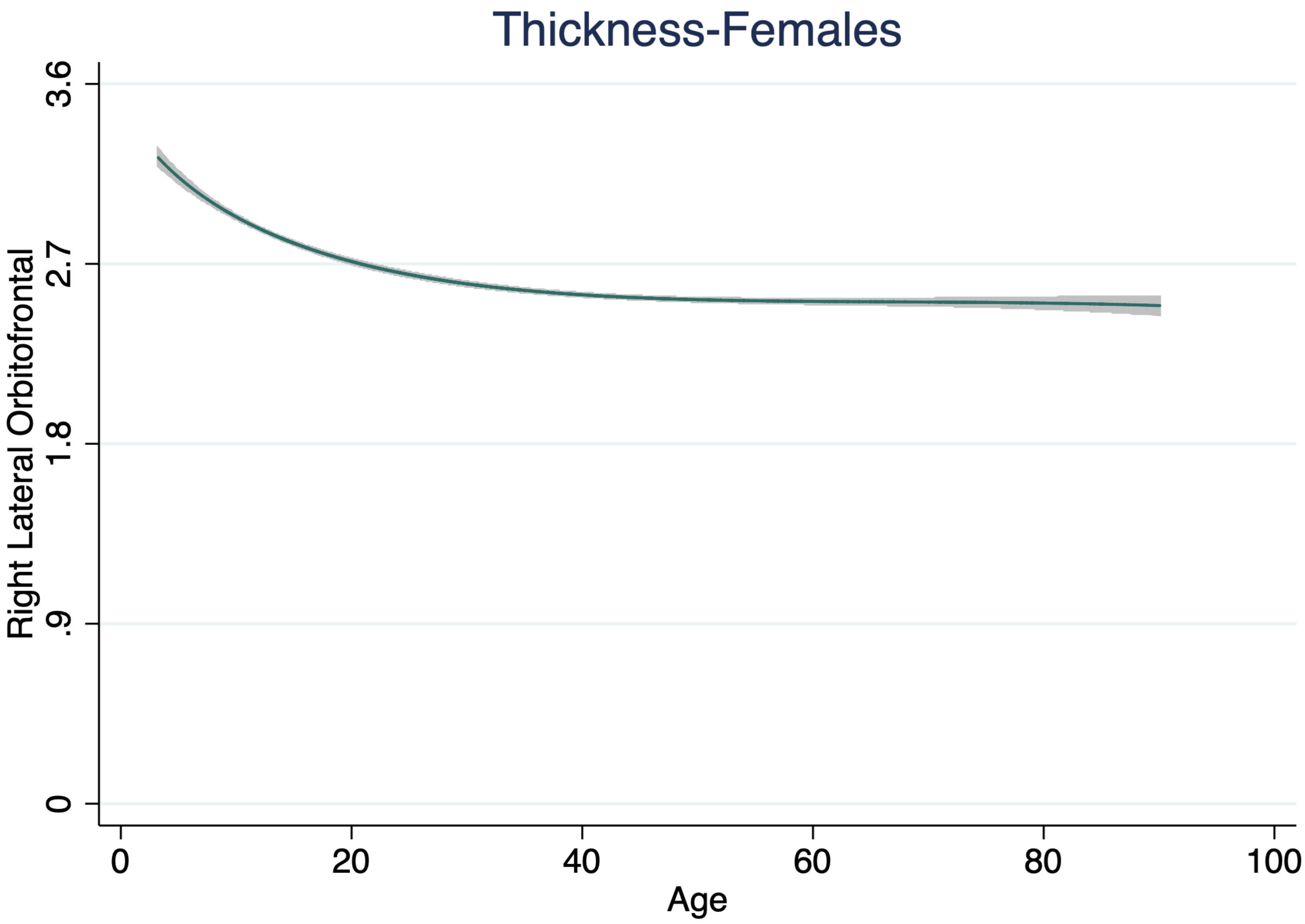
Thickness-Females
Right Lateral Orbitofrontal
3.6
2.7
1.8
.9
0
0
20
40
60
80
100
Age

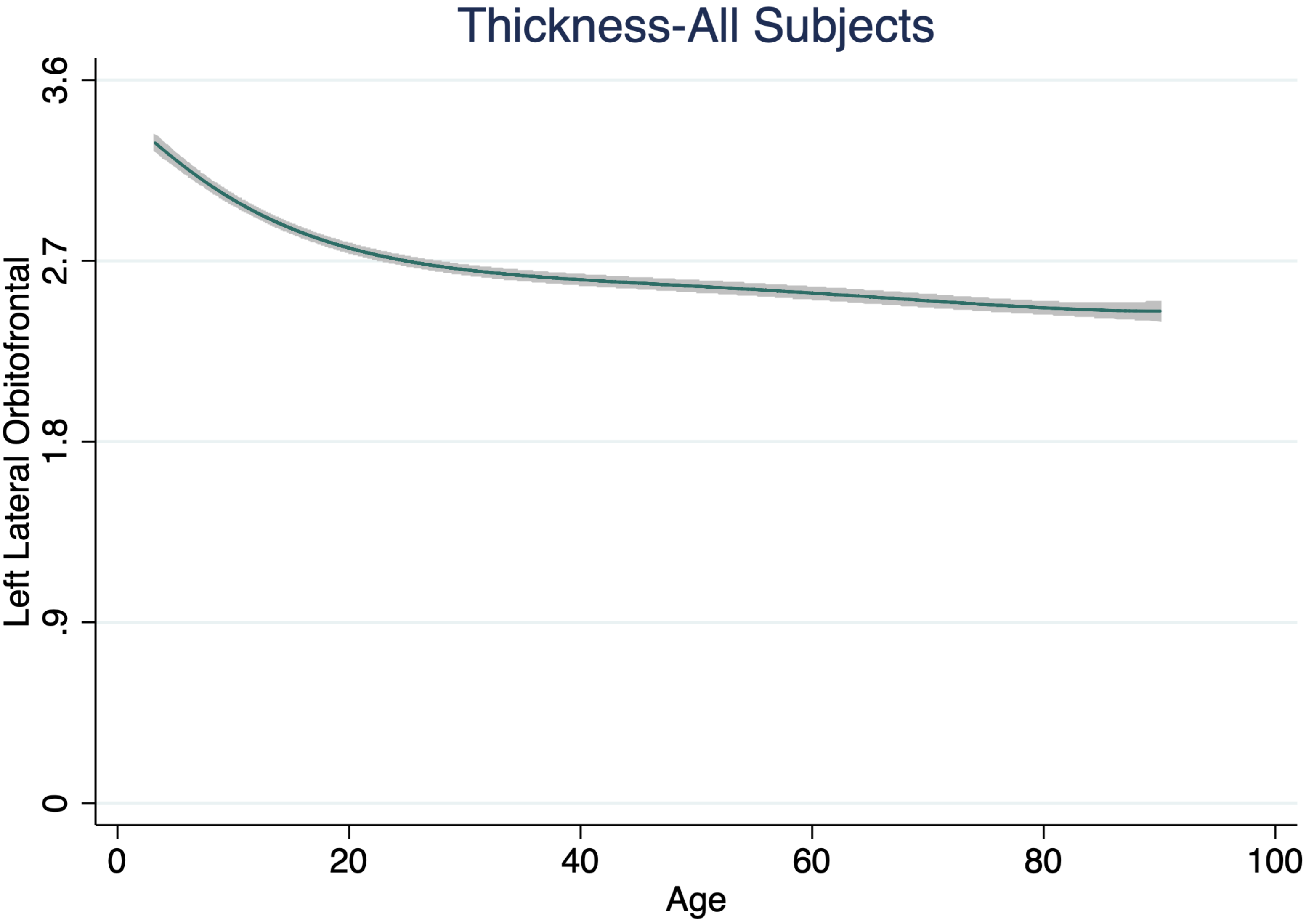

Thickness-All Subjects
Left Lateral Orbitofrontal
3.6
2.7
1.8
.9
0
0
20
40
60
80
100
Age

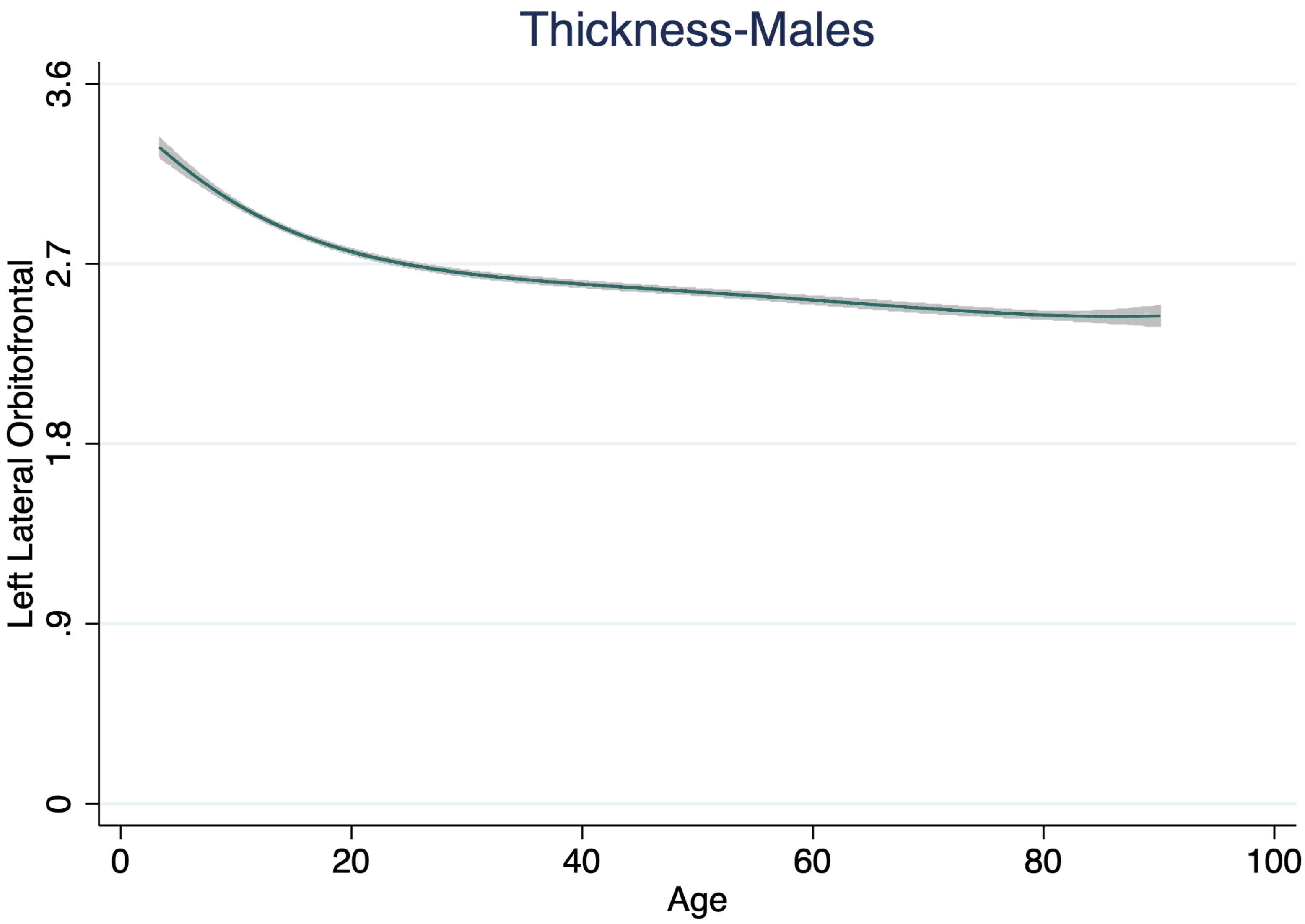
Thickness-Males
Left Lateral Orbitofrontal
3.6
2.7
1.8
.9
0
0
20
40
60
80
100
Age

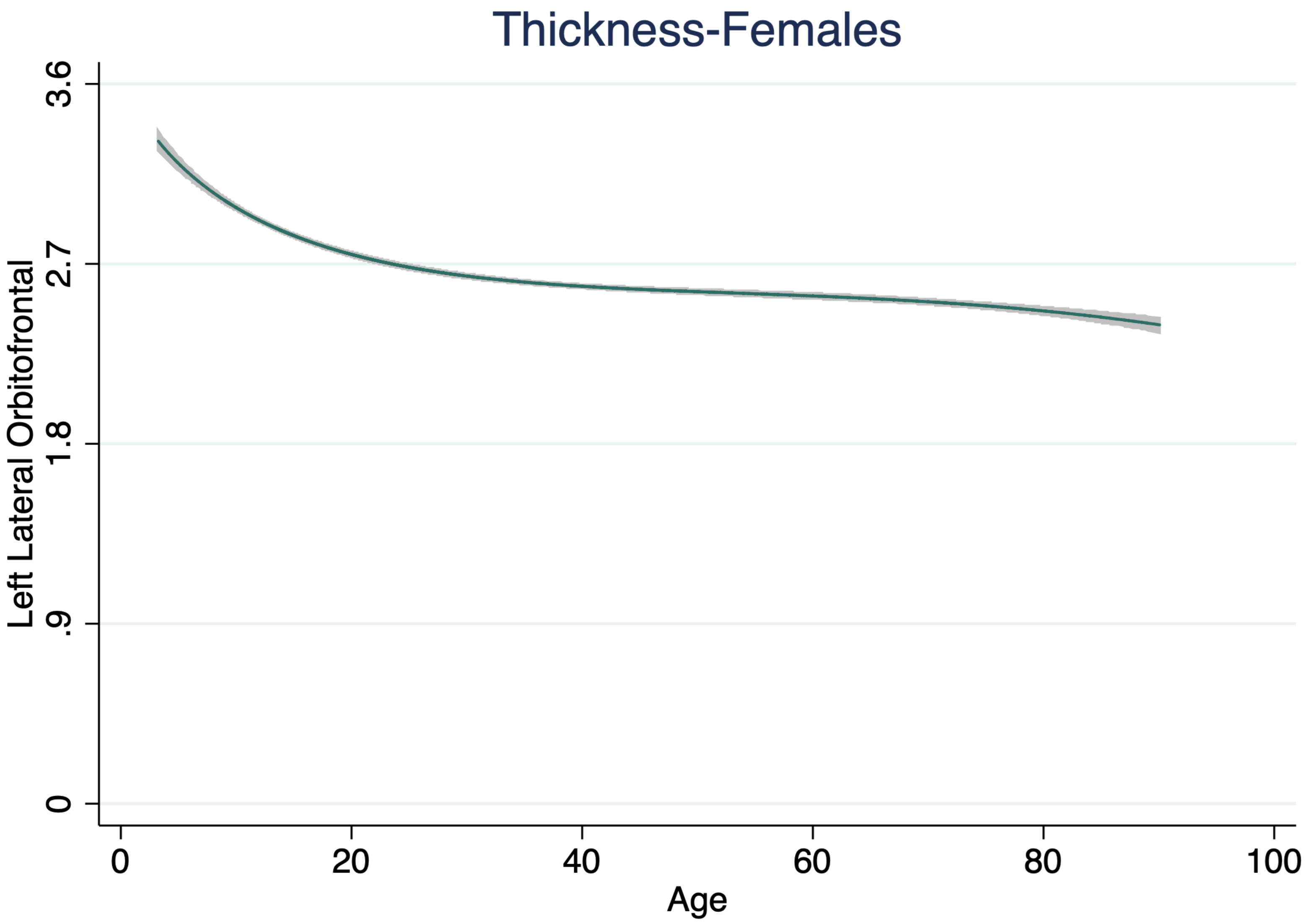
Thickness-Females
Left Lateral Orbitofrontal
3.6
2.7
1.8
.9
0
0
20
40
60
80
100
Age

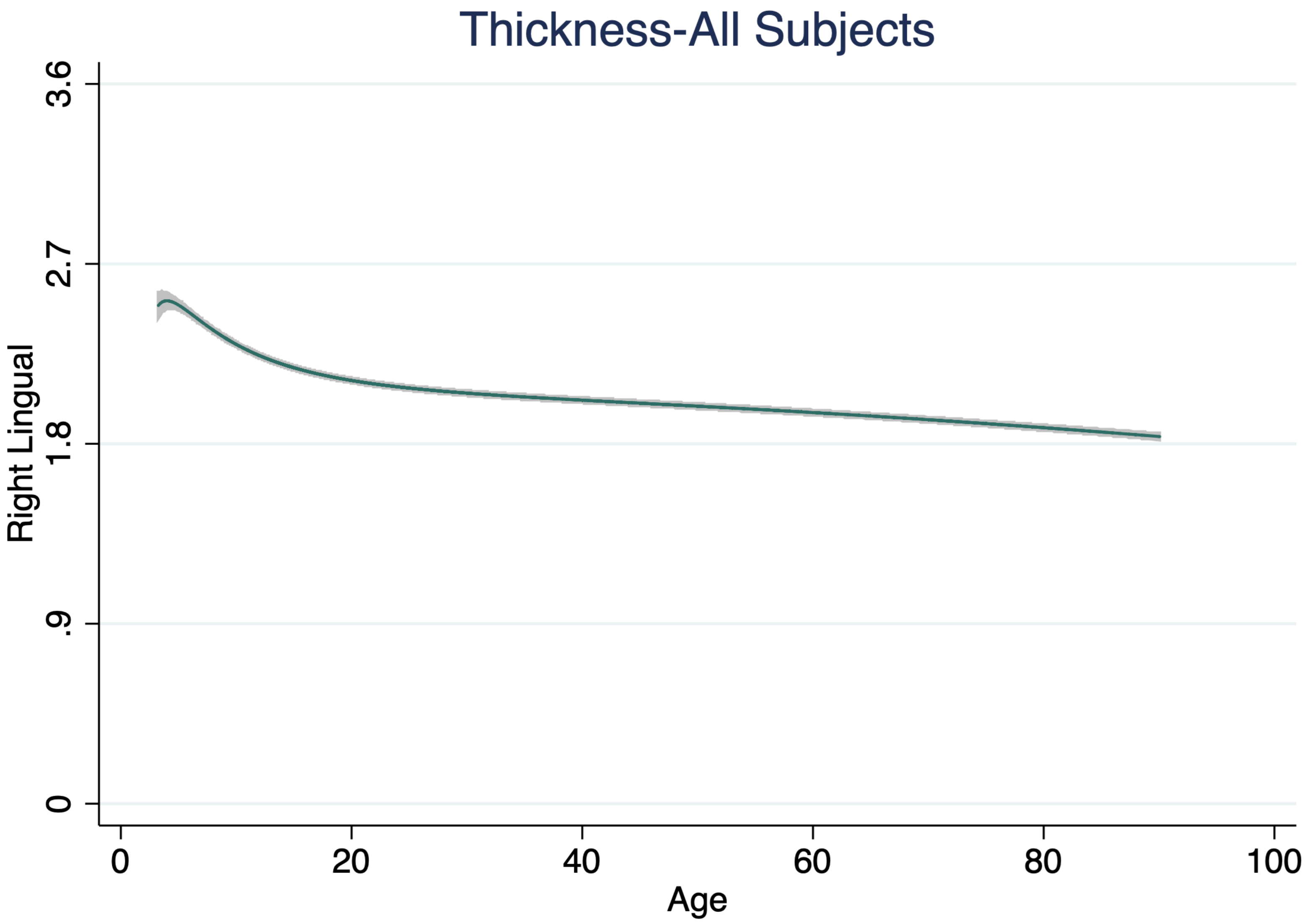
Thickness-All Subjects
Right Lingual
3.6
2.7
1.8
.9
0
0
20
40
60
80
100
Age

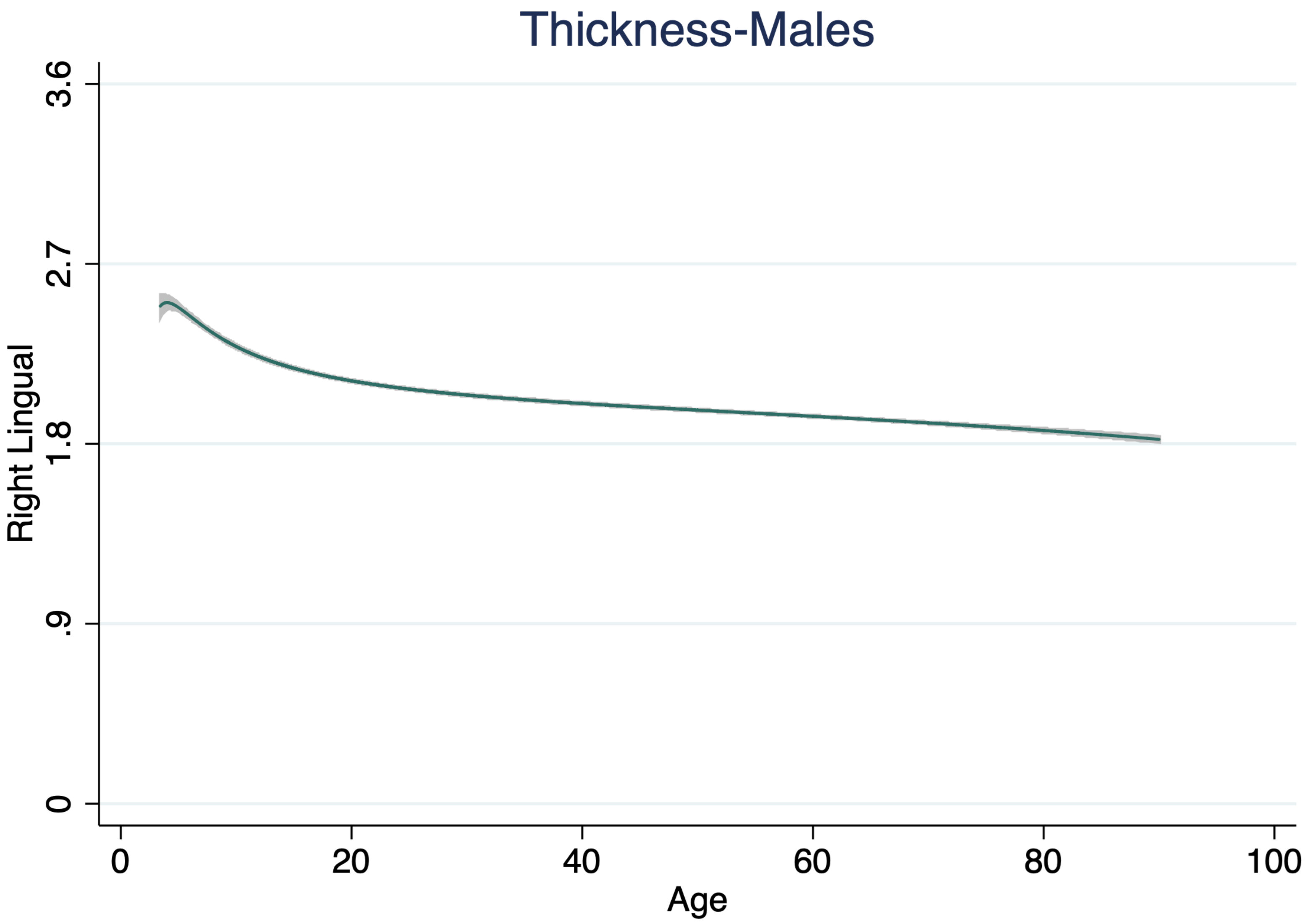

Thickness-Males
Right Lingual
3.6
2.7
1.8
.9
0
0
20
40
60
80
100
Age

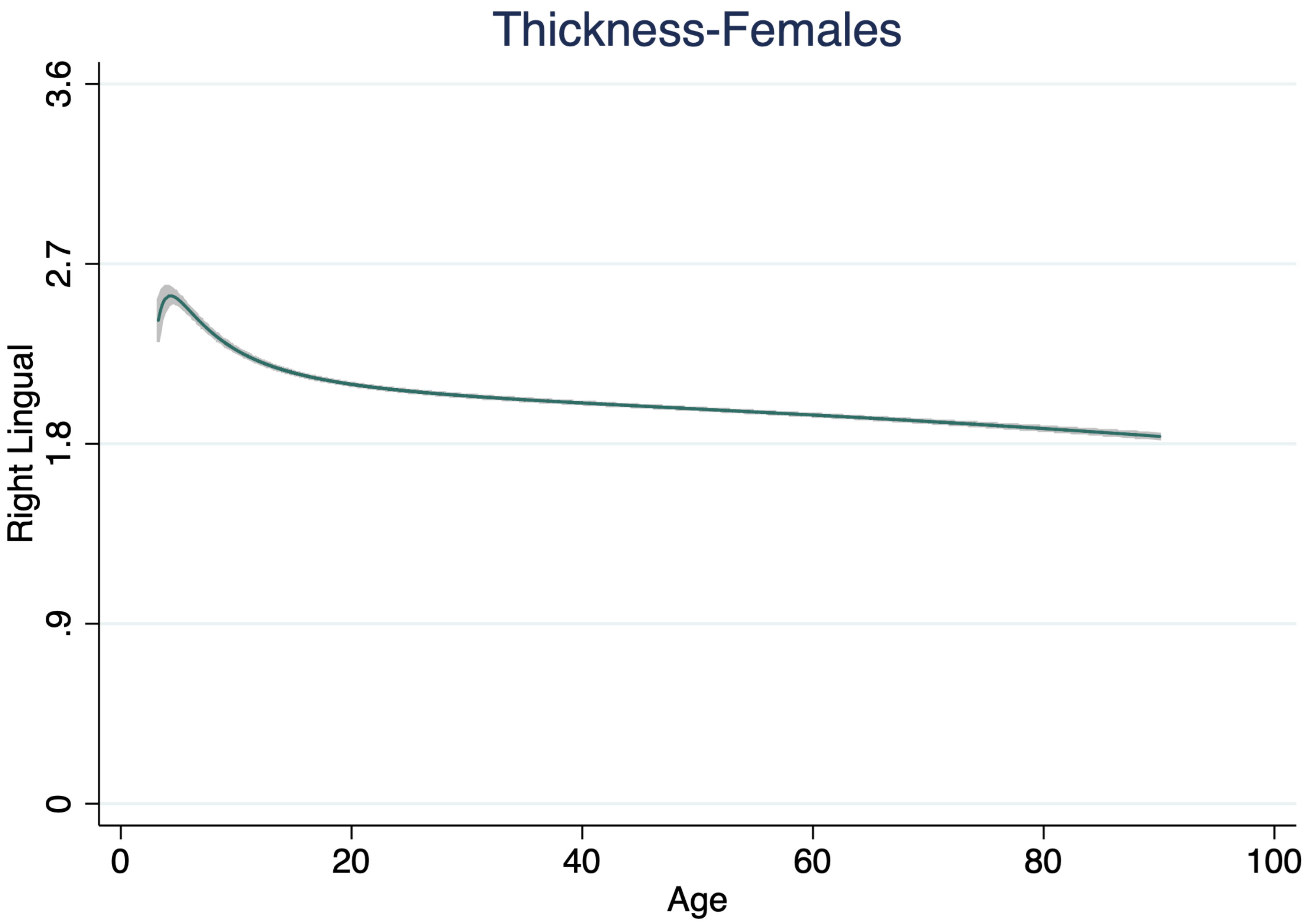

Thickness-Females
3.6
2.7
1.8
.9
0
Right Lingual
0
20
40
60
80
100
Age

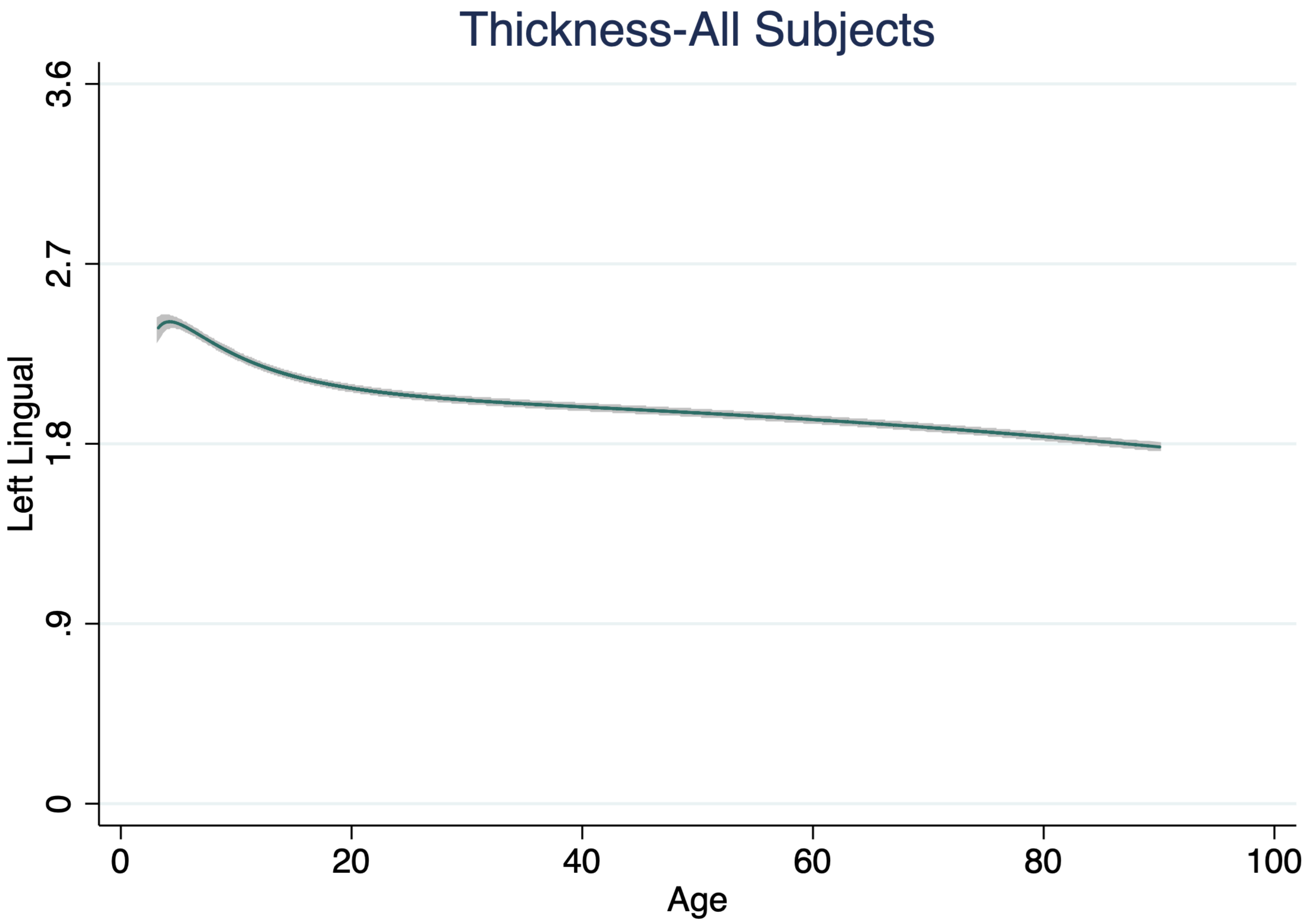

Thickness-All Subjects
Left Lingual
3.6
2.7
1.8
.9
0
0
20
40
60
80
100
Age

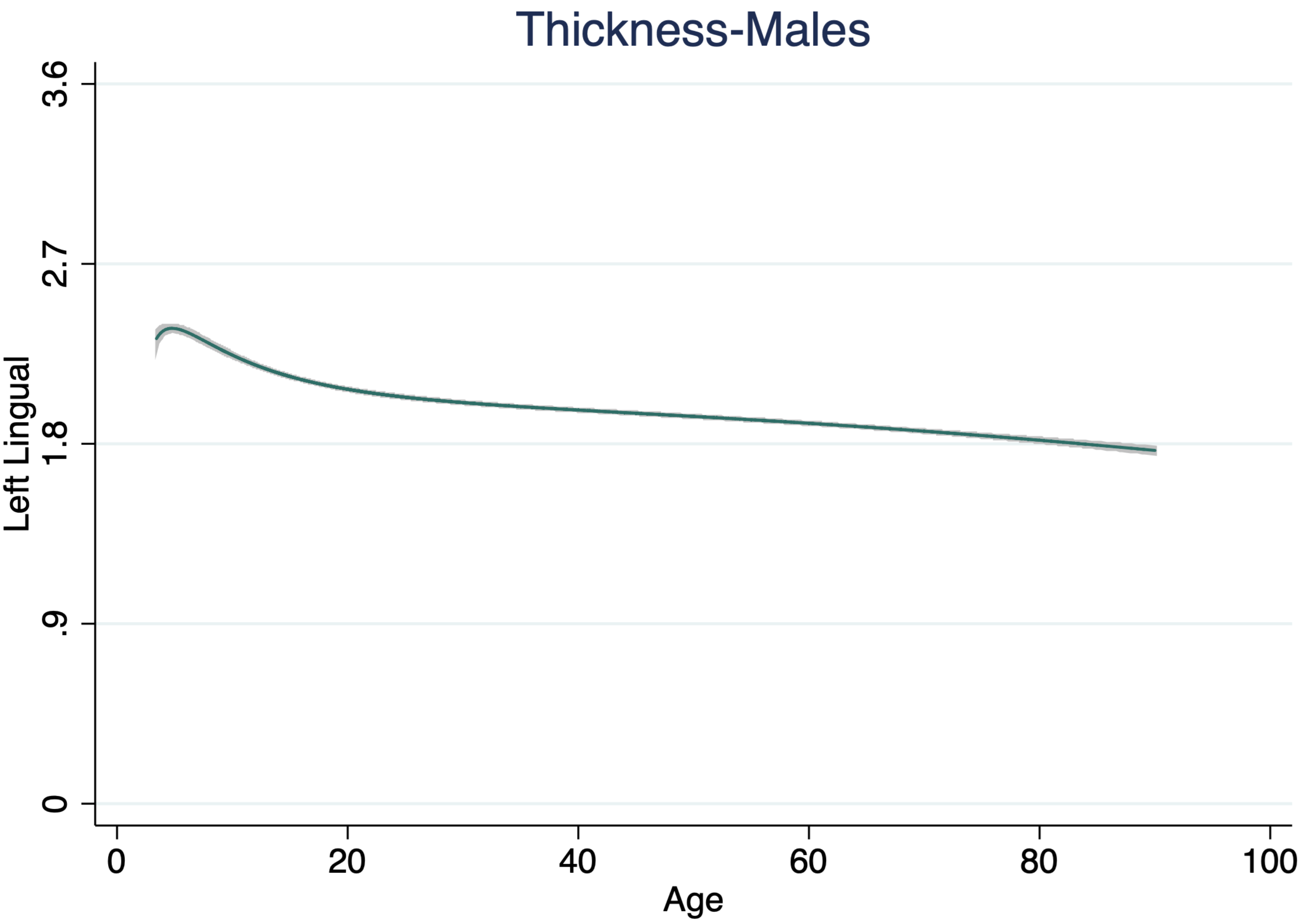

Thickness-Males
Left Lingual
3.6
2.7
1.8
.9
0
0
20
40
60
80
100
Age

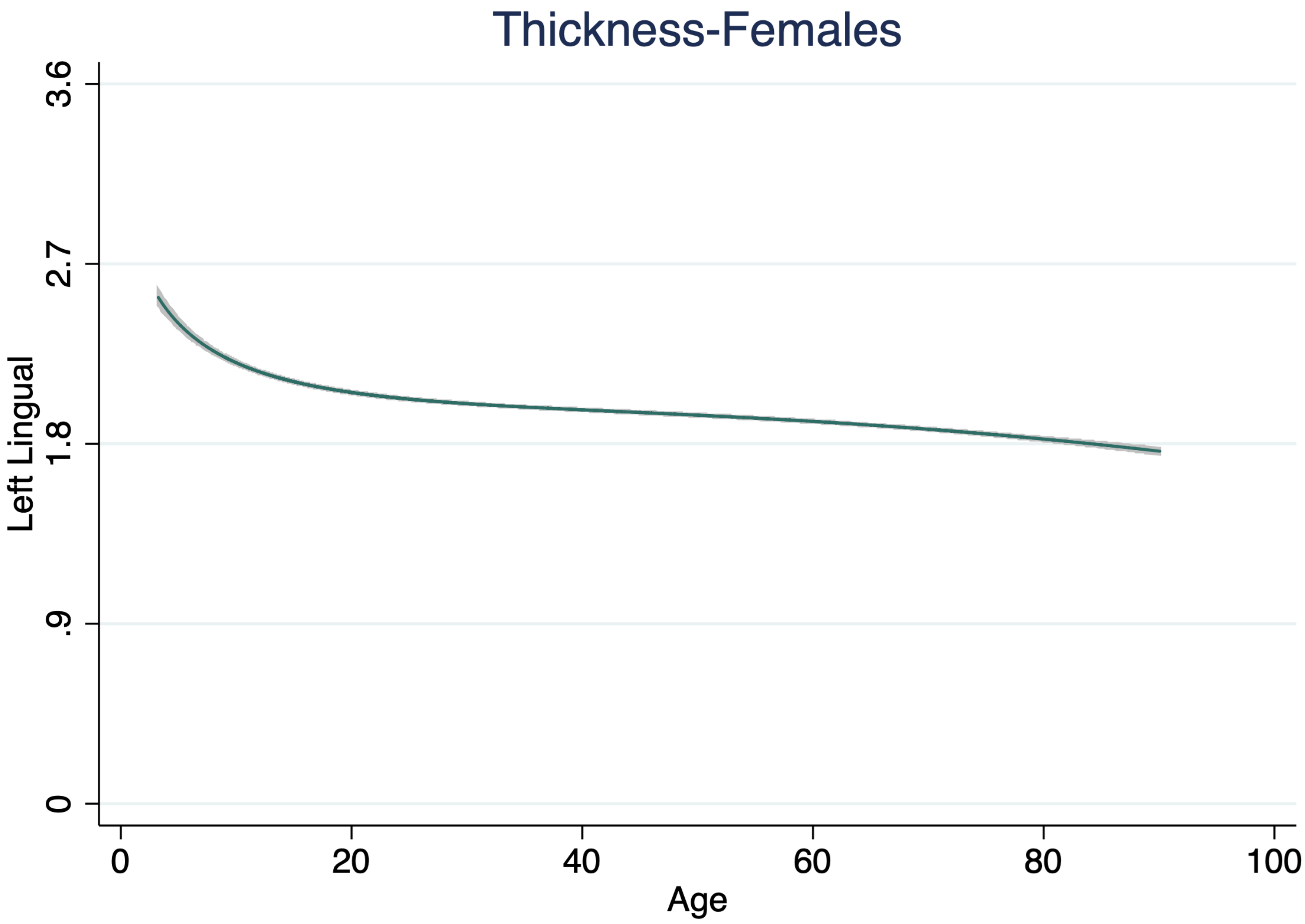
Thickness-Females
Left Lingual
0
.9
1.8
2.7
3.6
0
20
40
60
80
100
Age

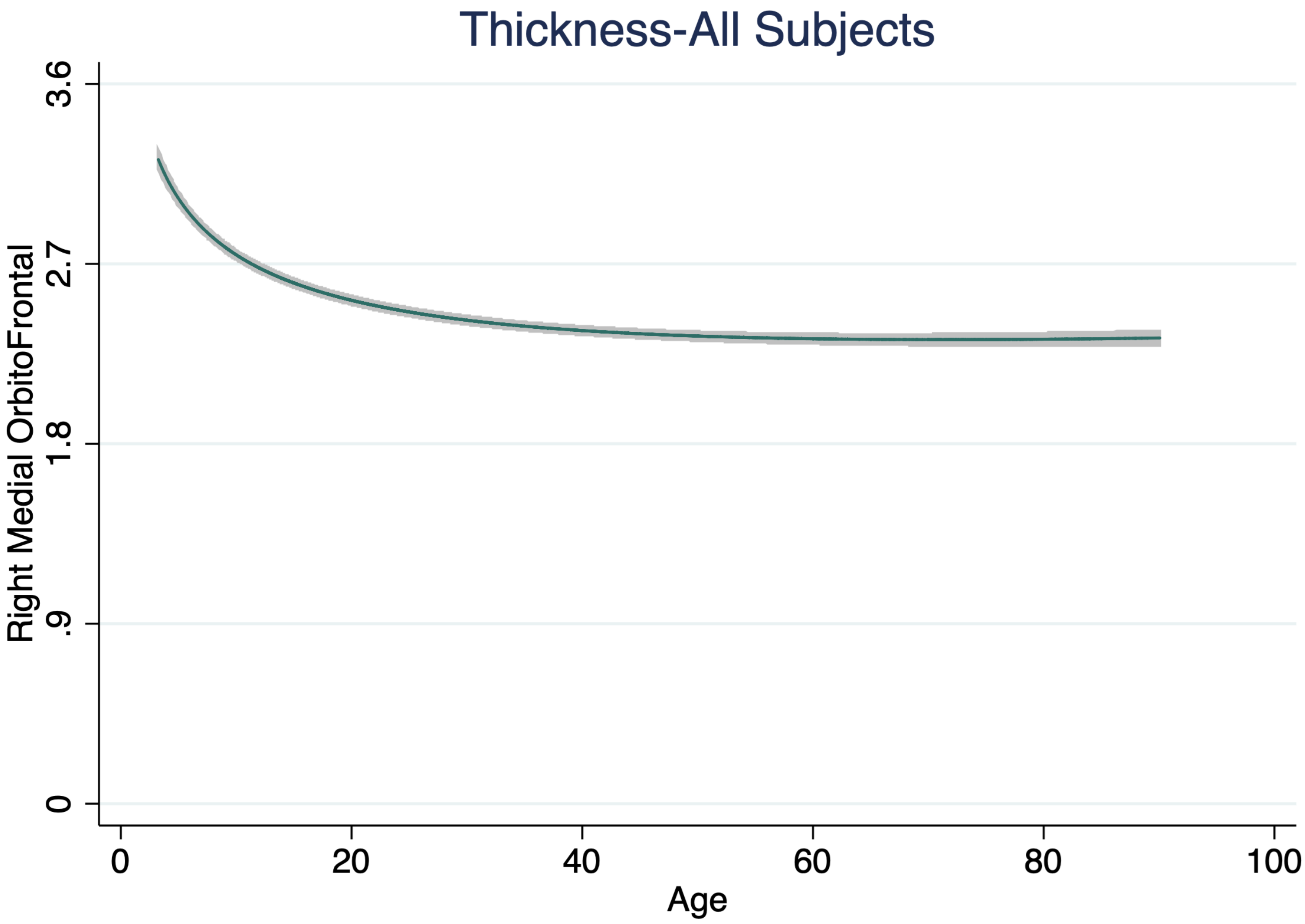

Thickness-All Subjects
Right Medial OrbitoFrontal
3.6
2.7
1.8
.9
0
0
20
40
60
80
100
Age

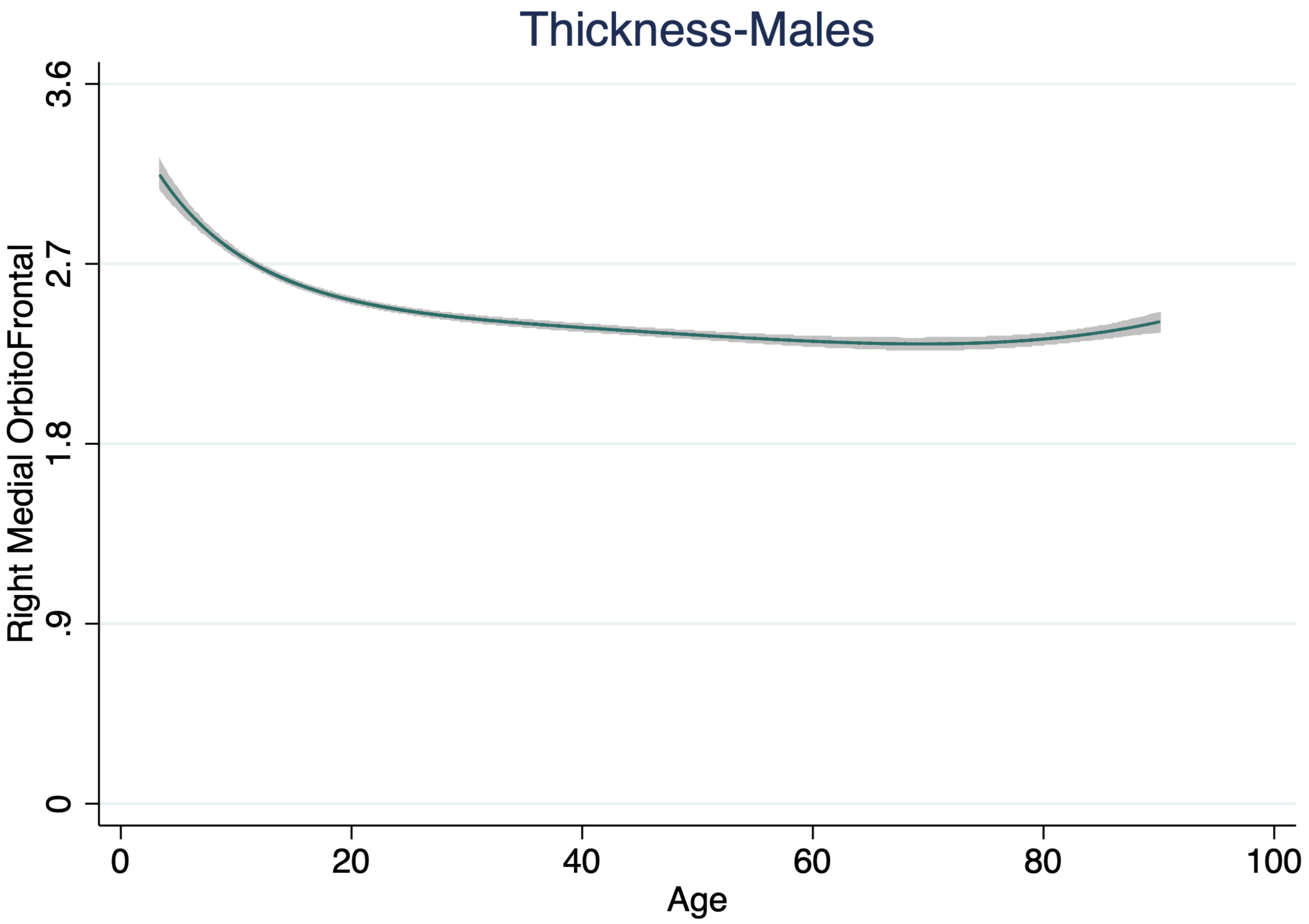

Thickness-Males
Right Medial OrbitoFrontal
3.6
2.7
1.8
.9
0
0
20
40
60
80
100
Age

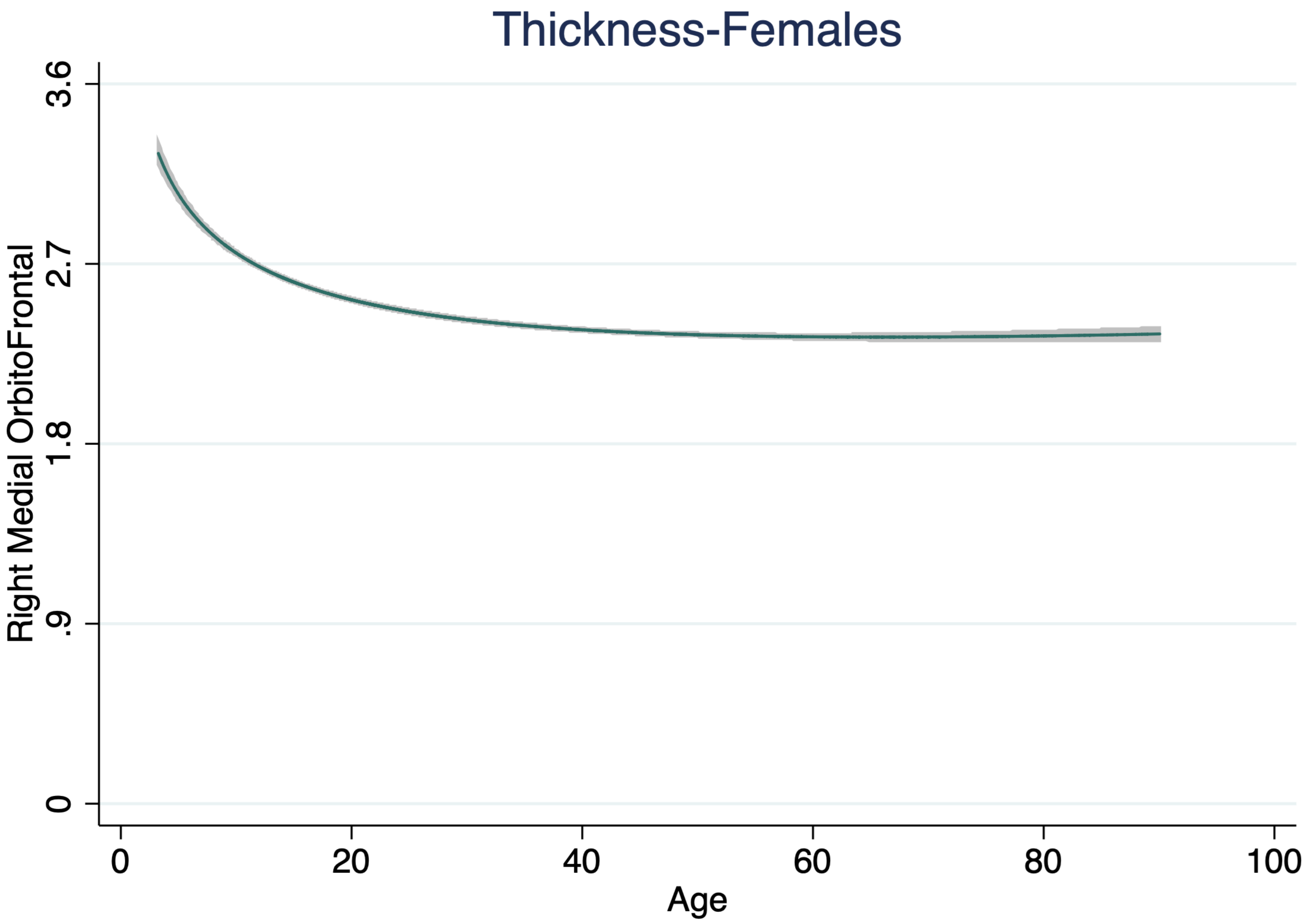

Thickness-Females
Right Medial OrbitoFrontal
3.6
2.7
1.8
.9
0
0
20
40
60
80
100
Age

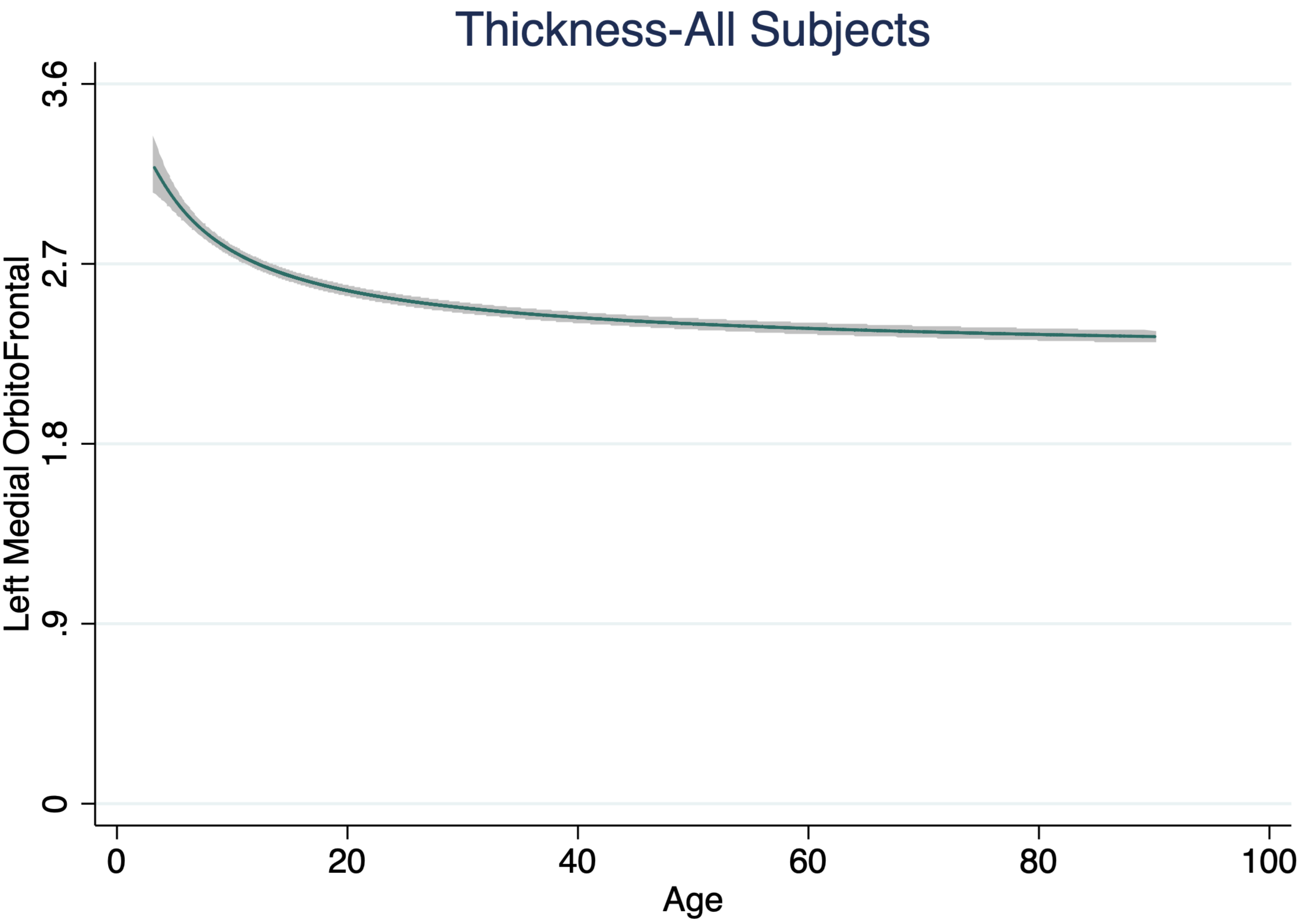
Thickness-All Subjects
Left Medial OrbitoFrontal
3.6
2.7
1.8
.9
0
0
20
40
60
80
100
Age

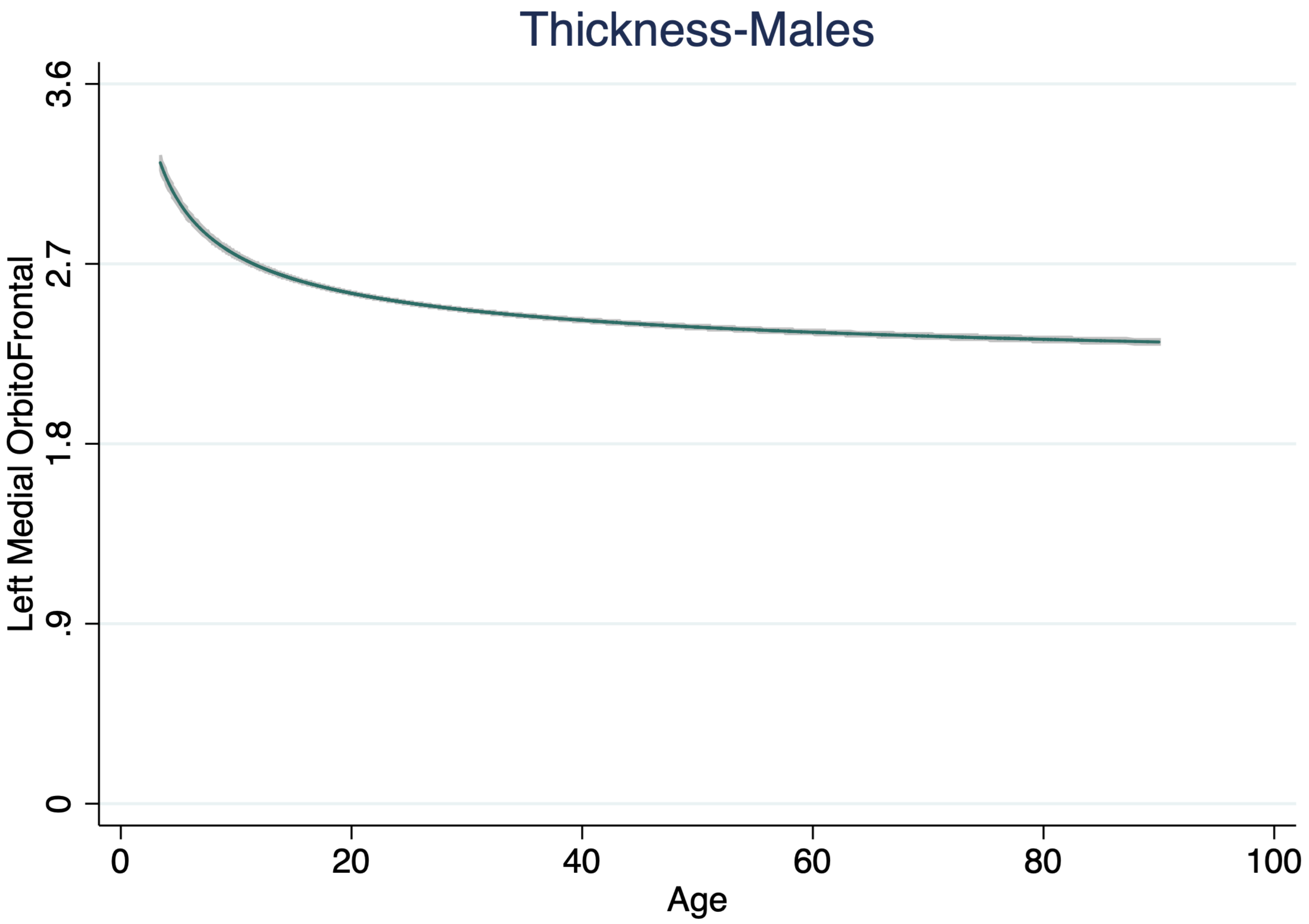
Thickness-Males
Left Medial OrbitoFrontal
3.6
2.7
1.8
.9
0
0
20
40
60
80
100
Age

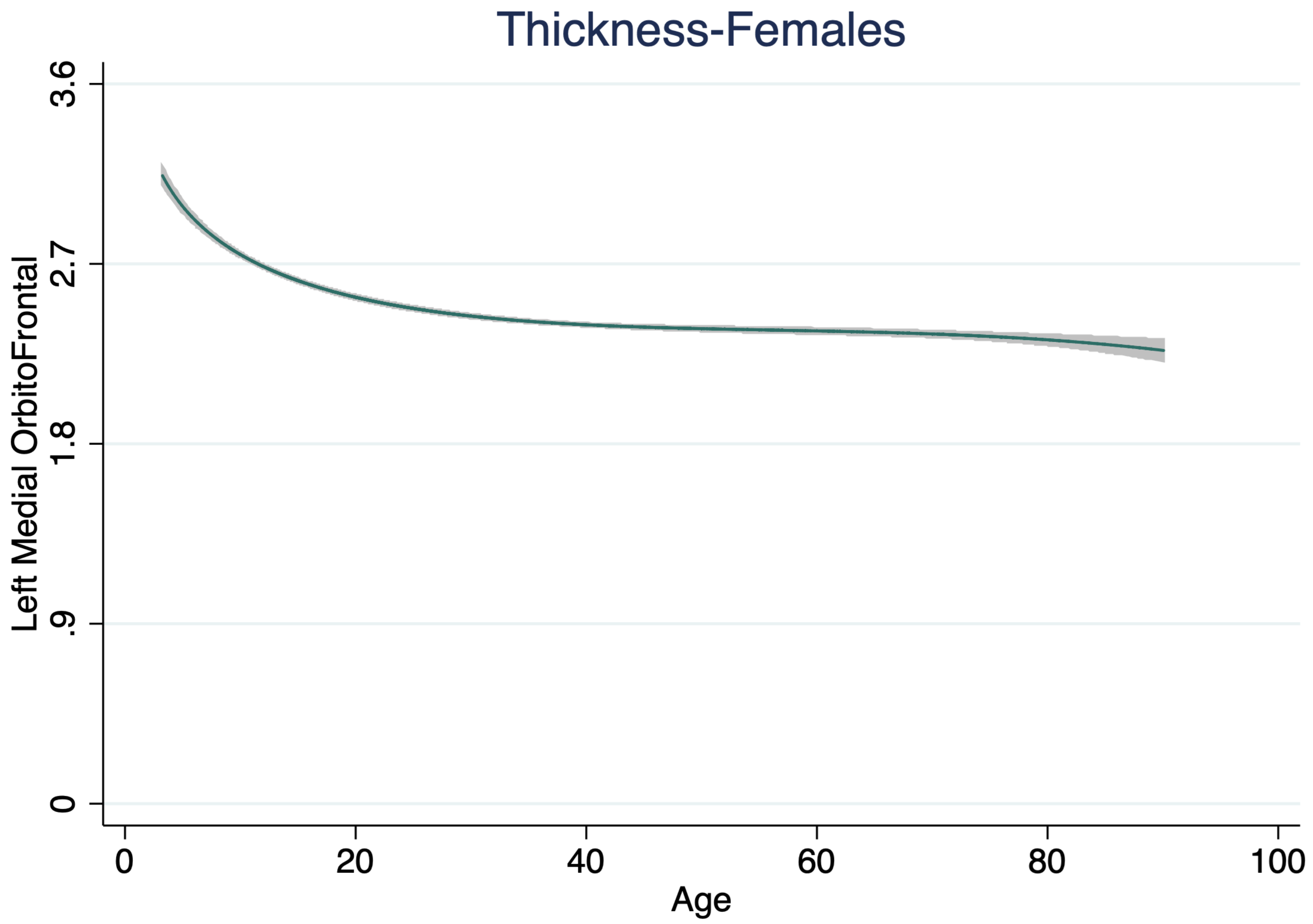

Thickness-Females
Left Medial OrbitoFrontal
3.6
2.7
1.8
.9
0
0
20
40
60
80
100
Age

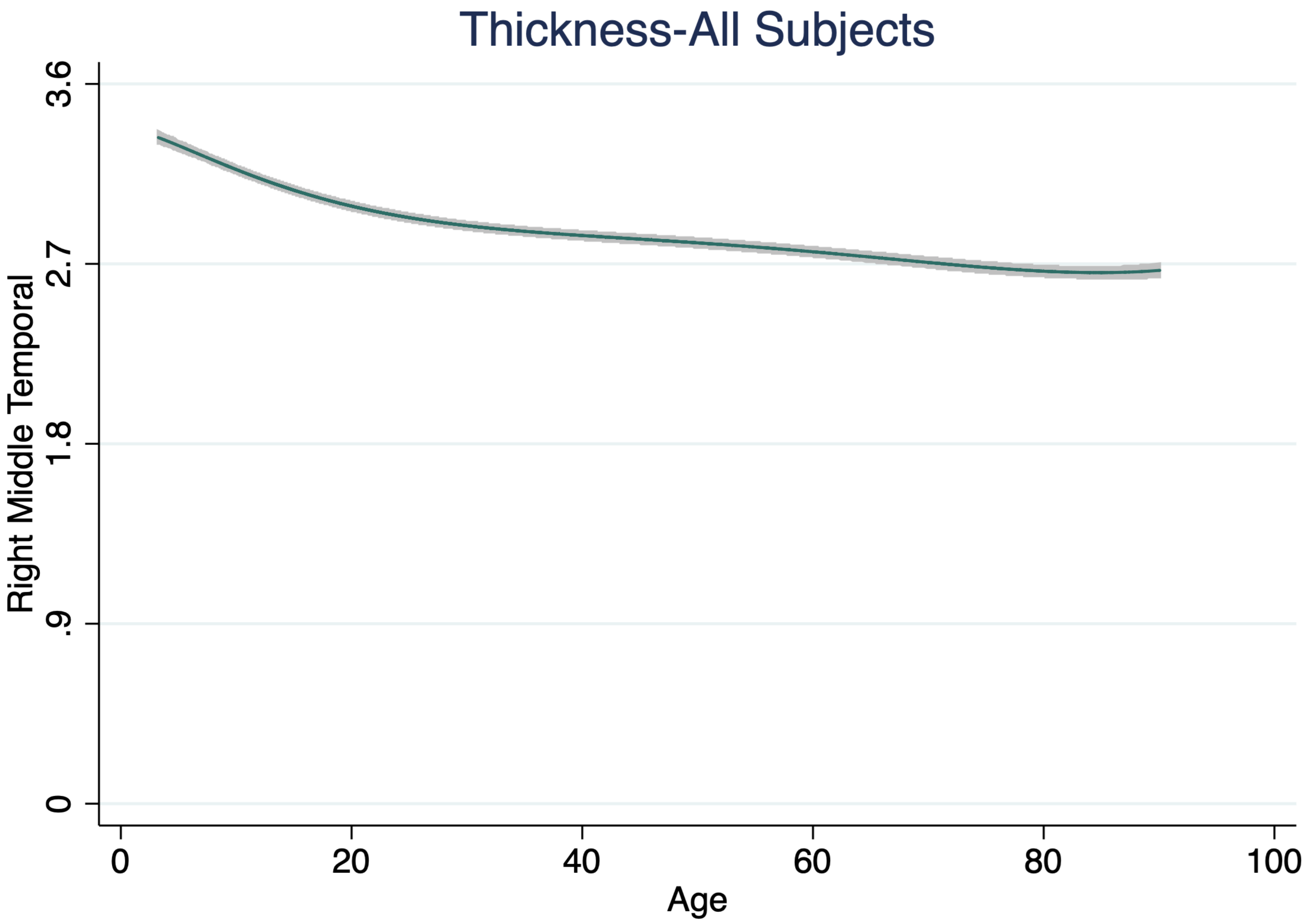

Thickness-All Subjects
Right Middle Temporal
3.6
2.7
1.8
.9
0
0
20
40
60
80
100
Age

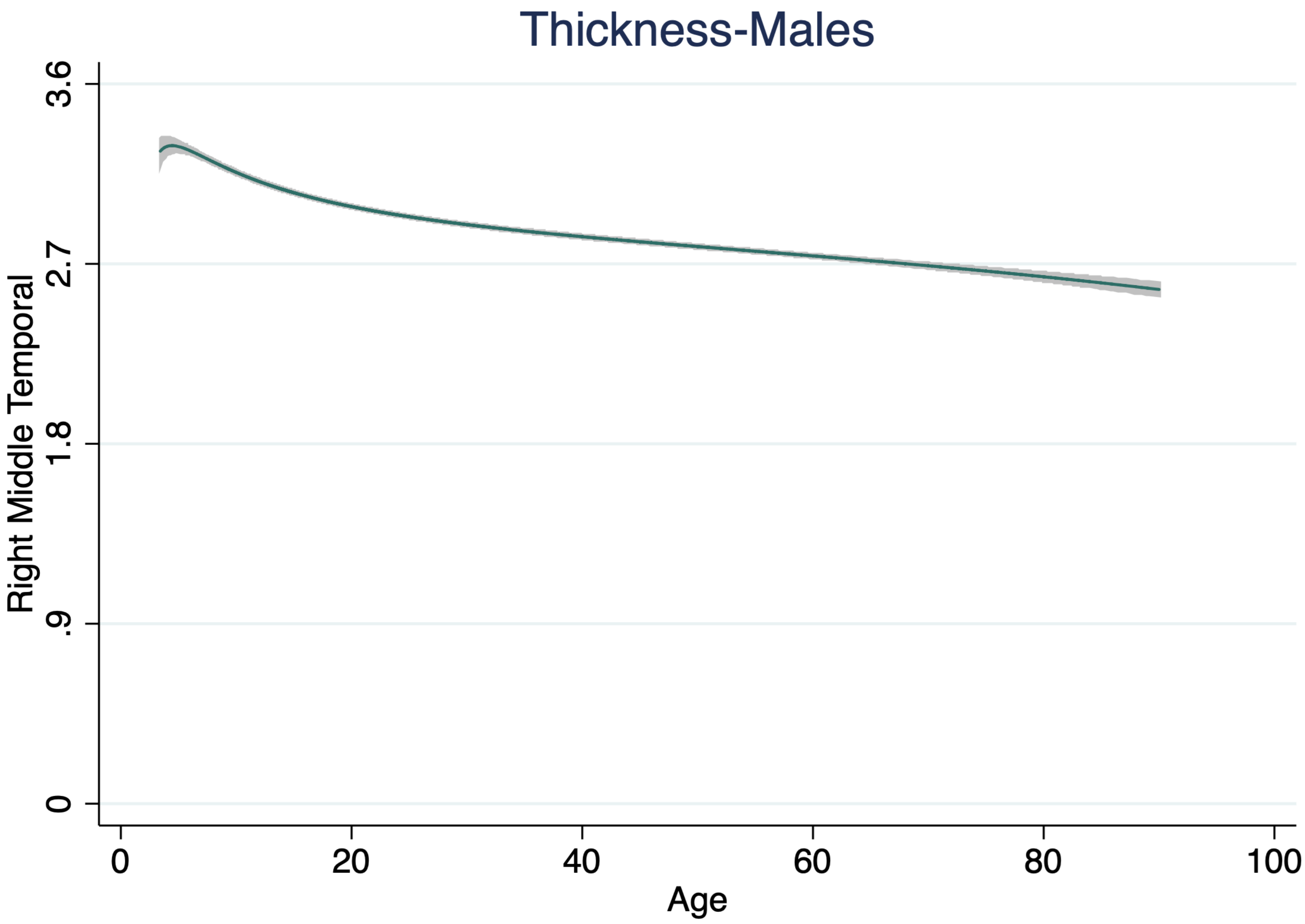
Thickness-Males
Right Middle Temporal
0
.9
1.8
2.7
3.6
0
20
40
60
80
100
Age

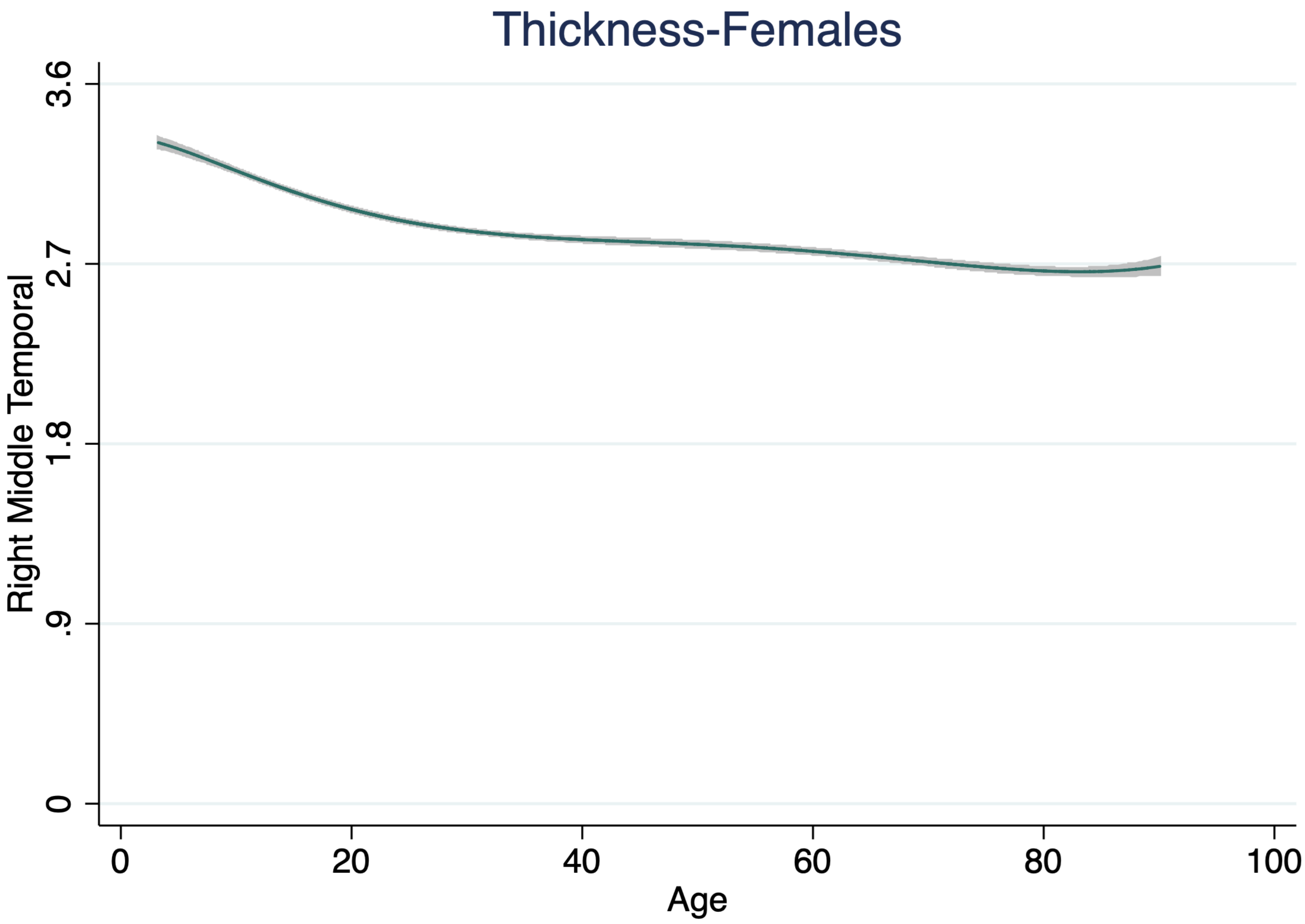

Thickness-Females
Right Middle Temporal
3.6
2.7
1.8
.9
0
0
20
40
60
80
100
Age

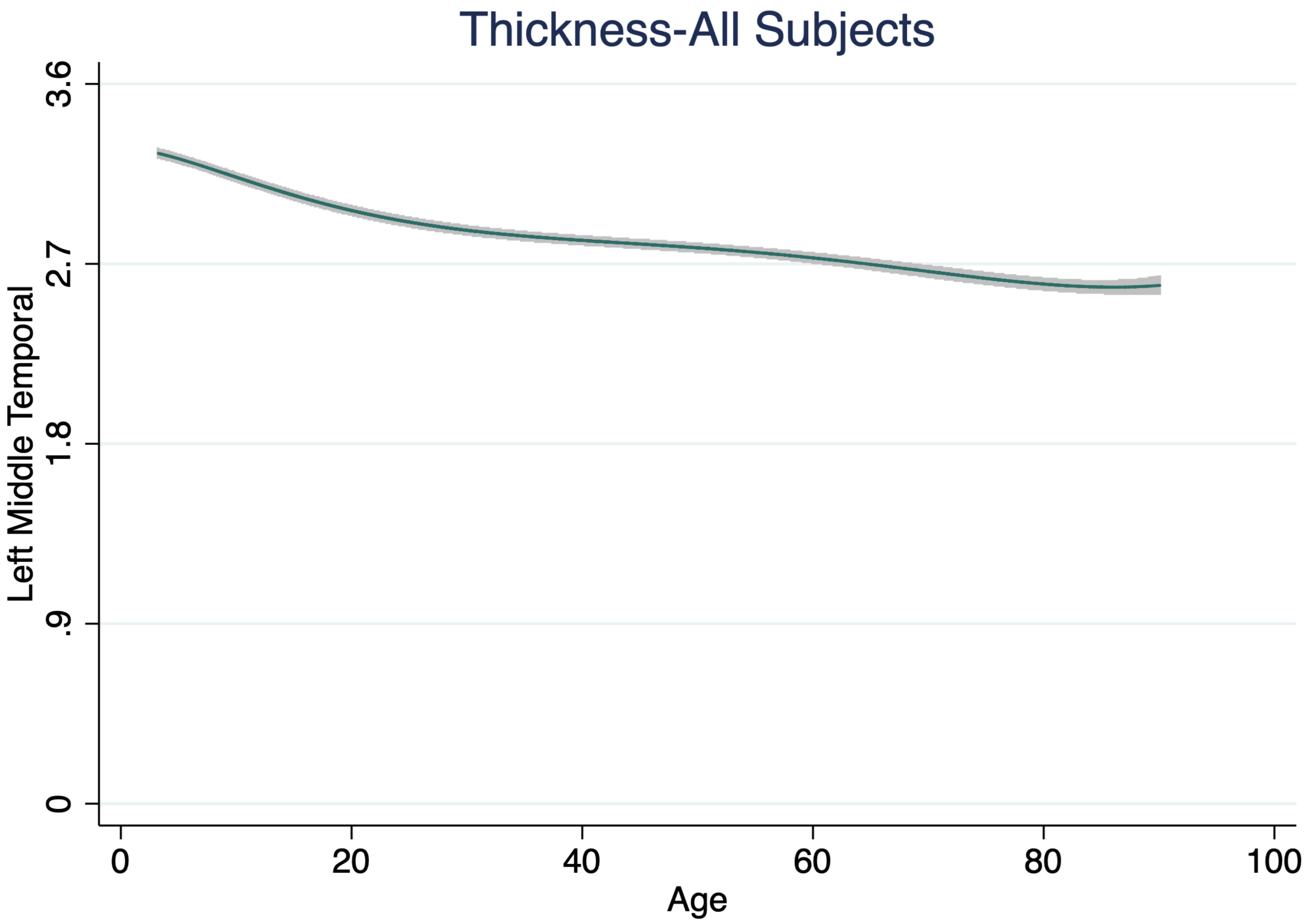

Thickness-All Subjects
Left Middle Temporal
3.6
2.7
1.8
.9
0
0
20
40
60
80
100
Age

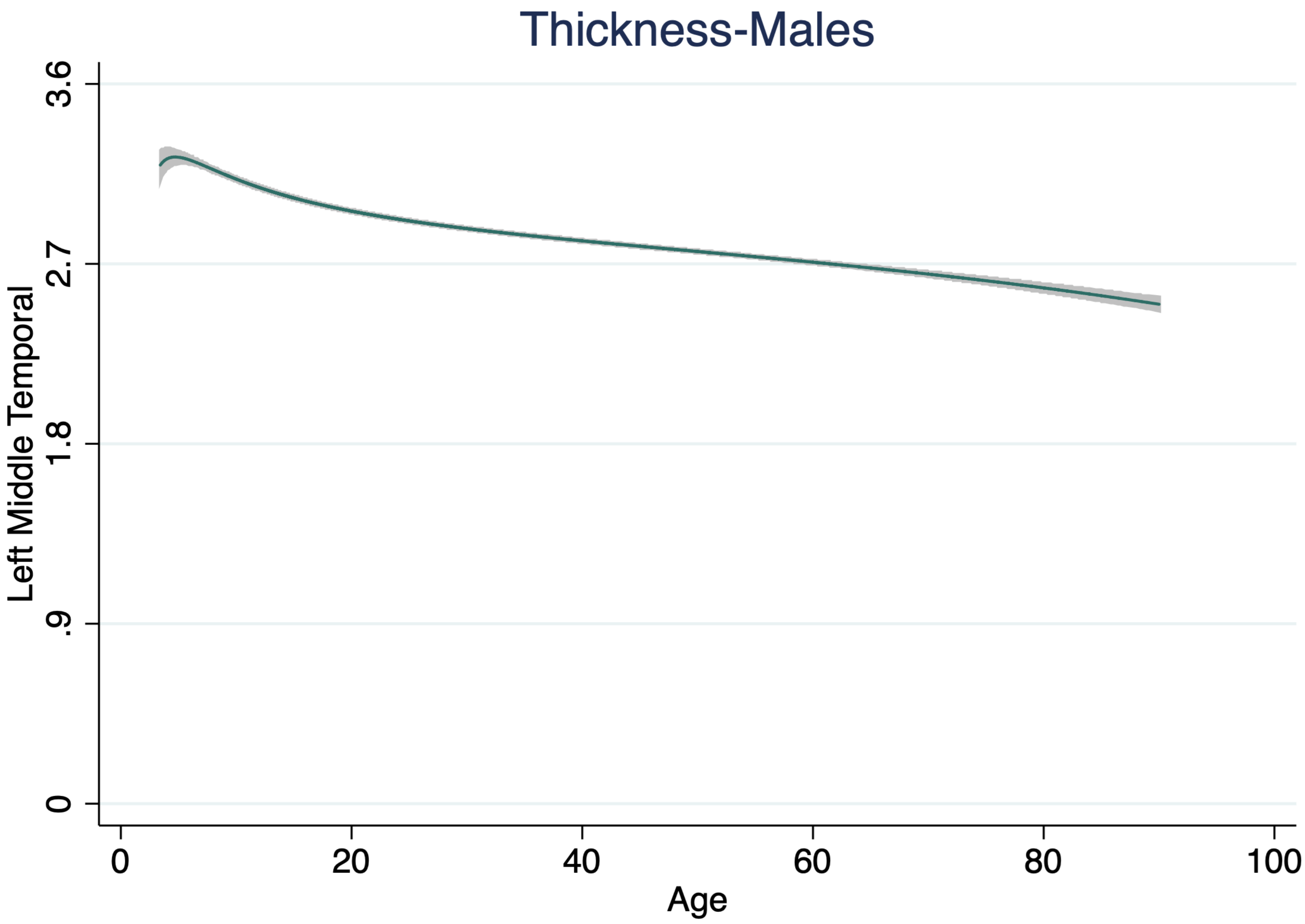

Thickness-Males
Left Middle Temporal
0
.9
1.8
2.7
3.6
0
20
40
60
80
100
Age

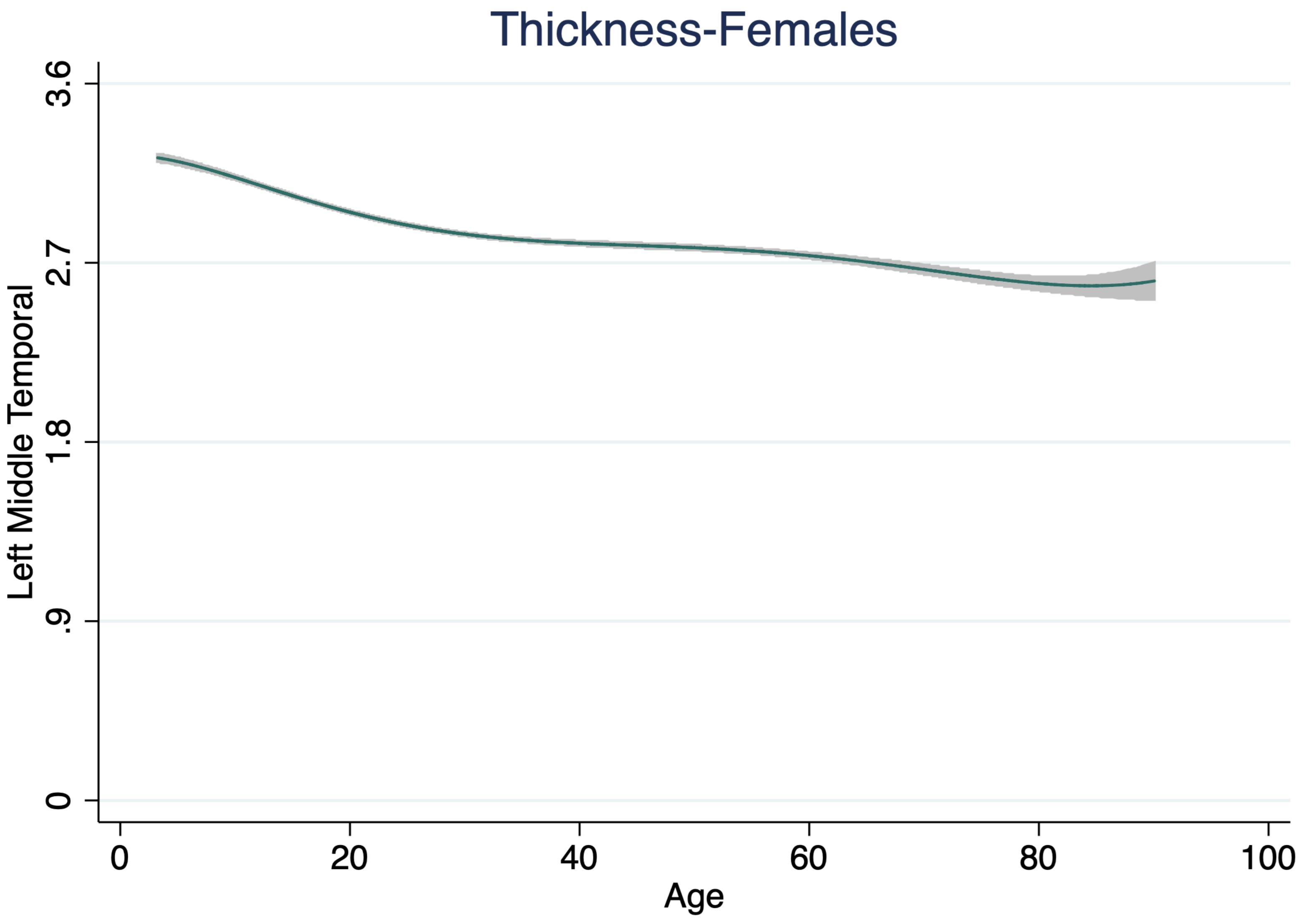
Thickness-Females
Left Middle Temporal
3.6
2.7
1.8
.9
0
0
20
40
60
80
100
Age

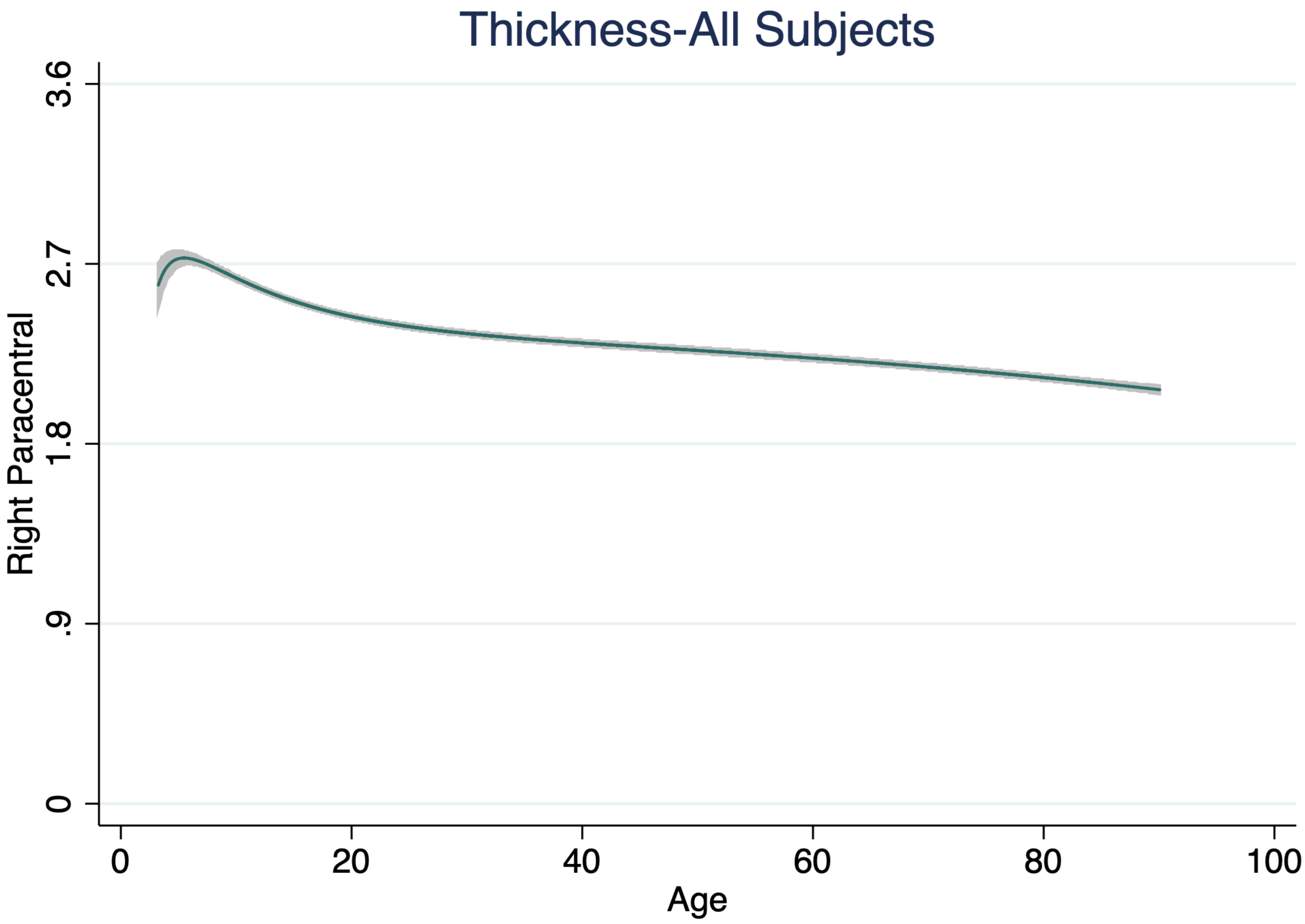

Thickness-All Subjects
Right Paracentral
3.6
2.7
1.8
.9
0
0
20
40
60
80
100
Age

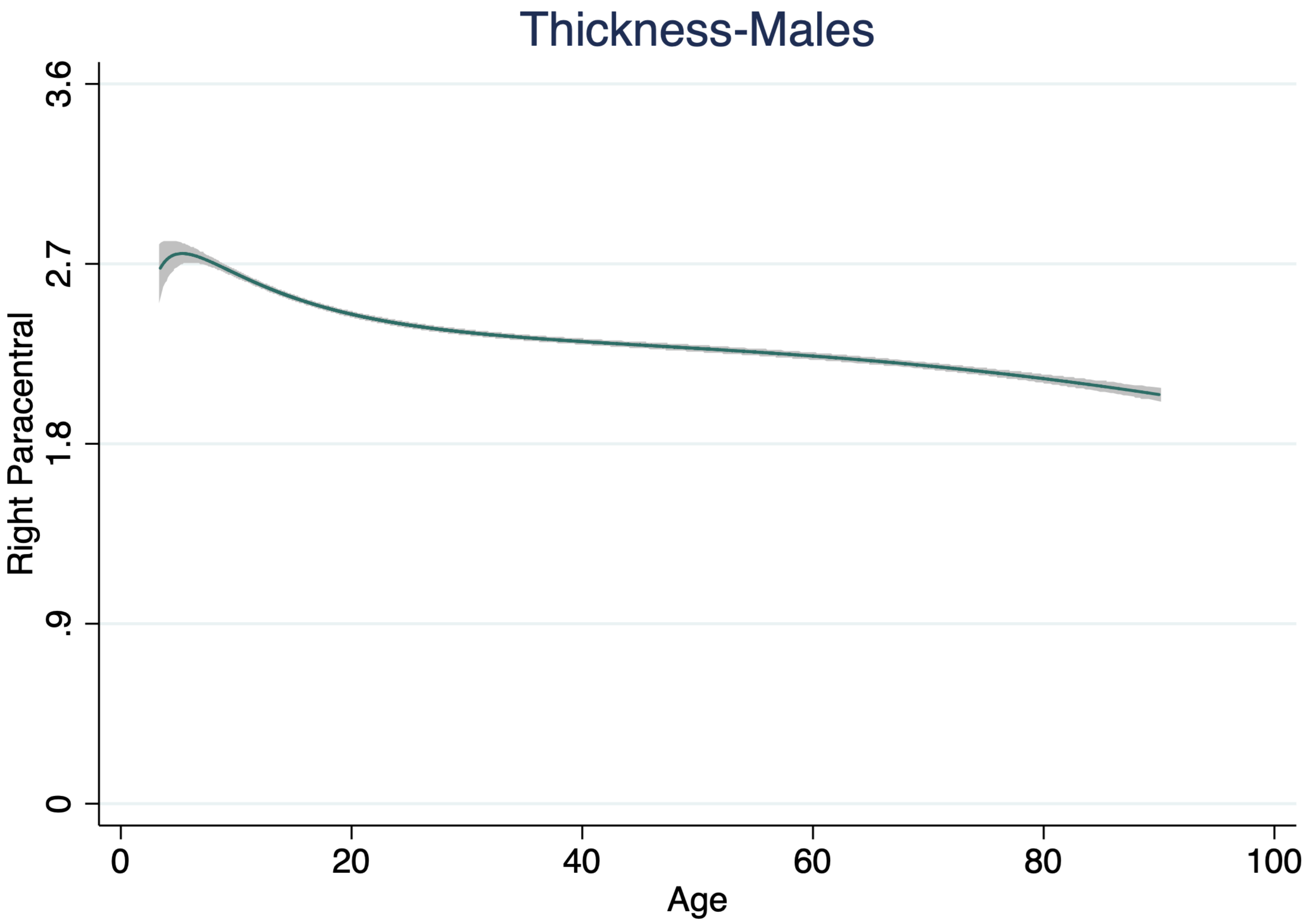

Thickness-Males
Right Paracentral
3.6
2.7
1.8
.9
0
0
20
40
60
80
100
Age

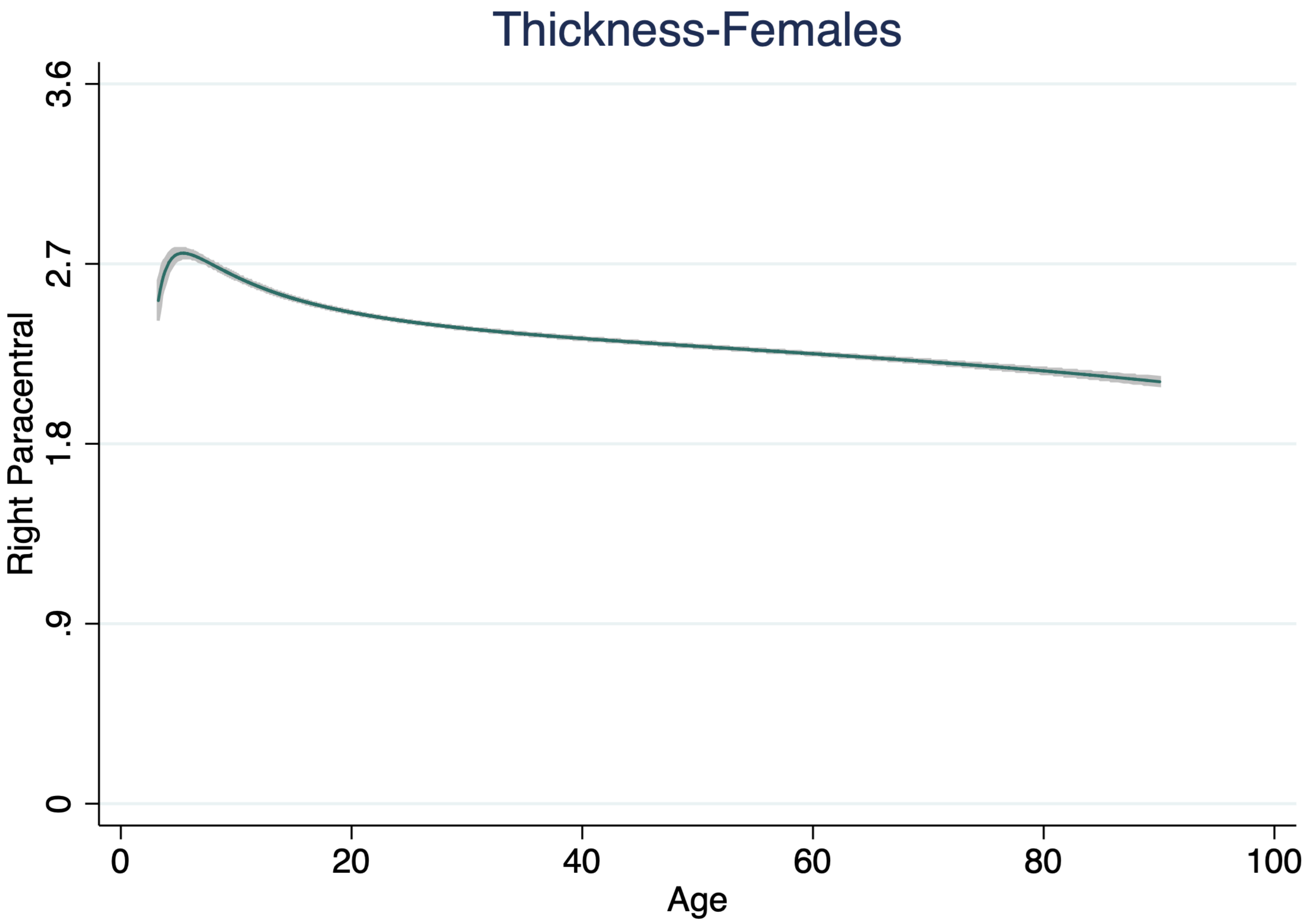

Thickness-Females
Right Paracentral
3.6
2.7
1.8
.9
0
0
20
40
60
80
100
Age

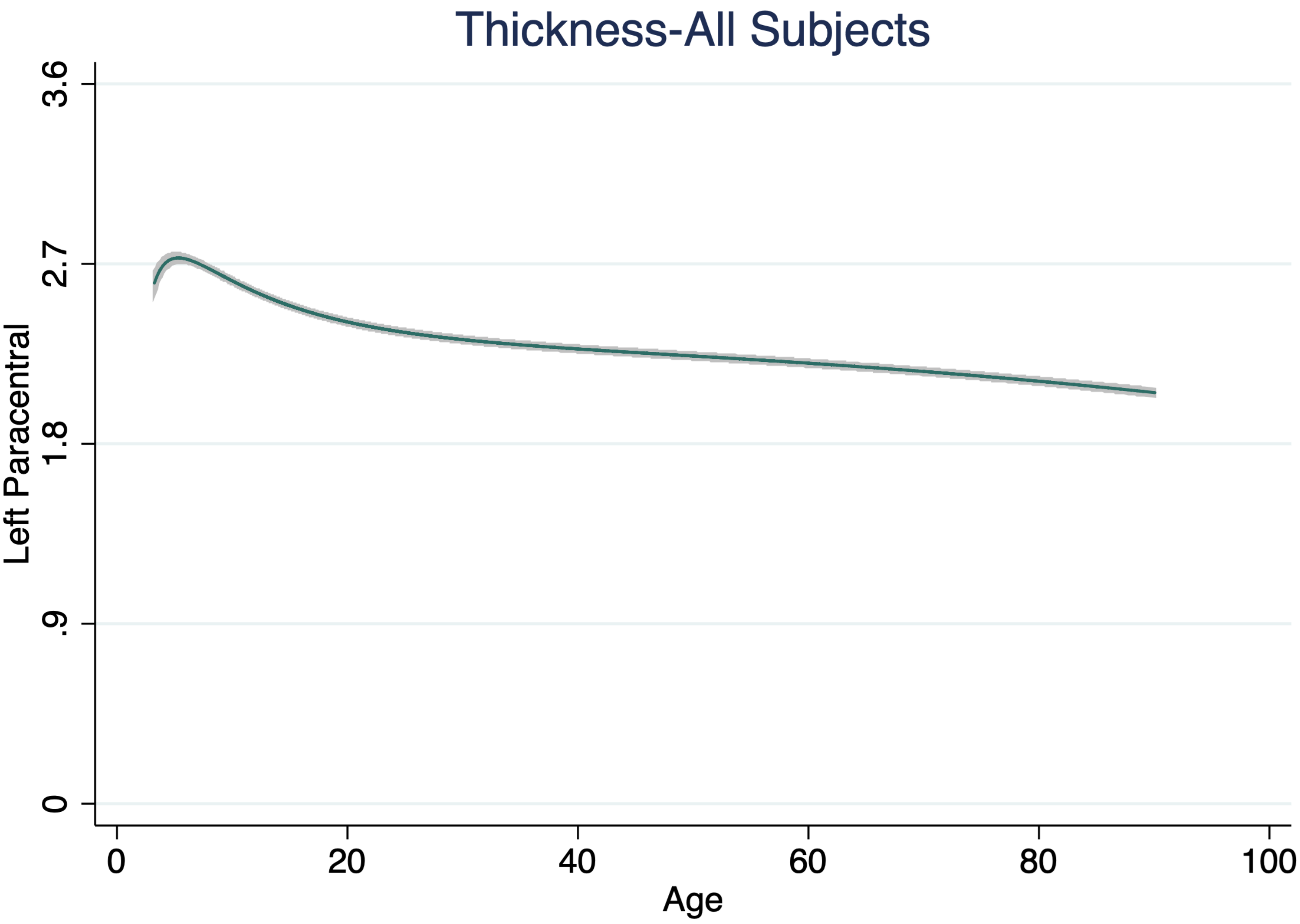

Thickness-All Subjects
Left Paracentral
3.6
2.7
1.8
.9
0
0
20
40
60
80
100
Age

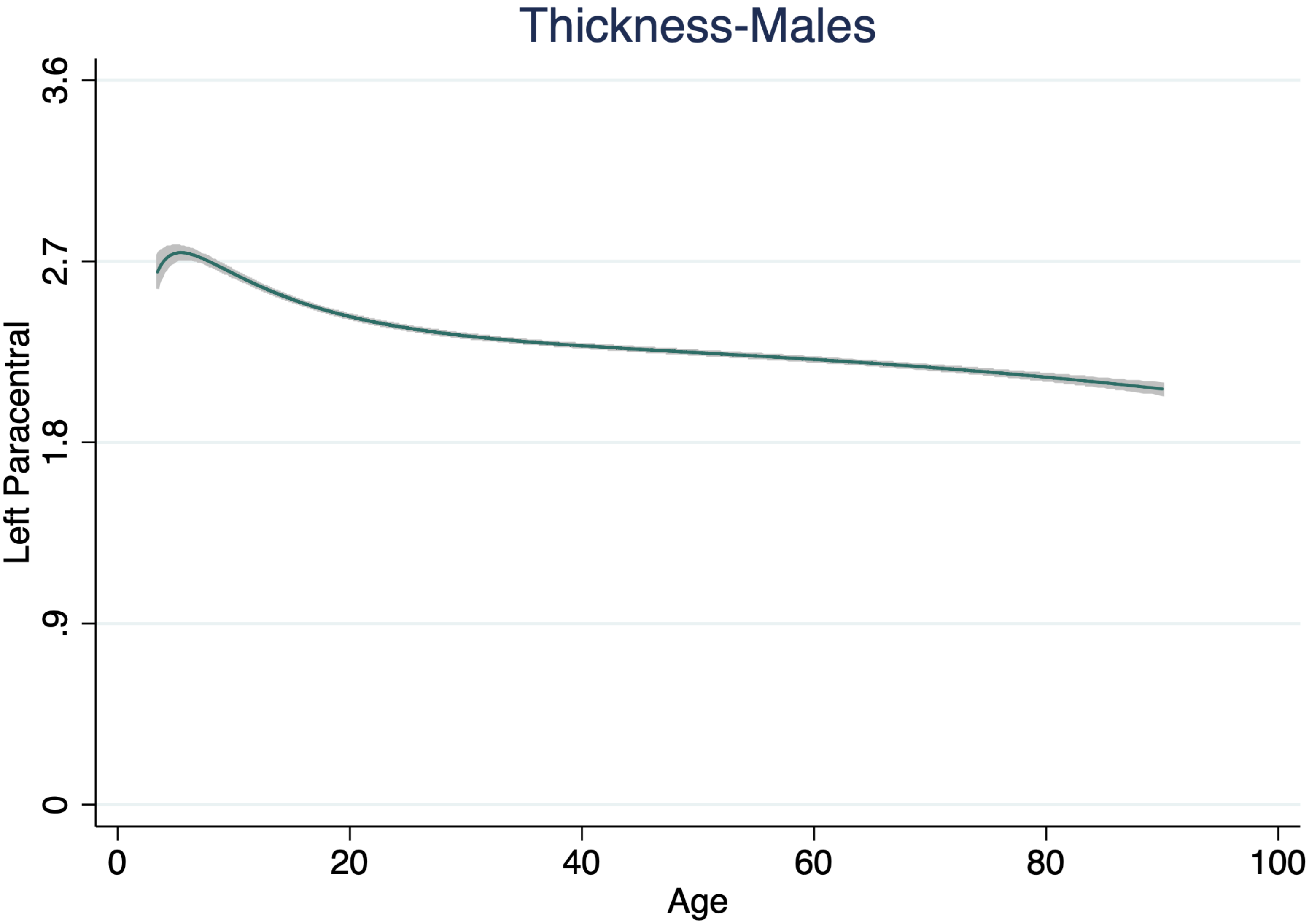
Thickness-Males
Left Paracentral
3.6
2.7
1.8
.9
0
0
20
40
60
80
100
Age

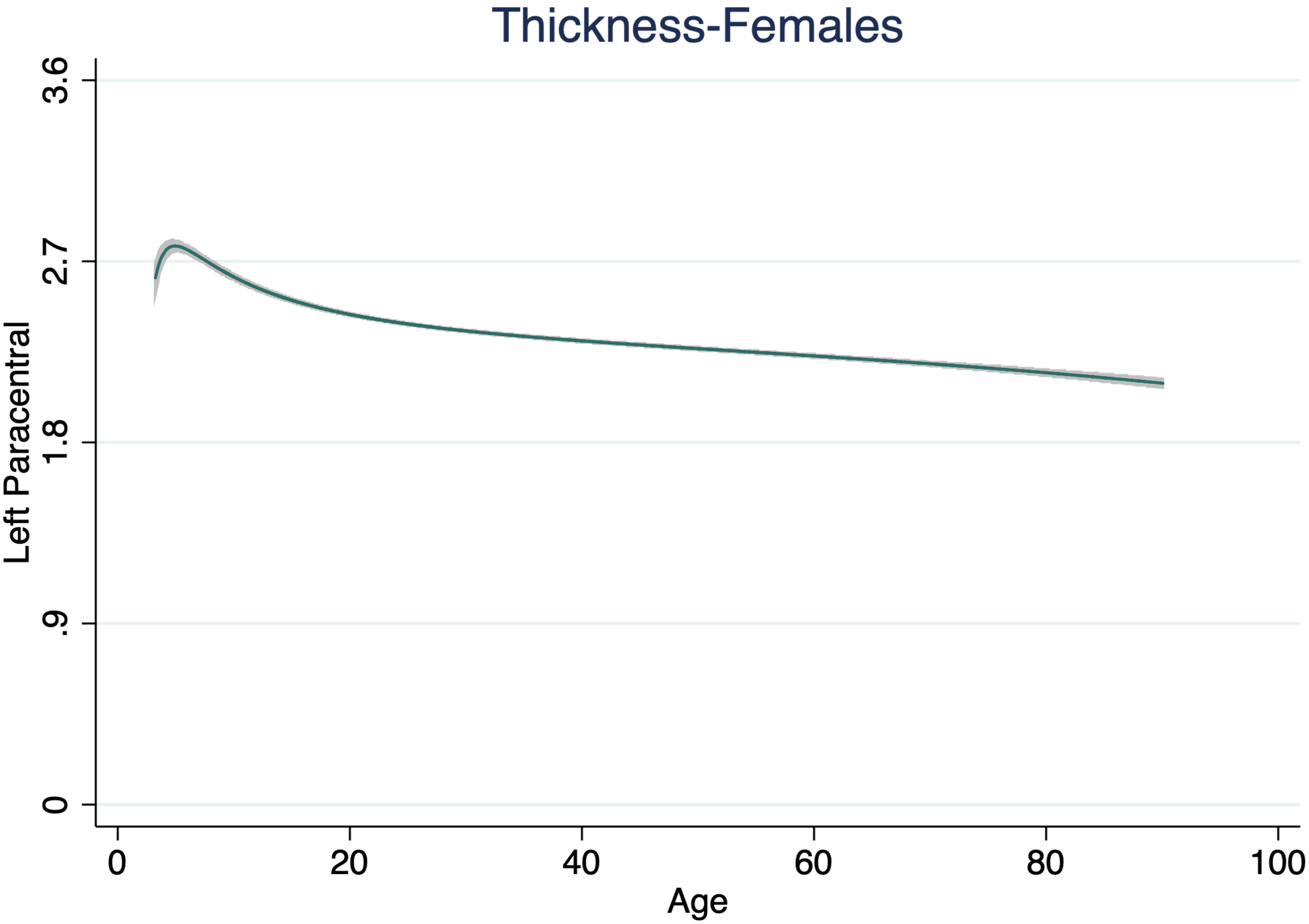

Thickness-Females
Left Paracentral
3.6
2.7
1.8
.9
0
0
20
40
60
80
100
Age

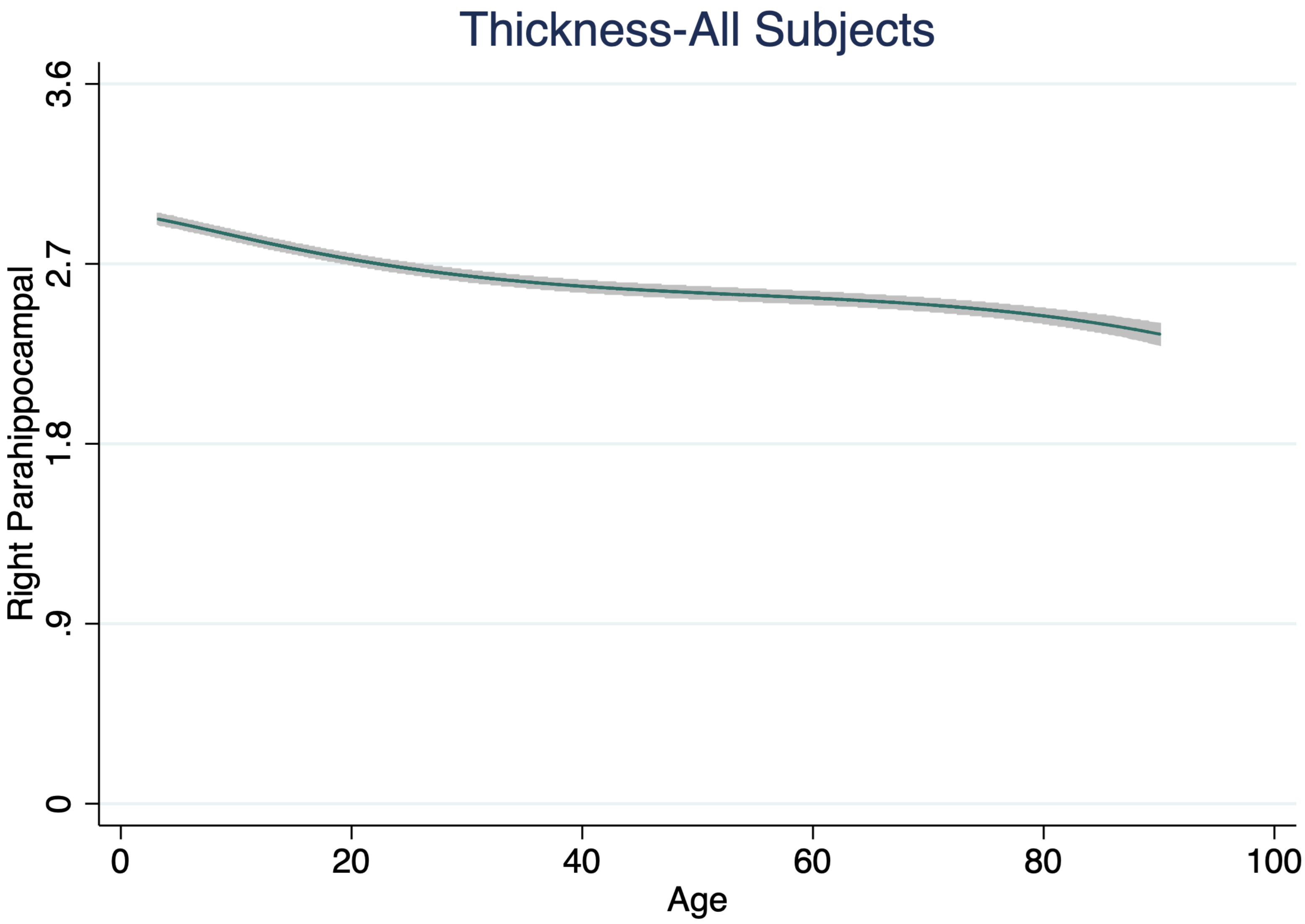
Thickness-All Subjects
Right Parahippocampal
3.6
2.7
1.8
.9
0
0
20
40
60
80
100
Age

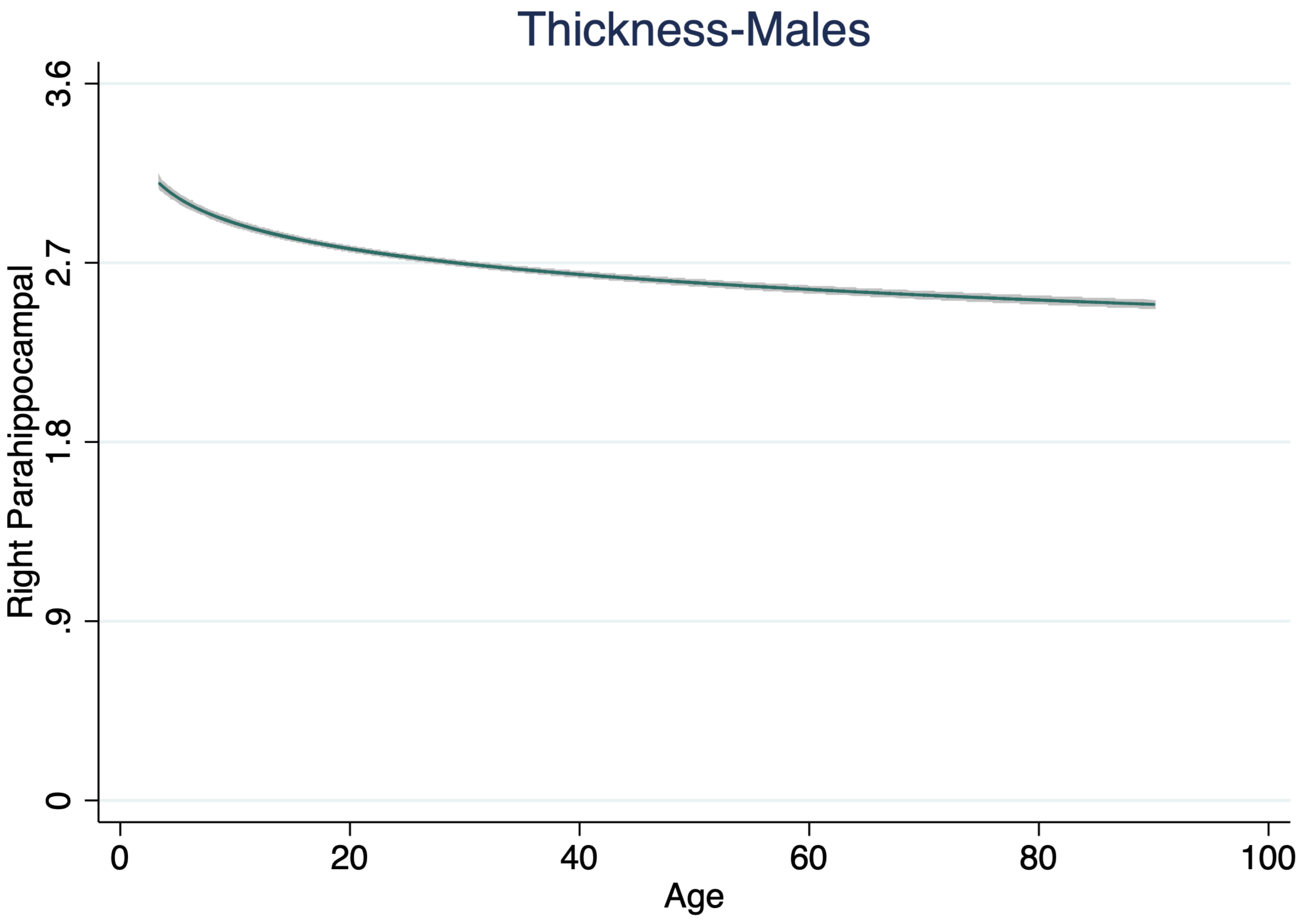

Thickness-Males
Right Parahippocampal
3.6
2.7
1.8
.9
0
0
20
40
60
80
100
Age

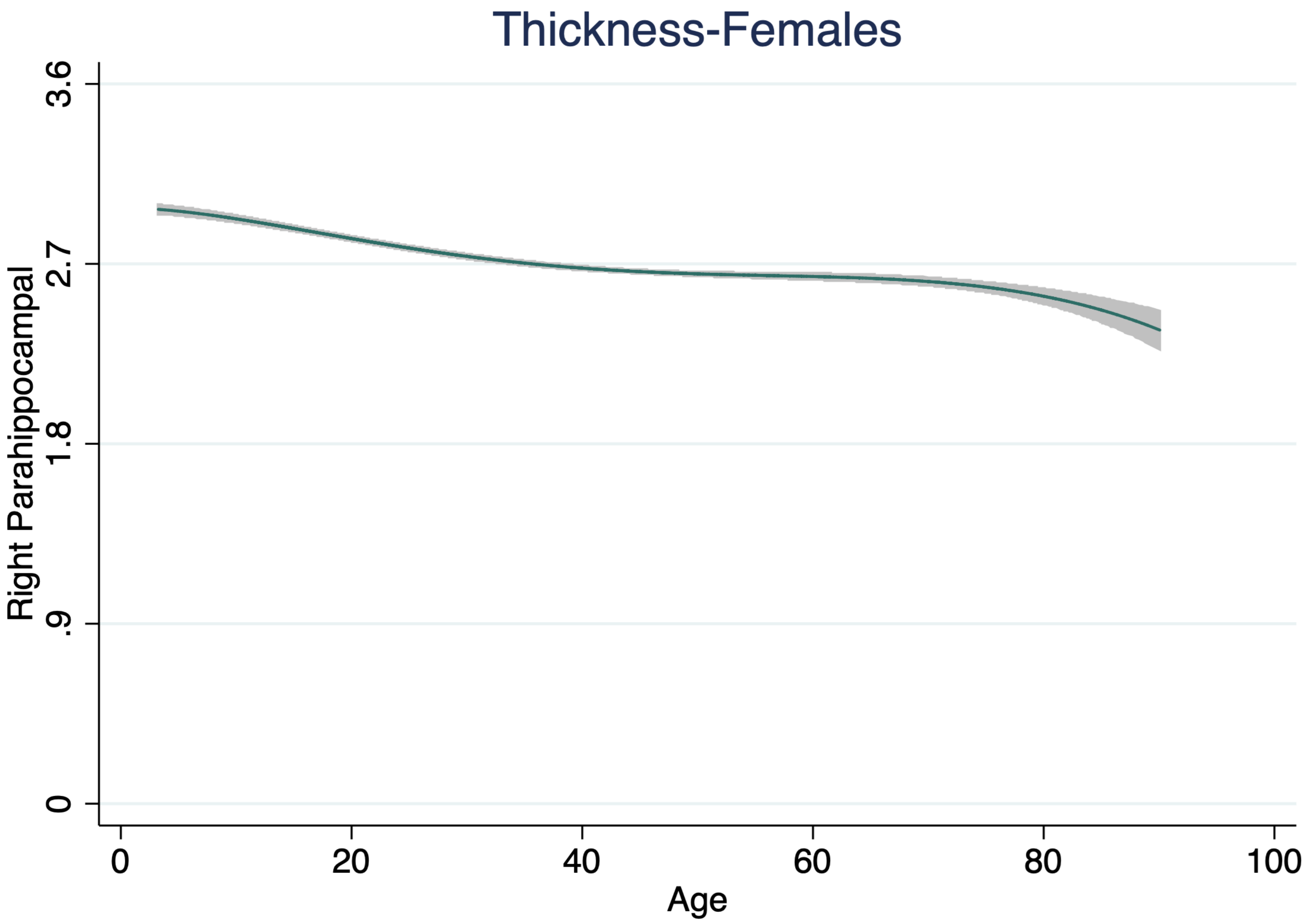

Thickness-Females
Right Parahippocampal
3.6
2.7
1.8
.9
0
0
20
40
60
80
100
Age

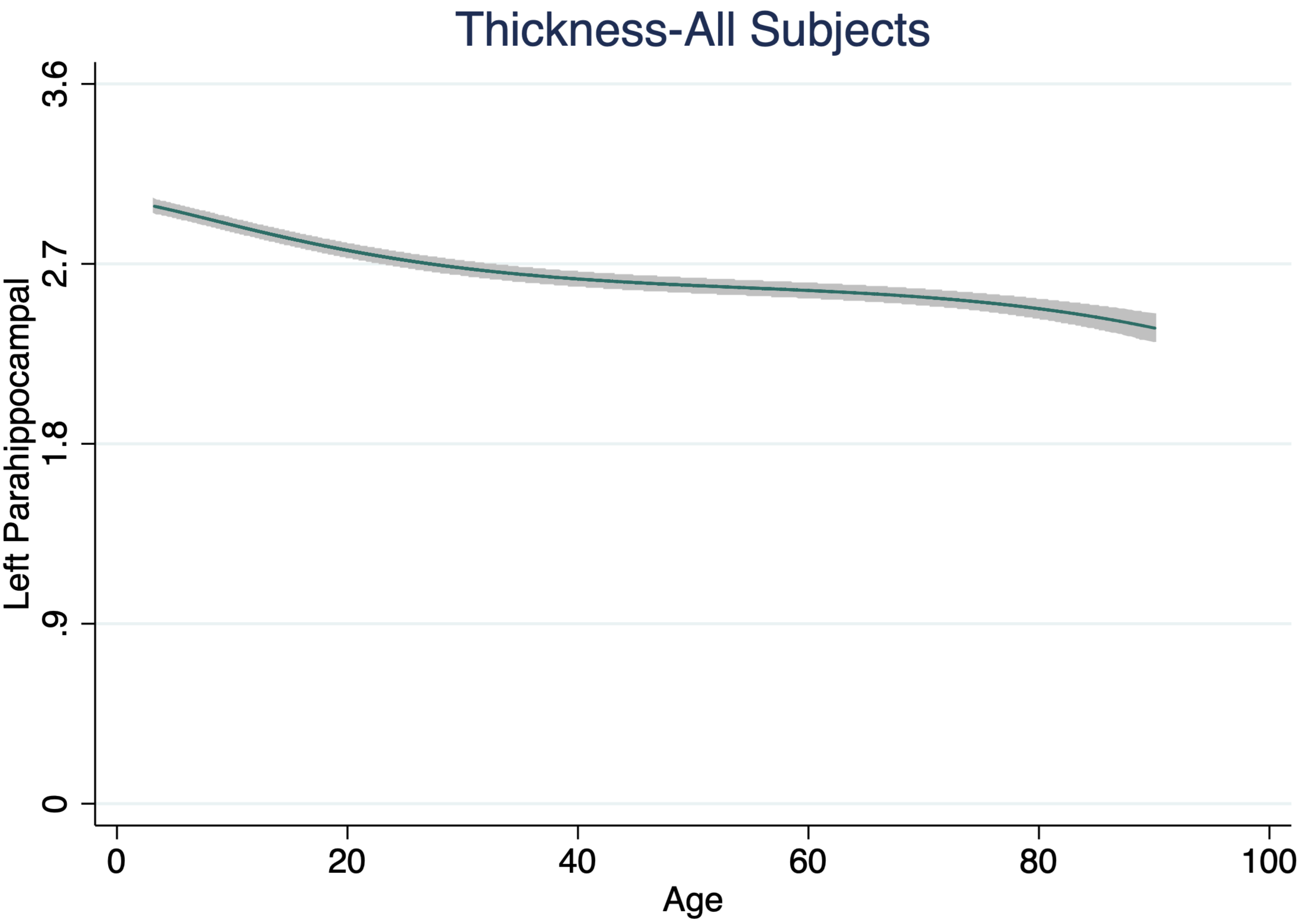
Thickness-All Subjects
Left Parahippocampal
3.6
2.7
1.8
.9
0
0
20
40
60
80
100
Age

Thickness-Males

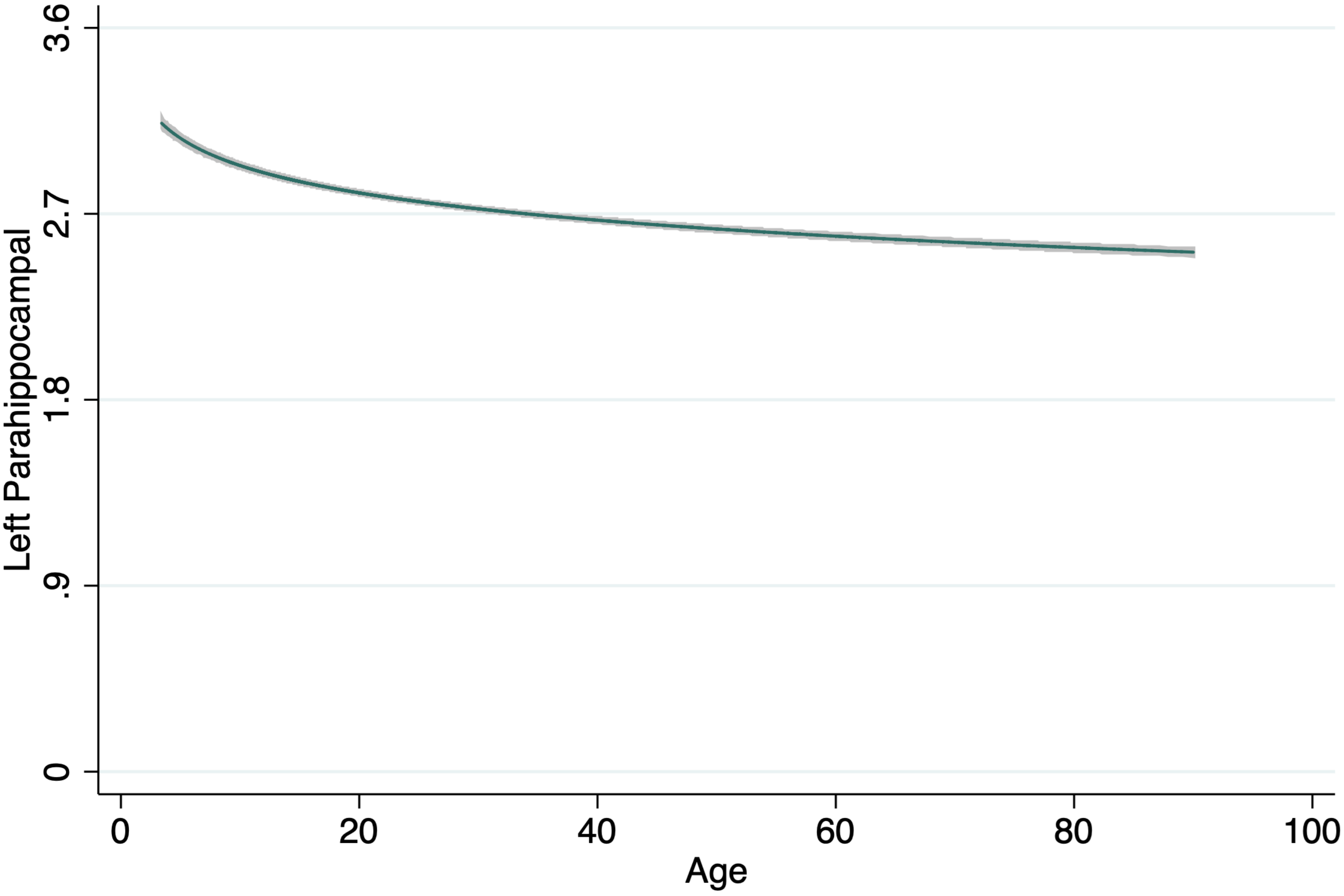

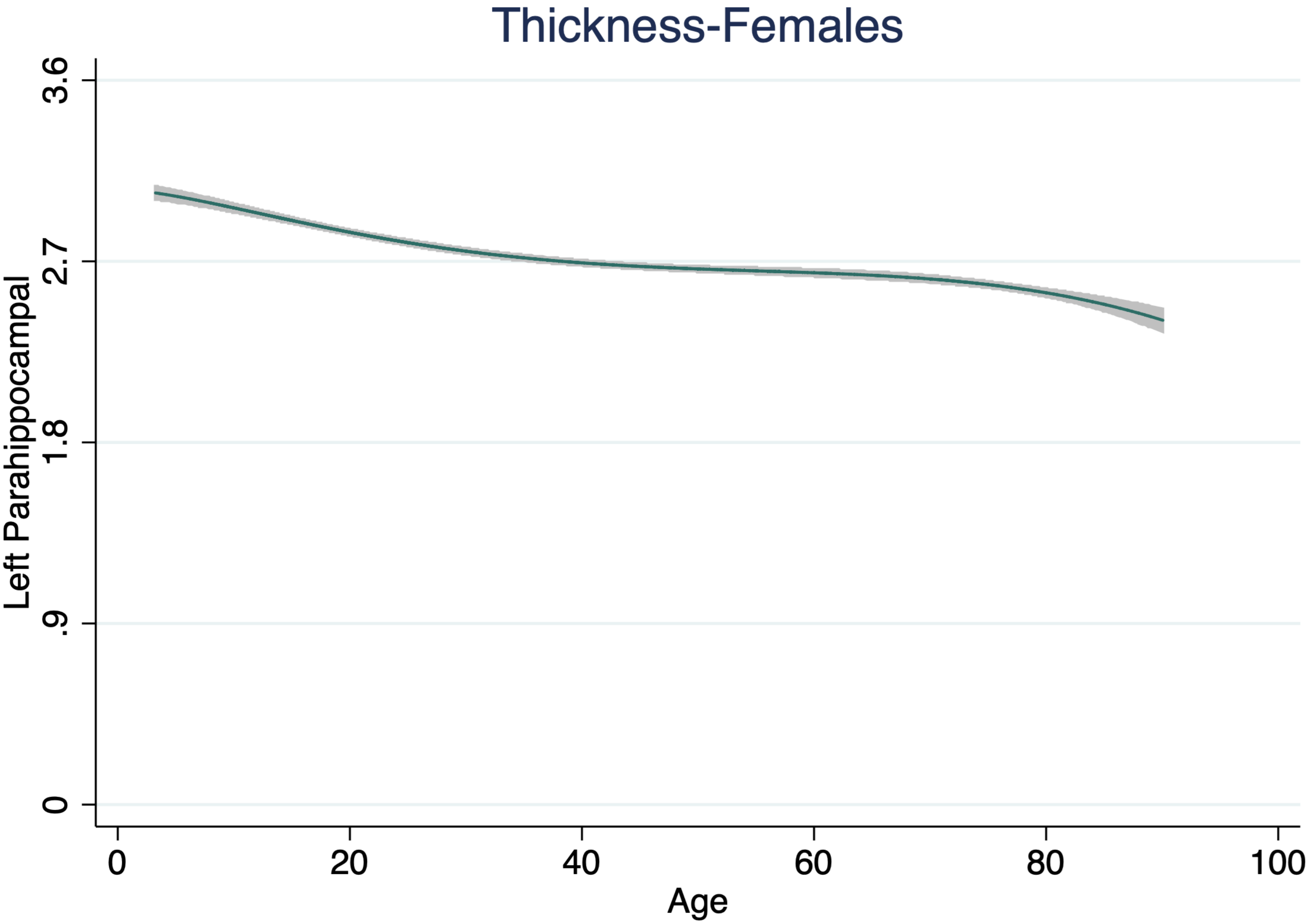

Thickness-Females
Left Parahippocampal
0
.9
1.8
2.7
3.6
0
20
40
60
80
100
Age

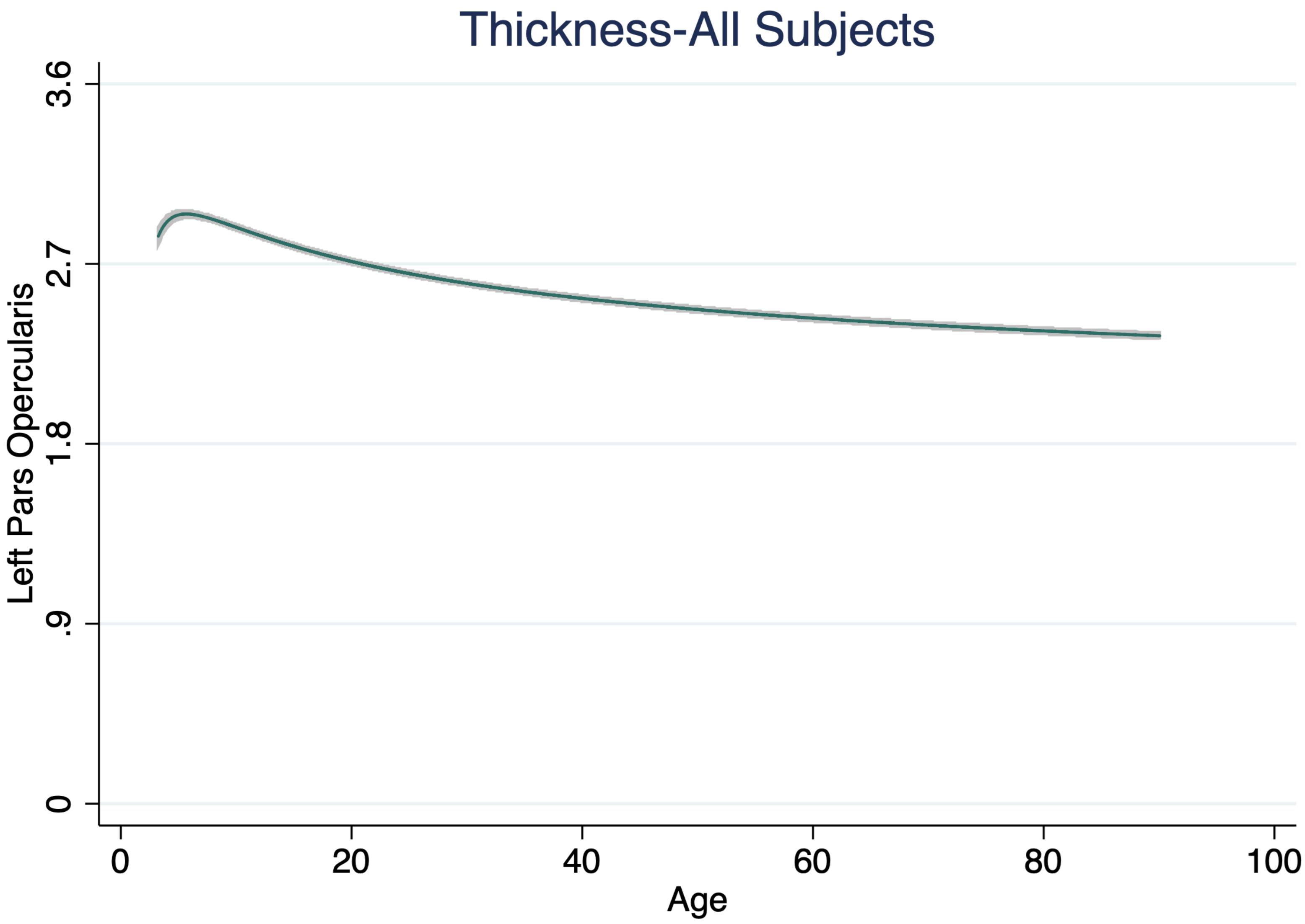

Thickness-All Subjects
Left Pars Opercularis
3.6
2.7
1.8
.9
0
0
20
40
60
80
100
Age

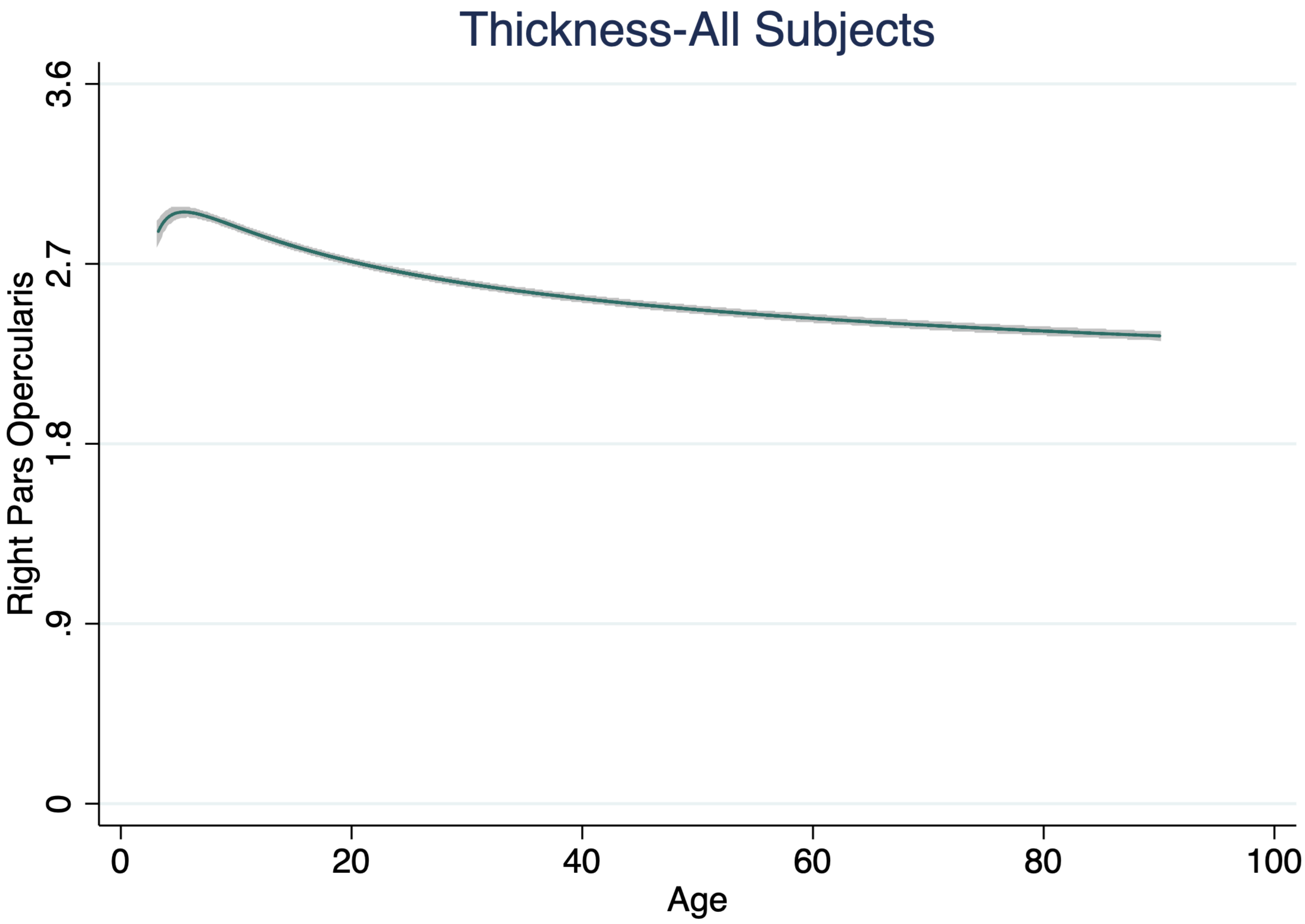
Thickness-All Subjects
Right Pars Opercularis
3.6
2.7
1.8
.9
0
0
20
40
60
80
100
Age

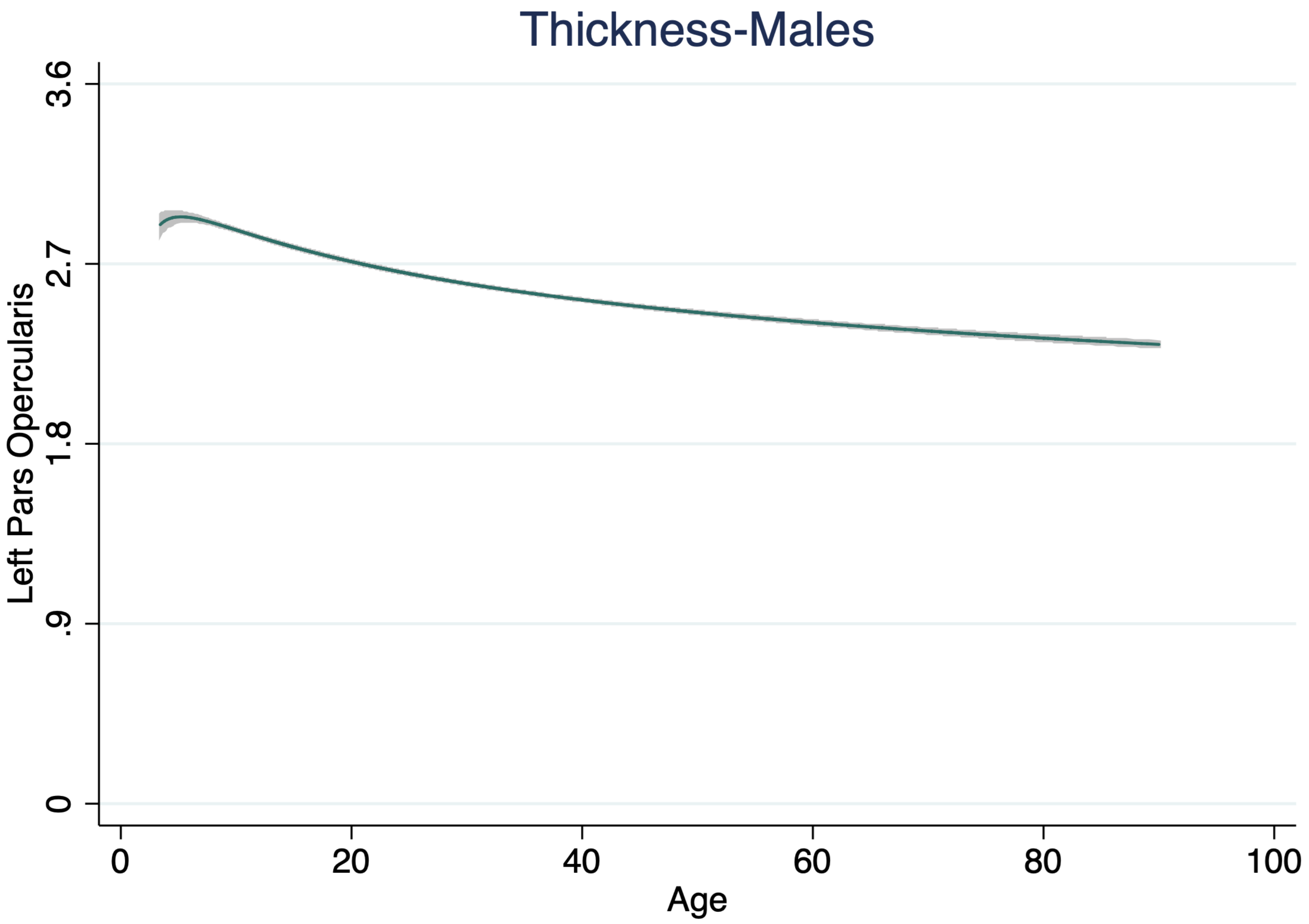

Thickness-Males
Left Pars Opercularis
3.6
2.7
1.8
.9
0
0
20
40
60
80
100
Age

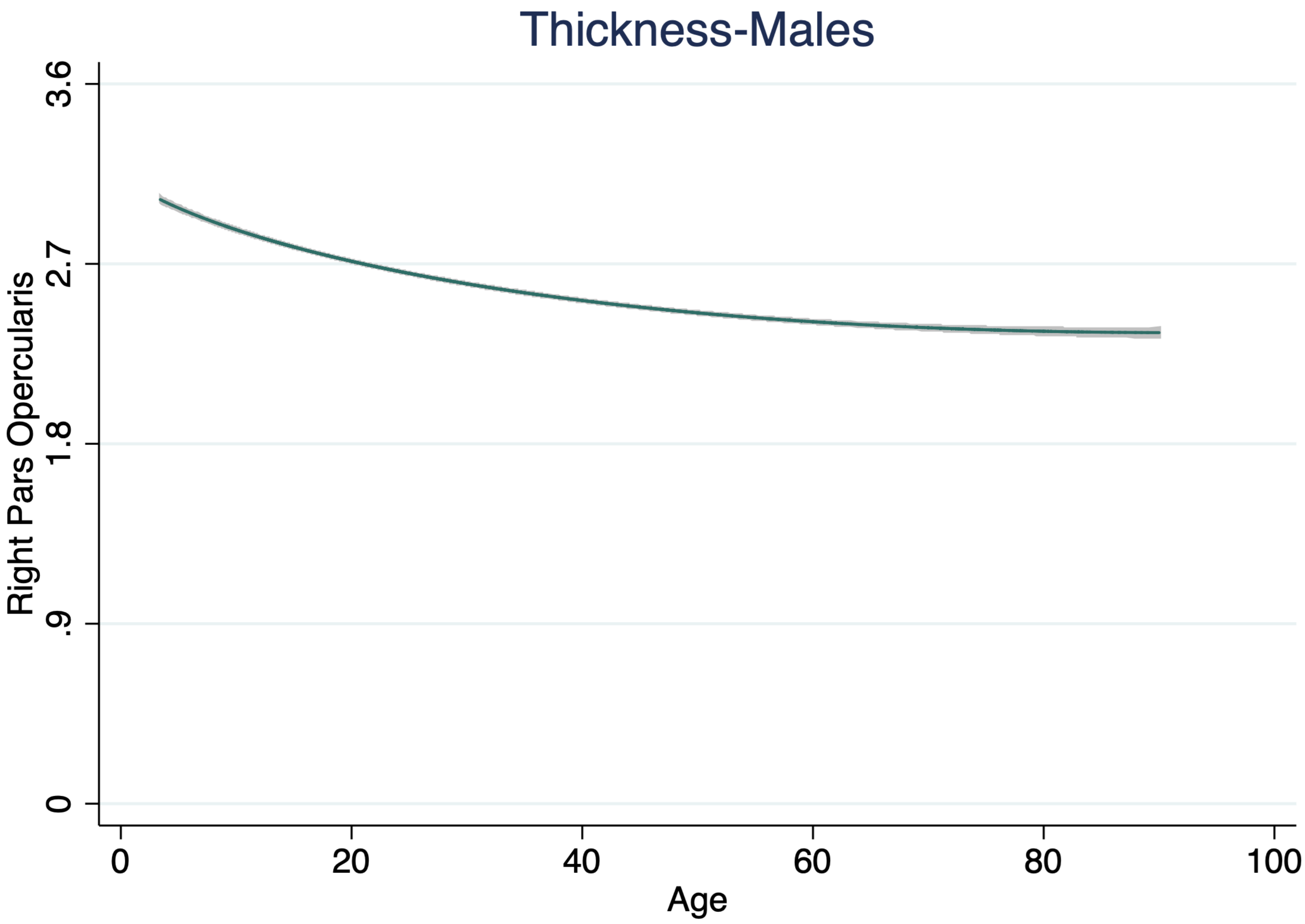
Thickness-Males
Right Pars Opercularis
3.6
2.7
1.8
.9
0
0
20
40
60
80
100
Age

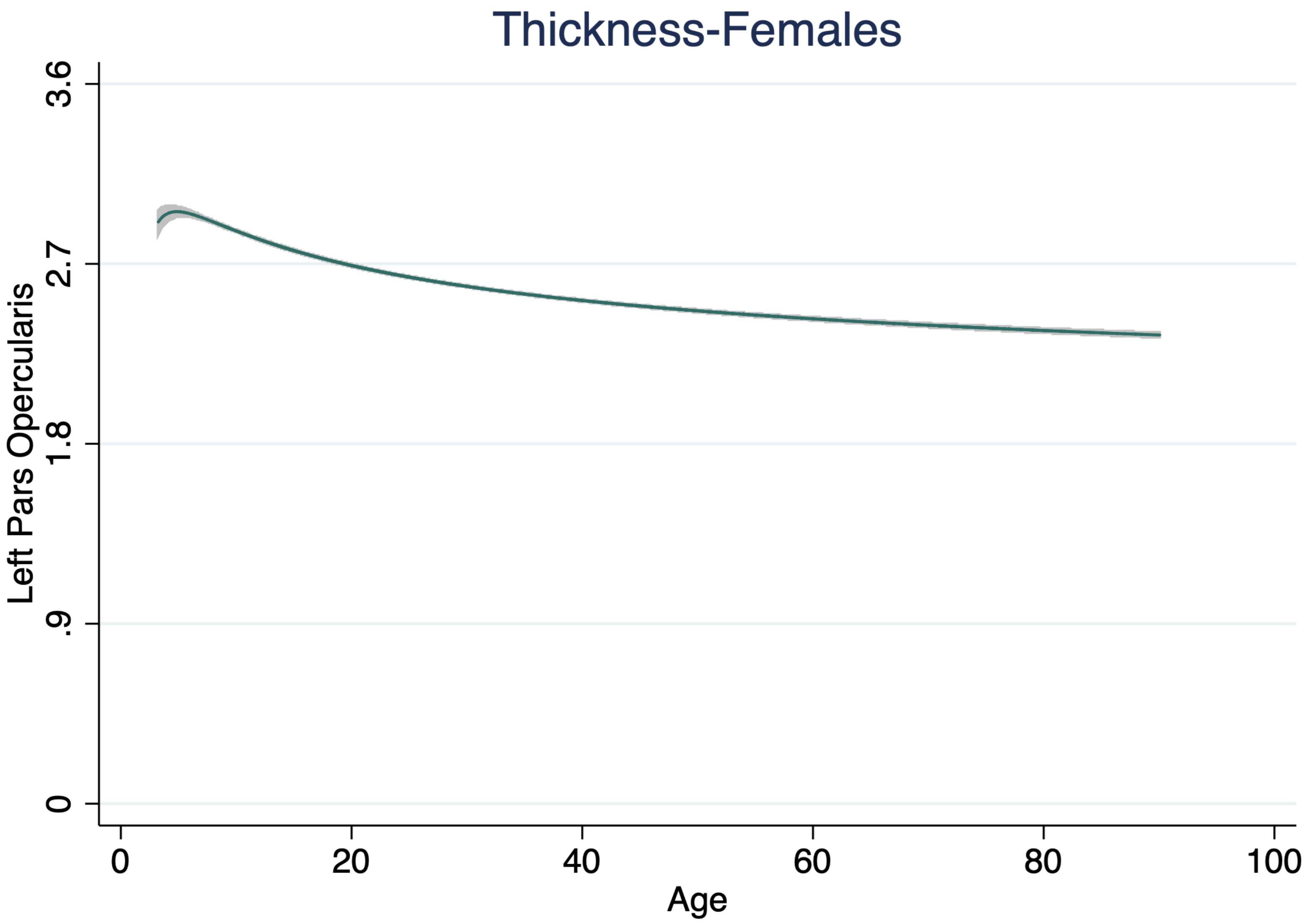

Thickness-Females
Left Pars Opercularis
3.6
2.7
1.8
.9
0
0
20
40
60
80
100
Age

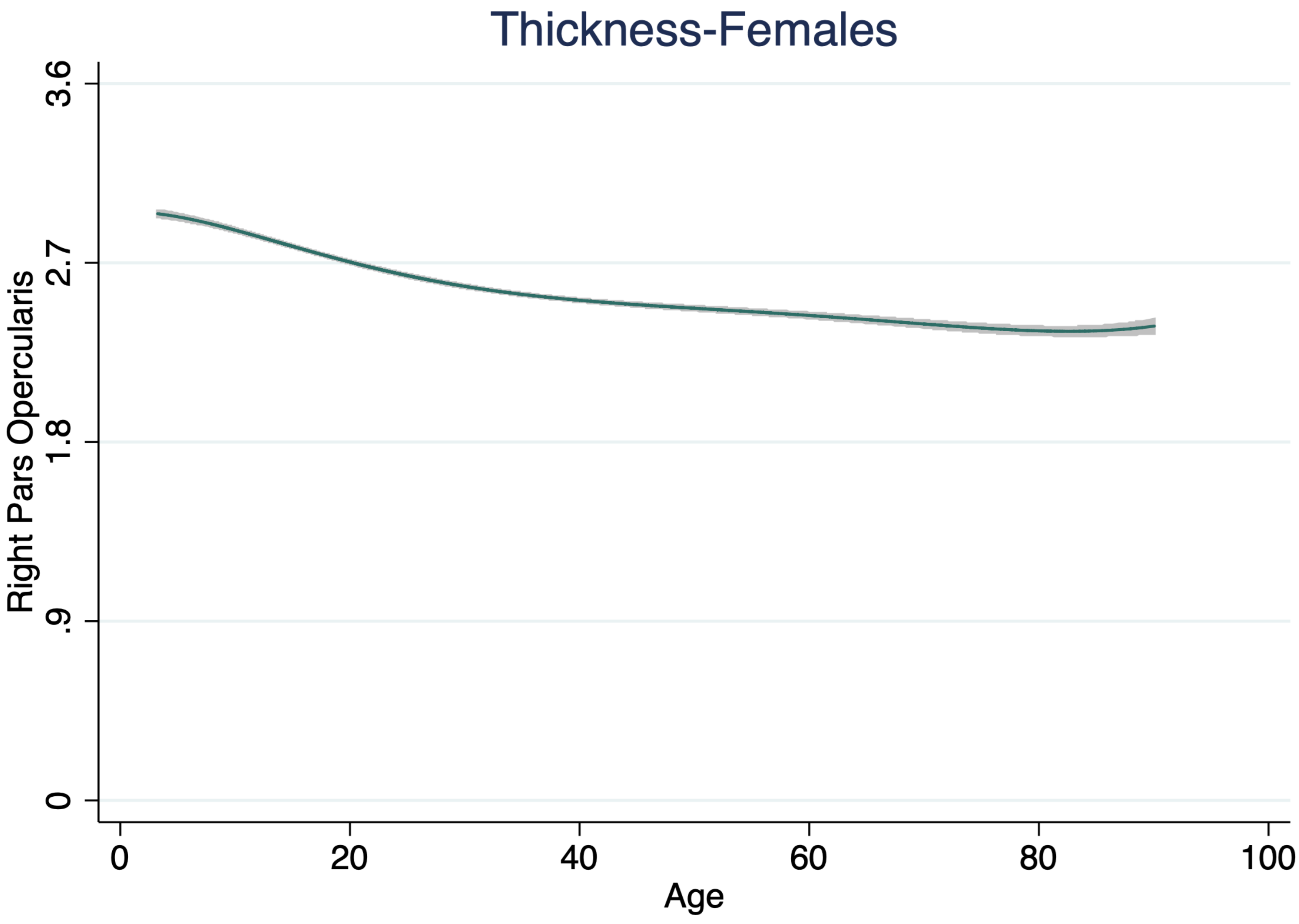
Thickness-Females
Right Pars Opercularis
3.6
2.7
1.8
.9
0
0
20
40
60
80
100
Age

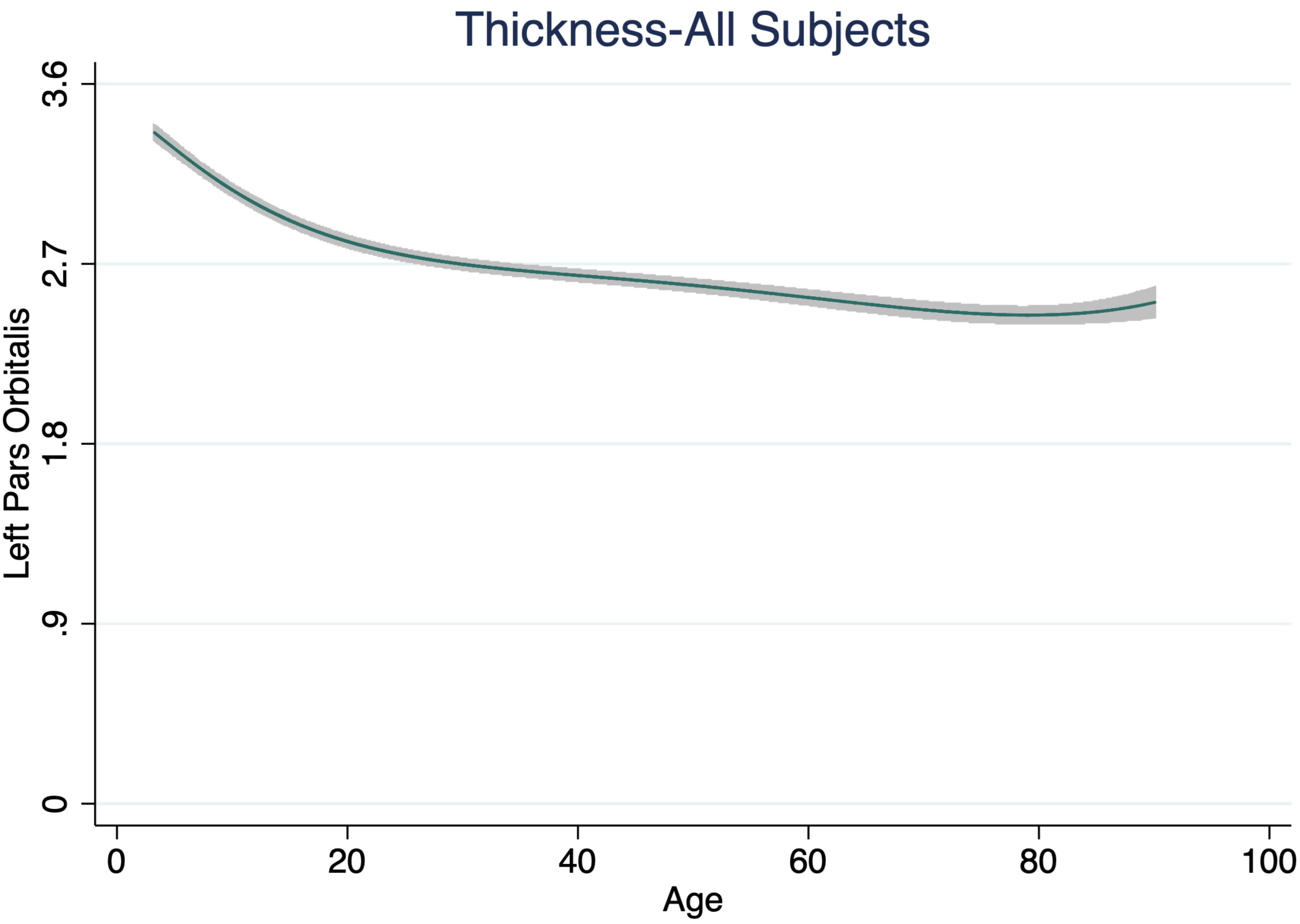

Thickness-All Subjects
Left Pars Orbitalis
0
.9
1.8
2.7
3.6
0
20
40
60
80
100
Age

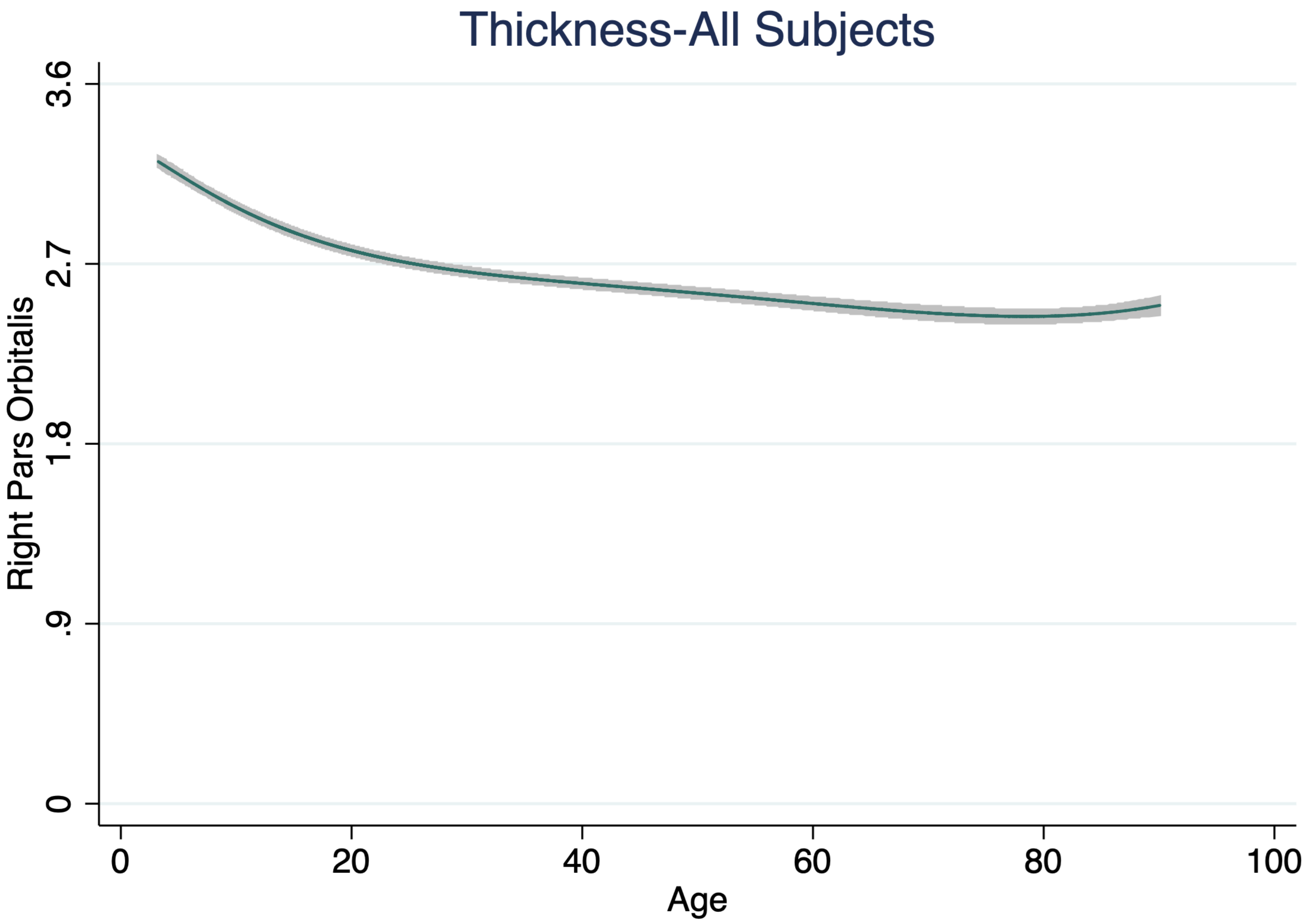

Thickness-All Subjects
Right Pars Orbitalis
3.6
2.7
1.8
.9
0
0
20
40
60
80
100
Age

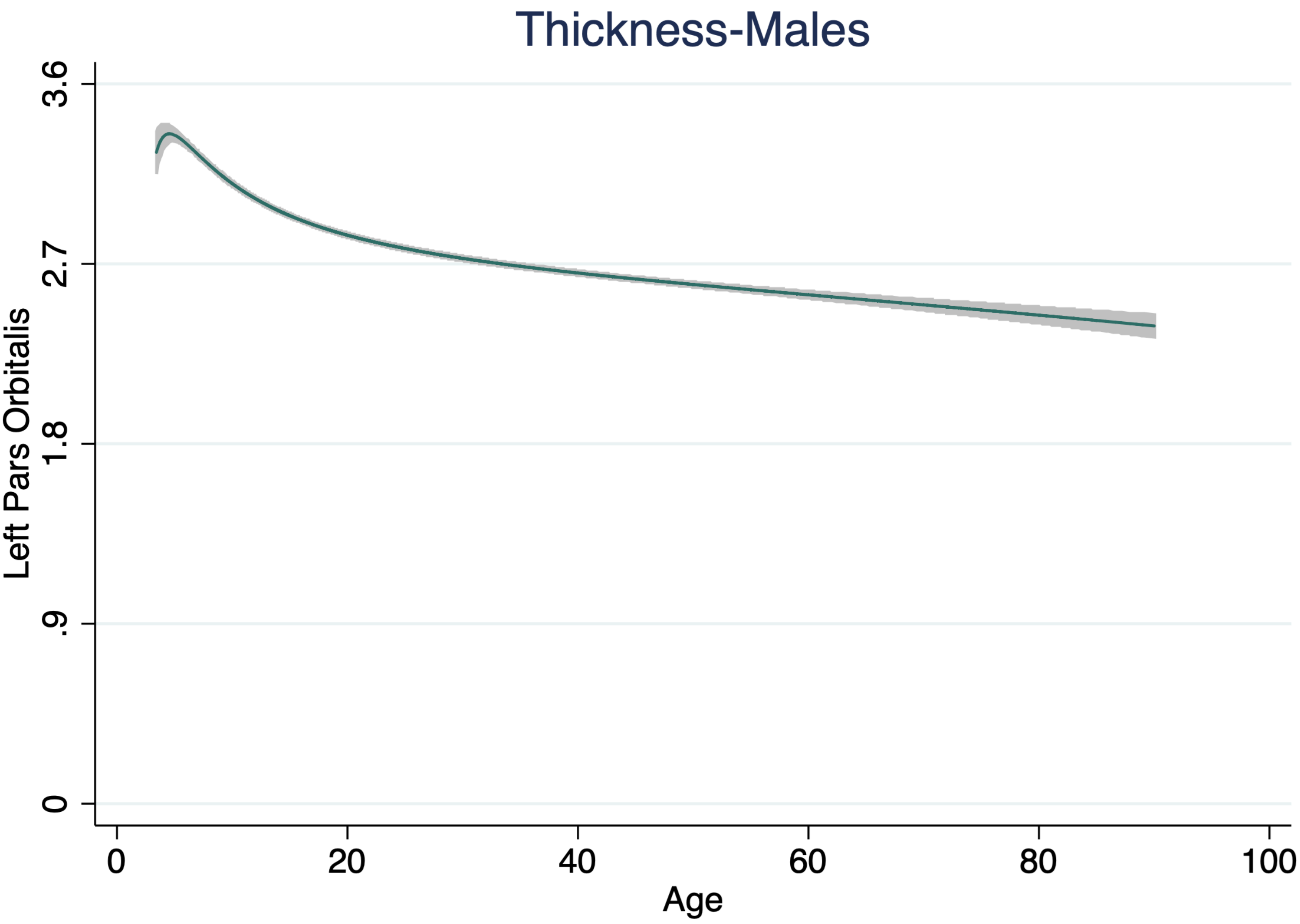
Thickness-Males
Left Pars Orbitalis
3.6
2.7
1.8
.9
0
0
20
40
60
80
100
Age

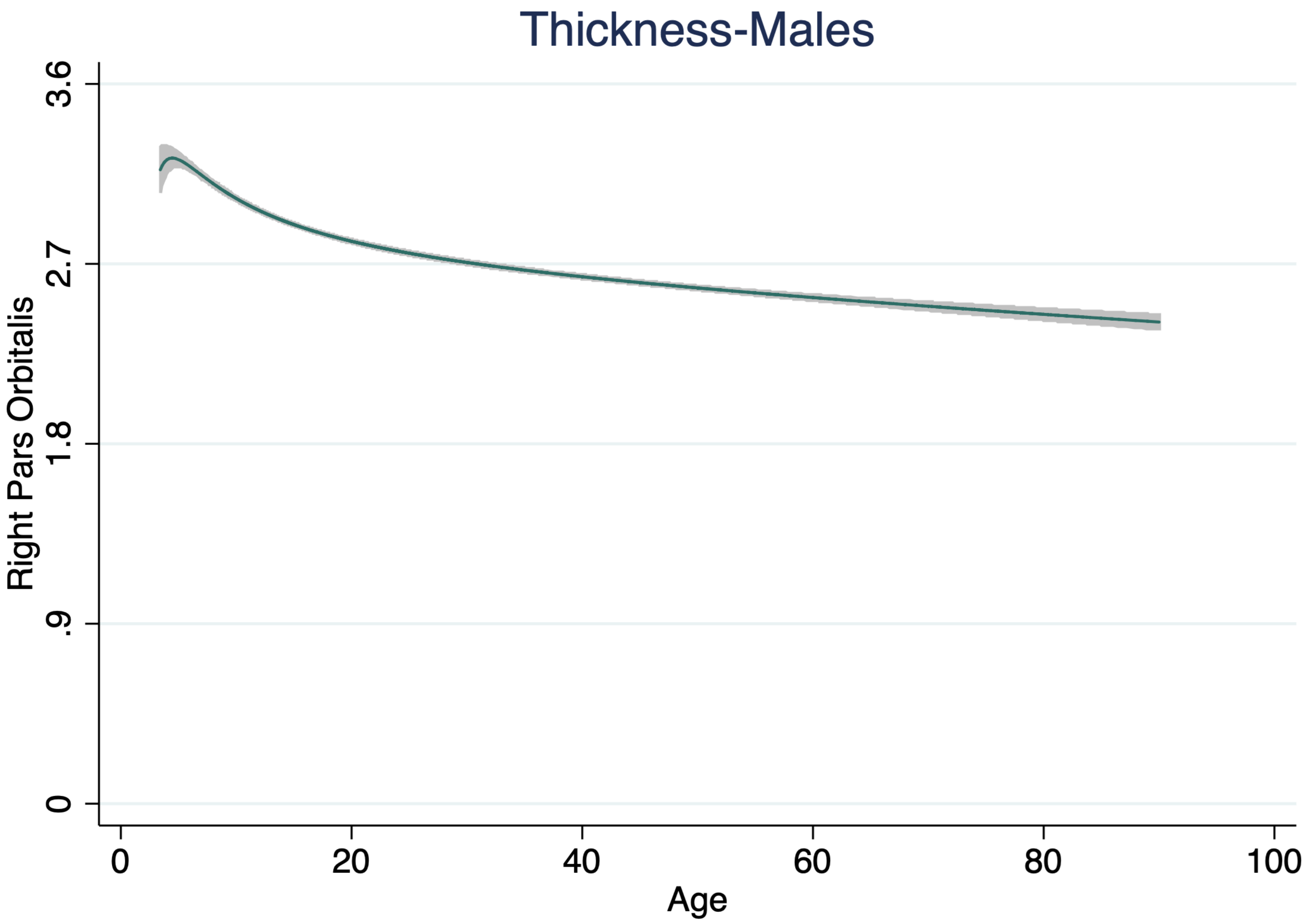
Thickness-Males
Right Pars Orbitalis
3.6
2.7
1.8
.9
0
0
20
40
60
80
100
Age

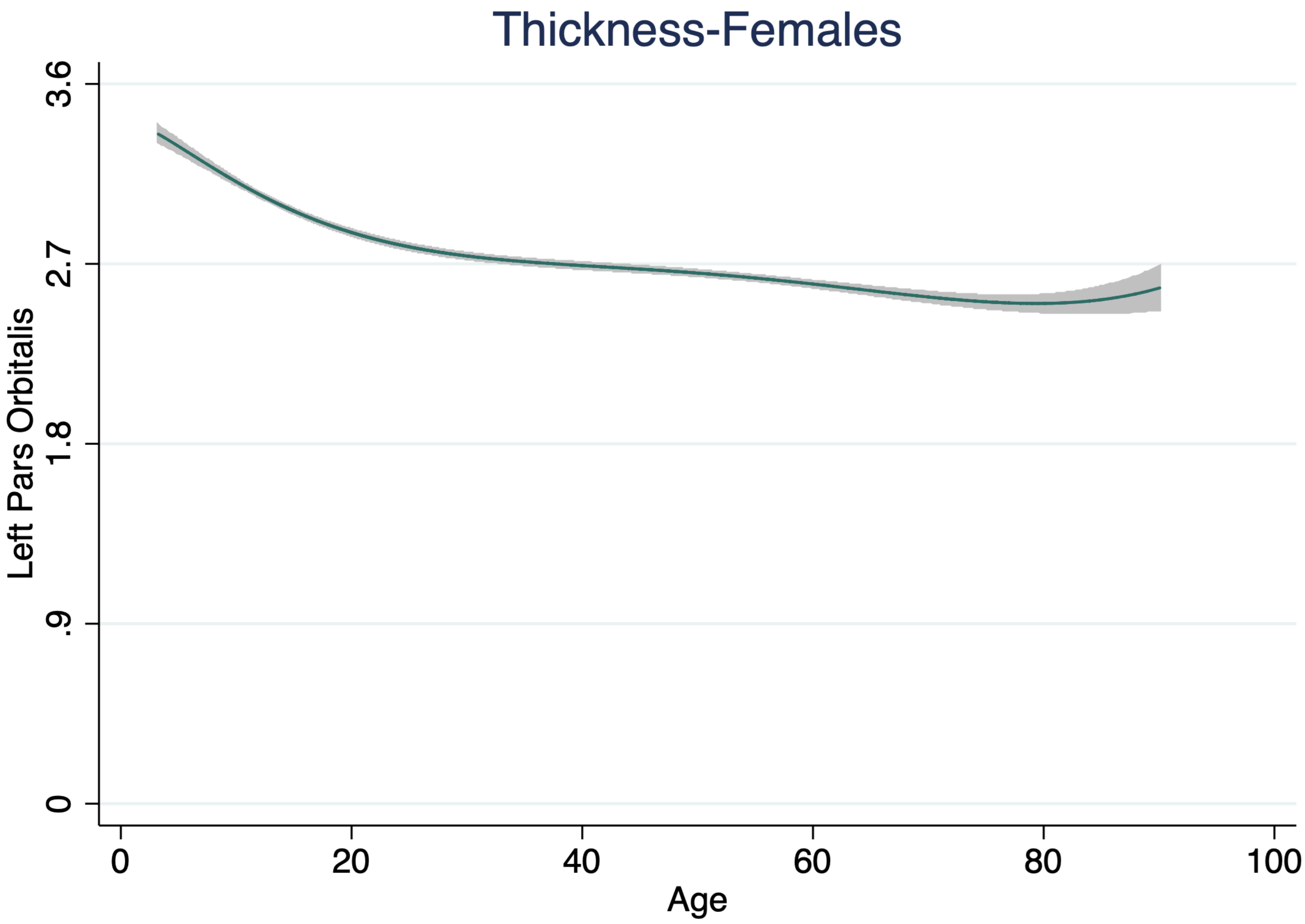

Thickness-Females
Left Pars Orbitalis
3.6
2.7
1.8
.9
0
0
20
40
60
80
100
Age

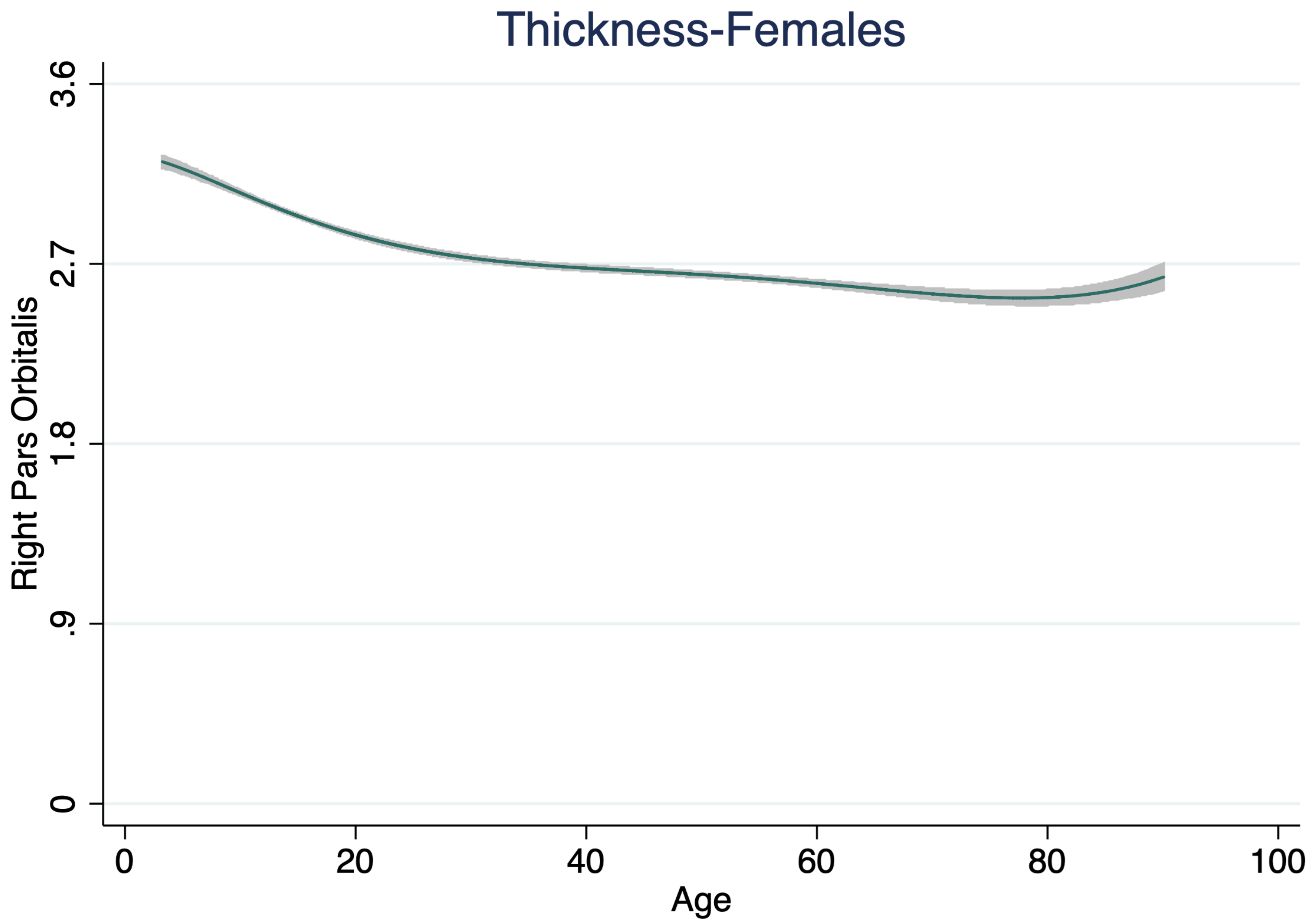
Thickness-Females
Right Pars Orbitalis
3.6
2.7
1.8
.9
0
0
20
40
60
80
100
Age

Thickness-All Subjects

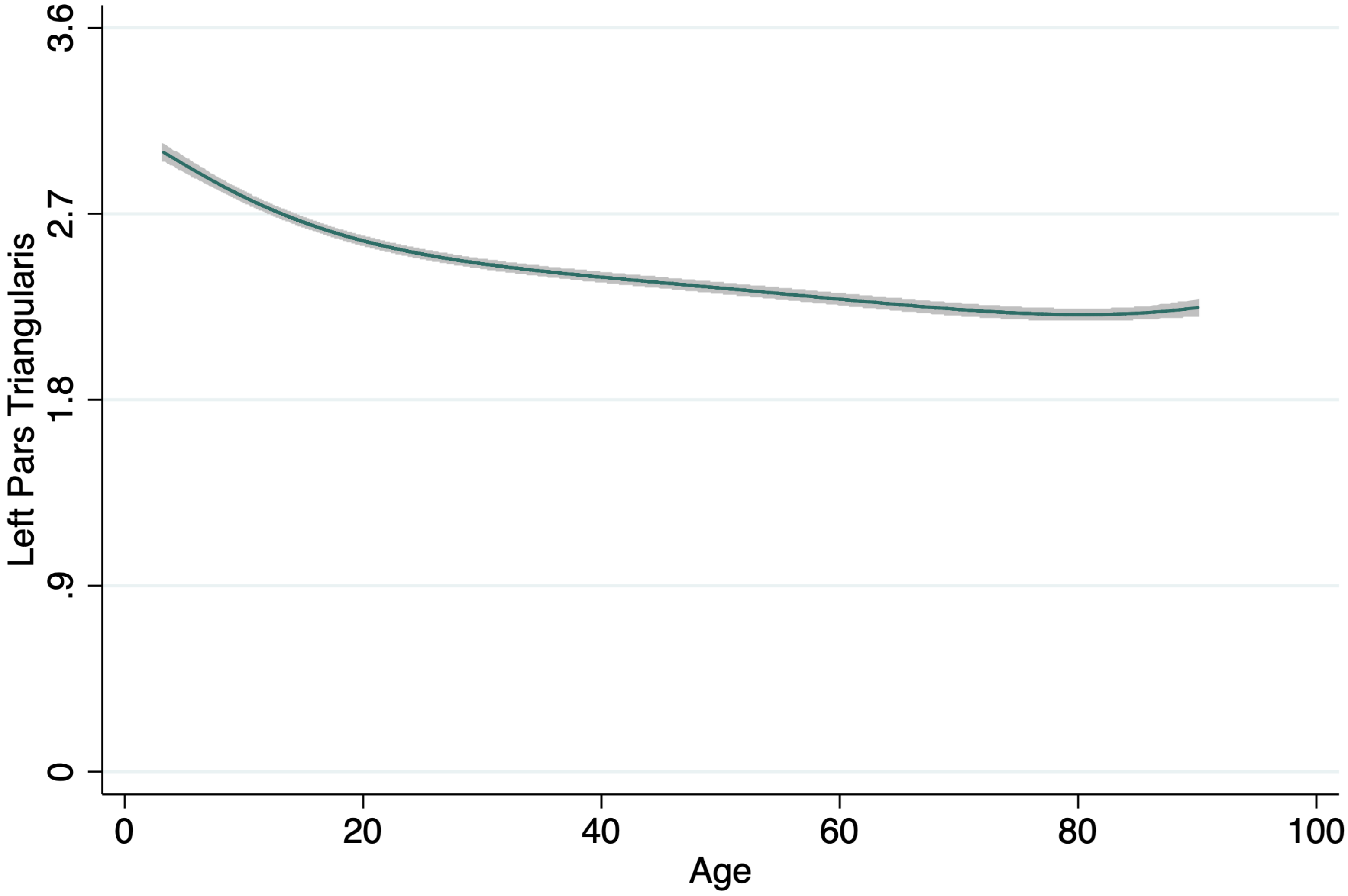

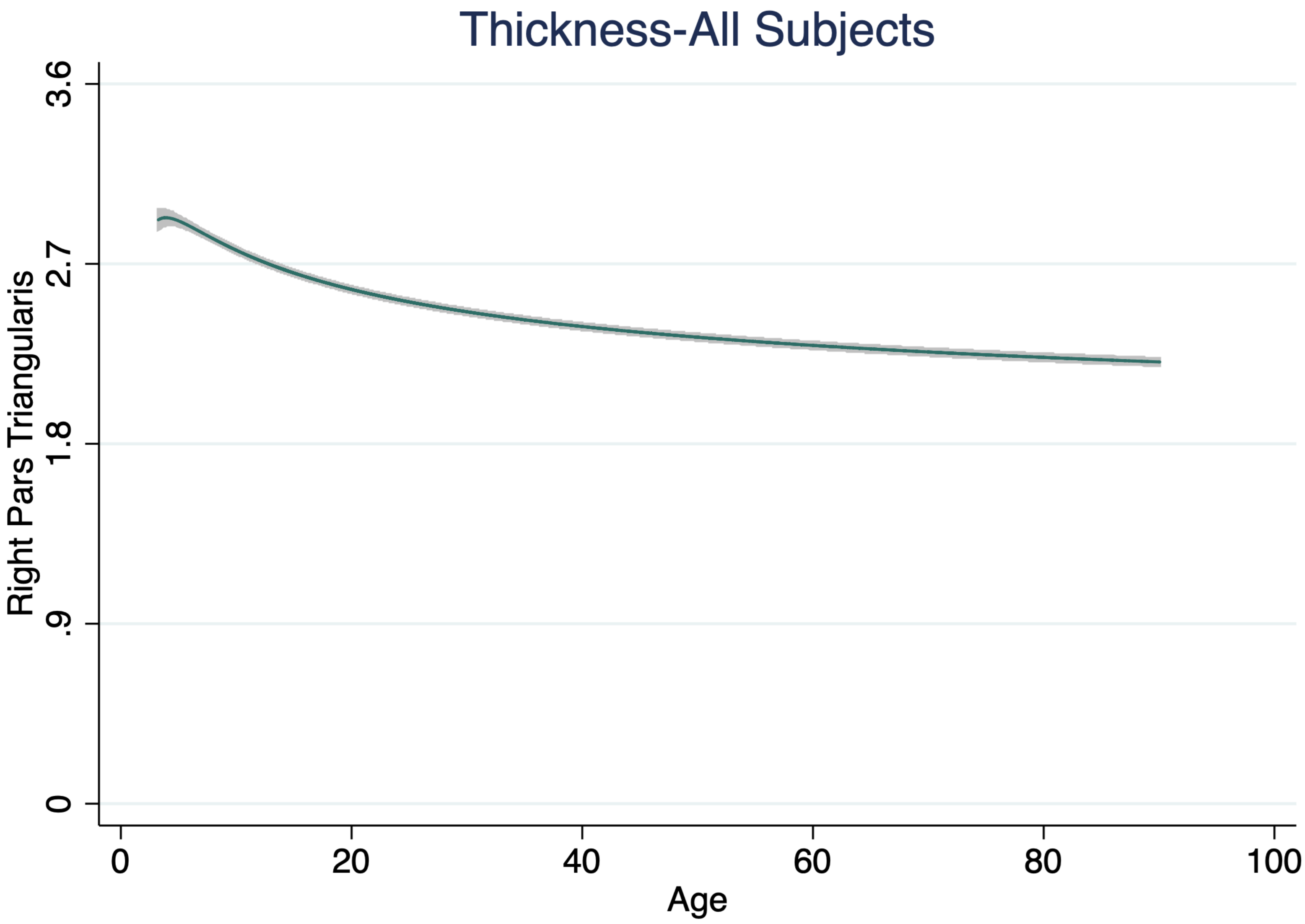

Thickness-All Subjects
Right Pars Triangularis
3.6
2.7
1.8
.9
0
0
20
40
60
80
100
Age

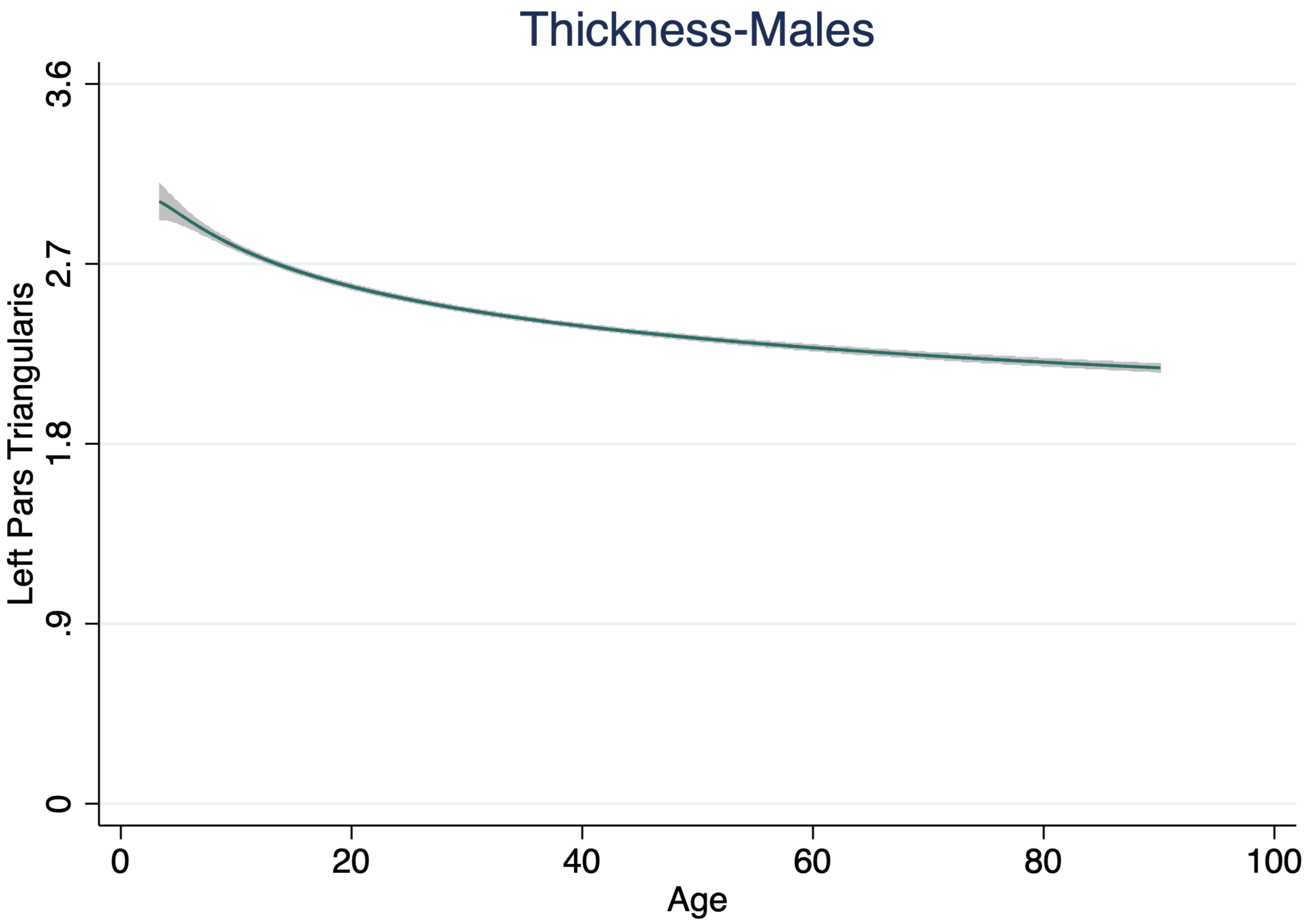
Thickness-Males
Left Pars Triangularis
3.6
2.7
1.8
.9
0
0
20
40
60
80
100
Age

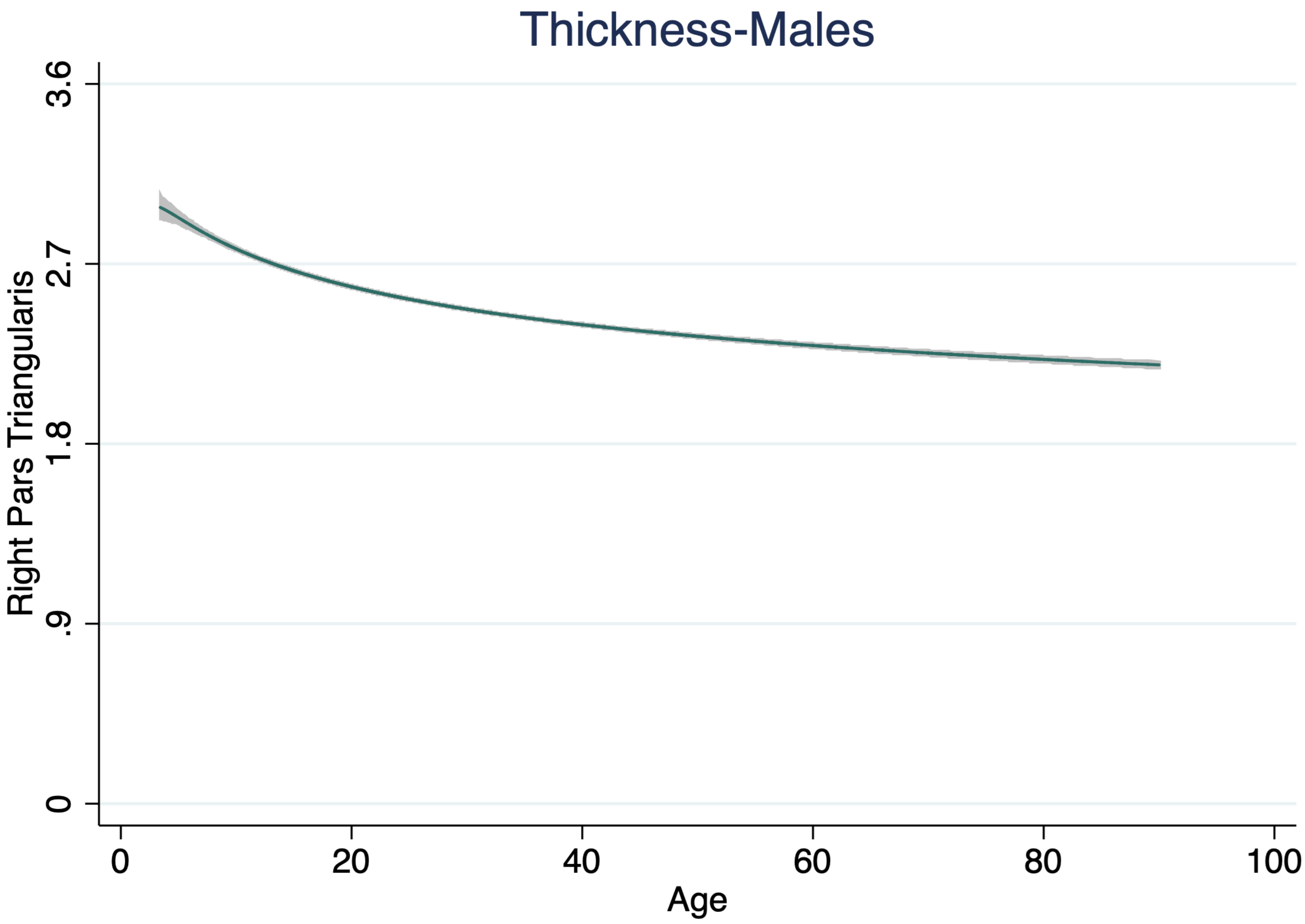

Thickness-Males
Right Pars Triangularis
3.6
2.7
1.8
.9
0
0
20
40
60
80
100
Age

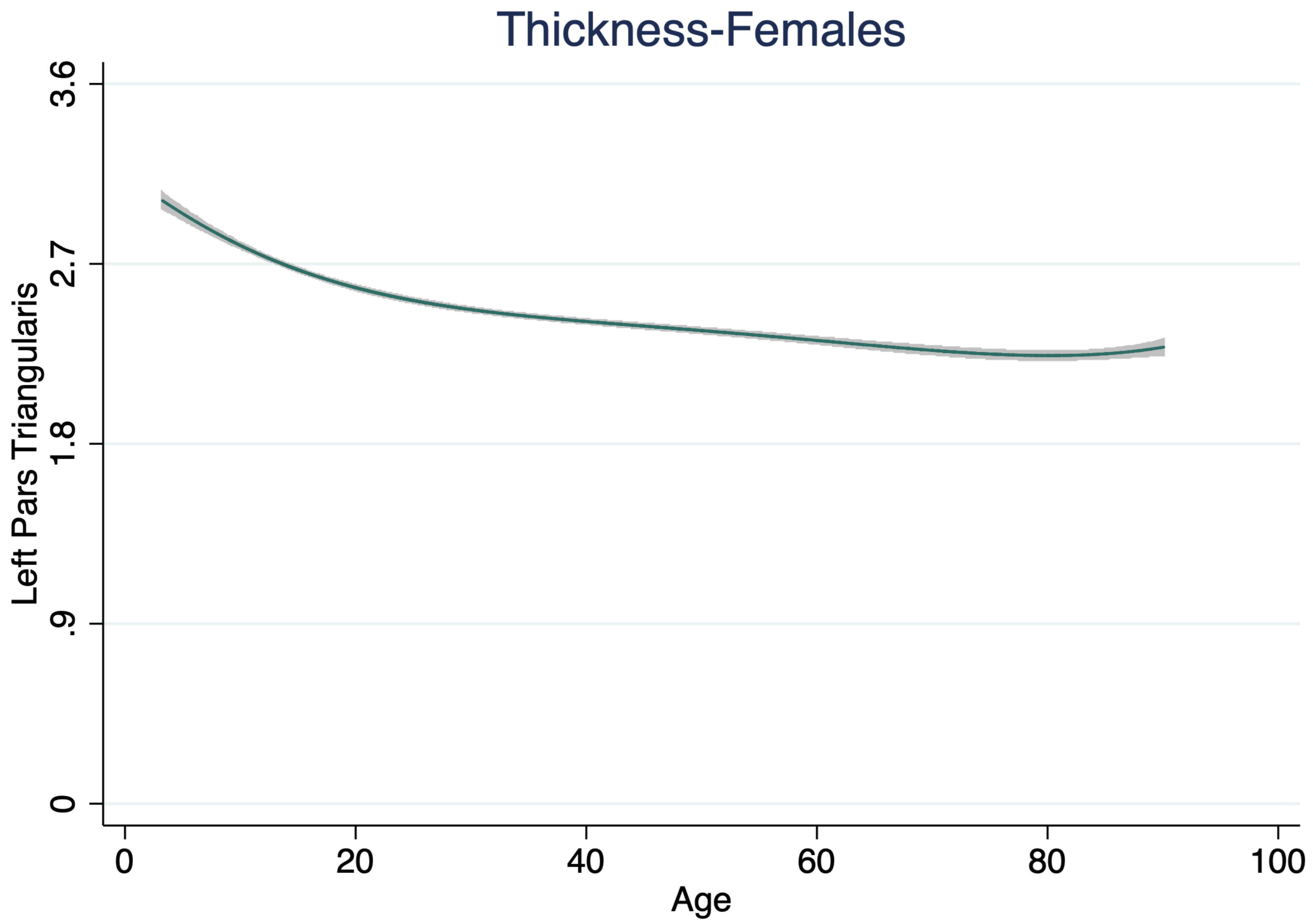
Thickness-Females
Left Pars Triangularis
3.6
2.7
1.8
.9
0
0
20
40
60
80
100
Age

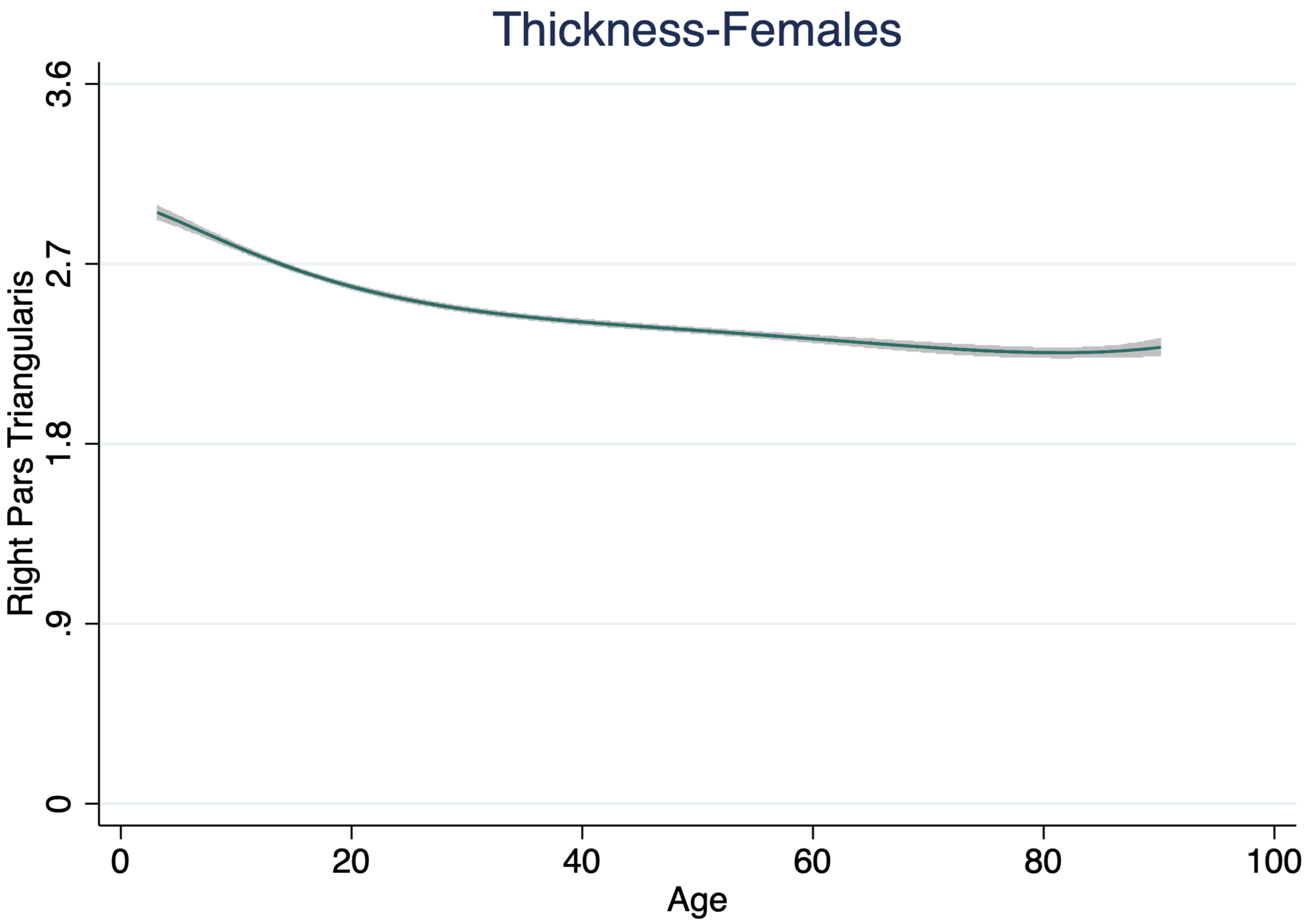
Thickness-Females
Right Pars Triangularis
3.6
2.7
1.8
.9
0
0
20
40
60
80
100
Age

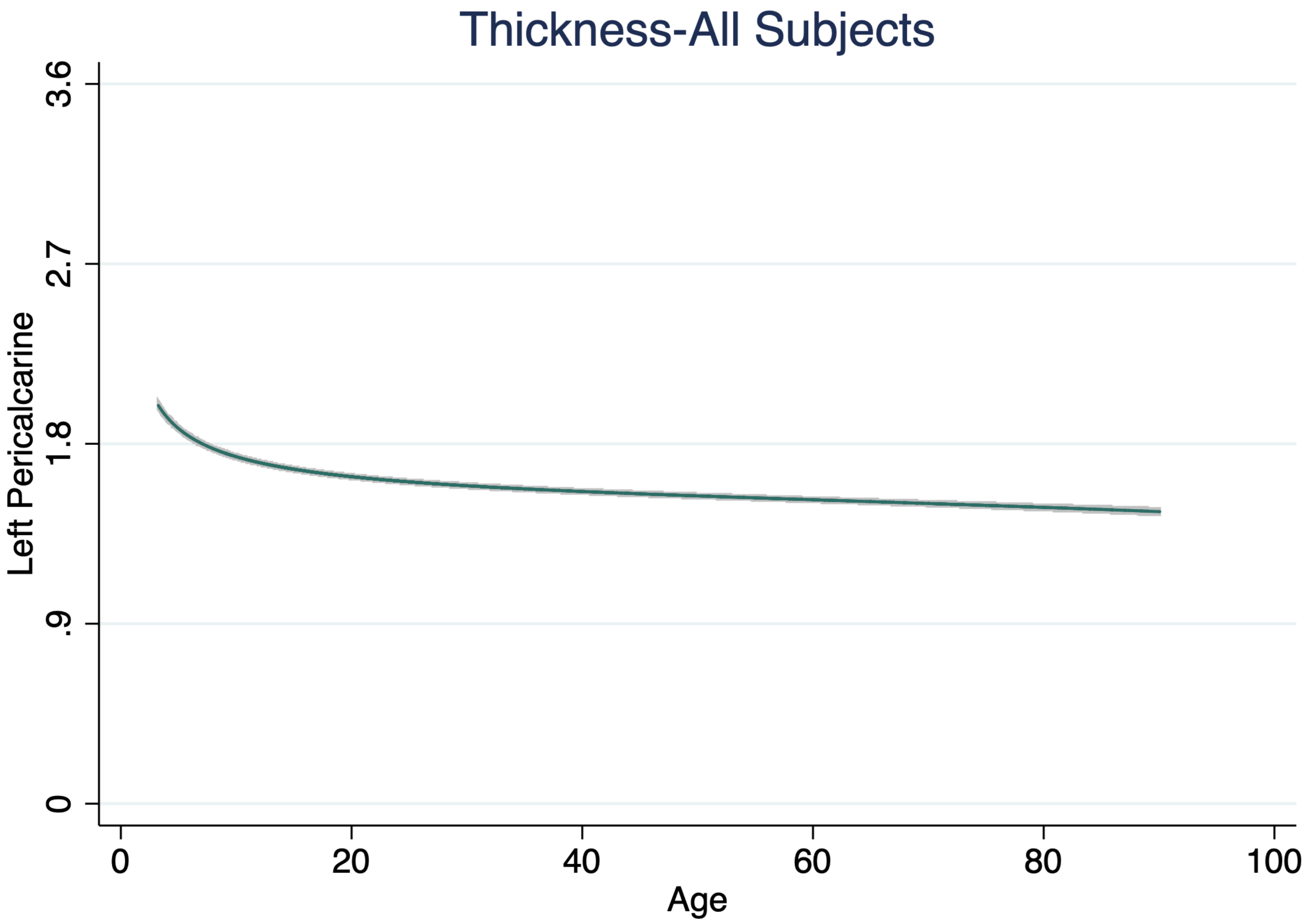

Thickness-All Subjects
Left Pericalcarine
3.6
2.7
1.8
.9
0
0
20
40
60
80
100
Age

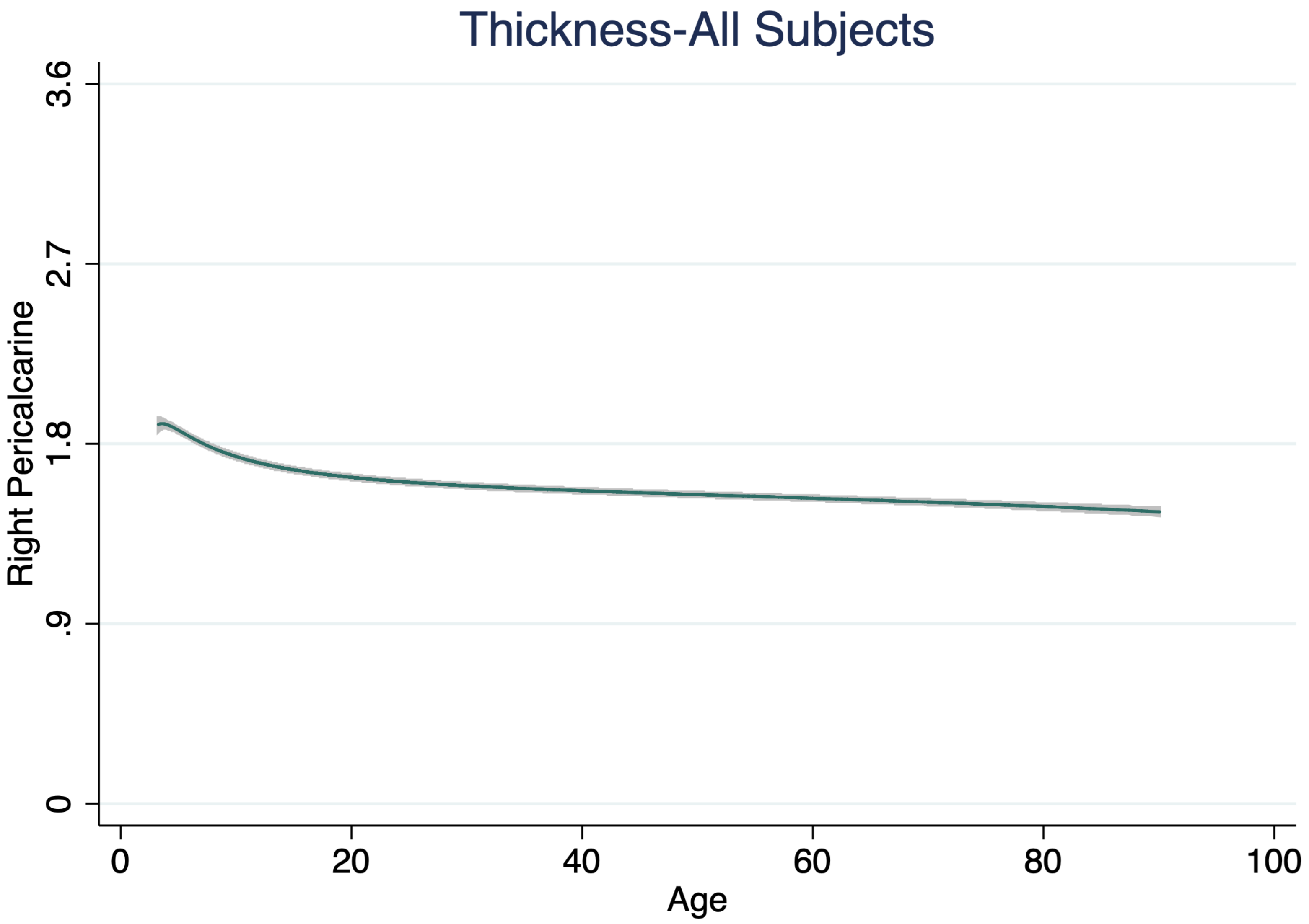
Thickness-All Subjects
Right Pericalcarine
3.6
2.7
1.8
.9
0
0
20
40
60
80
100
Age

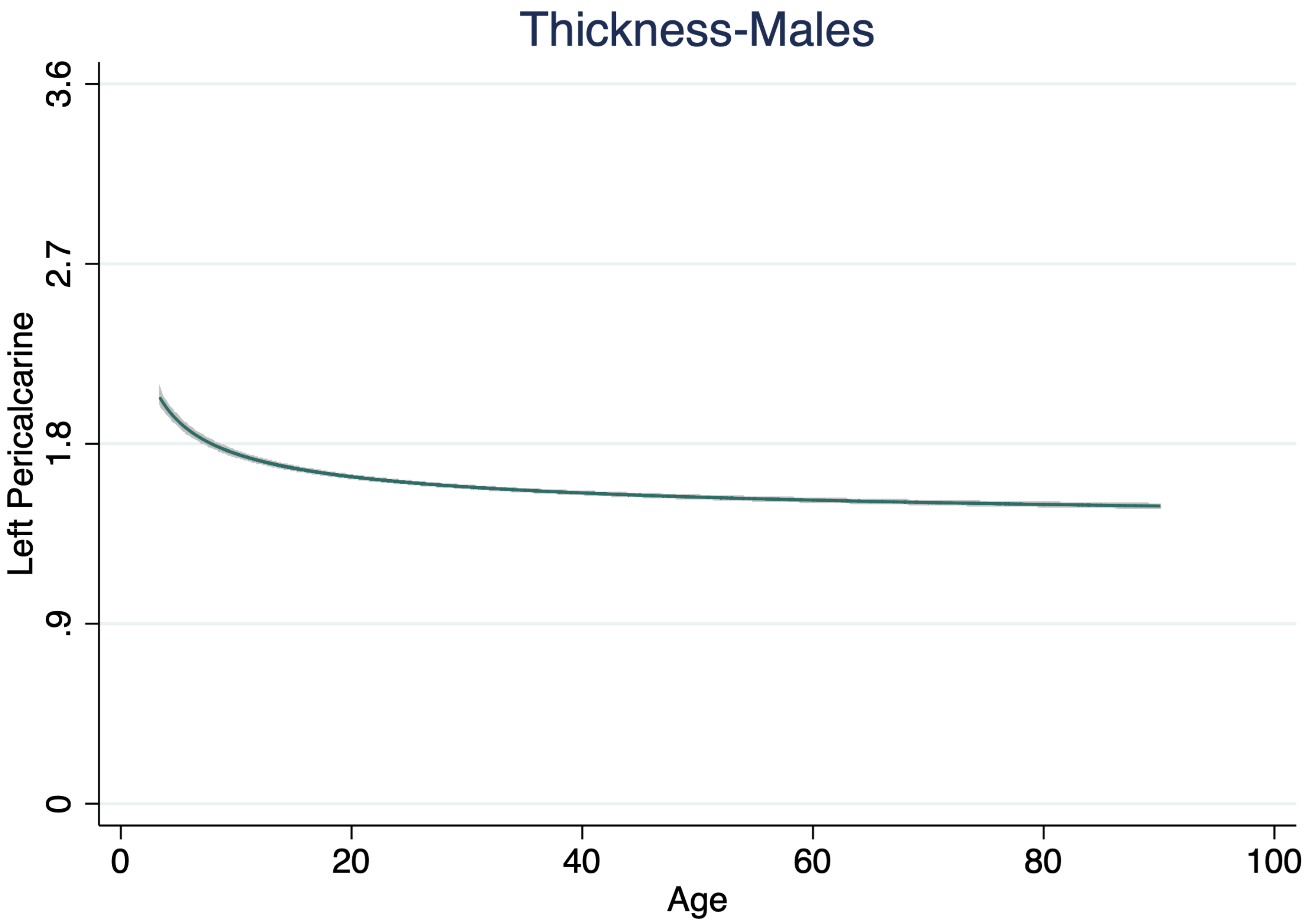

Thickness-Males
3.6
2.7
1.8
.9
0
Left Pericalcarine
0
20
40
60
80
100
Age

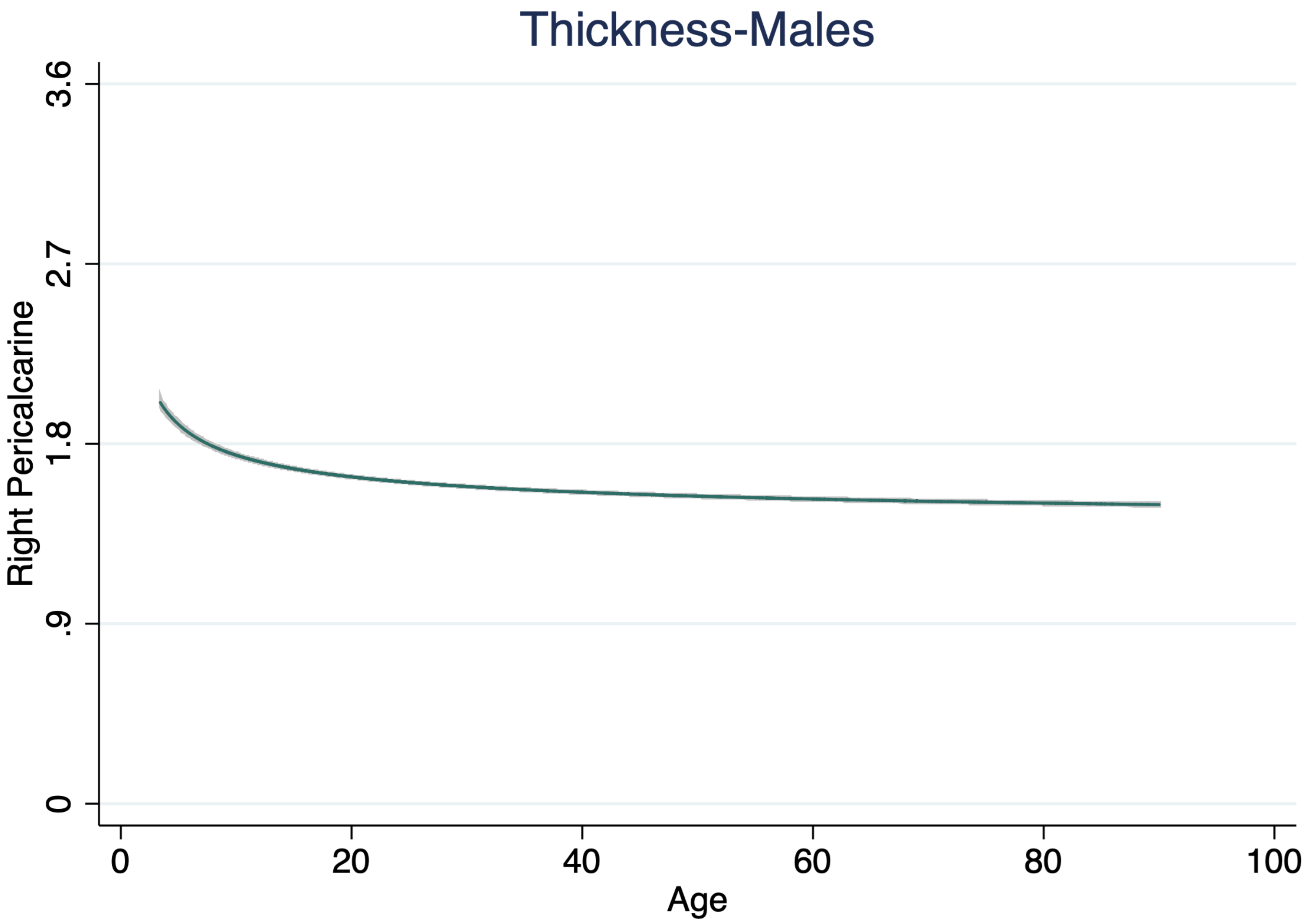
Thickness-Males
Right Pericalcarine
3.6
2.7
1.8
.9
0
0
20
40
60
80
100
Age

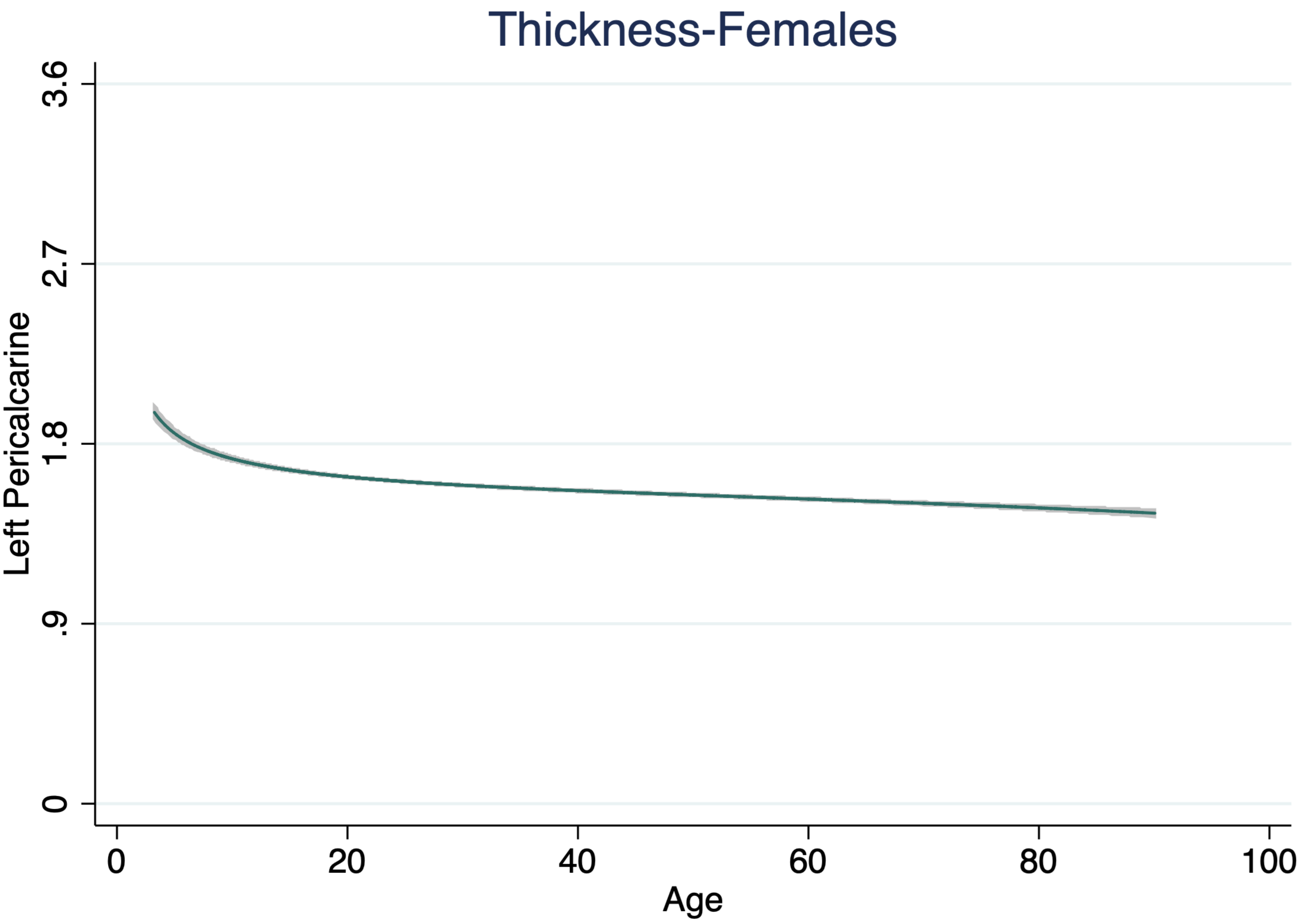

Thickness-Females
Left Pericalcarine
3.6
2.7
1.8
.9
0
0
20
40
60
80
100
Age

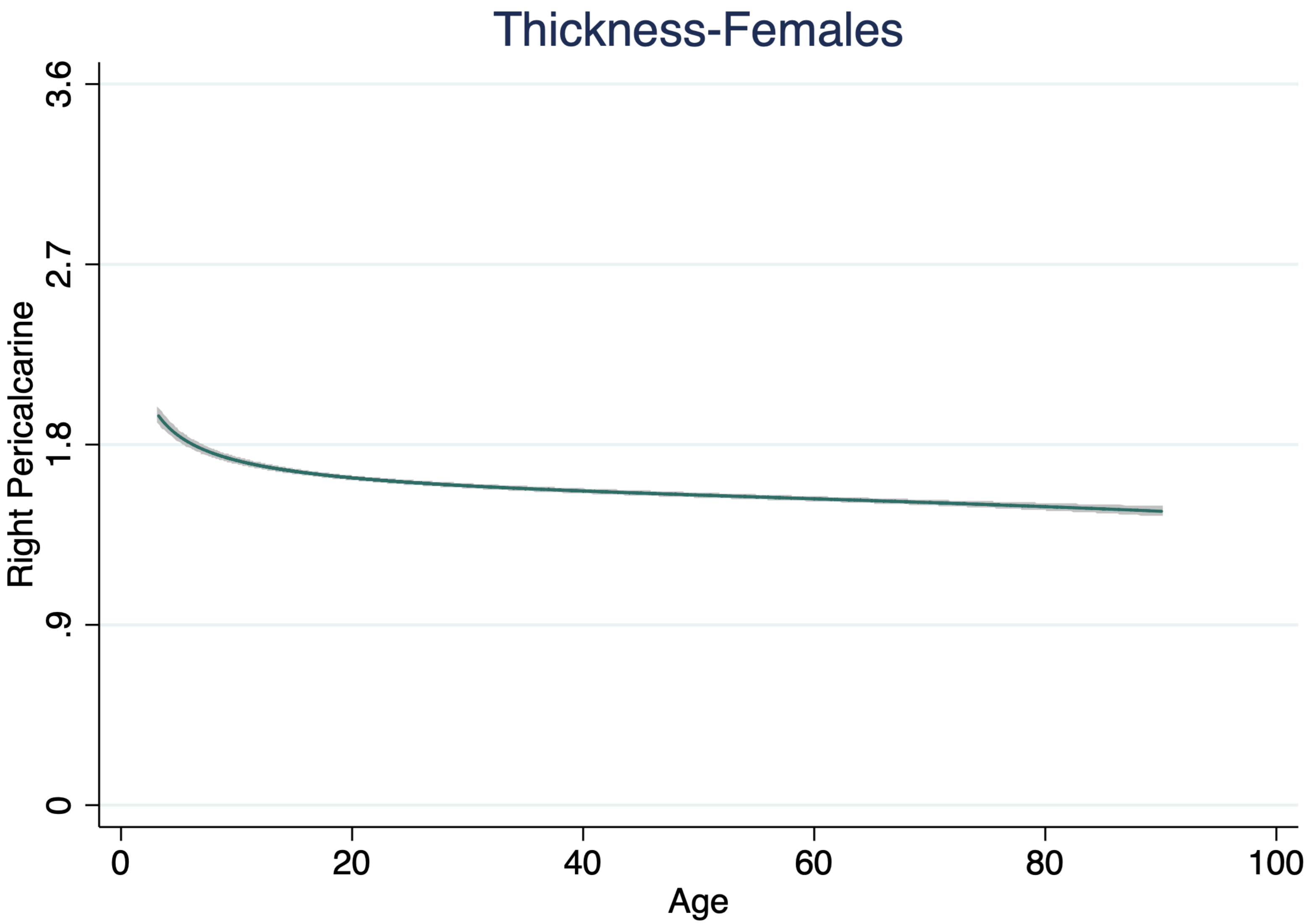

Thickness-Females
Right Pericalcarine
3.6
2.7
1.8
.9
0
0
20
40
60
80
100
Age

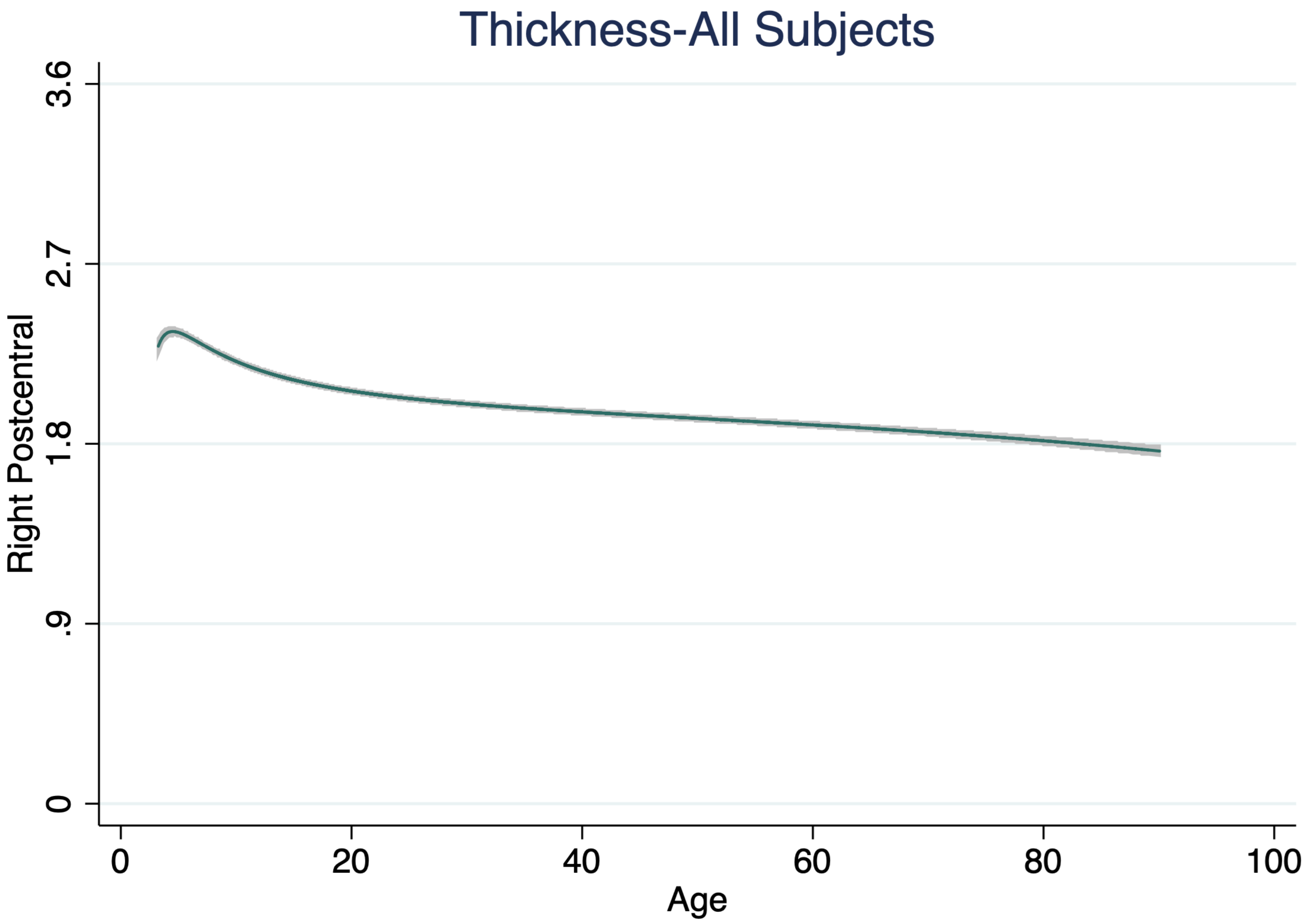
Thickness-All Subjects
Right Postcentral
3.6
2.7
1.8
.9
0
0
20
40
60
80
100
Age

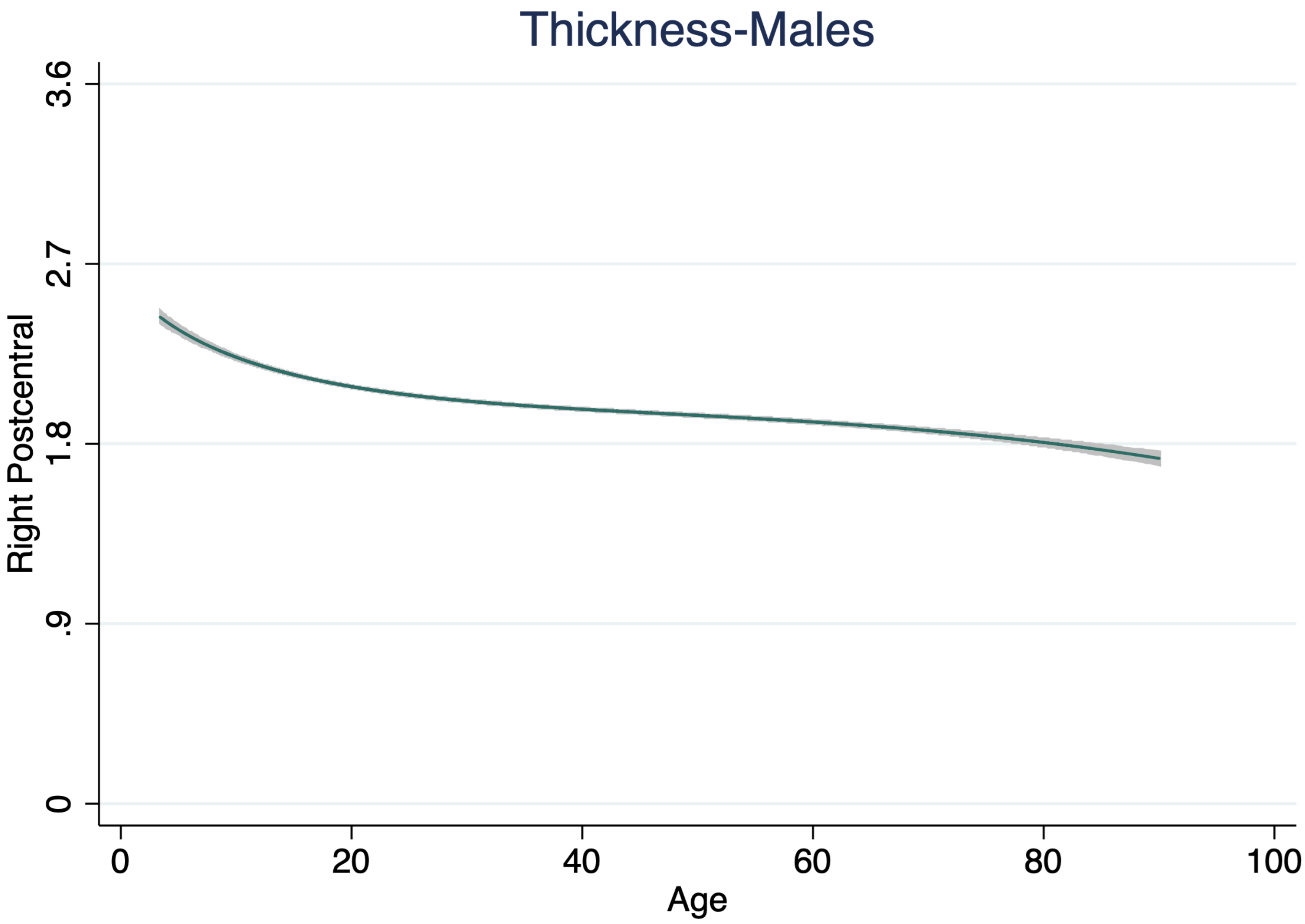
Thickness-Males
Right Postcentral
3.6
2.7
1.8
.9
0
0
20
40
60
80
100
Age

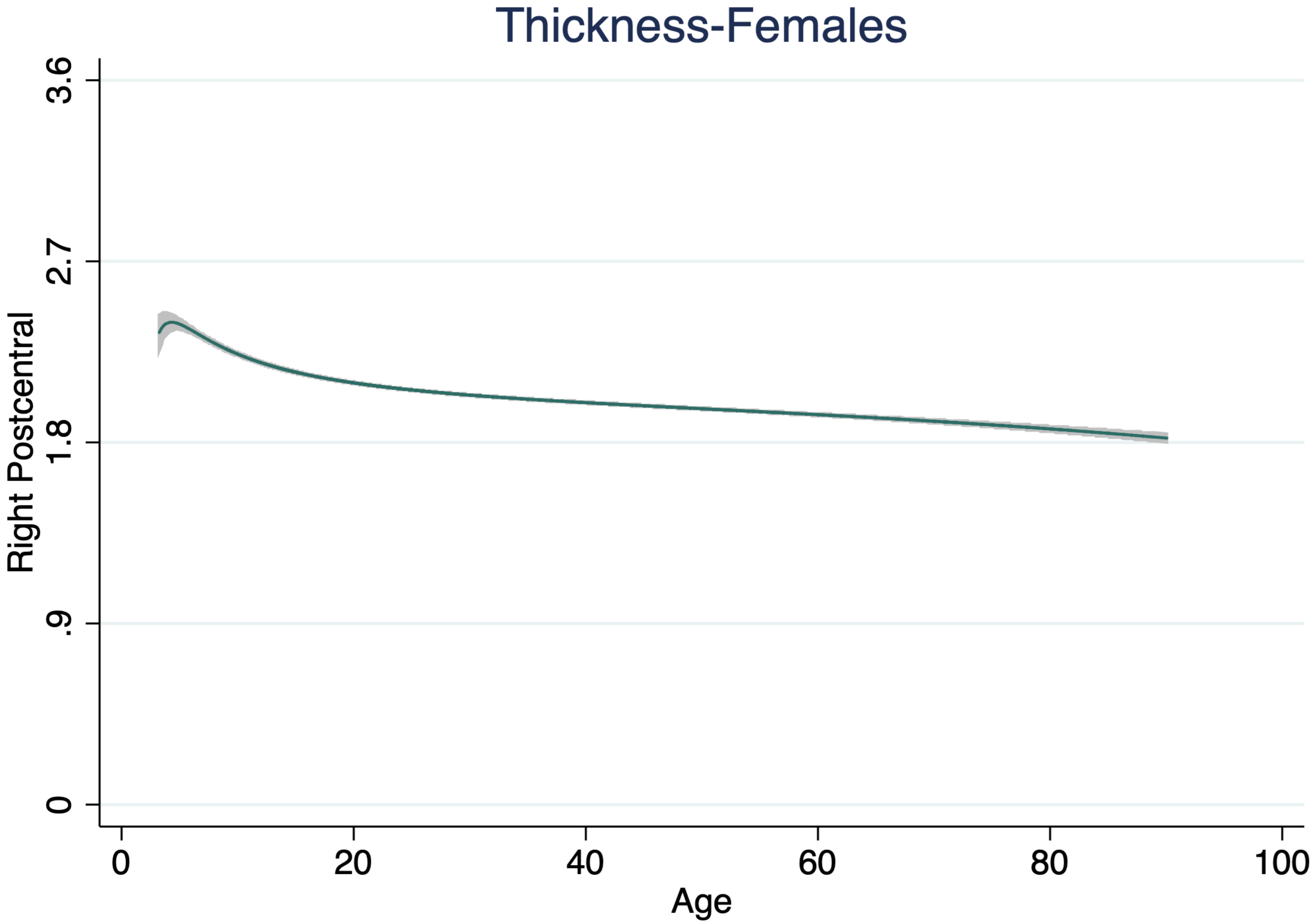
Thickness-Females
Right Postcentral
3.6
2.7
1.8
.9
0
0
20
40
60
80
100
Age

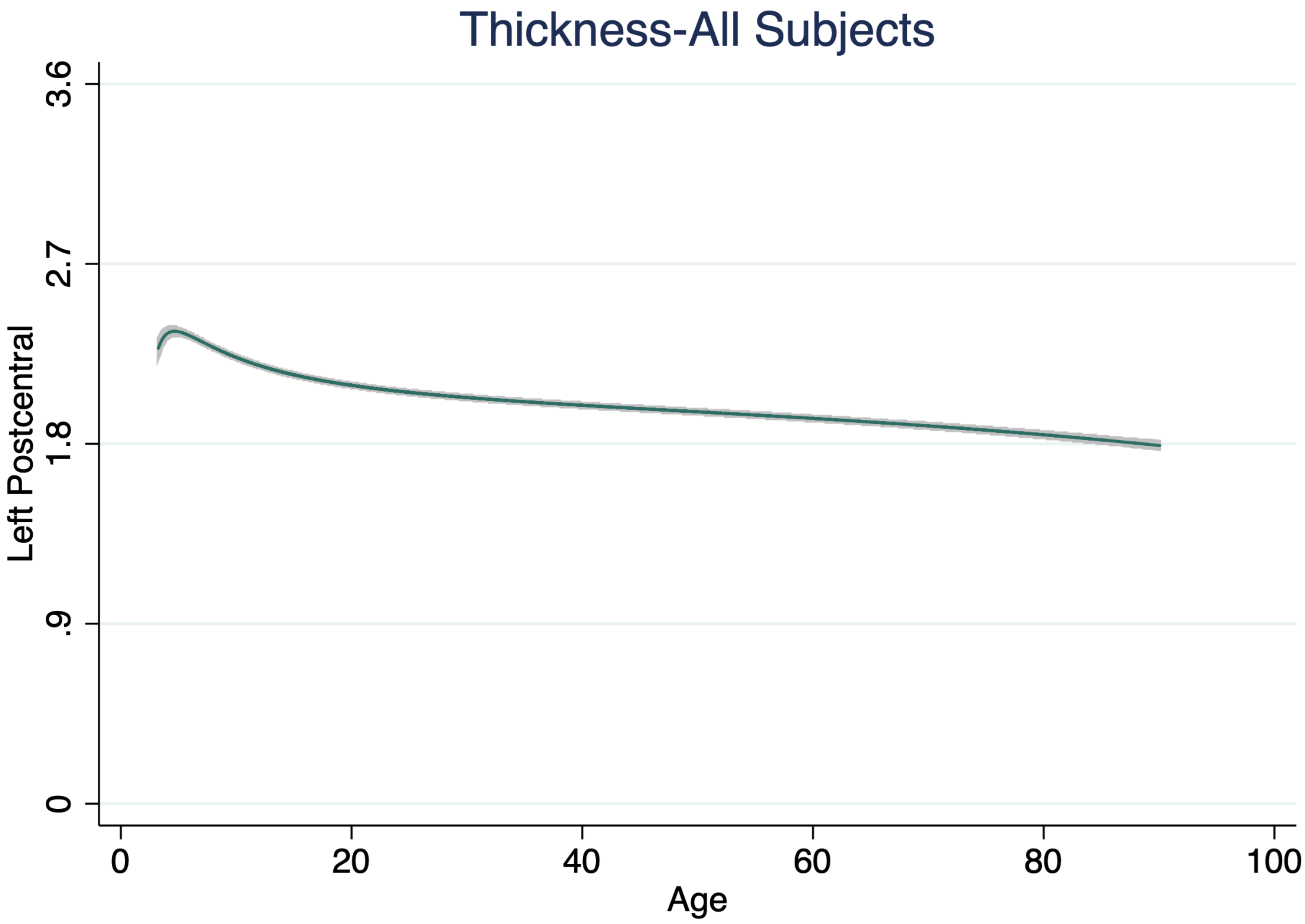

Thickness-All Subjects
Left Postcentral
3.6
2.7
1.8
.9
0
0
20
40
60
80
100
Age

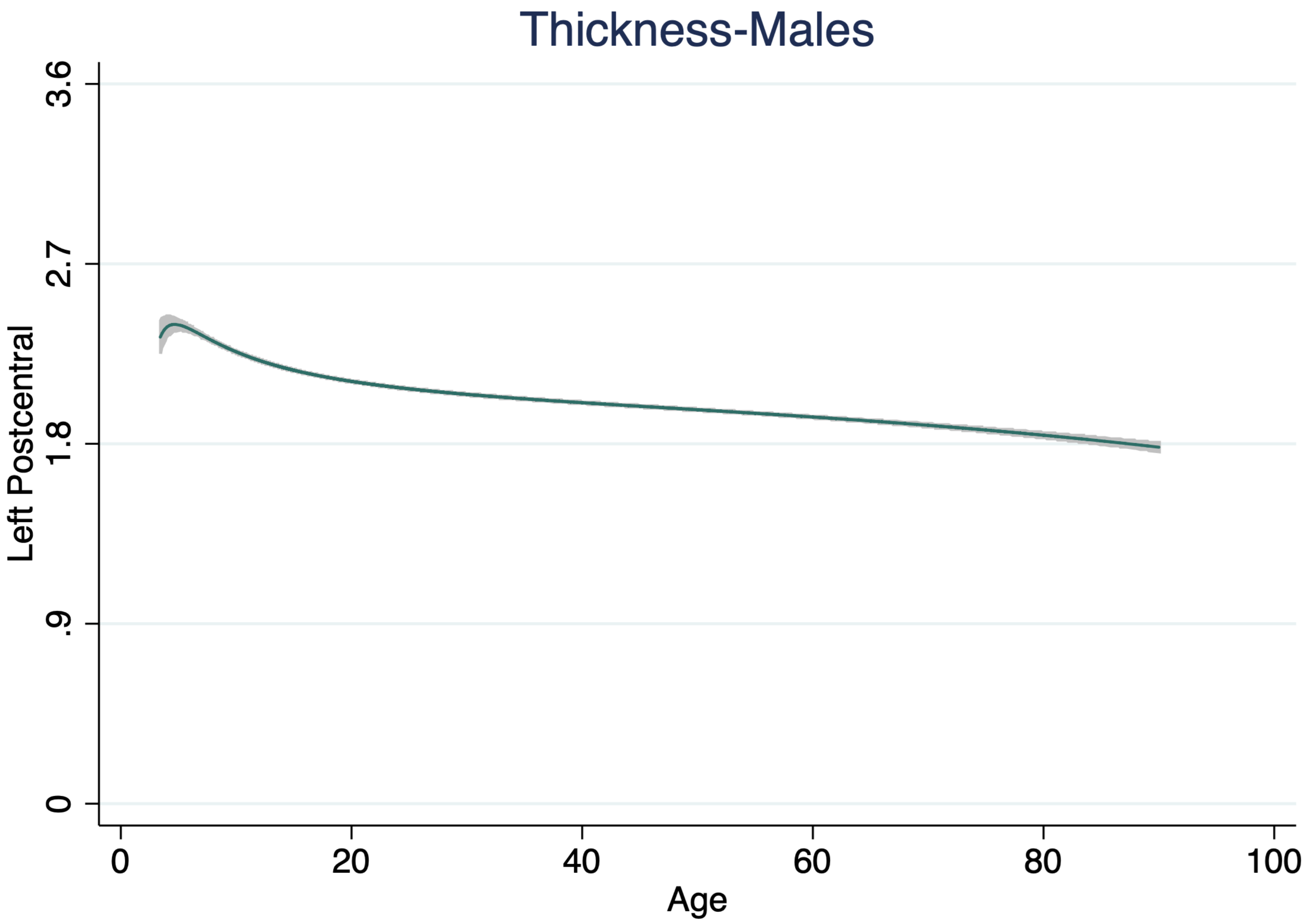
Thickness-Males
Left Postcentral
3.6
2.7
1.8
.9
0
0
20
40
60
80
100
Age

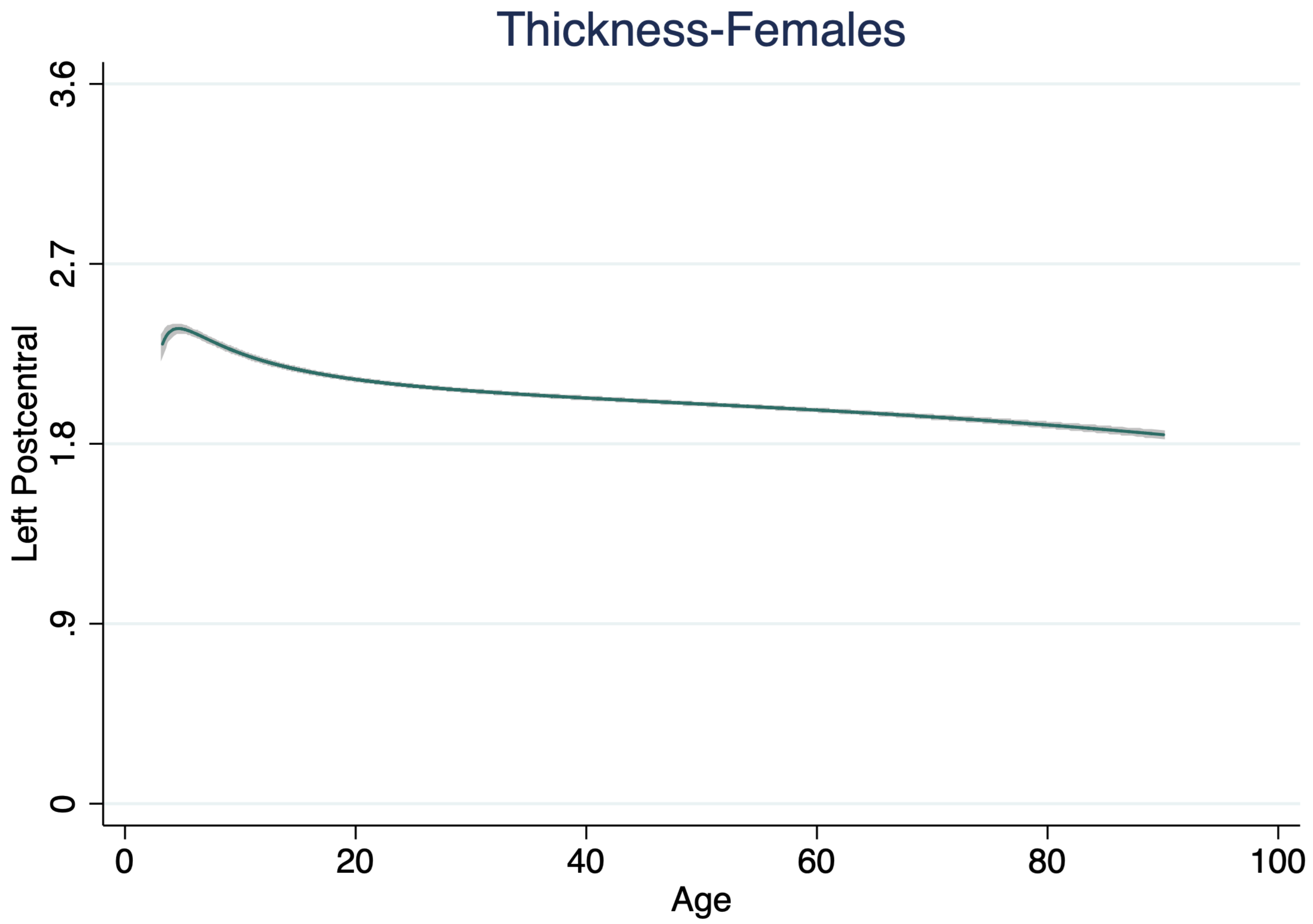

Thickness-Females
Left Postcentral
3.6
2.7
1.8
.9
0
0
20
40
60
80
100
Age

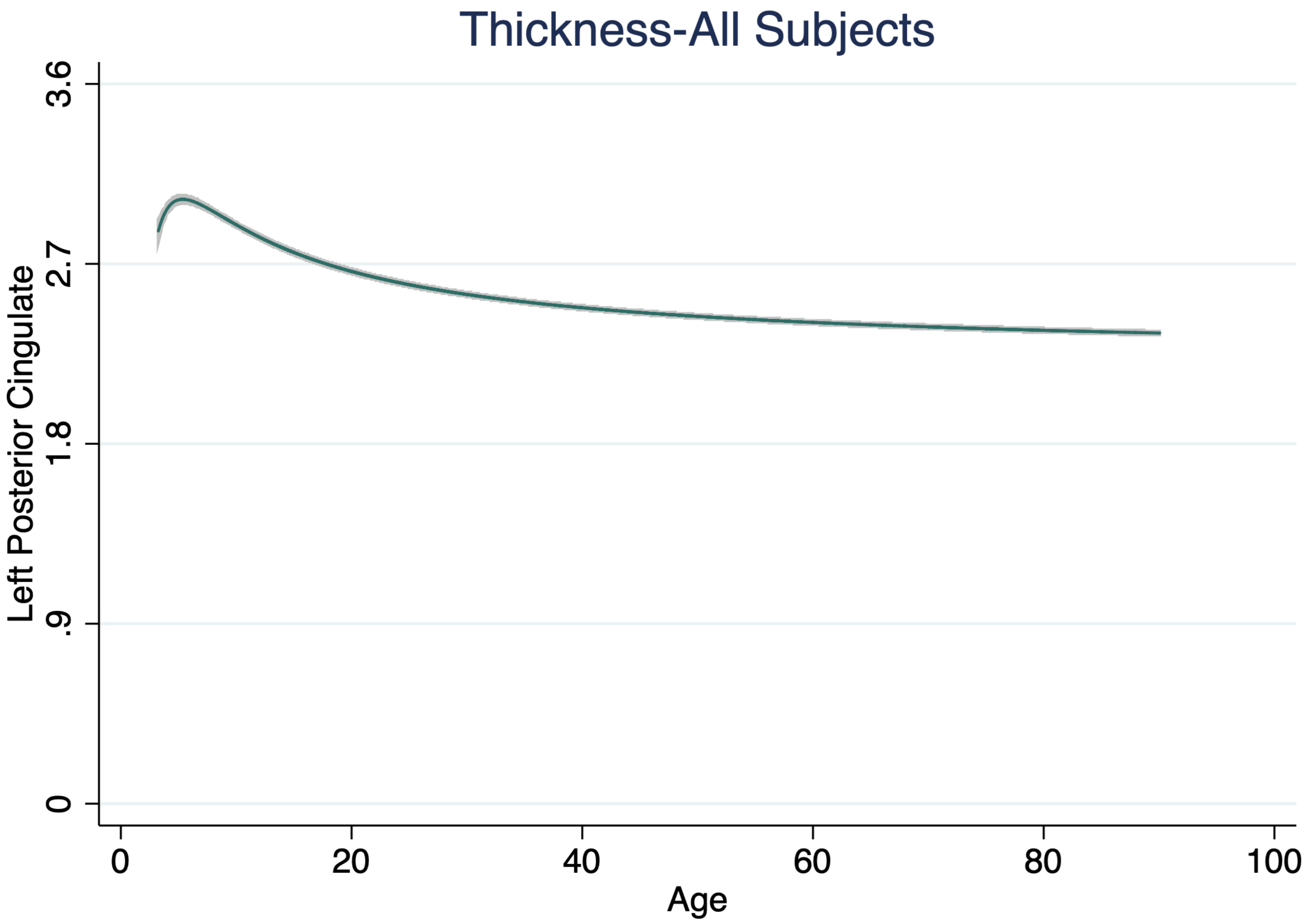

Thickness-All Subjects
Left Posterior Cingulate
3.6
2.7
1.8
.9
0
0
20
40
60
80
100
Age

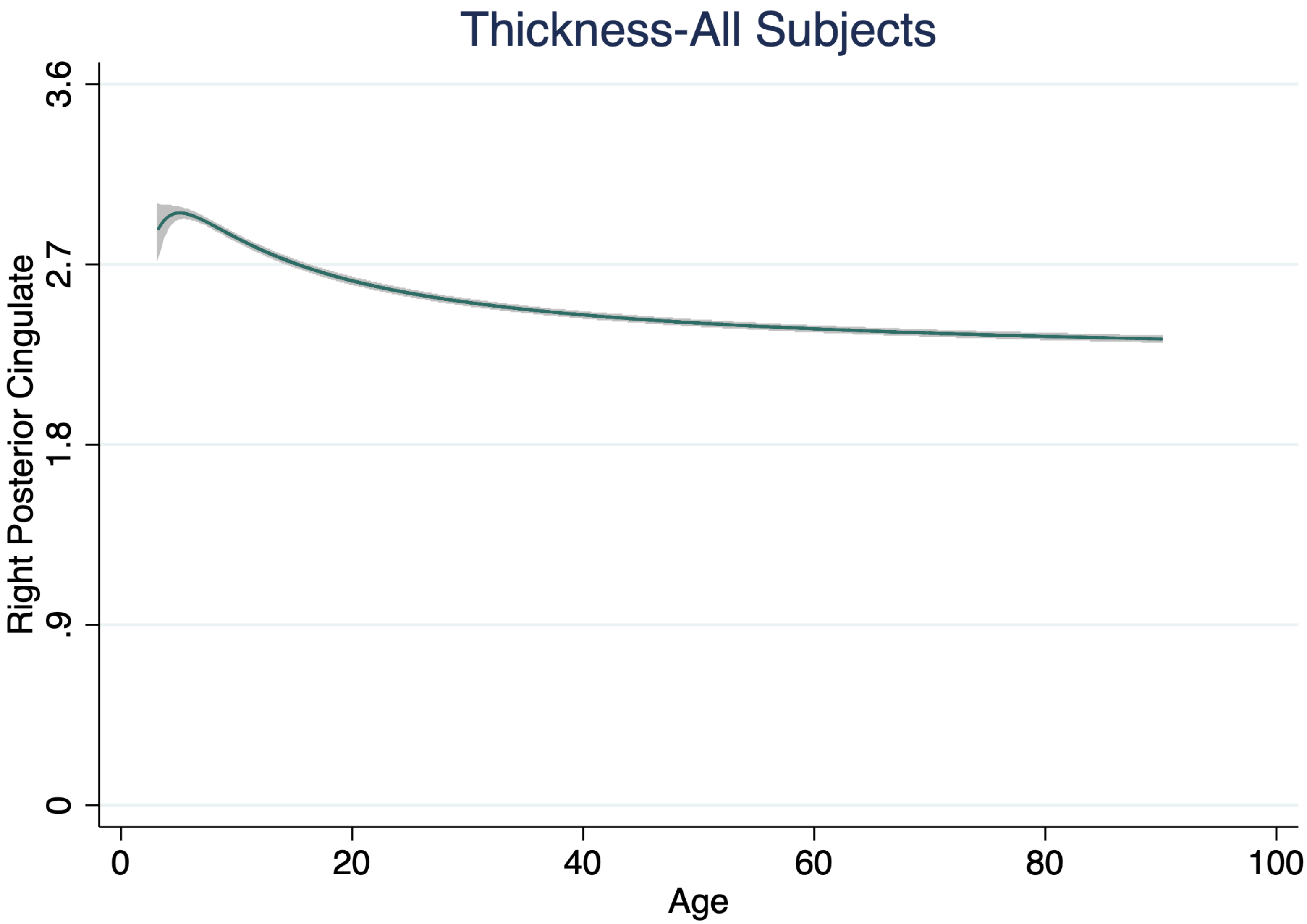

Thickness-All Subjects
Right Posterior Cingulate
3.6
2.7
1.8
.9
0
0
20
40
60
80
100
Age

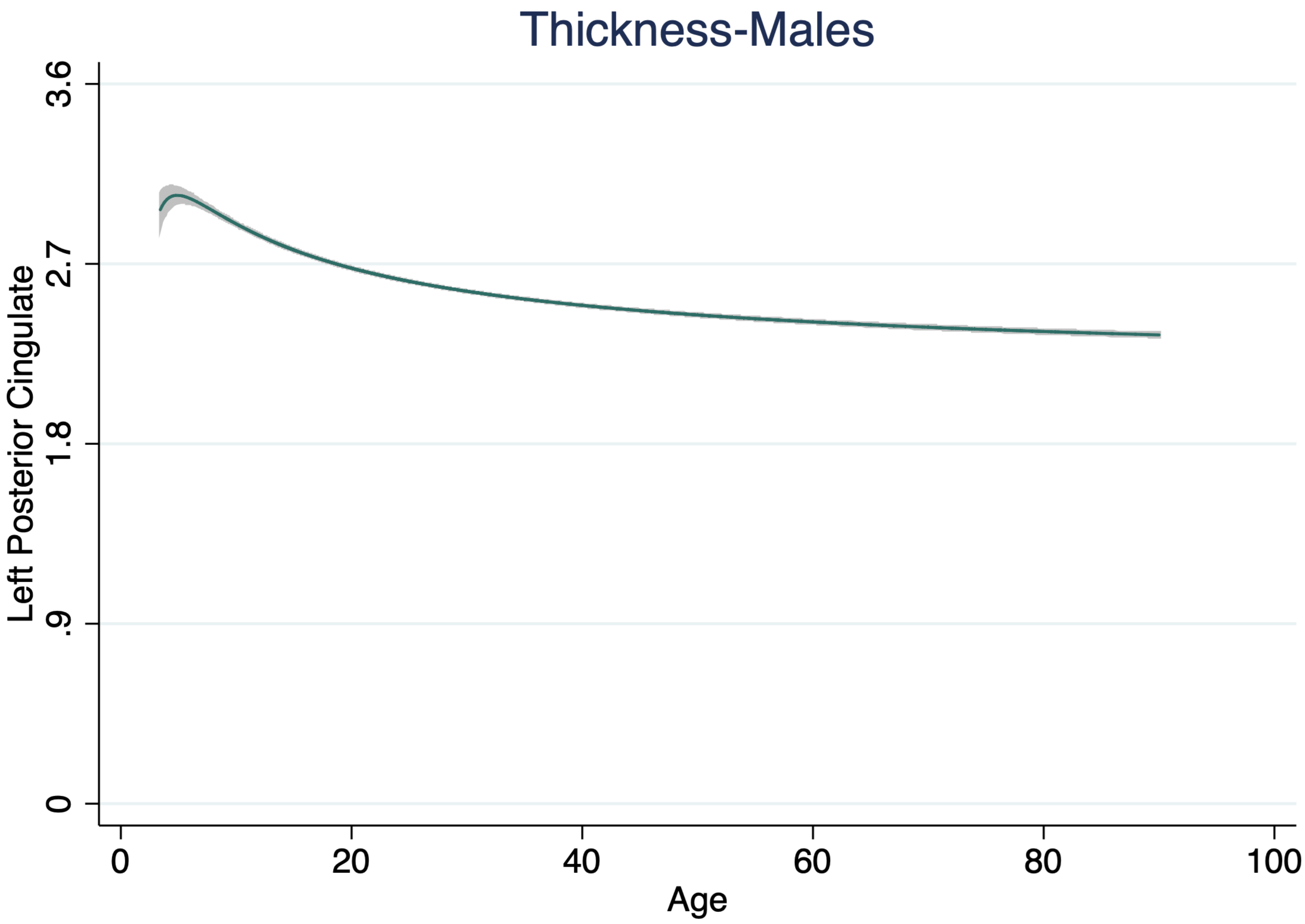
Thickness-Males
Left Posterior Cingulate
3.6
2.7
1.8
.9
0
0
20
40
60
80
100
Age

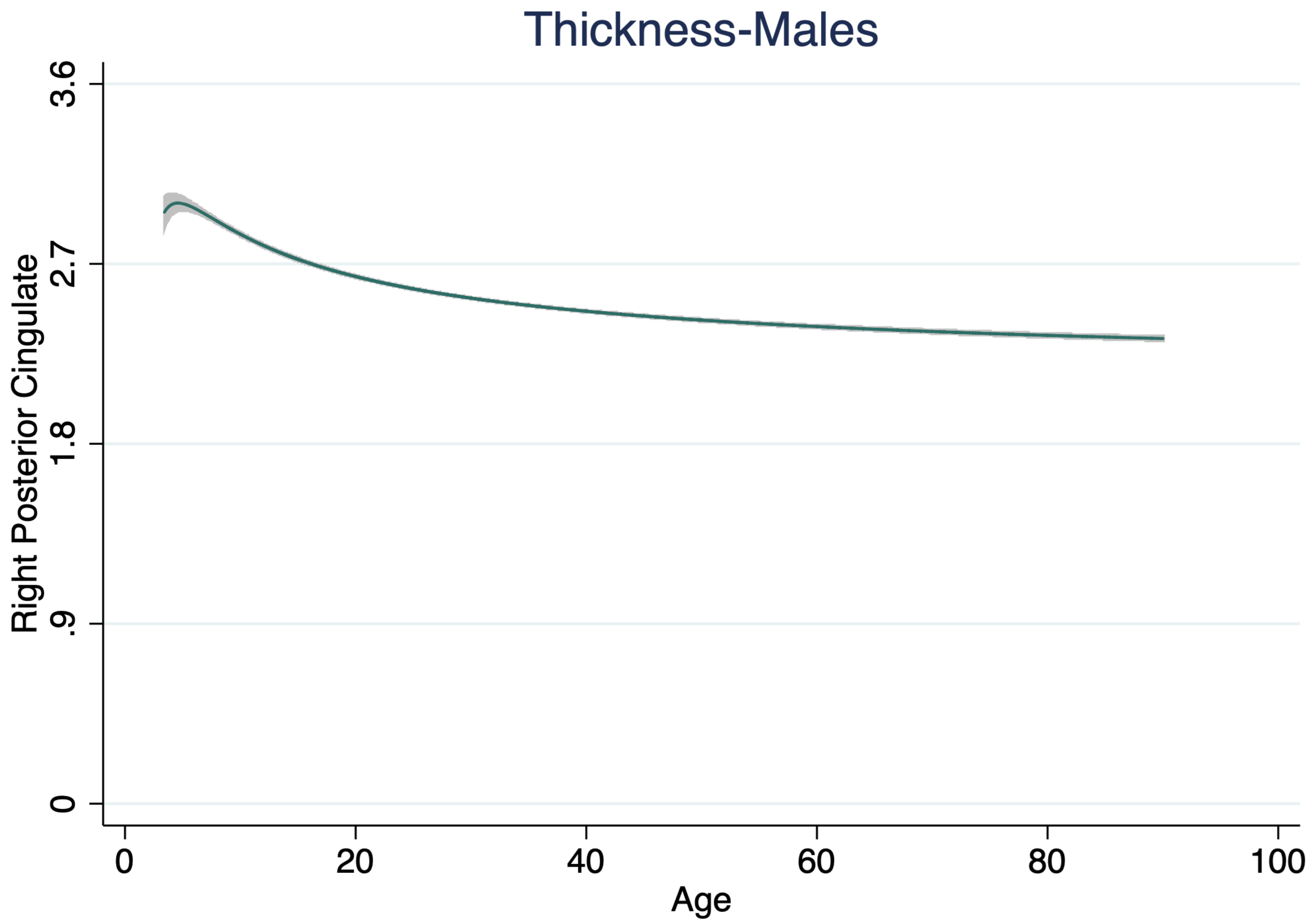
Thickness-Males
Right Posterior Cingulate
3.6
2.7
1.8
.9
0
0
20
40
60
80
100
Age

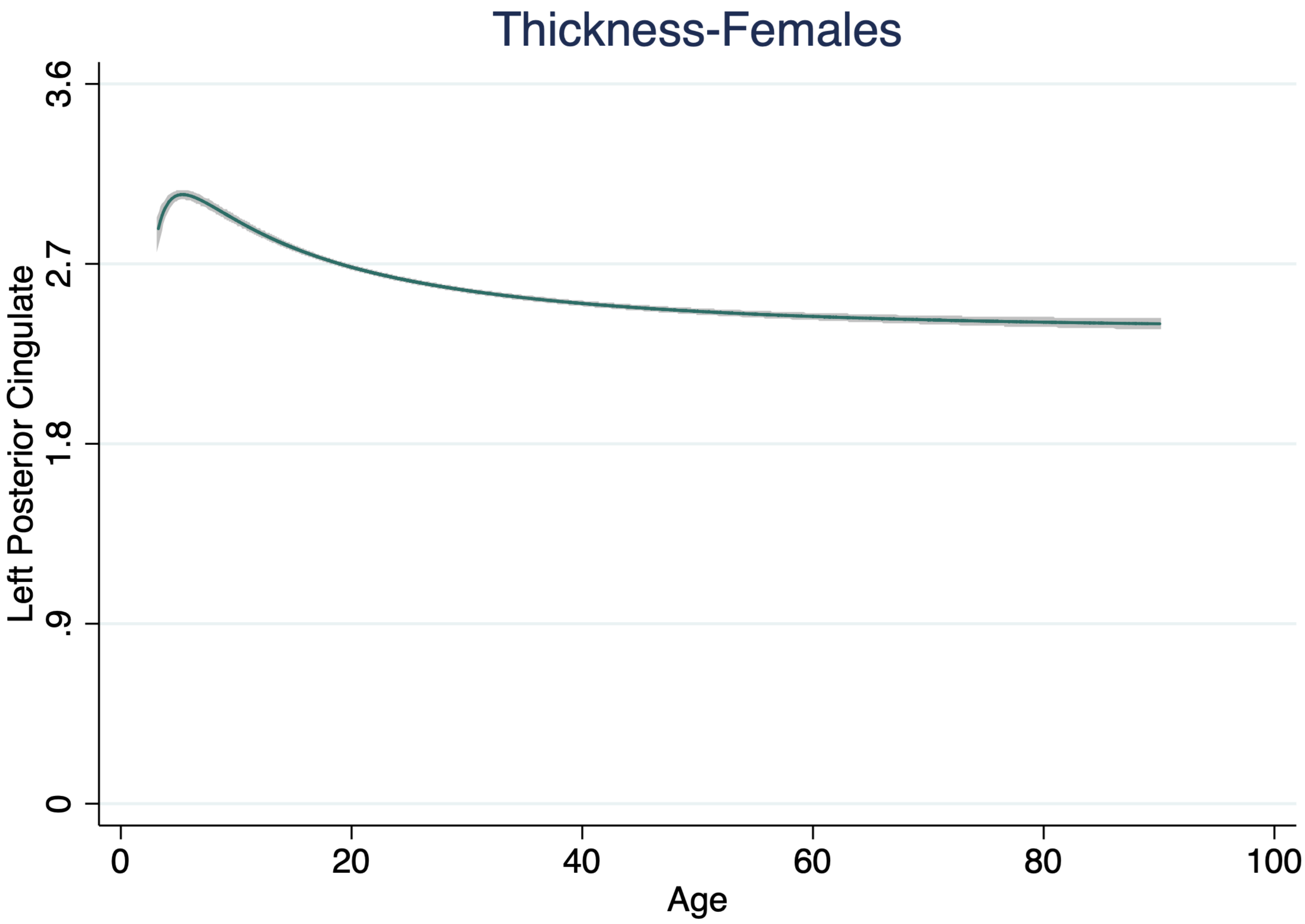

Thickness-Females
Left Posterior Cingulate
3.6
2.7
1.8
.9
0
0
20
40
60
80
100
Age

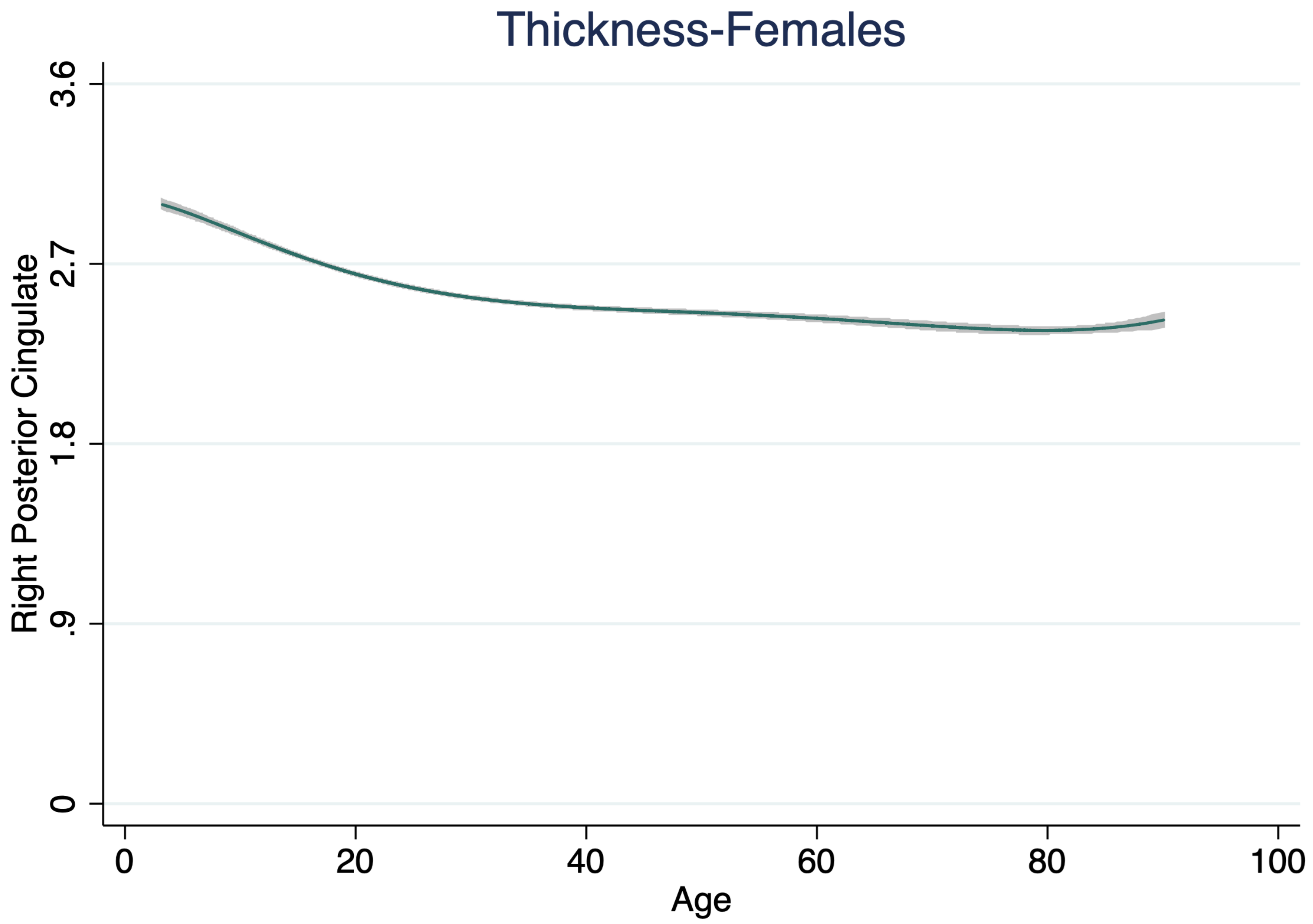
Thickness-Females
Right Posterior Cingulate
3.6
2.7
1.8
.9
0
0
20
40
60
80
100
Age

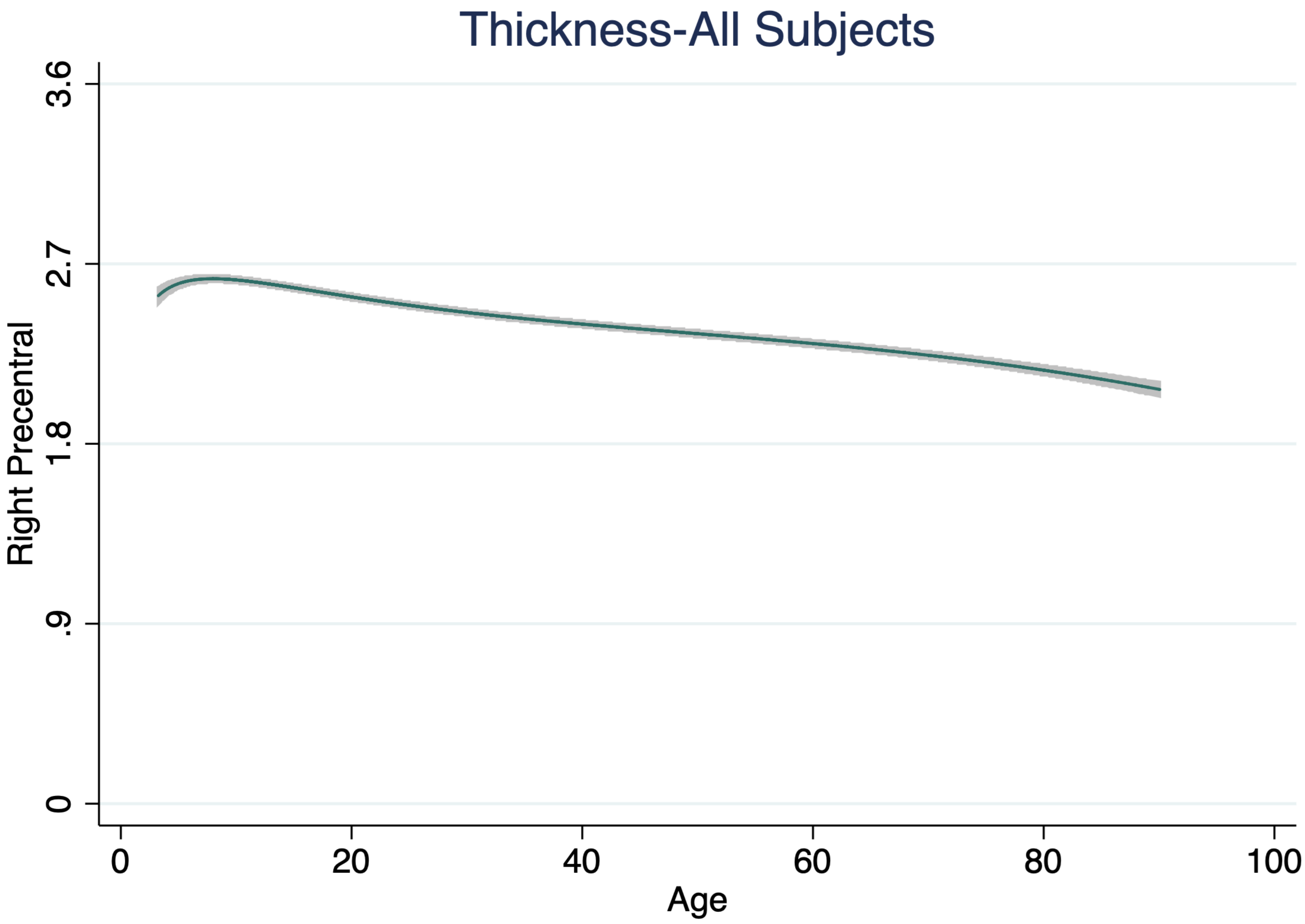

Thickness-All Subjects
Right Precentral
3.6
2.7
1.8
.9
0
0
20
40
60
80
100
Age

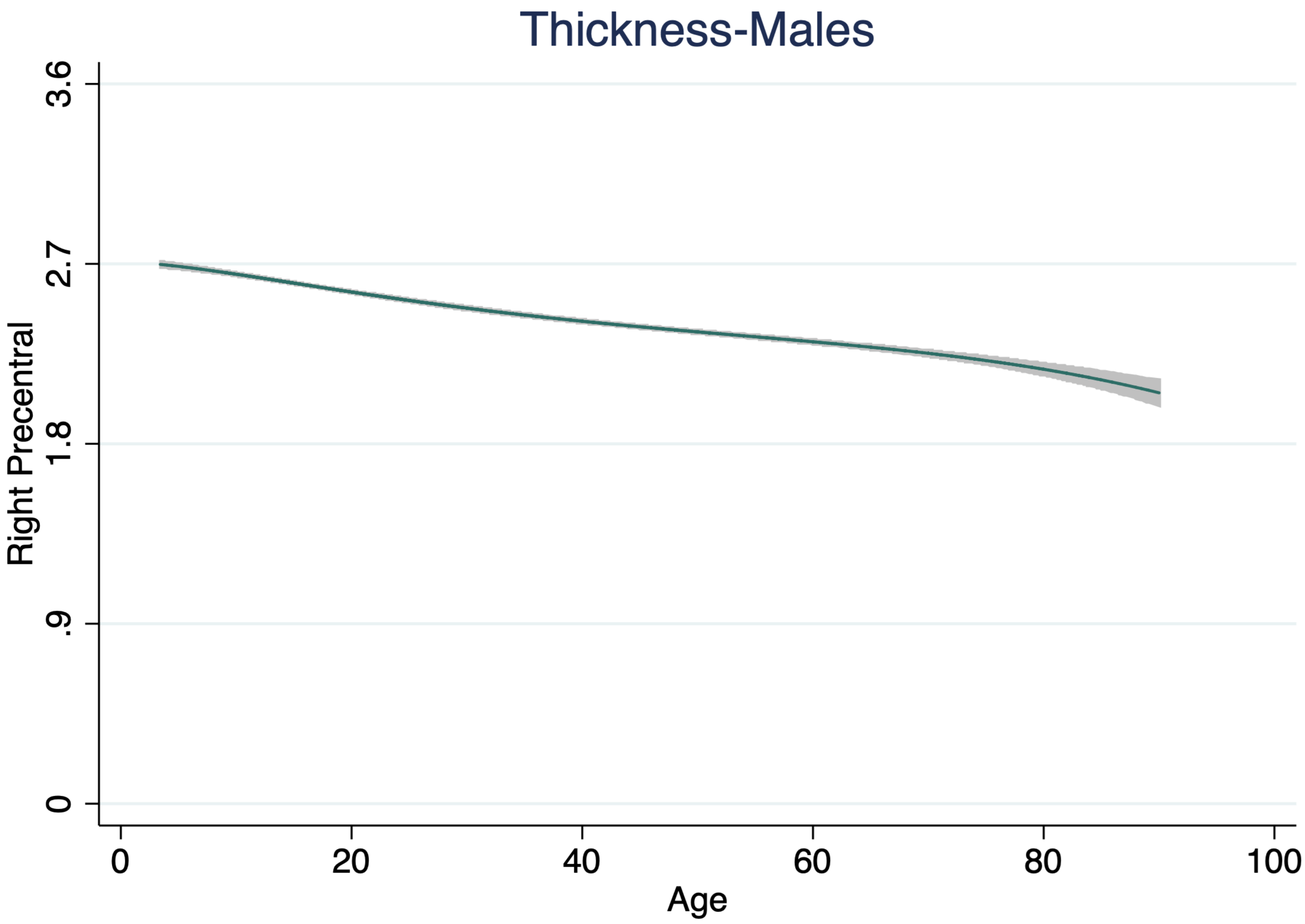

Thickness-Males
Right Precentral
3.6
2.7
1.8
.9
0
0
20
40
60
80
100
Age

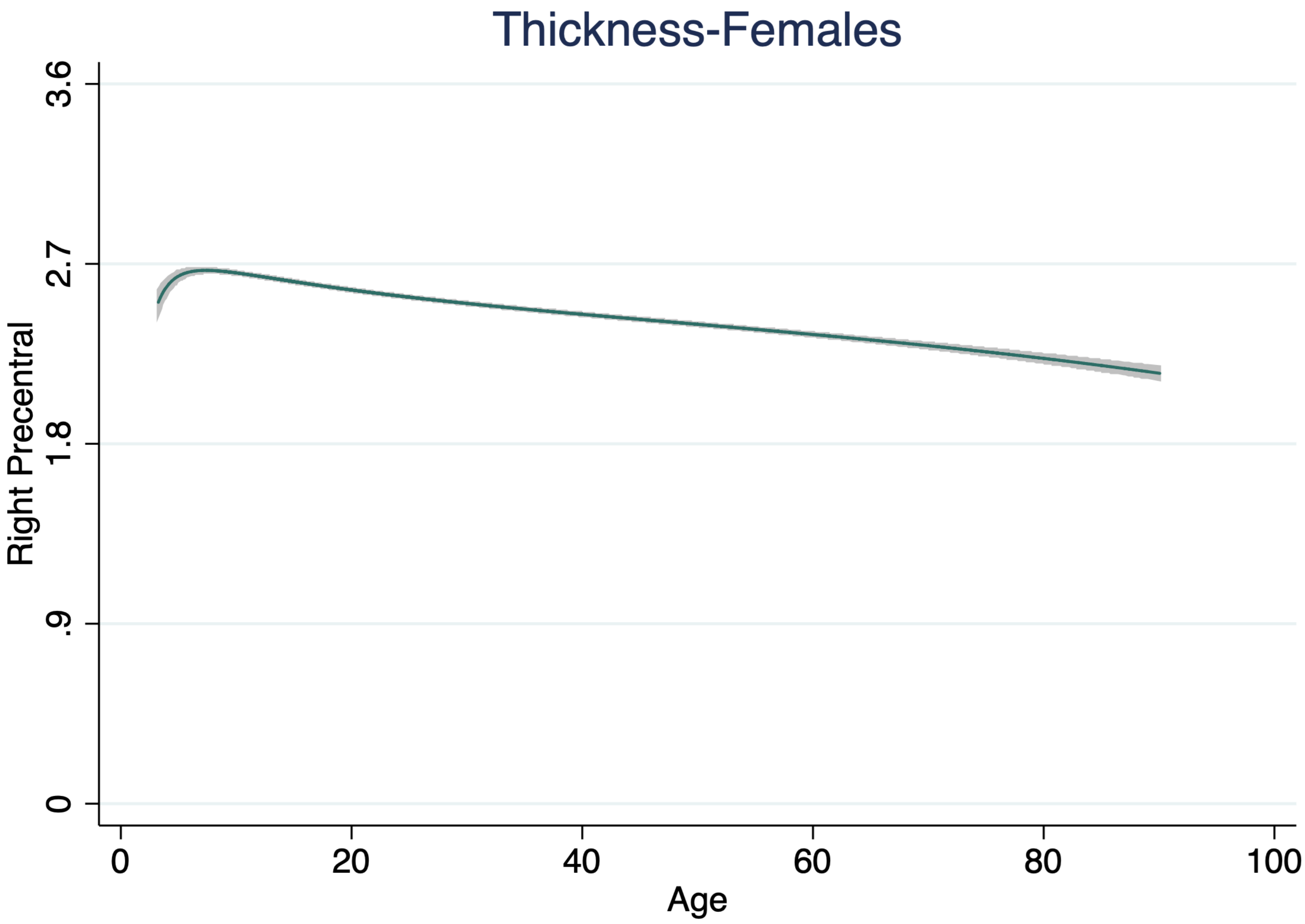

Thickness-Females
Right Precentral
3.6
2.7
1.8
.9
0
0
20
40
60
80
100
Age

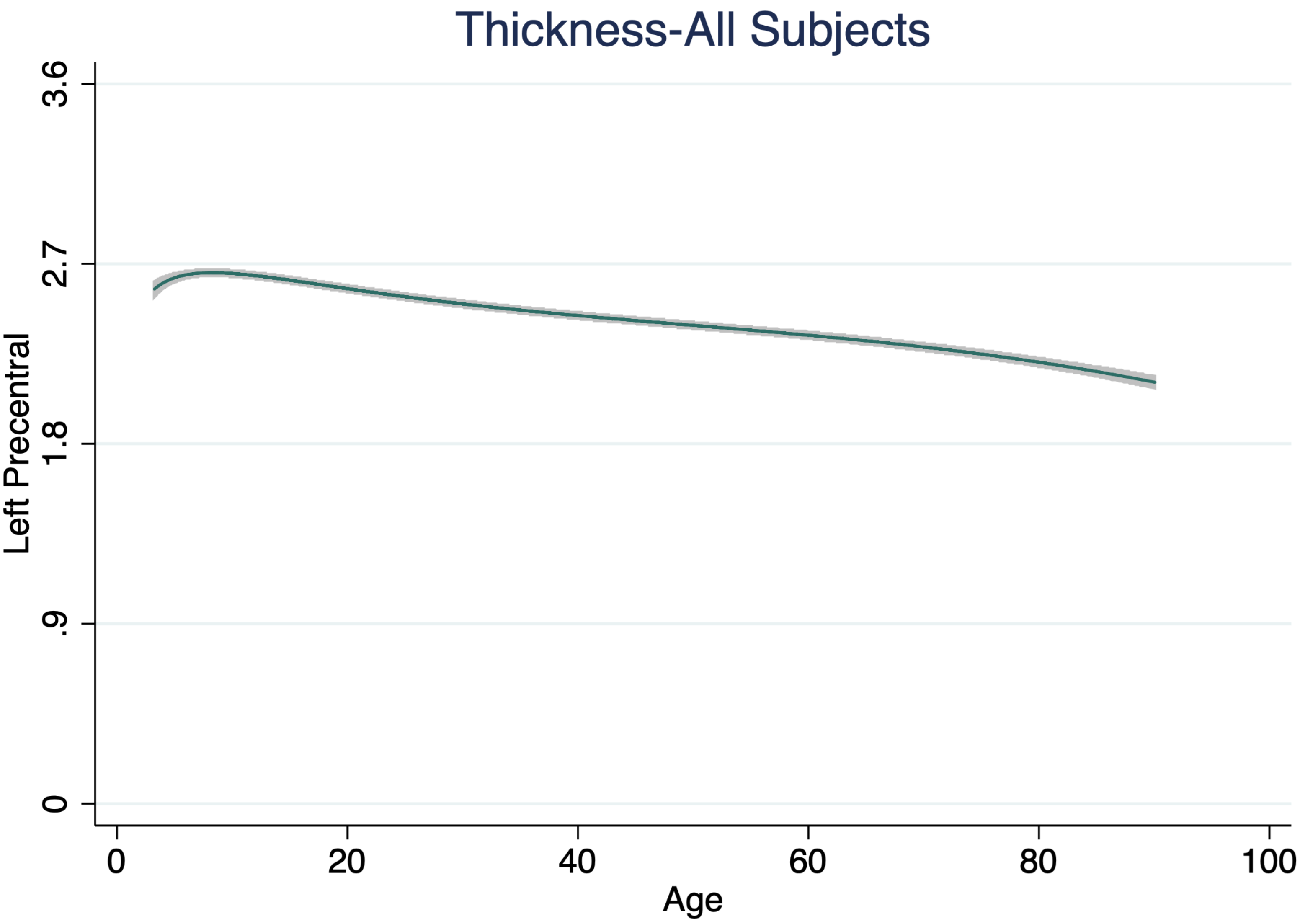

Thickness-All Subjects
Left Precentral
3.6
2.7
1.8
.9
0
0
20
40
60
80
100
Age

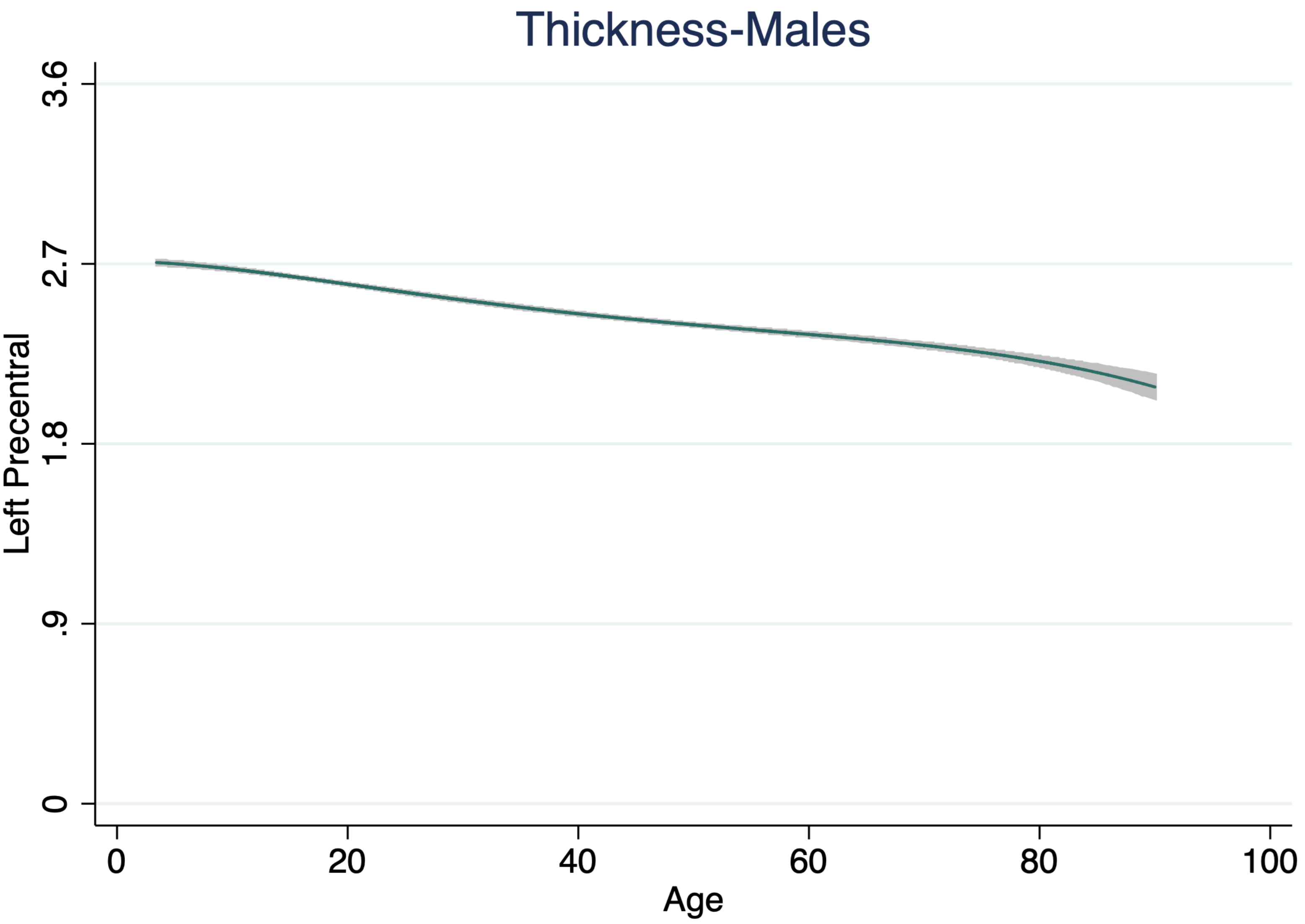
Thickness-Males
Left Precentral
3.6
2.7
1.8
.9
0
0
20
40
60
80
100
Age

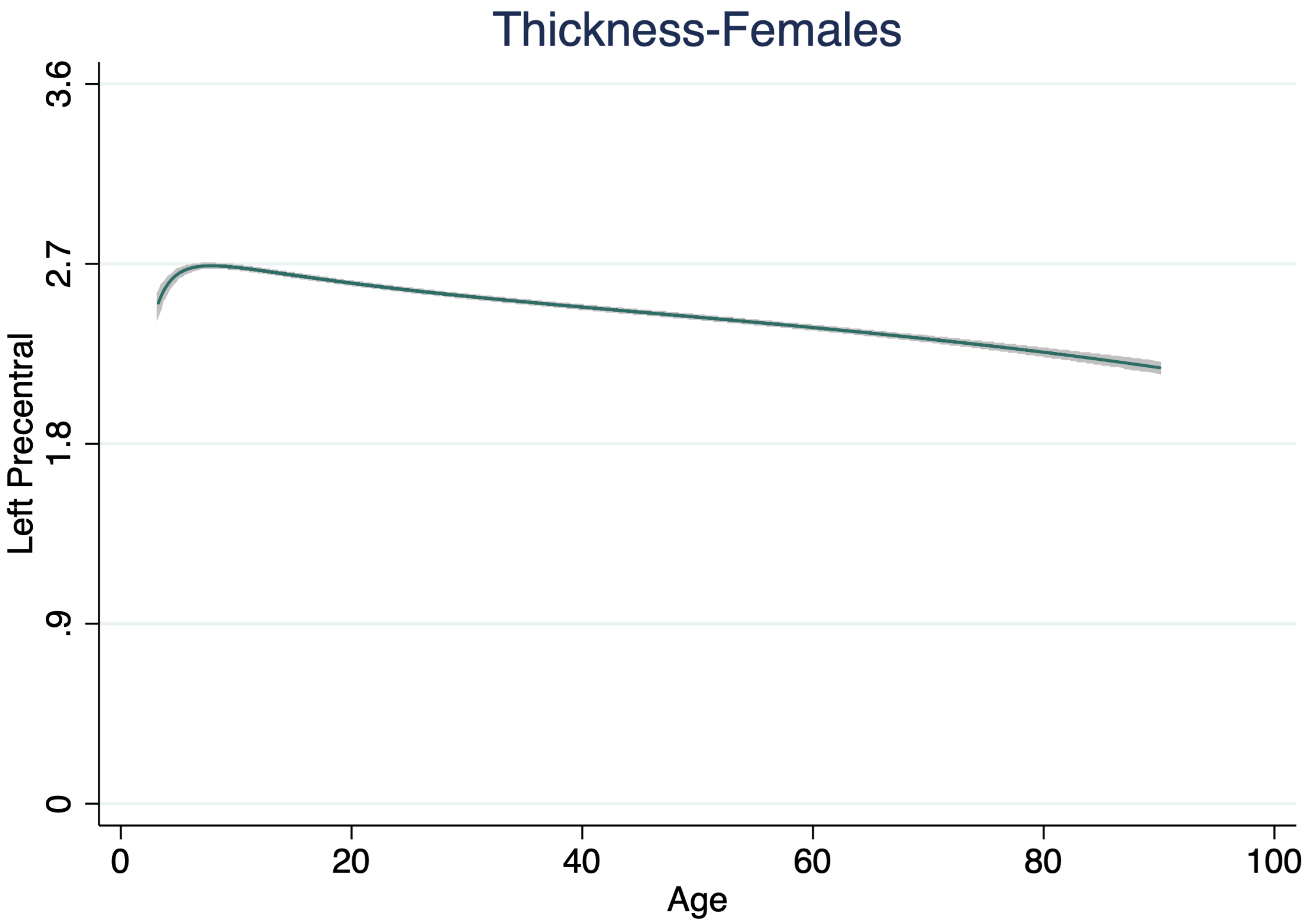

Thickness-Females
Left Precentral
3.6
2.7
1.8
.9
0
0
20
40
60
80
100
Age

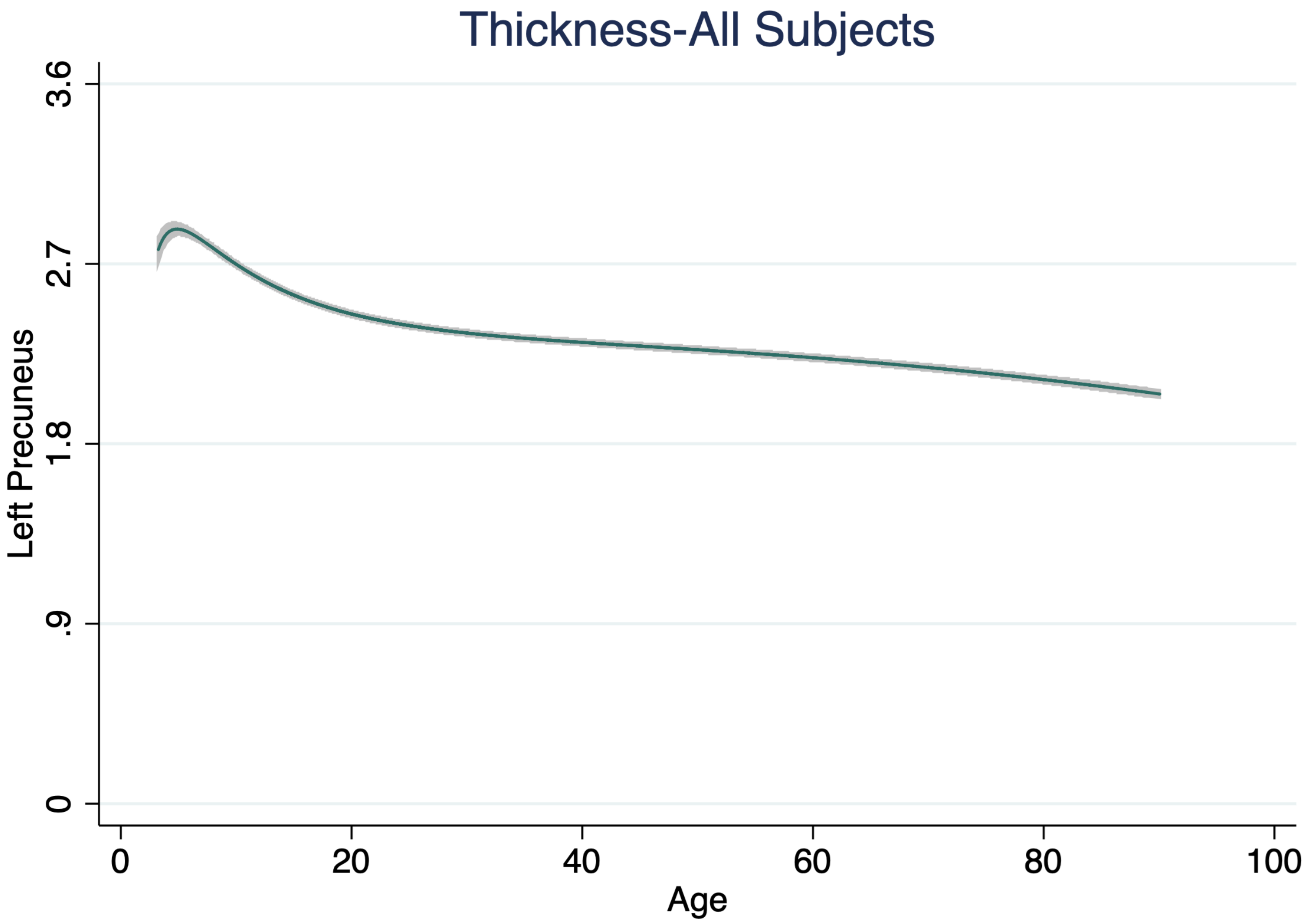

Thickness-All Subjects
Left Precuneus
3.6
2.7
1.8
.9
0
0
20
40
60
80
100
Age

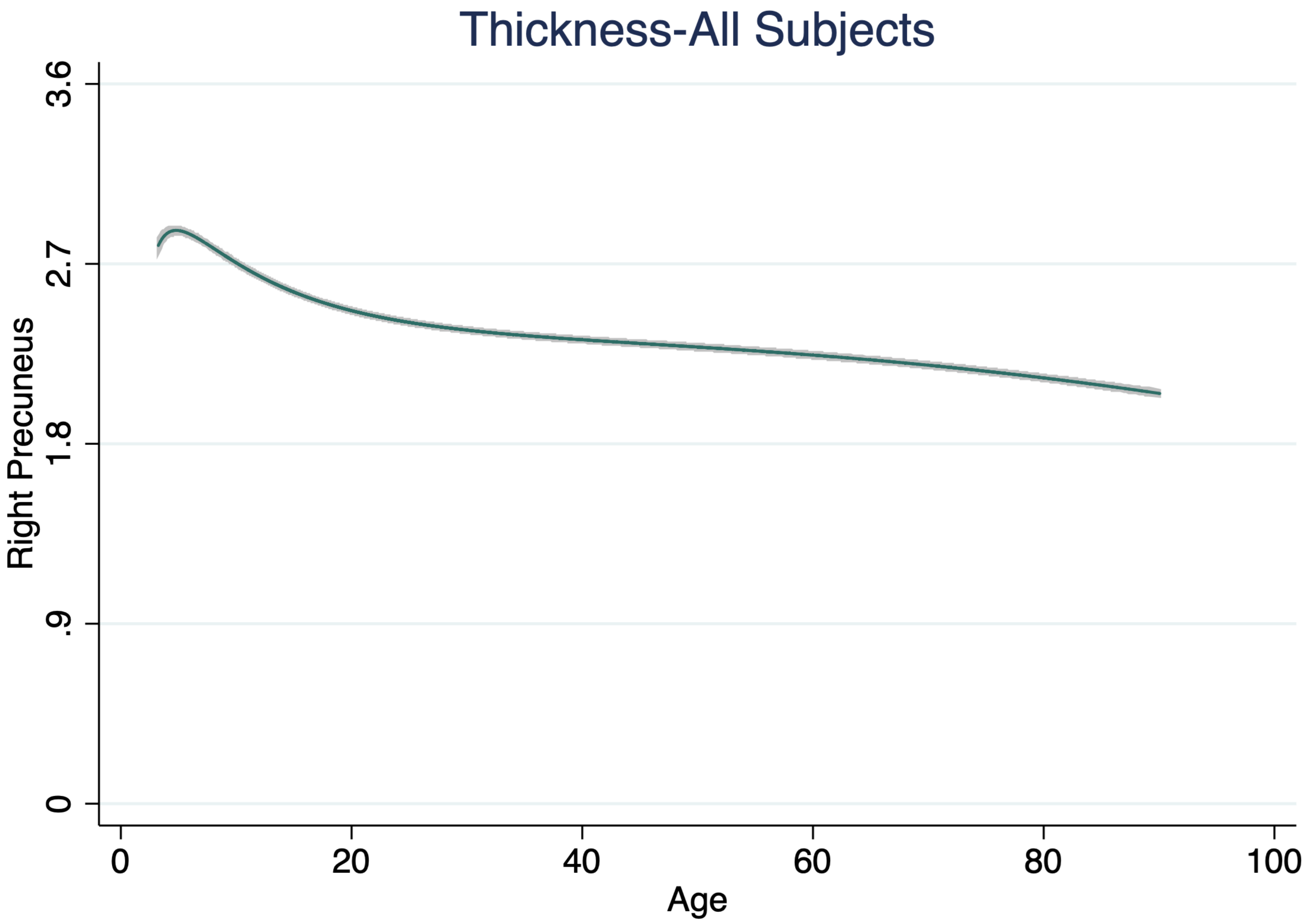

Thickness-All Subjects
Right Precuneus
0
.9
1.8
2.7
3.6
0
20
40
60
80
100
Age

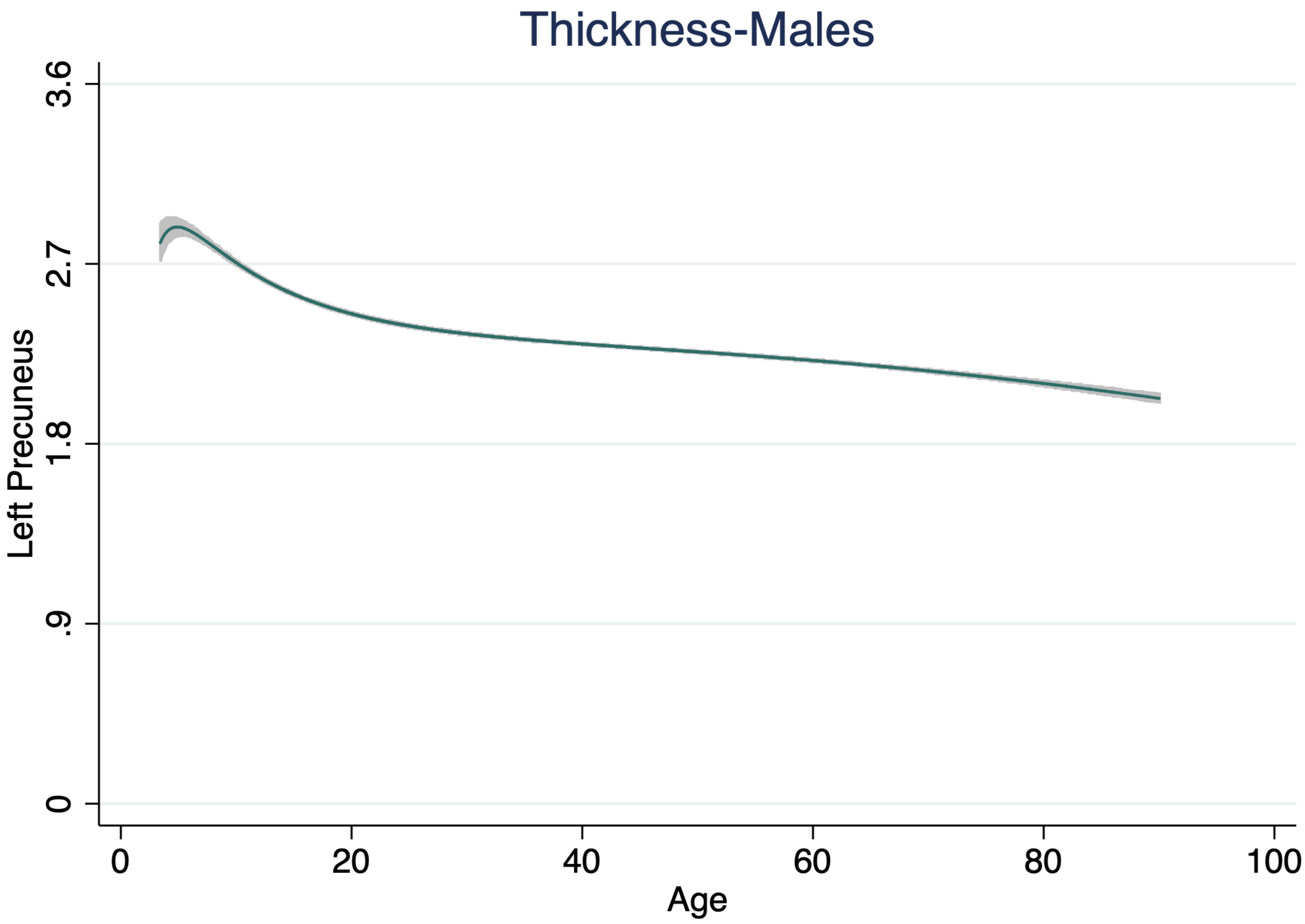

Thickness-Males
Left Precuneus
3.6
2.7
1.8
.9
0
0
20
40
60
80
100
Age

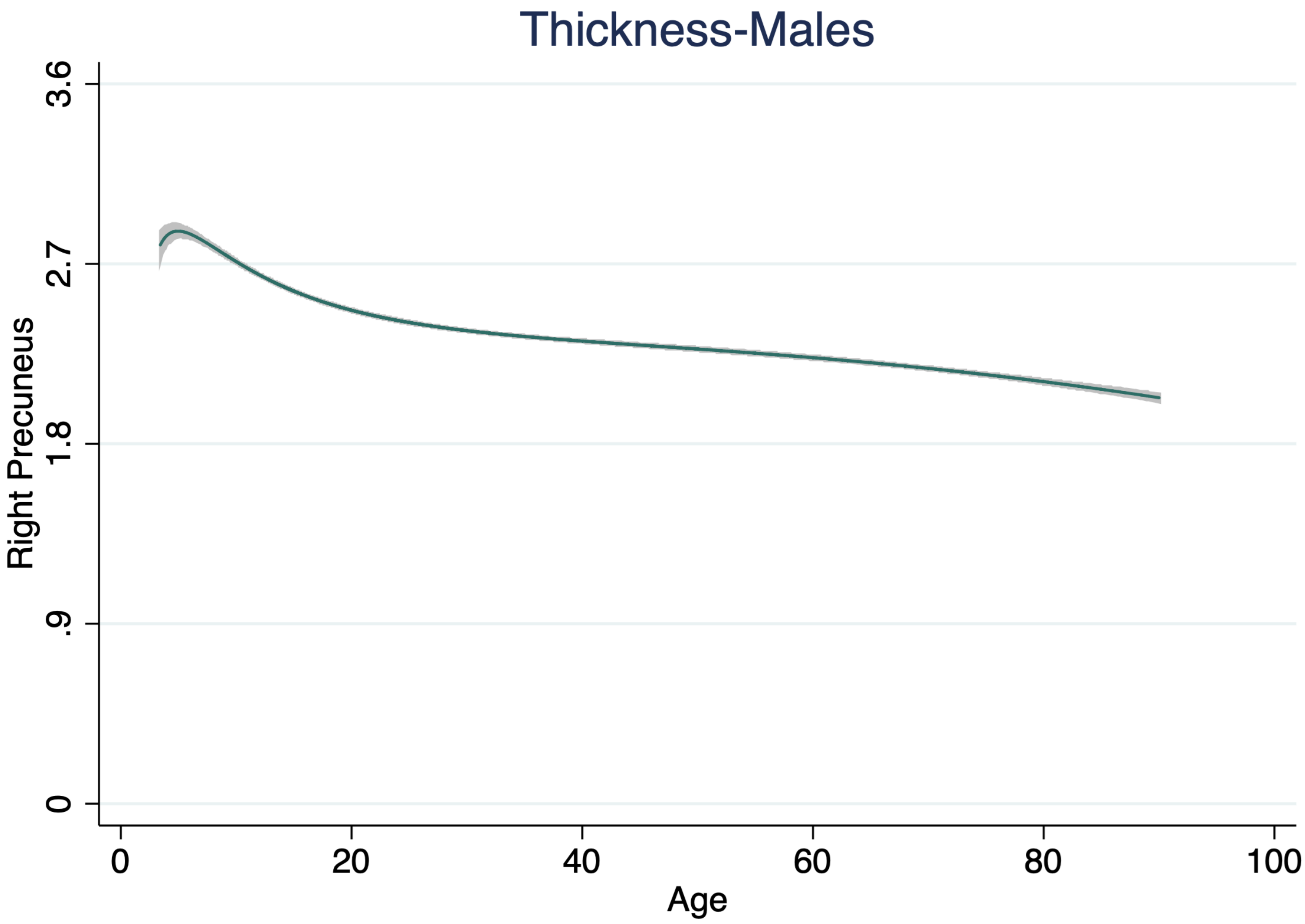

Thickness-Males
Right Precuneus
3.6
2.7
1.8
.9
0
0
20
40
60
80
100
Age

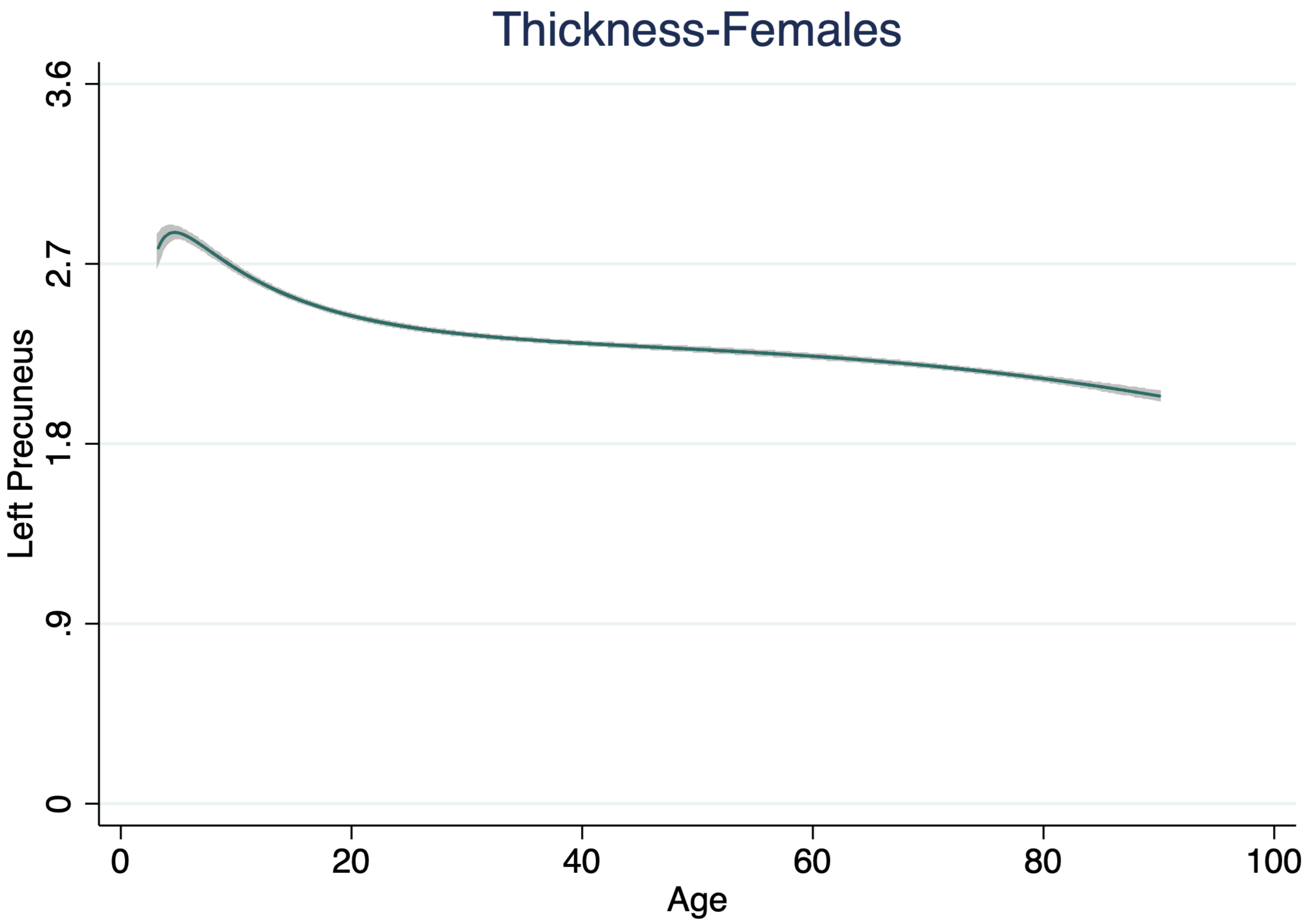

Thickness-Females
Left Precuneus
3.6
2.7
1.8
.9
0
0
20
40
60
80
100
Age

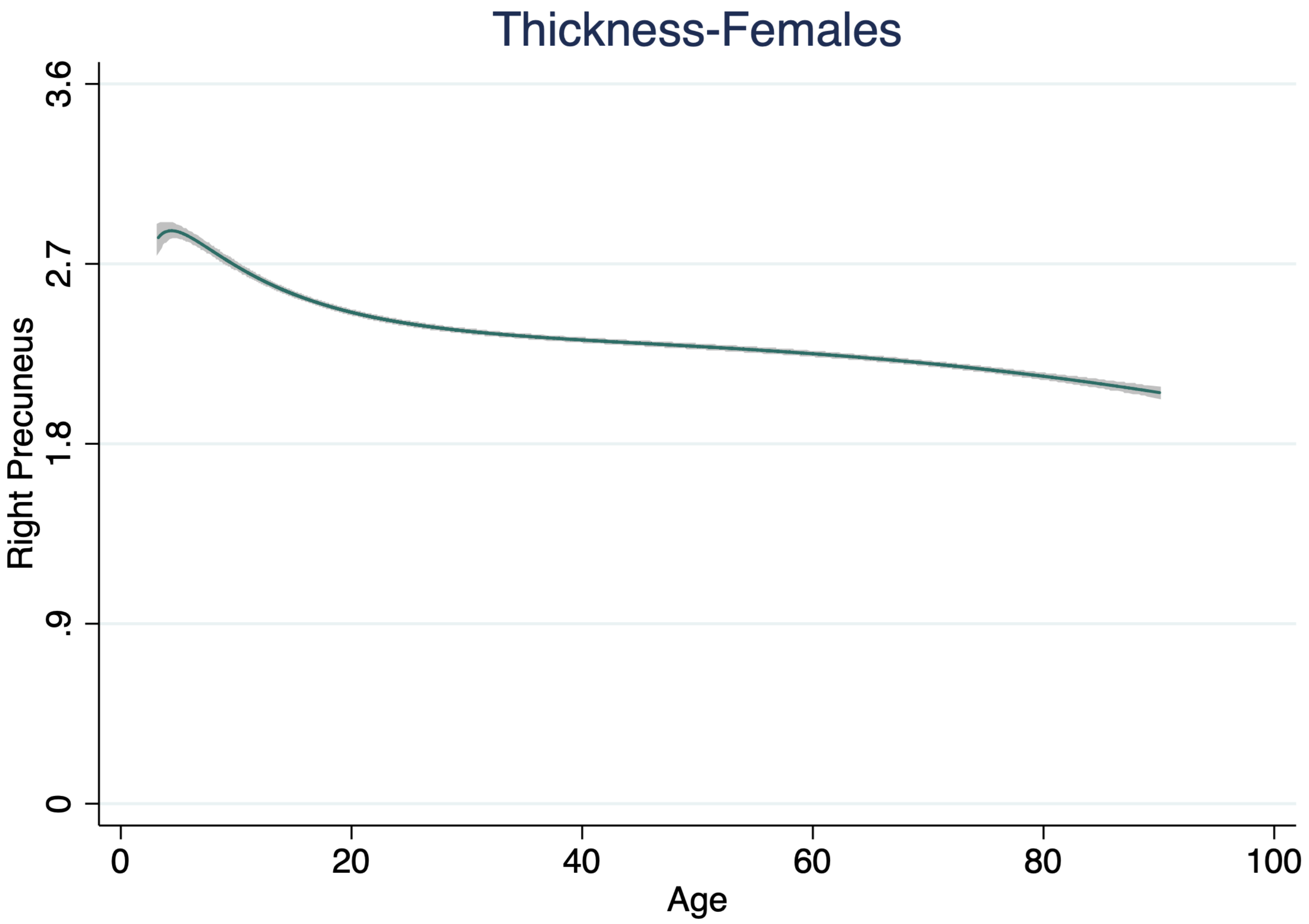
Thickness-Females
Right Precuneus
3.6
2.7
1.8
.9
0
0
20
40
60
80
100
Age

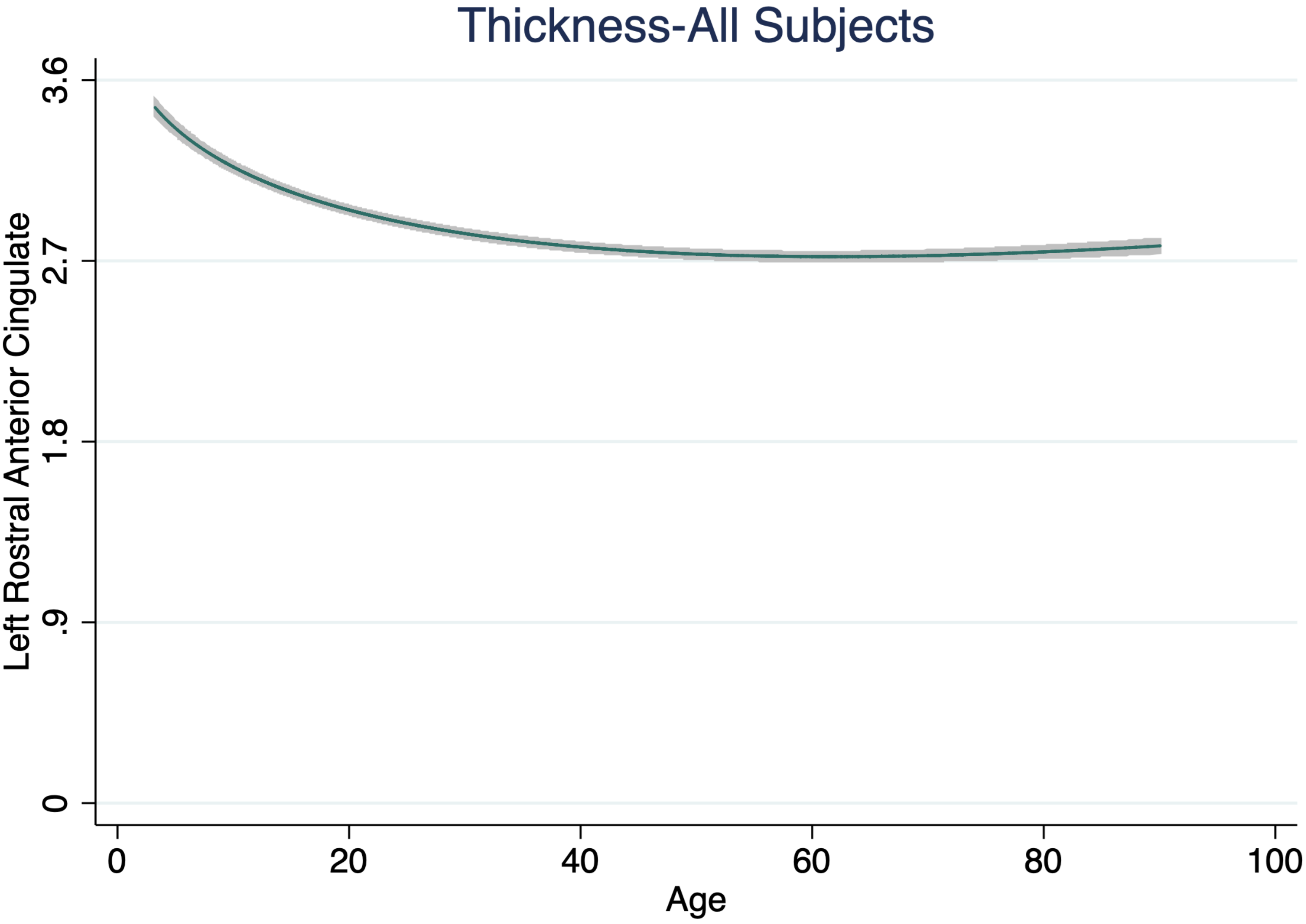

Thickness-All Subjects
Left Rostral Anterior Cingulate
3.6
2.7
1.8
.9
0
0
20
40
60
80
100
Age

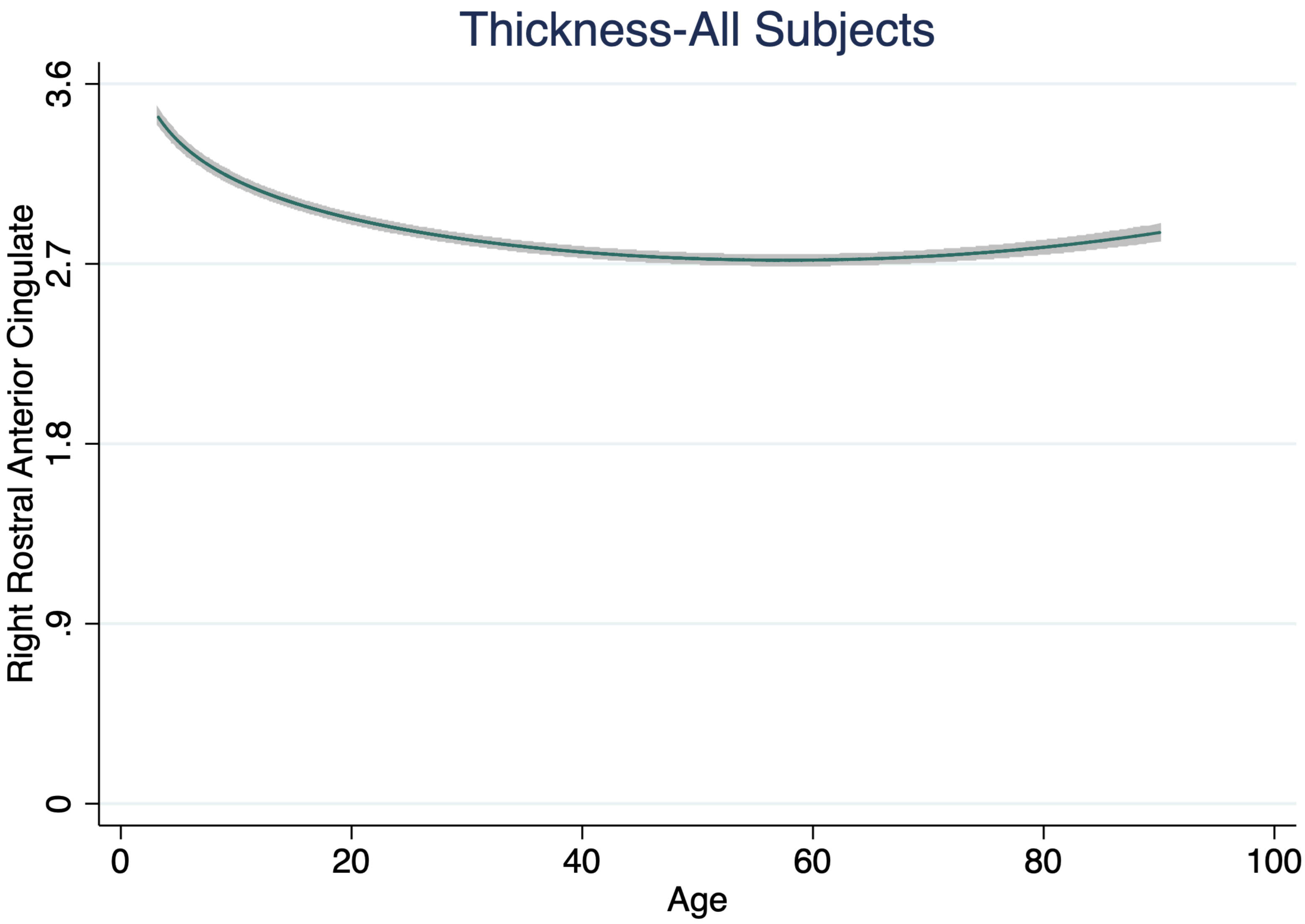
Thickness-All Subjects
Right Rostral Anterior Cingulate
3.6
2.7
1.8
.9
0
0
20
40
60
80
100
Age

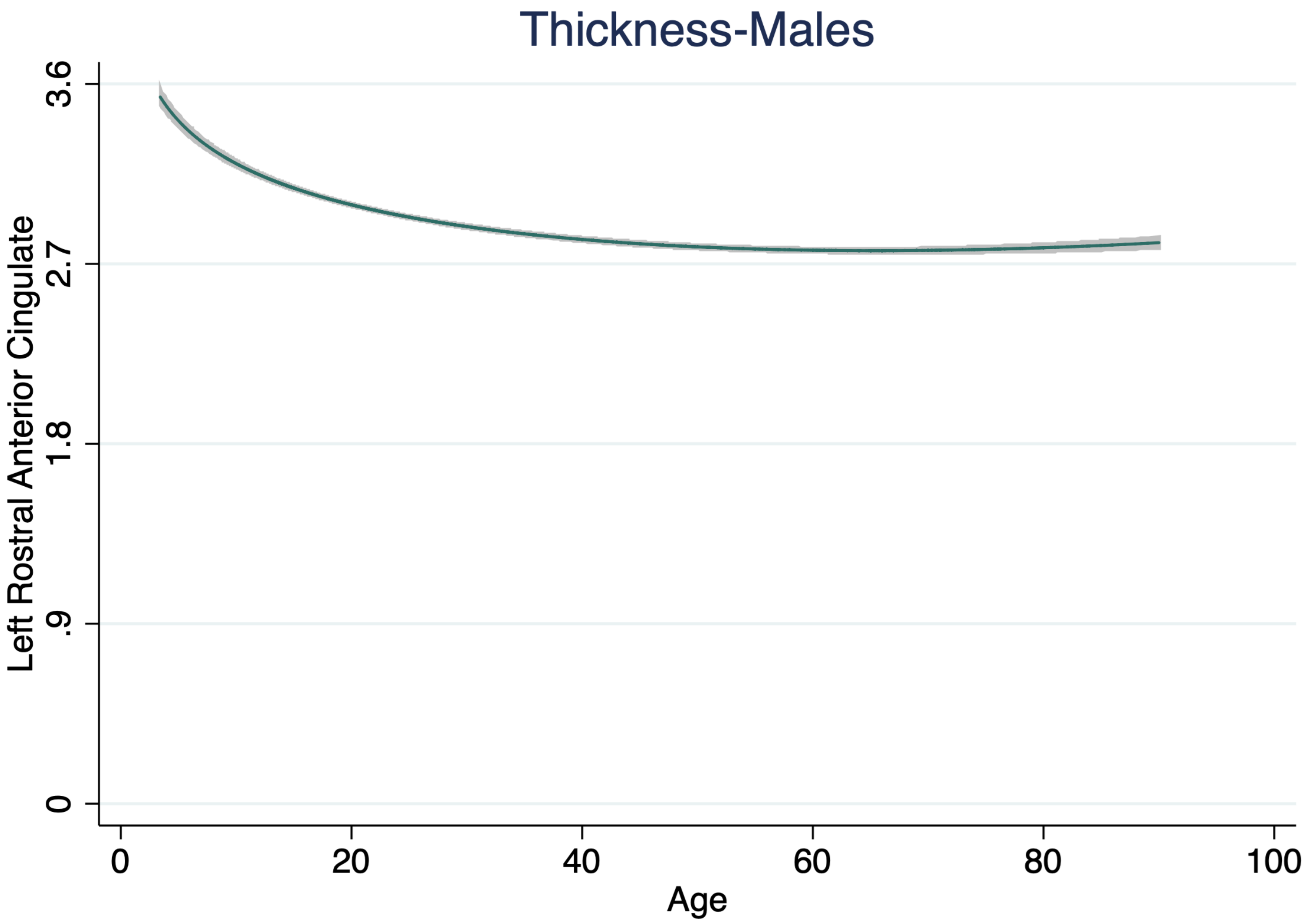
Thickness-Males
Left Rostral Anterior Cingulate
0
.9
1.8
2.7
3.6
0
20
40
60
80
100
Age

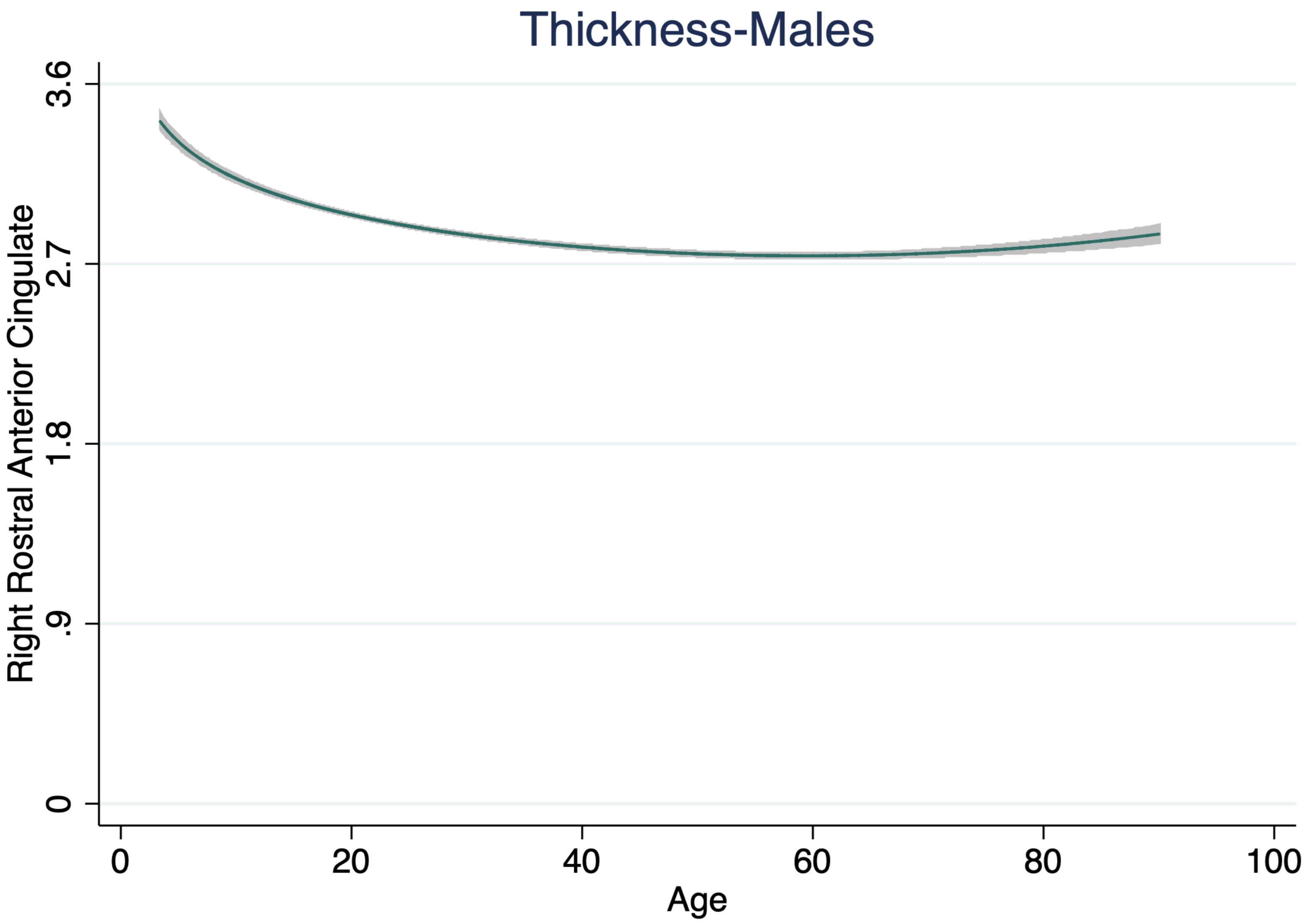

Thickness-Males
Right Rostral Anterior Cingulate
3.6
2.7
1.8
.9
0
0
20
40
60
80
100
Age

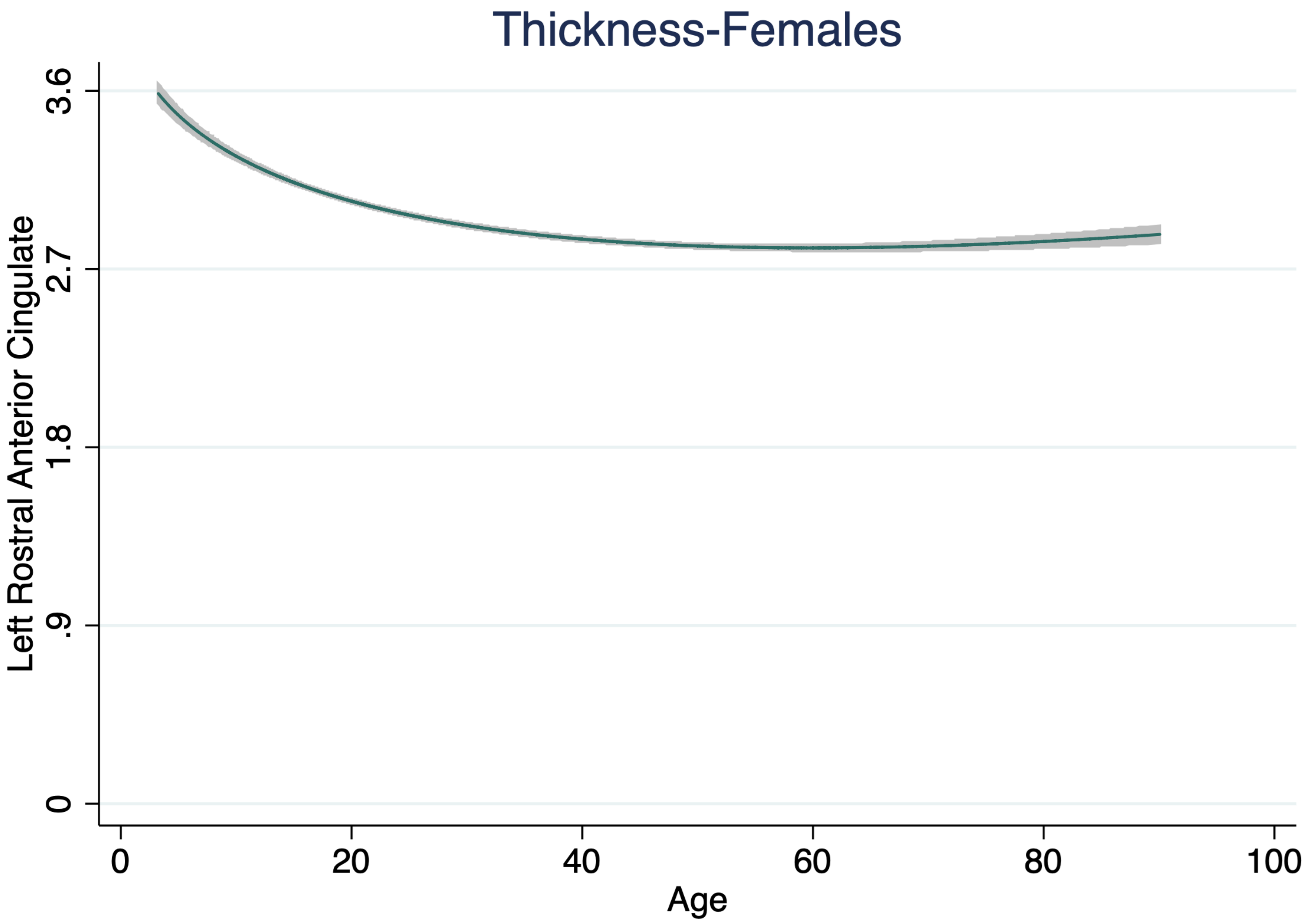

Thickness-Females
Left Rostral Anterior Cingulate
3.6
2.7
1.8
.9
0
0
20
40
60
80
100
Age

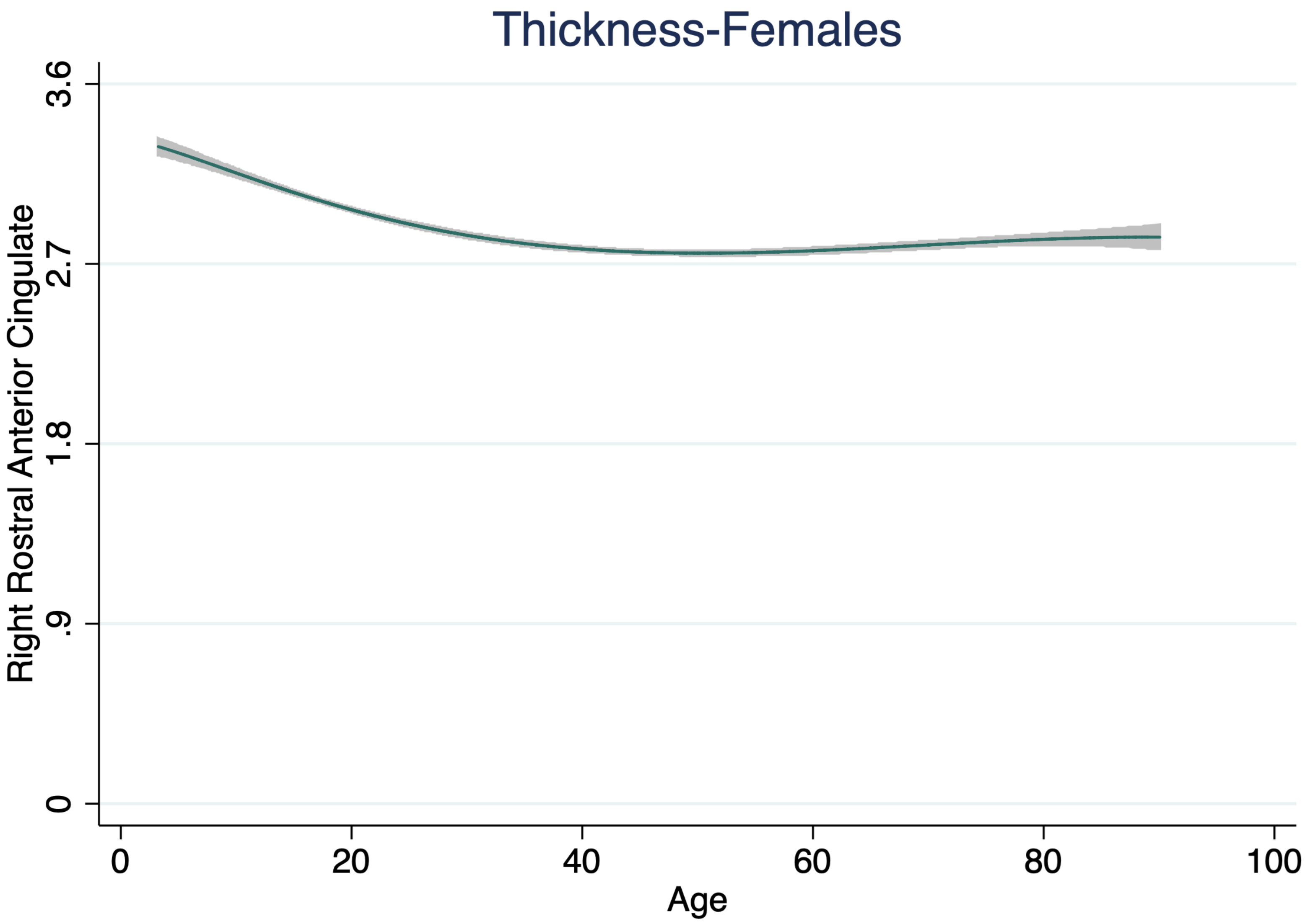

Thickness-Females
Right Rostral Anterior Cingulate
3.6
2.7
1.8
.9
0
0
20
40
60
80
100
Age

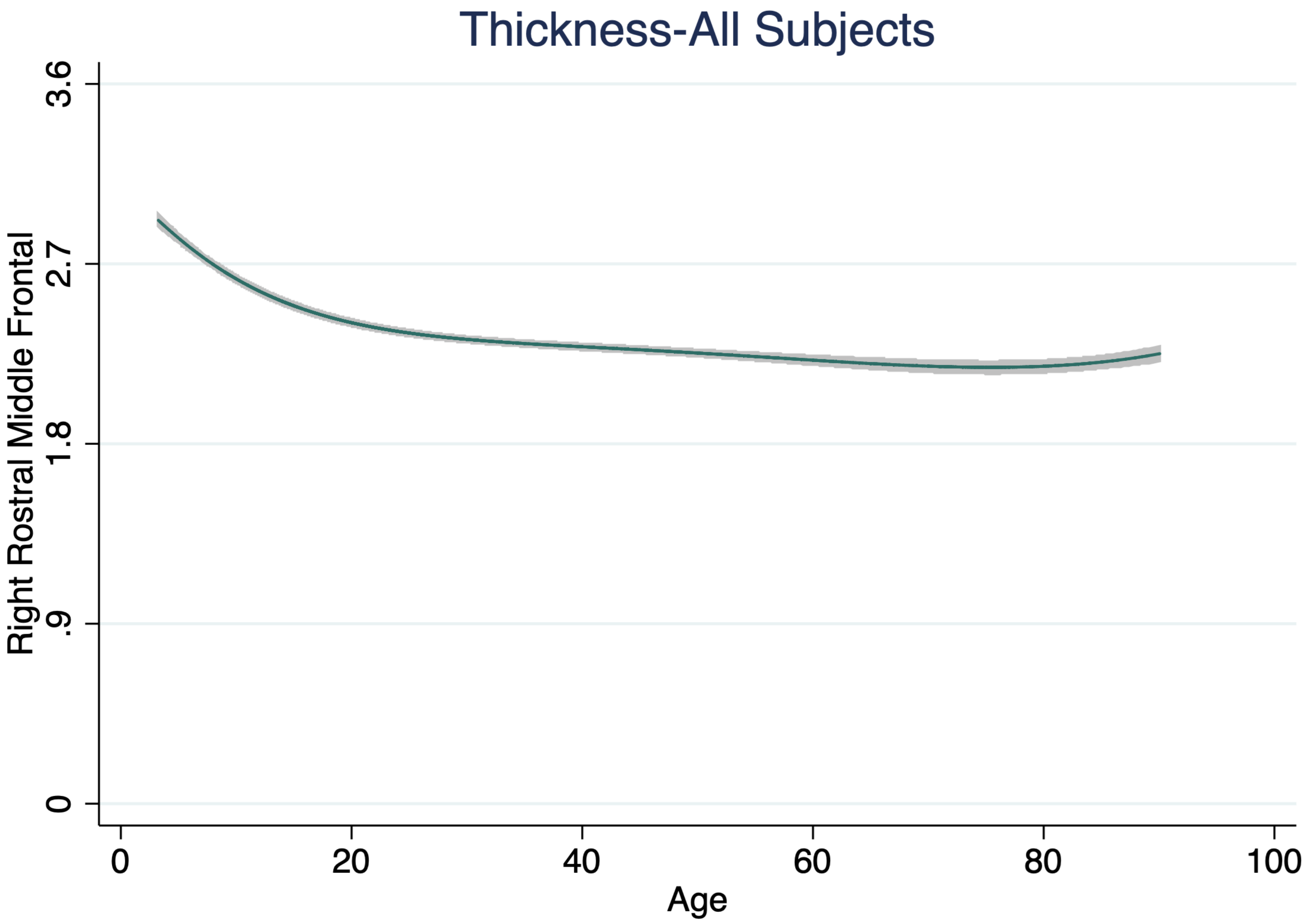
Thickness-All Subjects
Right Rostral Middle Frontal
3.6
2.7
1.8
.9
0
0
20
40
60
80
100
Age

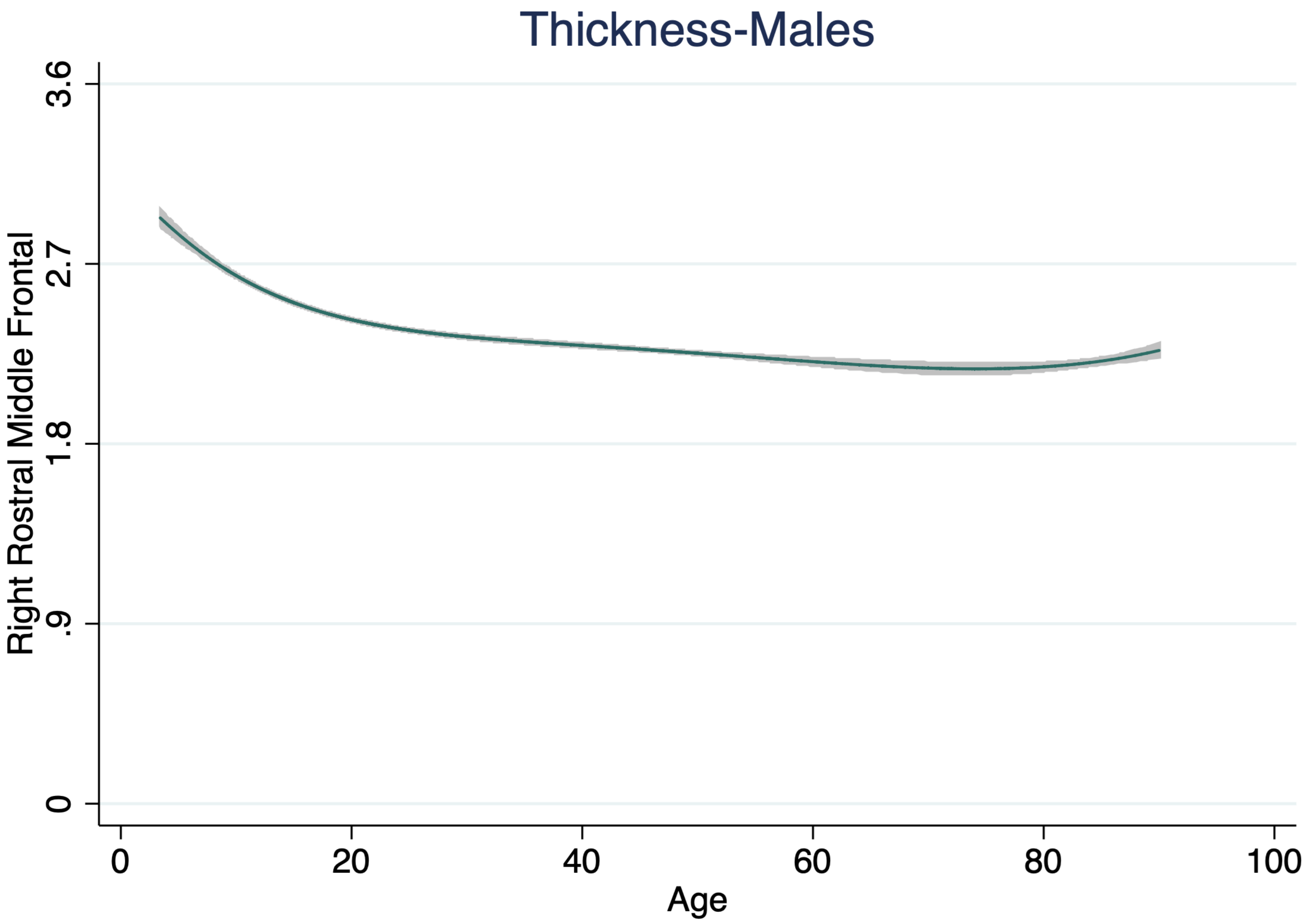

Thickness-Males
Right Rostral Middle Frontal
3.6
2.7
1.8
.9
0
0
20
40
60
80
100
Age

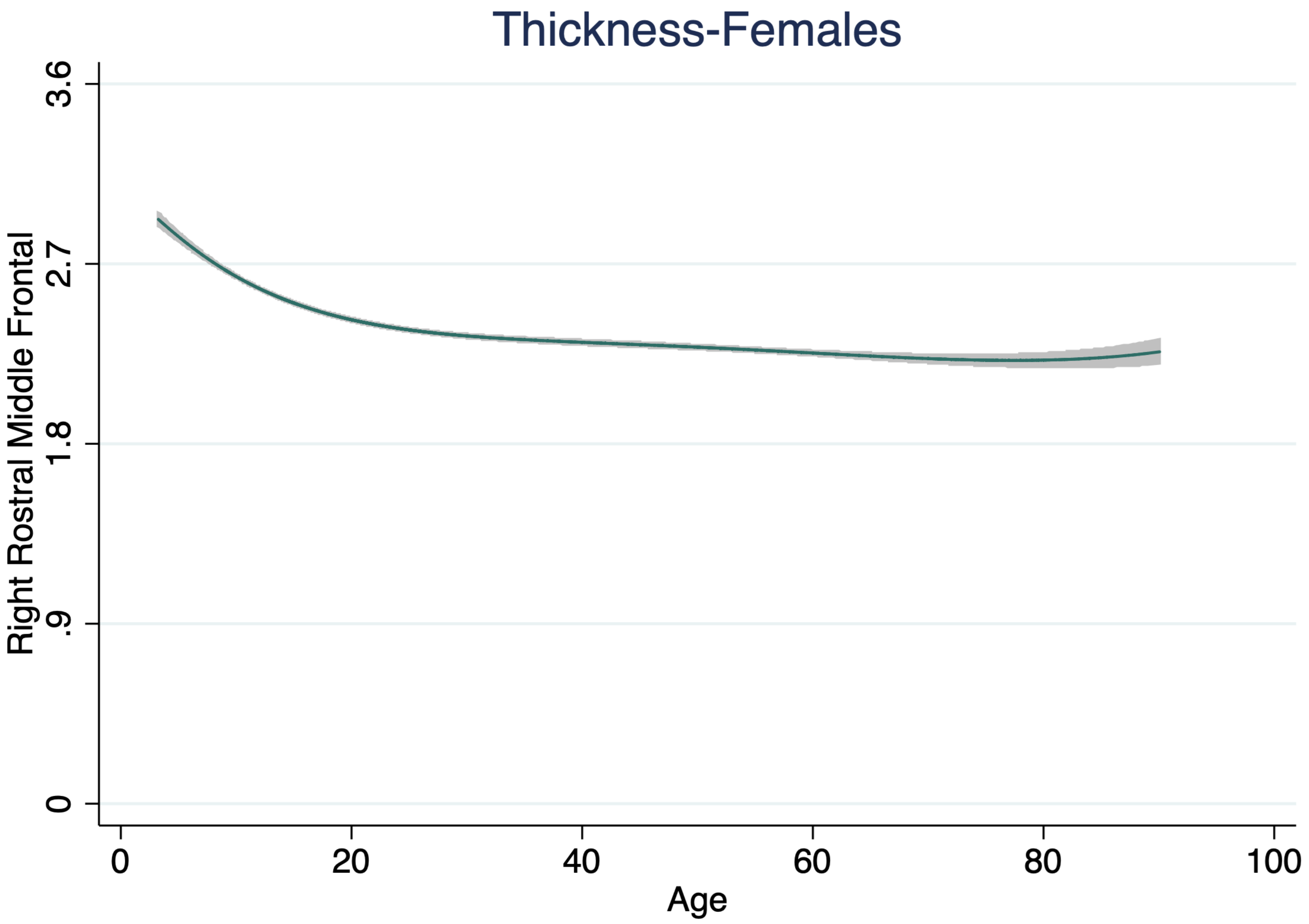

Thickness-Females
Right Rostral Middle Frontal
3.6
2.7
1.8
.9
0
0
20
40
60
80
100
Age

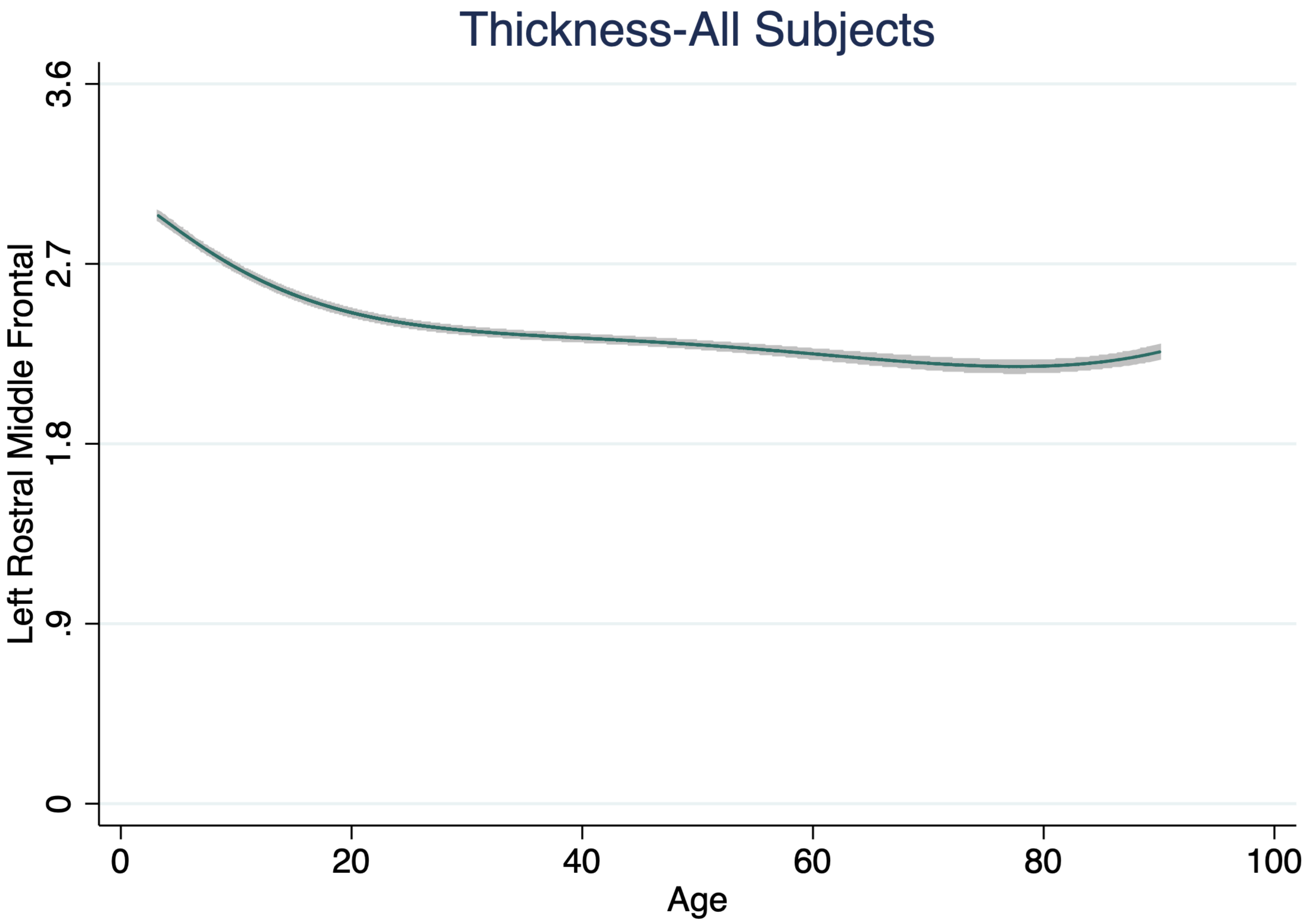
Thickness-All Subjects
Left Rostral Middle Frontal
0
.9
1.8
2.7
3.6
0
20
40
60
80
100
Age

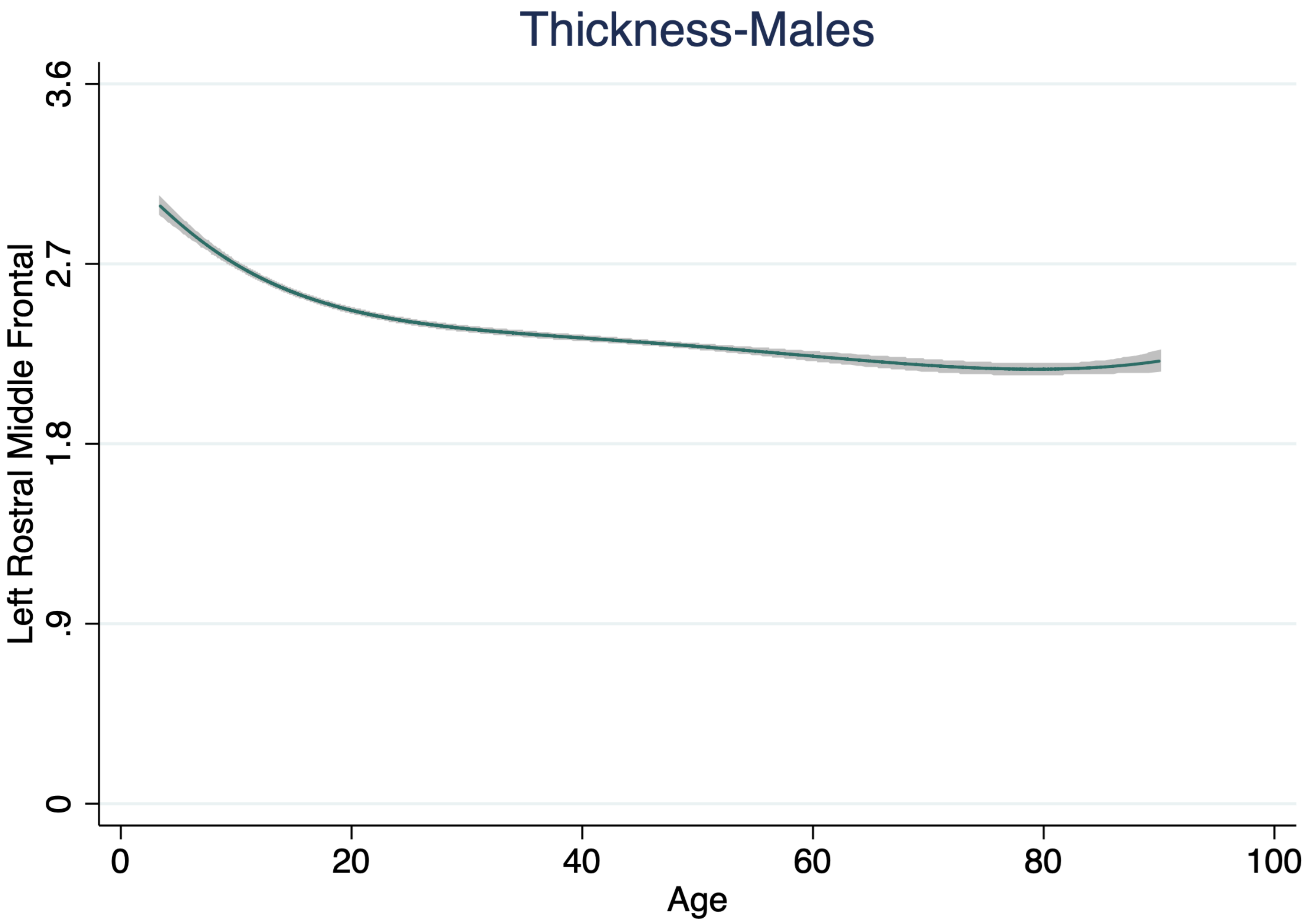
Thickness-Males
Left Rostral Middle Frontal
3.6
2.7
1.8
.9
0
0
20
40
60
80
100
Age

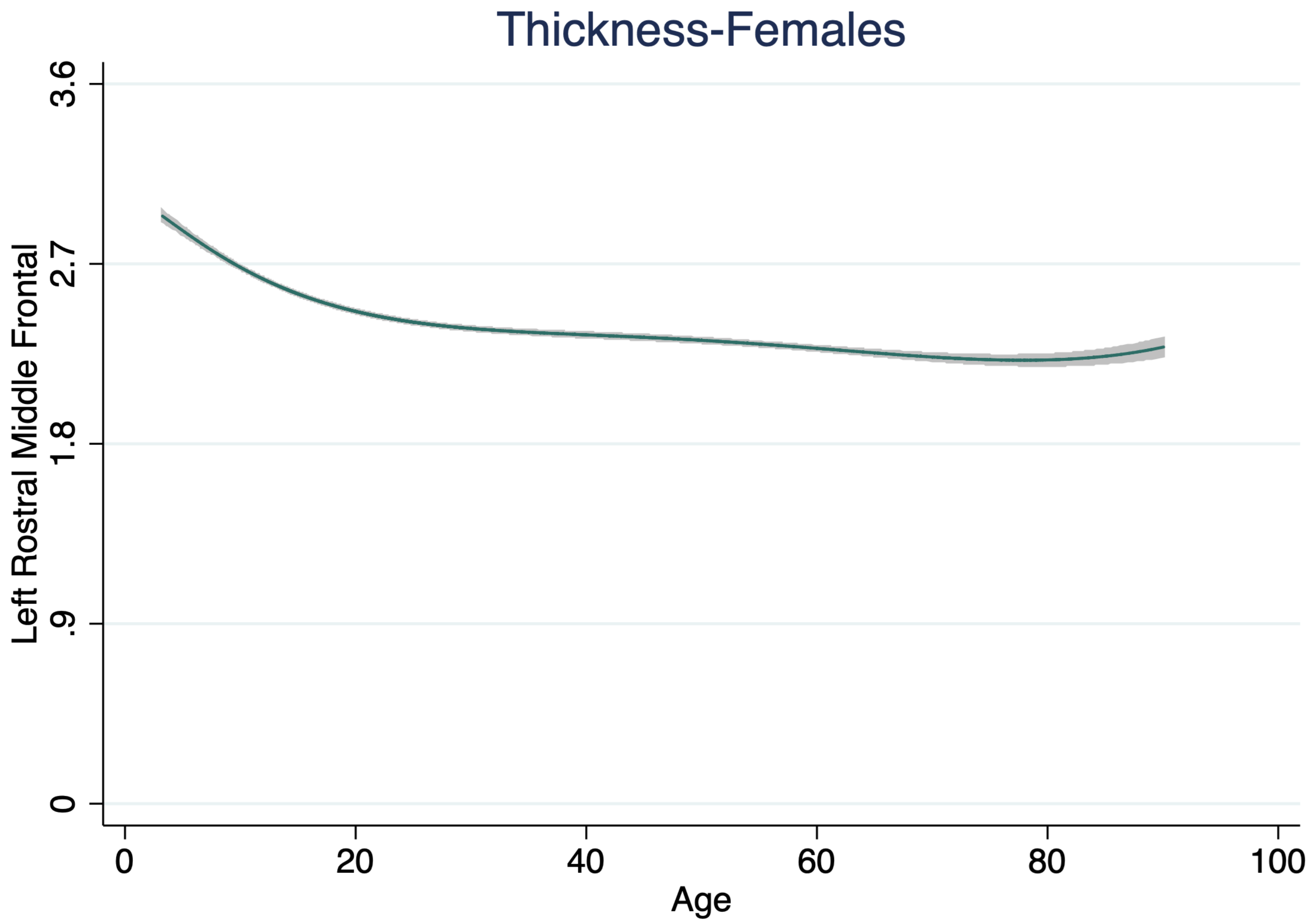

Thickness-Females
Left Rostral Middle Frontal
0
.9
1.8
2.7
3.6
0
20
40
60
80
100
Age

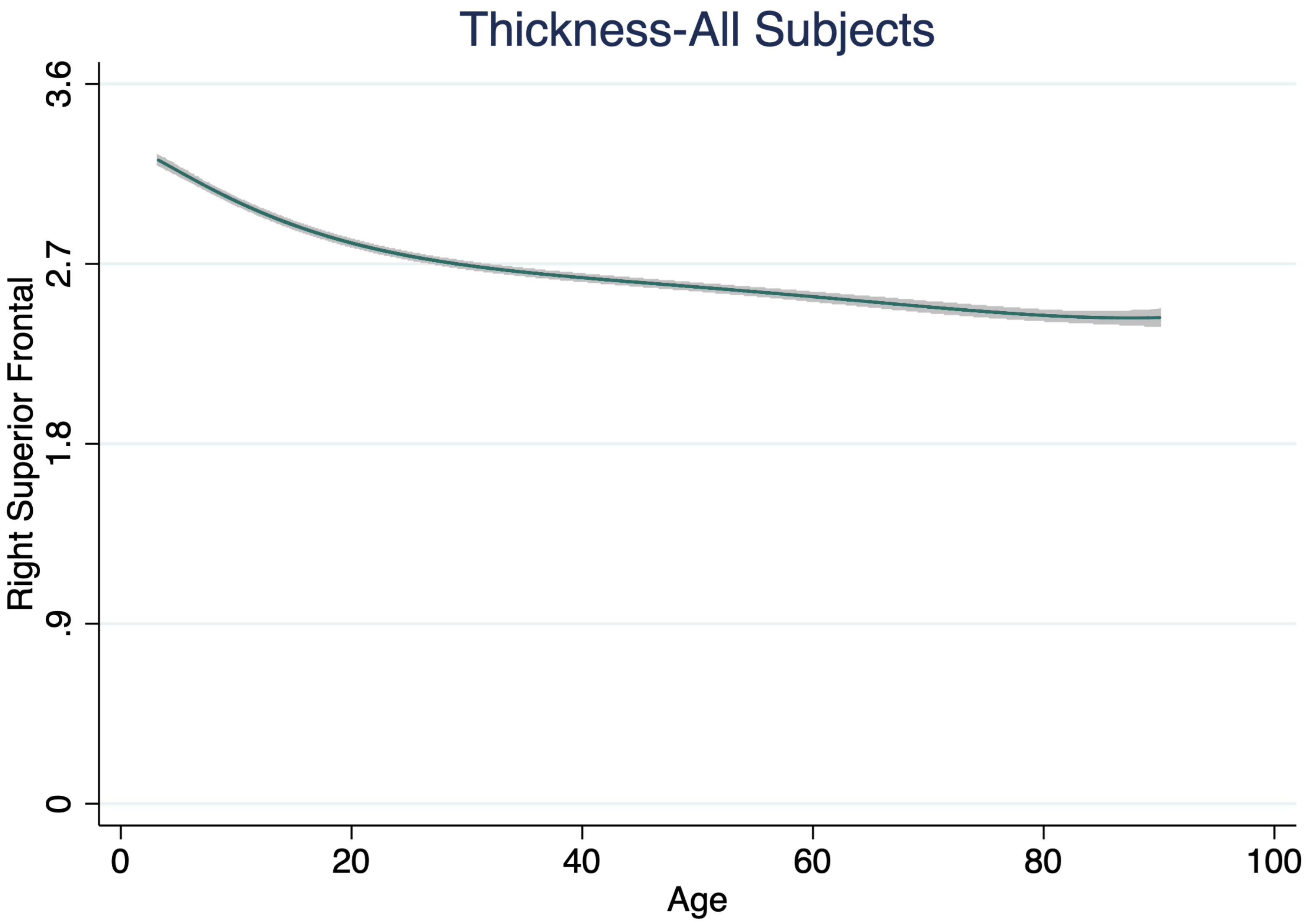

Thickness-All Subjects
Right Superior Frontal
3.6
2.7
1.8
.9
0
0
20
40
60
80
100
Age

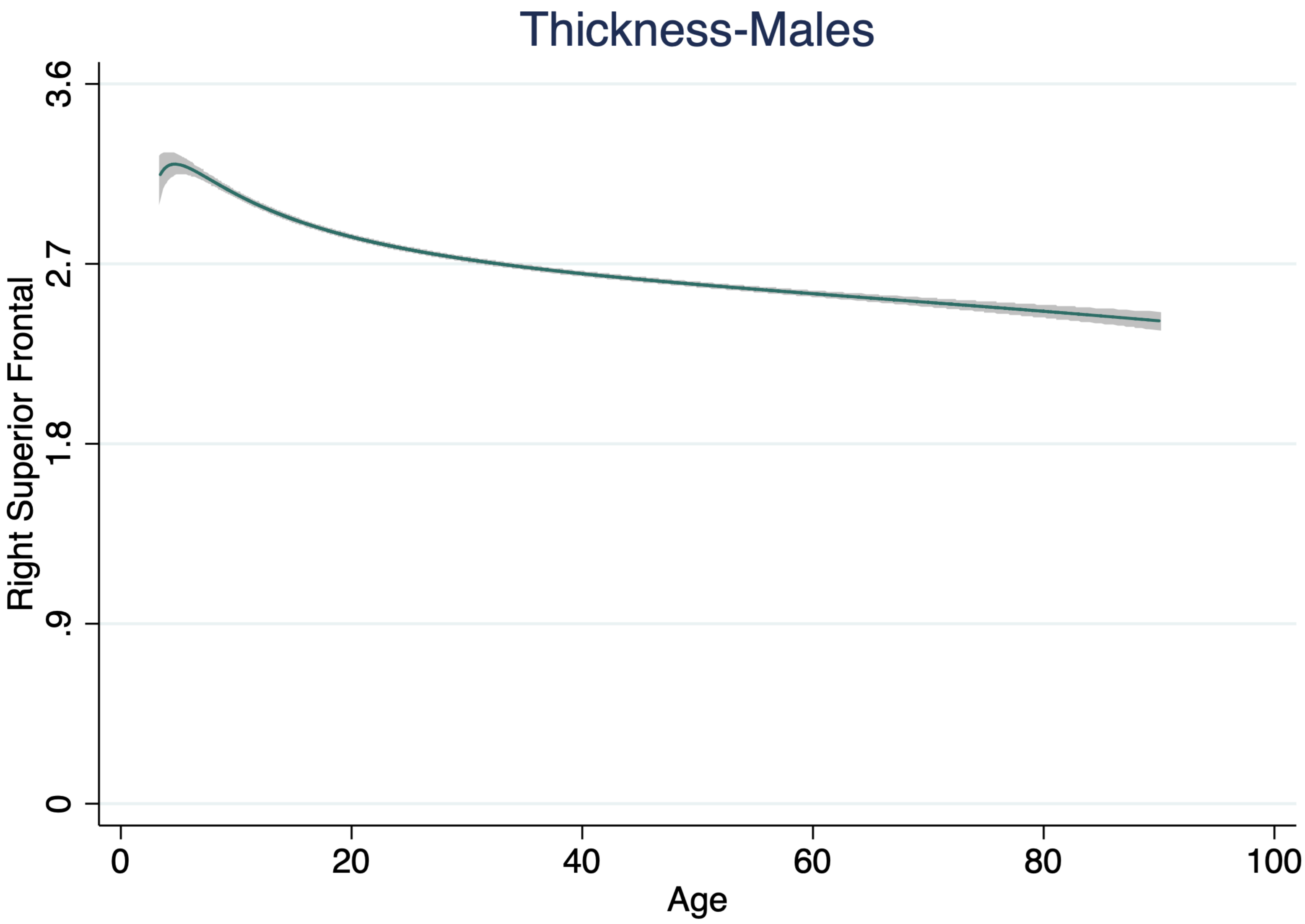
Thickness-Males
Right Superior Frontal
3.6
2.7
1.8
.9
0
0
20
40
60
80
100
Age

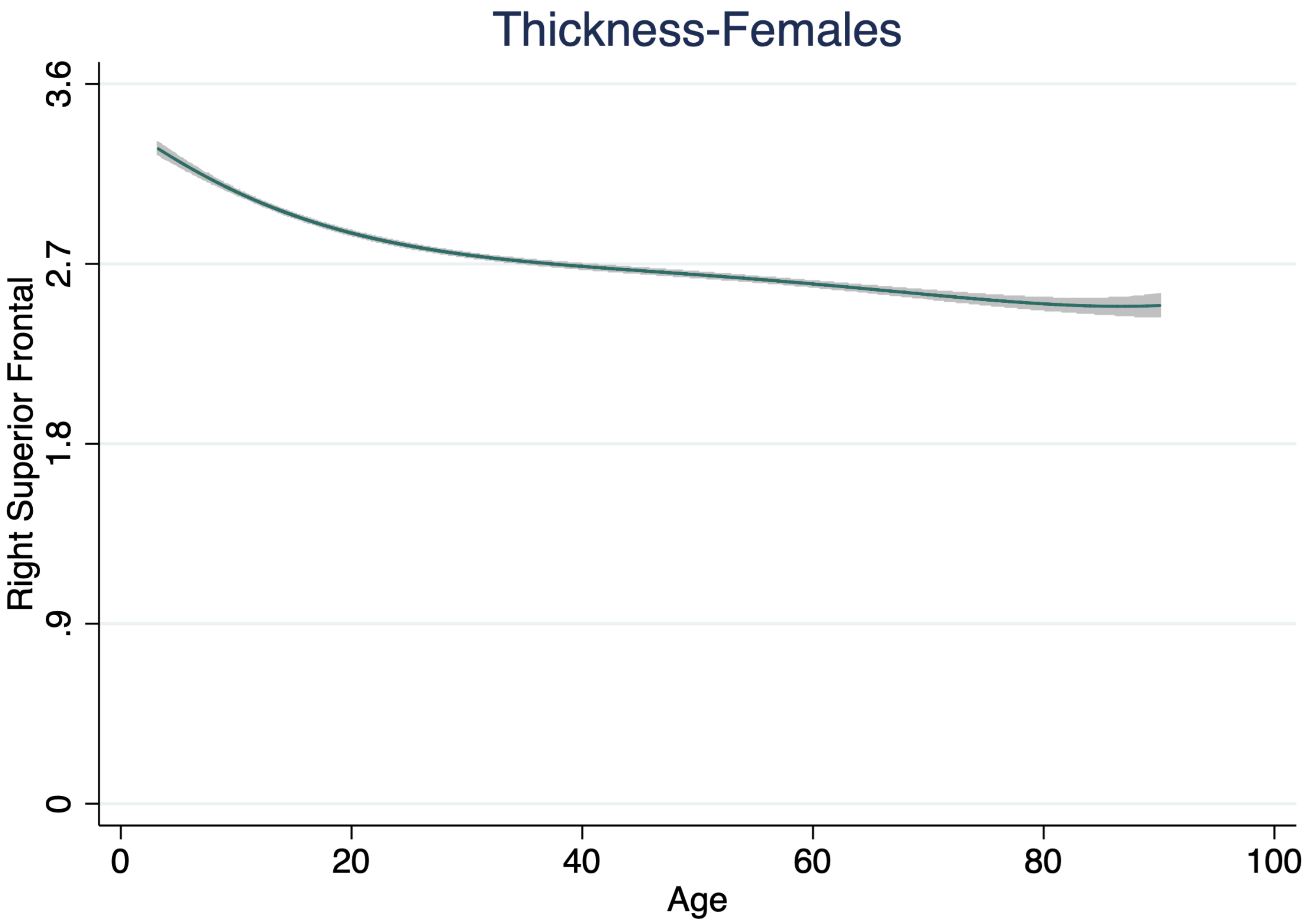
Thickness-Females
Right Superior Frontal
3.6
2.7
1.8
.9
0
0
20
40
60
80
100
Age

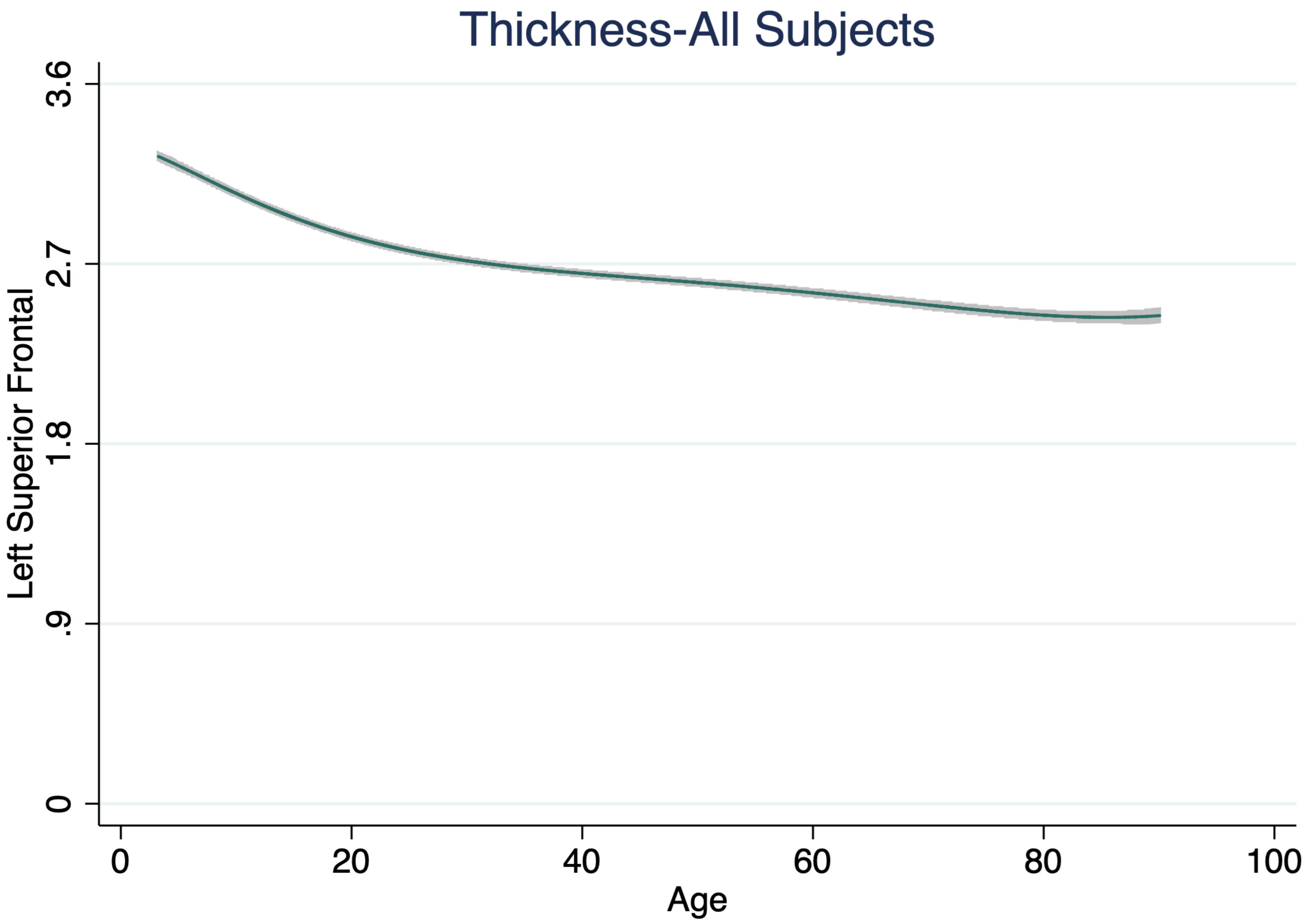

Thickness-All Subjects
Left Superior Frontal
3.6
2.7
1.8
.9
0
0
20
40
60
80
100
Age

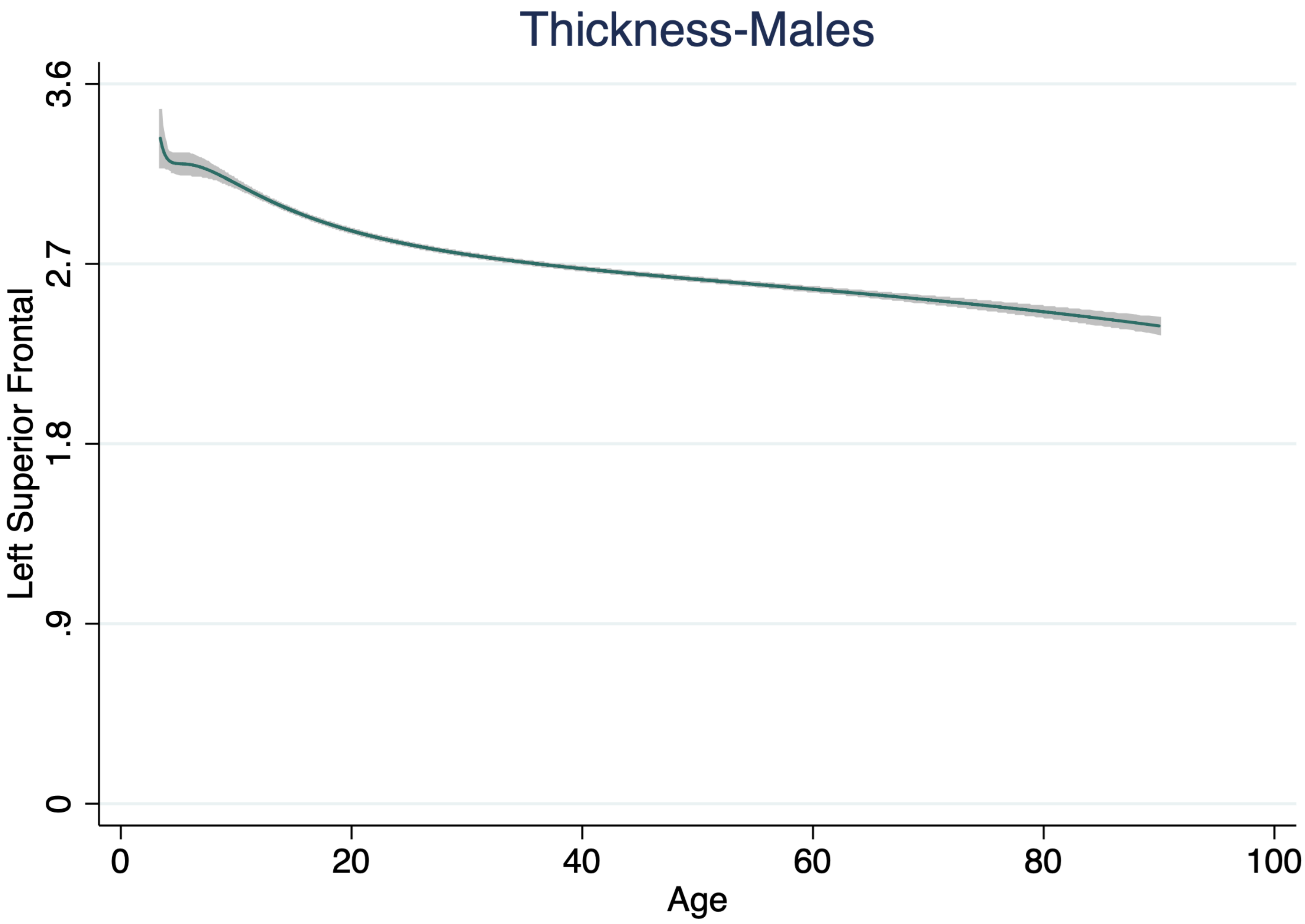

Thickness-Males
Left Superior Frontal
3.6
2.7
1.8
.9
0
0
20
40
60
80
100
Age

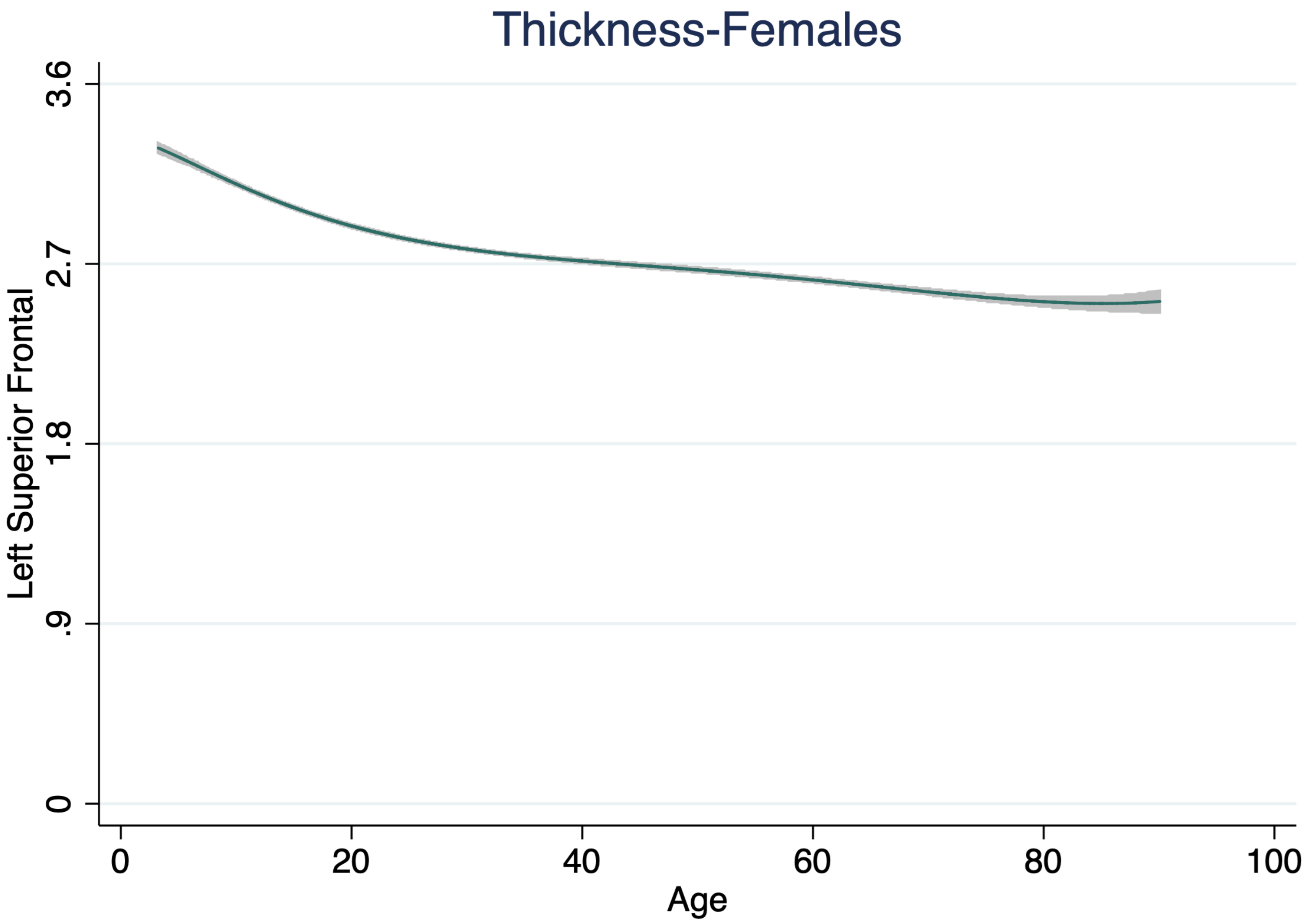
Thickness-Females
Left Superior Frontal
3.6
2.7
1.8
.9
0
0
20
40
60
80
100
Age

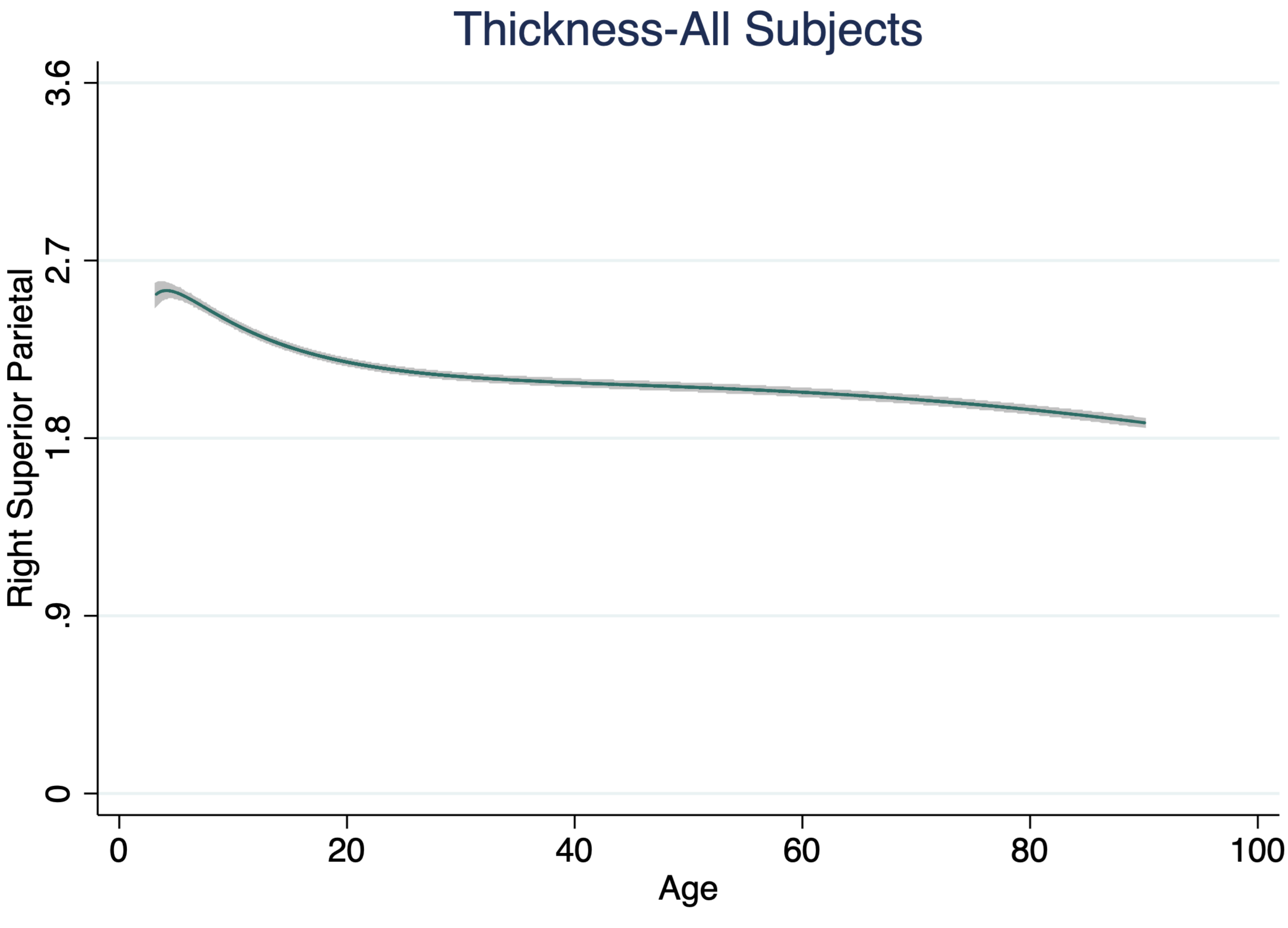

Thickness-All Subjects
Right Superior Parietal
0
.9
1.8
2.7
3.6
0
20
40
60
80
100
Age

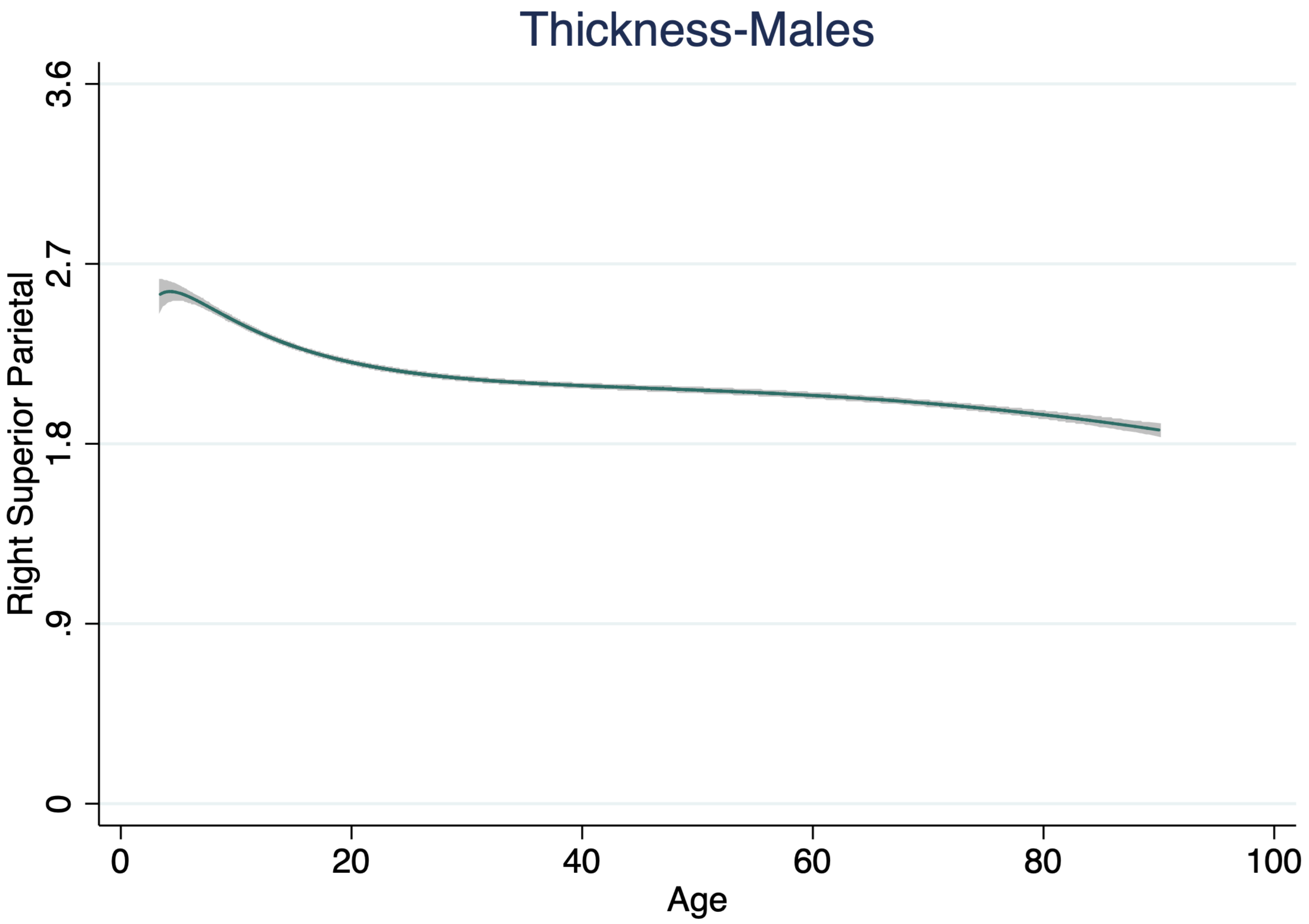
Thickness-Males
Right Superior Parietal
3.6
2.7
1.8
.9
0
0
20
40
60
80
100
Age

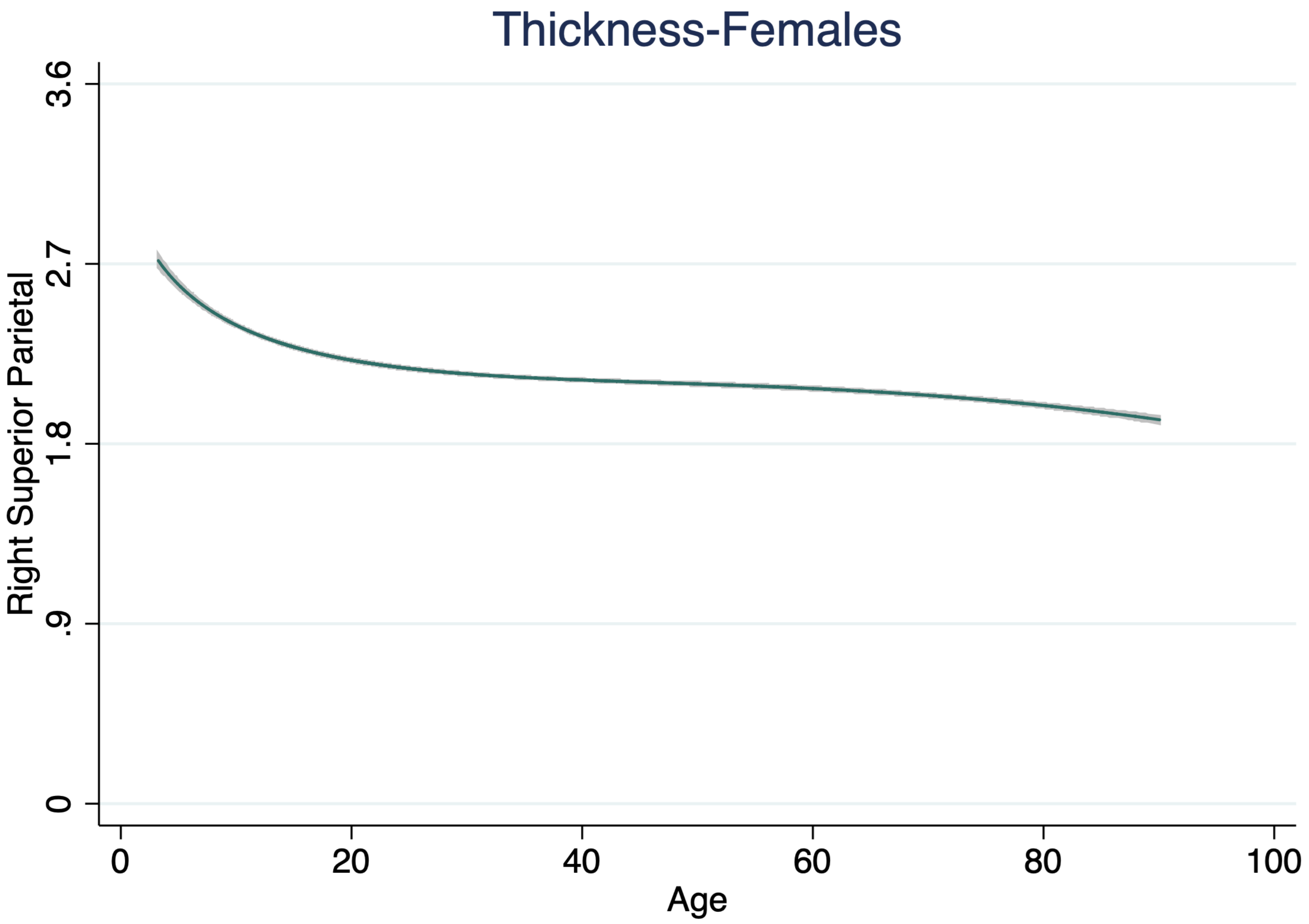
Thickness-Females
Right Superior Parietal
3.6
2.7
1.8
.9
0
0
20
40
60
80
100
Age

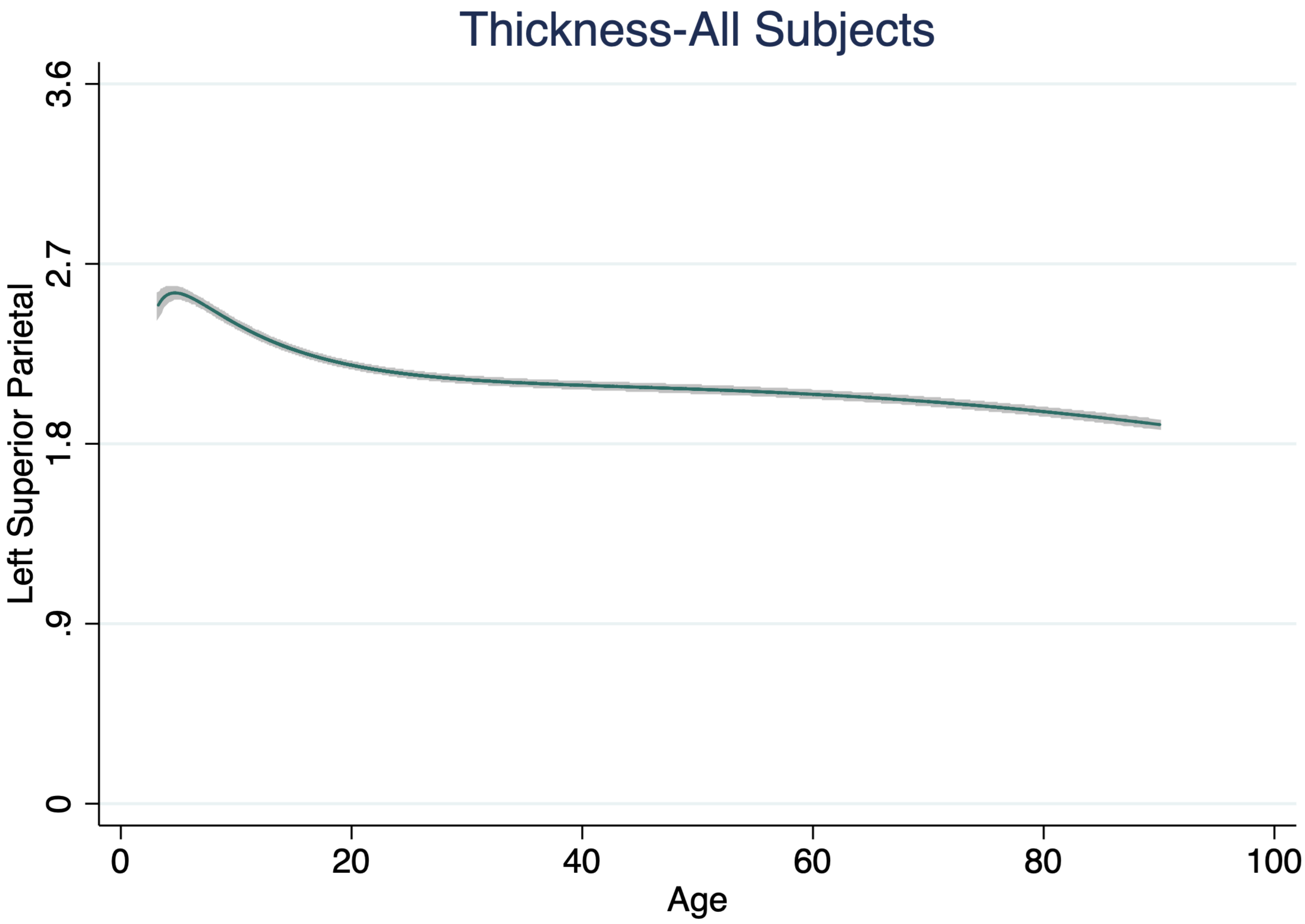

Thickness-All Subjects
Left Superior Parietal
3.6
2.7
1.8
.9
0
0
20
40
60
80
100
Age

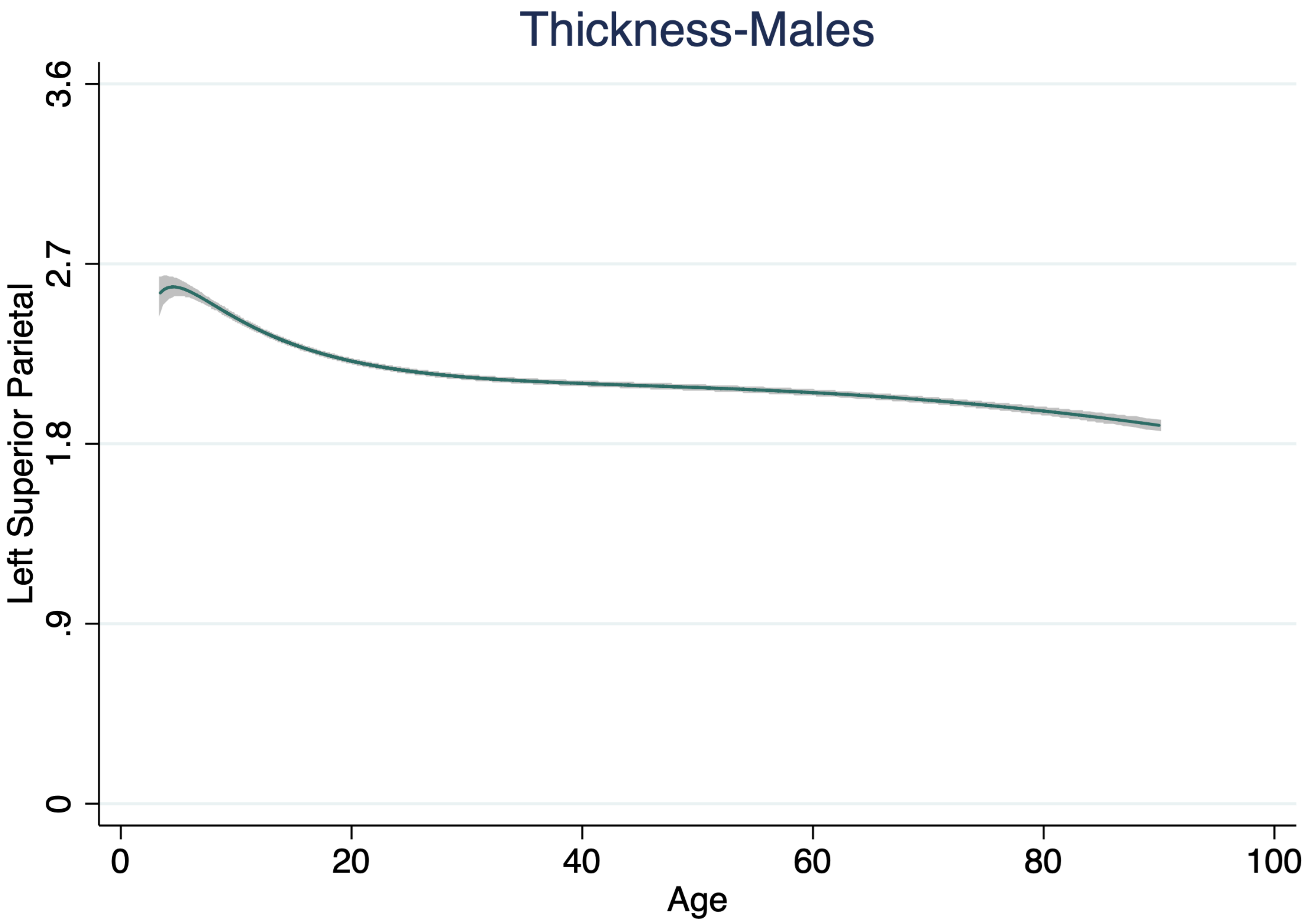

Thickness-Males
Left Superior Parietal
3.6
2.7
1.8
.9
0
0
20
40
60
80
100
Age

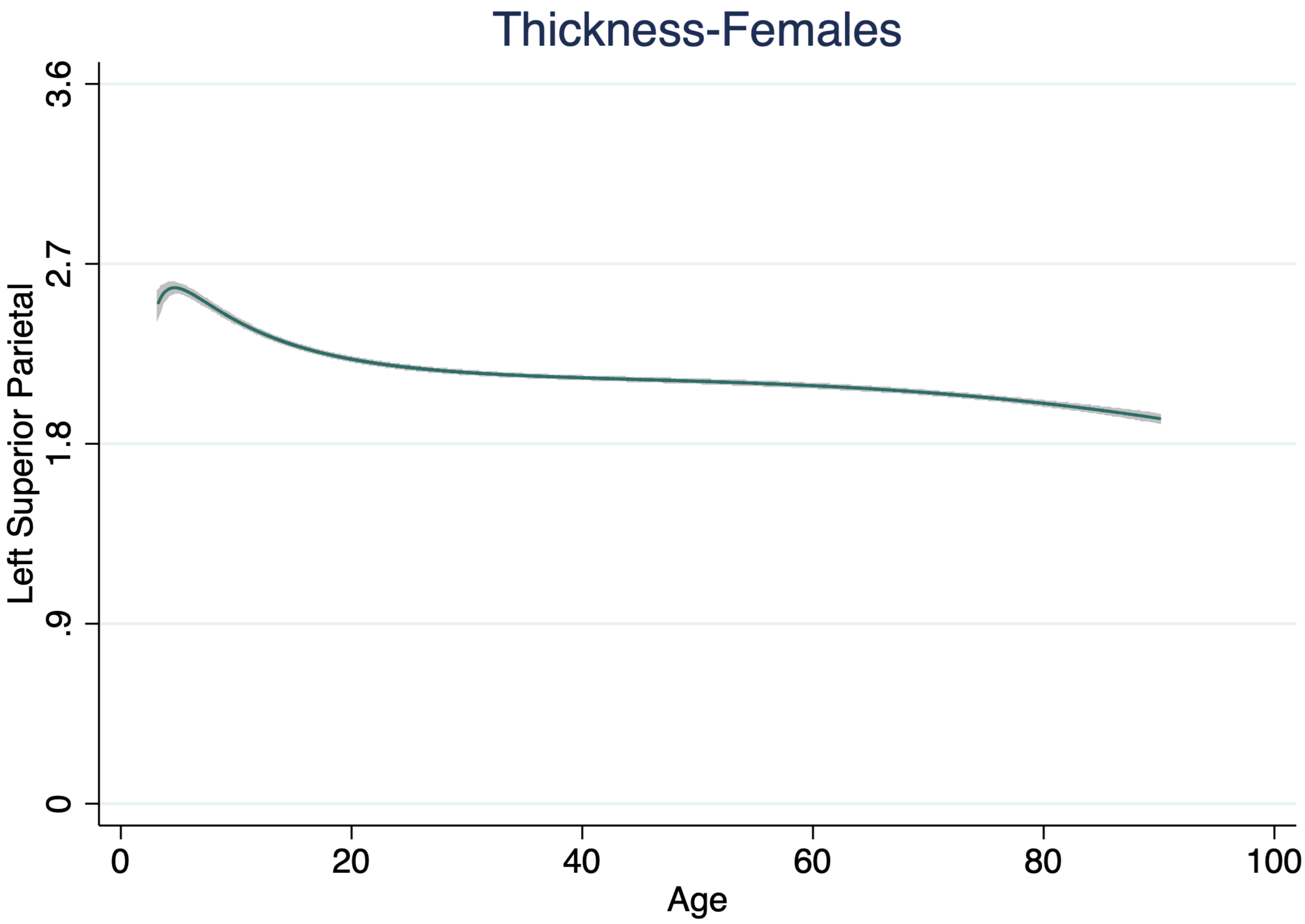
Thickness-Females
Left Superior Parietal
3.6
2.7
1.8
.9
0
0
20
40
60
80
100
Age

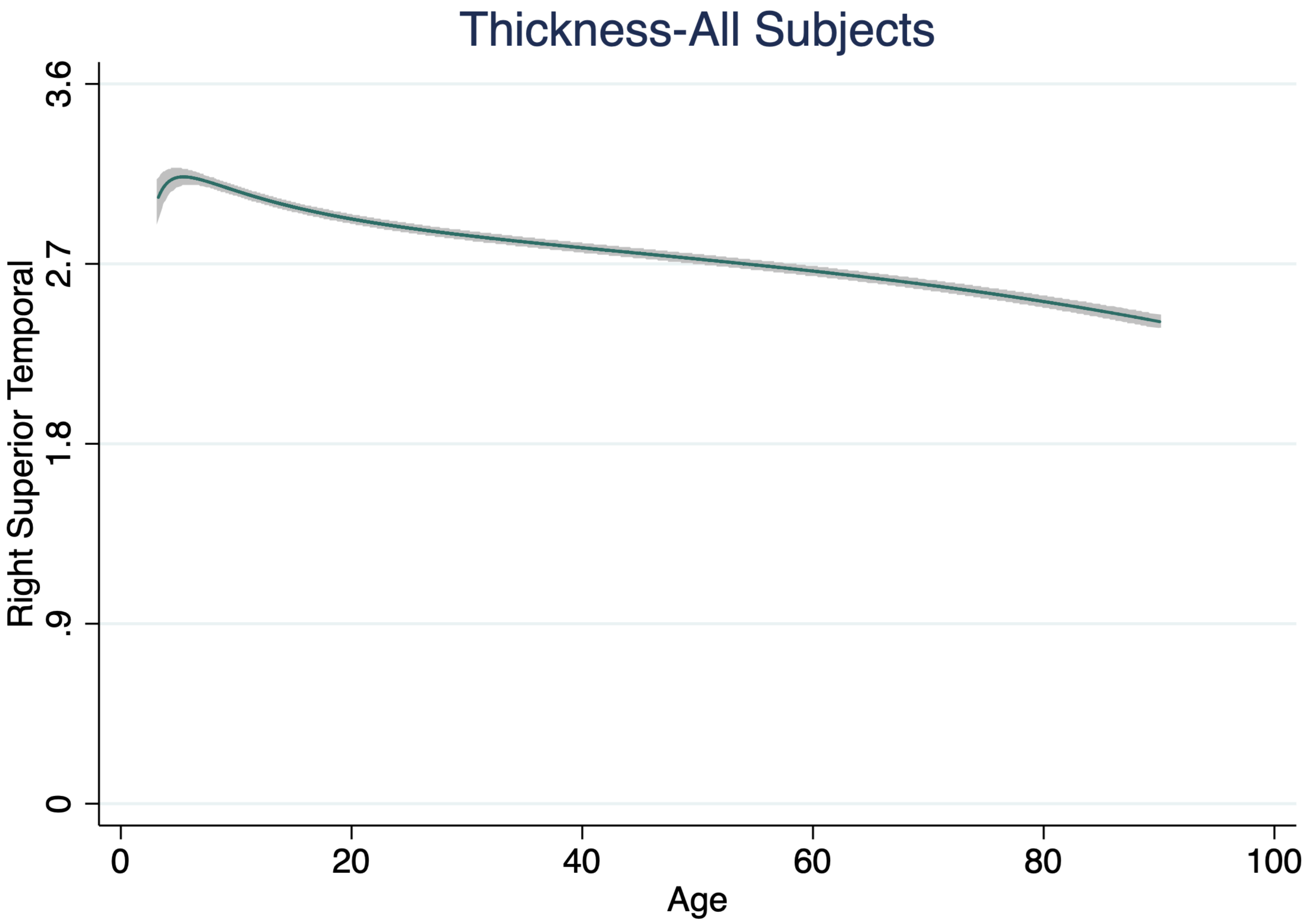
Thickness-All Subjects
Right Superior Temporal
0
.9
1.8
2.7
3.6
0
20
40
60
80
100
Age

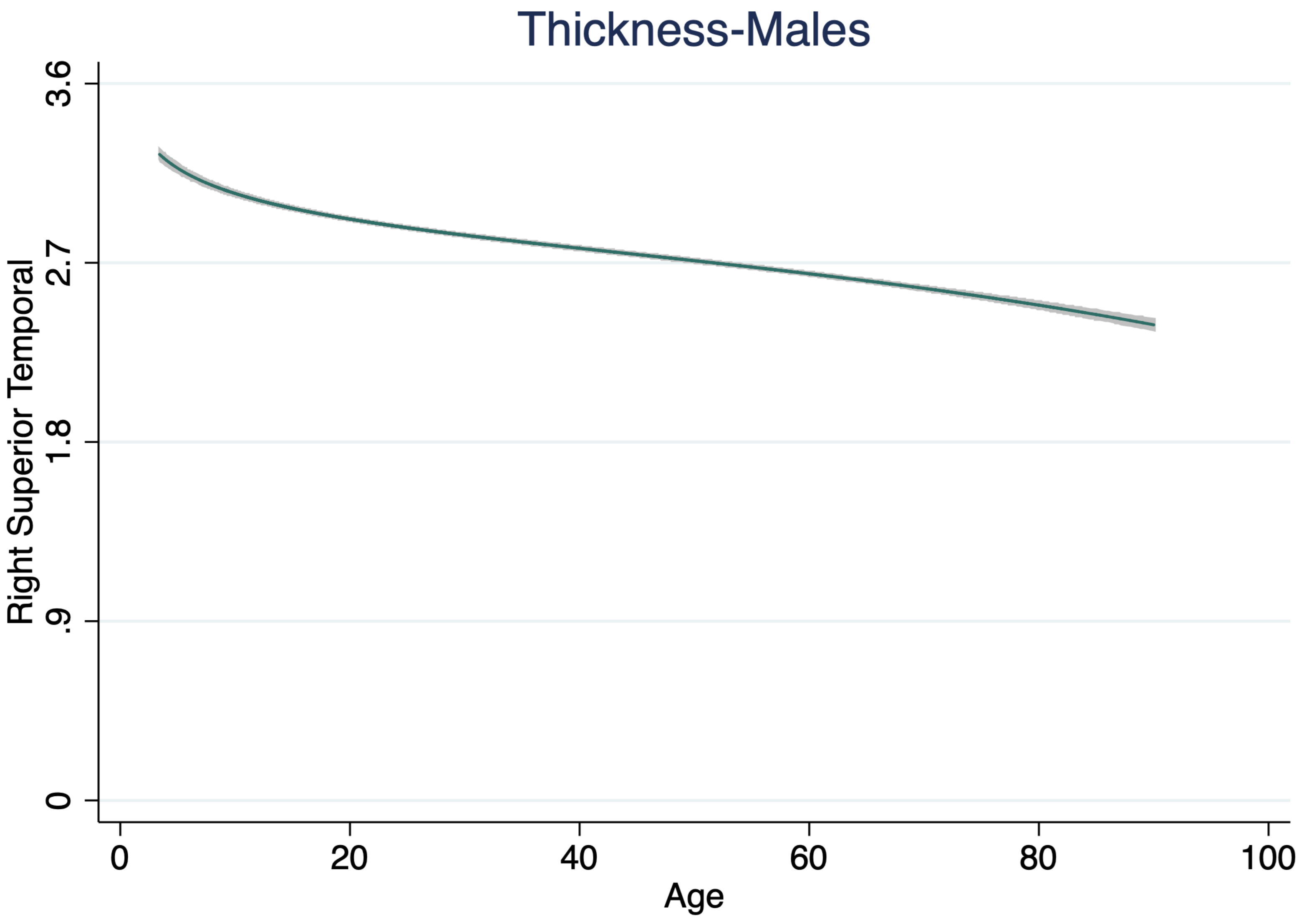
Thickness-Males
Right Superior Temporal
0
.9
1.8
2.7
3.6
0
20
40
60
80
100
Age

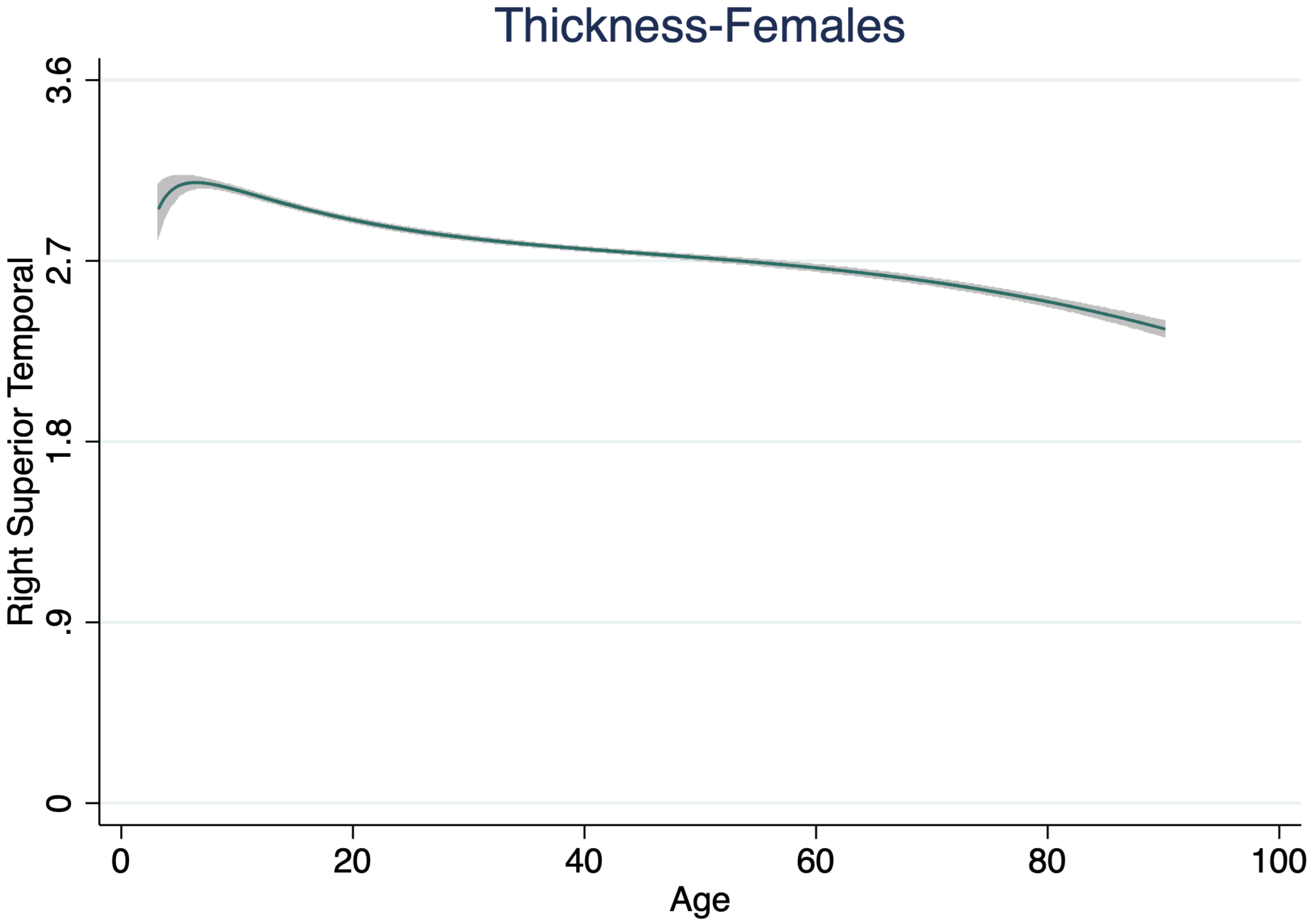

Thickness-Females
Right Superior Temporal
3.6
2.7
1.8
.9
0
0
20
40
60
80
100
Age

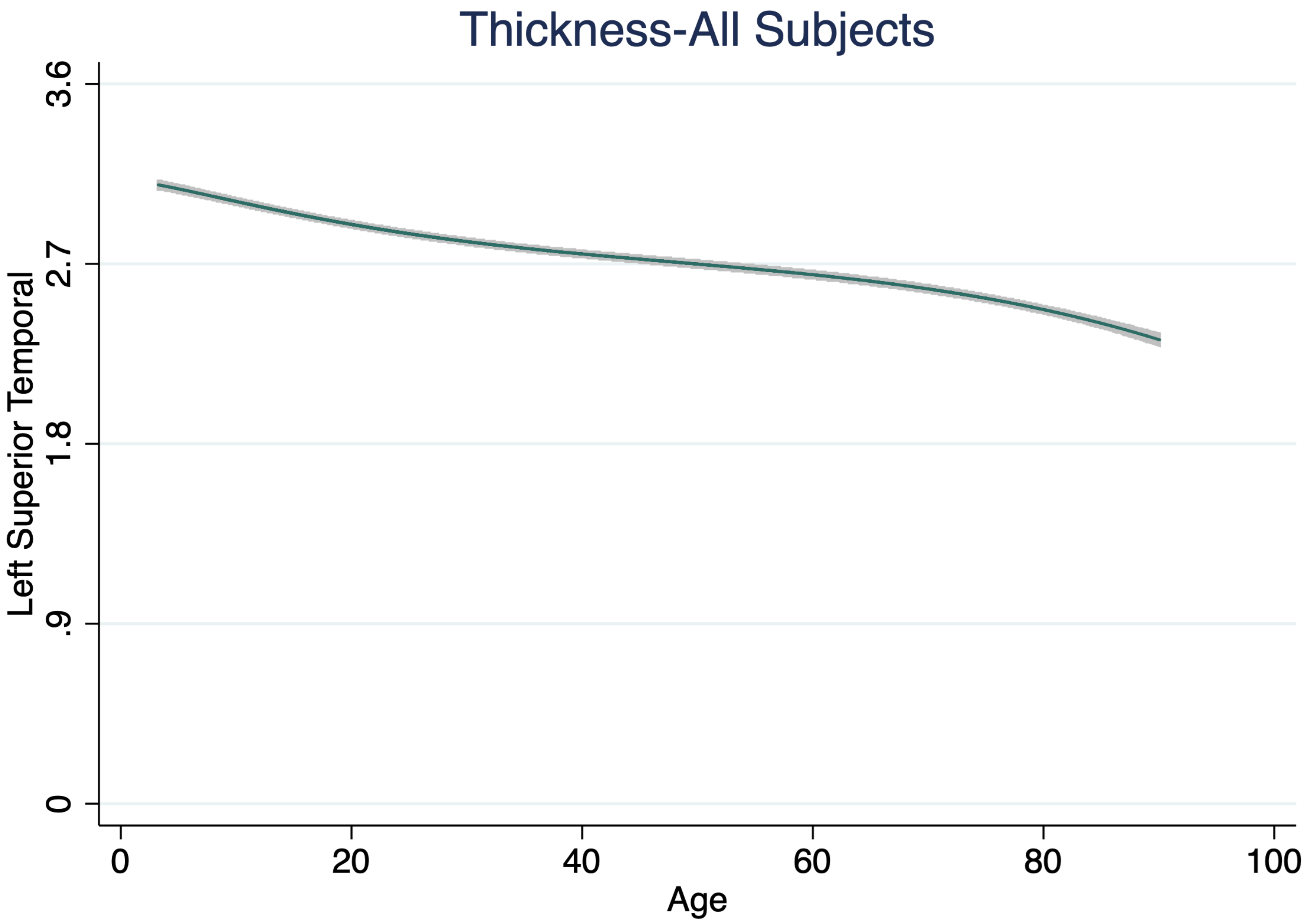

Thickness-All Subjects
Left Superior Temporal
3.6
2.7
1.8
.9
0
0
20
40
60
80
100
Age

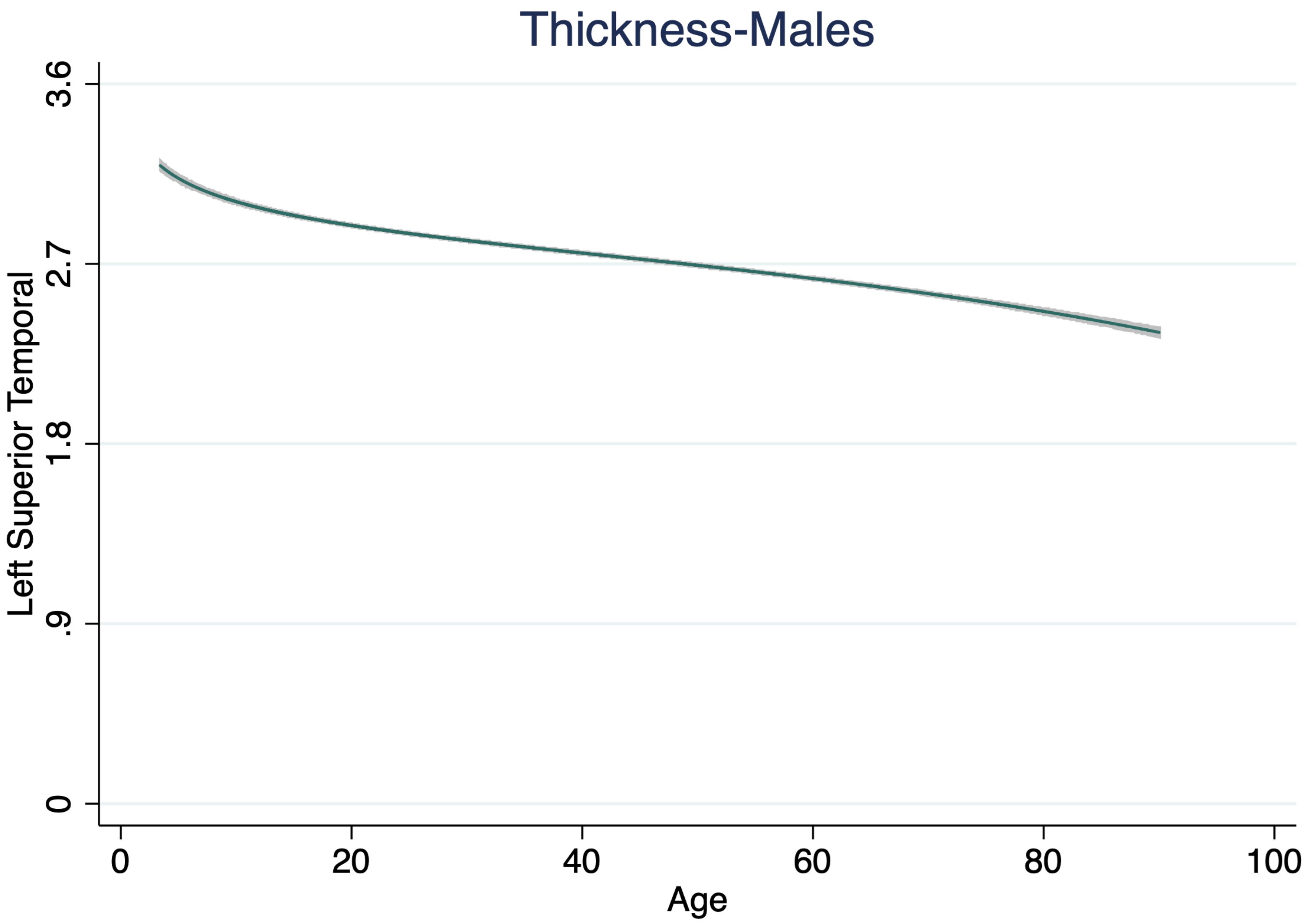

Thickness-Males
Left Superior Temporal
0
.9
1.8
2.7
3.6
0
20
40
60
80
100
Age

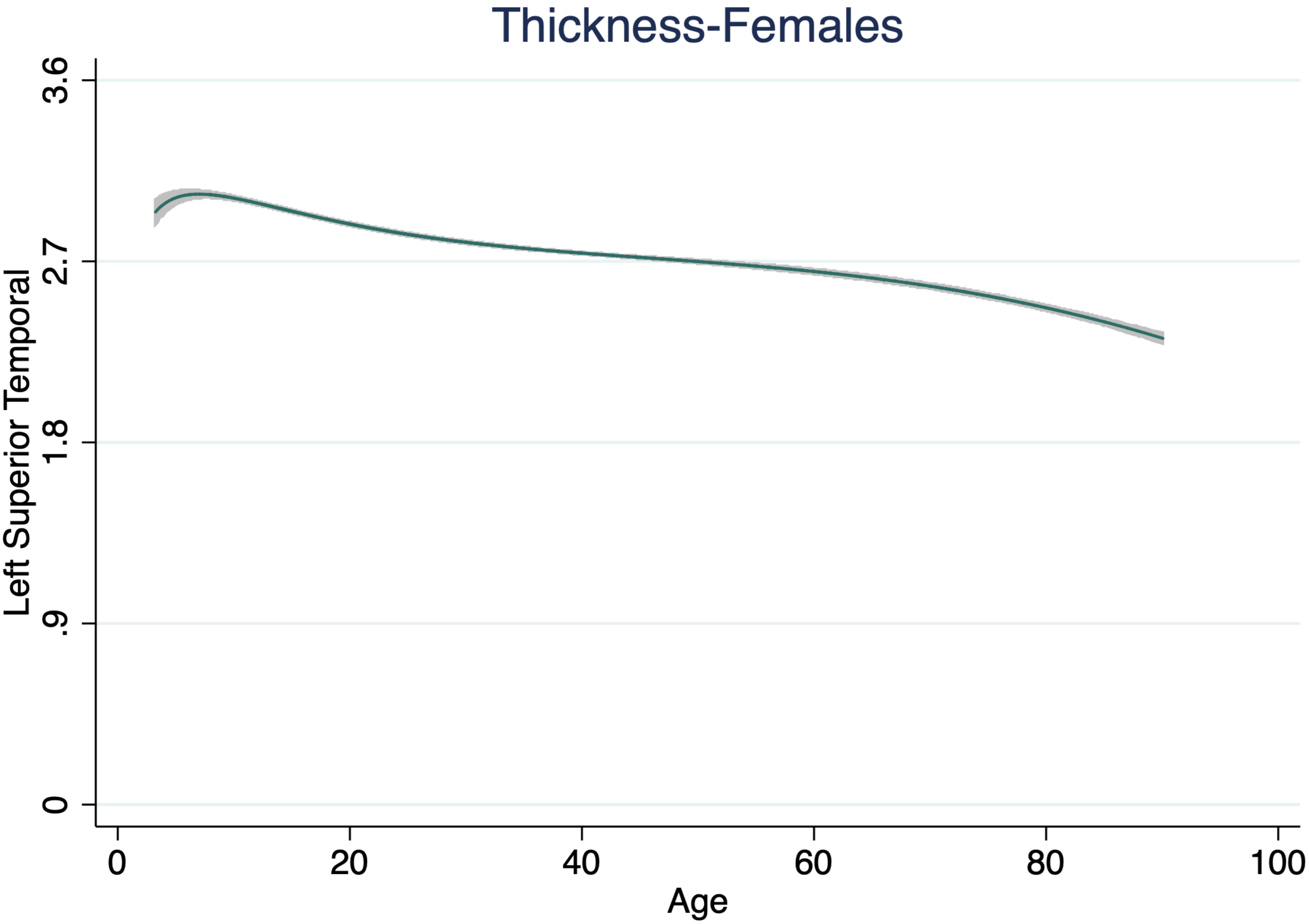
Thickness-Females
Left Superior Temporal
3.6
2.7
1.8
.9
0
0
20
40
60
80
100
Age

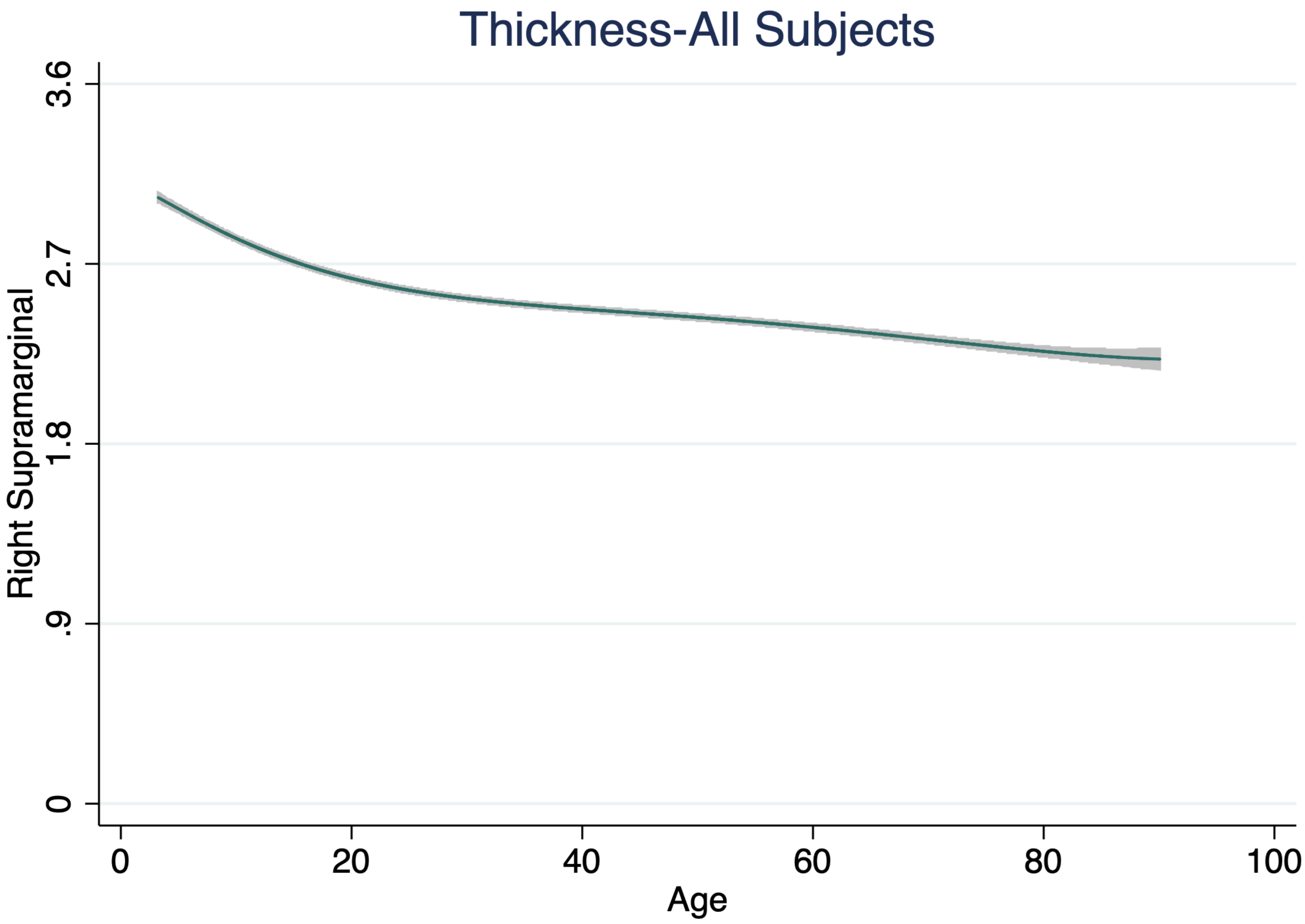
Thickness-All Subjects
Right Supramarginal
3.6
2.7
1.8
.9
0
0
20
40
60
80
100
Age

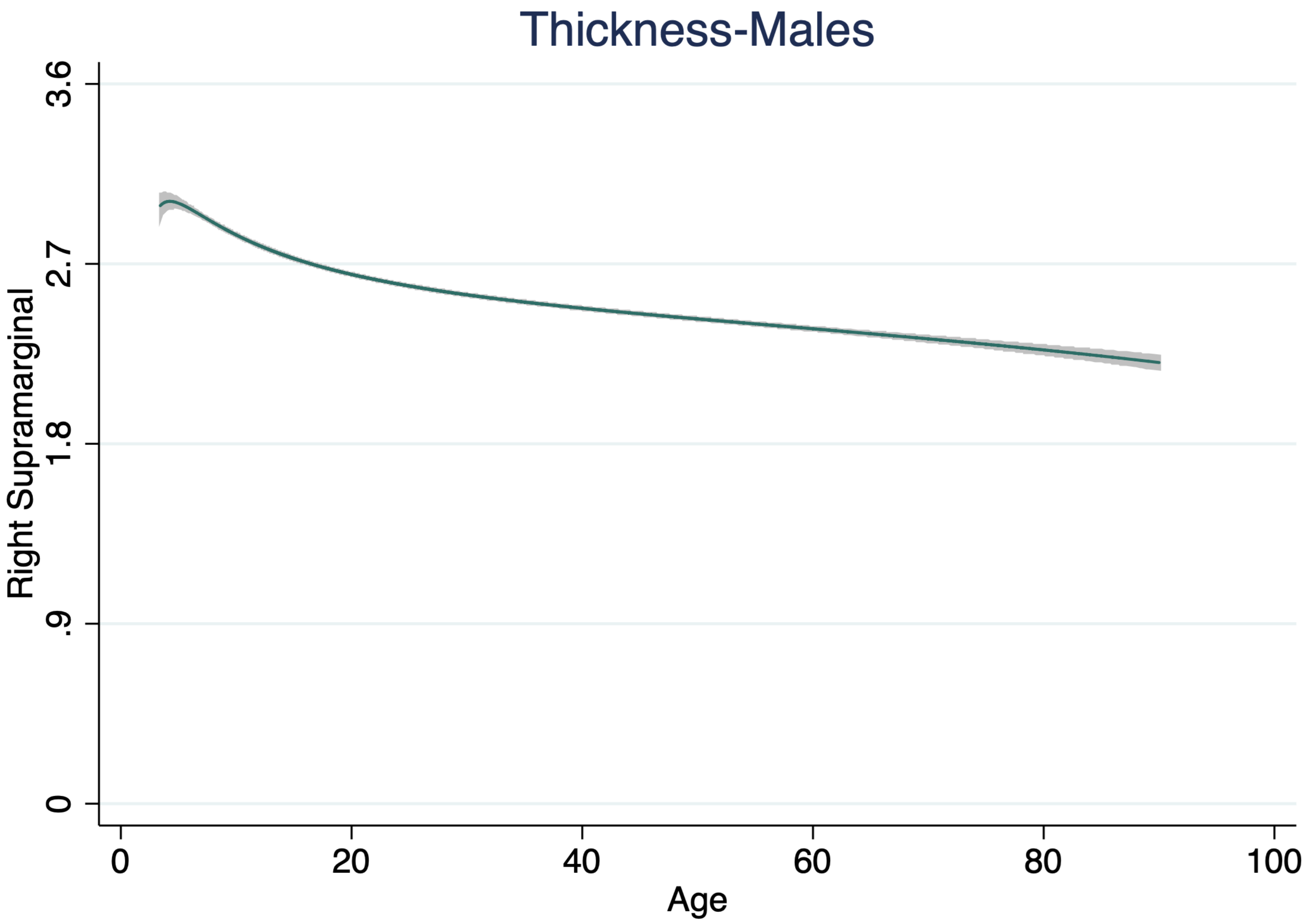

Thickness-Males
Right Supramarginal
3.6
2.7
1.8
.9
0
0
20
40
60
80
100
Age

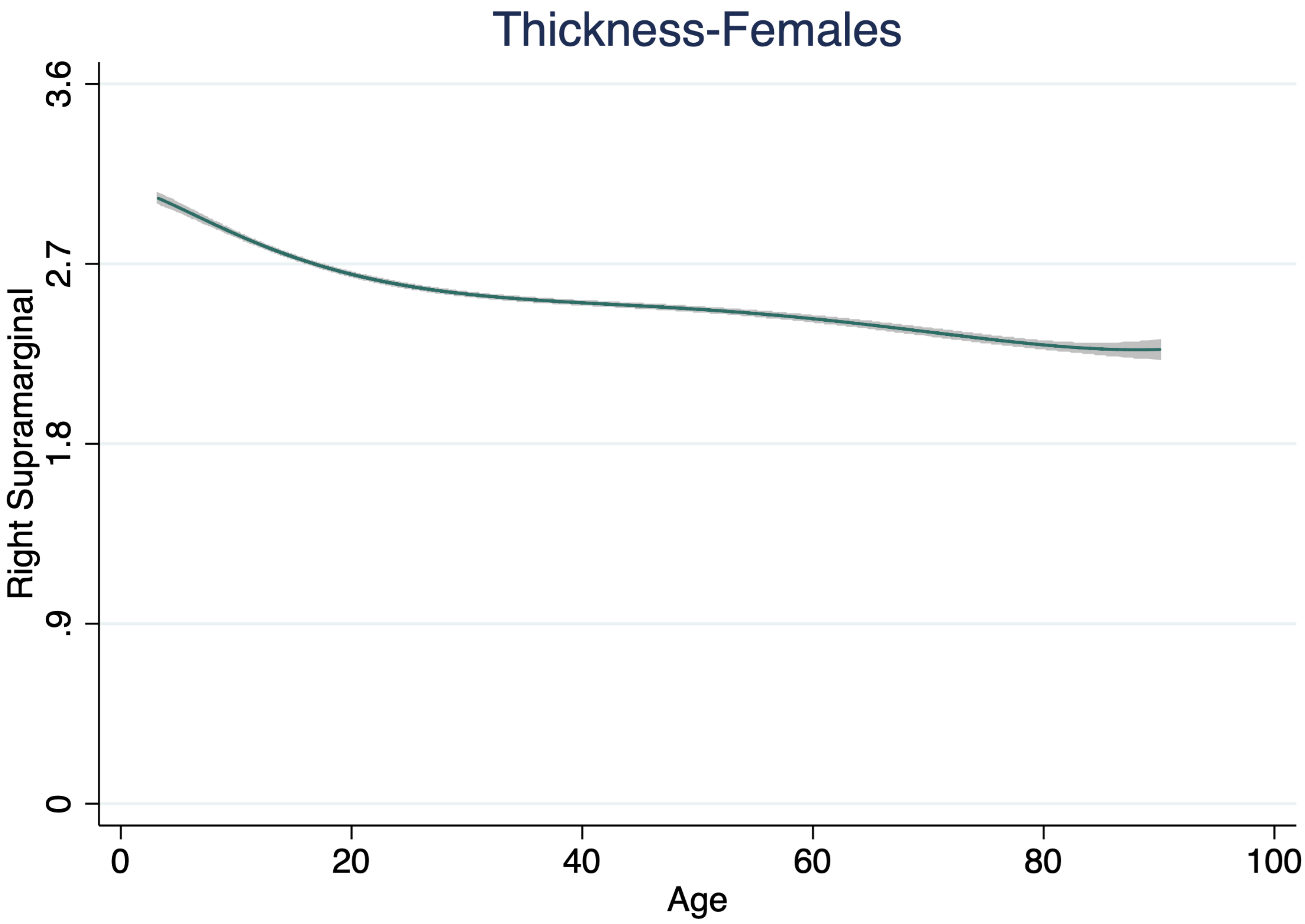

Thickness-Females
Right Supramarginal
3.6
2.7
1.8
.9
0
0
20
40
60
80
100
Age

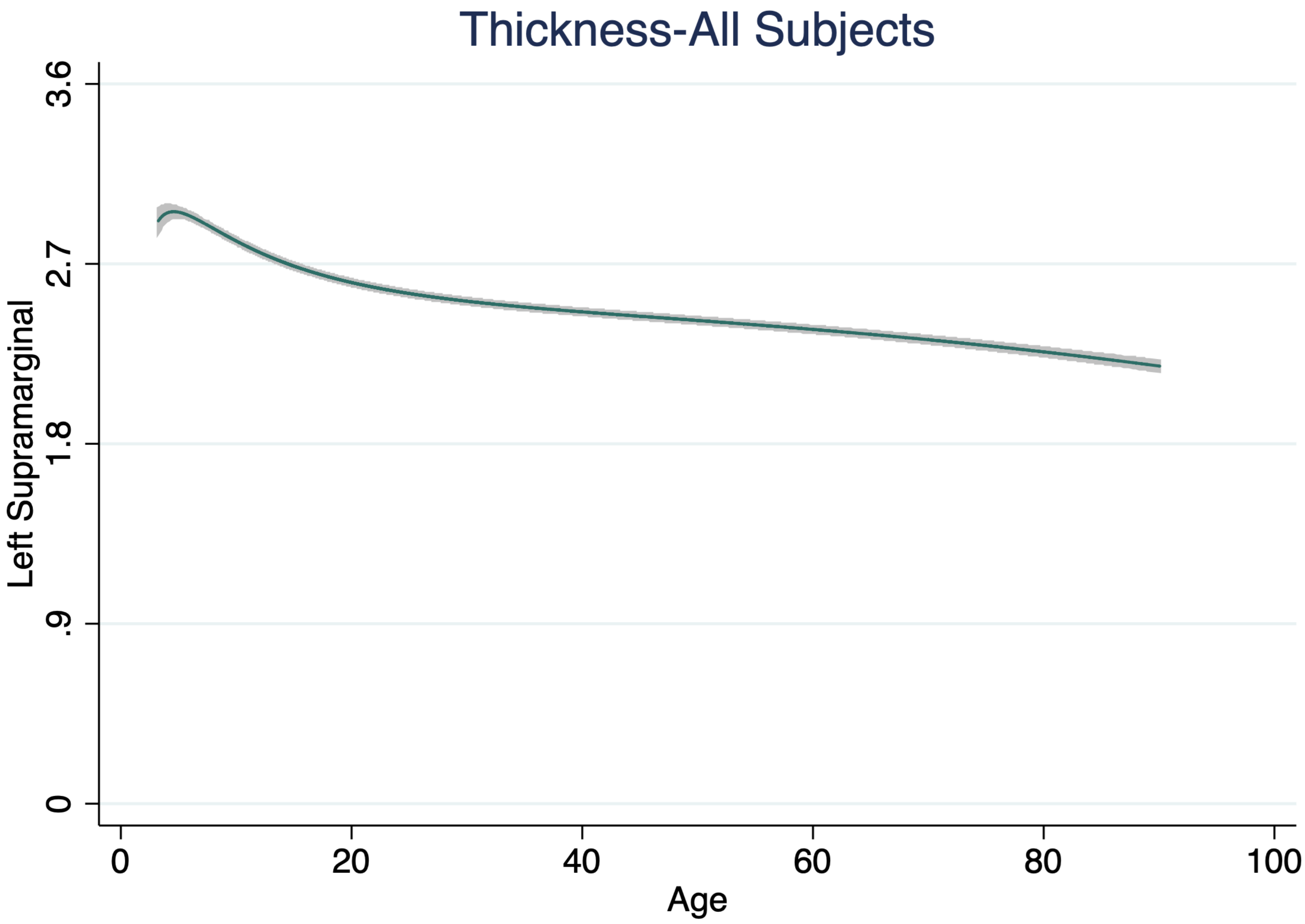

Thickness-All Subjects
Left Supramarginal
3.6
2.7
1.8
.9
0
0
20
40
60
80
100
Age

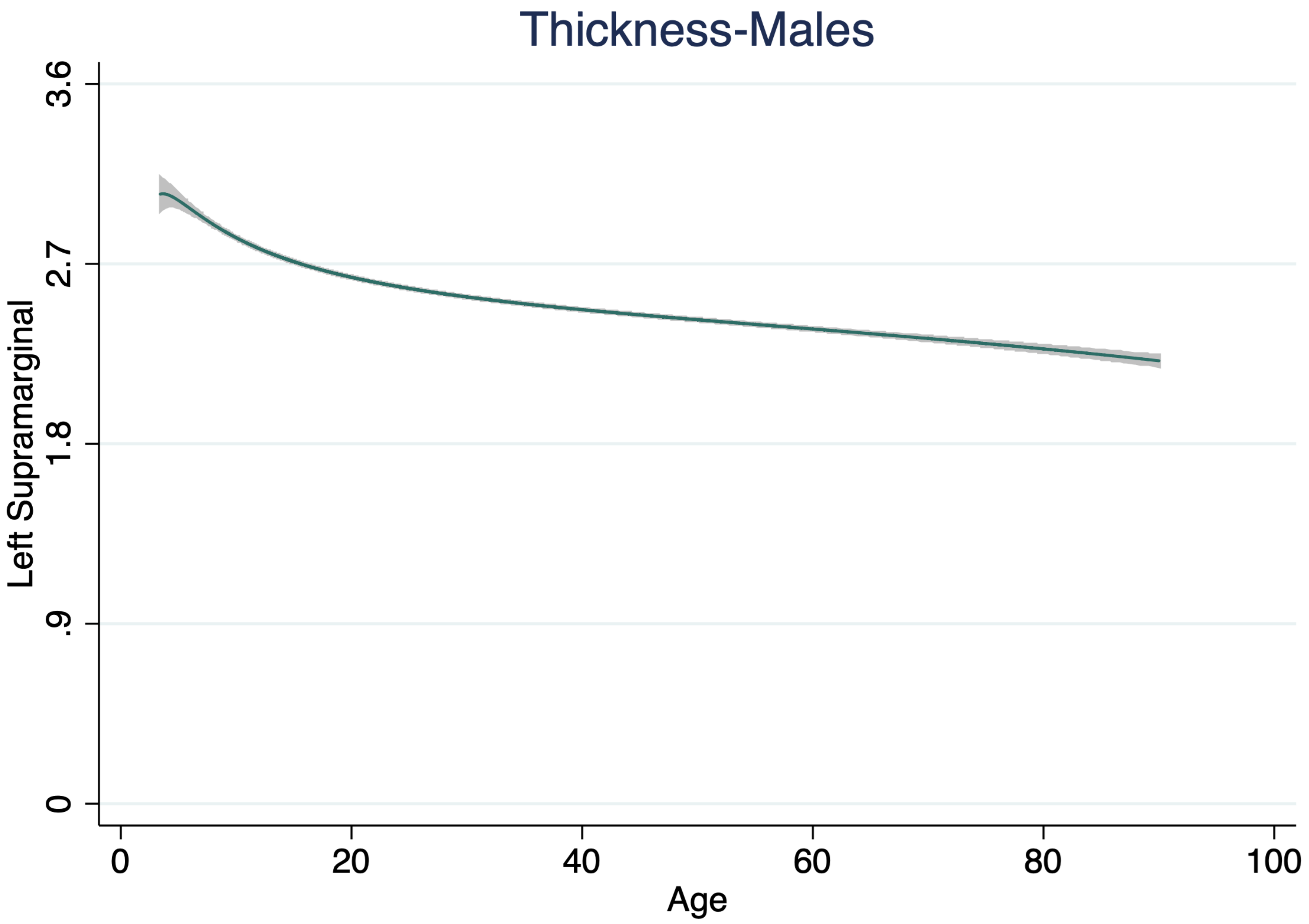
Thickness-Males
Left Supramarginal
3.6
2.7
1.8
.9
0
0
20
40
60
80
100
Age

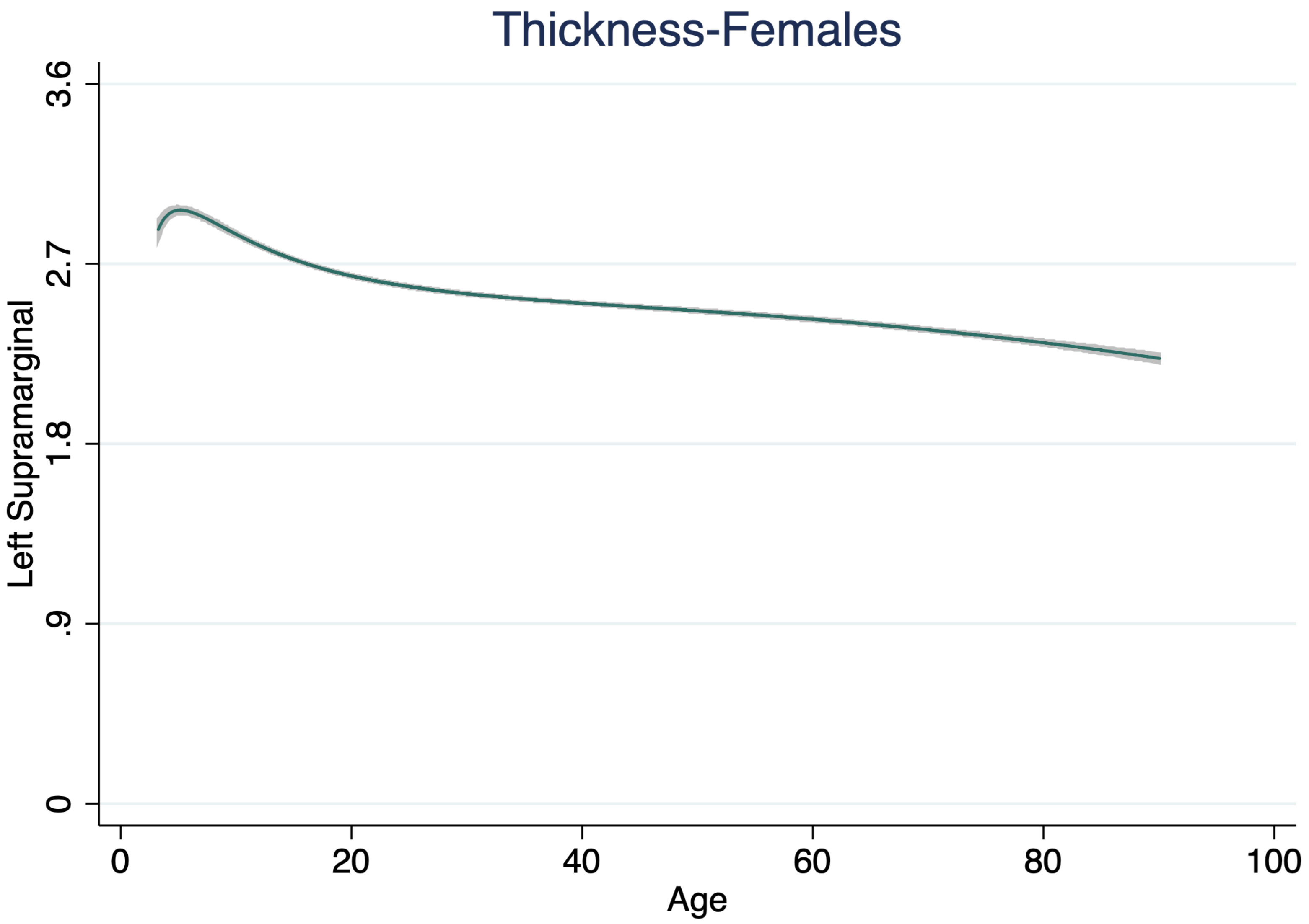
Thickness-Females
Left Supramarginal
3.6
2.7
1.8
.9
0
0
20
40
60
80
100
Age

Thickness-All Subjects

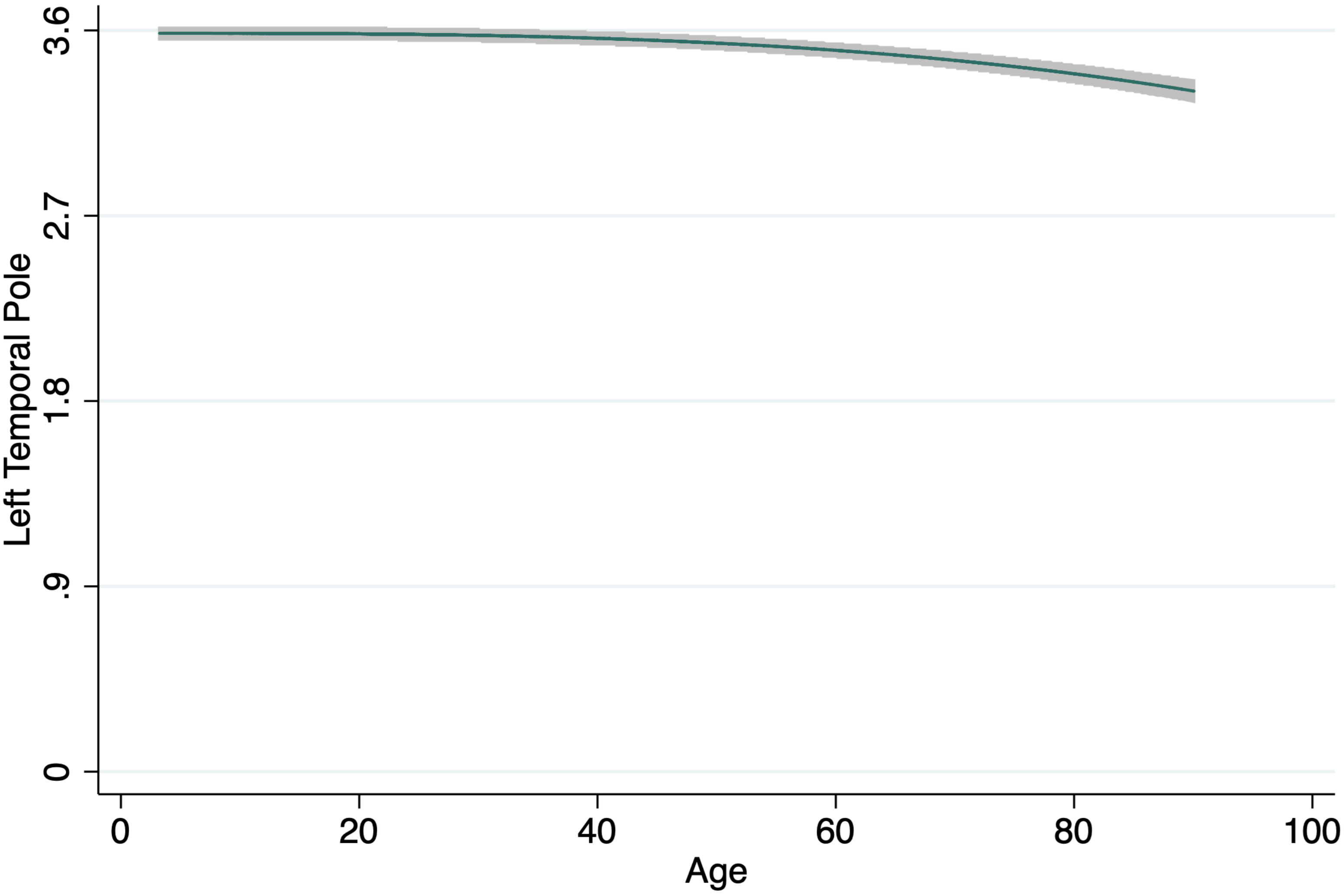

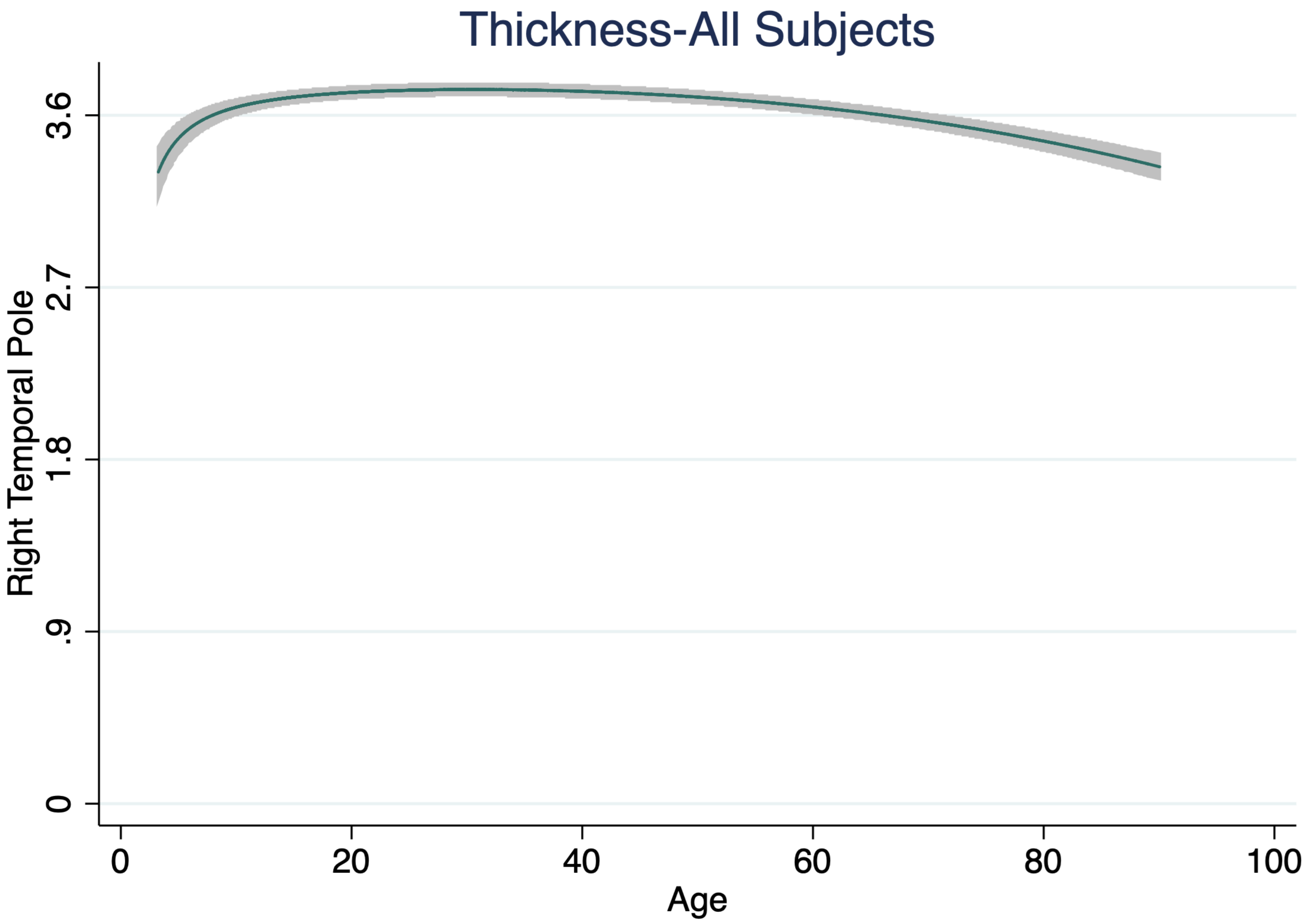

Thickness-All Subjects
Right Temporal Pole
3.6
2.7
1.8
.9
0
0
20
40
60
80
100
Age

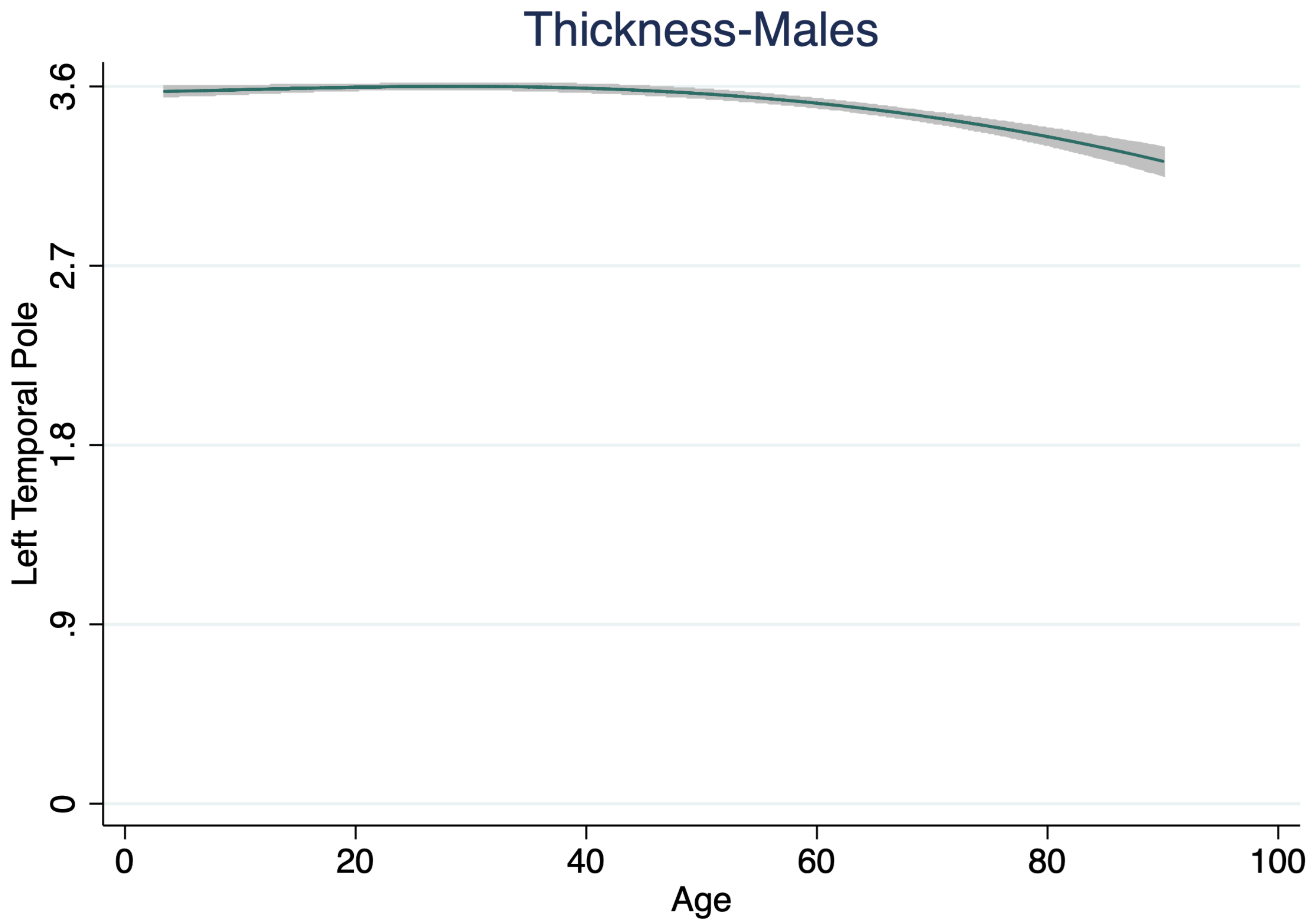

Thickness-Males
Left Temporal Pole
3.6
2.7
1.8
.9
0
0
20
40
60
80
100
Age

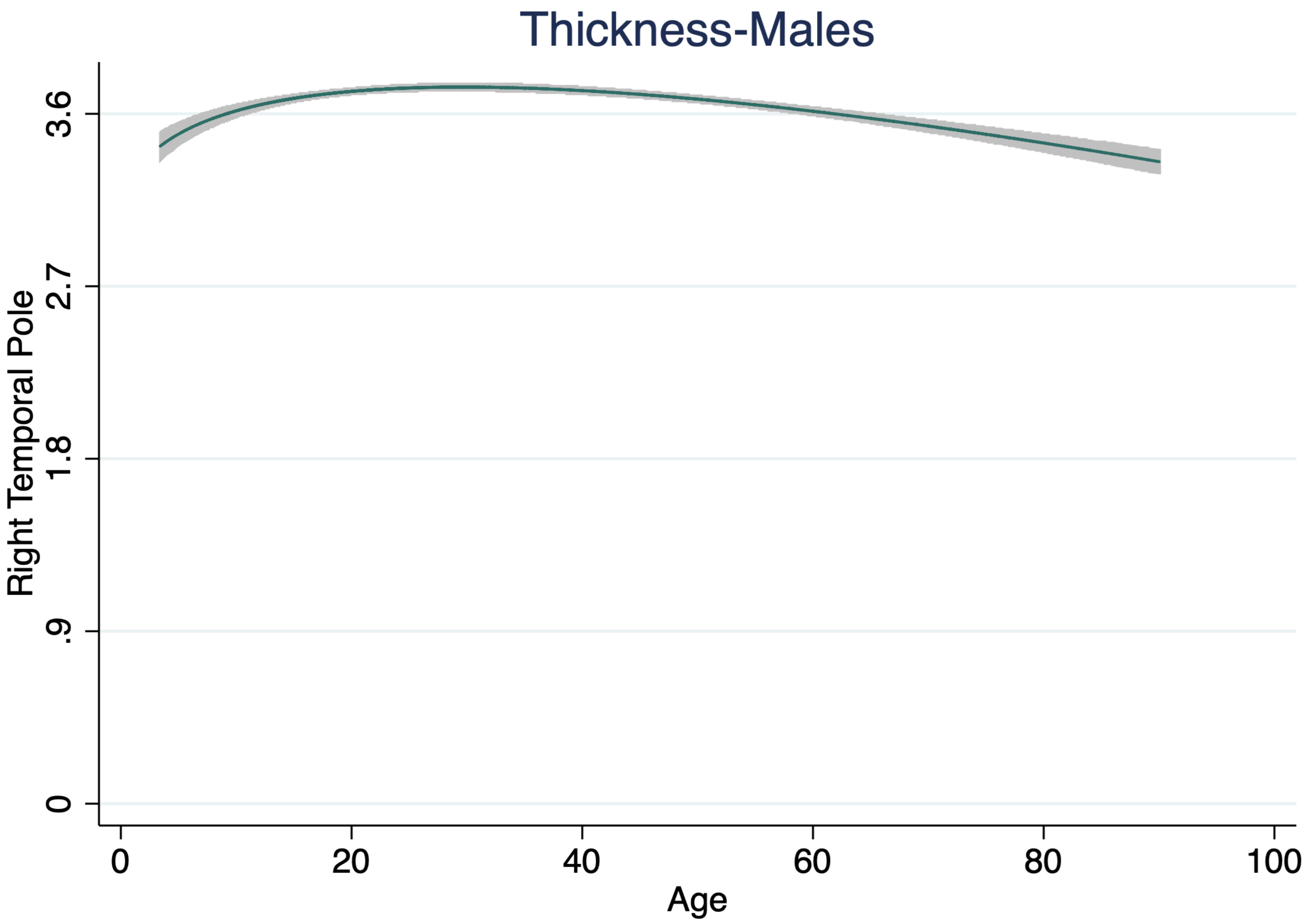

Thickness-Males
Right Temporal Pole
0
.9
1.8
2.7
3.6
0
20
40
60
80
100
Age

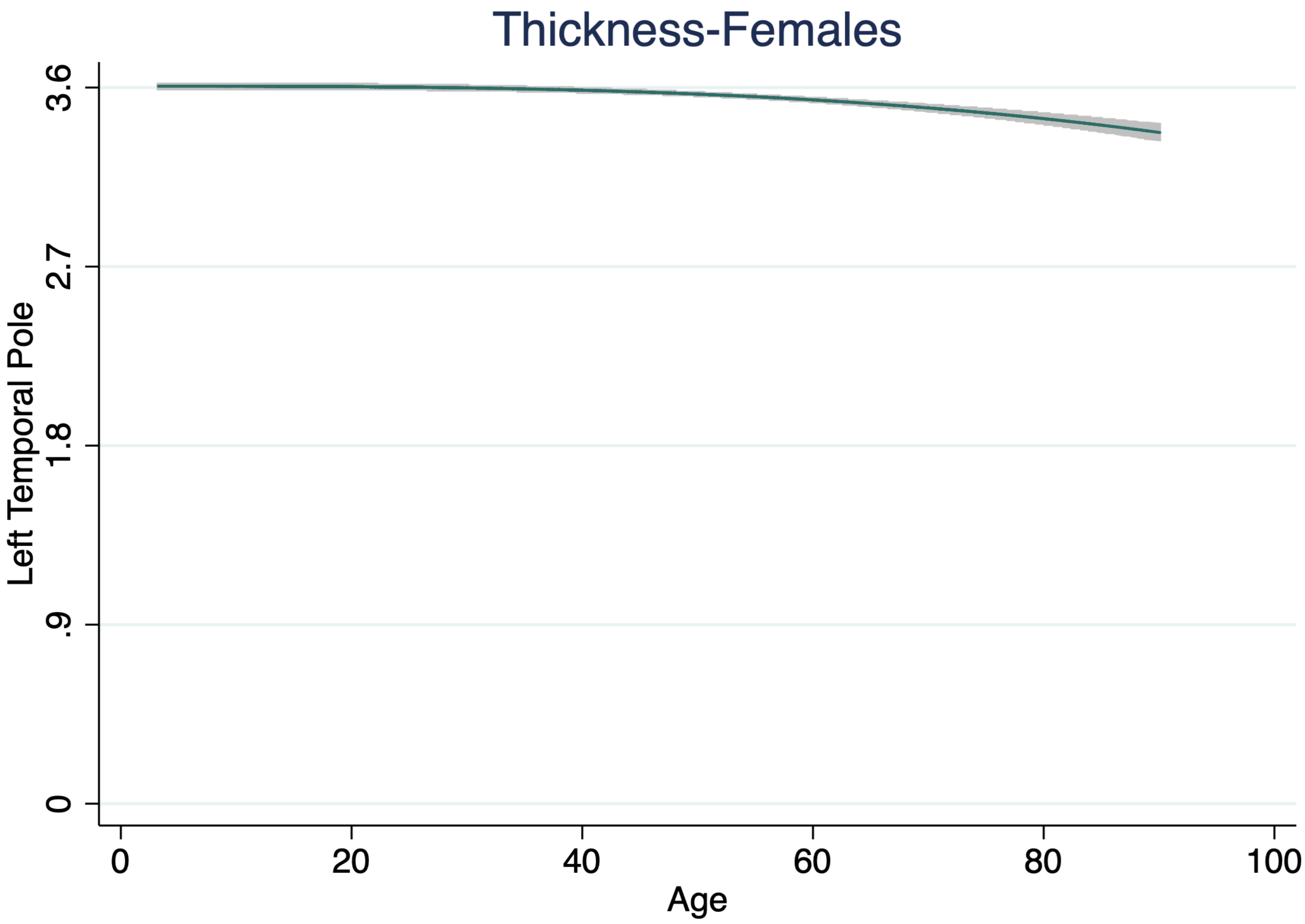

Thickness-Females
Left Temporal Pole
3.6
2.7
1.8
.9
0
0
20
40
60
80
100
Age

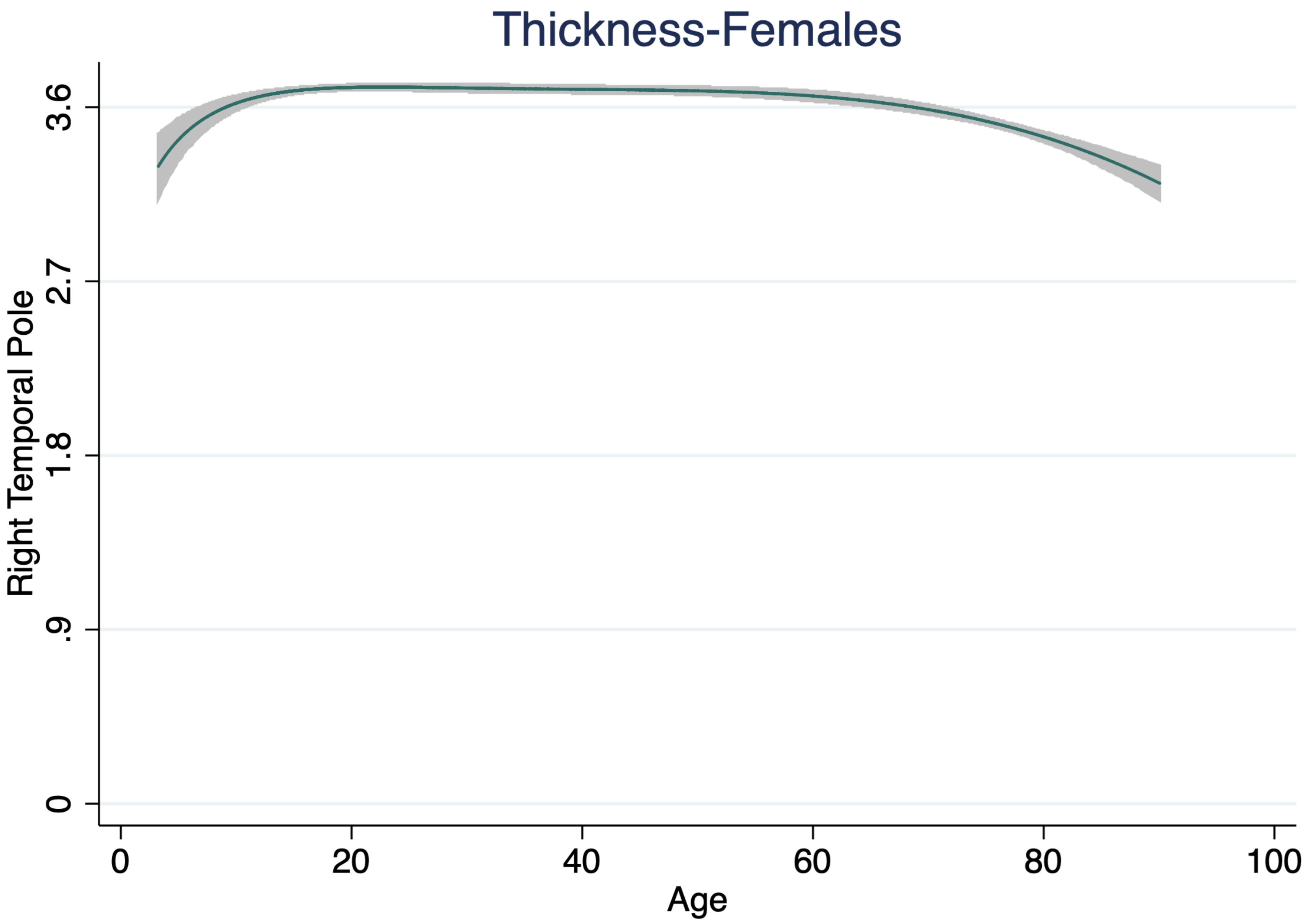

Thickness-Females
Right Temporal Pole
3.6
2.7
1.8
.9
0
0
20
40
60
80
100
Age

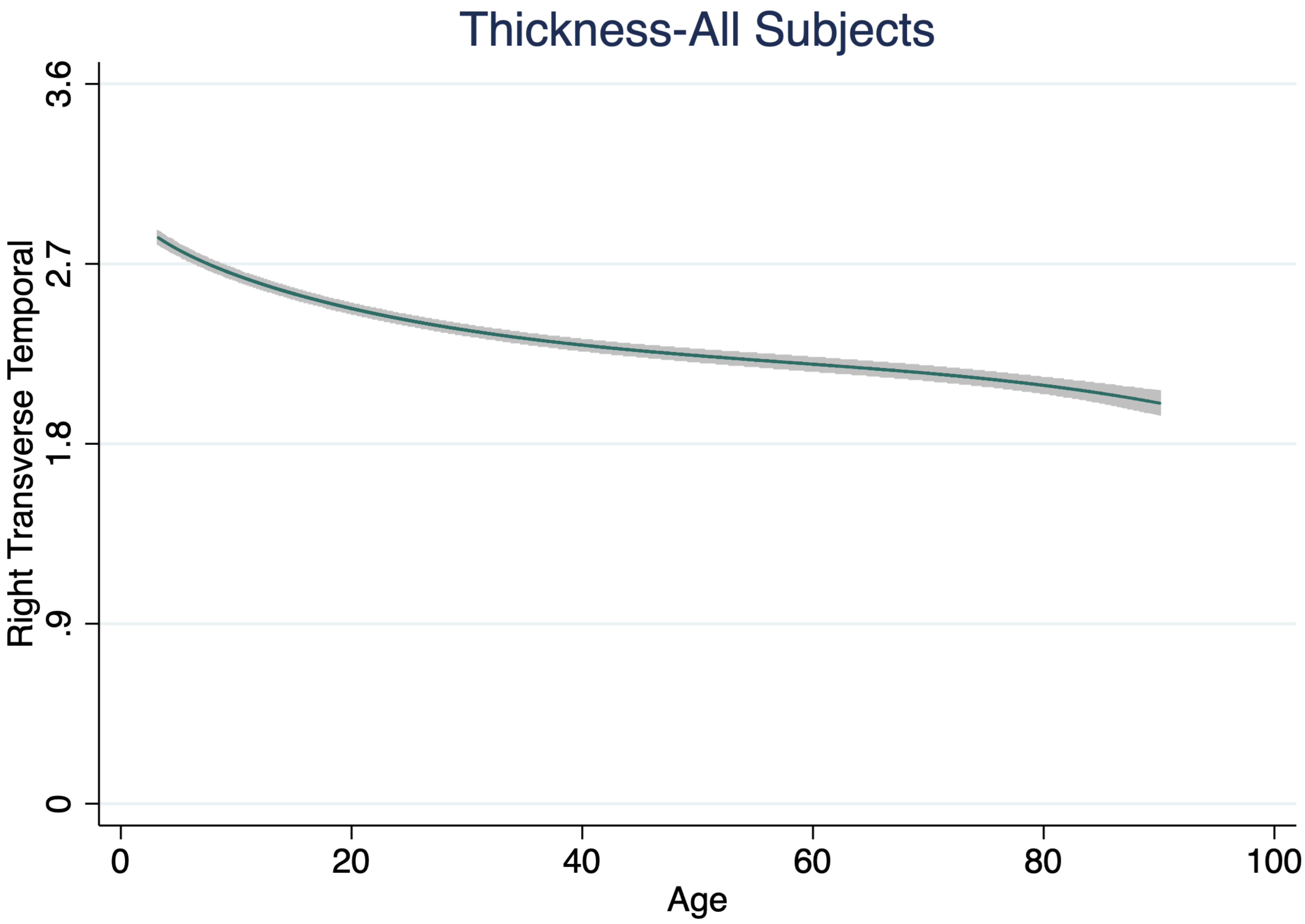

Thickness-All Subjects
Right Transverse Temporal
3.6
2.7
1.8
.9
0
0
20
40
60
80
100
Age

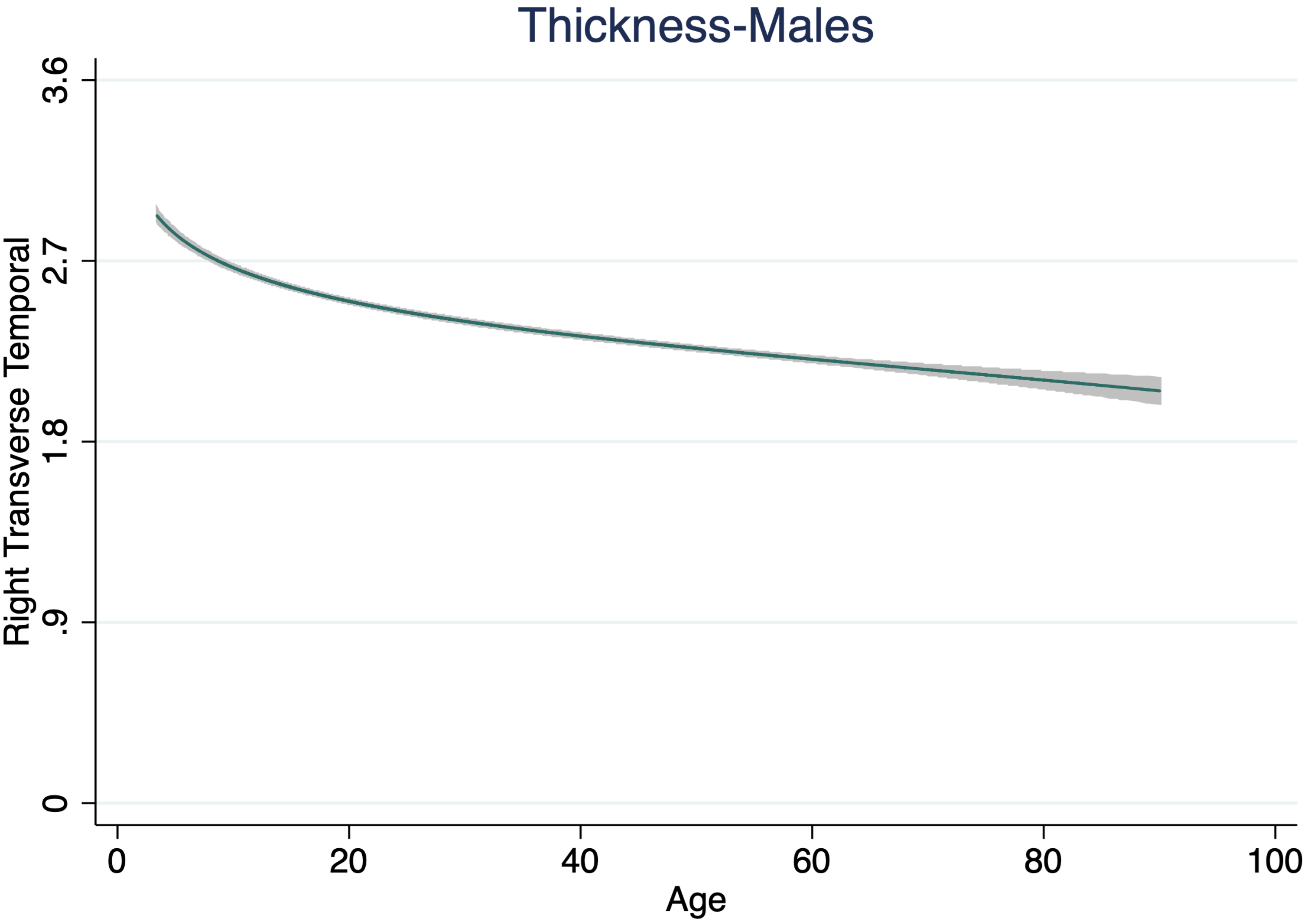

Thickness-Males
Right Transverse Temporal
3.6
2.7
1.8
.9
0
0
20
40
60
80
100
Age

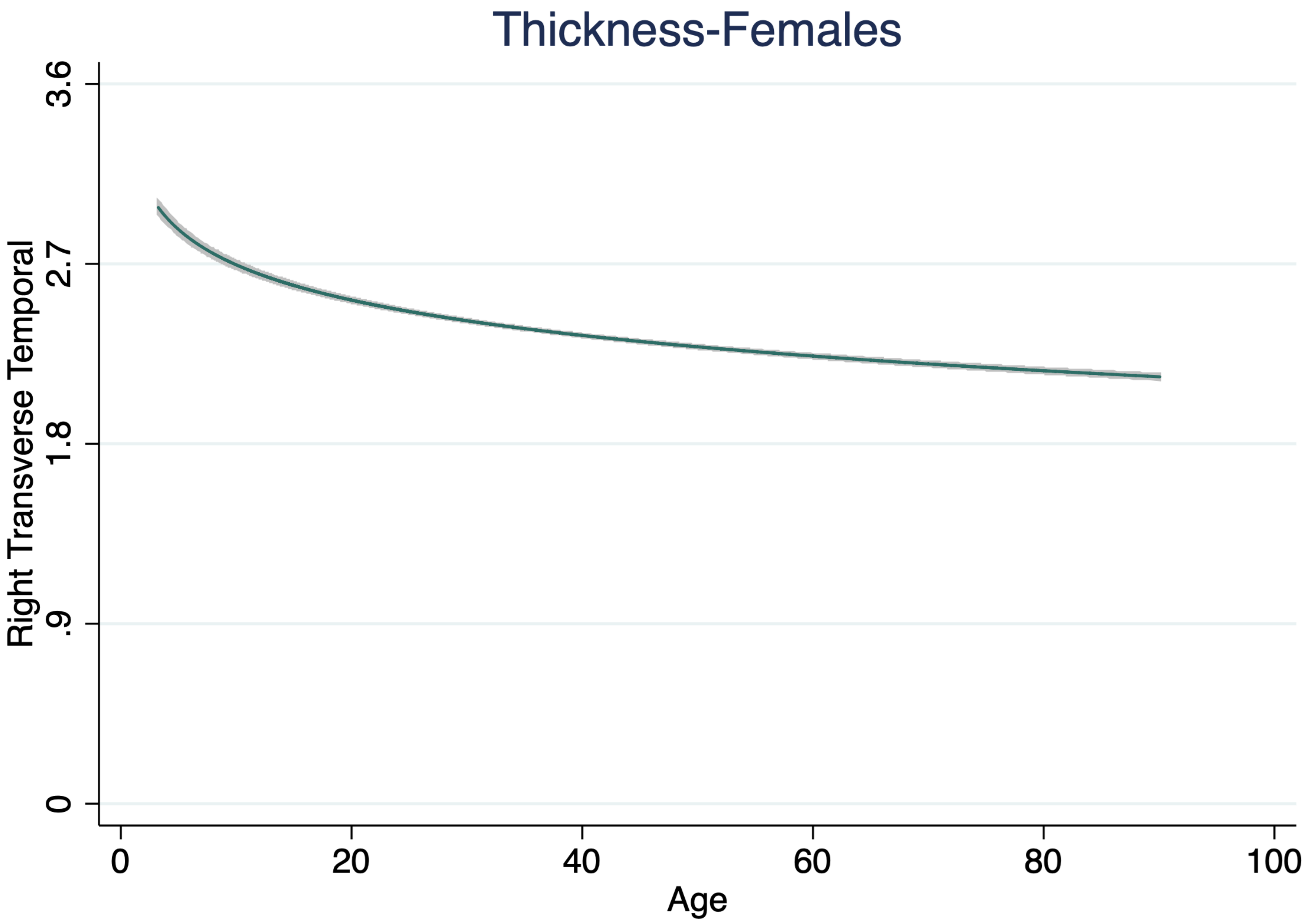

Thickness-Females
Right Transverse Temporal
3.6
2.7
1.8
.9
0
0
20
40
60
80
100
Age

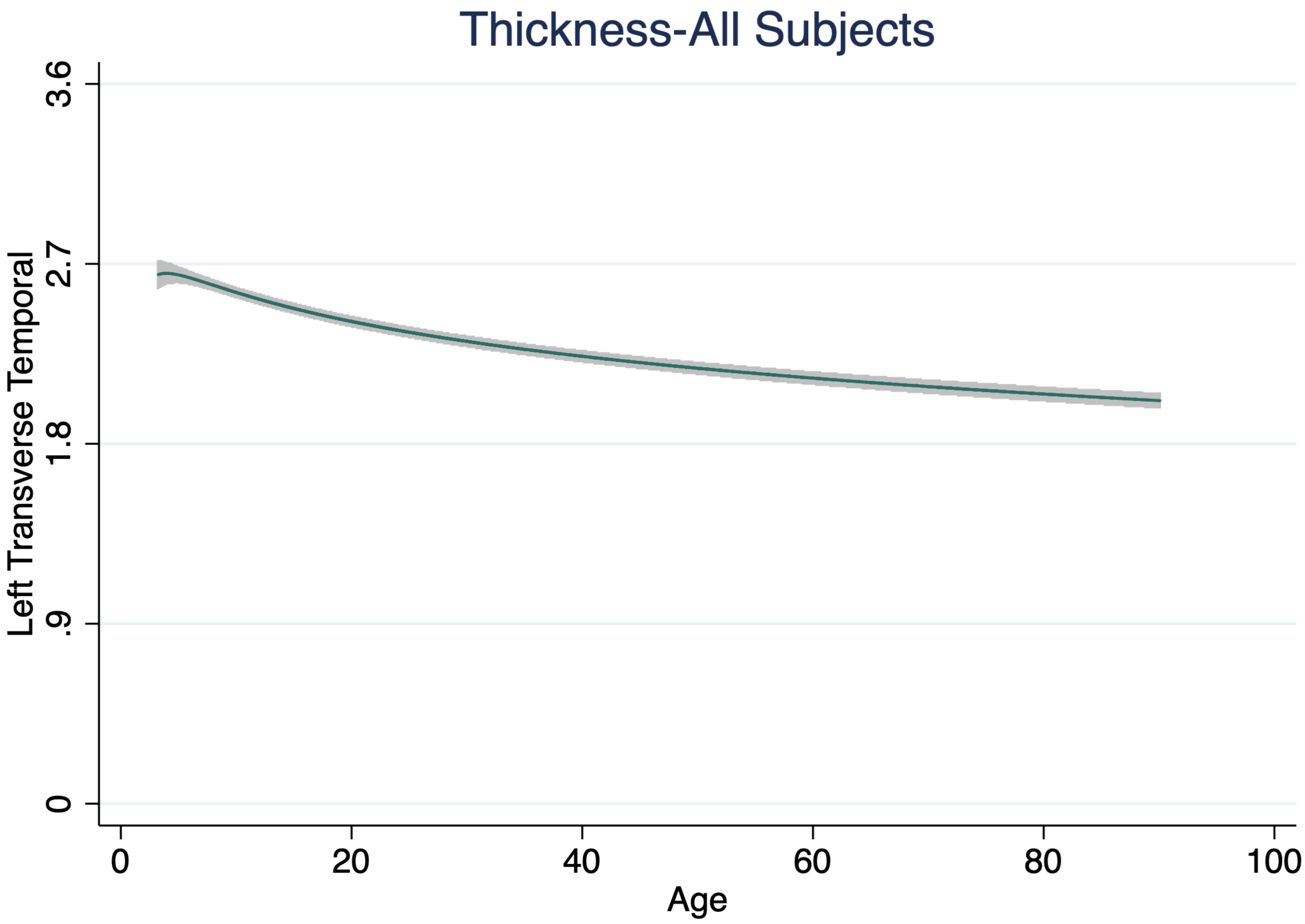
Thickness-All Subjects
Left Transverse Temporal
3.6
2.7
1.8
.9
0
0
20
40
60
80
100
Age

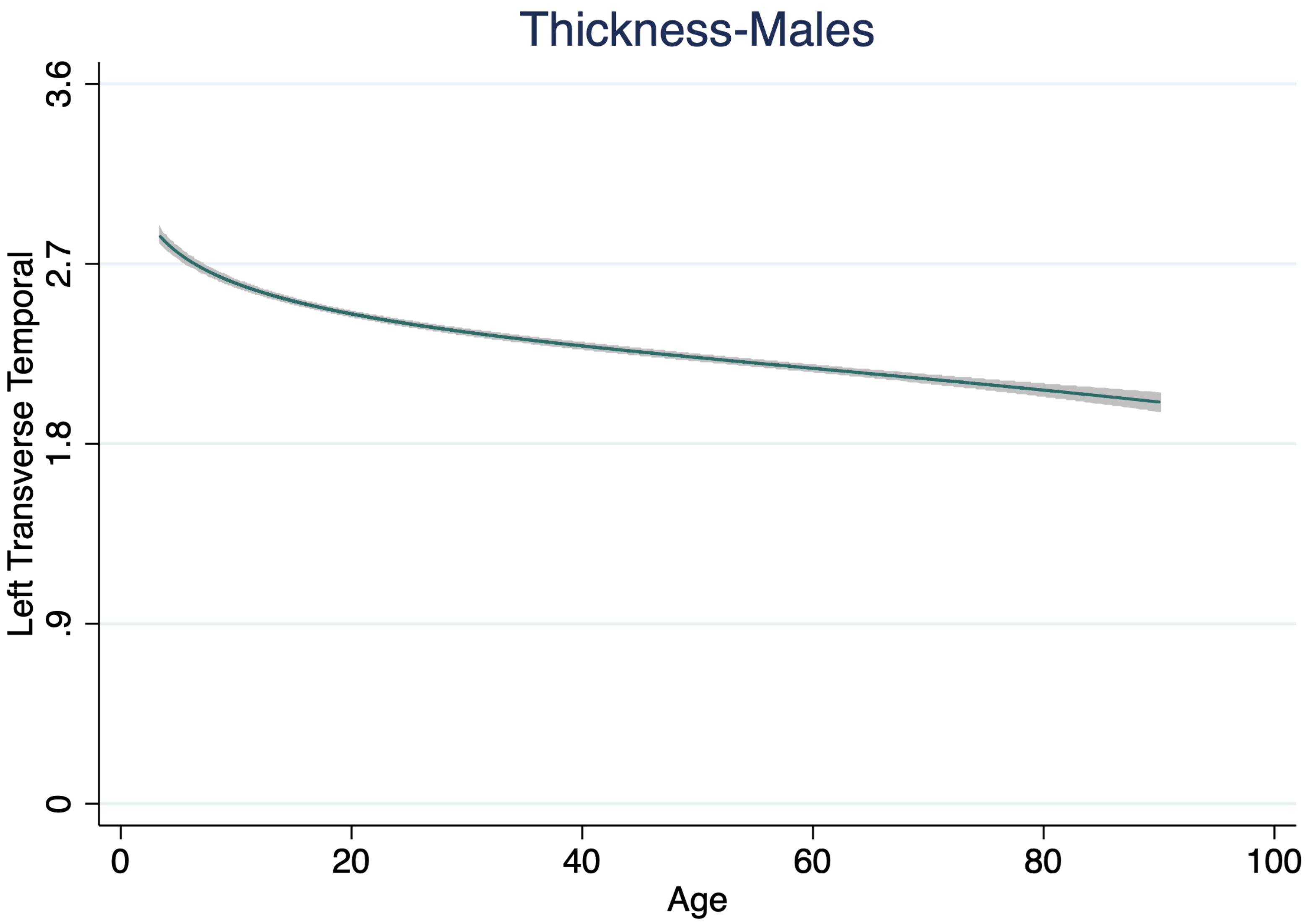
Thickness-Males
Left Transverse Temporal
3.6
2.7
1.8
.9
0
0
20
40
60
80
100
Age

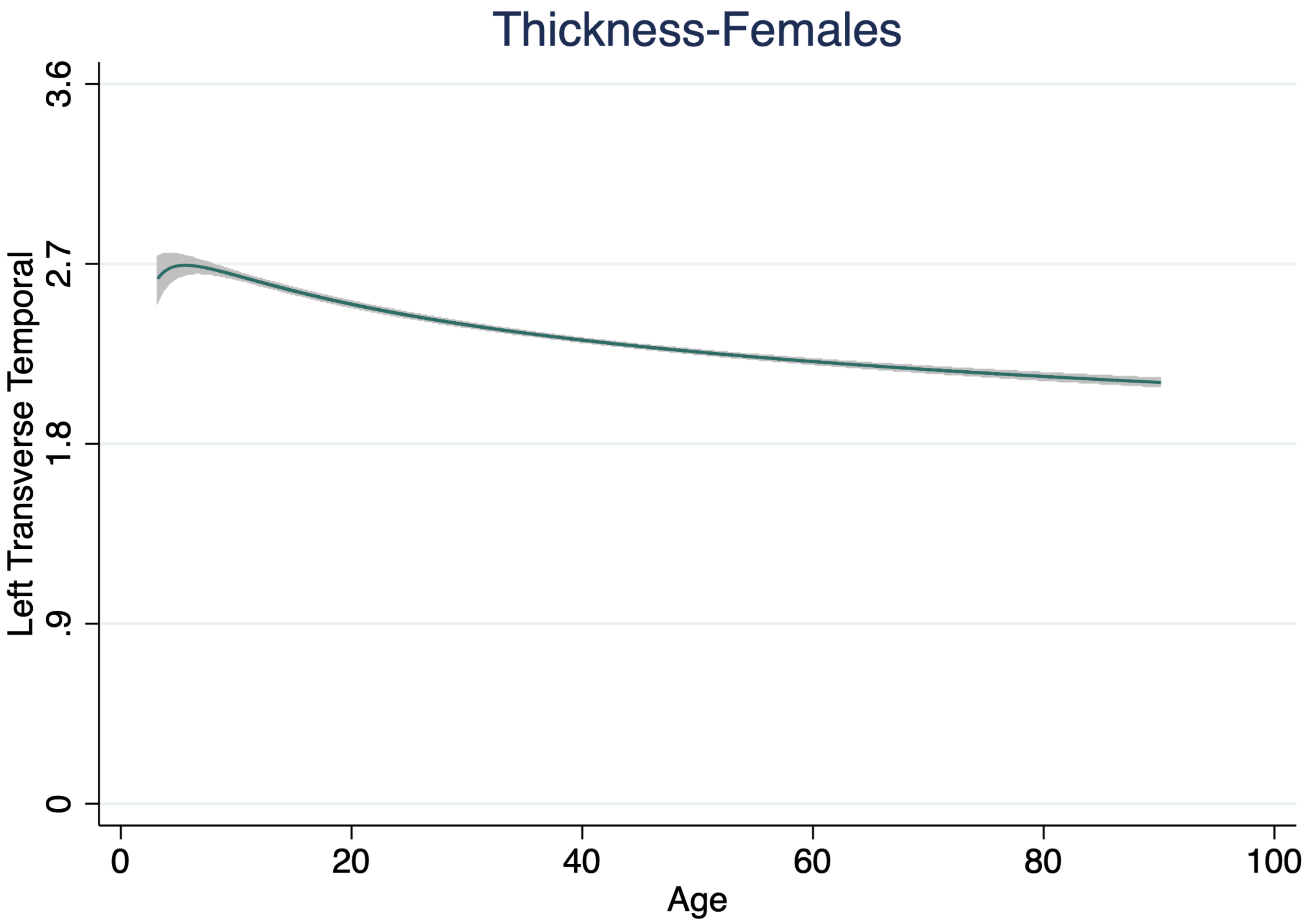

Thickness-Females
Left Transverse Temporal
0
.9
1.8
2.7
3.6
0
20
40
60
80
100
Age
