## Supplementary File 2 for "Cortical Thickness Trajectories across the Lifespan: Data from 17,075 healthy individuals aged 3-90 years"

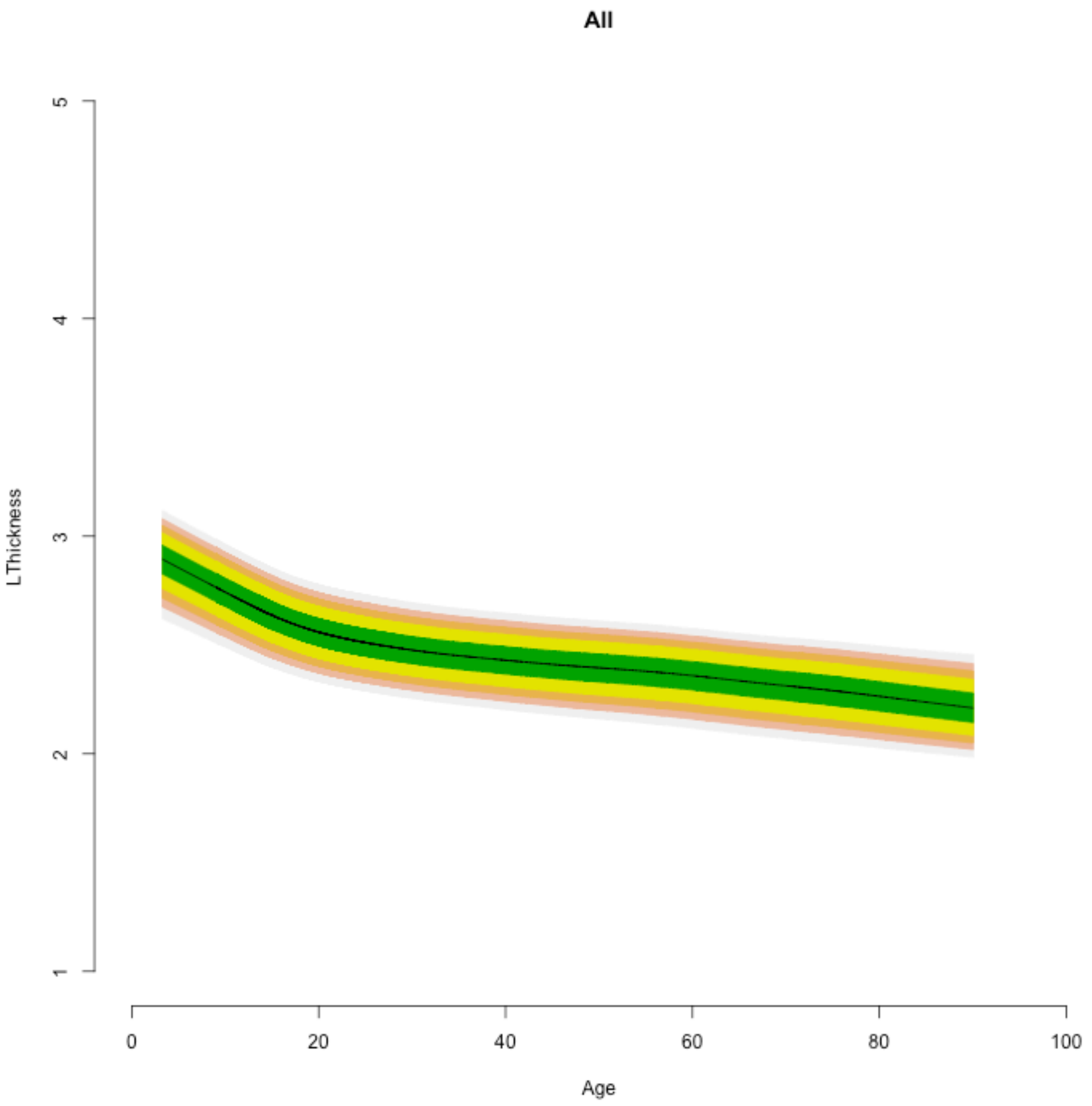

All
LThickness
Age
5
4
3
2
1
0
20
40
60
80
100

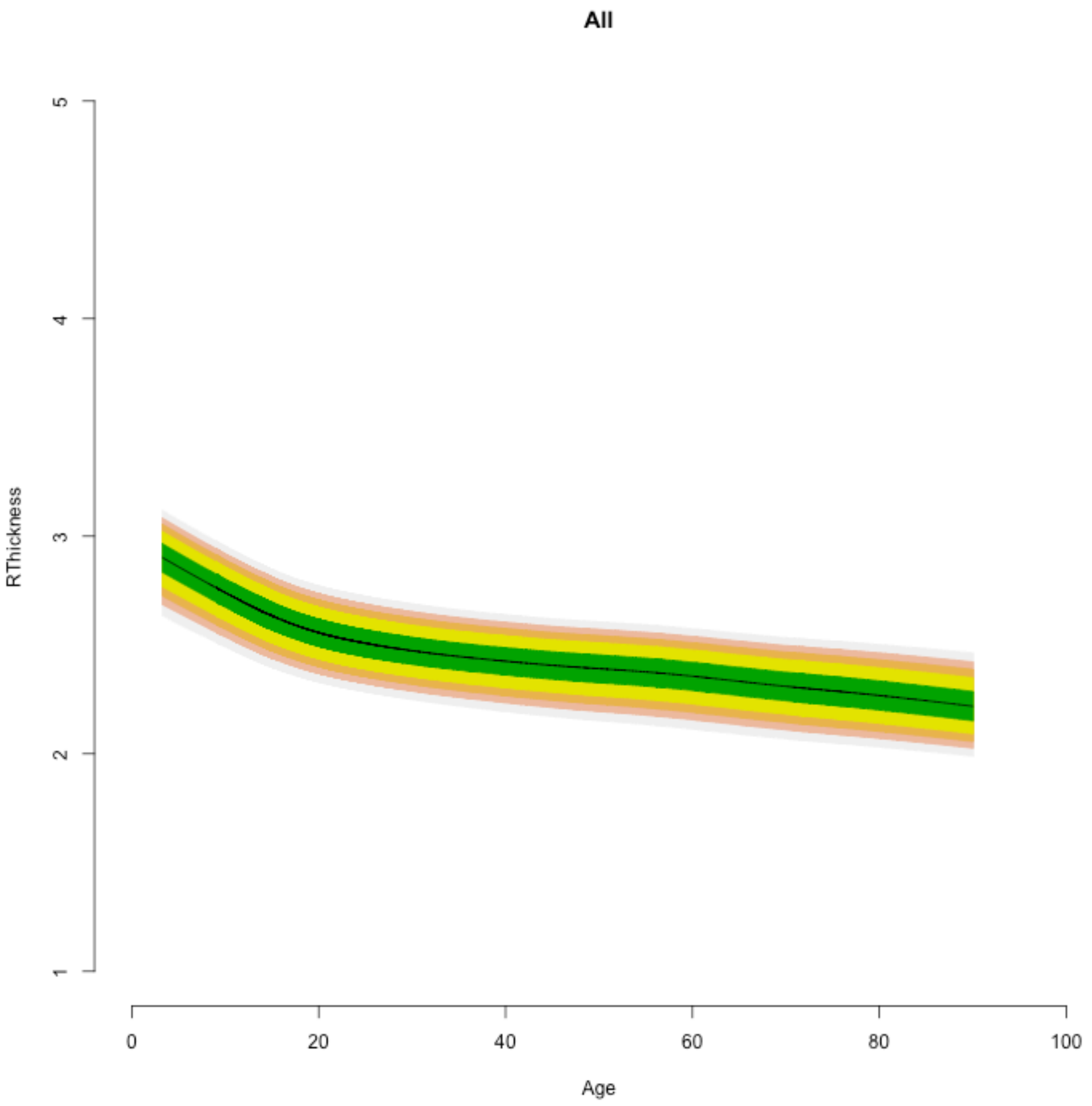
All
RThickness
5
4
3
2
1
0
20
40
60
80
100
Age

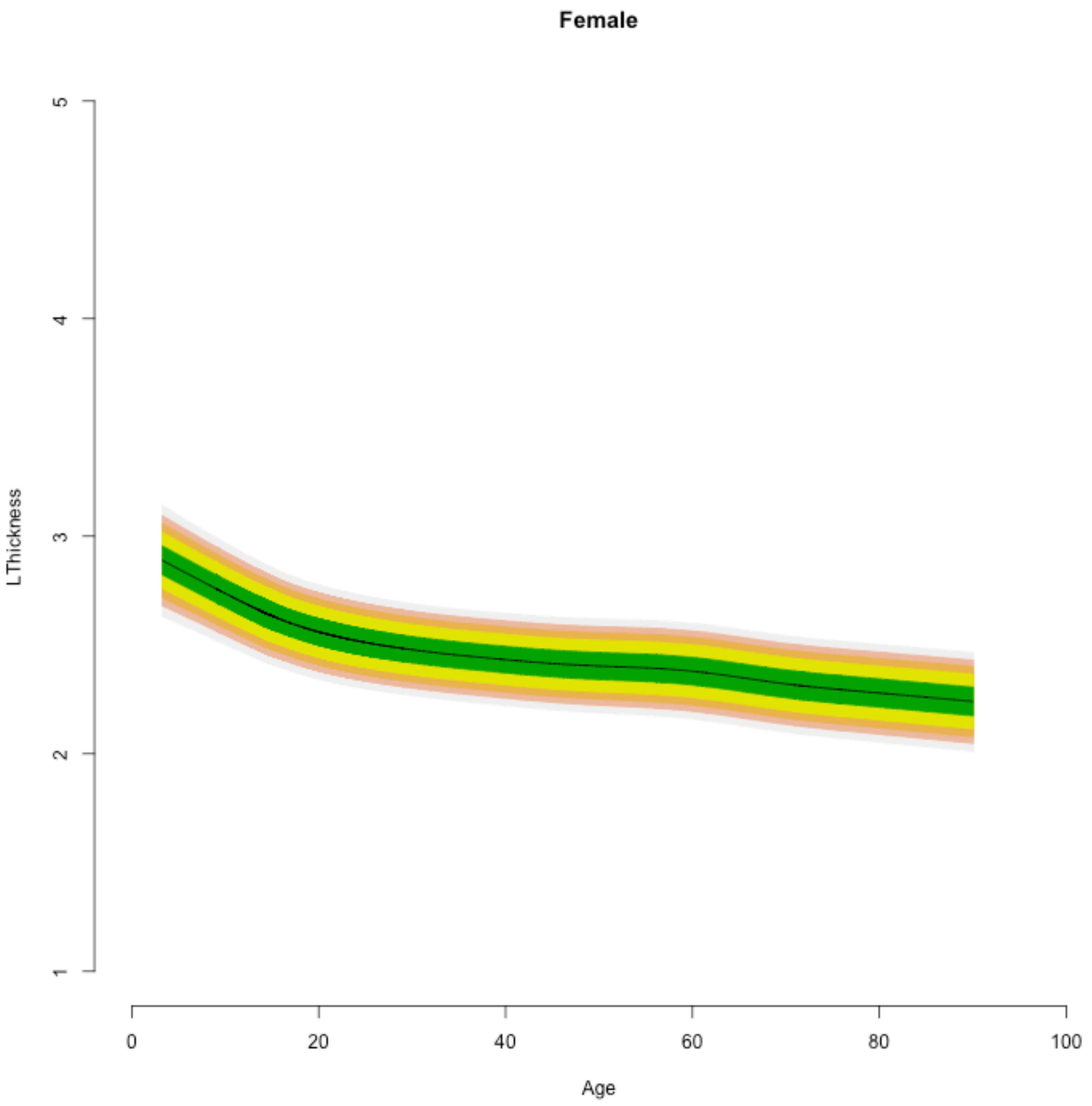

Female
LThickness
5
4
3
2
1
0
20
40
60
80
100
Age

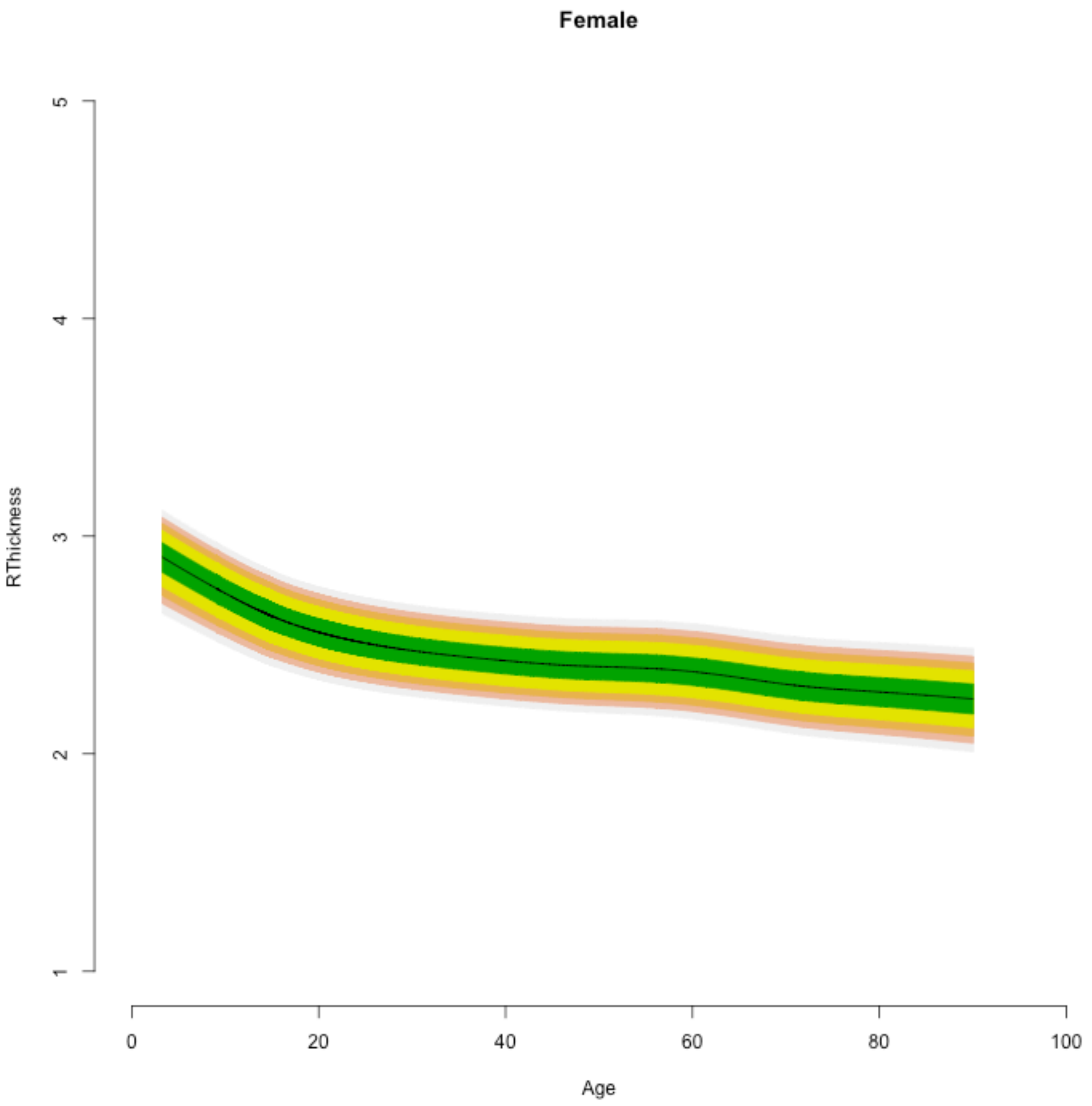
Female
RThickness
5
4
3
2
1
0
20
40
60
80
100
Age

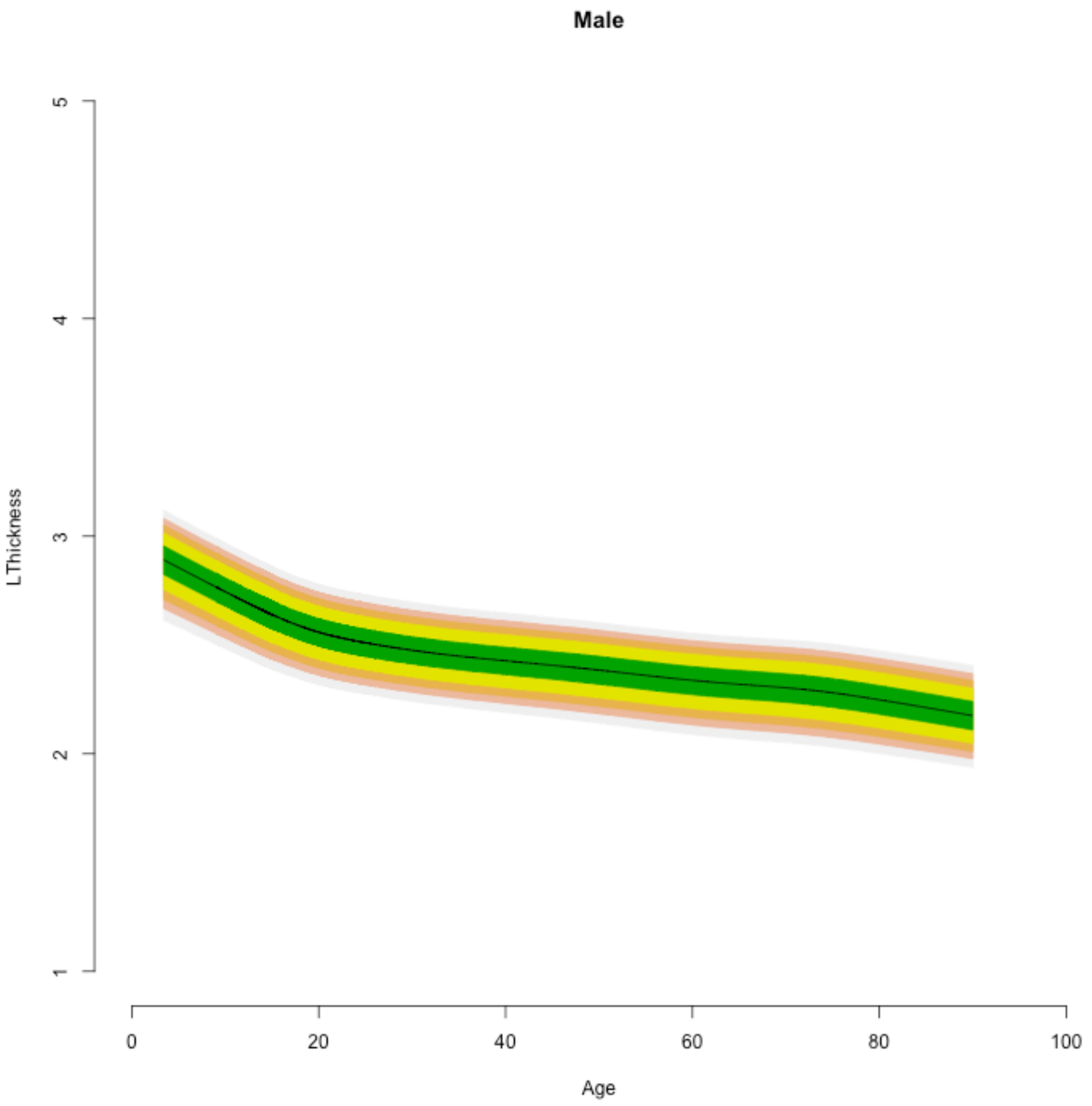

Male
LThickness
1
2
3
4
5
0
20
40
60
80
100
Age

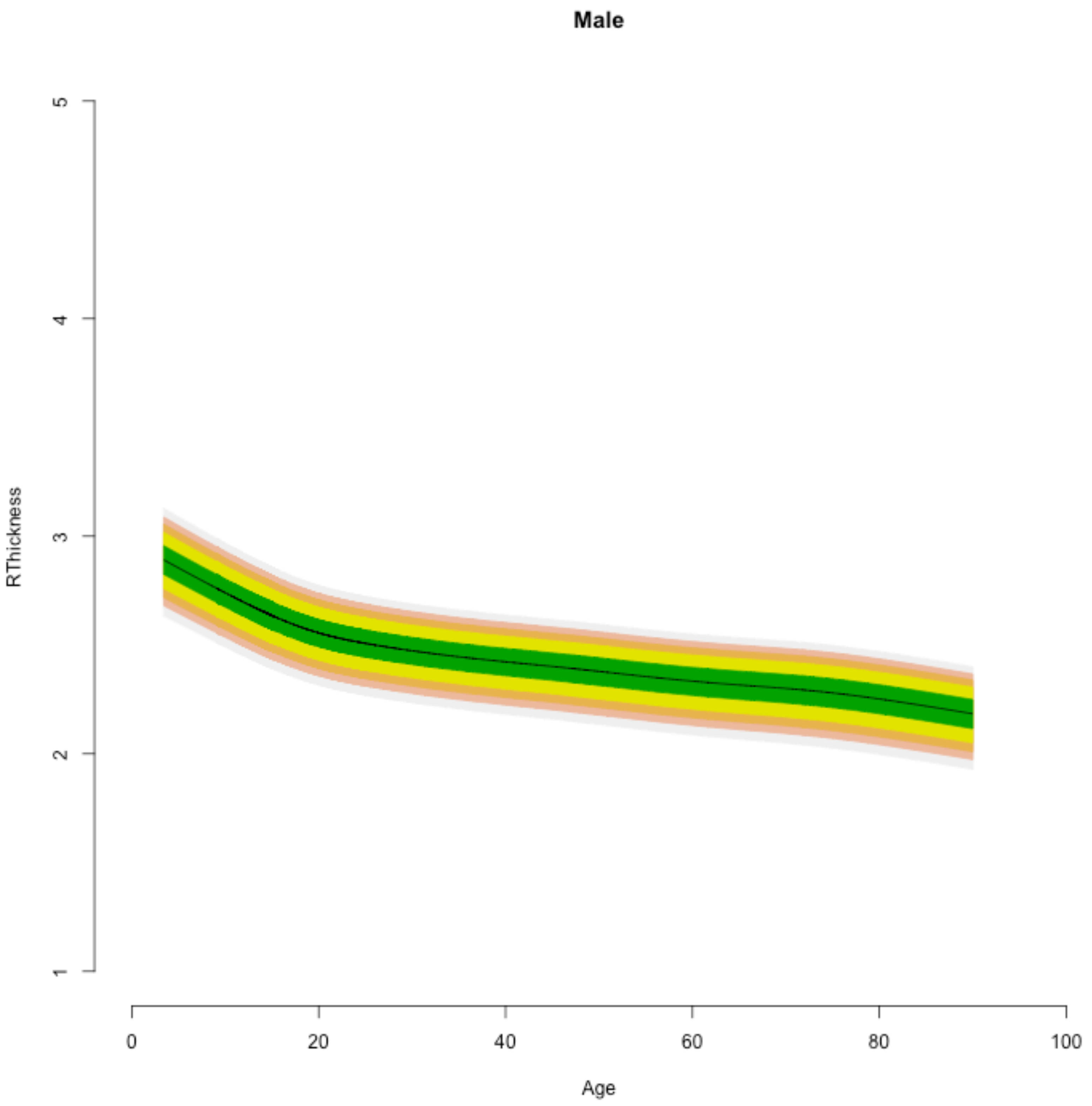

Male
RThickness
5
4
3
2
1
0
20
40
60
80
100
Age

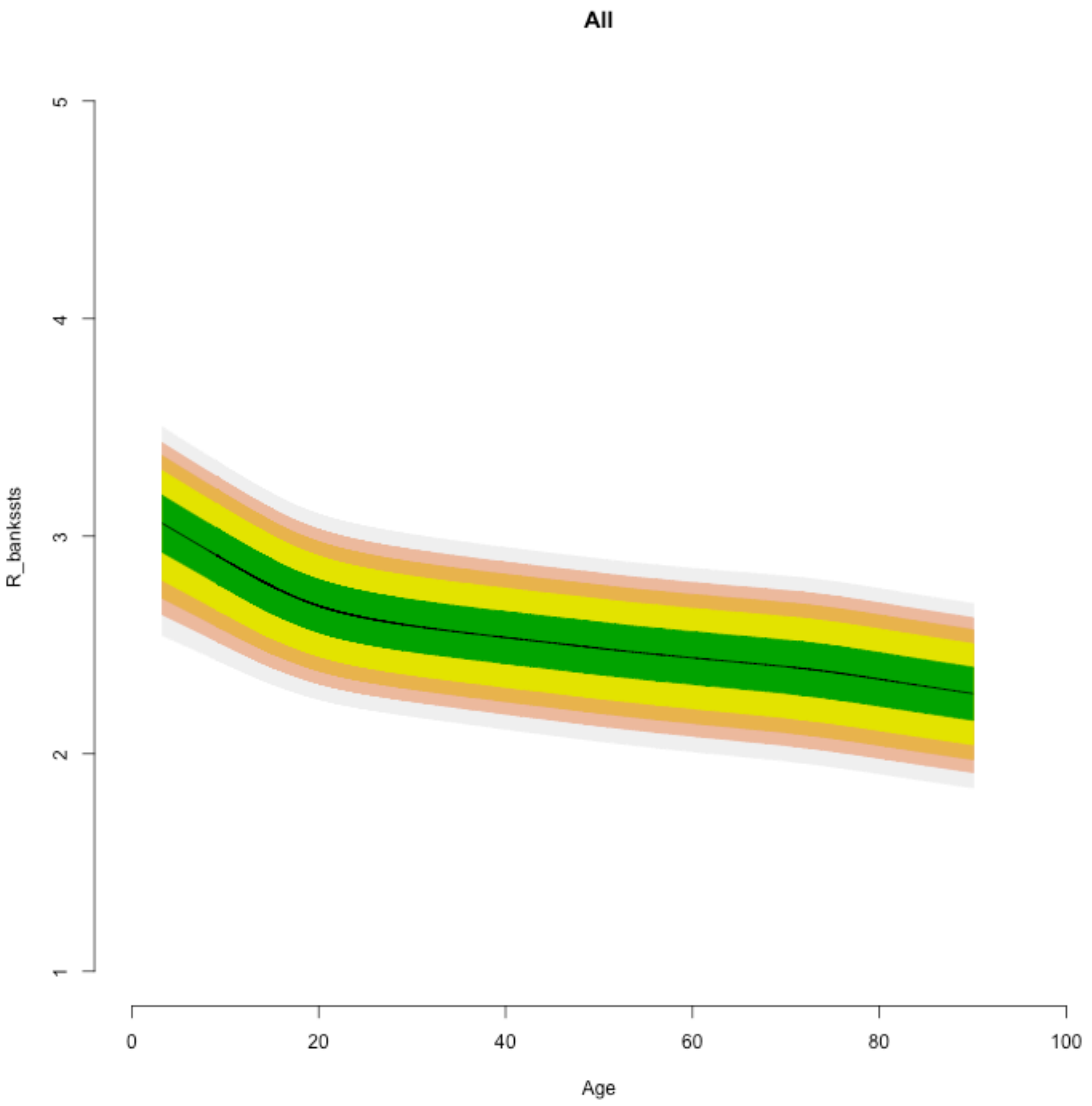
All
R_bankssts
1
2
3
4
5
0
20
40
60
80
100
Age

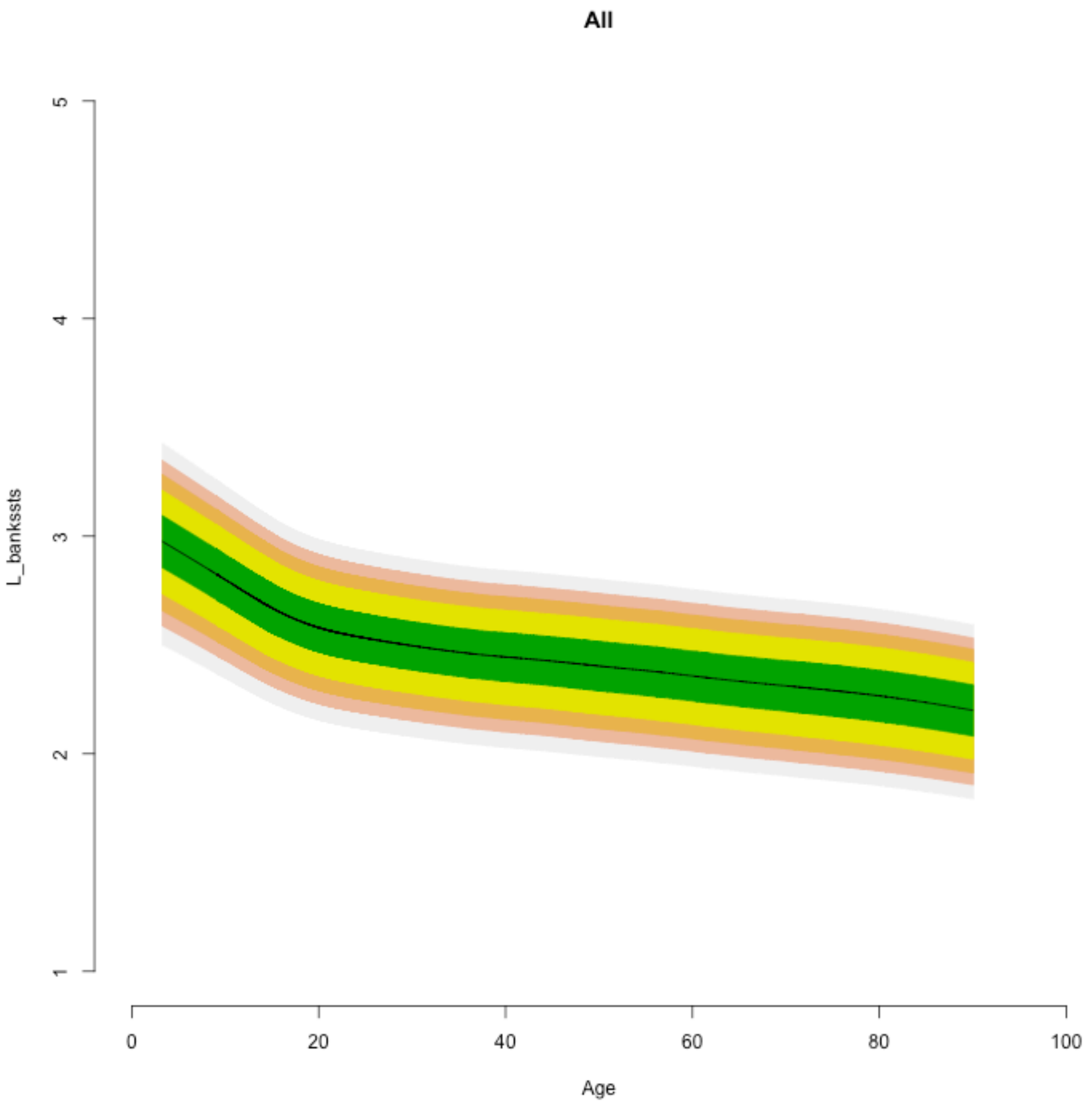
All
L_bankssts
1
2
3
4
5
0
20
40
60
80
100
Age

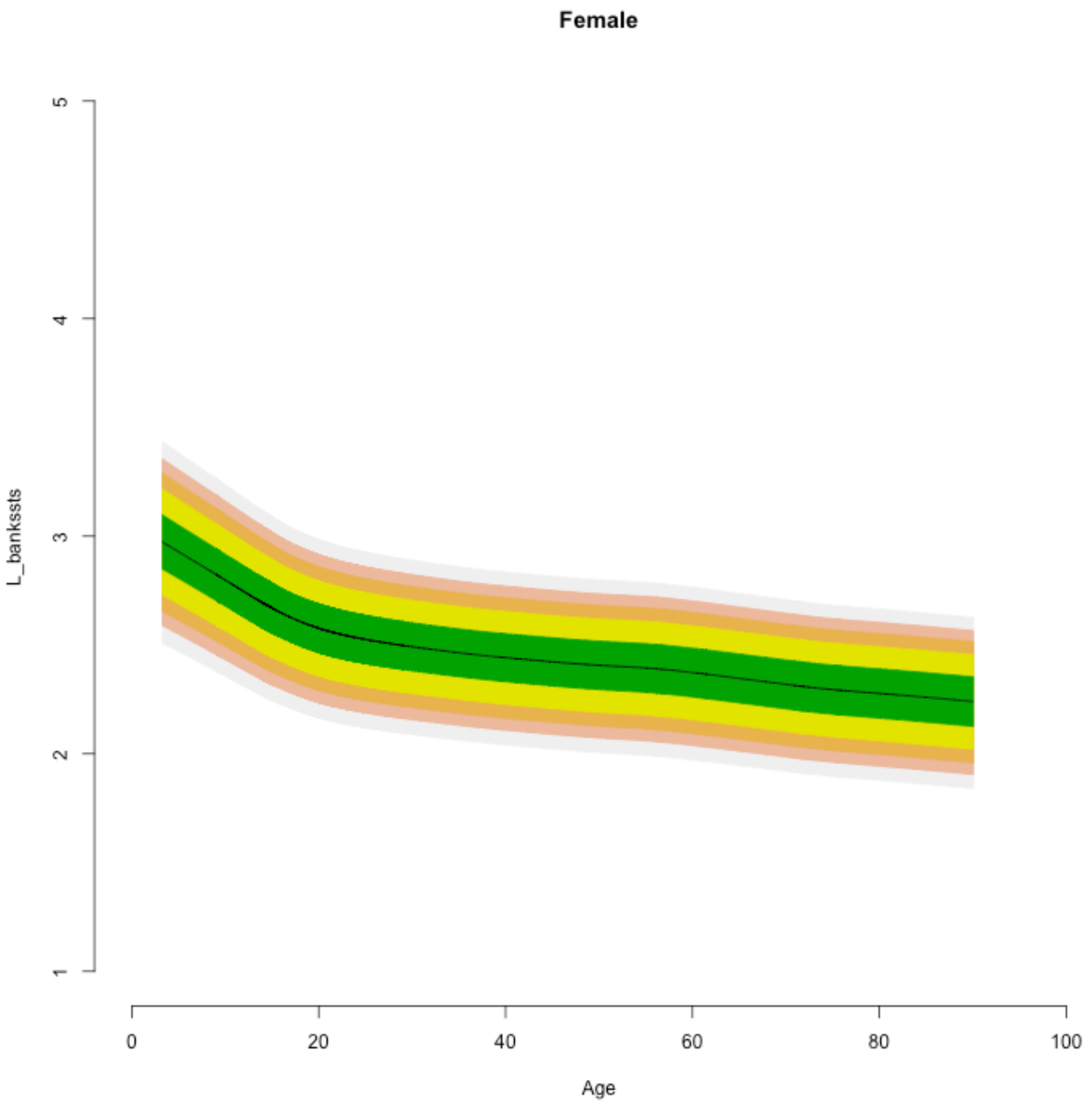
Female
L_bankssts
1
2
3
4
5
Age
0
20
40
60
80
100

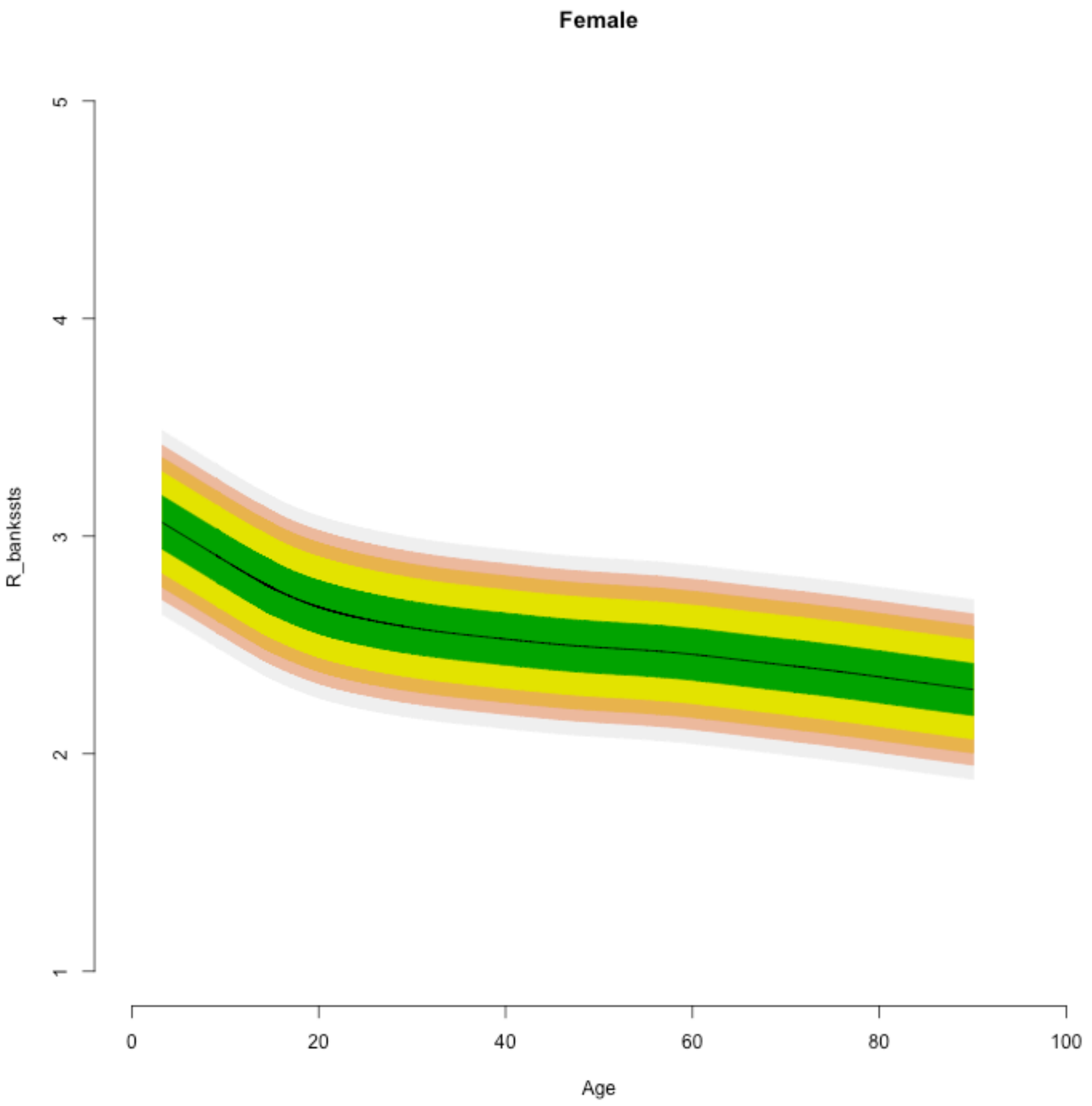

Female
R_banksts
1
2
3
4
5
0
20
40
60
80
100
Age

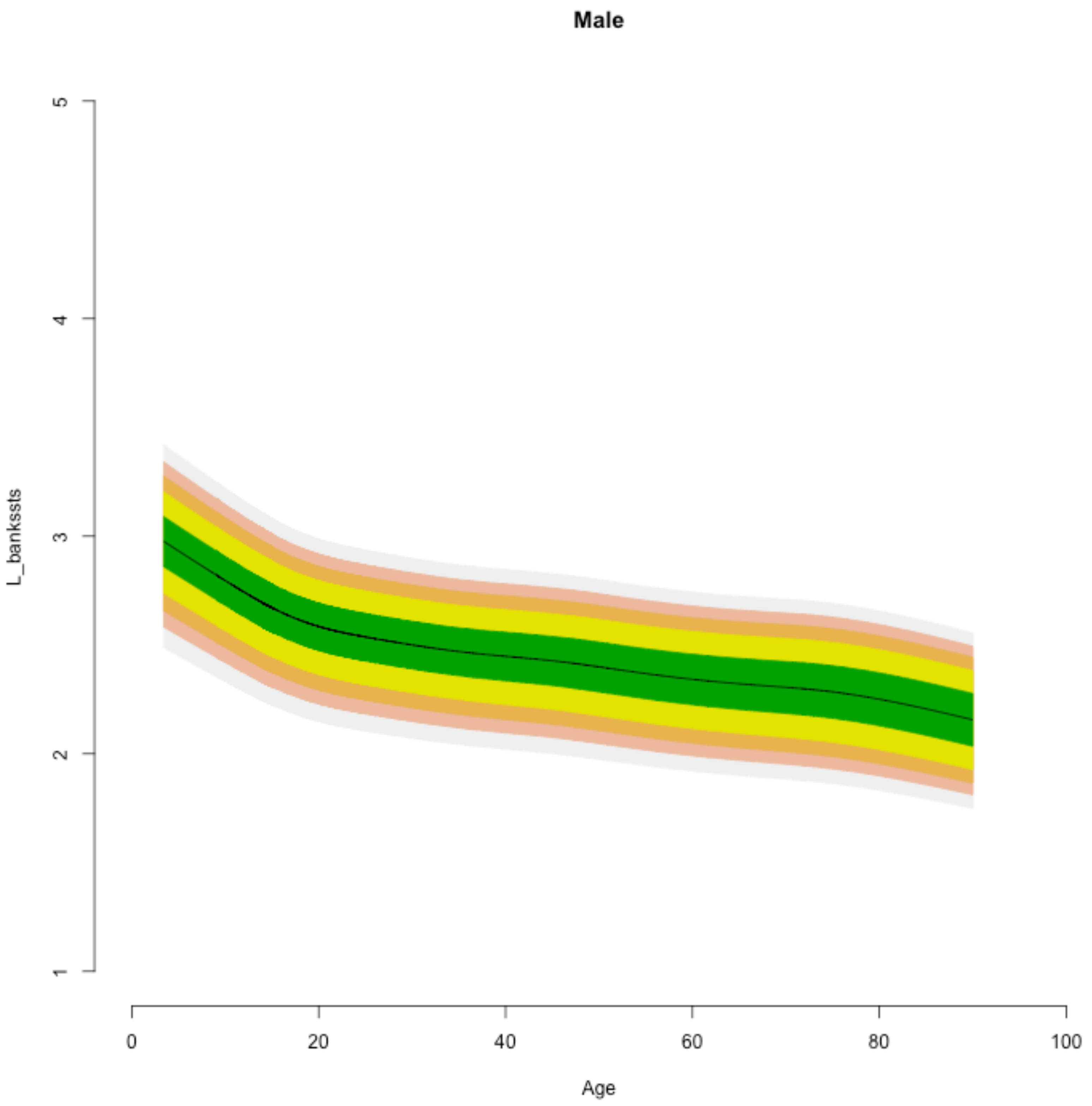

Male
L_bankssts
1
2
3
4
5
0
20
40
60
80
100
Age

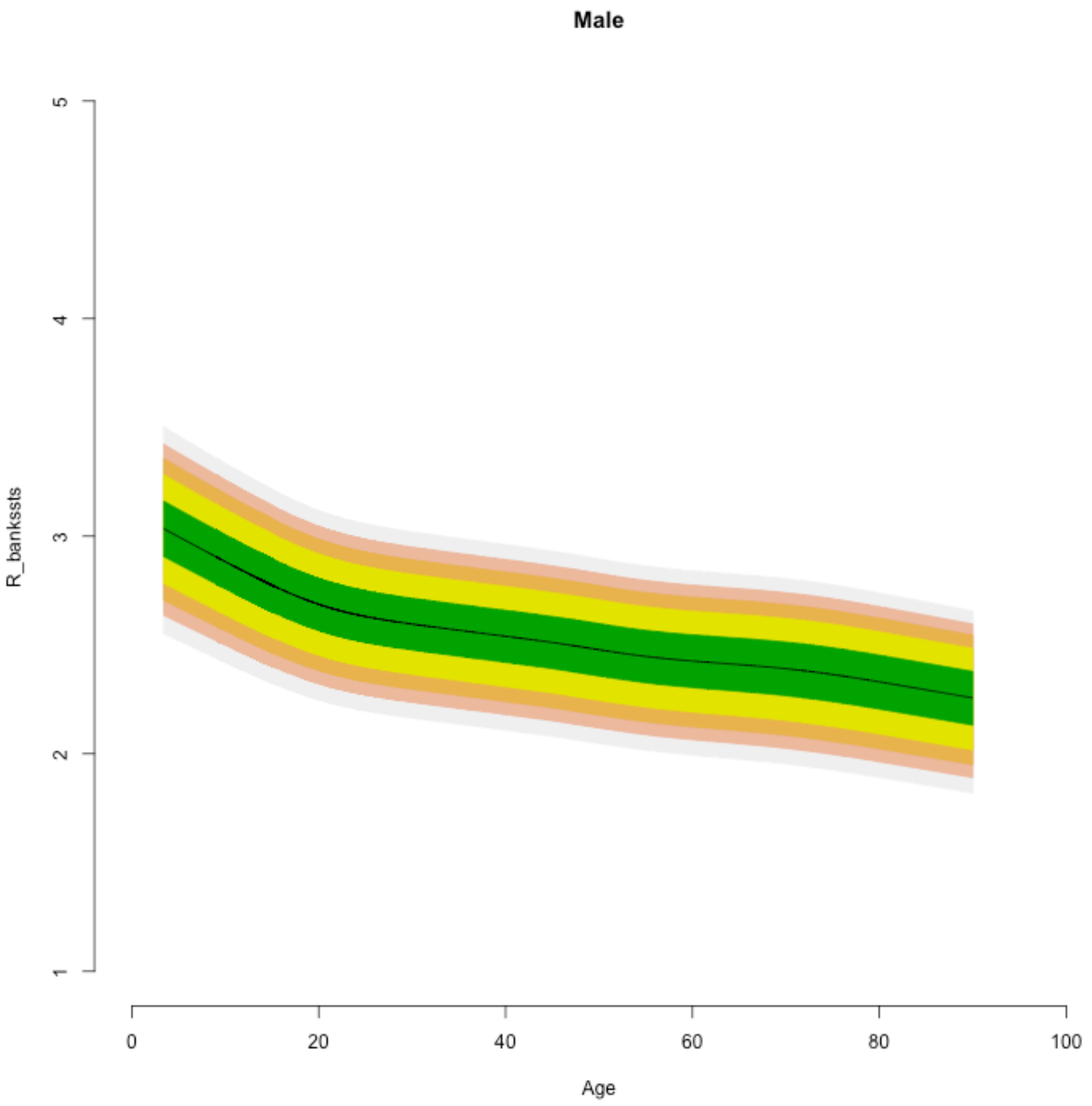
Male
R_bankssts
5
4
3
2
1
0
20
40
60
80
100
Age

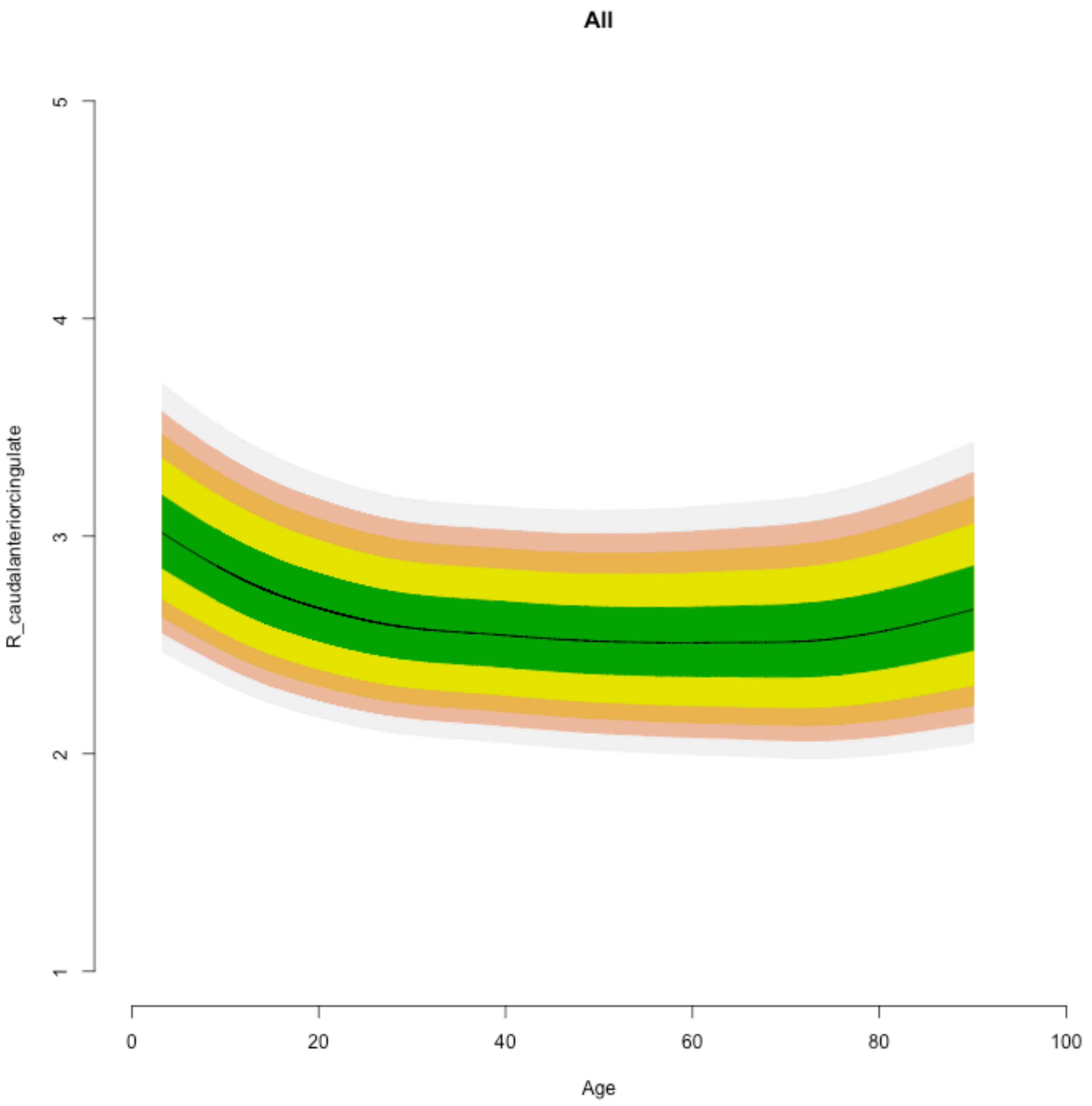
All
R_caudalanteriorcingulate
1
2
3
4
5
0
20
40
60
80
100
Age

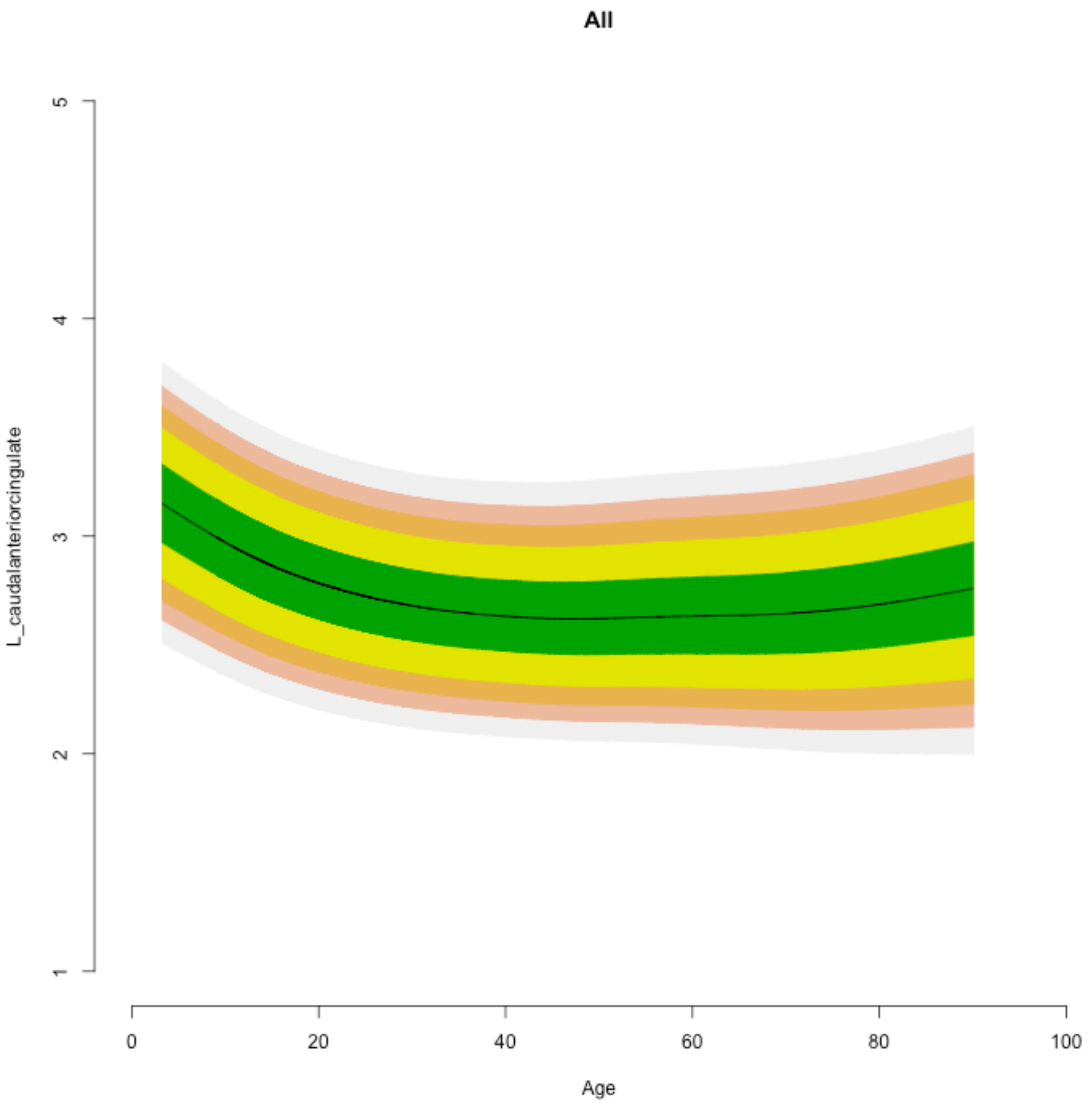
All
L_caudalanteriorcingulate
5
4
3
2
1
0
20
40
60
80
100
Age

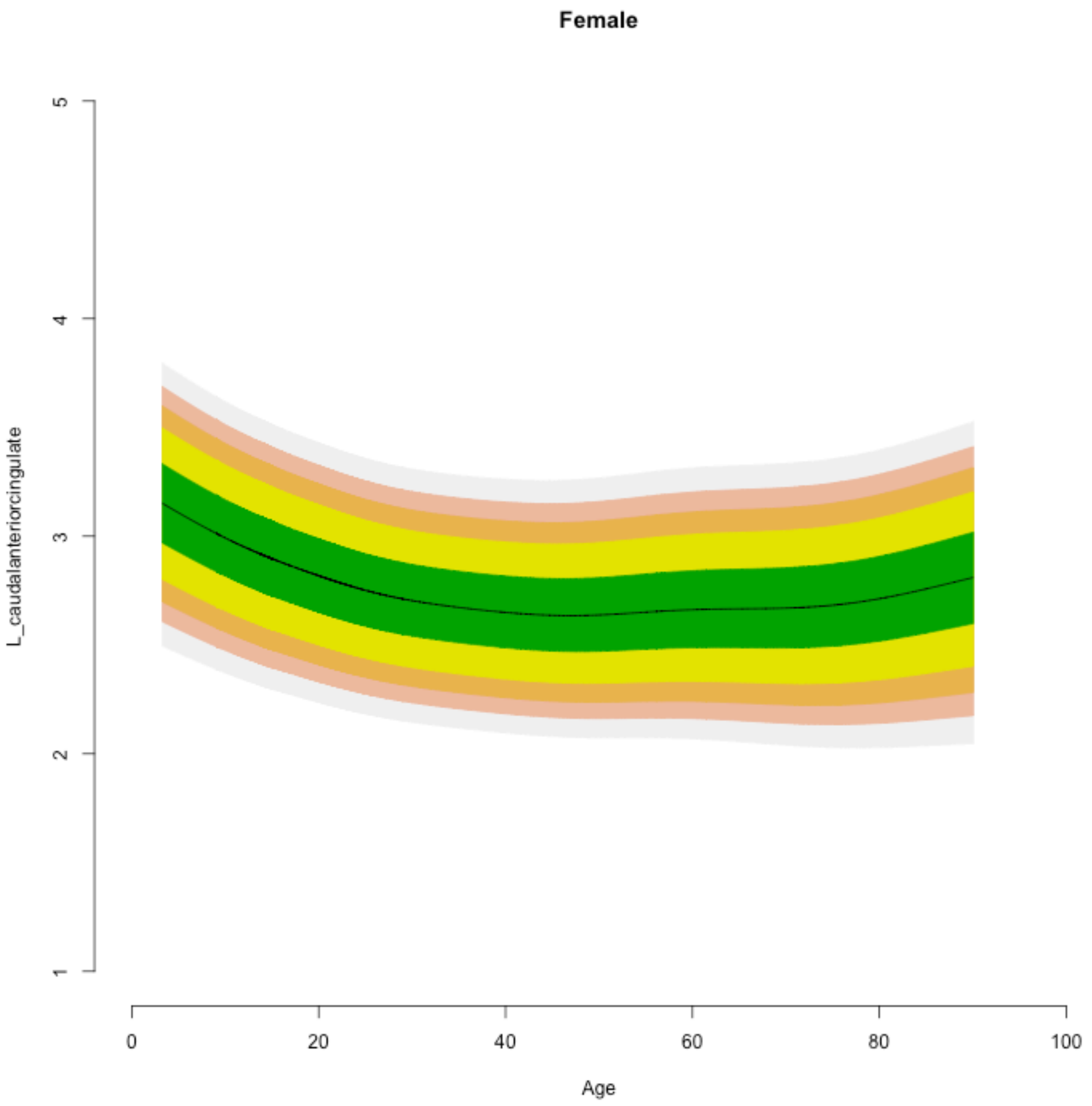

Female
L_caudalanteriorcingulate
5
4
3
2
1
0
20
40
60
80
100
Age

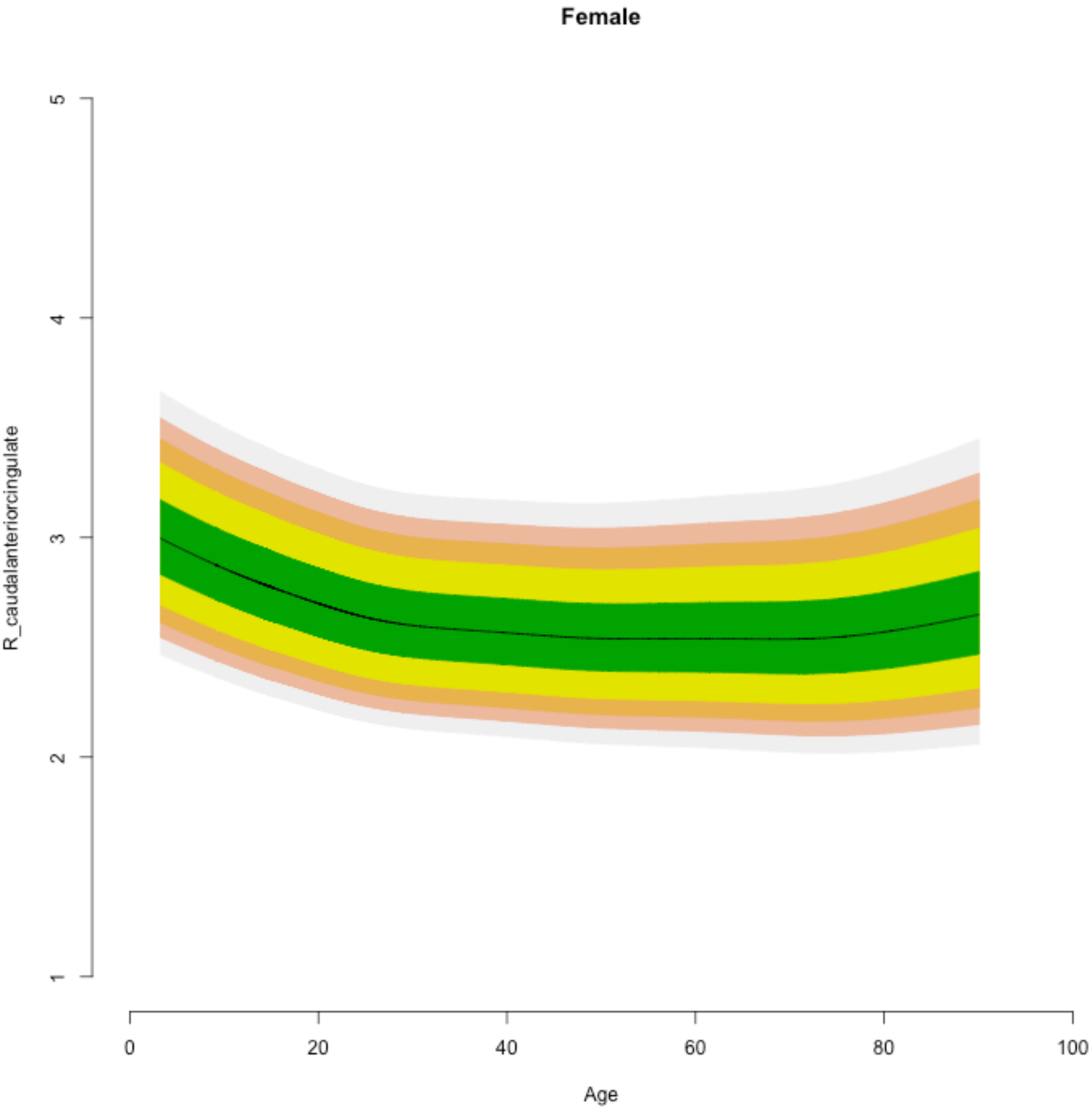
Female
R_caudalanteriorcingulate
Age
1
2
3
4
5
0
20
40
60
80
100

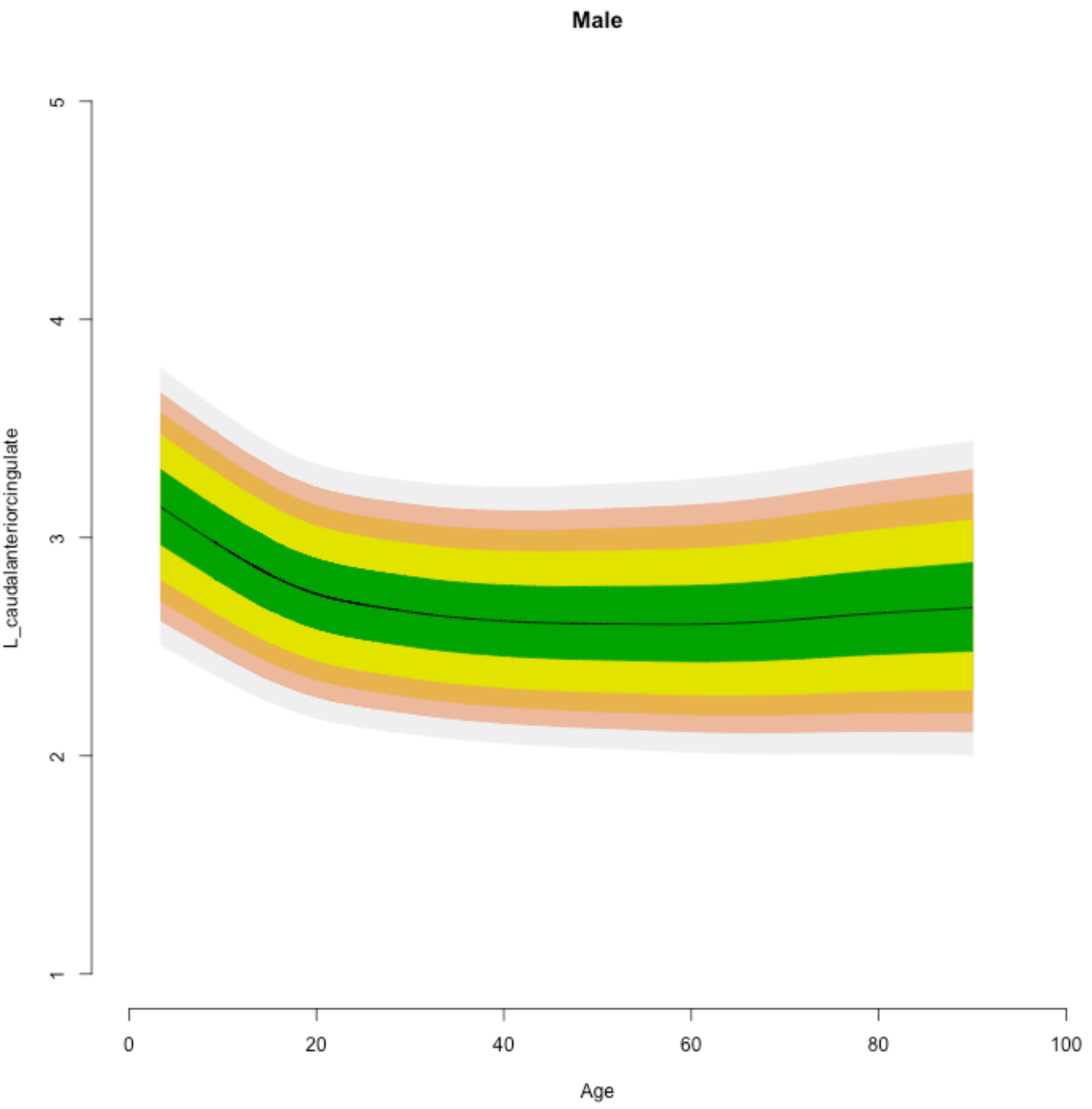
Male
L_caudalanteriorcingulate
1
2
3
4
5
0
20
40
60
80
100
Age

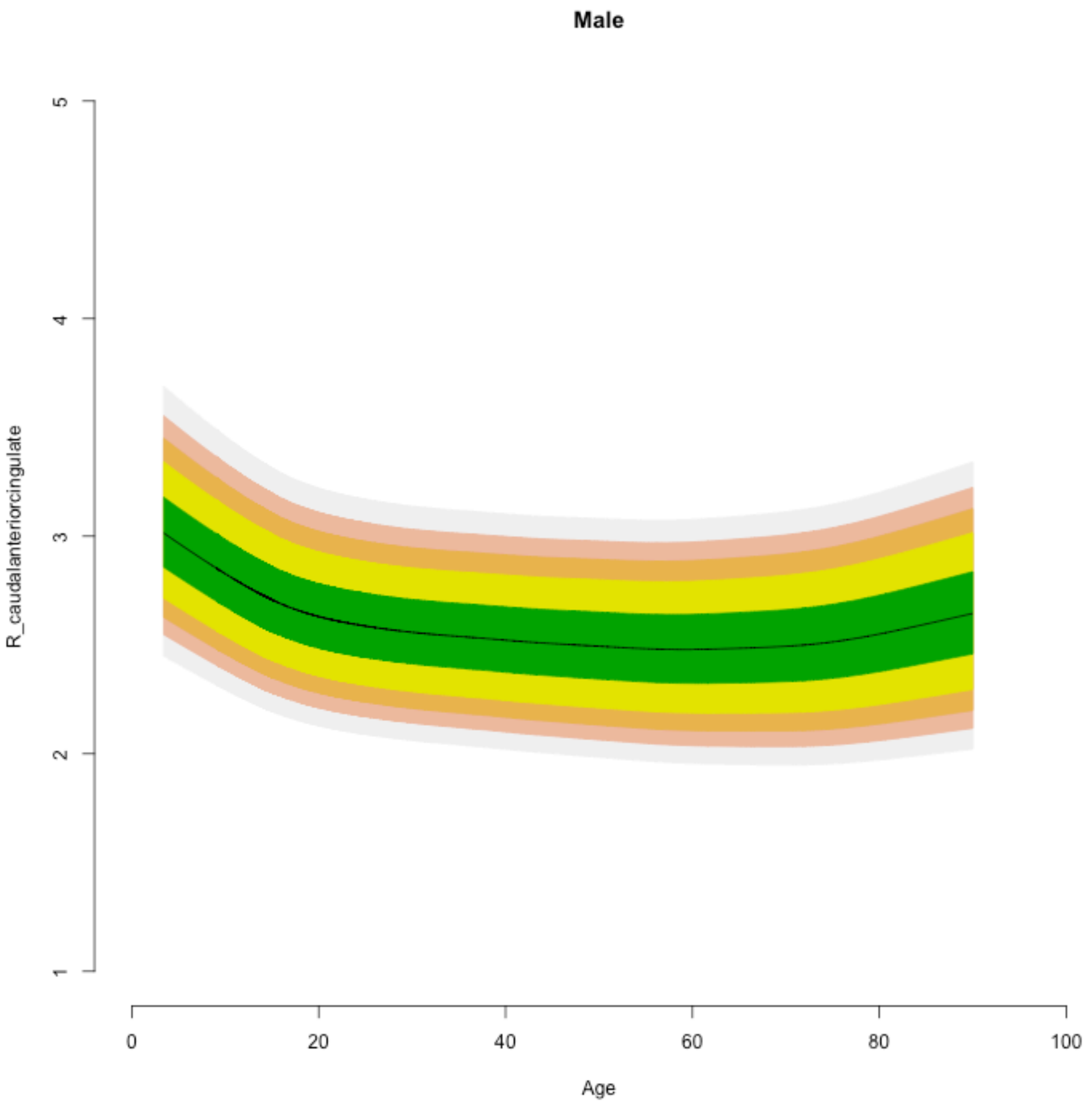

Male
R_caudalanteriorcingulate
5
4
3
2
1
0
20
40
60
80
100
Age

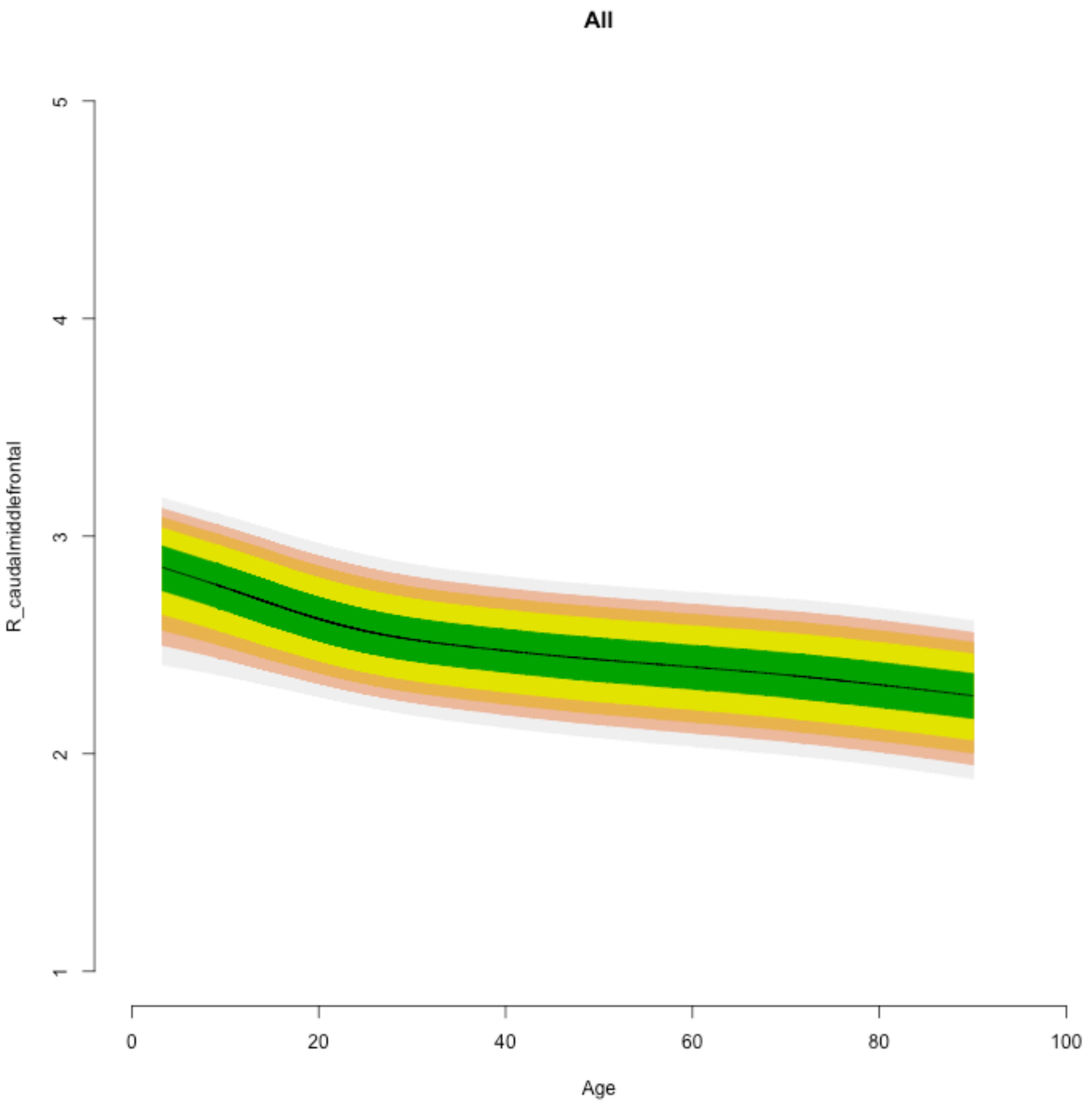
All
R_caudalmiddlefrontal
1
2
3
4
5
0
20
40
60
80
100
Age

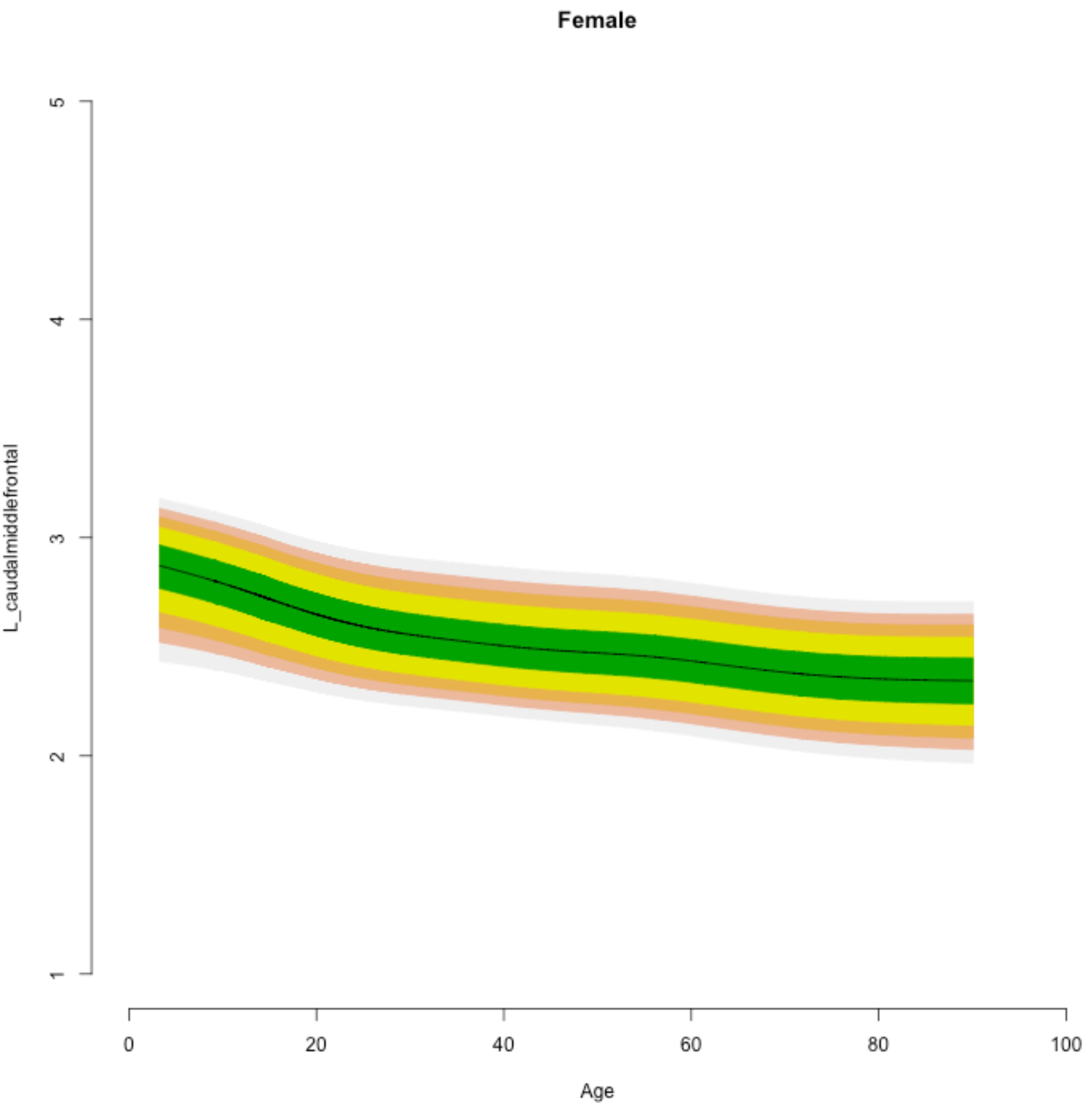
Female
L_caudalmiddlefrontal
1
2
3
4
5
0
20
40
60
80
100
Age

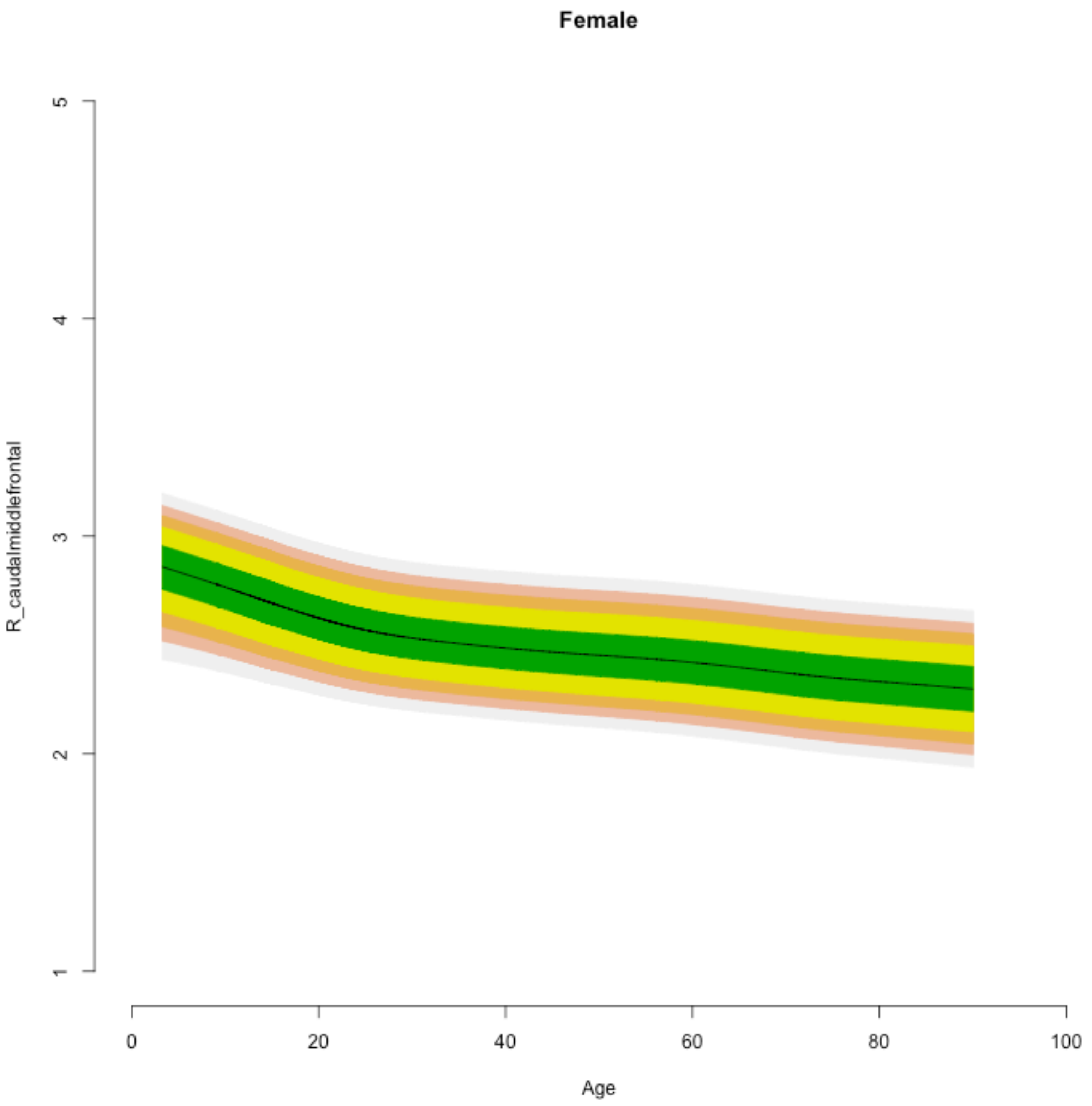
Female
R_caudalmiddlefrontal
1
2
3
4
5
0
20
40
60
80
100
Age

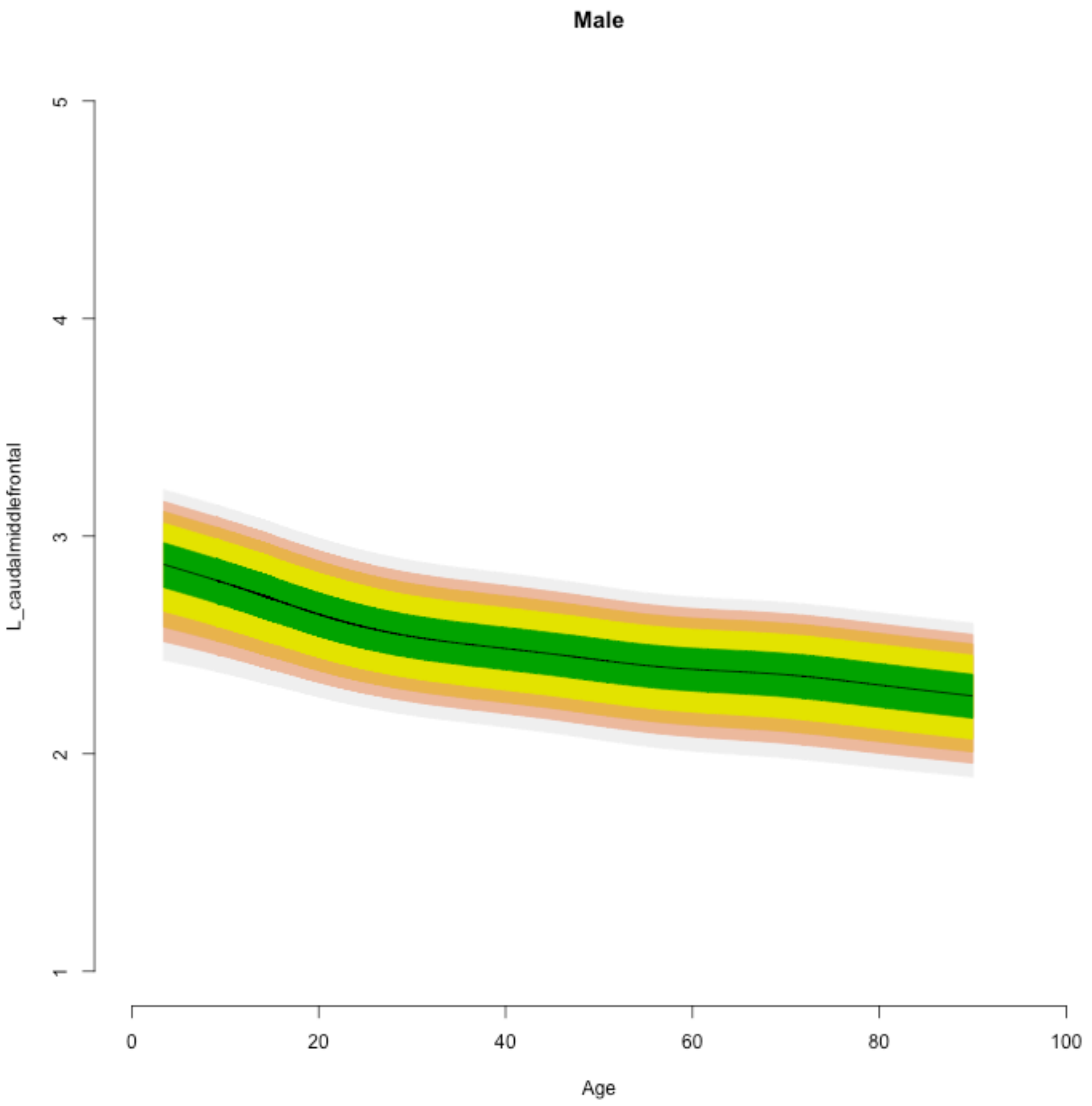

Male
L_caudalmiddlefrontal
5
4
3
2
1
0
20
40
60
80
100
Age

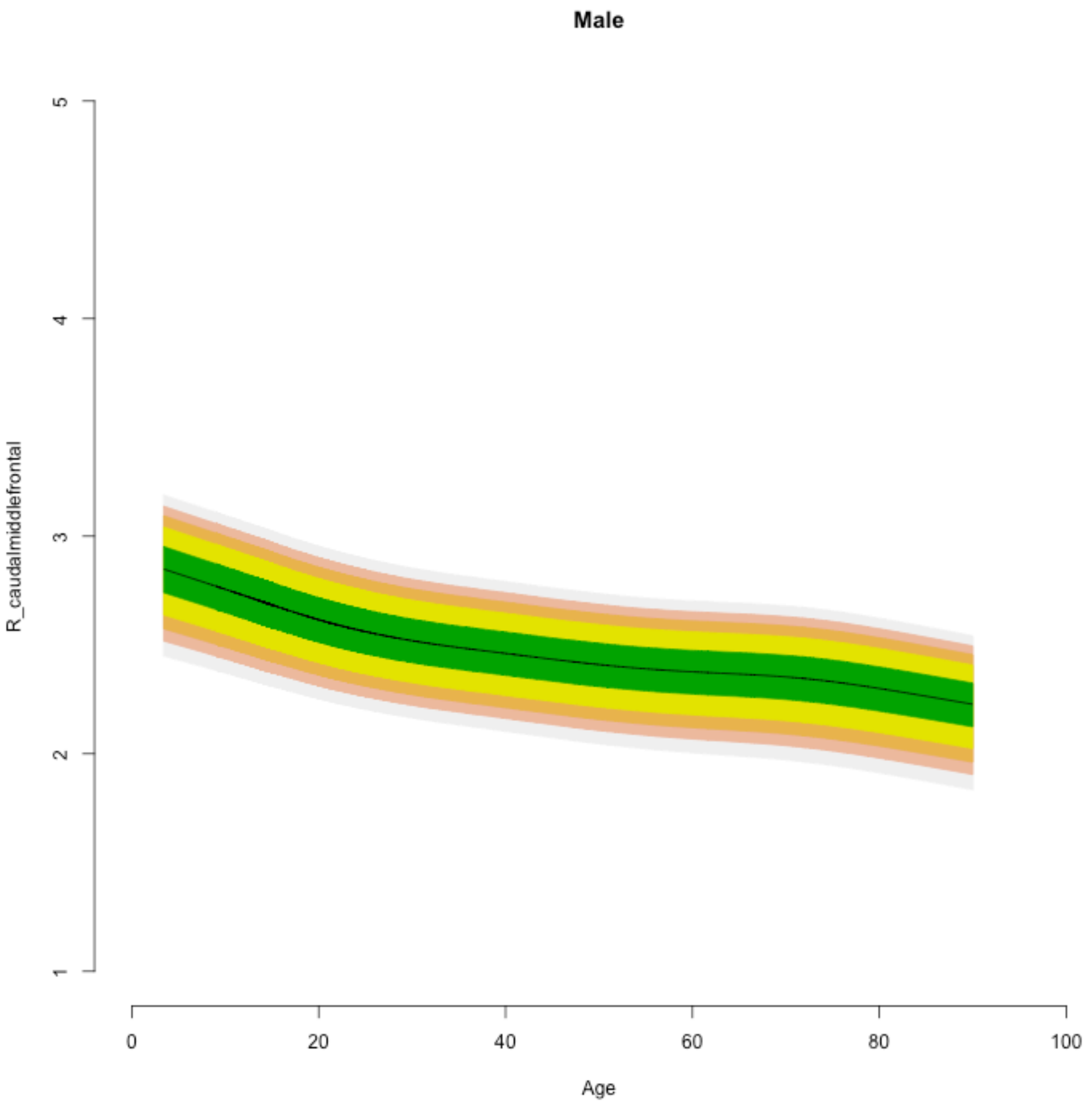

Male
R_caudalmiddlefrontal
1
2
3
4
5
0
20
40
60
80
100
Age

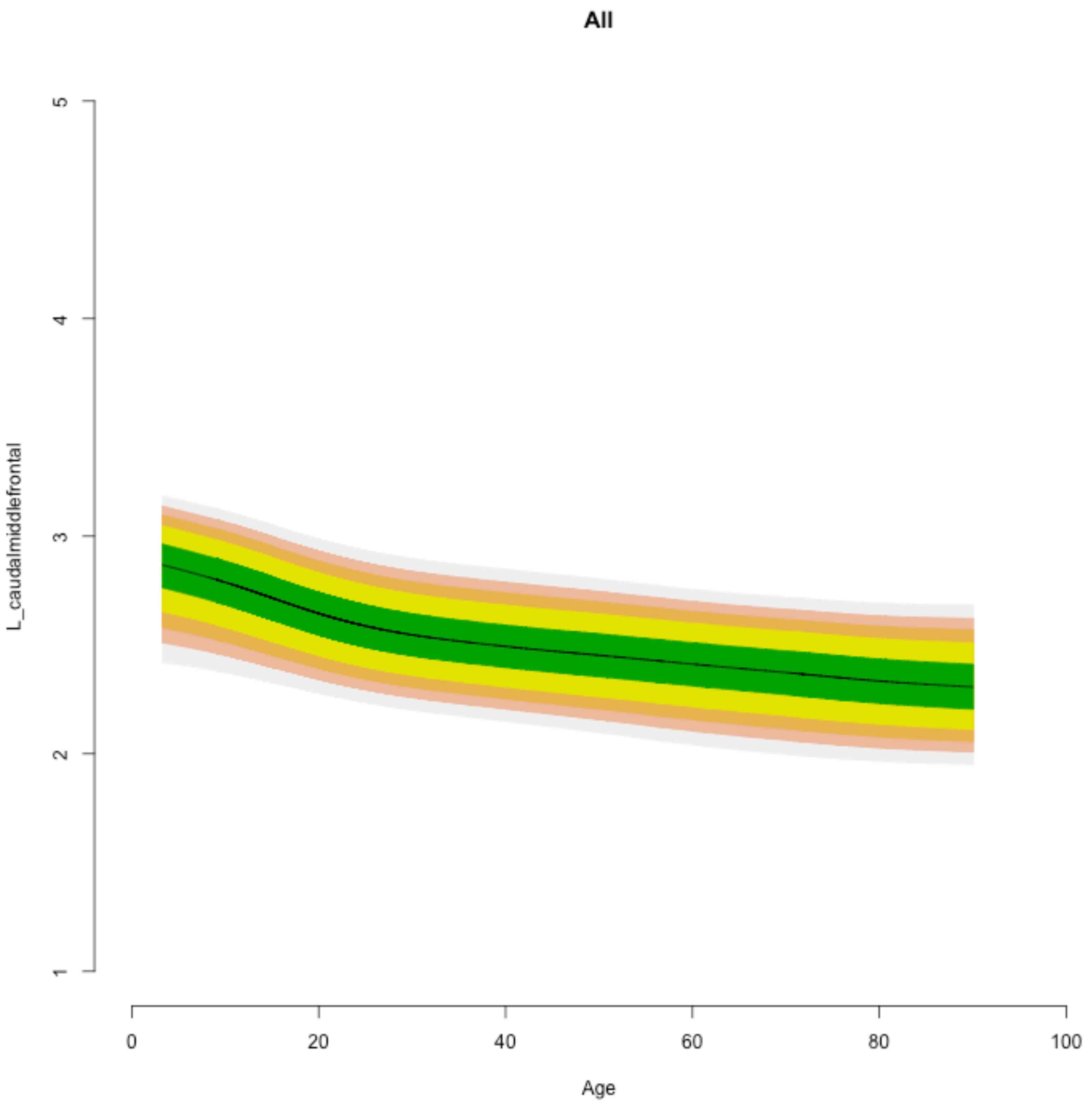

All
L_caudalmiddlefrontal
1
2
3
4
5
0
20
40
60
80
100
Age

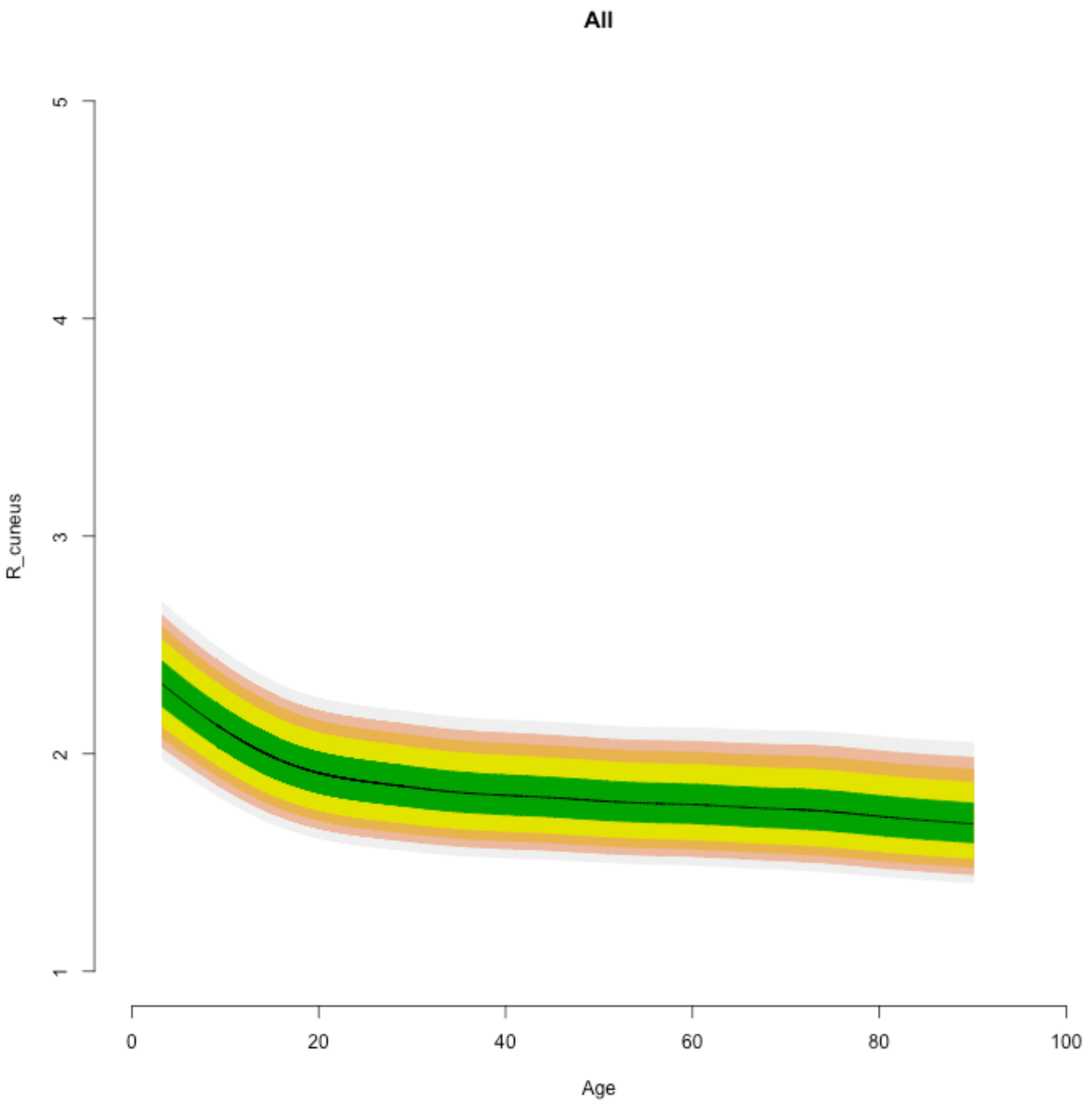
All
R_cuneus
1
2
3
4
5
0
20
40
60
80
100
Age

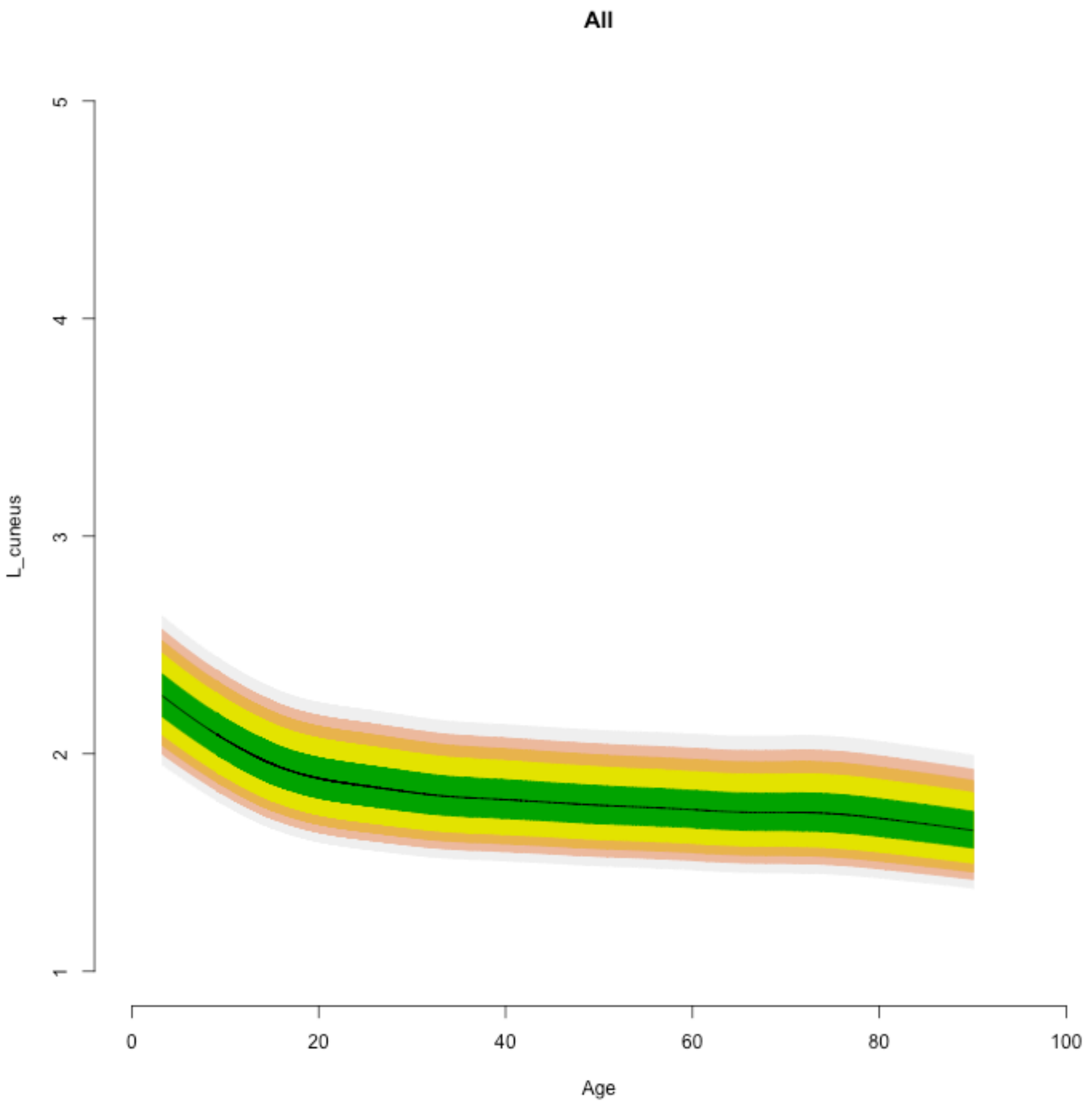
All
L_cuneus
1
2
3
4
5
0
20
40
60
80
100
Age

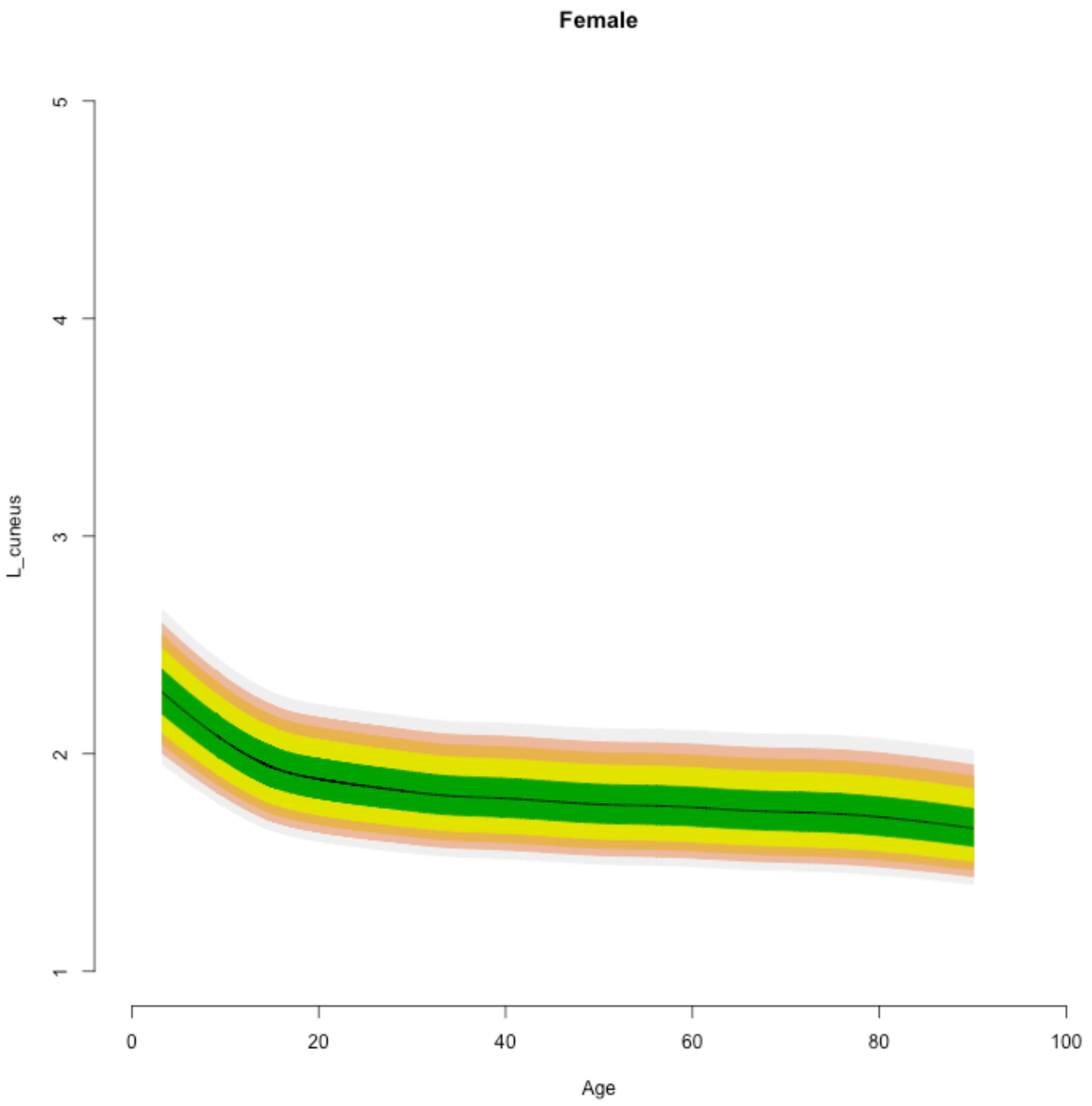

Female
L_cuneus
1
2
3
4
5
0
20
40
60
80
100
Age

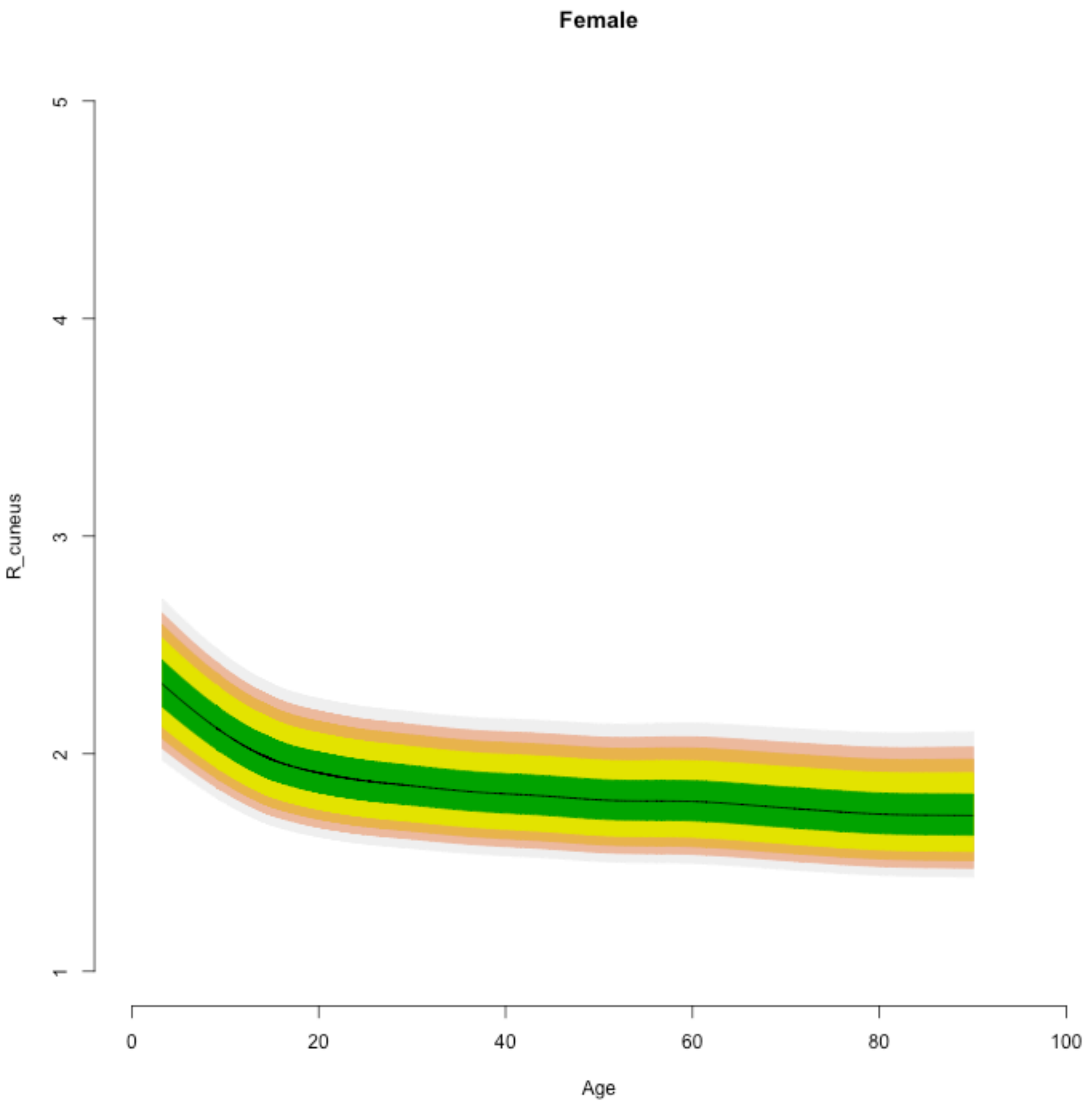

Female
R_cuneus
1
2
3
4
5
0
20
40
60
80
100
Age

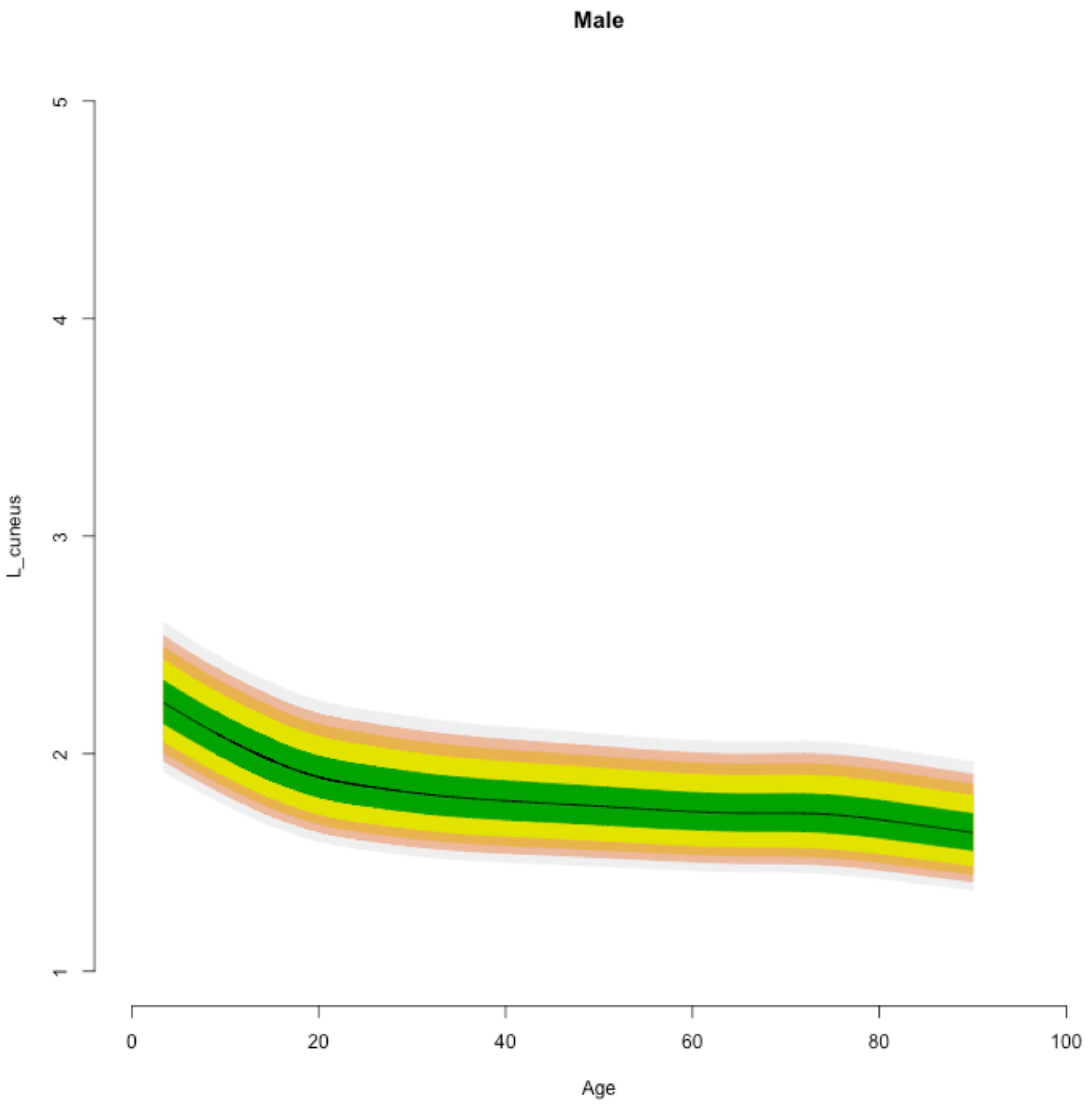
Male
L_cuneus
1
2
3
4
5
0
20
40
60
80
100
Age

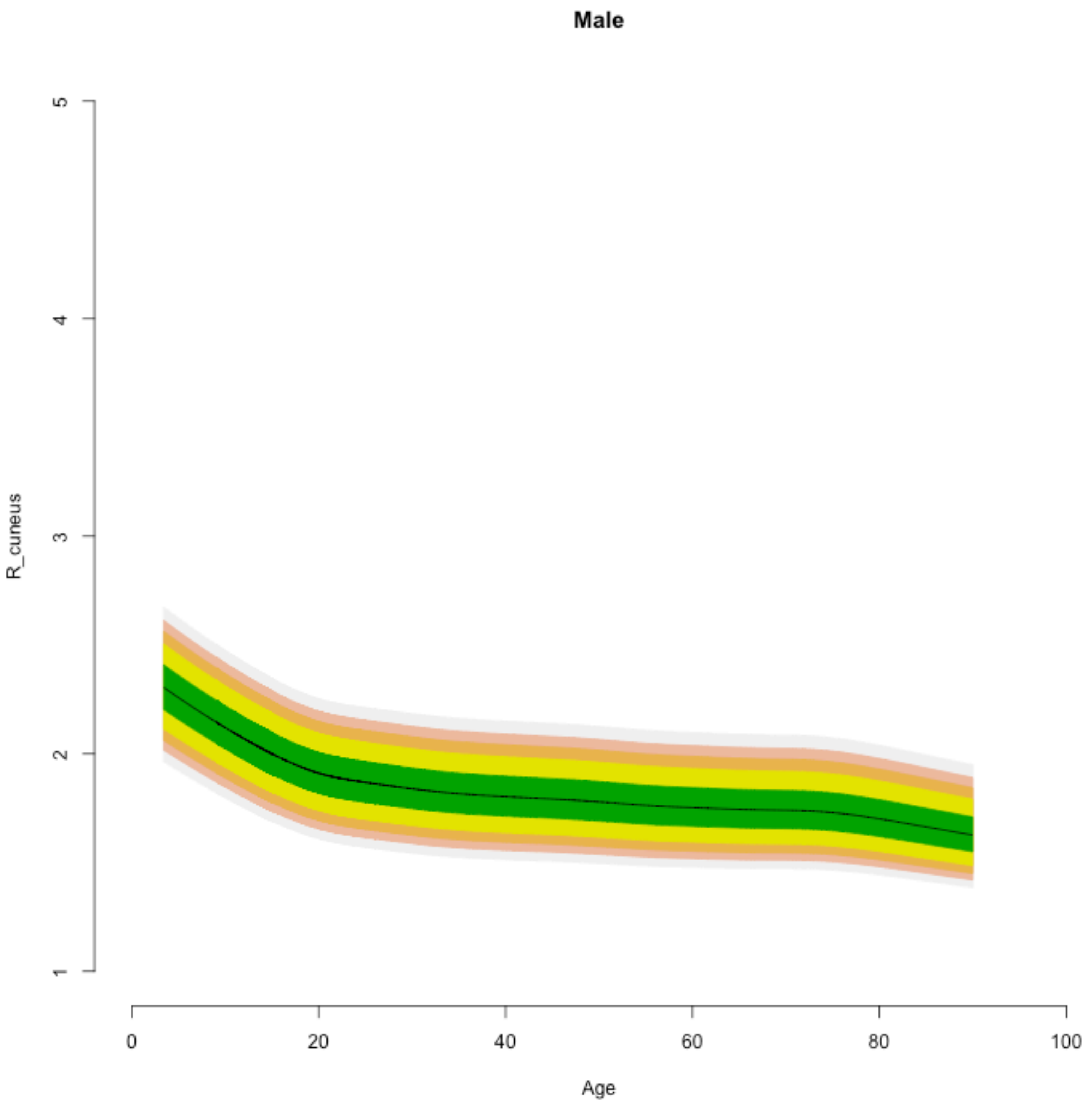

Male
R_cuneus
1
2
3
4
5
0
20
40
60
80
100
Age

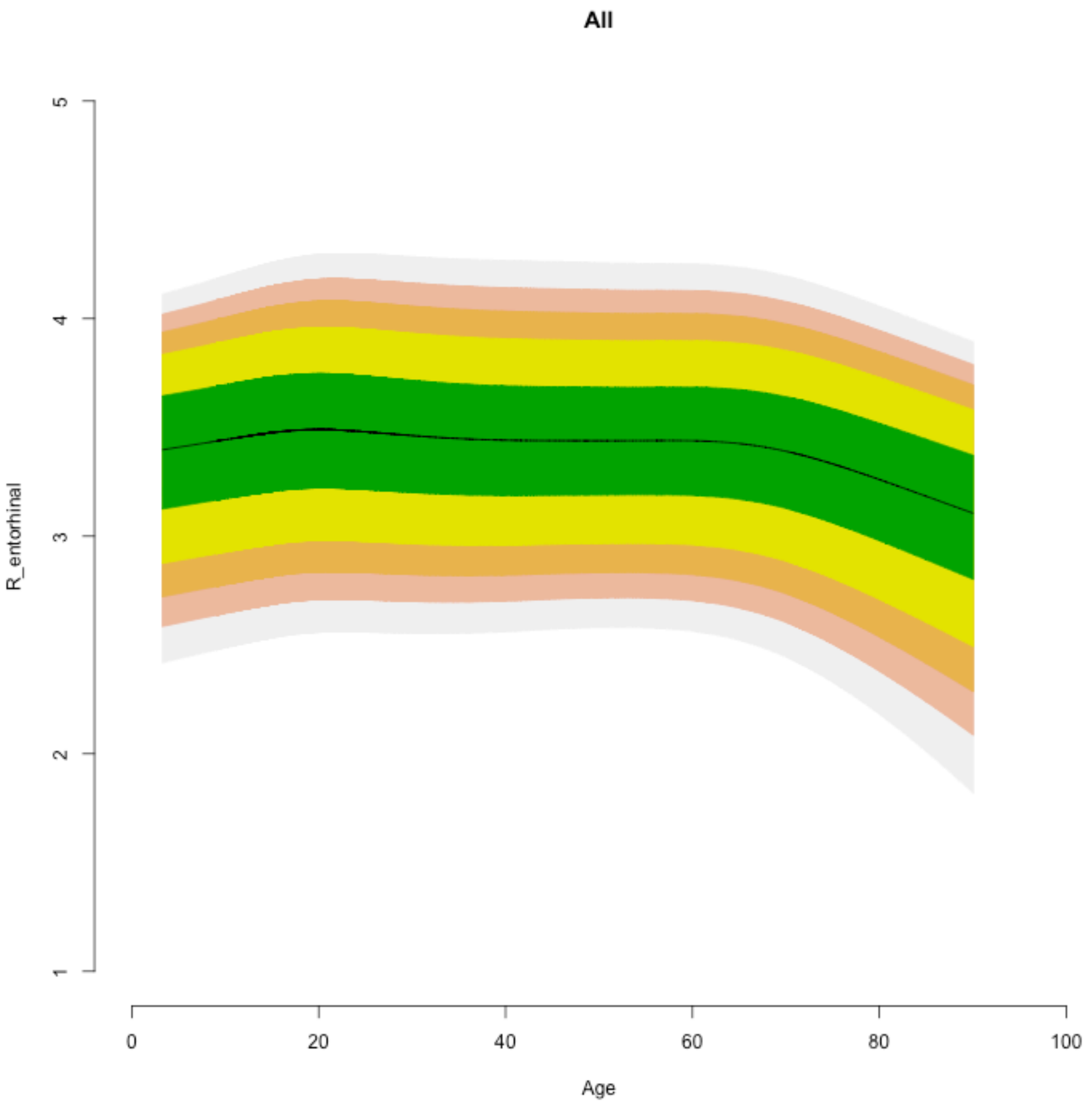

All
R_entorhinal
1
2
3
4
5
0
20
40
60
80
100
Age

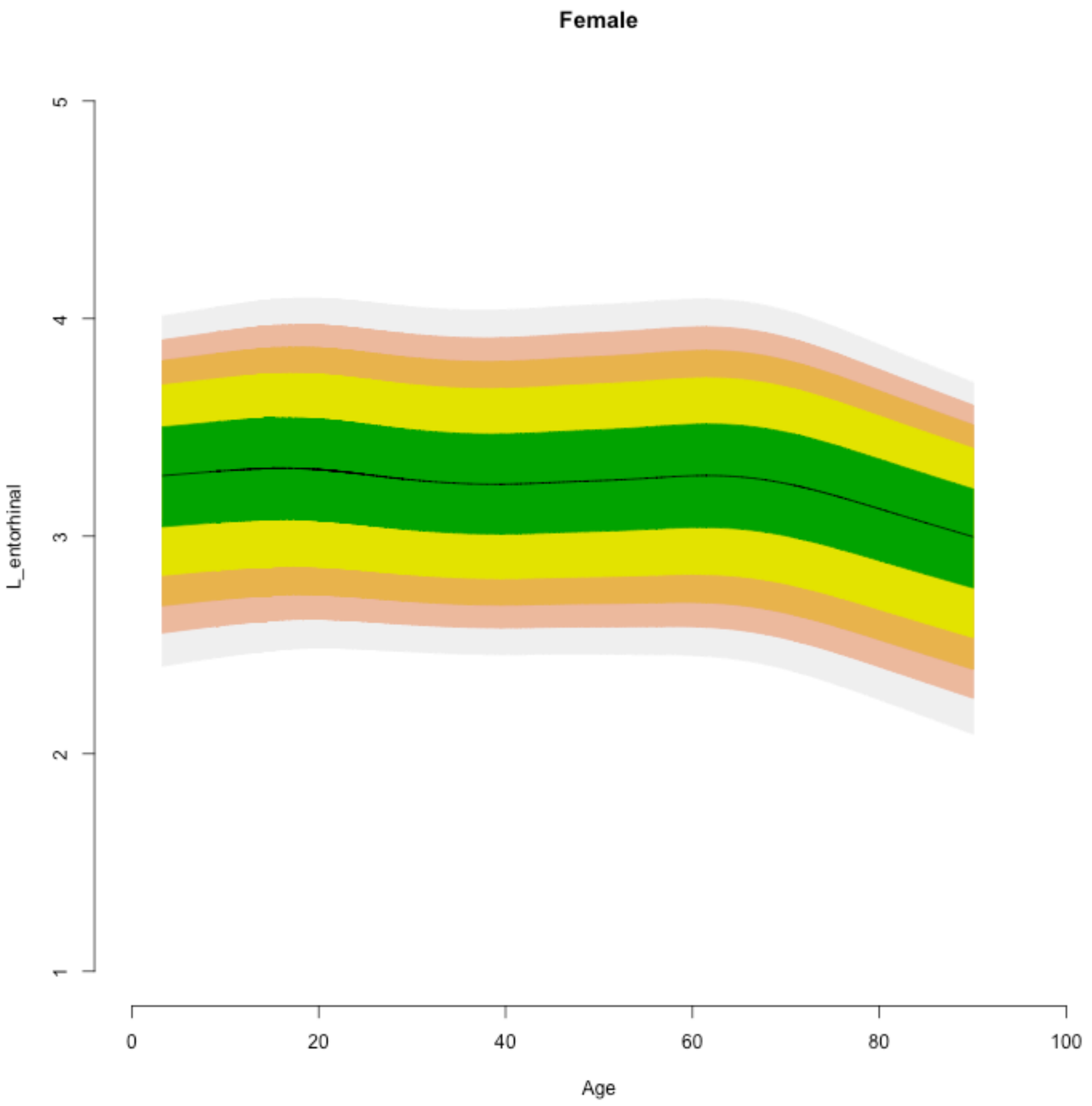

Female
L_entorhinal
1
2
3
4
5
0
20
40
60
80
100
Age

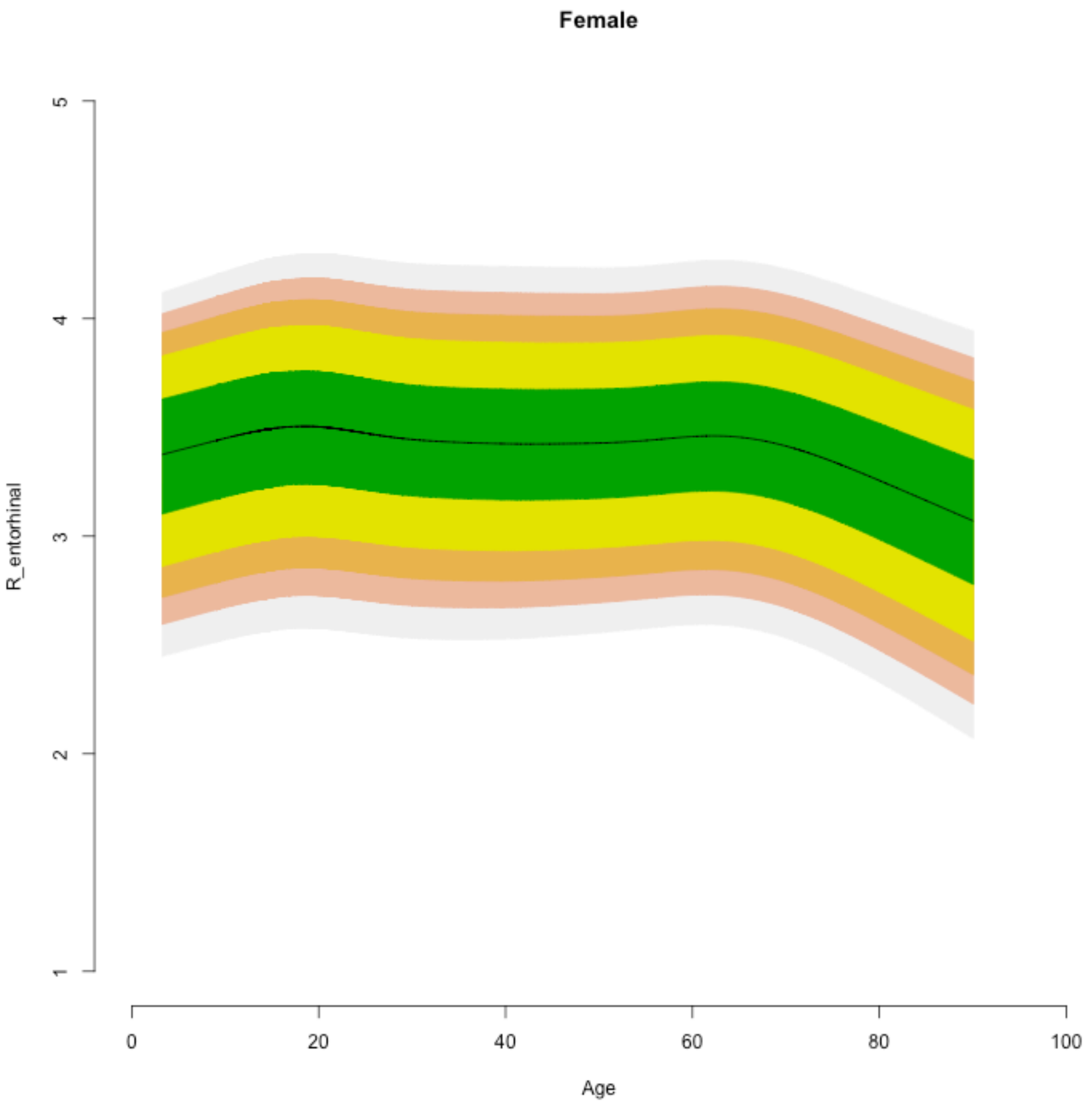
Female
R_entorhinal
1
2
3
4
5
0
20
40
60
80
100
Age

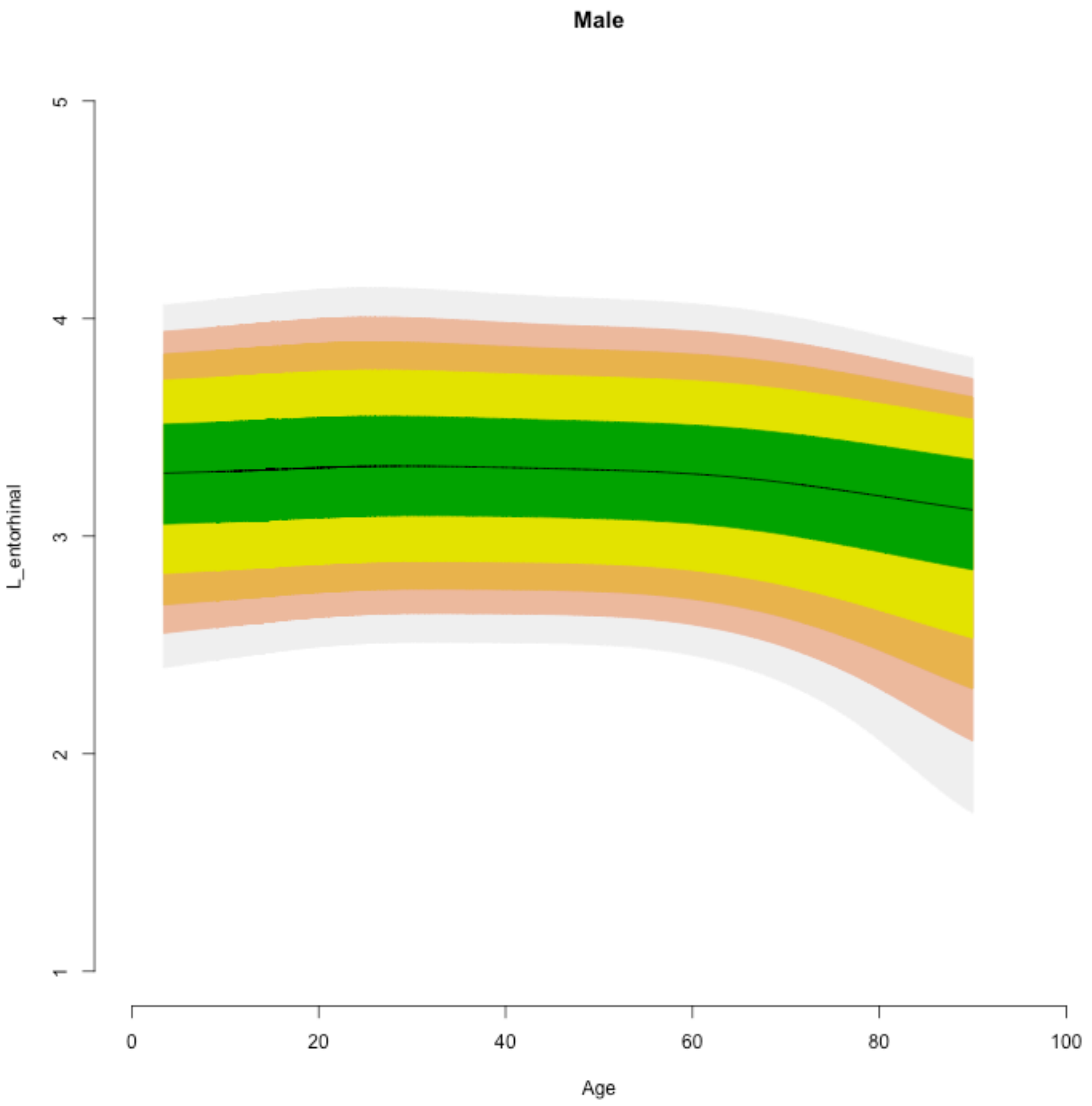

Male
L_entorhinal
1
2
3
4
5
0
20
40
60
80
100
Age

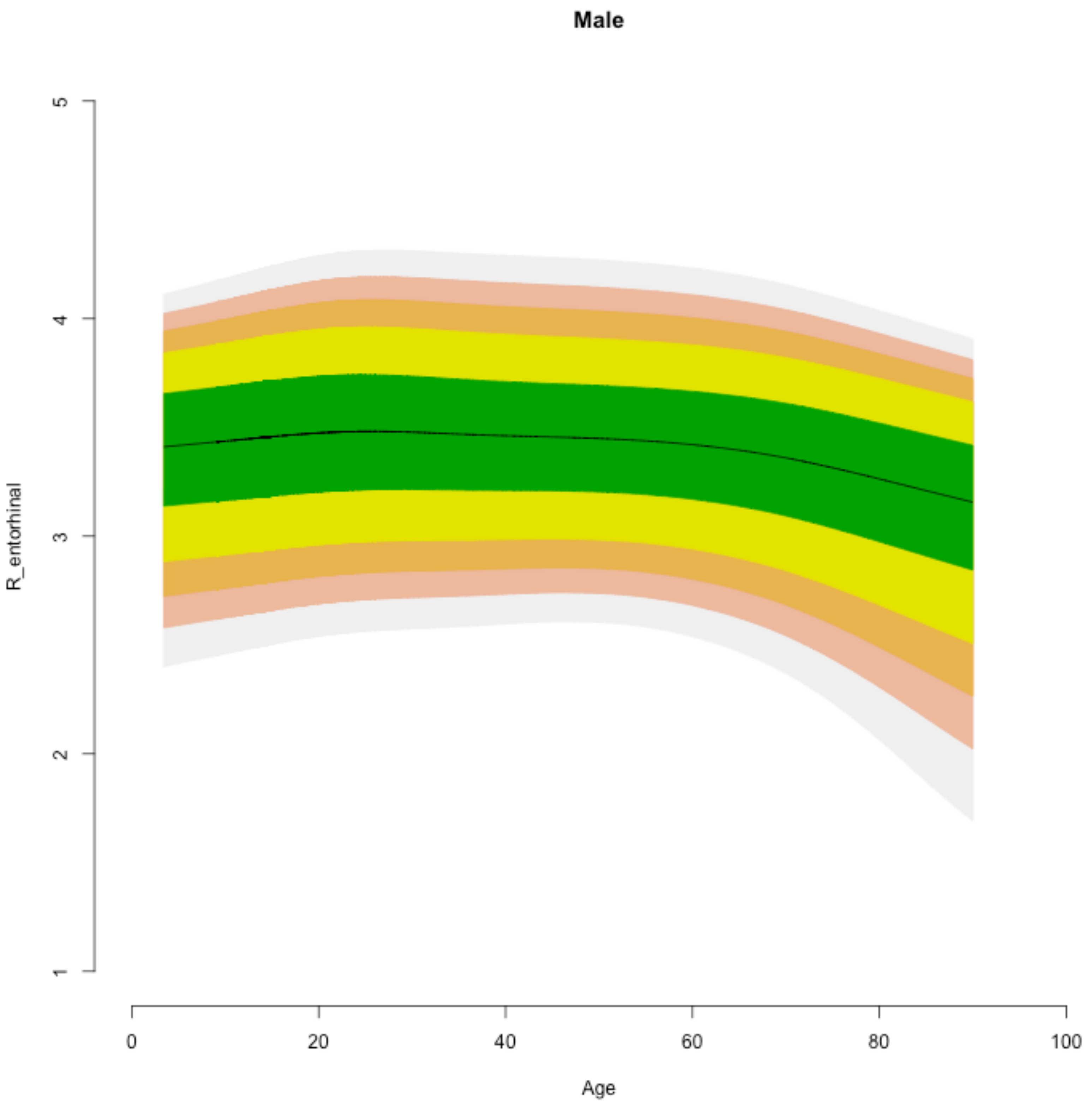

Male
R_entorhinal
1
2
3
4
5
0
20
40
60
80
100
Age

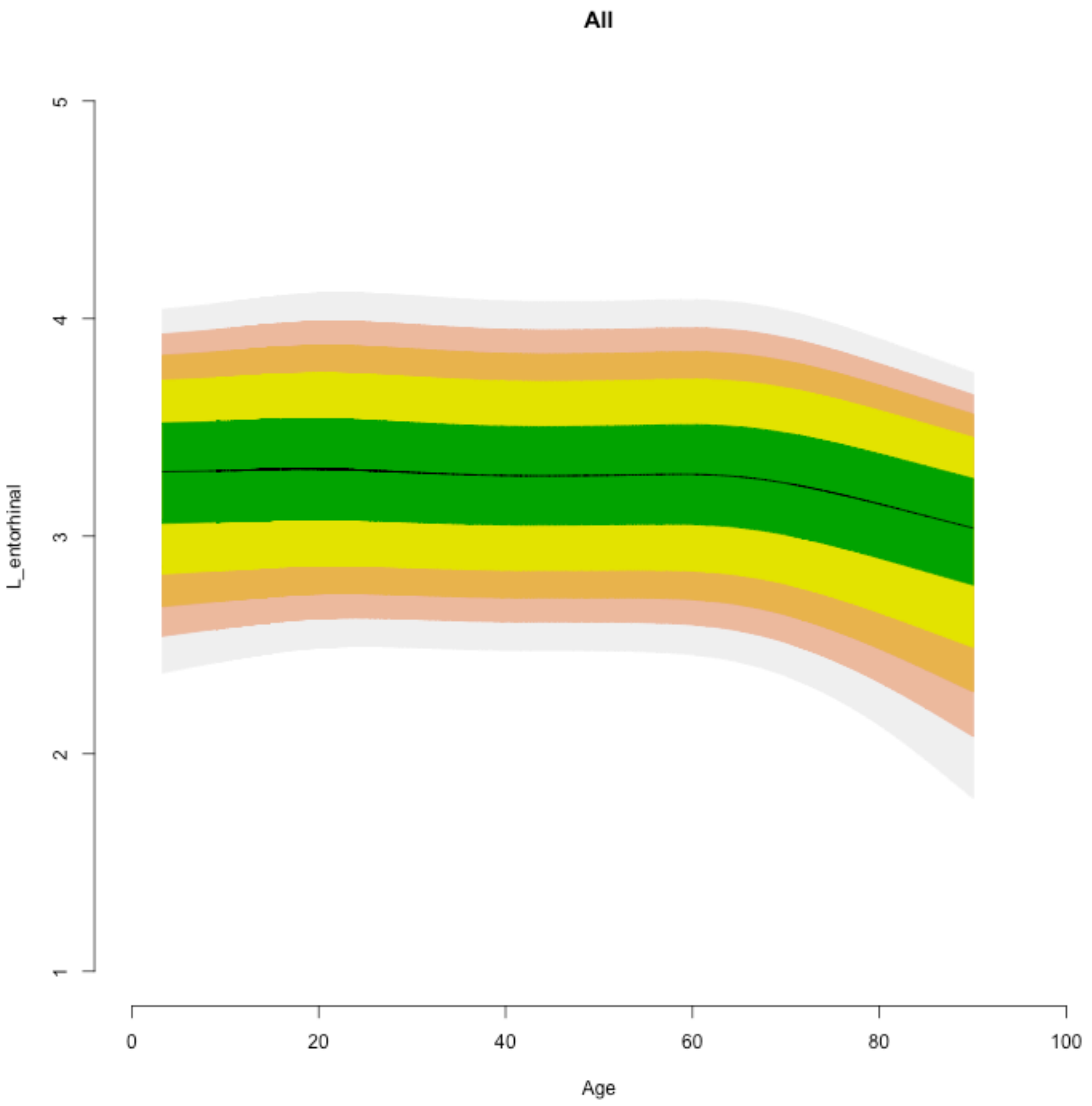

All
L_entorhinal
1
2
3
4
5
0
20
40
60
80
100
Age

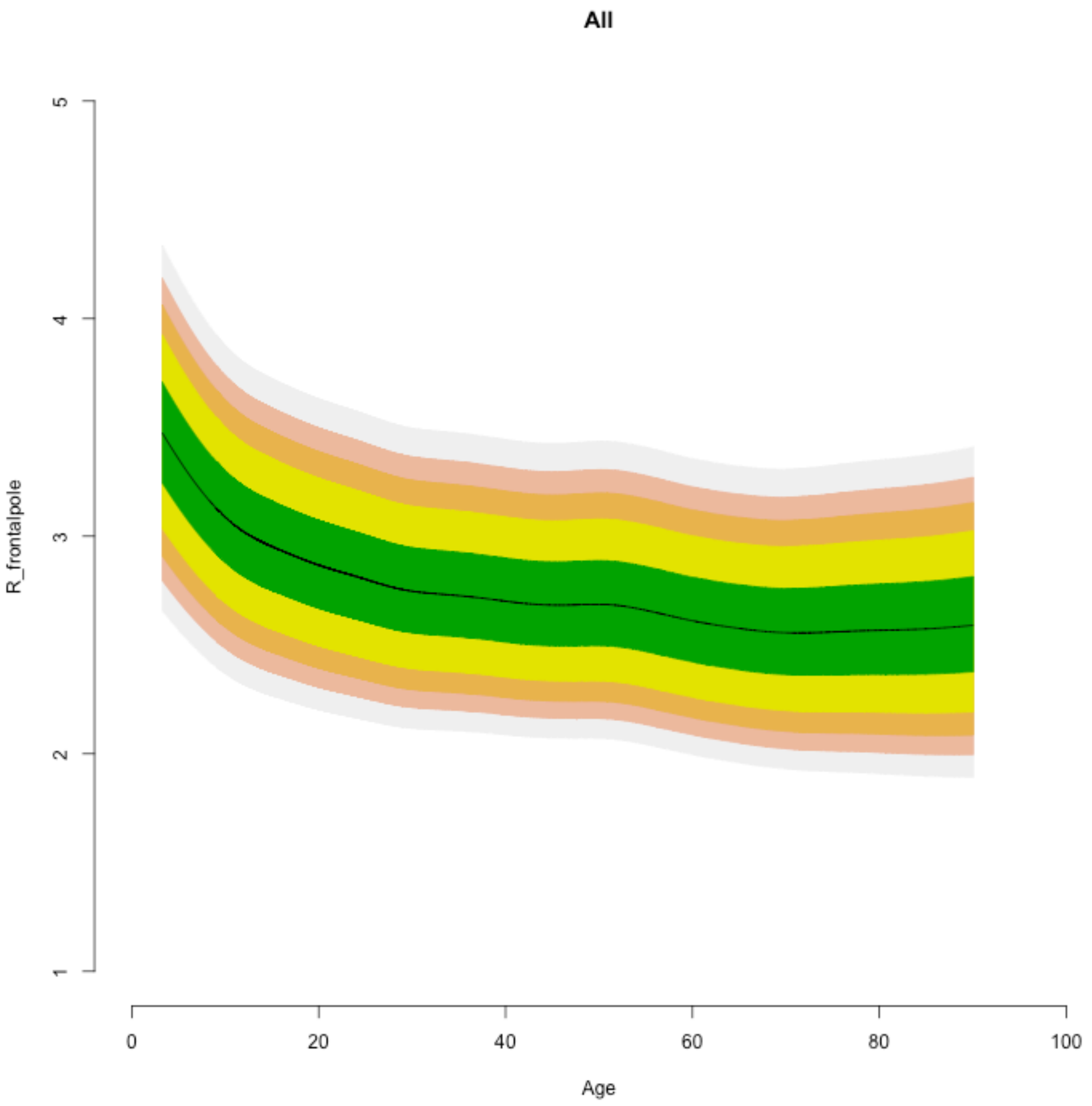

All
R_frontalpole
1
2
3
4
5
0
20
40
60
80
100
Age

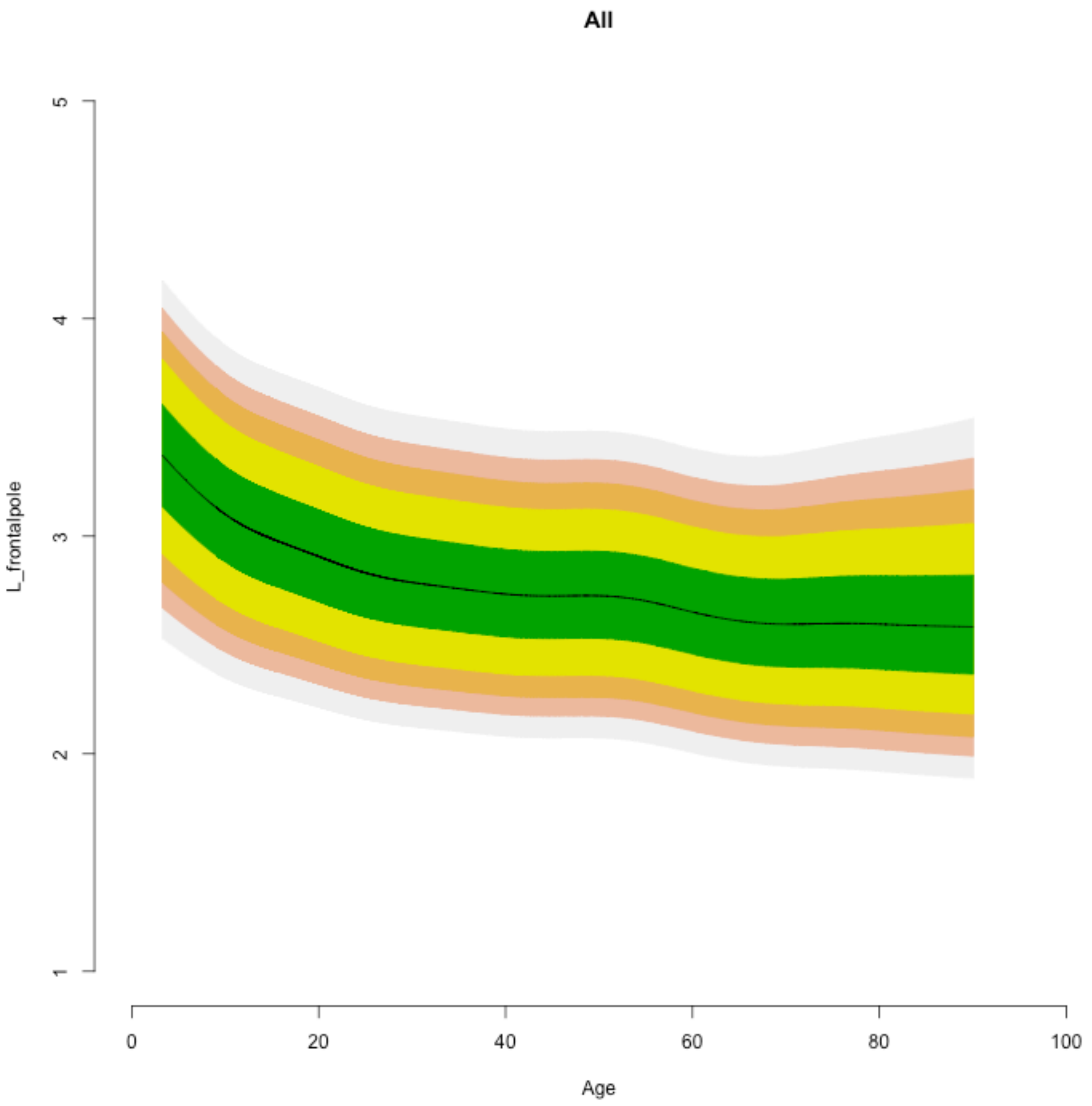

All
L_frontalpole
1
2
3
4
5
0
20
40
60
80
100
Age

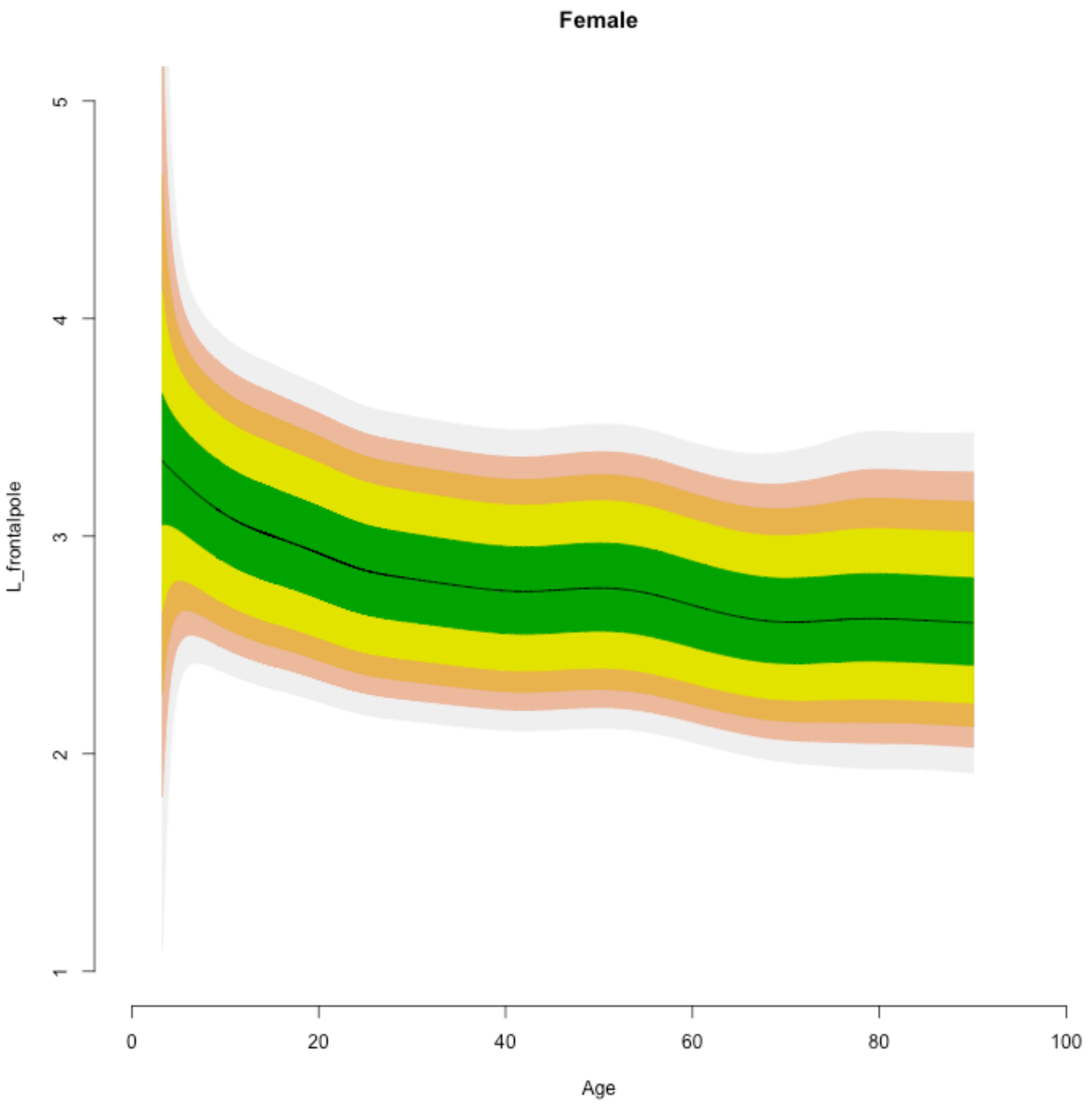

Female
L_frontalpole
1
2
3
4
5
0
20
40
60
80
100
Age

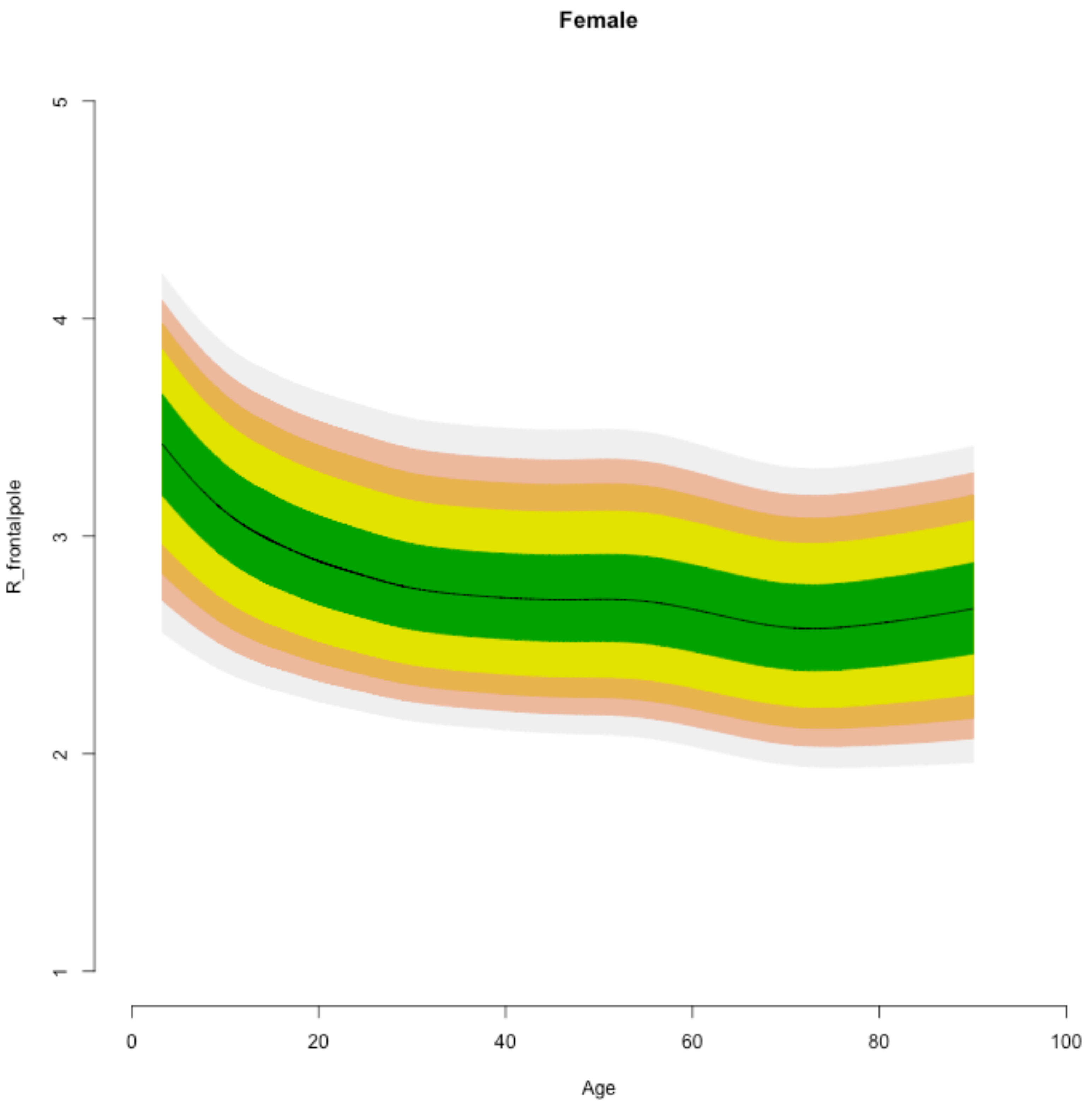

Female
R_frontalpole
1
2
3
4
5
0
20
40
60
80
100
Age

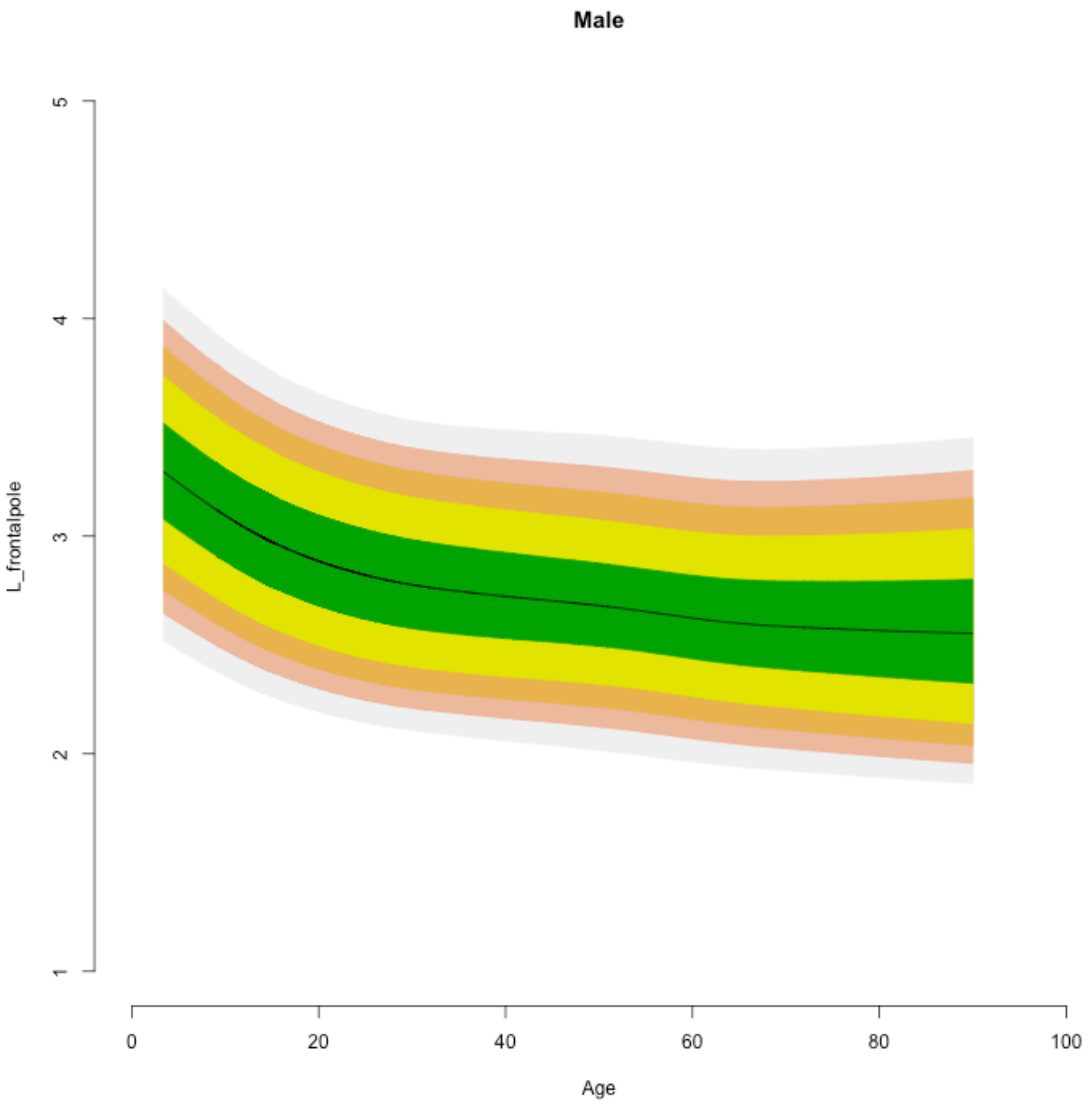
Male
L_frontalpole
Age
1
2
3
4
5
0
20
40
60
80
100

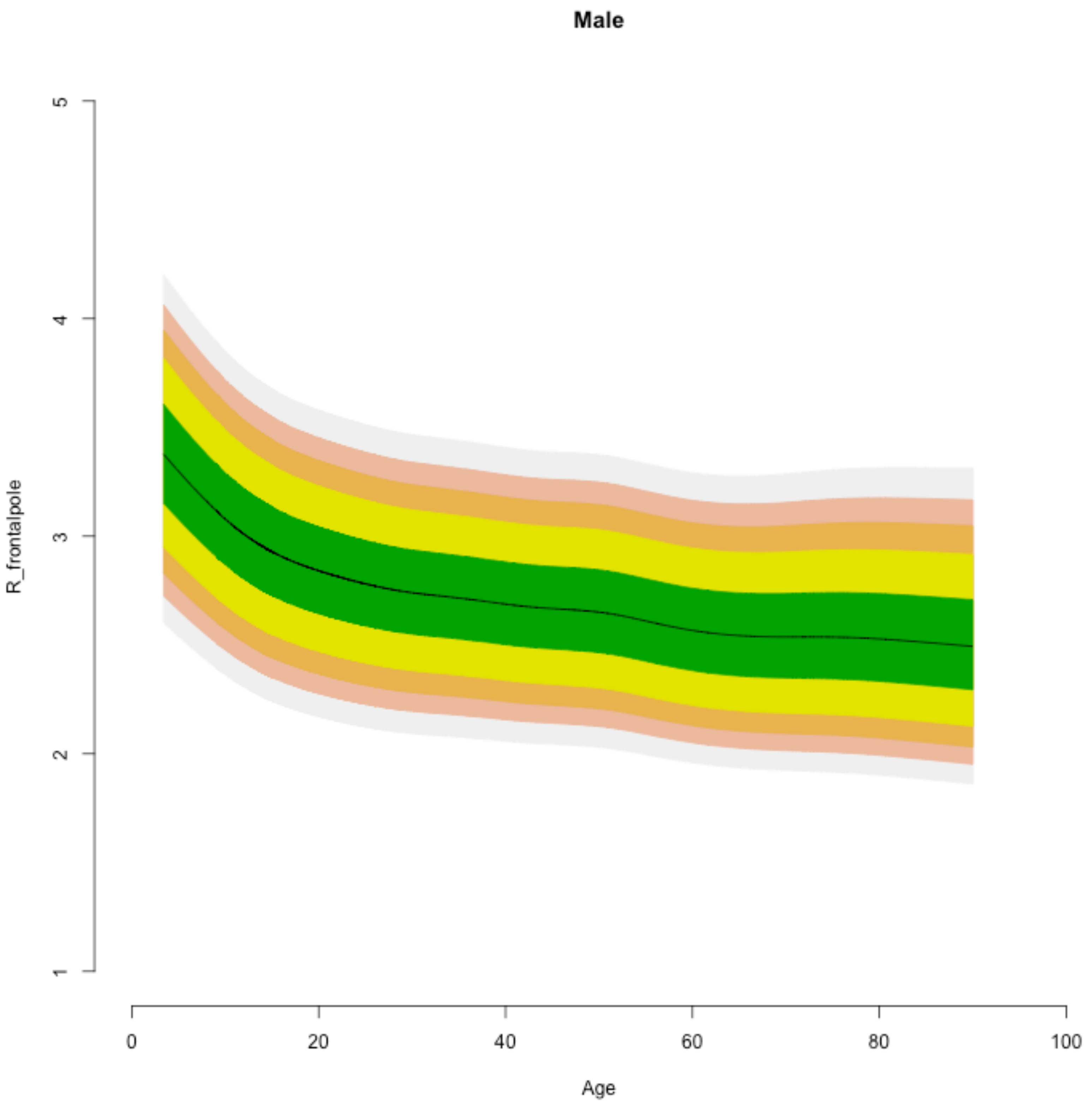

Male
R_frontalpole
1
2
3
4
5
0
20
40
60
80
100
Age

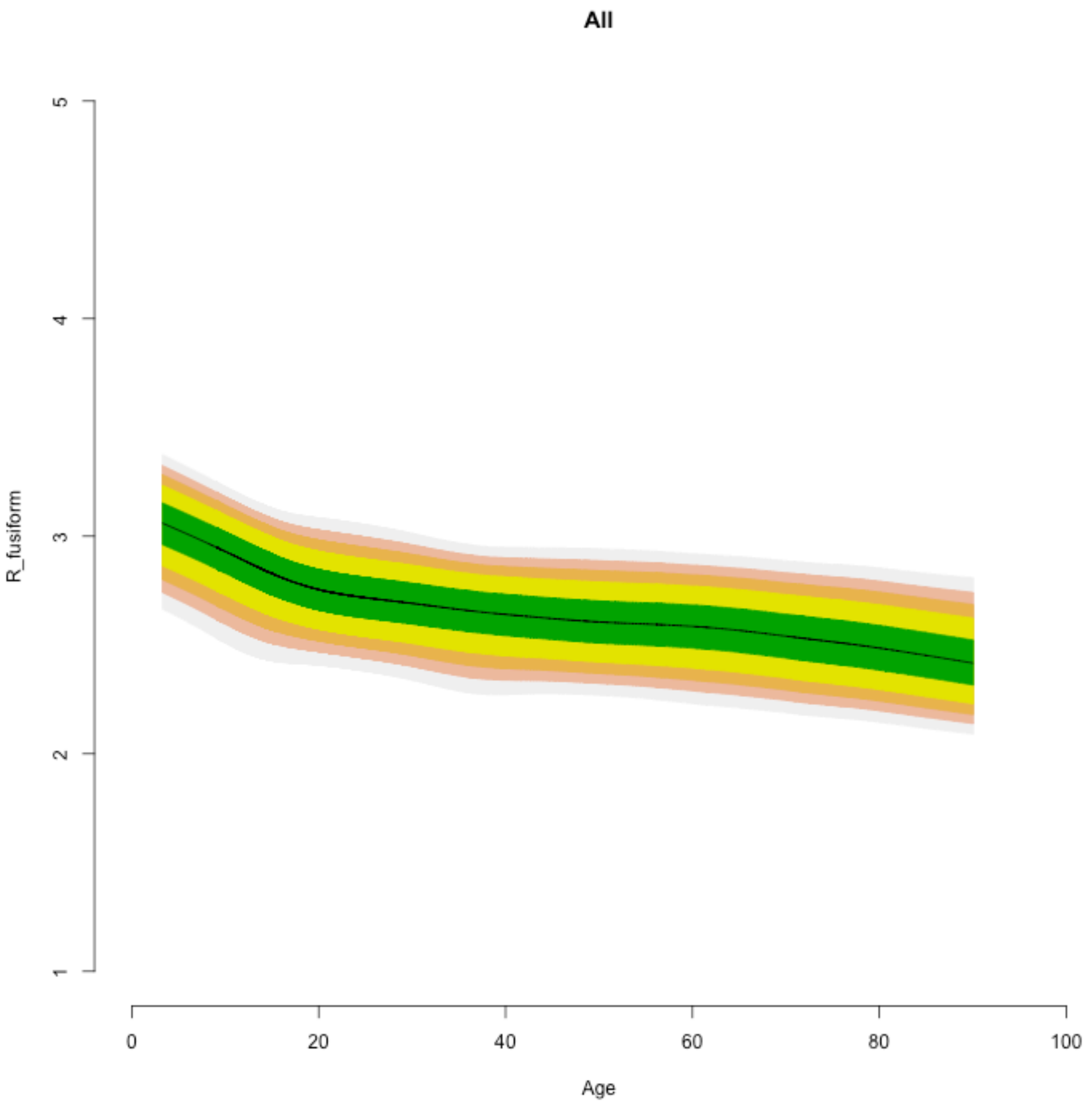
All
R_fusiform
5
4
3
2
1
0
20
40
60
80
100
Age

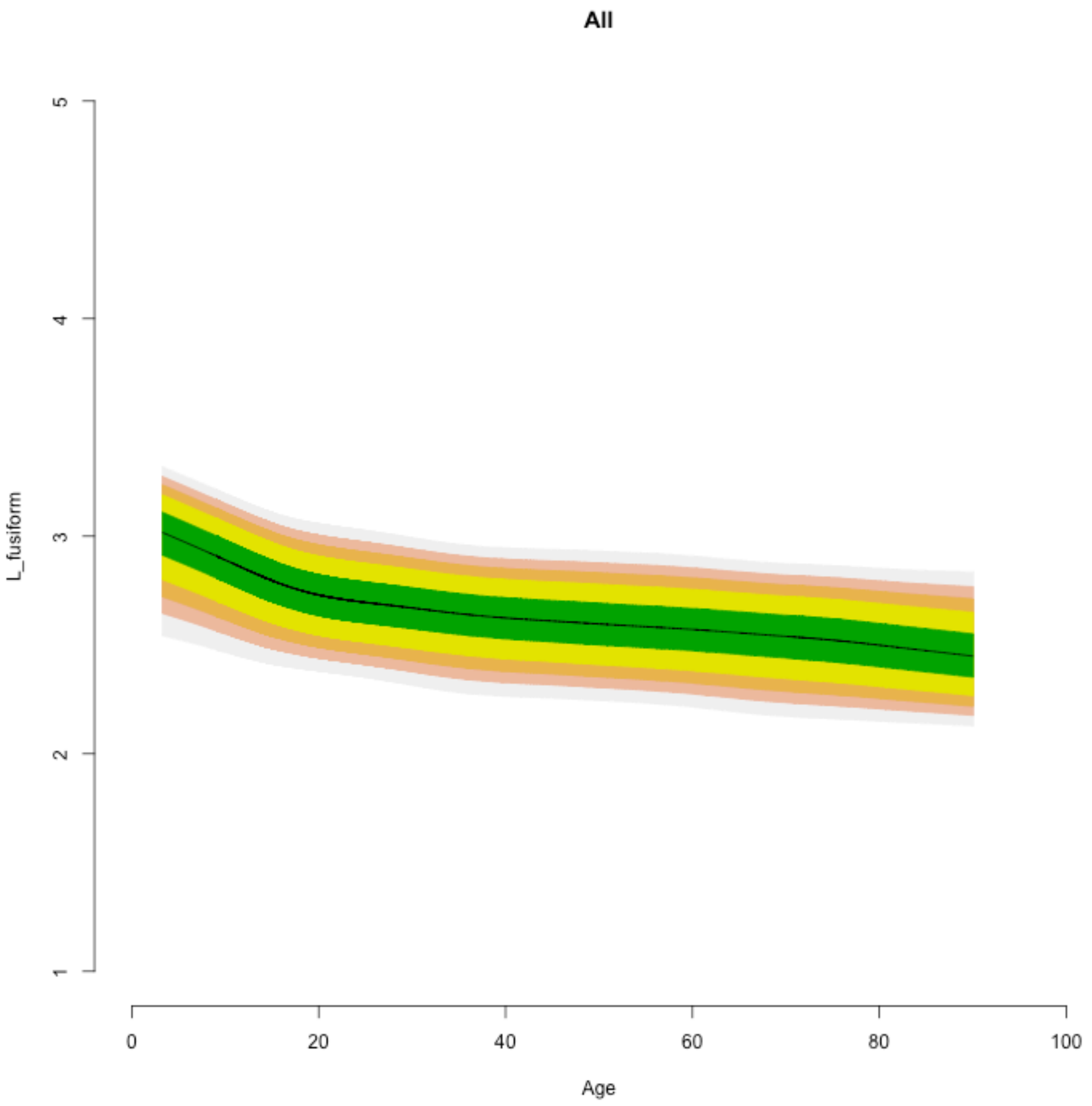
All
L_fusiform
1
2
3
4
5
0
20
40
60
80
100
Age

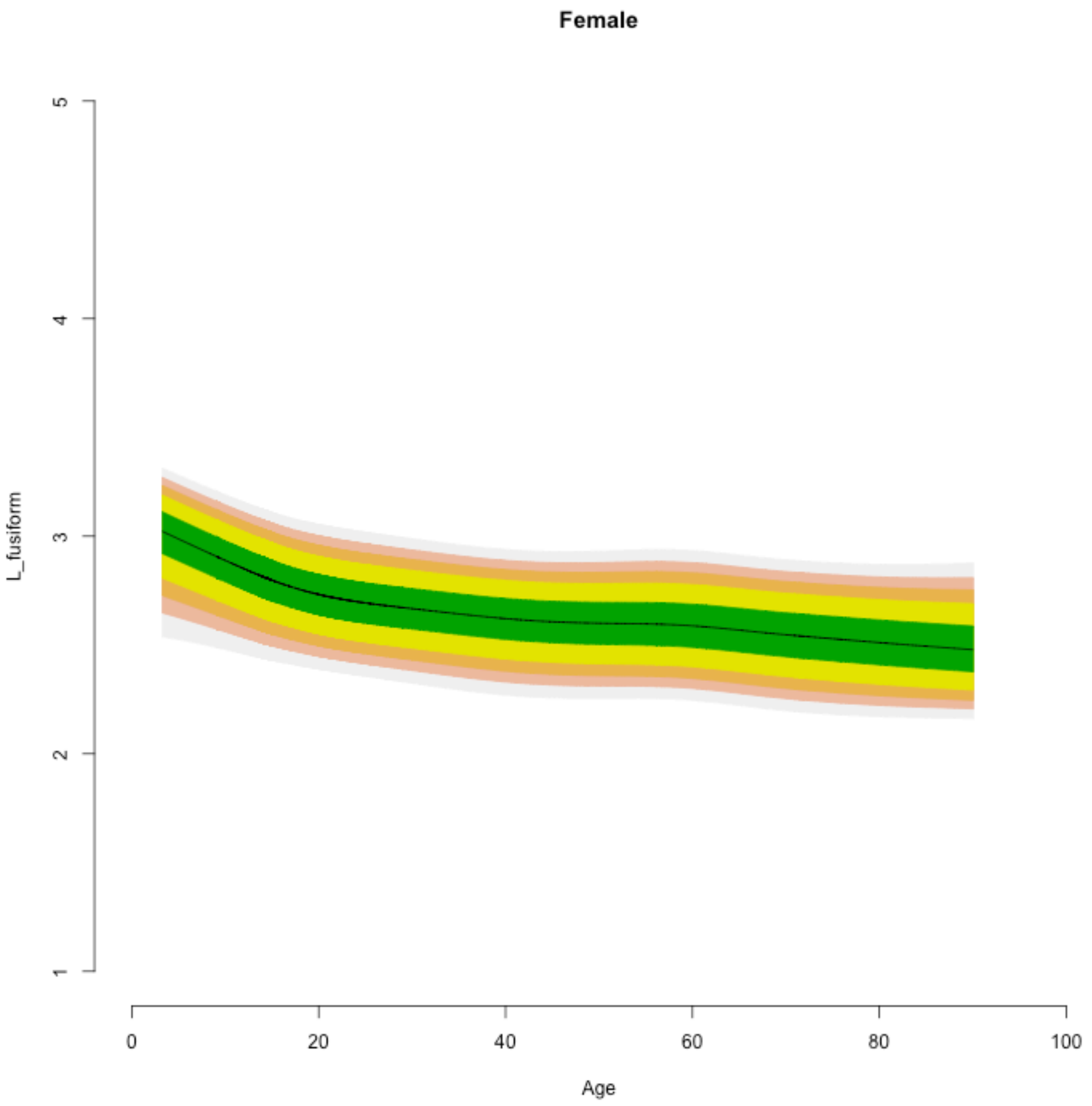

Female
L_fusiform
1
2
3
4
5
0
20
40
60
80
100
Age

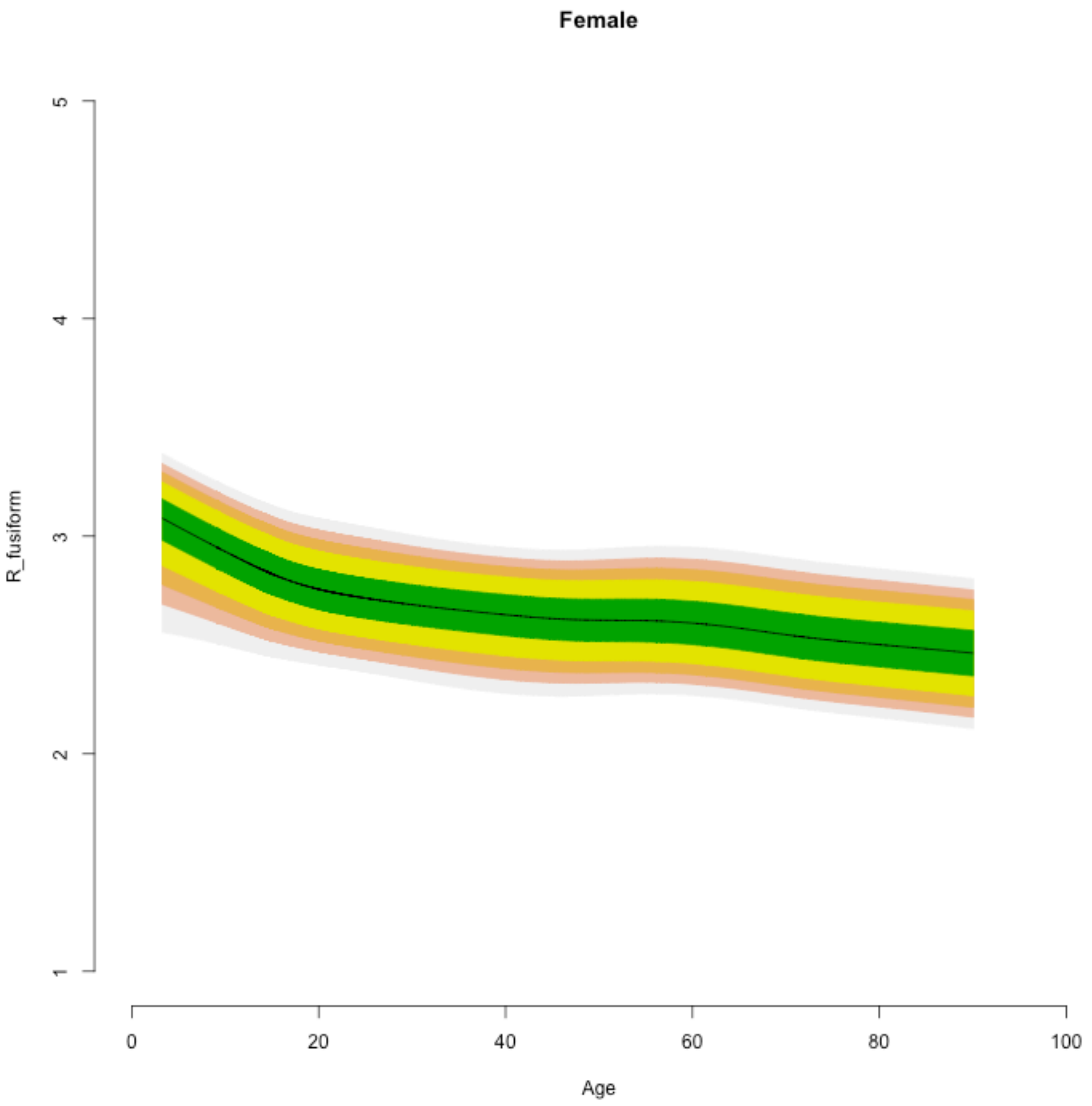

Female
R_fusiform
1
2
3
4
5
0
20
40
60
80
100
Age

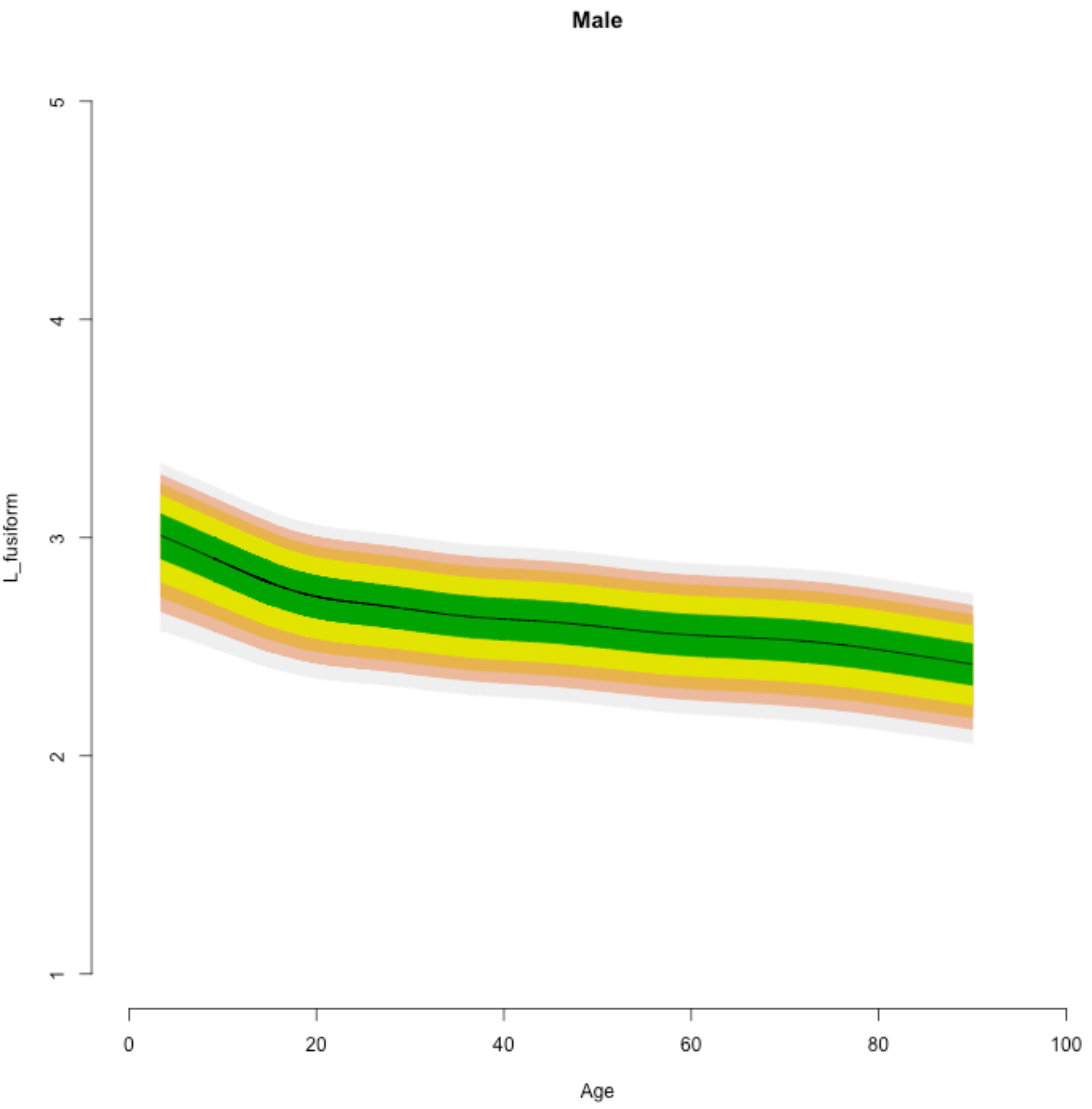

Male
L_fusiform
1
2
3
4
5
0
20
40
60
80
100
Age

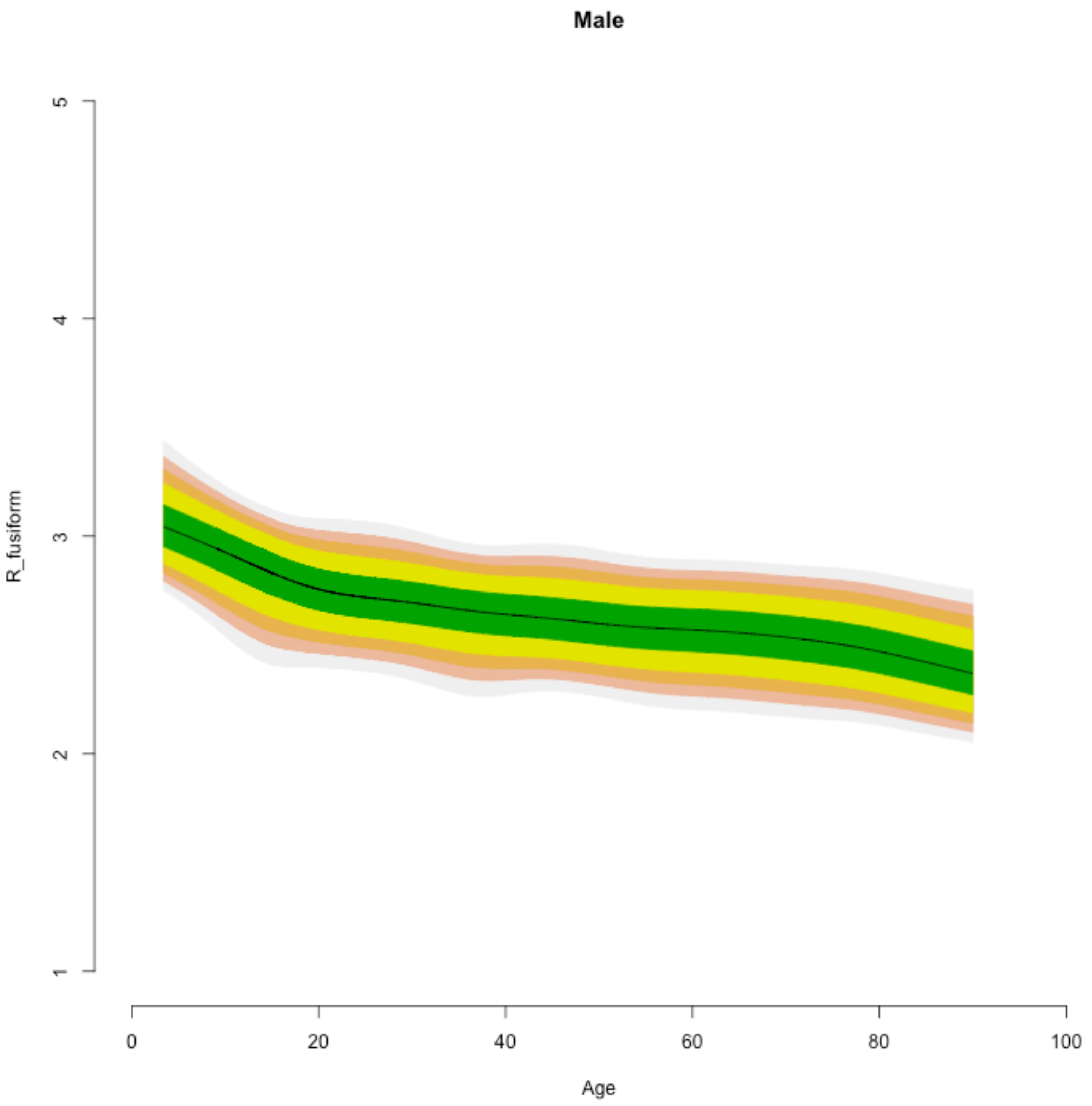
Male
R_fusiform
1
2
3
4
5
0
20
40
60
80
100
Age

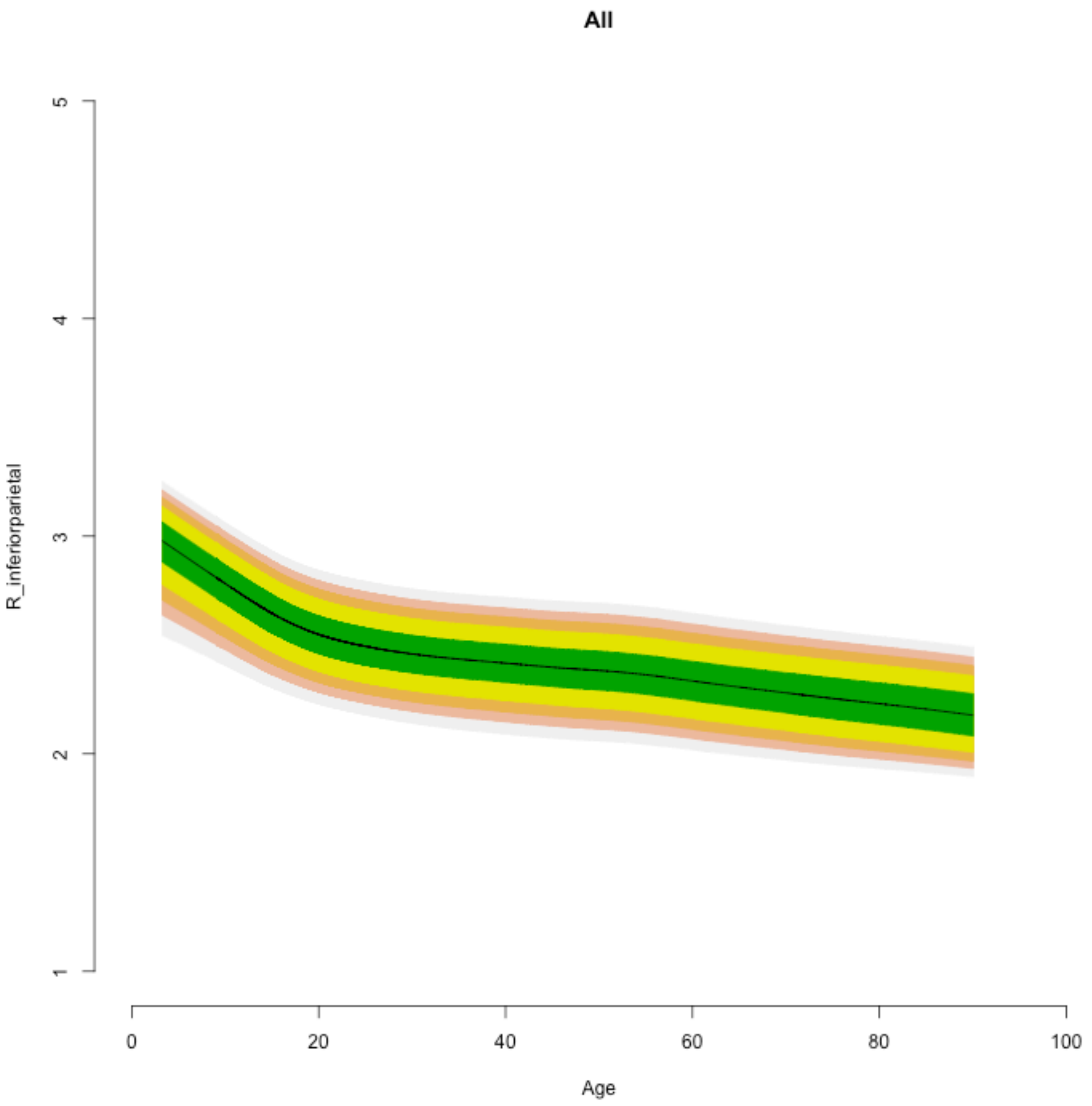

All
R_inferiorparietal
5
4
3
2
1
0
20
40
60
80
100
Age

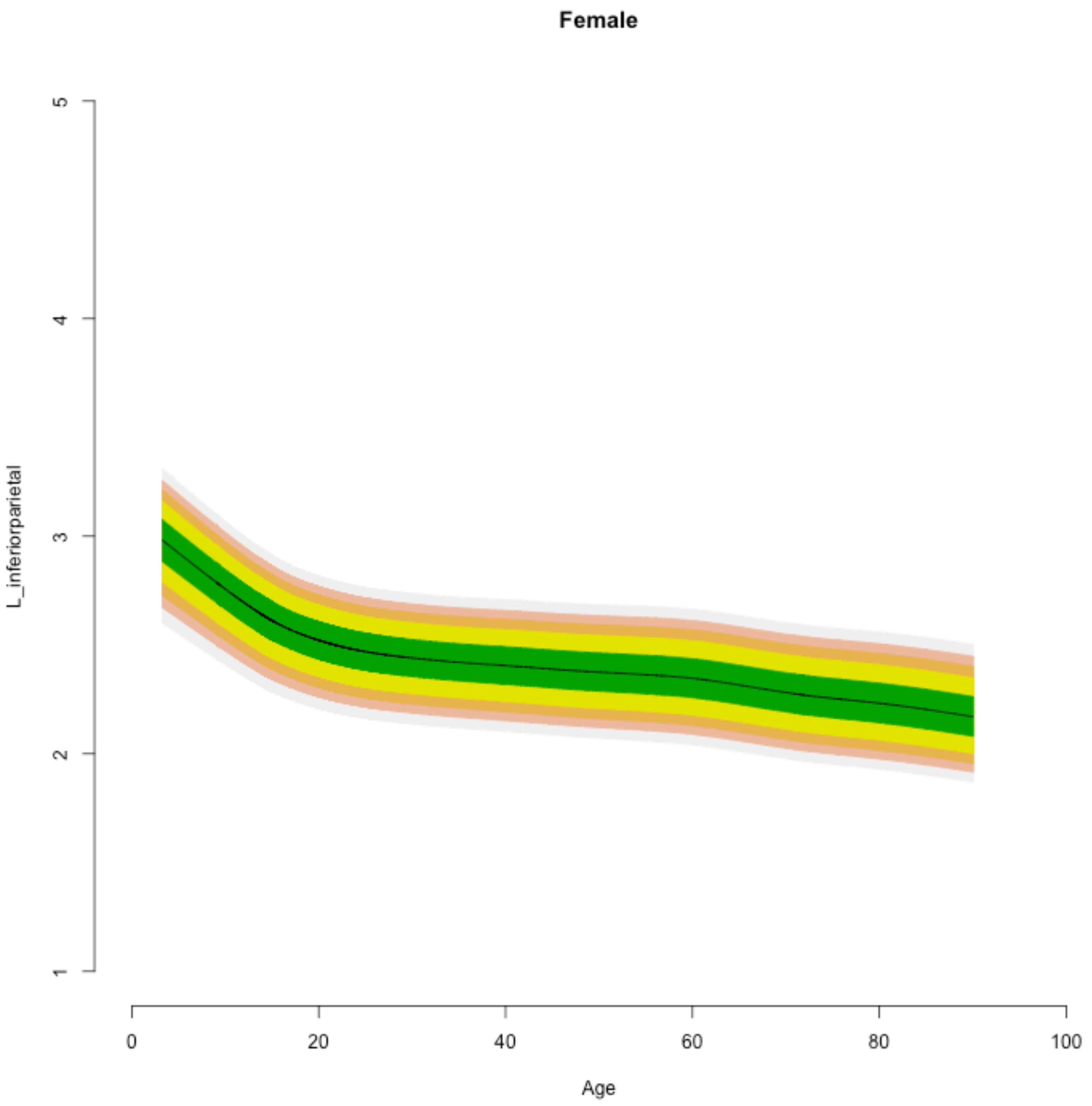

Female
L_inferiorparietal
1
2
3
4
5
0
20
40
60
80
100
Age

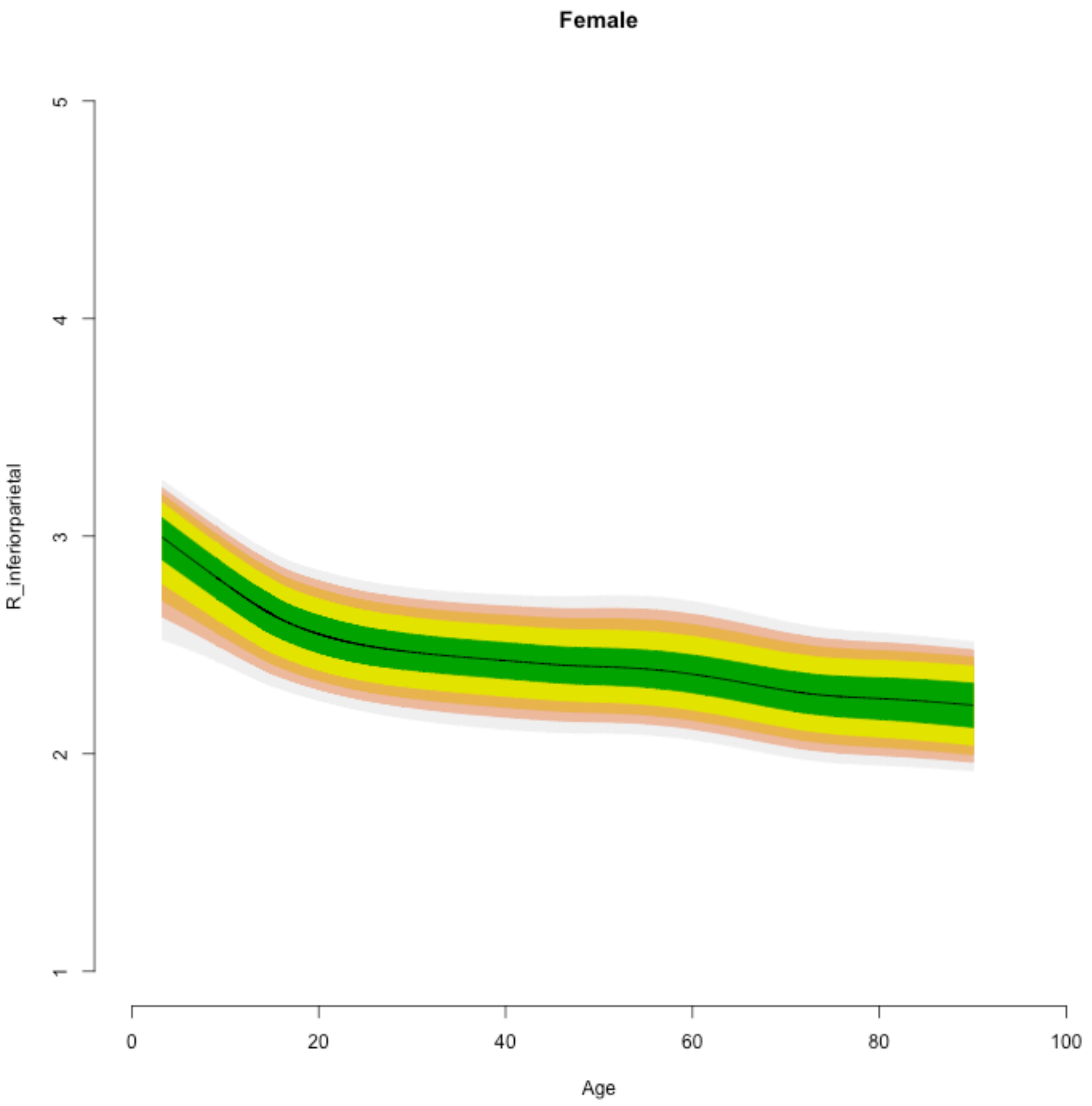
Female
R_inferiorparietal
Age
1
2
3
4
5
0
20
40
60
80
100

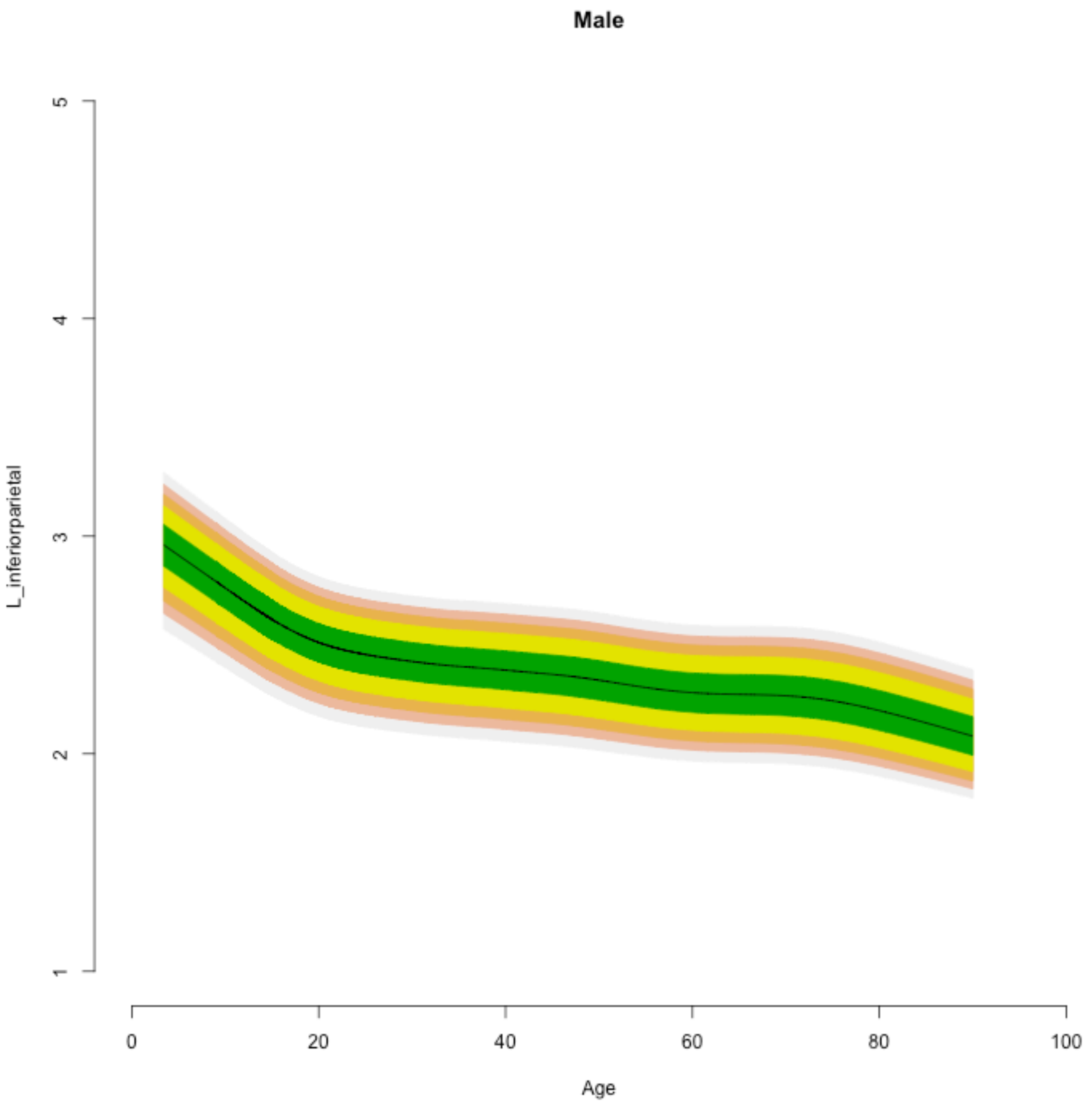

Male
L_inferiorparietal
1
2
3
4
5
0
20
40
60
80
100
Age

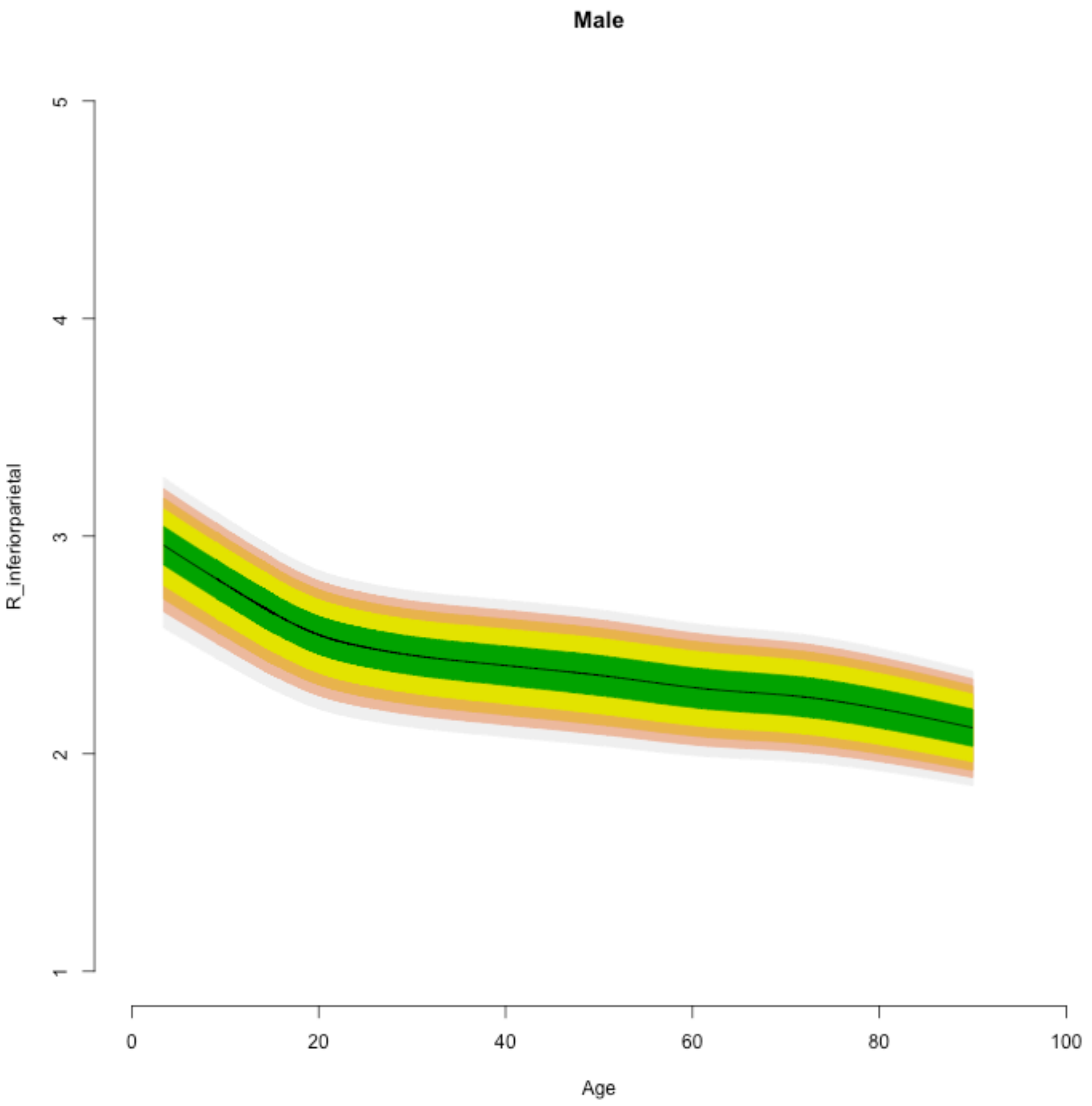
Male
R_inferiorparietal
5
4
3
2
1
0
20
40
60
80
100
Age

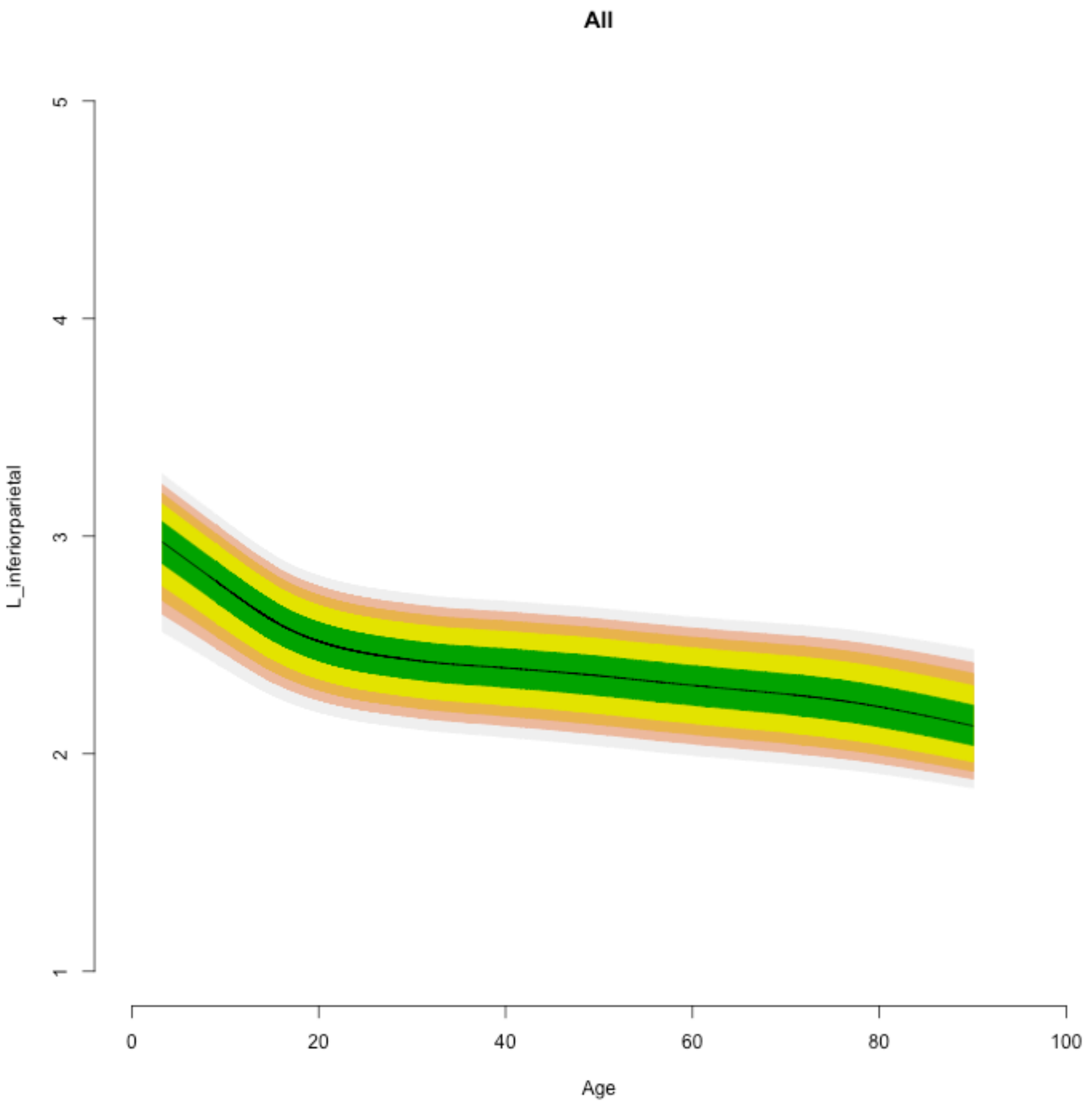

All
L_inferiorparietal
5
4
3
2
1
0
20
40
60
80
100
Age

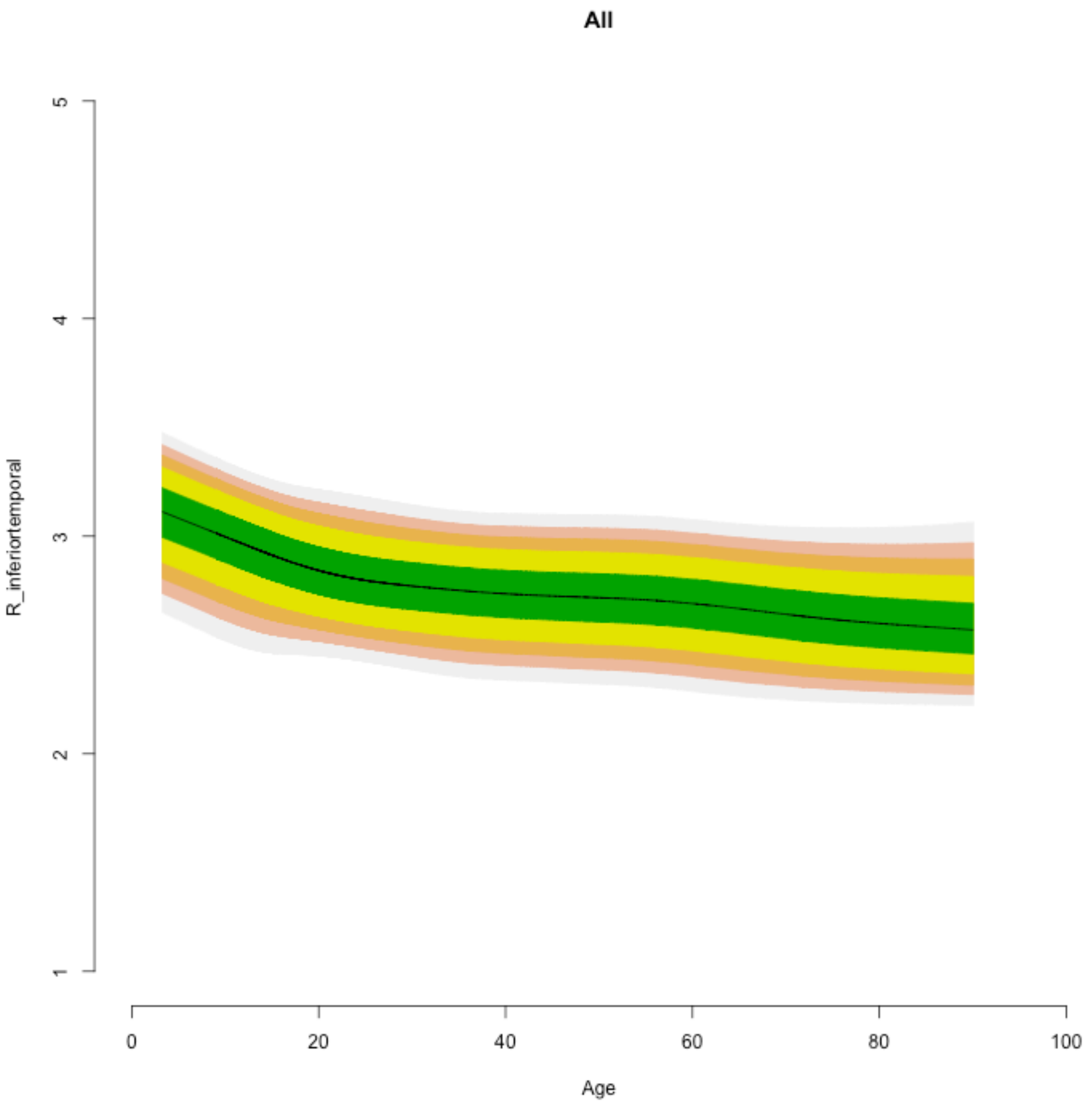
All
R_inferiortemporal
1
2
3
4
5
0
20
40
60
80
100
Age

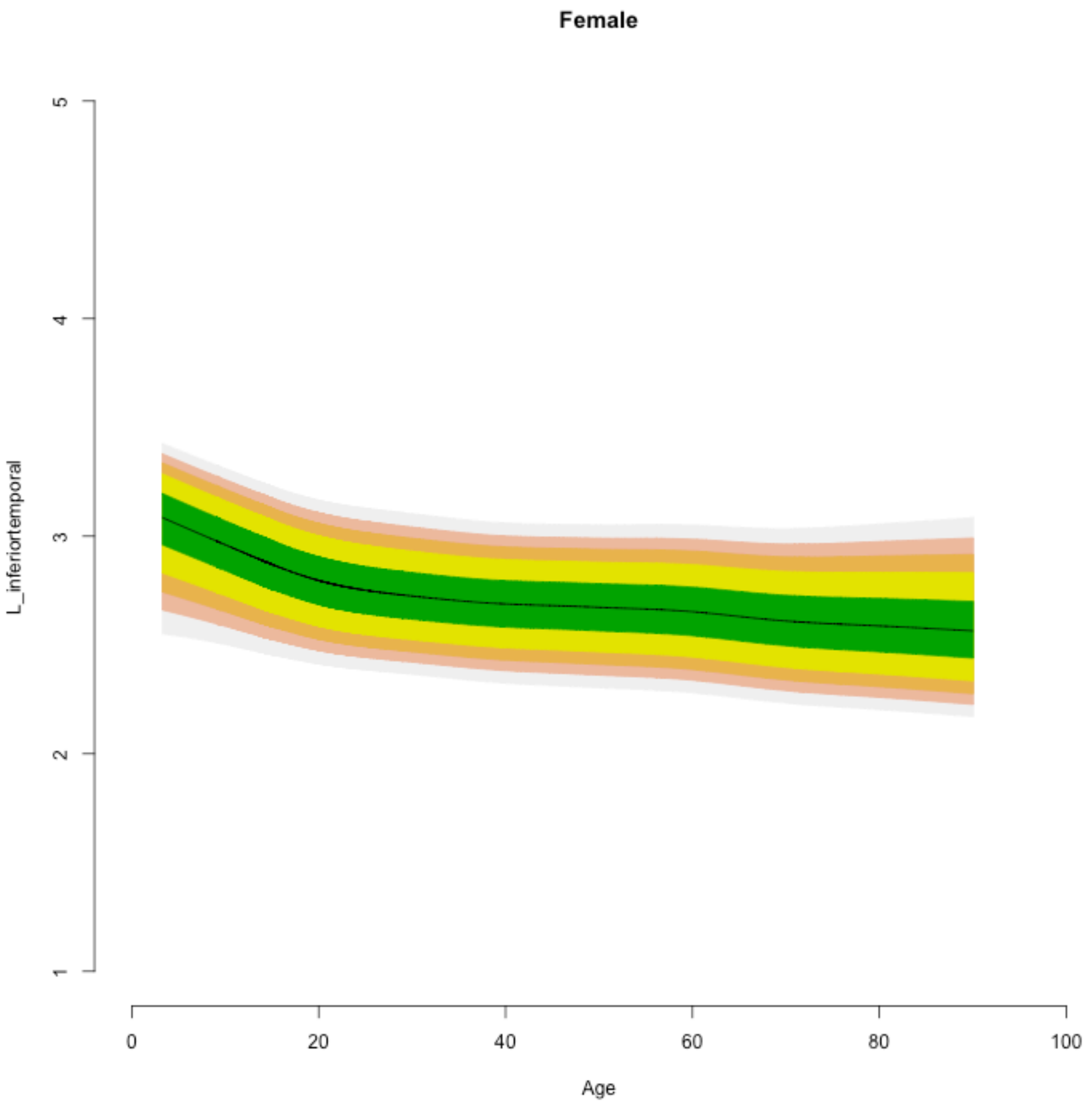

Female
L_inferiortemporal
1
2
3
4
5
0
20
40
60
80
100
Age

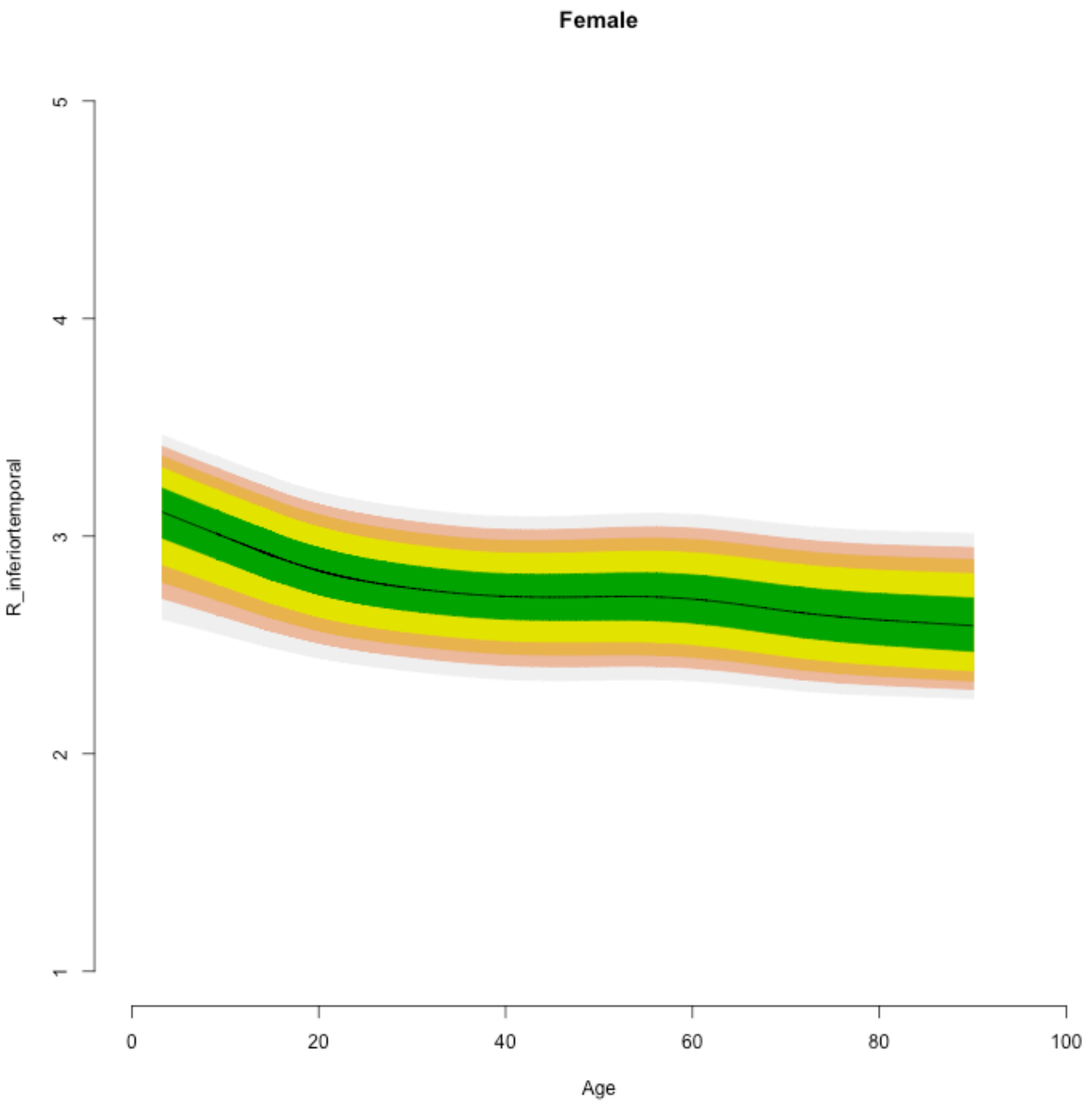

Female
R_inferiortemporal
1
2
3
4
5
0
20
40
60
80
100
Age

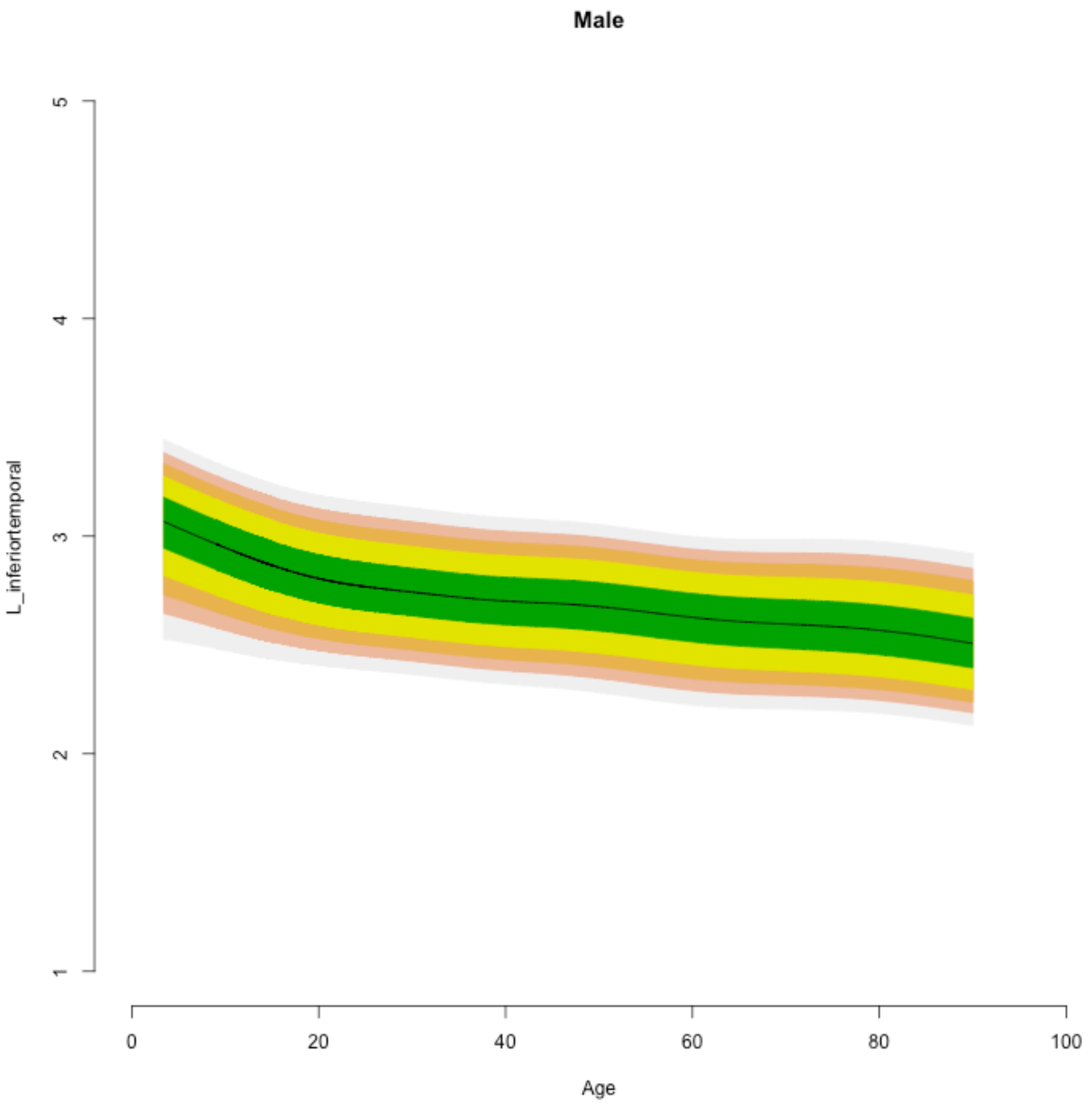

Male
L_inferiortemporal
1
2
3
4
5
0
20
40
60
80
100
Age

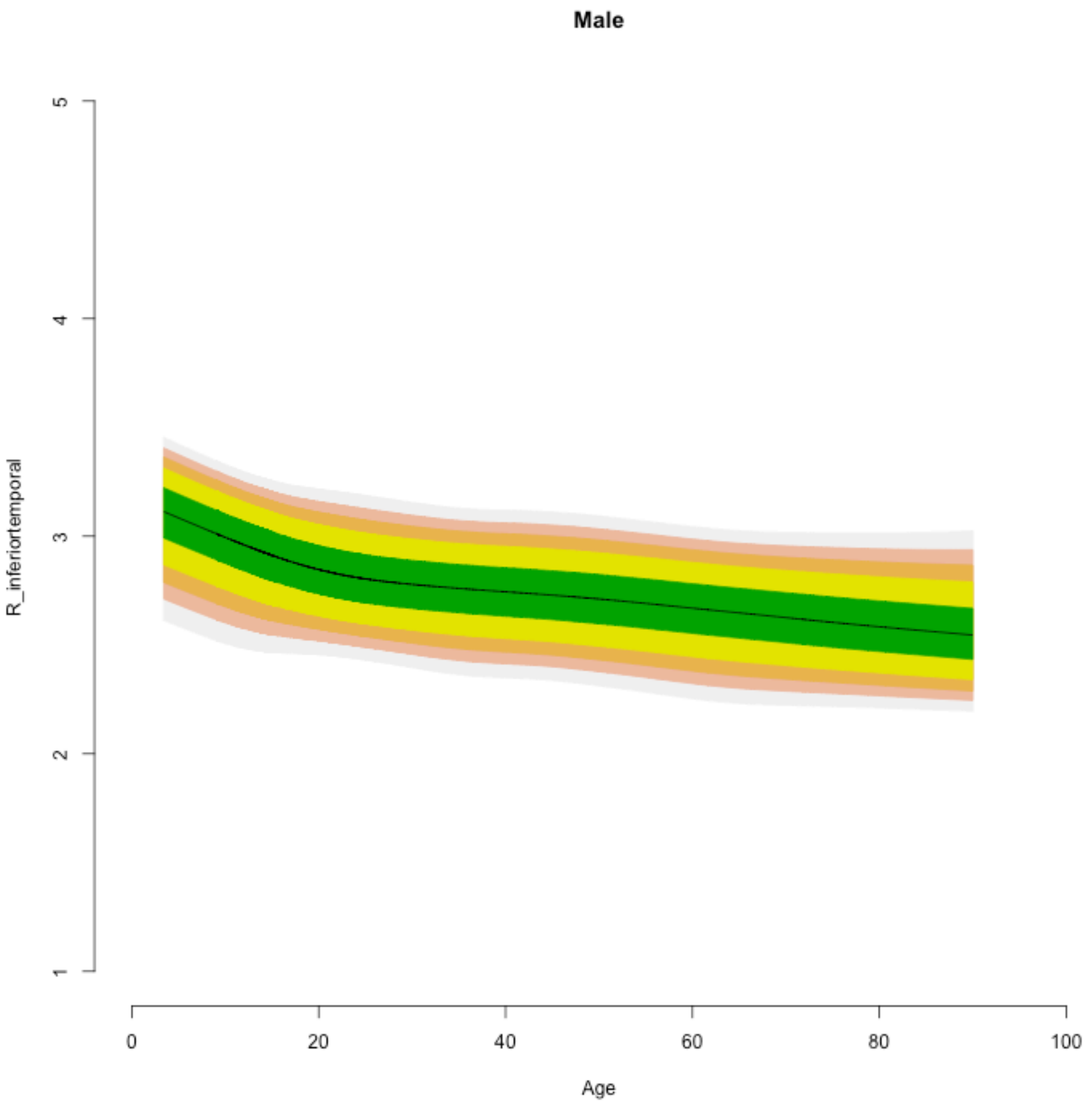

Male
R_inferiortemporal
1
2
3
4
5
0
20
40
60
80
100
Age

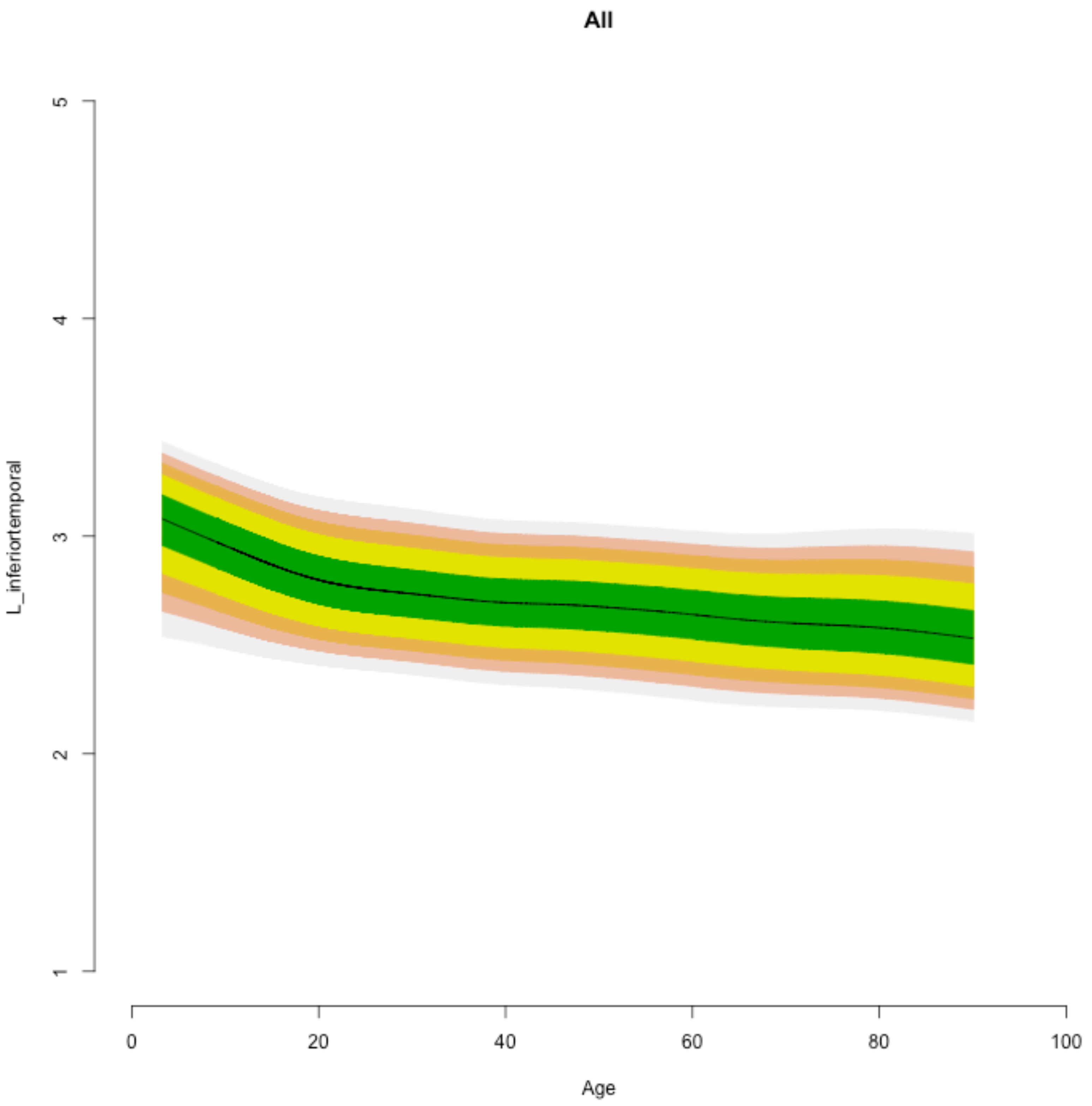
All
L_inferiortemporal
1
2
3
4
5
0
20
40
60
80
100
Age

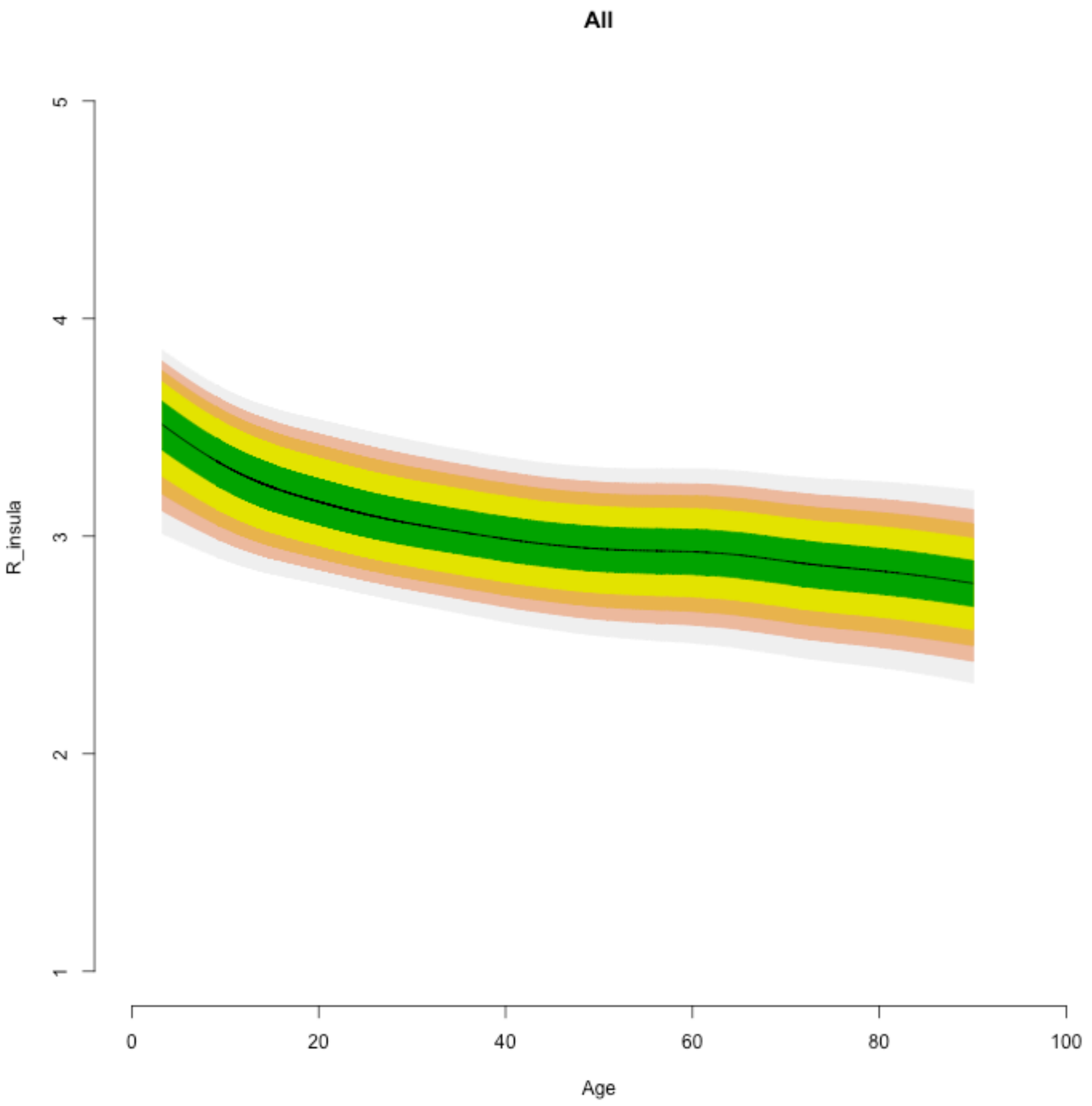
All
R_insula
5
4
3
2
1
0
20
40
60
80
100
Age

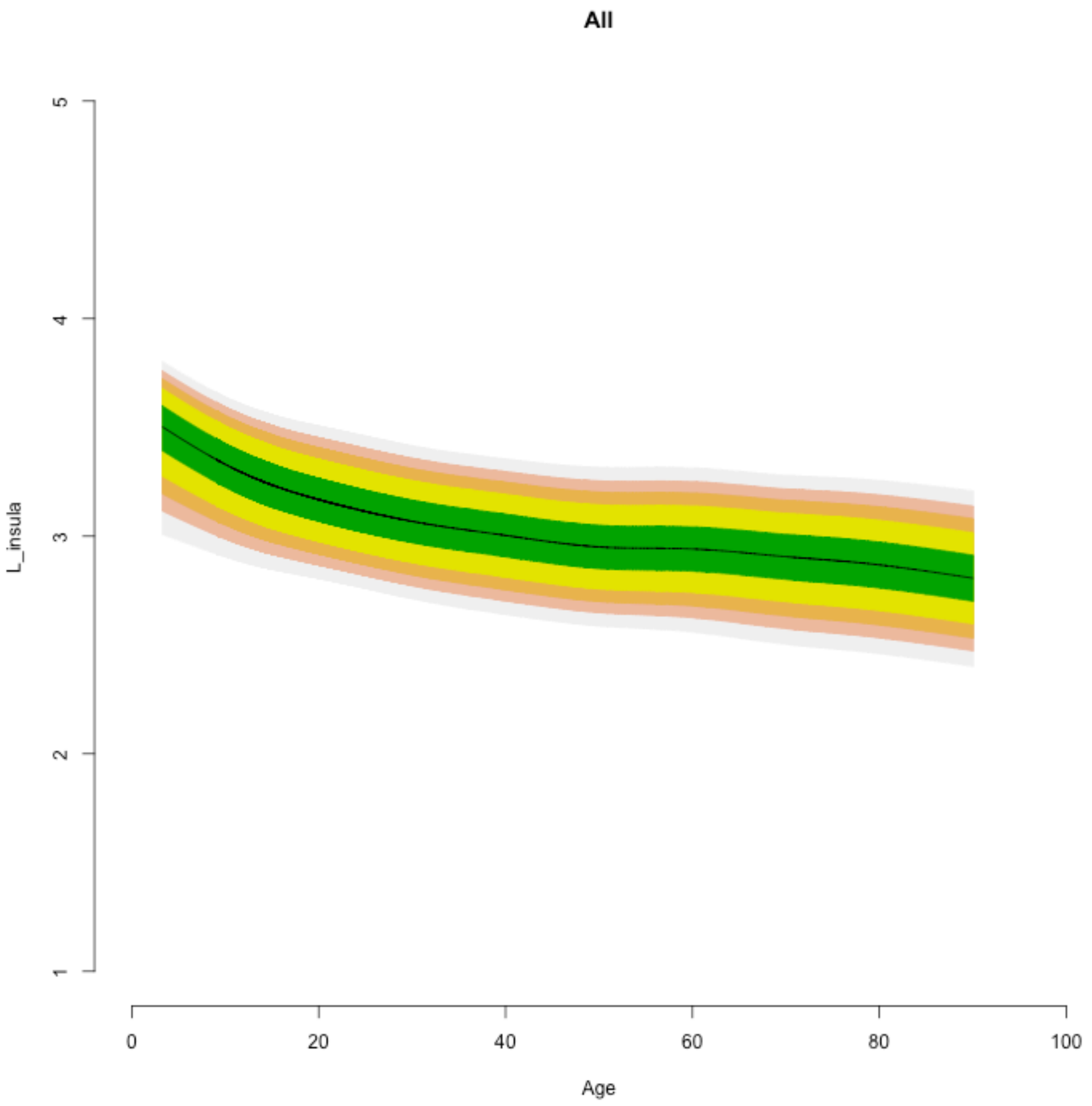

All
L_insula
1
2
3
4
5
0
20
40
60
80
100
Age

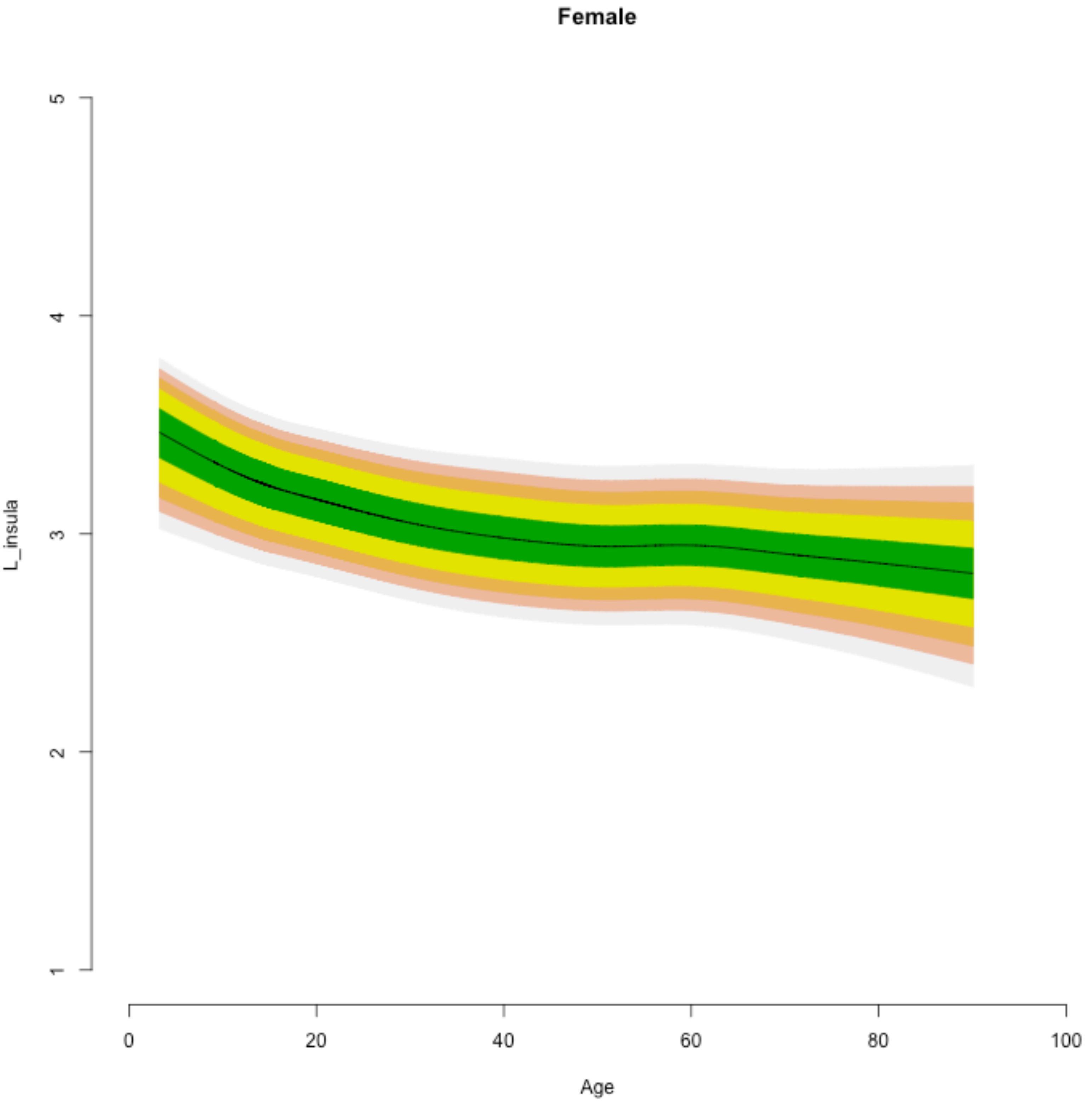
Female
L_insula
5
4
3
2
1
0
20
40
60
80
100
Age

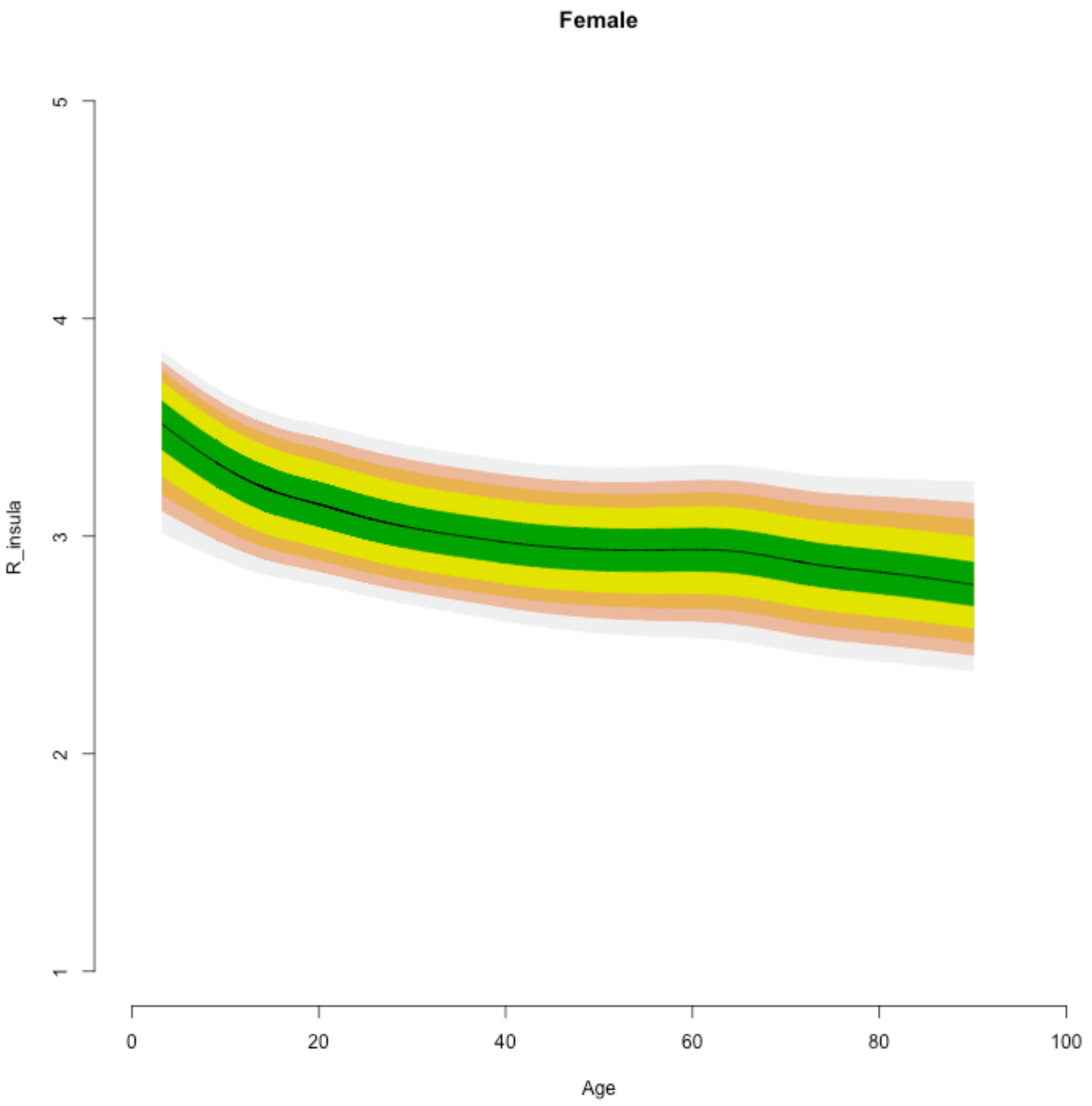

Female
R_insula
5
4
3
2
1
0
20
40
60
80
100
Age

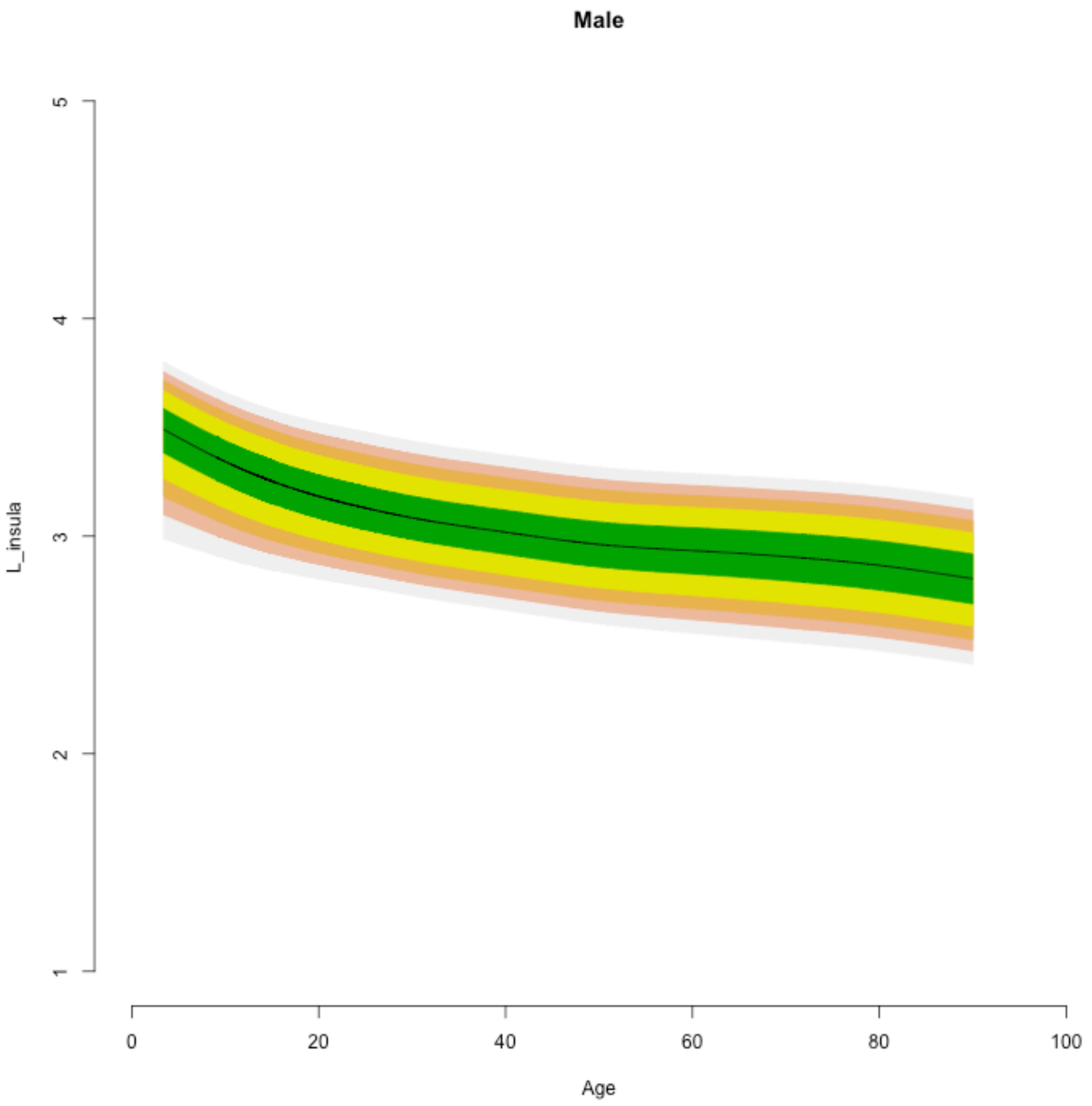

Male
L_insula
1
2
3
4
5
0
20
40
60
80
100
Age

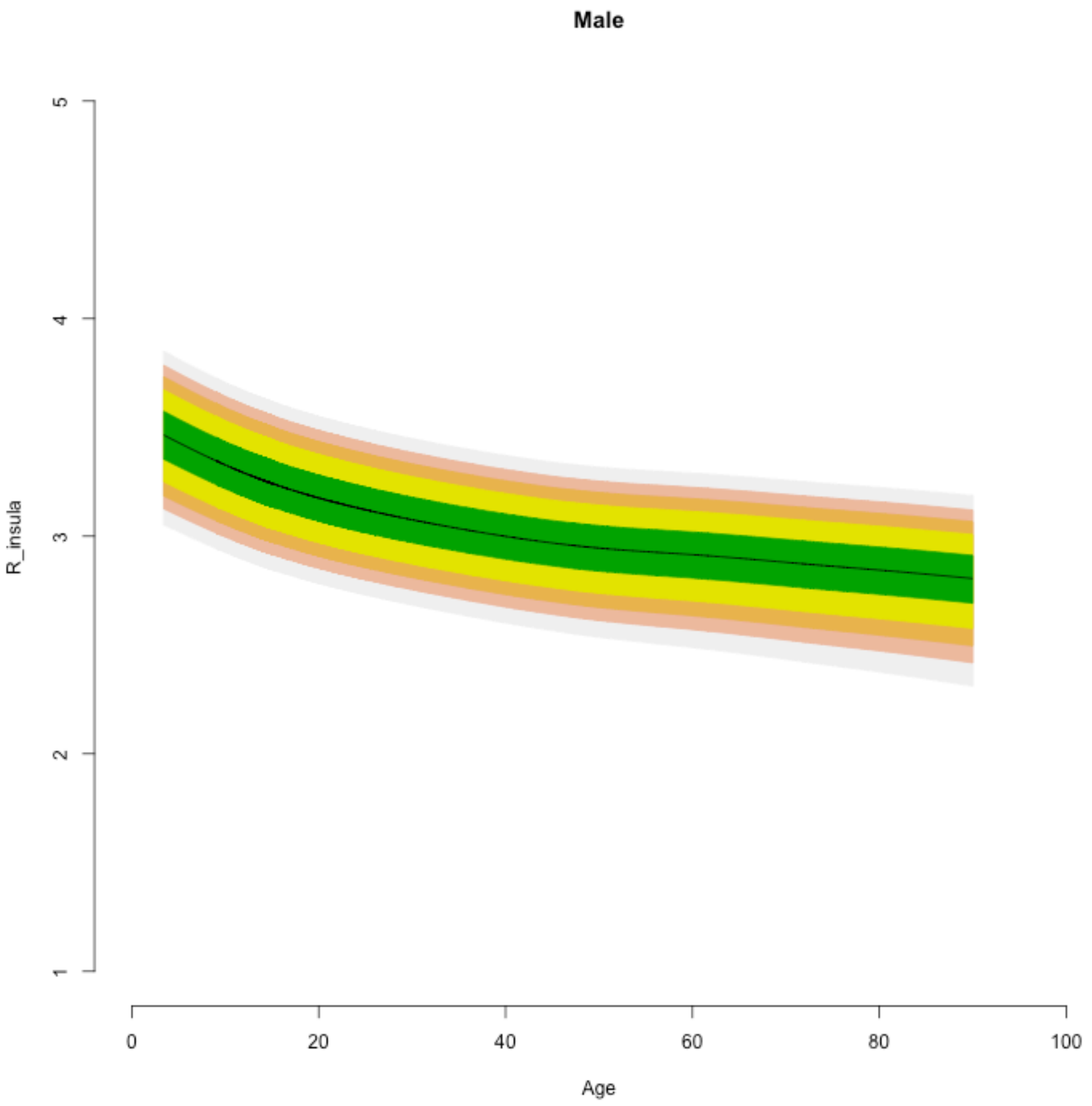

Male
R_insula
5
4
3
2
1
0
20
40
60
80
100
Age

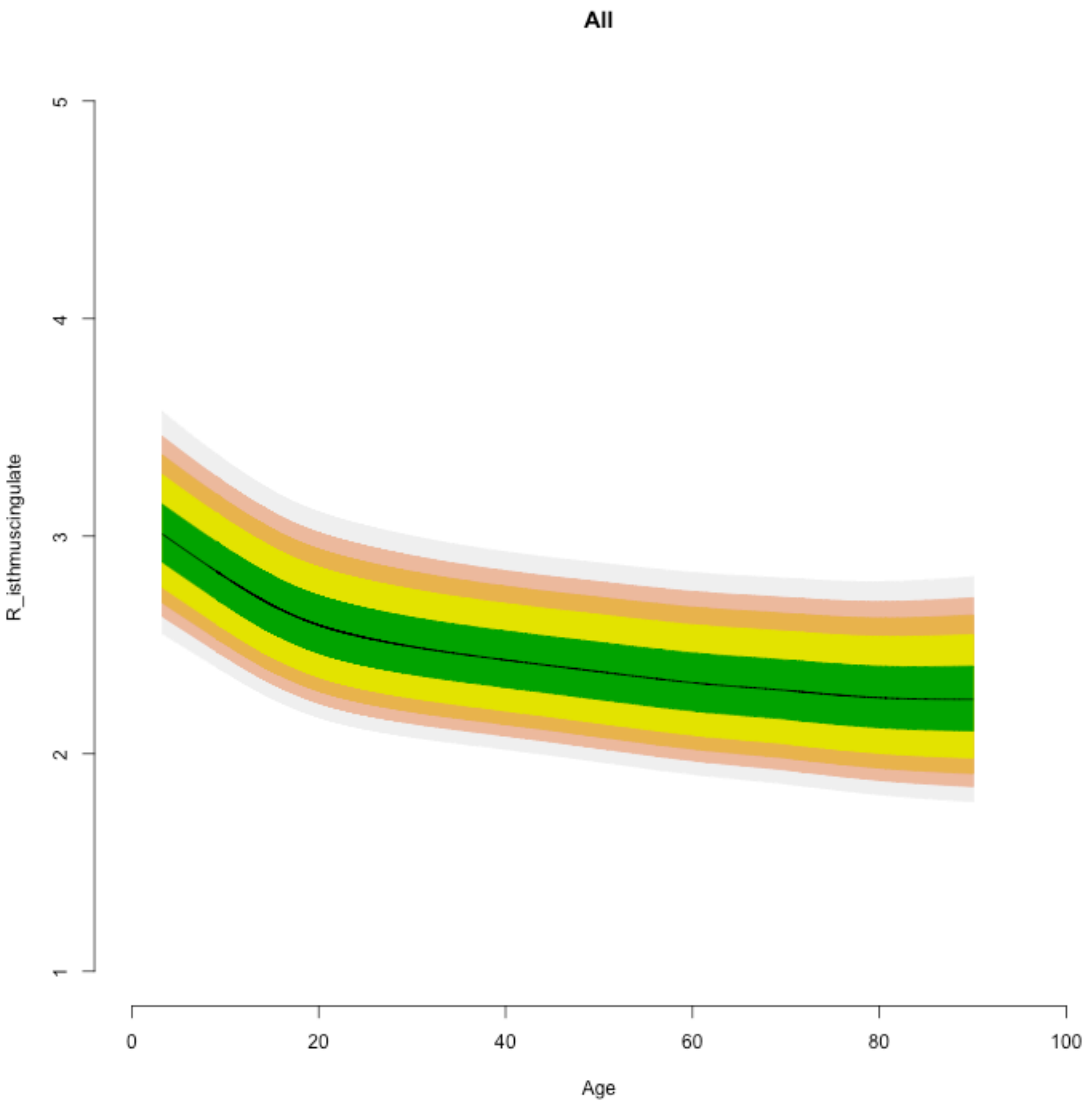

All
R_isthmuscingulate
Age
1
2
3
4
5
0
20
40
60
80
100

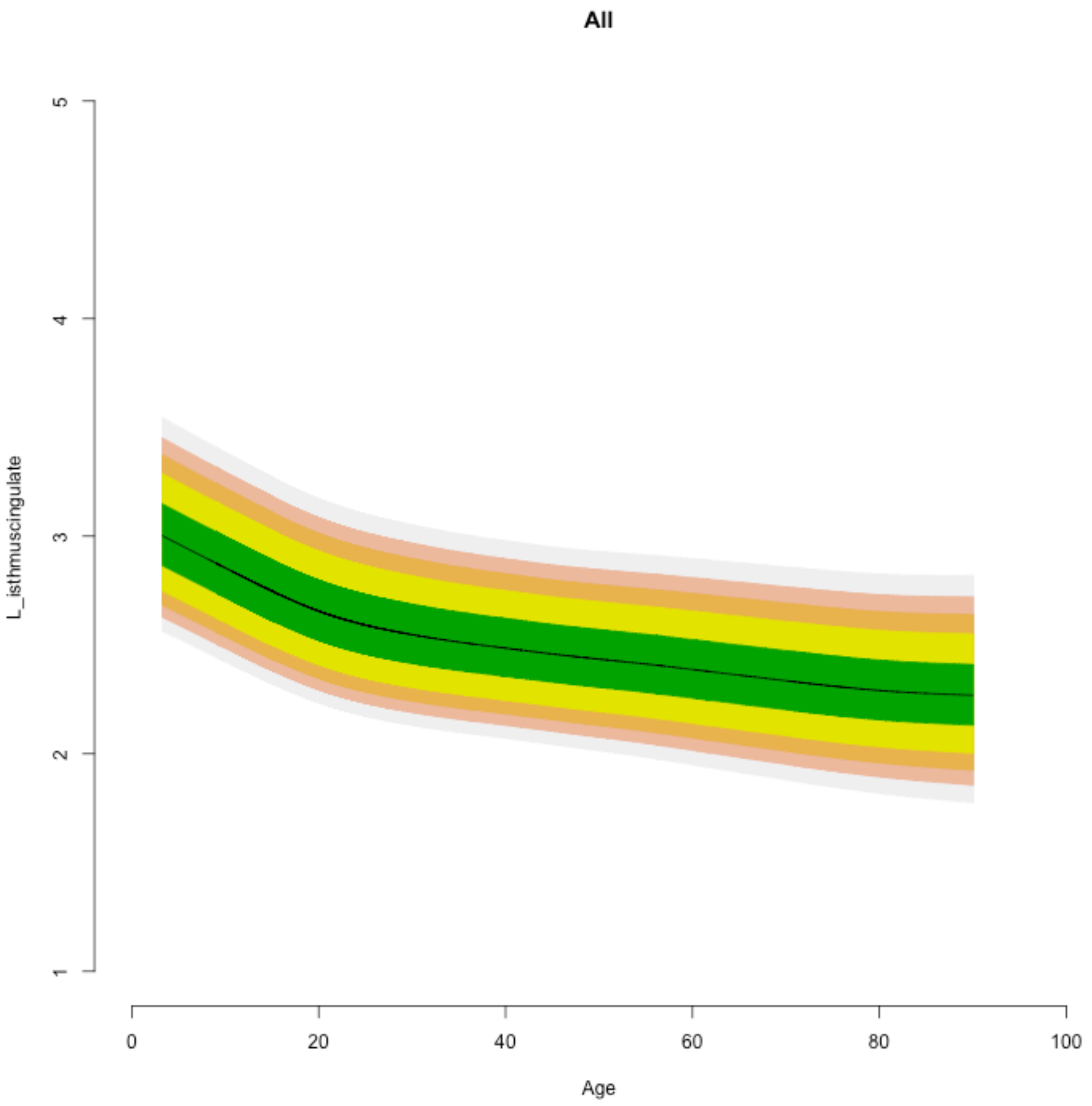

All
L_isthmuscingulate
1
2
3
4
5
0
20
40
60
80
100
Age

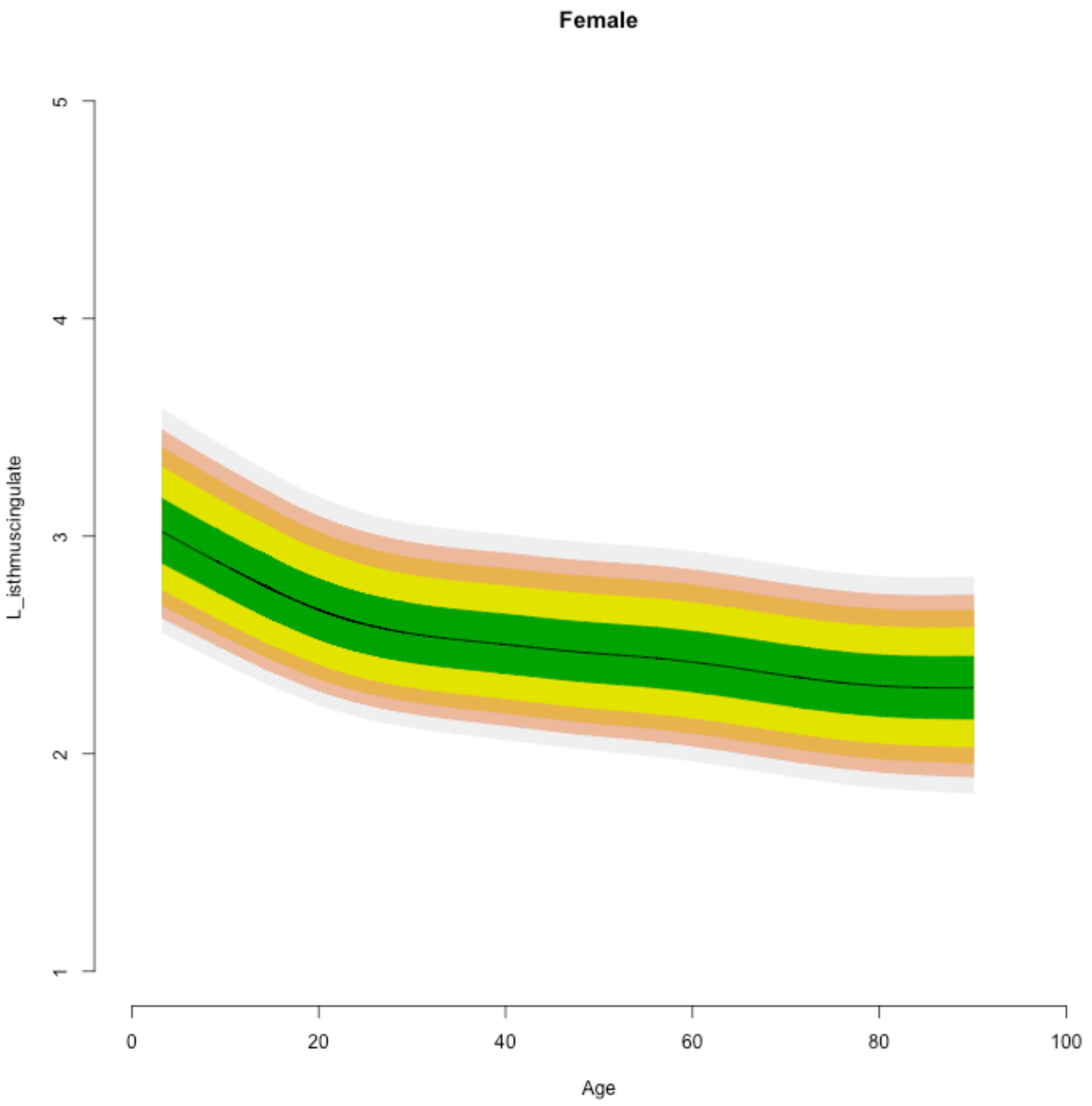
Female
L_isthmuscingulate
5
4
3
2
1
0
20
40
60
80
100
Age

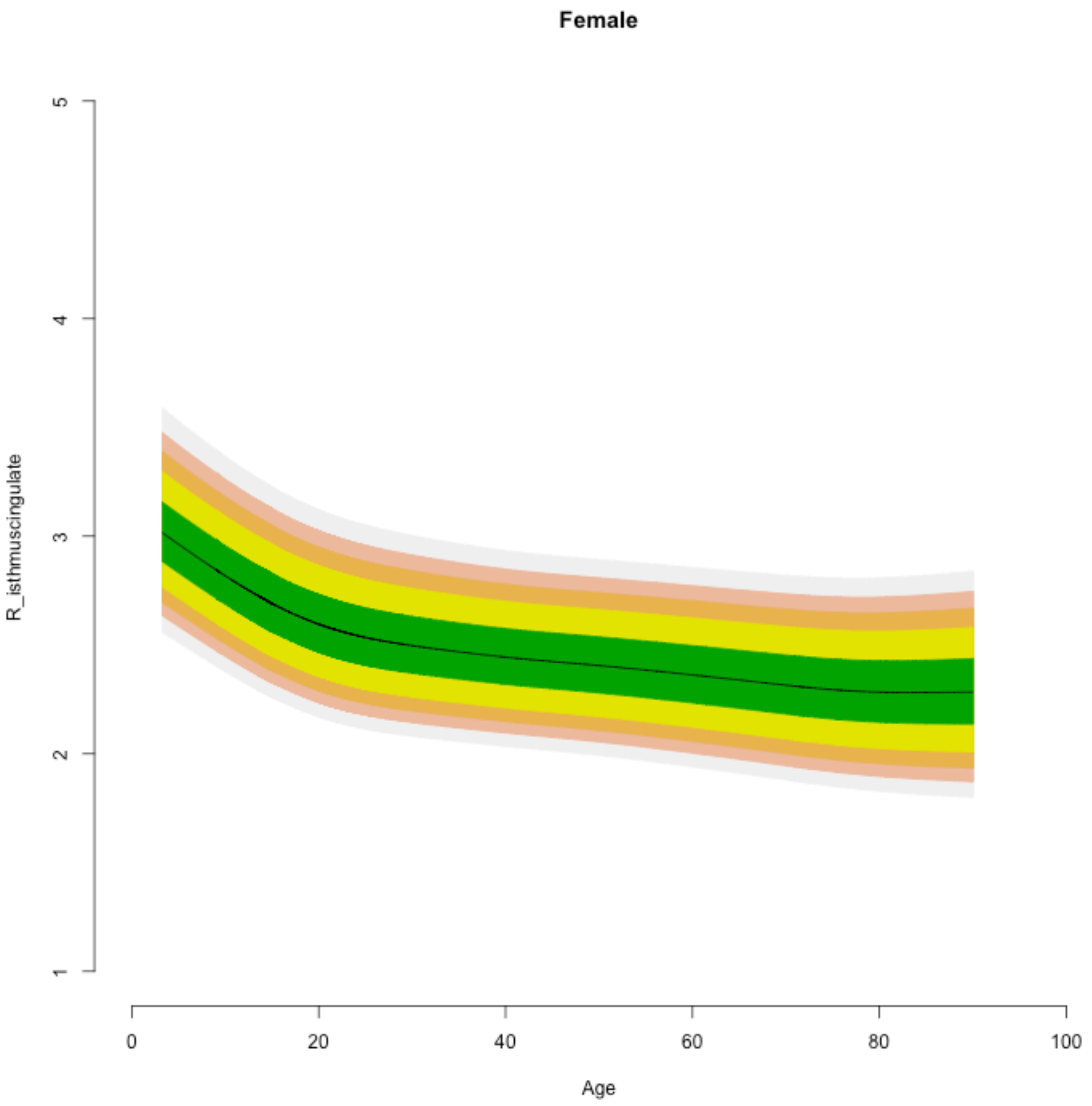

Female
R_isthmuscingulate
1
2
3
4
5
0
20
40
60
80
100
Age

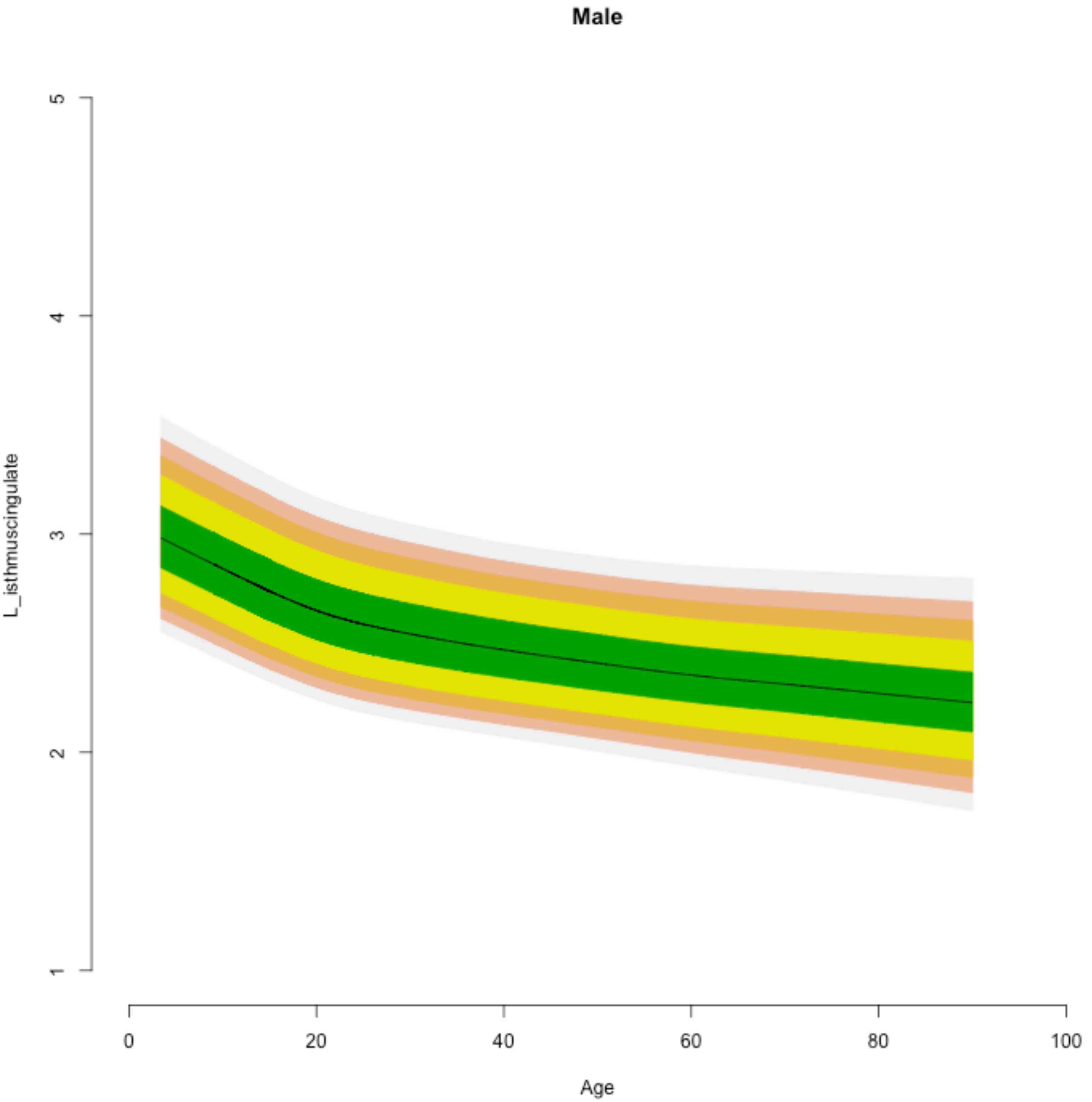
Male
L_isthmuscingulate
1
2
3
4
5
0
20
40
60
80
100
Age

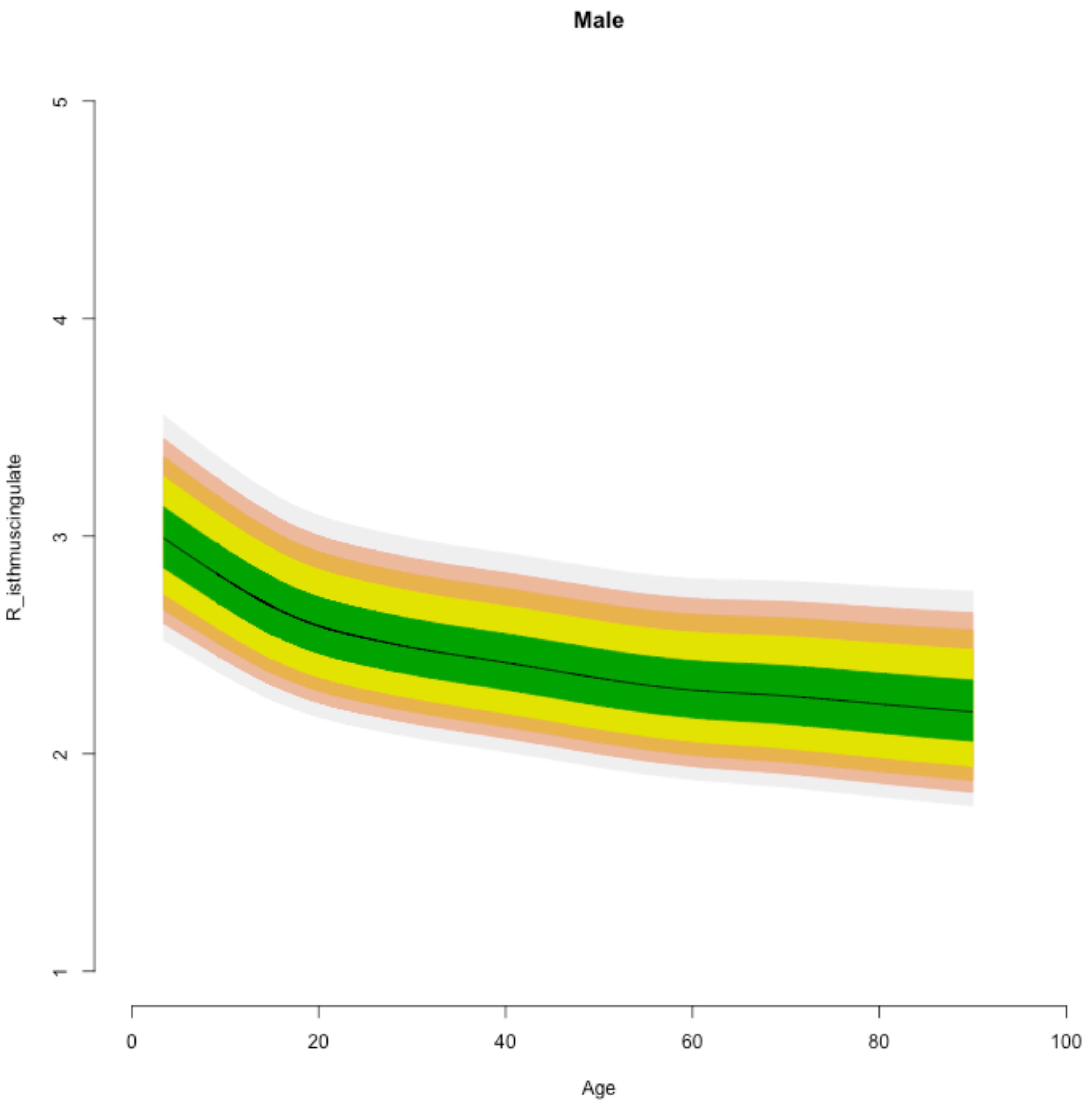
Male
R_isthmuscingulate
Age
1
2
3
4
5
0
20
40
60
80
100

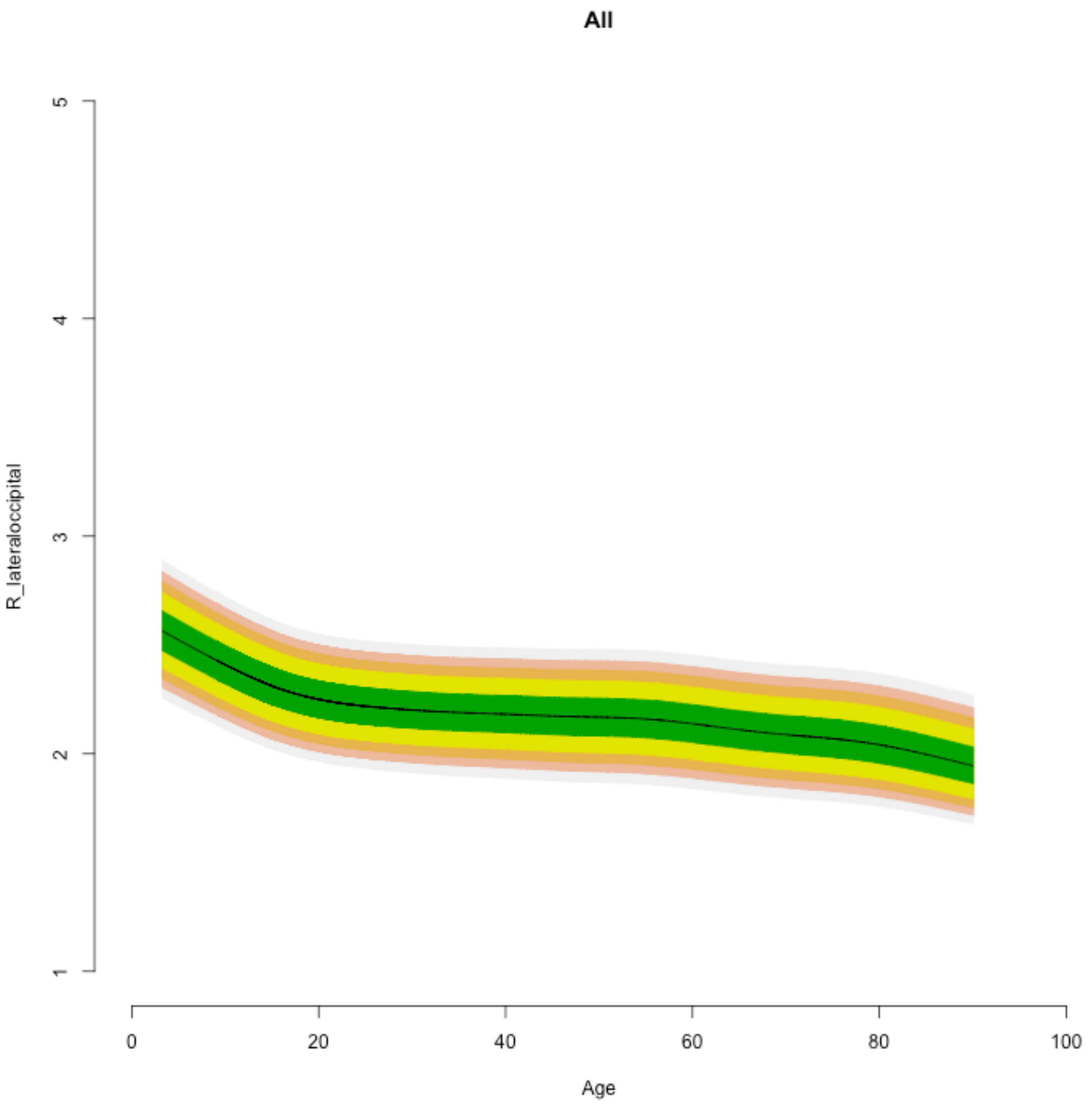
All
R_lateraloccipital
1
2
3
4
5
0
20
40
60
80
100
Age

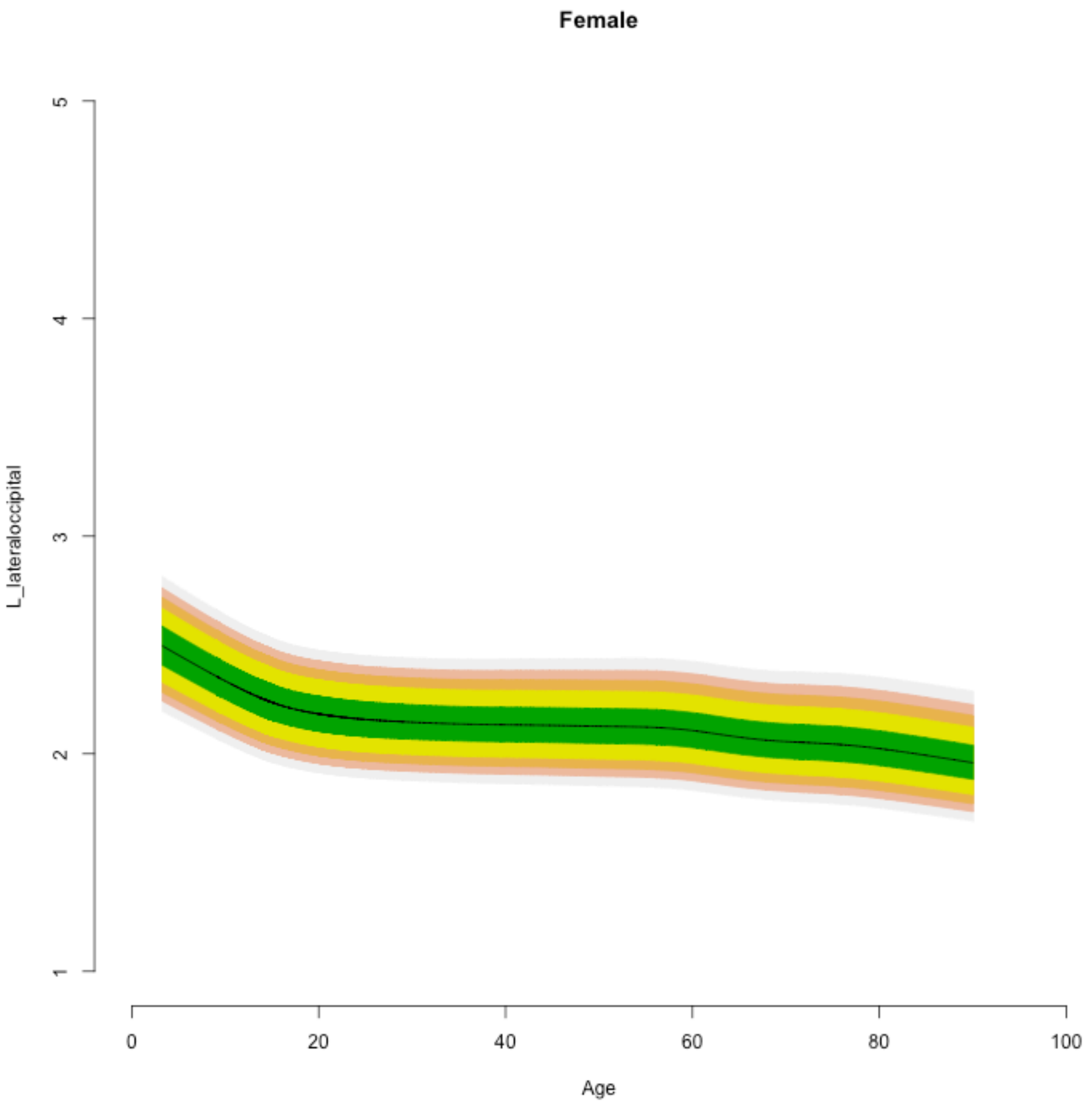

Female
L_lateraloccipital
1
2
3
4
5
0
20
40
60
80
100
Age

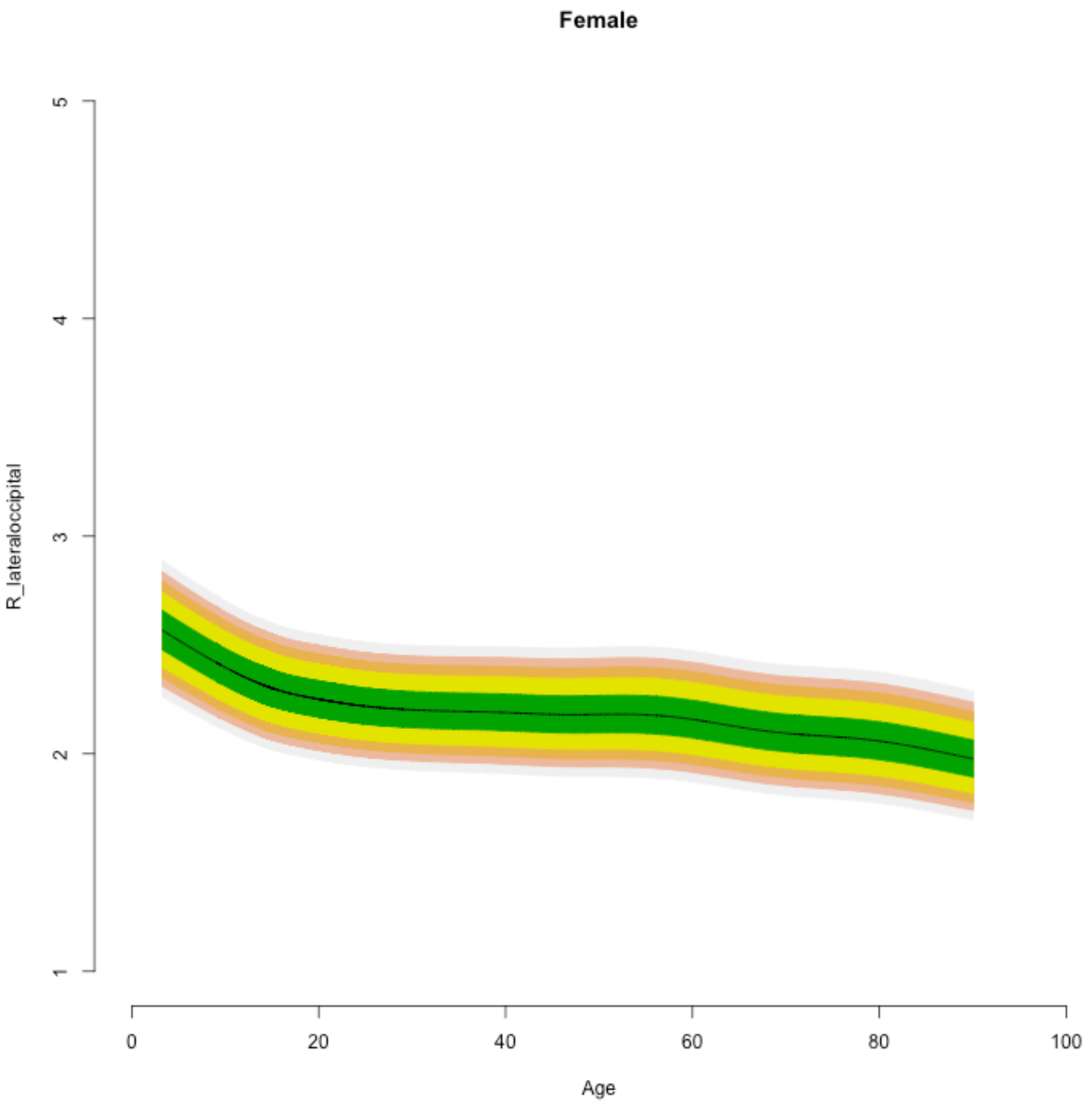

Female
R_lateraloccipital
1
2
3
4
5
0
20
40
60
80
100
Age

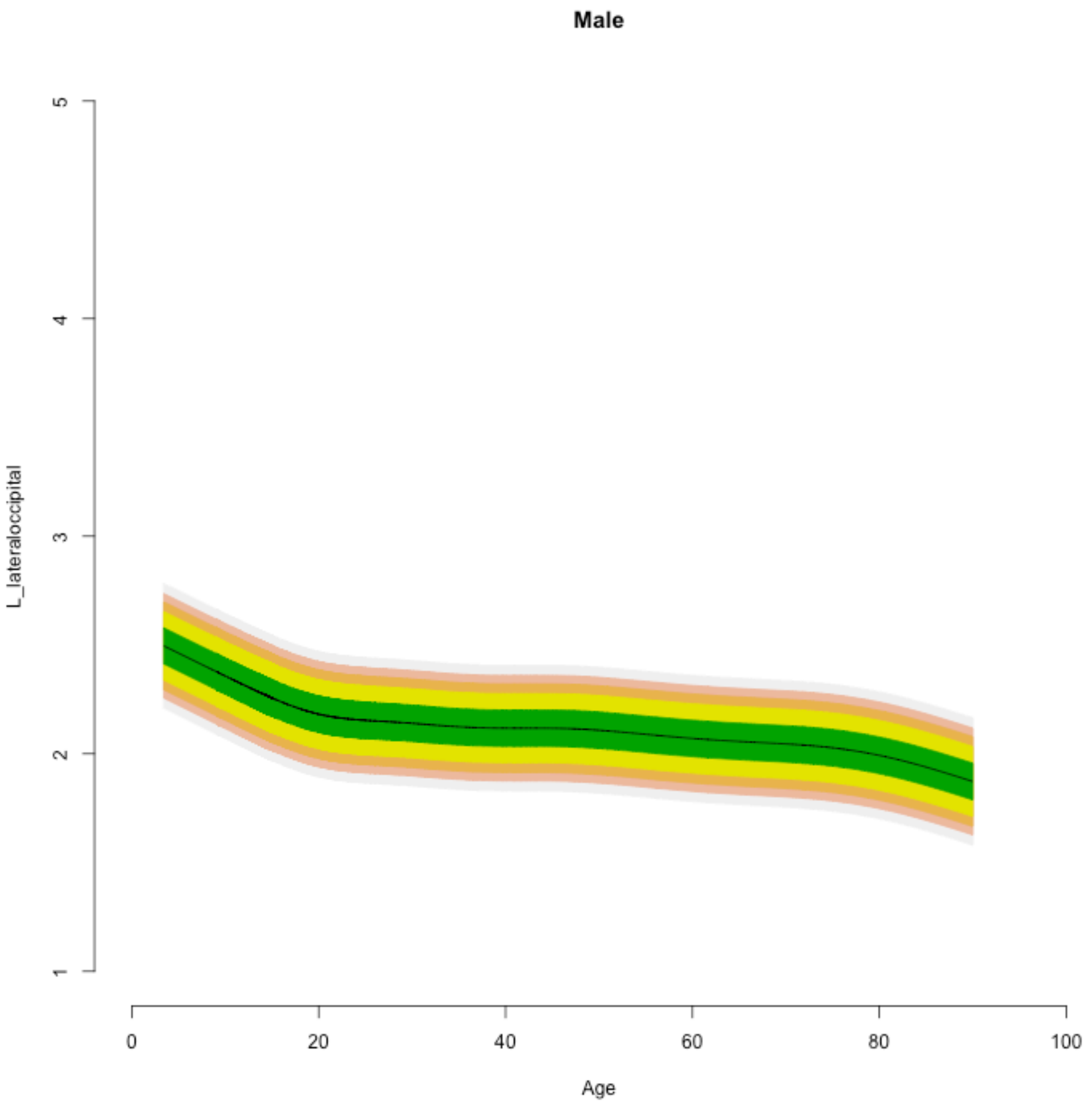

Male
L_lateraloccipital
1
2
3
4
5
0
20
40
60
80
100
Age

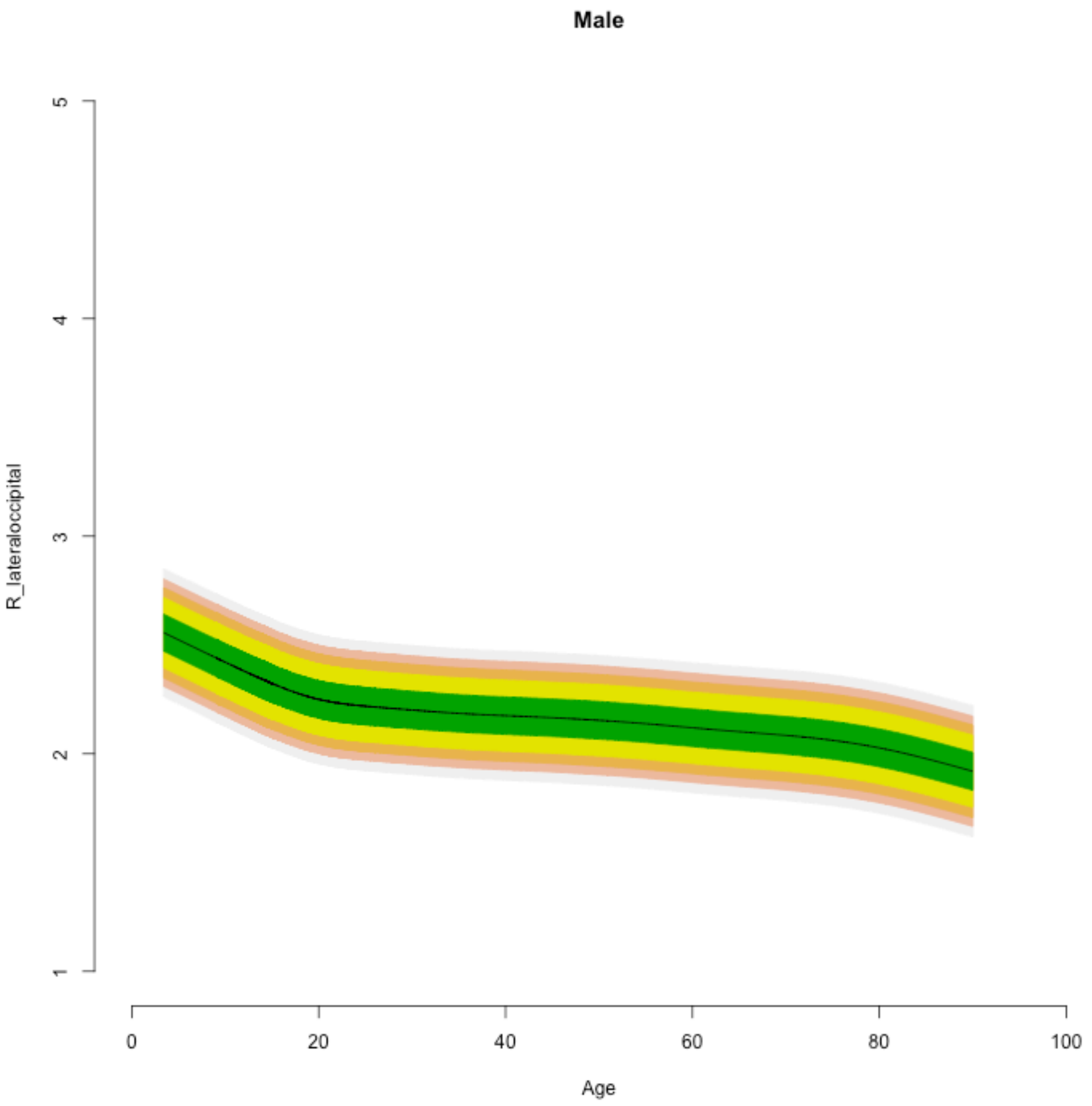

Male
R_lateraloccipital
1
2
3
4
5
0
20
40
60
80
100
Age

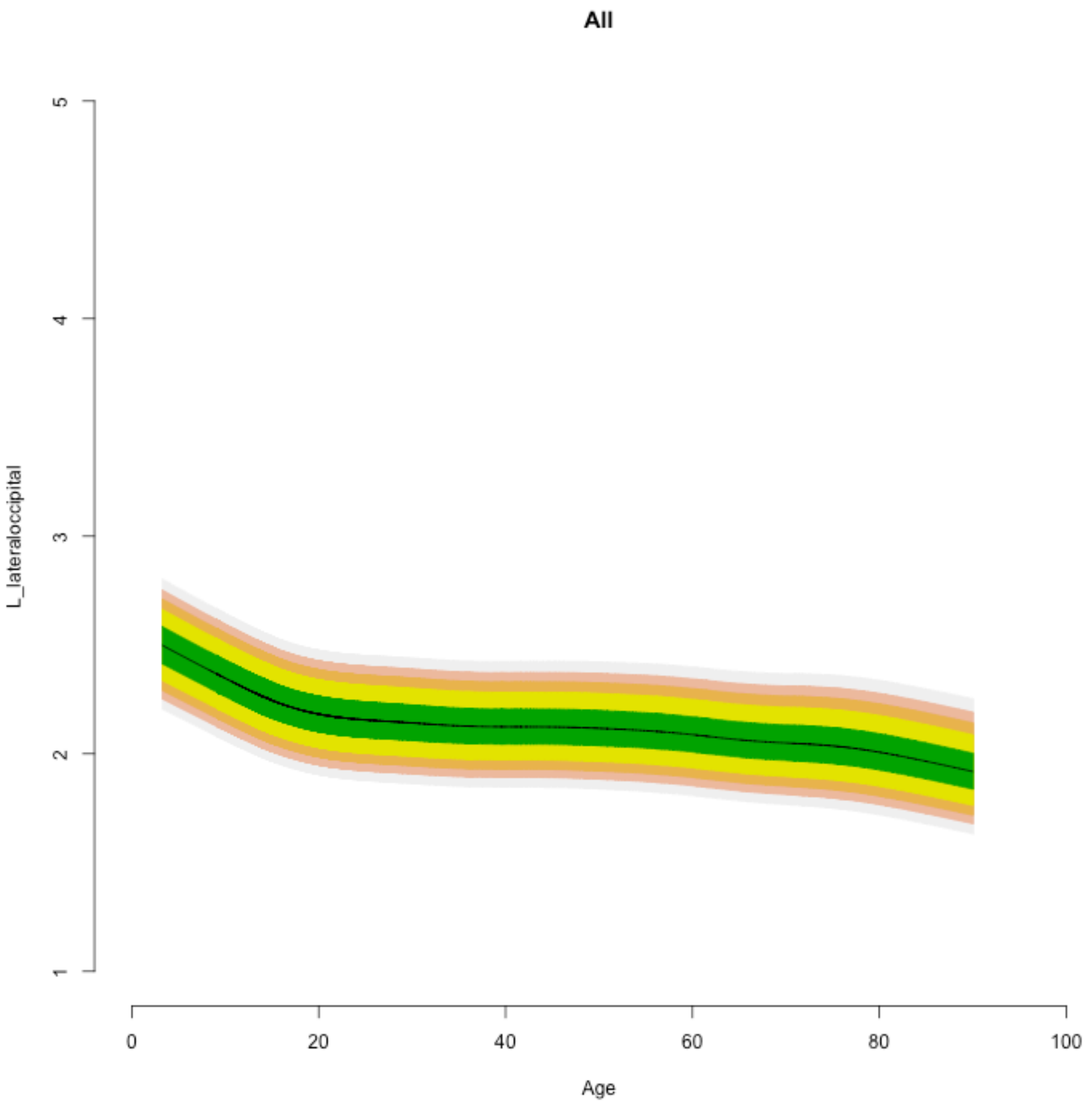
All
L_lateraloccipital
1
2
3
4
5
0
20
40
60
80
100
Age

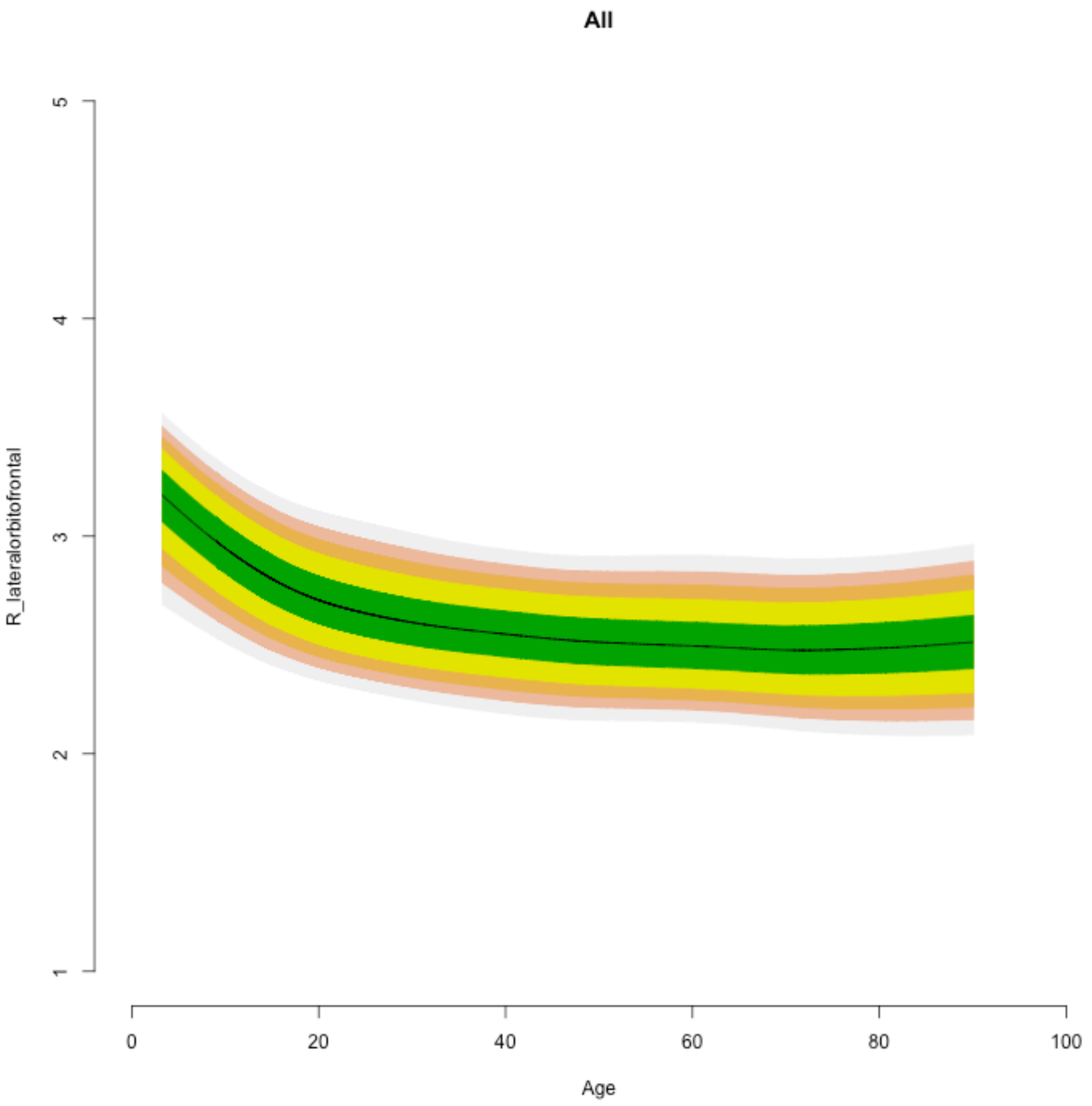
All
R_lateralorbitofrontal
5
4
3
2
1
0
20
40
60
80
100
Age

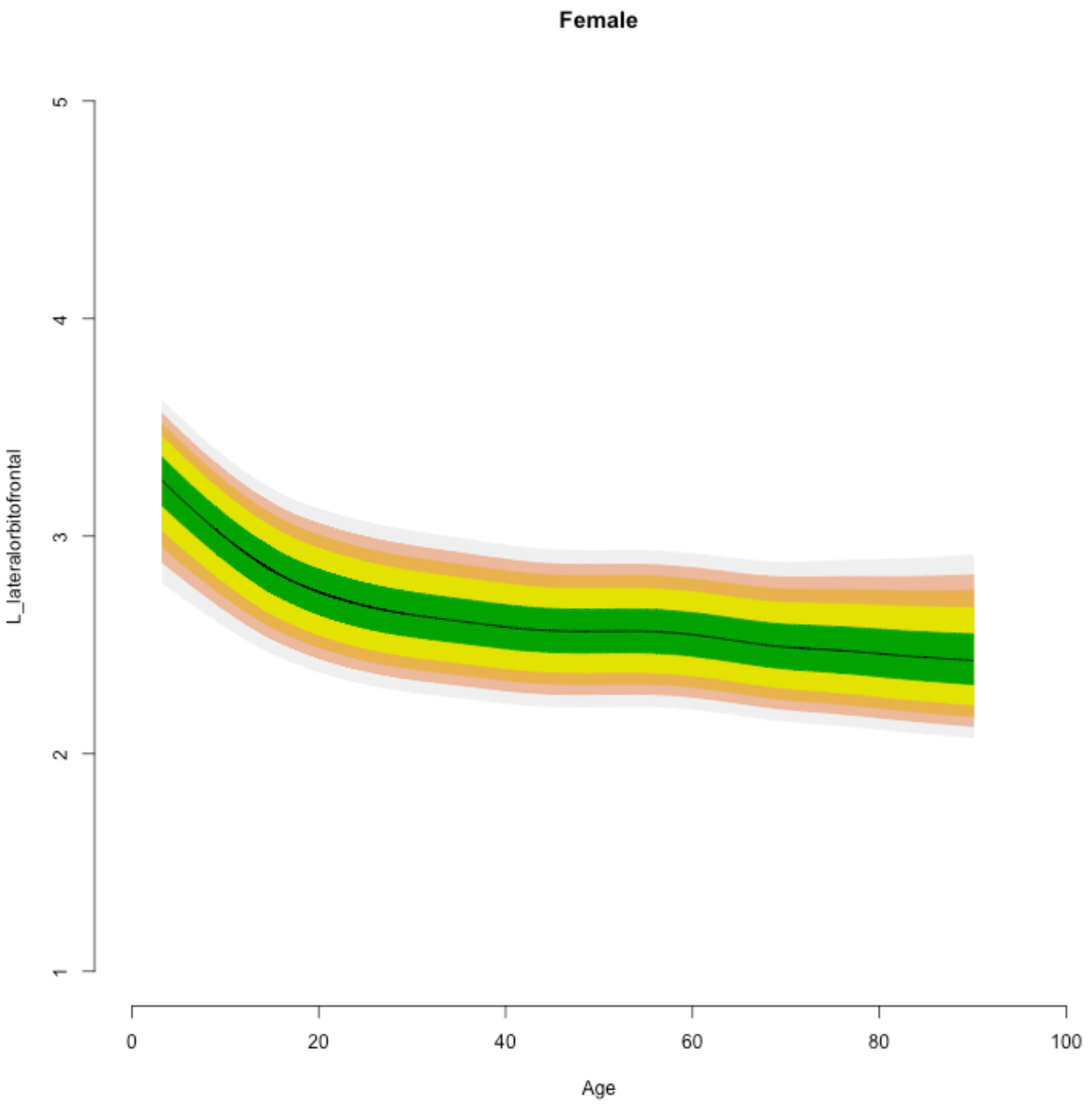

Female
L_lateralorbitofrontal
5
4
3
2
1
0
20
40
60
80
100
Age

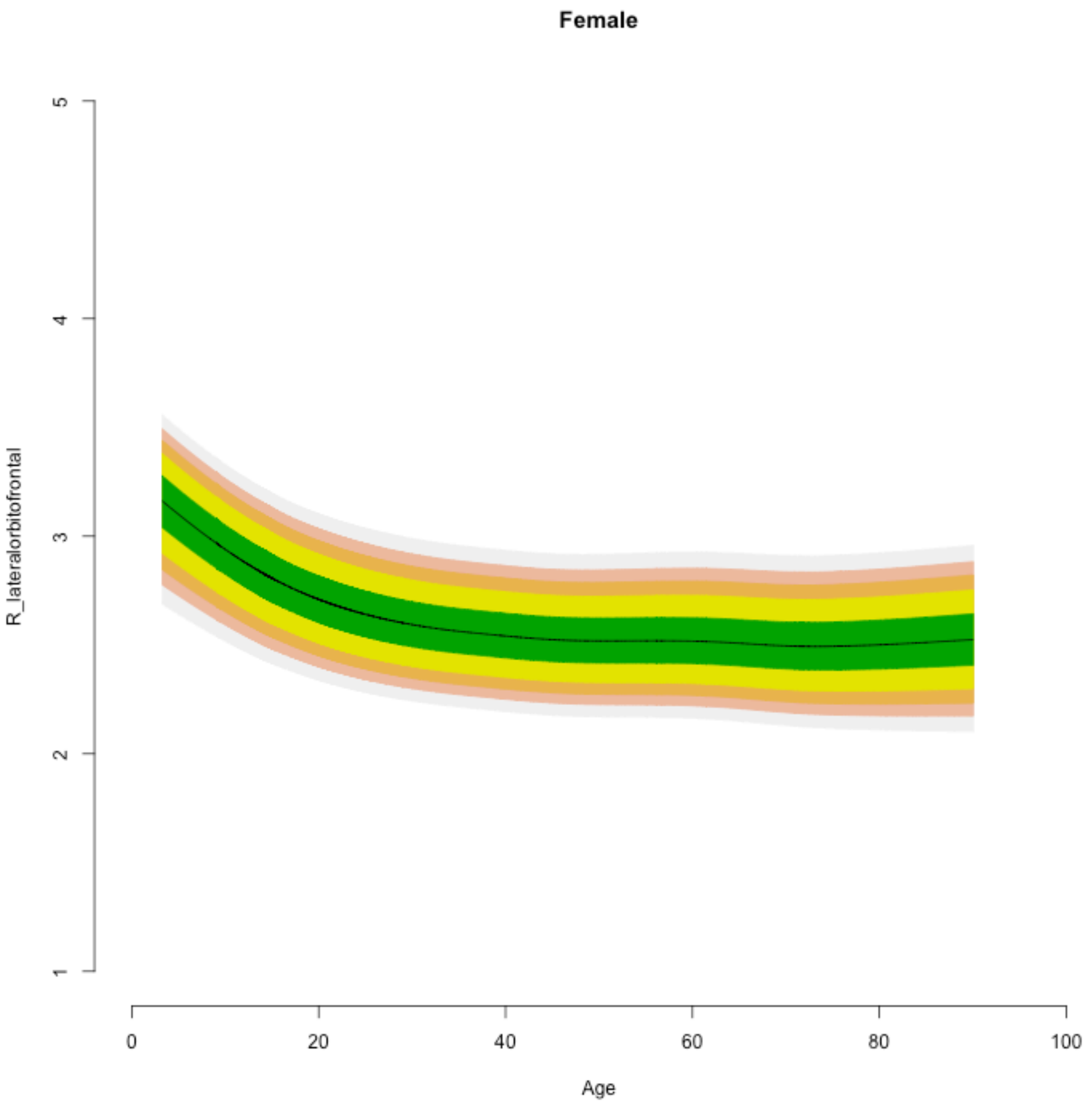
Female
R_lateralorbitofrontal
1
2
3
4
5
0
20
40
60
80
100
Age

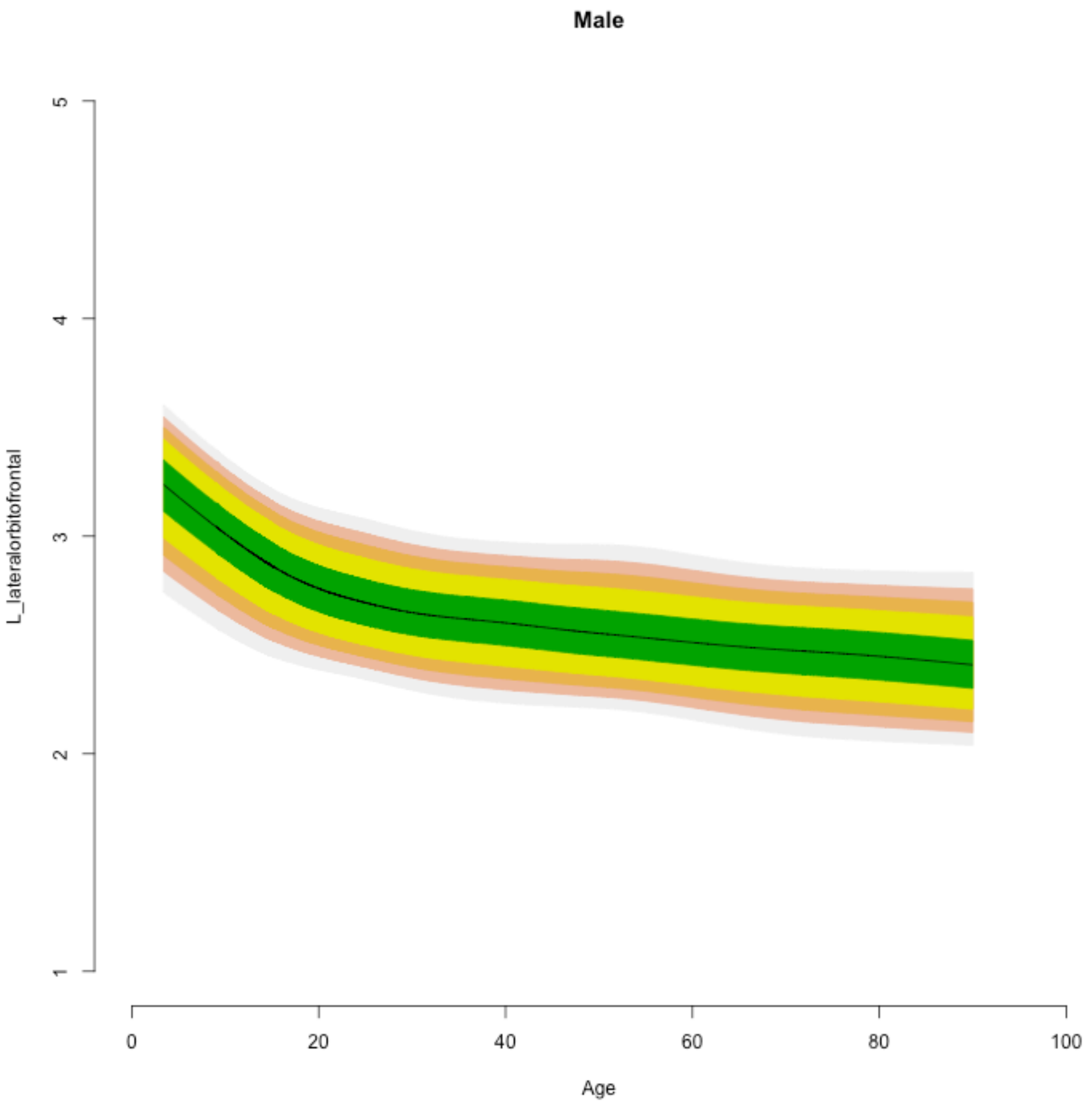
Male
L_lateralorbitofrontal
1
2
3
4
5
0
20
40
60
80
100
Age

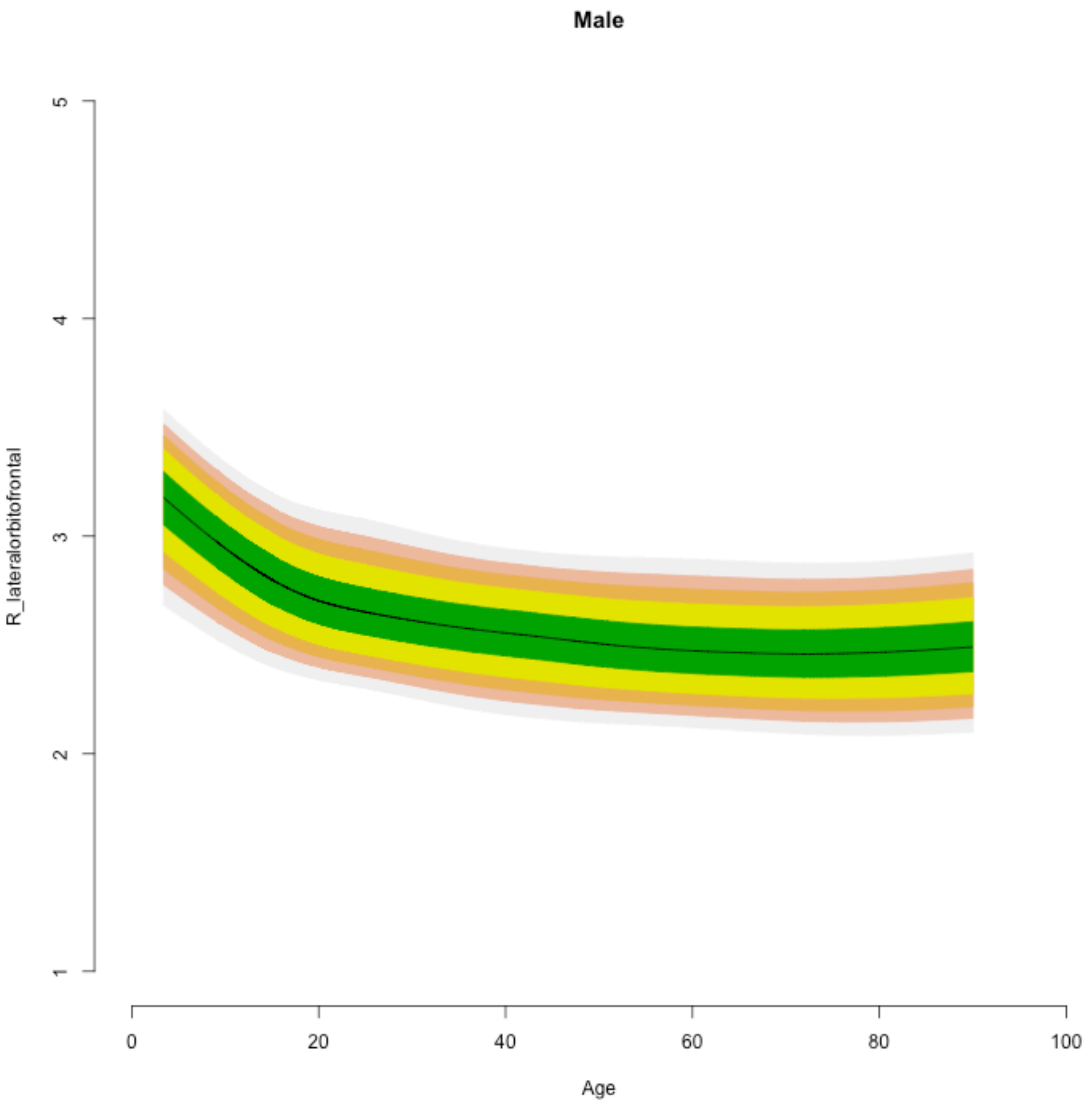

Male
R_lateralorbitofrontal
1
2
3
4
5
0
20
40
60
80
100
Age

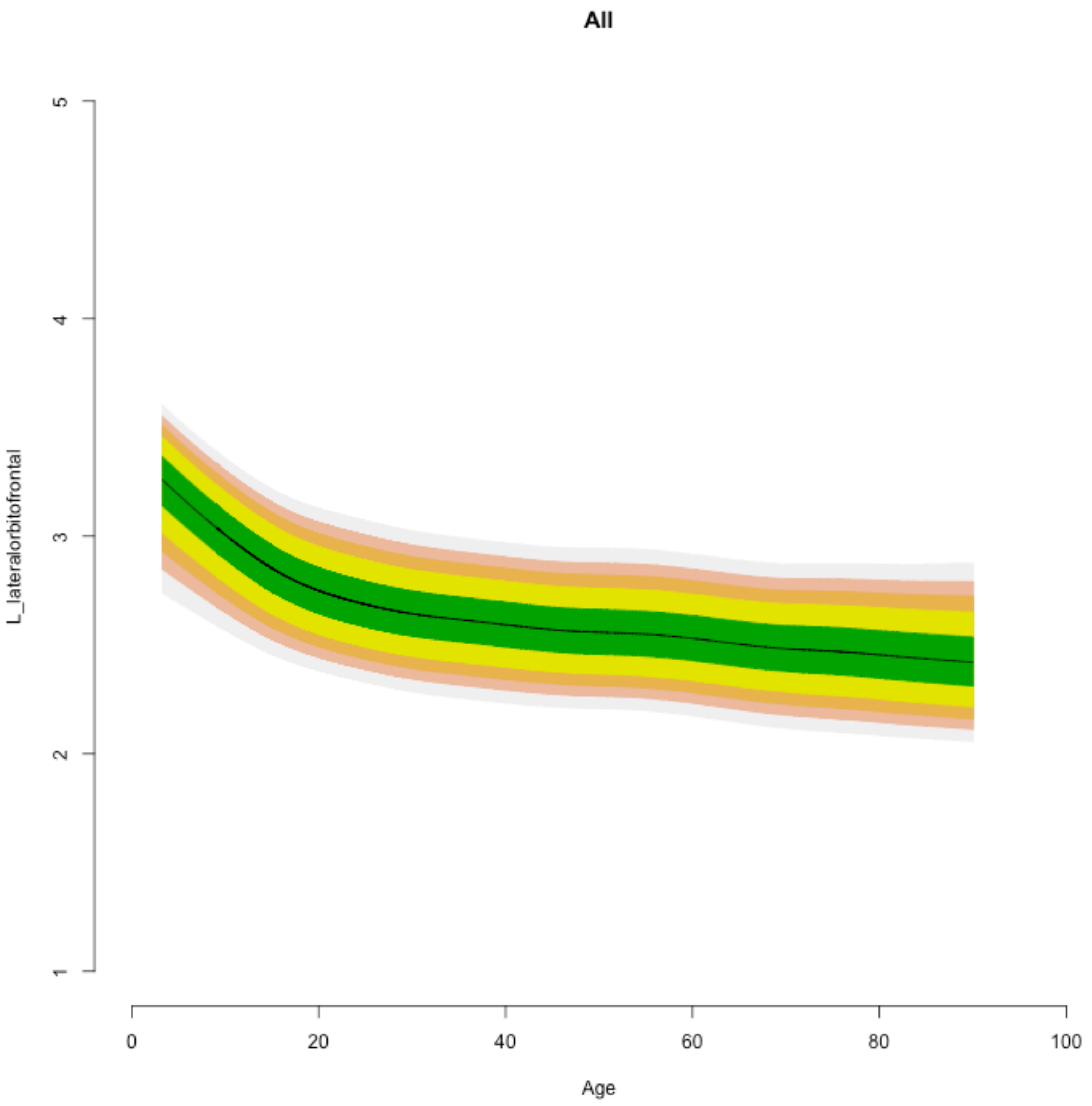
All
L_lateralorbitofrontal
5
4
3
2
1
0
20
40
60
80
100
Age

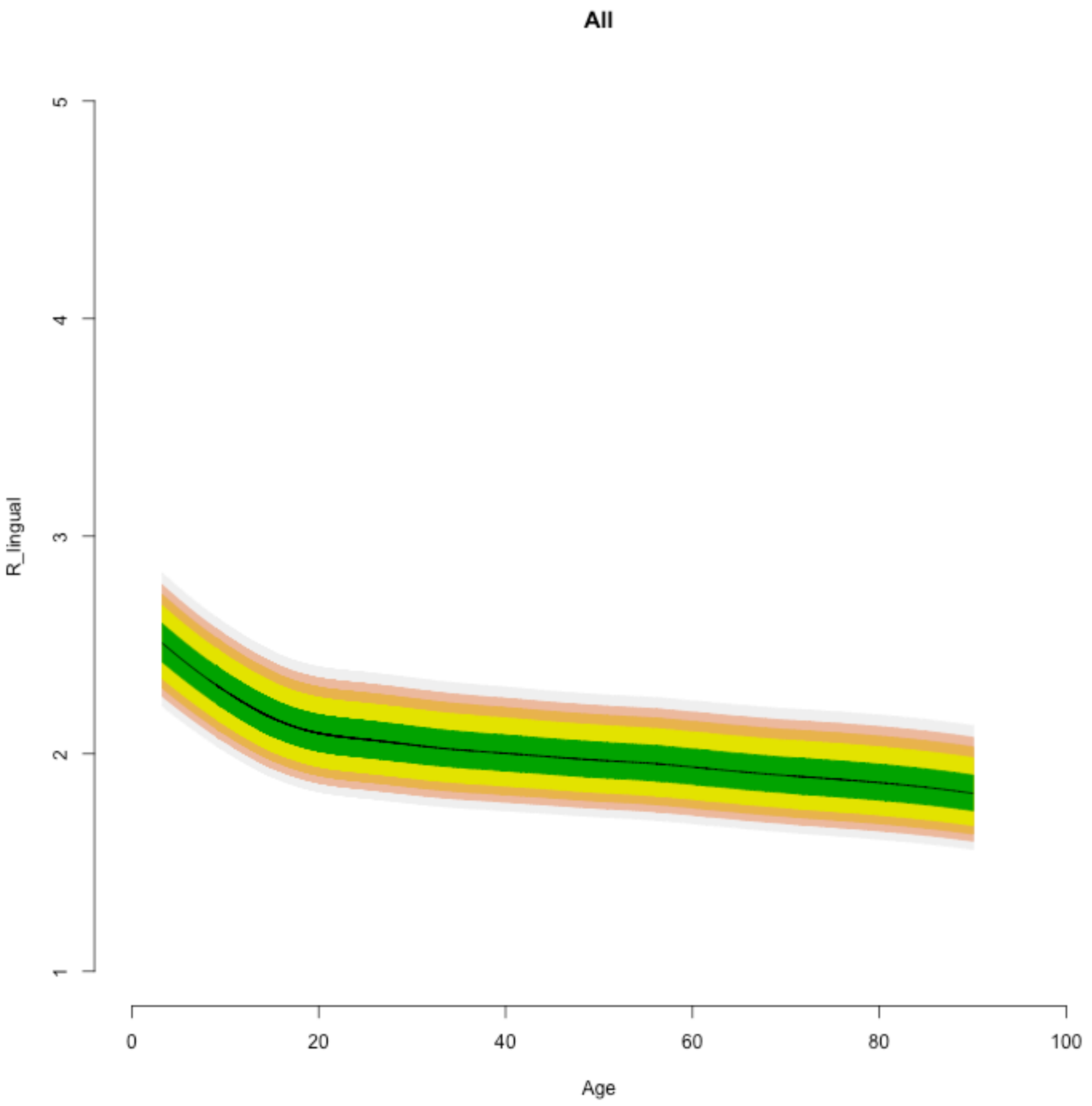
All
R_lingual
5
4
3
2
1
0
20
40
60
80
100
Age

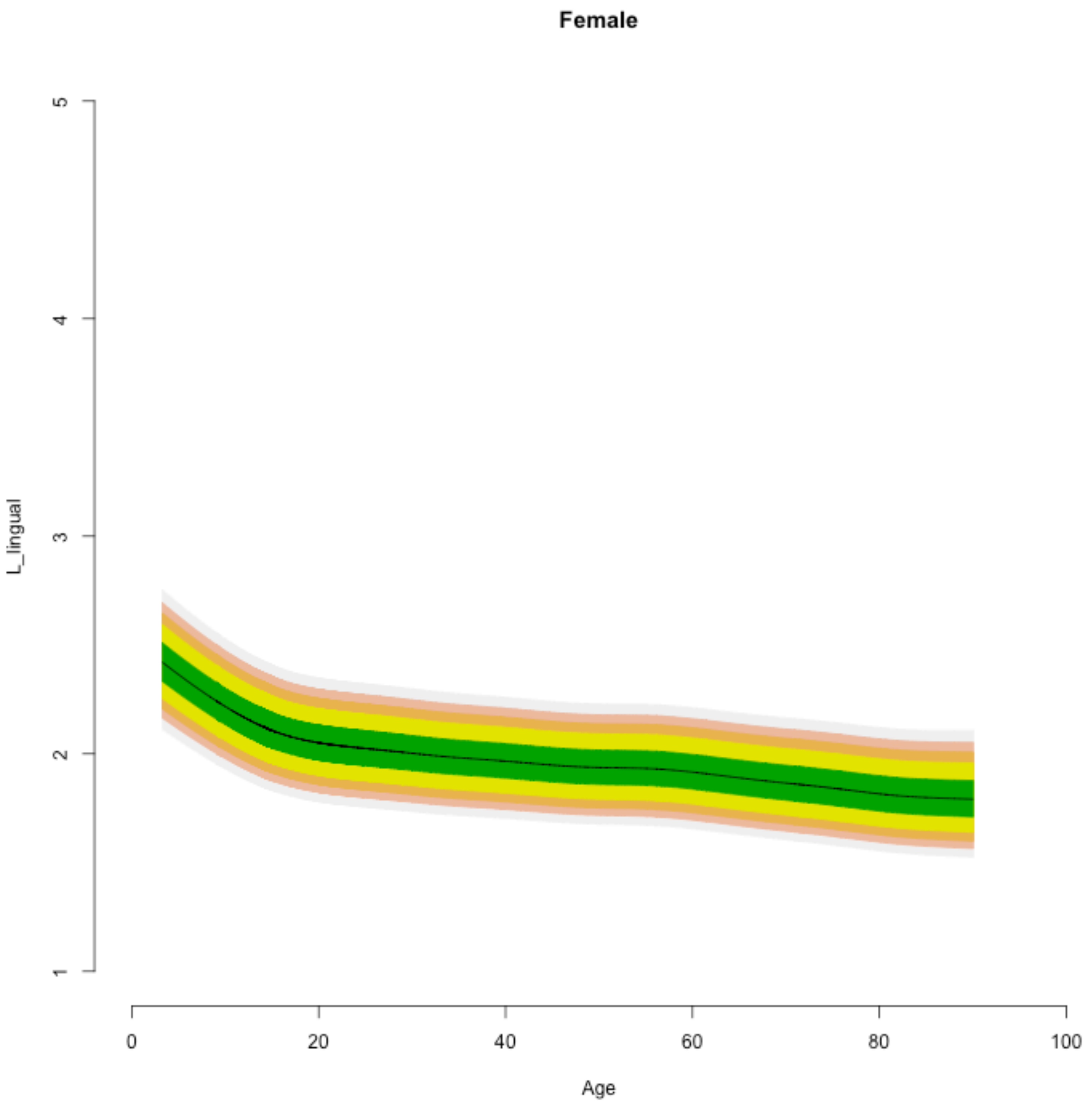
Female
L_lingual
1
2
3
4
5
0
20
40
60
80
100
Age

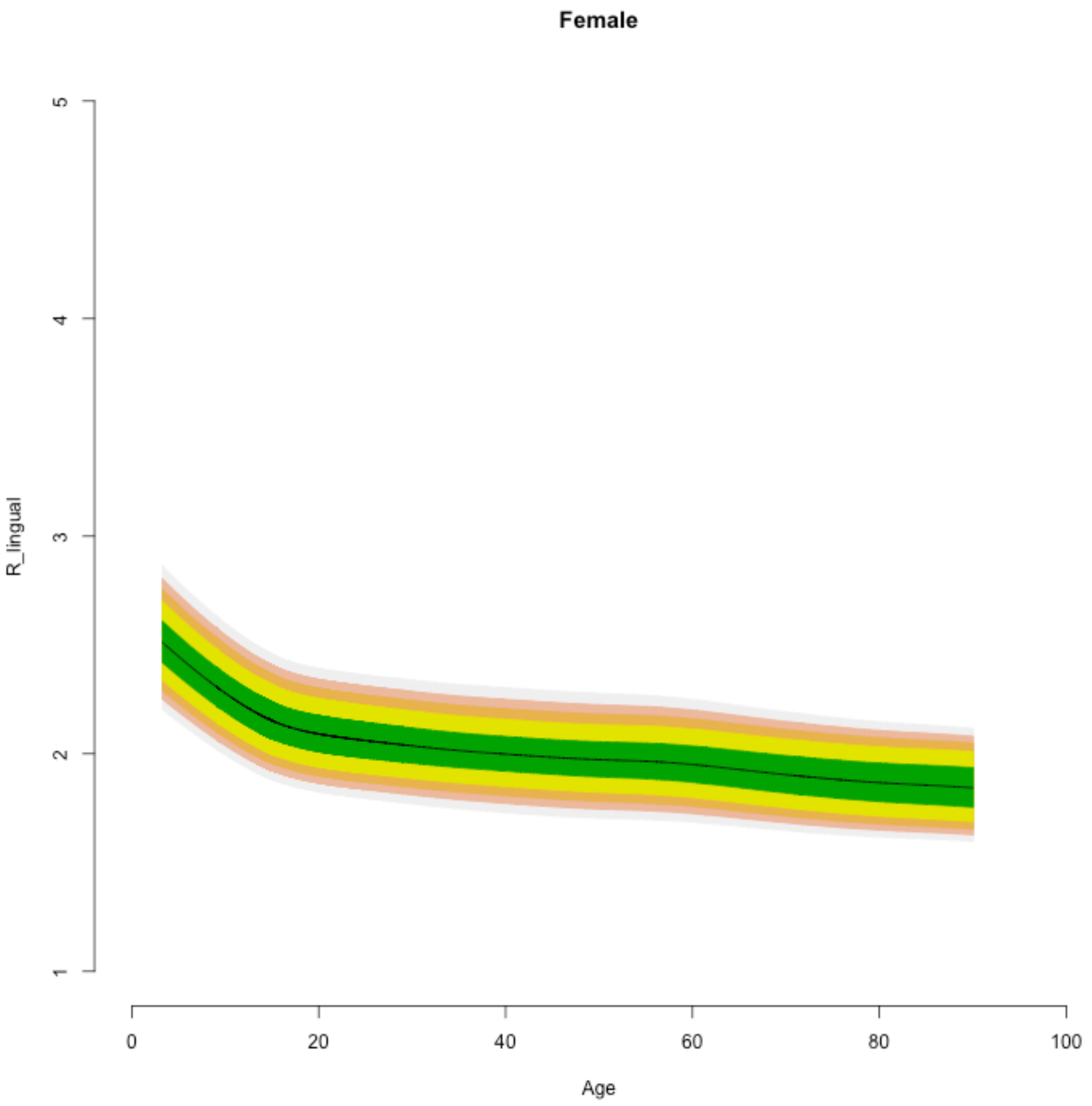

Female
R_lingual
1
2
3
4
5
0
20
40
60
80
100
Age

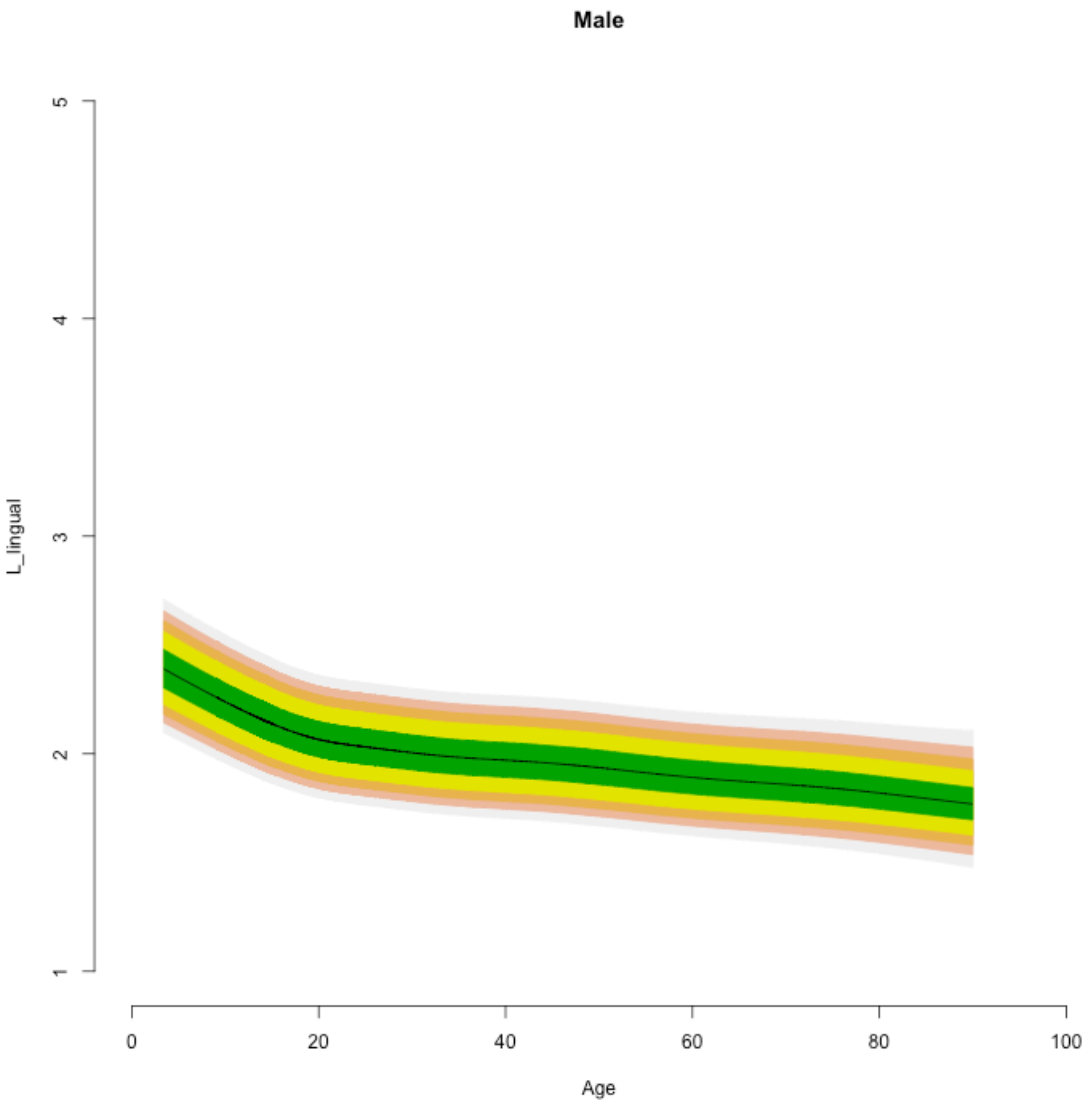

Male
L_lingual
1
2
3
4
5
0
20
40
60
80
100
Age

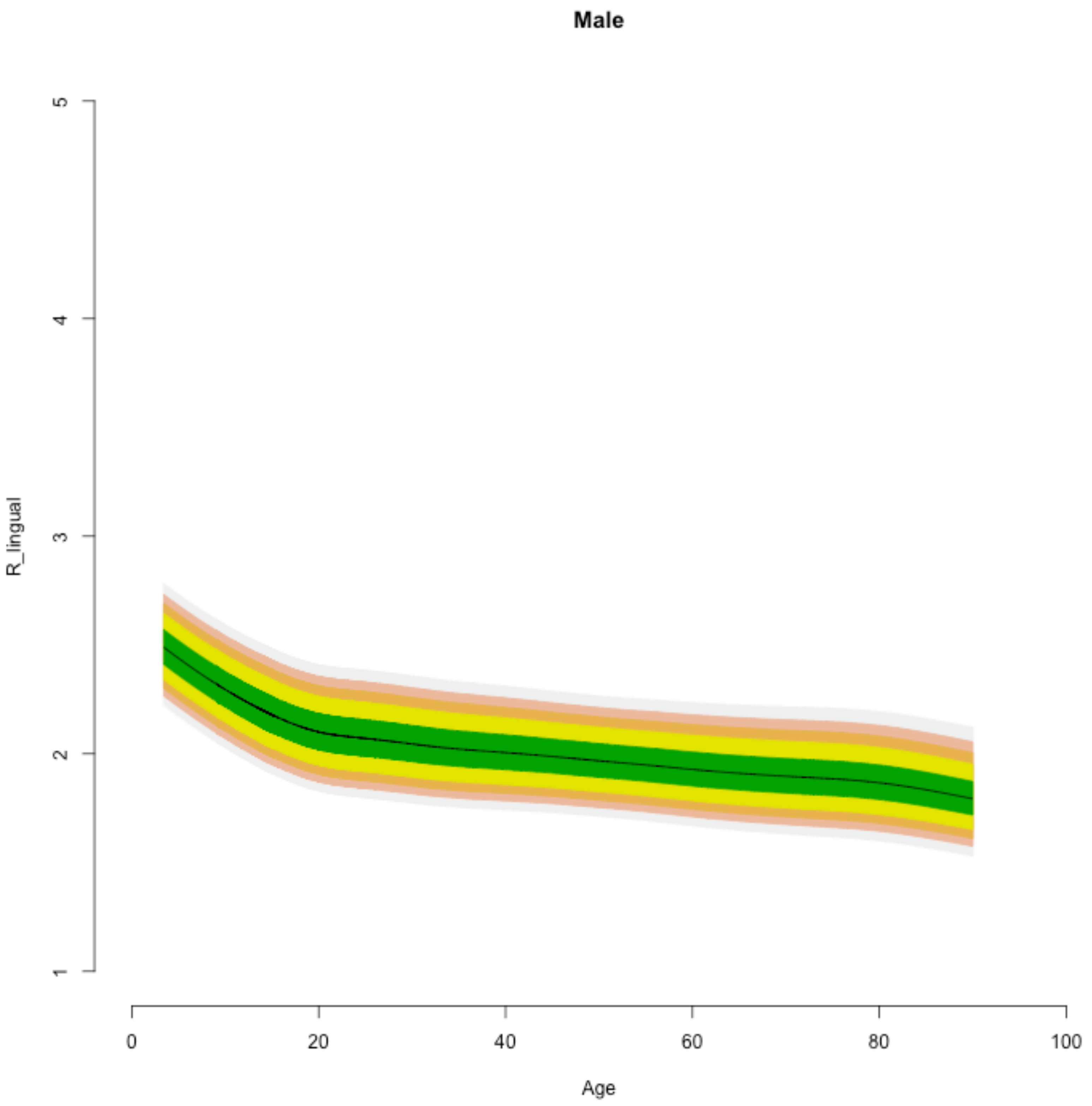

Male
R_lingual
5
4
3
2
1
0
20
40
60
80
100
Age

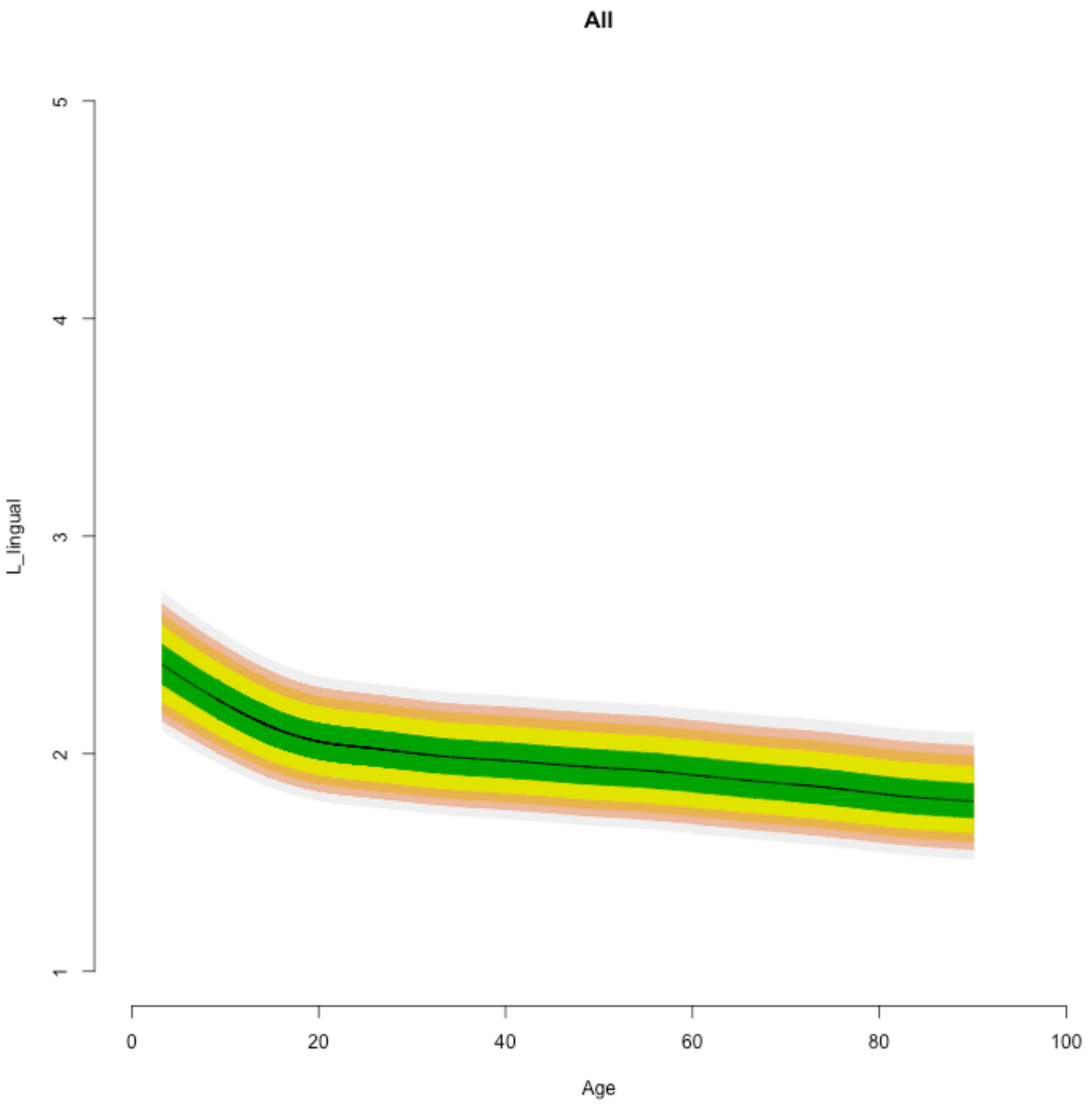

All
L_lingual
1
2
3
4
5
0
20
40
60
80
100
Age

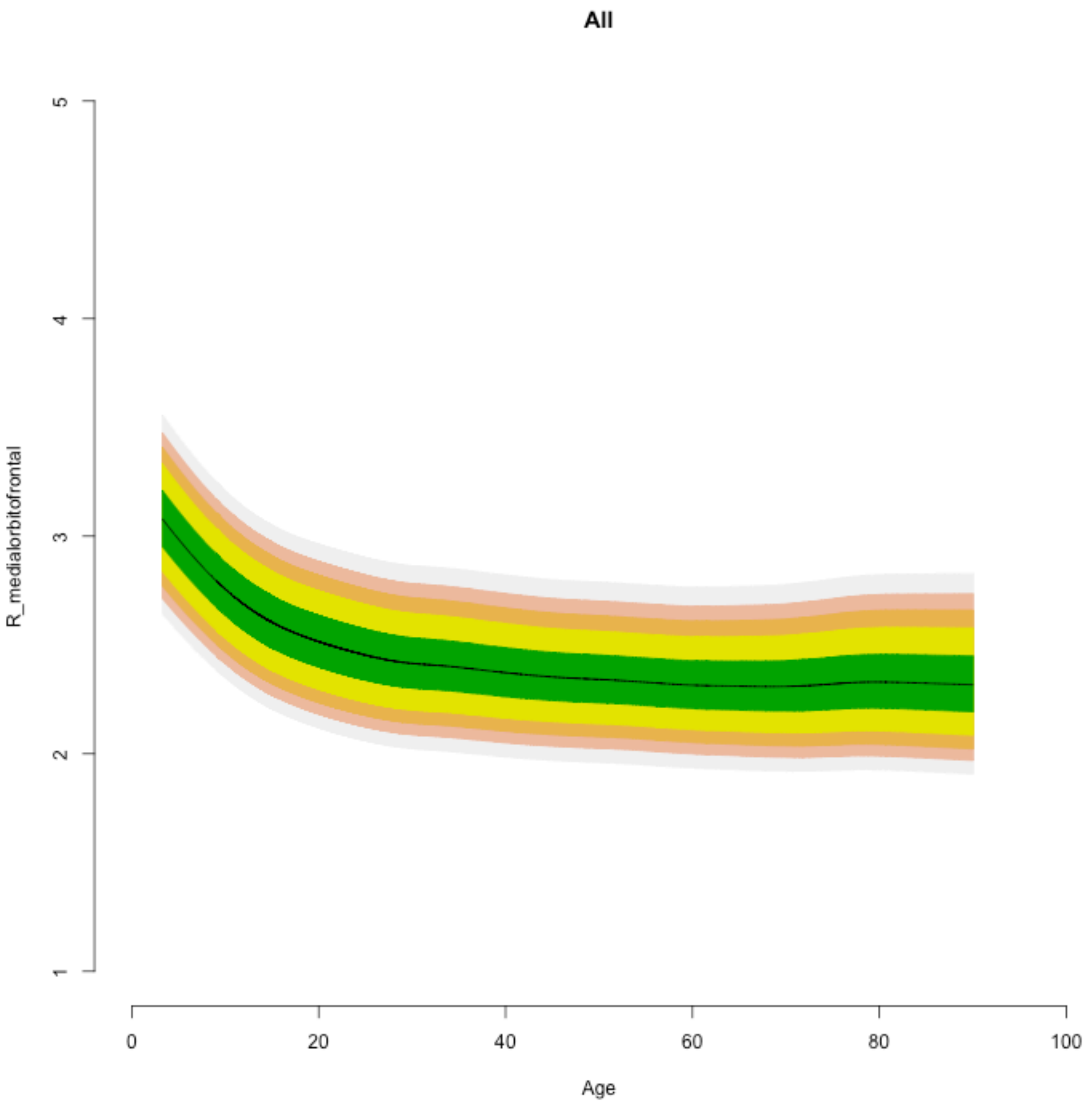
All
R_medialorbitofrontal
1
2
3
4
5
0
20
40
60
80
100
Age

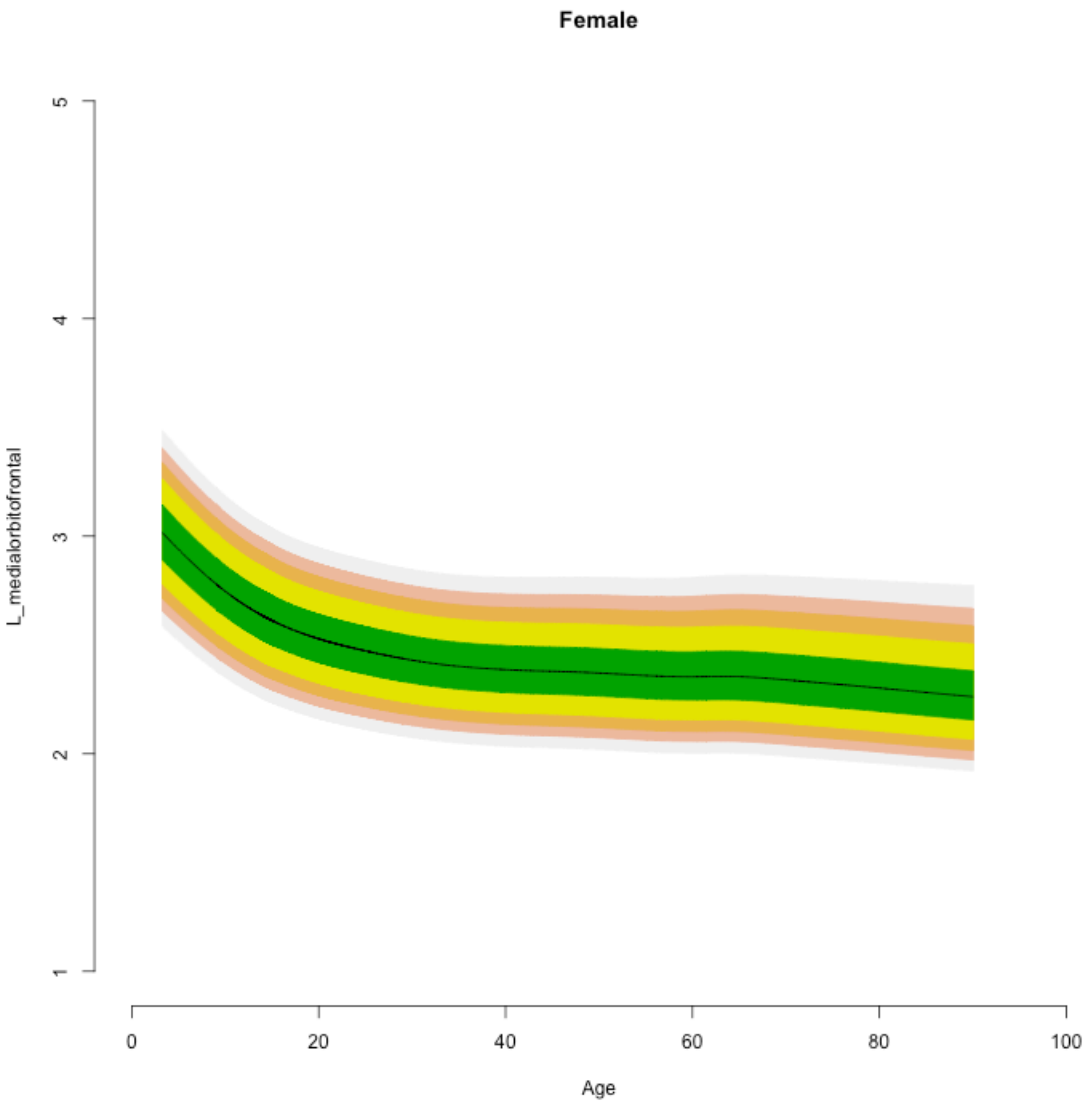
Female
L_medialorbitofrontal
1
2
3
4
5
0
20
40
60
80
100
Age

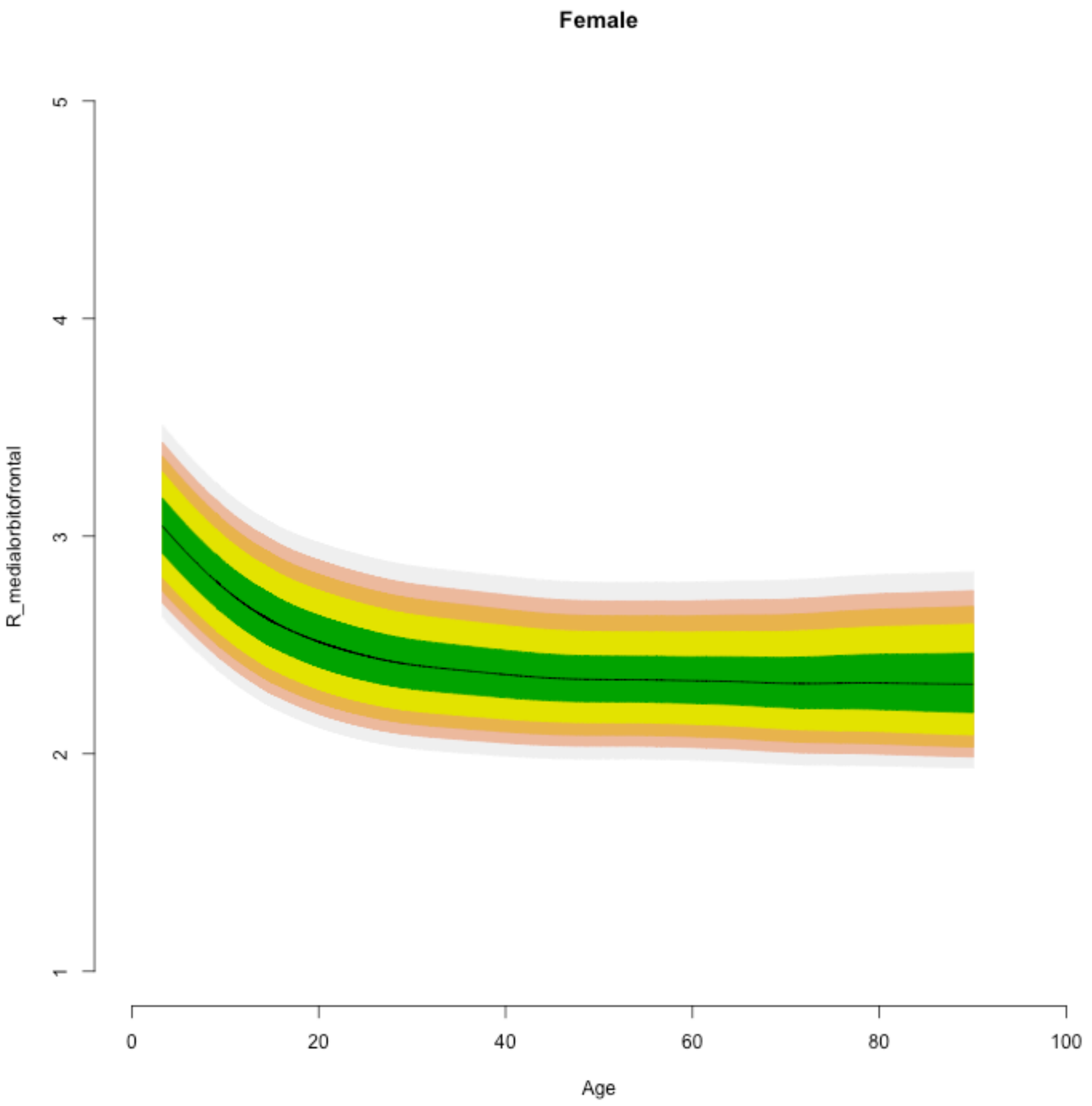
Female
R_medialorbitofrontal
5
4
3
2
1
0
20
40
60
80
100
Age

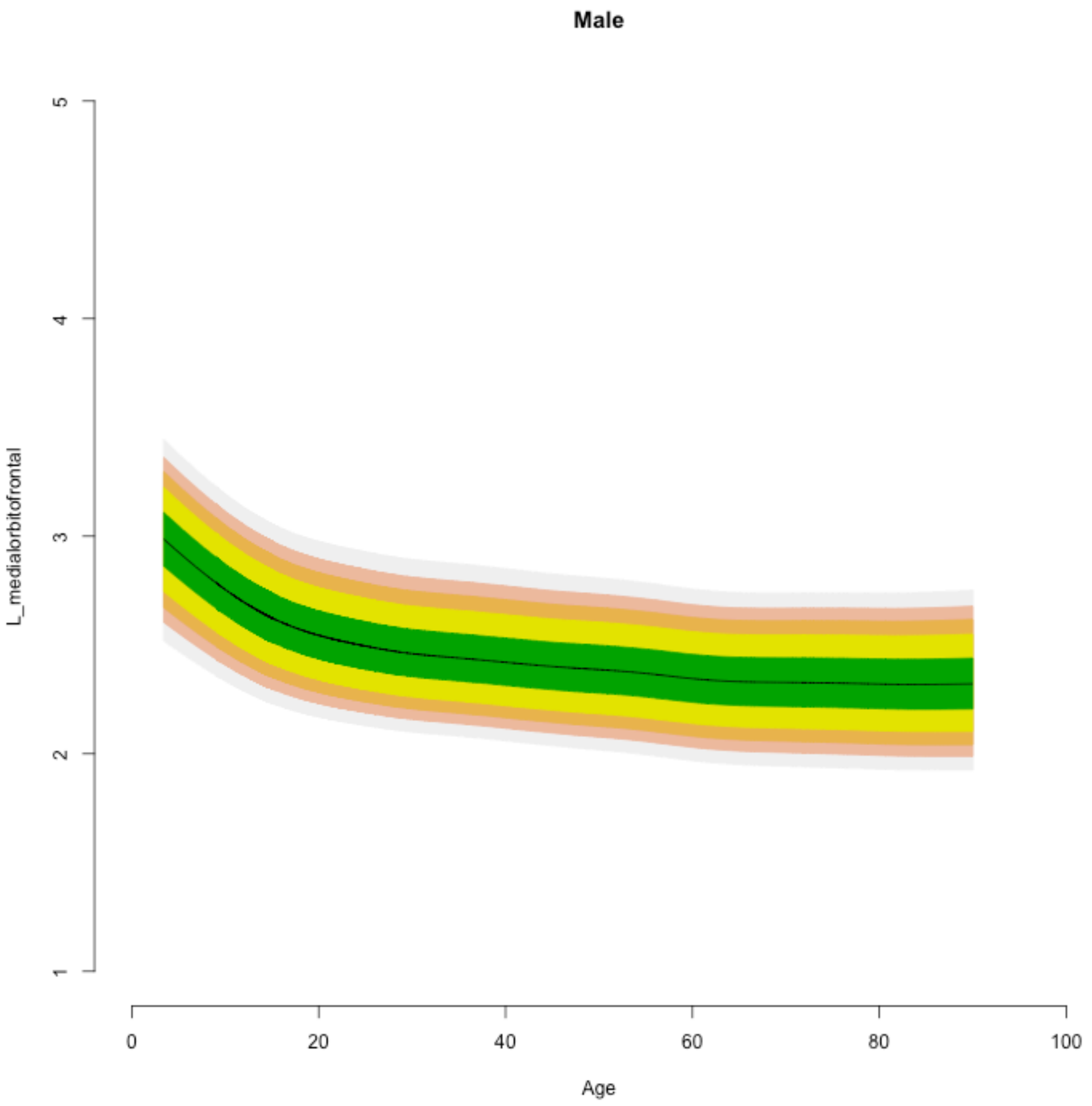

Male
L_medialorbitofrontal
1
2
3
4
5
0
20
40
60
80
100
Age

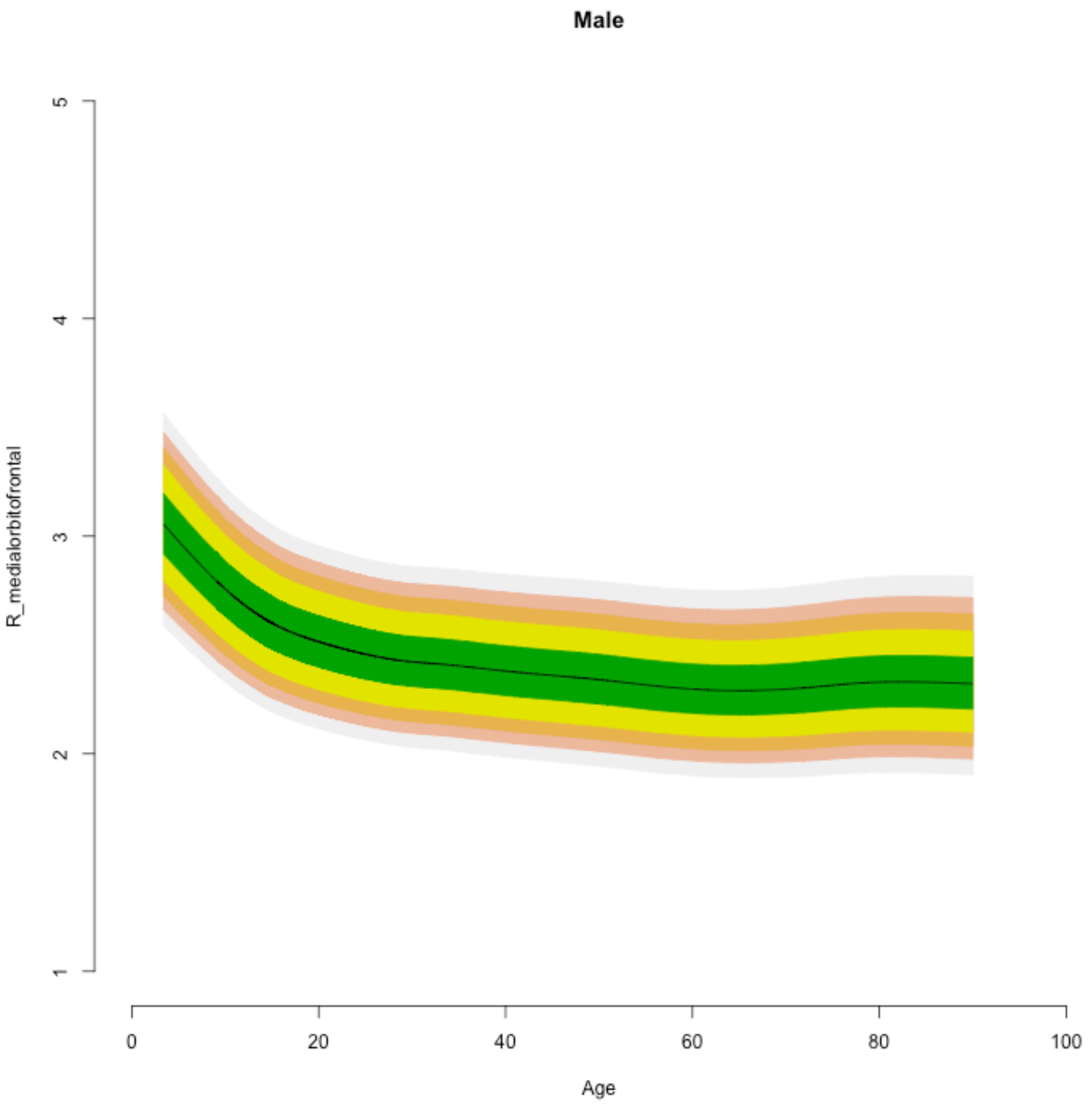

Male
R_medialorbitofrontal
1
2
3
4
5
0
20
40
60
80
100
Age

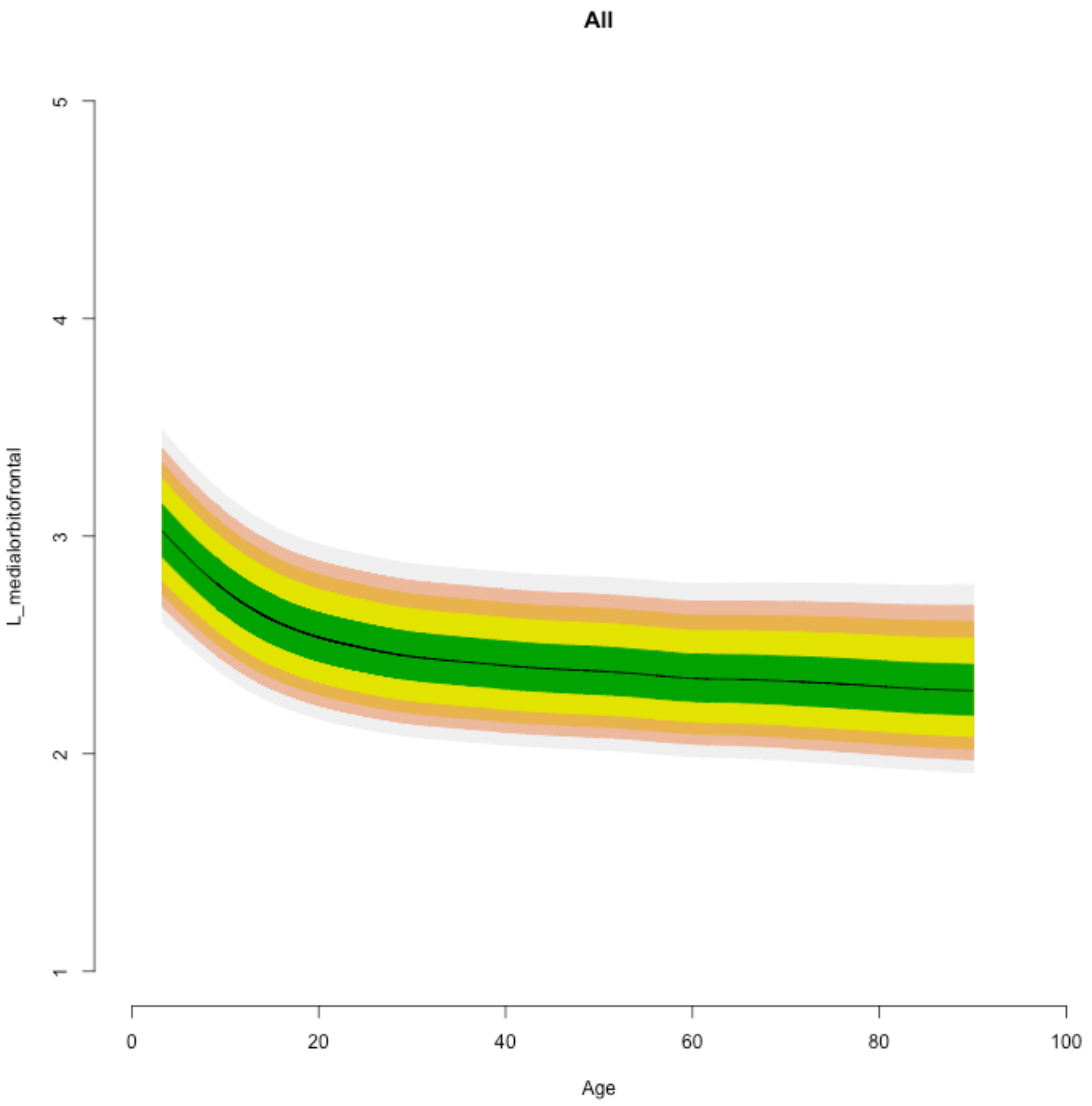

All
L_medialorbitofrontal
1
2
3
4
5
0
20
40
60
80
100
Age

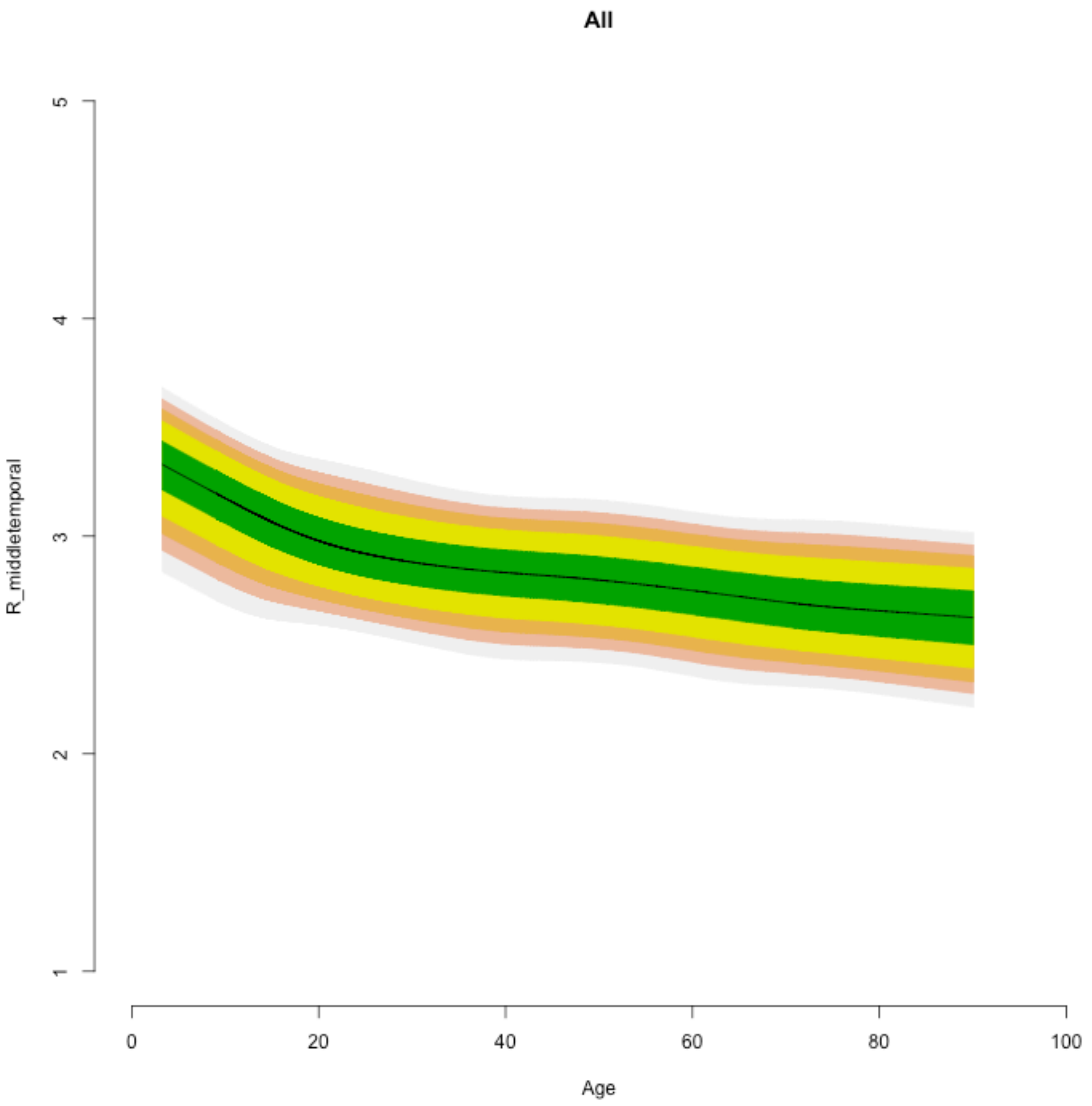
All
R_middletemporal
5
4
3
2
1
0
20
40
60
80
100
Age

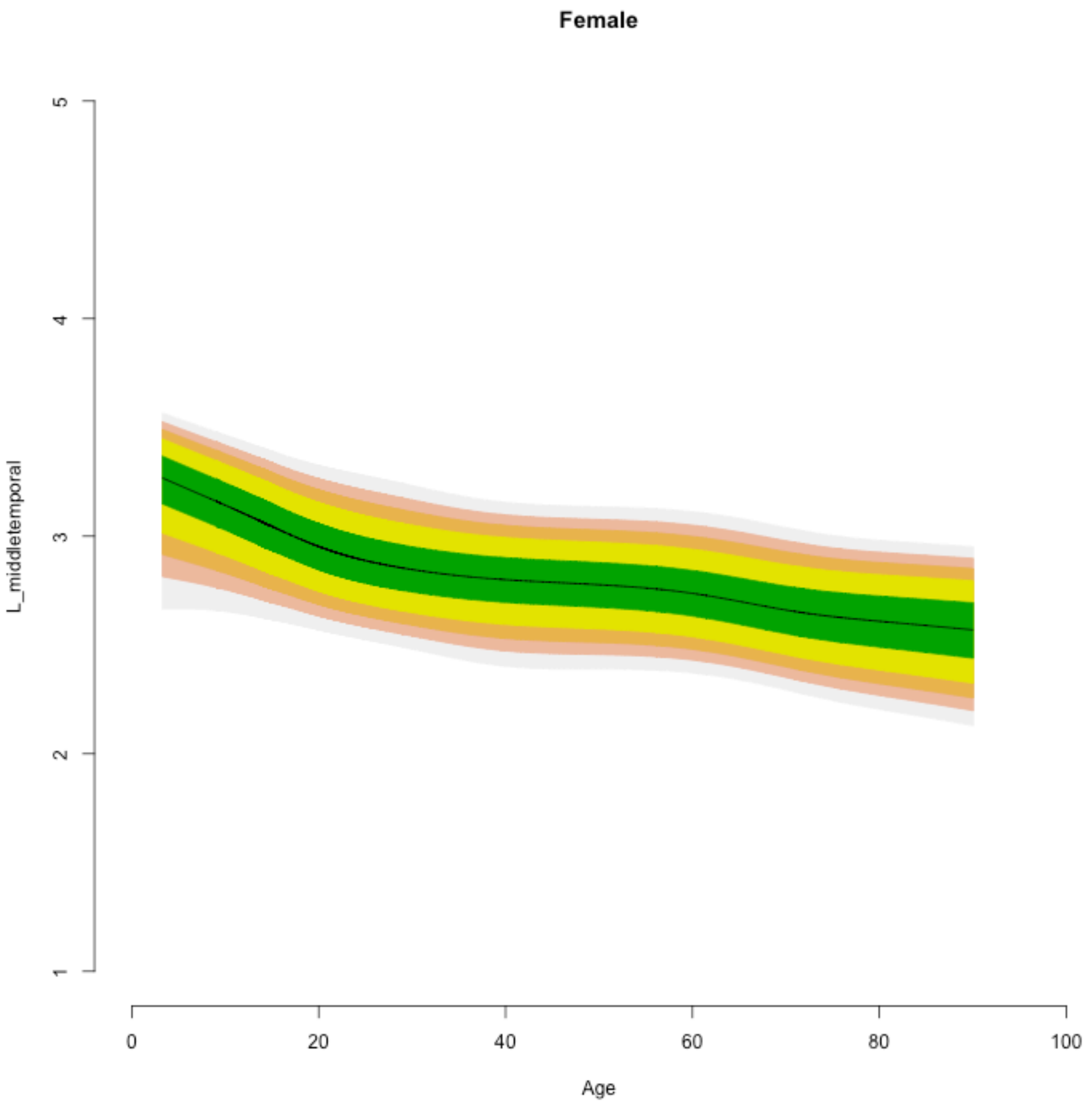

Female
L_middletemporal
5
4
3
2
1
0
20
40
60
80
100
Age

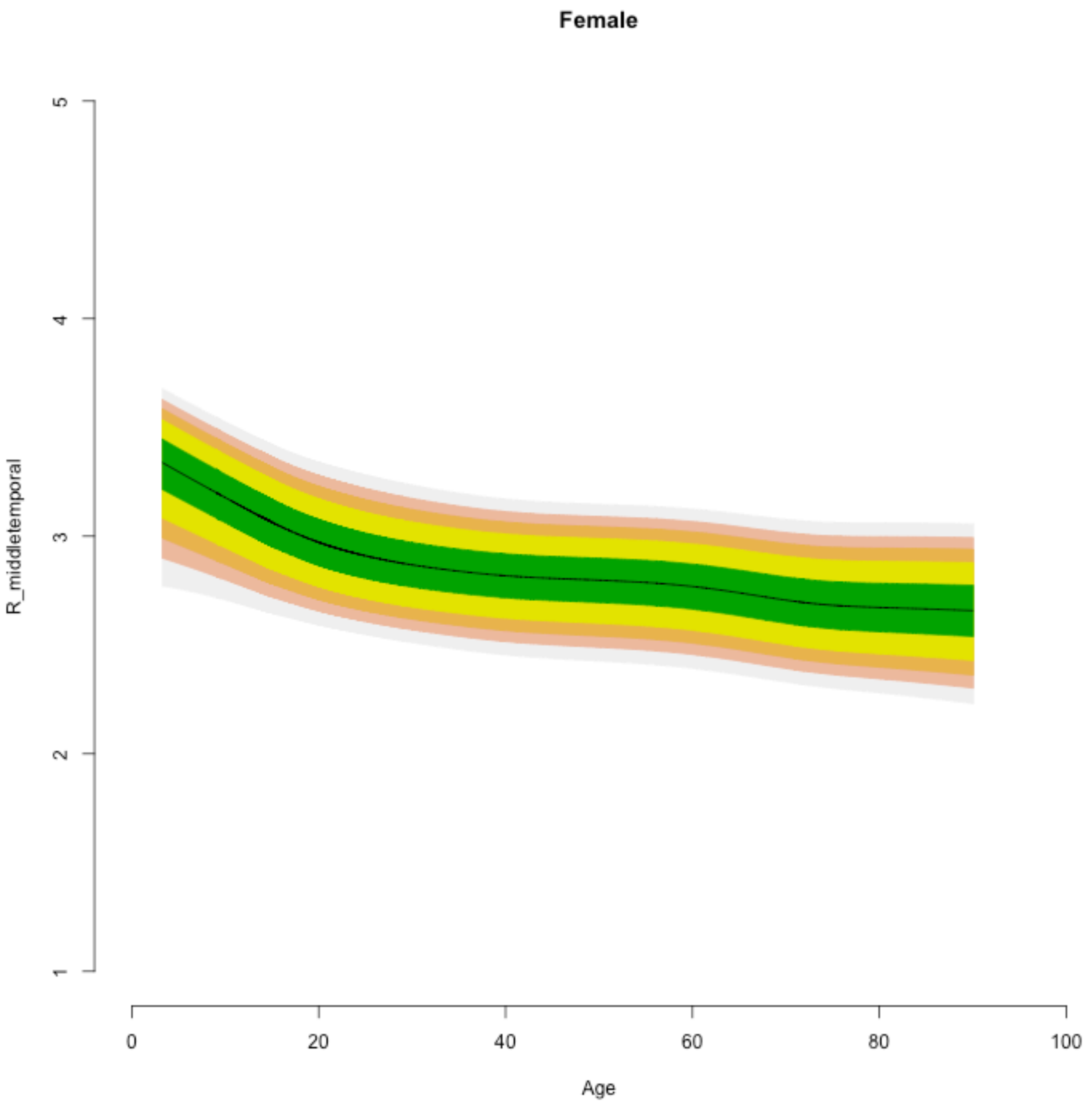
Female
R_middletemporal
1
2
3
4
5
0
20
40
60
80
100
Age

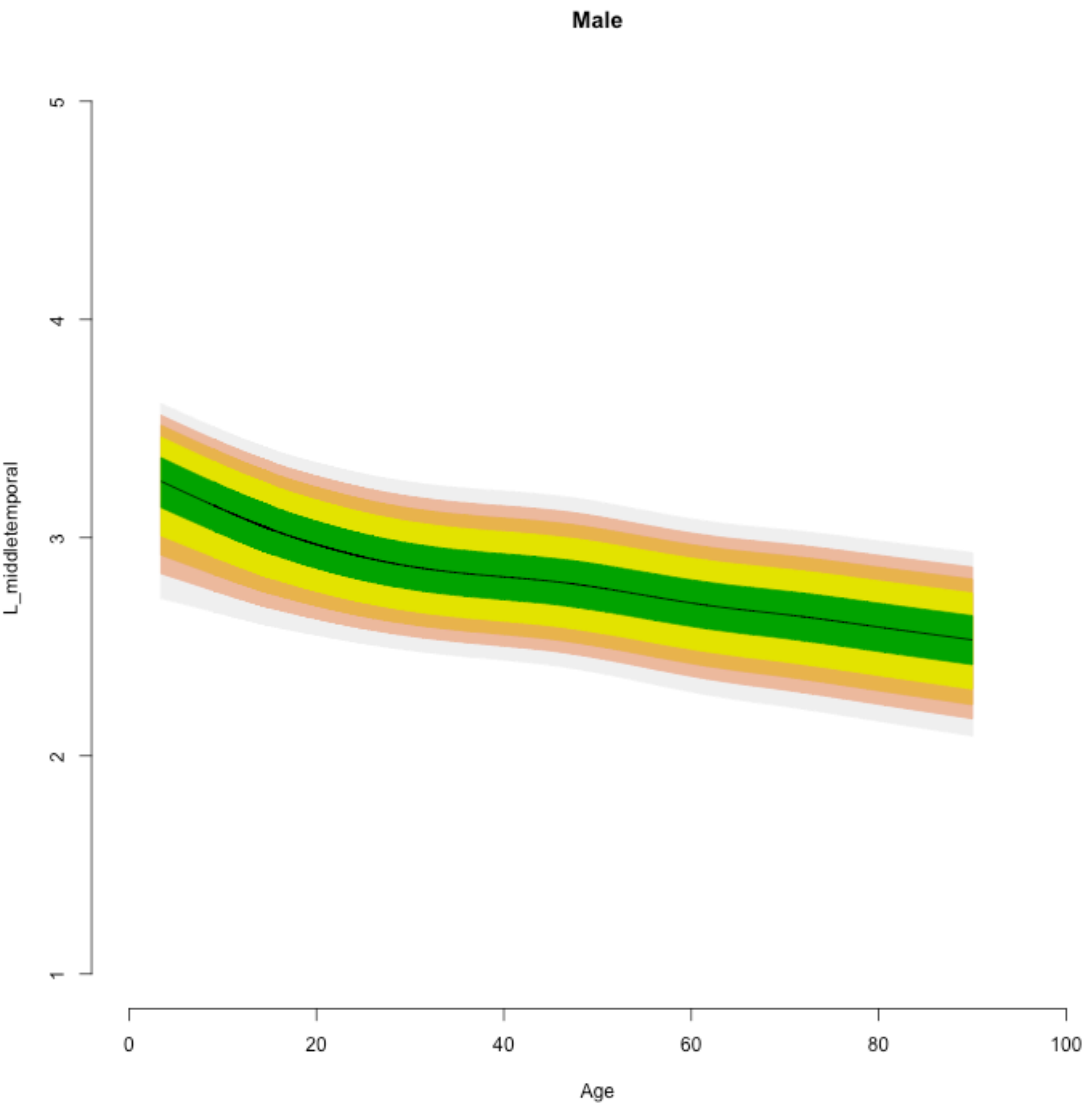

Male
L_middletemporal
5
4
3
2
1
0
20
40
60
80
100
Age

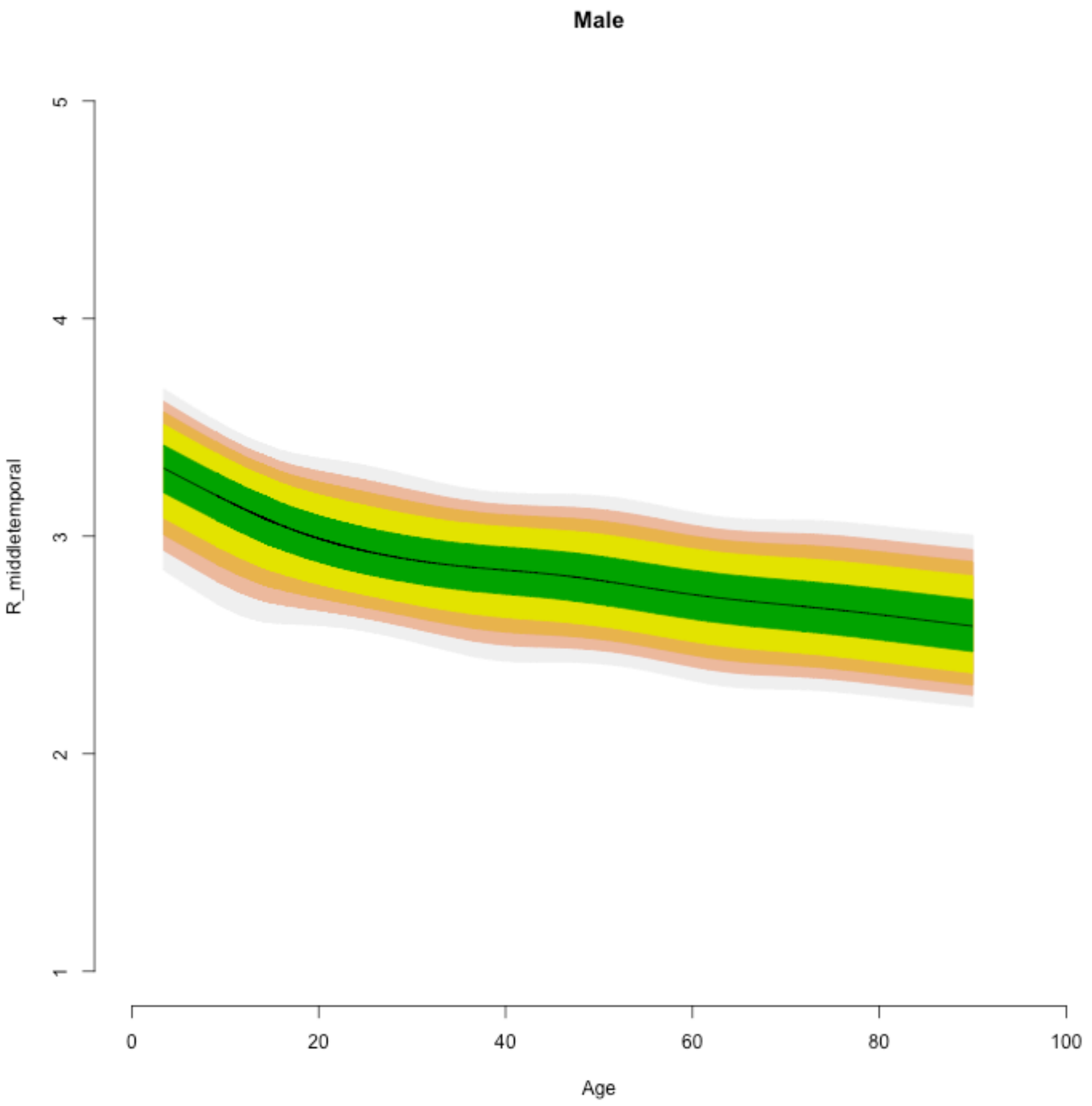

Male
R_middletemporal
5
4
3
2
1
0
20
40
60
80
100
Age

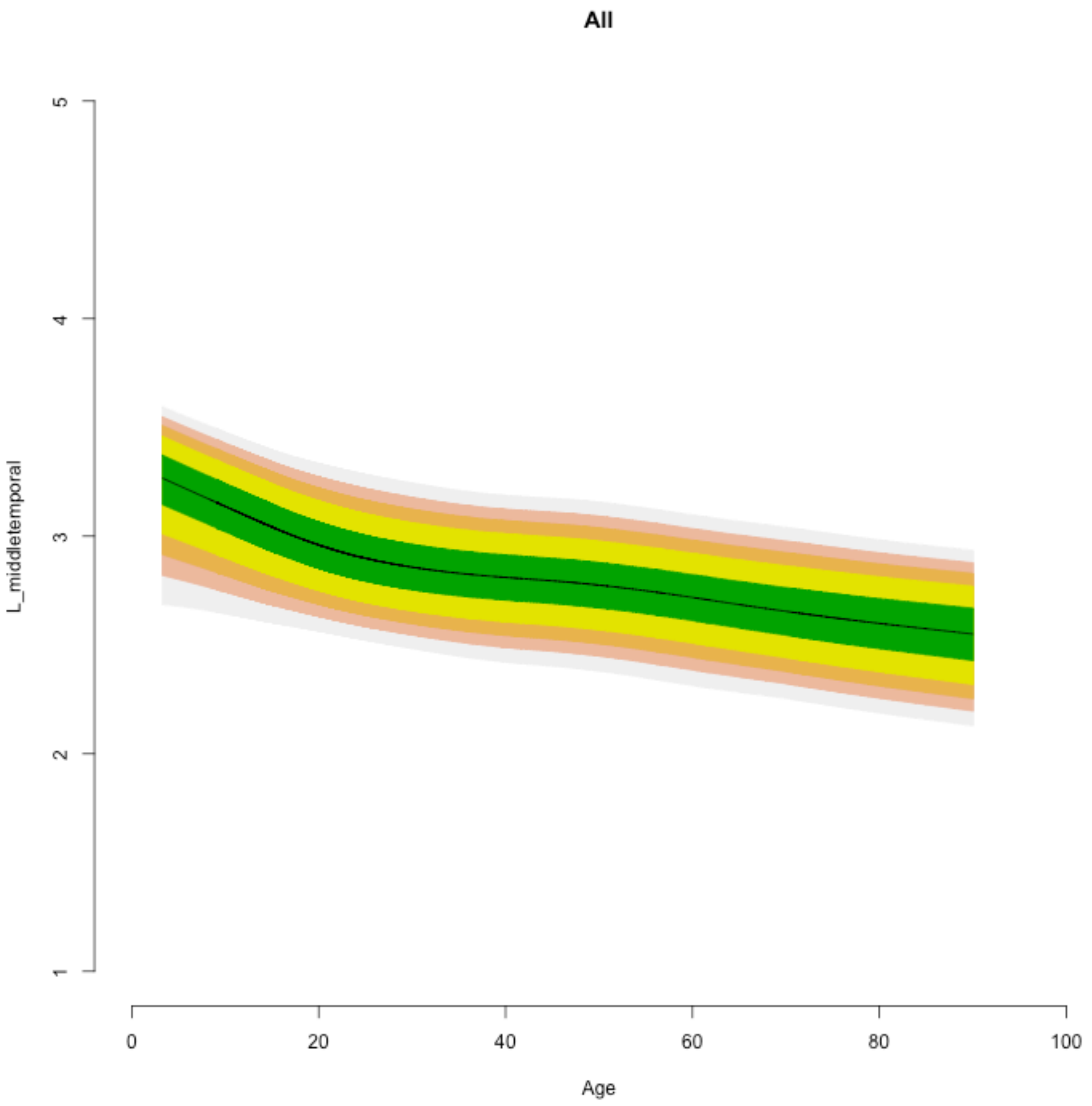
All
L_middletemporal
1
2
3
4
5
0
20
40
60
80
100
Age

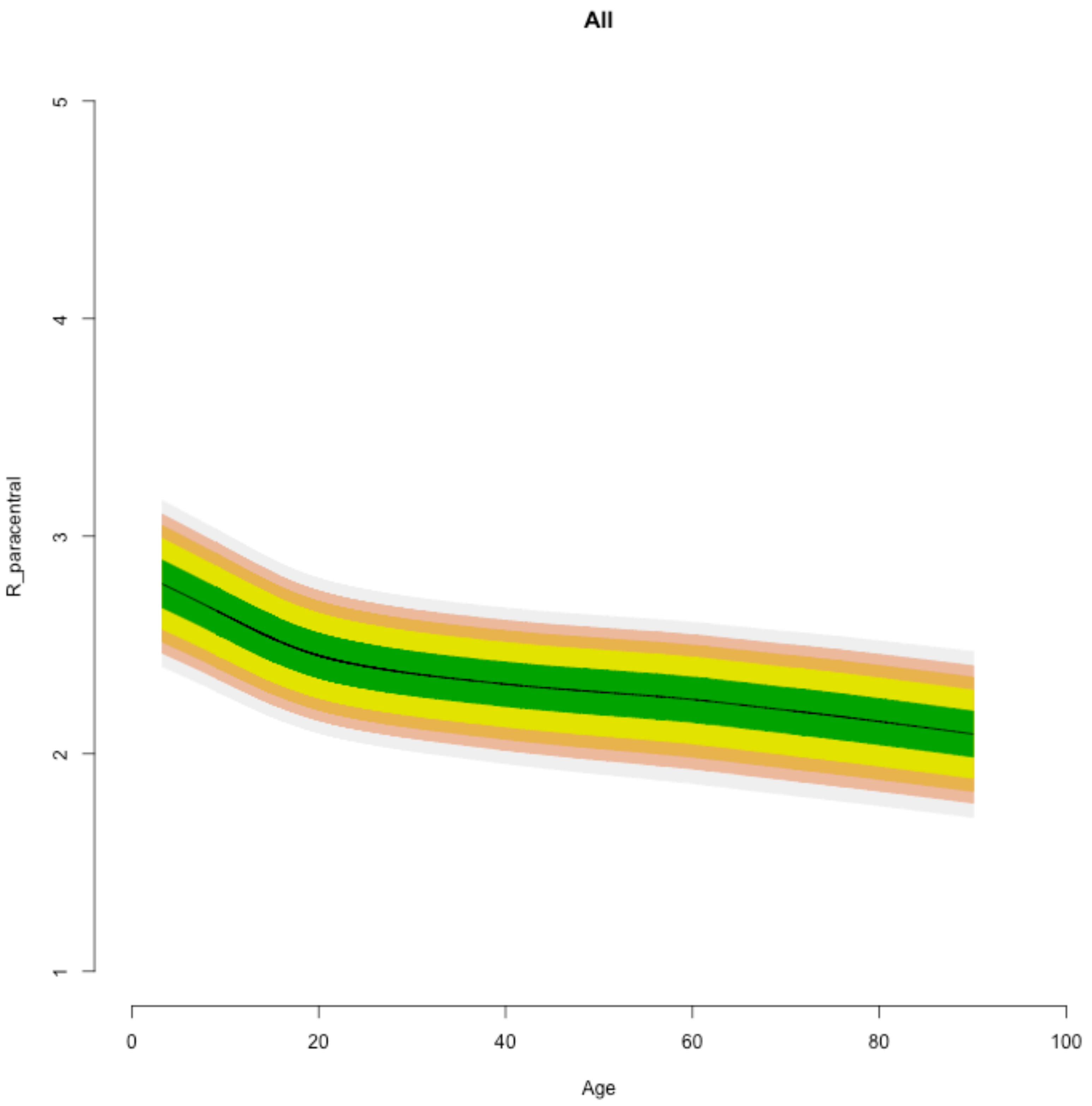
All
R_paracentral
1
2
3
4
5
0
20
40
60
80
100
Age

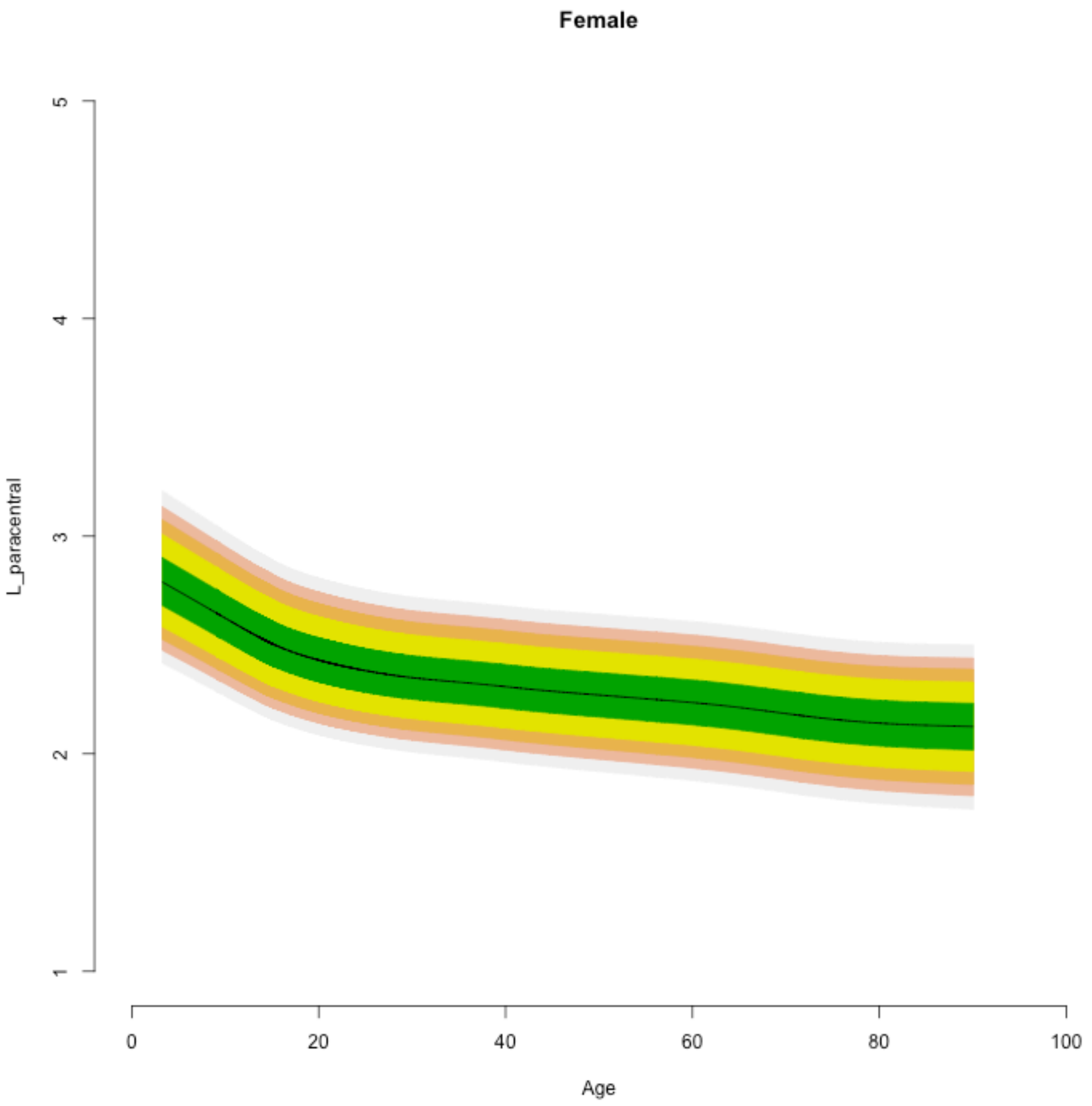

Female
L_paracentral
5
4
3
2
1
0
20
40
60
80
100
Age

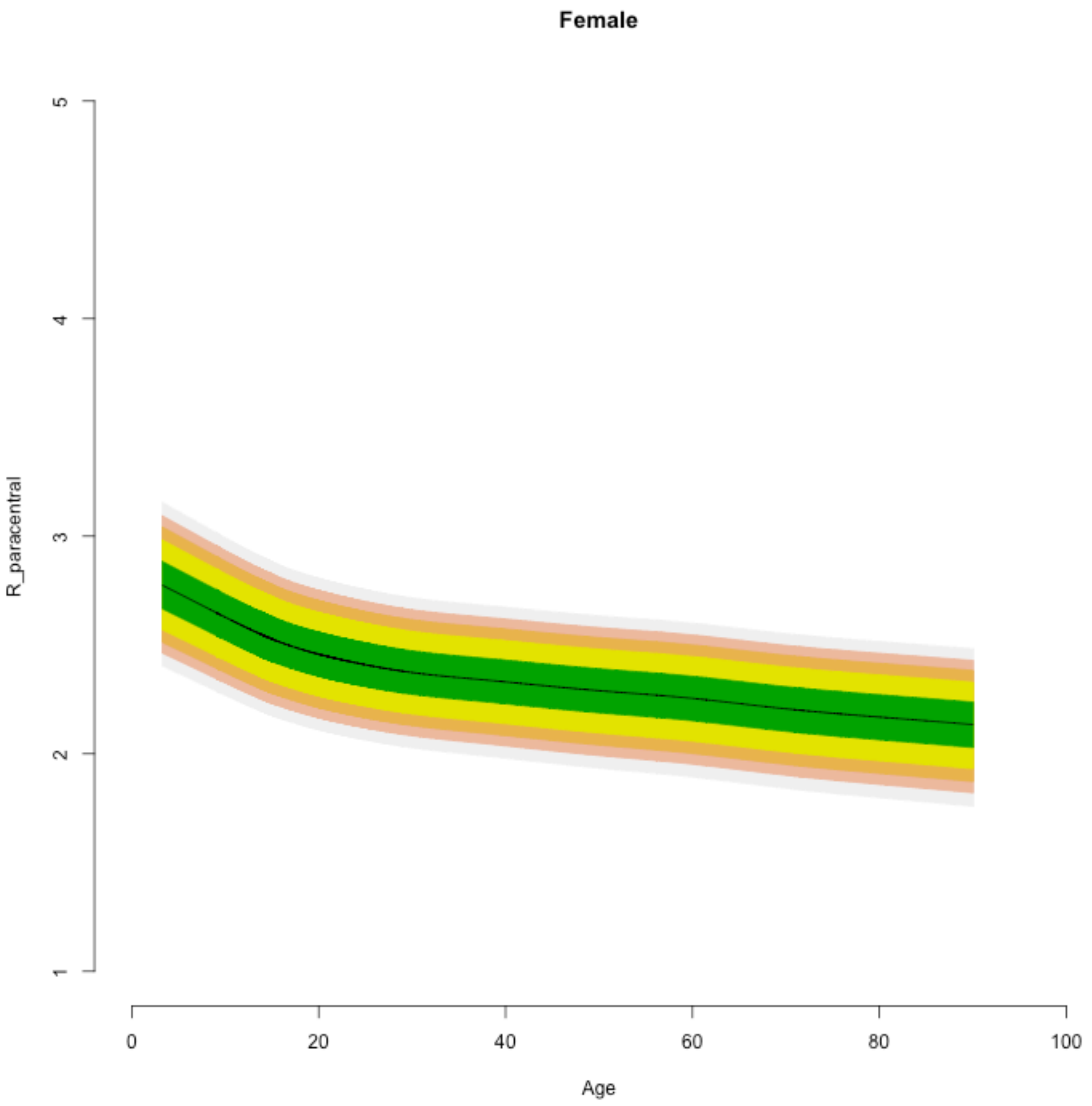
Female
R_paracentral
1
2
3
4
5
0
20
40
60
80
100
Age

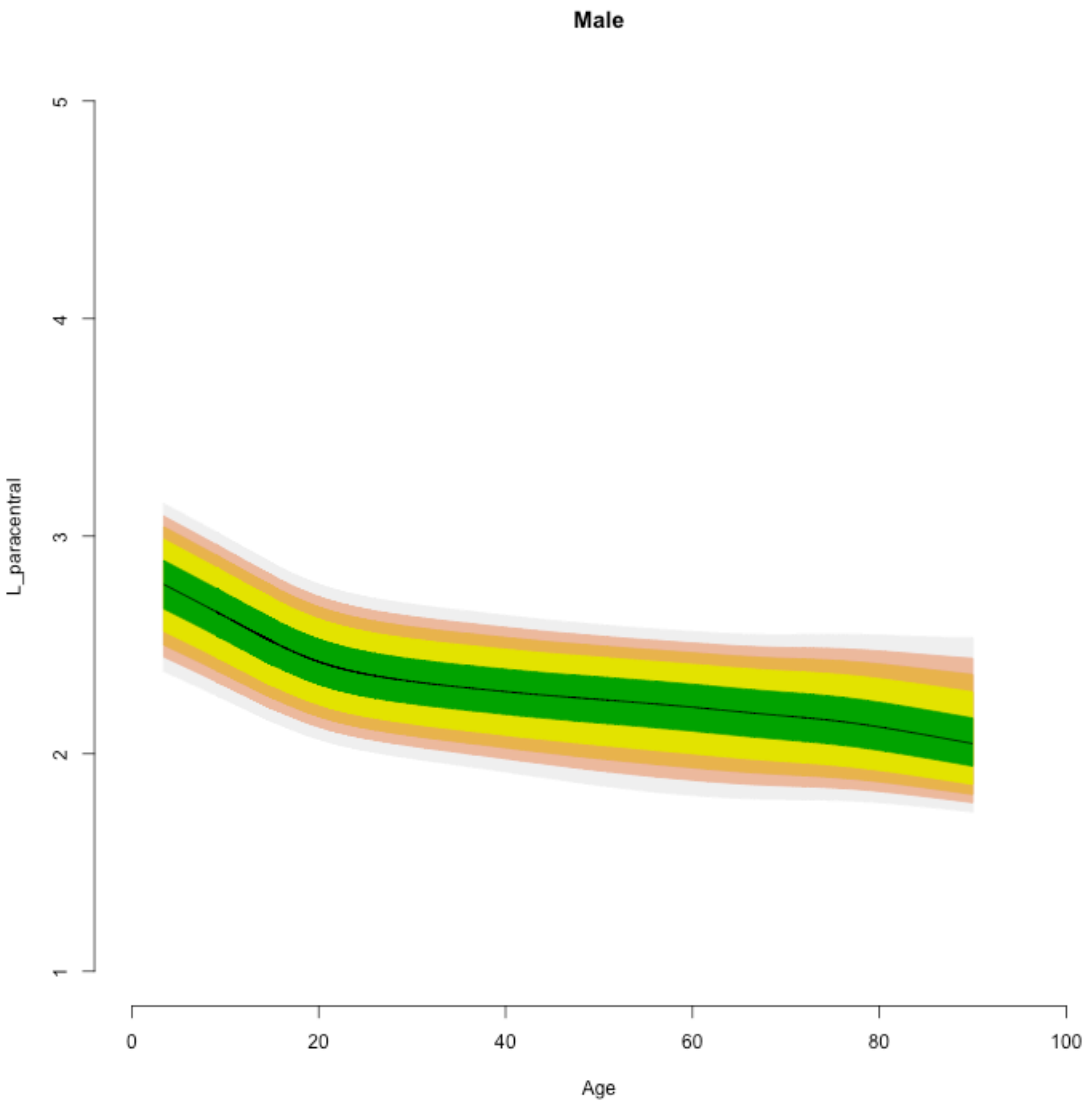

Male
L_paracentral
1
2
3
4
5
0
20
40
60
80
100
Age

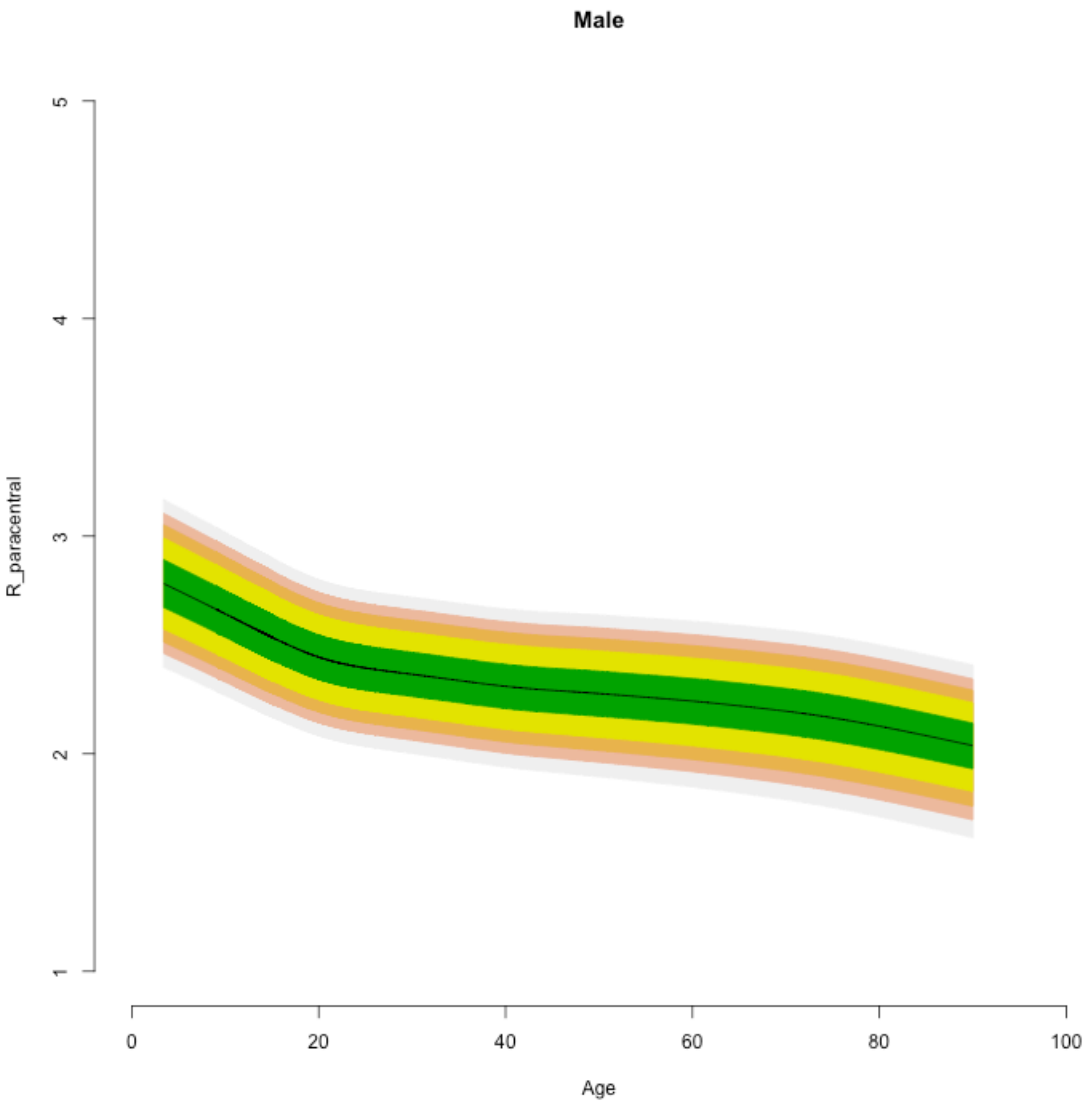

Male
R_paracentral
1
2
3
4
5
0
20
40
60
80
100
Age

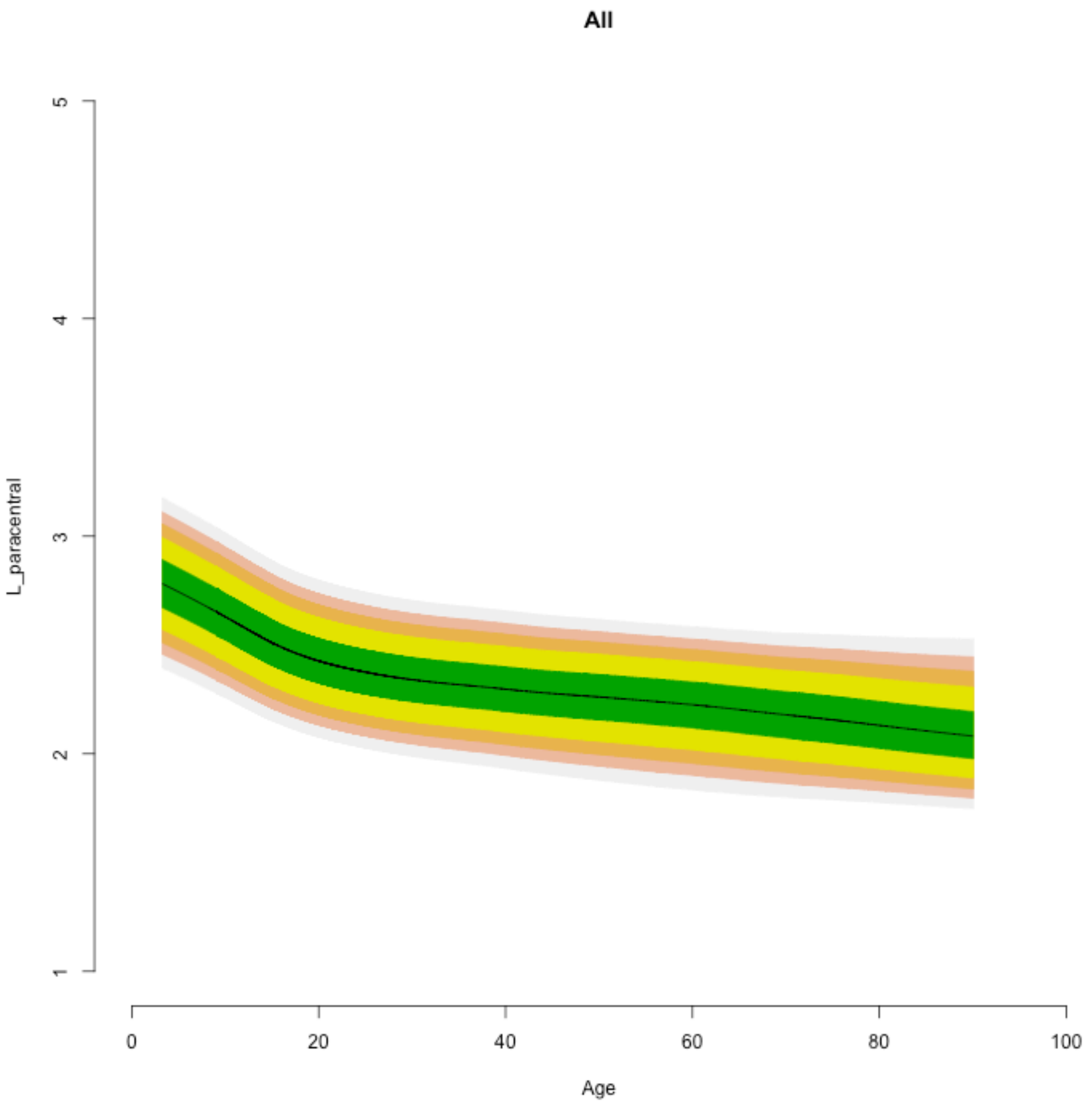
All
L_paracentral
1
2
3
4
5
0
20
40
60
80
100
Age

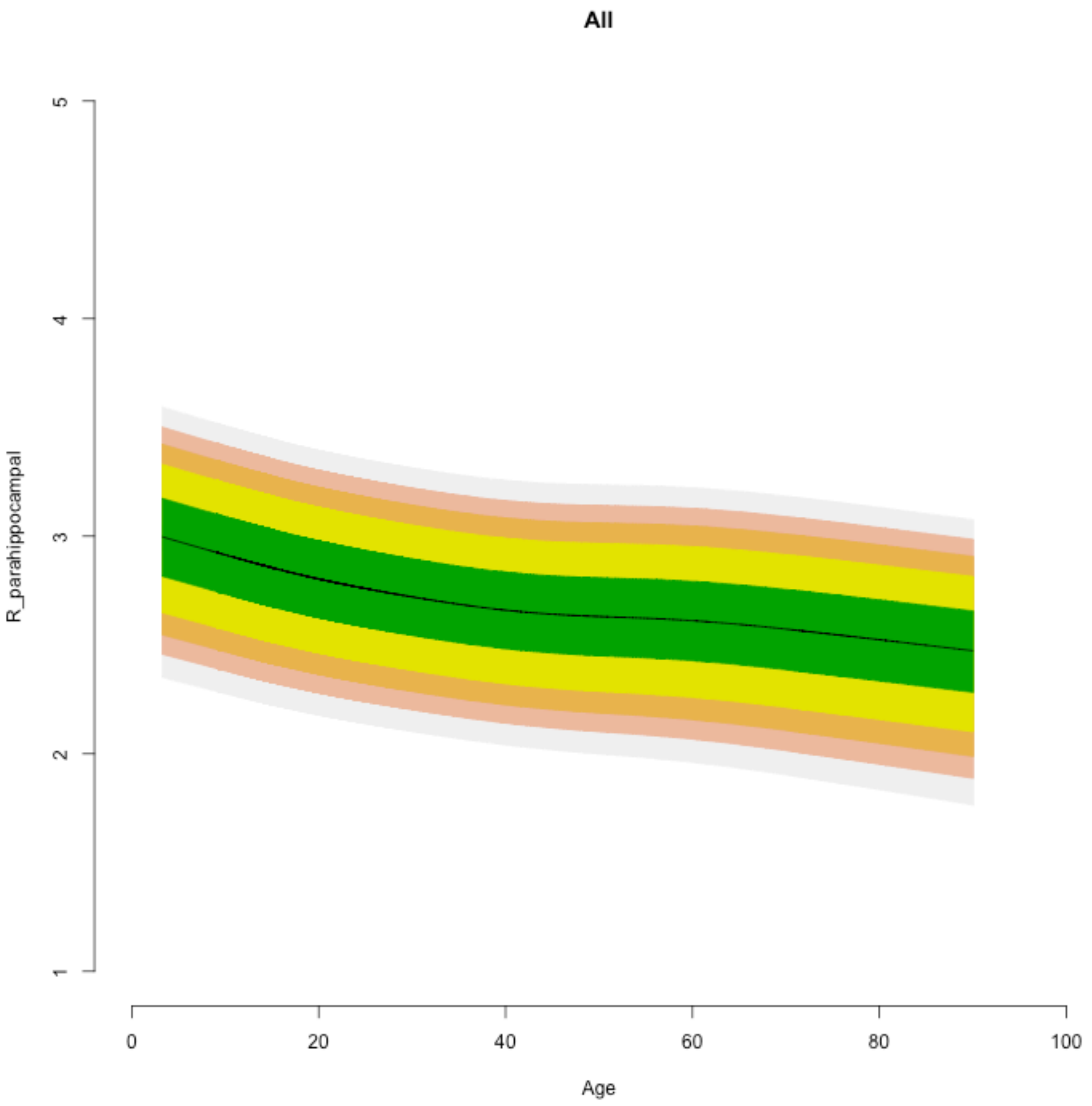

All
R_parahippocampal
1
2
3
4
5
0
20
40
60
80
100
Age

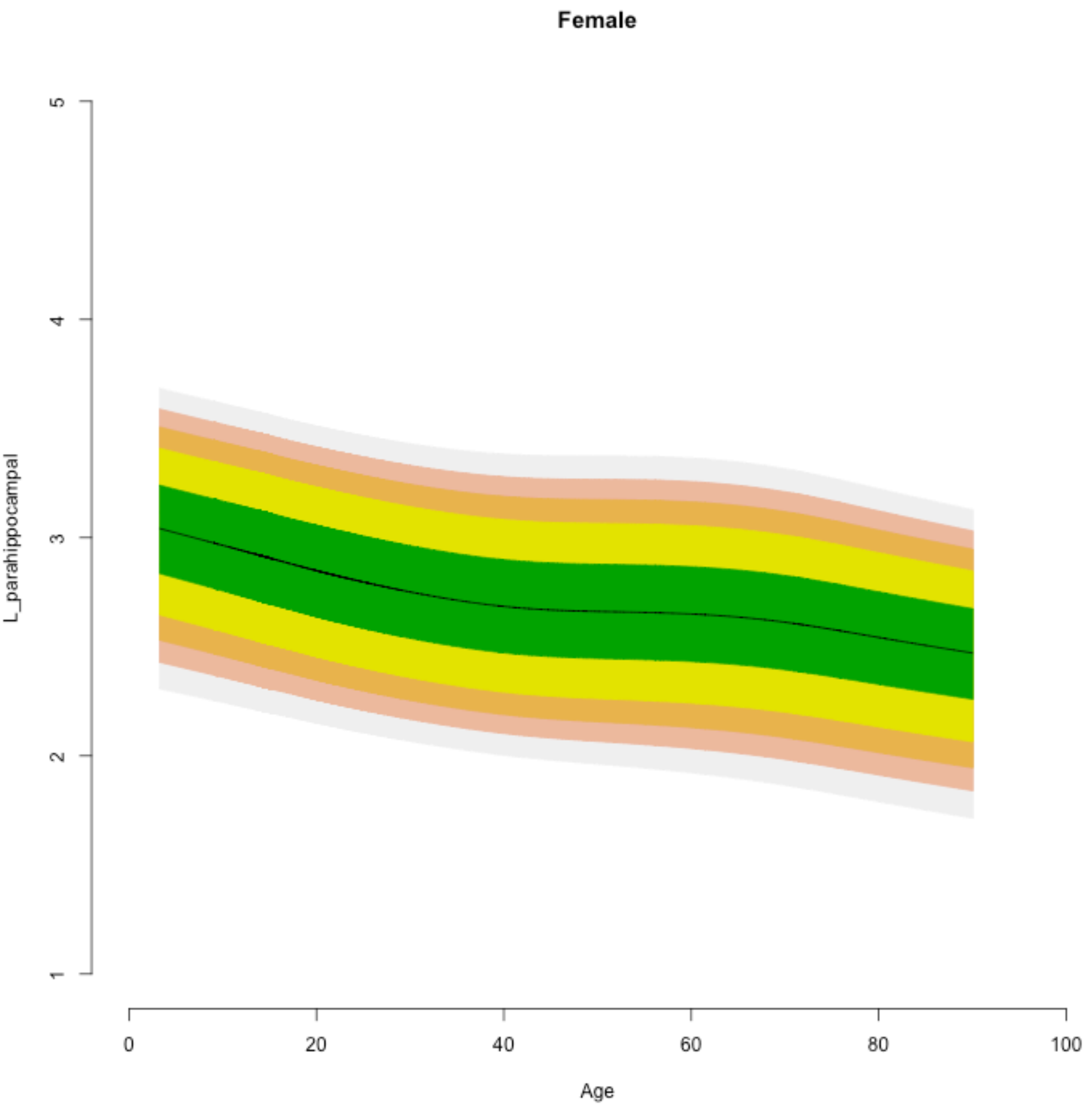
Female
L_parahippocampal
1
2
3
4
5
0
20
40
60
80
100
Age

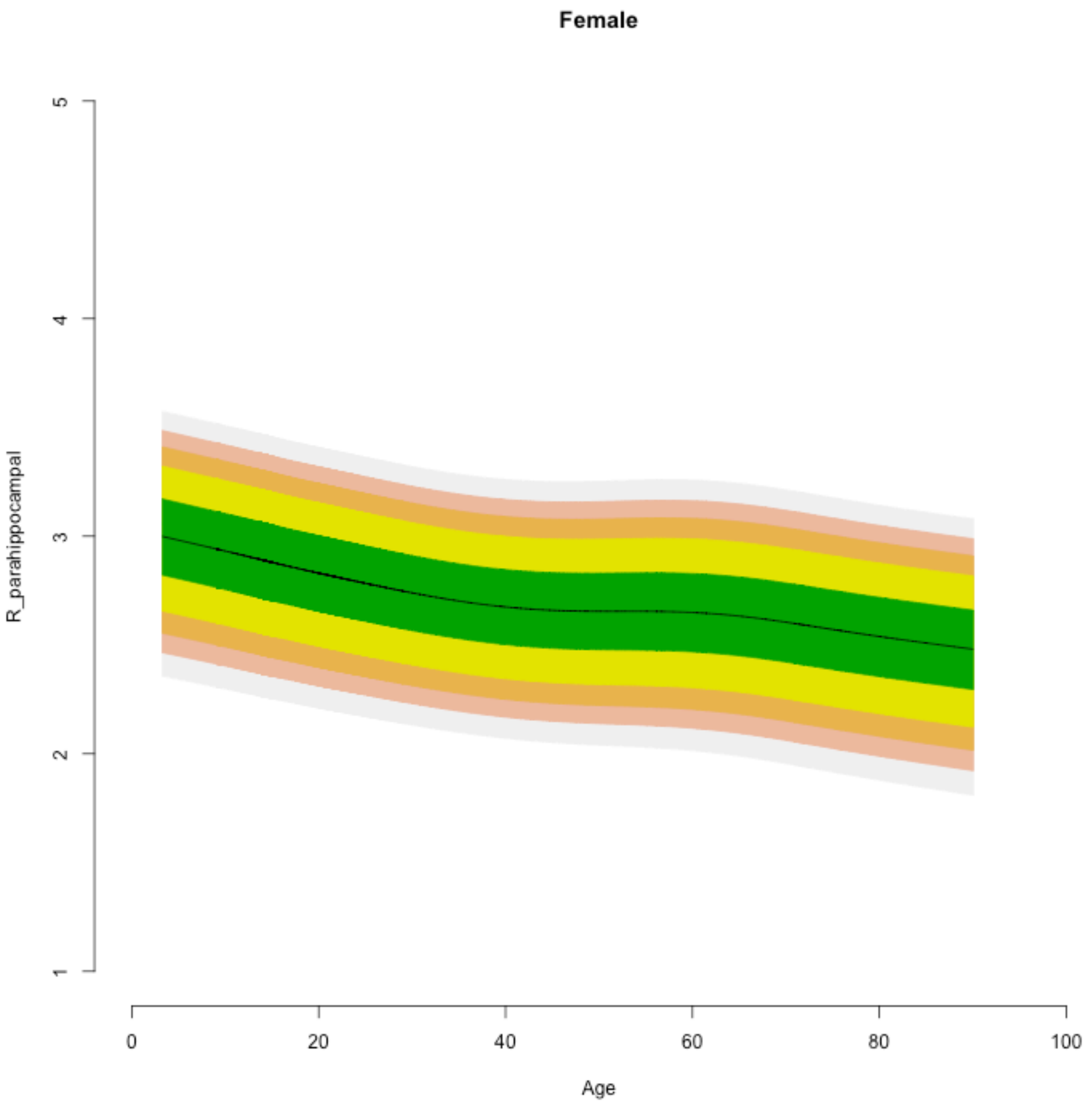

Female
R_parahippocampal
1
2
3
4
5
0
20
40
60
80
100
Age

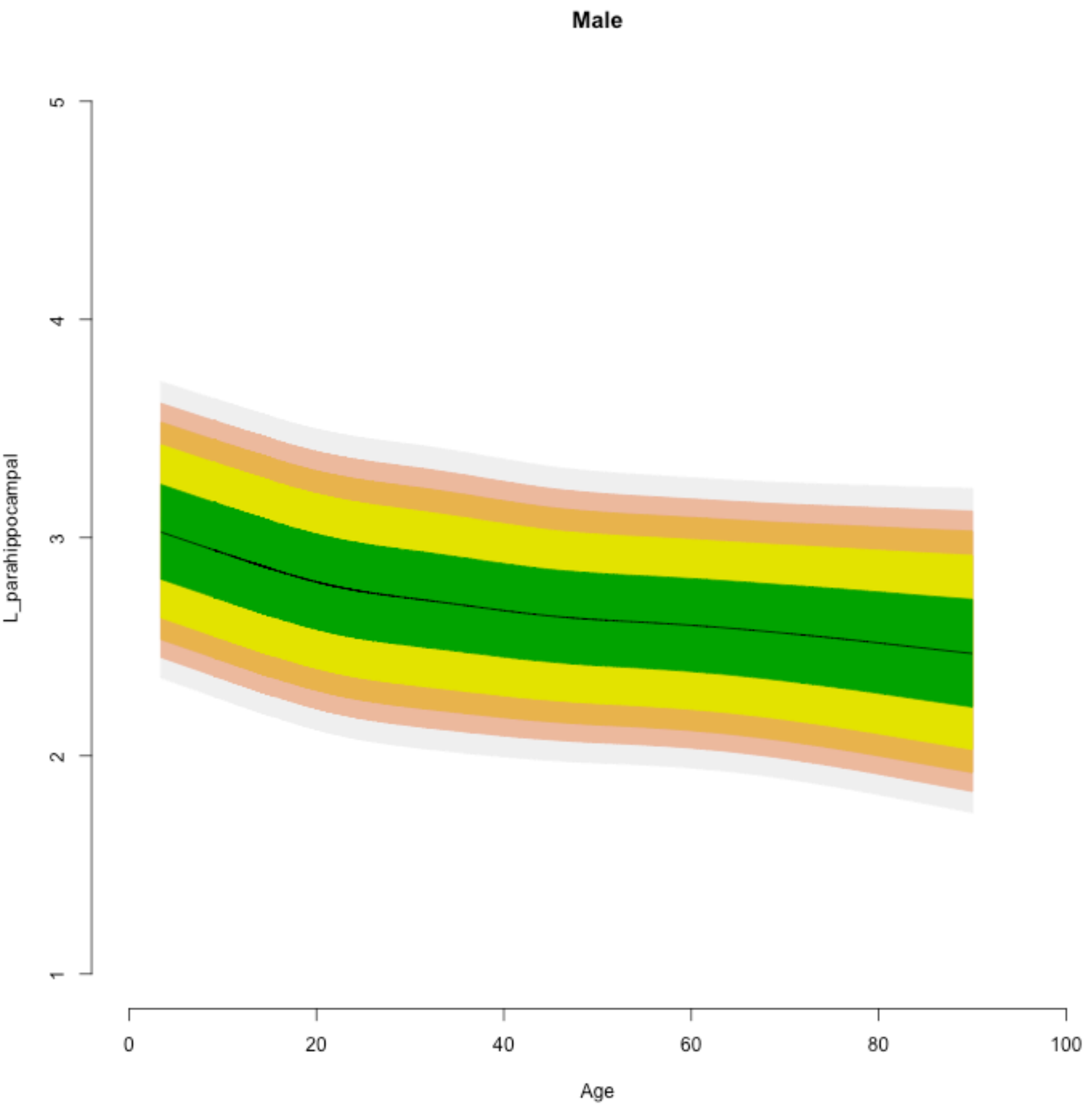
Male
L_parahippocampal
1
2
3
4
5
0
20
40
60
80
100
Age

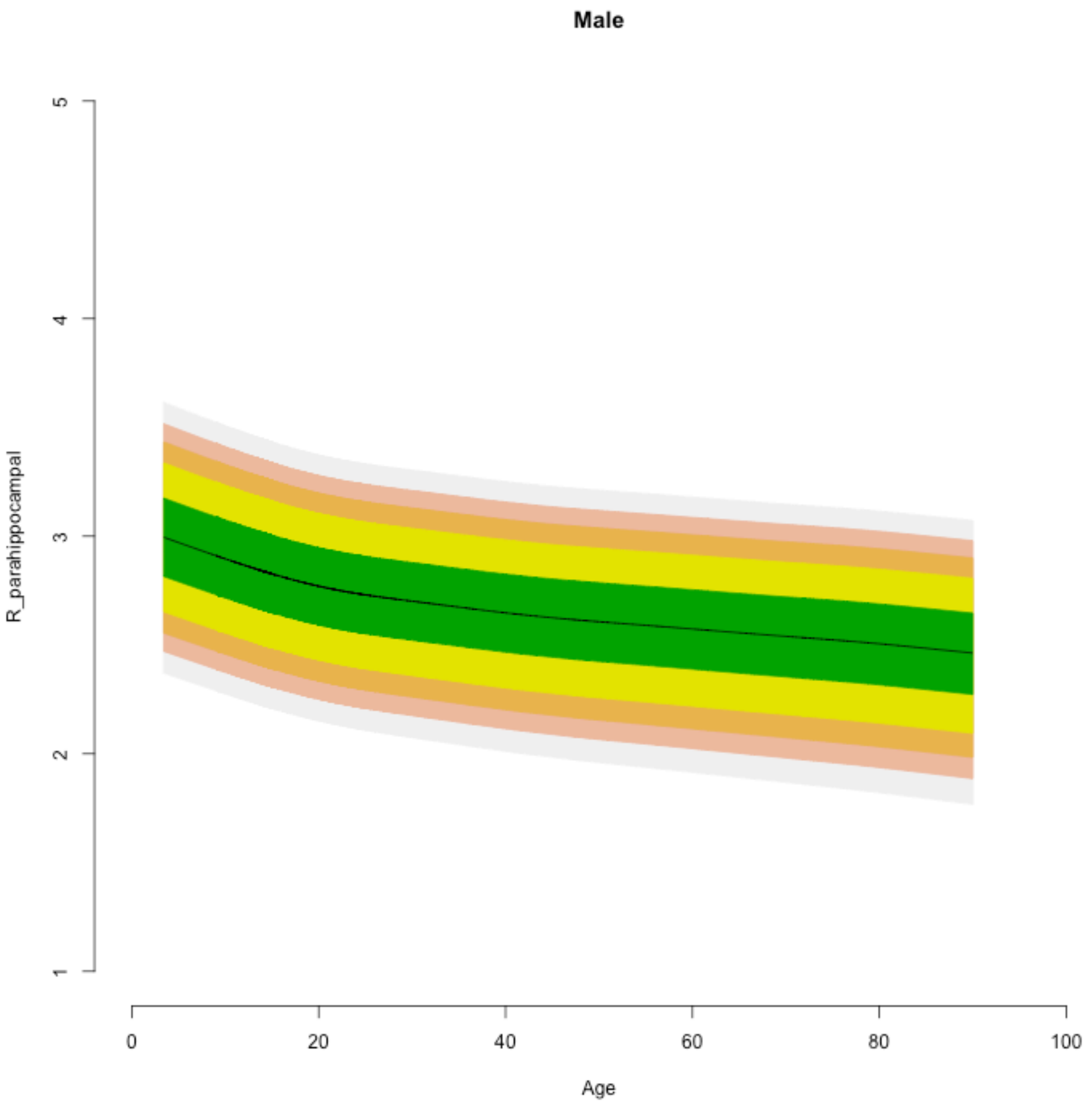
Male
R_parahippocampal
1
2
3
4
5
0
20
40
60
80
100
Age

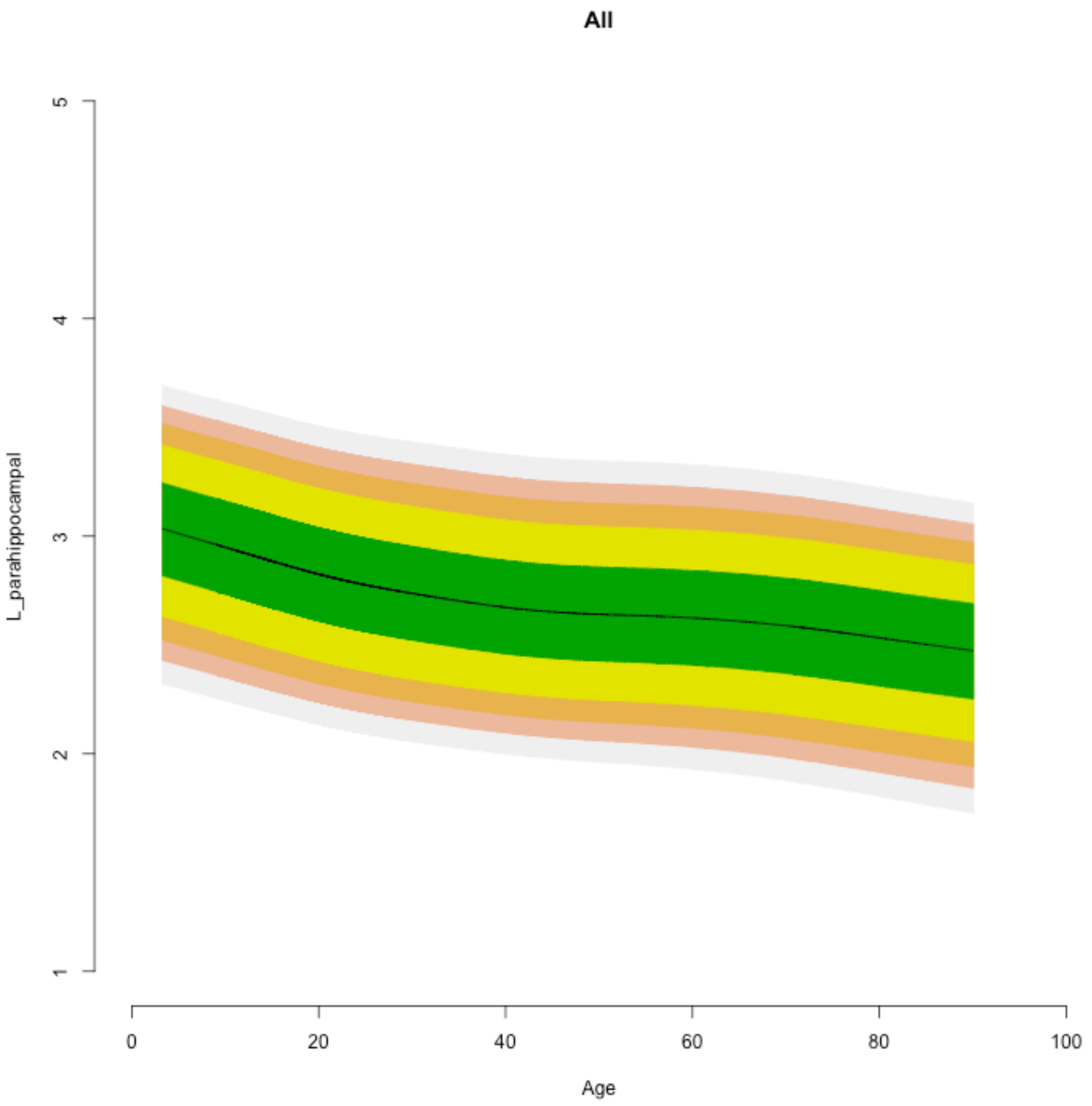

All
L_parahippocampal
1
2
3
4
5
0
20
40
60
80
100
Age

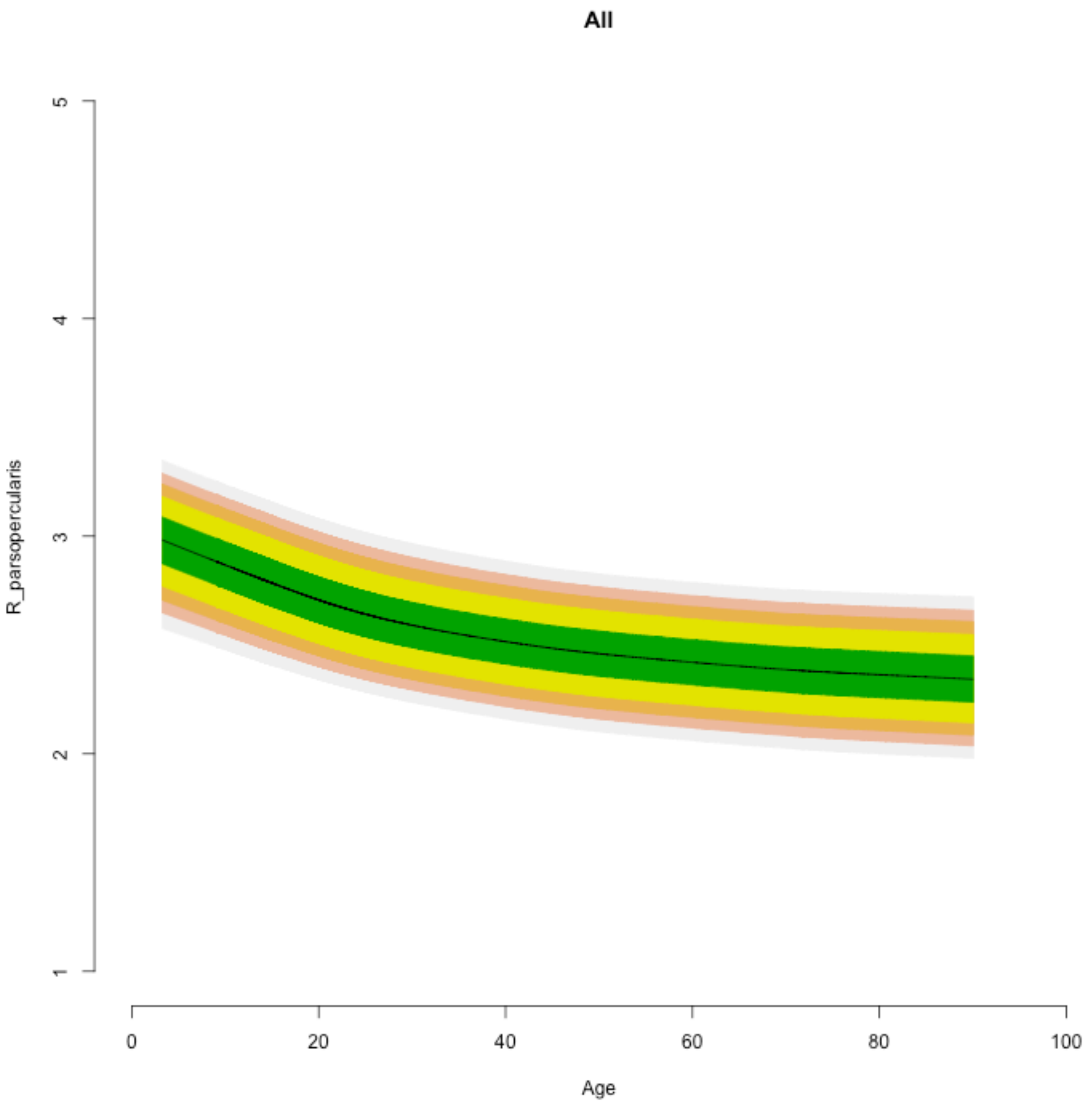
All
R_parsopercularis
1
2
3
4
5
0
20
40
60
80
100
Age

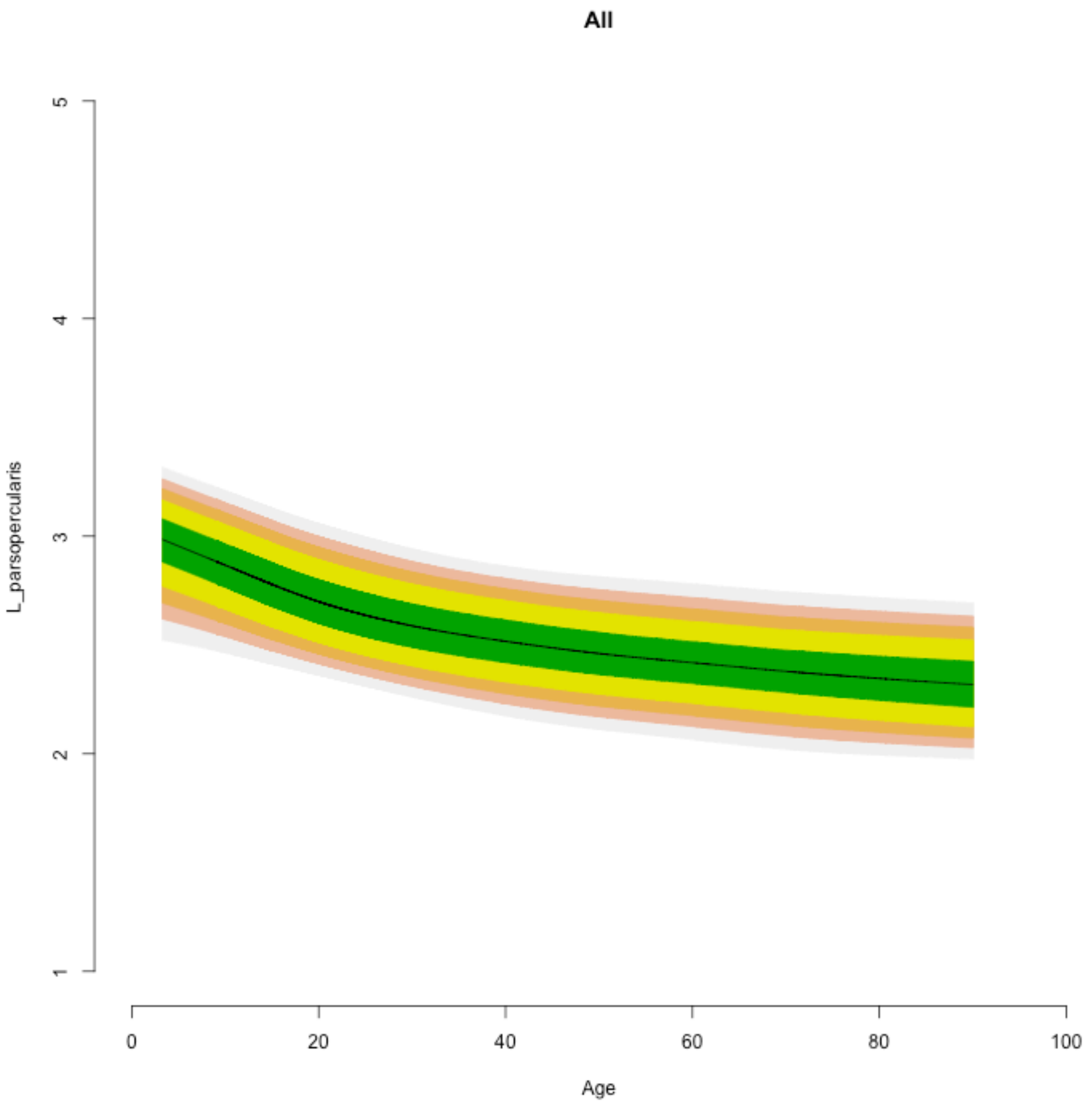
All
L_parsopercularis
1
2
3
4
5
0
20
40
60
80
100
Age

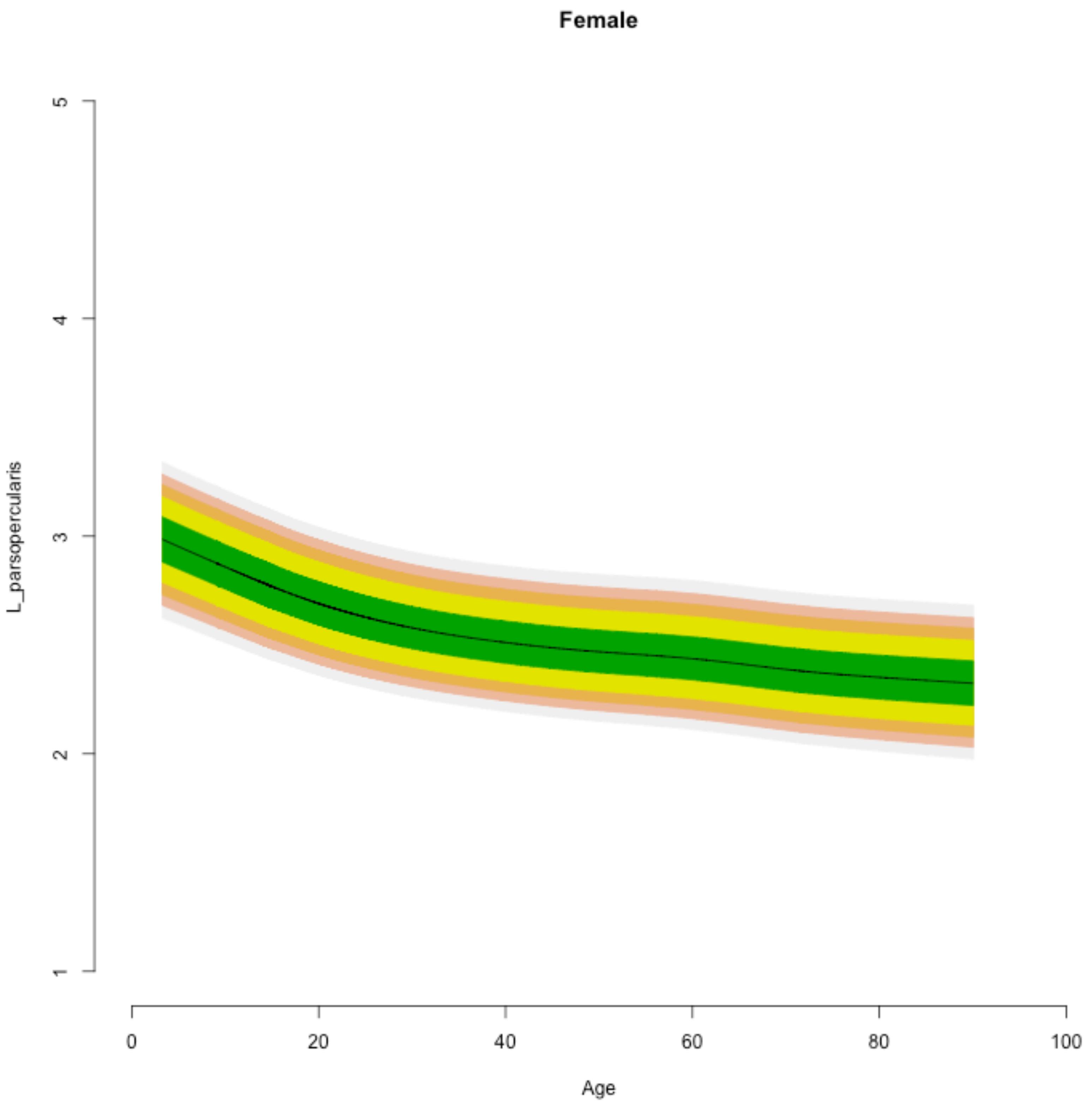
Female
L_parsopercularis
1
2
3
4
5
Age
0
20
40
60
80
100

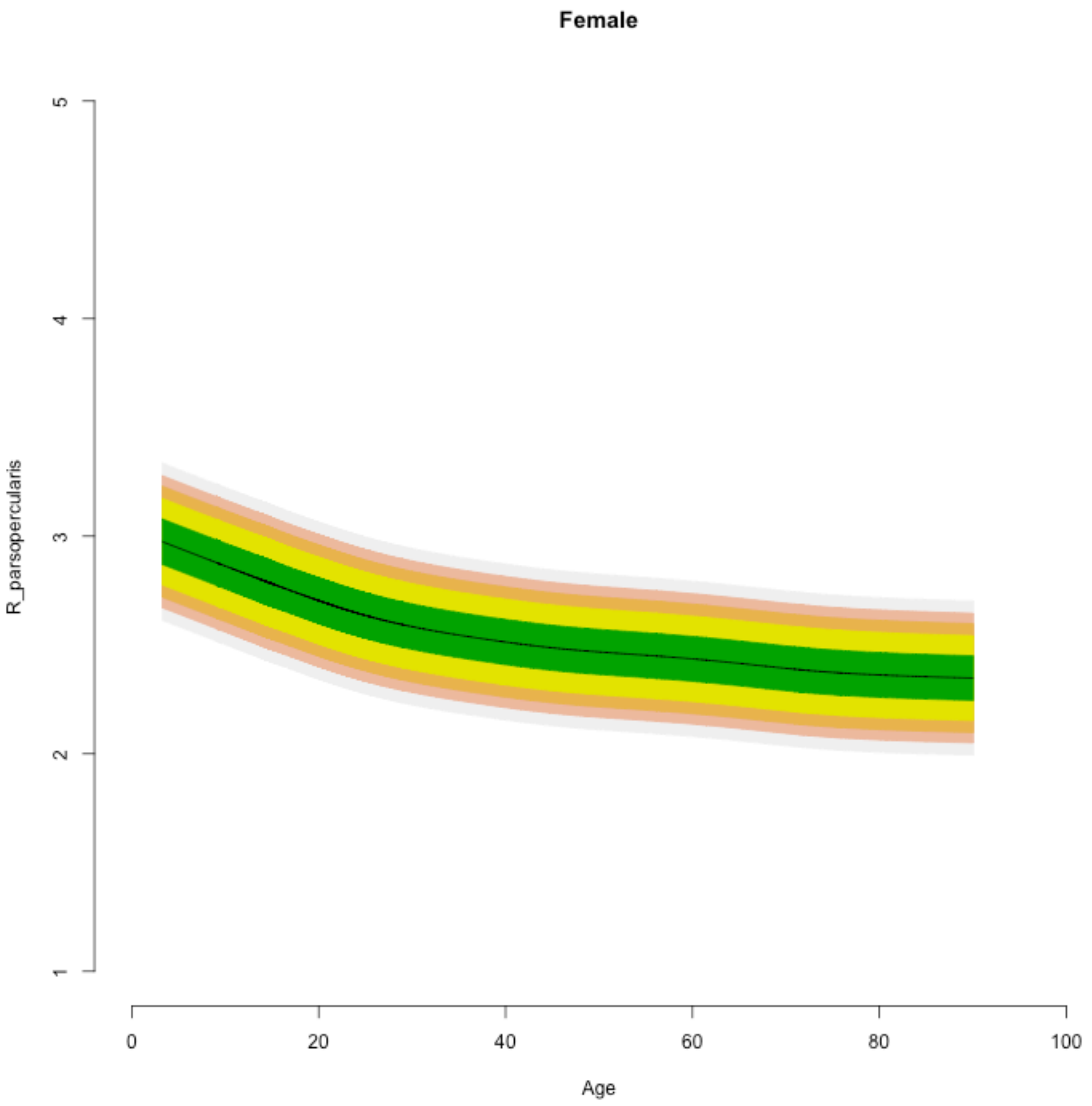
Female
R_parsopercularis
5
4
3
2
1
0
20
40
60
80
100
Age

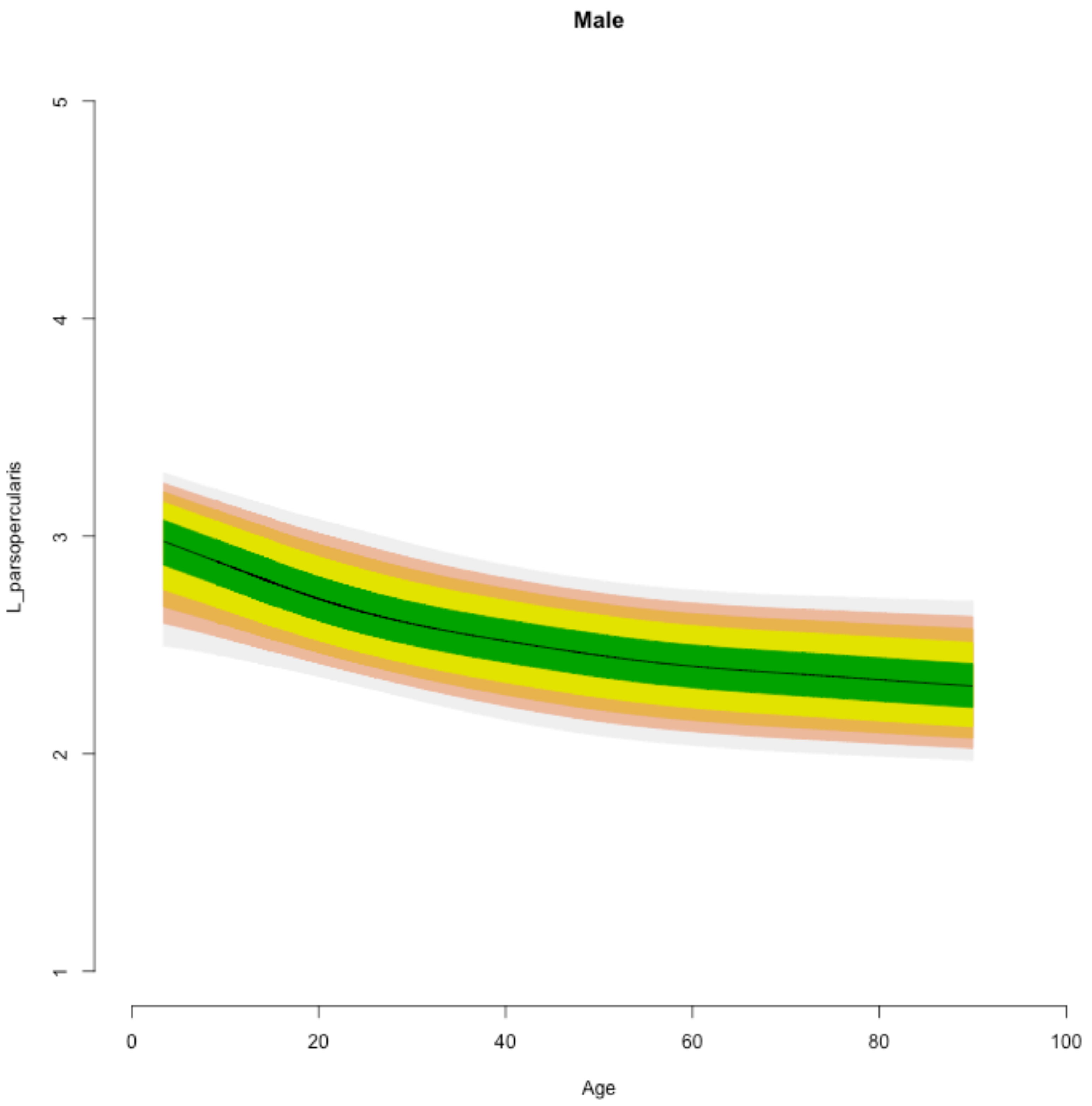

Male
L_parsopercularis
5
4
3
2
1
0
20
40
60
80
100
Age

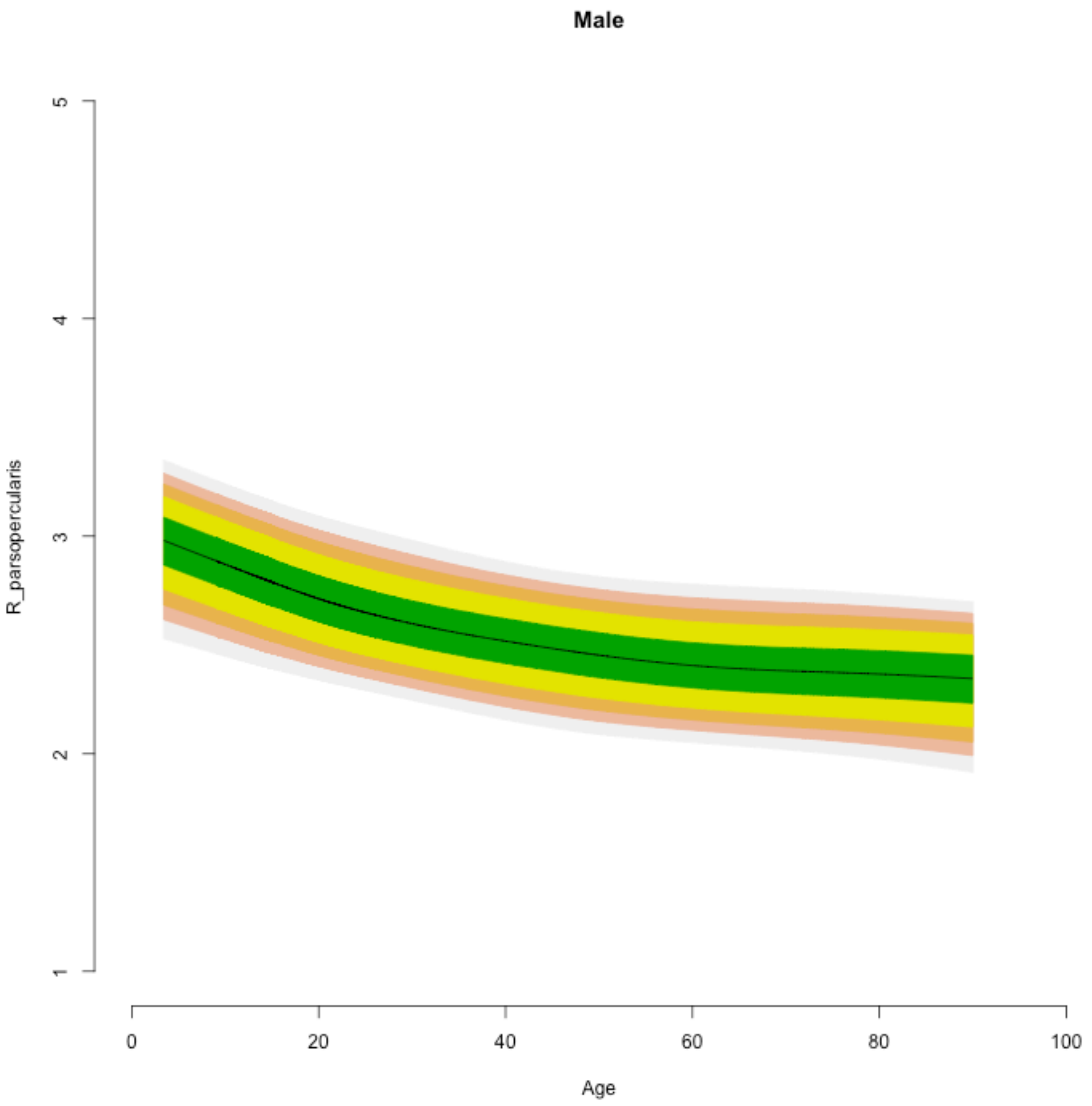

Male
R_parsopercularis
5
4
3
2
1
0
20
40
60
80
100
Age

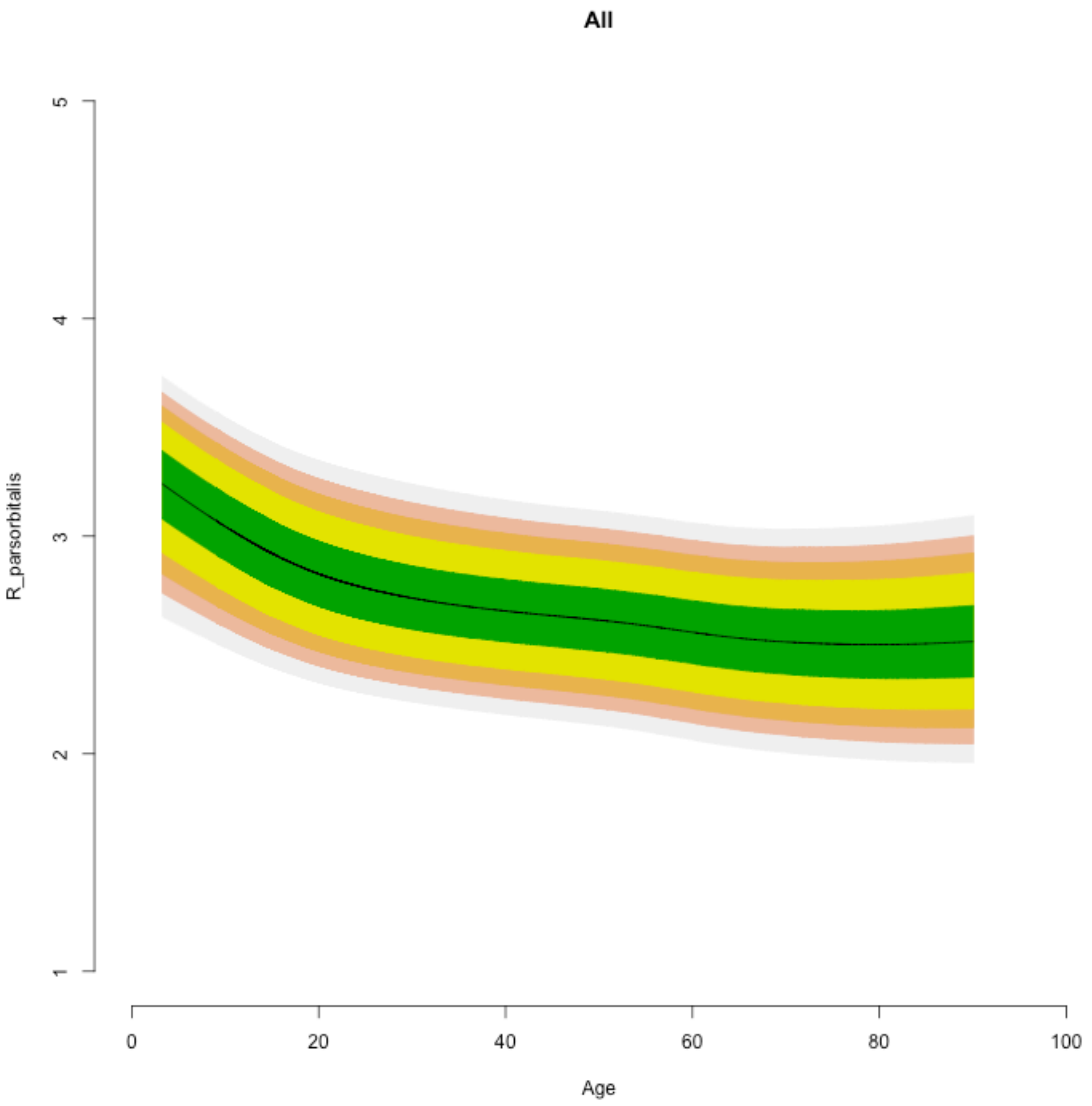

All
R_parsorbitalis
Age
1
2
3
4
5
0
20
40
60
80
100

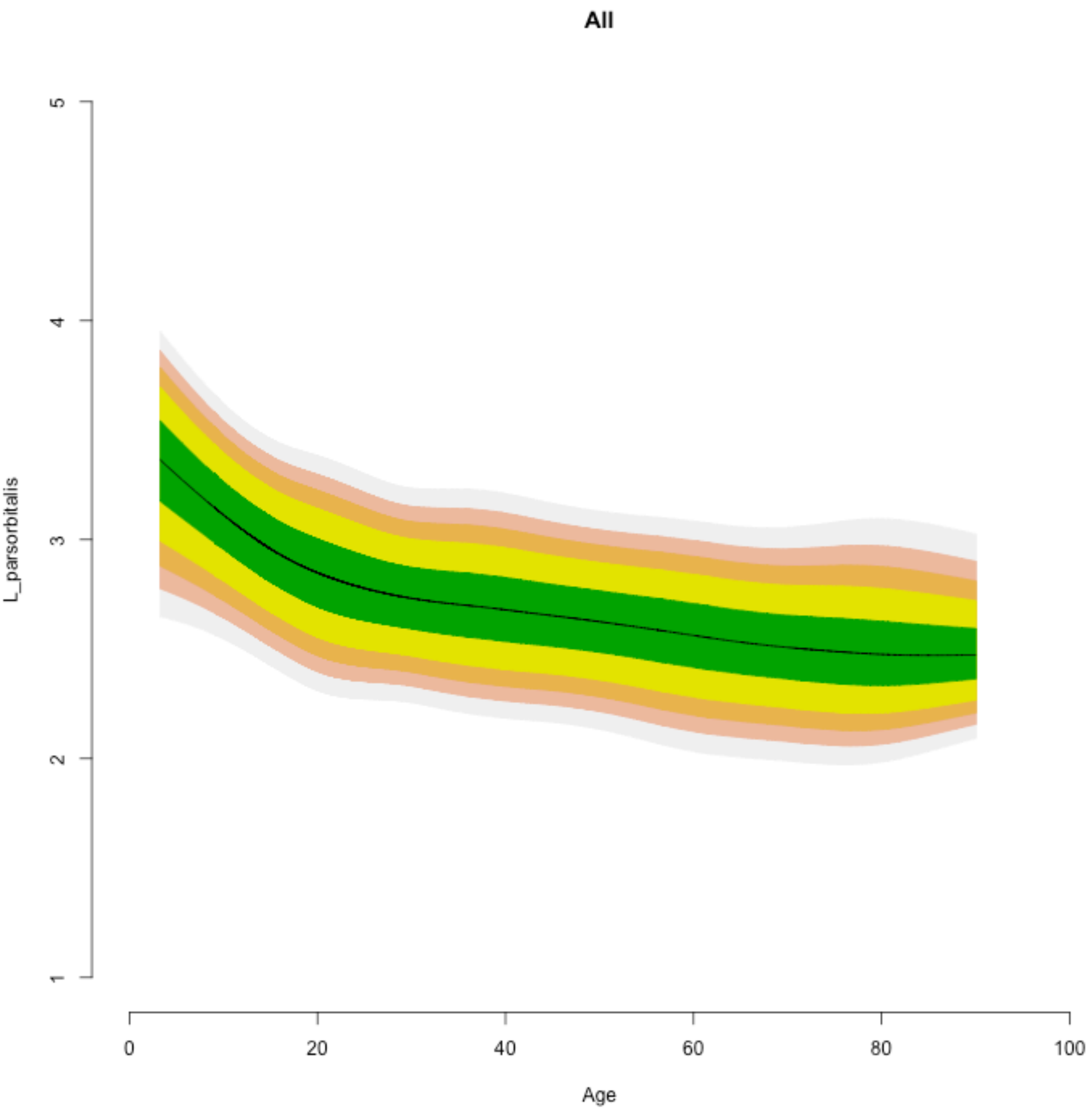
All
L_parsorbitalis
Age
1
2
3
4
5
0
20
40
60
80
100

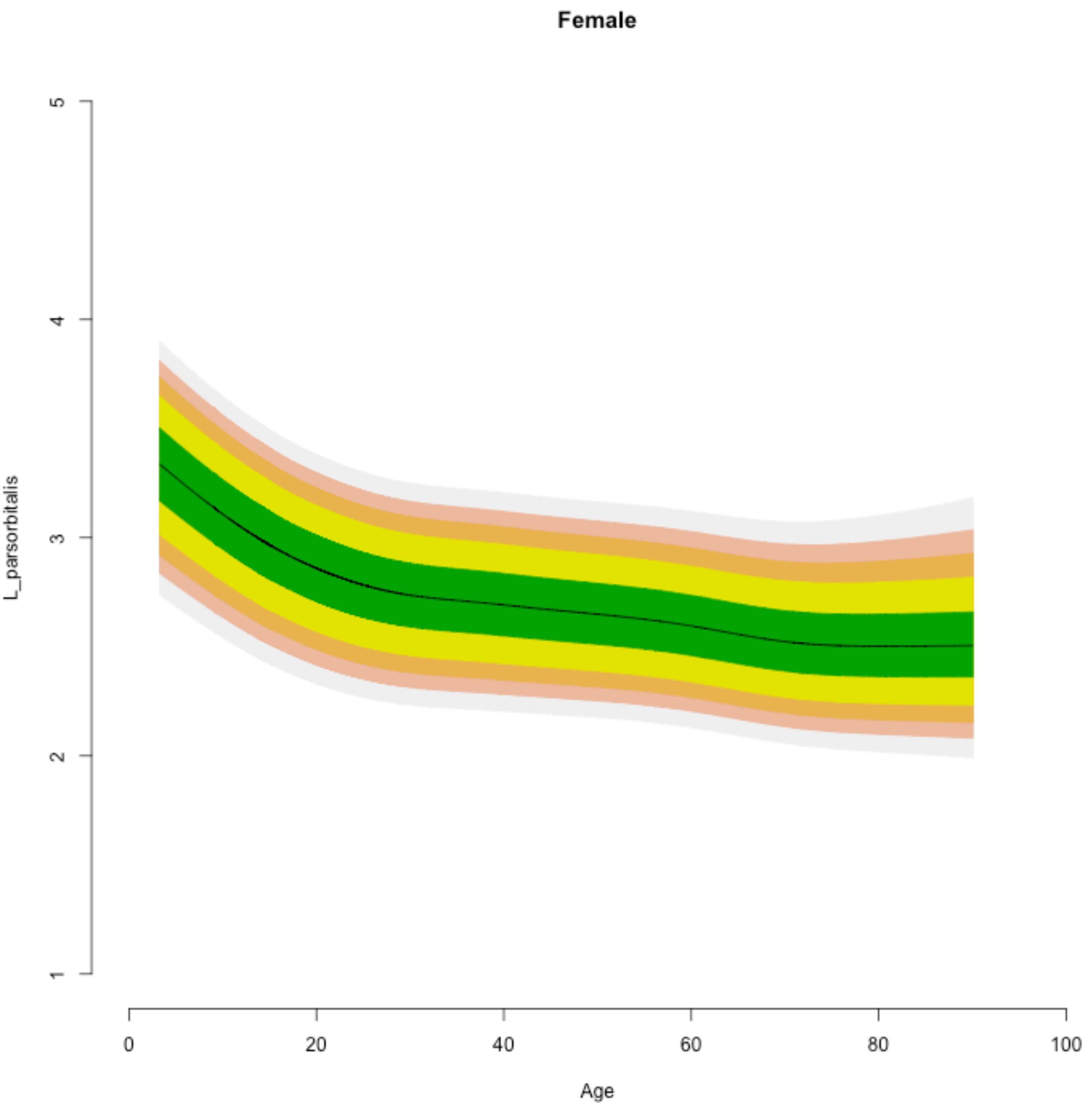

Female
L_parsorbitalis
5
4
3
2
1
0
20
40
60
80
100
Age

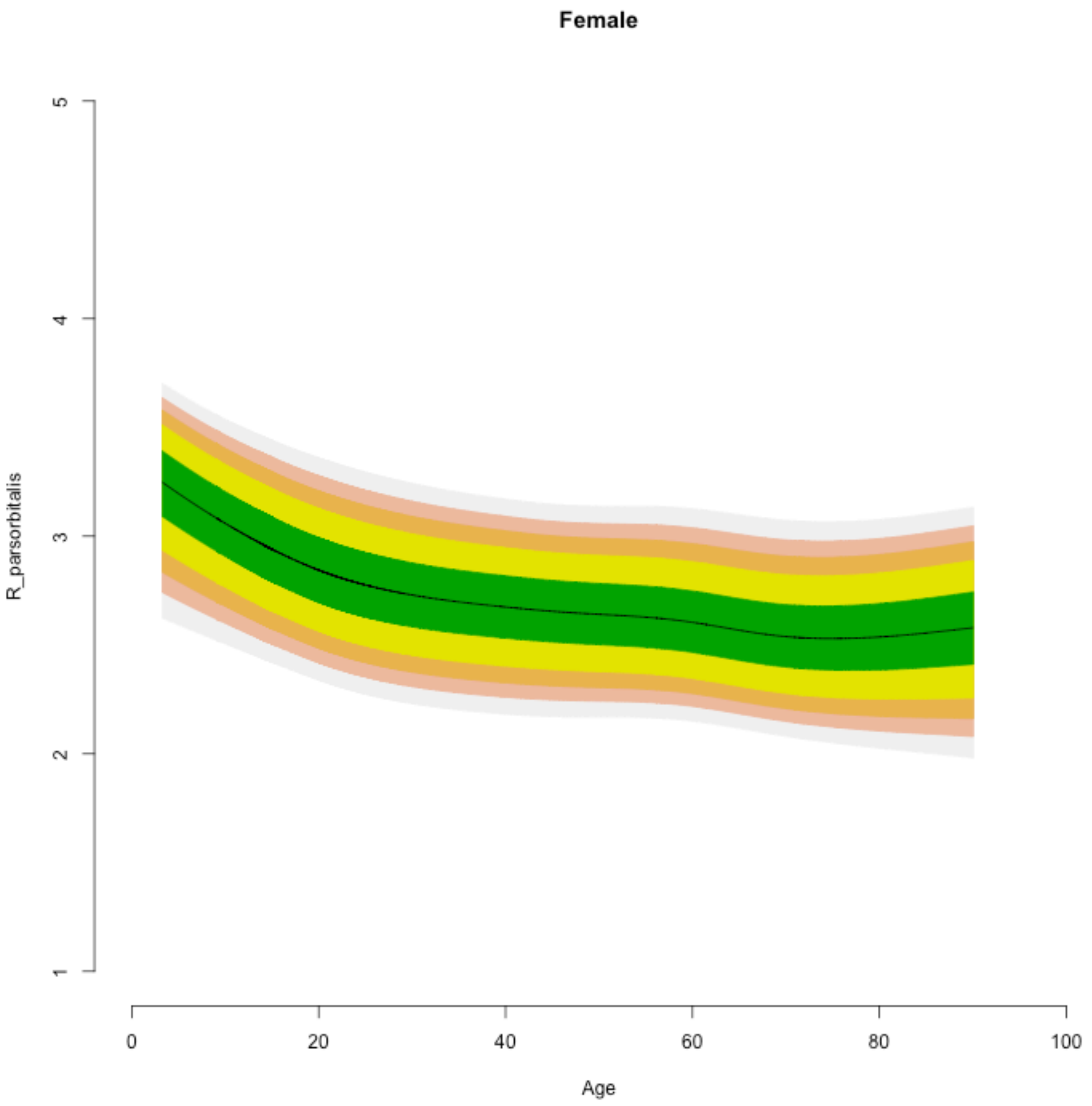

Female
R_parsorbitalis
1
2
3
4
5
0
20
40
60
80
100
Age

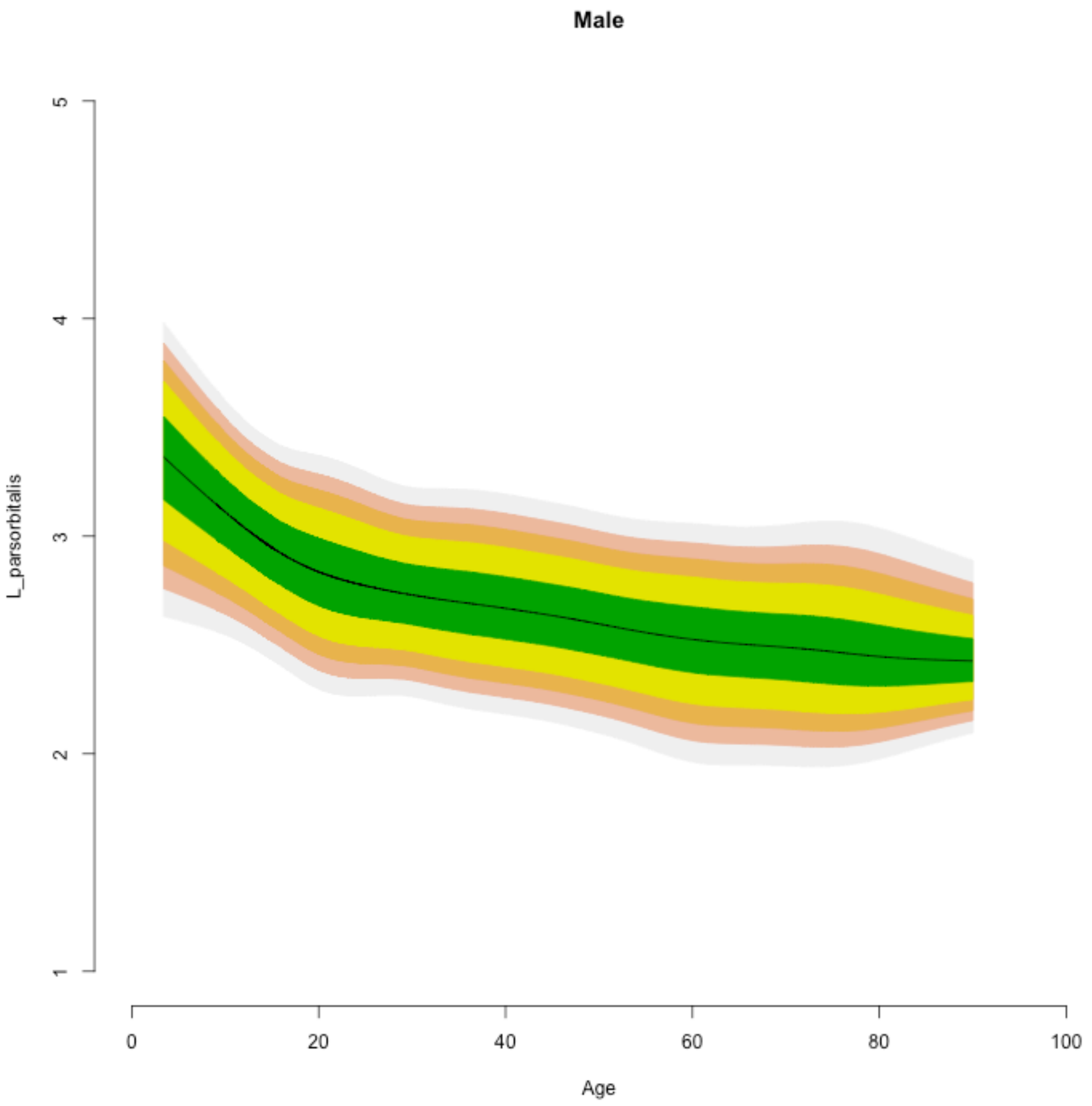

Male
L_parsorbitalis
5
4
3
2
1
0
20
40
60
80
100
Age

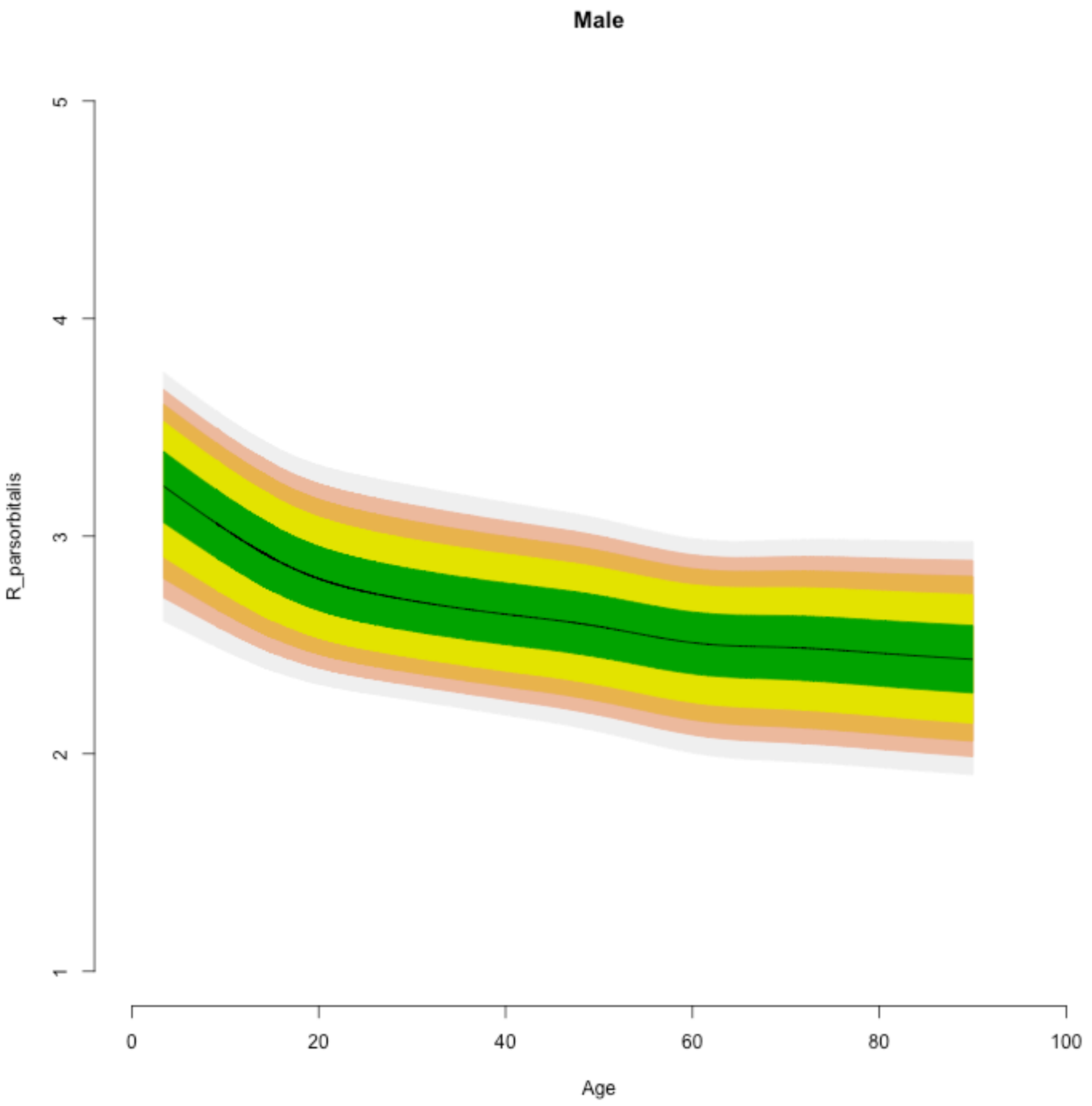

Male
R_parsorbitalis
Age
1
2
3
4
5
0
20
40
60
80
100

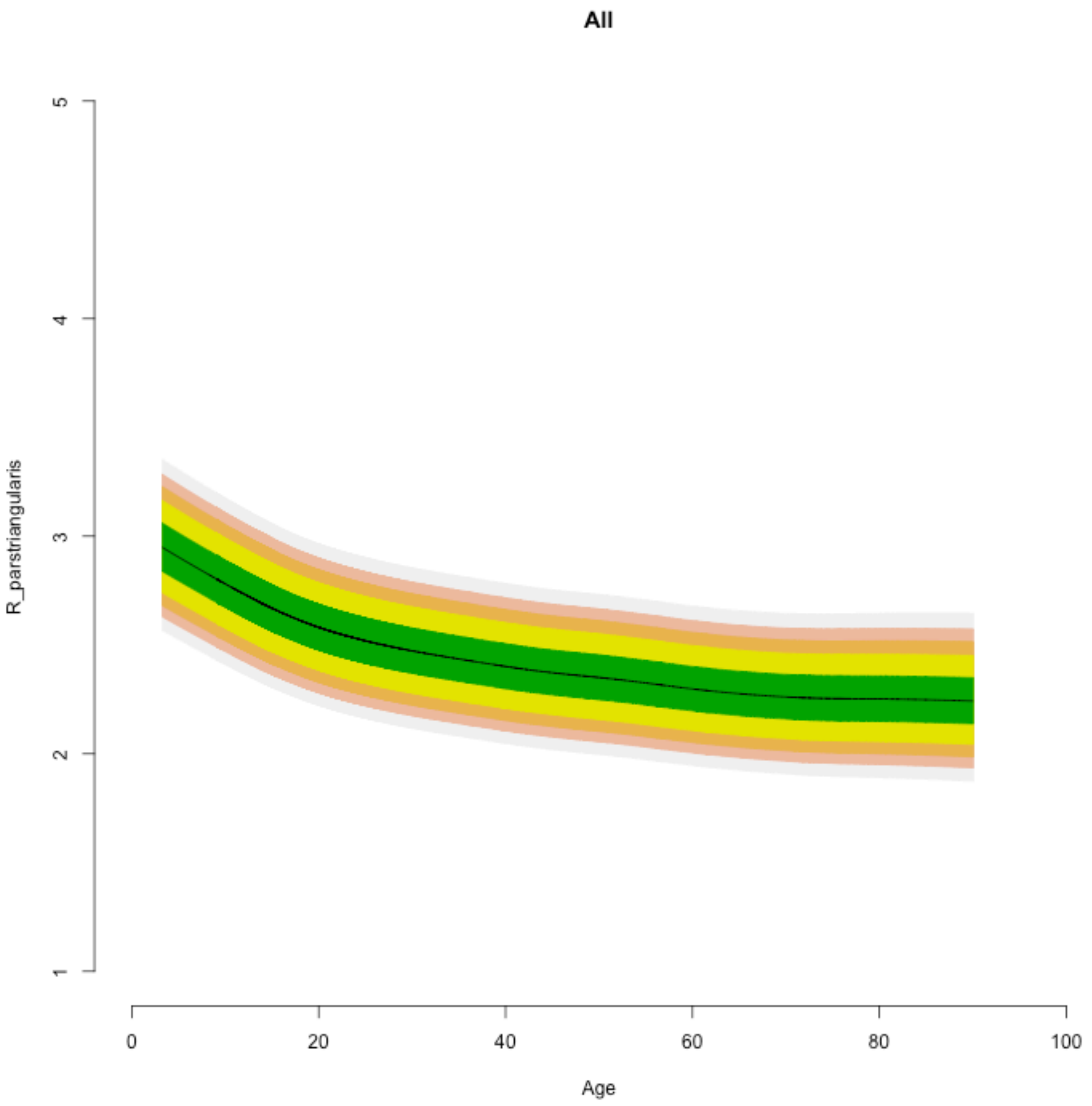
All
R_parstriangularis
1
2
3
4
5
0
20
40
60
80
100
Age

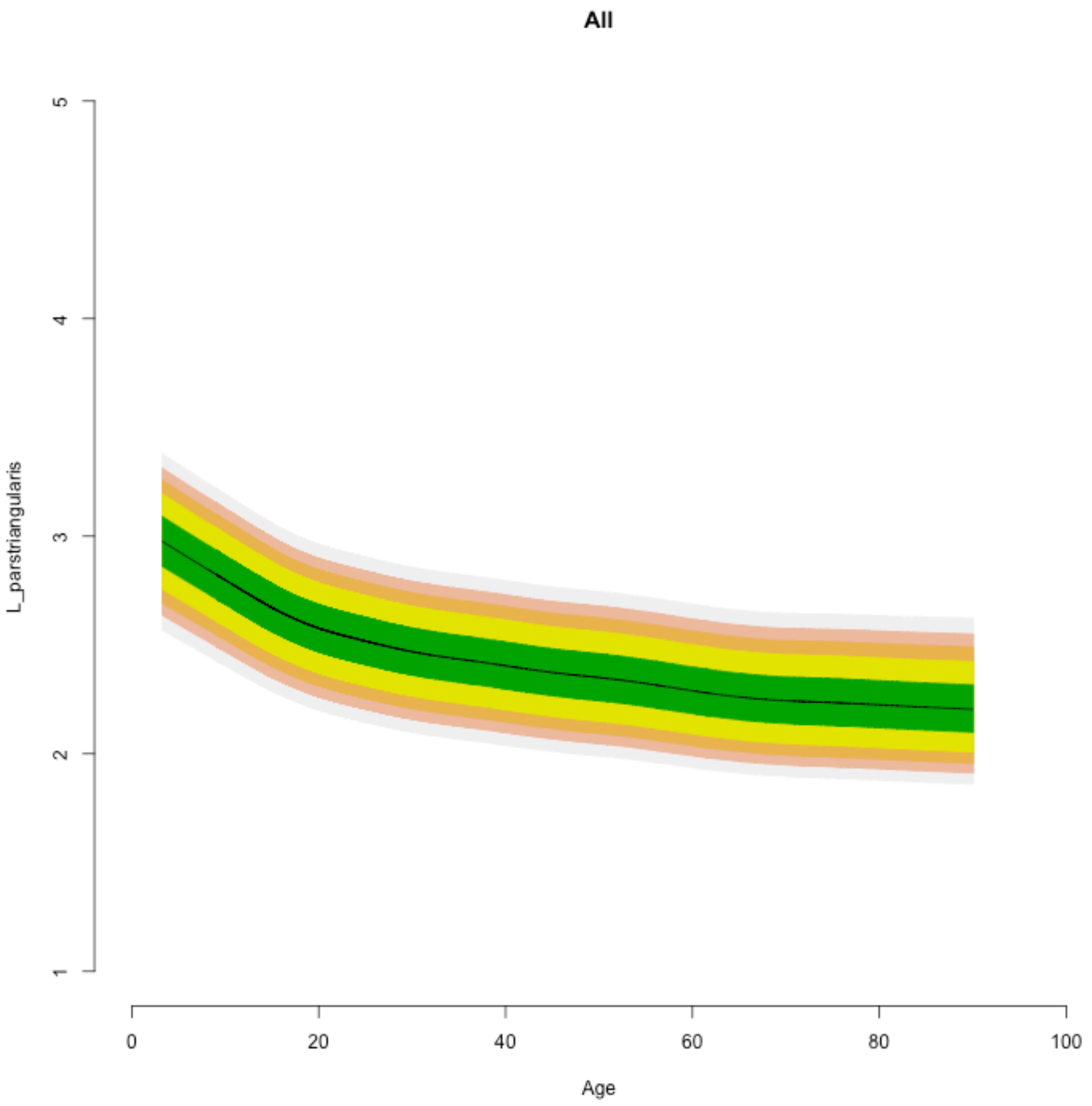
All
L_parstriangularis
1
2
3
4
5
0
20
40
60
80
100
Age

Female

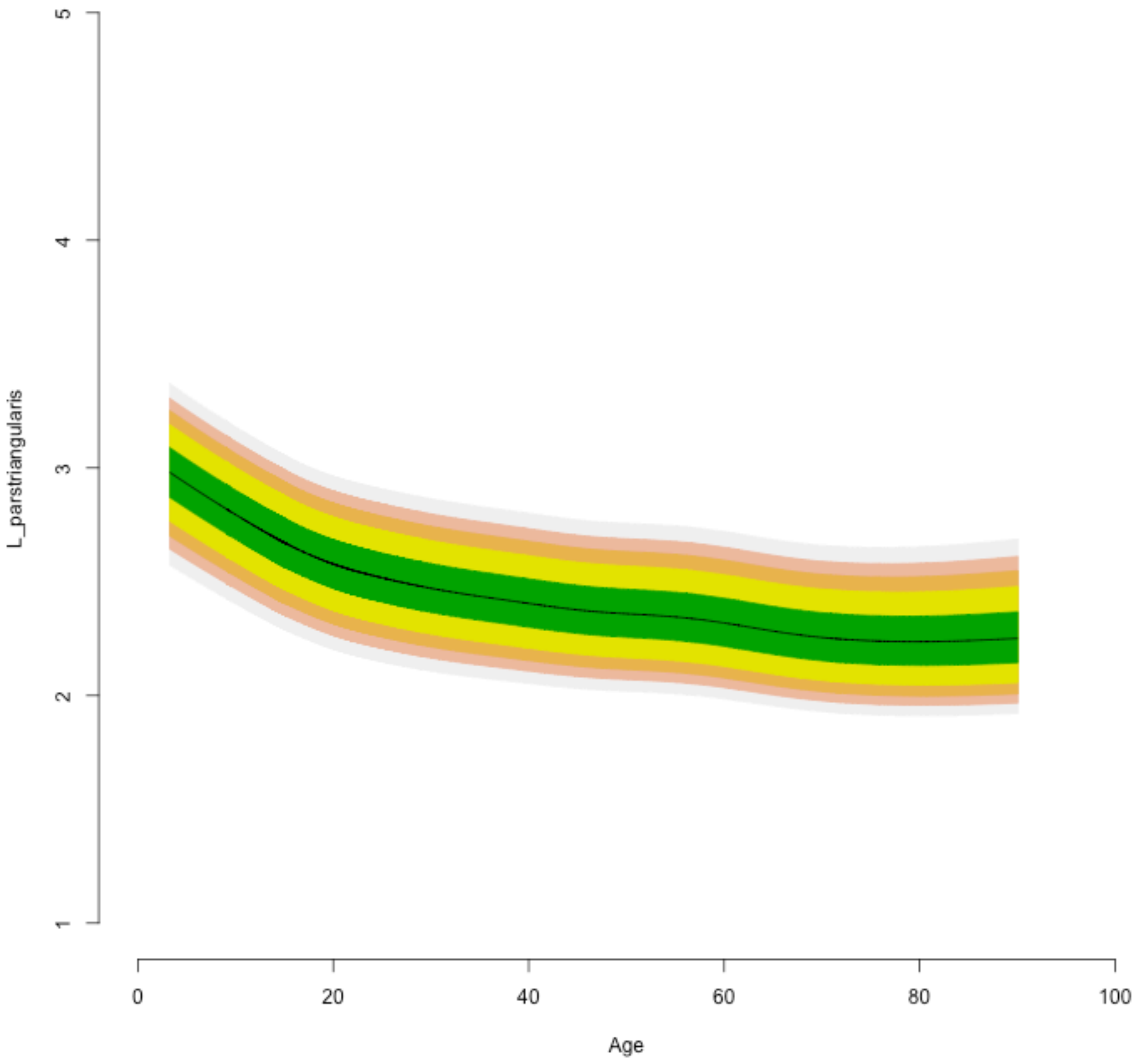

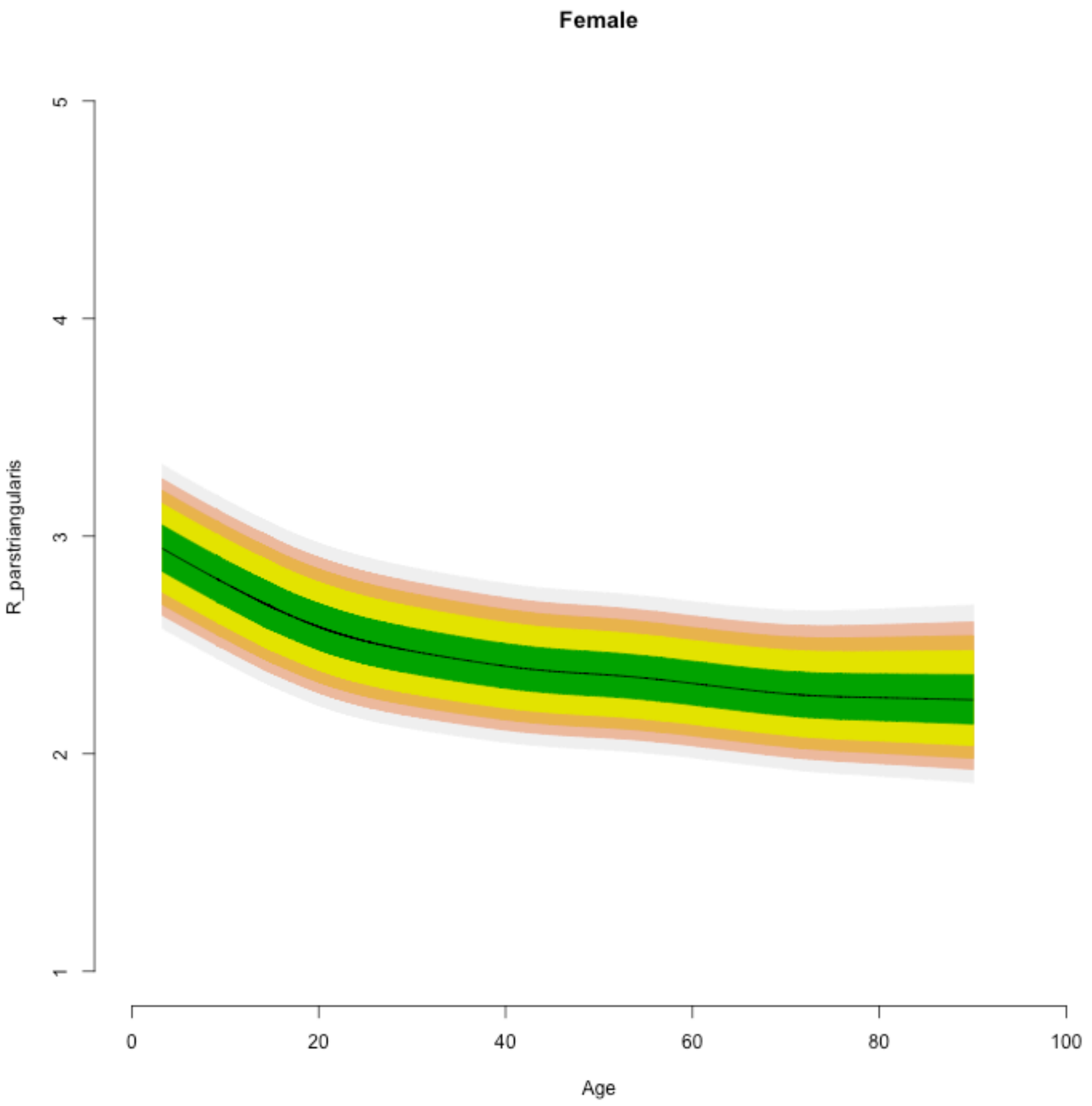

Female
R_parstriangularis
1
2
3
4
5
0
20
40
60
80
100
Age

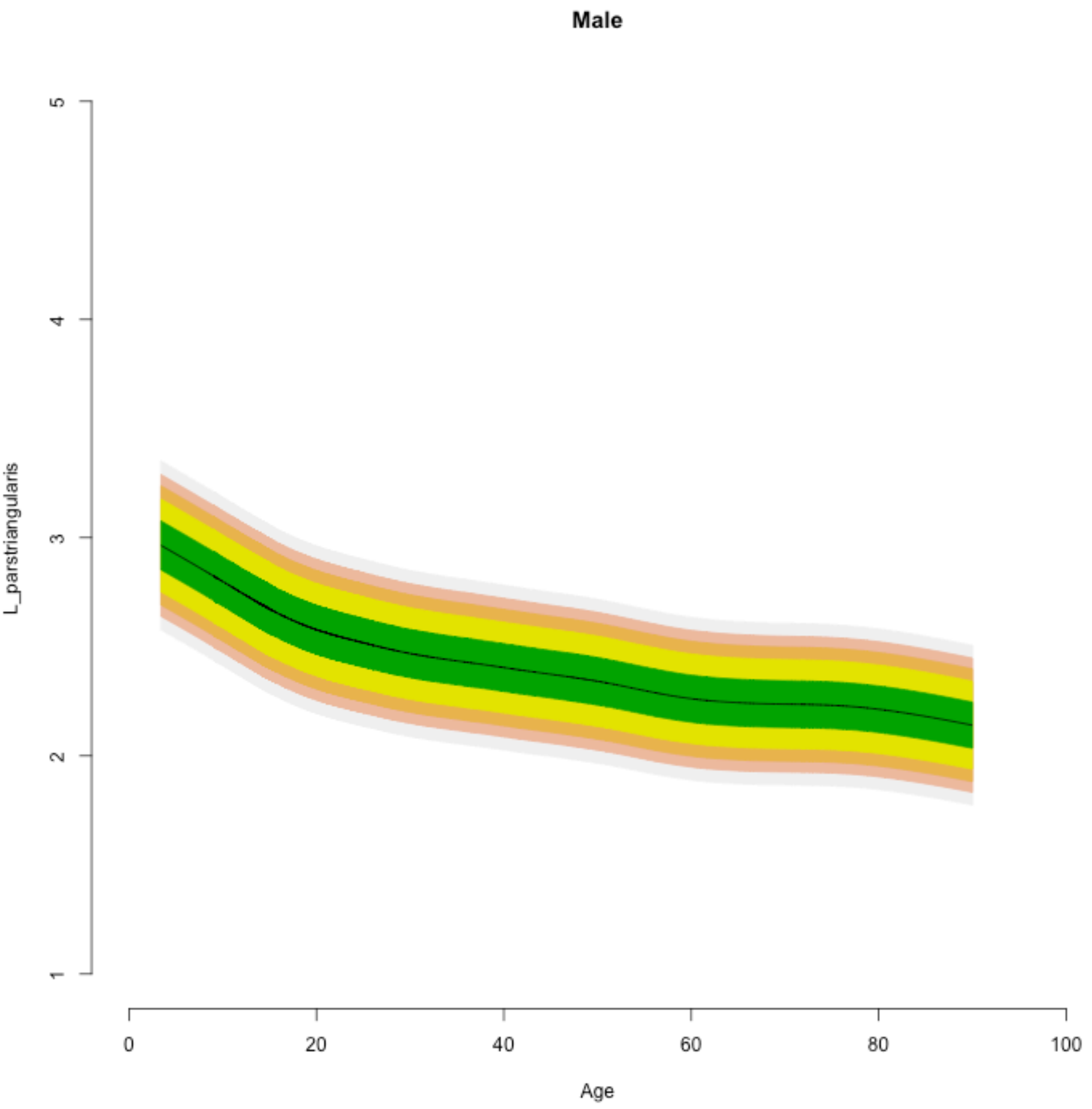

Male
5
4
3
2
1
L_parstriangularis
0
20
40
60
80
100
Age

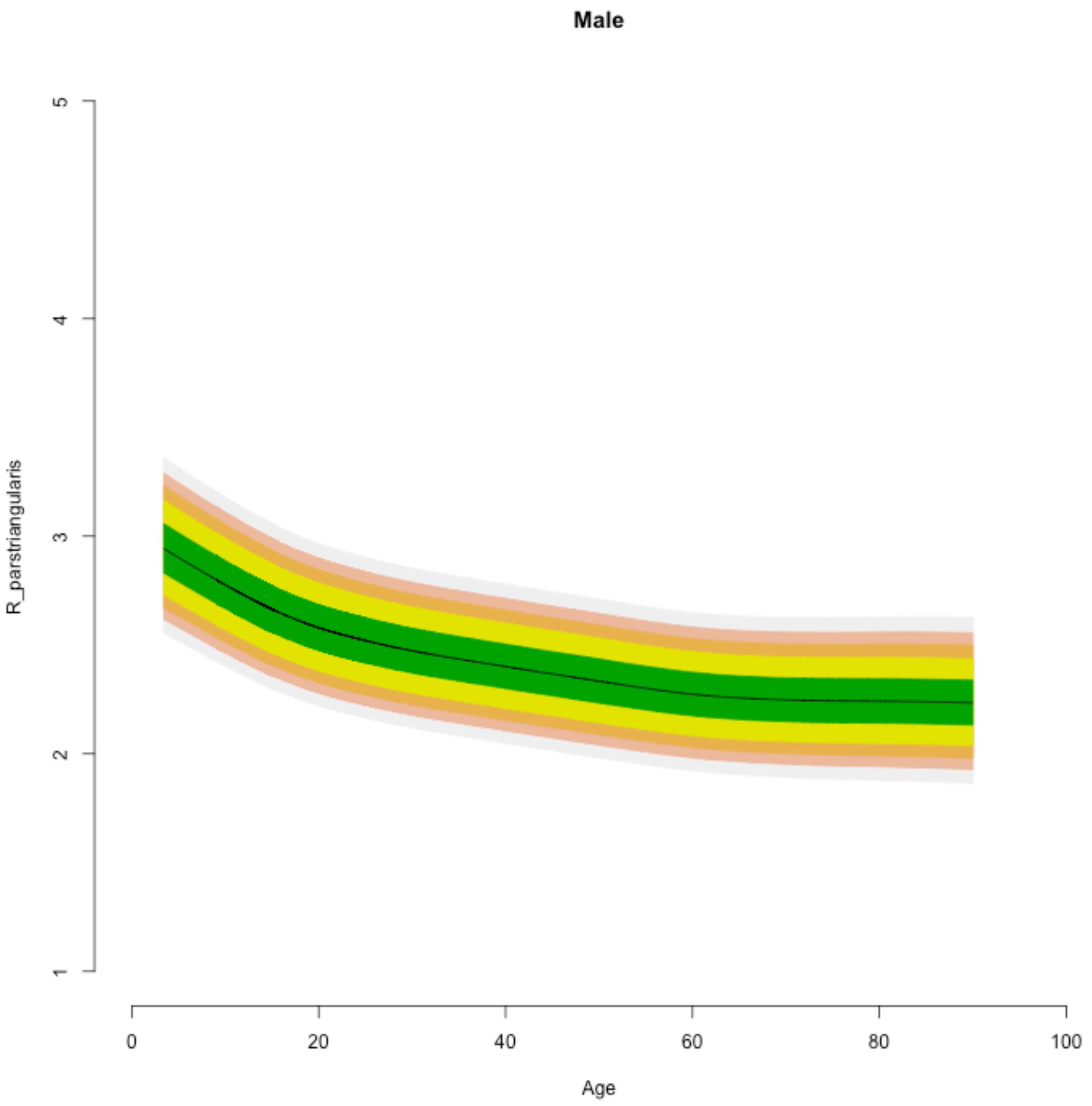
Male
R_parstriangularis
1
2
3
4
5
0
20
40
60
80
100
Age

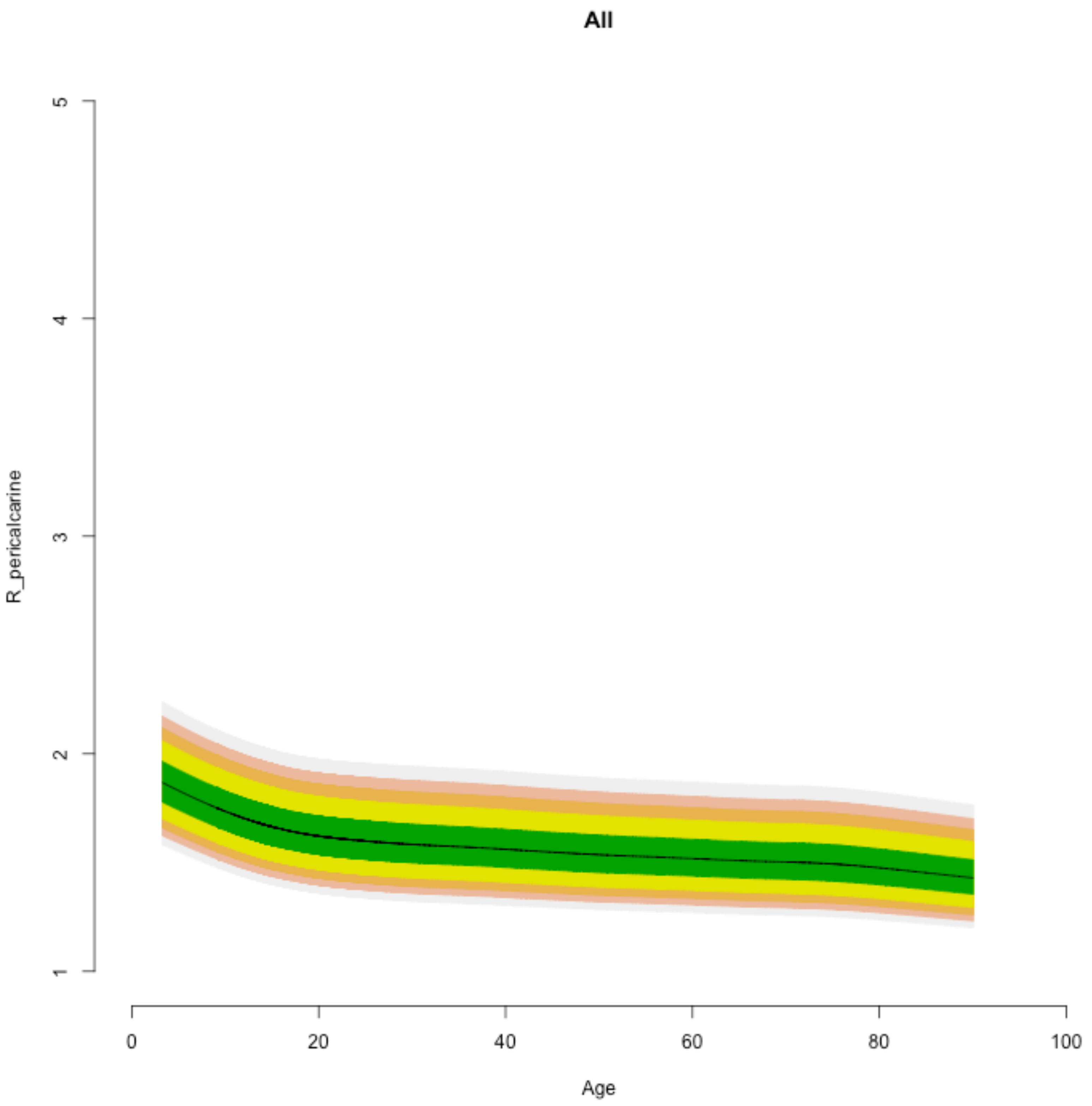

All
R_pericalcarine
1
2
3
4
5
0
20
40
60
80
100
Age

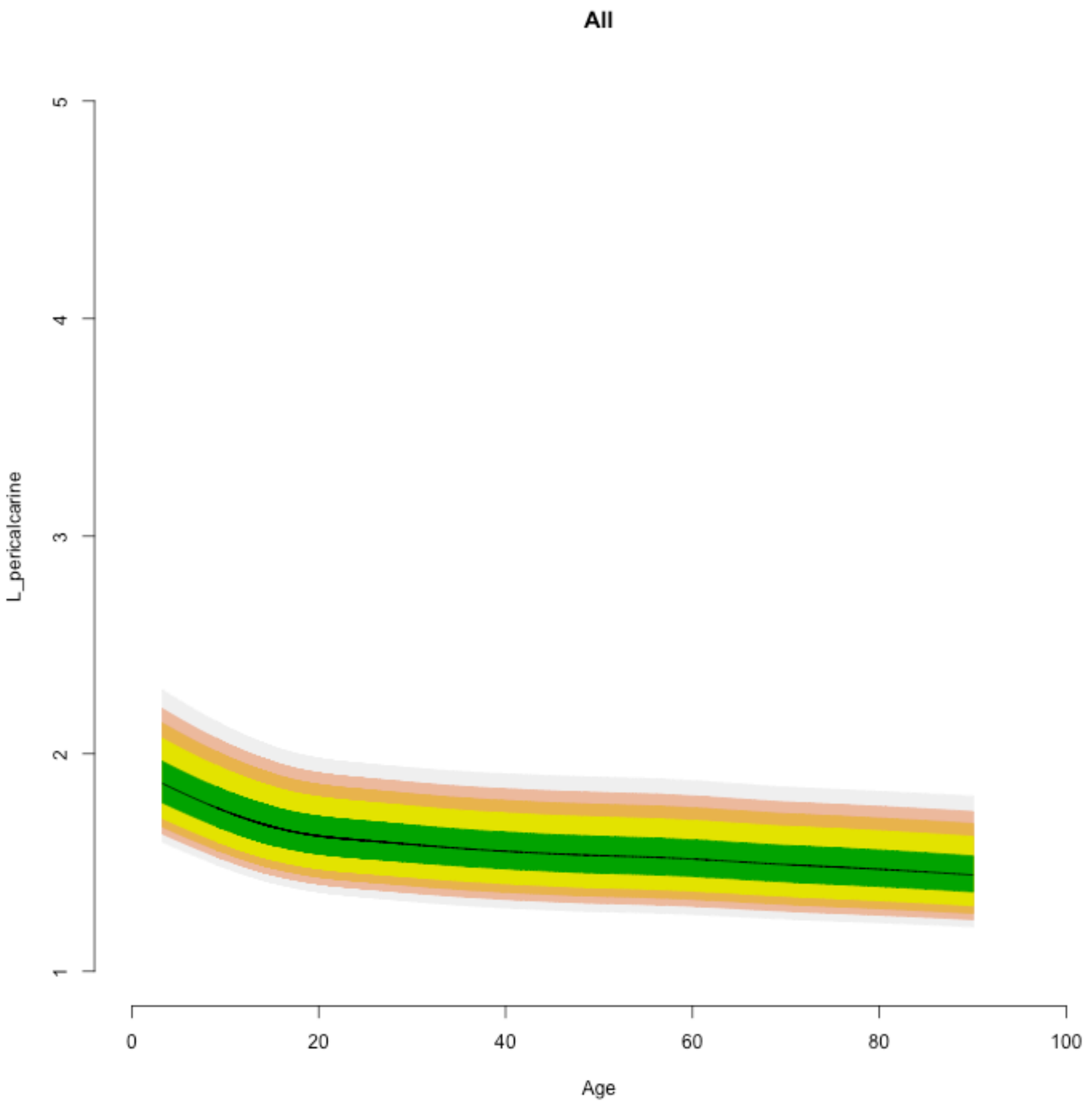

All
L_pericalcarine
5
4
3
2
1
0
20
40
60
80
100
Age

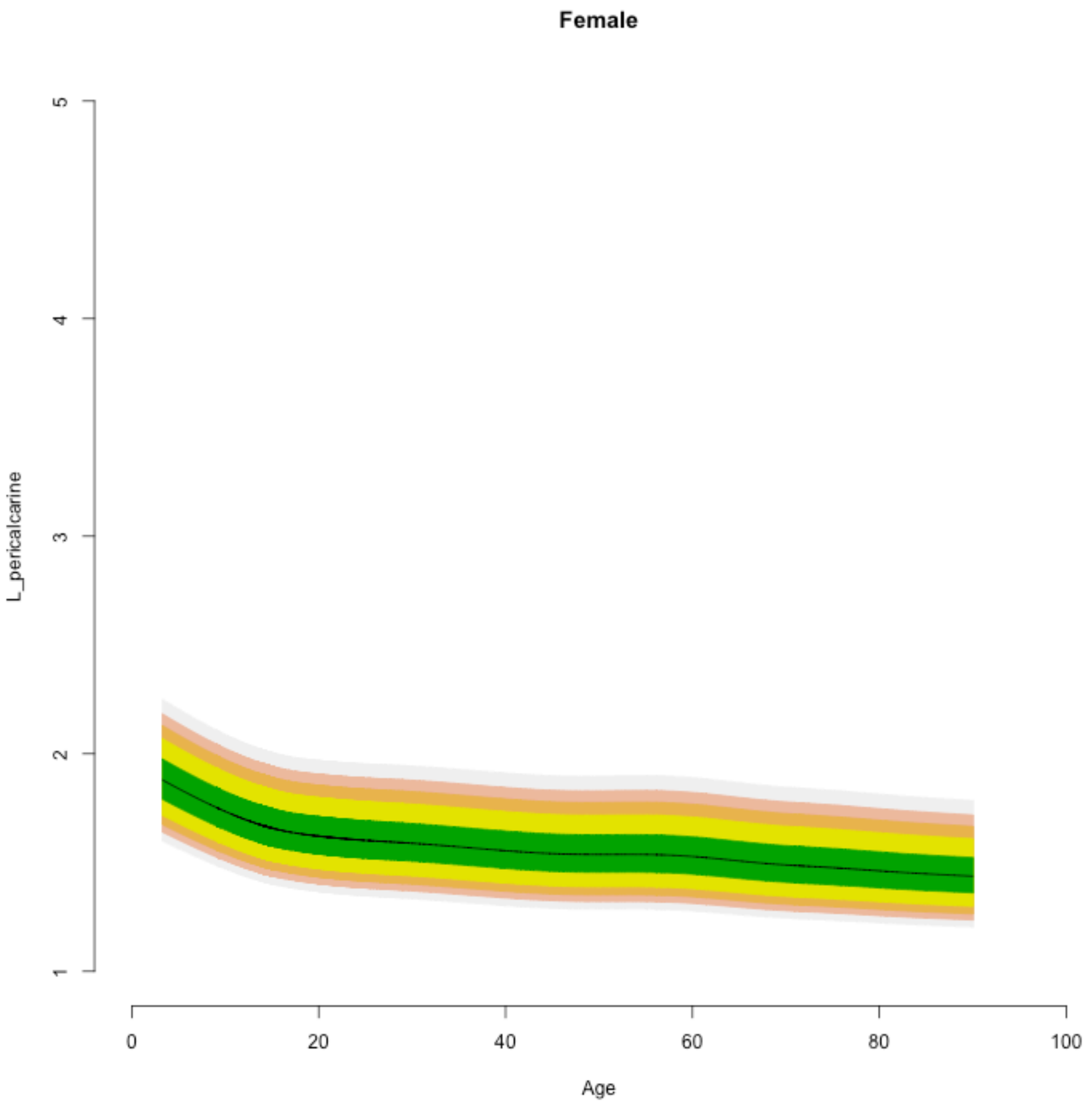

Female
L_pericalcarine
5
4
3
2
1
0
20
40
60
80
100
Age

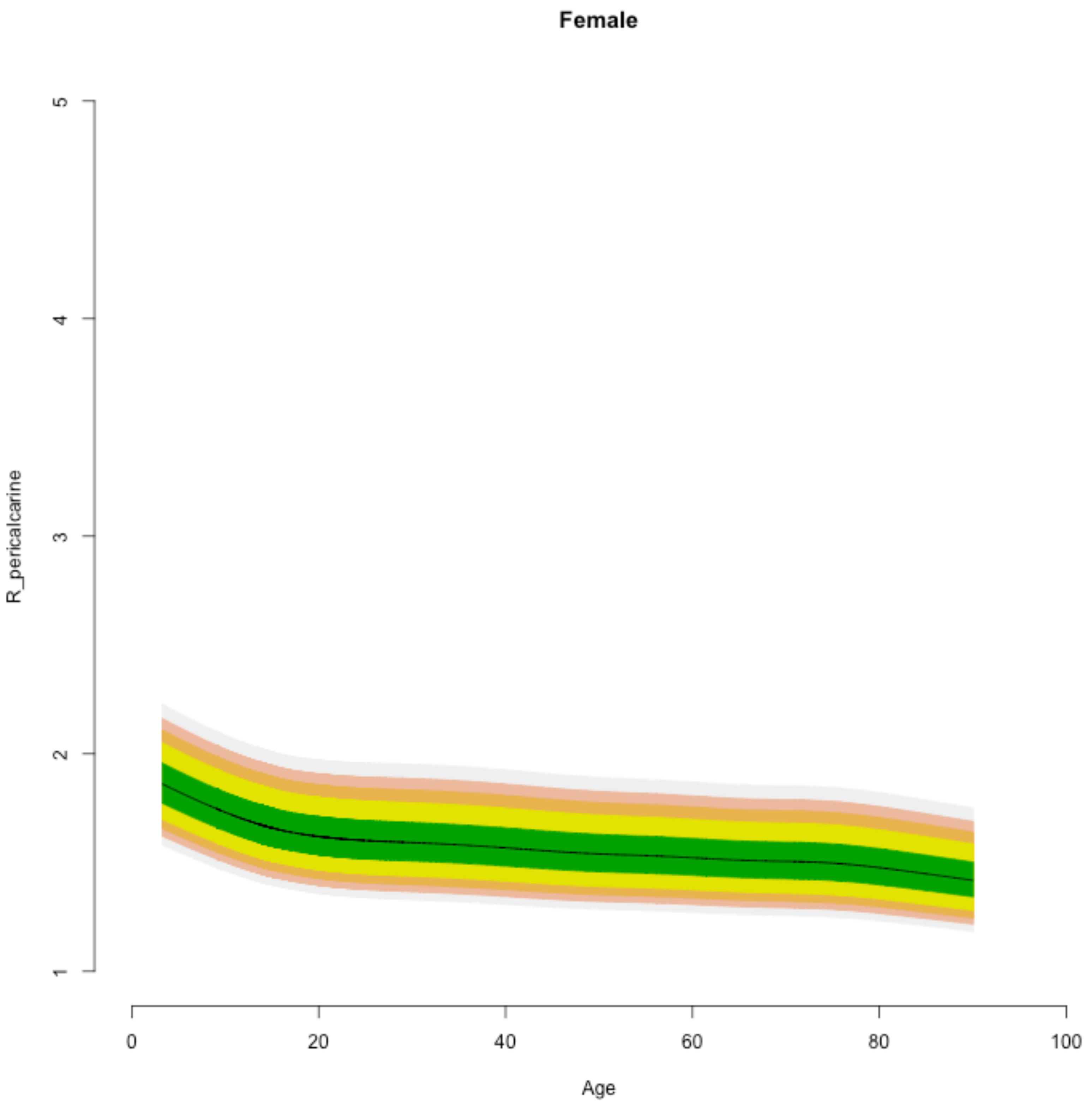
Female
R_pericalcarine
1
2
3
4
5
0
20
40
60
80
100
Age

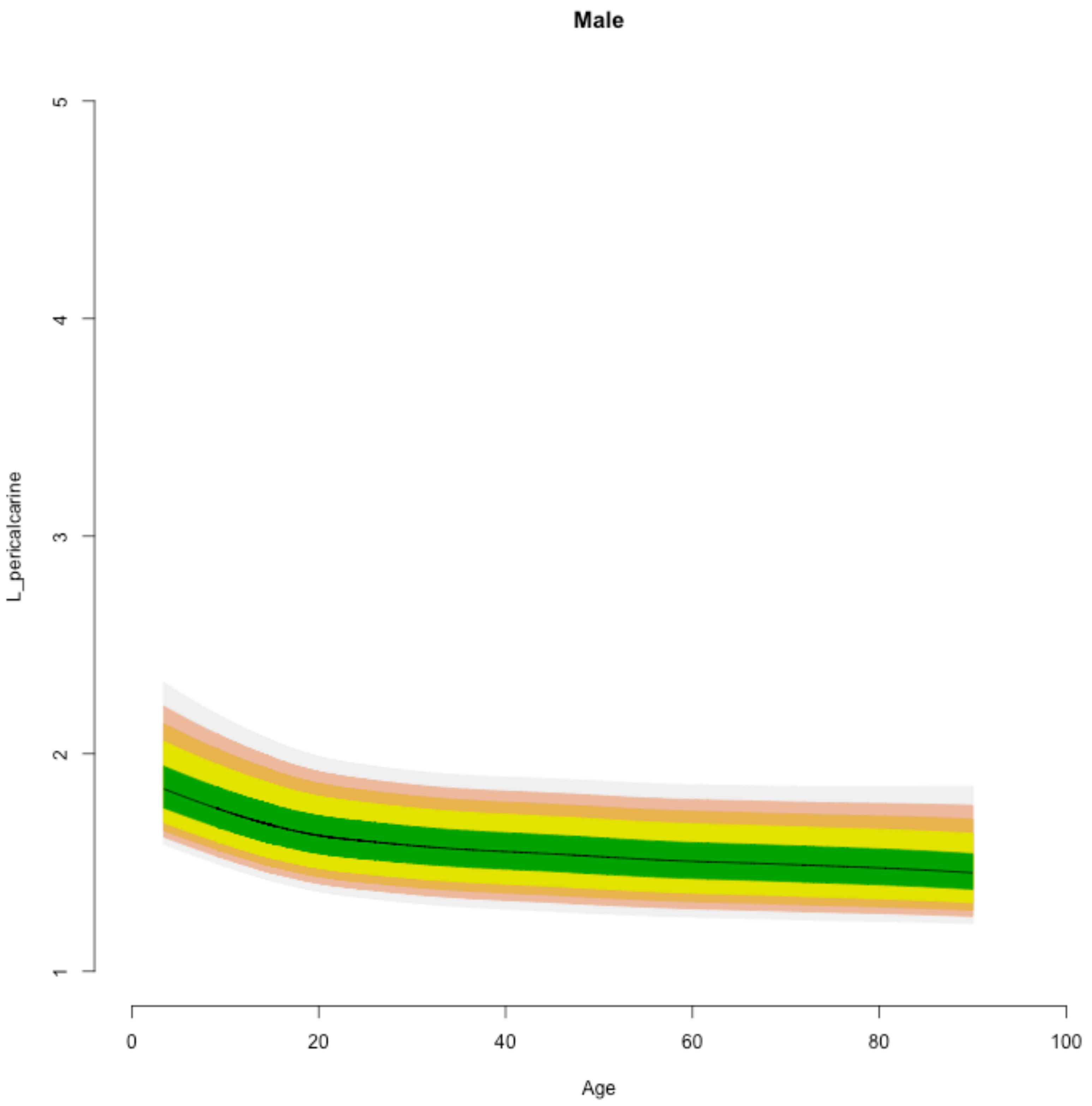

Male
L_pericalcarine
Age
1
2
3
4
5
0
20
40
60
80
100

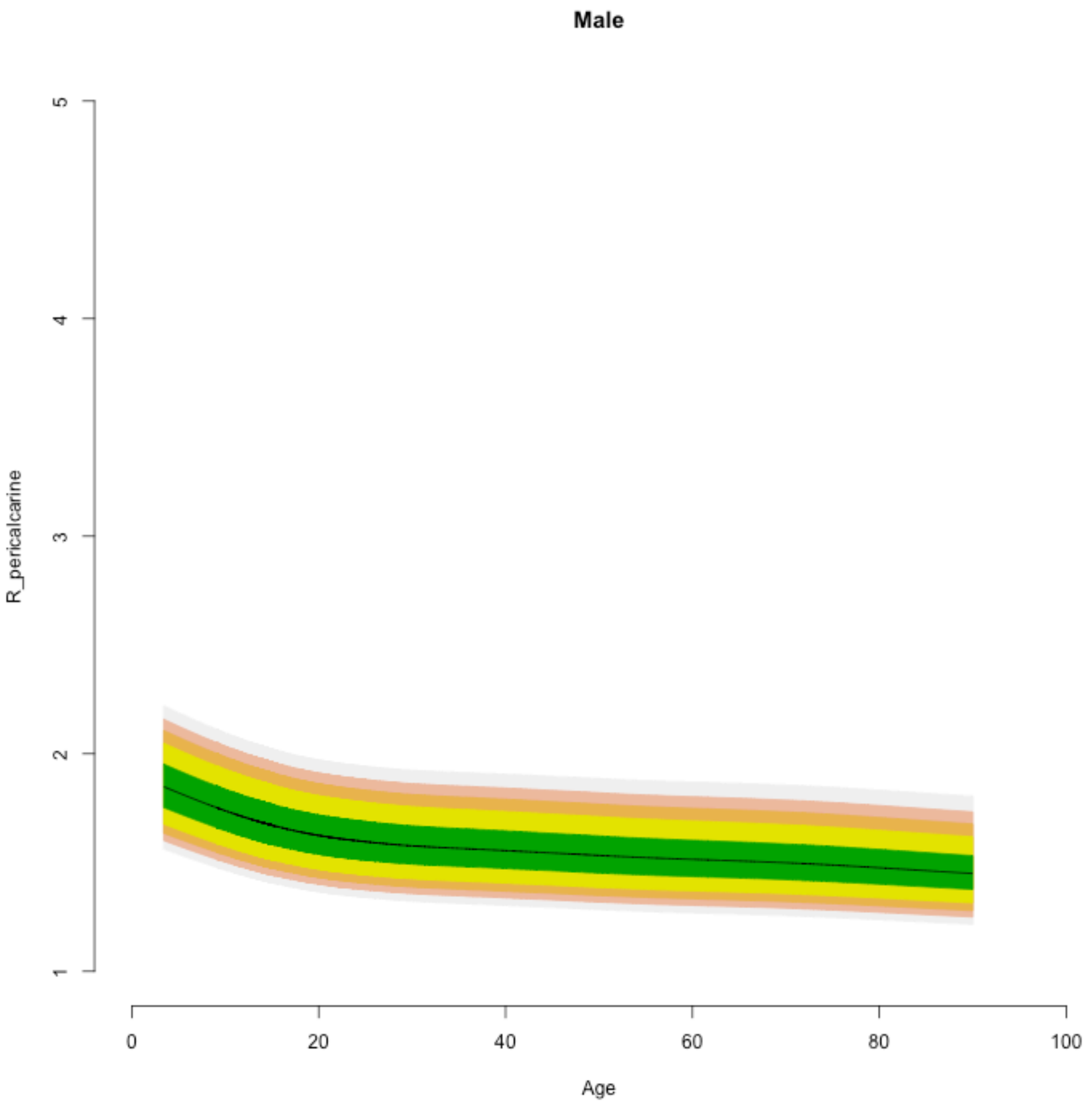
Male
R_pericalcarine
1
2
3
4
5
Age
0
20
40
60
80
100

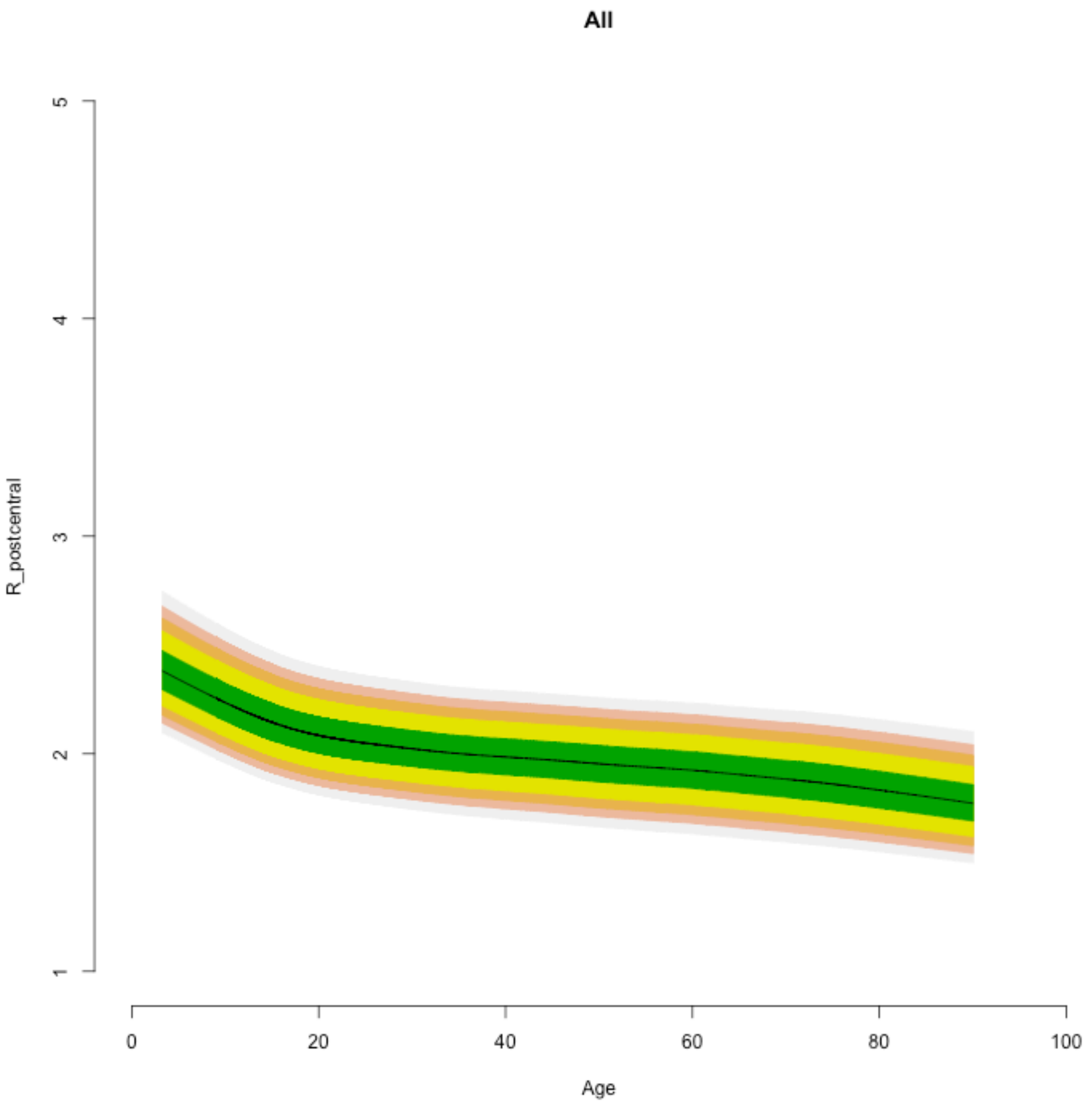

All
R_postcentral
5
4
3
2
1
0
20
40
60
80
100
Age

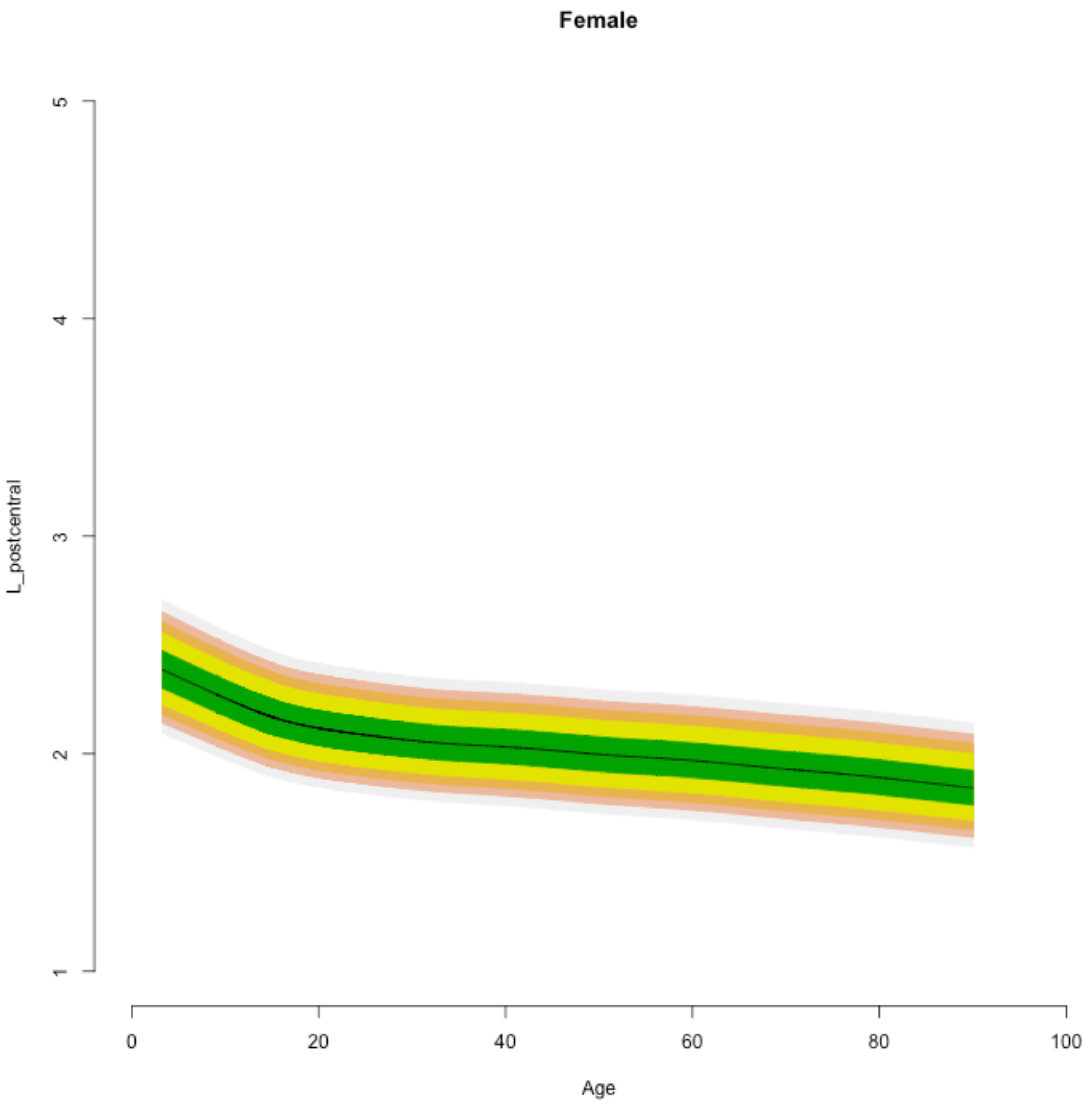
Female
L_postcentral
5
4
3
2
1
0
20
40
60
80
100
Age

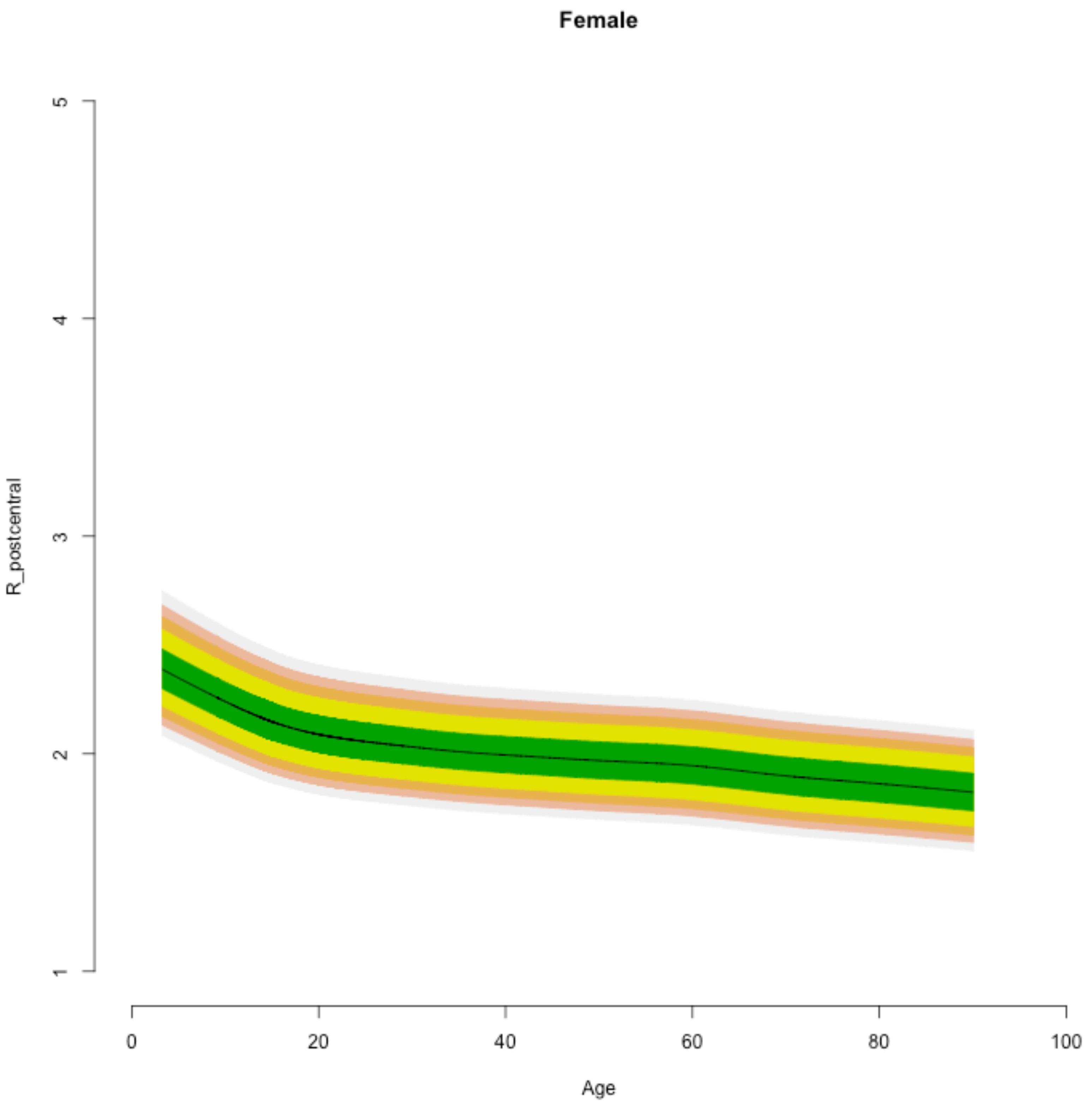

Female
R_postcentral
1
2
3
4
5
0
20
40
60
80
100
Age

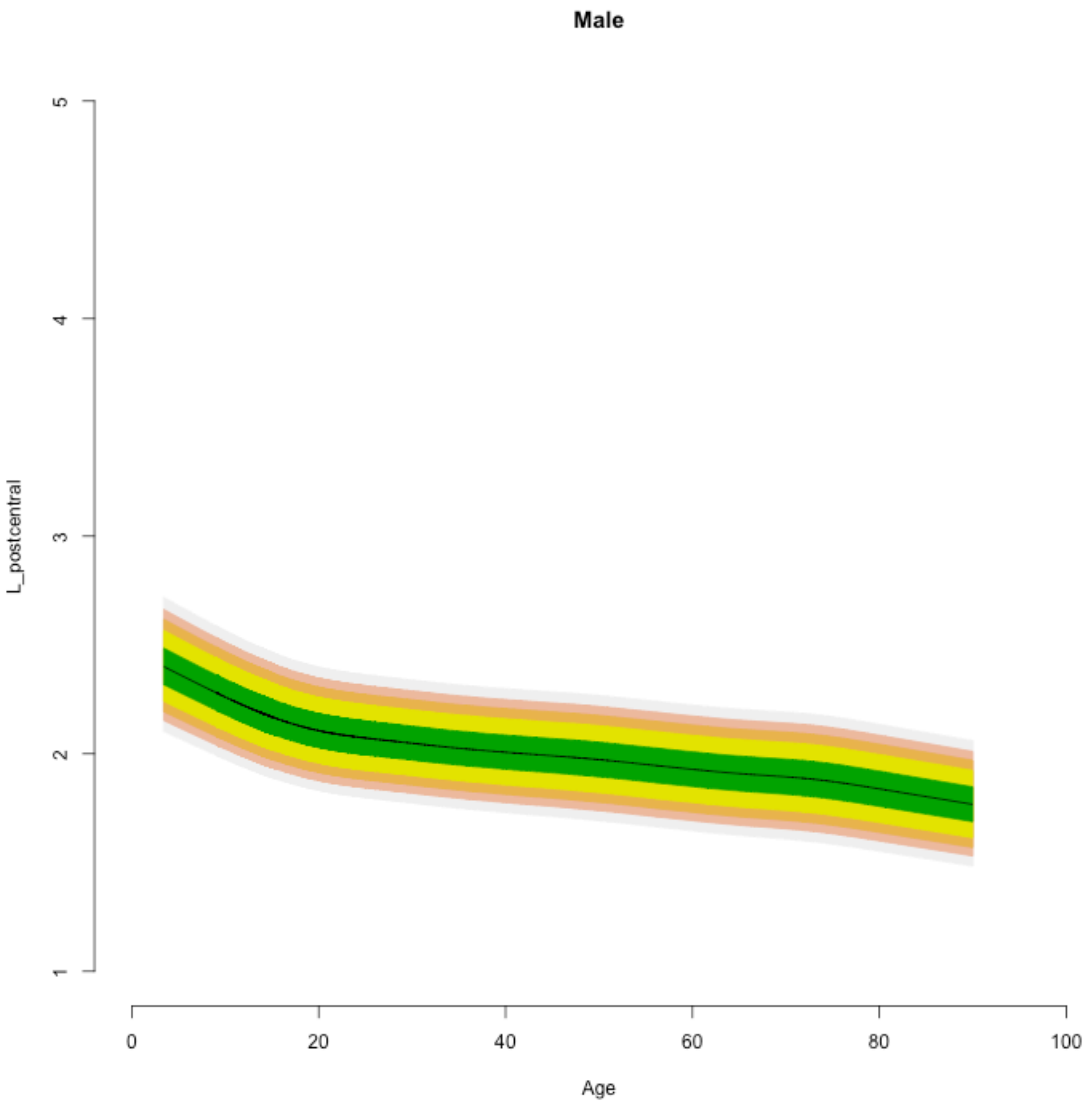

Male
L_postcentral
5
4
3
2
1
0
20
40
60
80
100
Age

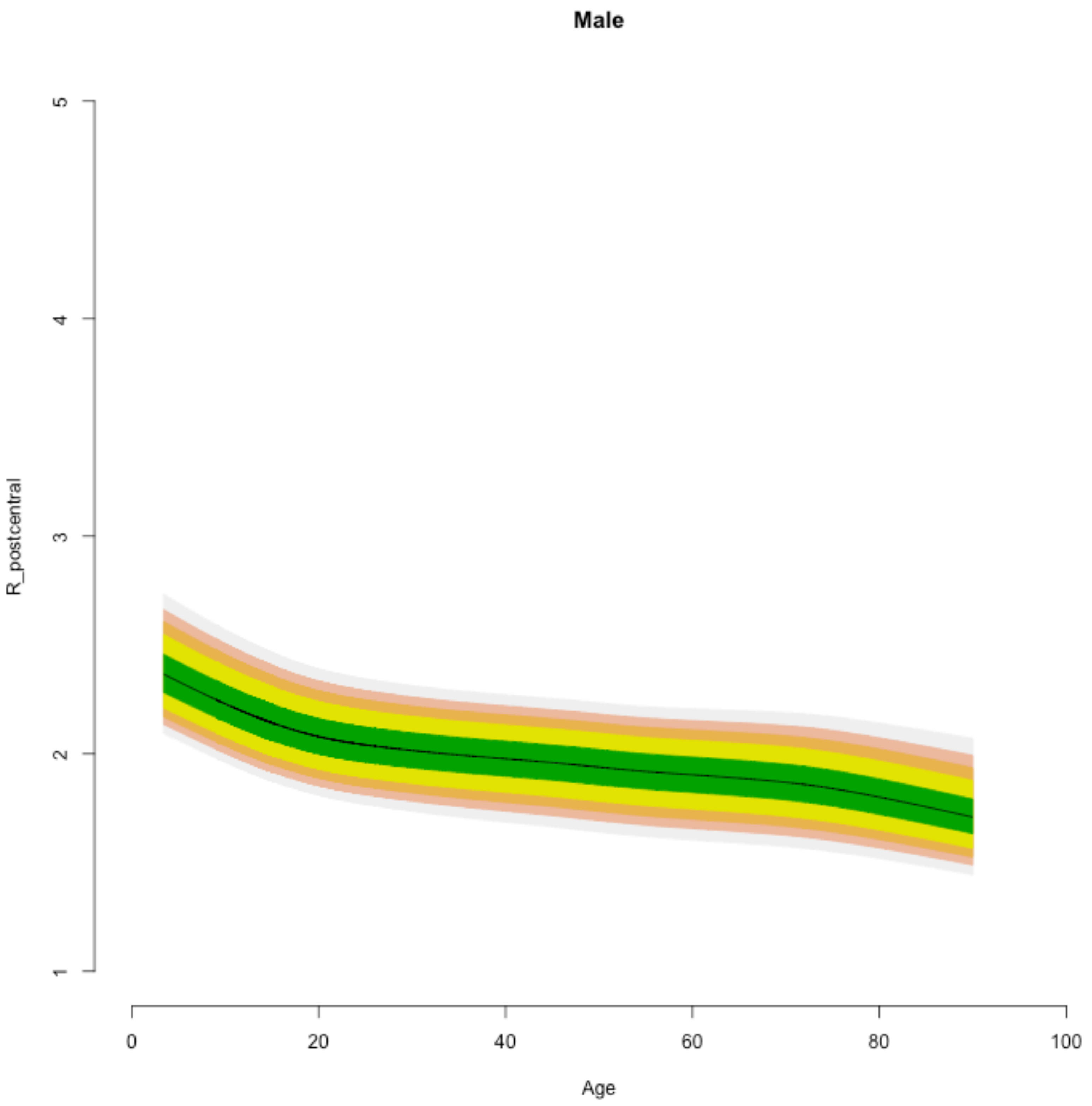

Male
R_postcentral
1
2
3
4
5
0
20
40
60
80
100
Age

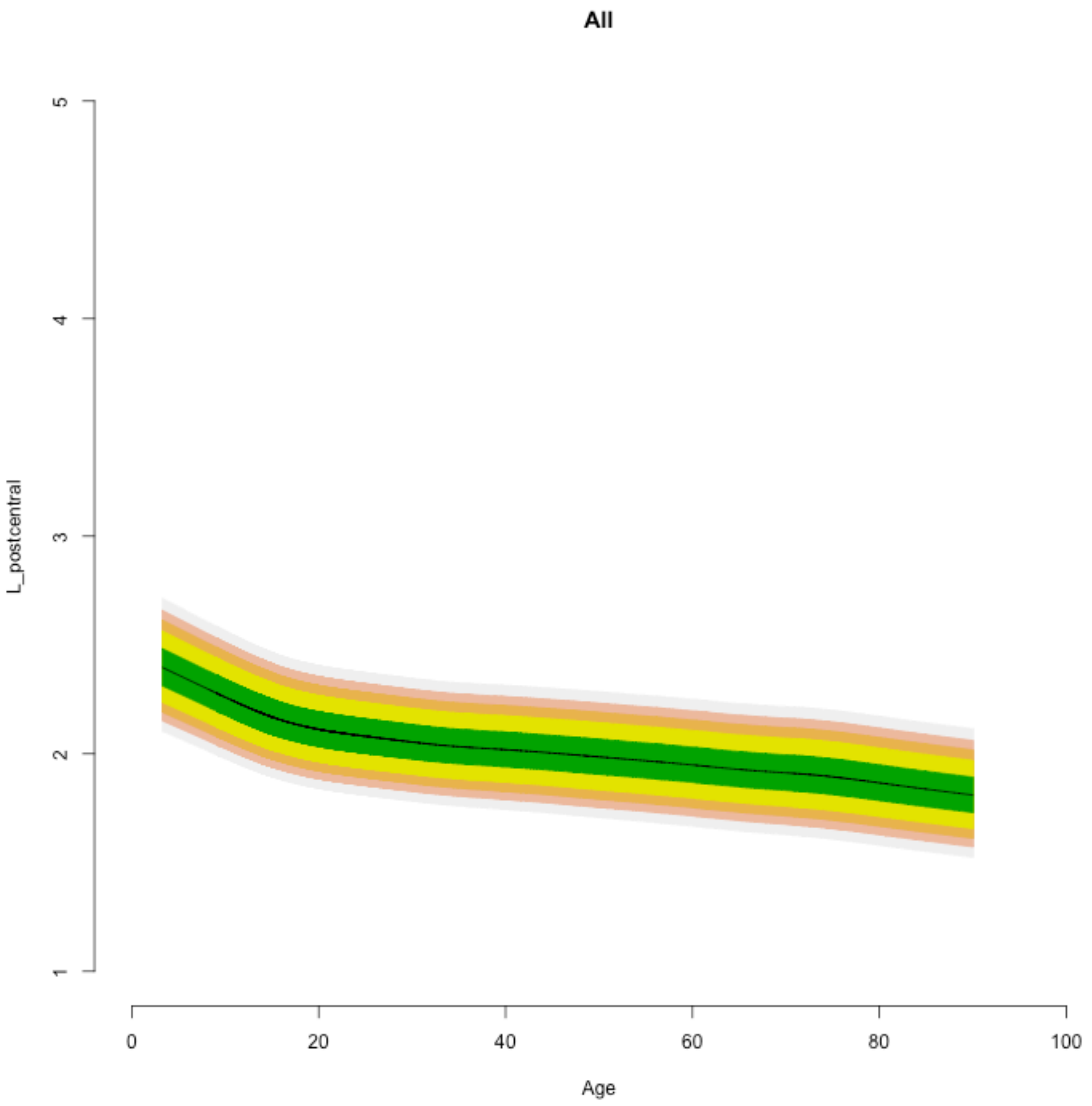
All
L_postcentral
1
2
3
4
5
0
20
40
60
80
100
Age

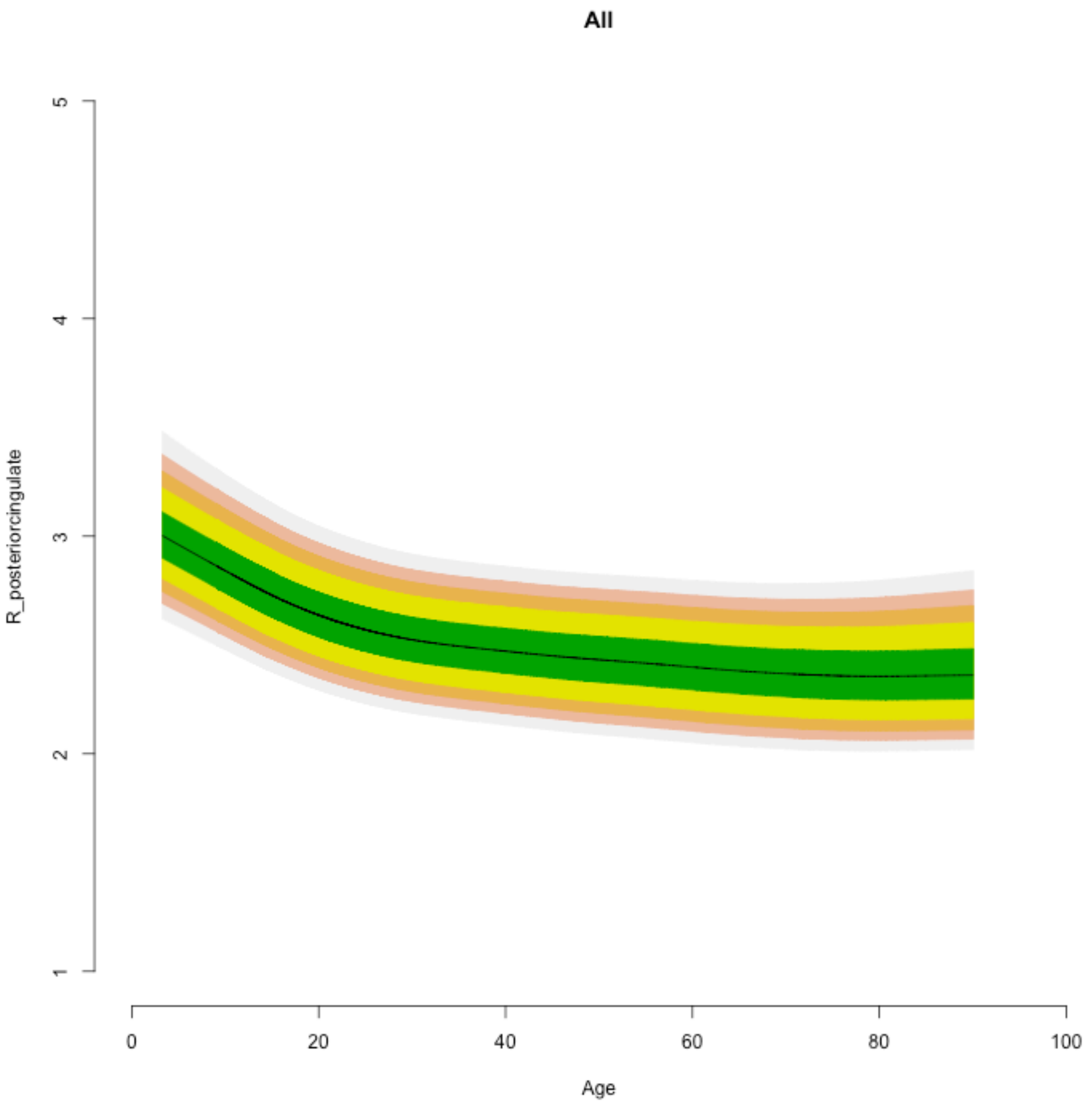
All
R_posteriorcingulate
5
4
3
2
1
0
20
40
60
80
100
Age

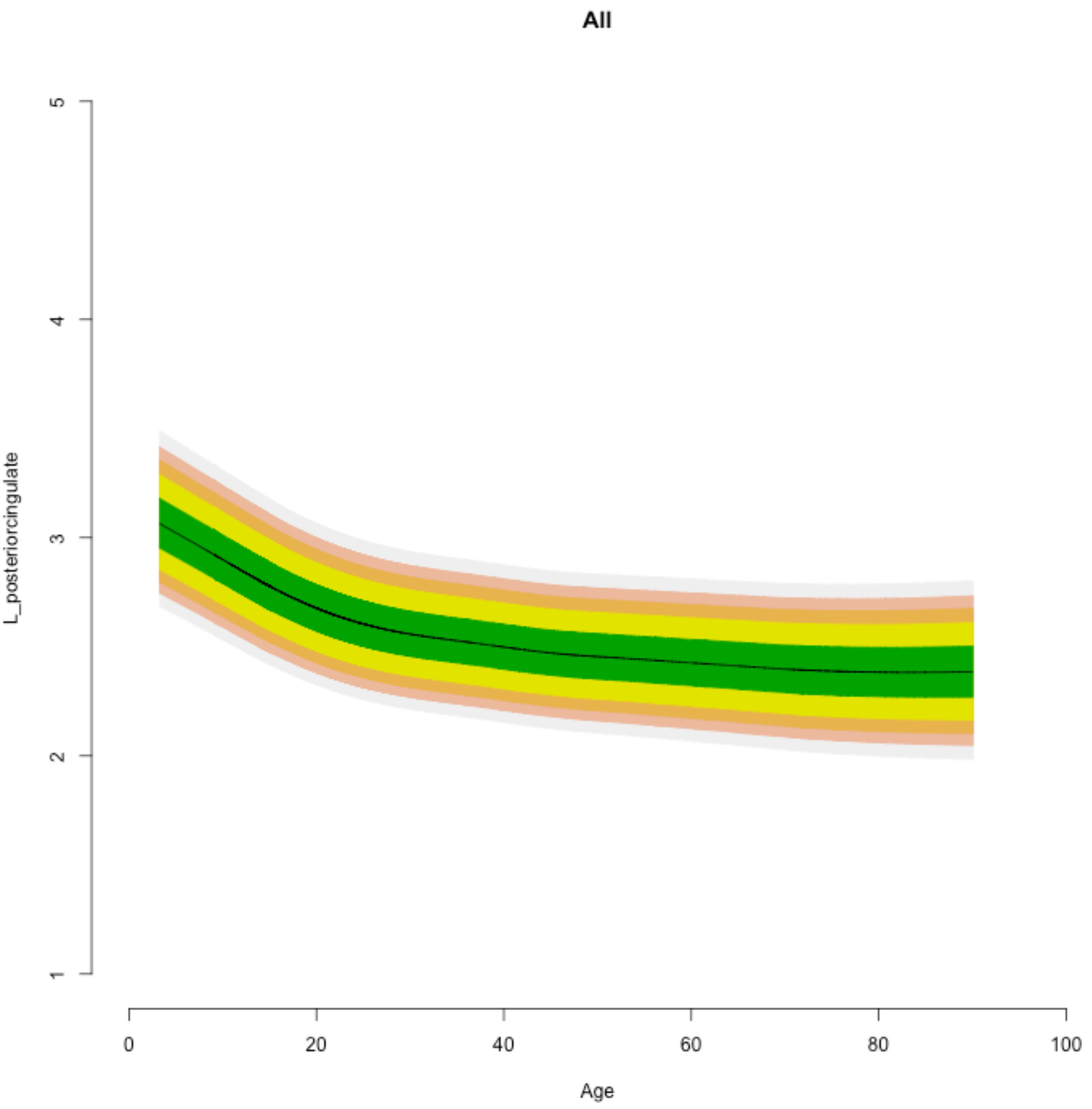
All
L_posteriorcingulate
1
2
3
4
5
0
20
40
60
80
100
Age

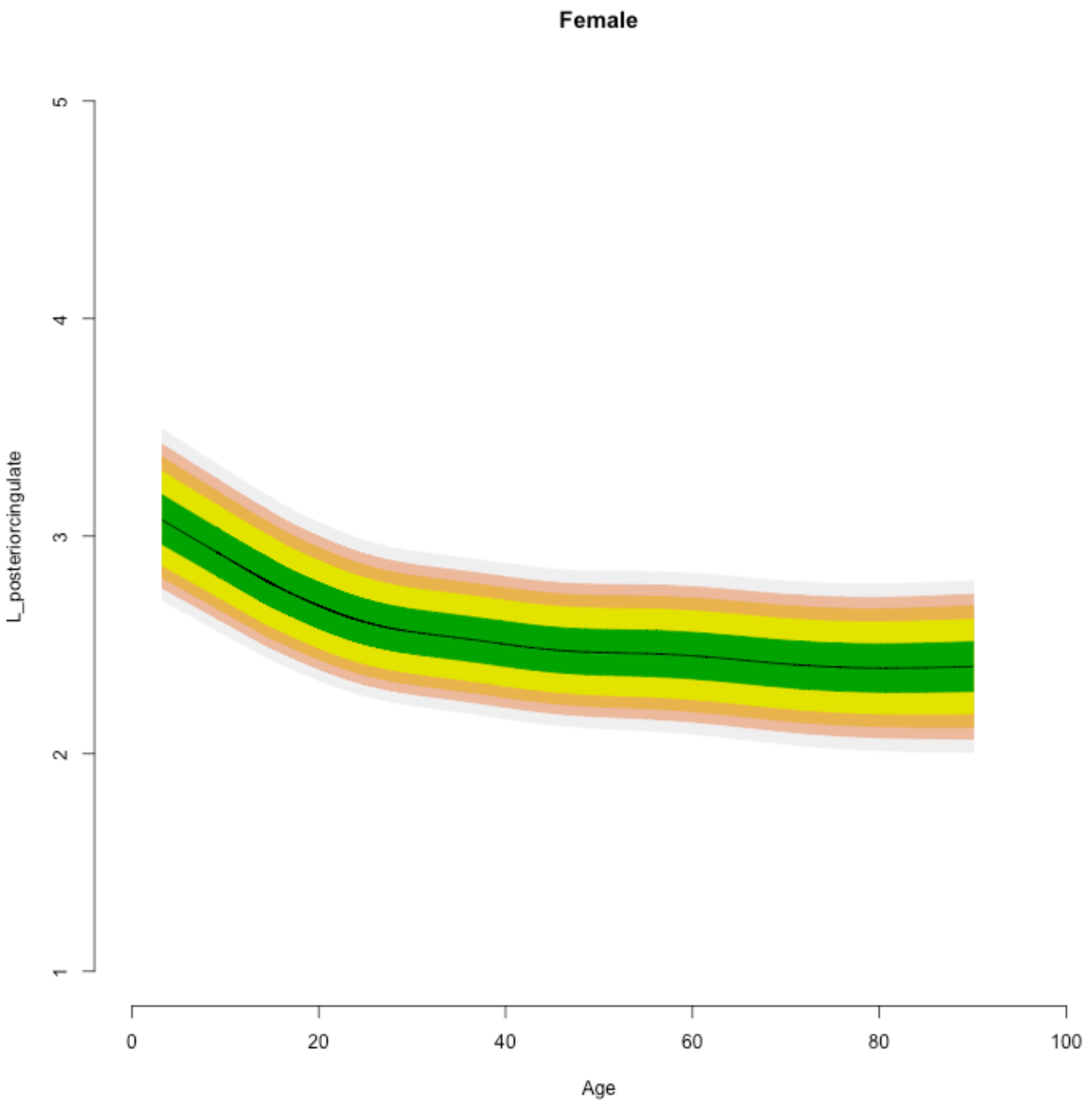

Female
L_posteriorcingulate
1
2
3
4
5
0
20
40
60
80
100
Age

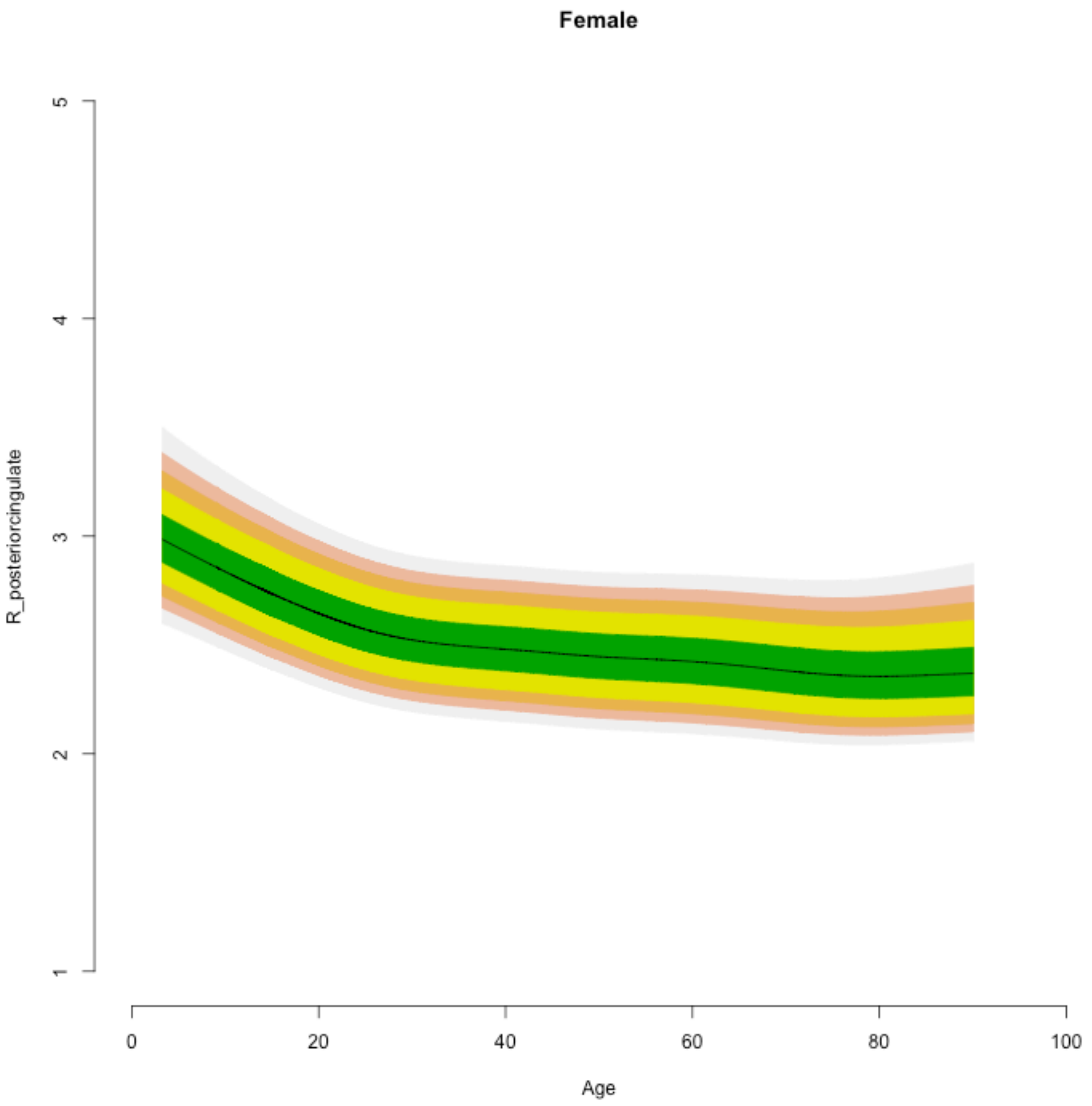
Female
R_posteriorcingulate
5
4
3
2
1
0
20
40
60
80
100
Age

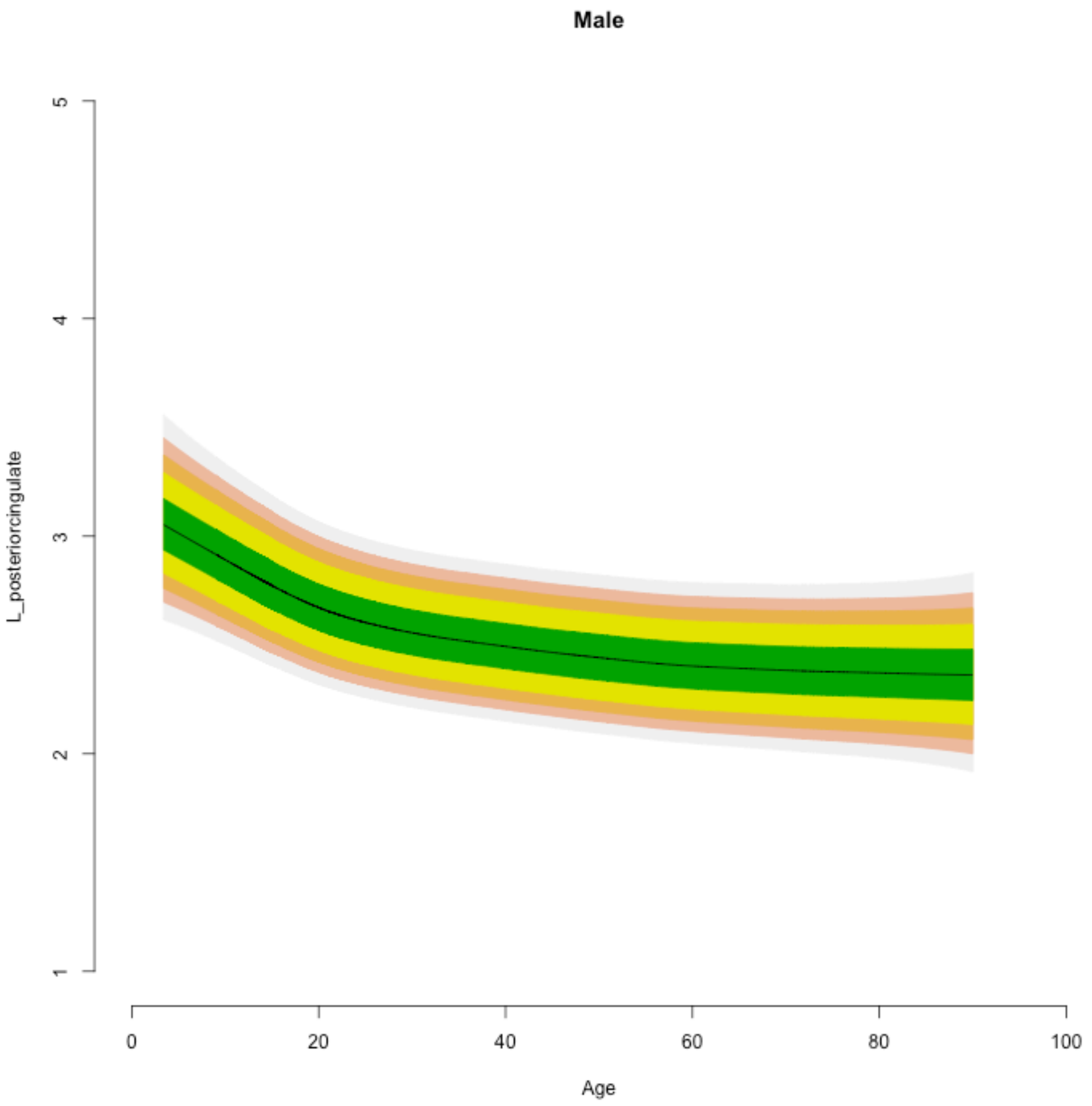

Male
L_posteriorcingulate
1
2
3
4
5
0
20
40
60
80
100
Age

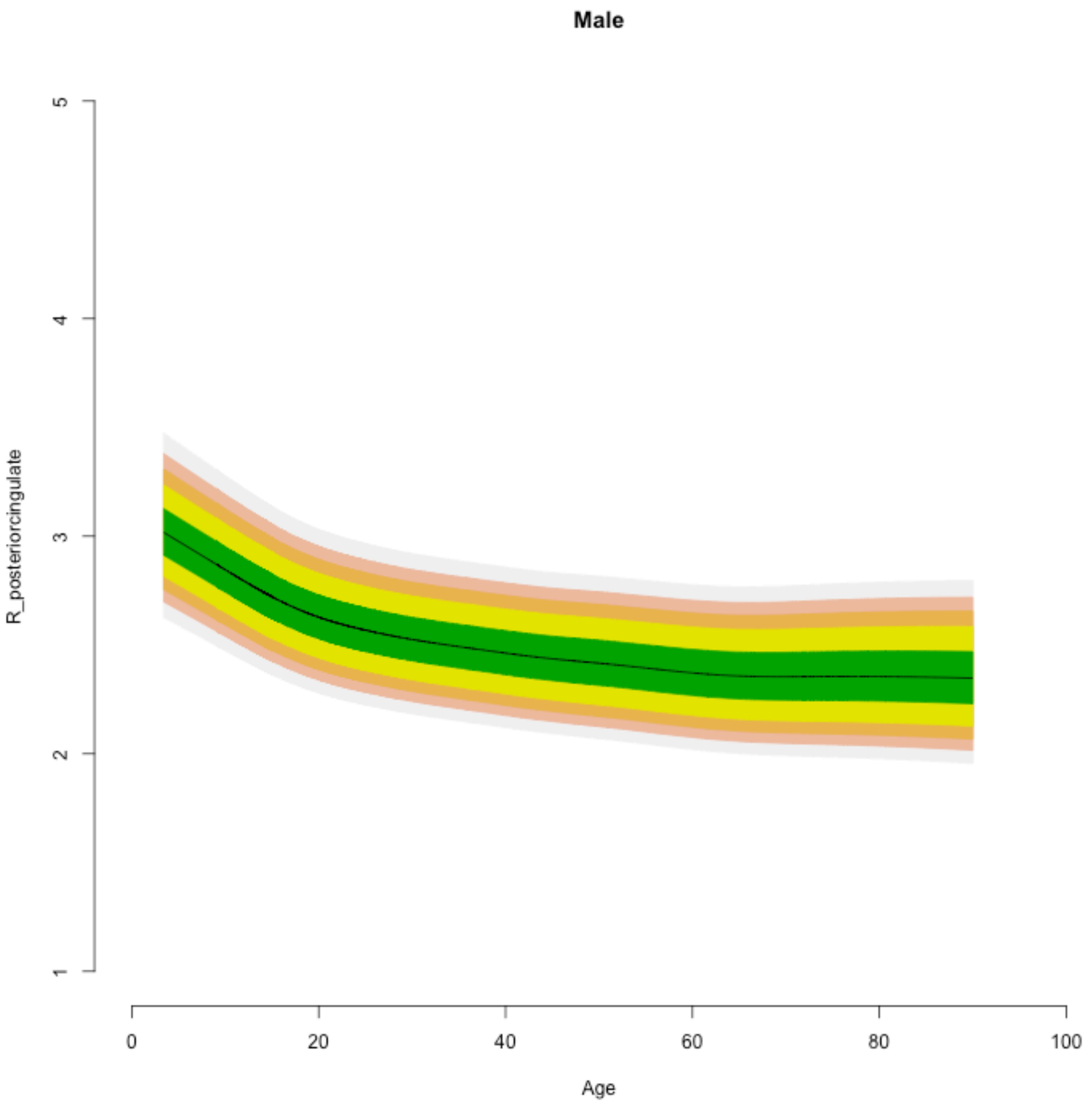
Male
R_posteriorcingulate
1
2
3
4
5
0
20
40
60
80
100
Age

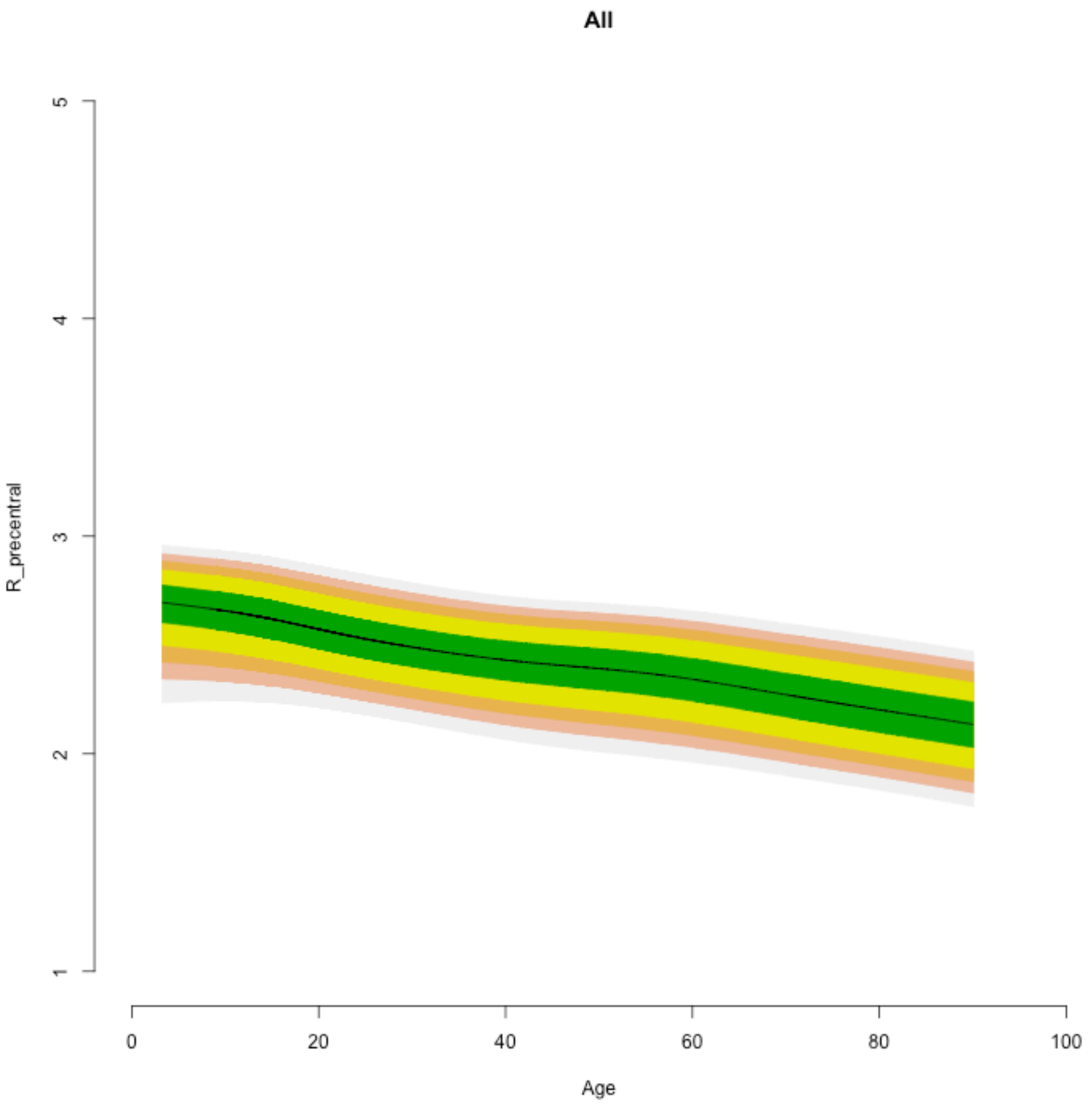
All
R_precentral
1
2
3
4
5
0
20
40
60
80
100
Age

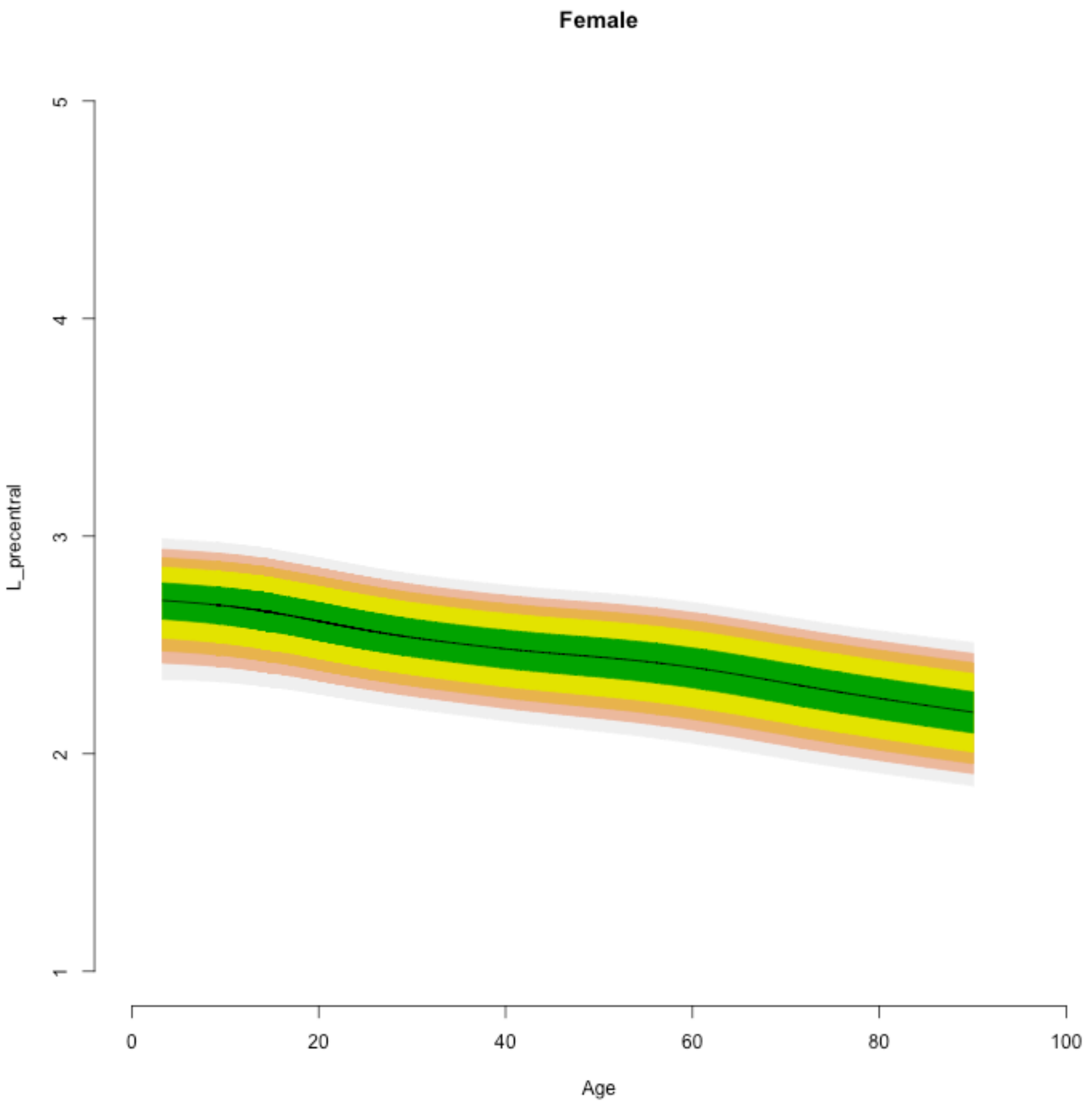

Female
L_precentral
1
2
3
4
5
0
20
40
60
80
100
Age

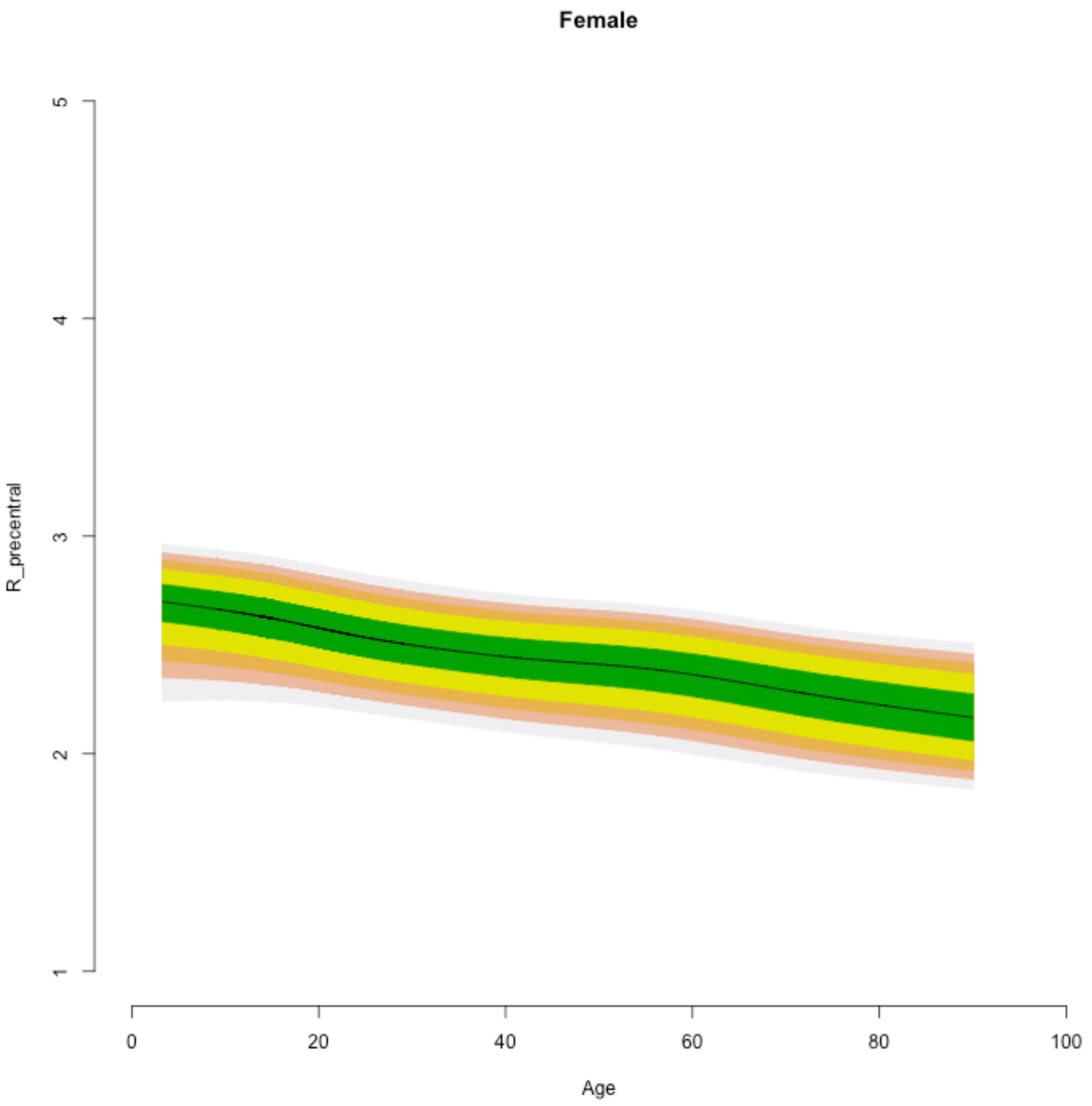

Female
R_precentral
1
2
3
4
5
0
20
40
60
80
100
Age

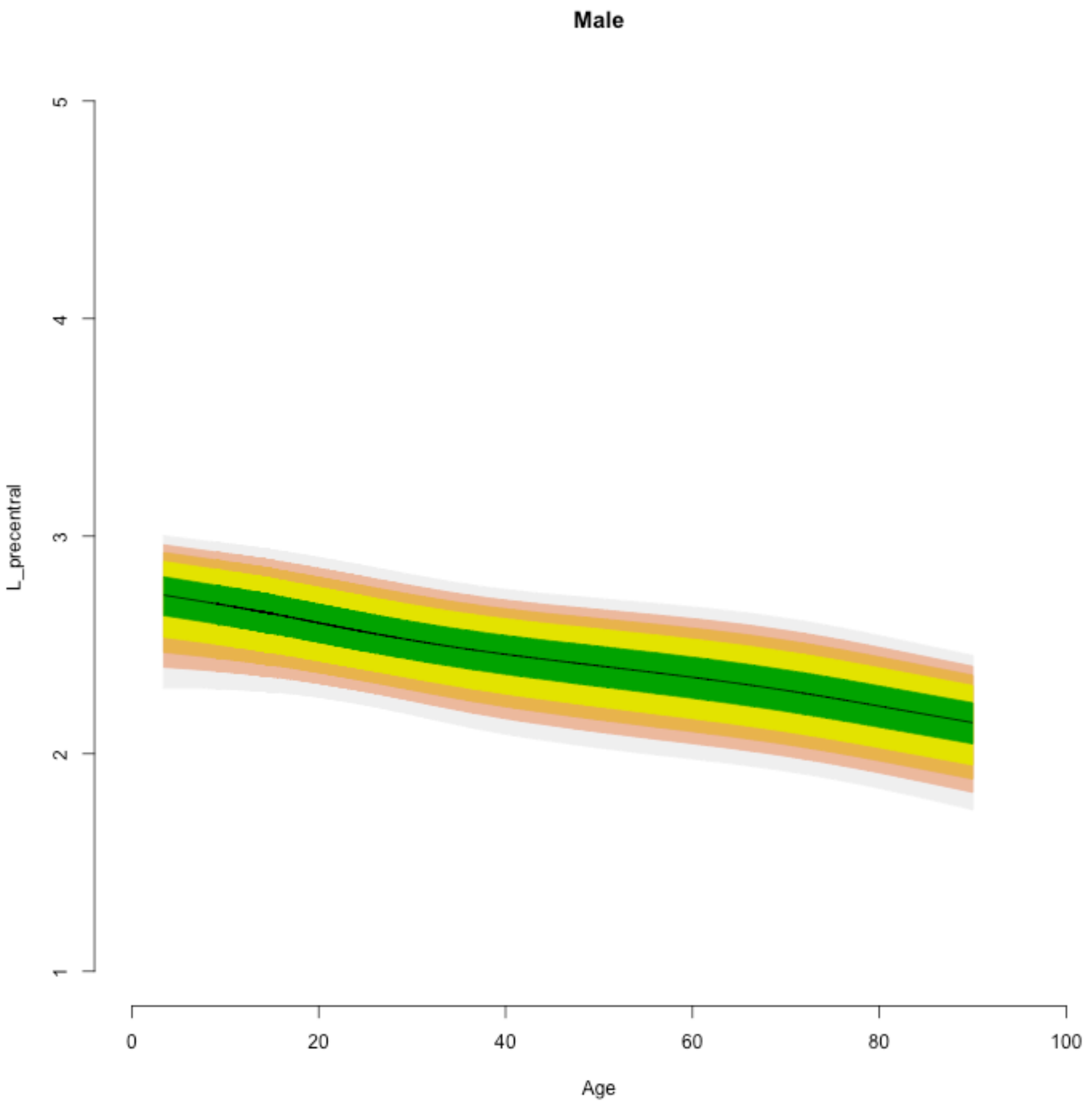

Male
L_precentral
1
2
3
4
5
0
20
40
60
80
100
Age

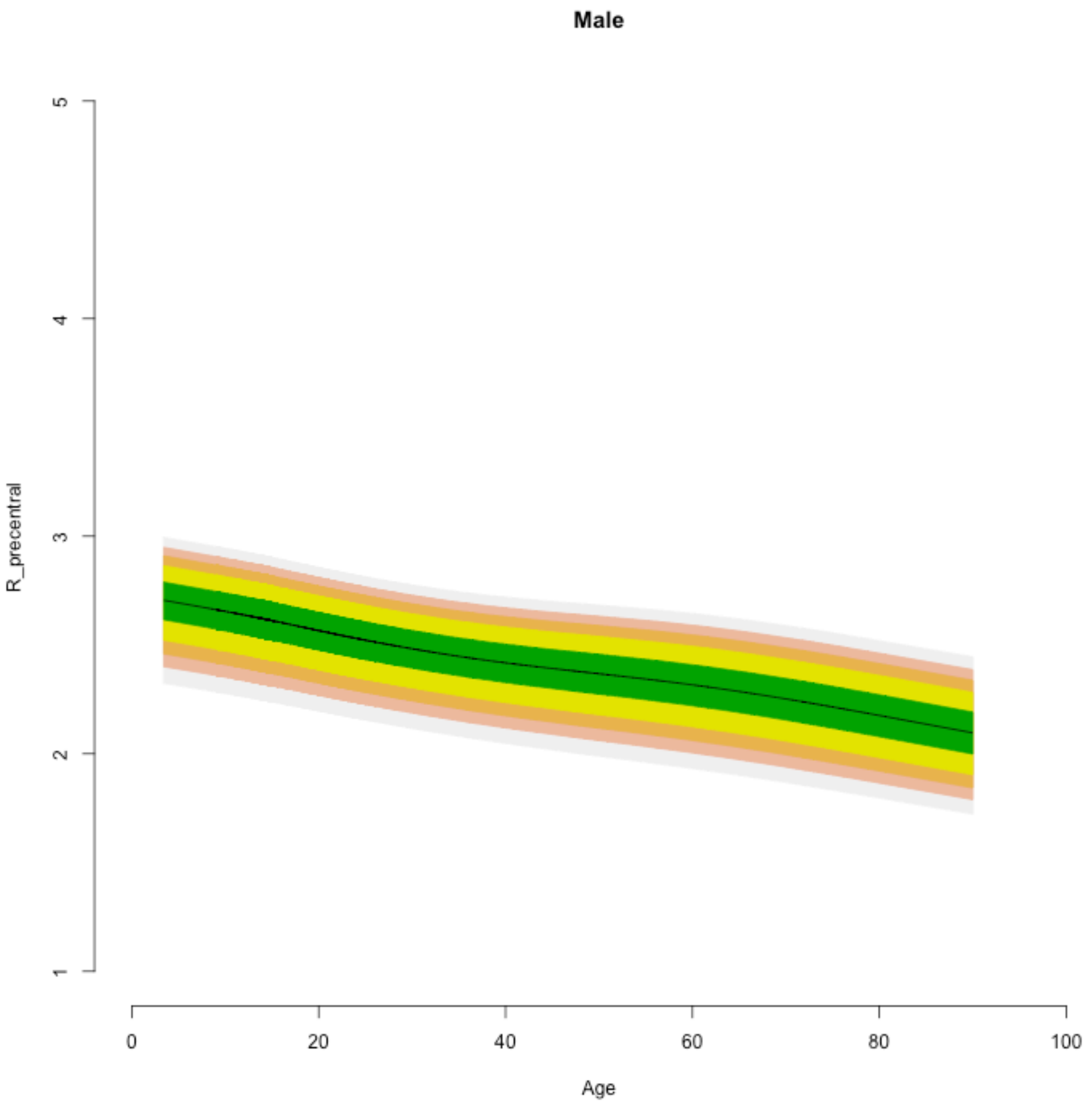
Male
R_precentral
5
4
3
2
1
0
20
40
60
80
100
Age

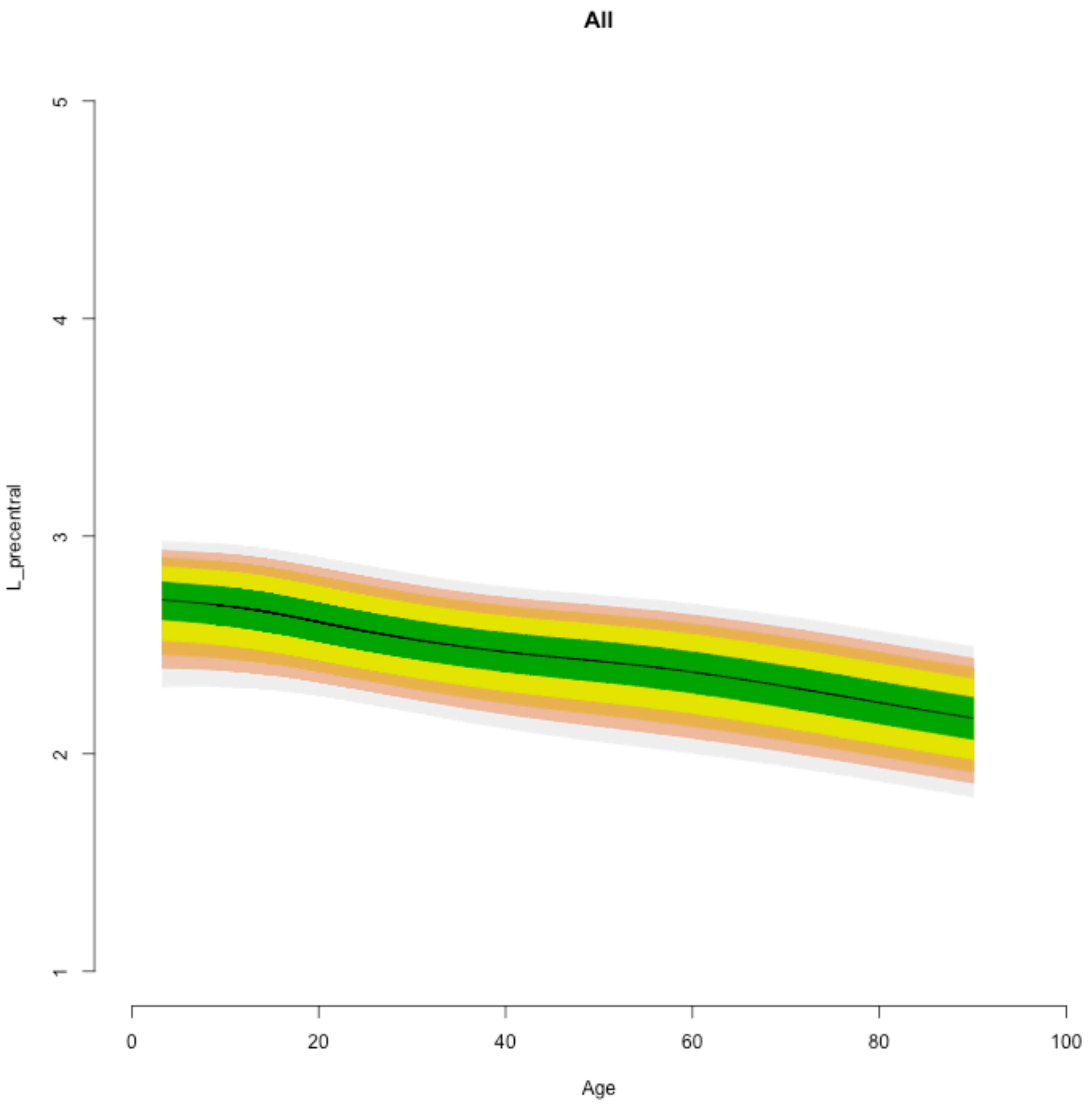

All
L_precentral
1
2
3
4
5
0
20
40
60
80
100
Age

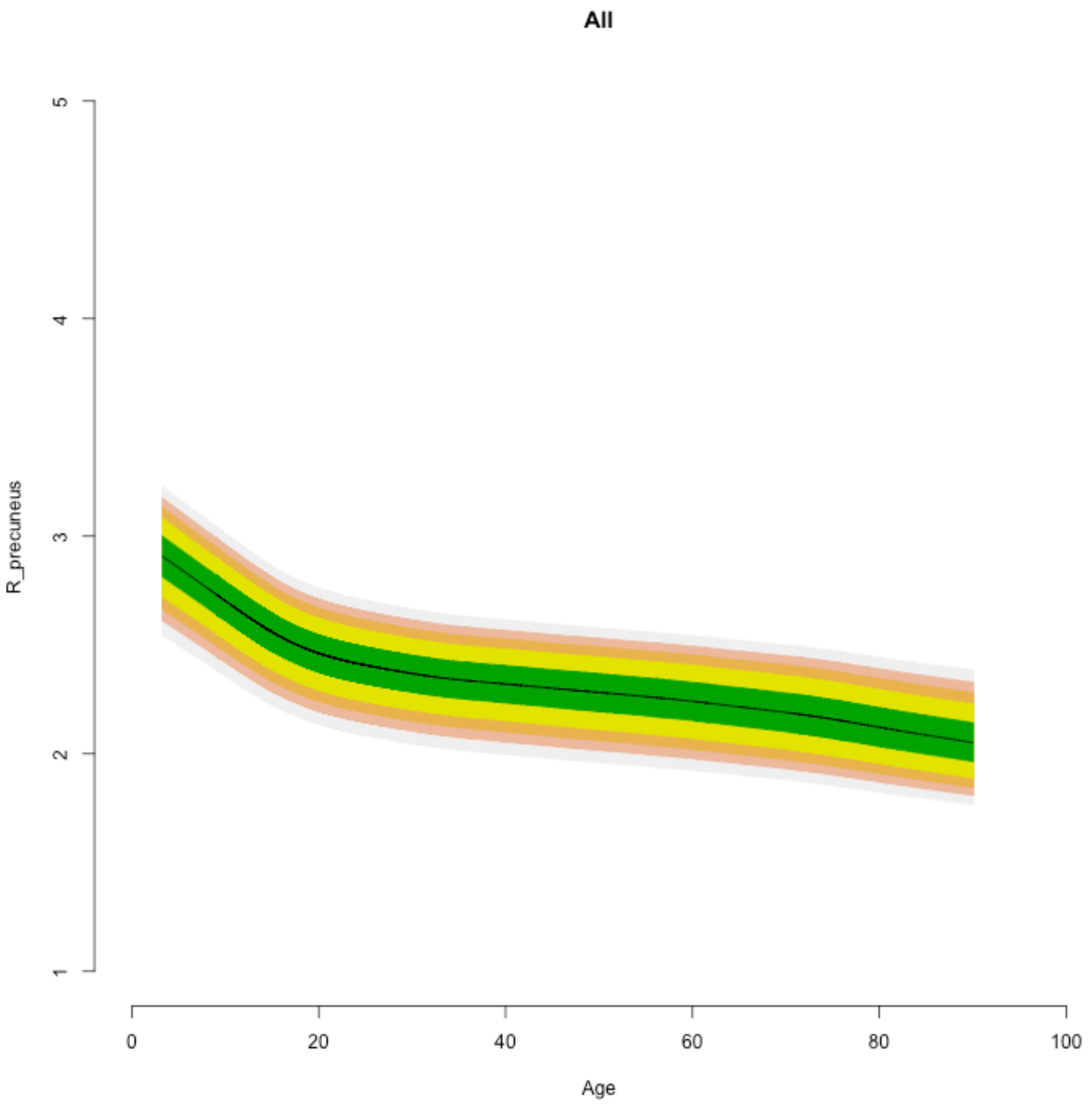
All
R_precuneus
5
4
3
2
1
0
20
40
60
80
100
Age

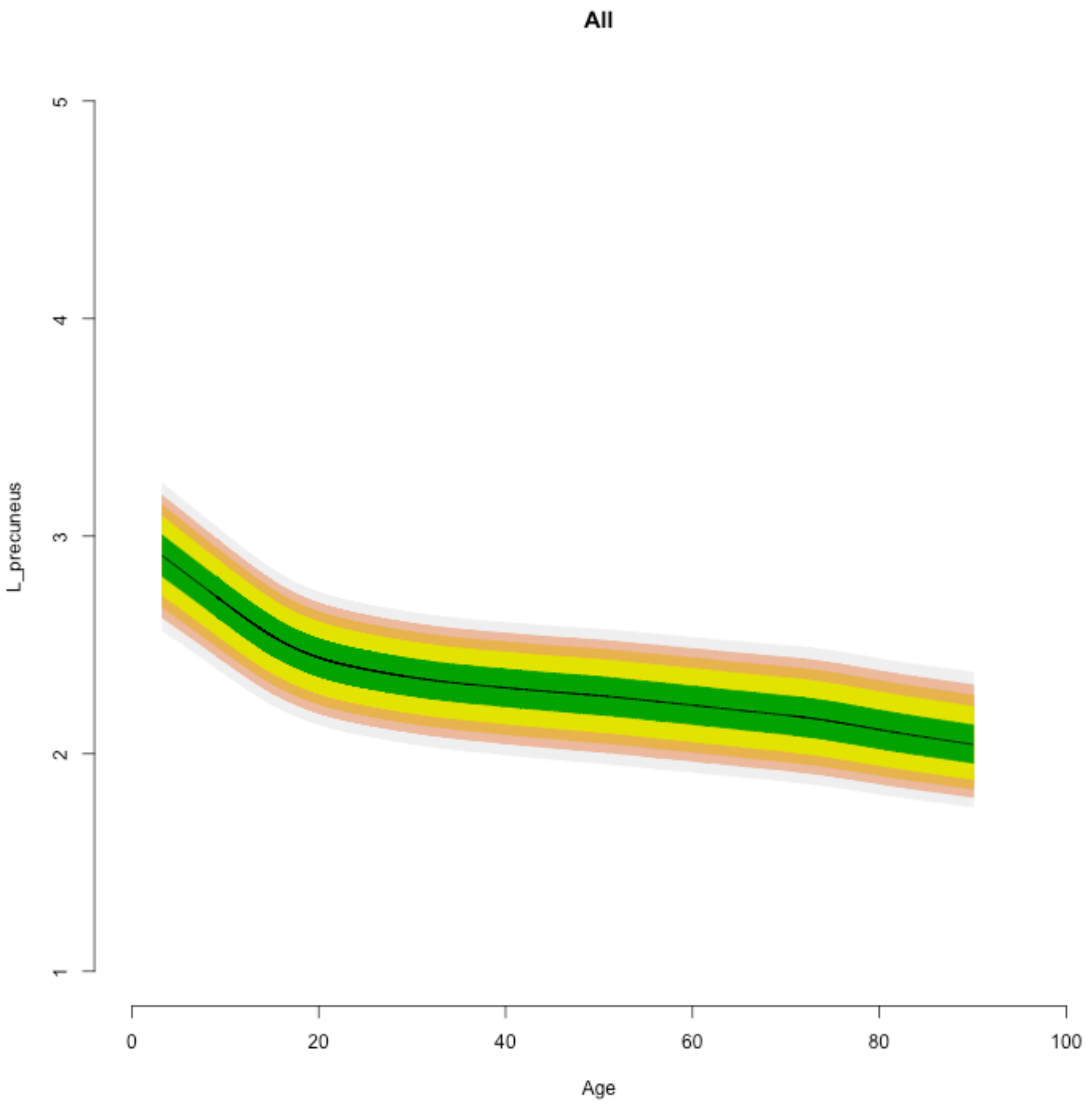
All
L_precuneus
1
2
3
4
5
0
20
40
60
80
100
Age

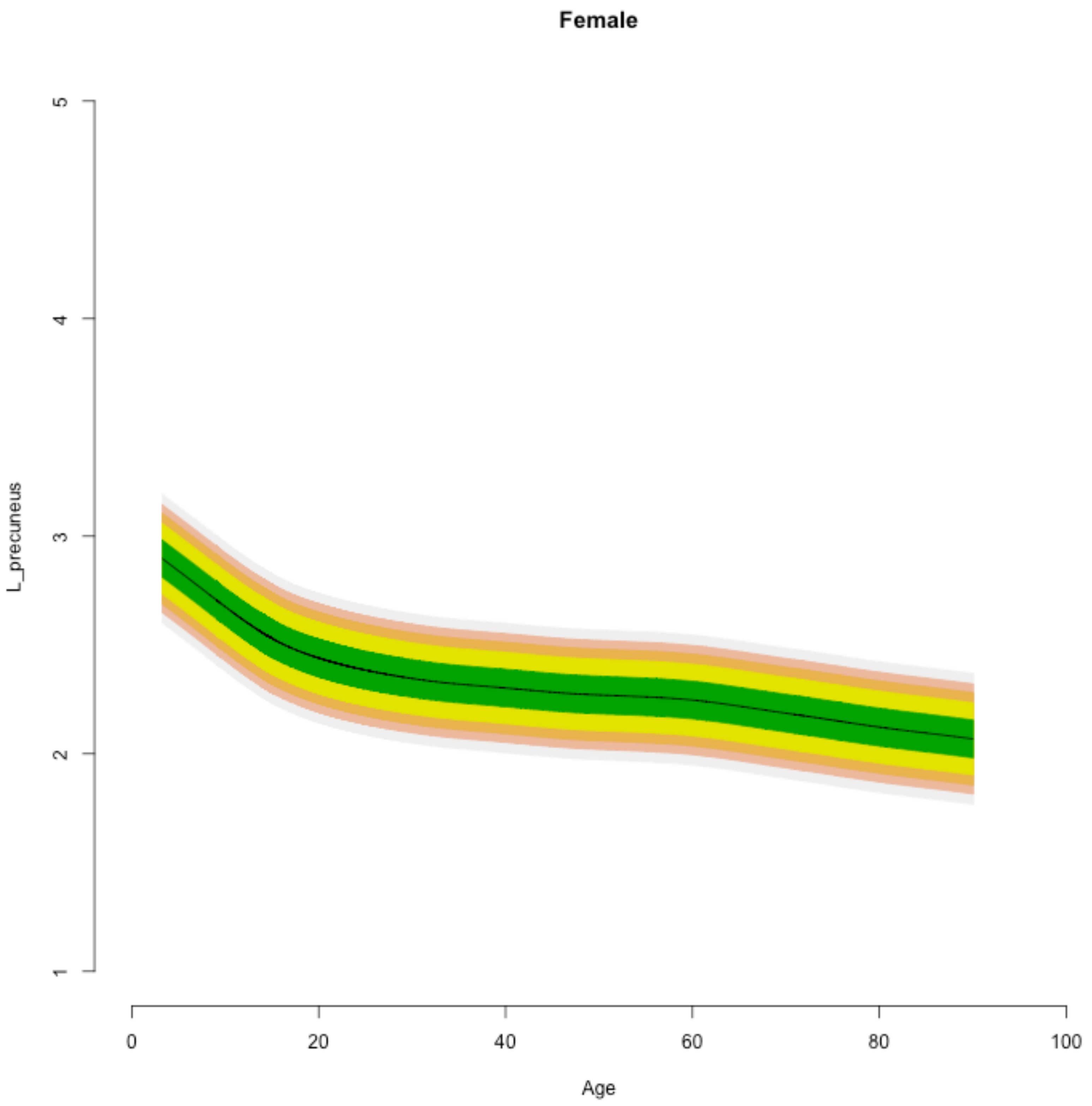
Female
L_precuneus
1
2
3
4
5
0
20
40
60
80
100
Age

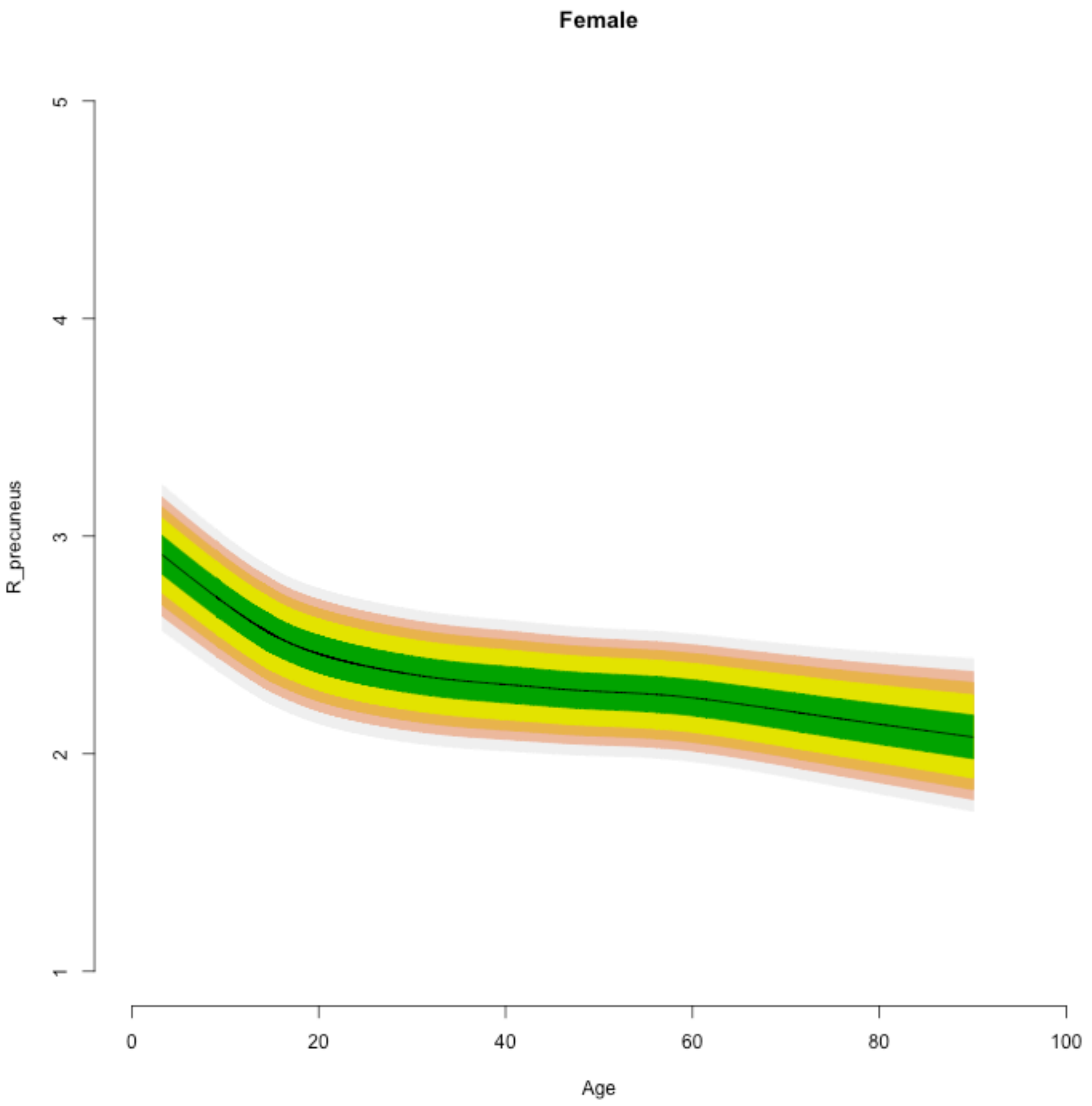
Female
R_precuneus
1
2
3
4
5
0
20
40
60
80
100
Age

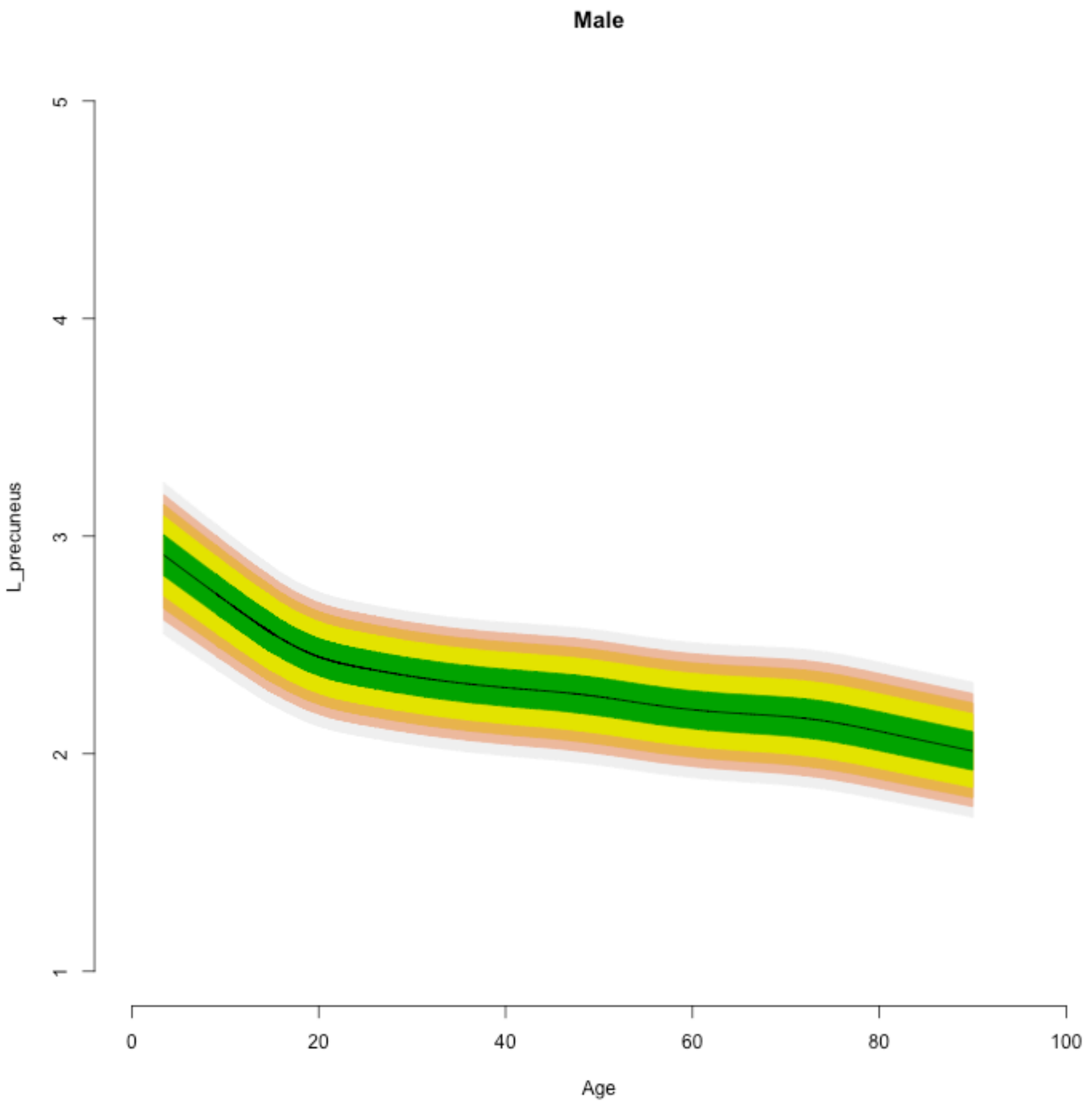
Male
L_precuneus
5
4
3
2
1
0
20
40
60
80
100
Age

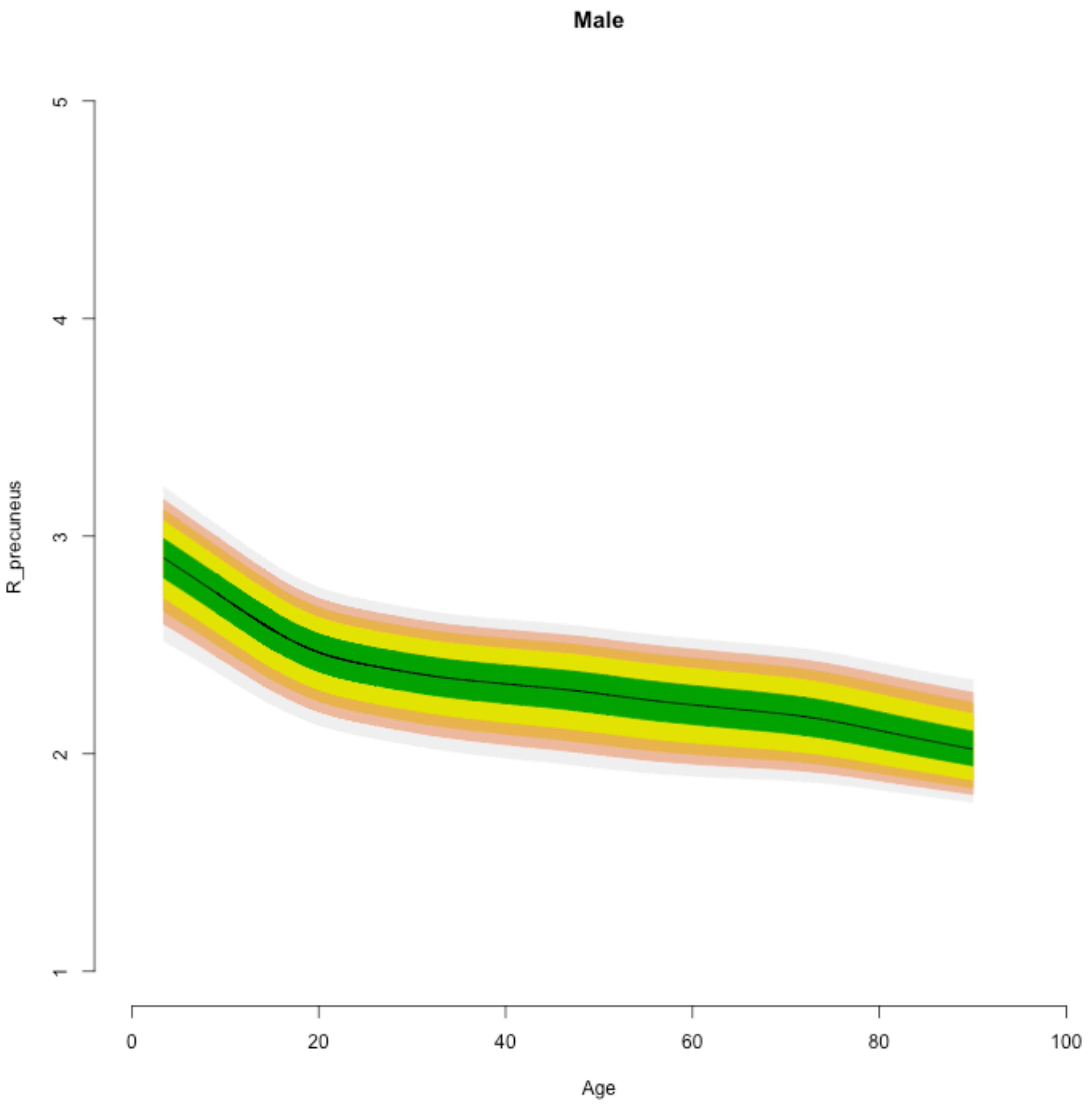

Male
R_precuneus
1
2
3
4
5
0
20
40
60
80
100
Age

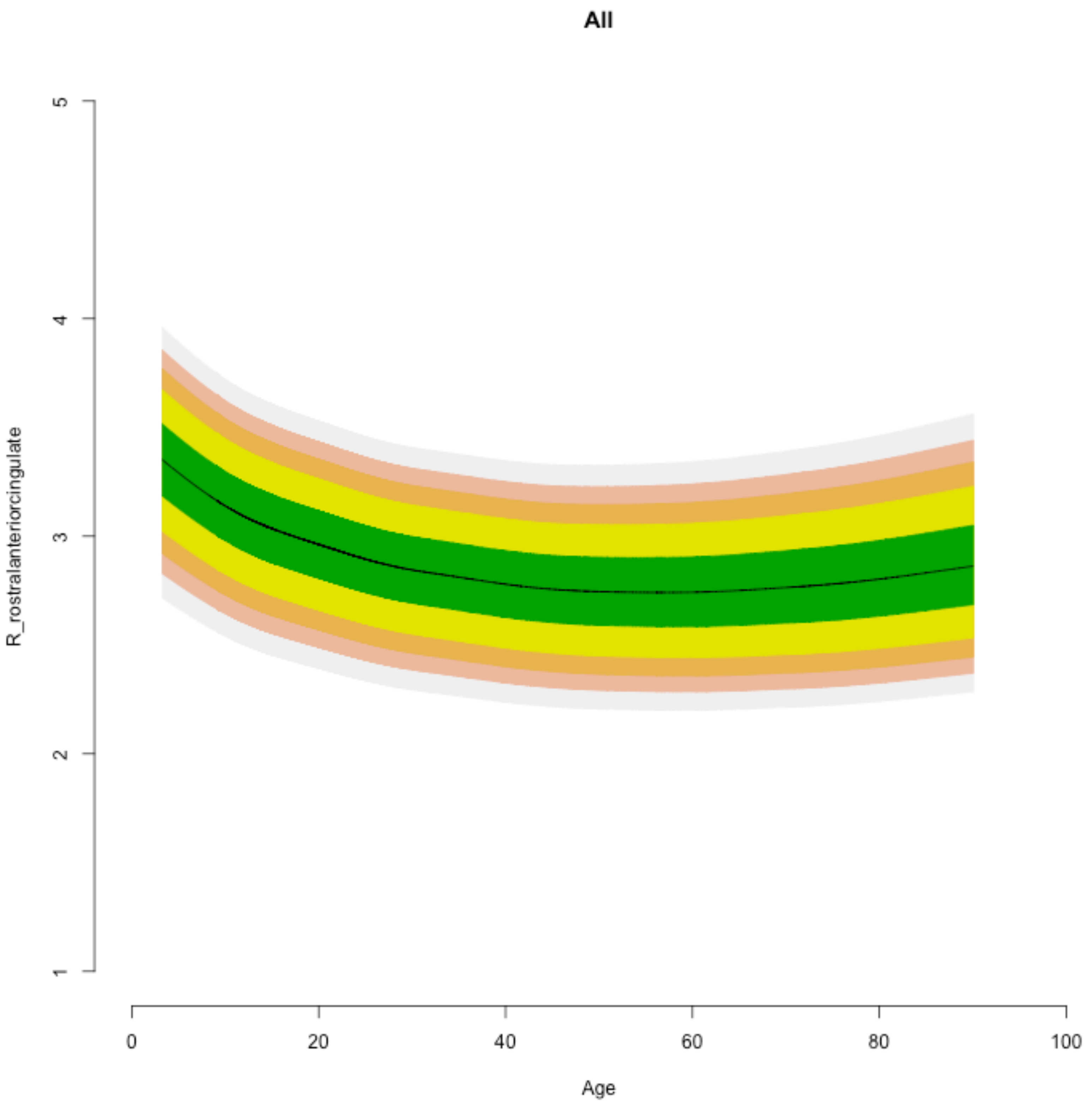

All
R_rostralanteriorcingulate
1
2
3
4
5
0
20
40
60
80
100
Age

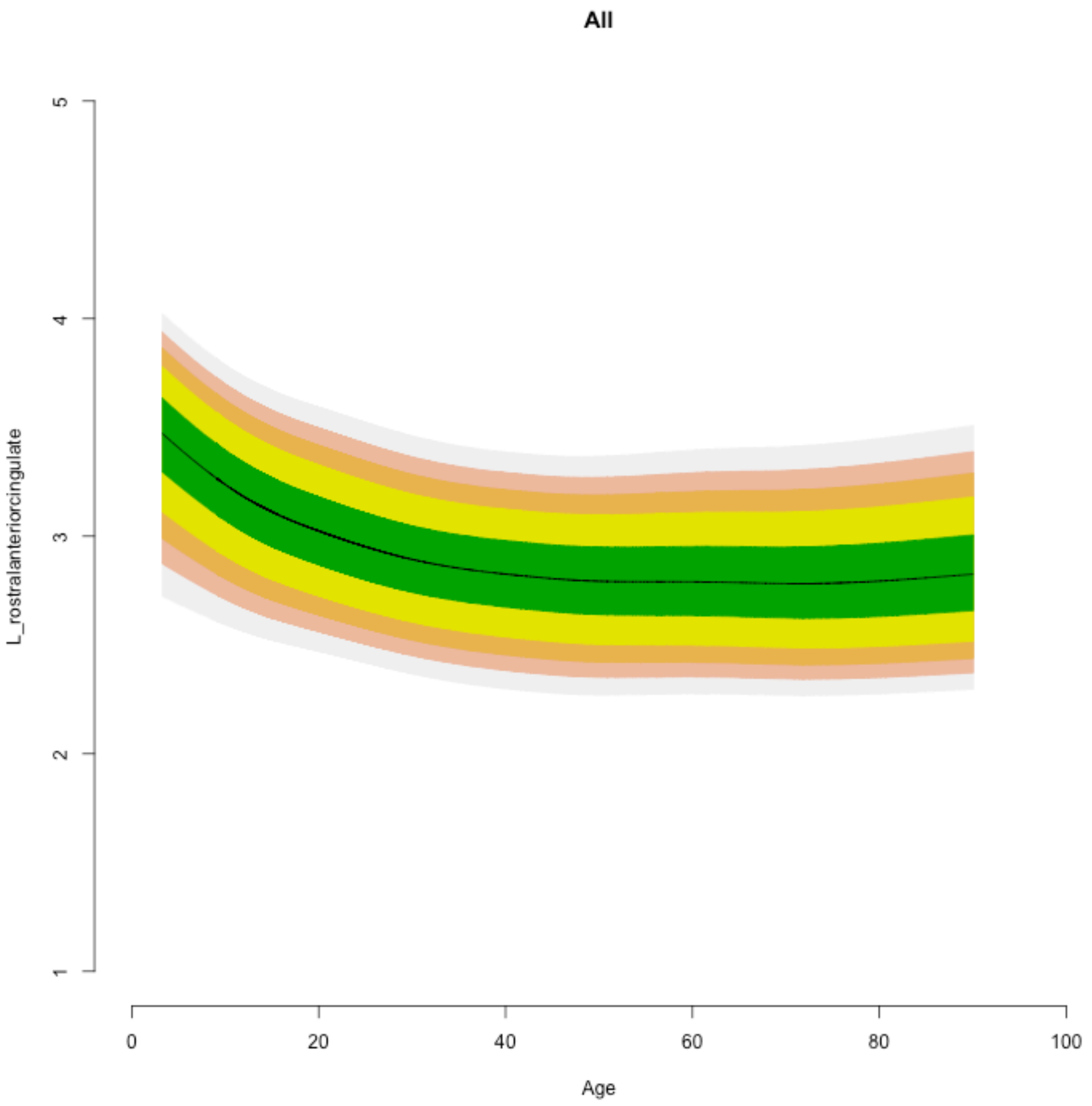
All
L_rostralanteriorcingulate
1
2
3
4
5
0
20
40
60
80
100
Age

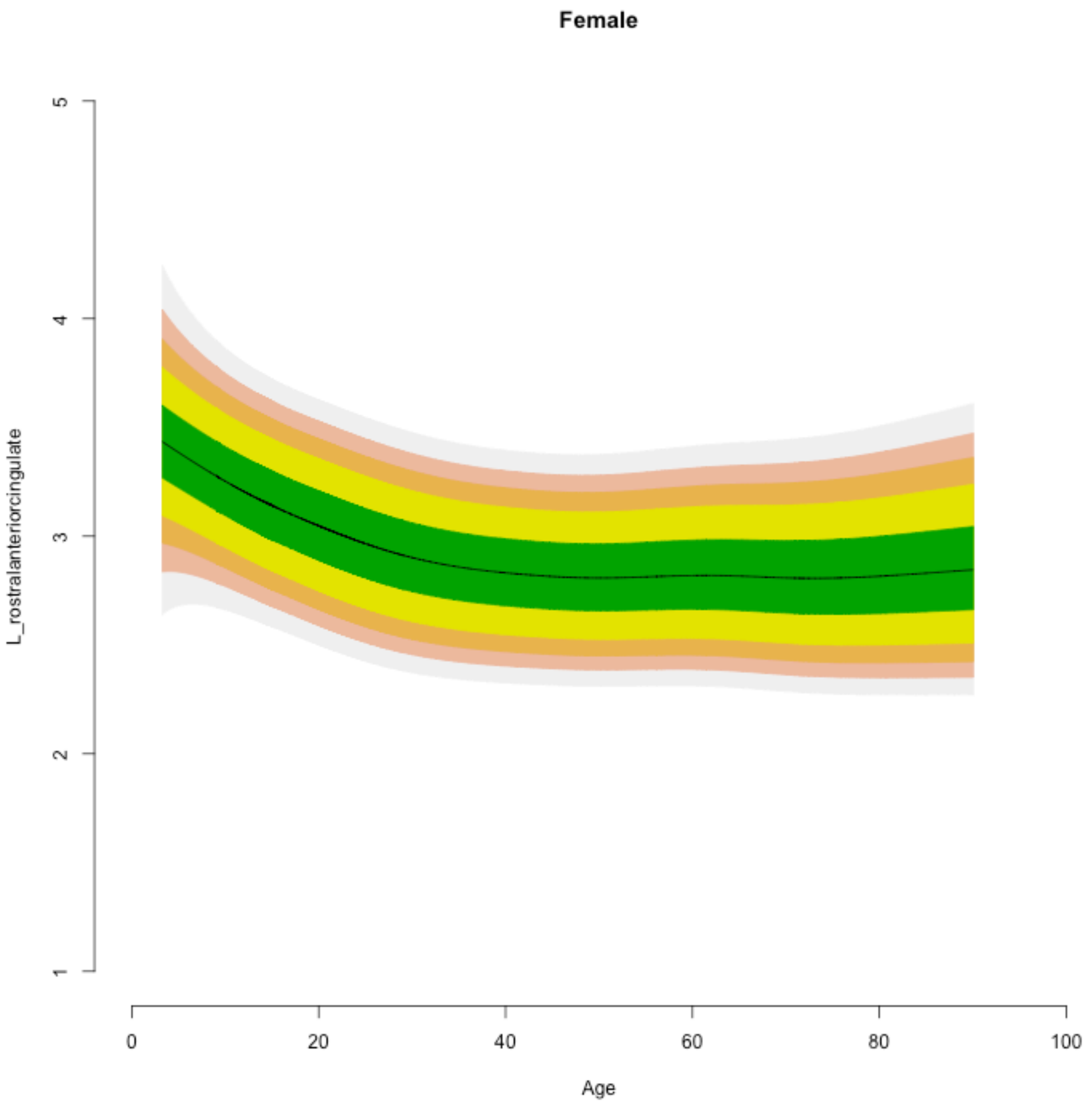

Female
L_rostralanteriorcingulate
Age
1
2
3
4
5
0
20
40
60
80
100

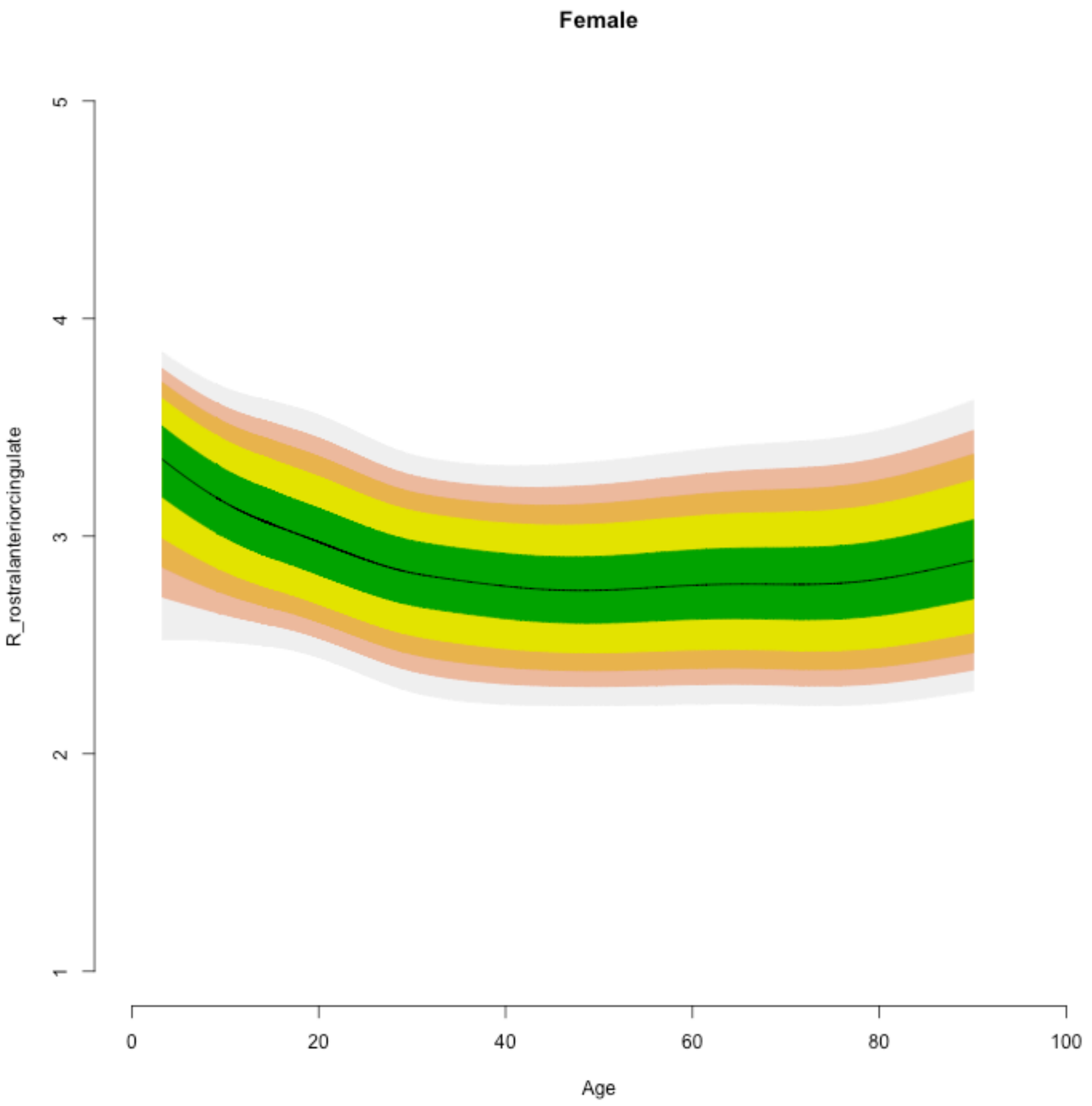
Female
R_rostralanteriorcingulate
1
2
3
4
5
0
20
40
60
80
100
Age

**Male**

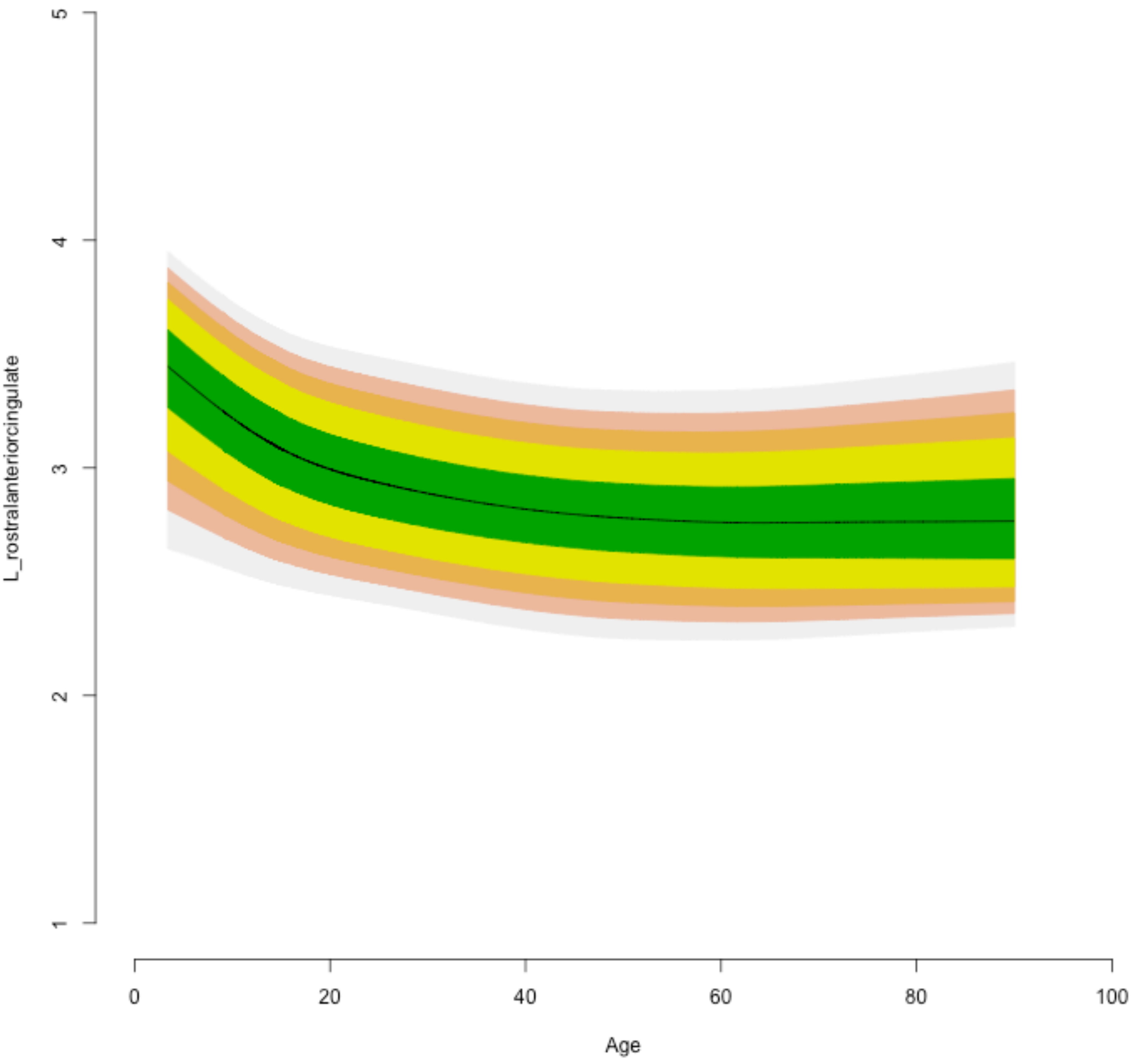

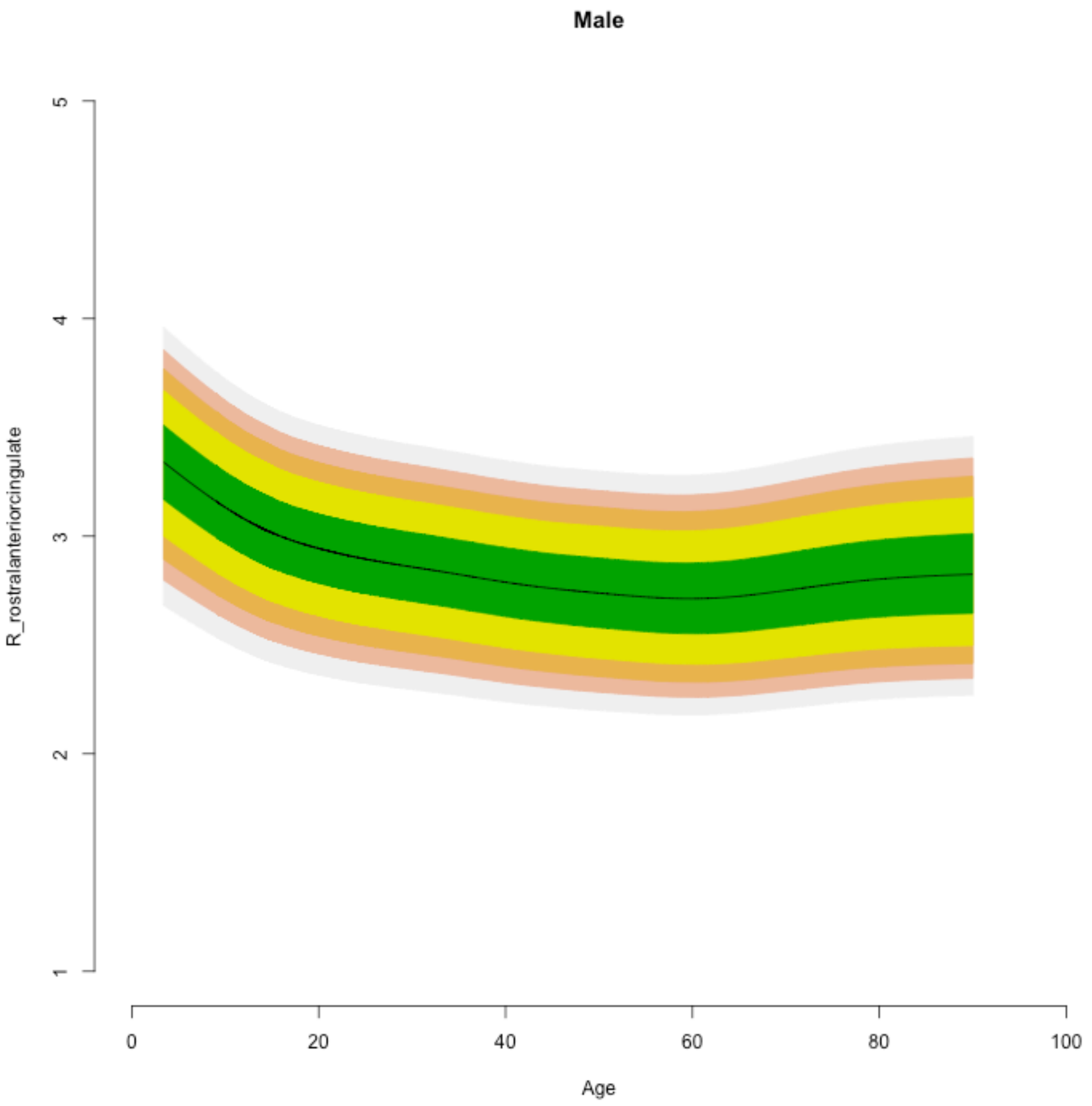

Male
5
4
3
2
1
R_rostralanteriorcingulate
0
20
40
60
80
100
Age

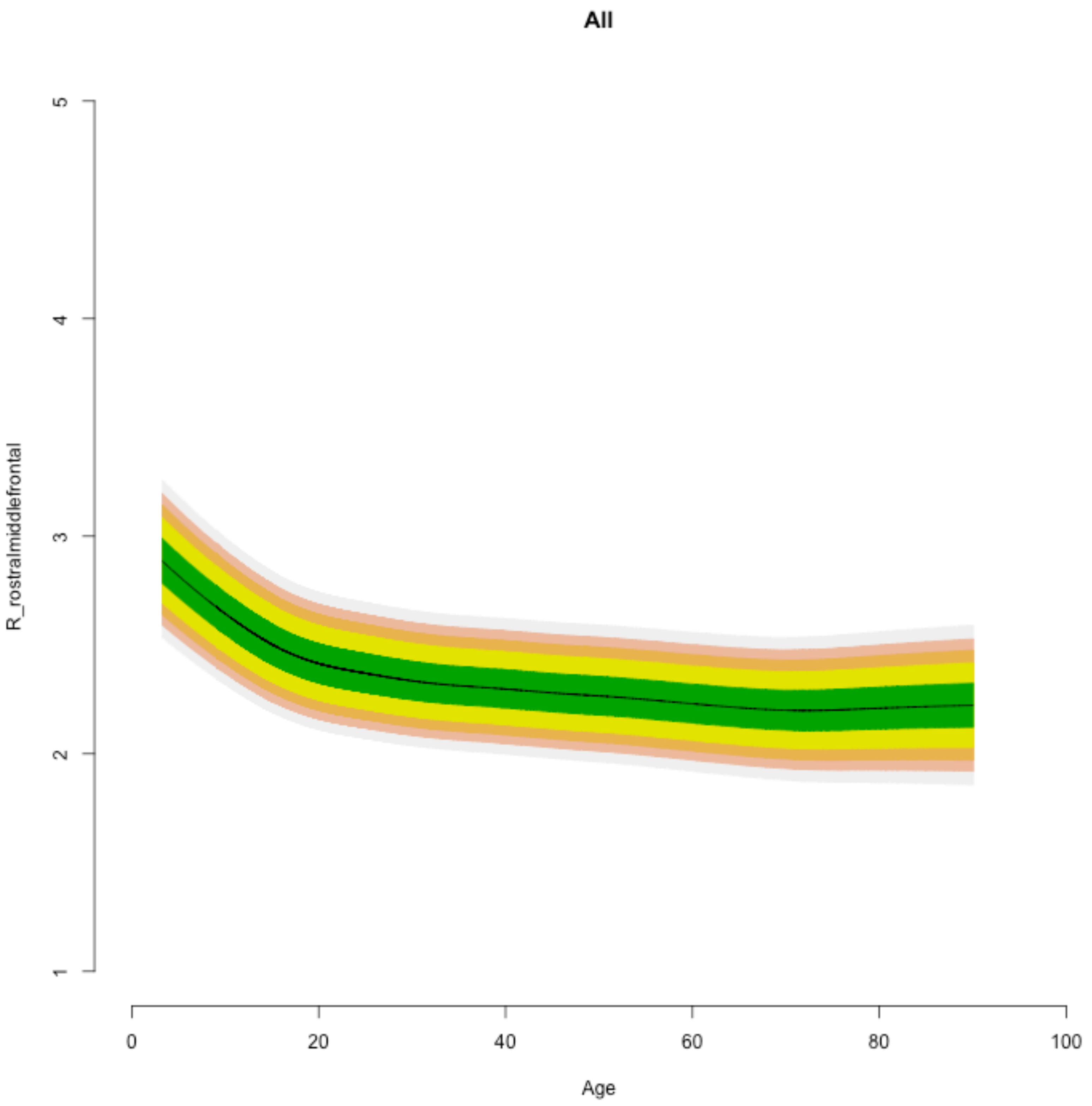
All
R_rostralmiddlefrontal
1
2
3
4
5
0
20
40
60
80
100
Age

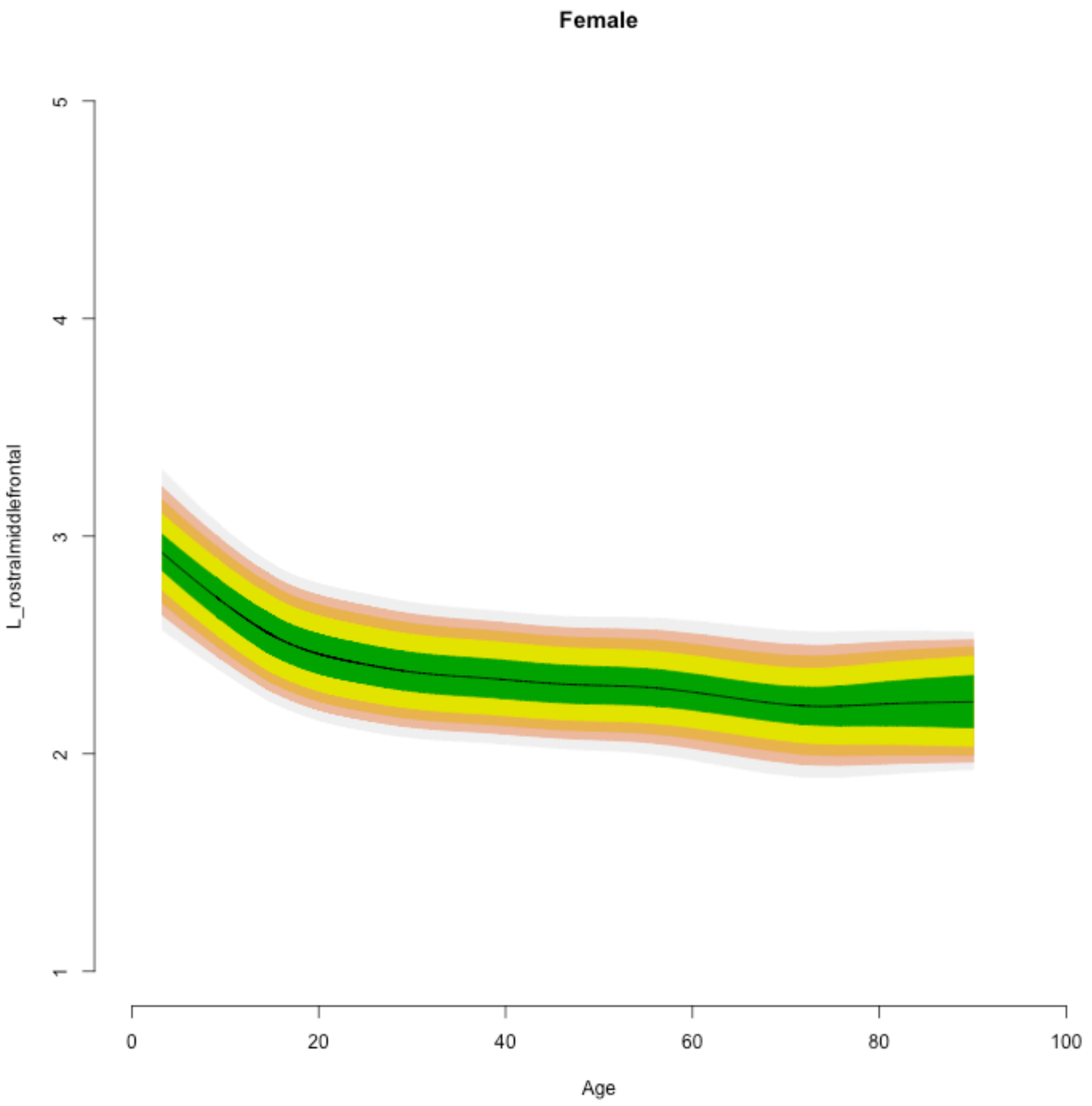
Female
L_rostralmiddlefrontal
5
4
3
2
1
0
20
40
60
80
100
Age

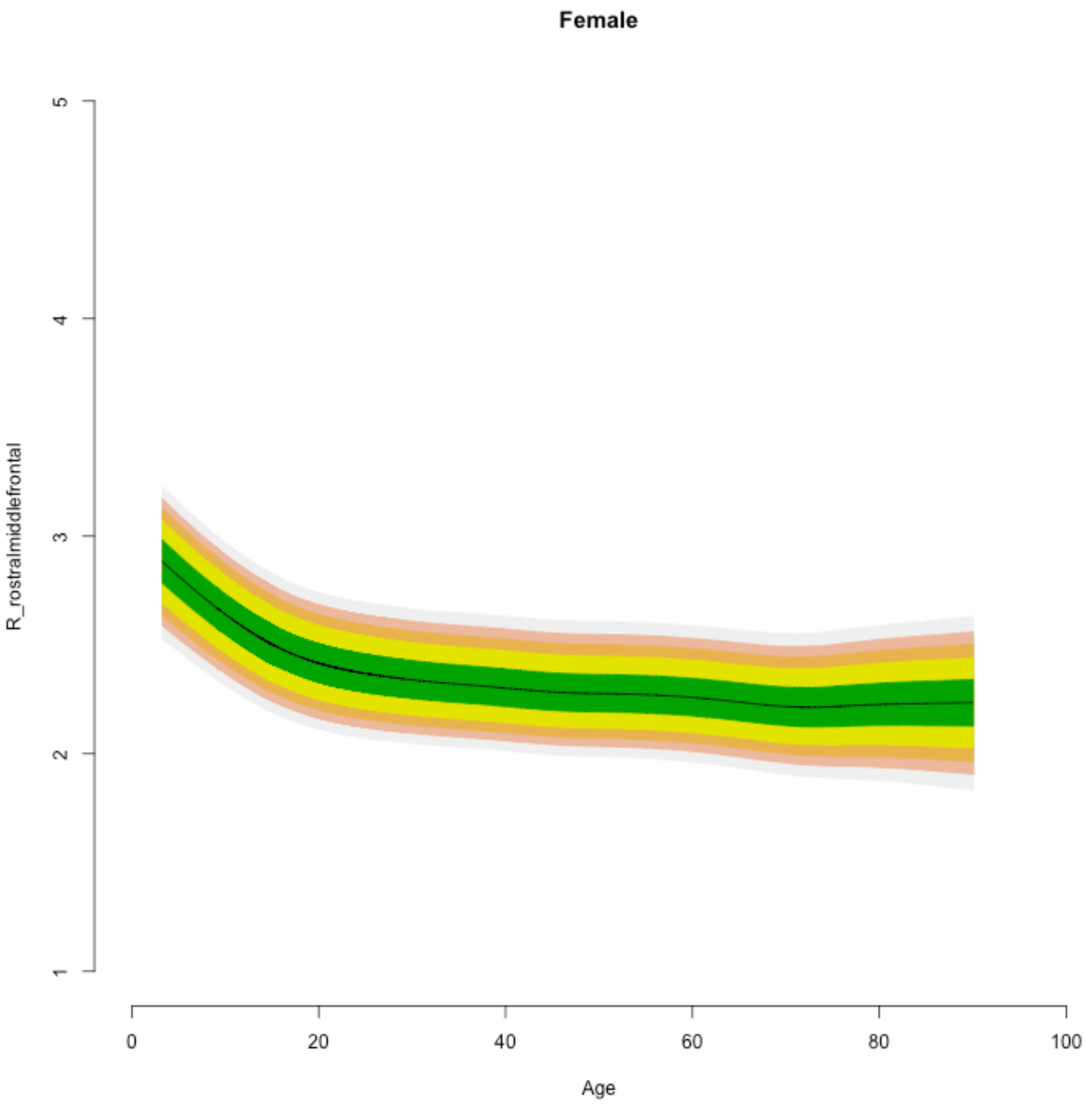
Female
R_rostralmiddlefrontal
Age
1
2
3
4
5
0
20
40
60
80
100

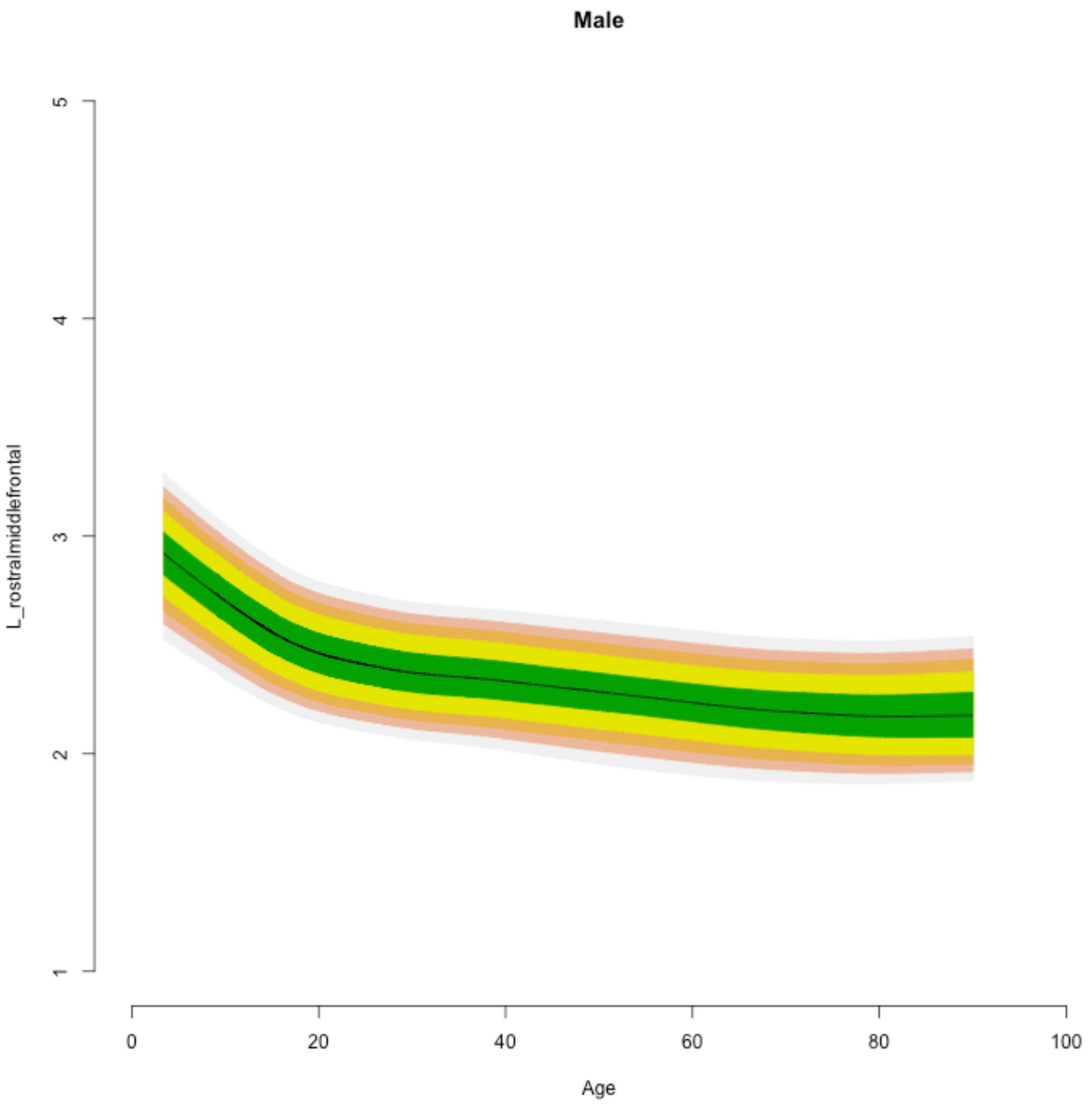
Male
L_rostralmiddlefrontal
1
2
3
4
5
0
20
40
60
80
100
Age

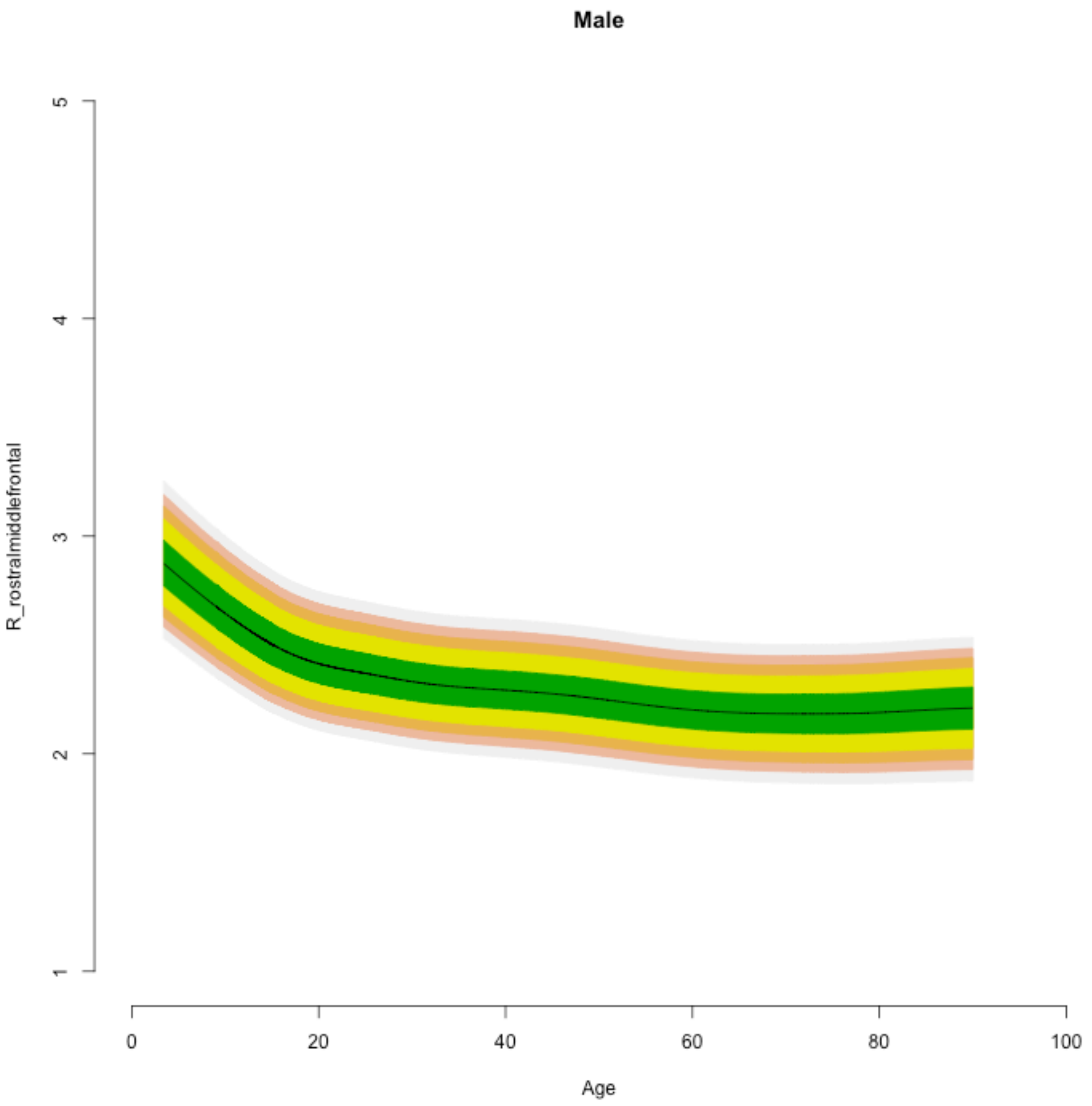
Male
R_rostralmiddlefrontal
5
4
3
2
1
0
20
40
60
80
100
Age

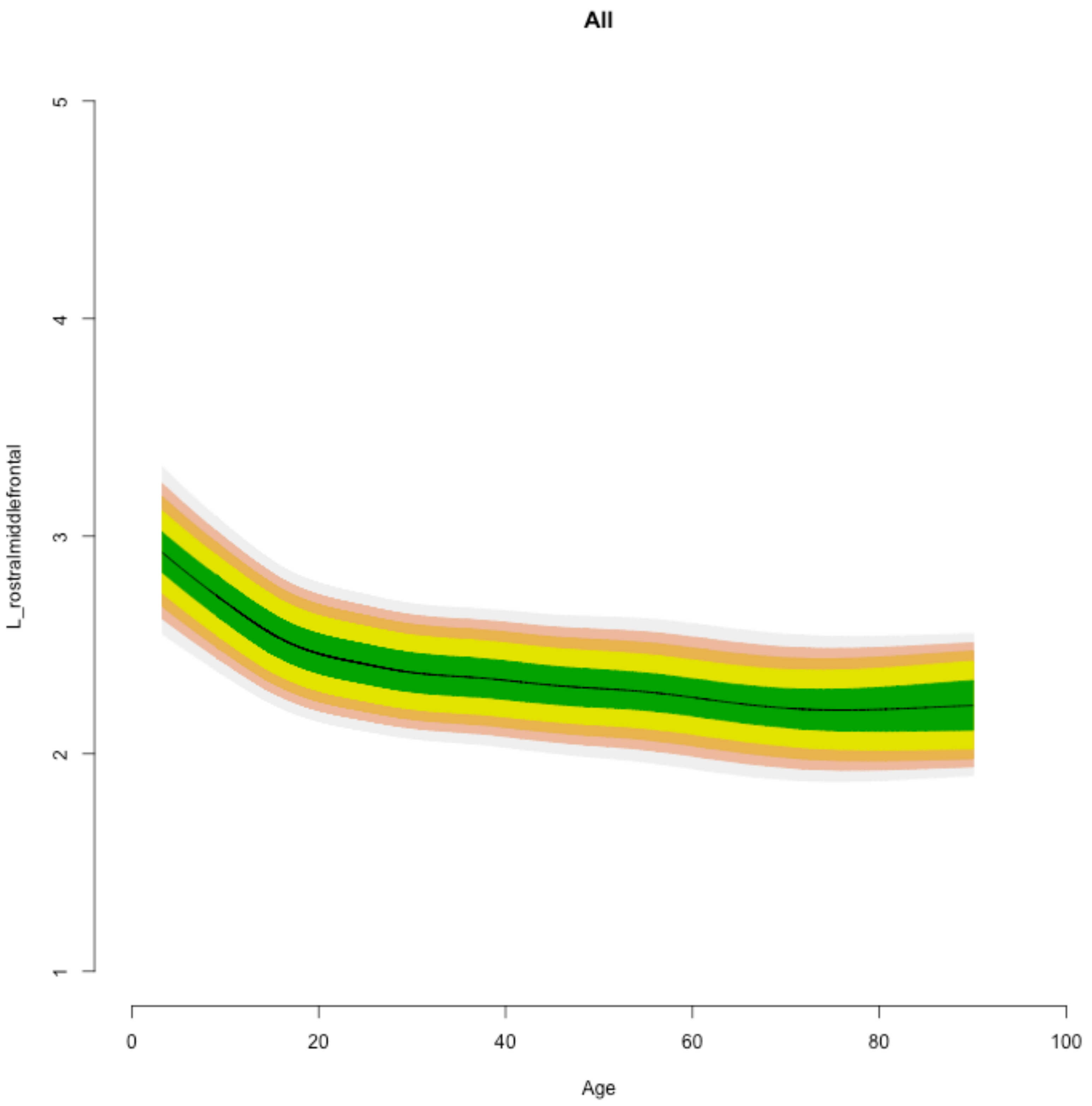
All
L_rostralmiddlefrontal
1
2
3
4
5
0
20
40
60
80
100
Age

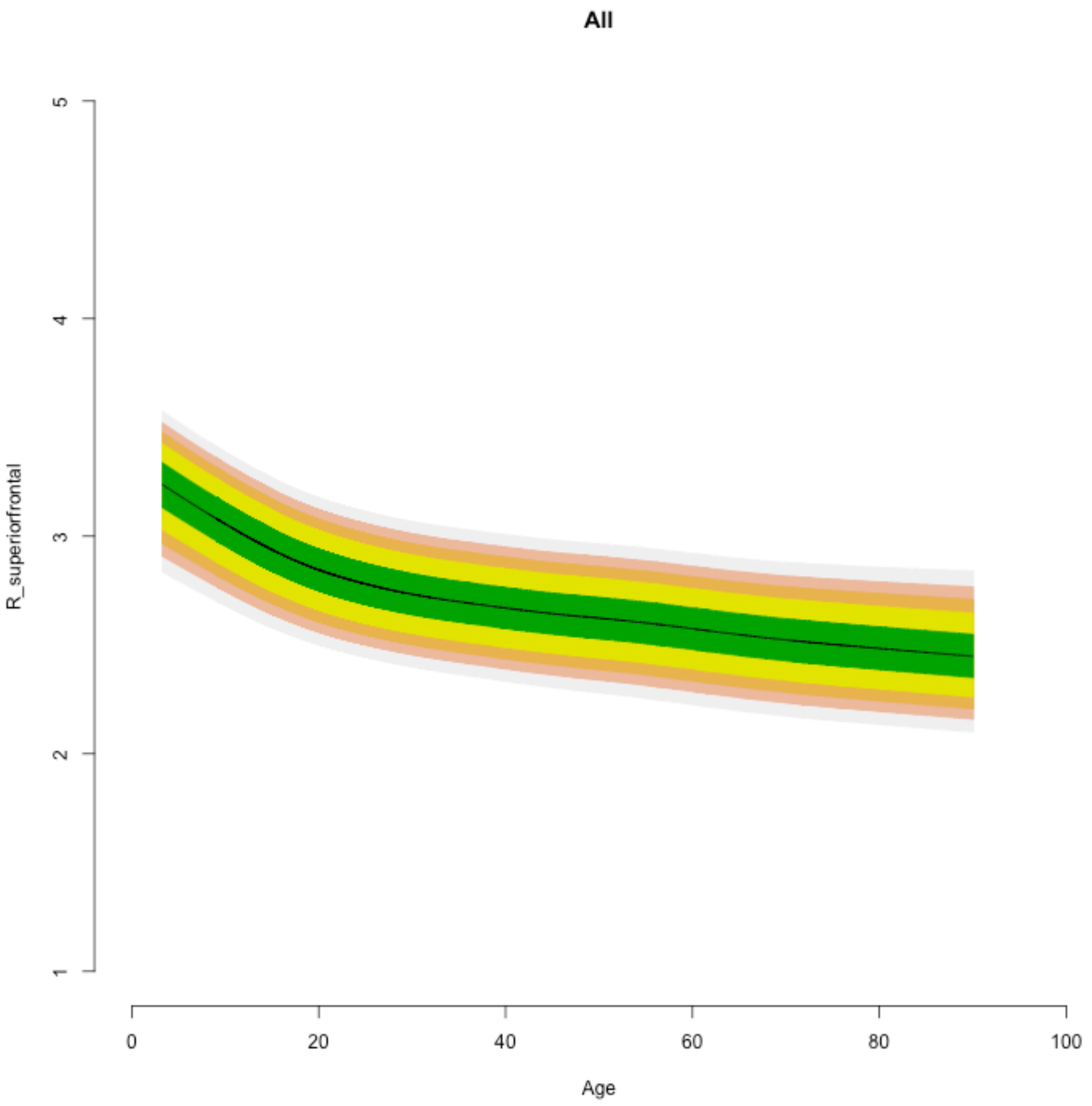
All
R_superiorfrontal
5
4
3
2
1
0
20
40
60
80
100
Age

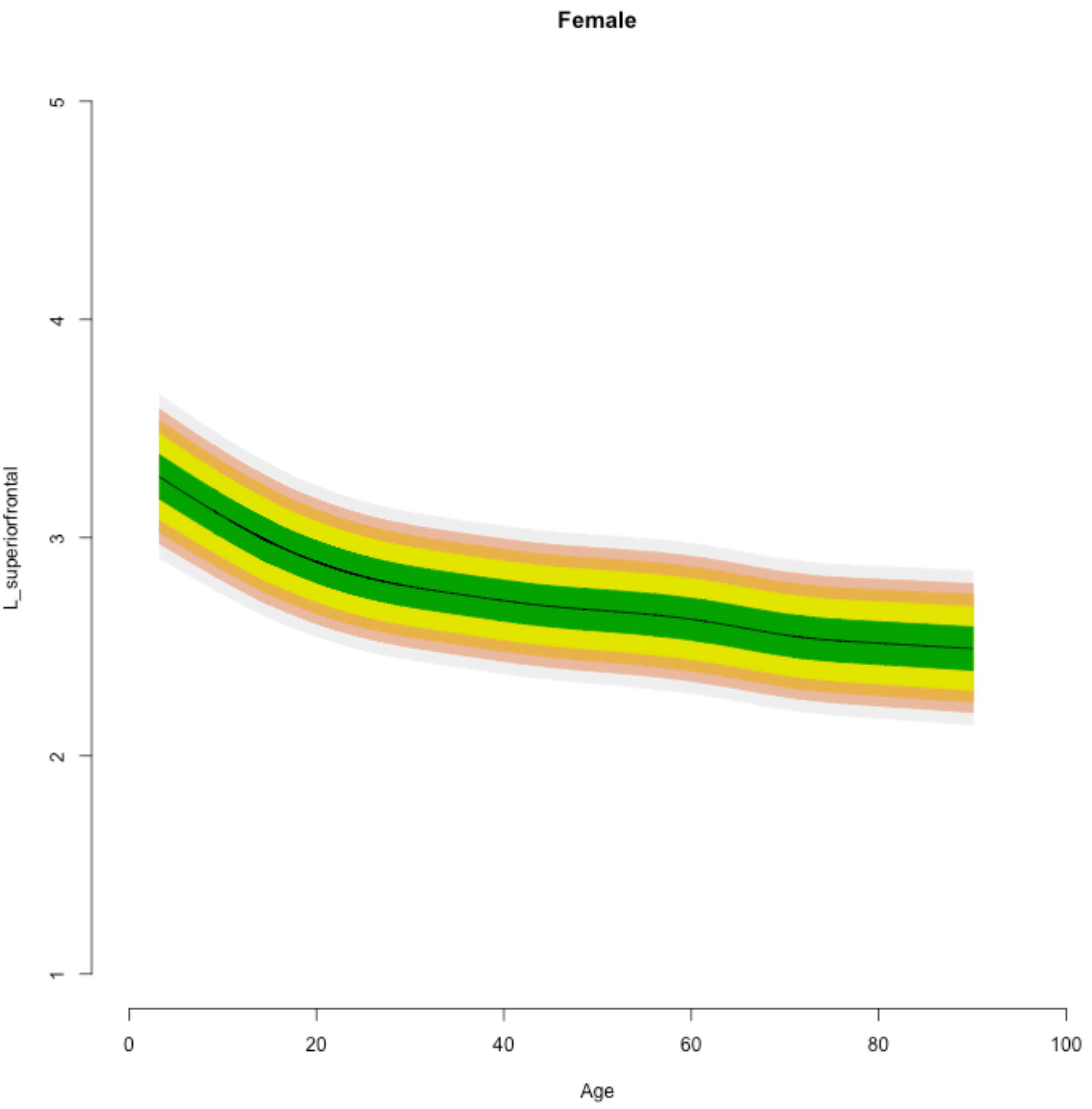

Female
L_superiorfrontal
1
2
3
4
5
0
20
40
60
80
100
Age

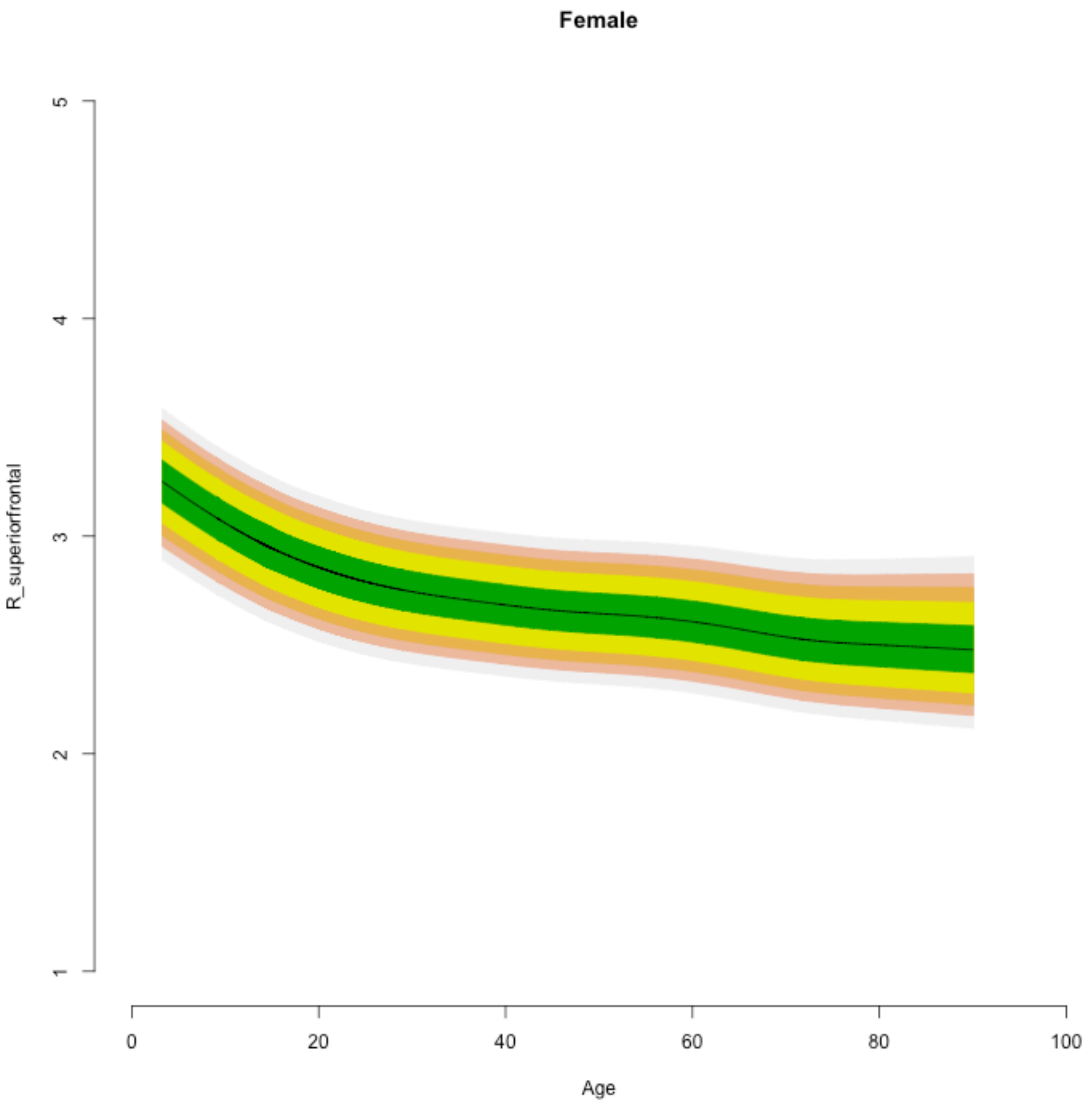
Female
R_superiorfrontal
1
2
3
4
5
0
20
40
60
80
100
Age

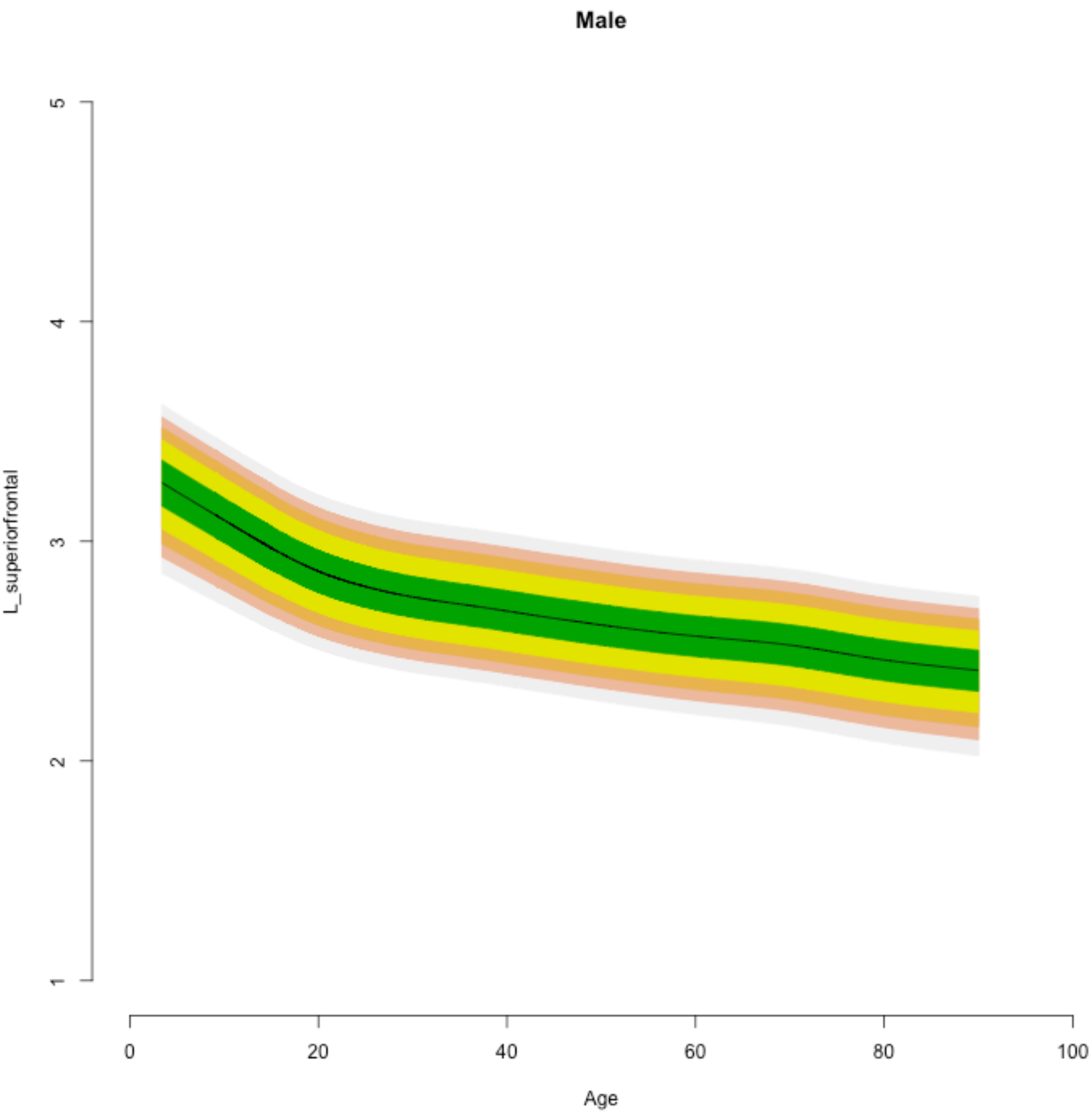

Male
L_superiorfrontal
1
2
3
4
5
0
20
40
60
80
100
Age

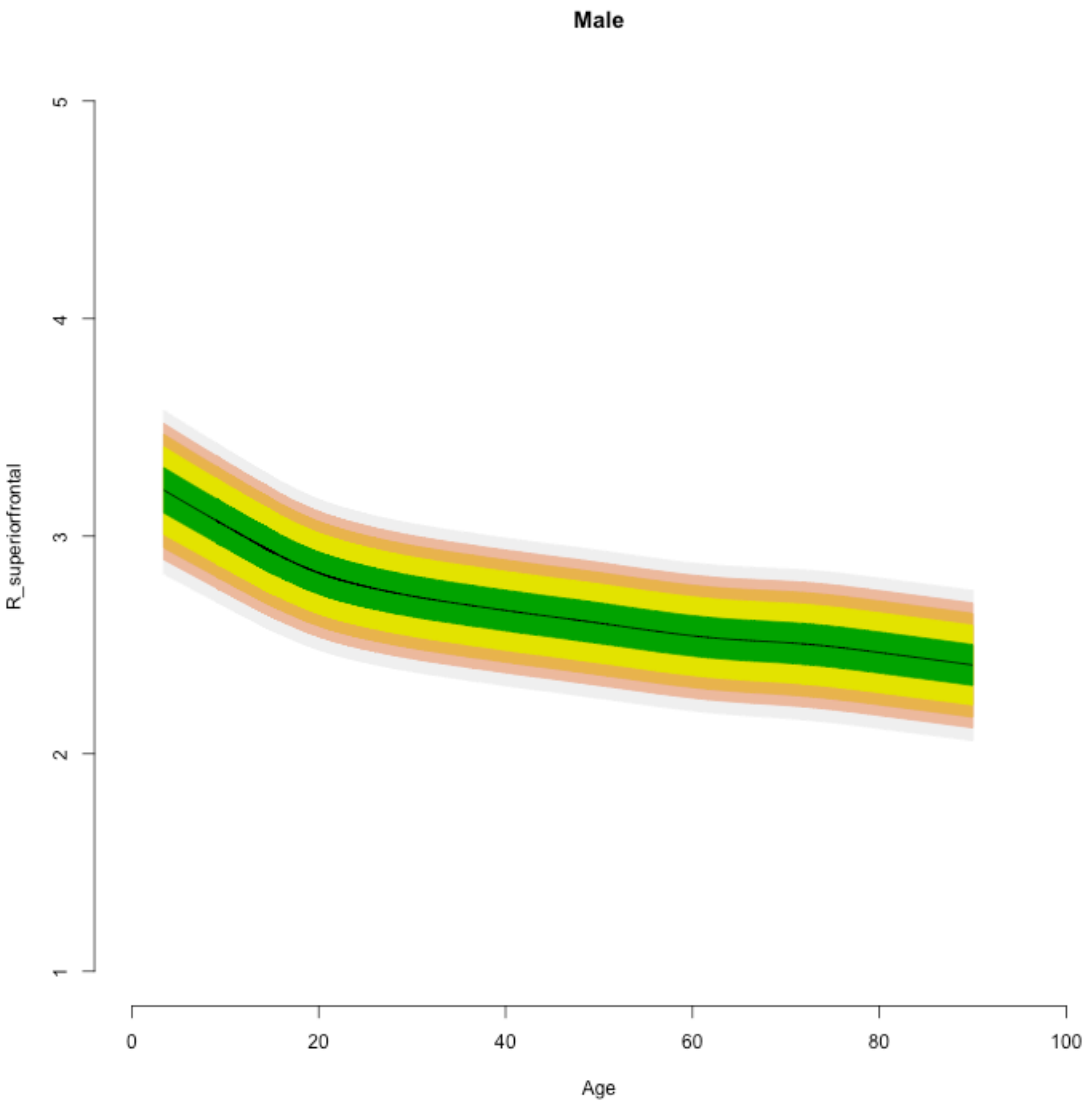

Male
R_superiorfrontal
5
4
3
2
1
0
20
40
60
80
100
Age

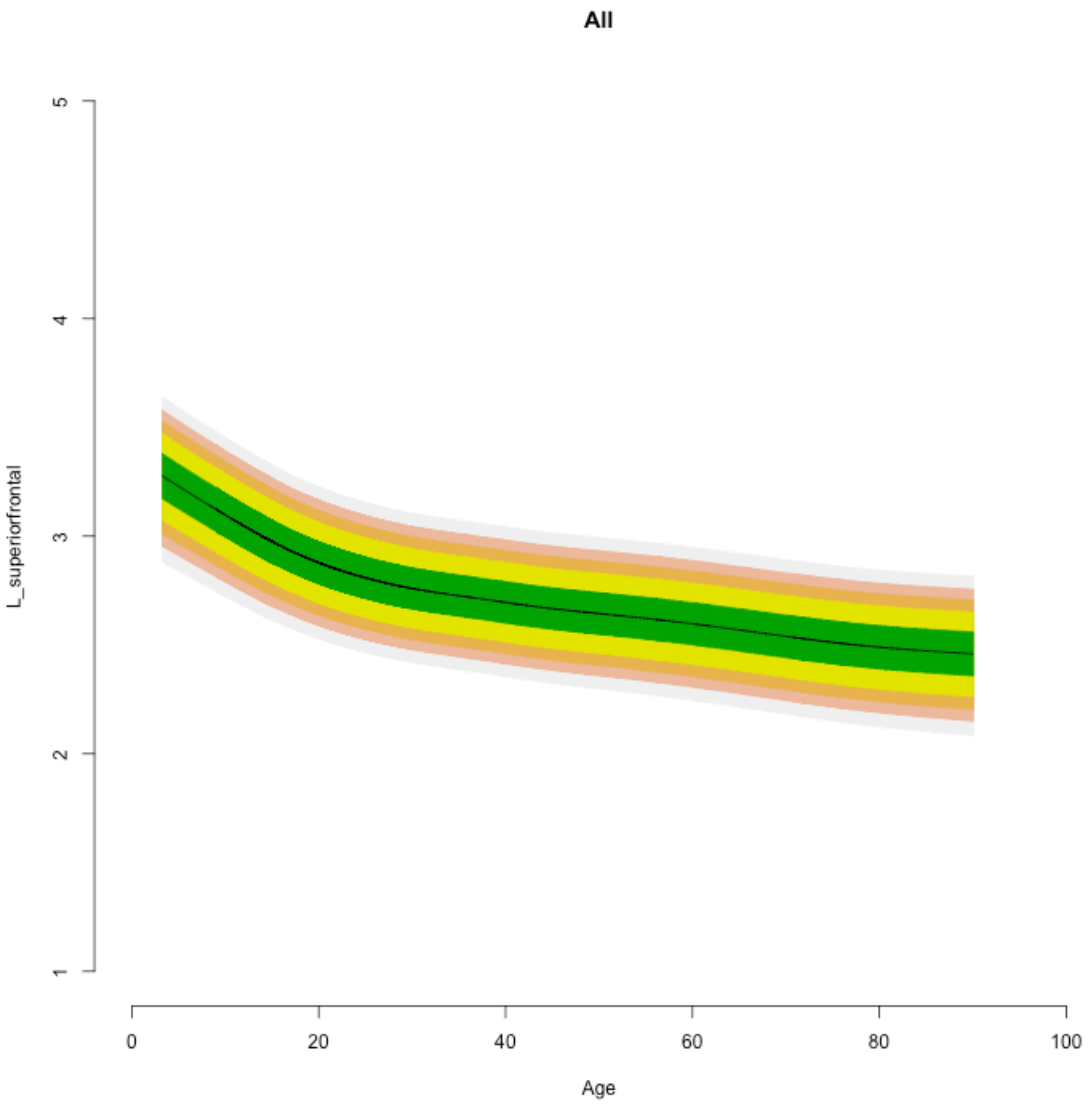

All
L_superiorfrontal
1
2
3
4
5
0
20
40
60
80
100
Age

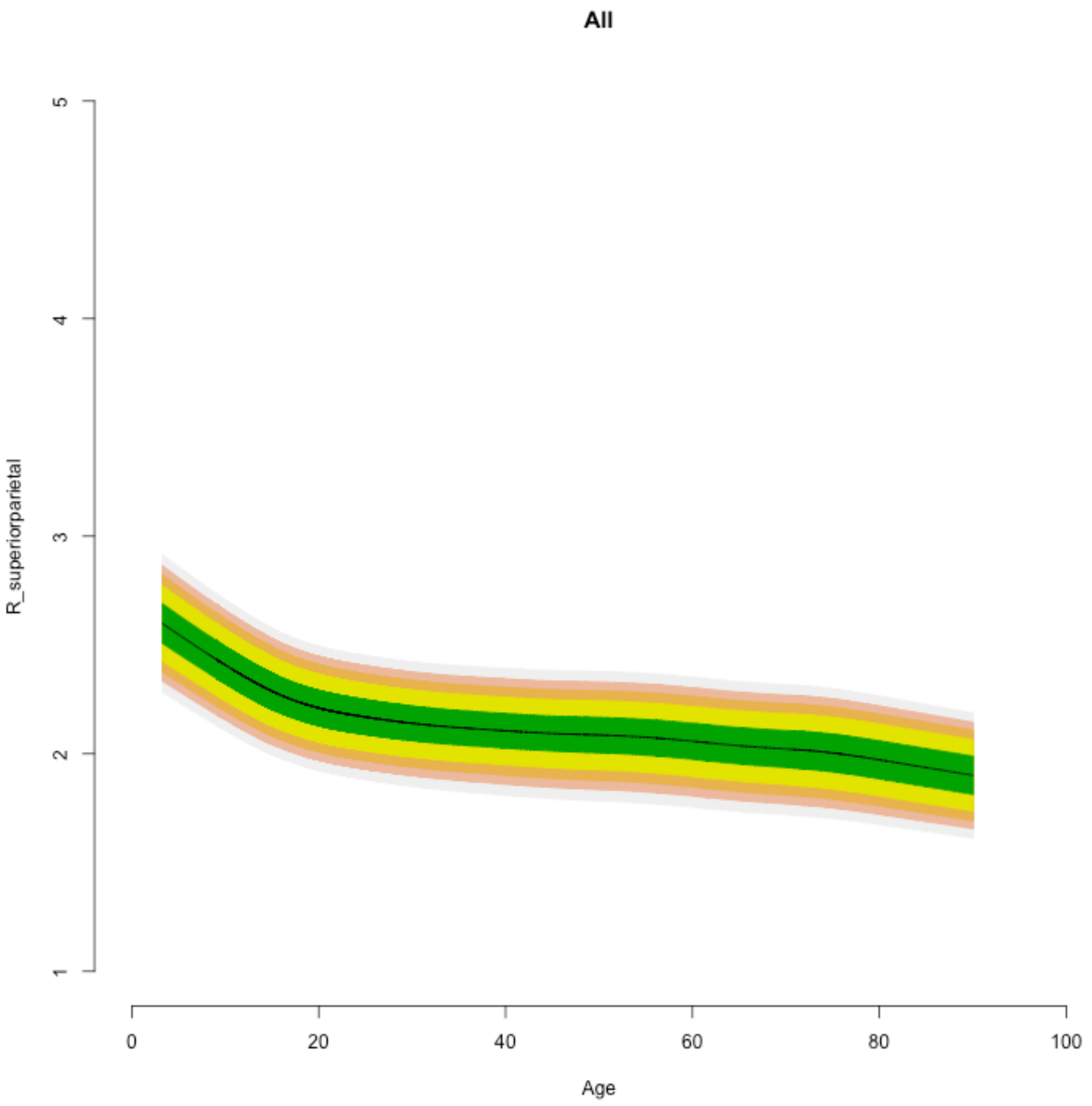
All
R_superiorparietal
1
2
3
4
5
0
20
40
60
80
100
Age

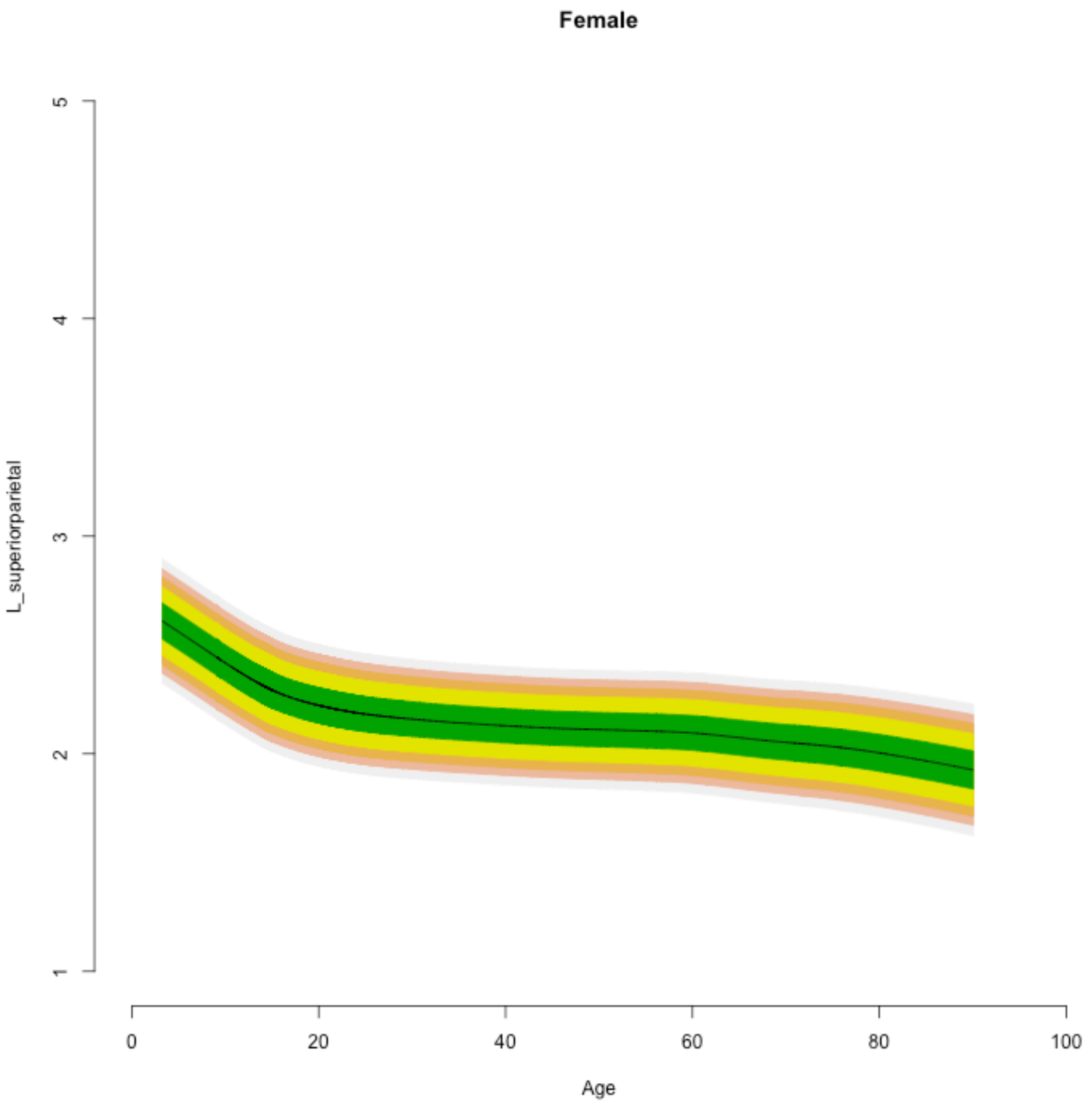

Female
L_superiorparietal
1
2
3
4
5
0
20
40
60
80
100
Age

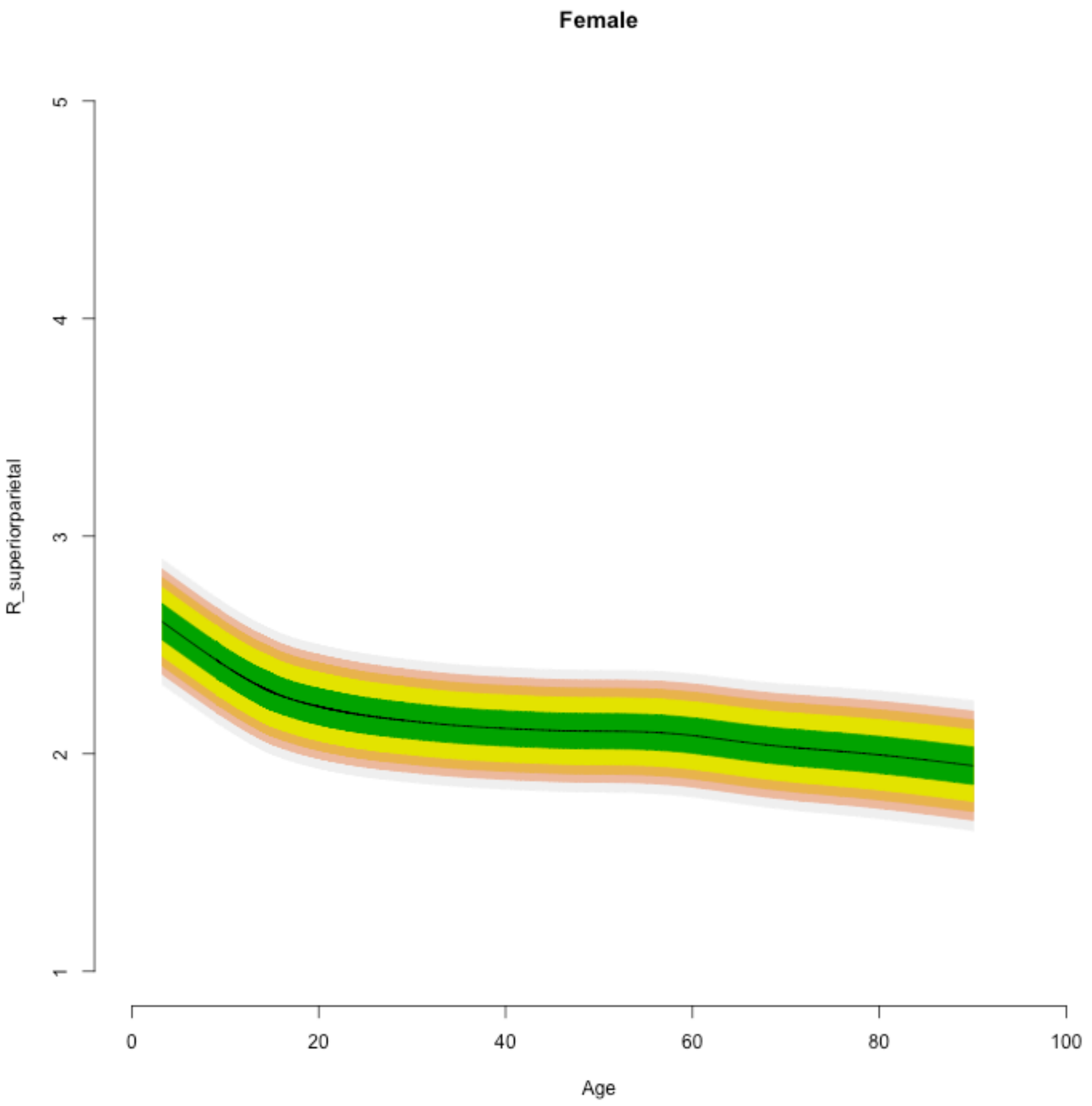

Female
R_superiorparietal
1
2
3
4
5
0
20
40
60
80
100
Age

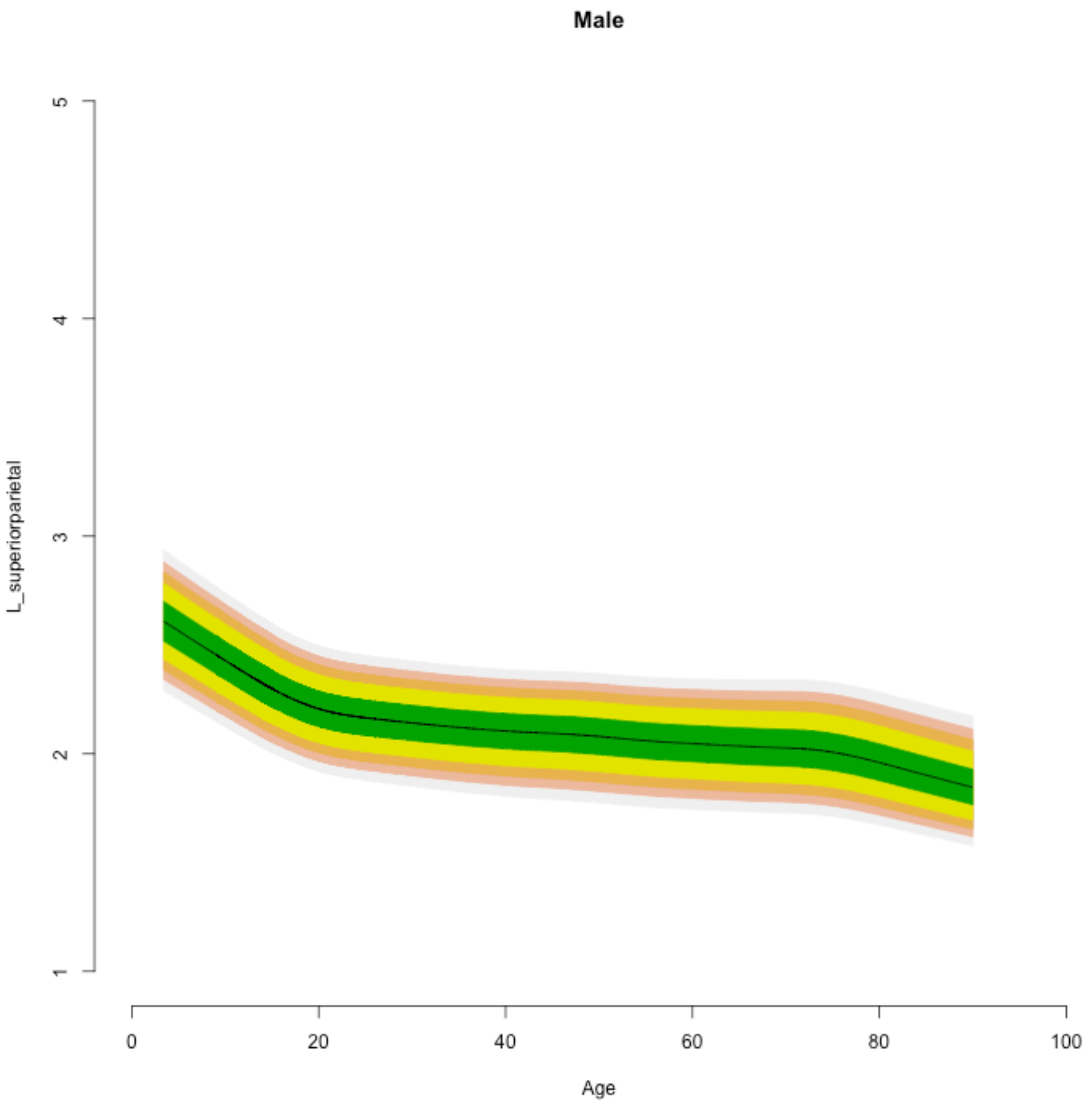

Male
L_superiorparietal
1
2
3
4
5
0
20
40
60
80
100
Age

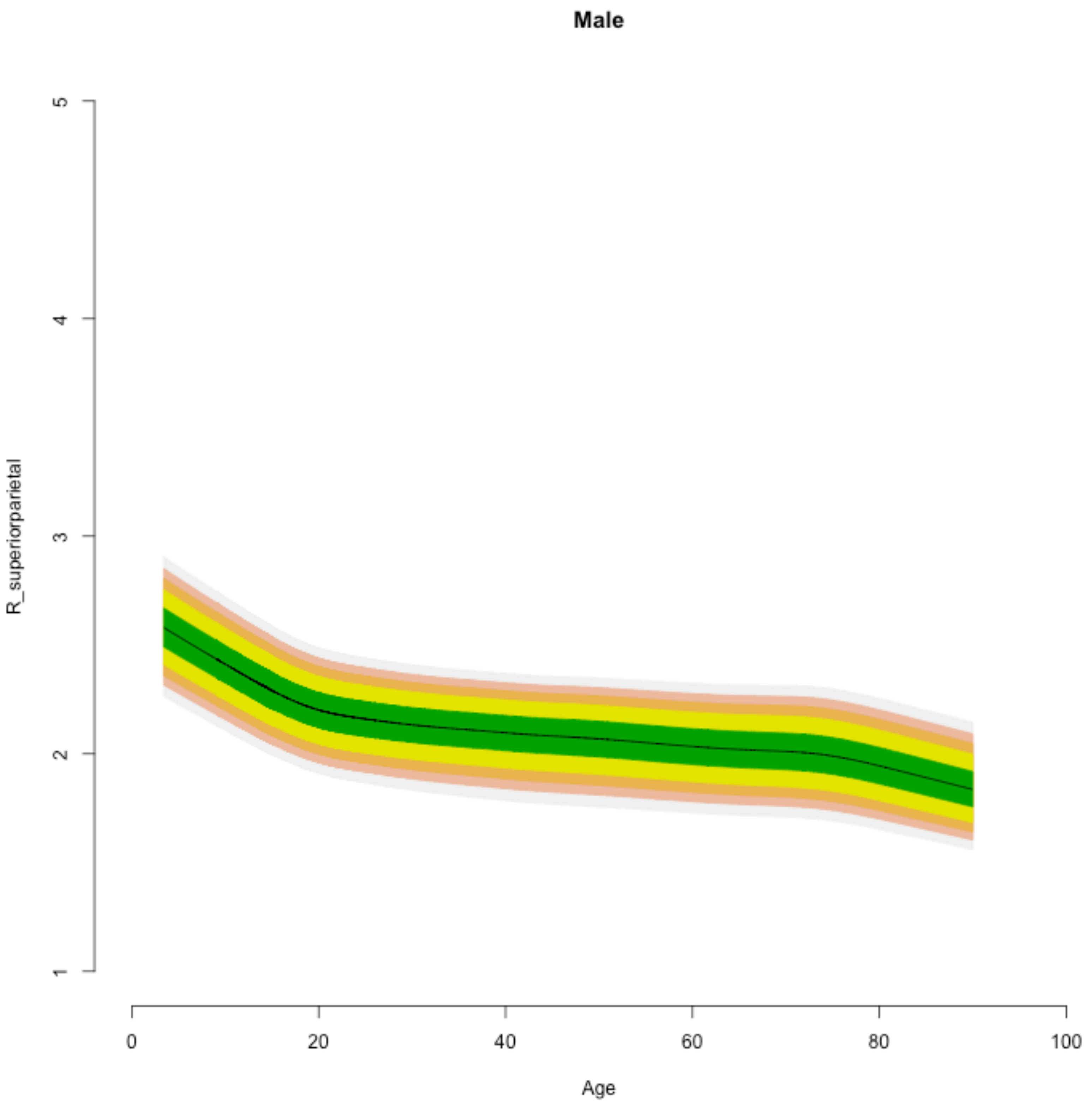
Male
R_superiorparietal
5
4
3
2
1
0
20
40
60
80
100
Age

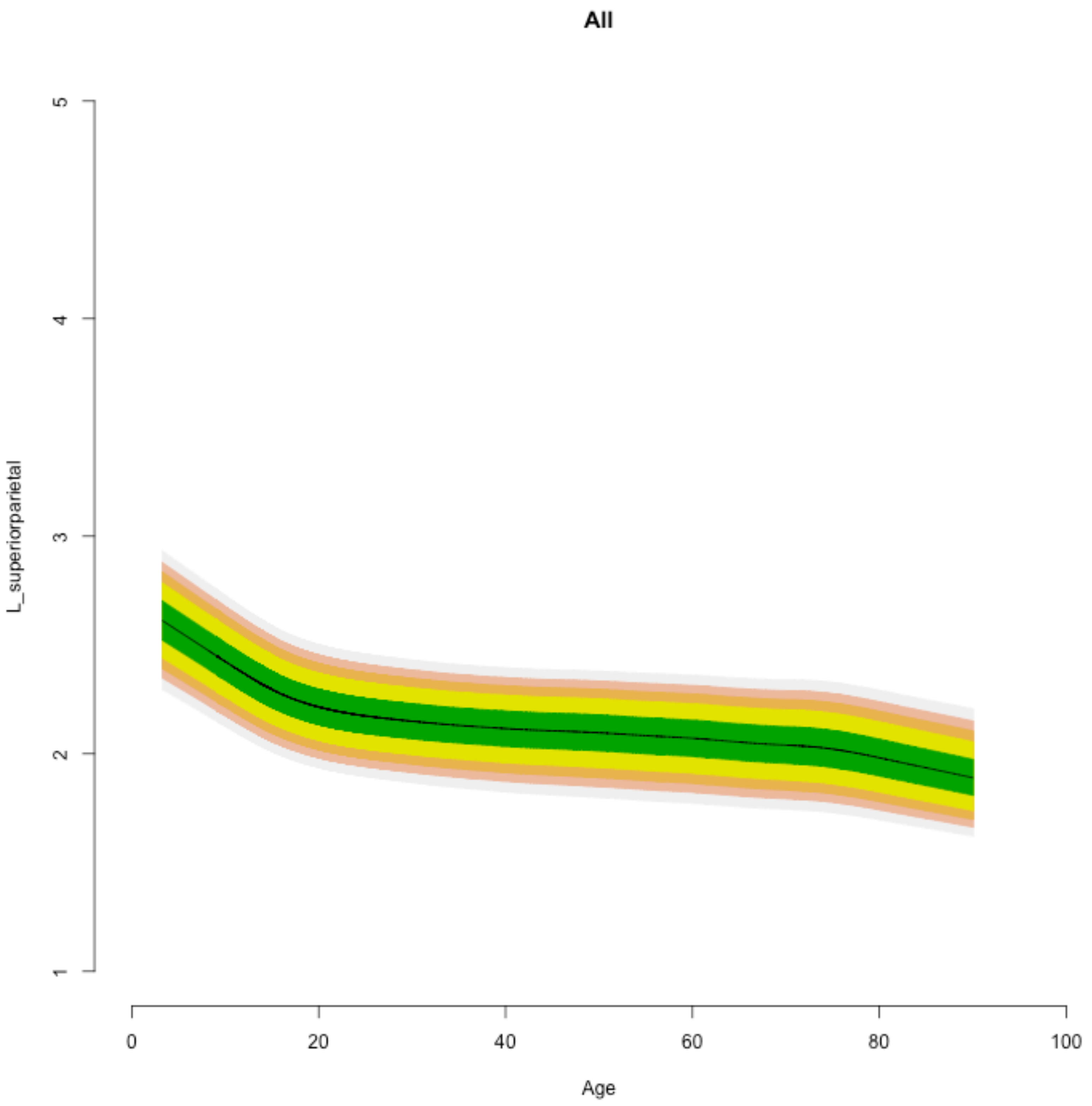
All
L_superiorparietal
5
4
3
2
1
0
20
40
60
80
100
Age

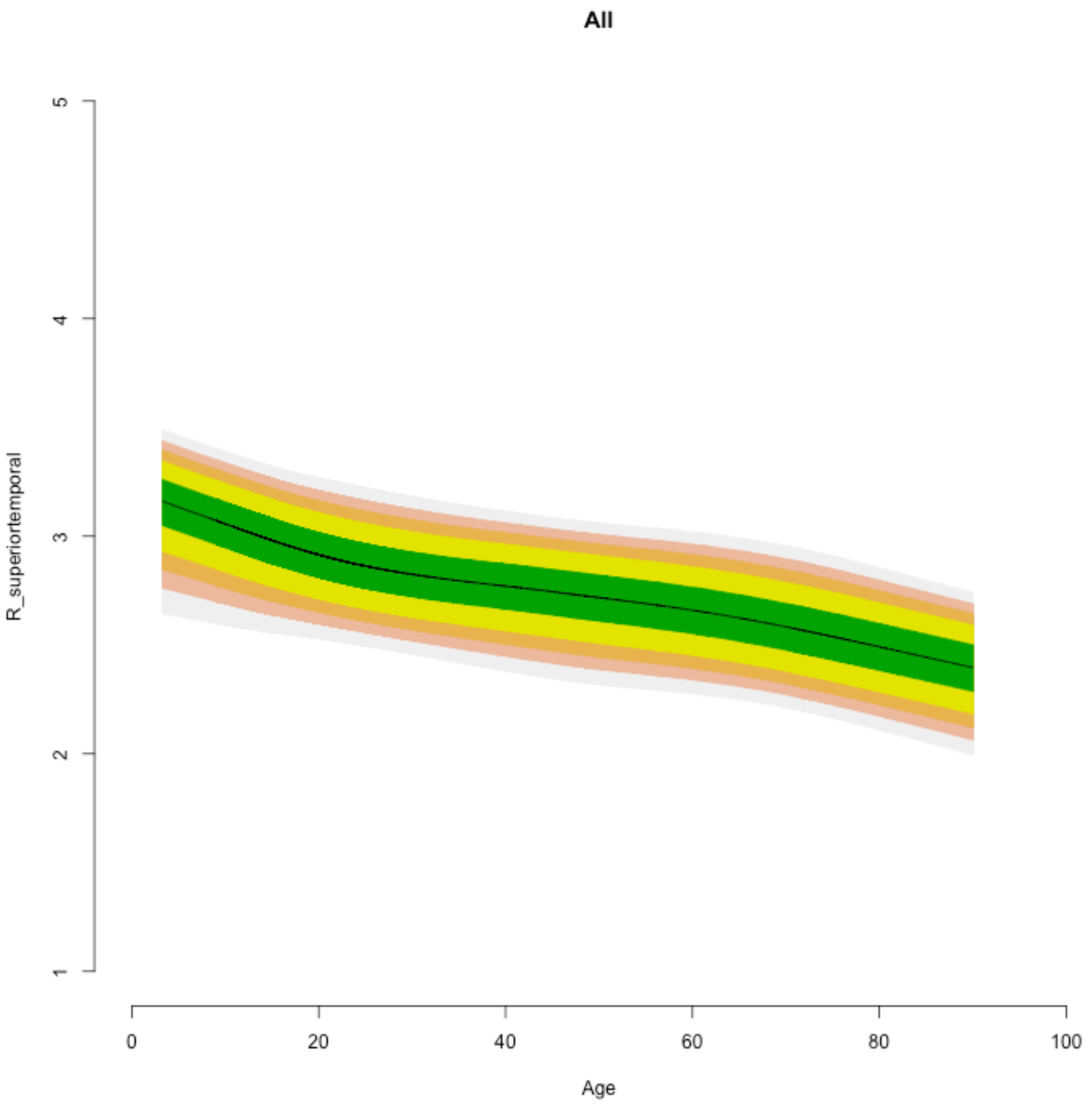

All
R_superiortemporal
5
4
3
2
1
0
20
40
60
80
100
Age

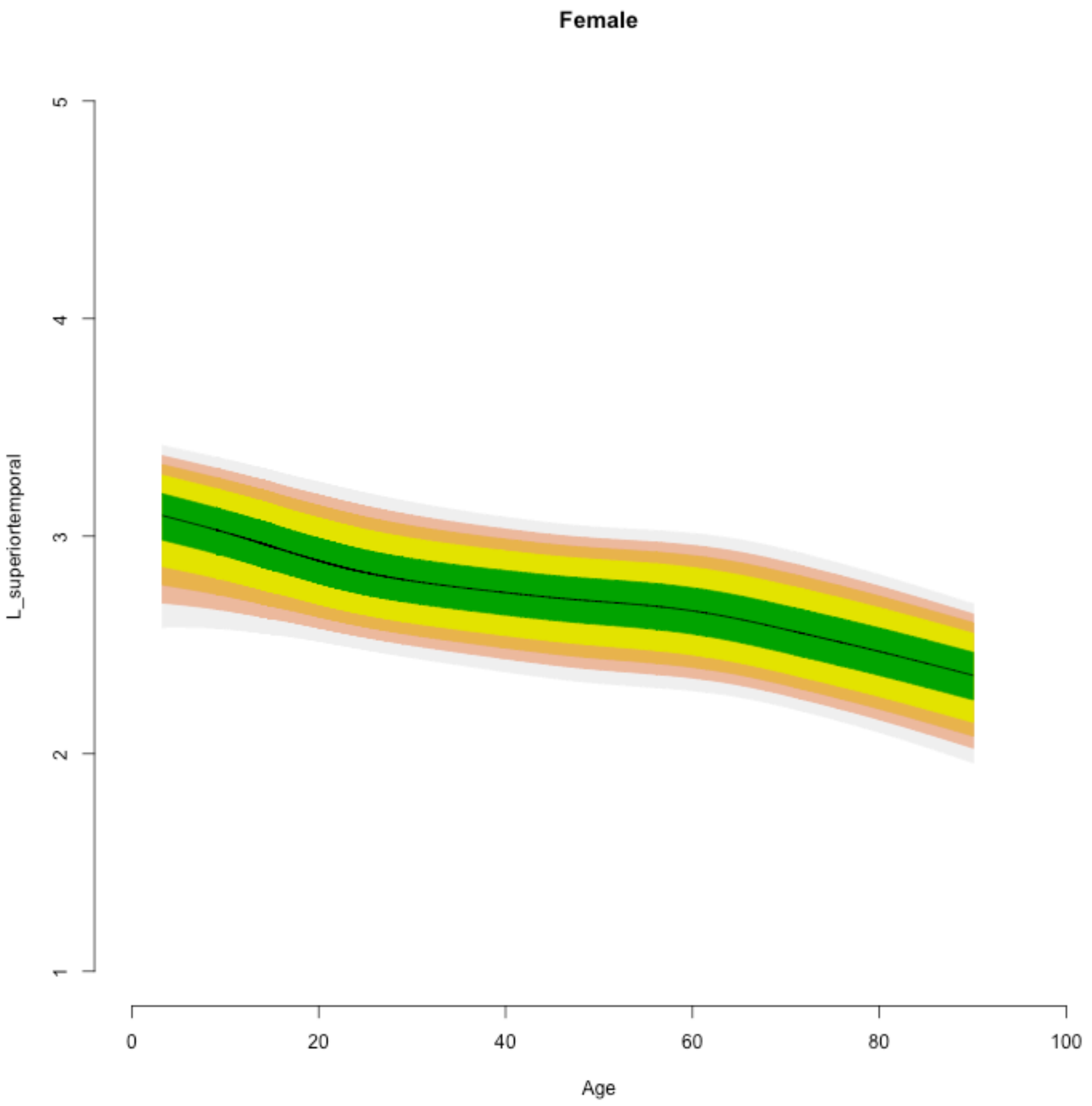
Female
L_superiortemporal
1
2
3
4
5
0
20
40
60
80
100
Age

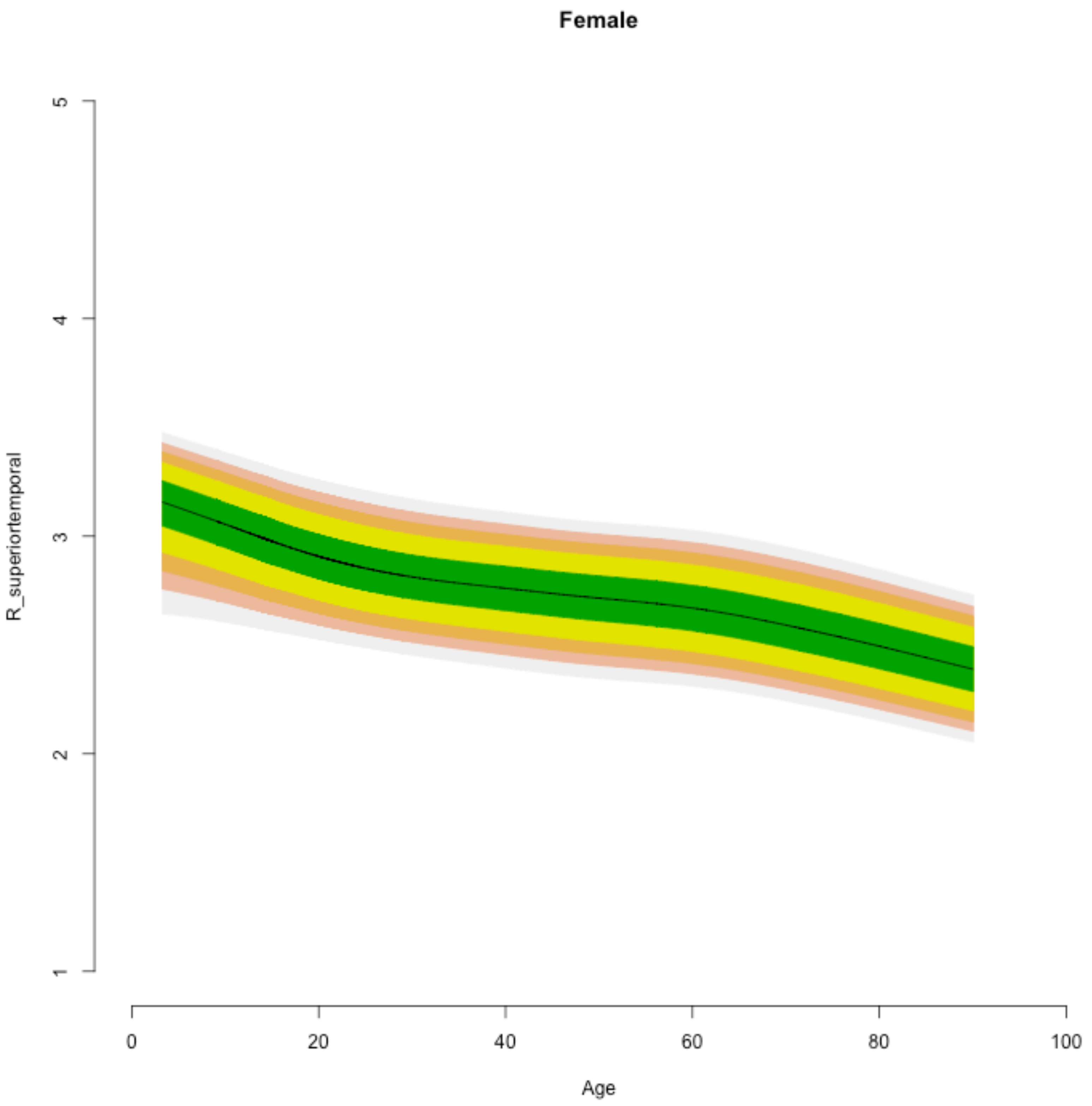
Female
R_superiortemporal
1
2
3
4
5
0
20
40
60
80
100
Age

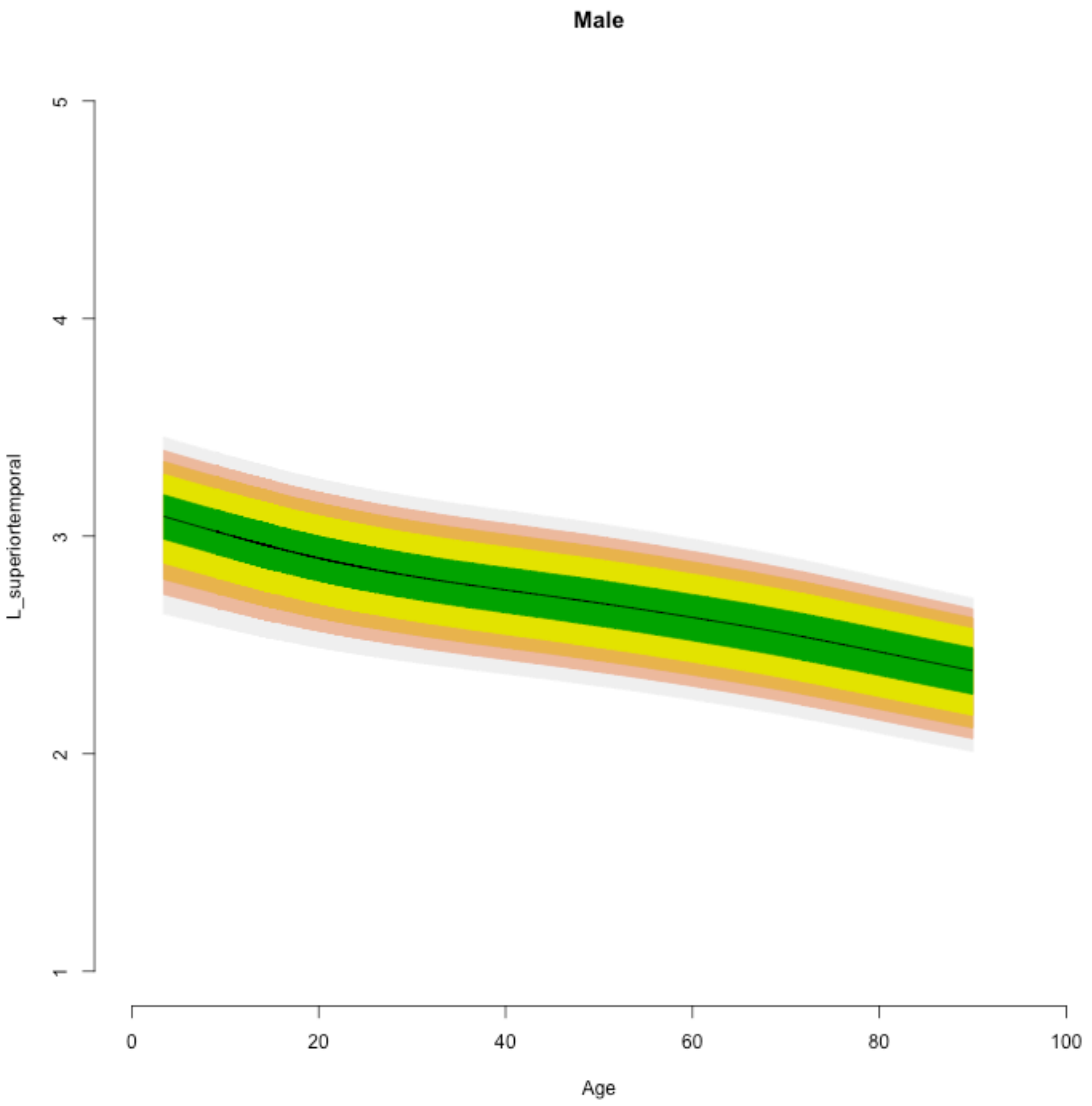
Male
L_superiortemporal
1
2
3
4
5
0
20
40
60
80
100
Age

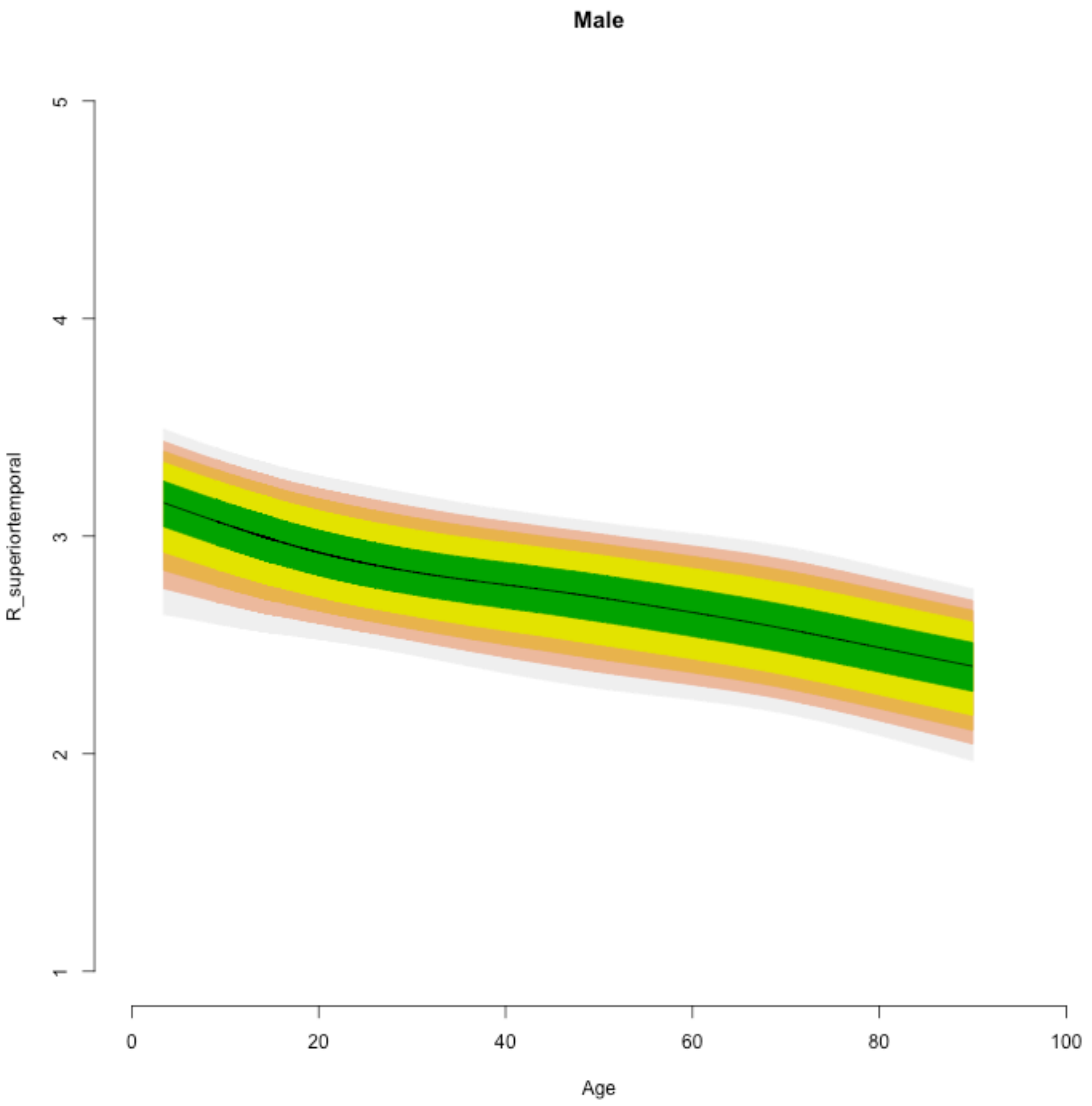
Male
R_superiortemporal
1
2
3
4
5
0
20
40
60
80
100
Age

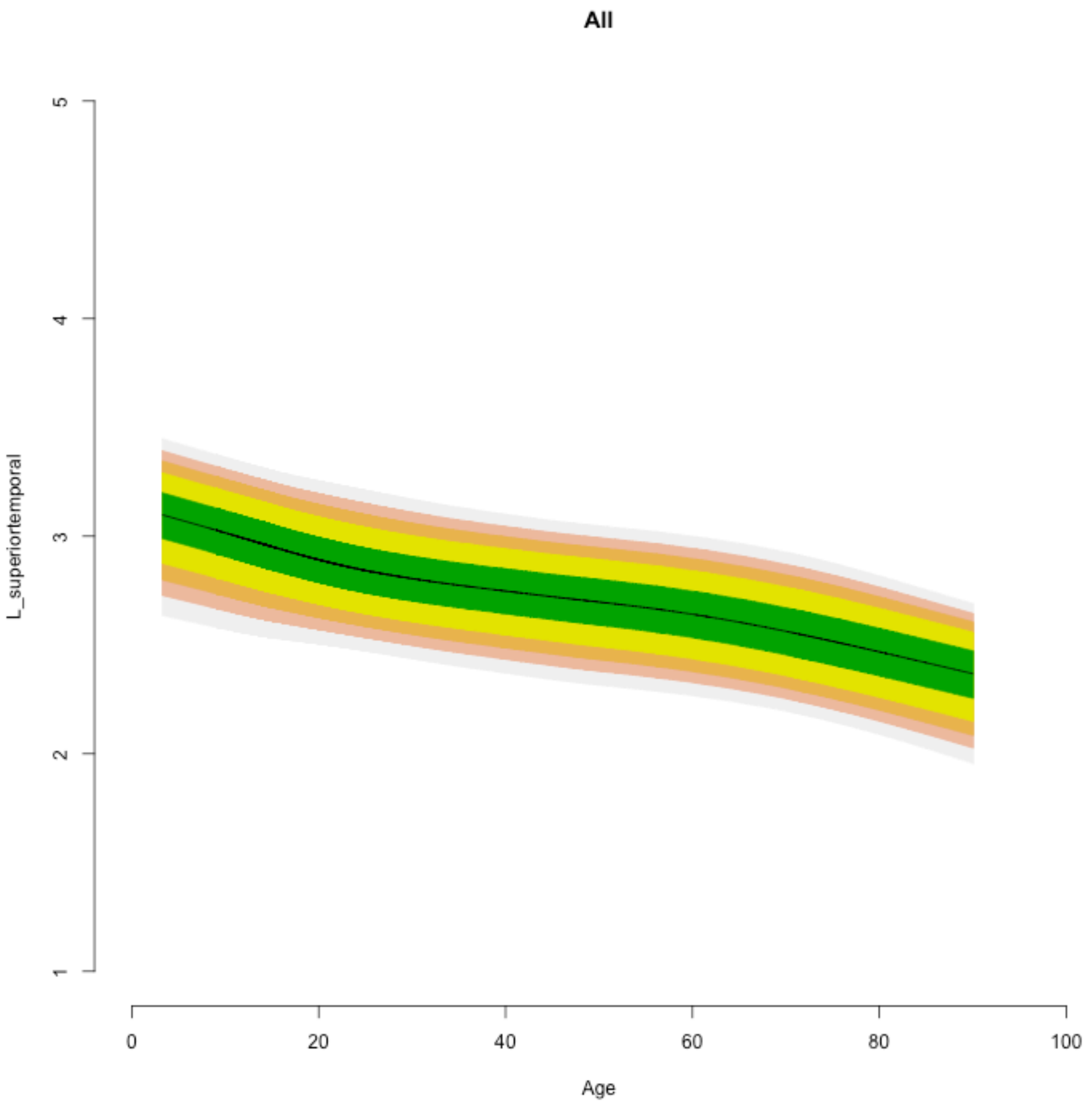

All
L_superiortemporal
1
2
3
4
5
0
20
40
60
80
100
Age

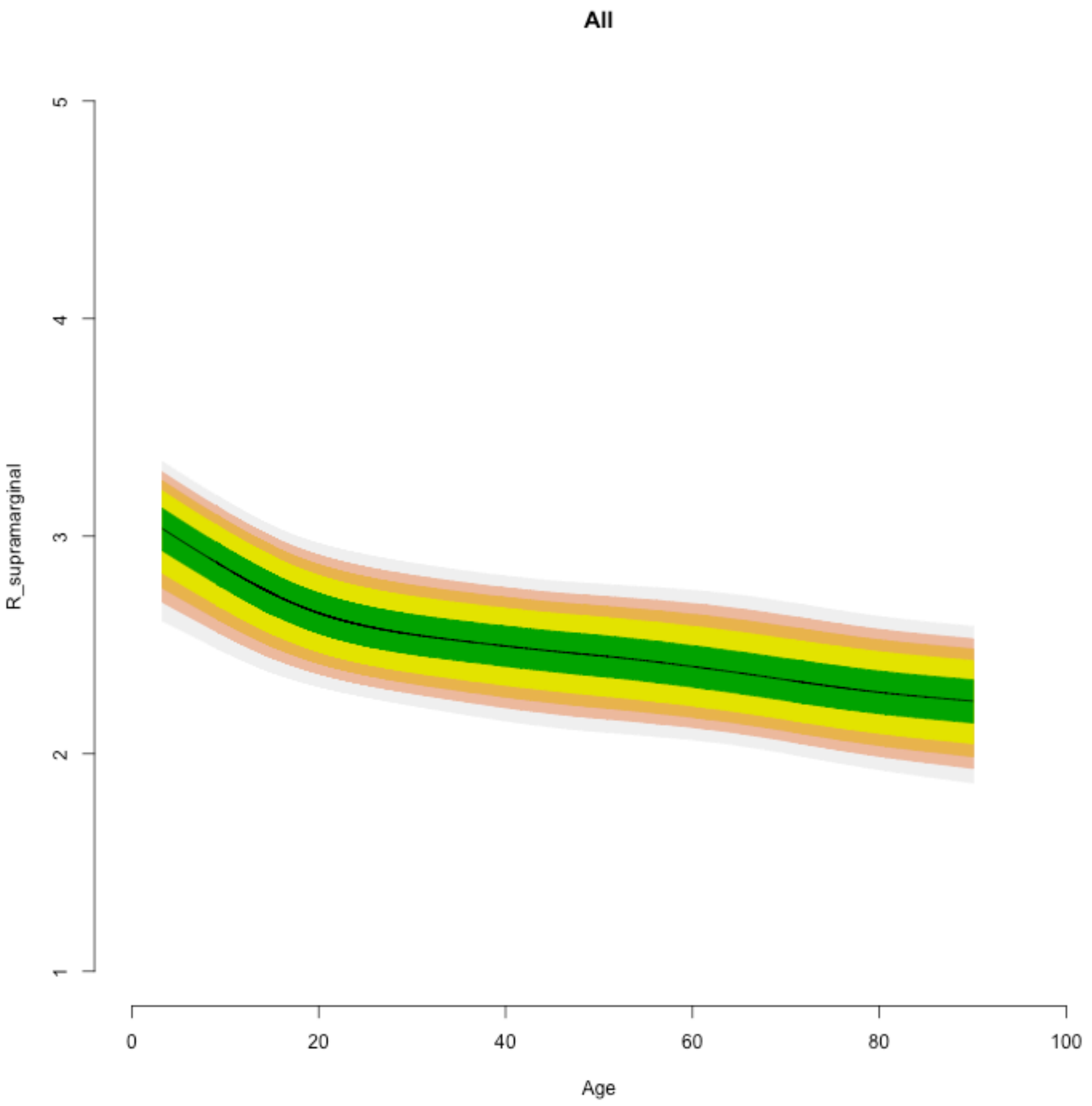

All
R_supramarginal
1
2
3
4
5
0
20
40
60
80
100
Age

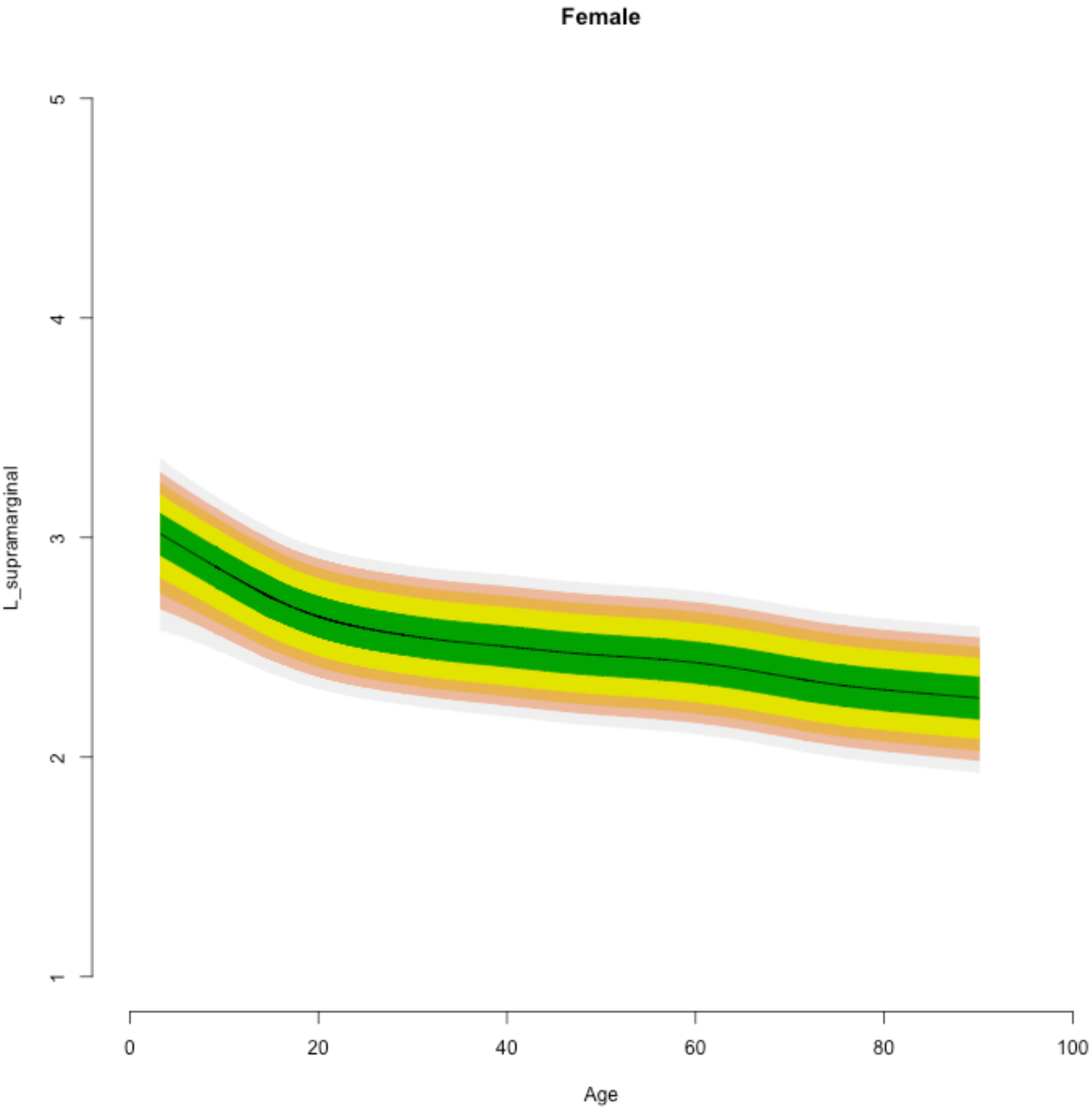
Female
L_supramarginal
Age
1
2
3
4
5
0
20
40
60
80
100

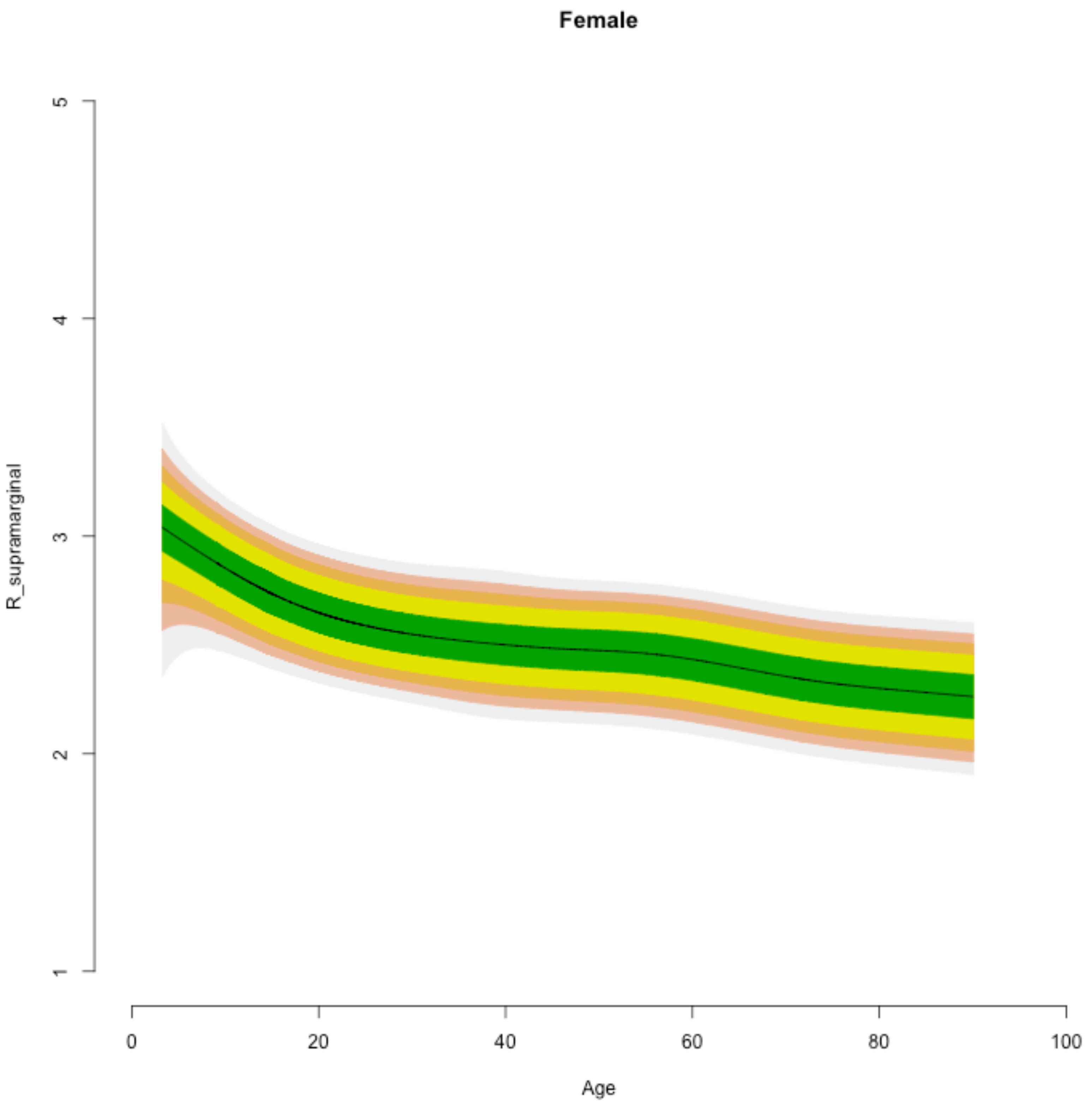
Female
R_supramarginal
1
2
3
4
5
0
20
40
60
80
100
Age

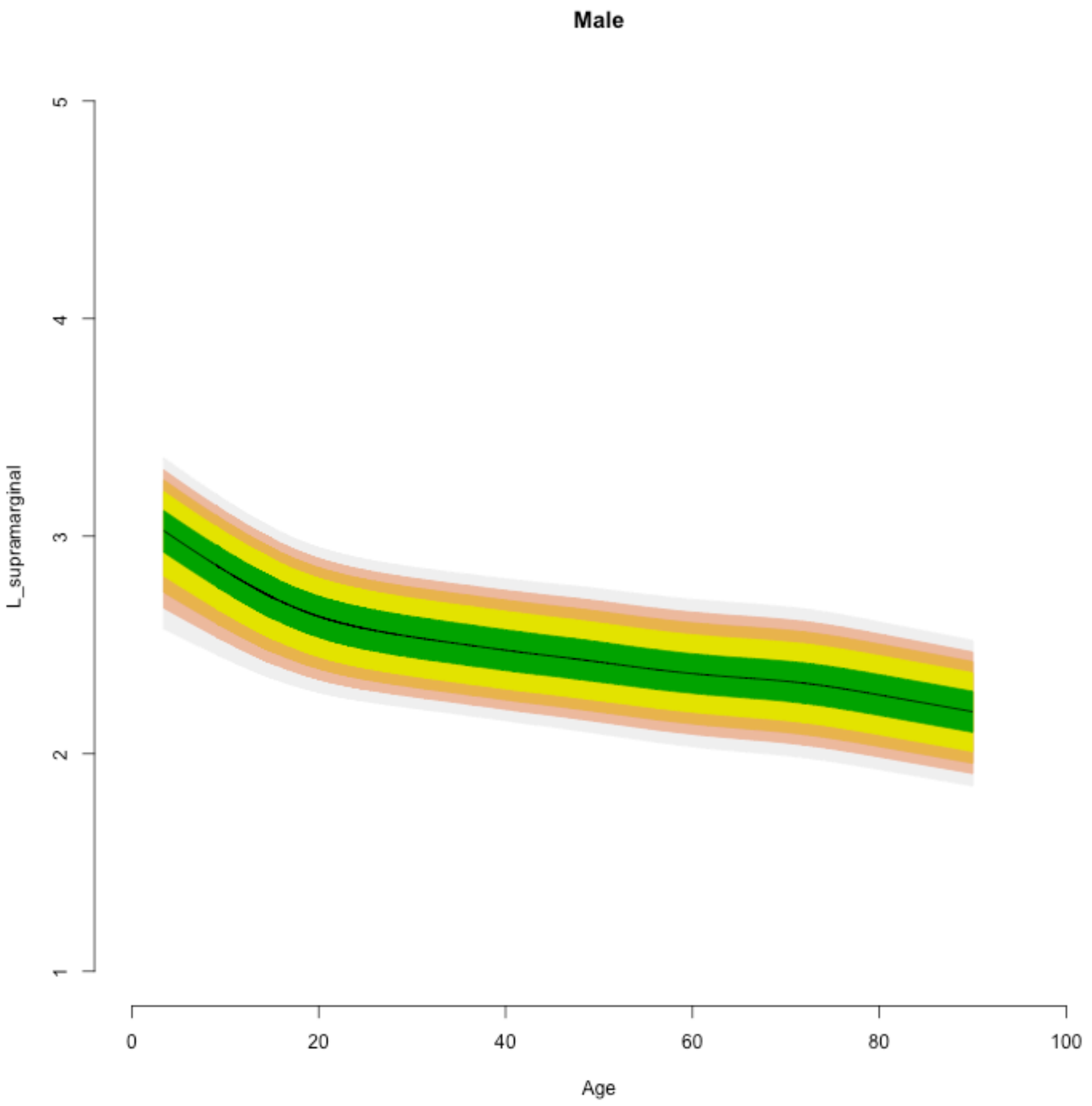
Male
L_supramarginal
Age
1
2
3
4
5
0
20
40
60
80
100

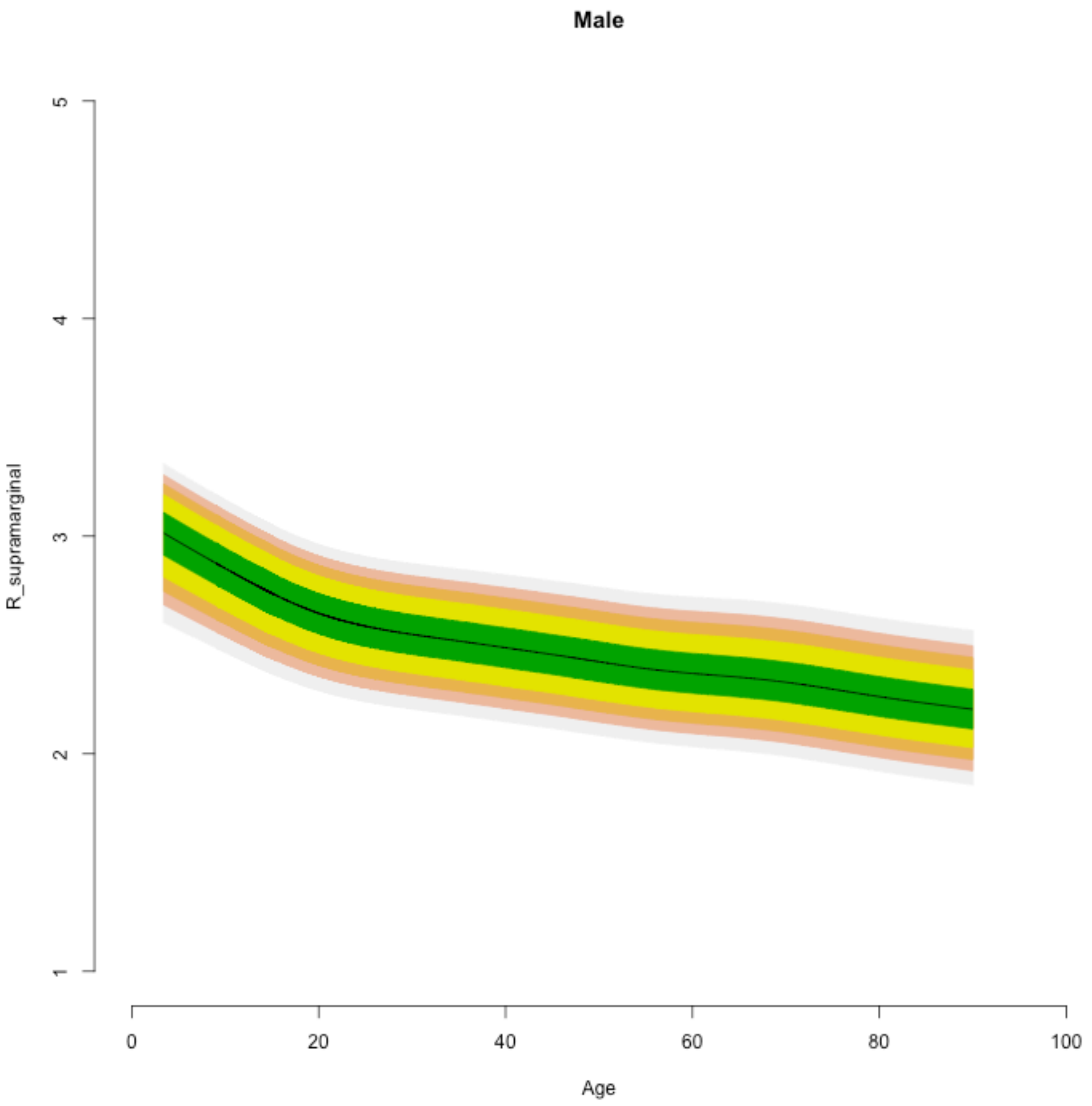

Male
R_supramarginal
5
4
3
2
1
0
20
40
60
80
100
Age

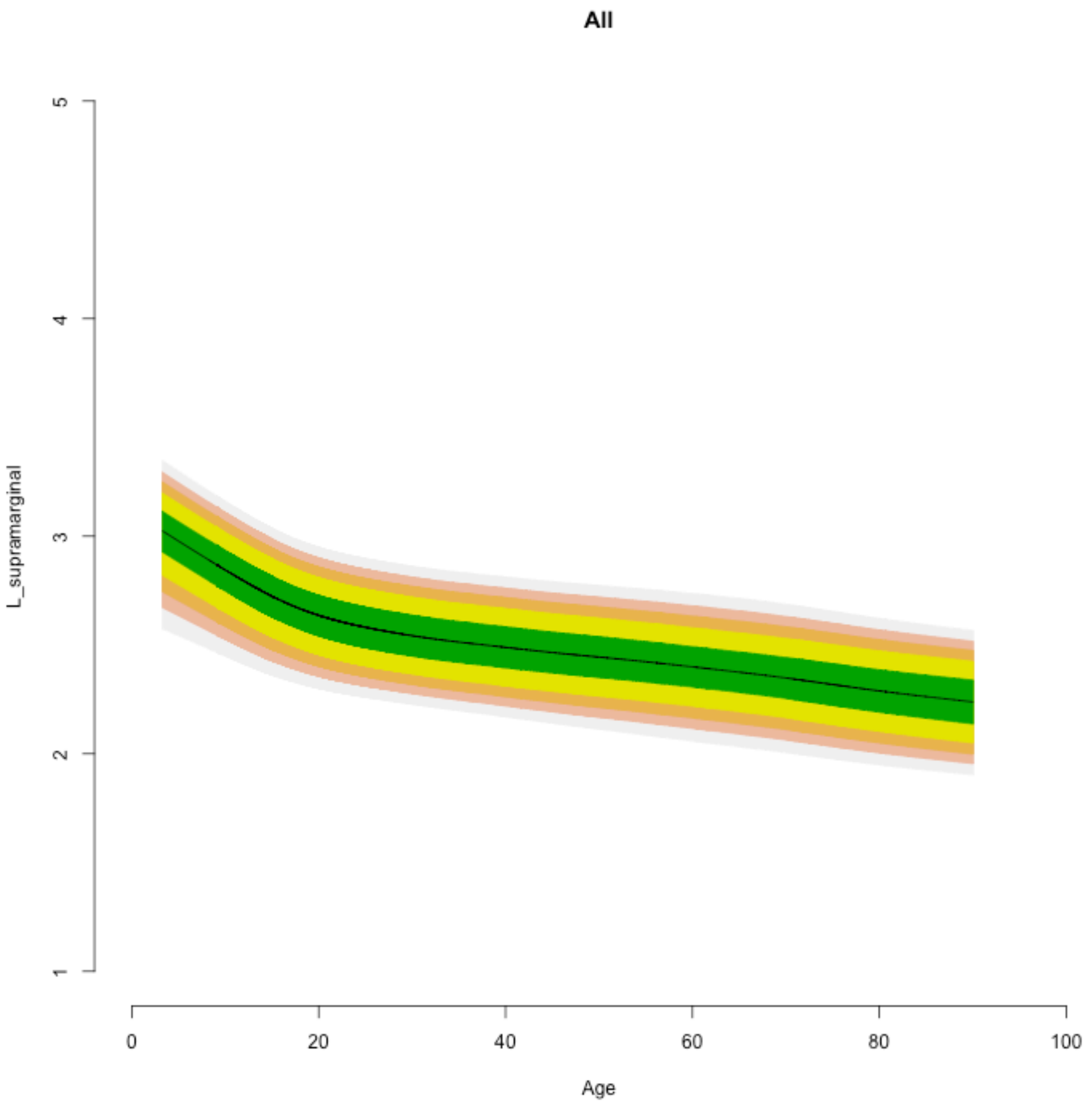

All
L_supramarginal
1
2
3
4
5
0
20
40
60
80
100
Age

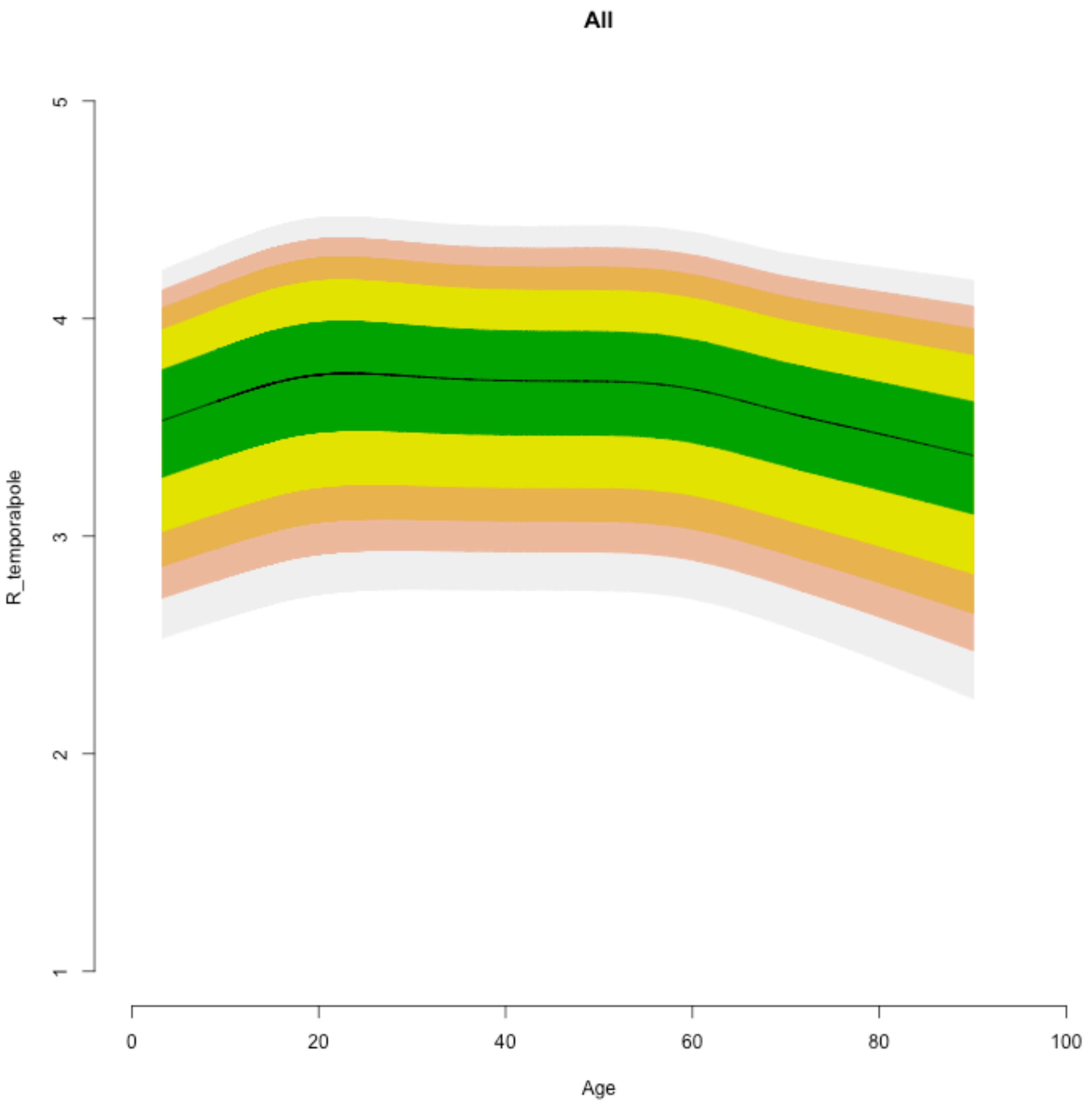
All
R_temporalpole
1
2
3
4
5
0
20
40
60
80
100
Age

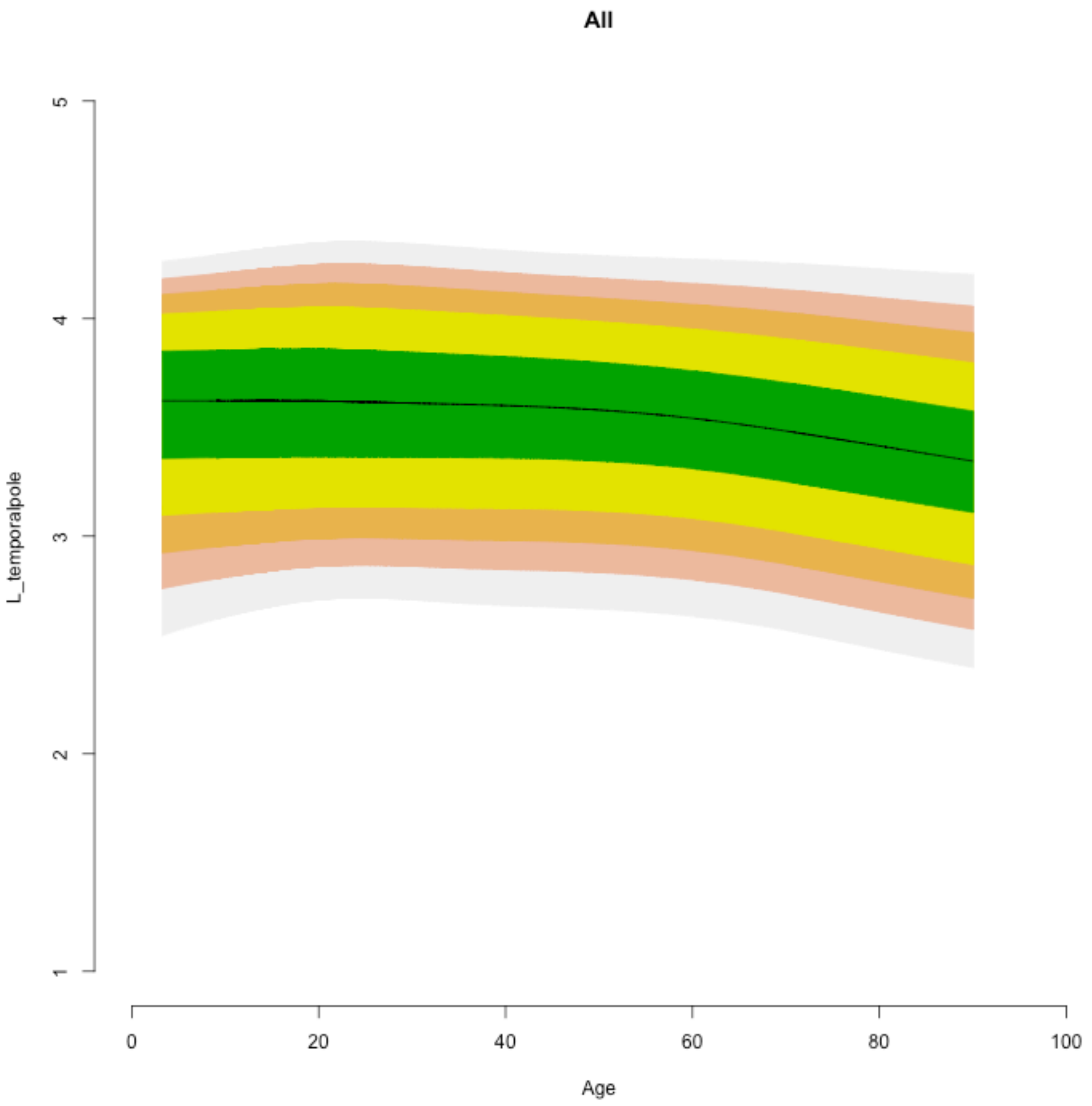
All
L_temporalpole
Age
0
20
40
60
80
100
1
2
3
4
5

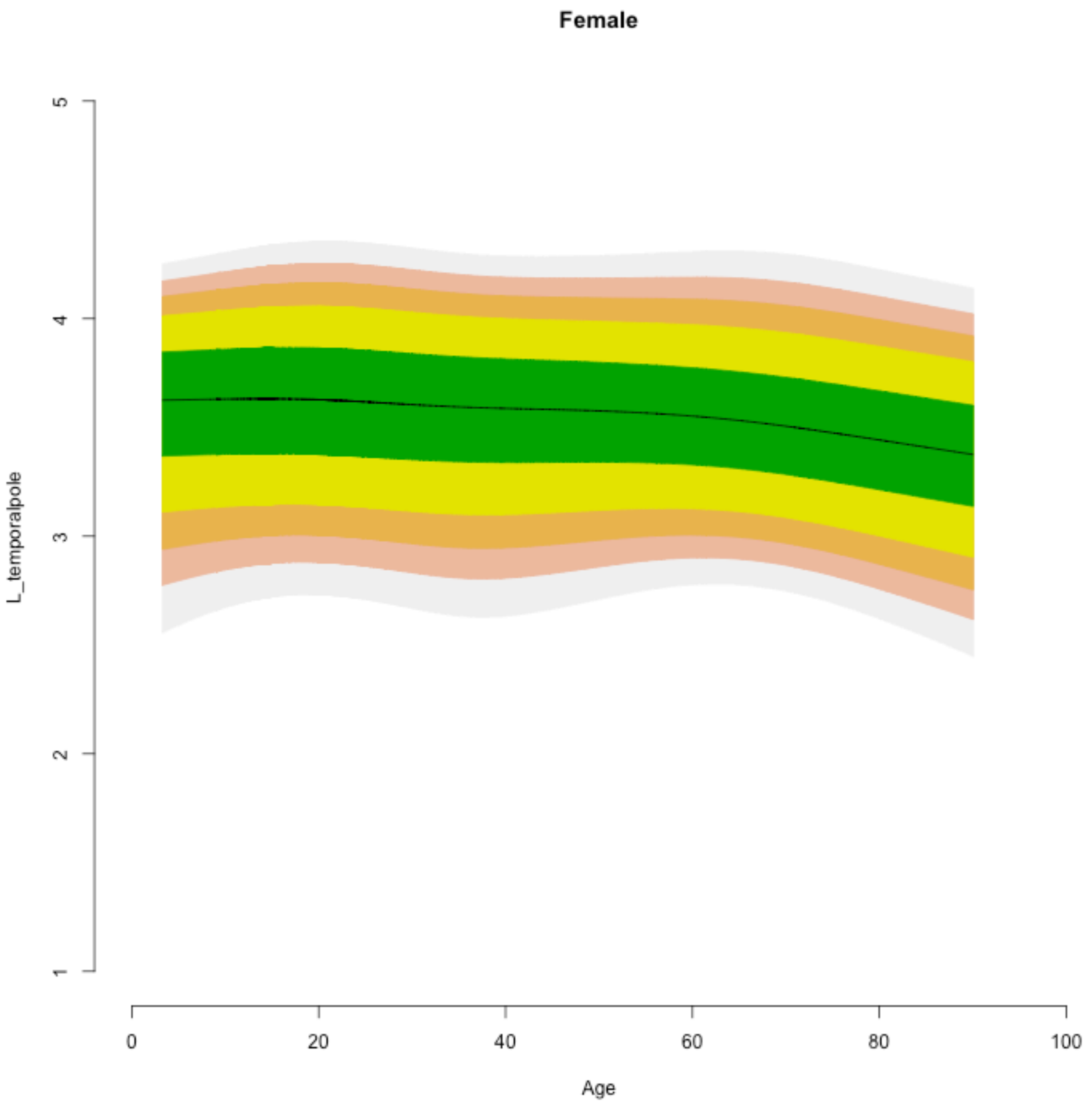

Female
L_temporalpole
Age
1
2
3
4
5
0
20
40
60
80
100

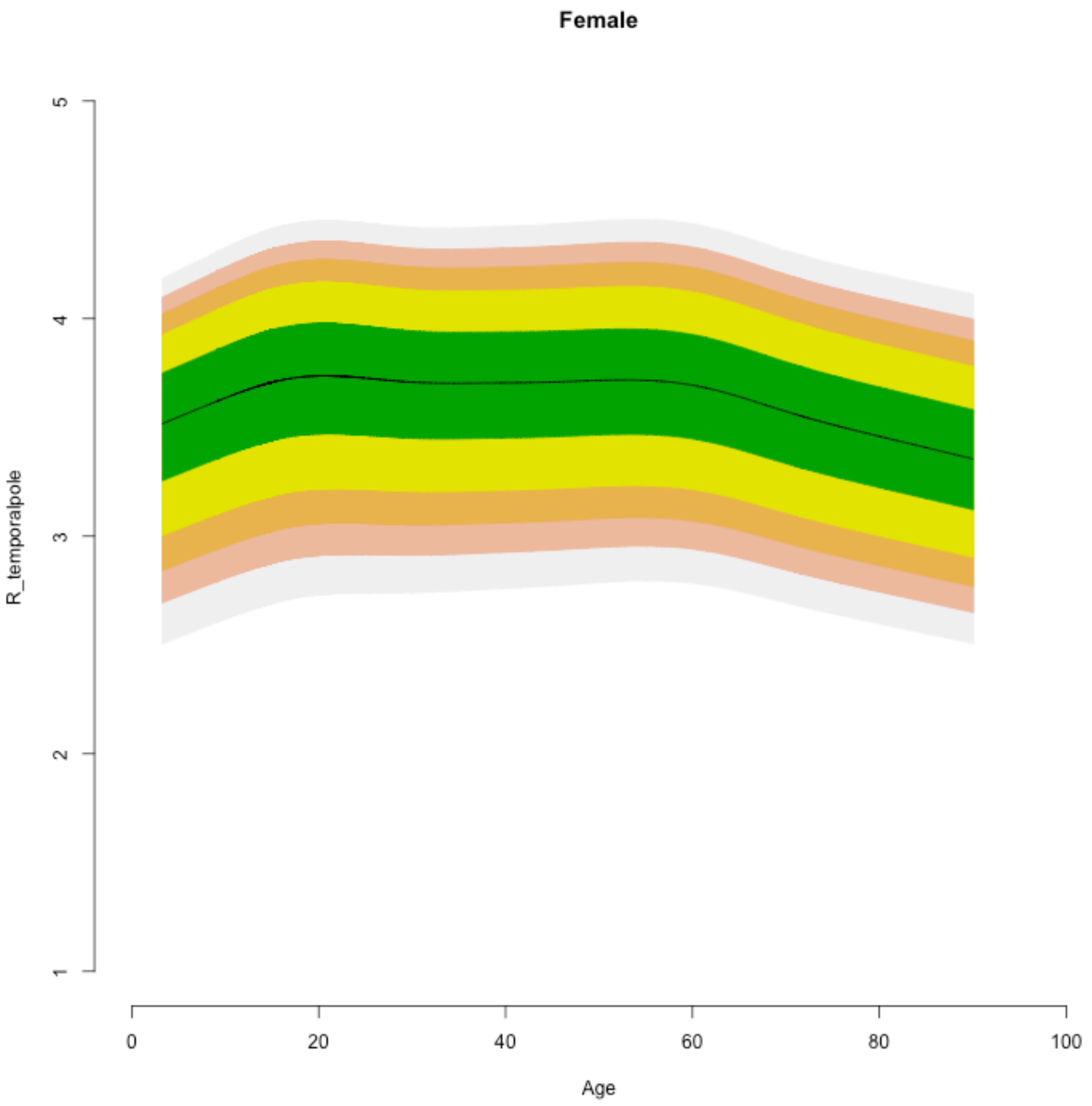
Female
R_temporalpole
5
4
3
2
1
0
20
40
60
80
100
Age

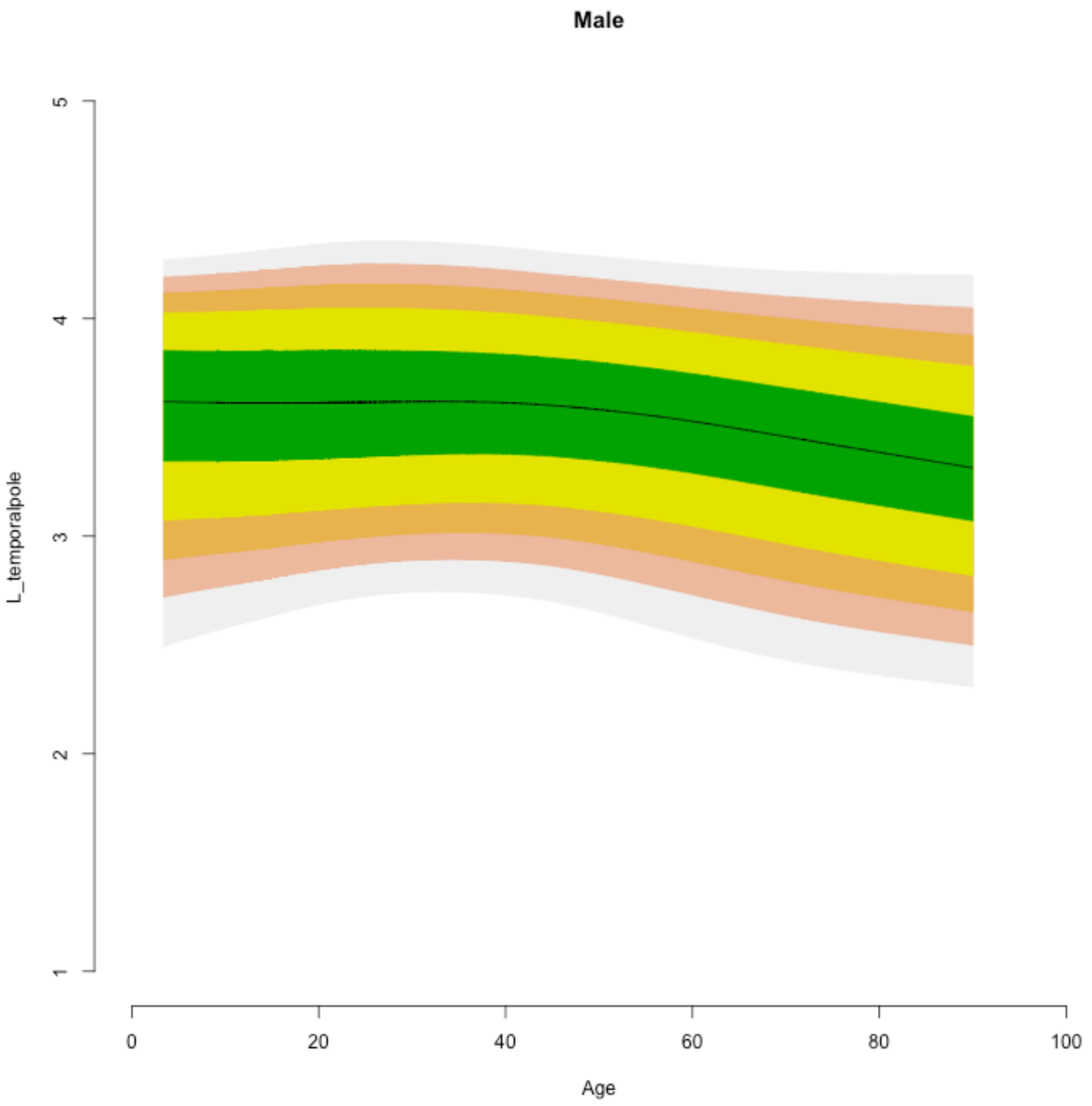
Male
L_temporalpole
1
2
3
4
5
0
20
40
60
80
100
Age

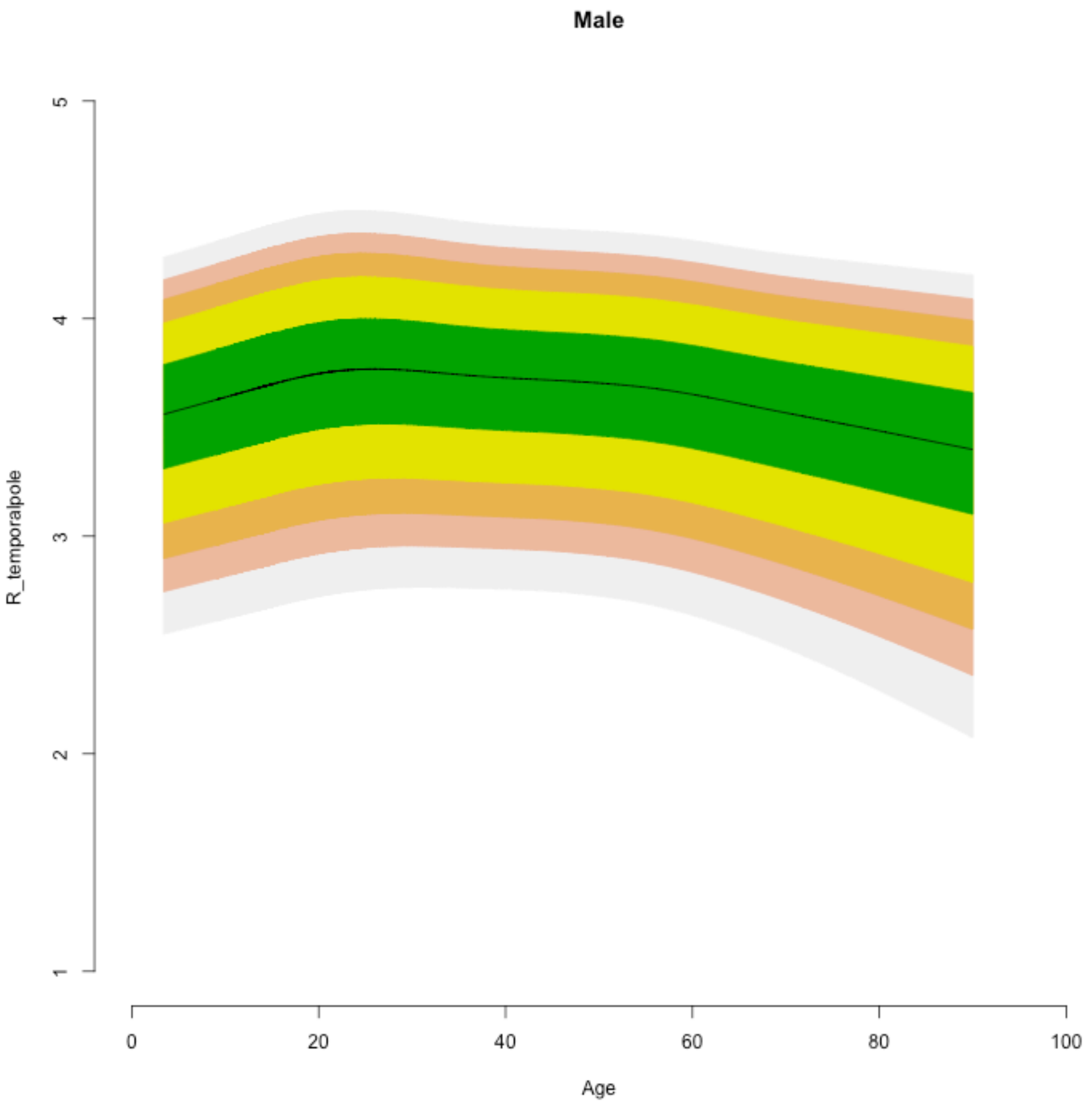
Male
R_temporalpole
1
2
3
4
5
0
20
40
60
80
100
Age

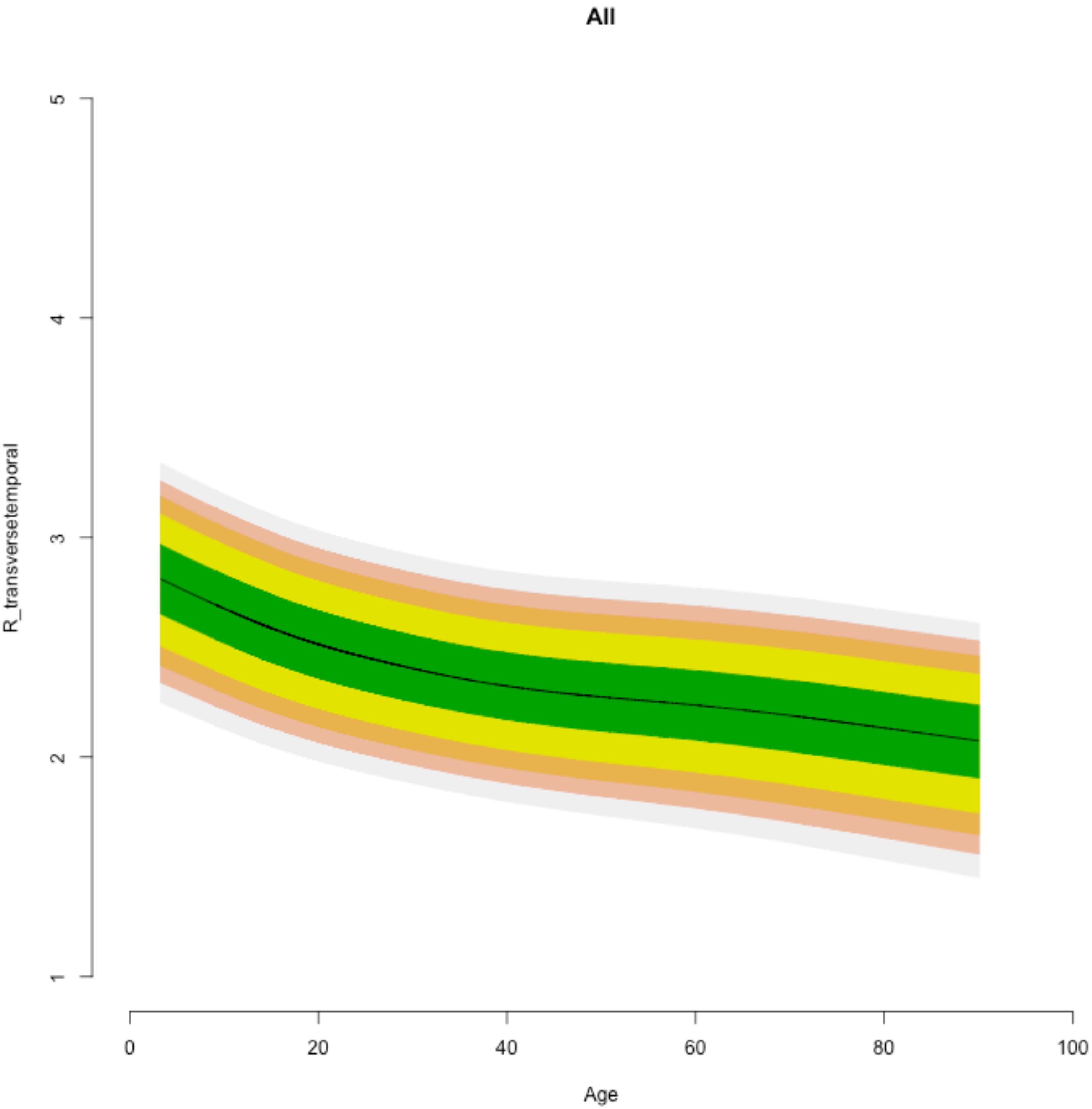

All
R_transversetemporal
5
4
3
2
1
0
20
40
60
80
100
Age

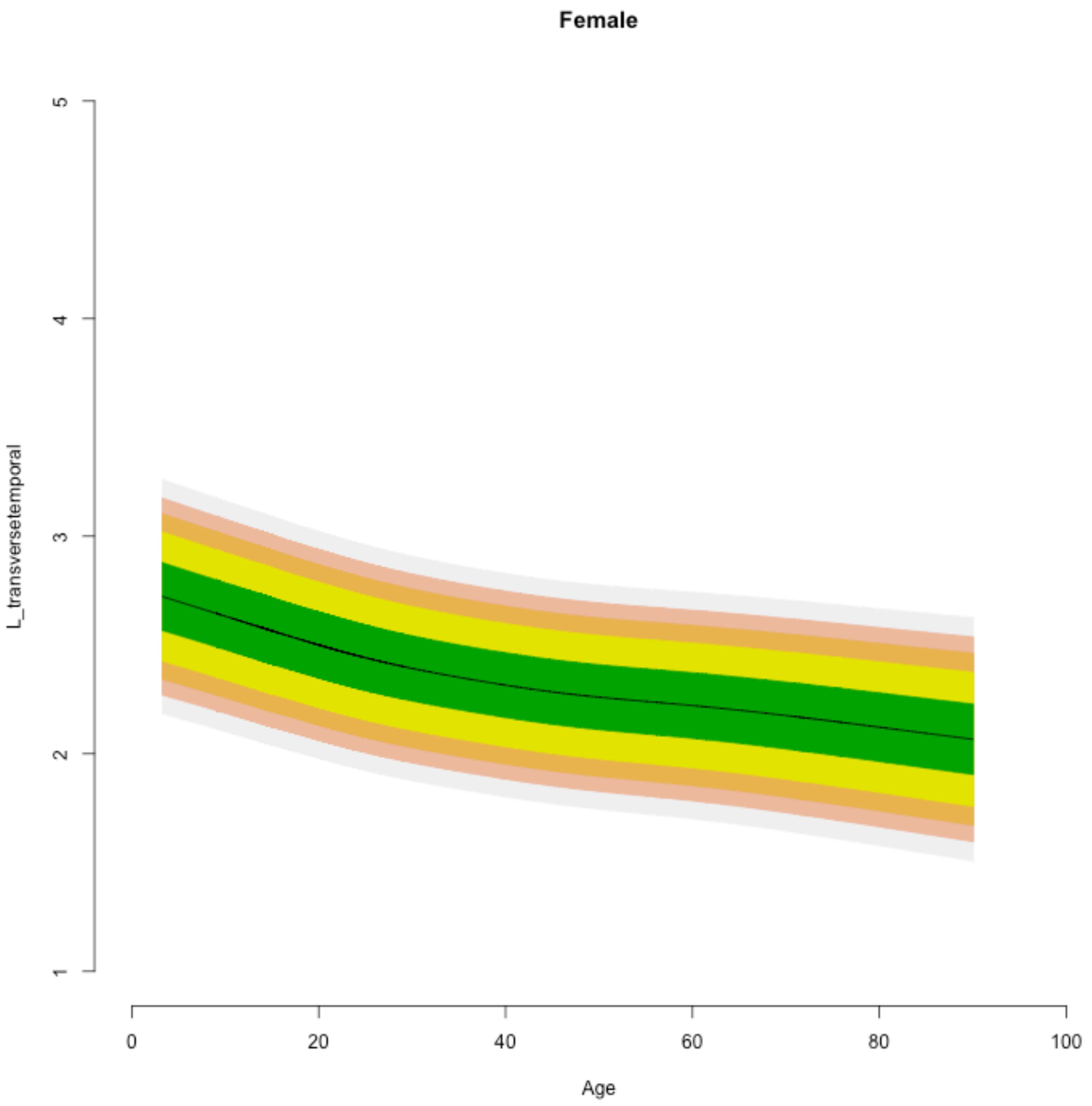
Female
L_transversetemporal
1
2
3
4
5
0
20
40
60
80
100
Age

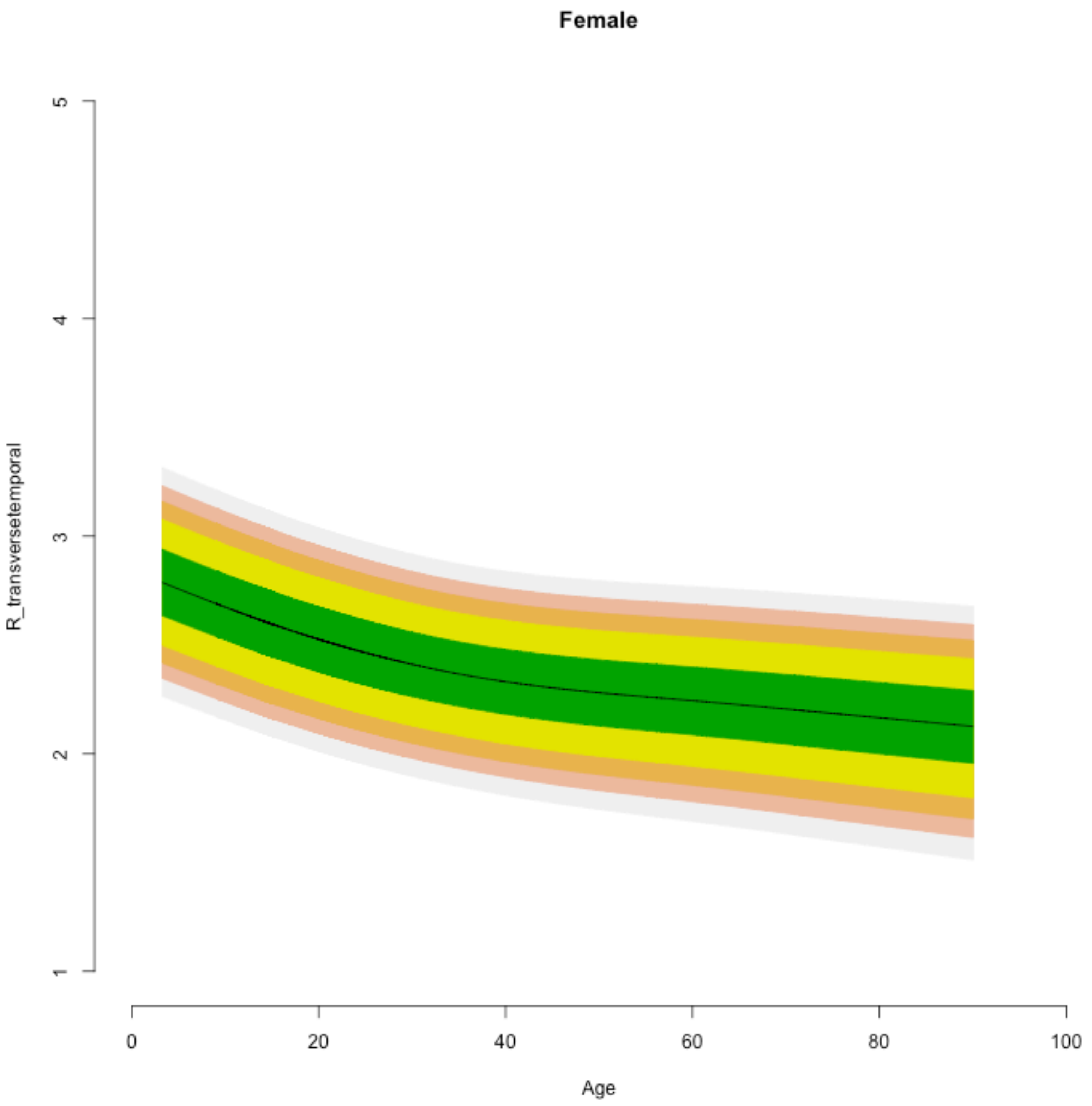
Female
R_transversetemporal
1
2
3
4
5
0
20
40
60
80
100
Age

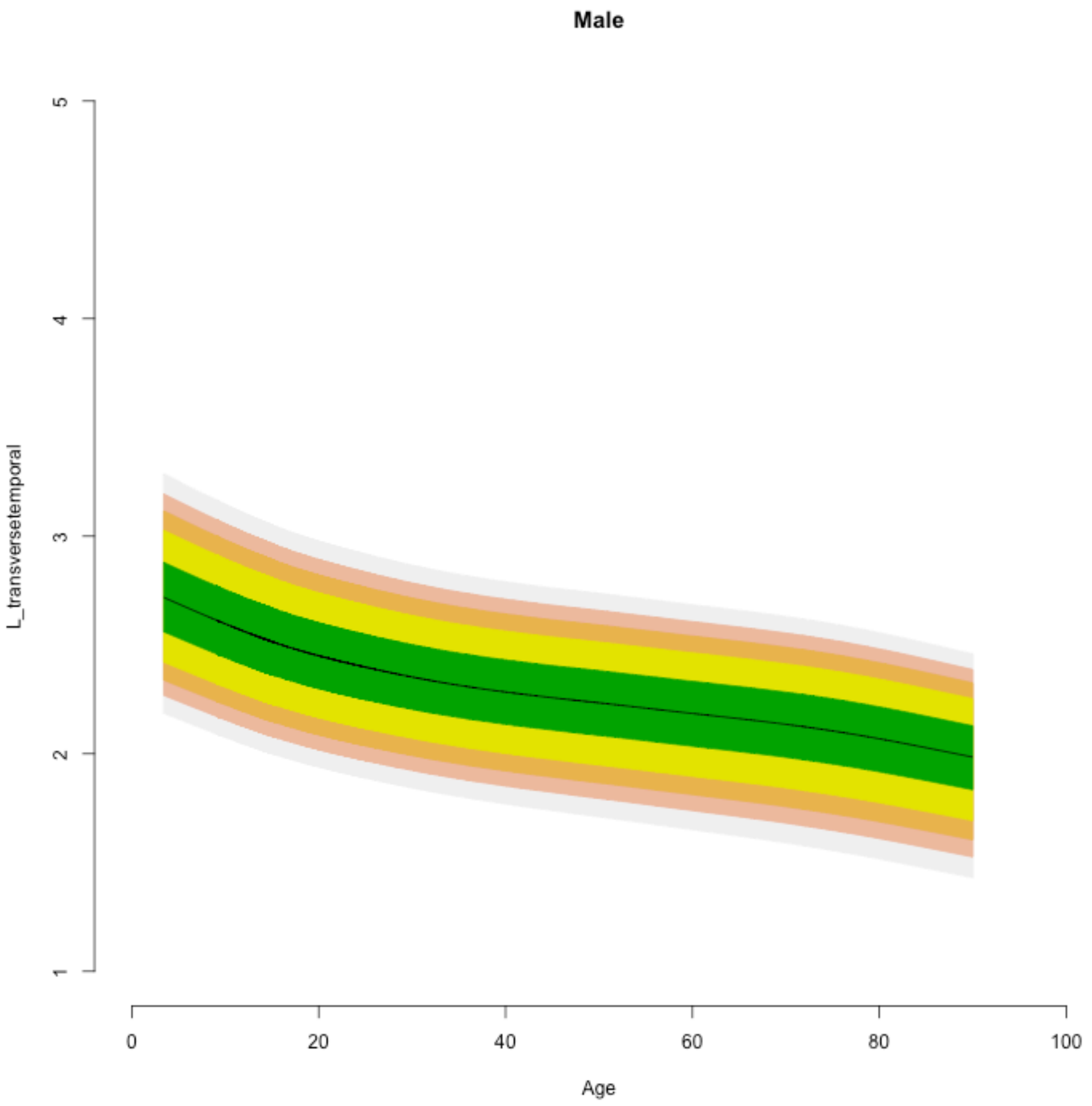
Male
L_transversetemporal
1
2
3
4
5
0
20
40
60
80
100
Age

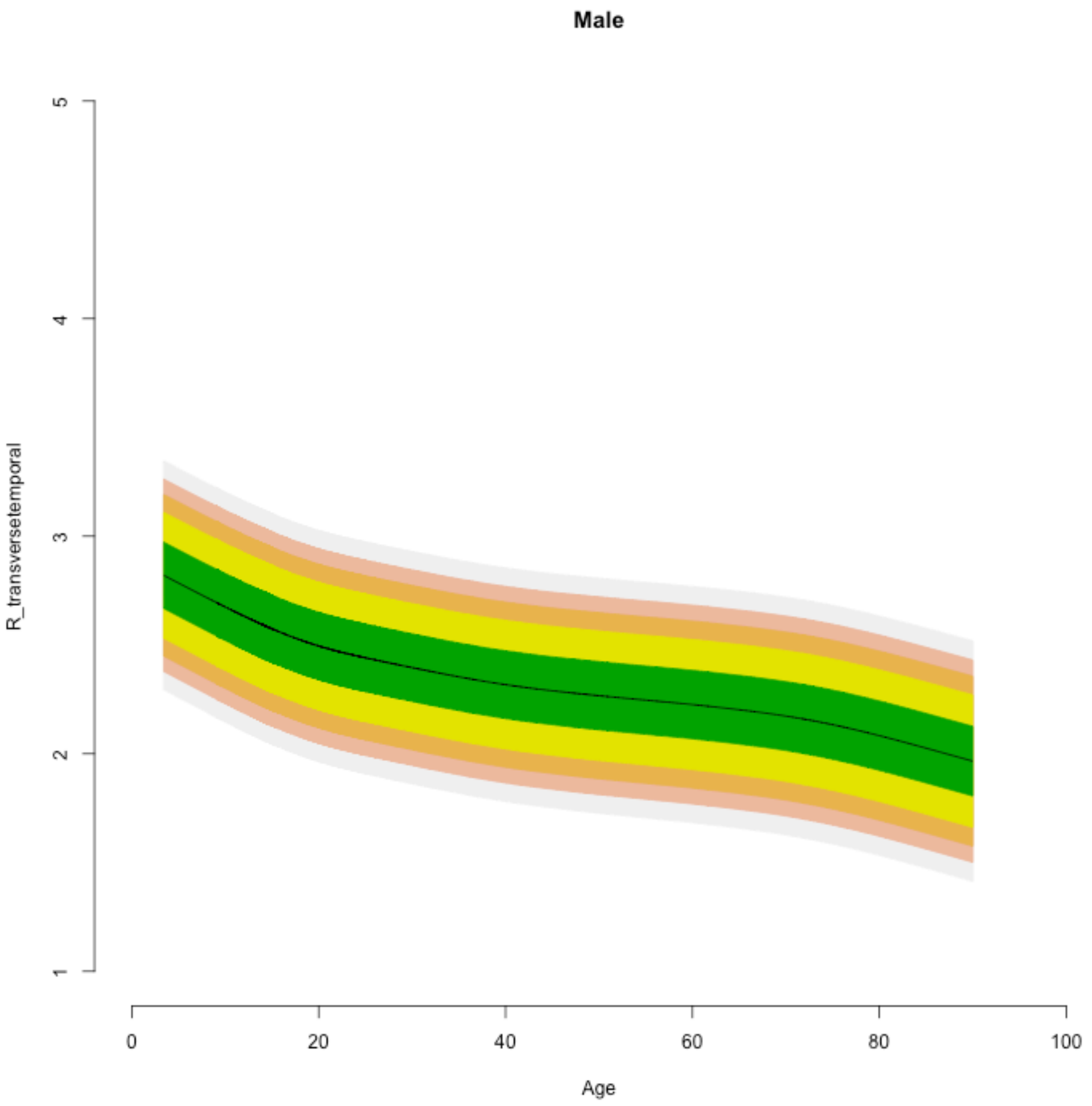

Male
R_transversetemporal
1
2
3
4
5
0
20
40
60
80
100
Age

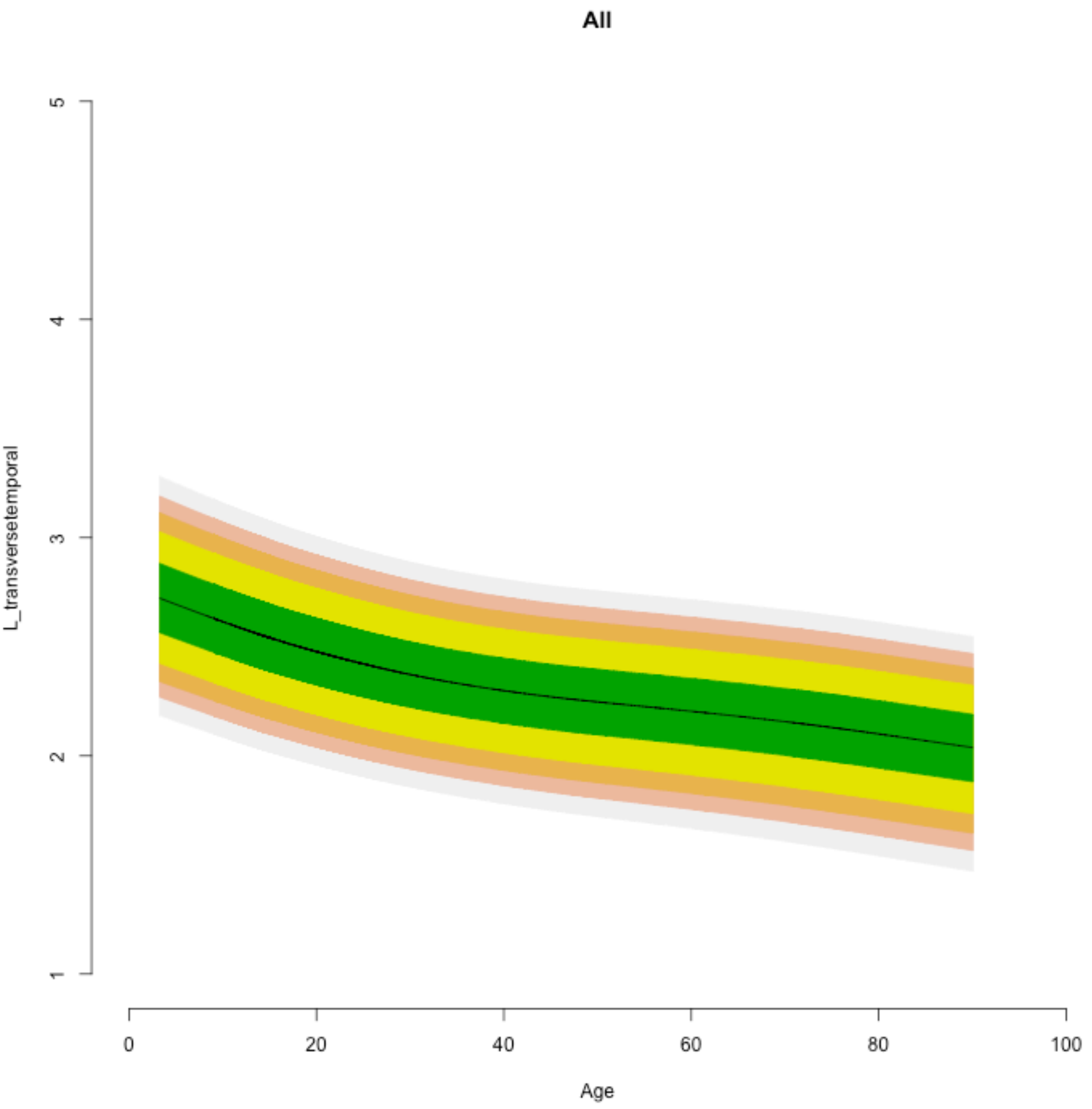
All
L_transversetemporal
1
2
3
4
5
0
20
40
60
80
100
Age
